# The limit of life at extremely low water activity: Lithium-concentration ponds in a solar saltern (Salar de Atacama, Chile)

**DOI:** 10.1101/2024.12.31.629924

**Authors:** C. Demergasso, P Avendaño, C. Escuti, R. Veliz, G. Chong, C. Pedrós-Alió

**Affiliations:** Centro de Biotecnología, Universidad Católica del Norte, Antofagasta, Chile; Departamento de Ciencias Geológicas, Universidad Católica del Norte, Antofagasta, Chile; Department of Systems Biology, Centro Nacional de Biotecnología, CSIC, Madrid, Spain

**Keywords:** lithium solar salterns, Proteobacteria, Haloarchaea, extremophiles, hypersaline, Atacama, water activity

## Abstract

Extremely hyper-saline ponds from an industrial lithium-concentration process in solar salterns in the Atacama Desert were studied to determine the limits of life at very low water activity. Water activity (a_w_) of 0.61 is the lowest a_w_ value for growth of living beings recorded to date. Xerophilic (sometimes called osmophilic) filamentous fungi and yeasts are predominant in high-sugar foods with such low a_w_ values. Some microorganisms are capable of growth at that water activity level. By contrast, high-salt environments are almost exclusively populated by prokaryotes, notably the *Halobacteria* class and some *Bacteroidetes*, capable of growing in saturated NaCl (a_w_ 0.75). The lowest a_w_ that can be achieved by the addition of NaCl is 0.75 (saturation point for NaCl). Crystallizer ponds in Li^+^ concentration plants reach down to water activity levels around 0.1. The aim of this study was to determine how far along the salinity gradient could life be found. Cell counts were attempted by epifluorescence microscopy and qPCR with bacterial and archaeal universal primers. Biomass for DNA extraction was obtained by an optimized protocol involving dialysis of brines previously fixed with ethanol. Prokaryotic diversity was studied by DNA extraction, PCR, qPCR and 16S rRNA amplicon sequencing in different ponds along the salinity gradient. Archaeal DNA was found in the lower salinity ponds, while bacterial DNA was found along the whole gradient. Bacterial cDNA was retrieved from ponds down to an a_w_ of 0.2. Moreover, bacteria could be grown in enriched cultures from most ponds.

## INTRODUCTION

Life is known to require liquid water. A water activity (a_w_) of 0.61 was previously considered to be the lowest a_w_ value for growth of living beings (Grant, 2004) and a new limit 0.585 a_w_ for microbial cell differentiation and division was established in *Aspergillus penicillioides*, with a theoretically determined limit of 0.565 (Stevenson and Hallsworth, 2014; Stevenson et al., 2015a; Stevenson et al., 2015b). This water activity is found for example in foods with high sugar content, where some fungi and yeasts are able to grow. More natural environments with low water activity are salt lakes and ponds. Many high salinity ponds form naturally, and microorganisms have had ample time to adapt to such conditions. Other extremely high salinity and Mg^2+^ rich systems include the deep lakes found at the bottom of the Mediterranean Sea (Hallsworth et al., 2007; Edgcomb et al., 2016; La Cono et al., 2019; Fisher et al., 2021) and underground brines in potash mine (Payler et al., 2019). These systems, however, are difficult to access and study.

Human beings have developed solar salterns where water is evaporated in successive ponds of increasing salt concentration. These are particularly convenient systems to study as they provide a whole gradient of salinity in a reduced space. As salinity increases, several salts precipitate when their solubility product is exceeded: Ca^2+^ carbonate, Ca^2+^ sulfate and Na^+^ chloride. Most salterns stop at this point to collect the precipitated common salt, where salinity is about ten times that of seawater (and a_w_ is 0.75).

In some special cases, however, the process of evaporation and concentration continues up to salinities around 20 times that of seawater. In this case, K^+^ salts (sulfate and chloride) are recovered first and finally, Li^+^ chloride reaches a concentration that can be commercially harvested to follow up the process to produce Li^+^ carbonate and Li^+^ hydroxide. This is the case of the Li^+^ recovering plants in the Salar de Atacama, Northern Chile, Salar de Uyuni in Bolivia (Cubillos et al., 2019), Salar de Hombre Muerto in Argentina (Kesler et al., 2012), and other saline deposits in Qaidam Basin, China (Stober et al., 2023).

In Salar de Atacama, at 2300 masl, solar radiation is intense, the atmosphere is very dry, and the wind common. Thus, evaporation proceeds at about 1050 and 4450 mm year^-1^ in winter and summer, respectively (Marazuela et al., 2019a).. Rainfall, on the contrary, rarely exceeds 100 mm per year (ranging from <10 mm·yr^−1^ in the salt flat nucleus to >160 mm·yr^−1^ in the eastern mountains (Marazuela et al., 2019a)).

The source used for Li^+^ extraction is the aquifer under the salt crust which originated from rainfall and snowfall in the high Andes and then percolates and runs downhill towards the west. Rock weathering contributes to the enrichment of different minerals and determines the composition of the ground water (Urrutia et al., 2018). This water reappears under the flat surface of the salar in a mixing zone along the eastern margin of the core saltern (Marazuela et al., 2019b). To the west of this zone the brines increase in salt concentration due to evaporation as well as halite dissolution (Urrutia et al., 2018). Thus, the western part of the salar has a water table with brines of heterogeneous chemical composition and salinity found at different depths (Marazuela et al., 2018).

The industrial operation pumps brines from different wells at different depths to optimize the final concentration of Li^+^. The brines are transferred from one pond to another as evaporation proceeds and salinity increases. Along this gradient many properties of the brine change dramatically, temperature doubles, and the chemical composition changes as different salts precipitate.

The most important factor for life, availability of liquid water, also decreases as salinity increases by evaporation. Our purpose was to find until what point in the salinity gradient were there living beings. While NaCl-dominated hypersaline environments are habitats for a rich variety of salt-adapted microbes, there are contradictory indications about the presence of life in salt-rich environments (Hallsworth et al., 2007; Yakimov et al., 2015; La Cono et al., 2019; Payler et al., 2019; Fisher et al., 2021). Recently, bacterial and archaeal DNA was claimed to have been retrieved from such Li^+^ concentration ponds (Cubillos et al., 2019). However, these ponds have a water activity far below the accepted limit of life (Hallsworth et al., 2007; Lee et al., 2018). Here we analyze a gradient of salinity encompassing the whole industrial process and determine the bacteria and archaea present from cDNA as well as rRNA and cultures.

## MATERIALS AND METHODS

### System studied

The Salar de Atacama Basin is the largest basin (about 2900 km^2^) within the Pre-Andean Depression and is separated from the Precordillera by an abrupt relief called the El Bordo Escarpment (Bevacqua, 1992; Spiro and Chong, 1996; Carmona et al., 2000; Pueyo et al., 2017; Marazuela et al., 2020). The Salar the Atacama system has several different lakes, lagoons and wetlands in its interior. There are two saltern operations dedicated to the production of Li^+^ salts (Fig. 1). For the present study we collected samples from the salt flats operated by Sociedad Química y Minera de Chile (SQM) between 2021 and 2022. Preliminary data were collected at the Sociedad Chilena de Litio (presently Albemarle Corporation) salterns between 2000 and 2016. These salterns consist of a number of ponds for water evaporation. Brine is pumped from hundreds of wells to the surface (Marazuela et al., 2019b), and the brine transferred from one pond to another as salinity increases. Finally, Li^+^ chloride reaches a commercially valuable concentration in the last ponds. We sampled five wells (BH 1-5) and 12 ponds (P1-12) with salinities ranging from 36 to 70%. The number of P and BH follows the a_w_ order inside each group. Several wells were selected to represent the different types of chemistry in the water table: i) high Li^+^ and Mg^2+^ and high SO_4_^2-^ (well BH4, depth 30 m); ii) high Li^+^ and Mg^2+^ and low SO_4_^2-^ (BH5, 150 m); iii) high Na^+^ (BH2, 30 m); iv) high sulfate (BH1, 15 m), v) high Mg^2+^ (BH3, 67 m) (Supplementary Fig. 1). BH4 receives infiltrations from salt stockpiles associated with ponds where bischofite precipitates, and BH5 is close to evaporation ponds and salt stockpiles.

**Figure 1.**
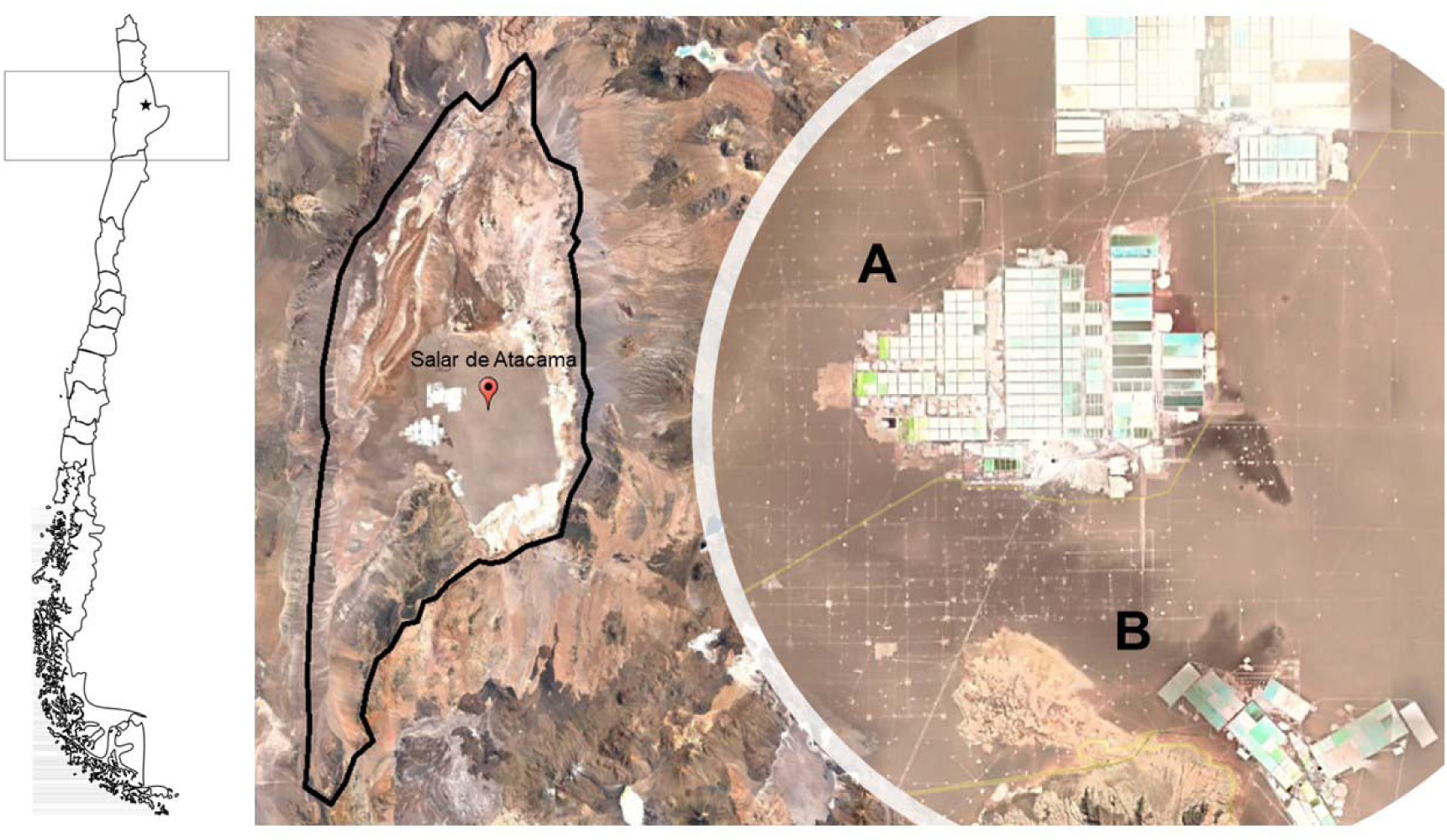
Satellite image of the Salar de Atacama. A) the SQM lithium plant sampled in 2021. B) The Albemarle plant sampled in previous years. (Google Earth Pro version 7.3.3.7786).

### Sampling and physico-chemical measurements

Ponds were sampled with stainless steel dippers about 30 m from the shores of the ponds in 20 L plastic jerrycans (Fig. 2). These were rinsed with distilled water, ethanol, distilled water again, and the brine itself, before collecting the sample and taking it to the lab. Due to the high salinities, contamination is highly unlikely. Due to COVID-19 pandemic, scientists were not allowed in the SQM compound. Samples were collected by SQM personnel and sent to the lab in Antofagasta where the following measurements were carried out: conductivity (mS cm^-1^) with a Thermo Orion 5 Star Portable PH/ORP/ISE/Cond/DO multiparameter, pH with pH-indicator strips and salinity (% NaCl) with an Atago S-28 refractometer.

**Figure 2.**
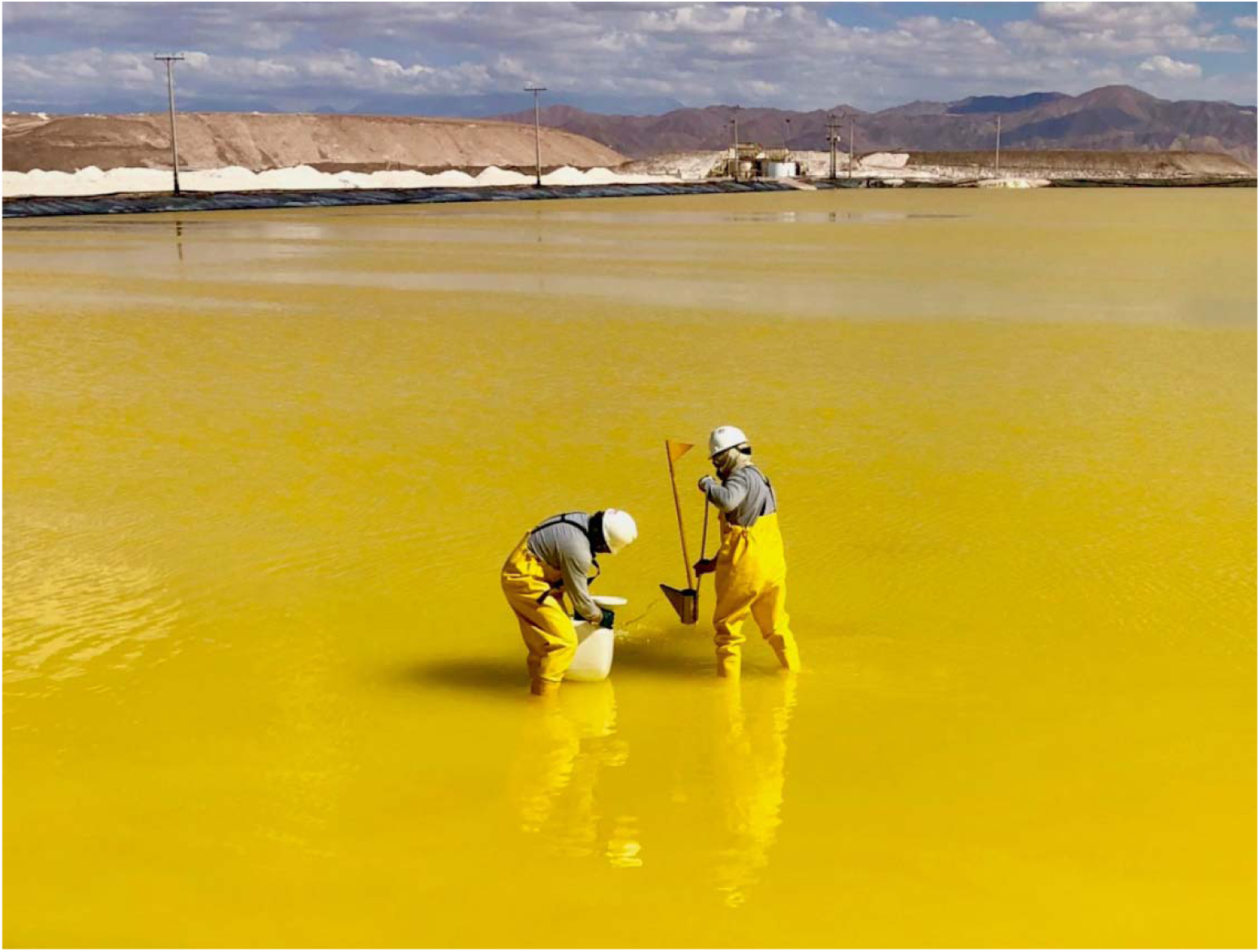
Sampling in a lithium concentrating pond.

Brines were prefiltered though 0.22 µm pore nitrocellulose filters (MF-Millipore – Merck) and the following ions analyzed. Na^+^, K^+^, Ca^2+^, Mg^2+^, Li^+^ ions were determined with a Varian SpectrAA 220 atomic absorption spectrometer. Cl^-^ was determined by the Mohr titration method, boric acid by acid-base titration and sulfate by gravimetry. Since temperature could not be measured *in situ* due to the COVID situation mentioned, we do not provide values. However, we show the characteristic increase in temperature with salinity for the 2009 samples (Supplementary Figure 6).

### Water activity

In 2021-2022 water activity was estimated by three different methods. First, SQM reported an estimate using chemical modelling with software gPROMS® (Process Systems Enterprise Ltd.). It was also estimated from the JL correlation described in Lukes (Lukes, 1993) and with the Pitzer equations implemented in the PhreeQC software (Parkhurst and Appelo, 2013).

In the samples taken in 2009 at the Sociedad Chilena de Litio salterns (presently Albemarle Corporation) we determined water activity of each pond at 20 and 30°C, using a Novasina IC II water activity machine fitted with an alcohol-resistant humidity sensor and eVALC alcohol filter (Novasina, Pfäffikon, Switzerland), as described previously (Hallsworth and Nomura, 1999). This equipment was calibrated using saturated salt solutions of known water activity (Winston and Bates, 1960). Values were determined three times using replicate solutions made up on separate occasions. The variation of replicate values was within ±0.002 a_w_. These results are shown in Supplementary Figure 6).

### Concentration of biomass

Due to the very high viscosities of the brines, only very small volumes could be filtered practically. These volumes were insufficient for satisfactory cell counts and DNA extractions. Thus, we developed a method to dilute the brine without affecting the potential microorganisms. Five (for DNA) to ten (for RNA) liters of brine were fixed with ethanol 1:1 and 300 to 500 mL aliquots placed in dialysis bags. The bags were immersed in 5 L containers with demineralized or distilled water and allowed to equilibrate until a conductivity of < 25 mS cm^-1^ was obtained in the bathing water (6 to 8 hours depending on the initial conductivity of the brine). The whole process was carried out in a hood (Supplementary Fig. 7). The dialysis product was filtered through a 0.22 µm pore nitrocellulose membrane (Merck) and the filter with the retained cells was stored in a lysis buffer (50 mM Tris–HCl pH 8.3, 40 mM EDTA and 0.75 M sucrose) at −20°C for subsequent DNA extraction, and in a commercial solution for RNA stabilization and store RNALater (Invitrogen) at −80°C for the RNA extraction.

Several controls were carried out to make sure the procedure did not cause either contamination or loss of biomass (see Supplementary Materials section C for details). First, the reactants for the DNA extraction (buffers, enzymes, detergents, etc.) were extracted. All the reagent batches failed to show amplification by conventional PCR (Supplementary section C-1, Fig. 8). We also carried out qPCR and sequenced the V4 by Illumina, for assessing the number of copies per mL which were always below the detection level of the technique (100 copies per mL Supplementary Table 2), and for the identification of the present taxa, respectively.

Next, the same procedure was applied to the demineralized and distilled water used for dialysis (Supplementary materials: section C-2, Fig. 9 and Table 4). In this case, values around 10^4^ copies mL^-1^ were obtained in all batches for bacteria. For archaea, only one batch provided a similar number of copies but all the other ones were below detection levels. This water does not enter in contact with the samples, since it remains outside the dialysis bags. However, we sequenced several batches of this water and discarded all the sequences that were identical in the water and the samples (as we did with reagent controls). These amounted to 2.1% of all the sequences. The number of reads for each well and pond and the controls received from the 16S sequencing service are included in Supplementary materials: section E and Fig. 14.

Third, we carried out a positive control. A known number of cells from enriched consortia suspended in a 100 g L^-1^ KCl solution were subjected to the same fixation, dialysis and filtration procedure as the samples. This was to show both that the protocol was not detrimental to the microorganisms and that the fixation with ethanol was effective (Supplementary section C-3). The number of cells recovered after dialysis was always close to 100% (Supplementary Table 6). And the taxonomic composition was virtually identical before and after the procedure (Supplementary Fig. 10). Fourth, the technique of total RNA extraction and cDNA synthesis was tested using an enriched consortium suspended in a KCl solution and in brines (section C-4, Supplementary Fig. 11 and Table 7).

### Cell abundance

When possible (lower salinity samples), a 100 mL volume of fixed and dialyzed brine was vacuum filtered using a 0.22 µm pore membrane for epifluorescence microscopy. A fraction of the membrane was stained with DAPI (4,6-diamidino-2-phenylindole prepared at 2 mg ml^-1^) for 15 minutes in a dark chamber. Cells were visualized by epifluorescence microscopy using an OLYMPUS IX-81 microscope, equipped with a Fluoview FV-1000 spectral Confocal module.

Cell counts were also carried out by qPCR (see DNA extraction protocol below). Quantification of 16S rRNA amplicons was done with the SYBR-Fluorophore Green Kit (Bioline) and a real-time PCR machine Rotor-Gene Q (Qiagen) and probes for total bacterial and total archaea described previously (Remonsellez et al., 2009).

### Nucleic acid extraction

The protocol carried out included: two steps of freeze and thaw at −80°C and 65°C, cell disruption by sonication for 10 min at 0.5 cycle and 30% amplitude, a cell lysis with 1 mg ml^-1^ Lysozyme at 37°C for 1 hour and a second cell lysis with 0.5 mg ml^-1^ of Proteinase K and 1% SDS at 55°C for 45 minutes, continued with the commercial kit HP PCR Template Preparation Kit by ROCHE. The quantity (ng µL^-1^) and quality of the DNA (A260/A280 and A260/A230 ratios) were measured with a NANODROP-1000 spectrophotometer. RNA extraction was performed using TRIzol Reagent – Ambion, DNA was digested using the RQ1 restriction enzyme (Promega) and cDNA was synthesized using the Sensiscript Reverse Transcription Kit and Random primers – Qiagen according to the protocol established by the manufacturer.

### Sequencing

Sequencing of samples and blanks (DNA extraction reagents and water for dialysis process) were carried out according to the following procedure. The hypervariable V4 region of the 16S rRNA gene was amplified from each sample using for each sample uniquely barcoded reverse primers (806R: GTGYCAGCMGCCGCGGTAA) and a common forward primer (515F: GGACTACNVGGGTWTCTAAT). Both the reverse and the forward primers were extended with the sequencing primer pads, linkers, and Illumina adapters (Caporaso et al., 2012). The PCR was performed using MyFi™ Mix (Bioline Meridian, Cat. No. BIO-25050) on LightCycler 96 (Roche) in the final volume of 40 μL. Amplicons were quantified using the Quant-It PicoGreen dsDNA Assay kit (ThermoFisher Scientific, Cat. No. P7589), according to the manufacturer’s protocol. Equal amounts of amplified DNA (120 ng) from each sample were pooled into a sequencing library followed by removing DNA fragments smaller than 120 bp (unused primers and dimer primers) with UltraClean PCR Clean-Up Kit (MoBio, Cat. No. 12500). The final amplicon concentration was quantified by qPCR with KAPA Library Quantification Kit for Illumina Platforms (KAPA Biosystems, Cat. No. KK4854) in the presence of the set of six DNA standards (KAPA Biosystems, Cat. No. KK4905). Subsequently, the library was diluted to a concentration of 4 nM and denatured with 0.1 N NaOH. The library was sequenced at the Microbiome Core at the Steele Children’s Research Center, University of Arizona, using the MiSeq platform (Illumina) and custom primers (Caporaso et al., 2012). Due to the peculiarities of our samples in terms of relatively low diversity and biased GC content, a PhiX Sequencing Control V3 was used as recommended by Illumina (Illumina, Cat. No. FC-110-3001). The raw sequencing data were demultiplexed and barcodes trimmed using idemp script.

### Taxonomic and phylogenetic analysis

Demultiplexed fastq files were received and subjected to primer removal, filtered by sequence quality (for keeping Ph quality over 30), denoised, merged, and chimaera removal using the DADA2 pipeline (Callahan et al., 2016). All filtered-merged sequences were assigned to amplicon sequence variants (ASV) by the DADA2 pipeline. The representative reads were mapped to the SILVA database (release 138) for taxonomy and abundance data (Quast et al., 2013). Then we used the Phyloseq (version 1.42.0) pipeline to (a) remove the ASVs found in controls (Supplementary Tables 3 and 5; 3.3% of all the sequences were removed for this reason): (b) eliminate taxa with one read, (c) remove taxa with less than 0.005% mean relative abundance across all read counts, (d) remove ASVs that were not observed more than twice in at least 10% of the samples; e) eliminate samples having less than 1000 reads and f) all samples were rarefied to 1756 reads per sample (see ASVs removed in Supplementary Table 9).

### Statistical analysis of microbial diversity

The abundance of Amplicon Sequences Variants (ASV) in different samples was analyzed using Primer-7 (Primer-E) software (Clarke and Gorley, 2015). We used the fourth root transformation to homogenize the amounts of ASVs and thus reduce the dominance effect. A similarity matrix (resemblance) was constructed using the Bray-Curtis method (Bray and Curtis, 1957) included in Primer V7 software (Clarke and Gorley, 2015). We used non-metric multidimensional scaling (NMDS), included in Primer-7, to build a restricted arrangement of ASVs according to a taxonomic level (phylum, order, family and genus). ASV numbers, Shannon (H’) and Simpson diversity indexes were calculated using Primer-7 (Clarke and Gorley, 2015).

Nucleotide sequence accession numbers at the DNA Data Bank of Japan (DDBJ) repository are the following: (Supplementary Table 10).

### Culture studies

For the lower salinity ponds, between 5 and 100 mL of brine was filtered through 0.22 µm pore size filters thus obtaining a natural inoculum in the filter and sterile brine. For the higher salinity ponds, filtration was not practical. In this case brines were centrifuged at 10285 x g for 20 minutes. In this case the supernatant provided the sterile brine and the pellet the natural inoculum (Supplementary Fig. 12). In all cases, the sterile brines were supplemented per liter with LB medium (10X, 100 mL), trace element solution (1 mL) and vitamins (10 mL) which means a dilution to 11.1% in the salt content and the resulting increase in a_w_. See Supplementary section D for composition of the different solutions. Each supplemented brine was divided in two flasks, one receiving an inoculum from the same pond and the other left without inoculum as a control. The flasks were incubated aerobically with stirring at 40 °C for ten days when the cultures were checked for turbidity.

Additional cultures were incubated in anaerobiosis without stirring. In this case, the sterile brines were also amended with lactate and FeSO_4_ in addition to LB medium, vitamins and trace metals (Supplementary Section D). The brines were bubbled with nitrogen gas for 4 minutes. Incubation was at 40 °C for 10 days when cultures were checked for the appearance of iron sulfide precipitates indicative of anaerobic sulfate reduction.

Both aerobic and anaerobic cultures were re-inoculated in fresh brine every 15 to 20 days and allowed to grow again. Eventually, cultures were gradually transferred to a synthetic brine with a much higher a_w_ to speed and facilitate culturing. The composition of this synthetic brine can be seen in the same supplementary section D (Supplementary Table 8).

## RESULTS

### Physico-chemical parameters

The essential parameter, water activity, is not trivial to determine. We used different models to estimate it. The correlation between Lukes (Lukes, 1993) and Pitzer in the PhreeQC (Parkhurst and Appelo, 1999) models estimates was very good (r^2^ = 0.964, see Supplementary Figure 4). The modelled estimated values were also correlated (r^2^ = 0.969) with Li concentrations (Fig. 3). In addition, water activity was determined experimentally with the method of Hallsworth and Nomura (Hallsworth and Nomura, 1999) in 2009 with samples obtained from the Li^+^ extraction process in Sociedad Chilena del Litio (Demergasso et al., 2007). As can be seen (Supplementary Fig. 6), the a_w_ values obtained experimentally correlated (r^2^ = 0.895) with the determined Li^+^ concentration.

**Figure 3.**
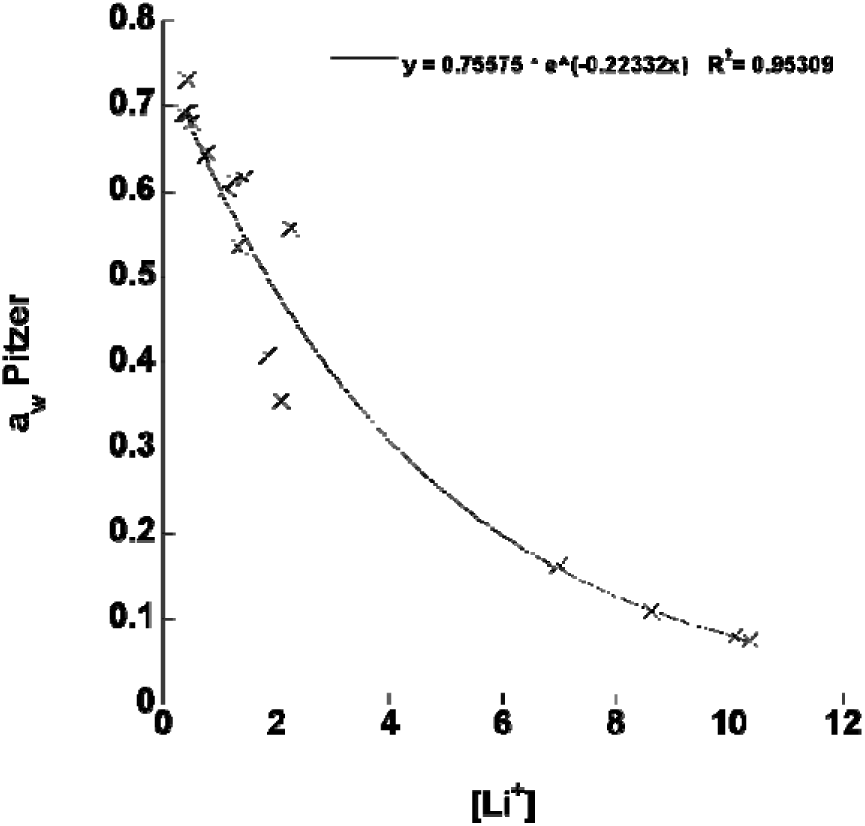
Exponential correlation between a_w_ estimated by Pitzer equation versus [Li^+^] in the studied system. [Li+] means mol/m^3^.

Despite the fact that different techniques and different estimating software were used and that samples came from different salterns in different years, it can be seen that the relationship between water activity and Li^+^ concentration was remarkably similar (Fig. 3, and Supplementary Fig. 6, respectively).

As salinity increased along the gradient of ponds, water activity decreased. pH also decreased, although measurements at these high salinities are problematic. However, both electrode measurements and litmus paper determination coincided fairly well.

Twelve ponds were selected along the gradient. Cation composition in brine samples was dominated by Na^+^, K^+^, Mg^2+^ and Li^+^, and anions were dominated by Cl^-^, SO_4_^2-^ and B(OH)_3_ was also present. The highest concentrations of Li^+^ coincided with the lowest concentrations of Na^+^ (K^+^ showed a profile similar to that of Na^+^) and medium levels of Mg^2+^ (Fig. 4; actual data in Supplementary Table 1). The ternary plots depict the evolution of well brines due to evaporation and precipitation (halite, carnallite, and bischofite) processes in the ponds that mostly affected [Na^+^], [Mg^2+^], [Li^+^] as well as [Cl^-^], [SO_4_^2-^] and [B(OH)_3_] (Fig. 4). Cl^-^ was always the predominant anion and smaller changes in SO_4_^2-^ and B(OH)_3_ occurred in the process (Fig. 4).

**Figure 4.**
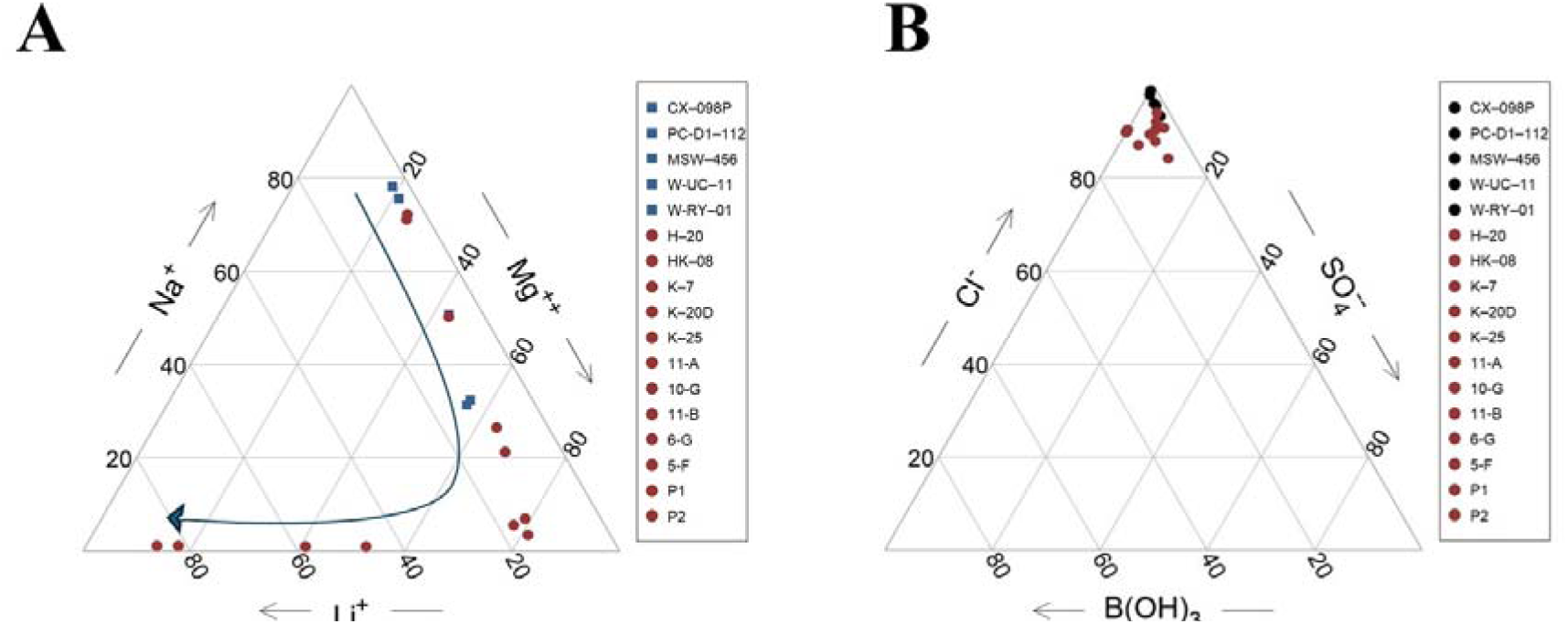
Ternary diagrams for brines collected from SQM concentration process in Salar de Atacama. Wells (blue) and evaporation ponds (red). The blue line represents the evolution of brines lacking Na^+^, following precipitation of NaCl, into Mg^2+^ and Li^+^ intermediate brines, and then to a Li^+^ rich brine following evaporation and precipitation of carnallite and bischofite.

The different ions showed very interesting changes in concentrations as salinity increased. Na^+^ and K^+^ decreased and practically disappeared (Fig. 5A). On the other hand, Li^+^ and different species of boron increased along the gradient (Fig. 5B). Mg^2+^ and Ca^2+^ showed complementary changes with maxima for Mg^2+^ at intermediate salinities and minima for Ca^2+^ (Fig. 5D). Finally, chloride increased during the whole process and sulfate had a maximum at intermediate a_w_ and then practically disappeared (Fig. 5C).

**Figure 5.**
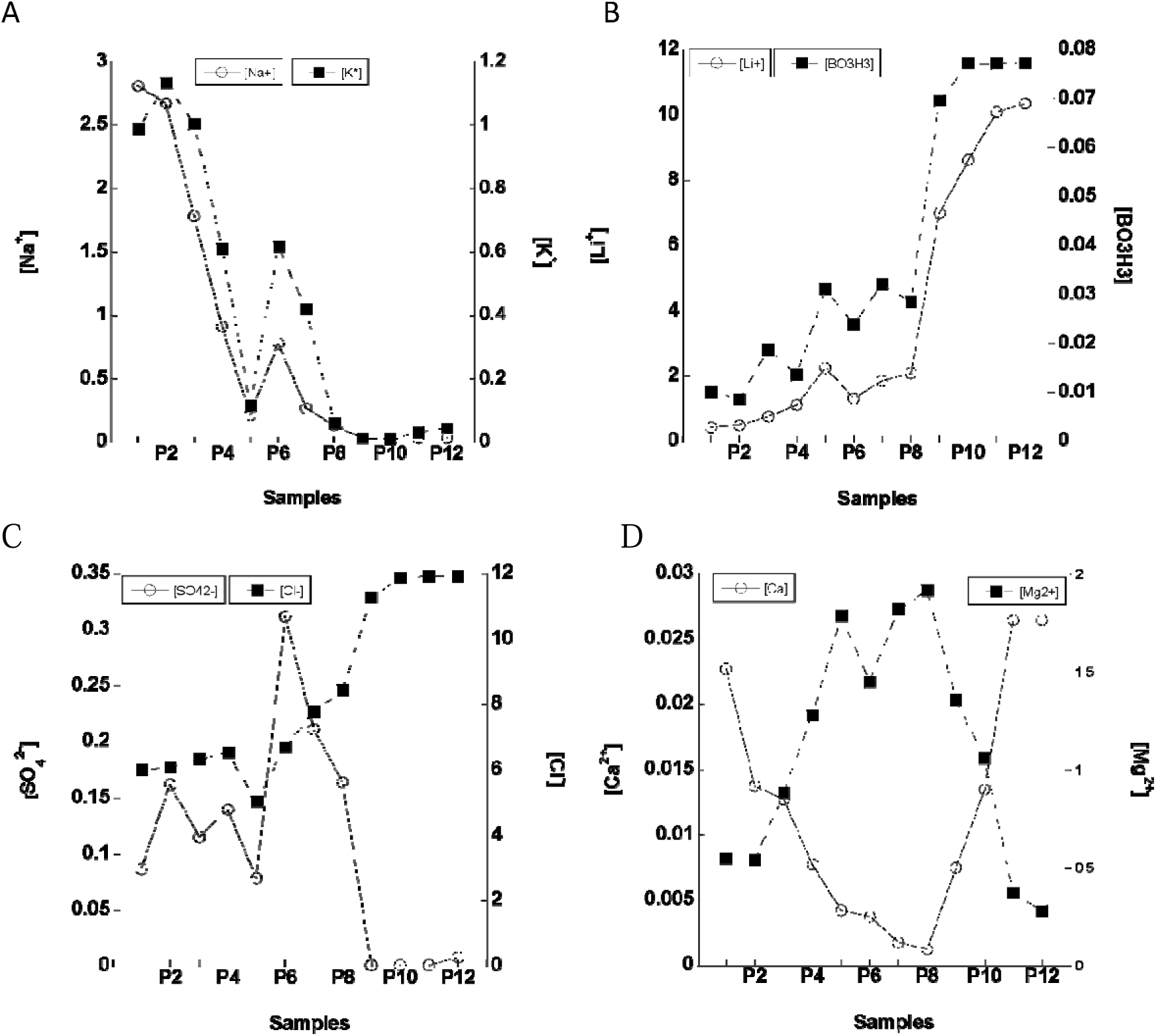
Molar ion concentration along the a_w_ gradient. The x axis indicates the pond number ordered by a_w_. [Li+] and others mean mol/m^3^.

A principal component analysis was done with the physicochemical characterization of the well and pond brines to complement with the dynamic of chemical components not included in Figs. 4 and 5 and physicochemical parameters (pH, EC and a_w_, among others, Supplementary Figure 2). Interestingly, the analyses showed the difference between the ORP of wells (< −100 mV) and ponds (80-250 mV) from underground and surface environments, respectively (Supplementary Figure 3).

### Abundance of microorganisms

qPCR determinations from DNA retrieved bacteria from almost all the gradient while archaea appeared only in the lowest salinity wells and ponds (Fig. 6A). In Fig. 6B the copies per mL retrieved from bacterial DNA and RNA (cDNA) are shown together. Biological activity, as shown by amplification of cDNA, was present consistently down to an a_w_ of 0.731. For higher salinities activity could be seen only in four out of 12 ponds despite the presence of DNA. The pond with the highest salinity where we found cDNA was P9 with an a_w_ of 0.268.

**Figure 6.**
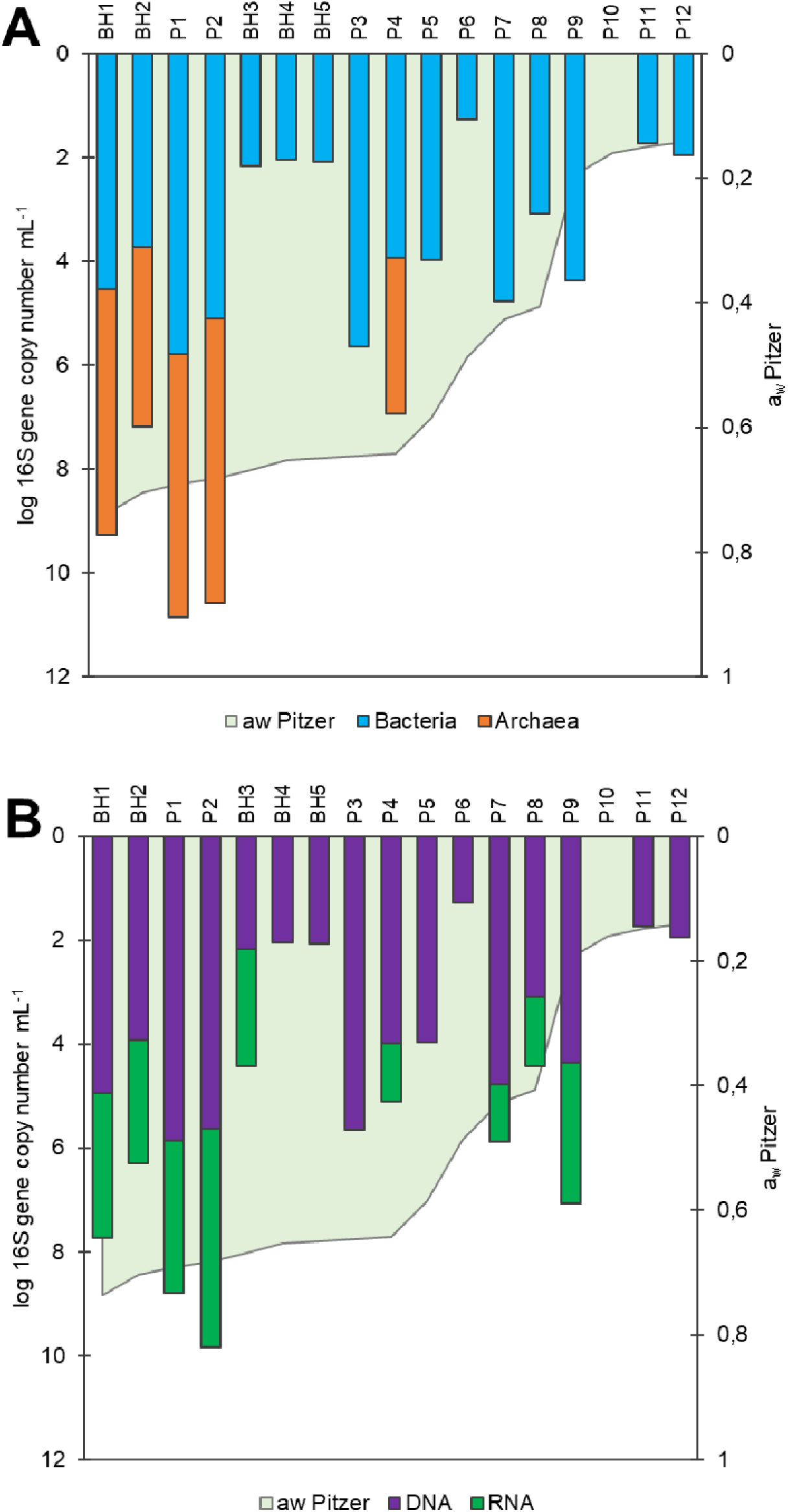
**A.** copies per mL from retrieved bacterial (blue) and archaeal DNA (orange) by qPCR. **B.** Copies per mL retrieved from bacterial DNA (purple) and cDNA (green). No archaeal sequences were obtained from cDNA. The green shaded area shows water activity estimated by Pitzer equation.

### Microbial diversity

The number of ASVs and diversity and evenness indices were higher in the borehole samples than in most ponds (see Fig. 7 and Supplementary Section F) (e.g., general Simpson index including Bacteria and Archaea, and Bacterial Shannon index p<0.05, Supplementary Table 11). In addition, a decrease in the bacterial richness was observed when Li^+^ increased and a_w_ decreased, and on the contrary, an increase was evidenced when Mg^2+^ increased (Supplementary Figure 15). Archaea were only present at low [Li^+^] but were distributed over the [Mg^2+^] gradient (Fig. 7).

**Figure 7.**
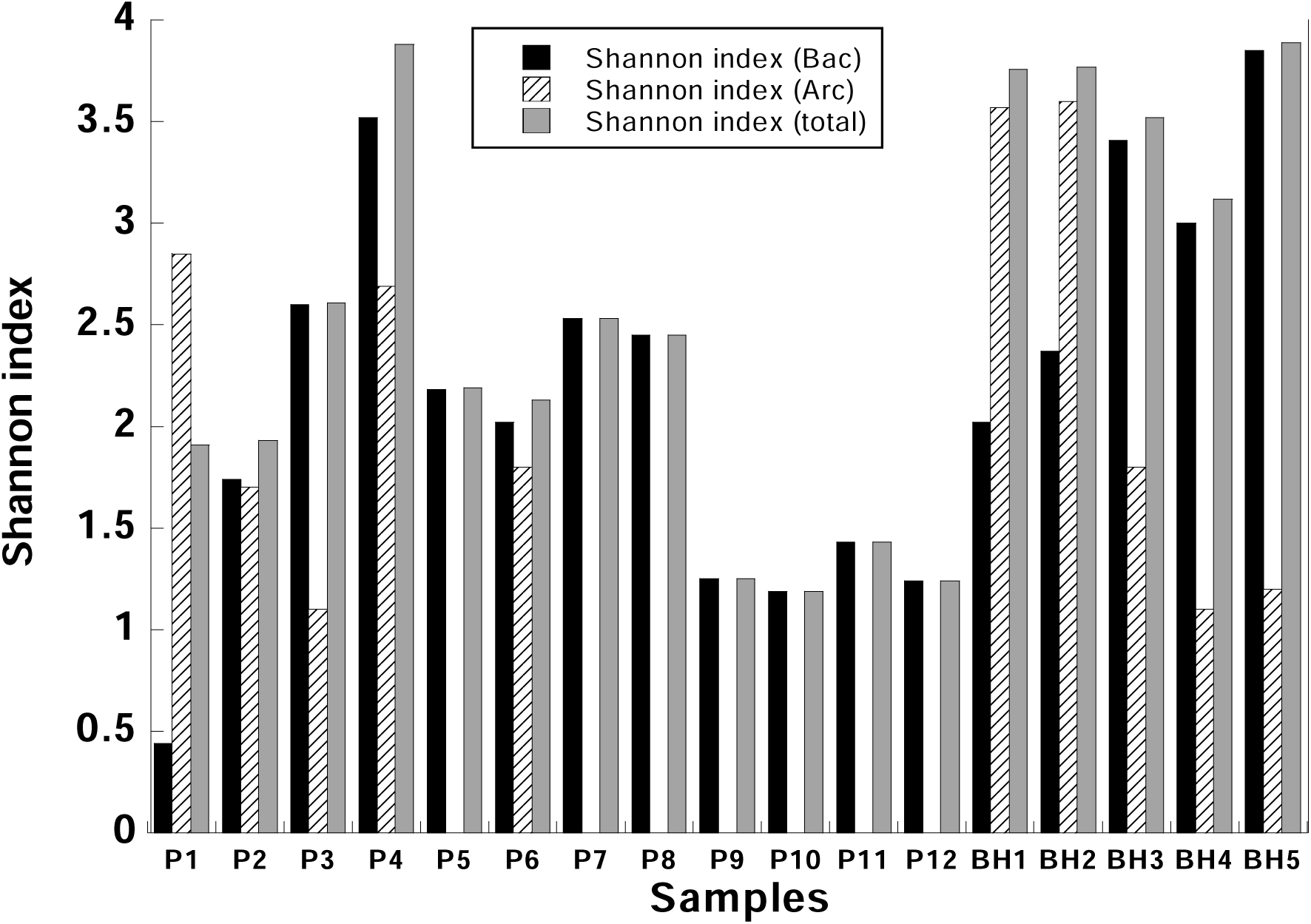
Shannon index in ponds (P1-12) and wells (BH1-5) of SQM Li production process ordered by a_w_. inside each kind of samples. The species richness and evenness indices are in Supplementary Table 11.

### Taxonomic composition

Figure 8 provides an overview of the changes in taxonomy along the gradient. Archaea were only present at the beginning and disappeared or became extremely rare later on. Bacteria, on the other hand, were present throughout. Most classes decreased and disappeared as the maximal concentration of Mg^2+^ was reached and did not reappear further along the gradient. Some (*Bacilli* and *Bacteroidia*) maintained a very low presence throughout the gradient. Finally, *Alphaproteobacteria* and *Gammaproteobacteria* increased in relative abundance until an a_w_ of approximately 0.6 and then maintained their presence to the end of the gradient. The more common bacteria were *Gammaproteobacteria* and *Alphaproteobacteria* (Fig. 8 and Supplementary Fig. 16 A and B for family and genus, respectively). Most of the former were previously classified as Betaproteobacteria: *Burkholderiales* (Fig. 8B), while the *Alphaproteobacteria* were mostly *Caulobacterales* and *Sphingomonadales*. There were also some *Bacillales* in the most saline ponds. This distribution can be compared to physicochemical parameters in the NMDS diagram (Fig. 9). There seemed to be three main types of communities: those of wells 1 and 2 rich in *Halobacteria* class, those from more diverse wells 3 to 5 with a variety of groups and those of the ponds, where *Burkholderiales* and *Caulobacterales* predominate (Fig. 10). The presence of *Burkholderiales* at the lowest a_w_ ponds was intriguing. Figure 11 shows the most abundant genera in this order. At the lowest a_w_ the main genera were *Ralstonia*, *Aquabacterium, Hydrogenophaga*, and *Burkholderia-Caballeronia-Paraburkholderia*.

**Figure 8.**
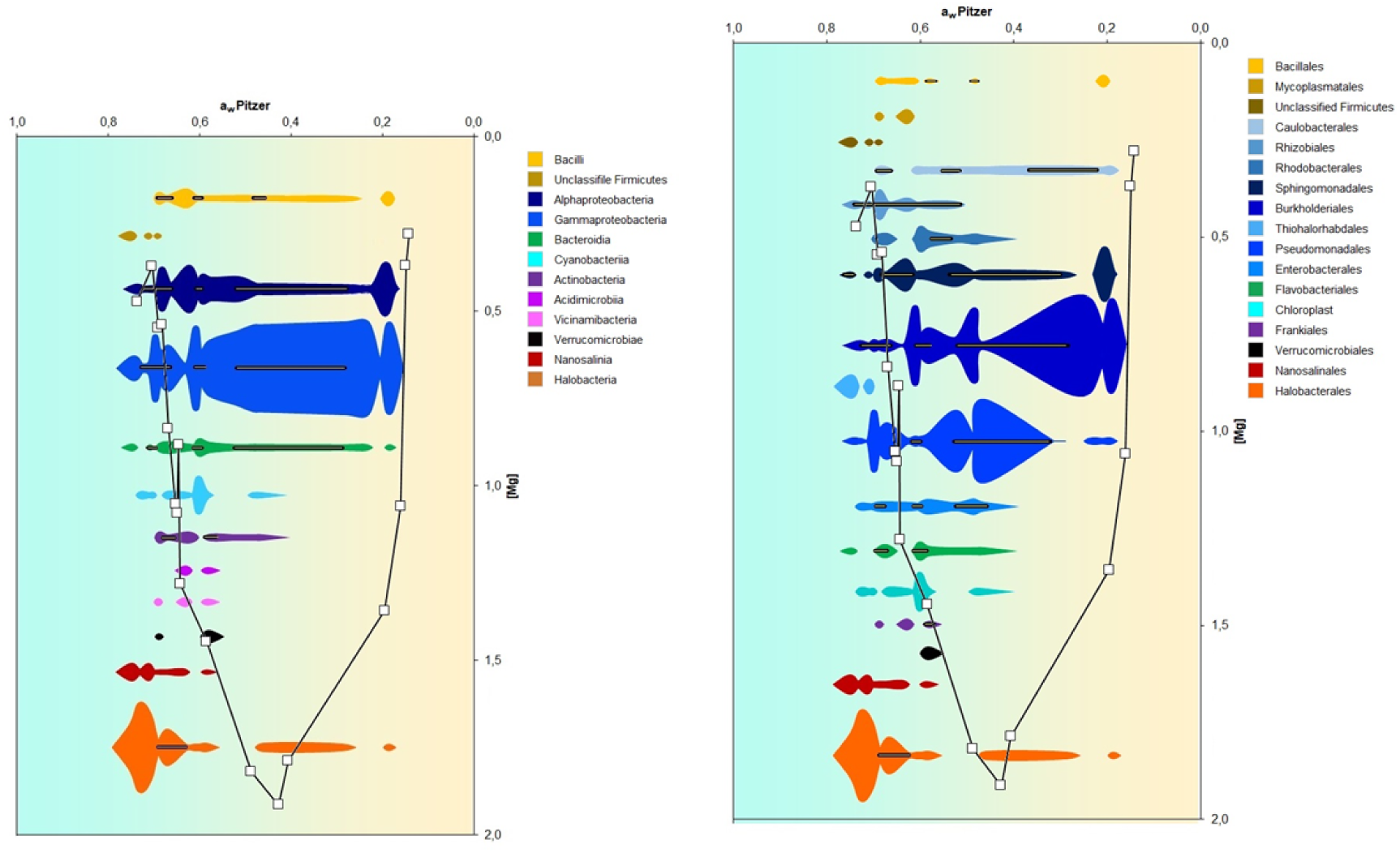
Abundance profiles of Bacteria and Archaea classes (A) and orders (B) versus a_w_ (background color gradient), which coincides with the steps of the industrial evaporation process, and Mg^2+^ concentration (shown by the continuous line and empty square symbols). The horizontal lines within colored violins indicate samples where sequences were obtained from cDNA. [Mg^++^] means mol/m^3^.

**Figure 9.**
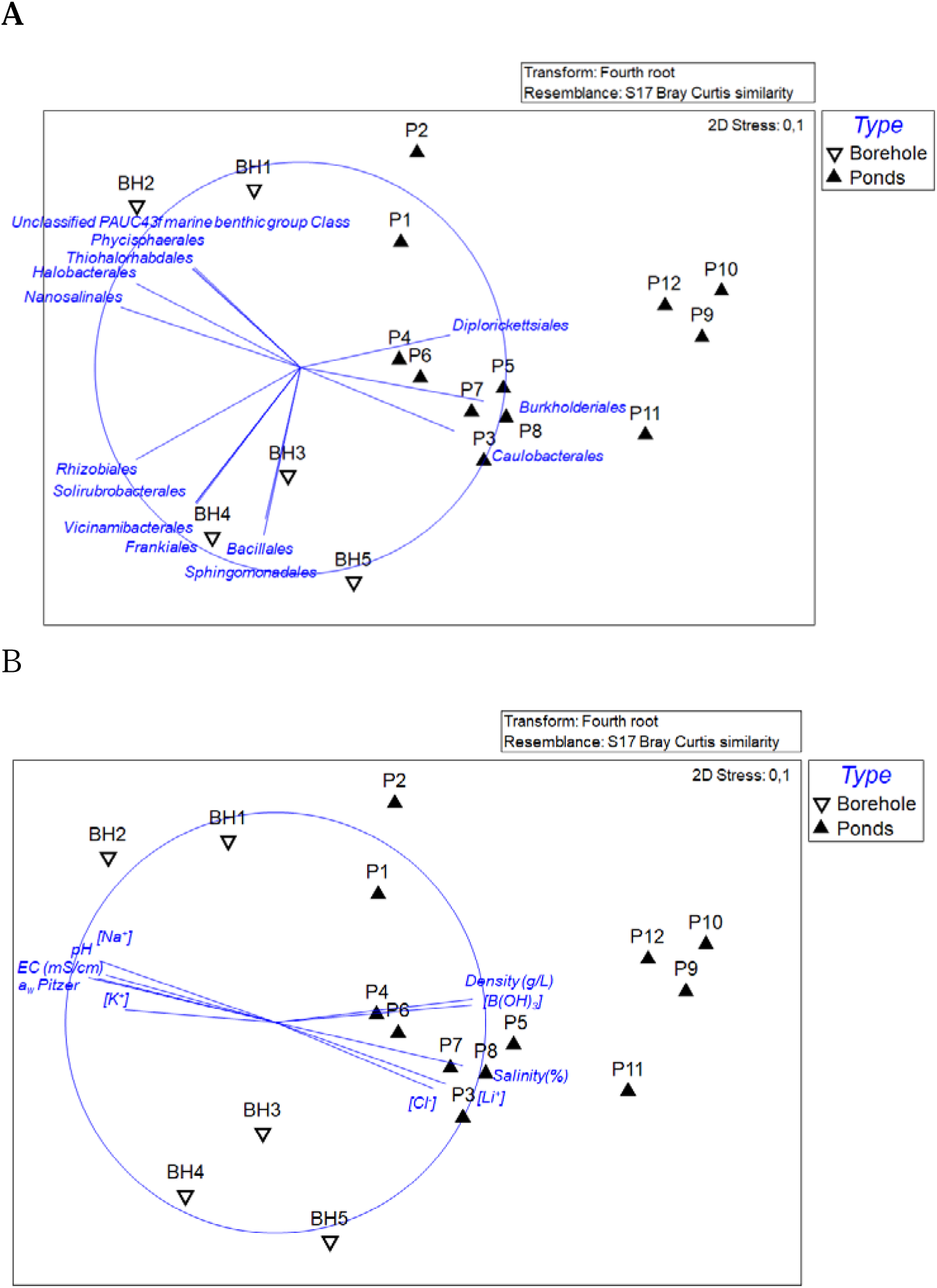
Non-metric multidimensional scaling (NMDS) of Bray-Curtis similarities for fourth root transformed order abundance data with vectors of orders whose relative abundance has a Spearman correlation >0.7 with ordination axes (A), and vectors of physichochemical parameters which have Spearman correlation >0.7 (B). [Li+] and others means mol/m^3^.

**Figure 10.**
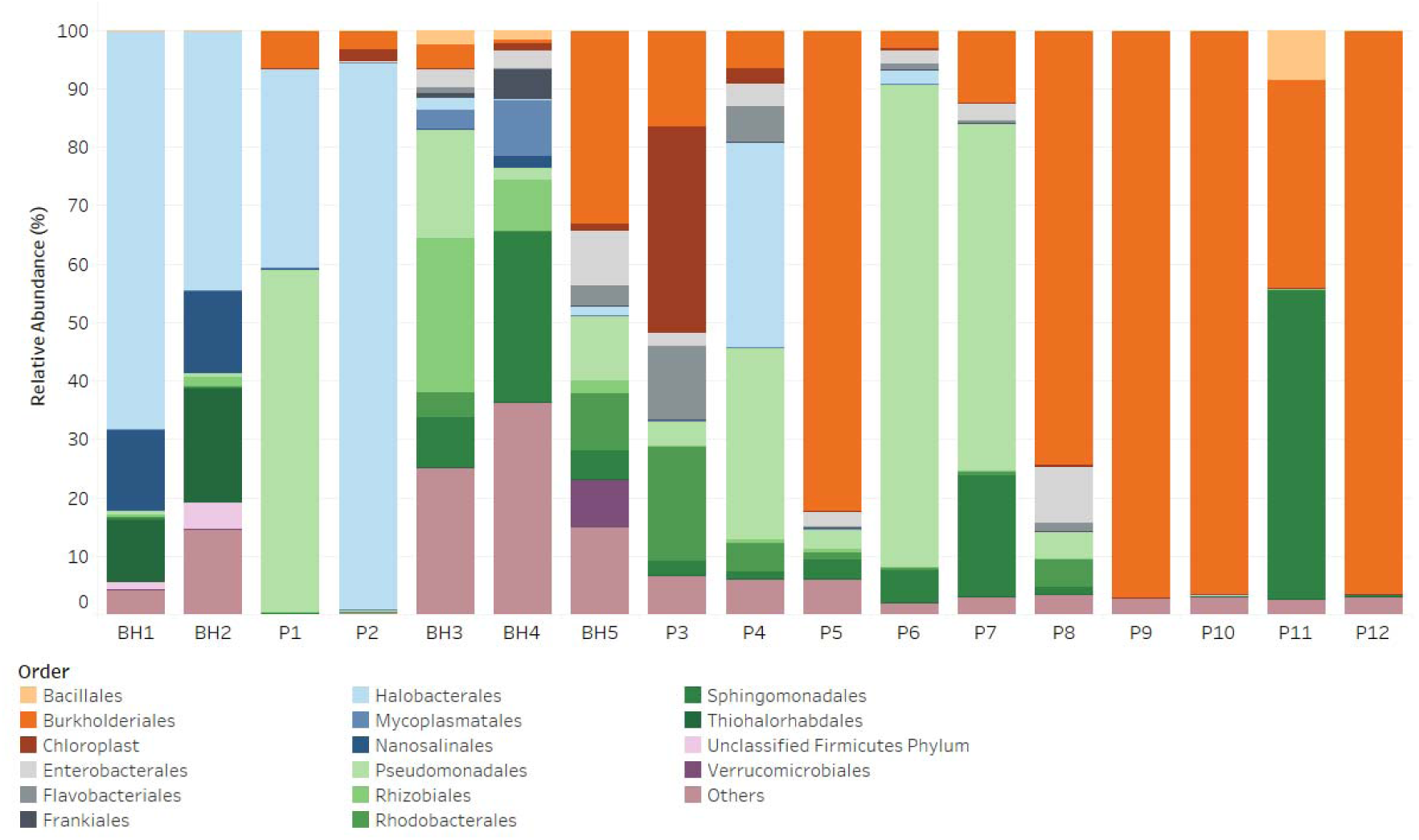
Relative abundance of Bacteria and Archaea orders in wells (BH) and evaporation ponds (P). The wells and ponds are ordered by a_w_ estimated by Pitzer equation.

**Figure 11.**
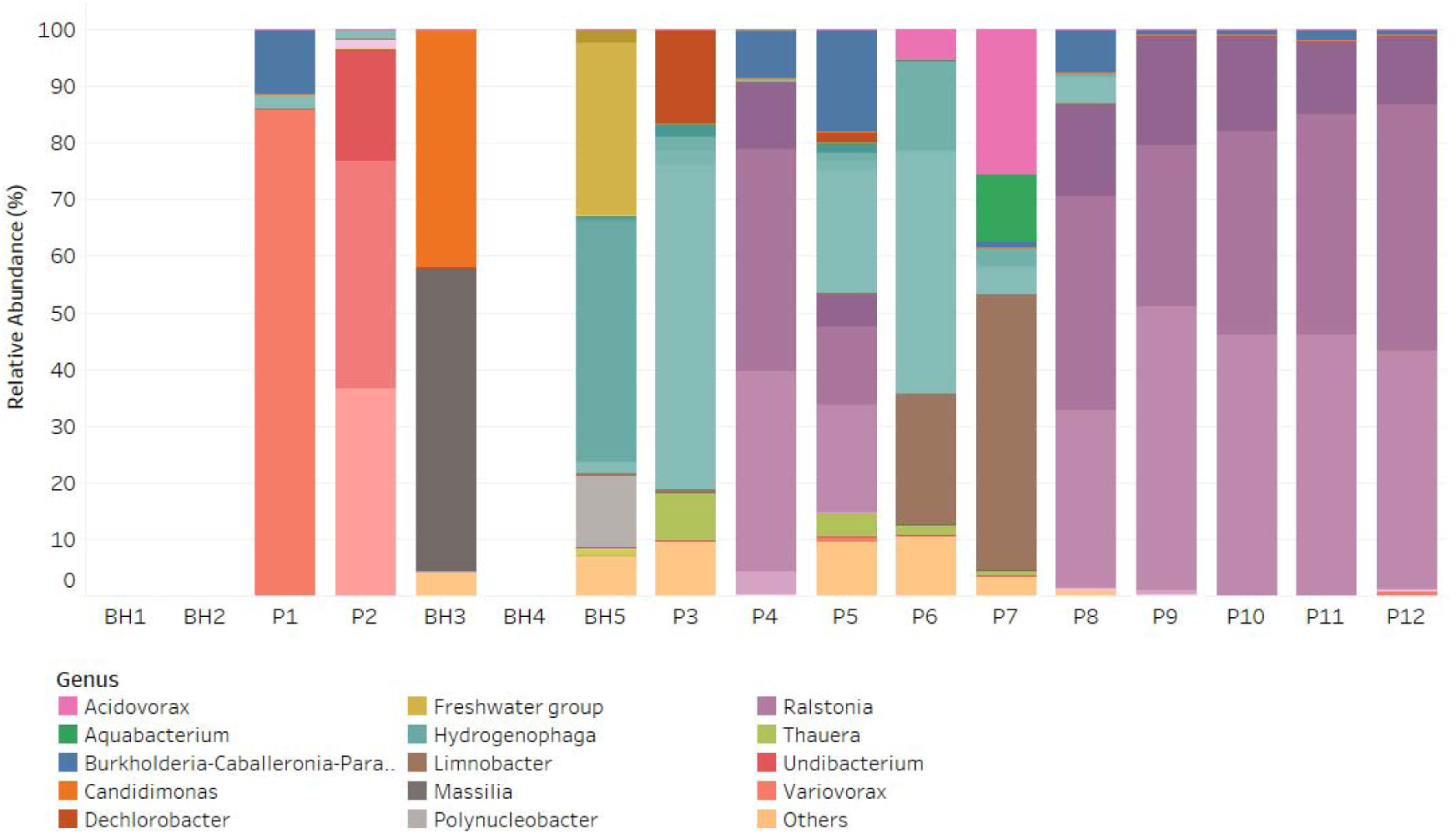
Relative abundance of genera from the *Burkholderiales* order in wells (BH) and evaporation ponds (P). The wells and ponds are ordered by a_w_. Different ASVs are plotted in different shades of the genus color.

The wells had brines with rather different composition and, they had very different communities among themselves and from the first ponds (Fig. 10 for orders and Suppl Fig. 16 for families and genera). BH1 and BH2 were very rich in Archaea (Fig. 8), more specifically *Halobacteriales* from *Nanosalinaceae*, *Halomicrobiaceae, Haloferacaceae*, and *Halobacteriaceae* families (Supplementary Fig. 16). The Halobacteraceae were also abundant in ponds P1, P2, and P4, but they were rare in the remaining wells and ponds. These two wells had the largest Na^+^ chloride concentrations of all the wells (Supplementary Fig. 1) and, thus, it makes sense that the community was dominated by *Halobacteria* phylum.

BH1 and BH2 were also enriched in *Thiohalorhabdales* (Fig. 10, a group of basal *Gammaproteobacteria* that are sulfur-oxidizing and halophilic. Unlike the chemo organoheterotrophic *Halobacteria* class, these bacteria are chemolithoautotrophic (Sorokin and Merkel, 2023). But this group disappeared in the ponds.

Each one of the other three wells had its own particular chemistry. The three were enriched in Mg^2+^ and Li^+^, but the proportions of ions were different. Their community composition was also different: *Rhizobiales* and *Pseudomonadales* were dominant in BH3, *Sphingomonadales* in BH4 and *Rhodobacterales*, *Burkholderiales* and *Pseudomonadales* in BH5 (Fig. 10).

The NMDS graphs confirm that BH1 and BH2 were closer to each other (Fig. 9B) and coincidental with Na^+^ and K^+^ concentrations, while the other three wells showed a different position in the diagram, midway between the different physicochemical parameters (Fig. 9B). Clearly, different wells bring brines with different chemical and community compositions.

The pond communities changed substantially with salinity and density, as well as Li^+^, chloride, and borate concentrations (Fig. 9B). As we already mentioned, P1 and P2 were closer to the BH1 and BH2 wells. But the remaining ponds spread out from the center to the right of the NMDS diagram, showing progressive dissimilarity in their taxonomic composition (Fig. 9A). *Pseudomonadales* were dominant in P4, P6 and P7, while *Burkholderiales* were dominant in P5 and P8 to P12 (Fig. 10). In the four ponds with the lowest a_w_ only *Bacillales* and *Sphingomonadales* were present in one of them aside from *Burkholderiales*.

### Enrichment cultures

The procedure to obtain enrichment cultures is schematized in Supplementary Fig. 12. Samples were either filtered (when a_w_ was relatively high) or centrifuged (for the low a_w_ ponds). This provided, on the one hand, sterile brine that was used as culture medium and, on the other hand, concentrated cells that were used as inoculum. Some were incubated in aerobiosis with shaking and some in anaerobiosis without shaking (see methods). After 10 days, turbidity was observed in all experimental flasks except for those with brines from P1 and P2 (Supplementary Fig. 13). None of the uninoculated controls showed any turbidity. The presence of cells was checked by epifluorescence microscopy. Despite the difficulties of the autofluorescence of the samples that precluded quantitative counts, cells were seen in all experimental flasks.

Next, cultures were transferred to fresh medium and allowed to grow again. This was repeated a third time. Turbidity was observed again in all experimental flasks but in none of the uninoculated cultures.

Having shown that growth did actually take place in the original brines (OB), we designed an artificial brine (AB) to accelerate growth and simplify the experimental manipulations (see supplementary method for composition). Four enrichment cultures (from the aerobic enrichments) produced growth under these new conditions: those derived from P5, P6, P7, and P8. Cultures P6, P8, and P7 were pooled in a new one called “consortium” to further reduce the number of cultures and flasks to process. Currently, the P6 and Consortium enrichments are kept in artificial brine. In the case of the anaerobic cultures, turbidity and the presence of iron sulfide precipitate were observed in a third of the brines, and growth occurred until the third transfer in the original enriched brine. It was not possible to recover these cultures in modified Bataglia medium (Supplementary Section D).

A large variety of taxa grew in these enriched cultures (Fig. 12). Predominant orders enriched were *Burkholderiales*, *Pseudomonadales*, *Enterobacterales*, *Staphylococcales*, and *Sphingomonadales* in aerobic cultures, and *Desulfovibrionales* and *Paenibacillales* in anaerobic cultures.

**Figure 12.**
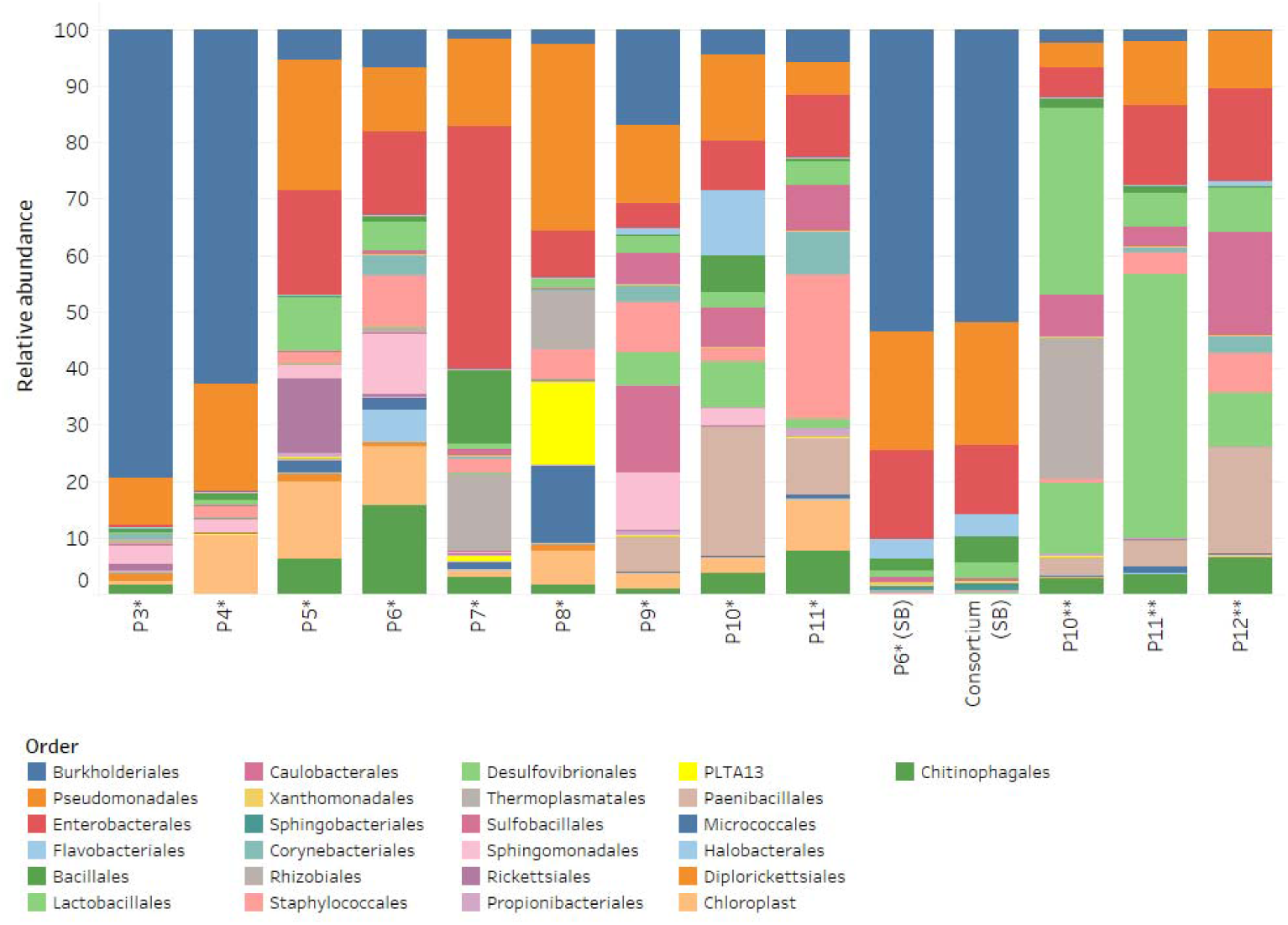
Relative abundance of the different taxa in the enriched microbial communities at the order level. “*” Aerobic culture, “**” Anaerobic culture, “SB” Synthetic brine.

Finally, we compared the taxa retrieved from the enrichment cultures with those found in cDNA in the ponds. More than 60% of the microbial orders (62%) and families (63%) identified in cultures were also present in cDNA from the ponds where the cultures were obtained (data not shown). That percentage decreased at the genus taxonomical level (41%). The best match between cDNA and cultures was observed for *Burkholderiales*, *Pseudomonadales* and *Staphylococcales* orders.

## DISCUSSION

### Chemical composition along the gradient

The LiCl_2_ concentration process (LiCP from now on) entails an extremely unique series of chemical conditions. As water evaporates ion concentrations increases and different salts reach their solubility product at different ponds. Thus, in addition to decreasing values of a_w_, the consequence is a different ionic composition in each pond. Ions in high concentrations have complex effects on molecules. This so-called specific ion effects are still a matter of debate and the mechanisms are not well understood (Mazzini and Craig 2016, (Gregory et al., 2022). Franz Hofmeister, in the nineteenth century, was the first to order ions in a series of increasing effects on the salting out of proteins. This has been interpreted in terms of some ions being chaotropic (they disrupt proteins and thus keep them in solution) and others kosmotropic (favoring precipitation of proteins). This series is somewhat parallel to the lyotropic series, related to the heat of hydration of the ions. The latter departs from the Hofmeister series in some details (Mazzini and Craig, 2016). In the lyotropic series cations are ordered from more kosmotropic to more chaotropic as Li, Na, K, Rb, Cs, Ca, Sr, Ba and anions as I, NO_3_, Br, Cl, F, PO_4_, SO_4_ (Mazzini and Craig, 2016; Mazzini and Craig, 2018; Gregory et al., 2022). In some papers Mg^2+^ is considered to be as chaotropic as Ca^2+^ but in others it is considered to compensate the chaotropicity of Ca^2+^. Moreover, when different ions are found together and when their concentrations increase towards saturation their physicochemical behavior is even more complex (see the case of Mg^2+^ in Fig. 5, and Supplementary Section A). Models are used to estimate the composition and behavior of solutions such as these ones. But, they also have many limitations, for example an ionic strength of 10 M was the limit reported for the applicability of thermodynamic models (Pitzer model) in fractional crystallization of salts from seawater (Keller et al., 2021) and the highest ionic strength registered here was over that limit (13.6 M was the highest registered in the samples). The necessity of developing new models for a more realistic description of fractional characterization of Mg^2+^ in complex salts has already been identified after recognizing the Pitzer overestimation of Mg^2+^ interactions with other ions in Mg^2+^ containing solutions (Keller et al., 2021). Given this confusing situation, it is almost impossible to predict how each brine along the gradient is going to affect living beings, but it is evident that each pond presents a different problem to microorganisms, that will suffer changes in their metabolism and structure (Duda et al. 2004).

The initial brines in the LiCP have an a_w_ similar to that of crystallizer ponds in common solar salterns. However, it is difficult to find possible analogues for the other ponds. Oren (Oren, 2013) reviewed the evidence for life in hypersaline environments rich in Mg^2+^ and Ca^2^, that are the closest to the LiCP. Some of the most extreme brines are the MgCl_2_ deep-sea hypersaline, anoxic basins (DHABs) at the bottom of the Mediterranean Sea (Wallmann et al., 1997; Wallmann, 2001; Hallsworth et al., 2007; La Cono et al., 2019). About eight such basins have been discovered so far along the Mediterranean Ridge. Several are thalassohaline and, thus, rich in NaCl, but Discovery, Kryos and Hephaestus are rich in MgCl_2_ (La Cono et al., 2019). The latter has a brine with an a_w_ = 0.395 and a Mg concentration of 4.7 M. No signs of life could be retrieved from this brine and it was deemed to be sterile. Indication of active cells (as mRNA) were present up to 2.97 M MgCl_2_. This limit was found about 1.5 m down the interface between seawater and the brine where the a_w_ was 0.652. The maximal Mg^2+^ concentration in the LiCP was 1.9 M. Therefore, despite the strong chaotropic effect of Mg^2+^, the concentrations found here would in principle not preclude biological activity. A different matter is the a_w_ since we found values around 0.4 in these Mg^2+^ rich ponds. Thus, microbes were challenged not only with high concentrations of Mg^2+^ but also with lower a_w_. In the last half of the gradient the main salt became LiCl_2_, both ions are considered to be kosmotropic and, therefore, microorganisms able to survive here would have a kind of “relief” from the chaotropic ions but a greater challenge because of the lower water activity.

Another rather extreme environment is Don Juan Pond in the Wright Valley of Antarctica. This pond is a CaCl_2_ solution with concentrations up to 7.2 M Ca^2+^ and 4.2 Cl^-^. This changes with season and it may go down to 0.45. Microbial mats have been found near the shore, but no conclusive evidence of life in the water column exists (Oren, 2013). After the results in the present paper, it would be worth making an effort similar to that made in the DHABs to either confirm or discard the presence of living beings in this very peculiar environment.

### The microbial community

We estimated cell abundance from qPCR data. Since many bacteria have more than one copy of the 16S rRNA gene, these numbers will most likely be overestimates. Even so, there was a clear decrease in numbers with decreasing a_w_. The initial ponds had 10^5^ to 10^6^ copies mL^-1^, a range similar to that of many fresh and marine waters. Numbers then decreased to 10^4^ for most ponds in the middle of the gradient. These concentrations are similar to those from the deep ocean and from many groundwaters (Griebler and Lueders, 2009). The last ponds, finally, only had 100 copies mL^-1^. Some brines also have these low concentrations of cells (REF). We could not retrieve copies from pond P10 even though pond P9, with a similar a_w_ did have 10^4^ copies mL^-1^. Also, pond P6 showed a much lower value than those before and after it along the gradient. Likely, the qPCR did not work properly in these cases.

We found both bacteria and archaea present in the brines. The archaea were found only in the wells with higher NaCl concentration and in a couple of ponds with the lowest salinity. These were all *Halobacteria* as could be expected. Archaea, however, were absent from the rest of the gradient. This is the opposite of what is usually found in solar salterns destined to precipitate common salt, where the proportion of archaea increases from seawater to the crystallizer ponds (Pedrós-Alió, 2005). In principle, we would have expected to find only archaea in the Li^+^ concentration ponds and not bacteria. However, the opposite was true.

A shift from a cosmotropic (NaCl-dominated hypersaline) to a chaotropic saline environment could explain the dynamic of the archaeal taxa in the process. The predominance of archaeal taxa with high GC content (59-70%, except for *Haloquadratum* 48%) observed in wells and ponds at the highest a_w_ (Figs. 8-10) is a common feature in saline environments and it was considered an adaptation strategy (Baliga et al., 2004; Oren, 2013). However, the high GC content was suggested to be a problem for archaeal survival in Mg^2+^ rich saline environments because of the additional stabilizing effect on the already stable DNA. Oren et al. (Oren, 2013) proposed that this Mg^2+^ over-stabilization could prevent DNA replication and transcription in chaotropic saline environments. Intracellular Mg^2+^ concentration measured in *Halobacterium* was higher compared to *E. coli* (Oren, 2013) but we did not find similar information for other halobacteria taxa. Interestingly, a recent work has observed a significant correlation between GC content and halophilicity in bacterial taxa but not in archaeal ones and suggested that it is not an adaptative strategy but a result of biases in DNA repairing systems after breaks induced by high salinity (Hu et al., 2023).

In principle, we could expect a smooth turnover of taxa along a gradient of a_w_ as the one studied here. However, this was clearly not the case. There were two sources of disruption for such a potentially smooth turnover. First, the wells provided brines with rather different composition and, as could be seen, brought in rather different communities to the first ponds. The second source of perturbations to the smooth turnover was the changing chemical composition along the gradient of a_w_. As mentioned, each range of salinity presents the bacteria with a different chemical composition and with different proportions of chaotropic and kosmotropic ions. This has been illustrated in the case of Mg^2+^ in Fig. 8. As the concentration of Mg^2+^ increased, most bacterial groups disappeared from the ponds and finally only a few Proteobacteria dominated the community.

The taxonomy of the bacteria was not that of known halophiles. In fact, *Ralstonia* and *Pseudomonas* are some of the most widespread genera in many different habitats. If we did not have the estimates of activity discussed in the next section, a plausible hypothesis would be that these bacteria had been “pickled” by the high salt content and that, for some reason, some *Ralstonia* and *Pseudomonas* ASVs were those that better resisted under these conditions. It must be remembered that the total number of cells was down to 100 mL^-1^ at the ponds with lowest a_w_. Again, this would be compatible with a progressive loss of ASVs along the gradient, with only some of the most resistant remaining.

In their study of a Li^+^ crystallizer pond, Cubillos et al. (Cubillos et al., 2018) found that most bacterial sequences belonged to *Xanthomonadaceae* and *Staphylococcaceae*. The best known genera of both families are commensals and/or pathogens of animals and plants and their presence in this environment seems peculiar. We did not find any of these two families in our samples. Moreover, since only proportional composition but no indication of abundance was provided in that paper it is difficult to determine the significance of these results (Cubillos et al., 2018). These authors also found an *Halobacteria* population in the lithium concentrated pond (estimated a_w_ by Pitzer equation = 0.221), while we could not find any Archaea along most of the gradient. Their results must be considered with care, however. Cubillos et al. (2018) did not have a satisfactory protocol to concentrate DNA and had to resort to filter 1 mL of brine and carry out nested PCR. Moreover, no report of negative controls was presented. Both issues rise the concern that contamination was possible.

In a second study, Cubillos et al. (2019) successfully isolated in pure culture two *Bacillus* strains from the crystallizer. Neither one was able to grow at the *in situ* concentration of Li^+^, but they did grow up to 1.44 M Li^+^ although they grew considerably faster without Li^+^ in the growth medium. This result would be consistent with the fact that *Bacillus* spores are resistant to many factors. The *Bacillus* might have been pickled in the brines but would have remained alive. Certainly, members of the genus *Bacillus* are good candidates for this strategy. We also found Bacillaceae along the gradient including the most concentrated ponds, although in very low proportion.

### The activity

This situation of bacteria able to tolerate the low a_w_ in a resting stage would have been our most likely explanation for the bacterial community found. However, we have two different estimates of activity. Amplification of cDNA and enrichment cultures.

We could not find cDNA from archaea in any of the samples. In the crystallizers of NaCl solar salterns, the archaea show very low growth rates with long doubling times, up to 72 days (Pedros-Alió et al., 2000). Perhaps the slow growth here prevented us from amplifying cDNA from Archaea. On the other hand, we were able to retrieve bacterial cDNA from the ponds and wells with the highest a_w_ (BH1 – BH3 and P1 and P2) and failed to do so in those with lowest a_w_ (P10 – P12). This could be expected. It was somewhat unexpected, however, that two of the wells and some of the ponds with a_w_ above 0.58 (P3, P5 and P6) did not produce cDNA. It is interesting to note that P5 and P6 are among the samples with the highest Mg^2+^ concentrations (Fig. 5). However, several of these samples provided very little amounts of nucleic acids and our methods were at their limit of detection. But what was really surprising was to retrieve cDNA from ponds P7 to P9, with a_w_ between 0.45 and 0.2. Hallsworth et al. (Hallsworth et al., 2007), among others, argued that 16S rRNA was probably more resistant to degradation under conditions of high Mg^2+^ that at physiological conditions. This could certainly be the case in ponds P7 – P9. If we did not have the enrichment cultures this would have been the most likely explanation.

However, we obtained growth from most wells and ponds. The controls were always negative and, moreover, the positive cultures were transferred and allowed to grow again three times in succession. We must conclude that there was growth at these very low a_w_. Apparently, *in situ* growth in the ponds was prevented by lack of carbon and energy sources and vitamins rather than by low a_w_. Thus, when supplemented, bacteria were able to grow despite the low availability of water.

The first reaction any microbiologist will have after seeing these results will be to assume some kind of contamination. We are convinced that contamination was not an issue for several reasons: a) The controls without inoculum never showed any turbidity; b) since all the cultures were incubated together, a potential contaminant would have been the same in all of them. Yet, different cultures developed different bacterial compositions; c) many of the bacteria that grew in the cultures were already present in the ponds and, moreover, several of them were found to be active with the cDNA criterion. Other bacteria present in the ponds did not grow and some of the bacteria grown in the cultures were not found in the ponds. But this is to be expected whenever running enrichment cultures from a natural system; and d) any contaminant would have been challenged with the same difficulties of low a_w_ than the autochthonous microbiota and, actually, it could be expected to be incapable of overgrowing the natural microbiota. In summary, it seems that some of the bacteria present in the high salinity ponds were able to grow at extremely low a_w_.

The lower limit of a_w_ for life has been proposed to be around 0.63 (Stevenson et al., 2015b). And we observed growth at a_w_ of 0.371. Actually, the brines were amended with 111 mL (LB medium plus minerals and vitamins) per L of brine. This obviously increased the a_w_ and likely “helped” the bacteria to grow. Still, some of the cultures grew at a_w_ definitely below the accepted threshold for life. How could bacteria grow under these extreme conditions? One possibility is that the brine medium was not homogeneous, and the bacteria had access to more water molecules that could be expected from a uniform medium in particular microniches. Certainly, often growth occurred attached to the walls or forming aggregates. Also, the added LB medium might not mix easily with the concentrated brines and form micelles with higher a_w_ where bacteria could grow. Finally, capsules were observed in some of the cultures (data not shown). Both micelles and capsules might provide a more aqueous microenvironment to the cells and permit enough diffusion to keep cellular metabolic activity (Woodward-Rowe et al., 2024). The ability to produce capsules, resist desiccation, and to inhabit xeric environments have been previously reported in *Sphingomonadales* (Zhu et al., 2015; Dorado-Morales et al., 2016; Porcar et al., 2018), one of the taxa enriched in our culturing efforts.

Another factor to consider is chaotropicity. As mentioned, according to the lyotropic series Li, Na, K, Rb, Cs, Ca, Sr and Ba are increasingly chaotropic, while Mg is generally considered chaotropic. In effect we saw a decrease in most taxa as Mg increased. Many of these did not reappear in the gradient, but conditions became less chaotropic after Mg salt precipitated. Li is kosmotropic as well as Cl. As Hallsworth et al. (2007) stated “chaotropicity rather than a_w_ is the limiting factor at MgCl_2_ concentrations around 2.2–2.5 M (that) is consistent with the effect of addition of kosmotropic solutes that further reduce a_w_, but nevertheless alleviate chaotropic stress (Oren, 1983).

## Acknowledgements

Funding. Research Support Project 32002137 from Minera Escondida Ltda.

We acknowledge the contribution of SQM for the support and permits for getting the samples and for their contribution to the system understanding. We thank Dr. John Hallsworth for determination of a_w_ in the 2009 samples.

## Data availability statement

The raw sequence data presented in the study are deposited in the DNA Data Bank of Japan (DDBJ) repository, BioProject: PRJNA1200060.

