## Supplementary information for "The limit of life at extremely low water activity: Lithium-concentration ponds in a solar saltern (Salar de Atacama, Chile)"

**(Salar de Atacama, Chile)**

**Demergasso C. ^1^, Avendaño, P.^1^, Escuti, C.^1^, Veliz, R. ^1^, Chong G. ^2^, Pedrós-Alió C.** ^3^

**Index**

A. **Physico-chemical supplements**

Characterization of the boreholes

Physico-chemistry of the brines

B. **Preliminary data 2009 (Temperature and measured aw)**
C. **Dialysis procedure and its controls**

General setup

Reactants used in qPCR

Water for dialysis

Assessment of fixation and dialysis procedures

Feasibility of RNA extraction after dialysis and cDNA amplification

D. **Culture media used for enrichments**

Recipes and analysis

E. **Reads and taxa retrieved**

Numbers of reads

Removed ASVs

Deposited sequences

F. **Additional information on the microbial diversity of samples and cultures**

Diversity indices

Species richness vs Mg and Li

Families and genera from boreholes and pond samples

**A. Water table depths at the sampled boreholes and main ion concentrations in the brines**

A-1. Characterization of the boreholes

**Supplementary Figure 1**. Characterization of the representative boreholes sampled in this study with different depths (in meters) and ion concentrations (molar).

Molar ion concentration

[SO42-]

| Depth (m) of the boreholes | [Li+] | [Mg2+] | [K+] | [Na+] |
| --- | --- | --- | --- | --- |
| BH1 15 | 0.437977 | 0.472578 | 0.424297 | 2.93995 |
| BH2 30 | 0.358738 | 0.370278 | 1.92225 | 2.69321 |
| BH3 67 | 0.736205 | 0.83587 | 0.636317 | 1.71497 |
| BH4 30 | 1.26495 | 1.05295 | 0.76266 | 1.05527 |
| BH5 150 | 1.43207 | 1.0779 | 0.556522 | 1.04787 |

|  | [SO42-] |
| --- | --- |
| MSW-456 | 0.106224 |
| PC-D1-112 | 0.0670408 |
| CX-098P | 0.147959 |
| W-RY-01 | 0.133673 |
| W-UC-11 | 0.0553061 |

A-2. Physicho-chemistry of the actual brines

**Supplementary Table 1.** Physicochemical parameters and molar ion concentrations of the boreholes and ponds. [Li+], and others means mol/m^3^.

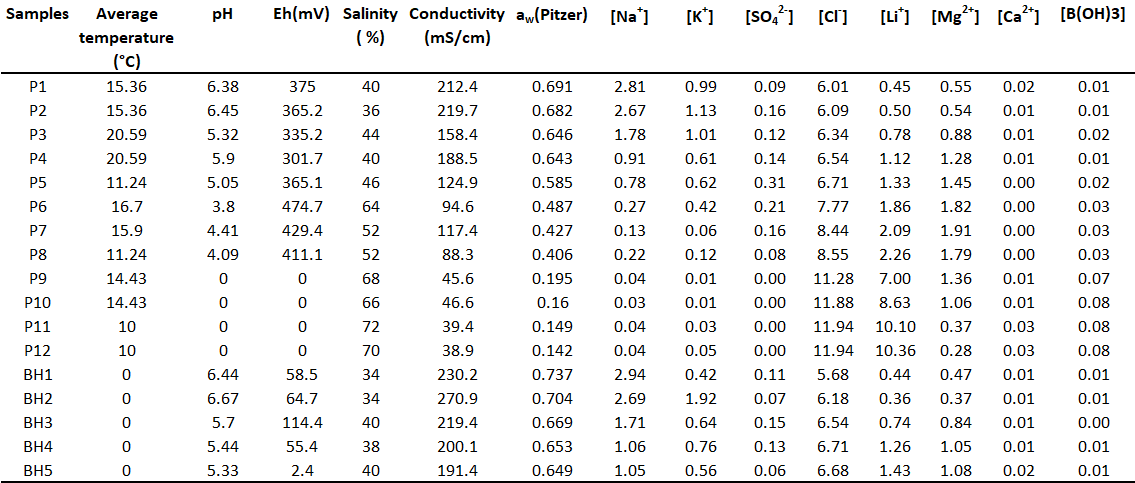

**Supplementary Figure 2.** Principal component analysis (PCA) of the physicochemical parameters of the ponds and boreholes from the SQM plant.

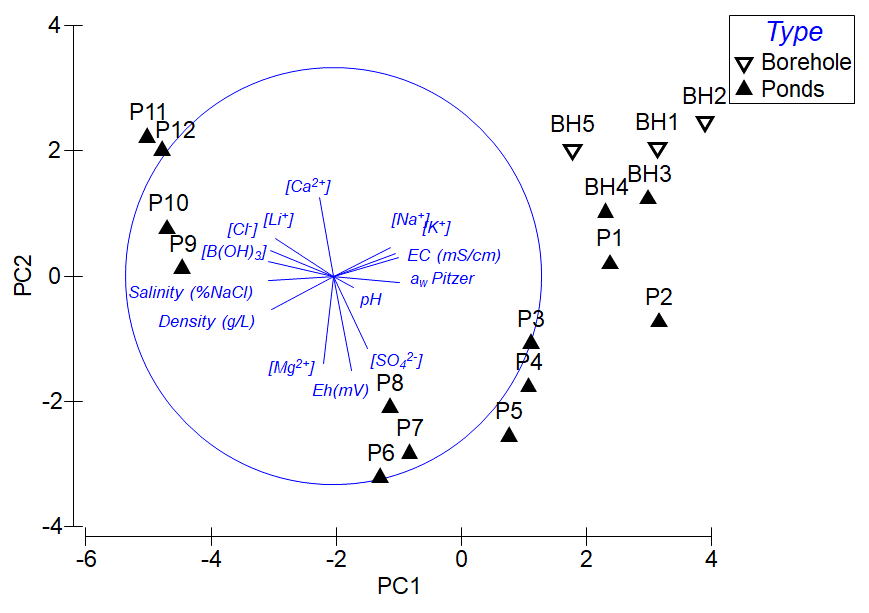

**Supplementary Figure 3.** ORP profile against samples ordered by a_w_. The characteristics of the most concentrated brines prevented the determination of pH and ORP using electrodes.

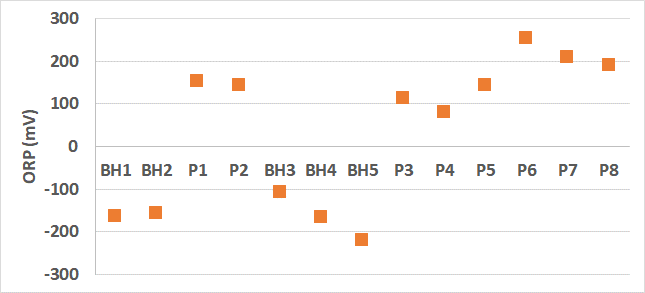

**Supplementary Figure 4**. Correlation between the two different models used in the industry to estimate a_w_ (Luke correlation and GProm model) and the Pitzer equation.

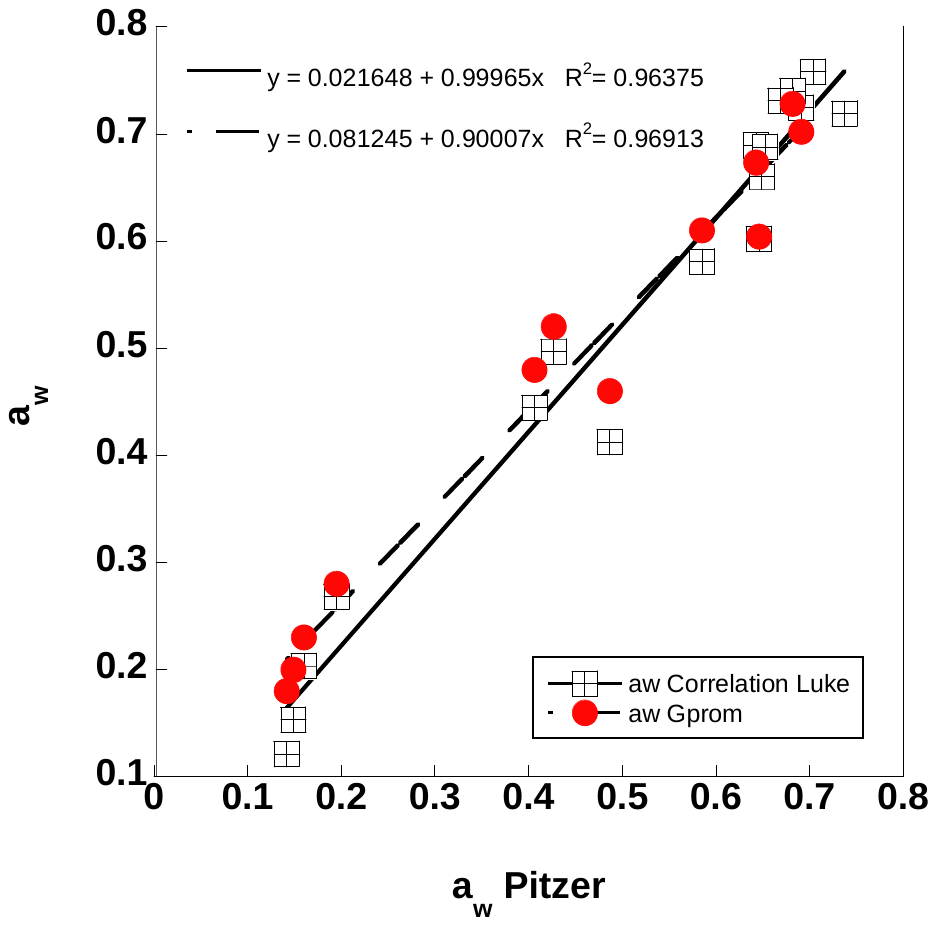

The dynamics of the ion concentrations in LCP obviously depend on the solubility product of the different salts formed by the variety of cations and anions. However, a Mg decrease from P9 to P12 due to carnallite and bischofite precipitation was observed when maximal but not positive level of SI for carnallite (-0.36) and bischofite (-0.8) were reached (Supplementary Fig. 5) according for Pitzer model. It highlighted the necessity of developing new models for a more realistic description of fractional characterization of Mg^2+^ in complex salts as has been previously established (Keller et al., 2021).

Keller, A., Burger, J., Hasse, H., and Kohns, M. (2021). Application of the Pitzer model for describing the evaporation of seawater. *Desalination* 503**,** 114866. doi: <https://doi.org/10.1016/j.desal.2020.114866>.

**Supplementary Figure 5**. Saturation index of carnallite and bischofite with respect of the ponds of the lithium extraction process ordered by a_w_.

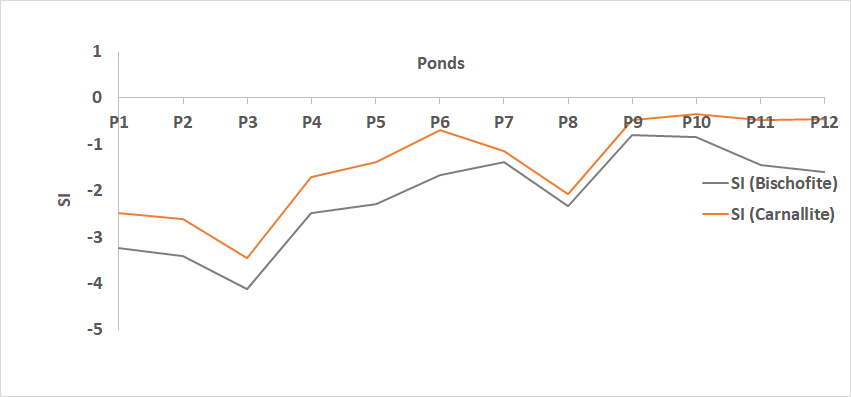

**B. Preliminary data from 2009 sampling at Sociedad Chilena del Litio (Albemarle)**

Temperature could not be determined *in situ* due to COVID limitations in this work. We show a profile obtained in ponds from Sociedad Chilena del Litio in 2009. Temperature always increases with salinity. In addition, water activity was determined experimentally with the method of Hallsworth and Nakamura in 2009. As can be seen, the relationship with lithium concentrations is remarkably similar to that in 2021 (Figure 3) despite the fact that it is a different saltern and a different year.

**Supplementary Figure 6**. Measured water activity and temperature profiles against Li concentration in the evaporation ponds from Sociedad Chilena del Litio determined in 2009. The regression of a_w_ vs lithium concentration was a_w_ = 0.675 * e^(-0.265 [Li]) (R^2^ = 0.895). [Li+] means mol/m^3^.

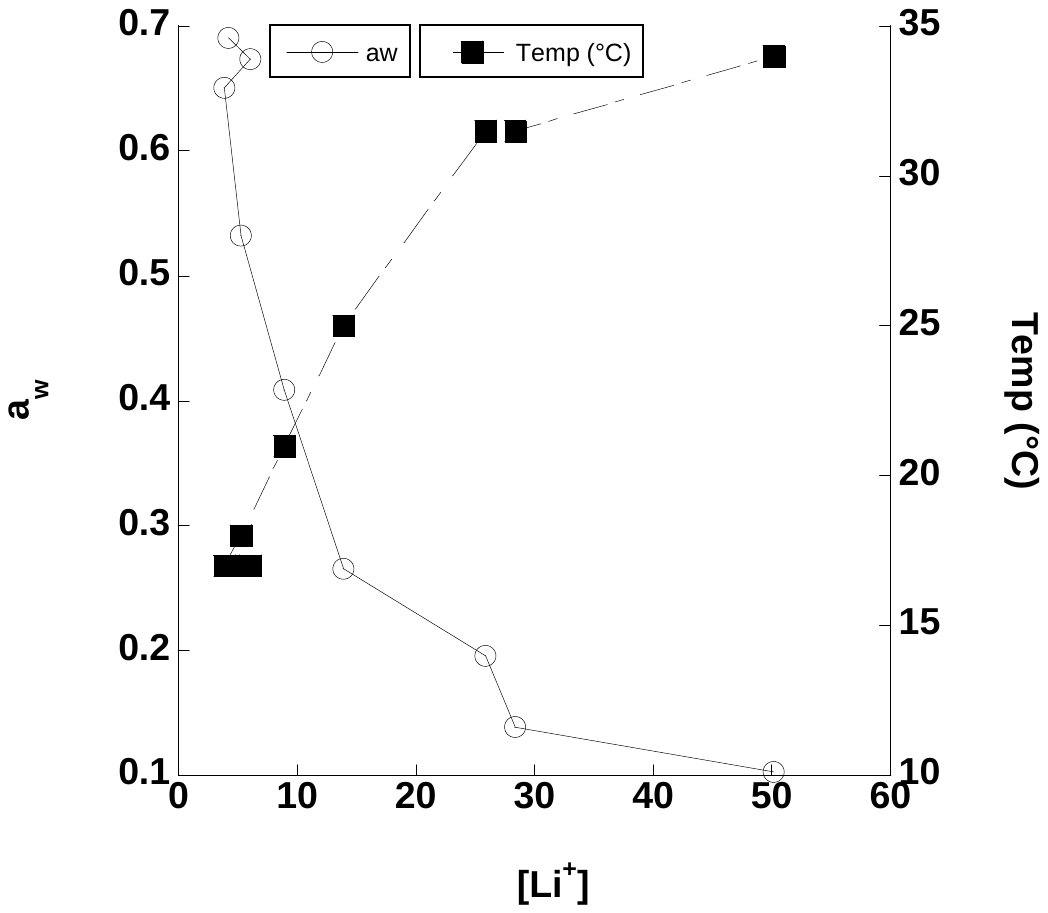

**C. Dialysis procedure and its controls**

C-0. General setup

**Supplementary Figure 7.** Left: beakers containing dialysis bags for brines from two different ponds. Right: measurement of conductivity in the demineralized water outside bags.

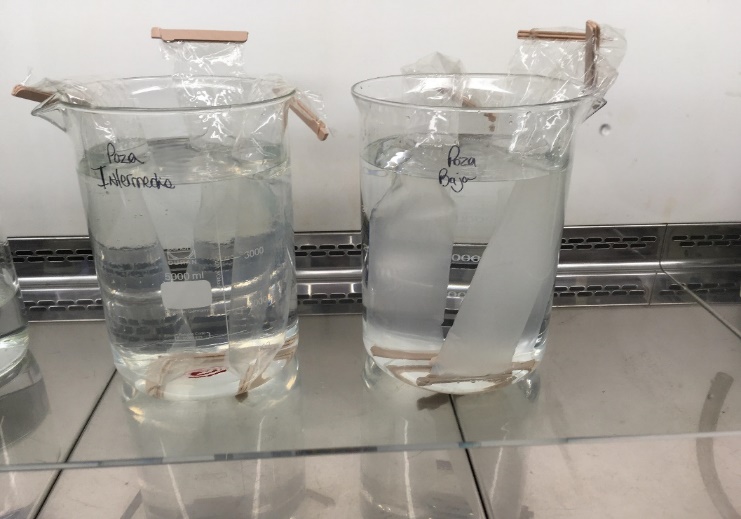

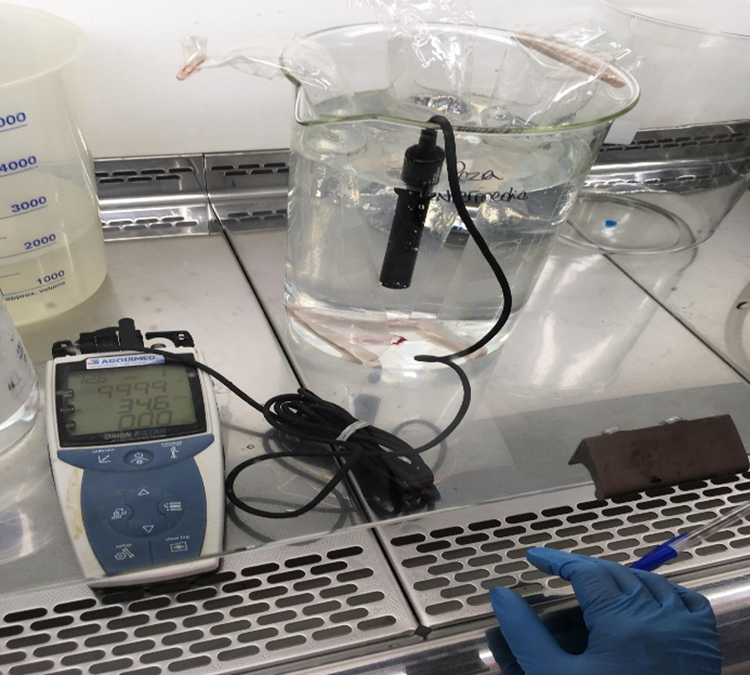

C-1. Control of reactants used in DNA extraction and amplification

Buffers, enzymes, and detergents were checked for the presence of contaminating microorganisms. The equivalent amounts of each reagent were combined and the DNA extracted. The PCR product was run in an 1% agarose electrophoresis gel (90 volts 35 minutes) with positive and negative controls (water). None of the batches assayed produced any DNA.

**Supplementary Figure 8**. Results of DNA extraction reagents by PCR of Bacteria and Archaea and verification by 1% agarose gel electrophoresis (90 volts, 35 min.) ("C+" positive control with DNA of known concentration, "C-" negative control with nuclease-free water (detection limit of the electrophoresis technique is 1E+03 copies µL^-1^).

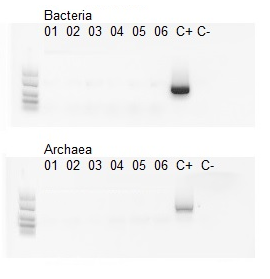

The extracts were analyzed by Q-PCR with bacterial and archaeal primers. All results were below the detection limit of the technique (1E+02 copies mL^-1^).

**Supplementary Table 2.** Quantification by Q-PCR of DNA from DNA Extraction Reagents (copies mL^-1^) (detection limit of the technique 1E+02 copies mL^-1^).

| **Assay** | **Bacteria**  **(copies mL^-1^)** | **Archaea**  **(copies mL^-1^)** |
| --- | --- | --- |
| 01 | 6.24E+00 | 1.48E+00 |
| 02 | 7.08E+00 | 1.24E+00 |
| 03 | 9.96E+00 | 1.08E+00 |
| 04 | 3.84E+00 | 1.15E+00 |
| 05 | 6.69E+00 | 2.40E+00 |
| 06 | 8.48E+00 | 2.06E+00 |

After sequencing a very small number of reads was amplified from these reactants (568 and 4230 reads in two assays), while most samples had numbers of reads above 100000. Despite this, we discarded all the sequences in the samples identical to those obtained here.

**Supplementary Table 3.** Abundance of taxa from 16S analysis present in DNA from DNA Extraction Reagents.

| **Genus** | **Reads numbers** | **Relative Abundance (%)** |
| --- | --- | --- |
| *Georgenia* | 1096 | 22.84 |
| *Halomonas* | 907 | 18.90 |
| *Vibrio* | 652 | 13.59 |
| *Ferroplasma* | 426 | 8.88 |
| *Unclassified* | 322 | 6.71 |
| *Pseudomonas* | 267 | 5.56 |
| *Acidithiobacillus* | 266 | 5.54 |
| *Bacillus* | 93 | 1.94 |
| *Leptospirillum* | 89 | 1.85 |
| *Ralstonia* | 65 | 1.35 |
| *Sphingomonas* | 63 | 1.31 |
| *Chryseoglobus* | 57 | 1.19 |
| *Aeromicrobium* | 54 | 1.13 |
| *Exiguobacterium* | 51 | 1.06 |
| *Actinotalea* | 37 | 0.77 |
| *Microcella* | 36 | 0.75 |
| *Allorhizobium* | 35 | 0.73 |
| *Paenibacillus* | 33 | 0.69 |
| *Alcaligenes* | 27 | 0.56 |
| *Listeria* | 26 | 0.54 |
| *Curvibacter* | 24 | 0.50 |
| *Sulfobacillus* | 21 | 0.44 |
| *Arthrobacter* | 18 | 0.38 |
| *Devosia* | 18 | 0.38 |
| *Limosilactobacillus* | 14 | 0.29 |
| *Bacteroides* | 13 | 0.27 |
| *Providencia* | 13 | 0.27 |
| *Klebsiella* | 12 | 0.25 |
| *Bradyrhizobium* | 11 | 0.23 |
| *Uruburuella* | 10 | 0.21 |
| *Mycobacterium* | 9 | 0.19 |
| *Fusibacter* | 8 | 0.17 |
| *Alloprevotella* | 7 | 0.15 |
| *Haemophilus* | 6 | 0.13 |
| *Acidiphilium* | 5 | 0.10 |
| *Agathobacter* | 4 | 0.08 |
| *Haloparvum* | 3 | 0.06 |
| **Total** | **4798** |  |

C-2. Water used for the dialysis process

Five liters of the demineralized and distilled water used in the dialysis were filtered through 0.22 µm filters. Then, the usual process of extraction of nucleic acids was carried out. The PCR product was run in an 1% agarose electrophoresis gel (90 volts 35 minutes) with positive and negative controls. All batches produced bacterial DNA and one also archaeal DNA. The limit of detection of this technique is 1e03 copies µL^-1^

**Supplementary Figure 9.** Results of DNA of water for dialysis by PCR of Bacteria and Archaea and verification by 1% agarose gel electrophoresis (90 volt for 35 min.) ("C+" positive control with DNA of known concentration, "C-" negative control with nuclease-free water (detection limit of the electrophoresis technique is 1E+03 copies µL^-1^).

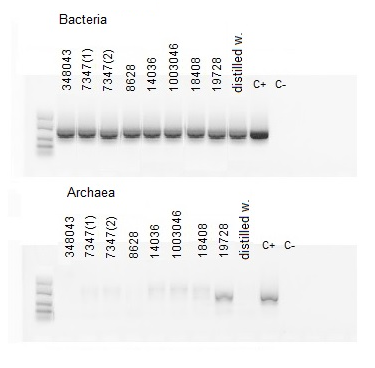

The Q-PCR retrieved bacteria and archaea in all samples above the limit of detection (1E+02 copies mL^-1^). In the case of archaea, values were barely above the limit in the case of bacteria the values were almost one order of magnitude above the limit.

**Supplementary Table 4.** Quantification by Q-PCR of DNA from DNA of water for dialysis (copies mL^-1^) (detection limit of the technique 1E+02 copies mL^-1^).

| **Batch** | **Bacterias**  **(copies mL^-1^)** | **Archaeas**  **(copies mL^-1^)** |
| --- | --- | --- |
| 348043 | 2.45E+04 | BDL |
| 7347 (1) | 1.44E+04 | BDL |
| 7347 (2) | 1.37E+04 | BDL |
| 8628 | 6.47E+03 | BDL |
| 14036 | 9.78E+04 | BDL |
| 1003046 | 1.09E+03 | BDL |
| 18408 | 2.79E+04 | BDL |
| 19728 | 9.28E+04 | 4.33E+02 |

After sequencing the numbers of reads were 38170 and 67222 in two assays. The taxa obtained appear in the following table:

**Supplementary Table 5.** Abundance of taxa from 16S analysis present in DNA from different batches of demineralized and distilled water used for the dialysis process.

| **Genus** | **Desmin. water** | |  | **Genus** | **Distilled water** | |
| --- | --- | --- | --- | --- | --- | --- |
|  | **Read numbers** | **Relative Abundance (%)** |  |  | **Read numbers** | **Relative Abundance (%)** |
| *Sphingomonas* | 46927 | 44,53 |  | *Unclassified* | 48341 | 41.33 |
| *Curvibacter* | 24233 | 22,99 |  | *Sphingomonas* | 36796 | 31,46 |
| *Blastomonas* | 16705 | 15,85 |  | *Bradyrhizobium* | 9212 | 7,88 |
| *Unclassified* | 5590 | 5,30 |  | *Afipia* | 8801 | 7,52 |
| *Bradyrhizobium* | 3371 | 3,20 |  | *Caulobacter* | 2862 | 2,45 |
| *Sediminibacterium* | 1505 | 1,43 |  | *Blastomonas* | 2093 | 1,79 |
| *Vibrionimonas* | 1199 | 1,14 |  | *Methylobacterium-M* | 1627 | 1,39 |
| *Pajaroellobacter* | 1140 | 1,08 |  | *Acidibacter* | 960 | 0,82 |
| *Ralstonia* | 914 | 0,87 |  | *Pedomicrobium* | 957 | 0,82 |
| *Herbaspirillum* | 622 | 0,59 |  | *Sediminibacterium* | 741 | 0,63 |
| *Afipia* | 515 | 0,49 |  | *Candidatus Obscuribacter* | 605 | 0,52 |
| *Methylobacterium-M* | 415 | 0,39 |  | *Novosphingobium* | 440 | 0,38 |
| *Rhodopseudomonas* | 399 | 0,38 |  | *Vibrionimonas* | 294 | 0,25 |
| *Reyranella* | 364 | 0,35 |  | *Undibacterium* | 254 | 0,22 |
| *Undibacterium* | 290 | 0,28 |  | *Polyangium* | 246 | 0,21 |
| *Pedomicrobium* | 232 | 0,22 |  | *Curvibacter* | 226 | 0,19 |
| *Caulobacter* | 212 | 0,20 |  | *Nevskia* | 198 | 0,17 |
| *Nevskia* | 200 | 0,19 |  | *Paucibacter* | 195 | 0,17 |
| *Variovorax* | 116 | 0,11 |  | *Acinetobacter* | 160 | 0,14 |
| *Mesorhizobium (C)* | 80 | 0,08 |  | *Bryobacter* | 141 | 0,12 |
| *Obscuribacter* | 65 | 0,06 |  | *Asinibacterium* | 133 | 0,11 |
| *Heliimonas* | 60 | 0,06 |  | *Fimbriiglobus* | 121 | 0,10 |
| *Acinetobacter* | 45 | 0,04 |  | *Pajaroellobacter* | 113 | 0,10 |
| *Bacillus* | 31 | 0,03 |  | *Methylocella* | 108 | 0,09 |
| *Acidithiobacillus* | 30 | 0,03 |  | *Nitrobacter* | 97 | 0,08 |
| *Tardiphaga* | 21 | 0,02 |  | *Rhodopseudomonas* | 94 | 0,08 |
| *Polyangium]brachysporum* | 18 | 0,02 |  | *Candidatus Ovatusbacter* | 92 | 0,08 |
| *Acidibacter* | 18 | 0,02 |  | *Vibrio* | 92 | 0,08 |
| *Citrobacter* | 18 | 0,02 |  | *Haliangium* | 77 | 0,07 |
| *Georgenia* | 16 | 0,02 |  | *Fusibacter* | 76 | 0,06 |
| *Nitrospirillum* | 11 | 0,01 |  | *Singulisphaera* | 76 | 0,06 |
| *Methylocella* | 10 | 0,01 |  | *Reyranella* | 60 | 0,05 |
| *Halomonas* | 7 | 0,01 |  | *Burkholderia-C* | 54 | 0,05 |
| *Haemophilus* | 5 | 0,00 |  | *Georgenia* | 54 | 0,05 |
| *Opitutus* | 4 | 0,00 |  | *Herbaspirillum* | 47 | 0,04 |
| *Vibrio* | 4 | 0,00 |  | *Acidithiobacillus* | 46 | 0,04 |
|  |  |  |  | *Leptothrix* | 42 | 0,04 |
|  |  |  |  | *Microcella* | 37 | 0,03 |
|  |  |  |  | *Legionella* | 35 | 0,03 |
|  |  |  |  | *Streptococcus* | 29 | 0,02 |
|  |  |  |  | *Bacillus* | 27 | 0,02 |
|  |  |  |  | *Rhodoferax* | 23 | 0,02 |
|  |  |  |  | *Dinghuibacter* | 22 | 0,02 |
|  |  |  |  | *Brevundimonas* | 21 | 0,02 |
|  |  |  |  | *Gemmata* | 21 | 0,02 |
|  |  |  |  | *Mesorhizobium* | 21 | 0,02 |
|  |  |  |  | *Mucilaginibacter* | 20 | 0,02 |
|  |  |  |  | *Jatrophihabitans* | 19 | 0,02 |
|  |  |  |  | *Pseudomonas* | 19 | 0,02 |
|  |  |  |  | *Phyllobacterium* | 18 | 0,02 |
|  |  |  |  | *Stenotrophomonas* | 18 | 0,02 |
|  |  |  |  | *Tardiphaga* | 18 | 0,02 |
|  |  |  |  | *Anaeromyxobacter* | 16 | 0,01 |
|  |  |  |  | *Massilia* | 15 | 0,01 |
|  |  |  |  | *Opitutus* | 12 | 0,01 |
|  |  |  |  | *Cavicella* | 11 | 0,01 |
|  |  |  |  | *Minicystis* | 11 | 0,01 |
|  |  |  |  | *Paenibacillus* | 7 | 0,01 |
|  |  |  |  | *Cutibacterium* | 6 | 0,01 |
|  |  |  |  | *Phenylobacterium* | 6 | 0,01 |
|  |  |  |  | *Aeromicrobium* | 5 | 0,00 |
|  |  |  |  | *Exiguobacterium* | 4 | 0,00 |
| **Total** | **105392** |  |  | **Total** | **116972** |  |

As mentioned, this water is on the outside of the dialysis bags and, therefore, in principle did not contaminate the samples. However, when identical sequences were found in the samples they were discarded.

C-3. Effectiveness of the fixation with ethanol and of the dialysis process

An enriched consortium 1 (CC1) fixed with ethanol was added in a 100 g L^-1^ KCl solution (conductivity 100 – 150 mS cm^-1^). Cells were then collected by filtration through a 0.22 µm pore size membrane and DNA was extracted and sequenced.

A second enriched consortium (CC2) fixed with ethanol was added in a 100 g L^-1^ KCl solution and dialyzed as usual and filtered for DNA extraction and analysis. In both case the DNA of the ethanol fixed CC was also obtained. The Q-PCR with bacterial primers was carried out. In all cases the recovery of both bacteria and archaea was higher than 98% (Supplementary Table 6).

**Supplementary Table 6. Abundance of Bacteria and Archaea in fixation with ethanol and dialysis efficiency controls.**

|  | **Bacteria**  **(copies mL^-1^)** | **Archaea**  **(copies mL^-1^)** | **Total**  **(copies mL^-1^)** | **% Recovery** |
| --- | --- | --- | --- | --- |
| CC1 (F) * | 2.27E+05 | 4.61E+08 | 1.61E+08 |  |
| CC1 (FF) * | 3.23E+05 | 8.96E+07 | 8.99E+07 | 98.16 |
| CC2 (F) | 1.02E+06 | 4.46E+05 | 1.46E+06 |  |
| CC2 (DF) | 2.80E+06 | 3.72E+05 | 3.17E+06 | 100 |
| CC3 (F) * | 1.65E+07 | 1.33E+05 | 1.66E+07 |  |
| CC3 (DF) * | 1.62E+07 | 1.30E+05 | 1.63E+07 | 99.89 |
| CC4 (F) | 8.20E+05 | 1.35E+04 | 8.33E+05 |  |
| CC4 (DF) | 8.02E+05 | 7.88E+03 | 8.10E+05 | 99.79 |
| CC5 (F) | 7.65E+05 | 2.10E+05 | 9.74E+05 |  |
| CC5 (DF) | 2.05E+06 | 7.24E+03 | 2.06E+06 | 100 |
| CC6 (F) | 4.40E+06 | 8.24E+05 | 5.22E+06 |  |
| CC6 (DF) | 3.89E+06 | 1.24E+06 | 5.13E+06 | 99.89 |
| CC7 (F) | 4.21E+06 | 2.37E+04 | 4.23E+06 |  |
| CC7 (DF) | 4.05E+06 | 2.50E+04 | 4.07E+06 | 99.75 |
| CC8 (F) | 8.72E+05 | 3.95E+05 | 1.27E+06 |  |
| CC8 (DF) | 8.31E+05 | 3.39E+05 | 1.17E+06 | 99.44 |
| *(F) Fixed with ethanol.*  *(FF) Fixed with ethanol and filtered.*  *(DF) Fixed with ethanol, dialyzed and filtered. (*) 16S sequencing results* | | | | |

DNA from CC1 and CC3 were sent for 16S sequencing to establish that, in addition to biomass recovering, the same taxa added to the KCl solution were fixed and recovered. As can be seen in Supplementary Figure 10 (A) equal amounts of taxa are recovered from CCs after fixation with 1:1 ethanol and (B) fixation and dialysis, , concluding that the process is effective and efficient.

**Supplementary Figure 10.** 16S sequencing results of CC1 and CC3 after ethanol fixation and dialysis assays for effectiveness assessment. (A) 1:1 ethanol fixation efficiency assay (B) dialysis process efficiency assay. ("F" is fixed with ethanol, "FF" is fixed and filtered through a 0.22 µm pore and "DF" is fixed, dialyzed and filtered through a 0.22 µm pore).

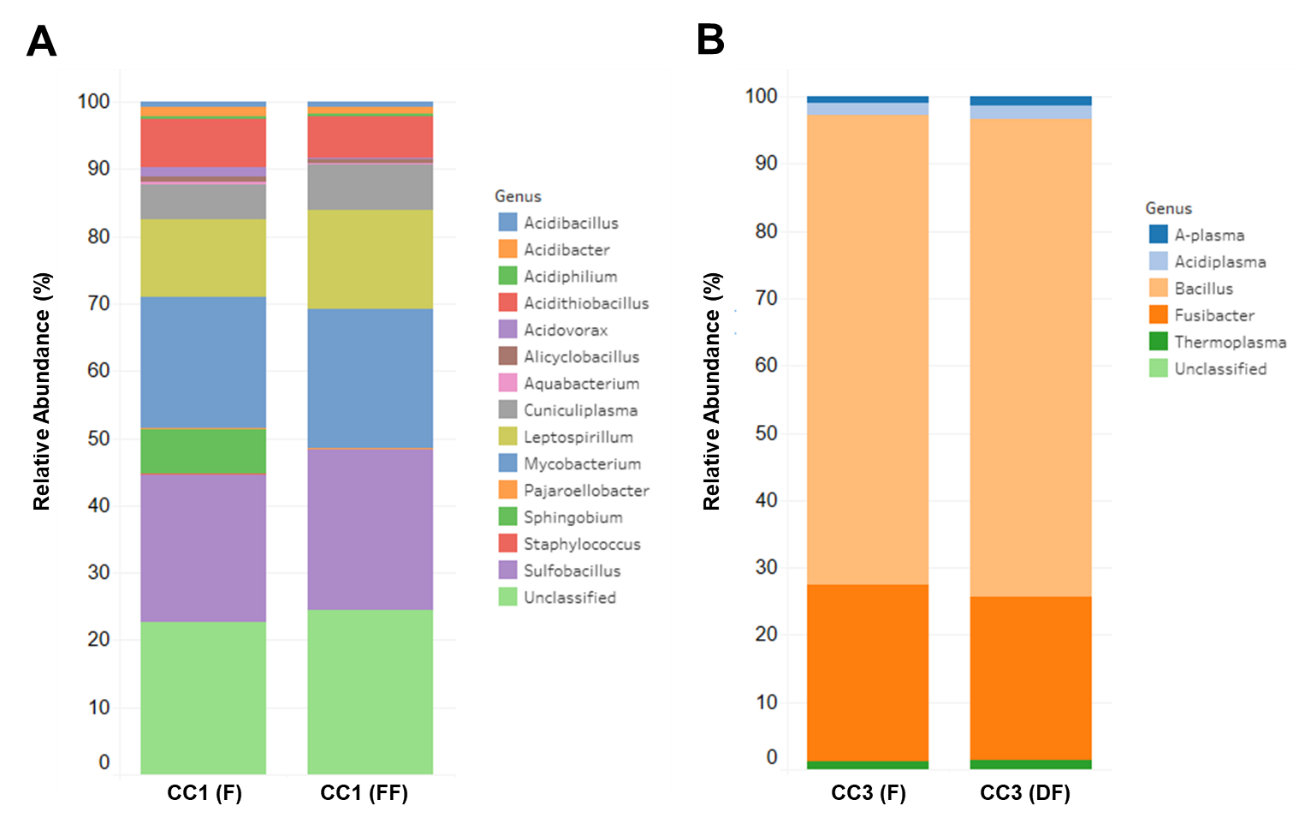

C-4. Feasibility of RNA extraction after dialysis and amplification

A 100 g/L KCl solution (conductivity 100 mS/cm) and a brine (halite type, ~200 mS cm^-1^) were inoculated with an ethanol fixed (1:1) bacterial enriched culture. The cell suspensions were dialyzed (to 20 mS cm^-1^) and filtered. RNA extractions were then carried out with the initial culture and the cells retrieved from both tests.

**Supplementary Table 7.** Quantification of total RNA extracted from saline solutions (KCl and brine).

| **Sample** | **Sample volume**  **(mL)** | **RNA Concentration**  **(ng ml^-1^)** | **Purity 260/280** |
| --- | --- | --- | --- |
| Enriched Consortium (CC) | 50 | 387,37 | 2,01 |
| CC fixed with ethanol in KCl and dialyzed | 5000 | 63,64 | 1,46 |
| CC fixed with ethanol in brine and dialyzed | 5000 | 42,61 | 2,04 |

The RNAs (A) extracted and cDNA (B) amplification were run in an 1% agarose electrophoresis gel (90 volts 35 minutes). The limit of detection of this technique is 1e03 copies µL^-1^

**Supplementary Figure 11.** Evidence of the cDNA amplification using random primers.

A	B

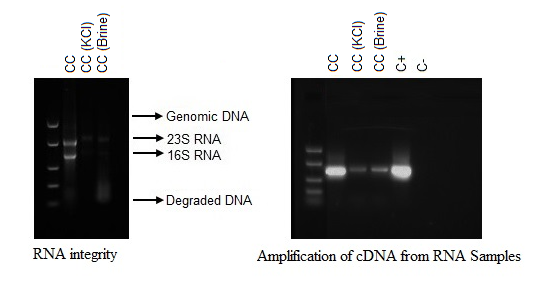

cDNA amplification was observed using a specific primer for Archaea. This test also confirmed that it is possible to amplify the Archaea Domain using Random Primers.

**D. Culture media**

**Trace Metals**

|  | **mg L^-1^** |
| --- | --- |
| Hydrochloric Acid | 1 mL |
| MnCl_2_ x 4 H_2_O | 100 |
| CoCl_2_ x 4 H_2_O | 120 |
| ZnCl_2_ | 30 |
| H_3_BO_3_ | 30 |
| NiCl_2_ x 6 H_2_O | 25 |
| CuCl_2_ x 6 H_2_O | 15 |
| NaMoO_4_ x 2 H_2_O | 25 |
| FeCl_2_ x 4 H_2_O | 15 |

**Vitamins**

|  | **mg L^-1^** |
| --- | --- |
| p- aminobenzoic acid | 5 |
| Biotin | 2 |
| Nicotinic acid | 5 |
| Calcium pantothenate | 5 |
| Thiamin | 5 |
| Pyridoxine | 10 |
| B12 | 0.1 |

**Medium Luria- Bertani (LB)**

A stock solution of the medium 10X should prepared.

|  | **g L^-1^** |
| --- | --- |
| Triptone | 10 |
| Yeast extract | 5 |

**Modified Battaglia culture medium**

|  | **g L^-1^** |
| --- | --- |
| KH_2_PO_4_ | 0.5 |
| NH_4_Cl | 2 |
| Na_2_SO_4_ | 1.4 |
| MgSO_4_ x 7H2O | 12.8 |
| NaCl | 0.6 |
| LiCl | 15 |
| H_3_BO_3_ | 10 |
| FeSO_4_ | 0.1 |
| Yeast extract | 1 |
| **Supplement** | **mL L^-1^** |
| Sodium lactate | 1 |
| Trace Metals | 1 |
| Vitamins | 10 |
| Adjust the pH to 6.5 and sterilize by filtration*.* | |

**Synthetic Brine**

|  | **g L^-1^** |
| --- | --- |
| KCl | 1.44 |
| NaCl | 0.6 |
| Na_2_SO_4_ | 1.4 |
| MgCl_2_ x 6 H_2_O | 5 |
| CaCl_2_ | 25 |
| LiCl | 15 |
| H_3_BO_3_ | 25 |
| MgSO_4_ x 7 H_2_O | 15 |
| **Supplement** | **mL L^-1^** |
| LB (10X) | 100 |
| Trace Metals | 1 |
| Vitamins | 10 |
| Adjust the pH to 6 and sterilize by filtration*.* | |

**Supplementary Table 8.** Results of the chemical analysis performed on the synthetic brine used to maintain the cultures (Estimated a_w_= 0.945).

|  | **g L^-1^** |
| --- | --- |
| Na^+^ | 1.1 |
| K^+^ | 1 |
| Cl^-^ | 45.4 |
| SO_4_^2-^ | 1.8 |
| Ca^2-^ | 0.1 |
| Mg^2-^ | 4.1 |
| Li^+^ | 5.7 |
| H_3_BO_3_ | 15 |

**Supplementary Figure 12**. Diagram of culturing experiments

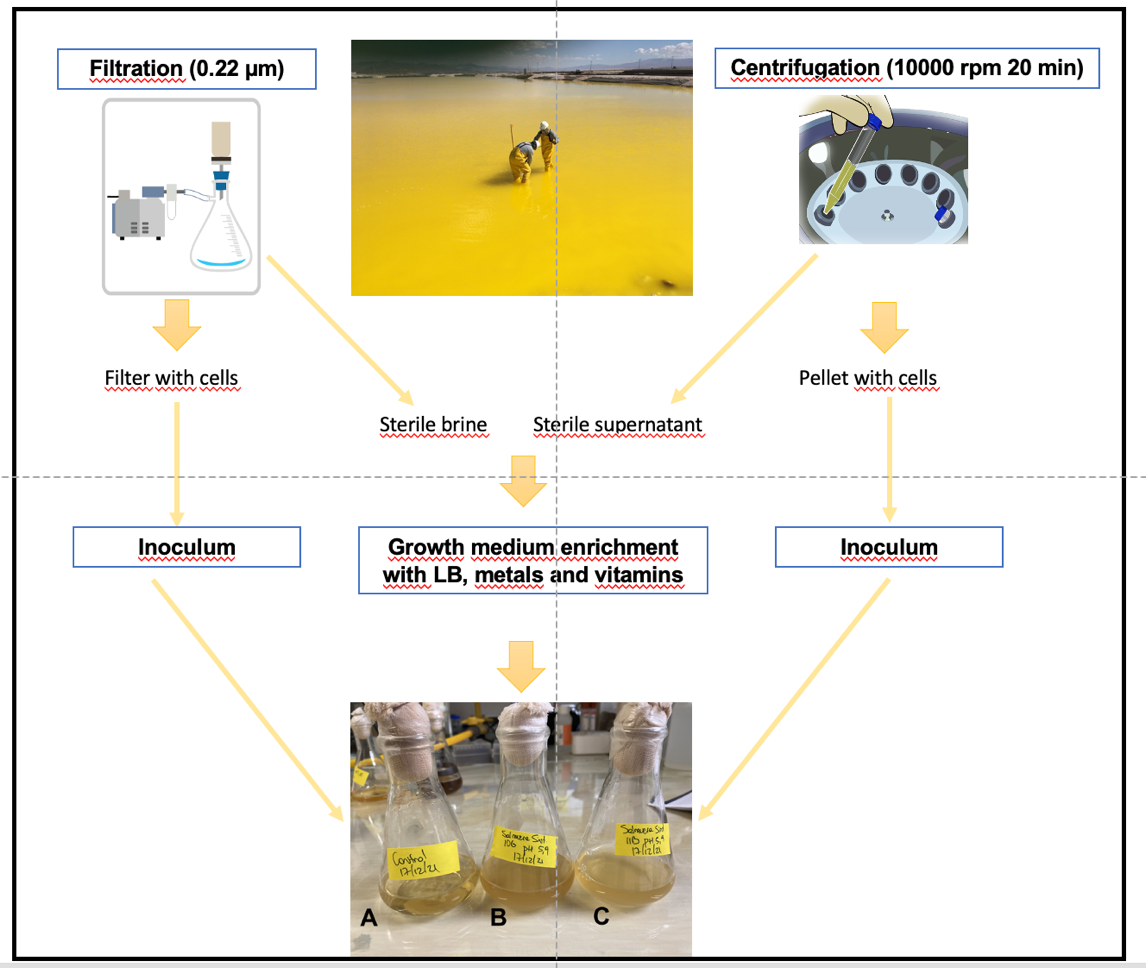

**Supplementary Figure 13.** Aerobic enrichment in LB medium prepared with filtrated brines from P5 and P6 ponds and inoculated by concentrated cells from the same ponds.

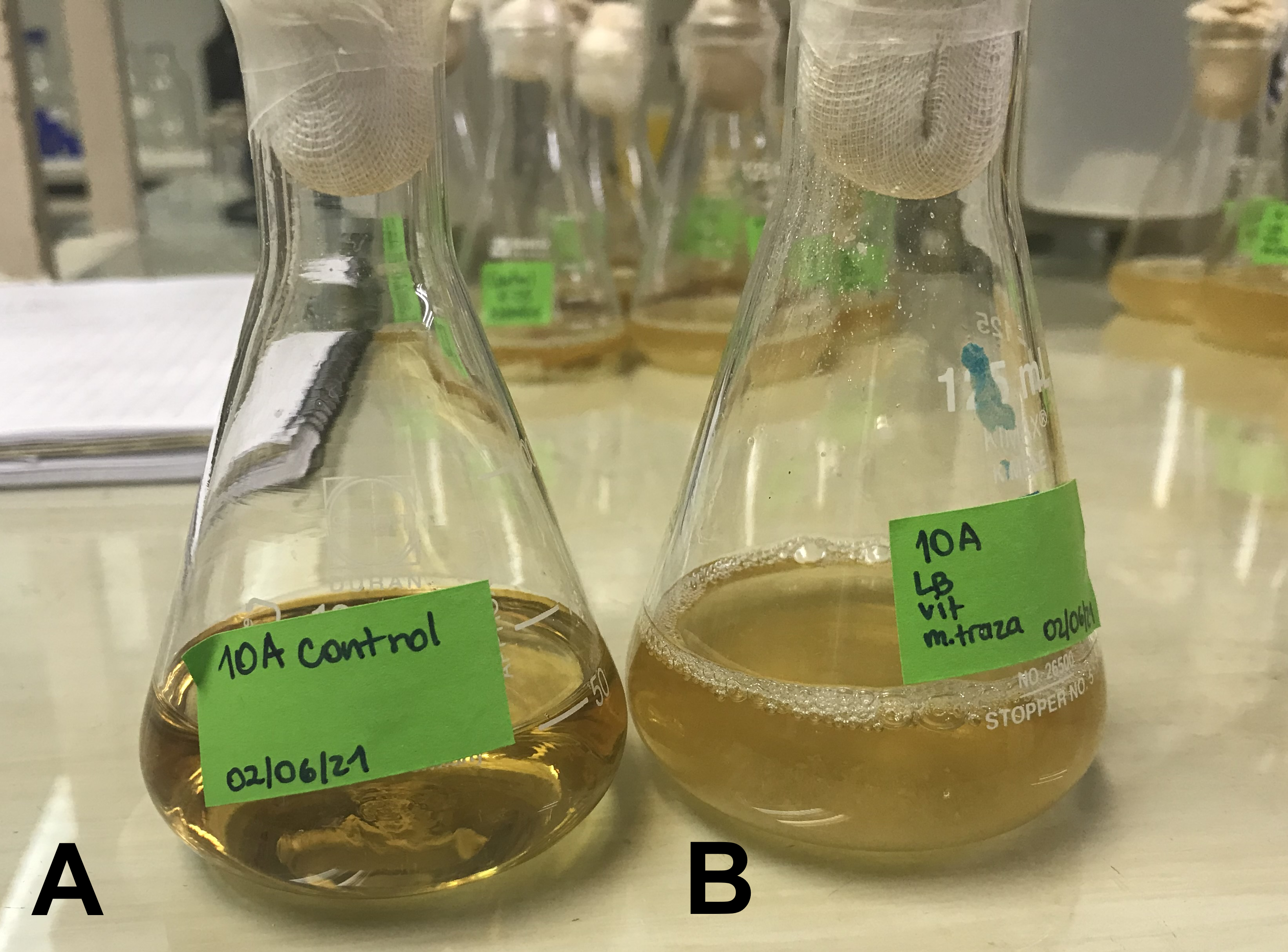

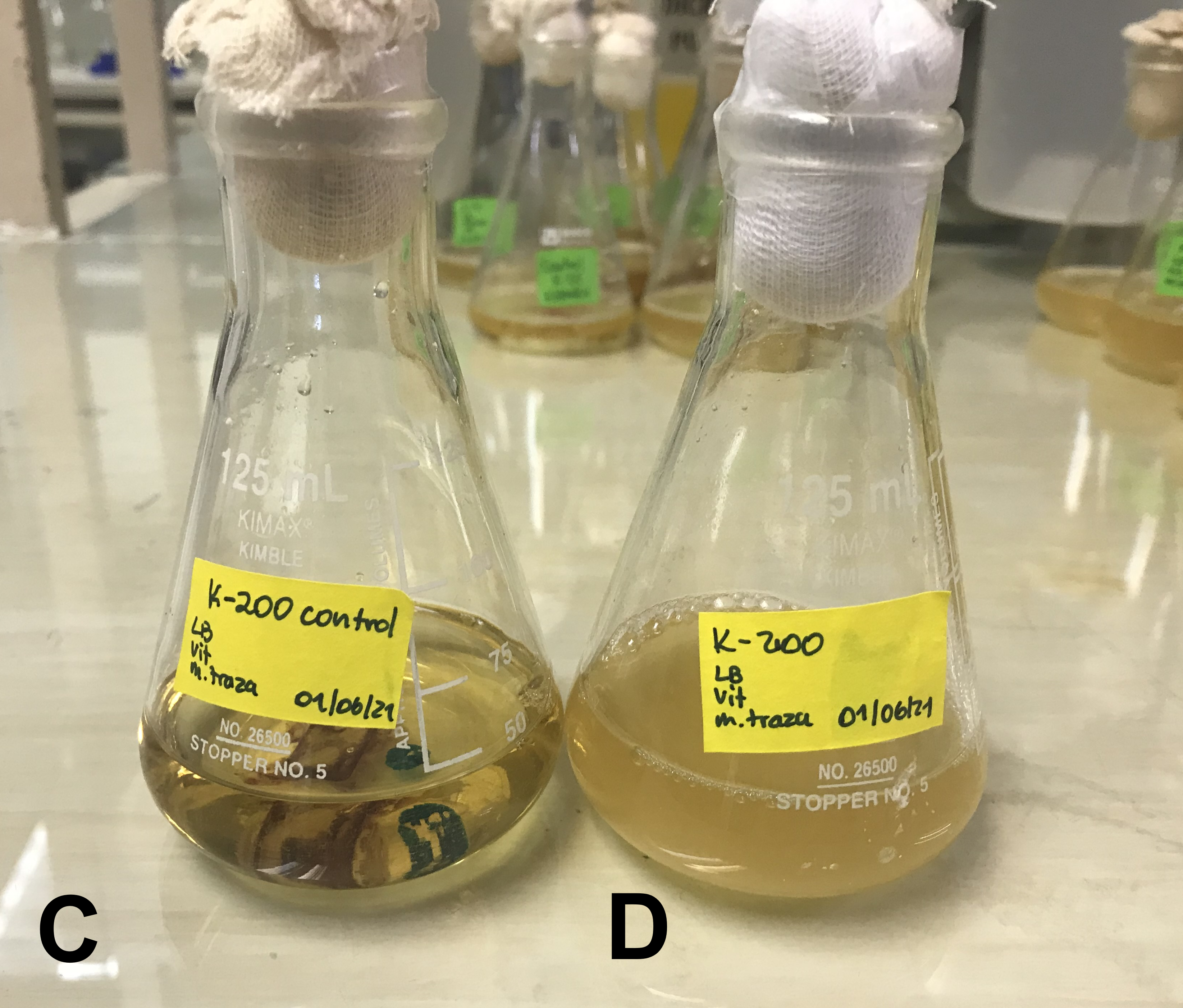

**E. Reads and taxa retrieved**

The unnormalized number of reads obtained from the different samples and controls are shown in the figure below. All borehole and pond samples provided number of reads at least two orders of magnitude higher than the controls. Remember that the demineralized water does not enter in contact with the dialyzed samples. The three most saline samples provided rather few reads, but as mentioned still two orders of magnitude above the controls.

**Supplementary Figure 14.** Number of reads received for each borehole (A) and pond (B) and their controls from the 16S sequencing by Illumina (raw data without quality filters).

|  | Reads |
| --- | --- |
| BH5 | 229498 |
| BH4 | 44110 |
| BH3 | 44843 |
| BH2 | 202839 |
| BH1 | 244022 |
| Desmin. Water | 262520 |
| Reagents | 14057 |
| P12 | 352216 |
| P11 | 232167 |
| P10 | 67926 |
| P9 | 180940 |
| P8 | 351913 |
| P7 | 442230 |
| P6 | 466740 |
| P5 | 554275 |
| P4 | 336549 |
| P3 | 142761 |
| P2 | 101076 |
| P1 | 123268 |
| Distilled water | 116972 |
| Desmin. water R2 | 67222 |
| Desmin. water R1 | 38170 |
| Reagents R2 | 4230 |
| Reagents R1 | 568 |

Then we used the Phyloseq (version 1.42.0) pipeline to (a) remove the ASVs found in controls (Supplementary Table 9), (b) eliminate taxa with one read, (c) remove taxa with less than 0.005% mean relative abundance across all read counts, (d) remove ASVs that were not observed more than twice in at least 10% of the samples; e) eliminate samples having less than 1000 reads and f) all sequences were rarefied to 1756 reads per sample.

**Supplementary Table 9.** List of ASV removed for bioinformatics processing of ponds (P) and boreholes (BH). This table is fairly long and has been provided as Anex 1.

**Supplementary Table 10.** Nucleotide sequence accession numbers at the DNA Sequences have been submitted to the Bank of Japan (DDBJ) repository.

| **Accession Number** | **Sample Name** | **Tax ID** | **BioProject** |
| --- | --- | --- | --- |
| SAMN45894863 | P1_brine | 1981201 | PRJNA1200060 |
| SAMN45894864 | P2_brine | 1981201 | PRJNA1200060 |
| SAMN45894865 | P3_brine | 1981201 | PRJNA1200060 |
| SAMN45894866 | P4_brine | 1981201 | PRJNA1200060 |
| SAMN45894867 | P5_brine | 1981201 | PRJNA1200060 |
| SAMN45894868 | P6_brine | 1981201 | PRJNA1200060 |
| SAMN45894869 | P7_brine | 1981201 | PRJNA1200060 |
| SAMN45894870 | P8_brine | 1981201 | PRJNA1200060 |
| SAMN45894871 | P9_brine | 1981201 | PRJNA1200060 |
| SAMN45894872 | P10_brine | 1981201 | PRJNA1200060 |
| SAMN45894873 | P11_brine | 1981201 | PRJNA1200060 |
| SAMN45894874 | P12_brine | 1981201 | PRJNA1200060 |
| SAMN45894875 | BH1_brine | 1981201 | PRJNA1200060 |
| SAMN45894876 | BH2_brine | 1981201 | PRJNA1200060 |
| SAMN45894877 | BH3_brine | 1981201 | PRJNA1200060 |
| SAMN45894878 | BH4_brine | 1981201 | PRJNA1200060 |
| SAMN45894879 | BH5_brine | 1981201 | PRJNA1200060 |
| SAMN45894880 | Ctrl_agua_dest | 256318 | PRJNA1200060 |
| SAMN45894881 | Ctrl_agua_desmin_R1 | 256318 | PRJNA1200060 |
| SAMN45894882 | Ctrl_agua_desmin_R2 | 256318 | PRJNA1200060 |
| SAMN45894883 | Ctrl_react_extrac_R1 | 256318 | PRJNA1200060 |
| SAMN45894884 | Ctrl_react_extrac_R2 | 256318 | PRJNA1200060 |
| SAMN45894885 | Ctrl_agua_demin_pozos | 256318 | PRJNA1200060 |
| SAMN45894886 | Ctrl_react_extrac_pozos | 256318 | PRJNA1200060 |
| SAMN45894887 | P1_cDNA | 1981201 | PRJNA1200060 |
| SAMN45894888 | P2_cDNA | 1981201 | PRJNA1200060 |
| SAMN45894889 | P3_cDNA | 1981201 | PRJNA1200060 |
| SAMN45894890 | P4_cDNA | 1981201 | PRJNA1200060 |
| SAMN45894891 | P5_cDNA | 1981201 | PRJNA1200060 |
| SAMN45894892 | P6_cDNA | 1981201 | PRJNA1200060 |
| SAMN45894893 | P7_cDNA | 1981201 | PRJNA1200060 |
| SAMN45894894 | P8_cDNA | 1981201 | PRJNA1200060 |
| SAMN45894895 | P9_cDNA | 1981201 | PRJNA1200060 |
| SAMN45894896 | P10_cDNA | 1981201 | PRJNA1200060 |
| SAMN45894897 | P11_cDNA | 1981201 | PRJNA1200060 |
| SAMN45894898 | P12_cDNA | 1981201 | PRJNA1200060 |
| SAMN45894899 | BH1_cDNA | 1981201 | PRJNA1200060 |
| SAMN45894900 | BH2_cDNA | 1981201 | PRJNA1200060 |
| SAMN45894901 | BH3_cDNA | 1981201 | PRJNA1200060 |
| SAMN45894902 | BH4_cDNA | 1981201 | PRJNA1200060 |
| SAMN45894903 | BH5_cDNA | 1981201 | PRJNA1200060 |
| SAMN45894904 | P3_aero | 1981201 | PRJNA1200060 |
| SAMN45894905 | P4_aero | 1981201 | PRJNA1200060 |
| SAMN45894906 | P5_aero | 1981201 | PRJNA1200060 |
| SAMN45894907 | P6_aero | 1981201 | PRJNA1200060 |
| SAMN45894908 | P7_aero | 1981201 | PRJNA1200060 |
| SAMN45894909 | P8_aero | 1981201 | PRJNA1200060 |
| SAMN45894910 | P9_aero | 1981201 | PRJNA1200060 |
| SAMN45894911 | P10_aero | 1981201 | PRJNA1200060 |
| SAMN45894912 | P11_aero | 1981201 | PRJNA1200060 |
| SAMN45894913 | P10_anaero | 1981201 | PRJNA1200060 |
| SAMN45894914 | P11_anaero | 1981201 | PRJNA1200060 |
| SAMN45894915 | P12_anaero | 1981201 | PRJNA1200060 |
| SAMN45894916 | ConsortiumSB | 1981201 | PRJNA1200060 |
| SAMN45894917 | P6SB | 1981201 | PRJNA1200060 |

**F. Additional information for samples and cultures microbial diversity**

**Supplementary Table 11.** Diversity indices in boreholes and ponds ordered by a_w_.

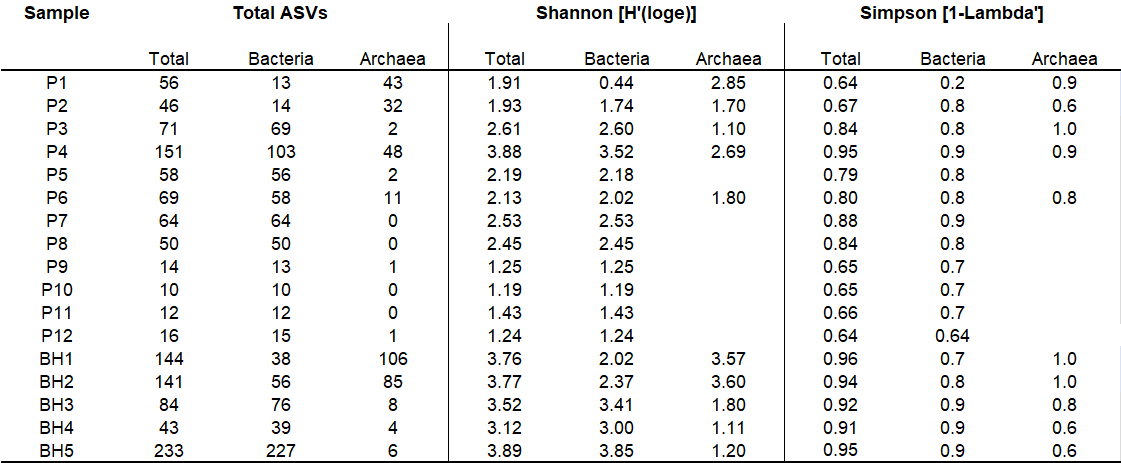

**Supplementary Figure 15**. Correlation between species richness and number of species and molar ion Mg^2+^ and Li^+^ concentrations. [Mg^2+^] and others mean molar concentration.

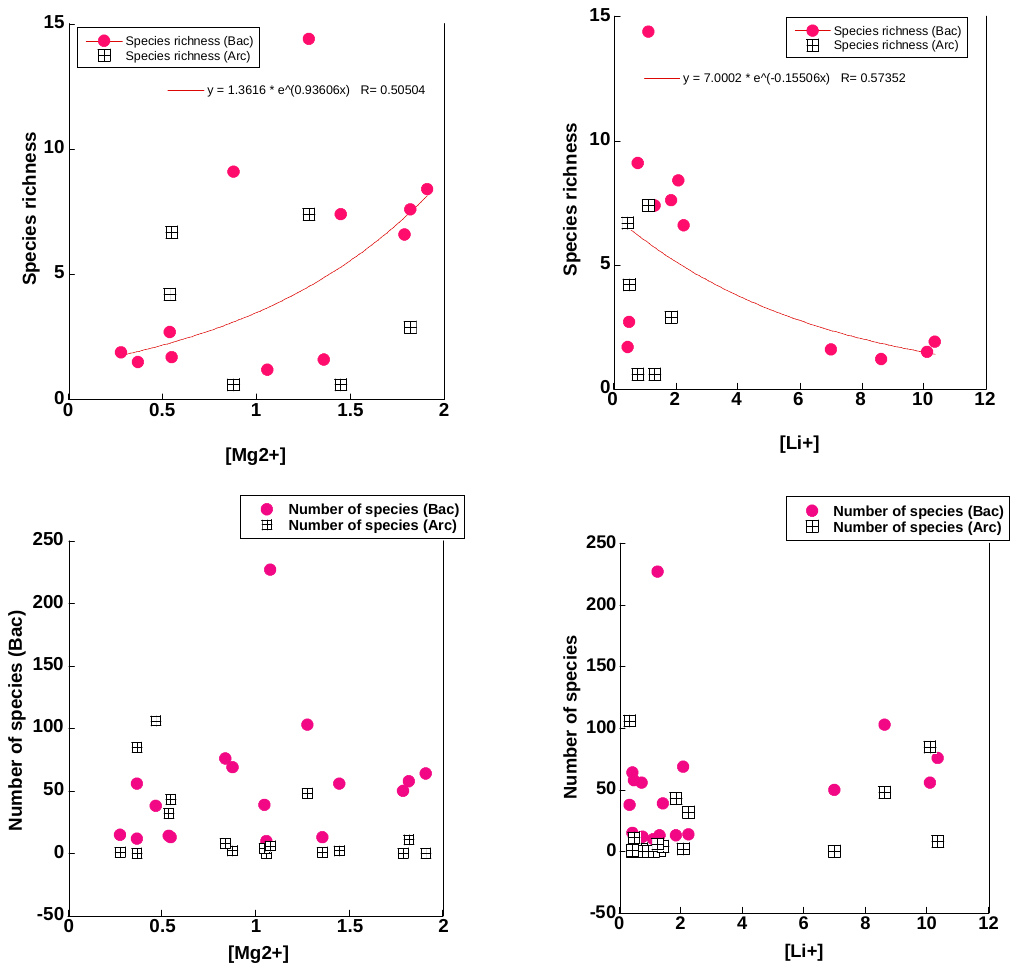

**Supplementary Figure 16.** Relative abundance of Bacteria and Archaea families (**A**) and genera (**B**) in boreholes (BH) and evaporation ponds (P). The boreholes and ponds are ordered by a_w_.

**A**

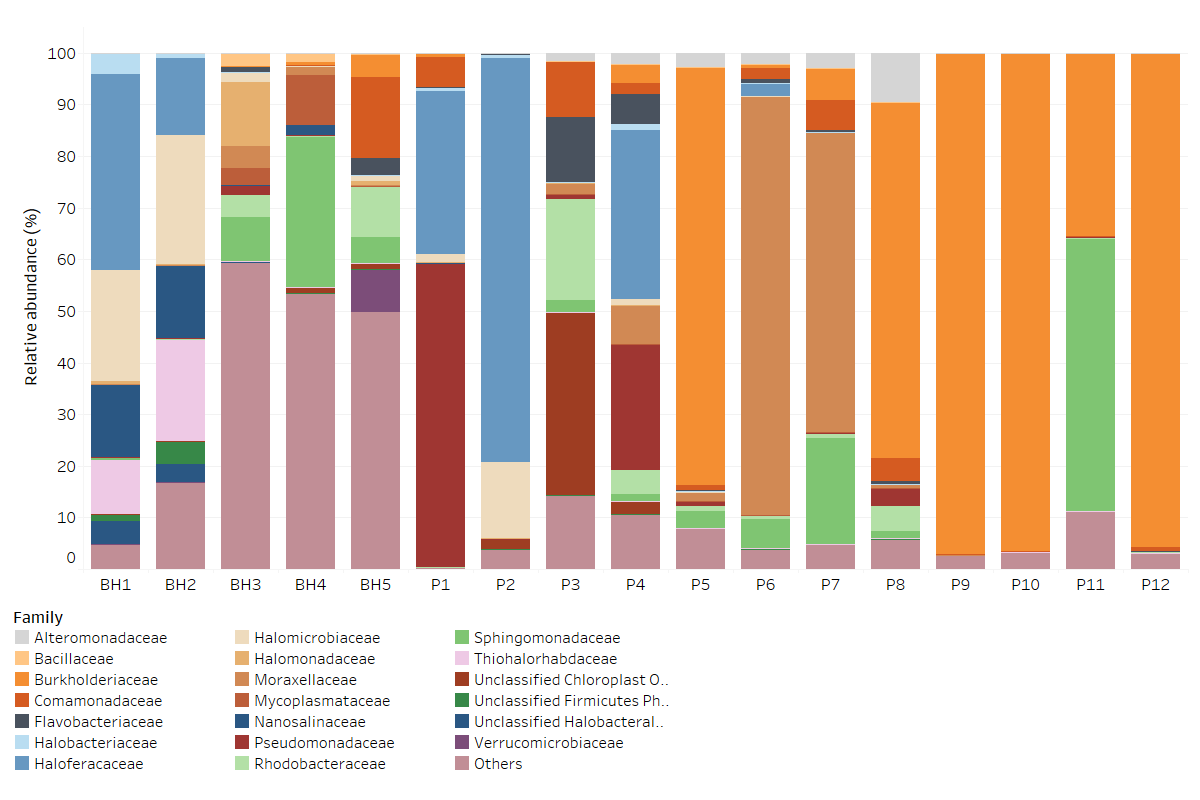

**B**
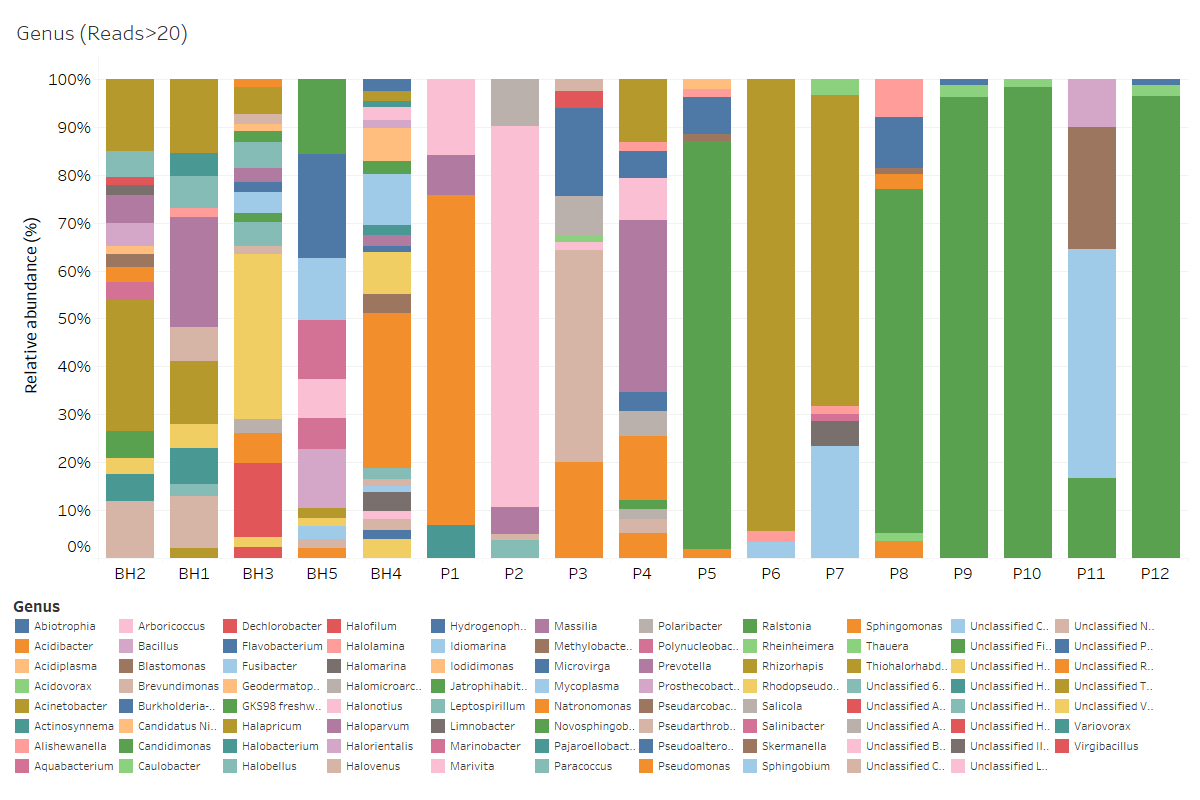
