## Supplementary Table 9 for "The limit of life at extremely low water activity: Lithium-concentration ponds in a solar saltern (Salar de Atacama, Chile)"

Supplementary Table 9. List of ASV removed for bioinformatics processing of ponds (P) and boreholes (BH).

| ASV ID | Sequence |
|---|---|
| P-asv1 | GCTAGCGTTATTCGGAATTACTGGGCGTAAAGCGCACGTAGGCGGCTTTGTAAGTTAGAGGTGAAAGCCTGGAGCTCAACTCCAGAATTGCCTTTAAGACTGCATCGCTTGAATCCAGGAGAGGTGAGTGGAATTCCGAGTGTAGAGGTGAAATTCGTAGATATTCGGAAGAACACCAGTGGCGAAGGCGGCTCACTGGACTGGTATTGACGCTGAGGTGCGAAAGCGTGGGG |
| P-asv2 | CCAAGCGTTAATCGGAATTACTGGGCGTAAAGCGTGCGCAGGCGGTTGTGCAAGACCGATGTGAAATCCCCGAGCTTAACTTGGGAATTGCATTGGTGACTGCACGGCTAGAGTGTGTCAGAGGGGGGTAGAATTCCACGTGTAGCAGTGAAATGCGTAGAGATGTGGAGGAATACCGATGGCGAAGGCAGCCCCCTGGGATAACACTGACGCTCATGCACGAAAGCGTGGGG |
| P-asv3 | GCTAGCGTTGTTCGGAATTACTGGGCGTAAAGCGCACGTAGGCGGCTTTGTAAGTCAGAGGTGAAAGCCTGGAGCTCAACTCCAGAACTGCCTTTGAGACTGCATCGCTTGAATCCAGGAGAGGTGAGTGGAATTCCGAGTGTAGAGGTGAAATTCGTAGATATTCGGAAGAACACCAGTGGCGAAGGCGGCTCACTGGACTGGTATTGACGCTGAAGTGCGAAAGCGTGGGG |
| P-asv4 | GCTAGCGTTGCTCGGAATTACTGGGCGTAAAGGGCGCGTAGGCGGACAGTTAAGTTGGGGGTGAAAGCCCGGGGCTCAACCTCGGAAATGCCTTCAATACTGGCTGTCTTGAGTACGGGAGAGGTGAGTGGAACTCCGAGTGTAGAGGTGAAATTCGTAGATATTCGGAAGAACACCAGTGGCGAAGGCGACTCACTGGCCCGTTACTGACGCTGAGGCGCGAAAGCGTGGGG |
| P-asv5 | GCAAGCGTTGTCCGGAATTATTGGGCGTAAAGCGCGCGCAGGCGGTCCTTTAAGTCTGATGTGAAAGCCCACGGCTCAACCGTGGAGGGTCATTGGAAACTGGGGGACTTGAGTGCAGAAGAGAAGAGTGGAATTCCACGTGTAGCGGTGAAATGCGTAGAGATGTGGAGGAACACCAGTGGCGAAGGCGACTCTTTGGTCTGTAACTGACGCTGAGGCGCGAAAGCGTGGGG |
| P-asv6 | GCGAGCGTTAATCGGATTTACTGGGCGTAAAGCGTGCGTAGGCGGCTTTTTAAGTCGGATGTGAAATCCCTGAGCTTAACTTAGGAATTGCATTCGATACTGGGAAGCTAGAGTATGGGAGAGGATGGTAGAATTCCAGGTGTAGCGGTGAAATGCGTAGAGATCTGGAGGAATACCGATGGCGAAGGCAGCCATCTGGCCTAATACTGACGCTGAGGTACGAAAGCATGGGG |
| P-asv7 | GCAAGCGTTATCCGGAATCACTGGGCGTAAAGGGTGCGTAGGCGGCCTAACAAGTCAGAGGTGAAAGGCATTGGCTCAACCAATGTAAGCCTTTGAAACTGTTGGGCTTGAGTGCAGGAGAGGTAAGTGGAATTCCTAGTGTAGAGGTGAAATTCGTAGATATTAGGAGGAACACCAGTGGCGAAGGCGACTTACTGGACTGTAACTGACGCTGAGGCACGAAAGCGTGGGG |
| P-asv8 | GCGAGCGTTAATCGGAATTACTGGGCGTAAAGCGTGCGCAGGCGGTTATGTAAGACAGATGTGAAATCCCCGGGCTCAACCTGGGAACTGCATTAGTGACTGCATAGCTGGAGTACGGCAGAGGGGGATGGAATTCCGCGTGTAGCAGTGAAATGCGTAGATATGCGGAGGAACACCGATGGCGAAGGCAATCCCCTGGGCCTGTACTGACGCTCATGCACGAAAGCGTGGGG |
| P-asv9 | GCTAGCGTTGCTCGGAATCACTGGGCGTAAAGGGTGCGTAGGCGGGTCTTTAAGTCAGGGGTGAAATCCTGGAGCTCAACTCCAGAACTGCCTTTGATACTGAAGATCTTGAGTTCGGGAGAGGTGAGTGGAACTGCGAGTGTAGAGGTGAAATTCGTAGATATTCGCAAGAACACCAGTGGCGAAGGCGGCTCACTGGCCCGATACTGACGCTGAGGCACGAAAGCGTGGGG |
| P-asv10 | GCAAGCGTTGTTCGGAATTACTGGGCGTAAAGCGTGCGTAGTCGGTATTGAGAGTCACGGGTGAAATCCCAGGGCTTAACCCTGGAACTGCCTGTGAGACCTCAGTACTAGAGTGTGAGAGGGGATAGTGGAATACCCAGTGTAGCGGTGAAATGCGTAGAGATTGGGTGGAACACCGGTGGCGAAGGCGGCTATCTGGCTCACAACTGACGATCAGGCACGAAAGCGTGGGG |
| P-asv11 | GCTAGCGTTGCTCGGAATCACTGGGCGTAAAGGGTGCGTAGGTGGATCTTTAAGTCAGGGGTGAAATCCTGGAGCTCAACTCCAGAACTGCCTTTGATACTGAGGATCTTGAGTTCGGGAGAGGTAAGTGGAACTGCGAGTGTAGAGGTGAAATTCGTAGATATTCGCAAGAACACCAGTGGCGAAGGCGGCTTACTGGCCCGATACTGACACTGAGGCACGAAAGCGTGGGG |
| P-asv12 | GCTAGCGTTGTTCGGAATTACTGGGCGTAAAGCGCACGTAGGCGGTTATTCAAGTCAGAGGTGAAAGCCCGGGGCTCAACCCCGGAACTGCCTTTGAAACTAGATGACTGGAATCTTGGAGAGGCGAGTGGAATTCCGAGTGTAGAGGTGAAATTCGTAGATATTCGGAAGAACACCAGTGGCGAAGGCGACTCGCTGGACAAGTATTGACGCTGAGGTGCGAAAGCGTGGGG |
| P-asv13 | GCAAGCGTTAATCGGAATTACTGGGCGTAAAGCGTGCGCAGGCGGTTTTGTAAGTCTGATGTGAAATCCCCGGGCTCAACCTGGGAACTGCATTGGAGACTGCAAGGCTAGAGTGTGTCAGAGGGGGGTAGAATTCCACGTGTAGCAGTGAAATGCGTAGAGATGTGGAGGAATACCGATGGCGAAGGCAGCCCCCTGGGATAACACTGACGCTCATGCACGAAAGCGTGGGG |
| P-asv14 | GCAAGCGTTGTCCGGAATTATTGGGCGTAAAGCGCGCGCAGGCGGTTCCTTAAGTCTGATGTGAAAGCCCCCGGCTCAACCGGGGAGGGTCATTGGAAACTGGGGAACTTGAGTGCAGAAGAGGAGAGCGGAATTCCACGTGTAGCGGTGAAATGCGTAGAGATGTGGAGGAACACCAGTGGCGAAGGCGGCTCTCTGGTCTGTAACTGACGCTGAGGCGCGAAAGCGTGGGG |
| P-asv15 | GCGAGCGTTAATCGGATTTACTGGGCGTAAAGCGTGCGTAGGCGGCTTCTTAAGTCGGATGTGAAATCCCTGAGCTTAACTTAGGAATTGCATTCGATACTGGGAAGCTAGAGTATGGGAGAGGATGGTAGAATTCCAGGTGTAGCGGTGAAATGCGTAGAGATCTGGAGGAATACCGATGGCGAAGGCAGCCATCTGGCCTAATACTGACGCTGAGGTACGAAAGCATGGGG |
| P-asv19 | GCGAGCGTTAATCGGATTTACTGGGCGTAAAGCGTGCGTAGGCGGCTTTTTAAGTCGGATGTGAAATCCCTGAGCTTAACTTAGGAGTTGCATTCGATACTGGGAAGCTAGAGTATGGGAGAGGATGGTAGAATTCCAGGTGTAGCGGTGAAATGCGTAGAGATCTGGAGGAATACCGATGGCGAAGGCAGCCATCTGGCCTAATACTGACGCTGAGGTACGAAAGCATGGGG |
| P-asv20 | GCAAGCGTTAATCGGAATTACTGGGCGTAAAGCGCGCGTAGGTGGTTTGTTAAGTTGAATGTGAAAGCCCCGGGCTCAACCTGGGAACTGCATCCAAAACTGGCAAGCTAGAGTATGGCAGAGGGTGGTGGAATTTCCTGTGTAGCGGTGAAATGCGTAGATATAGGAAGGAACACCAGTGGCGAAGGCGACCACCTGGGCTAATACTGACACTGAGGTGCGAAAGCGTGGGG |
| P-asv22 | GCGAGCGTTAATCGGATTTACTGGGCGTAAAGCGTGCGTAGGCGGCTTTTTAAGTCGGGTGTGAAATCCCTGAGCTTAACTTAGGAATTGCATTCGATACTGGGAAGCTAGAGTATGGGAGAGGATGGTAGAATTCCAGGTGTAGCGGTGAAATGCGTAGAGATCTGGAGGAATACCGATGGCGAAGGCAGCCATCTGGCCTAATACTGACGCTGAGGTACGAAAGCATGGGG |
| P-asv23 | TCGAGTGATGGCCGATCTTATTGGGCCTAAAGCGTCCGTAGCTGGCCGTGCAAGTCCGTCGGGAAATCCACCCGCTCAACGGGTGGGCGGCCGACGGAAACTGGACGGCTTGGGACCGGGAGGCTTGAAGGGTACGTCCGGGGTAGGAGTGAAATCCCGTAATCCTGGACGGACCACCGATGGGGAAACCACTTCAAGAGAACGGATCCGACAGTGAGGGACGAAAGCTAGGGT |
| P-asv26 | GCAAGCGTTAATCGGAATTACTGGGCGTAAAGCGCGCGTAGGTGGTTAGTTAAGTTGGATGTGAAATCCCCGGGCTCAACCTGGGAACTGCATTCAAAACTGACTGACTAGAGTATGGTAGAGGGTGGTGGAATTTCCTGTGTAGCGGTGAAATGCGTAGATATAGGAAGGAACACCAGTGGCGAAGGCGACCACCTGGACTGATACTGACACTGAGGTGCGAAAGCGTGGGG |
| P-asv27 | TCGAGTGATGTCCCATATTATTGGGCCTAAAGCGTCCGTAGCTGGCCAGGCAAGTCTATCGGGAAATCCACCTGCCCAACAGGTGGGCGTCCGGTAGAAACTGCCCGGCTTGGAACCGGAAGGCTCAGAGAGTACGTCCGGGGTAGGAGTGAAATCCCGTAATCCCGGACGGACTACCGATGGCGAAAGCACTCTGAGAAGACGGCTTCGACAGTGAGGGACGAAAGCTAGGGT |
| P-asv33 | CCAAGCGTTAATCGGAATTACTGGGCGTAAAGCGTGCGCAGGCGGTTGTGCAAGACCGATGTGAAATCCCCGAGCTTAACTTGGGAATTGCATTGGTGACTGCACGGCTAGAGTGTGTCAGAGGGGGGTAGAATTCCACGTGTAGCAGTGAAATGCGTAGAGATGTGGAGGAATACCGATGGCGAAGGCAGCCACCTGGGATAACACTGACGCTCATGCACGAAAGCGTGGGG |
| P-asv35 | CCAAGCGTTAATCGGAATTACTGGGCGTAAAGCGTGCGCAGGCGGTTGTGCAAGACCGATGTGAAATCCCCGAGCTTAACTTGGGAATTGCATTGGTGACTGCACGGCTAGAGTGTGTCAGAGGGGGGTAGAATTCCACGTGTAGCAGTGAAATGCGTAGAGATGTGGAGGAATACCGATGGCGAAGGCAGCACCCTGGGATAACACTGACGCTCATGCACGAAAGCGTGGGG |
| P-asv37 | GCTAGCGTTGTTCGGAATTACTGGGCGTAAAGCGCACGTAGGCGGCGATTTAAGTCAGAGGTGAAAGCCCGGGGCTCAACCCCGGAACTGCCTTTGAGACTGGATTGCTTGAATCCTGGAGAGGTGAGTGGAATTCCGAGTGTAGAGGTGAAATTCGTAGATATTCGGAAGAACACCAGTGGCGAAGGCGGCTCACTGGACAGGTATTGACGCTGAGGTGCGAAAGCGTGGGG |
| P-asv38 | GCTAGCGTTGTTCGGAATTACTGGGCGTAAAGCGCACGTAGGCGGCTATTCAAGTCAGAGGTGAAAGCCCGGGGCTCAACCCCGGAACTGCCTTTGAAACTAGATAGCTTGAATCCAGGAGAGGTGAGTGGAATTCCGAGTGTAGAGGTGAAATTCGTAGATATTCGGAAGAACACCAGTGGCGAAGGCGGCTCACTGGACTGGTATTGACGCTGAGGTGCGAAAGCGTGGGG |
| P-asv41 | GCAAGTGTTATCCGGAATCACTGGGCGTAAAGCGTCTGTAGGTGGTTTAATAAGTCAACTGTTAAATCTTGAAGCTCAACTTCAAAATCGCAGTCGAAACTATTAGACTAGAGTATAGTAGGGGTAAAGGGAATTTCCAGTGGAGCGGTGAAATGCGTAGAGATTGGAAAGAACACCGATGGCGAAGGCACTTTACTGGGCTATTACTGACACTCAGAGACGAAAGCTAGGGT |
| P-asv42 | GCAAGCGTTATCCGGAATCATTGGGTTTAAAGGGTCCGCAGGCGGTCAATTAAGTCAGAGGTGAAATCCAGTCGCTTAACGATTGAACTGCCTTTGATACTGGTTGACTTGAGTCATATGGAAGTAGATAGAATGTGTAGTGTAGCGGTGAAATGCTTAGATATTACACAGAATACCGATTGCGAAGGCAGTCTACTACGTATGTACTGACGCTCATGGACGAAAGCGTGGGG |
| P-asv45 | TCAAGTGATGTCCAATATTATTGGGCCTAAAGCGTCCGTAGCTGGCCGCGCAAGTCCGTCGGGAAATCCACCTGCCCAACAGGTGGGCGTCCGGCGGAAACTGTGTGGCTTGGAACCGGAAGGCTCAGAGGGTACGTCCGGGGTAGGAGTGAAATCCCGTAATCCCGGACGGACCACCGATGGCGAAAGCACTCTGAGAAGACGGCTTCGACAGTGAGGGACGAAAGCTAGGGT |
| P-asv49 | GCAAGCGTTAATCGGAATTACTGGGCGTAAAGCGTGCGCAGGCGGTTGTGTAAGACAGGTGTGAAATCCCCGGGCTTAACCTGGGAATTGCATTTGTGACTGCACGACTAGAGTGTGTCAGAGGGGGGTGGAATTCCACGTGTAGCAGTGAAATGCGTAGATATGTGGAGGAACACCAATGGCGAAGGCAGCCCCCTGGGATAACACTGACGCTCATGCACGAAAGCGTGGGG |
| P-asv51 | GCAAGCGTTAATCGGAATTACTGGGCGTAAAGCGTGCGCAGGCGGTTTTGTAAGACAGGCGTGAAATCCCCGGGCTCAACCTGGGAATTGCGCTTGTGACTGCAAGGCTGGAGTGCGGCAGAGGGGGATGGAATTCCGCGTGTAGCAGTGAAATGCGTAGATATGCGGAGGAACACCGATGGCGAAGGCAATCCCCTGGGCCTGCACTGACGCTCATGCACGAAAGCGTGGGG |
| P-asv54 | GCTAGCGTTATTCGGAATTACTGGGCGTAAAGCGCACGTAGGCGGACTGGAAAGTCAGAGGTGAAATCCCAGGGCTCAACCTTGGAACTGCCTTTGAAACTCCCAGTCTTGAGGTCGAGAGAGGTGAGTGGAATTCCGAGTGTAGAGGTGAAATTCGTAGATATTCGGAGGAACACCAGTGGCGAAGGCGGCTCACTGGCTCGATACTGACGCTGAGGTGCGAAAGCGTGGGG |
| P-asv58 | CCAAGCGTTAATCGGAATTACTGGGCGTAAAGCGTGCGCAGGCGGTTGTGCAAGACCGATGTGAAATCCCCGAGCTTAACTTGGGAATTGCATTGGTGACTGCACGGCTAGAGTGTGTCAGAGGGGGGTAGAATTCCACGTGTAGCAGTGAAATGCGTAGAGATGTGGAAGAATACCGATGGCGAAGGCAGCCCCCTGGGATAACACTGACGCTCATGCACGAAAGCGTGGGG |
| P-asv61 | CCAAGCGTTATCCGGAATCATTGGGTTTAAAGGGTCCGTAGGCGGCCTTATAAGTCAGTGGTGAAATCTCCTAGCTCAACTAGGAAACTGCCATTGATACTGTAGGGCTTGAATTTTTGTGAAGTAACTAGAATATGTAGTGTAGCGGTGAAATGCTTAGATATTACATGGAATACCAATTGCGAAGGCAGGTTACTAACAAACGATTGACGCTGATGGACGAAAGCGTGGGG |
| P-asv67 | GCAAGCGTTAATCGGAATTACTGGGCGTAAAGCGTGCGCAGGCGGTTATATAAGACAGATGTGAAATCCCCGGGCTCAACCTGGGAACTGCATTTGTGACTGTATAGCTAGAGTACGGCAGAGGGGGATGGAATTCCGCGTGTAGCAGTGAAATGCGTAGATATGCGGAGGAACACCGATGGCGAAGGCAATCCCCTGGGCCTGTACTGACGCTCATGCACGAAAGCGTGGGG |
| P-asv68 | TCGAGTGATGGCCGATCTTATTGGGCCTAAAGCGTCCGTAGCTGGCCTGGCAAGTCCGTCGGGAAATCCACCCGCTCAACGGGTGGGCGACCGACGGAAACTGCGAGGCTTGGGACCGGGAGGCTTGAAGGGTACGTCCGGGGTAGGAGTGAAATCCCGTAATCCTGGACGGACCACCGATGGGGAAACCACTTCAAGAGAACGGATCCGACAGTGAGGGACGAAAGCTAGGGT |
| P-asv69 | GTTAGCGTTGTTCGGAATTACTGGGCGTAAAGCGCACGTAGGCGGATCAGAAAGTTGGGGGTGAAATCCCAGGGCTCAACCCTGGAACGGCCTCCAAAACTCCTGGTCTTGAGTTCGAGAGAGGTGAGTGGAATTCCGAGTGTAGAGGTGAAATTCGTAGATATTCGGAGGAACACCAGTGGCGAAGGCGGCTCACTGGCTCGATACTGACGCTGAGGTGCGAAAGTGTGGGG |
| P-asv70 | GCGAGCGTTAATCGGATTTACTGGGCGTAAAGCGTGCGTAGGCGGCTTTTTAAGTCGGATGTGAAATCCCTGAGCTTAACTTAGGAATTGCATTCGATACTGGGAAGCTAGAGTATGGGAGAGGATGGTAGAATTCCAGGTGTAGCGGTGAAATGCGTAGAGATCTGGAAGAATACCGATGGCGAAGGCAGCCATCTGGCCTAATACTGACGCTGAGGTACGAAAGCATGGGG |

| | |
|---|---|
| P-asv71 | GCAAGCGTTAATCGGAATTACTGGGCGTAAAGCGTGCGCAGGCGGTTTTGTAAGTCTGATGTGAAATCCCCGGGCTCAACCTGGGAACTGCATTGGAGACTGCAAGGCTAGAGTGTGTCAGAGGGGGGTAGAAT<br>TCCACGTGTAGCAGTGAAATGCGTAGAGATGTGGAGGAATACCGATGGCGAAGGCAGCCACCTGGGATAACACTGACGCTCATGCACGAAAGCGTGGGG |
| P-asv74 | GCAAGCGTTAATCGGAATTACTGGGCGTAAAGCGTGCGCAGGCGGTTTTGTAAGTCTGATGTGAAATCCCCGGGCTCAACCTGGGAACTGCATTGGAGACTGCAAGGCTAGAGTGTGTCAGAGGGGGGTAGAAT<br>TCCACGTGTAGCAGTGAAATGCGTAGAGATGTGGAGGAATACCGATGGCGAAGGCAGCACCCTGGGATAACACTGACGCTCATGCACGAAAGCGTGGGG |
| P-asv75 | TCGAGTGATGGCCGATCTTATTGGGCCTAAAGCGTCCGTAGCTGGCCTGGCAAGTCCGTCGGGAAATCCACCCGCTCAACGGGTGGGCGGCCGACGGAAACTGCGAGGCTTGGGACCGGGAGGCTTGAAGGG<br>TACGTCCGGGGTAGGAGTGAAATCCCGTAATCCTGGACGGACCACCGATGGGGAAACCACTTCAAGAGAACGGATCCGACAGTGAGGGACGAAAGCTAGGGT |
| P-asv76 | TCAAGTGATGTCCCATATTATTGGGCCTAAAGCGTCCGTAGCTGGCCGGGCAAGTCCATCGGGAAATCCACCTGCCCAACAGGTGGGCGACCGGTGGAAACTGCCTGGCTTGGGATCGGAAGGCTCAGAGAGTA<br>CGTCCGGGGTAGGAGTGAAATCCCGTAATCCCGGACGGACTACCGATGGCGAAAGCACTCTGAGAAGACGACTCCGACAGTGAGGGACGAAAGCTCGGGT |
| P-asv78 | GCTAGCGTTGTTCGGAATTACTGGGCGTAAAGCGCACGTAGGCGGCGATTTAAGTCAGAGGTGAAAGCCCGGGGCTCAACCCCGGAACTGCCTTTGAGACTGGATTGCTAGAATCTTGGAGAGGCGAGTGGAAT<br>TCCGAGTGTAGAGGTGAAATTCGTAGATATTCGGAAGAACACCAGTGGCGAAGGCGGCTCGCTGGACAAGTATTGACGCTGAGGTGCGAAAGCGTGGGG |
| P-asv85 | TCGAGTGATGGCCGATCTTATTGGGCCTAAAGCGTCCGTAGCTGGCCTGGCAAGTCCGTCGGGAAATCCACCCGCTCAACGGGTGGGCGTCCGACGGAAACTGCCGGGCTTGGGACCGGGAGGCTTGAAGGG<br>TACGTCCGGGGTAGGAGTGAAATCCCGTAATCCTGGACGGACCACCGATGGGGAAACCACTTCAAGAGAACGGATCCGACAGTGAGGGACGAAAGCTAGGGT |
| P-asv86 | GCAAGCGTTAATCGGAATTACTGGGCGTAAAGCGCACGCAGGCGGTTGGATAAGTTAGATGTGAAAGCCCCGGGCTCAACCTGGGAATTGCATTTAAAACTGTCCAGCTAGAGTCTTGTAGAGGGGGGTAGAATT<br>CCAGGTGTAGCGGTGAAATGCGTAGAGATCTGGAGGAATACCGGTGGCGAAGGCGGCCCCCTGGACAAAGACTGACGCTCAGGTGCGAAAGCGTGGGG |
| P-asv88 | GCTAGCGTTGTTCGGAATTACTGGGCGTAAAGCGCGCGTAGGCGGTTACTCAAGTCAGAGGTGAAAGCCCGGGGCTCAACCCCGGAACTGCCTTTGAAACTAGGTGACTAGAATCTTGGAGAGGTCAGTGGAAT<br>TCCGAGTGTAGAGGTGAAATTCGTAGATATTCGGAAGAACACCAGTGGCGAAGGCGACTGACTGGACAAGTATTGACGCTGAGGTGCGAAAGCGTGGGG |
| P-asv89 | GCAAGCGTTAATCGGAATTACTGGGCGTAAAGCGCACGCAGGCGGCTTTTTAAGTCGGATGTGAAAGCCCCGGGCTCAACCTGGGAATTGCATTCGATACTGTAAAGCTAGAGTATGTGAGAGGGGGGTAGAAT<br>TCCAAGTGTAGCGGTGAAATGCGTAGAGATTTGGAGGAATACCAGTGGCGAAGGCGGCCCCCTGGCACAATACTGACGCTCAGGTGCGAAAGCGTGGGG |
| P-asv91 | GCTAGCGTTGTTCGGAATTACTGGGCGTAAAGCGCACGTAGGCGGCCATTCAAGTCAGAGGTGAAAGCCCGGGGCTCAACCCCGGAACTGCCTTTGAAACTAGATGGCTTGAATCTTGGAGAGGCGAGTGGAAT<br>TCCGAGTGTAGAGGTGAAATTCGTAGATATTCGGAAGAACACCAGTGGCGAAGGCGACTCGCTGGACAAGTATTGACGCTGAGGTGCGAAAGCGTGGGG |
| P-asv92 | GCAAGCGTTAATCGGAATTACTGGGCGTAAAGCGTGCGCAGGCGGTCCGCTAAGACAGATGTGAAATCCCCGGGCTTAACCTGGGAACTGCATTTGTGACTGGCGGGCTAGAGTATGGCAGAGGGGGGTAGAA<br>TTCCACGTGTAGCAGTGAAATGCGTAGAGATGTGGAGGAATACCGATGGCGAAGGCAGCCCCCTGGGCCAATACTGACGCTCATGCACGAAAGCGTGGGG |
| P-asv94 | GCAAGCGTTAATCGGAATTACTGGGCGTAAAGCGCGCGTAGGTGGTTTGTTAAGTTGGATGTGAAAGCCCCGGGCTCAACCTGGGAACTGCATCCAAAACTGGCAAGCTAGAGTATGGCAGAGGGTGGTGGAAT<br>TTCCTGTGTAGCGGTGAAATGCGTAGATATAGGAAGGAACACCAGTGGCGAAGGCGACCACCTGGGCTAATACTGACACTGAGGTGCGAAAGCGTGGGG |
| P-asv98 | TCAAGTGATGACCGATTTTATTGGGCCTAAAGCGTCCGTAGCTGGCCGCGCAAGTCCGTCGGGAAATCCACCCGCTTAACGGGTGGGCGTCCGGCGGAAACTGTGCGGCTTGGGACAGGAAGGCTCGAGGGGT<br>ACGTTCGGGGTAGGAGTGAAATCCCGTAATCCTGAACGGACCACCGATGGCGAAAGCACCTCGAGAGGACTGATCCGACAGTGAGGGACGAAAGCTAGGGT |
| P-asv99 | TCGAGTGATGTCCCATATTATTGGGCCTAAAGCGTCCGTAGCTGGCCAGGCAAGTCTATCGGGAAATCCACCTGCTCAACAGGTGGGCGTCCGGTAGAAACTGCCTGGCTTGGAACCGGAAGGCTCAGAGAGTA<br>CGTCCGGGGTAGGAGTGAAATCCCGTAATCCCGGACGGACTACCGATGGCGAAAGCACTCTGAGAAGACGGCTTCGACAGTGAGGGACGAAAGCTAGGGT |
| P-asv100 | ACGAGTGATGGCCGATATTATTGGGCCTAAAGCGTCCGTAGCTTGCTGTGTAAGTCCATTGGGAAATCCACGCGCTCAACGCGTGGGCGTCCGGTGGAAACTACACGGCTTGGGGCCGAGAGACTCGACGGGT<br>ACGTCCGGGGTAGGAGTGAAATCCTGTAATCCTGGACGGACCACCAATGGGGAAACCACGTCGAGAGACCGGACCCGACAGTGAGGGACGAAAGCCAGGGT |
| P-asv104 | GCAAGCGTTAATCGGAATTACTGGGCGTAAAGCGTGCGCAGGCGGTGATGTAAGACAGTTGTGAAATCCCCGGGCTCAACCTGGGAACTGCATCTGTGACTGCATCGCTGGAGTACGGCAGAGGGGGATGGAA<br>TTCCGCGTGTAGCAGTGAAATGCGTAGATATGCGGAGGAACACCGATGGCGAAGGCAATCCCCTGGGCCTGTACTGACGCTCATGCACGAAAGCGTGGGG |
| P-asv105 | GCTAGCGTTGCTCGGAATTACTGGGCGTAAAGGGAGCGTAGGCGGACATTTAAGTCAGGGGTGAAATCCCGGGGCTCAACCTCGGAATTGCCTTTGATACTGGGTGTCTTGAGTATGAGAGAGGTGTGTGGAAC<br>TCCGAGTGTAGAGGTGAAATTCGTAGATATTCGGAAGAACACCAGTGGCGAAGGCGACACACTGGCTCATTACTGACGCTGAGGCTCGAAAGCGTGGGG |
| P-asv109 | GCTAGCGTTGTTCGGAATTACTGGGCGTAAAGCGCACGTAGGCGGCTTTTTAAGTCAGGGGTGAAATCCCGGGGCTCAACCCCGGAACTGCCCTTGAAACTGGGAAGCTAGAATCTTGGAGAGGCGAGTGGAAT<br>TCCGAGTGTAGAGGTGAAATTCGTAGATATTCGGAAGAACACCAGTGGCGAAGGCGACTCGCTGGACAAGTATTGACGCTGAGGTGCGAAAGCGTGGGG |
| P-asv110 | TCGAGTGATGGCCGATCTTATTGGGCCTAAAGCGTCCGTAGCTGGCCGTGCAAGTCCGTCGGGAAATCCACCTGCTCAACGGGTGGGCGGCCGACGGAAACTGGACGGCTTGGGACCGGGAGGCTTGAAGGG<br>TACGTCCGGGGTAGGAGTGAAATCCCGTAATCCTGGACGGACCACCGATGGGGAAACCACTTCAAGAGAACGGATCCGACAGTGAGGGACGAAAGCTAGGGT |
| P-asv111 | GCAAGCGTTAATCGGAATTACTGGGCGTAAAGCGTGCGCAGGCGGTTTTGTAAGTCTGATGTGAAATCCCCGGGCTCAACCTGGGAACTGCATTGGAGACTGCAAGGCTAGAGTGTGTCAGAGGGGGGTAGAAT<br>TCCACGTGTAGCAGTGAAATGCGTAGAGATGTGGAAGAATACCGATGGCGAAGGCAGCCCCCTGGGATAACACTGACGCTCATGCACGAAAGCGTGGGG |
| P-asv115 | TCGAGTGATGGCCGATCTTATTGGGCCTAAAGCGTCCGTAGCTGGCCGTGCAAGTCCGTCGGGAAATCCACCCGCTCAACGGGTGGGCGGCCGACGGAAACTGGACGGCTTGGGACCGGGAGGCTTGAGGGG<br>TACGTCCGGGGTAGGAGTGAAATCCTGTAATCCTGGACGGACCACCGATGGGGAAACCACCTCAAGAGAACGGATCCGACAGTGAGGGACGAAAGCTAGGGT |
| P-asv116 | GCAAGCGTTAATCGGAATCACTGGGCGTAAAGCGCTCGTAGGTGGCTTTGTAAGTCAGGTGTGAAAGCCCACGGCTCAACCGTGGAATTGCACTTGATACTGCATTGCTTGAGTTCGGGAGAGGGTAGTGGAATT<br>CCAGGTGTAGGAGTGAAATCCGTAGAGATCTGGAGGAACACCAGTGGCGAAGGCGACTACCTGGACCGATACTGACACTGAGGAGCGAAAGCGTGGGG |
| P-asv118 | TCGAGTGATGGCCGATCTTATTGGGCCTAAAGCGTCCGTAGCTGGCCGTGCAAGTCCGTCGGGAAATCCACCCGCTCAACGGGTGGGCGGCCGACGGAAACTGGACGGCTTGGGACCGGGAGGCTTGAGGGG<br>TACGTCCGGGGTAGGAGTGAAATCCCGTAATCCTGGACGGACCACCGATGGGGAAACCACCTCAAGAGAACGGATCCGACAGTGAGGGACGAAAGCTAGGGT |
| P-asv119 | GCTAGCGTTGCTCGGAATTACTGGGCGTAAAGGGAGCGTAGGCGGACAGTTAAGTTAGAGGTGAAAGCCCAGGGCTCAACCTTGGAATTGCCTTTGATACTGGCTGTCTTGAGTATGGGAGAGGTGTGTGGAAC<br>TCCGAGTGTAGAGGTGAAATTCGTAGATATTCGGAAGAACACCAGTGGCGAAGGCGACACACTGGCCCATTACTGACGCTGAGGCTCGAAAGCGTGGGG |
| P-asv123 | TCGAGTGATGTCCCATATTATTGGGCCTAAAGCGTCCGTAGCTGGCCAGGCAAGTCTATCGGGAAATCCACCCGCCCAACGGGTGGGCGTCCGGTAGAAACTGCCCGGCTTGGAACCGGAAGGCTCAGAGAGTA<br>CGTCCGGGGTAGGAGTGAAATCCCGTAATCCCGGACGGACTACCGATGGCGAAAGCACTCTGAGAAGACGGCTTCGACAGTGAGGGACGAAAGCTAGGGT |
| P-asv124 | GCGAGCGTTAATCGGAATTACTGGGCGTAAAGCGTGCGCAGGCGGTTTTTTAAGATAGGCGTGAAATCCCCGGGCTCAACCTGGGAACTGCGCTTATGACTGGAAGACTCGAGTATGGCAGAGGGGGGTGGAAT<br>TCCACGTGTAGCAGTGAAATGCGTAGAGATGTGGAGGAACACCGATGGCGAAGGCAGCCCCCTGGGCCAATACTGACGCTCATGCACGAAAGCGTGGGG |
| P-asv126 | GCTAGCGTTGTTCGGAATTACTGGGCGTAAAGCGCACGTAGGCGGCTTTTTAAGTCAGGGGTGAAATCCCGGGGCTCAACCCCGGAACTGCCTTTGAAACTGGAGAGCTAGAATCTTGGAGAGGCGAGTGGAAT<br>TCCGAGTGTAGAGGTGAAATTCGTAGATATTCGGAAGAACACCAGTGGCGAAGGCGACTCGCTGGACAAGTATTGACGCTGAGGTGCGAAAGCGTGGGG |
| P-asv130 | GCAAGCGTTACTCGGAATCACTGGGCGTAAAGAGCGTGTAGGCGGGTAGTTAAGTCAGAAGTGAAATCCAATAGCTCAACTATTGAACTGCTTTTGAAACTGATTACCTAGAATATGGGAGAGGTAGATGGAATTT<br>CTGGTGTAGGGGTAAAATCCGTAGAGATCAGAAGGAATACCGATTGCGAAGGCGATCTACTGGAACATTATTGACGCTGAGACGCGAAAGCGTGGGG |
| P-asv131 | GCAAGCGTTACTCGGATTTACTGGGCGTAAAGCGCGCGCAGGCGGATCGGCAAGTCAGGTGTGAAATCTCGGGGCTCAACTCCGAAACTGCGCCTGAAACTATCGATCTTGAGTCTTGGCGGGGGAGGTGGAA<br>TATCGCATGTAGGGGTAAAATCCGTAGAGATGCGATGGAACACCAGAGGCGAAGGCGACCTCCTGGACAAGTACTGACGCTGAGGCGCGAAAGCGTGGGG |
| P-asv134 | GCAAGCGTTAATCGGAATTACTGGGCGTAAAGCGCGCGTAGGTGGTTTGTTAAGTTGAATGTGAAAGCCCCGGGCTCAACCTGGGAACTGCATCCAAAACTGGCAAGCTAGAGTATGGCAGAGGGTGGTGGAAT<br>TTCCTGTGTAGCGGTGAAATGCGTAGATATAGGAAGGAACATCAGTGGCGAAGGCGACCACCTGGGCTAATACTGACACTGAGGTGCGAAAGCGTGGGG |
| P-asv135 | GCAAGCGTTGTCCGGAATTATTGGGCGTAAAGCGCGCGCAGGCGGCATTGTAAGTCTTGTGTTTAAACGCGGGGCTCAACCCCGTGTCGCATGGGAAACTGCAAGGCTTGAGTGCAGAAGAGGAAAGTGGAATT<br>CCACGTGTAGCGGTGAAATGCGTAGAGATGTGGAGGAACACCAGTGGCGAAGGCGACTTTCTGGGCTGTAACTGACGCTGAGGCGCGAAAGCGTGGGG |
| P-asv146 | ACGAGTGATGGCCGATTTTATTGGGCCTAAAGCGTCCGTAGCTGGCCGAGCAAGTCTGTCGGGAAATCCAGTGGCTCAACCACTGGGCTGCCGGCGGAAACTGTTCGGCTTGGGACCGGAAGATCCGAGGGGT<br>ACGTTCGGGGTAGGAGTGAAATCCCGTAATCCTGAACGGACCACCGATGGCGAAAGCACCTCGGAAGCACGGATCCGACAGTGAGGGACGAAAGCTTGGGT |
| P-asv147 | GCAAGCGTTAATCGGAATTACTGGGCGTAAAGCGCGCGTAGGTGGTTTGTTAAGTTGAATGTGAAAGCCCCGGGCTCAACCTGGGAACTGCATCCAAAACTGGCAAGCTAGAGTATGGCAGAGGGTGGTGGAAT<br>TTCCTGTGTAGCGGTGAAATGCGTAGATATAGGAAAGAACACCAGTGGCGAAGGCGACCACCTGGGCTAATACTGACACTGAGGTGCGAAAGCGTGGGG |
| P-asv149 | GCGAGCGTTGTTCGGAATCACTGGGCGTAAAGCGTGCGTAGGCGGTTATACAAGTCAGAAGTGAAAGCCCAGGGCTTAACCCTGGAATTGCTTTTGAGACTGTATGGCTAGAGATCTAGAGAGGTTGGGGAAT<br>TCCGAGTGTAGAGGTGAAATTCGCAGATATTCGGAGGAACACCAGTGGCGTAGGCGCCCAACTGGCTAGATACTGACGCTGAGGCACGAAAGCGTGGGG |
| P-asv150 | TCGAGTGATGACCGATTTTATTGGGCCTAAAGCGTCCGTAGCTGGCCGCGCAAGTCCGTCGGGAAATCCACCCGCTTAACGGGTGGGCGTCCGGCGGAAACTGTGCGGCTTGGGACAGGAAGGCTCGAGGGGT<br>ACGTTCGGGGTAGGAGTGAAATCCCGTAATCCTGAACGGACCACCGATGGCGAAAGCACCTCGAGAGGACTGATCCGACAGTGAGGGACGAAAGCTGGGGT |
| P-asv153 | GCAAGCGTTAATCGGAATTACTGGGCGTAAAGCGCGCGTAGGTGGTTCAGTAAGTTGGAAGTGAAATCCCCGGGCTCAACCTGGGAACTGCTTTCAAAACTGCTGAGCTAGAGTACGGTAGAGGGTGGTGGAAT<br>TTCCTGTGTAGCGGTGAAATGCGTAGATATAGGAAGGAACACCAGTGGCGAAGGCGACCACCTGGACTGATACTGACACTGAGGTGCGAAAGCGTGGGG |
| P-asv154 | GCAAGCGTTAATCGGAATTACTGGGCGTAAAGCGTGCGCAGGCGGTTTTGTAAGACAGGCGTGAAATCCCCGGGCTCAACCTGGGAATTGCGCTTGTGACTGCAAAGCTGGAGTGCGGCAGAGGGGGATGGAA<br>TTCCGCGTGTAGCAGTGAAATGCGTAGATATGCGGAGGAACACCGATGGCGAAGGCAATCCCCTGGGCCTGCACTGACGCTCATGCACGAAAGCGTGGGG |
| P-asv155 | GCGAGCGTTAATCGGAATTACTGGGCGTAAAGCGTGCGCAGGCGGTTTTGTAAGACAGATGTGAAATCCCCGGGCTTAACCTGGGAACTGCGTTTGTGACTGCAAGGCTAGAGTACGGCAGAGGGGGGTGGAA<br>TTCCTGGTGTAGCAGTGAAATGCGTAGATATCAGGAGGAACACCGATGGCGAAGGCAGCCCCCTGGGCCTGTACTGACGCTCATGCACGAAAGCGTGGGG |
| P-asv157 | GCGAGCGTTAATCGGAATTACTGGGCGTAAAGCGTGCGCAGGCGGCTTTGCAAGACAGATGTGAAATCCCCGGGCTCAACCTGGGAACTGCATTTGTGACTGCAAGGCTAGAGTACGGTAGAGGGGGATGGAA<br>TTCCGCGTGTAGCAGTGAAATGCGTAGATATGCGGAGGAACACCAATGGCGAAGGCAATCCCCTGGACCTGTACTGACGCTCATGCACGAAAGCGTGGGG |

| | |
|---|---|
| P-asv159 | GCAAGCGTTAATCGGAATTACTGGGCGTAAAGCGCACGTAGGCGGTGTGTTAAGTTGGATGTGAAAGCCCCGGGCTCAACCTGGGAATTGCATTCAAAACTGGCACGCTAGAGTATGTGAGAGGGGGGTAGAATTCCAAGTGTAGCGGTGAAATGCGTAGAGATTTGGAGGAATACCAGTGGCGAAGGCGGCCCCCTGGCACAATACTGACGCTCAGGTGCGAAAGCGTGGGG |
| P-asv160 | ACGAGTGATGACCGATATTATTGGGCCTAAAGCGTCCGTAGCCGGCCAAACAAGTCCGTTGGGAAATCCACGCGCTCAACGCGTGGGCGTCCGGCGGAAACTGTTTGGCTTGGGGCCGGAAGACCTGAGGGGTACGTCCGGGGTAGGAGTGAAATCCCGTAATCCTGGACGGACCACCGTTGGCGAAAGCGCCTCAGGAAGACGGACCCGACGGTGAGGGACGAAAGCTAGGGT |
| P-asv161 | GCAAGCGTTAATCGGAATTACTGGGCGTAAAGCGCACGTAGGCGGTATGTTAAGTTGGATGTGAAAGCCCCGGGCTCAACCTGGGAATTGCATTCAAAACTGGCATGCTAGAGTATGTGAGAGGGGGGTAGAATTCCAAGTGTAGCGGTGAAATGCGTAGAGATTTGGAGGAATACCAGTGGCGAAGGCGGCCCCCTGGCACAATACTGACGCTGAGGTGCGAAAGCGTGGGG |
| P-asv162 | CCGAGCGTTGTCCGGATTTATTGGGCGTAAAGCGAGCGCAGGCGGTTAGATAAGTCTGAAGTTAAAGGCTGTGGCTTAACCATAGTACGCTTTGGAAACTGTTTAACTTGAGTGCAAGAGGGGAGAGTGGAATTCCATGTGTAGCGGTGAAATGCGTAGATATATGGAGGAACACCGGTGGCGAAAGCGGCTCTCTGGCTTGTAACTGACGCTGAGGCTCGAAAGCGTGGGG |
| P-asv164 | GCAAGCGTTAATCGGAATTACTGGGCGTAAAGCGCACGTAGGTGGTTTGTTAAGCTAGCTGTGAAATCCCCGGGCTCAACCTGGGCACTGCAGTTAGAACTGGCAAGCTAGAGTAGGGTAGAGGGGTGTGGAATTCCAGGTGTAGCGGTGAAATGCGTAGATATCTGGAGGAACATCAGTGGCGAAGGCGACACCCTGGACTCATACTGACACTGAGGTGCGAAAGCGTGGGG |
| P-asv165 | CCGAGTGGTAGTCGTTATTATTGGGCTTAAAGTGTTCGTAGCCTGCCATGCGAGTCCGCTGTGAAATCCACGCGCTTAACGTGTGGGCGTGCAGCGGATACTGTATGGCTAGGGACCGGACGGCGTTGAGGCTATTCCCAGGGTAGGAGTGAAATCCTAAAATCCTGGGAGGAGCACCTGTGGCGAAAGCGCTCAACGAGTACGGATCCGACGGTGAGGAACGAAAGCCAGGGG |
| P-asv166 | CCGAGTGATGGCCGATATTATTGGGCCTAAAGCGTCCGTAGCTGGCTAGGCAAGTCCGTTGGGAAATCTGCTCGCTTAACGAGCAGGCGTCCAGCGGAAACTGTGTAGCTTGGGACCGGAAGACCTGAGGGGTACGTCTGGGGTAGGAGTGAAATCCCGTAATCCTGGACGGACCGCCGGTGGCGAAAGCGCCTCAGGAGAACGGATCCGACAGTGAGGGACGAAAGCTAGGGT |
| P-asv167 | TCGAGTGATGGCCGATCTTATTGGGCCTAAAGCGTCCGTAGCTGGCCGTGCAAGTCCGTCGGGAAATCCACCCGCTCAACGGGTGGGCGGCCGACGGAAACTGGACGGCTTGGGACCGGGAGGCTTGAAGGGTACGTCCGGGGTAGGAGTGAAATCCCGTAATCCTGGACAGACCACCGATGGGGAAACCACTTCAAGAGAACGGATCCGACAGTGAGGGACGAAAGCTAGGGT |
| P-asv170 | ACGAGTGATGGCCGATATTATTGGGCCTAAAGCGTCCGTAGCTTGCTGTGTAAGTCCATTGGGAAATCCACGCGCTCAACGCGTGGGCGTCCGGTGGAAACTACACGGCTTGGGGCCGAGAGACTCGACGGGTACGTCCGGGGTAGGAGTGAAATCCCGTAATCCTGGACGGACCACCAATGGGGAAACCACGTCGAGAGACCGGACCCGACAGTGAGGGACGAAAGCCAGGGT |
| P-asv171 | CCGAGTGATGGCCGATATTATTGGGCCTAAAGCGTTCGTAGCTGGCCGGGCAAGTCCGTTGGGAAATCCGCGCGCTTAACGCGCGGGCGTCCAGCGGAAACTGTTCGGCTTGGGACCGGAAGACCTGAGGGGTACGTCCGGGGTAGGAGTGAAATCCCGTAATCCTGGACGGACCACCGGTGGCGAAAGCGCCTCAGGAGGACGGATCCGACAGTGAGGAACGAAAGCTAGGGT |
| P-asv172 | TCAAGTGATGACCGATTTTATTGGGCCTAAAGCGTCCGTAGCTGGCCGCGCAAGTCCGTCGGGAAATCCACCCGCTTAACGGGTGGGCGTCCGGCGGAAACTGTGCGGCTTGGGACAGGAAGGCTCGAGGGGTACGTTCGGGGTAGGAGTGAAATCCCGTAATCCTGAACGGACCACCGATGGCGAAAGCACCTCGAGAGGACTGATCCGACAGTGAGGGACGAAAGCTGGGGT |
| P-asv176 | GTTAGCGTTGTTCGGAATTACTGGGCGTAAAGCGCGCGTAGGCGGATCAGAAAGTTGGGGGTGAAATCCCAGGGCTCAACCCTGGAACGGCCTCCAAAACTCCTGGTCTTGAGTTCGAGAGAGGTGAGTGGAATTCCGAGTGTAGAGGTGAAATTCGTAGATATTCGGAGGAACACCAGTGGCGAAGGCGGCTCACTGGCTCGATACTGACGCTGAGGTGCGAAAGTGTGGGG |
| P-asv177 | GCAAGCGTTACTCGGAATCACTGGGCGTAAAGAGCATGTAGGCGGGTAAATAAGTTGGAAGTGAAATCCTATAGCTTAACTATAGAACTGCTTCCAAAACTGTTTACCTAGAGTATGGGAGAGGTAGATGGAATTTCTGGTGTAGGGGTAAAATCCGTAGAGATCAGAAGGAATACCGATTGCGAAGGCGATCTACTGGAACATTACTGACGCTCAGATGCGAAAGCGTGGGG |
| P-asv181 | TCGAGTGATGTCCAATATTATTGGGCCTAAAGCGTCCGTAGCTGGCCGCGCAAGTCCGTCGGGAAATCCACCTGCCCAACAGGTGGGCGTCCGGCGGAAACTGTGTGGCTTGGAACCGGAAGGCTCAGAGGGTACGTCCGGGGTAGGAGTGAAATCCCGTAATCCCGGACGGACCACCGATGGCGAAAGCACTCTGAGAAGACGGCTTCGACAGTGAGGGACGAAAGCTAGGGT |
| P-asv182 | TCGAGTGATGTCCCATATTATTGGGCCTAAAGCGTCCGTAGCTGGCCAGGCAAGTCTATCGGGAAATCCACCTGCCCAACAGGTGGGCGTCCGGTAGAAACTGCCCGGCTTGGAACCGGAAGGCTCAGAGAGTACGTCCGGGGTAGGAGTGAAATCCCGTAATCCCGGACAGACTACCGATGGCGAAAGCACTCTGAGAAGACGGCTTCGACAGTGAGGGACGAAAGCTAGGGT |
| P-asv189 | GCAAGCGTTAATCGGAATTACTGGGCGTAAAGCGCACGTAGGCGGTTTTTTAAGTCAGATGTGAAAGCCCCGGGCTCAACCTGGGAATTGCATTTGAAACTGGAAAACTAGAGTGTGTGAGAGGGGGGTAGAATTCCAAGTGTAGCGGTGAAATGCGTAGAGATTTGGAGGAATACCAGTGGCGAAGGCGGCCCCCTGGCACAACACTGACGCTCAGGTGCGAAAGCGTGGGG |
| P-asv192 | TCAAGTGATGTCCCATATTATTGGGCCTAAAGCGTCCGTAGCTGGCCGGGCAAGTCCATCGGGAAATCCACCTGCTCAACAGGTGGGCGACCGGTGGAAACTGCCTGGCTTGGGATCGGAAGGCTCAGAGAGTACGTCCGGGGTAGGAGTGAAATCCCGTAATCCCGGACGGACTACCGATGGCGAAAGCACTCTGAGAAGACGACTCCGACAGTGAGGGACGAAAGCTCGGGT |
| P-asv196 | GCAAGCGTTAATCGGAATTATTGGGCGTAAAGCGAGTGCAGACGGTTACTTAAGCCAGATGTGAAATCCCCAAGCTTAACTTGGGACGTGCATTTGGAACTGGGTGACTAGAGTGTGTCAGAGGGAGGTAGAATTCCACATGTAGCGGTGGAATGCGTAGAGATGTGGAAGAATACCGATGGCGAAGGCAGCCTCCTGGGATAACACTGACGTTGAGGCTCGAAAGCGTGGGG |
| P-asv197 | GCGAGCGTTAATCGGAATTACTGGGCGTAAAGCGTGCGCAGGCGGTTGTGTAAGACAGGTGTGAAATCCCCGGGCTTAACCTGGGAACTGCGCTTGTGACTGCACAGCTAGAGTACGGCAGAGGGGGGTGGAATTCCACGTGTAGCAGTGAAATGCGTAGAGATGTGGAGGAACACCGATGGCGAAGGCAGCCCCCTGGGCCTGTACTGACGCTCATGCACGAAAGCGTGGGG |
| P-asv198 | GCAAGCGTTAATCGGATTTACTGGGCGTAAAGCGCGCGTAGGTGGCCAATTAAGTCAAATGTGAAATCCCCGAGCTTAACTTGGGAATTGCATTCGATACTGGTTGGCTAGAGTATGGGAGAGGATGGTAGAATTCCAGGTGTAGCGGTGAAATGCGTAGAGATCTGGAGGAATACCGATGGCGAAGGCAGCCATCTGGCCTAATACTGACACTGAGGTGCGAAAGCATGGGG |
| P-asv199 | GCAAGCGTTAATCGGAATTACTGGGCGTAAAGCGCACGTAGGTGGTTTGTTAAGCCAGCTGTGAAATCCCCGGGCTCAACCTGGGCACTGCAGTTGGAACTGGCAAGCTAGAGTAGGGTAGAGGGGTGTGGAATTCCAGGTGTAGCGGTGAAATGCGTAGATATCTGGAGGAACATCAGTGGCGAAGGCGACACCCTGGACTCATACTGACACTGAGGTGCGAAAGCGTGGGG |
| P-asv200 | TCGAGTGATGGCCGATCTTATTGGGCCTAAAGCGTCCGTAGCTGGCCGTGCAAGTCCGTCGGGAAATCCACCCGCTCAACGGGTGGGCGGCCGACGGAAACTGGACGGCTTGGGACCGGGAGGCTTGAAGGGTACGTCCGGGGTAGGAGTGAAATCCTGTAATCCTGGACGGACCACCGATGGGGAAACCACTTCAAGAGAACGGATCCGACAGTGAGGGACGAAAGCTAGGGT |
| P-asv201 | GTTAGCGTTGTTCGGAATTACTGGGCGTAAAGCGCACGTAGGCGGATTGGAAAGTTGGGGGTGAAATCCCAGGGCTCAACCCTGGAACGGCCTCCAAAACTCCCAGTCTAGAGTTCGAGAGAGGTGAGTGGAATTCCGAGTGTAGAGGTGAAATTCGTAGATATTCGGAGGAACACCAGTGGCGAAGGCGGCTCACTGGCTCGATACTGACGCTGAGGTGCGAAAGTGTGGGG |
| P-asv202 | GCAAGTGTTATCCGGAATCACTGGGCGTAAAGCGTCTGTAGGTGGTTTGATAAGTCAACTGTTAAATCTTGAGGCTCAACCTCAAAATCGCGGTCGAAACTATTAGACTAGAGTATAGTAGGGGTAAAGGAATTTCCAGTGGAGCGGTGAAATGCGTAGATATTGGAAGGAACACCGATGGCGAAGGCACTTTACTGGGCTATTACTAACACTCAGAGACGAAAGCTAGGGT |
| P-asv203 | GCGAGCGTTAATCGGAATTACTGGGCGTAAAGCGTGCGCAGGCGGTTGTGTAAGACAGGTGTGAAATCCCCGGGCTTAACCTGGGAACTGCGCTTGTGACTGCACGGCTAGAGTACGGCAGAGGGGGGTGGAATTCCTGGTGTAGCAGTGAAATGCGTAGATATCAGGAGGAACACCGATGGCGAAGGCAGCCCCCTGGGCCTGTACTGACGCTCATGCACGAAAGCGTGGGG |
| P-asv205 | TCAAGTGATGTCCCATATTATTGGGCCTAAAGCGTCCGTAGCTGGCCGACCAAGTCTATCGGGAAATCCACCTGCCCAACAGGTGGGCGTCCGGTAGAAACTGGCCGGCTTGGAACCGGAAGGCTCAGAGAGTACGTCCGGGGTAGGAGTGAAATCCCGTAATCCCGGACGGACTACCGATGGCGAAAGCACTCTGAGAAGACGGATTCGACAGTGAGGGACGAAAGCTCGGGT |
| P-asv207 | TCGAGTGATGTCCCATATTATTGGGCCTAAAGCGTCCGTAGCTGGCCAGGCAAGTCTATCGGGAAATCCACCCGCTCAACGGGTGGGCGTCCGGTAGAAACTGCCCGGCTTGGAACCGGAAGGCTCAGAGAGTACGTCCGGGGTAGGAGTGAAATCCCGTAATCCCGGACGGACTACCGATGGCGAAAGCACTCTGAGAAGACGGCTTCGACAGTGAGGGACGAAAGCTAGGGT |
| P-asv209 | GTTAGCGTTGTTCGGAATTACTGGGCGTAAAGCGCGCGTAGGCGGGCTGTTAAGTCGGGGGTGAAATCCCGGGGCTCAACCCCGGAACGGCCCTCGATACTGGCAGCCTAGAGTTCGAGAGAGGTGAGTGGAATTCCGAGTGTAGAGGTGAAATTCGTAGATATTCGGAGGAACACCAGTGGCGAAGGCGGCTCACTGGCTCGATACTGACGCTGAGGTGCGAAAGCGTGGGG |
| P-asv213 | GCGAGCGTTGTCCGGAATTACTGGGCGTAAAGGGCTCGTAGGTGGTTTGTCGCGTCGTCTGTGAAATTCCGGGGCTTAACTCCGGGCGTGCAGGCGATACGGGCATAACTTGAGTACTGTAGGGGTAACTGGAATTCCTGGTGTAGCGGTGAAATGCGCAGATATCAGGAGGAACACCGATGGCGAAGGCAGGTTACTGGGCAGTTACTGACGCTGAGGAGCGAAAGCATGGGT |
| P-asv215 | GCAAGCGTTACTCGGAATCACTGGGCGTAAAGGACGCGTAGGCTGTTTAGCAAGTCAGATGTGAAATCCAATGGCTCAACCATTGAACTGCATTTGAAACTGCTAGACTAGAGTATGGGAGGGGGAGATGGAATTAGTGGTGTAGGGGTAAAATCCGTAGATATCACTAGGAATACCTAAAGCGAAGGCGATCTCCTGGAACATTACTGACGCTAAGGCGTGAAAGCGTGGGG |
| P-asv216 | GCTAGCGTTGCTCGGAATTACTGGGCGTAAAGGGCGCGTAGGCGGACAGTTTAGTCAGAGGTGAAAGCCCAGGGCTCAACCTTGGAATTGCCTTTGATACTGGCTGTCTTGAGTACGGGAGAGGTGAGTGGAACTCCGAGTGTAGAGGTGAAATTCGTAGATATTCGGAAGAACACCAGTGGCGAAGGCGACTCACTGGCCCGTTACTGACGCTGAGGCGCGAAAGCGTGGGG |
| P-asv219 | TCAAGTGATGTCCCATATTATTGGGCCTAAAGCGTCCGTAGCTGGCCGGGCAAGTCCATCGGGAAATCCACCTGCCCAACAGGTGGGCGACCGGTGGAAACTGCCTGGCTTGGGACCGGAAGGCTCAGAGAGTACGTCCGGGGTAGGAGTGAAATCCCGTAATCCCGGACGGACTACCGATGGCGAAAGCACTCTGAGAAGACGGCTCCGACAGTGAGGGACGAAAGCTCGGGT |
| P-asv220 | ACGAGTGATGGCCGATCTTATTGGGCCTAAAGCGTCCGTAGCCGGCCAGACAAGTCCGTTGGGAAATCGACGTGCCTAACACGTCGGCGTCCAGCGGAAACTGTCTGGCTTGGGGCCGGAAGACTCGAGGGGTACGTCCGGGGTAGGAGTGAAATCCCGTAATCCTGGACGGACCGCCGGTGGCGAAAGCGCCTCGAGAGGACGGACCCGACGGTCAGGGACGAAAGCTAGGGT |
| P-asv221 | TCAAGTGATGTCCCATCTTATTGGGCCTAAAGCGTCCGTAGCTGGCCAGGCAAGTCCATCGGGAAATCCACCTGCTCAACAGGTGGGCGACCGGTGGAAACTGCCCGGCTTGGAACCGGAAGGCTCAGAGAGTACGTCCGGGGTAGGAGTGAAATCCCGTAATCCCGGACGGACTACCGATGGCGAAAGCACTCTGAGAAGACGGATTCGACAGTGAGGGACGAAAGCTCGGGT |
| P-asv223 | GCAAGCGTTACTCGGAATCACTGGGCGTAAAGAGCGTGTAGGCGGATTTGTAAGTTGGAAGTGAAATCCTATGGCTCAACCATAGAACTGCTTCCAAAACTGCTCATCTAGAGTGTGGGAGAGGTAGATGGAATTTCTGGTGTAGGGGTAAAATCCGTAGAGATCAGAAGGAATACCGATTGCGAAGGCGATCTACTGGAACATAACTGACGCTGAGACGCGAAAGCGTGGGG |
| P-asv231 | TCGAGTGATGTCCAATATTATTGGGCCTAAAGCGTCCGTAGCTGGCCACGCAAGTCCGTCGGGAAATCCACCTGCCCAACAGGTGGGCGTCCGGCGGAAACTGTGTGGCTTGGGACCGGAAGGCTCAGAGGGTACGTCCGGGGTAGGAGTGAAATCCCGTAATCCCGGACGGACCACCGATGGCGAAAGCACTCTGAGAAGACGGCTCCGACAGTGAGGGACGAAAGCTAGGGT |
| P-asv233 | GCAAGCGTTAATCGGAATTACTGGGCGTAAAGCGCACGTAGGCGGTTTTTTAAGTCAGATGTGAAAGCCCCGGGCTCAACCTGGGAATTGCATTTGAAACTGGGGAACTAGAGTGTGTGAGAGGGGGGTAGAATTCCAAGTGTAGCGGTGAAATGCGTAGAGATTTGGAGGAATACCAGTGGCGAAGGCGGCCCCCTGGCACAACACTGACGCTCAGGTGCGAAAGCGTGGGG |
| P-asv234 | GCAAGCGTTGTCCGGATTTATTGGGTTTAAAGGGTGCGTAGGCGGGTCTTTAAGTCAGTGGTGAAAGCCTGCAGCTTAACTGTAGAACTGCCATTGATACTGGAGACCTTGAGTGTACTAGAGGTAGGCGGAATTTATGGTGTAGCGGTGAAATGCATAGATACCATAAAGAACACCGATAGCGTAGGCAGCTTACTGGAGTACAACTGACGCTGAGGCACGAAAGCATGGGG |
| P-asv235 | GCAAGCGTTAATCGGAATTACTGGGCGTAAAGCGCATGCAGGTGGTTTGTTAAGTCAGATGTGAAAGCCCTGGGCTCAACCTAGGAATCGCATTTGAAACTGACAAGCTAGAGTACTGTAGAGGGGGGTAGAATTTCAGGTGTAGCGGTGAAATGCGTAGAGATCTGAAGGAATACCGGTGGCGAAGGCGGCCCCCTGGACAGATACTGACACTCAGATGCGAAAGCGTGGGG |

| | |
|---|---|
| P-asv238 | GCTAGCGTTGCTCGGAATCACTGGGCGTAAAGGGCGCGTAGGCGGCTGATTTAGTCGAGGGTGAAAGCCCGTGGCTCAACCACGGAATGGCCTTCGATACTGGTTGGCTTGAGACCGGAAGAGGACAGCGGAA<br>CTGCGAGTGTAGAGGTGAAATTCGTAGATATTCGCAAGAACACCAGTGGCGAAGGCGGCTGTCTGGTCCGGTTCTGACGCTGAGGCGCGAAAGCGTGGGG |
| P-asv240 | GCAAGCGTTATCCGGAATGATTGGGCGTAAAGCGTCTGTAGGTGGTTCTTTAAGTCTACTGTCAAATACGTTGGCTCAACCAACGGCAGGCAGTGGAGTACTAAAGAACTAGAGTGCGGTAGAGGTAGAGGGAAT<br>CCCTAGCGTAACAGTGAAATGTGTAGATTTTAGGGAGAACACCAGCGGCGAAGGCGCTCTACTGGGCCGATACTGACACTGAGAGACGAAAGCTAGAGG |
| P-asv243 | CCAAGCGTTAATCGGAATTACTGGGCGTAAAGCGTGCGCAGGCGGTTGTGCAAGACCGATGTGAAATCCCCGGGCTTAACCTGGGAATTGCATTGGTGACTGCACGGCTAGAGTGTGTCAGAGGGGGGTAGAAT<br>TCCACGTGTAGCAGTGAAATGCGTAGAGATGTGGAGGAATACCGATGGCGAAGGCAGCCCCCTGGGATAACACTGACGCTCATGCACGAAAGCGTGGGG |
| P-asv244 | GCAAGCGTTACTCGGAATCACTGGGCGTAAAGAGCGTGTAGGCGGGTATATAAGTCAGAAGTGAAATCCAATAGCTTAACTATTGAACTGCTTTTGAAACTGTATACCTAGAATGTGGGAGAGGTAGATGGAATTT<br>CTGGTGTAGGGGTAAAATCCGTAGAGATCAGAAGGAATACCGATTGCGAAGGCGATCTACTGGAACATTATTGACGCTGAGACGCGAAAGCGTGGGG |
| P-asv245 | GCAAGCGTTAATCGGAATTACTGGGCGTAAAGCGTGCGCAGGCGGTTATGCAAGACAGATGTGAAATCCCCGGGCTCAACCTGGGAATTGCATTTGTGACTGCATGGCTAGAGTACGGTAGAGGGGGATGGAAT<br>TCCGCGTGTAGCAGTGAAATGCGTAGATATGCGGAGGAACACCGATGGCGAAGGCAATCCCCTGGACCTGTACTGACGCTCATGCACGAAAGCGTGGGG |
| P-asv247 | GCAAGCGTTGTTCGGAATCACTGGGCGTAAAGAATGCGTAGGCGGCTAGACAAGTCAGGAGTGAAATCCCAGGGCTCAACCCTGGAACTGCTTTTGAAACTGTCTTGCTAGAGTCCCGGAGGGGTTGGCGGAAT<br>TCCGAGTGTAGAGGTGAAATTCGCAGATATTCGGAGGAACACCAGTGGCGTAGGCGGCCAACTGGACGGGAACTGACGCTGAGGCATGAAAGTGTGGGG |
| P-asv250 | GCAAGCGTTAATCGGAATTACTGGGCGTAAAGCGTGCGCAGGCGGTGATGTAAGACAGGCGTGAAATCCCCGGGCTCAACCTGGGAATTGCGCTTGTGACTGCATCGCTGGAGTGCGGCAGAGGGGGATGGAA<br>TTCCGCGTGTAGCAGTGAAATGCGTAGATATGCGGAGGAACACCGATGGCGAAGGCAATCCCCTGGGCCTGCACTGACGCTCATGCACGAAAGCGTGGGG |
| P-asv254 | TCAAGTGATGTCCAATATTATTGGGCCTAAAGCGTCCGTAGCTGGCCGCGCAAGTCCGTCGGGAAATCCACCTGCCCAACAGGTGGGCGTCCGGCGGAAACTGTGTGGCTTGGAACCGGAAGGCTCAGAGGGT<br>ACGTCCGGGGTAGGAGTGAAATCCCGTAATCCCGGACAGACCACCGATGGCGAAAGCACTCTGAGAAGACGGCTTCGACAGTGAGGGACGAAAGCTAGGGT |
| P-asv260 | GCAAGCGTTATTCGGAATCACTGGGCGTAAAGGACGCGTAGGCGGGTTTGTAAGTCAGGTGTGAAATCCTACAGCTTAACTGTAGAACTGCACTTGAAACTACTAACCTAGAGTATGGGAGGGGGAGATGGAATT<br>AGTGGTGTAGGGGTAAAATCCGTAGATATCACTAGGAATACCTAAAGCGAAGGCGATCTCCTGGAACATTACTGACGCTAAGGCGTGAAAGCGTGGGG |
| P-asv261 | GCTAGCGTTGTTCGGAATTACTGGGCGTAAAGCGCACGTAGGCGGCTTTGTAAGTTAGAGGTGAAAGCCTGGAGCTCAACTCCAGAACTGCCTTTAAGACTGCATCGCTTGAATCCAGGAGAGGTGAGTGGAATT<br>CCGAGTGTAGAGGTGAAATTCGTAGATATTCGGAAGAACACCAGTGGCGAAGGCGGCTCACTGGACTGGTATTGACGCTGAGGTGCGAAAGCGTGGGG |
| P-asv262 | TCGAGTGATGGCCGATCTTATTGGGCCTAAAGCGTCCGTAGCTGGCCTGGCAAGTCCGTCGGGAAATCCACCCGCTCAACGGGTGGGCGGCCGACGGAAACTGCGAGGCTTGGGACCGAGAGGCTTGAAGGG<br>TACGTCCGGGGTAGGAGTGAAATCCCGTAATCCTGGACGGACCACCGATGGGGAAACCACTTCAAGAGAACGGATCCGACAGTGAGGGACGAAAGCTAGGGT |
| P-asv263 | CCGAGTGATGGCCGAAATTATTGGGCCTAAAGCGTCCGTAGCTGGCCGGACAAGTCCGTTGGGAAATCCGCGCGCTTAACGCGCGGGCGTCCAGCGGAAACTGTTCGGCTTGGGACCGGAAGACCTGAGGGGT<br>ACGTCCGGGGTAGGAGTGAAATCCCGTAATCCTGGACGGACCACCGGTGGCGAAAGCGCCTCAGGAGGACGGATCCGACAGTCAGGGACGAAAGCTAGGGT |
| P-asv264 | GCTAGCGTTATTCGGAATTACTGGGCGTAAAGCGCACGTAGGCGGATCAGAAAGTCAGAGGTGAAATCCCGGGGCTCAACCCCGGAACGGCCTTTGAAACTCCTGATCTTGAGTTCGAGAGAGGTGAGTGGAAT<br>TCCGAGTGTAGAGGTGAAATTCGTAGATATTCGGAGGAACACCAGTGGCGAAGGCGGCTCACTGGCTCGATACTGACGCTGAGGTGCGAAAGCGTGGGG |
| P-asv265 | CCGAGTGATGGCCGATATTATTGGGCCTAAAGCGTCCGTAGCTGGCTAGACAAGTCCGTTGGGAAATCTGCTCGCTTAACGAGCAGGCGTCCAGCGGAAACTGTGTAGCTTGGGACCGGAAGACCTGAGGGGTA<br>CGTCTGGGGTAGGAGTGAAATCCCGTAATCCTGGACGGACCGCCGGTGGCGAAAGCGCCTCAGGAGAACGGATCCGACAGTGAGGGACGAAAGCTAGGGT |
| P-asv270 | TCAAGTGATGTCCAATATTATTGGGCCTAAAGCGTCCGTAGCTGGCCGCGCAAGTCCGTCGGGAAATCCACCTGCTCAACAGGTGGGCGTCCGGCGGAAACTGTGTGGCTTGGAACCGGAAGGCTCAGAGGGTA<br>CGTCCGGGGTAGGAGTGAAATCCCGTAATCCCGGACGGACCACCGATGGCGAAAGCACTCTGAGAAGACGGCTTCGACAGTGAGGGACGAAAGCTAGGGT |
| P-asv272 | GCAAGCGTTACTCGGAATCACTGGGCGTAAAGAGCGTGTAGGCGGGTATATAAGTCAGAAGTGAAATCCAATAGCTCAACTATTGAACTGCTTTTGAAACTGTATACCTAGAATGTGGGAGAGGTAGATGGAATTT<br>CTGGTGTAGGGGTAAAATCCGTAGAGATCAGAAGGAATACCGATTGCGAAGGCGATCTACTGGAACATTATTGACGCTGAGACGCGAAAGCGTGGGG |
| P-asv274 | CCAAGCGTTAATCGGAATTACTGGGCGTAAAGCGTGCGCAGGCGGTTGTGCAAGACCGATGTGAAATCCCCGAGCTTAACTTGGGAATTGCATTGGTGACTGCACGGCTAGAGTGTGTCAGAGGGGGGTAGAAT<br>TCCACGTGTAGCAGTGAAATGCGTAGAGATGTGGAGGAATCCCGATGGCGAAGGCAGCCCCCTGGGATAACACTGACGCTCATGCACGTAAGCGTGGGG |
| P-asv279 | GCTAGCGTTGTTCGGAATCACTGGGCGTAAAGCGCACGTAGGCGGATTATTAAGTCAGGGGTGAAAGCCTGGAGCTCAACTCCAGAACTGCCTTTGATACTGATAGTCTCGAGTTCGGAAGAGGTTGGTGGAACT<br>GCGAGTGTAGAGGTGAAATTCGTAGATATTCGCAAGAACACCAGTGGCGAAGGCGGCCAACTGGTCCGATACTGACGCTGAGGTGCGAAAGCGTGGGG |
| P-asv281 | GCTAGCGTTGTTCGGAATTACTGGGCGTAAAGCGCACGTAGGCGGATATTTAAGTCAGGGGTGAAATCCCAGAGCTCAACTCTGGAACTGCCTTTGATACTGGGTATCTTGAGTATGGAAGAGGTGAGTGGAATT<br>CCGAGTGTAGAGGTGAAATTCGTAGATATTCGGAGGAACACCAGTGGCGAAGGCGGCTCACTGGTCCATTACTGACGCTGAGGTGCGAAAGCGTGGGG |
| P-asv286 | GCAAGCGTTATCCGGATTTATTGGGTTTAAAGGGTGCGTAGGCGGGGTATTAAGTCAGTGGTGAAATCTTTCGGCTCAACCGAGAAACTGCCATTGATACTGATACTCTTGAATTTGGTTGAAGTAGGCGGAATGA<br>GGCATGTAGCGGTGAAATGCATAGATATGCCTCAGAACACCGATTGCGAAGGCAGCTTACTAAGTCAATATTGACGCTGAGGCACGAAAGCGTGGGT |
| P-asv287 | TCGAGTGATGGCCGATCTTATTGGGCCTAAAGCGTCCGTAGCTGGCCGTGCAAGTCCGTTGGGAAATCCACCCGCTCAACGGGTGGGCGGCCGACGGAAACTGGACGGCTTGGGACCGGGAGGCTTGAAGGG<br>TACGTCCGGGGTAGGAGTGAAATCCCGTAATCCTGGACGGACCACCGATGGGGAAACCACTTCAAGAGAACGGATCCGACAGTGAGGGACGAAAGCTAGGGT |
| P-asv291 | CCGAGTGATGGCCGAAATTATTGGGCCTAAAGCGTTCGTAGCTGGCCGGACAAGTCCGTTGGGAAATCCGCGCGCTTAACGCGCGGGCGTCCAGCGGAAACTGTTCGGCTTGGGACCGGAAGACCTGAGGGGT<br>ACGTCCGGGGTAGGAGTGAAATCCTGTAATCCTGGACGGACCACCGGTGGCGAAAGCGCCTCAGGAGGACGGATCCGACAGTGAGGAACGAAAGCTAGGGT |
| P-asv292 | GCGAGCGTTGTTCGGAATTACTGGGCGTAAAGCGTATGTAGGCGGCTTCGTAAGTCAGATGTGAAAGCCCAGAGCTTAACTTTGGAACTGCATTTGAGACTGCGAGGCTTGAATATCGGAGGGGTTGGCGGAATT<br>CCCAGTGTAGCAGTGAAATGCGCAGATATTGGGAGGAACACCGATGGCGTAAGCAGCCAACTGGACGATTATTGACGCTGAGATACGAAAGCGTGGGG |
| P-asv294 | GCAAGCGTTACTCGGAATCACTGGGCGTAAAGAGCATGTAGGCGGATAGATAAGTTGGAAGTGAAATCCACTGGCTTAACCAGTGAACTGCTTCCAAAACTGTCTATCTAGAATATGGGAGAGGTAGATGGAATTT<br>CTGGTGTAGGGGTAAAATCCGTAGAGATCAGAAGGAATACCGATTGCGAAGGCGATCTACTGGAACATTATTGACGCTCAGATGCGAAAGCGTGGGG |
| P-asv296 | TCGAGTGATGGCCGATCTTATTGGGCCTAAAGCGTCCGTAGCTGGCCTGGCAAGTCCGTCGGGAAATCCACCCGCTCAACGGGTGGGCGACCGACGGAAACTGTCGGGCTTGGGACCGGGAGGCTTGAGGGG<br>TACGTCCGGGGTAGGAGTGAAATCCCGTAATCCTGGACGGACCACCGATGGGGAAACCACCTCAAGAGAACGGATCCGACAGTGAGGGACGAAAGCTAGGGT |
| P-asv299 | GCTAGCGTTGTTCGGAATTACTGGGCGTAAAGCGCACGTAGGCGGATACTTAAGTCAGGGGTGAAATCCCGGGGCTCAACCCCGGAACTGCCTTTGATACTGGGTATCTGGAGTCCGGAAGAGGTGAGTGGAAT<br>TCCGAGTGTAGAGGTGAAATTCGTAGATATTCGGAGGAACACCAGTGGCGAAGGCGGCTCACTGGTCCGGTACTGACGCTGAGGTGCGAAAGCGTGGGG |
| P-asv300 | GCTAGCGTTGTTCGGAATTACTGGGCGTAAAGCGCACGTAGGCGGCTTTTTAAGTCAGGGGTGAAATCCCGGGGCTCAACCCCGGAACTGCCTTTGAAACTGGAAAGCTAGAATACTGGAGAGGTGAGTGGAAT<br>TCCGAGTGTAGAGGTGAAATTCGTAGATATTCGGAAGAACACCAGTGGCGAAGGCGACTCACTGGACAGTTATTGACGCTGAGGTGCGAAAGCGTGGGG |
| P-asv303 | CCGAGCGTTGTCCGGATTTATTGGGCGTAAAGCGAGCGCAGGTGGTTTATTAAGTCTGGTGTAAAAGGCAGTGGCTCAACCATTGTATGCATTGGAAACTGGTAGACTTGAGTGCAGGAGAGGAGAGTGGAATTC<br>CATGTGTAGCGGTGAAATGCGTAGATATATGGAGGAACACCGGTGGCGAAAGCGGCTCTCTGGCCTGTAACTGACACTGAGGCTCGAAAGCGTGGGG |
| P-asv305 | GCTAGCGTTGTTCGGAATTACTGGGCGTAAAGCGCACGTAGGCGGCTATTCAAGTCAGAGGTGAAAGCCCGGGGCTCAACCCCGGAACTGCCTTTGAAACTAGATAGCTTGAATCTTGGAGAGGCGAGTGGAAT<br>TCCGAGTGTAGAGGTGAAATTCGTAGATATTCGGAAGAACACCAGTGGCGAAGGCGACTCGCTGGACAAGTATTGACGCTGAGGTGCGAAAGCGTGGGG |
| P-asv307 | GCAAGCGTTAATCGGAATTACTGGGCGTAAAGCGCGCGTAGGTGGTTCAGTAAGTTGGAAGTGAAATCCCCGGGCTCAACCTGGGAACTGCTTTCAAAACTGCTGAGCTAGAGTACGGTAGAGGGTAGTGGAAT<br>TTCCTGTGTAGCGGTGAAATGCGTAGATATAGGAAGGAACACCAGTGGCGAAGGCGACTACCTGGACTGATACTGACACTGAGGTGCGAAAGCGTGGGG |
| P-asv309 | GCAAGCGTTAATCGGAATCACTGGGCGTAAAGGGCACGCAGGCGGTTGGATAAGTCGGGTGTGAAAGCCCCGGGCTCAACCTGGGAATGGCATTCGATACTGTCTAGCTAGAGTCTGGTAGAGGGAAGTGGAA<br>TTTCCGGTGTAGCGGTGAAATGCGTAGATATCGGAAGGAACACCAGTGGCGAAGGCGACTTCCTGGACCAAGACTGACGCTGAGGTGCGAAAGCGTGGGT |
| P-asv314 | GCAAGCGTTGTTCGGAATCACTGGGCGTAAAGCGTACGTAGGCGGCTATTCAAGTCAGAAGTGAAAGCCTGGGGCTCAACCCCAGAACTGCTTTTGAAACTGGATGGCTAGAGTCCCGGAGAGGTTGGCGGAAT<br>TCCGAGTGTAGAGGTGAAATTCGCAGATATTCGGAGGAACACCAGTGGCGTAGGCGGCCAACTGGACGGGAACTGACGCTGAGGTACGAAAGCGTGGGG |
| P-asv315 | GCAAGCGTTAATCGGAATTACTGGGCGTAAAGCGCACGTAGGCGGTATGTTAAGTTGGATGTGAAAGCCCCGGGCTCAACCTGGGAATTGCATTCAAAACTGGCATGCTAGAGTATGTGAGAGGGGGGTAGAAT<br>TCCAAGTGTAGCGGTGAAATGCGTAGAGATTTGGAGGAATACCAGTGGCGAAGGCGGCCCCCTGGCACAATACTGACGCTCAGGTGCGAAAGCGTGGGG |
| P-asv318 | GCGAGCGTTAATCGGAATTATTGGGCGTAAAGCGAGTGTAGACGGTTATTTAAGCCAGATGTGAAATACCCGAGCCTAACTTGGGAGGTGCATATGGAACTGGGTAGCTAGAGTGTGTCAGAGGGAGGTAGAACT<br>CCACGTGTAGCAGTGAAATGCGTAGAGATGTGGAAGAATACCGATGGCGAAGGCAGCCTCCTGGGATAACACTGACGTTGAGGCTCGAAAGCGTGGGG |
| P-asv323 | TCGAGTGATGGCCGATCTTATTGGGCCTAAAGCGTCCGTAGCTGGCCGTGCAAGTCCGTCGGGAAATCCACCCGCTTAACGGGTGGGCGGCCGACGGAAACTGGACGGCTTGGGACCGGGAGGCTTGAGGGG<br>TACGTCCGGGGTAGGAGTGAAATCCCGTAATCCTGGACGGACCACCGATGGGGAAACCACCTCAAGAGAACGGATCCGACAGTGAGGGACGAAAGCTAGGGT |
| P-asv324 | GCTAGCGTTGTTCGGAATTACTGGGCGTAAAGCGCACGTAGGCGGCTTTTTAAGTCAGGGGTGAAATCCCGGGGCTCAACCCCGGAACTGCCCTTGAAACTGGAAAGCTAGAATACTGGAGAGGTGAGTGGAAT<br>TCCGAGTGTAGAGGTGAAATTCGTAGATATTCGGAAGAACACCAGTGGCGAAGGCGACTCACTGGACAGTTATTGACGCTGAGGTGCGAAAGCGTGGGG |
| P-asv325 | GTTAGCGTTGTTCGGAATTACTGGGCGTAAAGCGCACGTAGGCGGATCAGAAAGTTGGGGGTGAAATCCCGGGGCTCAACCCCGGAACGGCCTCCAAAACTCCTGGTCTTGAGTTCGAGAGAGGTGAGTGGAAT<br>TCCGAGTGTAGAGGTGAAATTCGTAGATATTCGGAGGAACACCAGTGGCGAAGGCGGCTCACTGGCTCGATACTGACGCTGAGGTGCGAAAGTGTGGGG |
| P-asv328 | GCAAGCGTTATCCGGAATGATTGGGCGTAAAGCGTCTGTAGGTGGTTCTTTAAGTCTACTGTCAAATACGTTGGCTCAACCAACGGCAGGCAGTAGAGTACTAAAGAACTAGAGTGCGGTAGAGGTAGAGGGAAT<br>CCCTAGCGTAACAGTGAAATGTGTAGATTTTAGGGGGAACACCAGCGGCGAAGGCGCTCTACTGGGCCGATACTGACACTGAGAGACGAAAGCTAGAGG |
| P-asv329 | GCAAGCGTTGTCCGGATTTATTGGGCGTAAAGCGAGCGCAGGCGGTTTCTTAAGTCTGATGTGAAAGCCCCCGGCTCAACCGGGGAGGGTCATTGGAAACTGGGAAACTTGAGTGCAGAAGAGGAGAGTGGAAT<br>TCCATGTGTAGCGGTGAAATGCGTAGATATATGGAGGAACACCAGTGGCGAAGGCGGCTCTCTGGTCTGTAACTGACGCTGAGGCTCGAAAGCGTGGGG |

| | |
|---|---|
| P-asv330 | GCAAGCGTTAATCGGAATTACTGGGCGTAAAGCGCGCGTAGGTTGTTAGTTAAGCGAGATGTGAAAGCCCCGGGCTTAACCTGGGAATTGCGTTTCGAACTGGCTGACTAGAGTACGGTAGAGGGTGGTGGAAT<br>TTCAGGTGTAGCGGTGAAATGCGTAGAGATCTGAAGGAACATCAGTGGCGAAGGCGACCACCTGGACTGATACTGACACTGAGGTGCGAAAGCGTGGGG |
| P-asv333 | GCAAGCGTTAATCGGAATTACTGGGCGTAAAGCGTGCGCAGGCGGTTTTGTAAGACAGGCGTGAAATCCCCGGGCTCAACCTGGGAATTGCGCTTGTGACTGCAAGGCTGGAGTGCGGCAGAGGGGGATGGAA<br>TTCCGCGTGTAGCAGTGAAATGCGTAGATATGCGGAAGAACACCGATGGCGAAGGCAATCCCCTGGGCCTGCACTGACGCTCATGCACGAAAGCGTGGGG |
| P-asv338 | GCAAGCGTTAATCGGAATTACTGGGCGTAAAGCGCGCGTAGGCGGTTTGATAAGTGAGATGTGAAAGCCCCGGGCTCAACCTGGGAACTGCATCTCATACTGTCAGGCTAGAGTACGGTAGAGGGGGGTAGAAT<br>TCCACGTGTAGCGGTGAAATGCGTAGAGATGTGGAGGAATACCAGTGGCGAAGGCGGCCCCCTGGATCGATACTGACGCTGAGGTGCGAAAGCGTGGGG |
| P-asv339 | GCAAGCGTTAATCGGAATTACTGGGCGTAAAGCGTGCGCAGGCGGTTCGGAAAGAAAGATGTGAAATCCCAGAGCTTAACTTTGGAACTGCATTTTTAACTACCGAGCTAGAGTGTGTCAGAGGGAGGTGGAATT<br>CCGCGTGTAGCAGTGAAATGCGTAGATATGCGGAGGAACACCGATGGCGAAGGCAGCCTCCTGGGATAACACTGACGCTCATGCACGAAAGCGTGGGG |
| P-asv341 | GCAAGCGTTAATCGGAATTACTGGGCGTAAAGCGCACGTAGGCGGTTAGTTAAGTTGGATGTGAAAGCCCCGGGCTCAACCTGGGAATTGCATTCAAAACTGGCTAACTAGAGTATGTGAGAGGGGGGTAGAATT<br>CCAAGTGTAGCGGTGAAATGCGTAGAGATTTGGAGGAATACCAGTGGCGAAGGCGGCCCCCTGGCACAATACTGACGCTGAGGTGCGAAAGCGTGGGG |
| P-asv344 | ACGAGTGATGGCCGATATTATTGGGCCTAAAGCGTCCGTAGCCGGCCGAACAAGTCCGTTGGGAAATCGACGCGCTCAACGCGTCGGCGTCCGGCGGAAACTGTTCGGCTTGGGGCCGGAAGACCTGAGGGGT<br>ACGTCCGGGGTAGGAGTGAAATCCCGTAATCCTGGACGGACCACCGGTGGCGAAAGCGCCTCAGGAAGACGGACCCGACGGTGAGGGACGAAAGCTAGGGT |
| P-asv346 | GCTAGCGTTGTTCGGAATTACTGGGCGTAAAGCGCACGTAGGCGGCGACACAAGTCAGAGGTGAAAGCCCGGGGCTCAACCCCGGAACTGCCTTTGAAACTAGGTTGCTAGAATCTTGGAGAGGTCAGTGGAAT<br>TCCGAGTGTAGAGGTGAAATTCGTAGATATTCGGAAGAACACCAGTGGCGAAGGCGACTGACTGGACAAGTATTGACGCTGAGGTGCGAAAGCGTGGGG |
| P-asv348 | CCAAGCGTTAATCGGAATTACTGGGCGTAAAGCGTGCGCAGGCGGTTTGTTAAGCGAGATGTGAAAGCCCCGGGCTCAACCTGGGAATTGCATTTCGAACTGGCAAACTAGAGTCTTGTAGAGGGGGGTAGAAT<br>TCCAGGTGTAGCGGTGAAATGCGTAGAGATCTGGAGGAATACCGGTGGCGAAGGCGGCCCCCTGGACAAAGACTGACGCTCAGGCACGAAAGCGTGGGG |
| P-asv350 | GTTAGCGTTGTTCGGAATTACTGGGCGTAAAGCGCACGTAGGCGGATCAGAAAGTTGGGGGTGAAATCCCAGGGCTCAACCCTGGAACGGCCTCCAAAACTCCTGGTCTTGAGTTCGAGAGAGGTGAGTGGAAT<br>TCCGAGTGTAGAGGTGAAATTCGTAGATATTCGGAAGAACACCAGTGGCGAAGGCGGCTCACTGGCTCGATACTGACGCTGAGGTGCGAAAGTGTGGGG |
| P-asv352 | GCAAGCGTTATTCGGCATGACTGGGCGTAAAGAGTCCGTAGGCGGTAAAATAAGTTTCTTGTTAAATTTTAAGACTTAATCTTAAGCTAGCAATTAATACTGTTTTACTAAGAGTTTAATACAGAAAAGTAGAATTTTA<br>TACGATAGAGGTGAAATTTGTAAATATATAAAGGAATGCCGACTGCGAAGGCAACTTTTTGGTATAAACTGACGTTGAGGGACGAAAGCGTGGGT |
| P-asv356 | GCGAGCGTTGTCCGGAATTACTGGGCGTAAAGAGCTCGTAGGTGGTTTGTCGCGTCGTCTGTGAAATTCCGGGGCTTAACTCCGGGCGTGCAGGCGATACGGGCATAACTTGAGTACTGTAGGGGAGACTGGA<br>ATTCCTGGTGTAGCGGTGAAATGCGCAGATATCAGGAGGAACACCGGTGGCGAAGGCGGGTCTCTGGGCAGTAACTGACGCTGAGGAGCGAAAGCATGGGG |
| P-asv357 | CCGAGTGATGGCCGATATTATTGGGCCTAAAGCGTCCGTAGCTGGCCGGACAAGTCCGTTGGGAAATCTGCTCGCTTAACGAGCAGGCGTCCAGCGGAAACTGTTCGGCTTGGGACCGGAAGACCTGAGGGGT<br>ACGTCCGGGGTAGGAGTGAAATCCCGTAATCCTGGACGGACCACCGGTGGCGAAAGCGCCTCAGGAAGACGGATCCGACAGTGAGGGACGAAAGCTAGGGT |
| P-asv359 | CCGAGTGATGGCCGATATTATTGGGCCTAAAGCGTCCGTAGCCTGCTGTGTAAGTCCATTGGGAAATCGACCAGCTCAACTGGTCGGCGTCCGGTGGAAACTACACAGCTTGGGGCCGAGAGACTCAACGGGTA<br>CGTCCGGGGTAGGAGTGAAATCCTATAATCCTGGACGGACCACCAATGGGGAAACCACGTTGAGAGACCGGACCCGACGGTGAGGGACGAAAGCCAGGGT |
| P-asv362 | CCAAGCGTTATCCGGAATCATTGGGTTTAAAGGGTCCGTAGGCGGCCTTATAAGTCAGTGGTGAAATCTCCTAGCTCAACTAGGAAACTGCCATTGATACTGTAGGGCTTGAATTTTTGTGAAGTAACTAGAATATG<br>TAGTGTAGCGGTGAAATGCTTAGATATTACATAGAATACCAATTGCGAAGGCAGGTTACTAACAAACGATTGACGCTGATGGACGAAAGCGTGGGG |
| P-asv368 | GCAAGCGTTAATCGGAATTACTGGGCGTAAAGCGCACGCAGGCGGTTGGTTAAGTCAGATGTGAAAGCCCCGGGCTCAACCTGGGAATTGCATTTGAAACTGGCCAACTAGAGTATGTGAGAGGGGGGTAGAAT<br>TCCAAGTGTAGCGGTGAAATGCGTAGAGATTTGGAGGAATACCAGTGGCGAAGGCGGCCCCCTGGCACAATACTGACGCTCAGGTGCGAAAGCGTGGGG |
| P-asv371 | GCTAGCGTTGCTCGGAATTACTGGGCGTAAAGGGCGCGTAGGCGGACAGTTTAGTCAGAGGTGAAAGCCCAGGGCTCAACCTTGGAATTGCCTTTGATACTGGCTGTCTTGAGTTCGGGAGAGGTGAGTGGAAT<br>GCCGAGTGTAGAGGTGAAATTCGTAGATATTCGGCGGAACACCAGTGGCGAAGGCGACTCACTGGCCCGATACTGACGCTGAGGCGCGAAAGCGTGGGG |
| P-asv372 | GCAAGCGTTAATCGGAATTACTGGGCGTAAAGCGCGCGTAGGCGGCTTGATAAGATGGGTGTGAAATCCCCGGGCTTAACCTGGGAACTGCATCCATAACTGTCTGGCTAGAGTACAGTAGAGGGTGGTGGAAT<br>TTCCTGTGTAGCGGTGAAATGCGTAGATATAGGAAGGAACACCAGTGGCGAAGGCGACCACCTGGACTGATACTGACGCTGAGGTGCGAAAGCGTGGGG |
| P-asv377 | TCGAGTGATGTCCCATATTATTGGGCCTAAAGCGTCCGTAGCTGGCCAGGCAAGTCTATCGGGAAATCCACCCGCCCAACGGGTGAGCGTCCGGTAGAAACTGCCCGGCTTGGAACCGGAAGGCTCAGAGAGTA<br>CGTCCGGGGTAGGAGTGAAATCCCGTAATCCCGGACGGACTACCGATGGCGAAAGCACTCTGAGAAGACGGCTTCGACAGTGAGGGACGAAAGCTAGGGT |
| P-asv378 | GCAAGCGTTGTTCGGAATCACTGGGCTTAAAGCGTATGTAGGCGGACTTGAAAGTCAGAAGTGAAATCCCCGGGCTTAACCCGGGAACTGCTTTTGAAACTCCAAGTCTAGAATCCTGGAGAGGTTGGGAGAATT<br>CCGAGTGTAGAGGTGAAATTCGCAGATATTCGGAGGAATACCAGTGGCGTAGGCGCCCAACTGGACAGGTATTGACGCTCAGATACGAAAGCGTGGGG |
| P-asv379 | GCTAGCGTTGTTCGGAATGACTGGGCGTAAAGGGCGCGTAGGCGGTCTTTTAAGTTAGGCGTGAAAGCCCCGGGCTTAACCCGGGACGTGCGCTTAAGACTGAAAGGCTTGAGTATGGAAGAGGTAAGTGGAAT<br>TCCTAGTGTAGAGGTGAAATTCGTAGATATTAGGAAGAACACCGGTGGCGAAGGCGGCTTACTGGTCCATTACTGACGCTGAGGCGCGAAAGTGTGGGG |
| P-asv380 | ACGAGTGATGGCCGATCTTATTGGGCCTAAAGCGTCCGTAGCCGGCCAGACAAGTCCGTTGGGAAATCGACGTGCCTAACACGTCGGCGTCCGGCGGAAACTGTCTGGCTTGGGGCCGGAAGACTCGAGGGGT<br>ACGTCCGGGGTAGGAGTGAAATCCCGTAATCCTGGACGGACCGCCGGTGGCGAAAGCGCCTCGAGAGGACGGACCCGACGGTCAGGGACGAAAGCTAGGGT |
| P-asv383 | GCAAGCGTTATCCGGATTTATTGGGTTTAAAGGGTCCGTAGGCGGACTTATAAGTCAGTGGTGAAAGCCTGTCGCTTAACGATAGAACTGCCATTGATACTGTAAGTCTTGAGTATATTTGAGGTAGCTGGAATAA<br>GTAGTGTAGCGGTGAAATGCATAGATATTACTTAGAACACCAATTGCGAAGGCAGGTTACCAAGATATAACTGACGCTGAGGGACGAAAGCGTGGGG |
| P-asv386 | ACGAGTGATGGCCGATATTATTGGGCCTAAAGCGTCCGTAGCTTGCTGTGTAAGTCCATTGGGAAATCGACGCGCTCAACGCGTCGGCGTCCGGTGGAAACTGTACAGCTTGGGGCCGAGAGACTCGACGGGTA<br>CGTCCGGGGTAGGAGTGAAATCCCGTAATCCTGGACGGACCACCAATGGGGAAACCACGTCGAGAGACCGGACCCGACAGTGAGGGACGAAAGCCAGGGT |
| P-asv389 | CCAAGTGATGGCCGATATTATTGGGCCTAAAGCGTCCGTAGCCAGCCGAACAGGTCCGTTGGGAAATCTGCTCGCTCAACGAGCAGGCGTCCGGCGGAAACCAGTCGGCTTGGGACCGAGAGGACCGAAGGGT<br>ACGTCCGGGGTAGGAGTGAAATCCCGTAATCCTGGACGGACCACCGATGGCGAAAGCACTTCGGTAGATCGGATCCGACGGTGAGGGACGAAAGCTAGGGT |
| P-asv390 | GCAAGCGTTAATCGGAATCACTGGGCGTAAAGGGCACGCAGGCGGTTGGATAAGTCGGGTGTGAAAGCCCCGGGCTCAACCTGGGAATGGCATTCGATACTGTCCAGCTAGAGTCTGGTAGAGGGAAGTGGAA<br>TTTCCGGTGTAGCGGTGAAATGCGTAGATATCGGAAGGAACACCAGTGGCGAAGGCGACTTCCTGGACCAAGACTGACGCTGAGGTGCGAAAGCGTGGGT |
| P-asv396 | GCTAGCGTTGTTCGGAATTACTGGGCGTAAAGCGAGCGTAGGTTGCGCCGCAAGTCAGATGTGAAATCCCAGGGCTTAACCCTGGAACTGCATTTGAAACTGCGGTGCTGGAATCTCAGAGGGGGTAGCGGAAT<br>TCCAAATGTAGGGGTGAAATCCGTAGATATTTGGAGGAACACCGGTGGCGAAGGCGGCTACCTGGATGAGTATTGACACTGAGGCTCGAAAGCGTGGGG |
| P-asv401 | GCAAGCGTTGTCCGGATTTATTGGGCGTAAAGCGAGCGCAGGCGGAAGAATAAGTCTGATGTGAAAGCCCTCGGCTTAACCGAGGAACTGCATCGGAAACTGTTTTTCTTGAGTGCAGAAGAGGAGAGTGGAAC<br>TCCATGTGTAGCGGTGGAATGCGTAGATATATGGAAGAACACCAGTGGCGAAGGCGGCTCTCTGGTCTGCAACTGACGCTGAGGCTCGAAAGCATGGGT |
| P-asv402 | CCAAGTGATGGCCGATATTATTGGGCCTAAAGCGTCCGTAGCCAGCCGAACAGGTCCGTCGGGAAATCTGCTCGCTCAACGAGCAGGCGTCCGGCGGAAACCAGTCGGCTTGGGACCAGAAGACCCGAGGGGT<br>ACGTCCGGGGTAGGAGTGAAATCCCGTAATCCTGGACGGACCACCGGTGGCGAAAGCGCCTCGGGAAGATGGATCCGACGGTGAGGGACGAAAGCTAGGGT |
| P-asv404 | GCTAGCGTTGTTCGGAATTACTGGGCGTAAAGGGCGCGTAGGCGGACCTGCAAGTCAGGGGTGAAATCCCGGGGCTCAACCTCGGAACTGCCCTTGAGACTGTTGGTCTAGAGTCTGTGAGAGGATGACGGAA<br>TACCCAGTGTAGAGGTGAAATTCGTAGATATTGGGTAGAACACCGGTGGCGAAGGCGGTCATCTGGCGCAGCACTGACGCTGAGGCGCGAAAGCGTGGGG |
| P-asv406 | GCAAGCGTTAATCGGAATTACTGGGCGTAAAGCGCGCGTAGGCGGTTTGATAAGTTGGATGTGAAAGCCCCGGGCTCAACCTGGGAATTGCATCCAAGACTGTCAGGCTAGAGTACGGGAGAGGGGGGTAGAA<br>TTCCACGTGTAGCGGTGAAATGCGTAGAGATGTGGAGGAATACCGGTGGCGAAGGCGGCCCCCTGGATCGATACTGACGCTGAGGTGCGAAAGCGTGGGG |
| P-asv407 | GCTAGCGTTGTTCGGAATTACTGGGCGTAAAGCGTGCGTAGGCGGCTTTGCAAGTAGAGGGTGAAATCCCGGGGCTTAACCTCGGAACTGCCTTCTAAACTGCATGGCTAGAATCCTGGAGAGGTGAGCGGAAT<br>TCCTAGTGTAGAGGTGAAATTCGTAGATATTAGGAAGAACACCAGTGGCGAAGGCGGCTCACTGGACAGGTATTGACGCTGAGGTACGAAAGCGTGGGG |
| P-asv408 | GCGAGCGTTGTCCGGAATCACTGGGCGTAAAGGGCGCGTAGGCGGCCGACTAAGTGTGTGGTGAAAGCCCGGGGCTCAACCCCGGGTCTGCCGTGCAAACTGGTTGGCTTGAGCACTGTAGAGGCAGGTGGA<br>ATTCCGGGTGTAGCGGTGGAATGCGTAGAGATCCGGAAGAACACCGGTGGCGAAGGCGGCCTGCTGGGCAGTTGCTGACGCTGAGGCGCGACAGCGTGGGG |
| P-asv416 | CCGAGTGATGGCCGATCTTATTGGGCCTAAAGCGTCCGTAGCCGGCCTCGCAAGTCCGTCGGGAAATCCAGTAGCCCAACTACTGGGCGTCCGGCGGAAACTCCGAGGCTTGGGACCGGAAGGCTCGAGGGGT<br>ACGTCCGGGGTAGGAGTGAAATCCCGTAATCCCGAACGGACCACCGATGGCGAAAGCACCTCGAGAAGACGGATCCGACGGTAAGGGACGAAAGCCAGGGT |
| P-asv417 | GCGAGCGTTATCCGGATTTATTGGGTTTAAAGGGTGCGTAGGCGGCTTAGTAAGTCGGTGGTTAAAGGTAGTCGCTTAACGATTTTACATGCCACCGATACTGTTGAGCTTGAGTTATTTGGAGGCAGCTAGAATT<br>TCCGGTGTAGCGGTGAAATGCATAGATACCGGAAGGAATACCAATTGCGAAGGCAGGTTGCTACAAATATACTGACGCTGATGCACGAAAGCGTGGGG |
| P-asv421 | GCAAGCGTTACTCGGAATCACTGGGCGTAAAGAGCGTGTAGGCGGTTTGATAAGTTGGAAGTGAAATCCTATAGCTTAACTATAGAATTGCTTCCAAAACTGTTTAACTAGAATGTGGGAGAGGTAGATGGAATTT<br>CTGGTGTAGGGGTAAAATCCGTAGATATCAGAAGGAATACCTATTGCGAAGGCGATCTACTGGAACATAATTGACGCTGAGACGCGAAAGCGTGGGG |
| P-asv426 | GCAAGCGTTAATCGGAATTACTGGGCGTAAAGCGTGCGCAGGCGGTTATGCAAGACAGAGGTGAAATCCCCGGGCTCAACCTGGGAACTGCCTTTGTGACTGCATAGCTAGAGTACGGTAGAGGGGGATGGAAT<br>TCCGCGTGTAGCAGTGAAATGCGTAGATATGCGGAAGAACACCGATGGCGAAGGCAATCCCCTGGACCTGTACTGACGCTCATGCACGAAAGCGTGGGG |
| P-asv427 | GCTAGCGTTATTCGGAATTACTGGGCGTAAAGCGCACGTAGGCGGCTTTGTAAGTTAGAGGTGAAAGCCTGGAGCTCAACTCCAGAATTGCCTTTAAGACTGCATCGCTTGAATCCAGGAGAGGTGAGTGGAATT<br>CCGAGTGTAGAGGTGAAATTCGTAGATATTCGGAAGAACTCCAGTGGCGAAGGCGGCTCACTGGACTGGTATTGACGCTGAGGTGCGTAAGCGTGGGG |
| P-asv429 | GCGAGCGTTAATCGGAATTACTGGGCGTAAAGCGTGCGCAGGCGGTTTTTTAAGATAGGCGTGAAATCCCCGGGCTCAACCTGGGAACTGCGCTTATGACTGGAAGACTAGAGTATGGCAGAGGGGGGTGGAAT<br>TCCACGTGTAGCAGTGAAATGCGTAGAGATGTGGAGGAACACCGATGGCGAAGGCAGCCCCCTGGGCCAATACTGACGCTCATGCACGAAAGCGTGGGG |
| P-asv430 | GCAAGCGTTATCCGGAATGATTGGGCGTAAAGCGTCTGTAGGTGGCTTTTTAAGTTCGCCGTCAAATCCCAGGGCTCAACCCTGGACAGGCGGTGGAAACTACCAAGCTGGAGTACGGTAGGGGCAGAGGGAAT<br>TTCCGGTGGAGCGGTGAAATGCGTAGAGATCGGAAAGAACACCAACGGCGAAAGCACTCTGCTGGGCCGACACTGACACTGAGAGACGAAAGCTAGGGG |

| | |
|---|---|
| P-asv431 | GCGAGCGTTAATCGGAATTACTGGGCGTAAAGCGTGCGCAGGCGGTTTTGTAAGACAGGCGTGAAATCCCCGGGCTCAACCTGGGAACTGCGCTTGTGACTGCAAGGCTAGAGTACGGCAGAGGGGGGTGGAATTCCACGTGTAGCAGTGAAATGCGTAGAGATGTGGAGGAACACCAATGGCGAAGGCAGCCCCCTGGGTCGATACTGACGCTCATGCACGAAAGCGTGGGT |
| P-asv433 | TCGAGTGATGGCCGATCTTATTGGGCCTAAAGCGTTCGTAGCTGGCTAGTCAAGTCCGTTGGGAAATCGACGTGCCTAACACGTCGGCGTCCAGCGGAAACTGACTTGCTTGGGACCGGAAGACTCGAGGGGTACGTCCGGGGTAGGAGTGAAATCCTGTAATCCTGGACGGACCACCAATGGCGAAAGCACCTCGAGAGCACGGATCCGACAGTGAGGAACGAAAGCTGGGGT |
| P-asv436 | CCGAGTGATGGCCGATATTATTGGGCCTAAAGCGTCCGTAGCCGGCCGGACAAGTCCGTTGGGAAATCGACGCGCTCAACGCGTCGGCGTCCAGCGGAAACTGTCCGGCTTGGGGCCGGAAGACCTGAGGGGTACGTCCGGGGTAGGAGTGAAATCCTGTAATCCTGGACGGACCACCAATGGGGAAACCACCTCAGGAAGACGGACCCGACGGTGAGGGACGAAAGCTAGGGT |
| P-asv437 | GCAAGCGTTAATCGGAATTACTGGGCGTAAAGCGTACGTAGGTGGTTTGTTAAGCCAGCTGTGAAATCCCCGGGCTCAACCTGGGCACTGCAGTTGGAACTGGCAAGCTAGAGTAGGGTAGAGGGGTGTGGAATTCCAGGTGTAGCGGTGAAATGCGTAGATATCTGGAGGAACATCAGTGGCGAAGGCGACACCCTGGACTCATACTGACACTGAGGTACGAAAGCGTGGGG |
| P-asv444 | GCAAGCGTTAATCGGAATTACTGGGCGTAAAGCGCGCGTAGGCGGTTTGTTAAGTCGGATGTGAAAGCCCCGGGCTCAACCTGGGAACTGCACCCGATACTGGCCGACTAGAGTACGAGAGAGGGAGGTAGAATTCCACGTGTAGCGGTGAAATGCGTAGATATGTGGAGGAATACCGGTGGCGAAGGCGACCTCCTGGCTCGATACTGACGCTGAGGTGCGAAAGCGTGGGG |
| P-asv449 | GCTAGCGTTGTTCGGAATTACTGGGCGTAAAGGGCGCGTAGGCGGCTTTGTAAGTCAGATGTGAAAGCCCCGGGCTCAACCTGGGAACTGCATTTGATACTGCAAGGCTAGAGATCGAGAGAGGAAAGTGGAATTACGAGTGTAGAAGTGAAATTCGTAGATATTCGTAGGAACACCAGTGGCGAAGGCGACTTTCTGGCTCGATACTGACGCTAAAGCGCGAAAGCGTGGGG |
| P-asv450 | GCGAGCGTTAATCGGAATTACTGGGCGTAAAGCGTGCGCAGGCGGTTTGGCAAGTCAGATGTGAAATCCCCGAGCTCAACTTGGGAACTGCGTTTGAAACTGCCAGACTAGAATATGTCAGAGGGGGGTAGAATTCCACGTGTAGCAGTGAAATGCGTAGAGATGTGGAGGAATACCAATGGCGAAGGCAGCCCCCTGGGATAATATTGACGCTCATGCACGAAAGCGTGGGG |
| P-asv452 | GCAAGCGTTGTCCGGAATTATTGGGCGTAAAGCGCGCGCAGGCGGTCTCTTAAGTCTGATGTGAAAGCCCACGGCTCAACCGTGGAGGGTCATTGGAAACTGGGAGACTTGAGTGTAGGAGAGGAAAGTGGAATTCCACGTGTAGCGGTGAAATGCGTAGAGATGTGGAGGAACACCAGTGGCGAAGGCGACTTTCTGGCCTATAACTGACGCTGAGGCGCGAAAGCGTGGGG |
| P-asv456 | CCGAGTGATGGCCGATATTATTGGGCCTAAAGCGTTCGTAGCTGGCCGGACAAGTCCGTTGGGAAATCCGCGCGCTTAACGCGCGGGCGTCCAGCGGAAACTGTTCGGCTTGGGACCGGAAGACCTGAGGGGTACGTCCGGGGTAGGAGTGAAATCCCGTAATCCTGGACGGACCACCGGTGGCGAAAGCGCCTCAGGAGGACGGATCCGACAGTGAGGAACGAAAGCTAGGGT |
| P-asv459 | CCGAACGTTATTCGGAATTACTGGGCTTAAAGGGTGCGTAGGCGGCCATGCAAGTCAGATGTGAAATCCCACGGCTCAACCGTGGAACTGCGTTTGAAACTGCATGGCTTGAGGGAGATAGGGGTGAGCGGAACTGATGGTGGAGCGGTGAAATGCGTTGATATCATCAGGAACACCGGTGGCGAAGGCGGCTCACTGGGTCTCTTCTGACGCTGAGGCACGAAAGCTAGGGG |
| P-asv3564 | GCAAGCGTTGTTCGGATTTATTGGGCGTAAAGAGCTCGTAGGCGGCTTGTCGCGTCGACTGTGAAAACCCACGGCCCAACCGTGCGGTCTGCAGCCGATACGGGCAGGCTAGAGTTCGGTAGGGGAGACTGGAATTCCTGGTGTAGCGGTGAAATGCGCAGATATCAGGAGGAACACCGGTGGCGAAGGCGGGTCTCTGGGCCGATACTGACGCTGAGGAGCGAAAGCGTGGGG |
| P-asv3565 | GCAAGCGTTGTTCGGAATTACTGGGCGTAAAGGGTTCGTAGGTGGCTCGCTAAGTCAGACGTGAAATCCCTCAGCTTAACTGGGGAACTGCGTCTGAGACTGGCAAGCTTGAGTGCAGGAGAGGAACGCGGAATTCCAGGTGTAGCGGTGAAATGCGTAGATATCTGGAGGAACACCGGTGGCGAAGGCGGCGCTCTGGACTGCAACTGACACTGATGAACGAAAGCCAGGGG |
| P-asv3566 | GCGAGCGTTAATCGGAATTACTGGGCGTAAAGCGTGCGTAGACGGTGATGTAAGTCGGGTGTGAAAGCCCCGGGCTCAACCTGGGAATTGCATTCGAGACTGCATTGCTTGGGTGCGGAAGAGGGAAGCGGAATTTCCGGTGTAGCGGTGAAATGCGTAGATATCGGAAGGAACACCAGTGGCGAAAGCGGCTTCCTGGTCCGACACCGACGTTCAGGCACGAAAGCGTGGGG |
| P-asv3567 | GCGAGCGTTAATCGGAATTACTGGGCGTAAAGCGTGCGTAGACGGTTACGTAAGTCGGATGTGAAATCCCCGGGCTCAACCTGGGAATTGCATTCGAGACTGTGTAGCTGGGGTGCGGAAGAGGGAAGCGGAATTTCCGGTGTAGCGGTGAAATGCGTAGATATCGGAAGGAACATCAGTGGCGAAAGCGGCTTCCTGGTCCAGCACCGACGTTCAGGCACGAAAGCGTGGGG |
| P-asv3568 | GCGAGCGTTGTTCGGAATTATTGGGCGTAAAGGGCGCGTAGGCGGTACGGTAAGTCACCTGTGAAACCTCTGGGCTTAACTCAGAGCCTGCAGGCGAAACTACCGTGCTGGAGGGTGGGAGAGGTGCGTGGAACTGCGAGTGTAGAGGTGAAATTCGTAGATATTCGCAAGAACACCAGTGGCGAAGGCGGCTCACTGGCCCGGTACTGACGCTGAGGTGCGAAAGCGTGGGG |
| P-asv3569 | GCGAGCGTTGTTCGGAATTACTGGGCGTAAAGAGTGTGTAGGCGGTGTTCCCAGTCTGTTGTGAAATCTCCCGGCTTAACTGGGAGGGTGCAGCGGAAACTAGAATGCTAGAGGGTGGGAGAGGTAAGCGGAATTCCTGGTGTAGCGGTGAAATGCGTAGATATCAGGAGGAACACCTGTGGTGTAGACAGCTTACTGGACCACTTCTGACGCTGAGACACGAAAGCGTGGGT |
| P-asv3570 | GCAAGCGTTGTTCGGAGTTACTGGGCGTAAAGGGTGTGTAGGCGGTTCTCTAAGTCTGGTGTGAAATCTCCCGGCTCAACTGGGAGGGTGCGCCGAAAACTGGAGGGCTAGAGTGCGGGATGGGGGAGTGGAATTCCTGGTGTAGCGGTGAAATGCGCAGATATCAGGAGGAACACCTGTGGTGTAGACGGCTCTCTGGACCGCAACTGACGCTGAAACACGAAAGCGTGGGT |
| P-asv3571 | CCTAGCGTTGTCCGGATTTATTGGGCGTAAAGAGCTCGTAGGCGGTTTGGTGAGTCGGGTGTGAAATCTCCACGCTCAACGTGGAGGAGCCACCCGATACTGCCATGACTAGAGTCCGGTAGGGGAGTGTGGAATTCCCGGTGTAGCGGTGAAATGCGCAGATATCGGGAGGAACACCAGTAGCGAAGGCGACACTCTGGGCCGGTACTGACGCTGAGGAGCGAAAGCGTGGGG |
| P-asv3572 | GCAAGCGTTGTTCGGAATTATTGGGCGTAAAGGGCGCGTAGGCGGTCTGGTAAGTTTGATGTGAAATCCCATGACTCAACCATGGAACTGCATCTCATACTGCCGGACTTGAGTCTGGGAGAGGGAAGTGGAATTCCCGGTGTAGCGGTGAAATGCGTAGATATCGGGAGGAACACCTGTGGCGAAAGCGGCGCACTGGACCATAACTGACGCTGAGGCGCGAAAGCTAGGGG |
| P-asv3573 | GCAAGCGTTGTTCGGAATTACTGGGCGTAAAGGGCGCGTAGGCGGTCCTTTAAGTCAGGCGTGAAAGCCCTGGGCTCAACCTGGGAATGGCGCTTGATACTGAAGGACTAGAGAGTGGGAGAGGTTGGTGGAATTCCCAGTGTAGAGGTGAAATTCGTAGATATTGGGAAGAACACCGGTGGCGAAGGCAGCCTCCTGGGTCAACACTGACGCTCATGCACGAAAGCGTGGGG |
| P-asv3574 | GCGAGCGTTGTCCGGAATCACTGGGCGTAAAGGGCGCGTAGGTGGCTCGTTAAGTGGCTGGTGAAATCCCGGGGCTCAACTCCGGGGCTGCCGGTCAGACTGGCGGGCTAGAGCACGGTAGGGGCAGATGGAATTCCCGGTGTAGCTGTGGAATGCGTAGATATCGGGAAGAACACCAGTGGCGAAGGCGTTCTGCTGGGCCGTTGCTGACACTGAGGCGCGACAGCGTGGGG |
| P-asv3575 | GCGAGCGTTGTCCGGAATTATTGTGCGTAAAGGGCTCGTAGGCGGTTTGTCGCGTCGGGAGTGAAATCCACTGGCTTAACTGGTGGCTTGCTTTCGATACGGGCAGACTGGAGGTATGCAGGGGAGAACGGAATTCCTGGTGTAGCGGTGAAATGCGCAGATATCAGGAGGAACACCGGTGGCGAAGGCGGTTCTCTGGGCATGTCCTGACGCTGAGGAGCGAAAGTGTGGGG |
| P-asv3576 | GCGAGCGTTGTCCGGAATTACTGGGCGTAAAGAGCTTGTAGGTGGTTTGTCGCGTTGTTCGTGAAAACTCACAGCTCAACTGTGGGCGTGCGGGCGATACGGGCAGACTAGAGTACTGCAGGGGAGACTGGAATTCCTGGTGTAGCGGTGGAATGCGCAGATATCAGGAGGAACACCGGTGGCGAAGGCGGGTCTCTGGGCAGTAACTGACGCTGAGAAGCGAAAGCGTGGGG |
| P-asv3577 | GCAAGCGTTGTCCGGATTCATTGGGCGTAAAGAGCTCGTAGGCGGTTCGATAAGTCGGGTGTGAAACCTCCAGGCTCAACCTGGAGCCGCCACCCGATACTGTCGTGACTAGAGTCCAGTAGGGGAGCGTGGAATTCCTGGTGTAGCGGTGAAATGCGCAGATATCAGGAGGAACACCAGCGGCGAAGGCGGCGCTCTGGGCTGGAACTGACGCTGAGGAGCGAAAGCGTGGGG |
| P-asv3578 | GCTAGCGTTGCTCGGAATCACTGGGCGTAAAGGGCGCGTAGGCGGCCATTCAAGTCGGGGGTGAAAGCCCAGGGCTCAACCCTGGAATTGCCTTCGATACTGTTTGGCTAGAGACCGGAAGAGGTTAGTGGAACTGCGAGTGTAGAGGTGAAATTCGTAGATATTCGCAAGAACACCAGTGGCGAAGGCGGCTAACTGGTCCGGTTCTGACGCTGAGGCGCGAAAGCGTGGGG |
| P-asv3579 | GCGAGCGTTGTCCGGATTTATTGGGCGTAAAGAGCTCGTAGGCGGTTTGGTAGGTCTGGTGTGAAATCTCCACGCTCAACGTGGAGGGGCCACCCGATACTGCCATGACTGGAGTCCGGTAGAGGAGTGTGGAATTCCCGGTGTAGCGGTGAAATGCGCAGATATCGGGAGGAACACCCGTAGCGAAGGCGGCACTCTGGGCCGGTACTGACGCTGAGGAGCGAAAGCGTGGGG |
| P-asv3580 | GCAAACGTTGCTCGGAATTACTGGGCGTAAAGCGCGTGTAGGCGGACCTGCAAGTCGGACGTGAAAGCCCTGGGCTCAACCCAGGAACTGCATCCGAAACTGCGGGCCTGGAGTGGTGGAGAGGGAAGCAGAATTCCTGGTGTAGAGGTGAAATTCGTAGATATCAGGAGGAACACCGGTGGCGAAGGCGGCTTCCTGGACACTTACTGACGCTGAGACGCGAAAGCGTGGGG |
| P-asv3581 | GCAAGCGTTGTTCGGAATTATTGGGCGTAAAGGGCGCGTAGGCGGTGCTGTAAGTCTGTGGTGAAATCTCTGGGCTCAACTCAGAGTCTGCCATAGAAACTGCAGTGCTGGAGTGTGGGAGAGGTGAGTGGAATTCCCGGTGTAGCGGTGAAATGCGTAGATATCGGGAGGAACACCTGTGGCGAAAGCGGCTCACTGGACCACAACTGACGCTGGGGCGCGAAAGCTAGGGG |
| P-asv3582 | GCGAGCGTTGTCCGGAATTATTGGGCGTAAAGGGCATGCAGGCGGTCTGCTAAGTTCAGAGTGACAGTCATCAGCTTAACTGGTGAAGTACTTTGAAAACTGGTGGACTTGAGAGCGAGAGAGGGACATGGAATTCCGGGTGGAGCGGTGAAATGCGTAGAGATCCGGAGGAACACCAAGGCGAAGGCAGTGTCCTGGCTCGTTTCTGACGCTGAGGTGCGAAAGCTAGGGG |
| P-asv3583 | GCGAGCGTTAATCGGAATTACTGGGCGTAAAGCGCGCGTAGGCGGCTTTGCAAGTCGGGTGTGAAATCCCCAGGCCTAACCTGGGAACTGCATTCGAGACTGCATGGCTAGAGTATCGGAGAGGGAAGTGGAATTTCAGGTGTAGCGGTGAAATGCGTAGATATCTGAAGGAACATCAGTGGCGAAAGCGACTTCCTGGACGAATACTGACGCTCATGTGCGAAAGCGTGGGG |
| P-asv3584 | GCAAGCGTTGTTCGGAATTACTGGGCGTAAAGGGTGCGTAGGCGGGTCTGTAAGTCGAGGTTGAAATATCCGGGCTCAACTCGGGGCACGGCCTCGATACTGCAGATCTCGAGTCGAGAAGAGGGTAGTGGAATTCCCGGTGTAGCGGTGAAATGCGTAGATATCGGGAGGAACACCAGTGGCGAAGGCGGCTACCTGGTCTTGTACTAACGCTGATGCACGAAAGCTGGGGG |
| P-asv3585 | GCAAGCGTTGTCCGGAATTATTGGGCGTAAAGAGCTCGTAGGCGGTCTGTCGCGTCGAATGTGAAAACCCGGGGCTCAACCCCGGGCCTGCATTCGATACGGGCAGACTAGAGTTCGGTAGGGGAGTCTGGAATTCCTGGTGTAGCGGTGAAATGCGCAGATATCAGGAGGAACACCGGTGGCGAGGGCGGGACTCTGGGCCGATACTGACGCTGAGGAGCGAAAGCGTGGGG |
| P-asv3586 | CCGAGCGTTAGGCGGAATTACTGGGCGTAAAGCGTGCGCAGGCGGTTCGTTGTGTCGGTGGTGAAAGCCCCGGGCTTAACCTGGGAATGGCCATGGAAACTGGCGAACTGGAGTGTGGCAGAGGGGGGTGGAATTCCGCGTGTAGCAGTGAAATGCGTAGAGATGCGGAGGAACACCGATGGCGAAGGCAGCCCCCTGGGCTAGCACTGACGCTCAGGCACGAAAGCGTGGGG |
| P-asv3587 | GCGAGCGTTGTCCGGAATTATTGGGCGTAAAGGGCTCGTAGGCGGTTTGTCGCGTCGGGAGTGAAAACCAGGTGCTTAACACCTGGCCTGCTTCCGATACGGGCAGACTAGAGGTATTCAGGGGAGAACGGAATTCCTGGTGTAGCGGTGAAATGCGCAGATATTAGGAGGAACACCGGTGGCGAAGGCGGTTCTCTGGGAATGACCTGACGCTGAGGAGCGAAAGTGTGGGG |
| P-asv3588 | GCGAACGTTGTTCGGAATTACTGGGCGTAAAGCGCGTGTAGGCGGCAACGCAAGTCGGGCGTGAAATCCCACGGCTCAACCGTGGAAGTGCACTCGAAACTGCGTAGCTAGAGTCCTGGAGAGGAAGGTGGAATGCTTGGTGTAGAGGTGAAATTCGTAGATATCAAGCGGAACACCGGTGGCGAAGGCGGCCTTCTGGACAGTGACTGACGCTGAGACGCGAAAGCGTGGGG |
| P-asv3589 | GCGAGCGTTATCCGGATTCACTGGGTTTAATGGGTGCGTAGGTGGGGAAGTAAGTCAGTGGTGAAATCTCCGTGCTTAACATGGAAACTGCCATTGATACTATTTTTCTCGAATTTTCCGGAGGTTAGCGGAATATGTCATGTAGCGGTGAAATGCTTAGATATGACATAGAACACCGATTGCGGAGGCAGCTAACTACAGGGATATTGACACTGAGGCACGAAAGCGTGGGG |
| P-asv3590 | GCTAGCGTTGCTCGGAATCACTGGGCGTAAAGCGCACGTAGGCGGCTTTTTAAGTCAGGGGTGAAATTCTGGAGCCCAACTCCAGAACTGCCCTTGATACTGAGAGGCTCGAGTCCGGGAGAGGTGAGTGGAACTGCGAGTGTAGAGGTGAAATTCGTAGATATTCGCAAGAACACCAGTGGCGAAGGCGGCTCACTGGCCCGGTACTGACGCTGAGGTGCGAAAGCGTGGGG |
| P-asv3591 | GCAAGCGTTGTTCGGAATTACTGGGCGTAAAGCGCGGGCAGGTGGCCCGTTAAGTCTGCGGTGAAATCCCGGGGCTCAACTCCGGATATGCCGCGGATACTGGCGGGCTAGAGACTGGTAGAGGCGAGTGGAATTCCAGGTGTAGCGGTGGAATGCGTAGATATCTGGAAGAACACCGGTGGCGAAGGCGGCTCGCTGGGCCAGTTCTGACACTCATCCGCGAAAGCGTGGGG |
| P-asv3592 | GCCAGCGTTGCTCGGAATTACTGGGCGTAAAGGGTTCGTAGGTGGTGCGACAAGTTGGAAGTGAAATCTCTGGGCTTAACCCAGAGGCTGCTTTCAAAACTGCCGTGCTCGAGTGCGAGAGAGGTCAGTGGAATTGCAGGTGTAGCGGTGAAATGCGTAGATATTGGAAAGAACACCGGTGGCGAAGGCGGCAACCTGGTCCTTGACTGACGCTGAGGCGCGAAAGCGTGGGG |

| | |
|---|---|
| P-asv3593 | GCGAGCGTTGTCCGGAGTTACTGGGCGTAAAGGGCACGCAGGCGGTCGTGTAAGTTTGGGGTTACAGCTCCCGGCTCAACTGGGAGAGCGCCTCGAAGACTGCAGGACTTGAGGGCCAGAGAGGGACACAGAATTCCGGGTGTAGTGGTGAAATGCGTAGATATCCGGAGGAATACCAAAGGCGAAGGCAGTGTCCTGGCTGGTTACTGACGCTGAGGTGCGAAAGCTAGGGG |
| P-asv3594 | GCAAGCGTTACTCGGAATTACTGGGCGTAAAGCGTGCGTAGGTGGTGGTTTAAGTCAGATGTGAAAGCCCCGGGCTCAACCTGGGAACTGCATTTGATACTGGATCACTAGAGTGCGGTAGAGGATAGCGGAATTCCCGGTGTAGCAGTGAAATGCGTAGAGATCGGGAGGAACATCCGTGGCGAAGGCGGCTATCTGGGCCAGCACTGACACTGAGGCACGAAAGCGTGGGG |
| P-asv3595 | GCGAGCGTTGTCCGGAATTACTGGGCGTAAAGAGCTCGTAGGCGGTTTGTCACGTCGTCTGTGAAAACCCGAGGCTTAACCTCGGGCCTGCAGGCGATACGGGCAGACTTGAGTACTGTAGGGGAGACTGGAATTCCTGGTGTAGCGGTGGAATGCGCAGATATCAGGAGGAACACCGGTGGCGAAGGCGGGTCTCTGGGCAGTAACTGACGCTGAGGAGCGAAAGCGTGGGT |
| P-asv3596 | GCGAGCGTTGACCGGAATCACTGGGCGTAAAGGGCGCGTAGGCGGCTTGGTAAGGGGGCGGTGAAAGCCCGGGGCTCAACCCCGGGTCTGCCGGACCGACTGCCGAGCTTGAGCACACTAGGGGCGGGTGGAATTCCGGGTGTAGCGGTGGAATGCGTAGAGATCCGGAAGAACACCGGTGGCGAAGGCGGCCCGCCGGGGTGTAGCTGACGCTGATGCGCGACAGCGTGGGG |
| P-asv3597 | GCGAGCGTTAATCGGAATTACTGGGCGTAAAGCGTGCGCAGGCGGTTATGTAAGACAGATGTGAAATCCCCGGGCTCAACCTGGGAACTGCATTAGTGACTGCATAGCTGGAGTACGGCAGAGGGGGATGGAATTCAGCGTGTAGCAGTGAAATGCGTAGATATGCGGAGGAACACCGATGGCGAAGGCAATCCCCTGGGCCTGTACTGACGCTCATGCACGAAAGCGTGGGG |
| P-asv3598 | GCAAGCGTTGTTCGGAATTACTGGGCGTAAAGGGCTAGTAGGTGGCCAACTAAGTCAGACGTGAAATCCCTCAGCTTAACTGGGGAACTGCTTCTGATACTGGATGGCTTGAGTTTGGGAGAGGGATGCGGAATTCCAGGTGTAGCGGTGAAATGCGTAGATATCTGGAGGAACACCGGTGGCGAAGGCGGCATCCTGGACCAACACTGACACTGAATAGCGAAAGCCAGGGG |
| P-asv3599 | GCAAGCGTTAATCGGAATCACTGGGCGTAAAGGGTGCGTAGGCGGTACGTTAGGTCTGTCGTGAAATCCCCGGGCTCAACCTGGGAATGGCGGTAGAAACCGGCGCACTAGAGTATGGGAGAGGGTGGTGGAATTCCAGGTGTAGCAGTGAAATGCGTAGAGATCTGGAGGAACATCAGTGGCGAAGGCGGCCAACTGGCCCAATACTGACGCTGAGGCACGAAAGCGTGGGG |
| P-asv3600 | GCGAGCGTTGTCCGGAATTACTGGGCGTAAAGGGCTCGTAGGTGGTTTGTCGCGACGTCTGTGAAATTCTGGGGCTTAACTCCGGGCGTGCAGGCGATACGGGCATAACTTGAGTGCTGTAGGGGTAACTGGAATTCCTGGTGTAGCGGTGAAATGCGCAGATATCAGGAGGAACACCGATGGCGAAGGCAGGTTACTGGGCAGTTACTGACGCTGAGGAGCGAAAGCATGGGT |
| P-asv3601 | GCAAGCGTTGTCCGGATTTATTGGGCGTAAAGCGCTCGCAGGCGGTTCGATGCGTCGTGCGTGAAATCTCTCGGCTCAACTGGGAGGGGTCGTGCGATACGGTCGGACTGGAGGCCAGTAGGGGGACGCGGAATTCCGGGTGTAGTGGTGGAATGCGTAGAGATCCGGAGGAACACCAGCGGCGAAGGCGGCGTCCTGGACTGGTACTGACGCTCAGGAGCGAAAGCGTGGGG |
| P-asv3602 | GCAAGCGCTAATCGGAATTACTGGGCGTAAAGCGCACGCAGGCGGTTTGTTAAGTCAGATGTGAAATCCCCGCGCTTAACGTGGGAACTGCATTTGAAACTGGCAAGCTAGAGTCTTGTAGAGGGGGGTAGAATTCCAGGTGTAGCGGTGAAATGCGTAGAGATCTGGAGGAATACCGGTGGCGAAGGCGGCCCCCTGGACAAAGACTGACGCTCAGGTGCGAAAGCGTGGGG |
| P-asv3603 | GTTAGCGTTGTTCGGAATGACTGGGCGTAAAGCGCGCGTAGGCGGATCGCAAAGTTGGGGGTGAAAGCCCAGGGCTCAACCCTGGAATTGCCTTCAAGACTCGCGGTCTGGAGTTCGAGAGAGGTGGGTGGAATTCCGAGTGTAGAGGTGAAATTCGTAGATATTCGGAGGAACACCAGTGGCGAAGGCGGCCCACTGGCTCGACACTGACGCTGAGGCGCGAAAGCGCGGGG |
| P-asv3604 | GCAAGCGTTAATCGGAATTACTGGGCGTAAAGCGTGCGCAGGCGGTTTGCTAAGACCGATGTGAAATCCCCGGGCTCAACCTGGGAACTGCATTGGTGACTGGCAGGCTAGAGTATGGCAGAGGGGGGTAGAATTCCACGTGTAGCAGTGAAATGCGTAGAGATGTGGAGGAATACCGATGGCGAAGGCAGCCCCCTGGGCCAATACTGACGCTCATTCACGAAAGCGTGGGG |
| P-asv3605 | GCAAGCGTTGTCCGGAATTATTGGTCGTAAAGAGCGCGTAGGCGGTTTGTCGCGTCTGCTGTGAAAACGCAAGACTTAACCTTGCGCCTGCAGAGGGTACGGGCAGACTAGAGTGCGGTAGGGGAGACTGGAATTCCTGGTGTAGCGGTGGAATGCGCAGATATCAGGAGGAACACCGATGGCGAAGGCAGGTCTCTGGGCCGTTACTGACGCTGAGGAGCGAAAGCATGGGG |
| P-asv3606 | TCCCTCTTTCCTCAGAATTACTGGGCGTAAAGCGTGCGCAGGCGGTTTTGTAAGTCTGATGTGAAATCCCCGGGCTCAACCTGGGAACTGCATTGGAGACTGCAAGGCTAGAGTGTGTCAGAGGGGGGTAGAATTCCACGTGTAGCAGTGAAATGCGTAGAGATGTGGAGGAATACCGATGGCGAAGGCAGCCCCCTGGGATAACACTGACGCTCATGCACGAAAGCGTGGGG |
| P-asv3607 | GCAAGCGTTACTCGGAATTACTGGGCGTAAAGCGTGCGTAGGTGGTGGTTTAAGTCCGTTGTGAAAGCCCTGGGCTCAACCTGGGAATTGCAGTGGATACTGGGTCACTAGAGTGTGGTAGAGGATAGTGGAATTTCCGGTGTAGCAGTGAAATGCGTAGAGATCGGAAGGAACATCTGTGGCGAAGGCGACTATCTGGGCCAACACTGACACTGAGGCACGAAAGCGTGGGG |
| P-asv3608 | CCGAGCGTTATTCGGAATCACTGGGCGTAAAGAGTACGTAGGCGGTTTGGTAAGTCAGGTGTGAAATCCCGGAGCTCAACTCCGGAACTGCATCCGATACTGCTAGACTAGAGTAATGGAGAGGCAGGTGGAATTCTCGGTGTAGCAGTGAAATGCGTGGATATCGAGAGGAAGACCAATGGCGAAGGCAGCCTGCTGGACATTTACTGACGCTGAGGTACGAAGGCCAGGGT |
| P-asv3609 | GCAAGCGTTAATCGGAATTACTGGGCGTAAAGGGTGCGTAGGTGGTTTAATAAGTTATCTGTGAAATTCCTGGGCTTAACCTGGGGTGGTCAGATAAGACTGTTAGACTCGAGTATAGGAGAGGGTAGTGGAATTTCCGGTGTAGCGGTGAAATGCGTAGAGATCGGAAGGAACACCAGTGGCGAAGGCGGCTACCTGGCCTAATACTGACACTGAGGCACGAAAGCGTGGGG |
| P-asv3610 | GCGAGCGTTGTTCGGAATCACTGGGCGTAAAGAGTGCGTAGGCGGCTTGATAAGTCAGGAGTGAAATCCCCAGGCTCAACCTGGGAATTGCTTTTGAAACTATCTGGCTAGAATCTCGGAGAGGTTGGCGGAATTCCAAGTGTAGAGGTGAAATTCGCAGATATTTGGAGGAACACCAGTGGCGTAAGCGGCCAACTGGACGAGTATTGACGCTGAGGCACGAAAGCGTGGGG |
| P-asv3611 | GCGAGCGTTGTTCGGAATCATTGGGCGTAAAGCGTATGTAGGCGGACAAGAAAGTCAGAAGTGAAATCCCAGGGCTTAACCCCGGAATTGCTTTTGAAACTCCTTGTCTAGAGTCCCGGAGGGGCCAGCGGAATTCCGAGTGTAGAGGTGAAATTCGCAGATATTCGGAGGAACACCAGTGGCGTAGGCGGCTGGCTGGACGGGAACTGACGCTGAGATACGAAAGCATGGGT |
| P-asv3612 | CCGAGCGTTATCCGGAATTATTGGGTTTAAAGGGTGCGTAGGCGGCGTGCTAAGTCAGGGGTGAAAGACGGTGGCTCAACCATCGCAGTGCCTTTGATACTGGCATGCTTGAATGTACTTGAGGTAGGCGGAATGTGGCAAGTAGCGGTGAAATGCATAGATATGCCACAGAACACCGATTGCGAAGGCAGCTTACCAAGGTATGATTGACGCTGAGGCACGAAAGCGTGGGG |
| P-asv3613 | GCAAGCGTTAATCGGAATTACTGGGCGTAAAGCGCACGTAGGTGGTTTGTTAAGCTAGCTGTGAAATCCCCGGGCTCAACCTGGGAATGGCGGTAGGAACCGGCGCACTAGAGTATGGGAGAGGGTGGTGGAATTCCAGGTGTAGCGGTGAAATGCGTAGAGATCTGGAGGAACATCAGTGGCGAAGGCGGCCACCTGGCCCAATACTGACGCTGAGGCACGAAAGCGTGGGG |
| P-asv3614 | GCTAGCGTTGTTCGGAATTACTGGGCGTAAAGCGAGTGTAGGCGGTTGCCCAAGTCAGGTGTGAAAGCCTTGAGCTCAACTCAAGAAATGCACTTGGTACTGGGTGGCTAGAGGACCGGAGAGGATAGTGGAATTCCCAGTGTAGTGGTGAAATACGTAGAGATTGGGAAGAACACCAGTGGCGAAGGCGGCTATCTGGACGGTTACTGACGCTAAGACTCGAAAGCGTGGGG |
| P-asv3615 | GCGAGCGTTGTCCGGATTTATTGGGTTTAAAGGGTGCGTAGGTGGCTATTTAAGTCAGTGGTGAAATACGGTTGCTCAACAATCGAGGTGCCATTGATACTGAATAGCTTGAAATAATTGGAGGCTGCCGGAATGGATGGTGTAGCGGTGAAATGCATAGATATCATCCAGAACACCGATTGCGAAGGCAGGTGGCTACGATTTGTTTGACACTGAGGCACGAAAGCATGGGG |
| P-asv3616 | GCAAGCGTTGTCCGGAATCACTGGGTGTAAAGGGTGTGTAGGCGGGATTGTAAGTCAGAGGTGAAATCTTGCGGCTCAACCGTAAAATTGCCTTTGATACTGCAGTTCTTGAGTCCCGGAGAGGTTGTTGGAATTCGTGGTGTAGCGGTGAAATGCGTAGATATCACGAGGAACACCAGAGGCGAAAGCGAACAACTGGACGGGTACTGACGCTAAGGCACGAAAGCGTGGGG |
| P-asv3617 | GCAAGCGTTGTCCGGAATTATTGGGCGTAAAGAGCTCGTAGGCGGTTTGTCGCGTCTGGTGTGAAATCCCGAGGCTCAACCTCGGGCTTGCATCGGGTACGGGCAGACTAGAGTACTGTAGGGGAGACTGGAATTCCTGGTGTAGCGGTGGAATGCGCAGATATCAGGAGGAACACCGATGGCGAAGGCAGGTCTCTGGGCAGTTACTGACGCTGAGGAGCGAAAGCATGGGG |
| P-asv3618 | GCGAACGTTATTCGGATTCACTGGGCTTAAAGCGCGTGTAGGCGGACACGCACGTCCGAGGCTGAAAGCCCCCGGCTCAACCGGGGAACGGGCCCGGATACGGCGTGTCTGGAGGGGCATAGGGGGAGCTGGAACTTCCGGTGGAGCGGTGAAATGCGTTGAGATCGGAAGGAACGCCCGTGGCGAAAGCGAGCTCCTGGATGCCTTCTGACGCTGAGACGCGAAAGCCAGGGG |
| P-asv3619 | GCAAGCATTGATCGGAATTACTGGGCGTAAAGGGCGCGTAGGCGGCAAGAAAAGTCAGATGTGAAATTCCAGGGCTCAACCCTGGAGCTGCATTTGAAACTATCTAGCTAGAGTATAGTTAGGGTAAACGGAATTCCACGTGTAGCGGTGAAATGCGTAGATATGTGGAAGAACACCGGTGGTGAAGACGGTTTACTGGGCTAAAACTGACGCTGAGGCGCGAAAGCATGGGG |
| P-asv3620 | GCTAGCGTTGCTCGGAATCACTGGGCGTAAAGCGCACGTAGGCGGCTTTTTAAGTCAGGGGTGAAATCCTGGAGCTCAACTCCAGAACTGCCTTTGATACTGAGGAGCTCGAGTCCGGGAGAGGTGAGTGGAACTGCGAGTGTAGAGGTGAAATTCGTAGATATTCGCAAGAACACCAGTGGCGAAGGCGGCTCACTGGCCCGGTACTGACGCTGAGGTGCGAAAGCGTGGGG |
| P-asv3621 | GCGAGCGTTGTCCGGATTTATTGGGCGTAAAGGGCTCGTAGGCGGTTCGTCGCGTCGGATGTGAAAACTCGGGGCTTAACTTCGAGCTTGCATTCGATACGGGCGTAACTAGAGGATGGCAGGGGAGACTGGAATTCCTGGTGTAGCGGTGGAATGCGCAGATATCAGGAGGAACACCGGTGGCGAAGGCGGGTCTCTGGGCCATTACTGACGCTGAGGAGCGAAAGCGTGGGG |
| P-asv3622 | GCAAGCGTTATCCGGATTTATTGGGTTTAAAGGGTCCGTAGGCGGGCCCGTAAGTCAGTGGTGAAATCTCACAGCTTAACTGTGAAACTGCCGTTGATACTGCGGGTCTTGAGTGTAGTTGAGGTAGCTGGAATGAGTAGTGTAGCGGTGAAATGCATAGATATTACTCAGAACACCAATTGCGAAGGCAGGTTACCAAGTTACAACTGACGCTGAGGGACGAAAGCGTGGGG |
| P-asv3623 | GCTAGCGTTACTCGGAATTATTGGGCGTAAAGCGTGCGTAGGCGGCTTCATAAGTTGGAAGTGAAAGCCCAGGGCTCAACCCTGGAATTGCTTTCAAAACTGTAAAGCTAGAATGTCAGAGGGGATAGCGGAATTCCTAGTGTAGAGGTGAAATTCGTAGATATTAGGAGGAACACCGGAGGCGAAAGCGGCTATCTGGCTGAACATTGACGCTGTGGCACGAAAGCGTGGGG |
| P-asv3624 | TCTCTCTTTCCTCGGATTTACTGGGCGTAAAGCGTGCGTAGGCGGCTTTTTAAGTCGGGTGTGAAATCCCTGAGCTTAACTTAGGAATTGCATTCGATACTGGGAAGCTAGAGTATGGGAGAGGATGGTAGAATTCCAGGTGTAGCGGTGAAATGCGTAGAGATCTGGAGGAATACCGATGGCGAAGGCAGCCATCTGGCCTAATACTGACGCTGAGGTACGAAAGCATGGGG |
| P-asv3625 | GCAAGCGTTAATCGGAATCACTGGGCGTAAAGGGTGCGTAGGCGGTACGTTAGGTCTGTCGTGAATCCCCCGGGCTCAACCTGGGAATGGCGATAGAAACCGGCGCACTAGAGTATGGGAGAGGGTGGTGGAATTCCAGGTGTAGCGGTGAAATGCGTAGAGATCTGGAGGAACATCAGTGGCGAAGGCGGCCACCTGGCCCAATACTGACGCTGAGGCACGAAAGCGTGGGG |
| P-asv3626 | GCAAGCGTTGTCCGGAATTATTGGGCGTAAAGCGCGCGCAGGCGGTCACTTAAGTCCATCTTAGAAGTGCGGGGCTTAACCCCGTGATGGGATGGAAACTGGGAGACTGGAGTATCGGAGAGGAAAGTGGAATTCCTAGTGTAGCGGTGAAATGCGTAGATATTAGGAAGAACACCGGTGGCGAAGGCGACTTTCTGGACGAAAACTGACGCTGAGGCGCGAAAGCGTGGGG |
| P-asv3627 | GCGAGCGTTATTCGGAATCACTGGGCGTAAAGCGTGCGTAGGCGGCCTAGCAAGTTAGAAGTGAAAGCCCAGGGCTCAACCCTGGAATTGCTTTTAAAACTGCAAGGCTAGAATCTCGGAGAGGATAGCGGAATTTCCAGTGTAGCAGTGAAATGCGTAGATATTGGAAGGAACACCAGTGGCGTAAGCGGCTATCTGGACGAGTATTGACGCTGAGGCACGAAAGCGTGGGG |
| P-asv3628 | GCGAGCGTTAATCGGAATCACTGGGCGTAAAGGGCGCGTAGGCGGTTAGTTACGTCTGCCGTGAAATCCCCGGGCTCAACCTGGGAATGGCGGTGGAAACGGGCTGACTAGAGTATGGGAGAGGGTGATGGAATTCCAGGTGTAGCGGTGAAATGCGTATAGATCTGGAGGAACATCAGTGGCGAAGGCGGTCACCTGGCCCAATACTGACGCTGAGGCGCGAAATCGTGGGG |
| P-asv3629 | GCGAGCGTTATCCGGAATTATTGGGCGTAAAGAGCTCGTAGGCGGTTTGTCGCGTCTGCTGTGAAAGCCCGGGGCTTAACCCCGGGTGTGCAGTGGGTACGGGCAGACTAGAGTGCAGTAGGGGAGACTGGAATTCCTGGTGTAGCGGTGAAATGCGCAGATATCAGGAGGAACACCGATGGCGAAGGCAGGTCTCTGGGCTGTTACTGACGCTGAGGAGCGAAAGCATGGGG |
| P-asv3630 | GCGAACGTTGTTCGGAATCACTGGGCTTAAAGGGCGCGTAGGCGGCTTGTCAAGTCAGGGGTGAAATCTTTCGGCTTAACCGGAAAAGTGCCCTTGATACTGACAGGCTCGAGGGAGGTAGGGGCATGTGGAACTTCCGGTGGAGCGGTGAAATGCGTTGATATCGGAAGGAACGCCGGTGGCGAAAGCGACGTGCTGGACCTCTTCTGACGCTGAGGCGCGAAAGCTAGGGG |
| P-asv3631 | GCGAGCGTTGTTCGGATTTACTGGGCGTAAAGGGCGCGTAGGCGGCCCTCTTTGTCGCGTGTGAAAGCCCCGGGCTCAACCTGGGAATTGCGCGCGAAACGGGGGGGCTGGAGTTCGGGAGAGGAGCGTGGAATTCCCAGTGTAGAGGTGAAATTCATAGATATTGGGAAGAACACCGGTGGCGAAGGCGGCGCTCTGGACCGAGACTGACGCTGAGGCGCGAAAGCGTGGGG |

| | |
|---|---|
| P-asv3632 | GCTAGCGTTGTCCGGAATCATTGGGCGTAAAGCGCGTGTAGGCGGTTCGATTAGTCCGCTGTAAAAGTCGGGGGCTCAACCCTCGAAAGCCGGTGGATACTGTCGGGCTAGAGTGCGGAAGAGGCGAGTGGAATTCCTGGTGTAGCGGTGAAATGCGCAGATATCAGGAGGAACACCAATTGCGAAGGCAGCTCGCTGGGACGTTACTGACGCTGAGACGCGAAAGCGTGGGG |
| P-asv3633 | GCGAACGTTGTTCGGAATCACTGGGCTTAAAGGGCGCGTAGGCGGGCCATCAAGTCCGGGGTGAAAGCCCCCAGCTTAACTGGGGAAGTGCCTTGGATACTGGTGGTCTAGAGGGAGGTAGGGGCCTGTGGAACTTCGGGTGGAGCGGTGAAATGCGTTGATATCCGAAGGAACGCCGGTGGCGAAAGCGACGGGCTGGATCTCTACTGACGCTGAGGCGCGAAAGCCAGGGG |
| P-asv3634 | GCGAGCGTTAATCGGAATTACTGGGCGTAAAGCGTGCGCAGGCGGTGATGTAAGACCGATGTGAAATCCCCGGGCTTAACCTGGGAACTGCATTGGTGACTGCATTGCTAGAGTATGGCAGAGGGGGGTGGAATTCCACGTGTAGCAGTGAAATGCGTAGAGATGTGGAGGAACACCGATGGCGAAGGCAGCCCCCTGGGCCAATACTGACGCTCATGCACGAAAGCGTGGGG |
| P-asv3635 | GCCAGCGTTGCTCGGAATTACTGGGCGTAAAGGGTCTGTAGGCGGTGCGGCAAGTCGGAAGTGAAATCTCAGGGCTTAACTCTGGGGCTGCTCCCGAAACTGCCGTGCTAGAGTGTGAGATAGGCGAGTGGAATTGCAGGTGTAGCGGTGAAATGCGTAGATATCTGCAGGAACATCCGAGGCGAAAGCGGCTCGCTAGACCACAACTGACGCTGAGAGACGAAAGCTAGGGG |
| P-asv3636 | GCAAGCGTTGTTCGGAATCACTGGGCGTAAAGGGCGTGTAGGCGGTCCGCTAAGTCTGATGTGAAAGCCCGGGGCTCACCCCCGGAAGGGCATTGGAAACTGGCGGACTCGAGTCCCGGAGAGGAAGGTGGAATTCCTGGTGTAGCGGTGGAATGCGCAGATATCAGGAGGAACACCGGTGGCGAAGGCGGGATCCTGGGCCAGTACTGACGCTGAGGAGCGAAAGCGTGGGG |
| P-asv3637 | GCTAGCGTTGCTTGGAATTACTGGGCGTAAAGCGCACGTAGGCGGCTTTCTAAGTCGGGGGTGAAATCCTGGAGCTCAACTCCAGAACTGCCTTCGATACTGGAAAGCTCGAGTCCGGGAGAGGTGAGTGGAACTGCGAGTGTAGAGGTGAAATTCGTAGATATTCGCAAGAACACCAGTGGCGAAGGCGGCTCACTGGCCCGGTACTGACGCTGAGGTGCGAAAGCGTGGGG |
| P-asv3638 | GCTAGCGTTGCTCGGAATTACTGGGCGTAAAGGGCGCGTAGGCGGATGTTTAAGTCAGAGGTCAAATCCCAGGGCTCAACCTTGGAACTGCCTTTGATACTGAACGTCTAGAGTTCGGGAGAGGTGTGTGGAACGCCGAGTGTAGAGGTGAAATTCGTAGATATTCGGCAGAACACCAGTGGCGAAGGCGACACACTGGCCCGATACTGACGCTGAGGCGCGAAAGCGTGGGG |
| P-asv3639 | GCAAGCGTTGTCCGGAATTATTGGGCGTAAAGAGCTCGTAGGCGGTCTGTCACGTCGGCTGTGAAAACCTGGGGCTCAACTCCGGGCCTGCAGCCGATACGGGCAGACTAGAGTTCGGCAGGGGAGACTGGAATTCCTGGTGTAGCGGTGAAATGCGCAGATATCAGGAGGAACACCGGTGGCGAAGGCGGGTCTCTGGGCCGATACTGACGCTGAGGAGCGAAAGCGTGGGG |
| P-asv3640 | GCAAGCGTTGTTCGGAATTACTGGGCGTAAAGGGCGCGTAGGCGGTCTTGTAAGTCTTGTGTGAAATCCCTCGGCTTAACTGAGGAACTGCACGGGATACTGCGAGACTTGAGATCGGGAGAGGGAAGCGGAATTCCGGGTGTAGCGGTGAAATGCGTAGATATCCGGAGGAACACCGGTGGCGAAGGCGGCTTCCTGGACCGACACTGACGCTGAGGCGCGAAAGCTAGGGG |
| P-asv3641 | GCAAGCGTTGTCCGGAATTATTGGGCGTAAAGAGCGTGTAGGCGGCTTGATAAGTCTGCTGTGAAAGTCCAGGGCTCAACCCTGGAAAGCCGGTGGATACTGTCAAGCTAGAGTCCGGAAGGGGCGAGTGGAATTCCTGGTGTAGCGGTGAAATGCGCAGATATCAGGAGGAACACCAATGGCGAAGGCAGCTCGCTGGGACGTGACTGACGCTGAGACGCGAAAGCGTGGGG |
| P-asv3642 | GCGAGCGTTGTTCGGAATTATTGGGCGTAAAGGGCGCGTAGGCGGCGCGGTAAGTCACCTGTGAAACCTCTGGGCTCAACTCAGAGCCTGCAGGCGAAACTGCCGTGCTGGAGTGTGGGAGAGGTGCGTGGAATTCCCGGTGTAGCGGTGAAATGCGTAGATATCGGGAGGAACACCTGTGGCGAAAGCGGCGCACTGGACCACAACTGACGCTGAGGCGCGAAAGCTAGGGG |
| P-asv3643 | GCGAGCGTTAATCGGAATTACTGGGCGTAAAGCGCGCGTAGACGGTTTTGTAAGTCAGATGTGAAATCCCTGGGCTCAACCTGGGAACTGCATTTGATACTGCATGGCTAGAGTATGGAAGAGGGAAGTGGAATTTCCGGTGTAGCGGTGAAATGCGTAGATATCGGAAGGAACACCAGTGGCGAAAGCGACTTCCTGGTCCAATACTGACGTTCATGTGCGAAAGCGTGGGG |
| P-asv3644 | GCAAGCGTTGTCCGGAATTATTGGGCGTAAAGAGCTCGTAGGCGGCTCAGTTAGTCGGGTGTGAAATTCTCAGGCTCAACCTGGGGACGCCACCCGATACTGCTGTGGCTAGAGTCCAGTAGAGGAGTGTGGAATTCCCGGTGTAGCGGTGAAATGCGCAGATATCGGGAGGAACACCAACGGCGAAGGCAGCACTCTGGGCTGGTACTGACGCTGAGGAGCGAAAGCGTGGGT |
| P-asv3645 | GCGAGCGTTAATCGGAATTACTGGGCGTAAAGCGTGCGCAGGCGGTTTTGTAAGTCAGATGTGAAATCCCCGAGCTCAACTTGGGAACTGCGTTTGAAACTACAAGACTAGAATATGTCAGAGGGGGGTAGAATTCCACGTGTAGCAGTGAAATGCGTAGAGATGTGGAGGAACACCAATGGCGAAGGCAGCCCCCTGGGATAATATTGACGCTCATGCACGAAAGCGTGGGG |
| P-asv3646 | GCAAGCGTTGTCCGGATTTACTGGGCGTAAAGCGCCCGCAGGCGGCTTGTTAAGTTCAAGGTGAAGTCTCCCGGCTTAACTGGGAGGGTGCCTCGAAAACTGGCAGACTTGAGGTGAGGAGAGGGATGCGGAATTCCAGGTGTAGCGGTGAAATGCGTAGATATCTGGAGGAACACCAGTGGCGAAGGCGGCATCCTGGCCTTATCCTGACGCTCAGGGGCGAAAGCGTGGGG |
| P-asv3647 | GCTAGCGTTGTTCGGAATTACTGGGCGTAAAGCGCACGTAGGCGGATTTGTAAGTAGGGGGTGAAATCCCCAGGCTCAACCTGGGAACTGCCTTCTAGACTGCAAGTCTGGAGATCAGGAGAGGCGAGTGGAATACCGAGTGTAGAGGTGAAATTCGTAGATATTCGGTGGAACACCAGTGGCGAAGGCGACTCGCTGGACTGATACTGACGCTGAGGTGCGAAAGCGTGGGG |
| P-asv3648 | GCAAGCGTTGTCCGGAATTATTGGGCGTAAAGAGCTCGTAGGCGGTCTGTTGCGTCCGCTGTGAAATCTCGGGGCTTAACCCTGGACTTGCTGCGGATACGGGCAGACTAGAGGTAGGTAGGGGAGAATGGAATTCCCGGTGTAGCGGTGAAATGCGCAGATATCGGGAGGAACACCGGTGGCGAAGGCGGTTCTCTGGGCCTTACCTGACACTGAGGAGCGAAAGCGTGGGG |
| P-asv3649 | GCAAGCGTTGCCCGGAATCATTGGGCGTAAAGAGCGTGTAGGCGGCCAGAAAAGTCCGCTGTGAAAACTCGAGGCTCAACTTCGAGCTGTCGGTGGAAACTTTCTGGCTAGAGTCCGGAAGAGGAGAATGGAATTCCTGGTGTAGCGGTGAAATGCGCAGATATCAGGAAGAACACCCGTGGCGAAGGCGGTTCTCTGGGACGGTACTGACGCTGAGACGCGAAAGCGTGGGG |
| P-asv3650 | GCGAGCGTTGTTCGGAATTATTGGGCGTAAAGGGTGCGTAGGCGGTTCCGTAAGTCCTGTGTGAAATCTTCGGGCTCAACTCGAAGTCTGCACGGGAAACTGCGGGGCTTGAGTGTGGGAGAGGTGAGTGGAACTCCCGGTGTAGCGGTGAAATGCGTAGATATCGGGAGGAACACCTGTGGCGAAAGCGGCTCACTGGACCACAACTGACGCTGATGCACGAAAGCTGGGGG |
| P-asv3651 | GCAAGCGTTGTCCGGAATCATTGGGCGTAAAGCGCGCGTAGGCGGCCCGGTAAGTCCGCTCTGAAAGTCCAGGGCTCAACCCTGGAAAGCGGGTGGATACTGCCGGGCTAGAGTCCGGAAGAGGCGAGTGGAATTCCTGGTGTAGCGGTGGAATGCGCAGATATCAGGAGGAACACCAATAGCGAAGGCAGCTCGCTGGGACGGTACTGACGCTAAGGCGCGAAAGCGTGGGG |
| P-asv3652 | GCAAGCGTTGTTCGGAATTACTGGGCGTAAAGCGCATGCAGGTGGCTGAGTAAGTCGAATGTGGAAGCCCCGGGCTCAACTCGGGAAATGCATCCGAAGCTGCTCAGCTAGAGTCCCGGAGGGGAGGGCAGAATTCCCAATGTAGGGGTGAAATCCGTAGATATTGGGAGGAATACCGGTGGCGAAGGCGGCCCTCTGGACGGTGACTGACACTCAGATGCGAAAGCGTGGGG |
| P-asv3653 | ACTAGCGTTGTTCGGATTTACTGGGCGTAAAGCGCGCGTAGGCGGATCGCTAAGTCAGAGGTGAAATCCCAGGGCTCAACCCTGGAACTGCCTTTGATACTGACGATCTGGAGATCGGGAGAGGTTGGTGGAATTCCGAGTGTAGAGGTGAAATTCGTAGATATTCGGAGGAACACCAGAAGCGAAGGCGGCCAACTGGCCCGATACTGACGCTGAGGCGCGAAAGCGTGGGG |
| P-asv3654 | GCAAGCGTTGTCCGGAATCATTGGGCGTAAAGAGCGTGTAGGCGGCCCGATAAGTCCGCTGTGAAAGTCGAGGGCTCAACCCTCGAATGCCGGCGGATACTGTCGGGCTAGAGTCCGGAAGAGGCGAGTGGAATTCCTGGTGTAGCGGTGAAATGCGCAGATATCAGGAGGAACACCAACGGCGAAGGCAGCTCGCTGGGACGGTACTGACGCTGAGACGCGAAAGCGTGGGG |
| P-asv3655 | GCGAGCGTTAATCGGAATTACTGGGCGTAAAGCGCGCGTAGGCGGCTTTGCAAGTCGGGTGTGAAATCCCCAGGCTTAACCTGGGAACTGCATTCGAGACTGCATTGCTAGAGTATGGGAGAGGGAAGTGGAATTTCCGGTGTAGCGGTGAAATGCGTAGATATCGGAAGGAACATCAGTGGCGAAAGCGACTTCCTGGACCAATACTGACGCTCATGTGCGAAAGCGTGGGG |
| P-asv3656 | GCAAGCGTTGTCCGGATTTATTGGGCGTAAAGAGCGTGTAGGCGGCCAGACAGGTCCGTCGTGAAAACTAGAGGCTTAACCTCTAGACGTCGATGGAAACCGTCTGGCTAGAGTCCGGAAGAGGAGAATGGAATTCCTGGTGTAGCGGTGAAATGCGCAGATATCAGGAAGAACACCCGTGGCTAAGGCGGTTCTCTAGTACGGTACTGACGCTGAGACGCGAAAGCGTGGGG |
| P-asv3657 | GCAAGCGTTGTCCGGAATCATTGGGCGTAAAGAGCGCGTAGGCGGTCCGATAAGTCGGTTGTGAAAACTGGGGGCTCAACCCTCAGAGGCCGACCGATACTGTCGGGCTAGAGTCCGGAAGAGGCGAGTGGAATTCCTGGTGTAGCGGTGAAATGCGCAGATATCAGGAGGAACACCAATGGCGAAGGCAGCTCGCTGGGACGGTACTGACGCTGAGGCGCGAAAGCGTGGGG |
| P-asv3658 | GCAAGCGTTGTCCGGATTTATTGGGCGTAAAGCGCGTGTAGGCGGCCAGGTAGGTCGGTTGTGAAAACTGGAGGCTCAACCTCCAGACGTCGACCGAAACCATCTGGCTAGAGTCCGGAAGAGGAGAGTGGAATTCCTGGTGTAGCGGTGAAATGCGCAGATATCAGGAAGAACACCGATGGCGAAGGCAGCTCTCTGGGACGGTACTGACGCTGAGACGCGAAAGCGTGGGG |
| P-asv3659 | GCTAGCGTTGTTCGGAATTACTGGGCGTAAAGCGCGCGCAGGCGGTTTTCCAAGTCAGTGGTGAAAGCCCGGAGCTCAACTCCGGAACTGCCATTGAAACTGTTAAACTTGAGGACGAGAGAGGTGAGTGGAATTCCCAGTGTAGAGGTGAAATTCGTAGATATTGGGAAGAACACCAGTGGCGAAGGCGGCTCACTGGCTCGTTTCTGACGCTCAGGCGCGATAGCGTGGGG |
| P-asv3660 | GCGAGCGTTGTCCGGAATTACTGGGCGTAAAGAGCTTGTAGGCGGTCTGTCACGTCTTCTGTGAAAACTTGGGGCTTAACCTTAAGCTTGCAGGGGATACGGGCAGACTAGAGTACTTCAGGGGAGACTGGAATTCCTGGTGTAGCAGTGAAGTGCGTAGAGATCGGGAGGAACATCAGTGGCGAAGGCGGCCGCCTGGACCAGCACTGACACTGAGGCACGAAAGCGTGGGG |
| P-asv3661 | GCAAGCGTTGTCCGGAATCATTGGGCGTAAAGAGCGTGTAGGCGGCCCGATCAGTCCGCTGTGAAAGTCCAGGGCTCAACCCTGGAATGCCGGTGGATACTGTCGGGCTAGAGTCCGGAAGGGGCGAGTGGAATTCCTGGTGTAGCGGTGAAATGCGCAGATATCAGGAGGAACACCAATGGCGAAGGCAGCTCGCTGGGACGTGACTGACGCTGAGACGCGAAAGCGTGGGG |
| P-asv3662 | GCAAGCGTTACTCGGAATTATTGGGCGTAAAGGGCGCGTAGGCGGCCGCGTGCGTCAGAGGTGAAATCCCCCGGCTTAACCGGGGACCCGCCTCTGATACGGCGCGGCTTGAGTCCGGGAGAGGGGAGCAGAATTCCCAGTGTAGCGGTGAAATGCGTAGATATTGGGAGGAATACCGGTGGCGAAGGCGGCTCCCTGGACCGGTACTGACGCTGAGGCGCGAAAGCGTGGGT |
| P-asv3663 | GCAAGCGTTGTCCGGAATTATTGGGCGTAAAGAGCTCGTAGGCGGTCTGTCACGTCGGATGTGAAAACTCGGGGCTCAACCCCGAGCCTGCATTCGATACGGGCAGACTAGAATTCGGTAGGGGAGACTGGAATTCCTGGTGTAGCGGTGAAATGCGCAGATATCAGGAGGAACACCGGTGGCGAAGGCGGGTCTCTGGGCCGATATTGACGCTGAGGAGCGAAAGCGTGGGG |
| P-asv3664 | GCTAGCGTTGTCCGGATTTACTGGGCGTAAAGAGCGCGCAGGCGGTCGTTCAAGTCGAGTGTGAAAGCCCCCGGCTCAACTGGGGAGGGTCACTCGATACTGATCGACTTGAAGGCGGGAGAGGGAAGCGGAATTCCCGGTGTAGTGGTGAAATGCGTAGATATCGGGAGGAACACCAGTGGCGAAGGCGGCTTTCTGGCCCGTTCTTGACGCTGAGGCGCGAAAGCTAGGGT |
| P-asv3665 | GCGAACGTTGTTCGGAATCACTGGGCTTAAAGGGCGCGTAGGCGGGTTCGCAAGTCTGTGGTGAAAGCCTCCAGCTTAACTGGAGAAGTGCCGTGGATACTGCGAATCTTGAGGAGGGTAGGGGCATCTGGAACGGTCGGTGGAGCGGTGAAATGCGTTGATATCGACCGGAACTCCGGTGGCGAAGGCGAGGTGCTGGACCCTTTCTGACGCTGAGGCGCGAAAGCCAGGGG |
| P-asv3666 | GCTAGCGTTGCTCGGAATTACTGGGCGTAAAGCGCACGTAGGCGGCTTCTTAAGTCGGTGGTGAAATCCTGGAGCTCAACTCCAGAACTGCCTTCGATACTGGGAAGCTCGAGTCCGGGAGAGGTGAGTGGAACTGCGAGTGTAGAGGCGAAATTCGTAGATATTCGCAAGAACACCAGTGGCGAAGGCGGCTCACTGGCCCGGTACTGACGCTGAGGTGCGAAAGCGTGGGG |
| P-asv3667 | GCGAGCGTTGTCCGGAATCACTGGGCGTAAAGGGCGCGTAGGCGGCTCATTAAGTGGCTGGTGAAATCCCGGGGTTCAACTCCGGGGCTGCCGGTCAGACTGGTGGGCTAGAGCACGGTAGAGGCGGATGGAATTCCCGGTGTAGCGGTGGAATGCGTAGATATCGGGAAGAATACCGGTGGCGAAGGCGTTCCGCTGGGCCGTTGCTGACGCTGAGGCACGACAGCGTGGGG |
| P-asv3668 | GCGAACGTTATTCGGAATCACTGGGCTTAAAGCGCGTGTAGGCGGATGGCCGCGTCGGTCGCTCAAAGCCCCCGGCTCAACCGGGGAACCGGCGCCGAAACGGGCCGTCTGGAGGGACGTAGGGGGACCTGGAACTTCCGGTGGAGCGGTGAAATGCGTTGAGATCGGAAGGAACGCCCGCGGCGAAAGCGAGGTCCTGGACGTCACCTGACGCTGAGACGCGAAAGCCAGGGG |
| P-asv3669 | GCAAGCGTTGTCCGGATTTATTGGGCGTAAAGAGCGCGCAGGCGGTTCGTTAAGTCCGAGGTGAAATCTCCCGGCTCAACTGGGAGGGGTCTTCGGATACTGGCGGACTTGAGGAGTGCAGAGGTGAGTGGAATTCTCGGTGTAGTGGTGAAATGCGTAGATATCGAGAGGAACACCAGCGGCGAAGGCGGCTCACTGGGCACCACCTGACGCTGAGGCGCGAAAGCGTGGGG |
| P-asv3670 | GCTAGTGTTGTTCGGAATTACTGGGCGTAAAGGGCGCGCAGGCGGCTTGGCAAGTCGAGTGTGAAATCCCTTGGCTTAACTGAGGAATAGCGCTCGAAACTACCTCGCTGGAGGGCGGGAGAGGGAAGTGGAACTCTCGGTGTAGGGGTGAAATCTGTAGATATCGAGAGGAACACCGGTGGCGAAGGCGGCTTCCTGGCCCGTACCTGACGCTGAGGCGCGAAAGCGTGGGG |

| | |
|---|---|
| P-asv3671 | GCGAGCGTTAATCGGAATTACTGGGCGTAAAGCGTGCGCAGGCGGTTTCGTAAGACAGATGTGAAATCCCCGGGCTCAACCTGGGAACTGCATTTGTGACTGCGGGGCTAGAGTGTGTCAGAGGGAGGTGGAATTCCGCGTGTAGCAGTGAAATGCGTAGAGATGCGGAGGAACACCGATGGCGAAGGCAGCCTCCTGGGATAACACTGACGCTCAGGCACGAAAGCGTGGGG |
| P-asv3672 | TCAAGCGTTGTTCGGATTCATTGGGCGTAAAGGGTGCGTAGGTGGCGCGGTAAGTCGGGTGTGAAATTTCGGAGCTTAACTCCGAAACTGCATTCGATACTGCCGTGCTTGAGGACTGGAGAGGAGACTGGAATTTACGGTGTAGCAGTGAAATGCGTAGAGATCGTAAGGAAGACCAGTGGCGAAGGCGGGTCTCTGGACAGTTCCTGACACTGAGGCACGAAGGCCAGGGG |
| P-asv3673 | GCGAACGTTATCCGGATTTACTGGGCGTAAAGCGCATGCAGGCGGTTTCTTAAGTCGGACGTGAAAGCTCCTGGCTCAACTGGGAGAGGTCGTTCGAAACTGGGGAACTTGAGGATGGTAGAGGAGAGTGGAATTCCCGGTGTAGTGGTGAAATGCGTAGATATTGGGAGGAACACCAGTGGCGAAGGCGGCTCTCTGGGCCATTCCTGACGCTAAGATGCGAAAGCCAGGGG |
| P-asv3674 | GCAAGCGTTGTTCGGAATTACTGGGCGTAAAGGGCTCGTAGGTGGCCAACTAAGTCAGACGTGAAATCCCCAGGCTTAACCTGGGAACTGCGTCTGATACTGGATGGCTTGAATTCGGGAGAGGGATGCAGAATTCCAGGTGTAGCGGTGAAATGCGTAGATATCTGGAGGAATACCGGTGGCGAAGGCGGCATCCTGGACCGGCATTGACGCTGAGGAGCGAAAGCCAGGGG |
| P-asv3675 | GCCAGCGTTATTCGGAATTACTGGGCGTAAAGCGTGCGCAGGCGGTTACTTAAGTCAGGTGTGAAGGCCCTGGGCTTAACCTAGGAATTGCGCTTGAGACTGGGTAGCTTTGAGTGTGGCAGAGGGAGGTGGAATTCCACGTGTAGCAGTGAAATGCGTAGAGATGTGGAGGAACACCGATGGCGAAGGCAGCCTCCTGGGCCAGCACTGACGCTCATGCACGAAAGCGTGGGG |
| P-asv3676 | GCAAGCGTTGTTCGGAATCACTGGGCGTAAAGGGCGTGTAGGCGGTCAGGTAAGTGGGACGTGAAATGCCCCGGCTCACCCGGGGACCGGCGTCTCATACTGCTTGGCTTGAGTATGGGAGAGGATGAGGGAATTCCTGGTGTAGCGGTGAAATGCGTAGATATTCGGAAGAACACCAGTGGCGAAGGCGGCTCACTGGACGCGTATTGACGCTGAGGTACGAAAGCGTGGGG |
| P-asv3677 | GCAAGCGTTGTTCGGAATTACTGGGCGTAAAGGGTTCGTAGGTGGCTTGCTAAGTCAGACGTGAAATCCCACGGCTCAACTGTGGAAGAGCACTCGAAACTGCGTGGCTAGAGTCCTGGAGAGGAAGGTGGAATACTTGGTGTAGAGGTGAAATTCGTAGATATCAAGTGGAACACCAGTGGCGAAGGCGGCCTTCTGGACAGTGACTGACGCTGAGACGCGAAAGCGTGGGG |
| P-asv3678 | GCTAGCGTTGCTCGGAATCACTGGGCGTAAAGGGCGCGTAGGCGGCCATTCAAGTCGGGGGTGAAAGCCTGTGGCTCAACCACAGAATTGCCTTCGATACTGTTTGGCTTGAGTTTGGTAGAGGTTGGTGGAACTGCGAGTGTAGAGGTGAAATTCGTAGATATTCGCAAGAACACCAGTGGCGAAGGCGGCCAACTGGACCAACACTGACGCTGAGGCGCGAAAGCGTGGGG |
| P-asv3679 | GCAAGCGTTGTCCGGAATTATTGGGCGTAAAGAGCTCGTAGGCGGCTTGTCACGTCGGTTGTGAAAGCCCGGGGCTTAACCCCGGGTCTGCAGTCGATACGGGCAGGCTAGAGTTCGGTAGGGGAGATCGGAATTCCTGGTGTAGCGGTGAAATGCGTAGATATCAGGAGGAACACCGGTGGCGAAGGCGGATCTCTGGGCCGATACTGACGCTGAGGAGCGAAAGCGTGGGG |
| P-asv3680 | GCAAGCGTTGTCCGGATTTATTGGGCGTAAAGCGAGTGCAGGCGGCTCGATAAGTCTGATGTGAAAGCCTTCGGCTCAACCGGAGAATTGCATCAGAAACTGTCGAGCTTGAGTACAGAAGAGGAGAGTGGAACTCCATGTGTAGCGGTGAAATGCGTAGATATATGGAAGAACACCGGTGGCGAAGGCGGCTCTCTGGTTTGTTACTGACGCTGAGGCTCGAAAGCATGGGT |
| P-asv3681 | CCAGGCGTTATCCGGATTTATTGGGTTTAAAGGGAGCGTAGGCGGATTATTAAGTCAGTGGTGAAAGACGGTGGCTCAACCATCGTTAGCCATTGAAACTGGTAGTCTTGAGTGCAGACAGGGATGCTGGAACTCGTGGTGTAGCGGTGAAATGCTTAGATATCATGATGAACTCCGATCGCGAAGGCAGGTGTCCGGGCTGCAACTGACGCTGAGGCTCGAAAGTGTGGGT |
| P-asv3682 | CCGGGCGTTATCCGGATTTATTGGGTTTAAAGGGAGCGTAGGCTGTCTATTAAGCGTGTTGTGAAATTTACCGGCTCAACTGGTGGCTTGCAGCGCGAACTGGTCGACTTGAGTATGCAGGAAGTAGGCGGAATTCATGGTGTAGCGGTGAAATGCTTAGATATCATGACGAACTCCGATTGCGCAGGCAGCTTACTGTAGCATAACTGACGCTGATGCTCGAAAGTGCGGGT |
| P-asv3683 | GCAAGCGTTAATCGGAATTACTGGGCGTAAAGCGCGCGTAGGTGGTTCAGCAAGTTGGATGTGAAAGCCCCGGGCTCAACCTGGGAACTGCATCCAAAACTACTGAGCTAGAGTACGGTAGAGGGTGGTGGAATTTCCTGTGTAGCGGTGAAATGCGTAGATATAGTAAGGAACACCAGTGGCGAAGGCGACCACCTGGACTGATACTGACACTGAGGTGCGAAAGCGTGGGG |
| P-asv3684 | GCGAGCGTTAATCGGAATTACTGTGCGTAAAGCGTGCGCAGGCGGTTGTGCAAGACAGGTGTGAAATCCCCGGGCTTAACCTGGGAACTGCACTTGTGACTGCACGGCTAGAGTACGGTAGAGGGGGATGGAATTCCGCGTGTAGCAGTGAAATGCGTAGATATGCGGAGGAACACCGATCGCGAAGGCAATCCCCTGGACCTGTACTGACGCTCATGCATGAAAGCGTGGGG |
| P-asv3685 | CCGAGCGTTGTCCGGAATCATTGGGCGTAAAGGGTACGTAGGCGGGTATTTAAGTTAGAAGTCAAAGGCTATAGCTCAACTATAGTAAGCTTCTAAAACTGAATACCTTGAGAGATGGAAGGGAAAGTGGAATTCCTAGTGTAGCGGTGGAATGCGCAGATATTAGGAAGAATACCGGTGGCGAAGGCGACTTTCTGGCCATTTTCTGACGCTGAGGTACGAAAGCGTGGGT |
| P-asv3686 | GCGAGCGTTAATCGGAATTACTGGGCGTAAAGCGTGCGCAGGCGGTTATGTAAGACAGATGTGAAATCCCCGGGCTCAACCTGGGAACTGCATTAGTAACTGCATAGCTGGAGTACGGCAGAGGGGGATGGAATTCCGCGTGTAGCAGTGAAATGCGTAGATATGCGGAGGAACACCGATGGCGAAGGCAATCCCCTGGGCCTGTACTGACGCTCATGCACGAAAGCGTGGGG |
| P-asv3687 | CCAAGCGTTATCCGGATTTATTGGGTTTAAAGGGTGCGTAGGCGGCCTGTTAAGTCAGGGGTGAAAGACGGTGGCTCAACCATCGCAGTGCCCTTGATACTGATGGGCTTGAATGGACTAGAGGTAGGCGGAATGTGACAAGTAGCGGTGAAATGCATAGATATGTCACAGAACACCGATTGCGAAGGCAGCTTACTATGGTCTTATTGACGCTGAGGCACGAAAGCGTGGGG |
| P-asv3688 | GCAAGCGTTGTCCGGAATTATTGGGCGTAAAGCGCGCGCAGGCGGTCATTTAAGTCTGGTGTTTAAACCTTGGGCTCAACCTAAGGTCGCACTGGAAACTGGGTGACTTGAGTACAGAAGAGGAAAGTGGAATTCCACGTGTAGCGGTGAAATGCGTAGATATGTGGAGGAACACCAGTGGCGAAGGCGACTTTCTGGGCTGTAACTGACGCTGAGGCGCGAAAGCGTGGGG |
| P-asv3689 | GCAAGCGTTAATCGGAATTACTGGGCGTAAAGCGCGCGTAGGTGGTTTAGTCAGTCAGATGTGAAAGCCCTGGGCTCAACCTAGGAATTGCACCTGATACTGCTAGGCTAGAGTACAGTAGAGGGGAGTGGAATTTCCTGTGTAGCGGTGAAATGCGTAGATATAGGAAGGAACACCAGTGGCGAAGGCGACTCTCTGGACTGATACTGACACTGAGGTGCGAAAGCGTGGGG |
| P-asv3690 | GCGAGCGTTAATCGGAATTACTGGGCGTAAAGCGAGCGCAGACGGTTACTTAAGCAGGATGTGAAATCCCCGGGCTTAACCTGGGAACTGCGTTCTGAACTGGGTGACTAGAGTGTGTCAGAGGGAGGTAGAATTCCACGTGTAGCAGTGAAATGCGTAGAGATGTGGAGGAATACCGATGGCGAAGGCAGCCTCCTGGGATAACACTGACGTTCATGCCCGAAAGCGTGGGT |
| P-asv3691 | GCAAGCGTTAATCGGAATTACTGGGCGTAAAGCGCGCGTAAGTAGTTTGTTAAGTTGGATGTGAAAGCCCCGGGCTCAACCTGGGAACTGCATTCAAAACTGACAAGCTAGAGTATGGTAGAGGGTGGTGGAATTTCCTGTGTAGCGGTGAAATGCGTAGATATAGGAAGGAACACCAGTGGCGAAGGCGACCACCTGGACTGATACTGACACTGAGGTGCGAAAGCGTGGGG |
| P-asv3692 | GCAAGCGTTGTCCGGAATTATTGGGCGTAAAGAGCTCGTAGGCGGTTTGTCGCGTCTGCTGTGAAAACGCAAGGCTTAACCTTGCGCCTGCAGTGGGTACGGGCAGACTAGAGTGCGGTAGGGGAGACTGGAATTCCTGGTGTAGCGGTGGAATGCGCAGATATCAGGAGGAGCACCGATGGCGAAGGCAGGTCTCTGGGCCGTTACTGTCGCTGAGGAGCGAAAGCATGGGG |
| P-asv3693 | GCAAGCGTTAATCGGAATTACTGGGCGTAAAGCGTGTGTAGGCGGTTCGGAAAGAAAGATGTGAAATCCCAGGGCTCAACCTTGGAACTGCATTTTAACTGCCGAGCTGGAGTATGTCAGAGGGGGGTAGAATTCCACGTGTAGCAGTGAAATGCGTAGATATGTGGAGGAATACCGATGGCGAAGGCAGCCCCCTGGGATAATACTGACGCTCAGACACGAAAGCGTGGGG |
| P-asv3694 | GCAAGCATTAATCGGAATTATTGGGCGTAAAGGGCGCGTAGGCGGGATTAAAAGTCAGATGTGAAATTCCGAAGCTCAACTTCGGAGCCGCATTTGAAACTGCAATTCTTGAGGGATGGCGGAGGAAACGGAATTCCACGTGTAGCGGTGAAATGCGTAGATATGTGGAAGAACACCTGTGGCGAAAGCGGTTTTCTAGCTTTTACCTGACGCTGAGGCGCGAGAGCAAGGGG |
| P-asv3695 | GCGAGCGTTACCCGGATTTACTGGGCGTAAAGGGCGTGTAGGCGGTTTTCTAAGTCCGGGGCTAAAGACCAGGGCTCAACCCTGGGTTTGCCTTGGATACTGGAAAGCTCGACGGCTGGAGGGGGCAGCGGAATTTCCGGAGTAGCGGTGAAATGCGCAGATACCGGAAGGAACGCCAATAGCGAAGGCAGCTGCCTGGACAGTACGTGACGCTGAGGCGCGAAAGCGTGGGG |
| P-asv3696 | GCAAGCGTTGTCCGGAATGACTGGGCGTAAAGCGCGCGCAGGCGGTCTATCACGTCTGGGGTGAAAGTCCAGAGCTCAACTCTGGGATGGCCTTGGAAACGGGTGGACTTGAGGGTCGGAGAGGTCAGGGGAATTCCACGTGTAGCGGTGAAATGCGTAGAGATGTGGAGGAATACCAGTGGCGAAGGCGCCTGACTGGACGAAACCTGACGCTGAGGCGCGAAGGCGTGGGG |
| P-asv3697 | GCGAGCGTTGTCCGGAATTATTGGGCGTAAAGAGCTTGTAGGCGGCTTGTCGCGTCTGCTGTGAAAACCCGGGGCTTAACCCCGGGCTTGCAGTGGGTACGGGCAGGCTAGAGTGTGGTAGGGGAGACTGGAATTCCTGGTGTAGCGGTGGAATGCGCAGATATCAGGAGGAACACCGATGGCGAAGGCAGGTCTCTGGGCCATTACTGACGCTGAGAAGCGAAAGCGTGGGG |
| P-asv3698 | GCAAGCGTTATCCGGATTTATTGGGTTTAAAGGGTCCGTAGGTGGACCATTAAGTCAGAAGTGAAATCCCGCGGCTCAACCGCGGAACTGCTTTTGATACTGGTGGTCTTGAGTGTGCTCGAAGTTGGCGGAATATGACATGTAGCGGTGAAATGCTTAGATATGTCATAGAACACCGATTGCGAAGGCAGCCAACTAGGTCACAACTGACACTGAGGGACGAAAGCGTGGGG |
| P-asv3699 | TCGAGTGATGGCCGATATTATTGGGCCTAAAGCGTCCGTAGCCGGTCATGGAAGTCCGTTGGGAAATCTGCTCGCTCAACGAGCAGGCGTCCGGCGGAAACTCCGTGACTTGGGACCGAGAGAACCGAAGGGTACGTCTGGGGTAGGAGTGAAATCCCGTAATCCTGGACGGACCACCGATGGCGAAAGCACTTCGGTAGATCGGATCCGACGGTGAGGGACGAAAGCTAGGGT |
| P-asv3700 | GCAAGCGTTAACCGGATTTACTGGGCGTAAAGGGTGCGTAGGCGGATTGGTAAGTCAGATGTGAAATTTCGGTACTCAATACCGAAGCTGCATTTGAAACTGCCTATCTAGAGGGTAGATGGAGAAAGCGGAATTCCATGTGTAGCGGTGAAATGCGTAGATATATGGAAGAACACCTGTGGCGTAAGCGGCTTTCTAATTTATTCCTGACGCTGAGGCACGAGAGTGTGGGG |
| P-asv3701 | TCGAGTGATGACCGATGTTATTGGGCCTAAAGCGTCCGTAGCTGGCCGCGCAAGTCCGTCGGGAAATCCACCCGCTTAACGGGTGGGCGTCCGGCGGAAACTGCACGGCTTGGGACAGGAAGGCTCGAGGGGTACGTTCGGGGTAGGAGTGAAATCCCGTAATCCTGAACGGACCGCCGATGGCGAAAGCACCTCGAGAGGACTGATCCGACGGTGAGGGACGAAAGCTGGGGT |
| P-asv3702 | TCAAGCGTTATCCGGATTCACTGGGCGTAAAGGGTGCGTAGGCGGTGTTATTAGTCGGGTGTTAAATCCTGGGGCTCAACCTCAGAATCGCATCCGAAACGGTAATACTAGAAGAAGTAAGAGGTACACAGAACTCTCGGTGTAGGGGTGAAATCCGTTGATATCGAGGGGAATACCAAATGCGAAGGCAGTGTACTGGGACTTTCTTGACGCTGAGGCACGAAAGCGTGGGT |
| P-asv3703 | GCAAGCGTTGTCCGGAATCATTGGGCGTAAAGAGTTCGTAGGTGGTATGTTAAGTCTGGTGTTAAAGCCCGAAGCTCAACTTCGGTTCGGCACTGGATACTGGCAAACTTGAATGTGGTAGAGGTAAAGGGAATTCCTGGTGTAGCGGTGAAATGCGTAGATATCAGGAGGAACATCGGTGGCGTAAGCGCTTTACTGGGCCATAATTGACACTGAGGAACGAAAGCCGGGGT |
| P-asv3704 | CCAAGCGTTATCCGGAATTACTGGGTGTAAAGCGTCTGTAGGTGGTTTGGAAAGTTTCGTGTGAAACTTCAGGGCTCAACCCTGAATTGTGCGAAAAACTATCAAACTTGAGTGTGGGAGAGGCAAGCAGAATAGTAAGAGTAGGGGTGCAATCCGTTGATACTTACTAGAATACCAAAAGCGAAGGCAGCTTGCTGGAACATTACTGACACTGAGAGACGAAAGCGTGGGG |
| P-asv3705 | CCGAGCGTTATCCGGAATCATTGGGTTTAAAGGGTCCGTAGGCGGCTATATAAGTCAGTGGTGAAATCTGTTAGCTCAACTAACAAATTGCCATTGATACTGTATAGCTTGAATAATTGTGAAGTAGCTAGAATATGTAGTGTAGCGGTGAAATGCTTAGATATTACATGGAATACCAATTGCGAAGGCAGGTTACTAACAATTTATTGACGCTGATGGACGAAAGCGTGGGG |
| P-asv3706 | GCGAGCGTTGTTCGGAATCACTGGGCGTAAAGCGTGCGTAGGCGGACGCGCAAGTCAGGAGTGAAATCCAGGGGCTCAACCTCTGAACTGCTCTTGAAACTGCACGTCTAGAGTCCCGGAGGGGCCAGCGGAATTCCGAGTGTAGAGGTGAAATTCGCAGATATTCGGAGGAACACCAGTGGCGTAGGCGGCTGGCTGGACGGGAACTGACGCTCAGGCACGAAAGCGTGGGG |
| P-asv3707 | GCAAGCGTTATCCGGATTTATTGGGTTTAAAGGGTGCGCAGGCGGCTTTATAAGTCAGTGGTGAAATCTTCCCGCTCAACGGGAAAACTGCCATTGATACTGTAGAGCTAGAGTACAGATGGGGTAGGCGGAATGTAGCATGTAGCGGTGAAATGCTTAGATATGCTACAGAACACCAATCGCGAAGGCAGCTTACCAAGCTGGTACTGACGCTCAGGCACGAAAGCGTGGGT |
| P-asv3708 | GCTAGCGTTGTTCGGAATTACTGGGCGTAAAGCGCACGTAGGCGGCTTTGCAAGTTAGAGGTGAAAGCCCGGGGCTCAACCCCGGAATTGCCTTTAAGACTGCATCGCTTGAATCCGGGAGAGGTGAGTGGAATTCCGAGTGTAGAGGTGAAATTCGTAGATATTCGGAAGAACACCAGTGGCGAAGGCGGCTCACTGGACCGGTATTGACGCTGAGGTGCGAAAGCGTGGGG |
| P-asv3709 | GCGAGCGTTATCCGGAATTATTGGGCGTAAAGAGCTCGTAGGCGGTTTGTCACGTCTGCTGTGAAAGCCCAAGGCTTAACCTTGGGTTTGCAGTGGGTACGGGCAGGCTGGAGTGCAGTAGGGGAGACTGGAATTCCTGGTGTAGCGGTGAAATGCGCAGATATCAGGAGGAACACCGATGGCGAAGGCAGGTCTCTGGGCTGTTACTGACGCTGAGGAGCGAAAGCATGGGG |

| | |
|---|---|
| P-asv3710 | GCAAGCGTTATCCGGATTTATTGGGTTTAAAGGGTCCGTAGGCGGATCTGTAAGTCAGTGGTGAAATCTCATAGCTTAACTATGAAACTGCCATTGATACTGCAGGTCTTGAGTAAGGTAGAAGTGGCTGGAATAAGTAGTGTAGCGGTGAAATGCATAGATATTACTTAGAACACCAATTGCGAAGGCAGGTCACTATGTCTTAACTGACGCTGATGGACGAAAGCGTGGGG |
| P-asv3711 | GCAAGCGTTGTCCGGATTTACTGGGCGTAAAGGGTGCGTAGGCGGCCTGACAAGTCAGAAGTTAAAGCCCTCGGCTCAACCGAGGTAAGCTTTTGAAACTGTTAGGCTTGAGTACAGGAGAGGAGAGTGGAATTCCTAGTGTAGCGGTGAAATGCGTAGATATTAGGGGGAACACCAGTGGCGAAGGCGACTCTCTGGACTGTAACTGACGCTGAGGCACGAAAGCGTGGGG |
| P-asv3712 | GCAAGCGTTGTCCGGAATCATTGGGTGTAAAGGGTGCGTAGGCGGGCTAGTAAGTCTGGGGTGAAATCCCACCGCTCAACGGTGGAACTGCCTTGGATACTGCTAGTCTTGAGTATGTCAGAGGTCGGTGGAATTCGTGGTGTAGCGGTGAAATGCTTAGATATCACGAAGAACACCAGTGGTGAAGACGGCCGGCTGGGACATAACTGACGCTGAGGCACGAAAGCGTGGGG |
| P-asv3713 | ACGAGTGATGGCCGATATTATTGGGCCTAAAGCGTTCGTAGCTGGCCGAACAAGTCCGTTGGGAAATCGACGCGCCCAACGCGTCGGCGTCCAGCGGAAACTGTTCGGCTTGGGGCCGGAGGACTCGAGGGGTACGTCCGGGGTAGGAGTGAAATCCCGTAATCCTGGACGGACCACCGGTGGCGAAAGCGCCTCGAGAGGACGGACCCGACAGTGAGGGACGAAAGCCAGGGT |
| P-asv3714 | GCAAGCGTTGTTCGGAATTATTGGGCGTAAAGGGCGTGTAGGTGGTCTGCCAAGTCAGGTGTGAAATCCCACGGCTCAACCGTGGAACTGCACTTGAAACTGGTGGACTCGAGTGTCGGAGGGGGCAGCGGAATTCCCGGTGTAGAGGTGAAATTCGTAGATATCGGGAGGAACACCAGTGGCGAAGGCGGCTGCCTGGCCGATGACTGACACTGAGACGCGAAAGCGTGGGG |
| P-asv3715 | GCAAGCGTTGTTCGGAATTATTGGGCGTAAAGCGCGTGTAGGCGGCTTTGCAAGTCGGGTGTGAAAGCCCTCAGCTCAACTGAGGAAGTGCGCCCGAAACTGCAGAGCTTGAGTGCCGGAGAGGGTGGCGGAATTCCCCAAGTAGAGGTGAAATTCGTAGATATGGGGAGGAACACCGGTGGCGAAGGCGGCCACCTGGACGGTAACTGACGCTGAGACGCGAAAGCGTGGGT |
| P-asv3716 | GCAAGCGTTGTCCGGAATTATTGGGCGTAAAGCGCGCGCAGGTGGTTTCTTAAGTCTGATGTGAAAGCCCCCGGCTCAACCGGGGAGGGTCATTGGAAACTGGGAGACTTGAGTGCAGAAGAGGATAGTGGAATTCCAAGTGTAGCGGTGAAATGCGTAGAGATTTGGAGGAACACCAGTGGCGAAGGCGACTGTCTGGTCTGTAACTGACACTGAGGCGCGAAAGCGTGGGG |
| P-asv3717 | GCGAGCGTTAATCGGAATTACTGGGCGTAAAGGGCACGCAGGTGGTGTATTAAGTCGGGTGTGAAATCCCCGGGCTCAACCCGGGAACGGCATCTGATACTGGTACGCTTGAGTTTTGTAGAGGGGGGTGGAATTCCAGGTGTAGCGGTGAAATGCGTAGAGATCTGGAGGAATACCGGTGGCGAAGGCGGCCCCCTGGACAGAGACTGACACTGAGGTGCGAAAGCGTGGGG |
| P-asv3718 | GCGAGCGTTAATCGGAATTACTGGGCGTAAAGCGCGCGTAGGCGGCGCGGTCAGTCAGGTGTGAAAGCCCCGGGCTTAACCTGGGAATGGCACTTGATACTGCCGCGCTGGAGTCTGGGAGAGGGAAGGGGAATTCCGGGTGTAGCGGTGAAATGCGTAGATATCCGGAGGAACACCGGTGGCGAAGGCGCCTTCCTGGCCCAAGACTGACGCTGAGGTGCGAAAGCGTGGGG |
| P-asv3719 | GCGAGCGTTAATCGGAATTACTGGGCGTAAAGCGTGCGCAGGCGGCTCTTCAAGTCAGGTGTGAAAGCCCCGGGCTTAACCTGGGAATTGCGCTTGAAACTGGGGGGCTAGAGTGTGGCAGAGGGAGGTGGAATTCCTGGTGTAGCGGTGAAATGCGTAGATATCAGGAGGAACACCGATGGCGAAGGCAGCCTCCTGGGCCAACACTGACGCTCAGGCACGAAAGCGTGGGG |
| P-asv3720 | GCTAGCGTTGTTCGGAATTACTGGGCGTAAAGCGCACGTAGGCGGCCTTGTAAGTTGGGGGTGAAATCCCGGAGCTCAACTCCGGAACTGCCTTCAAAACTACAAAGCTAGAGGATGTGAGAGGAGAGTGGAATTCCGAGTGTAGAGGTGAAATTCGTAGATATTCGGAAGAACACCAGTGGCGAAGGCGACTCTCTGGCACATTTCTGACGCTGAGGTGCGAAAGCGTGGGG |
| P-asv3721 | GCAAGCGTTGTCCGGATTTATTGGGTTTAAAGGGTGCGTAGGTGGCTTGTTAAGTCTGGGGTGAAAGGCAGTGGCTCAACCACTAGCAGGGCCCTGGATACTGGCAGGCTCGAGTGCGCGTGAGGCCGGTGGAACGGGTGGTGTAGCGGTGAAATGCATAGATACCATCCAGAACACCGATTGCGGAGGCAGCCGGCTAACGCGCAACTGACACTGAGGCACGAAAGCGTGGGG |
| P-asv3722 | CCGAGTTGTCGGGACGATTATTGGGCCTAAAGCATCCGTAGCCTGTTCTGCAAGTCCTCCGTTAAATCCACCTGCTTAACAGTTGGGCTGCGGGGGATACTACAGCGCTAGGAGGCGGGAGAGGCAAGCGGTACTCAATGGGTAGGGGTAAAATCCGCTGATCCATTGAAGACCACCAGTGGCGAAGGCGGCTTGCCAGAACGCGCTCGACGGTGAGGGATGAAAGCTGGGGG |
| P-asv3723 | GCGAGCGTTAATCGGAATTACTGGGCGTAAAGGGCGCGTAGGCGGTTCGATAAGTCGGGTGTGAAAGCCCCGGGCTCAACCTGGGAATTGCACTCGATACTGTCGGGCTGGAGTGTGGTAGAGGGAAGCGGAATTCCGGGTGTAGCGGTGAAATGCGTAGATATCCGGAGGAACACCAGTGGCGAAGGCGGCTTCCTGGACCAACACTGACGCTGAGGCGCGAAAGCGTGGGG |
| P-asv3724 | GCTAGCGTTGCTCGGAATCACTGGGCGTAAAGGGTGCGTAGGCGGGTCTTTAAGTCAGAGGTGAAAGCCTGGAGCTCAACTCCAGAACTGCCTTTGATACTGAGGATCTTGAGTATGGGAGAGGTGAGTGGAACTGCGAGTGTAGAGGTGAAATTCGTTGATATTCGCAAGAACACCAGTGGCGAAGGCGGCTCACTGGCCCATAACTGACGCTGAGGCACGAAAGCGTGGGG |
| P-asv3725 | GCAAGCGTTAAGCGGAATTATTGGGCGTAAAGCGAGTGCAGACGGTTACTTAAGCCAGATGTGAAATCCCCAAGCTTAACTTGGGGCGTGCATTTGGAACTGGGTGACTAGAGTGTGTCAGAGGGAGGTAGAATTCCACATGTAGCGGTGGAATGCGTAGAGATGTGGAGGAATACCGATGGCGAAGGCAGCCTCCTGGGATAACACTGACGTTGAGGCTCGAAAGCGTGGGG |
| P-asv3726 | GCAAGCGTTATCCGGAATCACTGGTCGTAAAGGGTGCGTAGGCGGCCTAACAAGTCAGAGGTGAAAGGCATTGGCTCAACCAATGTAAGCCTTTGAAACTGTTGGGCTTGAGTGCAGGAGAGGTAAGTGGAATTCCTAGTGTAGAGGTGAAATTCGTAGATATTAGGAGGAACACCAGTGGCGAAGGCGACTTACTGGACTGTAACTGACGCTGAGGCACGAAAGCGTGGGG |
| P-asv3727 | GCAAGCTTTATCCGGAATCATTGGGTTTAAAGGGTCCGTAGGCGGGTGAGTAAGTCAGTGGTGAAAGTCTGTTGCTTAACAATAGAATTGCCATTGATACTGCTTGTCTTGAATAGCTATGAAGTAGTTAGAATGTGCAGTGTAGCGGTGAAATGCATAGATATTGCACAGAATACCAATTGCGAAGGCAGATTACTAATAGTTTATTGACGCTGATGGACGAAAGCGTGGGG |
| P-asv3728 | GCAAGCGTTAATCGGAATTACTGGGCGTAAAGCGCACGCAGGCGGTTGGATAAGTTAGACGTGAAAGCCCCGGGCTCAACCTGGGAATTGCATTTAAAACTGTCCAGCTAGAGTCTTGTAGAGGGGGGTAGAATTCCAGGTGTAGCGGTGAAATGCGTAGAGATCTGGAGGAATACCGGTGGCGAAGGCGGCCCCCTGGACAAAGACTGACGCTCAGGTGCGAAAGCGTGGGG |
| P-asv3729 | GCAAGCGTTGCTCGGTATCACTGGGCTTAAAGGGCGCGTAGGCGGTGCGGTGCGGTTTCGGTGAAATCCCTCGGCCCAACCGAGGAACAGCCGGAGCAACGGCCGCGCTAGACGGCATCAGGGGACAGTGGAACTCCCGGTGGAGCGGTGAAATGCGTAGATATCGGGAGGAACGCCGGTGGCGAAAGCGACTGTCTGGGGTGCATGTGACGCTGAGGCGCGAAAGCCAGGGG |
| P-asv3730 | GCAAGCGTTGTCCGGATTTACTGGGCGTAAAGCGCGCGCAGGTGGATGGATCAGTCCGGGGTGAAACCTCACGGCTCAACTGTGAGTGGTCCCCGGATACTATCCGTCTTGAGGCGCCTAGAGGAGAGCGGAATTCCCGGTGTAGTGGTGGAATGCGTAGATATCGGGAAGAACACCAGTGGCGAAAGCGGCTCTCTGGGGGCGATCTGACACTGAGGCGCGAAAGCGTGGGG |
| P-asv3731 | GCTAGCGTTGTTCGGAATTACTGGGCGTAAAGCGCGCGCAGGCGGTTTTTCAAGTCAGTGGTGAAAGCCCGGAGCTCAACTCCGGAACTGCCATTGAAACTGTTAAACTTGAGGATGAGAGAGGTGAGTGGAATTCCCAGTGTAGAGGTGAAATTCGTAGATATTGGGAAGAACACCGGTGGCGAAGGCGGCTCACTGGCTCATTACTGACGCTCAGGCGCGACAGCGTGGGG |
| P-asv3732 | GCAAGCGTTGTTCGGAATCACCGGGCTTAAAGCGCGTGTAGGCGGCCCGTCAAGTACCTTGTGAAATCCCTCGGCTCAACCGGGGAATGGCTGGGTATACTGACGGGCTTGAGGTGGGTAGGGGTGCCTGGAACTCTAGGTGGAGCGGTGAAATGCGTAGATATCTAGAGGAACGCCCGTGGCGAAAGCGGGGCACTGGGCCTTACCTGACGCTAAGACGCGAAAGCGTGGGG |
| P-asv3733 | GCAAGCGTTGTCCGGAATCACTGGGCGTAAAGGGCGCGTAGGCGGCTCATAAAGGCTGTGGTGAAAGTTCGGGGCTCAACCTCGAATCGGCCGTGGCGACTTGTGGGCTGGAGCACTGTAGAGGCAGGTGGAATTCCGGGTGTAGCGGTGGAATGCGTAGAGATCCGGAAGAACACCAGTGGCGAAGGCGGCCTGCTGGGCAGTGGCTGACGCTGAGGCGCGACAGCGTGGGG |
| P-asv3734 | GCGAGCGTTGTTCGGATTTACTGGGCGTAAAGGGCGCGTAGGCGGCGGGTCAAGTCGAGTGTGAAAGCCCCGGGCTCAACCTGGGAGGTGCGCTCGGGACTGGTCTGCTAGAGTTCGGGAGAGGAGCGTGGAATTCCCAGTGTAGAGGTGAAATTCGTAGATATTGGGAAGAACACCGGTGGCGAAGGCGGCGCTCTGGACCGAGACTGACGCTGAGGCGCGAAAGCGTGGGG |
| P-asv3735 | GCAAGCGTTGTCCGGAATCATTGGGCGTAAAGCGCGTGTAGGCGGTCCGGTCAGTCGGCTGTGAAAGTCCAGGGCTCAACCCTGGGATGCCGGTCGATACTGCCGGACTAGAGTCCGGAAGAGGCGAGTGGAATTCCCGGTGTAGCGGTGAAATGCGCAGATATCGGGAGGAACACCAATGGCGAAGGCAGCTCGCTGGGACGGTACTGACACTGAGACGCGAAAGCGTGGGG |
| P-asv3736 | GCGAGCGTTGTTCGGAATTATTGGGCGTAAAGCGCGTGTAAGCGGCTCCTTAAGTCTGATGTGAAAGCCCCGGGCTCAACCTGGGAAGTGCATTGGAAACTGGGGAACTTGAATACGGGAGGGGGTAGTGGAATTCCTGGTGTAGGAGTGAAATCCGTAGATATCAGGAGGAACACCGGTGGCGAAGGCGGCTACCTGGACCGATATTGACGCTGAGACGCGAAAGCGTGGGG |
| P-asv3737 | CCGAGCGTTGTCCGGAATTACTGGGCGTAAAGAGCTCGTAGGTGGTTTGTCGCGTTGTCCGTGAAAACTCACAGCTCAACTGTGGGCGTGCGGGCGATACGGGCAGACTGGAGTACTGCAGGGGAGACTGGAATTCCTGGTGTAGCGGTGGAATGCGCAGATATCAGGAGGAACACCGGTGGCGAAGGCGGGTCTCTGGGCAGTAACTGACGCTGAGGAGCGAAAGCGTGGGG |
| P-asv3738 | GCAAACGTTGCTCGGAATCATTGGGCGTAAAGCGCGCGTAGGCGGCACAGCAAGTTGGATGTGAAAGCCCTCGGCTTAACCGAGGAAGTGCATTCAAAACTACTGAGCTTGAGTACAAAAGAGGATCGCGGAATTCCCGGTGTAGAGGTGAAATTCGTAGATATCGGGAGGAACACCAGCGGCGAAGGCGGCGATCTGGGTTGATACTGACGCTGAGGTGCGAAAGCGTGGGG |
| P-asv3739 | GCAAGCGTTGTTCGGAATTACTGGGCGTAAAGGGCTCGTAGGCGGCCAACTAAGTCGGACGTGAAATCCCTCAGCTTAACTGGGGAACTGCGTCCGATACTGGATGGCTTGGATTTGGGAGAGGGATGCGGAATTCCAGGTGTAGCGGTGAAATGCGTAGATATCGGGAGGAACACCTGTGGCGAAAGCGGCGCACTGGACCATAACTGACGCTGAGGCGCGAAAGCTAGGGG |
| P-asv3740 | GCGAGCGTTGTCCGGATTCATTGGGCGTAAAGCGTGCGTAGGCGGCTGCGCAAGTCGGTTGTGAAACCCCGGGGCTCAACCCCGGAACTGCATCCGATACTGCGTGGCTAGAGGTAGGTAGGGGAGATCGGAATTCCTGGTGTAGCGGTGAAATGCGCAGATATCAGGAGGAACACCGGTGGCGAAGGCGGATCTCTGGGCCTTACCTGACGCTCAGGCACGAAAGCTAGGGG |
| P-asv3741 | GCAGGCGTTAATCGGAATTACTGGGCGTAAAGCGTGCGCAGGCGGCTCCTTAAGTCAGATGTGAAAGCCCCGGGCTTAACCTGGGAACTGCGTTTGAAACTGGGGAGCTAGAGTGTGGGAGAGGGGGGTGGAATTCCAGGTGTAGCAGTGAAATGCGTAGAGATCTGGAGGAACACCGATGGCGAAGGCAGCCCCCTGGGCTAACACTGACGCTCATGCACGAAAGCGTGGGG |
| P-asv3742 | GCAAGCGTTGTCCGGATTTATTGGGCGTAAAGAGCGTGTAGGCGGCCAGGCAGGTCCGCTGTGAAAACTCGAGGCTCAACCTCGAGAGGTCGGTGGAAACCGTTTGGCTAGAGTCCGGAAGAGGAGAGTGGAATTCCCGGTGTAGCGGTGAAATGCGCAGATATCGGGAAGAACACCCGTGGCGAAGGCGGCTCTCTGGGACGGTACTGACGCTGAGACGCGAAAGCGTGGGG |
| P-asv3743 | GCGAGCGTTGTCCGGAGTCACTGGGCGTAAAGGGTGCGTAGGCGGTCGTGCGCGTGGATGGTGAAATCTTCCGGCTCACCCGGGAGGGGTCCGTCCAGACGGCAGGACTTGAGGGTCGGAGAGGGGCGTGGAATTCCGGGTGGAGCGGTGAAATGCGTAGAGATCCGGAGGAACACCGACGGCGTAGGCAGCGCCCTGGACGACACCTGACGCTGAGGCACGAAAGCTGGGGG |
| P-asv3744 | GCAAGCGTTGTCCGGAATTATTGGGCGTAAAGAGCGTGTAGGCGGTCCGGTAAGTCGGTTGTGAAAGTCCAGGGCTCAACCCTGGGATGCCGATCGATACTGCCGGACTAGAGTCCGGAAGAGGCGAGTGGAATTCCCGGTGTAGCGGTGAAATGCGCAGATATCGGGAGGAACACCAATGGCGAAGGCAGATCGCTGGGACGGTACTGACGCTGAGACGCGAAAGCGTGGGG |
| P-asv3745 | GCAAGCGTTGTCCGGATTTATTGGGTTTAAAGGGTGCGTAGGTGGCCCTGCAAGTCAGTGGTGAAATATCTCAGCTTAACTGAGAGGGTGCCATTGATACTGCAGGGCTTGAGTACAGATGAGGTAGGCGGAATTGACGGTGTAGCGGTGAAATGCATAGATATCGTCAAGAACACCGATAGCGAAGGCAGCTTACCAAGCTGTAACTGACACTGAGGCACGAAAGTGTGGGG |
| P-asv3746 | GCTAGCGTTGTTCGGATTTACTGGGCGTAAAGCGCGCGTAGGCGGCTTTTCAAGTCAGGGGTGAAATCCCACGGCTCAACCGTGGAATTGCCTTTGAAACTGTAAAGCTAGAGTGTCGGAGGGGATAGCGGAATTGCTAATGTAGAGGTGAAATGCGCAGATATCAGGAGGAACACCGGTGGCGAAGGCGGTTCTCTGGGCCTTACCTGACGCTGAGGAGCGAAAGCGTGGGG |
| P-asv3747 | ACCAGCGTTGTCCGGATTTACTGGGCGTAAAGAGCGCGCAGGCGGTCGTTCAAGTCGAGTGTGAAAGCCCCCGGCTCAACTGGGGAGGGTCACTCGATACTGATCGACTTGAAGGCAGGAGAGGGAAGCGGAATTCCCGGTGTAGTGGTGAAATGCGTAGATATCGGGAGGAACACCAGTGGCGAAGGCGGCTTCCTGGCCTGTTCTTGACGCTGAGGCGCGAAAGCTAGGGG |
| P-asv3748 | GCCAGCGTTGTCCGGATTCATTGGGCGTAAAGCGCGCGTAGGCGGCCGAGCAGGTCGCCCGTGACAGCCCCCGGCTCAACCGGGGACCGTCGGGCGAAACCGCCCGGCTCGAGGCCGGTAGGGGGAAGCGGAATTCCCGGTGTAGTGGTGGAATGCGTAGAGATCGGGAGGAACACCCGTGGCGAAGGCGGCTTCCTGGCCCGAGCCTGACGCTGAGGCGCGAAAGCGTGGGG |

| | |
|---|---|
| P-asv3749 | GCGAGCGTTGTTCGGATTTACTGGGCGTAAAGGGCGCGCAGGCGGCGATTTTTGTCAGGCGTGAAAGCCCCGGGCTCAACCTGGGAACTGCGCTTGGGACGGGATCGCTAGAGATCGGGAGAGGAGCGTGGAATTCCCAGTGTAGAGGTGAAATTCGTAGATATTGGGAAGAACACCGGTGGCGAAGGCGGCGCTCTGGACCGATACTGACGCTGAGGCGCGAAAGCGTGGGG |
| P-asv3750 | GCGAGCGTTAATCGGAATTACTGGGCGTAAAGCGTGCGCAGGCGGTTTTGTAAGCCGGATGTGAAATCCCCGGGCTTAACCTGGGAATGGCATTCGGGACTGCAAGGCTGGAGTGCGGCAGAGGAGACTGGAATTCCTGGTGTAGCAGTGAAATGCGTAGATATCAGGAGGAACACCGATGGCGAAGGCAGGTCTCTGGGCTGACACTGACGCTCATGCACGAAAGCGTGGGG |
| P-asv3751 | GCCAGCGTTGTTCGGTATTACTGGGTGTAAAGGGTTCGTAGGCGGTGTAGCAAGTCGGGAGTGAAATCTCTGGGCTTAACCCAGAGGCTGCTTCCGAAACTGCTGTGCTGGAGTGTGGGAGAGGCGCGTGGAATTGCAGGTGTAGCGGTGAAATGCGTAGATATCTGCAGGAACACCCGTGGCGAAGGCGGCGCGCTGGACCACTACTGACGCTGAGGAACGAAAGCTAGGGG |
| P-asv3752 | ACGAGCGTTGTCCGGATTTATTGGGCGTAAAGCGCGCGTAGGCGGCCCGGAAAGTCGGTTGTGAAAACCCGGGGCTCAACCCCGGGCCTGCATCCGAAACTTCCGGGCTGGAGGTAGGTAGGGGAGATCGGAATTCCTGGTGTAGCGGTGAAATGCGCAGATATCAGGAGGAACACCGGTGGCGAAGGCGGATCTCTGGGCCTTACCTGACGCTGAGGCGCGAAAGCTAGGGG |
| P-asv3753 | GCTAGCGTTGTCCGGAATCATTGGGCGTAAAGAGCGCGTAGGCGGTCCGGTAGGTCTGCCGTGAAAGCCGGGGGCTCAACCCTTGGATGCCGGTGGATACTGCCGGGCTAGAGTCCGGAAGAGGCGAGTGGAATTCCTGGTGTAGCGGTGAAATGCGCAGATATCAGGAAGAACACCGATGGCGAAGGCAGCTCGCTGGGACGGAACTGACGCTGAGGCGCGAAAGCGTGGGG |
| P-asv3754 | GCCAGCGTTGTTCGGAATTACTGGGTGTAAAGGGTTCGTAGGCGGTGTGGCAAGTCACGTGTGAAATCCCCAGGCTCAACCTGGTGATGCCACGTGATACTGCCATGGCTTGAGTCCGGTAGGGGACCATGGAATTCCTGGTGTAGCGGTGAAATGCGCAGATATCAGGAGGAACACCAGTGGCGAAGGCGGTGGTCTGGGCCGGTACTGACGCTGAGGAGCGAAAGCGTGGGG |
| P-asv3755 | GCAAGCGTTACCCAGATTTATTGGGCGTAAAGCGAGCGCAGGCGGTCTTTTAAGTCTAATGTGAAAGCCTTCGGCTCAACCGAAGAAGTGCATTGGAAACTGGGAGACTTGAGTGCAGAAGAGGACAGTGGAACTCCATGTGTAGCGGTGAAATGCGTAGATATATGGAAGAACACCAGTGGCGAAGGCGGCTGTCTGGTCAGTAACTGACGCTGAGGCTCGAAAGCATGGGT |
| P-asv3756 | GCGAGCGTTGTCCGGAGTTACTGGGCGTAAAGGGTGCGCAGGCGGCCTACCGCGTGGGGGGTGAAATCGCCCGGCTTAACCGGGCGGGTGCTTTCCAGACGGGTAGGCTTGAGGGGCGGAGAGGGGCGCGGAATTCCGGGTGGAGCGGTGAAATGCGTAGAGATCCGGAAGAACACCGACGGCGAAGGCAGCGCCCTGGACGCCATCTGACGCTCGGGCACGAAAGCGCGGGG |
| P-asv3757 | GCTAGCGTTGTCCGGATTTATTGGGCGTAAAGAGCGTGTAGGCGGCCAGGTAGGTCTGCTGTGAAAACTCGAGGCTCAACCTCGAGACGTCGGCGGAAACCATCTGGCTAGAGTCCGGAAGAGGAGAGTGGAATTCCTGGTGTAGCGGTGAAATGCGCAGATATCAGGAAGAACACCCGTGGCGAAGGCGGCTCTCTGGGACGGTACTGACGCTGAGACGCGAAAGCGTGGGG |
| P-asv3758 | GCGAGCGTTGTTCGGAATTACTGGGCGTAAAGGGCGTGCAGGCGGTGCGGTAAGTTGGGCGTGAAATCCCCGGGCTTAACCCGGGGGGCCGTTCAAGACTGCCGTGCTAGGGTGCGGGAGAGGGCAGTGGAATTCCCGGTGTAGCGGTGAAATGCGTAGATATCGGGAGGAACACCTGCGGCGAAGGCGGCTGCCTGGACCGTAACCGACGCTGAGGCGCGAAAGCTAGGGG |
| P-asv3759 | GCAAGCGTTGTCCGGAATCATTGGGCGTAAAGCGCGTGTAGGCGGCCAGATAAGTCCGTTCTGAAAGCCCGGGGCTCAACCCCGGGAGGCGGATGGATACTGTTTGGCTCGAGTCCGGAAGAGGCGAGTGGAATTCCCGGTGTAGCGGTGAAATGCGCAGATATCGGGAGGAACACCAATGGCGAAGGCAGCTCGCTGGGACGGTACTGACGCTGAGACGCGAAAGCGTGGGG |
| P-asv3760 | GCAAACGTTGTCCGGATTCATTGGGCGTAAAGAGCGTGTAAGCGGCCAGGTAGGTCGGTTGTGAAAACTGGAGGCTCAACCTTCAGACGTCGACCGAAACCATCTGGCTAGAGTCCGGGAGAGGAGAGTGGAATTCCTGGTGTAGCGGTGAAATGCGCAGATATCAGGAAGAACACCCGTGGCGAAAGCGGCTCTCTGGAACGGTACTGACGCTGAGACGCGAAAGCGTGGGG |
| P-asv3761 | GCGAGCGTTGTTCGGAATTATTGGGCGTAAAGGGTGCGTAGGCGGTGCCACAAGTCACTTGTGAAATCCCCAGGCTTAACCTGGGGCCTGCAGGCGAAACTGTGGTGCTGGAGTATGGGAGAGGTGCGTGGAATTCCCGGTGTAGCGGTGAAATGCGTAGATATCGGGAGGAACACCTGTGGCGAAAGCGGCGCACTGGACCATAACTGACGCTGAGGCGCGAAAGCTAGGGG |
| P-asv3762 | GCAAGCGTTAAGCGGAATCACTGGGCGTAAAGGGTGCGTAGGCGGTACGTTAGGTCTGTCGTGAAATCCCCAGGCTCAACCTGGGAATGGCGGTAGAAACCGGCGCACTAGAGTATGGGAGAGGGTGGTGGAATTCCAGGTGTAGCGGTGAAATGCGTAGAGATCTGGAGGAACATCAGTGGCGAAGGCGGCCACCTGGCCCAATACTGACGCTGAGGCACGAAAGCGTGGGG |
| P-asv3763 | GCGAGCGTTATCCGTATTTATTGGGTTTAAAGGGAGCGTAGGCGGTCTTATAAGTCAGTGGTGAAATCTCGTTGCTTAACAACGACCGTGCCATTGATACTGTAGGACTTGAGTACAGTTGCCGTTGGCGGAATGTGTCATGTAGCGGTGAAATGCATAGATATGACACAGAACACCGATTGCGAAGGCAGCTGATGAAACTGTAACTGACGCTGAGGCTCGAAAGCGTGGGG |
| P-asv3764 | CCTGGTGTTATCCGGATTTATTGGGTTTAAAGGGAGCGTAGGCCGTAGATTAAGTGTGTTGTGAAATGTAGGCGCCCAACGTCTGCCTTGCAGCGCAAACTGGTTTACTTGAGTACGCGCAACGCAGGCGGAATTCGTCGTGTAGCGGTGAAATGCTTAGATATGACGAAGAACTCCTATTGCGAAGGCAGCTTGCGGGAGCGTTACTGACGCTGAAGCTCGAAAGTGCGGGT |
| P-asv3765 | GCGAGCGTTAATCGGAATTACTGGGCGTAAAGCGGGCGCAGACGGTTACTTAAGCAGGATGTGAAATCCCCGGGCTCAACCTGGGAACTGCGTTCTGAACTGGGTGACTAGAGTGTGTCAGAGGGAGGTAGAATTCCACGTGTAGCAGTGAAATGCGTAGAGATGTGGAGGAATACCGATGGCGAAGGCAGCCTCCTGGGATAACACTGACGTTCATGCCCGAAAGCGTGGGT |
| P-asv3766 | CCAAACGTTGTTCGGAATCACTGGGCTTAAAGGGCGCGTAGGCGGCGAATCAAGTCAGGGGTGAAAGCCGGCAGCTTAACTGTCGAAGTGCCTTTGATACTGATTCGCTCGAGGGAGATAGGGGTCTGCGGAACTTCTGGTGGAGCGGTGAAATGCGTTGATATCAGAAGGAACGCCGGTGGCGAAAGCGGCGGACTGGATCTCTTCTGACGCTGAGGCGCGAAAGCTAGGGG |
| P-asv3767 | CCGAGCGTTGTCCGGAATCATTGGGCGTAAAGGGTACGTAGGCGGGTTTTTAAGTTAGAAGTCAAAGGCTATAGCTCAACTATAGTAAGCTTCTAAAACTGGAGACCTTGAGTAATGGAAGGGAAAGTGGAATTCCTAGTGTAGCGGTGGAATGCGCAGATATTAGGAGGAATACCGGTGGCGAAGGCGACTTTCTGGCCATTAACTGACGCTGAGGTACGAAAGCGTGGGT |
| P-asv3768 | TCAAGTGATGTCCCATATTATTGGGCCTAAAGCGTCCGTAGCTGGCCGGGCAAGTCCATCGGAAAATCCACCTGCCCAACAGGTGGGCGACCTGTGGAAACTGCCTGGCTTGGGATCGGAAGGCTCAGAGAGTACGTCCGGGGTAGGAGTGAAATCCCGTAATCCCGGACGGACTACCGATGGCGAAAGCACTCTGAGAAGACGACTCCGACAGTGAGGGACGAAAGCTCGGGT |
| P-asv3769 | GCTAGCGTTGTTCGGAATTACTGGGCGTAAAGCGCACGTAGGCGGATCGGAAAGTTGGGGGTGAAATCCCGGGGCTCAACCTCGGAACTGCCTCCAAAACTATCGGTCTTGAGTTCGAGAGAGGTGAGTGGAATTCCGAGTGTAGAGGTGAAATTCGTAGATATTCGGAGGAACACCAGTGGCGAAGGCGGCTCACTGGCTCGATACTGACGCTGAGGTGCGAAAGCGTGGGG |
| P-asv3770 | GCGAACGTTACCCGGAATCACTGGGCGTAAAGGGCGTGTAGGCGGCTCGGTATGTCGTTTGTGAAAGCCCCCTGCTAAACGGGGGGAGGTCAAGCGAAACTGCCGGGCTAGAGGGCCAGAGGGGGAAGTGGAACTACCAGAGTAGCGGTGGAATGCGTAGAGACTGGTAGGAACCTCGATAGCGAAGGCAACTTCCTGGCTGGTTCCTGACGCTGAGGCGCGAAAGCGTGGGG |
| P-asv3771 | GCAAGCGTTAATCGGAATTACTGGGCGTAAAGCGCGCGTAGGCGGTTTGATAAGCAGGATGTGAAAGCCCCGGGCTTAACCTGGGAATGGCATTCTGAACTGTCTGACTGGAGTATGGTAGAGGGGTGTGGAATTTCCGGTGTAGCGGTGAAATGCGTAGATATCGGAAGGAACACCAGTGGCGAAGGCGACACCCTGGACCAATACTGACGCTGAGGTGCGAAAGCGTGGGG |
| P-asv3772 | GCTAGCGTTGTTCGGAATCACTGGGCGTAAAGGGCGCGTAGGCGGCTTTATAAGTCGGGGGTGAAAGCCTGTGGCTCAACCACAGAATTGCCTTCGATACTGTAGAGCTTGAGACCGGAAGAGGTAAGTGGAACTGCGAGTGTAGAGGTGAAATTCGTAGATATTCGCAAGAACACCAGTGGCGAAGGCGGCTTACTGGTCCGGTTCTGACGCTGAGGCGCGAAAGCGTGGGG |
| P-asv3773 | GCAAGCGTTGTCCGGAATCACTGGGCGTACAGGGTGCGCAGGCGGACGTGCAAGTCTGACGTGAAAGCCCTGAGCTCAACTCAGGAAGTGCGTCGGAAACTGCATGTCTTGAGGGCCAGAGAGGCCGGTGGAATTCGCGGTGGAGCGGTGAAATGCGTAGATATCGCGAGGAACACCCGTGGCGAAAGCGACCGGCTGGATGGACCCTGACGCTGAGGCACGAAAGCGTGGGT |
| P-asv3774 | TCAAGTGGTAGTCATTATTATTGGGTTTAAAGTGTTCGTAGCCGGACCGGCAAGTCCGCTGTGAAATCCACATGCTTAACATGTGGGCGTGCAGCAGATACTGCTGGTCTAGGAACCGGACGGCGTCGAGGCTATTCCAAGGGTAGGGGTGAAATCCTAAAATCCTTGGAGGAGCACCTGTGGCGAAAGCGCTCGACGAGTACGGATTCGACGGTGAGGAACGAAAGCTAGAGG |
| P-asv3775 | GCTAGCGTTATTCGGAATTACTGGGCGTAAAGCGCACGTAGGCGGCTTTGTAAGTTAGAGGTGAAAGCCCGGGGCTCAACTCCGGAATTGCCTTTAAGACTGCATCGCTTGAATCCAGGAGAGGTGAGTGGAATTCCGAGTGTAGAGGTGAAATTCGTAGATATTCGGAAGAACACCAGTGGCGAAGGCGGCTCACTGGACTGGTATTGACGCTGAGGTGCGAAAGCGTGGGG |
| P-asv3776 | CCGAGTGATGACCGCTATTATTGGGCCTAAAGCGTCCGTAGCCGGCCGGACGGGTCCATCGGGAAATCCACGTGCCCAACGCGTGGGCGACCGGTGGAAACCATCCGGCTTGGGGCCGGAAGACCCAGGGGGTACGTCCGGGGTAGGAGTGAAATCCCGTAATCCTGGAGGGACCGCCGATGGCGAAAGCACCCTGGGAAGACGGACCCGACGGTGAGGGACGAAAGCTGGGGT |
| P-asv3777 | CCAAGTGATGGCCGATATTATTGGGCCTAAAGCGTCCGTAGCCGGCCGAACAAGTCCGTCGGGAAATCCACGCGCTCAACGCGTGGGCGTCCGGCGGAAACTGTTCGGCTTGGGACCGGGAGACCAGAGGGGTACGTCCGGGGTAGGAGTGAAATCCCGTAATCCTGGACGGACCACCGGTGGCGAAAGCGCCTCTGGAGAACGGATCCGACGGTGAGGGACGAAAGCCAGGGT |
| P-asv3778 | GCTAGCGTTGCTCGGAATTACTGGGCGTAAAGGGCGCGTAGGCGGACATTTAAGTCAGAGGTGAAAGCCCAGGGCTCAACCTTGGAATTGCCTTTGATACTGGATGTCTTGAGTTCGGGAGAGGTGAGTGGAATGCCGAGTGTAGAGGTGAAATTCGTAGATATTCGGCGGAACACCAGTGGCGAAGGCGACTCACTGGCCCGATACTGACGCTGAGGCGCGAAAGCGTGGGG |
| P-asv3779 | GCAAGCGTTGTTCGGAATTACTGGGCGTAAAGCGCGGGCAGGCGGTCTGTTAAGTCGTTTGTGAAAATCCGGGGCTCAACCCCGTGAAATGCATACGATACTGGCAGGCTGGAGACTGGTAGAGGCTAGCGGAATTCCTGGTGTAGCGGTGGAATGCGTAGATATCAGGAAGAACACCTGTGGCGAAGGCGGCTAGCTGGGCCAGATCTGACGCTCATCCGCGAAAGCGTGGGG |
| P-asv3780 | CCGAGCGTTATCCGGAATTATTGGGCGTAAAGGGTCCGCAGGCGGTCCTATTAGTCTTGTGTGAAATCTCCCGGCTTAACTGGGAAAACGCATAGGAAACGGTAGGACTAGAGGATGTGAGAGGTATGTGGAACTCATGGTGTAGGGGTGAAATCCGTTGATATCATGGGGAACACCAAAAGCGAAGGCAACATACTGGCACATTCCTGACGCTCAGGGACGAAAGCATGGGT |
| P-asv3781 | GCAAGCGTTGCTCGGAATCACTGGGCGTAAAGGGGAGGCAGGCGGTCCTGTGTGTCTGGGGTGAAAGCCCGGAGCTTAACTCCGGATGTGCCTCAGAAACTACAGGACTAGAGTTCTGGAGAGGTTAGCGGAATTCCCGGTGTAGCGGTGAAATGCGTAGAGATCGGGAGGAACACCAGCGGCGAAGGCGGCTAACTGGACAGATACTGACGCTCAACTCCGAAAGCATGGGG |
| P-asv3782 | CCAAGCGTTACCCGGAATCATTGGGTGTAAAGGATGCGTAGGTGGATTTTTAAGTCAGGTATGAAAGACCGGAGCTCAACTCCGTGTTTGTGCTTGAAACTGGAAATCTGGAATCAGGGAGAGGTTAGCGGAATTCTAAGTGTAGGGGTGCAATCCGTAGATACTTAGAAGAACACCAAAAGCGAAGGCAGCTAACTGGAACTGTATTGACACTGAGGCATGAAAGCGTGGGG |
| P-asv3783 | GCAAGCGTTGTCCGGAATCACTGGGTGTAAAGGGTGTGTAGGCGGACCTGTAAGTCGGAGGTGAAAGCCCTCAGCTCAACTGAGGAATTGCCCCCGATACTGCAGGTCTTGAGTCCCGGAGGGGCTACCGGAATTCGTGGTGTAGCGGTGAAATGCGTAGATATCACGAGGAACACCAGAGGTGAAGACGGGTAGCTGGACGGGTACTGACGCTGAGGCACGAAAGCGTGGGG |
| P-asv3784 | GCAAACGTTGTTCGGAATCATTGGGCGTAAAGCGCGCGTAGGCGGCCAGTTAGGTCGGGTGTGAAATCCCGGGGCTCAACCTCGGAACTGCGCTCGAAACCGGCTGGCTTGAGTCCCAGAGGGAGAAGTGGAATTCCTGGTGTAGGGGTGAAATCCGTAGATATCAGGAGGAACACCAGTGGCGAAGGCGACTTCTTGGCTGAGGACTGACGCTGAGGTGCGAAAGCGTGGGT |
| P-asv3785 | GCGAGCGTTAATCGGAATTACTGGGCGTAAAGCGTGCGTAGGCGGCTTGGTAAGCCGGTTGTGAAAGCCCCGGGCTCAACCTGGGAATGGCATCCGGAACTGTCAGGCTAGAGTGCAGGAGAGGAAGGTAGAATTCCCGGTGTAGCGGTGAAATGCGTAGAGATCGGGAGGAATACCAGTGGCGAAGGCGGCCTTCTGGACTGACACTGACGCTGAGGTACGAAAGCGTGGGT |
| P-asv3786 | CCAAGCGTTATCCGGATTCATTGGGTTTAAAGGGTCCGTAGGCGGATTTTTAAGTCAGTGGTGAAAGCCGACAGCTCAACTGTCGAACTGCCATTGATACTGGAAATCTTGAGTACAAATGAAGTAGGCGGAATGAGTCATGTAGCGGTGAAATGCATAGATATGACTCAGAACACCAATTGCGAAGGCAGCTTACTAACATGTAACTGACGCTGAGGGACGAAAGCGTGGGG |
| P-asv3787 | GCAAGCGTTGTTCGGAATCATTGGGCGTACAGGGTGCGCAGGCGGCCTGGCCAGTCCGATGTGAAAGCCCGGGGCTCAACTCCGGAATCGCATTGGAAACTGTCAGGCTTGAGTCCGAGAGGGGCCAGTGGAATTCCTGGTGTAGGGGTGAAATCCGTAGAGATCAGGAGGAACACCGGCGGCGAAAGCGACTGGCTGGATCGGGACTGACGCTGAGGCACGAAAGCGTGGGT |

| | |
|---|---|
| P-asv3788 | CCAAGTGATGGCCGATATTATTGGCCTAAAGCGTCCGTAGCCTGCTGTGTAAGTCCATTGGGAAATCGACCAGCTCAACTGGTCGGCGTCCGGTGGAAACTACACAGCTTGGGGCCGAGAGACTCAACGGGTA<br>CGTCTGGGGTAGGAGTGAAATCCTATAATCCTAGACGGACCACCAATGGGGAAACCACGTTGAGAGACCGGACCCGACGGTGAGGGACGAAAGCCAGGGT |
| P-asv3789 | GCAAGCGTTACTCGGAATCACTGGGCGTAAAGGGTGTGTAGGCGGAATATTAAGTCTCGAGTGAAATGCCATAGCTCAACTATGGAACTGCTTGGGAAACTGATAATCTAGAGTGTGGGAGAGGTAGATGGAATT<br>CCCGGTGTAGGGGTAAAATCCGTAGAGATCGGGAGGAATACCCATTGCGAAGGCGATTTACTGGAACACAACTGACGCTGAGGCACGAAAGCATGGGG |
| P-asv3790 | GCAAACGTTGCTCGGAATCATTGGGCGTAAAGCGCACGTAGGCGGCACAGCAAGTTGGATGTGAAAGCCCAGGGCTCAACCTTGGAGGTGCATTCAAAACTGCTGAGCTTGAGTTCCGGAGGGGCCCAGAGAAT<br>TCCCGGTGTAGAGGTGAAATTCGTAGATATCGGGAGGAATACCAGTGGCGAAGGCGCTGGGCTGGACGGATACTGACGCTGAGGTGCGAAAGCGTGGGG |
| P-asv3791 | GCTAGCGTTGTTCGGATTTACTGGGCGTAAAGCGTGCGTAGGCGGCTTATCAAGTCAGAAGTGAAAGCCCGGGGCTCAACTCCGGAACTGCTTTTGAGACTGGTAGGCTGGAGTTCGGGAGAGGGTAGTGGAAT<br>TCCCAGTGTAGAGGTGAAATTCGTAGATATTGGGAAGAACACCGGTGGCGAAGGCGGCTGCCTGGCCCGAAACTGACGCTGAGGCACGAAAGCGTGGGG |
| P-asv3792 | GCAAGCGTTATCCGGAATGATTGGGCGTAAAGCGTCTGTAGGTGGCTTGAAAAGTCTCCTGTCAAAGATCAGGGCTTAACCCTGGGCCGGCAGGAGAAACTCTCAGGCTAGAGTTTGGTAGGGGCAGAGGGAAT<br>TCCCGGTGGAGCGGTGAAATGCGTAGAGATCGGGAGGAACACCAAAGGCGAAAGCACTCTGCTGGGCCATTACTGACACTGAGAGACGAAAGCGAGGGG |
| P-asv3793 | GCTAGCGTTGTTCGGAATTACTGGGCGTAAAGCGCACGTAGGCGGATATTTAAGTCAGGGGTGAAATCCCGCAGCTCAACTGCGGAACTGCCTTTGATACTGGGTATCTTGAGTATGGAAGAGGTGAGTGGAATT<br>GCGAGTGTAGAGGTGAAATTCGTAGATATTCGCAGGAACACCAGTGGCGAAGGCGGCTCACTGGTCCATAACTGACGCTGAGGTGCGAAAGCGTGGGG |
| P-asv3794 | GCAAGCGTTAATCGGAATTACTGGGCGTAAAGCGCACGTAGGCGGTTCAGTCAGCCAGCCGTGAAATCCCCGGGCTCAACCTGGGAATTGCGGTTGGAACTGCTGAGCTAGAGTATGGTAGAGGGGTGTGGAA<br>TTCCAGGTGTAGCGGTGAAATGCGTAGATATCTGGAGGAACATCAGTGGCGAAGGCGACACCCTGGACCAATACTGACGCTGAGGTGCGAAAGCGTGGGG |
| P-asv3795 | GCAAGCGTTGTCCGGATTTATTGGGCGTAAAGCGCGCGCAGGCGGCTATGTAAGTCTGGTGTTAAAGCCCGGGGCTCAACCCCGGTTCGCATCGGAAACTGTGTAGCTTGAGTGCAGAAGAGGAAAGCGGTATT<br>CCACGTGTAGCGGTGAAATGCGTAGAGATGTGGAGGAACACCAGTGGCGAAGGCGGCTTTCTGGTCTGTAACTGACGCTGAGGCGCGAAAGCGTGGGG |
| P-asv3796 | GCGAGCGTTATCCGGATTTATTGGGTTTAAAGGGTGCGTAGGTTGTTCGGTAAGTCAGCGGTGAAACCTGAGCGCTCAACGTTCAGCCTGCCGTTGAAACTGCCGGGCTTGAGTTCAGTGGCGGCAGGCGGAAT<br>TCGTGGTGTAGCGGTGAAATGCATAGATATCACGAGGAACTCCGATTGCGAAGGCAGCTTGCCATACTGCGACTGACACTGAAGCACGAAGGCGTGGGT |
| P-asv3797 | CCGAGCGTTATTCGGAATCACTGGGCTTAAAGCGTACGTAGGCTGGTATATAGGCATTTTGTGAAATCCCTCGGCTCAACCGAGGAACTGCAGAGTGAACCATATACCTTGAGACAGACAGAGGTCGTTGGAACG<br>ATTGGTGGAGTGGTGAAATGCGTTGATATCAATCGGAACGCCAATGGAGAAGTCAAGCGACTGGGTCTGTTCTGACGCTGAGGTACGAAAGCGTGGGT |
| P-asv3798 | TCAAACGTTGTCCGGAATTATTGGGCGTAAAGGGCGCGTAGGCGGCCGGGTAAGTGAGATGTGAAAGCTCTGGGCTCAACCCAGGAATTGCATTTCATACTGCTCGGCTTGAGTTCGGTAGAGGAGGGCGGAAT<br>TCCAGAAGTAGCGGTGAAATGCGTTGATATCTGGAGGAACACCCGTGGCGAAGGCGGCTCTCTGGTCCGATACTGACGCTGAGGCGCGAAAGGTAGGGG |
| P-asv3799 | GCAAGCGTTGTTCGGAGTTACTGGGCGTAAAGAGTGTGTAGGCGGTTCTCTAAGTTCGGTGTGAAATCTCCCGGCTCAACTGGGAGGGTGCGCCGAGAACTGGAGGGCTCGAGTATGGGAGAGGAAAGCGGAA<br>TTCCTGGTGTAGCGGTGAAATGCGTAGATATGTGGCGGAACACCGATGGCGAAGGCAGCCCCCTGGGCTGAGATTGACGCTCAGGCACGAAAGCGTGGGG |
| P-asv3800 | GCGAGCGTTGTCCGGATTTATTGGGCGTAAAGAGCTCGTAGGCGGTTTGGTAAGTCGGATGTGAAATCTCCACGCTCAACGTGGAGGGGCCATTCGAGACTGCTATGACTAGAGTCCGGTAGAGGAATGTGGAA<br>TTCCCGGTGTAGCGGTGAAATGCGCAGAGATCGGGAGGAACACCCGTAGCGAAGGCGGCATTCTGGGCCGGTACTGACGCTGAGGAGCGAAAGCGTGGGG |
| P-asv3801 | GCGAACGTTGTTCGGAATCACTGGGCTTAAAGGGCGCGTAGGCGGGTGCTCAAGTCCGGGGTGAAATCCCACAGCTCAACTGTGGAACTGCCTCGGATACTGGGCGTCTCGAGGGTGGCAGGGGTATGCGGAA<br>CTATCGGTGGAGCGGTGAAATGCGTTGATATCGATAGGAACTCCGGTGGCGAAGGCGGCTTACTGGTCCACTTCTGACGCTGAGGCGCGAAAGCCAGGGG |
| P-asv3802 | GCGAGCGTTGTTCGGAATCACTGGGCGTAAAGCGCGTGTAGGCGGCCGGGCAAGTCGGATGTGAAATCCCGAGGCTCAACCTCGGAACTGCATTCGAAACTGCTCGGCTCGAGTCCCGGAGAGGATGGCAGAA<br>TTCCCGGTGTAGGGGTGAAATCCGTAGATATCGGGAGGAATACCAGTGGCGAAGGCGGCCATCTGGACGGTGACTGACGCTGAGACGCGAAAGCGTGGGG |
| P-asv3803 | GCTAGCGTTGTTCGGAATTACTGGGCGTAAAGGGCGCGTAGGCGGCTAAATAAGTCAAAGGTGAAATCCCTCGGCTCAACTGAGGAACTGCCTTTGAAACTGTTTGGCTTGAGGCCGGGAGGGGGTAGCGGAAT<br>TCCCAATGTAGCGGTGAAATGCGTAGATATTGGGAGGAACACCAGTGGCGAAGGCGGCTACCTGGACCGGTACTGACGCTGAGGCGCGAAAGCGTGGGT |
| P-asv3804 | GCGAGCGTTGTCCGGAGTTACTGGGCGTAAAGGGTGCGCAGGCGGCCGCTTAAGTTTGTGGTGAAATCTCCTGGCTTACCTGGGAGGGTGCCACGAAGACTGGGGGGCTTGAGGCTCGGGGAGGGGCGTGGA<br>ATTCCGGGTGGAGCGGTGAAATGCGTAGAGATCCGGAGGAACACCGACGGCGTAAGCAGCGCCCTAAACGAGACCTGACGCCCAGGCACGAAAGCTGGGGG |
| P-asv3805 | GCAAGCGTTGTCCGGAATTATTGGGCGTAAAGAGCTCGTAGGCGGTCTGTTGCGTCCGCTGTGAAATCTCGGGGCTTAACCCCGAGCCTGCAGTGGATACGGGCAGACTAGAGGTAGGTAGGGGAGAATGGAA<br>TTCCTGGTGTAGCGGTGAAATGCGCAGATATCAGGAGGAACACCGGTGGCGAAGGCGGTTCTCTGGGCCTTACCTGACACTGAGGAGCGAAAGCGTGGGG |
| P-asv3806 | GCTAGCGTTGTTCGGATTTACTGGGCGTAAAGCGCGTGTAGGCGGGTCGTTAAGTTGGGGGTGAAATCCCGAGGCTCAACCTCGGAACTGCCTTCAAAACTGGCGACCTGGACTCAAGTAGAGACAAGTGGAAT<br>TGCGAGTGTAGAGGTGAAATTCGTAGATATTCGCAGGAACACCAGTGGCGAAGGCGACTTGCTGGACTTGTAGTGACGCTGAGACGCGAAAGCGTGGGG |
| P-asv3807 | GCAAGCGTTGTCCGGAATCATTGGGCGTAAAGAGCGTGTAGGCGGCCCGACTAGTCCACTGTGAAAGTCCAGGGCTCAACCCTGGAATGCCGGTGGATACTGTCGGGCTAGAGTCCGGAAGGGGCGAGTGGAA<br>TTCCTGGTGTAGCGGTGAAATGCGCAGATATCAGGAGGAACACCAATGGCGAAGGCAGCTCGCTGGGACGTGACTGACGCTGAGACGCGAAAGCGTGGGG |
| P-asv3808 | GCAAGCGTTGTTCGGAATTATCGGGCGTAAAGGGCGCGTAGGCGGCGCGACAAGTCACCTGTGAAATCCCCGGGCTTAACCCGGGGCCTGCAGGCGAAACTGTCGTGCTGGAGTATGGGAGAGGTGCGTGGA<br>ATTCCCGGTGTAGCGGTGAAATGCGTAGATATCGGGAGGAACACCTGTGGCGAAAGCGGCGCACTGGACCATAACTGACGCTGAGGCGCGAAAGCTAGGGG |
| P-asv3809 | ACGAACGTTATTCGGAATCACTGGGCTTAAAGGGTGCGTAGGCTGCGCGGAAAGTTGGGTGTGAAATCCCTCGGCTCAACCGAGGAACTGCGCCCAAAACTACCGTGCTCGAGGGAGACAGAGGTGAGCGGAA<br>CTTAGGGTGGAGCGGTGAAATGCGTTGATATCCTAAGGAACACCCGTGGCGAAAGCGGCTCACTGGGTCTCTTCTGACGCTGAGGCACGAAAGCTAGGGT |
| P-asv3810 | GCAAGCGTTATCCGGATTCACTGGGTTTAAAGGGTGCGTAGGTGGGCAGTTAAGTCAGTGGTGAAATCTCCGGGCTCAACCCGGAAACTGCCATTGATACTATCTGTCTTGAATATCGTGGAGGTAAGCGGAATA<br>TGTCATGTAGCGGTGAAATGCTTAGATATGACATAGAACACCAATTGCGAAGGCAGCTTACTACCCGAATATTGACACTGAGGCACGAAAGCGTGGGG |
| P-asv3811 | GCGAGCGTTGTTCGGAATTATTGGGCGTAAAGGGCGCGTAGGCGGTGCCACAAGTCACCTGTGAAACCTCTGGGCTCAACTCAGAGCCTGCAGGCGAAACTGTGGTGCTGGAGTGTGGGAGAGGTGCGTGGAA<br>TTCCCGGTGTAGCGGTGAAATGCGTAGATATCGGGAGGAACACCTGTGGCGAAAGCGGCGCACTGGACCACAACTGACGCTGAGGCGCGAAAGCCAGGGG |
| P-asv3812 | GCAAGCGTTGTTCGGAATTACTGGGCGTAAAGGGCGCGTAGGCGGCCCGCTAAGTCGAACGTGAAATCCCCGGGCTCAACCCGGGAACTGCGTCCGATACTGGCAGGCTTGAATCCGGGAGAGGGATGCGGA<br>ATTCCAGGTGTAGCGGTGAAATGCGTAGATATCTGGAGGAACACCGGTAGCGAAGGCGGCATCCTGGACCGGTATTGACGCTGAGGCGCGAAAGCCAGGGG |
| P-asv3813 | GCAAGCGTTGTCCGGAATCATTGGGCGTAAAGCGCGCGTAGGCGGCCCTGTAAGTCCGTTGTGAAAGTCAAAGGCTCAACCTTTGAAAGCCGATGGATACTGCAGGGCTAGAGTCCGGAAGAGGCGAGTGGAA<br>TTCCTGGTGTAGCGGTGGAATGCGCAGATATCAGGAGGAACACCAATAGCGAAGGCAGCTCGCTGGGACGGTACTGACGCTAAGGCGCGAAAGCGTGGGG |
| P-asv3814 | CCGAGTGGTCGGGACGTTTACTGGGCCTAAAGCATCCGTAGCCGGTTCTGCAAGTCTTCCGTTAAATCCACCTGCTTAACAGATGGGCTGCGGAAGATACTACAGGGCTAGGAGGCGGGAGAGGCAAGCGGTAC<br>TCGATGGGTAGGGGTAAAATCCGTTGATCCATTGAAGACCACCAGTGGCGAAGGCGGCTTGCCAGAACGCGCTCGACGGTGAGGGATGAAAGCTGGGGG |
| P-asv3815 | CCAAGCGTTAATCGGAATTACTGGGCGTAAAGCGTGAGTAGGCGGTTTCGAAAGTCCGATGTGAAATCCCCGGGCTTAACCTGGGAACTGCGTTGGAAACTACGAGACTAGAGTGCGGCAGAGGGGGGTGGAAT<br>TCCAGGTGTAGCAGTGAAATGCGTAGATATCTGGAGGAACACCGATGGCGAAGGCAGCCCCCTGGGTTAGCACTGACGCTCATACACGAAAGCGTGGGG |
| P-asv3816 | GCGAGCGTTGTTCGGAATTATTGGGCGTAAAGGGCGCGTAGGCGGCGCGGTAAGTCACCTGTGAAATCTCCTGGCTCAACTGGGAGCTTGCAGGCGAAACTGCCGTGCTGGAGTGTGGGAGAGGTGCGTGGA<br>ATTCCTGGTGTAGCGGTGAAATGCGTAGATATCAGGAGGAACACCTGTGGCGAAAGCGGCGCACTGGACCACAACTGGCGCTGAGGCGCGAAAGTTAGGGG |
| P-asv3817 | CCGAGCGTTAGGCGGAATCACTGGGCTTAAAGCGTGTGTAGGCGGATTTGTAAGTACCTTGTGAAATCCCACGGCTTAACCGTGGAACTGCTTGGTATACTGCTTATCTTGGCCATCTAGGGGCGAGCGGAACA<br>AACGGTGGAGCGGTGAAATGCGTAGATATCGTTTGGAACGCCAATGGTGAAAACAGCTCGCTGGGGATGTGCCGACGCTGAGATACGAAAGCCAGGGG |
| P-asv3818 | GCTAGCGTTGCTCGGAATCACTGGGCGTAAAGCGCACGTAGGCGGATCGTTAAGTCAGGGGTGAAATCCTGGAGCTCAACTCCAGAACTGCCTTTGATACTGACGATCTCGAGTTCGGGAGAGGTGAGTGGAAC<br>TGCGAGTGTAAAGGTGAAATTCGTAGATATTCGCAAGAACACCAGTGGCGAAGGCGGCTCACTGGCCCGATACTGACGCTGAGGTGCGAAAGCGTGGGG |
| P-asv3819 | GCAAGCGTTATCCGGATTTACTGGGTTTAAAGGGTGCGTAGGTGGGTAAGTAAGTCAGTGGTGAAATCTCCGGGCTTAACCCGGAAACTGCCATTGATACTATTTATCTTGAATACTTTGGAGGTAAGCGGAATAT<br>GTCATGTAGCGGTGAAATGCTTAGATATGACATAGAACACCAATTGCGAAGGCAGCTTACTACACAGTGATTGACACTGAGGCACGAAAGCGTGGGG |
| P-asv3820 | GCTAGCGTTATTCGGATTTATTGGGCGTAAAGGGCGCGTAGGCGGTGCGGTAAGTCTGCGGTGAAATCTCCCGGCTCAACTGGGAGTCTGCCGTGGAAACTGCCGTGCTTGAGTGTGGGAGAGGTGAGTGGAA<br>TTCCCGGTGTAGCGGTGAAATGCGTAGATATCGGGAGGAACACCTGTGGCGAAAGCGGCTCACTGGACCACAACTGACGCTGAGGCGCGAAAGCCAGGGG |
| P-asv3821 | GCGAGCGTTGTTCGGAATTATTGGGCGTAAAGGGCGCGTAGGCGGTGCCACAAGTCACTTGTGAAATCCCCGGGCTTAACTCCGGGCCTGCAGGCGAAACTGTGGTGCTGGAGTGTGGGAGAGGTGCGTGGAA<br>TTCCCGGTGTAGCGGTGAAATGCGTAGATATCGGGAGGAACACCTGTGGCGAAAGCGGCGCACTGGACCATAACTGACGCTGAGGCGCGAAAGCTAGGGG |
| P-asv3822 | GCTAGCGTTGTCCGGAATCATTGGGCGTAAAGCGCGCGTAGGCGGCCCGGTAAGTCCGCTGTGAAAGTCAAAGGCTCAACCTTTGAATGTCAGCGGATACTGCCGGGCTAGAGTCCGGAAGAGGCGAGTGGAA<br>CTCCTGGTGTAGCGGTGGAATGCGCAGATATCAGGAAGAACACCAACGGCGAAGGCAGCTCGCTGGGACGGAACTGACGCTAAGGCGCGAAAGCGTGGGG |
| P-asv3823 | GCAAGCGTTGCTCGGAATCACTGGGCGTAAAGGGCGTGTAGGCGGCTTCGTGTGTCTGGCGTGAAAGCCCCGGGCTTACCCCGGGAAGTGCGTTGGAAACTGCGGAGCTAGAGTCTCGGAGAGGGAGGTGGA<br>ATTCCAGGTGTAGCGGTGAAATGCGTAGATATCTGGAAGAACACCGGTGGCGAAGGCGGCCTCCTGGACGGTGACTGACGCTGAGACGCGAAAGCGTGGGG |
| P-asv3824 | GCAAGCGTTAATCGGAATTACTGGGCGTAAAGCGTGCGTAGTCGGTTATCTAAGTCGGATGTGAAATCCCCGGGCTCAACCTGGGAATTGCATTCGATACTGGGTAGCTAGAGTTCGGCAGAGGGAAGCGGAAT<br>TTCGGGTGTAGCGGTGAAATGCGTAGATATCCGAAGGAACATCAGTGGCGAAAGCGGCTTCCTGGACCAGAACTGACGATCAGGCACGAAAGCGTGGGG |
| P-asv3825 | ACTAGCGTTGCTCGGAATCACTGGGCGTAAAGCGCACGTAGGCGGCTTTTTAAGTCAGGGGTGAAATCCTGGAGCTCAACTCCAGAACTGCCTTTGATACTGGAAAGCTCGAGTCCGGGAGAGGTGAGTGGAAC<br>TGCGAGTGTAGAGGTGAAATTCGTAGATATTCGCAAGAACACCAGTGGCGAAGGCGGCTCACTGGCCCGGTACTGACGCTGAGGTGCGAAAGCGTGGGG |
| P-asv3826 | GCAAGCGTTGTCCGGAATTATTGGGCGTAAAGAGCTCGTAGGTGGTCTGCCACGTCTGTTGTGAAAATCCAGGGCTTAACCCTGGACTTGCTGCGGATACGGGCAGACTAGAGGTAGGTAGGGGAGAATGGAAT<br>TCCCGGTGTAGCGGTGAAATGCGCAGATATCGGGAGGAACACCGGTGGCGAAGGCGGTTCTCTGGGCCTTACCTGACACTGAGGAGCGAAAGCGTGGGG |

| | |
|---|---|
| P-asv3827 | GCTAGCGTTGTTCGGAATTACTGGGCGTAAAGGGCGCGTAGGCGGCCGATTAAGTTAGATGTGAAAGTCCTGGGCTCAACCTGGGGACTGCATTTAATACTGGTTGGCTTGAATTCGGGAGAGGATAGCGGAATTCCCAGTGTAGAGGTGAAATTCGTAGATATCGGGAGGAACACCAGTGGCGAAGGCGGCGACCTGGACGGACACTGACGCTGAGACGCGAAAGCGTGGGG |
| P-asv3828 | GCAAGCGTTGTCCGGATTTATTGGGCGTAAAGAGCGTGTAGGCGGCCATGTAGGTCCGTTGCGAAAACTCGAGGCTCAACCTCGAGACGCCGATGGAAACCATGTGGCTAGAGTCCGGAAGAGGAGAGTGGAATTCCTGGTGTAGCGGTGAAATGCGCAGATATCAGGAAGAACACCCGTGGCTAAGGTGGCTCTCTAGTACGGTACTGACGCTGAGACGCGAAAGCGTGGGG |
| P-asv3829 | GCTAGCGTTGCTCGGAATTACTGGGCGTAAAGGGAGCGTAGGCGGATCGTTAAGTTGGGGGTGAAAGCCCGGGGCTCAACCTCGGAACTGCCTTCAATACTGGCGATCTTGAGTGTGAGAGAGGTGAGTGGAACTCCGAGTGTAGAGGTGAAATTCGTAGATATTCGGAAGAACACCAGTGGCGAAGGCGACTCACTGGCTCATTACTGACGCTGAGGCTCGAAAGCGTGGGG |
| P-asv3830 | GCAAGCGTTGCCCGGATTTATTGGGCGTAAAGGGCGCGCAGGCGGCTCGGCACGTCCGTTGTGAAATCGCCCGGCTCAACTGGGCGGGGTCAGCGGATACGGCCGAGCTGGAGCAAGCTAGGGGGCAATGGAATTCCCGGTGTAGTGGTGGAATGCGTAGATATCGGGAGGAACACCAGTGGCGAAGGCGGTTGCCTGGAGCTTTGCTGACGCTGAGGCGCGAAAGCGTGGGG |
| P-asv3831 | GCGAGCGTTGTCCGGAATTATTGGGCGTAAAGAGCTCGTAGGCGGCTTGTTGCGTCGACTGTGAAAACCCGCGGCTCAGCCGTGTGGCTTGCAGTCGATACGGGCAGGCTAGAGTTCGGTAGGGGAGACTGGAATTCCTGGTGTAGCGGTGAAATGCGCAGATATCAGGAGGAACACCGGTGGCGAAGGCGGGTCTCTGGGCCGATACTGACGCTGAGGAGCGAAAGCGTGGGG |
| P-asv3832 | GCGAGCGTTAATCGGAATTACTGGGCGTAGAGCGTACGTAGACGGTCAGATAAGTCGGGTGTGAAAGCCCCGGGCTCAACCTGGGAATTGCATTCGAGACTGTCTGGCTAGGGTGCGGAAGAGGGAAGCAGAATTCCCGGTGTAGCGGTGAAATGCGTAGATATCGGGAGGAATACCAGTGGCGAAAGCGGCTTCCTGGTCCAGCACCGACGTTCAGGTACGAAAGCATGGGG |
| P-asv3833 | GCAAGCGTTGTCCGGATTTATTGGGCGTAAAGAGCGTGTAGGCGGCCATACAGGTCCGTTGTGAAAACTGGAGGCTTAACCTCCAGACGTCGATGGAAACCGTATGGCTAGAGTCCGGAAGAGGAGAGTGGAATTCCTGGTGTAGCGGTGAAATGCGCAGATATCAGGAAGAACACCTATGGCGAAGGCAGCTCTCTGGGACGGTACTGACGCTGAGACGCGAAAGCGTGGGG |
| P-asv3834 | GCAAGCGTTATCCGGATTCACTGGGTTTAAAGGGAGCGCAGGCGGATTTGTAAGCCAGTGGTGAAAGGCTTAAGCTTAACTTGGGAACTGCCATTGATACTATAAGTCTTGAATGTTGTTGAGGTTTGCGGAATGGGTCATGTAGCGGTGAAATGCATAGATATCAGGAGGAACACCGGTGGCGAAGGCGGCCCTCTGGACGGATACTGACGCTGAGACGCGACAGCGTGGGG |
| P-asv3835 | GCAAGCGTTGTCCGGATTTATTGGGCGTAAAGAGCGTGTAGGCGGCCAGACAGGTCCGTTGTGAAAACTCGAGGCTCAACCTCGAGATGTCGATGGAAACCGTCTGGCTAGAGTCCGGAAGAGGAGTGTGGAATTCCTGGTGTAGCGGTGAAATGCGCAGATATCAGGAAGAACACCTATGGCGAAGGCAGCACTCTGGGACGGTACTGACGCTGAGACGCGAAAGCGTGGGG |
| P-asv3836 | GCAAGCGTTGTCCGGAATCATTGGGCGTAAAGAGCATGTAGGCGGCCCGGTCAGTCGGTTGTGAAAGTCAGGGGCTCAACCCTTGAATGCCGACCGATACTGCCGGGCTAGAGTCCGGAAGAGGCGAGTGGAATTCCTGGTGTAGCGGTGAAATGCGCAGATATCAGGAGGAACACCAATGGCGAAGGCAGCTCGCTGGGACGGTACTGACGCTGAGACGCGAAAGCGTGGGG |
| P-asv3837 | CCAAGCGTTGTTCGGATTCACTGGGCGTAAAGGGTGCGTAGGTGGTCGGGTAAGTCGGTTGTGAAATCTCGGAGCTCAACTCCGAAACTGCATTGGAAACTACCCGGCTGGAGGGTTGGAGGGGGGACTGGAATTCTCGGTGTAGCAGTGAAATGCGTAGATATCGAGAGGAACACCAGTGGCGAAGGCGAGTCCCTGGACAACTCCTGACACTGAGGCACGAAAGCTAGGGG |
| P-asv3838 | GCAAGCGTTGTTCGGAATTACTGGGCGTAAAGGGCTCGTAGGTTGCCAACTAAGTCAGACGTGAAATCCCTCAGCTTAACTGGGGAACTGCGTCTGATACTGGATGGCTTGAGATTGGGAGAGGGATGCGGAATTCCAGGTGTAGCGGTGAAATGCGTAGATATCTGGAGGAACACCGGTGGCGAAGGCGGCATCCTGGACCAACACTGACACTGAGGAGCGAAAGCCAGGGG |
| P-asv3839 | GCGAGCGTTATTCGGAGTTACTGGGAGTAAAGGGTGTGTAGGCGGTTCTTTAAGTTTGTTGTGAAATCTCCCGGCTTAACCGGGAGGGTGCGGCGAATACTGAGGGACTAGAGTGTGGGAGAGGAAAGTGGAATTCCTGGTGTAGCGGTGAAATGCGTAGATATCAGGAGGAACACCGGTGGTGTAGACGGCTTTCTGGACCATAACTGACGCTGAGACACGAAAGCGTGGGT |
| P-asv3840 | GCAAGCGTTGTTCGGAATTACTGGGCGTAAAGCGCGCGTAGGCGGCCCTGTAAGTCCGTTGTGAAAGTCAAAGGCTCAACCTCTGAAAGCCGATGGATACTGCAGGGCTAGAGTCCGGAAGAGGCGAGTGGAATTCCTGGTGTAGCGGTGGAATGCGCAGATATCAGGAGGAACACCAATAGCGAAGGCAGCTCGCTGGGACGGTACTGACGCTAAGGCGCGAAAGCGTGGGG |
| P-asv3841 | GCGAGCGTTGTTCGGAATTATTGGGCGTAAAGGGTGCGTAGGCGGTTTGGTAAGTCTTGTGTGAAATCTATGGGCTCAACTCATAGTCTGCACGAGAAACTGCCGGGCTTGAGTGTGGGAGAGGTGAGTGGAATTTCCGGTGTAGCGGTGAAATGCGTAGATATCGGAAGGAACACCTGTGGCGAAAGCGGCTCACTGGACCACAACTGACGCTGATGCACGAAAGCTAGGGG |
| P-asv3842 | GCTAGCGTTGCTCGGAATCATTGGGCGTAAAGGGCGCGTAGGGGACTCTTAAGTCGGGGGTGAAAGCCCAGGGTTCAACCCTGGAATTGCCTTCGATACTGAGAGTCTTGAGTTCGGAAGAGGTTGGTGGAACTGCGAGTGTAGAGGTGAAATTCGTAGATATTCGCAAGAACACCAGTGGCGAAGGCGGCCAACTGGTCCGATACTGACGCTGAGGCGCGAAAGCGTGGGG |
| P-asv3843 | GCGAGCGTTGTCCGGAATCACTGGGCGTAAAGGGCGCGTAGGCGGCTTGGTAAGGGGGCGGTGAAAGCCCGGGGCTCAACCCCGGGTCGGCCGGACCGACTGCCGAGCTTGAGCACACTAGGGGCGGGTGGAATTCCGGGTGTAGCGGTGGAATGCGTAGAGATCCGGAGGAACACCGGTGGCGAAGGCGGCCCGCCGGGGTGTGGCTGACGCTGAGGCGCGACAGCGTGGGG |
| P-asv3844 | GCTAGCGTTGTTCGGAATTACTGGGCGTAAAGCGCACGTAGGCGGATATTTAAGTCAGGGGTGAAATCCCAGAGCTCAACTCTGGAACTGCCTTTGATACTGGGTATCTTGAGTATGGAAGAGGTAAGTGGAATTGCGAGTGTAGAGGTGAAATTCGTAGATATTCGCAGGAACACCAGTGGCGAAGGCGGCTTACTGGTCCATAACTGACGCTGAGGTGCGAAAGCGTGGGG |
| P-asv3845 | GCAAGCGTTGTCCGGAATTACTGGGCGTAAAGAGCTCGTAGGTGGTTTGTCGCGTCGTTTGTGGAATACCGCAGCTTAACTGTGGGGTTGCAGGCGATACGGGCATAACTTGAGTGCTGTAGGGGAGACTGGAATTCCTGGTGTAGCGGTGGAATGCGCAGATATCAGGAGGAACACCGATGGCGAAGGCAGGTCTCTGGGCAGTAACTGACGCTGAGGAGCGAAAGCATGGGT |
| P-asv3846 | GCGAGCGTTGTCCGGAATTATTGGGCGTAAAGAGCTTGTAGGCGGTTTGTCGCGTCTGCTGTGAAAGACCGGGGCTTAACTCCGGTTCTGCAGTGGGTACGGGCAGACTAGAGTATGGTAGGGGAGACTGGAATTCCTGGTGTAGCGGTGAAATGCGCAGATATCAGGAGGAACACCGATGGCGAAGGCAGGTCTCTGGGCCATTACTGACGCTGAGAAGCGAAAGCATGGGG |
| P-asv3847 | CCGAGCGTTGTCCGGAATTATTGGGCGTAAAGAGCTCGTAGGCGGCGTGTCGCGTCTGCTGTGAAAGACCGGGGCTTAACTCCGGTTTTGCAGTGGATACGGGCATGCTAGAGGTAGGTAGGGGAGACTGGAATTCCTGGTGTAGCGGTGGAATGCGCAGATATCAGGAGGAACACCGGTGGCGAAGGCGGGTCTCTGGGCCTTACCTGACGCTGAGGAGCGAAAGCATGGGG |
| P-asv3848 | GCAAGCGTTAATCGGAATTACTGGGCGTAAAGCATGCGCAGGCGGTTTTGTAAGTCTGATGTAAAATCACCGGGCTCAACCTGGGAACTGCATTGGAGACTGCAAGGCTAGAGTGTGTCAGAGGGGGGTAGAATTCCACGTGTAGCAGTGAAATGCGTAGAGATGTGGAGGAATACCGATGGCGAAGGCAGCCCCCTGGGATAACACTGACGCTCATGCACGAAAGCGTGGGG |
| P-asv3849 | GCGAGCGTTAATCGGATTTACTGGGCGTAAAGCGTGCGTAGGCGGCTAATTGAGTCGGATGTGAAATCCCCGAGCTTAACTTGGGAATTGCATTCGATACTTGTTAGCTAGAGTGTGGGAGAGGATGGTAGAATTCCAGGTGTAGCGGTGAAATGCGTAGAGATCTGGAGGAATACCGATGGCGAAGGCAGCCATCTGGCCTAACACTGACGCTGAGGTACGAAAGCATGGGG |
| P-asv3850 | ACAAGCGTTATCCGGATTTATTGGGCGTAAAGCGCGTCTAGGTGGTTATGTAAGTCTGATGTGAAAATGCAGGGCTCAACTCTGTATTGCGTTGGAAACTGTATGACTAGAGTACTGGAGAGGTAAGCGGAACTACAAGTGTAGAGGTGAAATTCGTAGATATTTGTAGGAATGCCGATGGGGAAGCCAGCTTACTGGACAGATACTGACGCTAAAGCGCGAAAGCGTGGGT |
| P-asv3851 | GCTAGCGTTGCTCGGAATCACTGGGCGTAAAGGGCGCGTAGGCGGACTTTTAAGTCGGAGGTGAAAGCCCAGGGCTCAACCCTGGAATTGCCTTCGATACTGGGAGTCTTGAATTCGGAAGAGGTTGGTGGAACTGCAAGTGTAGAGGTGAAATTCGTAGATATTCGCAAGAACACCGGTGGCGAAGGCGGCCAACTGGTCCGATACTGACGCTGAGGCGCGAAAGCGTGGGG |
| P-asv3852 | CCGAGCGTTGTCCGGAATCATTGGGCGTAAAGGGTACGTAGGCGGCCTAGTAAGTTAGAAGTGAAATAATATAGCTCAACTATATAAAGCTTTTAAAACTGTTAGGCTTGAGAGATGAAAGGGAAAGTGGAATTCCTAGTGTAGCGGTGAAATGCGCAGATATTAGGAAGAATACCGGTGGCGAAGGCGACTTTCTGGTCATCATCTGACGCTGAGGTACGAAAGCGTGGGT |
| P-asv3853 | GCAAGCGTTGTCCGGAATTATTGGGCGTAAAGCGCGCGCAGGCGGCCTTTTAAGTCTGATGTGAAAGCCCCCGGCTCAACCGGGGAGGGTCATTGGAAACTGGAAGGCTTGAGTACAGAAGAGAAGAGTGGAATTCCATGTGTAGCGGTGAAATGCGTAGAGATGTGGAGGAACACCAGTGGCGAAGGCGACTCTTTGGTCTGTAACTGACGCTGAGGCGCGAAAGCGTGGGG |
| P-asv3854 | GCGAGCGTTAATCGGAATTACTGGGCGTAAAGGGAGCGCAGACGGTTACTTAAGCAGGATGTGAAATCCCCGGGCTCAACCTGGGAACTGCGTTCTGAACTGGGTGACTAGAGTGTGTCAGAGGGAGGTAGAATTCCACGTGTAGCAGTGAAATGCGTAGAGATGTGGAGGAATACCGATGGCGAAGGCAGCCTCCTGGGATAACACTGACGTTCATGCTCGAAAGCGTGGGT |
| P-asv3855 | GCGAGCGTTAATCGGAATTACTGGGCGTAAAGCGCGCGTAGGCGGTCTGATAAGCCGGTTGTGAAAGCCCCGGGCTCAACCTGGGAACGGCATCCGGAACTGTCAGGCTAGAGTGCAGGAGAGGAAGGTAGAATTCCCGGTGTAGCGGTGAAATGCGTAGAGATCGGGAGGAATACCAGTGGCGAAGGCGGCCTTCTGGTCAGACACTGACGCTGAGGTGCGAAAGCGTGGGT |
| P-asv3856 | GCGAACGTTGTTCGGAATAACTGGGCTTAAAGGGCGCGTAGGCGGGCTGACAAGTCTGGGGTGAAATCCCGCGGCTCAACCGTGGAACTGCCTTAGATACTGTCGGCCTCGAGGAAGGTAGGGGCATGCGGAACTAACGGTGGAGCGGTGAAATGCGTTGATATCGTTAGGAACTCCGGTGGCGAAAGCGGCGTGCTGGACCTTTTCTGACGCTGAGGCGCGAAAGCTAGGGG |
| P-asv3857 | GCAAGCGTTATCCGGATTTACTGGGTGTAAAGGGAGCGTAGACGGAATGGCAAGTCTGAAGTGAAATACCCGGGCTCAACCTGGGAACTGCTTTGGAAACTGTTGTTCTAGAGTGTTGGAGAGGTAAGTGGAATTCCTGGTGTAGCGGTGAAATGCGTAGATATCAGGAAGAACACCGGAGGCGAAGGCGGCTTACTGGACAATAACTGACGTTGAGGCTCGAAAGCGTGGGG |
| P-asv3858 | GCAAGCGTTACTCGGAATTACTGGGCGTAAAGCGTGCGTAGGTGGTCTGTTAAGTCTGATGTGAAAGCCCTGGGCTCAACCTGGGAATGGCATTGGAAACTGGCGGGCTAGAGTGCGGTAGAGGGTAGTGGAATTCCCGGTGTAGCAGTGAAATGCGTAGAGATCGGGAGGAACATTCGTGGCGAAGGCGGCTACCTGGACCAGCACTGACACTGAGGCACGAAAGCGTGGGG |
| P-asv3859 | GCTAGCGTTGTTCGGAATTACTGGGCGTAAAGCGCACGTAGGCGGCTTTGTAAGTTAGAGGTGAAAGCCTGGAGCTTAACTCCAGAACTGCCTTTAAGACTGCATCGCTTGAATCCGGGAGAGGTGAGTGGAATTCCGAGTGTAGAGGTGAAATTCGTAGATATTCGGAAGAACACCAGTGGCGAAGGCGGCTCACTGGACCGGTATTGACGCTGAGGTGCGAAAGCGTGGGG |
| P-asv3860 | CCGAGCGTTGTTCGGAATTATTGGGCGTAAAGCGGGTGTAGGTGGCTTAGCAAGTCAGATGTGAAAGCCCTGGGCTCAACCCAGGAAGTGCGTCTGATACTGCTTTGCTAGAGCACGGTAGAGGGAACTGGAATTCCAGGTGTAGGGGTGAAATCCGTAGATATCTGGAGGAATACCAATGGCGAAGGCAGGTTCCTGGTCCGTTGCTGACACTCAGACCCGAAAGCGTGGGG |
| P-asv3861 | GCTAGCGTTGTTCGGAATTACTGGGCGTAAAGGGCGCGTAGGCGGCCGGCTAAGTTAGATGTGAAAGTCCTGGGCTCAACCTGGGGACTGCATTTAATACTGGTTGGCTTGAATTCGGGAGAGGATAGCGGAATTCCCAGTGTAGAGGTGAAATTCGTAGATATTGGGAAGAACACCGGTGGCGAAGGCGGCTATCTGGACCGACATTGACGCTGAGGCGCGAAAGCGTGGGG |
| P-asv3862 | GCAAGCGTTATCCGGATTTACTGGGTTTAAAGAGTGCGTAGGCGGTGTTATAAGTCAGTGGTGAAAGGTCGTAGCTTAACTACGGAATTGCCATTGATACTGTAATGCTAGAGTGGGTACGAGGTACTTGGAATGCGTCATGTAGCGGTGAAATGCTTAGATATGACGTAGAACACCAAAAGCGAAGGCAGAGTACCAGTTAACCACTGACGCTGAGGCACGAAAGCGTGGGT |
| P-asv3863 | GCAAGCGTTAATCGGAATTACTGGGCGTAAAGAGTGCGTAGGCGGATAATTAAGTCATATGTGAAAGCCCCGGGCTTAACCTGGGAATTGCATCTGATACTGGTTATCTAGAGTATGGTAGAGGATAGTGGAATTTCTGGTGTAGCGGTGAAATGCGTAGATATCAGAAGGAACACCAGTGGCGAAGGCGACTGTCTGGGCCAATACTGACGCTGAGGTACGAAAGCGTGGGT |
| P-asv3864 | GCAAGCGTTGTCCGGAATTATTGGGCGTAAAGAGCTCGTAGGCGGTTTGTCGCGTCTGCTGTGAAAACCCGAGGCTCAACCTCGGGCCTGCAGTGGGTACGGGCAGACTAGAGTGTGGTAGGGGAGACTGGAATTCCTGGTGTAGCGGTGGAATGCGCAGATATCAGGAGGAACACCGATGGCGAAGGCAGGTCTCTGGGCCACAACTGACGCTGAGAAGCGAAAGCGTGGGG |
| P-asv3865 | GTTAGCGTTGTTCGGAATTACTGGGCGTAAAGCGCACGTAGGCGGATTGGAAAGTTGGGGGTGAAATCCCGGGGCTCAACCCCGGAACTGCCTCCAAAACTATCAGTCTAGAGTTCGAGAGAGGTGAGTGGAATTCCAAGTGTAGAGGTGAAATTCGTAGATATTTGGAGGAACACCAGTGGCGAAGGCGGCTCACTGGCTCGATACTGACGCTGAGGTGCGAAAGTGTGGGG |

| | |
|---|---|
| P-asv3866 | GCTAGCGTTGTTCGGAATTACTGGGCGTAAAGCGCACGTAGGCGGATCGTTAAGTGAGGGGTGAAATCCCAGGGCTCAACCCTGGAACTGCCTCTCATACTGGCGATCTCGAGTTCGAGAGAGGTGAGTGGAATTGCGAGTGTAGAGGTGAAATTCGTAGATATTCGCAGGAACACCAGTGGCGAAGGCGGCTCACTGGCTCGATACTGACGCTGAGGTGCGAAAGCGTGGGG |
| P-asv3867 | TCCCTCTTTCCTCTTCCTTCCTGGGCGTAAAGCGCGCGTAGGTGGTTAGTTAAGTTGGATGTGAAATCCCCGGGCTCAACCTGGGAACTGCATTCAAAACTGACTGACTAGAGTATGGTAGAGGGTGGTGGAATTTCCTGTGTAGCGGTGAAATGCGTAGATATAGGAAGGAACACCAGTGGCGAAGGCGACCACCTGGACTGATACTGACACTGAGGTGCGAAAGCGTGGGG |
| P-asv3868 | GCGAGCGTTGTCCGGATTTATTGGGCGTAAAGAGTTCGTAGGTGGTTTGTTAAGTTTGGTGTTAAAGATCGGGGCTCAACCCTGGGACTGCACTGAATACTGGCAGACTCGAGTGTGGTAGAGGCTAGTGGAATTCCCAGTGTAGCGGTGAAATGCGTAGATATTGGGAAGAACACCGGTGGCGTAGGCGACTAGCTGGGCCATAACTGACACTGAGGAACGAAAGCCAGGGG |
| P-asv3869 | GCAAGCGTTGTCCGGAATTACTGGGTGTAAAGGGAGTGTAGGCGGGACTGCAAGTCAGATGTGAAATTTATGGGCTCAACCCATAAACTGCATTTGAAACTGTGGTTCTTGAGTGAAGTAGAGGTTGGCGGAATTCCTAGTGTAGCGGTGAAATGCGTAGATATTAGGAGGAACATCAGTGGCGAAGGCGGCCAACTGGGCTTTTACTGACGCTGAGGCTCGAAAGCGTGGGG |
| P-asv3870 | GCGAGCGTTGTCCGGAATTATTGGGCGTAAAGGGCTCGTAGGCGGTTTGTCGCGTCGGGAGTGAAAACACCGGGCTTAACTCGGTGCTTGCTTCCGATACGGGCAGACTAGAGGTATGCAGGGGAGAACGGAATTCCTGGTGTAGCGGTGAAATGCGCAGATATCAGGAGGAACACCGGTGGCGAAGGCGGTTCTCTGGGCATTACCTGACGCTGAGGAGCGAAAGTGTGGGG |
| P-asv3871 | ACAAGTGATGGCCGATATTATTGGGCCTAAAGCGTCCGTAGCCGGCCAGACAAGTCCGTTGGGAAATCGACGTGCCTAACACGTCGGCGTCCAGCGGAAACTGTTTGGCTTGGGGCCAGAAGATCCGAGGGTACGTCCGGGGTAGGAGTGAAATCCCGTAATCCTGGACGGACCACCGGTGGCGAAAGCGCCTCGGAAAGATGGACCCGACGGTGAGGGACGAAAGCTGGGGT |
| P-asv3872 | GCGAGCGTTGTCCGGATTTACTGGGCGTAAAGGGAGCGTAGGCGGACTTTTAAGTGAGATGTGAAATACCCGGGCTCAACTTGGGTGCTGCATTTCAAACTGGAAGTCTAGAGTGCAGGAGAGGAGAATGGAATTCCTAGTGTAGCGGTGAAATGCGTAGAGATTAGGAAGAACACCAGTGGCGAAGGCGATTCTCTGGACTGTAACTGACGCTGAGGCTCGAAAGCGTGGGG |
| P-asv3873 | GCTAGCGTTGTTCGGAATTACTGGGCGTAAAGCGCGCGTAGGCGGATTGGAAAGTTGGGGGTGAAATCCCGGGGCTCAACCCCGGAACTGCCTCCAAAACTATCAGTCTGGAGTTCGAGAGAGGTGAGTGGAATTCCGAGTGTAGAGGTGAAATTCGTAGATATTCGGAGGAACACCAGTGGCGAAGGCGGCTCACTGGCTCGATACTGACGCTGAGGTGCGAAAGCGTGGGG |
| P-asv3874 | GCTAGCGTTGTTCGGAATTACTGGGCGTAAAGCGCACGTAGGCGGACTGTTAAGTCGGGGGTGAAATCCCGGGGCTCAACCCCGGAACTGCCTTCGATACTGGCAGTCTAGAGGACGGTAGAGGCTAGTGGAATGGCGAGTGTAGAGGTGAAATTCGTAGATATTCGCCGGAACACCAGTGGCGAAGGCGACTGGCTGGACCGTATCTGACGCTGAGGTGCGAAAGCGTGGGG |
| P-asv3875 | GCAAGCGTTAATCGGAATCACTGGGCGTAAAGCGCACGTAGGCGGCAACGTAAGTCAGATGTGAAAGCCCCGGGCTCAACCCGGGAACGGCATTTGAGACTGCGTCGCTCGAGTCTGAGAGAGGGGGGTGGAATTCCGGGTGTAGCGGTGAAATGCGTAGAGATCCGGAGGAACACCAGTGGCGAAGGCGGCCCCCTGGCTCAAGACTGACGCTGAGGTGCGAAAGCGTGGGG |
| P-asv3876 | CCGAGCGTTGTCCGGAGTTACTGGGCGTAAAGCGCGCGCAGGCGGCGGTGCTGGCCCGGCGTGAAAGCCCCCGGCTCAACCGGGGAGGGTCGTCGGGGACCGCACCGCTTGAGGGCGGTAGGGGCTGGTGGAATGCCTGGTGTAGTGGTGAAATGCGTAGAGATCAGGCGGAACACCCGTGGCGAAGGCGGCCAGCTGGGCCGTCCCTGACGCTGAGGCGCGAAGGCGTGGGG |
| P-asv3877 | CCAAACGTTATTCGGAATTATTGGGCTTAAAGGGTGCGTAGGCGGTGCAGTAAGTCGGGTGTGAAATCCCTCAGCTCAACTGAGGAACTGCGCCCGATACTGCTGTGCTTGAGGGAGATAGAGGTGAGCGGAACTGATGGTGGAGCGGTGAAATGCGTTGATATCATCAGGAACACCTGTGGCGAAAGCGGCTCACTGGATCTTTTCTGACGCTGAGGCACGAAAGCTAGGGG |
| P-asv3878 | GCAAGCGTTGTCCGGAATTATTGGGCGTAAAGAGCTCGTAGGCGGTTTGTCGCGTCTGCTGTGAAAACCTCAGGCTCAACCTGGGGCTTGCAGTGGGTACGGGCAGACTAGAGTGCGGTAGGGGTGACTGGAATTCCTGGTGTAGCGGTGGAATGCGCAGATATCAGGAGGAACACCGATGGCGAAGGCAGGTCACTGGGCCGCAACTGACGCTGAGGAGCGAAAGCATGGGG |
| P-asv3879 | GCGAGCGTTAATCGGAATTACTGGGCGTAAAGCGTACGTAGGCGGCCTATTAAGCAAGATGTGAAAGCCCCGGGCTCAACCTGGGAATAGCATTTTGAACTGGTAGGCTCGAGTTTTGAAGAGGGTGGTAGAATTTCCAGTGTAGCGGTGAAATGCGTAGATATTGGAAGGAATACCAGTGGCGAAGGCGGCCACCTGGTCAAAAACTGACGCTGAGGTACGAAAGCGTGGGG |
| P-asv3880 | GCAAGCGTTGTCCGGAATTATTGGGCGTAAAGAGCTCGTAGGCGGTCTGCCACGTCCGCTGTGAAAACCTAGGGCTTAACCCTGGGCTTGCAGTAGATACGGGCAGACTAGAGGTAGGTAGGGGAGAATGGAATTCCCGGTGTAGCGGTGAAATGCGCAGATATCGGGAGGAACACCGGTGGCGAAGGCGGTTCTCTGGGCCTTACCTGACGCTGAGGAGCGAAAGCGTGGGG |
| P-asv3881 | CCGAGCGTTATCCGGAATTATTGGGTTTAAAGGGTGCGTAGGCGGCATGTTAAGTCAGGGGTGAAAGACGATGGCTCAACCATCGCAGTGCCTTTGATACTGGCATGCTTGAATGTACATGAGGTAGGCGGAATGAGACAAGTAGCGGTGAAATGCATAGATATGTCTCAGAACACCGATTGCGAAGGCAGCTTACTAAAGTATGATTGACGCTGAGGCACGAAAGCGTGGGG |
| P-asv3882 | GCAAGCGTTGTCCGGAATTATTGGGCGTAAAGAGCTCGTAGGCGGTTTGTCGCGTCTGCTGTGAAAACGCGAGGCTCAACCTCGCGCCCGCAGTGGGTACGGGCAAACTAGAGTGCGGTAGGGGAGATTGGAATTCCTGGTGTAGCGGTGGAATGCGCAGATATCAGGAGGAACACCGATGGCGAAGGCAGATCTCTGGGCCGTAACCAACGCTGAGGAGCGAGAGCATGGGG |
| P-asv3883 | GCGAGCGTTAATCGGAATTACTGGGCGTAAAGCGCGCGTAGGTGGTTAATTAAGTCAGATGTGAAAGCCCAGGGCTCAACCTTGGAACTGCATTTGAAACTGGTTAACTAGAGTTTTGTAGAGGGTGGTAGAATTTCAGGTGTAGCGGTGAAATGCGTAGAGATCTGAAGGAATACCAGTGGCGAAGGCGGCCACCTGGACAAAGACTGACACTGAGGCGCGAAGGCGTGGGG |
| P-asv3884 | GCAAGCGTTAATCGGAATTACTGGGCGTAAAGCGCGCGTAGGTGGTTTGTTAAGTTGAATGTGAAATCCCCGGGCTCAACCTGGGAACTGCATCCAAAACTGGCAAGCTAGAGTATGGTAGAGGGTGGTGGAATTTCCTGTGTAGCGGTGAAATGCGTAGATATAGGAAGGAACACCAGTGGCGAAGGCGACCACCTGGACTGATACTGACACTGAGGTGCGAAAGCGTGGGG |
| P-asv3885 | TCGAGTGGTGGTCACTTTTATTGAGCCTAAAGCGTTCGTAGCCGGTCCTGTAAATCTTCAGATAAATTCTCCCGCTCAACGGGAGATCCTCTGAAGAGACTGTGGGACTAGGGACTGGGTGAGGTTGAATGTACTTCTGGGGTAGGGGTAAAATCCTGTAATCCTGGAAGGACAACCGGTGGCGAAGGCGTTCAACTAGAACAGATCCGACGGTGAGGAACGAGGGCTAGGGG |
| P-asv3886 | GCGAGCGTTAATCGGAATTACTGGGCGTAAAGCGTGCGCAGGCGGTTTTGTAAGACAGATGTGAAATCCCCGGGCTTAACCTGGGACCTGCATTTGTGACTGCAAAGCTAGAGTACGGCAGAGGGGGATGGAATTCCGCGTGTAGCAGTGAAATGCGTAGATATGCGGAGGAACACCGATGGCGAAGGCAATTCCCTGGGCCTGTACTGACGCTCATGCACGAAAGCGTGGGG |
| P-asv3887 | GCAAGCGTTGTCCGGAATCACTGGGCGTAAAGGGTGCGTAGGCGGTTCGGTAAGTTGCTGGTGAAATCCCGGGGCTCAACTCCGGGGCTGCCGGTACGACTGCCGGACTTGAGCACGGTAGGGGCAGATAGAATTCCCGGTGTAGCGGTGGAATGCGTAGATATCGGGAAGAATACCAGTGGCGAAGGCGTTCTGCTGGGCCGTTGCTGACGCTGAGGCACGACAGCGTGGGG |
| P-asv3888 | GCAAGCGTTGTTCGGAATCATTGGGCGTAAAGCGCGTGTAGGCGGCTTGGTAAGTCGGGTGTGAAATCCCGGGGCCCAACCCCGGAACTGCATTCGAAACTGCCTCGCTAGAGTCCCGGAGAGGAAGGCGGAATTCCCGGTGTAGAGGTGAAATTCGTAGATATCGGGAGGAACACCAGCGGCGAAGGCGGCCTTCTGGACGGTGACTGACGCTGAGACGCGAAAGCGTGGGG |
| P-asv3889 | GCGAGCGTTGTCCGGGTTTATTGGGCGTAAAGCGTGCGTAGGCGGCTGCGCAAGTCGGGTGTGAAACCCCGGGGCTCAACCCCGGAACTGCATCCGATACTGCGTGGCTAGAGGTAGGTAGGGGAGATCGGAATTCCTGGTGTAGCGGTGAAATGCGCAGATATCAGGAGGAACACCGGTGGCGAAGGCGGATCTCTGGGCCTTACCTGACGCTCAGGCACGAAAGCTAGGGG |
| P-asv3890 | TCAAGCGTTGTTCGGATTCACTGGGCGTAAAGGGTGCGTAGGTGGCAGGGTAAGTTTAATGTGAAATCTCCGGGCTTAACCCGGAAACTGCATTGAATACTATCCTGCTCGAGGGTTGGAGGGGGGACTGGAATTCTCGGTGTAGCAGTGAAATGCGTAGATATCGAGAGGAACACCAGTGGCGAAGGCGAGTCCCTGGACAACTCCTGACACTGAGGCACGAAAGCTAGGGG |
| P-asv3891 | GCAAACGTTGCTCGGAATCATTGGGCGTAAAGCGCACGTAGGCGGCTTCGCAAGTCGGATGTGAAATCCCTCGGCTCAACCGAGGACGTGCATCCGAAACTGCGGAGCTTGAGTACTGAAGAGGATCGCGGAATTCCCGGTGTAGAGGTGAAATTCGTAGATATCGGGAGGAACACCAGTGGCGAAGGCGGCGATCTGGGCAGTTACTGACGCTGAGGTGCGAAAGCGTGGGG |
| P-asv3892 | GCGAACGTTGTTCGGAATCACTGGGCTTAAAGGGCGCGTAGGCGGGCTGTTTAGTCGGGGGTGAAATCGTTCGGCCCAACCGGACAAGGGCCCCCGATACTGGCAGTCTCGAGGGAGGTAGGGGTTAGTGGAACTTCCGGTGGAGCGGTGAAATGCGTAGATATCGGAAGGAACGCCGGTGGCGAAAGCGGCTAACTGGACCTCTTCTGACGCTGAGGCGCGAAAGCTAGGGG |
| P-asv3893 | GCGAGCGTTATCCGGATTTACTGGGCGTAAAGCGCGTTAAGGCGGCCGACAAAGTTGGTGGTGAAATTTCAGGGCTCAACTCTGAAACTGCCTCCAATACTAGTCGGCTTGAGTGCGATAGAGGAAGGTGGAATTCCCGGTGTAGTAGTGAAATGCGTAGATATCGGGAGGAACACCAGTGGCGAAGGCGGCCTTCTGGCTCGCAACTGACGCTGTAGCGCGAAAGCGTGGGG |
| P-asv3894 | GCAAGCGTTGCTCGGAATCACTGGGCGTAAAGGGCGCGTAGGCGGGATGGCAAGTCGGTCGTGAAATCCCTGGGCTCAACTCAGGGGGGTCGGCCGATACTGCCATTCTTGAGGGATGCAGAGGAGACTGGAATTCCCGGTGTAGCGGTGAAATGTGTAGAGATCGGGAGGAACACCGGTGGCGAAGGCGGGTCTCTGGGCATTCCCTGACGCTGAGGCGCGAAAGCGTGGGG |
| P-asv3895 | GCTAGCGTTGTCCGGAATCATTGGGCGTAAAGAGCGTGTAGGCGGTTCGGTAAGTCCGCTGTAAAAGTCGGGGGCTCAACCCTCGAAAGCCGGTGGATACTGTCGGGCTAGAGTGCGGAAGAGGCGAGTGGAATTCCTGGTGTAGCGGTGAAATGCGCAGATATCAGGAGGAACACCAATTGCGAAGGCAGCTCGCTGGGACGTAACTGACGCTGAGACGCGAAAGCGTGGGG |
| P-asv3896 | GCAAGCGTTATCCAGATTTACTGGGCGTAAAGCGCGTGTAGGCGGCTGGTTAGGCGTGATGTGAAATCCTCGGGCTCAACCCGAAAAATGCATTGCAAACCGGCCTGGCTAGAGTGCAGGAGAGGGAAGCGGAATTCCAGGTGTAGCGGTGAAATGCGTAGATATCTGGAGGAACACCAGTGGCGAAGGCGGCTTCCTGGCCTGCAACTGACGCTGAGACGCGAAAGCGTGGGG |
| P-asv3897 | GCAAGCGTTGTCCGGATTTATTGGGCGTAAAGGGCCCGTAGGCGGCTTGATAAGTCGGGTGTGAAAACTCCAGGCTCAACCTGGAGACGCCACCCGATACTGTCATGGCTAGAGTCTGGTAGGGGAGTGCGGAATTCCTGGTGTAGCGGTGAAATGCGCAGATATCAGGAGGAACACCGGTGGCGAAGGCGGCACTCTGGGCCAGTACTGACGCTGAGGGGCGAAAGCGTGGGT |
| P-asv3898 | CCAAGCGTTGTTCGGATTCACTGGGCGTAAAGGGTGCGTAGGCGGTCGGGTAAGTCTGATGTGAAATCTCTGGGCTCAACCCAGAAATGGCATTGGATACTATTCGGCTTGAGGTCTGGAGGGGGGACTGGAATTCTCGGTGTAGCAGTGAAATGCGTAGATATCGAGAGGAACACCAGTGGCGAAGGCGAGTCCCTGGACAGGTCCTGACGCTGAGGCACGAAAGCTAGGGG |
| P-asv3899 | GCTAGCGTTGCTCGGAATTACTGGGCGTAAAGCGCACGTAGGCGGCTTCTTAAGTCGGTGGTGAAATCCTGGAGCTCAACTCCAGAACTGCCTTCGATACTGGGAAGCTCGAGTCCGGGAGAGGTGAGTGGAACTGCGAGTGTAGAGGTGAAATCCGTAGATATTCGCAAGAACACCAGTGGCGAAGGCGGCGATCTGGCCCGGTACTGACGCTGAGGTGCGAAAGCGTGGGG |
| P-asv3900 | GCGAGCGTTGTTCGGAATTACTGGGCGTAAAGGGCGCGTAGGCGGCCATCTTTGTCAGGCGTGAAAGCCCTGGGCTCAACCCAGGAATTGCGCTTGGGACGGGACGGCTTGAGTTCGGGAGAGGAGAGTGGAATTCCCAGTGTAGAGGTGAAATTCGTAGATATTGGGAAGAACACCGGTGGCGAAGGCGGCTCTCTGGACCGTAACTGACGCTGAGGCGCGAAAGCGTGGGG |
| P-asv3901 | GCGAACGTTGTTCGGAATCACTGGGCTTAAAGGGCGCGTAGGCGGGTGTCCAAGTCAGGGGTGAAATCCTCCAGCTTAACTGGAGAACTGCCTGTGATACTGGACGTCTGGAGTGAGGTAGGGGCATGTGGAACTTCCGGTGGAGCGGTGAAATGCGTAGATATCGGAAGGAACACCGGTGGCGAAGGCGGCATGCTGGACCTCAACTGACGCTGAGGCGCGAAAGCCAGGGG |
| P-asv3902 | GCGAGCGTTATTCGGAATTATTGGGCGTAAAGGGCGCGTAGGCGGCTCTGTAAGTCAAAGGTGAAATCCCTCGGCTCAACTGAGGAACTGCCTTTGAAACTGCAAGGCTTGAGGCCGGGAGGGGGTAGCGGAATTCCCAATGTAGCGGTGAAATGCGTAGATATTGGGAGGAACACCAGTGGCGAAGGCGGCTACCTGGACCGGTTCTGACGCTGAGGCGCGAAAGCGTGGGT |
| P-asv3903 | GCAAGCGTTGTCCGGAATTATTGGGCGTAAAGAGCTCGTAGGCGGTCTGTCACGTCGGCTGTGAAAACCCGGGGCTCAACCCCGGGCCTGCAGTCGATACGGGCAGACTAGAGTGCGGTAGGGGAGACTGGAATTCCTGGTGTAGCGGTGAAATGCGCAGATACCAGGAGGAACACCGGTGGCGAAGGCGGGTCTCTGGGCCGTAACTGACGCTGAGGAGCGAAAGCGTGGGG |
| P-asv3904 | GCAAGCGTTAATCGGAATTACTGGGCGTAAAGCGCACGCAGGCGGTCTGTCAAGTCGGATGTGAAATCCCCGGGCTCAACCTGGGAACTGCATTCGAGACTGGCAGGCTTGAGTCTTGTAGAGGGGGGTAGAATTCCAGGTGTAGCGGTGAAATGCGTAGAGATCTGGAGGGATACCGGTGGCGAAGGCGGCCCCCTGGACAAAGACTGACGCTCAGGTGCGAAAGCGTGGGG |

| | |
|---|---|
| P-asv3905 | GCAAACGTTATTCGGAATCACTGGGCTTAAAGGGCGCGTAGGCGGGCAACCAAGTCCGAGGTGAAATCCTCCAGCTTAACTGGAGAAGTGCCCCGGATACTGGTGGCCTCGAGGAGGGTAGGGGCATCTGGAACTAGCGGTGGAGCGGTGAAATGCGTTGATATCGCTAGGAACTCCGGTGGCGAAGGCGAGGTGCTGGACCCTTTCTGACGCTGAGGCGCGAAAGCCAGGGG |
| P-asv3906 | GCGAGCGTTATCCGGATTTATTGGGTTTAAAGGGTGCGCAGGCGGTCTGTTAAGTCAGCGGTCAAAGCCCGGGGCTCAACCCCGGCCCGCCGTTGAAACTGGCAGTCTCGAGTTGGAGAGAAGTATGCGGAATGCGCGGTGTAGCGGTGAAATGCATAGATATCGCGCAGAACTCCGATTGCGAAGGCAGCATGCCGGCTCCACACTGACGCTGAGGCACGAAAGCGTGGGT |
| P-asv3907 | CCAAGCGTTGTTCGGATTCATTGGGCGTAAAGGGTGCGTAGGCGGCGCGGTAAGTCGGGTGTGAAATTTCGGGGCTTAACTCCGAAACTGCATTCGATACTGCCGTGCTTGAGGACTGGAGAGGAGACTGGAATTTACGGTGTAGCGGTGAAATGCGTAGATATCGTAAGGAAGACCAGTGGCGAAGGCGGGTCTCTGGACAGTTCCTGACGCTGAGGCACGAAGGTCAGGGG |
| P-asv3908 | ACGAGCGTTGTCCGGAGTTACTGGGCGTAAAGGGCGCGCAGGCGGCCGCCCGCGTGTGTGGTGAAAGCTCCTGGCTTAACTAGGAGAGGTCCATGCAGACGGGGCGGCTTGAGCGTCGGAGAGGGGCATGGAATTCCGGGTGGAGCGGTGAAATGCGTAGAGATCCGGAGGAACACCAATGGCGAAGGCAGTGCCCTGGCCGACGGCTGACGCTCAGGCGCGAAAGCGTGGGG |
| P-asv3909 | GCAAGCGTTATTCGGAATTACTAGGCGTAAAGCGAGTGTAGGCGGACGCTTAAGTCCGACGTTAAATCTCCAGGCTTAACCTGGAGTCGTCGTCGGATACTGGGCGCCTCGAGTGAGGTAGGGGGCAGCGGAATTCCCGGTGTAGCGGTGGAATGCGTAGATATCGGGAGGAACACCTATGGCGAAAGCAGCTGCCTGGGCCTTTACTGACGCTAAGACTCGAAAGCCGGGGG |
| P-asv3910 | GCAAGCGTTAATCGGAATTACTGGGCGTAAAGCGTGCGTAGACGGTTAGACAAGTCGGGTGTGAAAGCCCCGGGCTCAACCTGGGAATTGCATTCGAGACAGTTTAGCTAGGGTACGGAAGAGGGAAGCGGAATTTCAGGTGTAGCGGTGAAATGCGTAGATATCTGAAGGAACACCAGTGGCGAAAGCGGCTTCCTGGTCCAGTACCGACGTTCAGGCACGAAAGCGTGGGG |
| P-asv3911 | GCAAGCGTTGTTCGGAATTACTGGGCGTAAAGGGCTCGTAGGTGGCCAACTAAGTCAGTCGTGAAATCCCTCGGCTTAACCGGGGAACTGCGTCTGATACTGGATGGCTTGAGTTCGGGAGAGGGATGCGGAATTCCAGGTGTAGCGGTGAAATGCGTAGATATCTGGAGGAACACCGGTGGCGAAGGCGGCATCCTGGACCGACACTGACACTGAGGAGCGAAAGCCAGGGG |
| P-asv3912 | GCCAGCGTTGTTCGGAATTACTGGGTGTAAAGGGTTCGTAGGCGGTGTGGCAAGTCGGGAGTGAAATCTCTGGGCTCAACTCAGAGACTGCTTCCGAAACTGCTGTGCTTGAGTGTGGGAGAGGCGCGTGGAATTGCAGGTGTAGCGGTGAAATGCGTAGATATCTGCAGGAACACCCGTGGCGAAAGCGGCGCGCTGGACCACTACTGACGCTGAGGAACGAAAGCTAGGGG |
| P-asv3913 | ACTAGCGTTGCTCGGAATTACTGGGCGTAAAGCGTGCGTAGGCGGTCCCGCAAGTCCGCTGTGAAAGCCCCGGGCTCAACCTGGGAATGGCAGTGGAAACTATGGGACTTGAATCTGGCAGAGGGGGGTGGAATGCCACGTGTAGCAGTGAAATGCGTAGAGATGTGGCGGAACACCGATGGCGAAGGCAGCCCCCTGGGCTAAGATTGACGCTCAGGCACGAAAGCGTGGGG |
| P-asv3914 | GCAAGCGTTAATCGGAATTACTGGGCGTAAAGCGCACGCAGGTGGCTAAGTCAGTCGATTGTGAAAGCCCTGGGCTTAACCTGGGAATTGCAGTCGATACTACTTAGCTCGAGTATAGGAGAGGGCAGTGGAATTCTCGGTGTAGCGGTGAAATGCGTAGATATCGAGAGGAACATCATTGGCGAAGGCGGCTGCCTGGCCCAATACTGACACTCAGGTGCGACAGCGTGGGG |
| P-asv3915 | GCGAGCGTTGTCCGGAATCACTGGGCGTAAAGGGCGCGTAGGTGGCGCGACAAGTCAGTGGTGAAAGCTCGACGCTCAACGTCGAGTCGGCCACTGATACTGTTGGGCTTGAGCACTGTAGAGGCTAATGGAATTCCGGGTGTAGCGGTGGAATGCGTAGAGATCCGGAAGAACACCGGTGGCGAAGGCGGTTAGCTGGGCAGTTGCTGACACTGAGGCGCGACAGCGTGGGG |
| P-asv3916 | GCGAGCGTTAATCGGAATTACTGGGCGTAAAGCGTGCGCAGGCGGTTCGTTGTGTCGGTGGTGAAATCCCCGGGCTTAACCTGGGAATGGCAGTGGAAACTAGAGGACTTGAATCCGGCAGAGGGGGGTGGAATGCCACGTGTAGCAGTGAAATGCGTAGATATGTGGCGGAACACCGATGGCGAAGGCAGCCCCCTGGGCTGAGATTGACGCTCAGGCACGAAAGCGTGGGG |
| P-asv3917 | GCAAGCGTTGTCCGGAATTATTGGGCGTAAAGAGCTCGTAGGCGGTTTGTCACGTCAGGGGTGAAATCCCGGAGCTCAACTTCGGAACTGCCCTTGATACAGCACGTCTTGAGTCCGGAAGAGGTGGGTGGAATTCCTAGTGTAGAGGTGAAATTCGTAGATATTAGGAAGAACACCGGTGGCGAAGGCGGCCCACTGGTCCGGTACTGACGCTGAGGTGCGAAAGCGTGGGG |
| P-asv3918 | ACGAGCGTTGTCCGGAGTTACTGGGCGTAAAGGGTGCGCAGGCGGCCTGGCGCGTCGCGAGTGAAATCTCCTGGCTCAACTGGGAGGGTGGTCGCGAGACGGCGAGGCTTGAGGCTCGGAGAGGGGCGTGGAATTCCGGGTGGAGCGGTGAAATGCGTAGAGATCCGGAGGAACACCGACGGCGAAGGCAGCGCCCTGGACGAGACCTGACGCTCAGGCACGAAAGCGCGGGG |
| P-asv3919 | GCAAGCGTTGTCCGGATTTATTGGGCGTAAAGAGCGTGTAGGCGGCCAGGTAGGTCTGCTGTGAAAACTCGAGGCTTAACCTCGAGATGTCGGCGGAAACCATCTGGCTAGAGTCCGGAAGAGGAGAGTGGAATTCCTGGTGTAGCGGTGAAATGCGCAGATATCAGGAAGAACACCCGTGGCGAAGGCGGCTCTCTGGGACGGTACTGACGCTGAGACGCGAAAGCGTGGGG |
| P-asv3920 | GCGAGCGTTGTCCGGAATTATTGGGCGTAAAGAGCTCGTAGGCGGCCTGTTGCGTCCGCTGTGAAATCCCGGGGCTTAACCCCGGGCTTGCAGTGGATACGGGCAGGCTGGAGGCAGGCAGGGGAGAACGGAATTCCCGGTGTAGCGGTGAAATGCGCAGATATCGGGAGGAACACCGGTGGCGAAGGCGGTTCTCTGGGCCTGTACTGACGCTGAGGAGCGAAAGCGTGGGG |
| P-asv3921 | GCAAGCGTTGTCCGGAATCATTGGGCGTAAAGCGCGCGTAGGTGGCTCTATAAGTCCGTTGTGAAAGTCAAAGGCTCAACCTTTGAAAGCCGATGGATACTGTAGAGCTAGAGTCCGGAAGAGGCGAGTGGAATTCCTGGTGTAGCGGTGGAATGCGCAGATATCAGGAGGAACACCAATAGCGAAGGCAGCTCGCTGGGACGTGACTGACACTAAGGCGCGAAAGCGTGGGG |
| P-asv3922 | GCAAACGTTGCTCGGAATTACTGGGCGTAAAGCGCGTGTAGGCGGATCGGAAAGTCGGGCGTGAAAGCCCTAGGCTCAACCTAGGAACTGCACCCGAAACTCCCAGTCTTGAGTGGTGGAGAGGGAAGTGGAATTCCTGGTGTAGAGGTGAAATTCGTAGATATCAGGAGGAACACCAGTGGCGAAGGCGGCTTCCTGGACACCAACTGACGCTGAGACGCGAAAGCGTGGGG |
| P-asv3923 | GCAAGCGTTGTTCGGAATCATTGGGCGTAAAGGGCGTGTAGGCGGTTTGTTAAGTCATGTGTGAAATCCCCCGGCTCAACCGAGGAACGACGCATGAAACTGGCAAGCTAGAGTACCGGAGAGGGGGGTGGAATTCCCGGTGTAGCGGTGAAATGCGTAGATATCGGGAGGAACACCTGTGGCGAAGGCGGCCCCCTGGTCGGATACTGACGCTGAGACGCGAAAGCGTGGGG |
| P-asv3924 | GCAAGCGTTGTTCGGAATCACTGGGCGTAAAGGGCGTGTAGGCGGTCAGGTAAGTGAGATGTGAAATGCCCCGGCTCACCCGGGGACCGGCATCTCATACTGCTTGGCTTGAGTACGGGAGAGGATGTGGAATTCCTGGTGTAGCGGTGAAATGCGTAGATATCAGGAGGAACACCGGTGGCGAAGGCGCACATCTGGCCCGATACTGACGCTGAGGCGCGAAAGCTAGGGG |
| P-asv3925 | GCGAGCGTTGTTCGGATTTACTGGGCGTAAAGGGCGCGTAGGCGGCCCTCTTTGTCGCGTGTGAAAGCCCCGGGCTCAACCTGGGAATTGCGCACGAAACGGGGGGGCTGGAGTTCGGGAGAGGAGCGTGGAATTCCCAGTGTAGAGGTGATATTCGTAGATATTGGGAAGAACACCGGTGGCGAAGGCGGCGCTCTGGACCGAGACTGACGCTGAGGCGCGAAAGCGTGGGG |
| P-asv3926 | GCAAGCGTTAATCGGAATTACTGGGCGTAAAGCGTGCGCAGGCGGTTTTGTAAGTCTGTCGTGAAATCCCCGGGCTCAACCTGGGAATGGCGATGGAGACTGCAAGGCTAGAATCTGGCAGAGGGGGGTAGAATTCCACGTGTAGCAGTGAAATGCGTAGAGATGTGGAGGAACACCGATGGCGAAGGCAGCCCCCTGGGTCAAGATTGACGCTCATGCACGAAAGCGTGGGG |
| P-asv3927 | GCAAACGTTGTTCGGAATTACTGGGCATAAAGAGCACGTAGGTGGTCCGCTAAGTCACGTGTGAAATCCCCCGGCTCAACCGGGGAATGGCGCGTGATACTGGCGGACTCGAGGCAGGTAGGGGTGAGCGGAACTCCAGGTGGAGCGGTGAAATGCGTAGATATCTGGAGGAACACCGGTGGCGAAAGCGGCTCACTGGACCTGATCTGACACTGAGGTGCGAAAGCTAGGGT |
| P-asv3928 | CCAAGCGTTATCCGGAATTATTGGGTTTAAAGGGTCCGCAGGCGGTCTATTTAGTCAGAGGTGAAATCCTACAGCTCAACTGTAGAATGGCCTTTGATACTGGTAGACTTGAGTCATTGTGAAGTGGTTAGAATGTGTGGTGTAGCGGTGAAATGCTTAGATATCACACAGAATACCCATTGCGAAGGCAGATCACTAACAATGTACTGACGCTCATGGACGAAAGCGTGGGG |
| P-asv3929 | GCAAGCGTTGTTCGGAATTATTGGGCGTAAAGCGCGTGTAGGCGGCTCTGCAAGTTGGGTGTGAAAGCCCTCGGCTCAACCGAGGAAGTGCGCCCAAAACTACAGAGCTTGAGTACCGGAGAGGATGGCGGAATACCCGGTGTAGAGGTGAAATTCGTAGATATCGGGTGGAACACCAGTGGCGAAGGCGGCCATCTGGACGGATACTGACGCTGAGACGCGAAAGCGTGGGG |
| P-asv3930 | GCAAGCGTTGTCCGGATTTATTGGGCGTAAAGAGCTCGTAGGCGGTTCGGTAAGTCGGATGTGAAATCTCCACGCTCAACGTGGAGGCGCCATTCGAAACTGCCATGACTTGAGTCCGGTAGGGGAGTGTGGAATTCCCGGTGTAGCGGTGAAATGCGCAGAGATCGGGAGGAACACCCGTAGCGAAGGCGGCACTCTGGGCCGGTACTGACGCTGAGGAGCGAAAGCGTGGGG |
| P-asv3931 | GCAAGCGTTGTTCGGAATTACTGGGCGTAAAGGGCGCGTAGGCGGCTTGGTCAGTCCCGTGTGAAATCCCTCGGCTCAACTGAGGAACTGCACGGGAAACTGCCTGGCTTGAGTTCGGGAGAGGGAAGCGGAATTCCGGGTGTAGCGGTGAAATGCGTAGATATCCGGAGGAACACCGGTGGCGAAGGCGGCTTCCTGGACCGACACTGACGCTGAGGCGCGAAAGCTAGGGG |
| P-asv3932 | CCGGGCGTTATCCGGATTTATTGGGTTTAAAGGGAGCGTAGGCCGCGCCTTAAGCGTGTTGTGAAATCCGGGTGCTCAACATCCGGCTTGCAGCGCGAACTGGGGCGCTTGAGTGCGCTGAAAGTAGGCGGAATTCGTGGTGTAGCGGTGAAATGCTTAGATATCACGAGGAACTCCGATTGCGAAGGCAGCCTACTGTAGCGCTACTGACGCTGATGCTCGAAAGCGTGGGT |
| P-asv3933 | GCGAGCGTTGTCCGGAATTATTGGGCGTAAAGAGCTTGTAGGCGGTTGGTCGCGTCTGCTGTGAAAGGCCGGGGCTTAACTCCGGTTTTGCAGTGGGTACGGGCTAACTAGAGTGCAGTAGGGGAGACTGGAATTCCTGGTGTAGCGGTGGAATGCACAGATATCAGGAGGAACACCGATGGCGAAGGCAGGTCTCTGGGCTGTAACTGACGCTGAGAAGCGAAAGCATGGGG |
| P-asv3934 | GCGAGCGTTGTCCGGATTTATTGGGCGTAAAGGGCTCGTAGGTGGTTGATCGCGTCGGAAGTGTAATCTTGGGGCTTAACCCTGAGCGTGCTTTCGATACGTGTTGACTTGAGGAAGGTAGGGGAGAATGGAATTCCTGGTGGAGCGGTGGAATGCGCAGATATCAGGAGGAACACCAGTGGCTAAGGCGGTTCTCTGGGCCTTTCCTGACGCTGAGGAGCGAAAGCGTGGGG |
| P-asv3935 | GCGAGCGTTAATCGGAATTACTGGGCGTAAAGCGCGCGTAGGCGGTCTGATAAGCCGGTTGTGAAAGCCCCGGGCTCAACCTGGGAACGGCATCCGGAACTGTCAGGCTAGAGTGCAGGAGAGGAAGGTAGAATTCCCGGTGTAGCGGTGAAATGCGTAGAGATCGGGAGGAGTACCAGTGGCGAAGGCGGCCTTCTGGACTGACACTGACGCTGAGGTGCGAAAGCGTGGGT |
| P-asv3936 | GCTAGCGTTGCTCGGAATCACTGGGCGTAAAGGGTGCGTAGGCGGGTCTTTAAGTCAGGGGTGAAATCCTGGAGCTCAACTCCAGAACTGCCTTTGATACTGAGGATCTTGAGTTCGGGAGAGGTGAGTGGAACTGCGAGTGTAGAGGTGAAATTCGTAGATATTCGCAAGAACACGAGTGGCGAAGGCGGCTCACTGGCCAGATACTGACGCTGAGGCACGAAAGCGTGGGG |
| P-asv3937 | GCGAACGTTAATCGGAATTACTGGGCGTAAAGCGCGTGTAGGCGGTTCGTTAAGTGGGATGTGAAAGCCCTGGGCTCAACCCAGGAATTGCATTCCAGACTGGCGGACTCGAGTGCAGTAGAGGGAGGTGGAATTCCAGGTGTAGCGGTGAAATGCGTAGGTATCTGGAGGAACATCGGTGGCGAAGGCGGCCTCCTGGACTGACACTGACGCTGAGACGCGAAAGCGTGGGG |
| P-asv3938 | GCAAGCGTTGTCCGGATTTATTGGGCGTAAAGCGAGCGCAGGCGGTCTTTTAAGTCTGATGTGAAAGCCCTCGGCTCAACCGAGGAAGGTCATTGGAAACTGGAAGACTTGAGTGCAGAAGAGGAGAGTGGAATTCCATGTGTAGCGGTGAAATGCGTAGATATATGGAGGAACACCAGTGGCGAAGGCGACTCTCTGGTCTGTAACTGACGCTGAGGCTCGAAAGCGTGGGG |
| P-asv3939 | GCTAGCGTTGCTCGGAATGACTGGGCGTAAAGGGCGCGTAGGCGGATGGCACAGTCAGATGTGAAATTCCCGGGCTTAACCTGGGGGCTGCATTTGATACGTGTTGTCTAGAGTGAGGAAGAGGGTTGTAGAATTCCCAGTGTAGAGGTGAAATTCGTAGATATTGGGAAGAACACCGGTGGCGAAGGCGGCAACCTGGTCCTTTACTGACGCTGAGGCGCGAAAGCGTGGGG |
| P-asv3940 | GCAAGCGTTAATCGGAATTACTGGGCGTAAAGAGTGCGCAGGCGGTTTTGCAAGACCGATGTGAAATCCCCGGGCTTAACCTGGGAACTGCATTGGTGACTGCAAGGCTAGAGTGTGTCAGAGGGAGGTGGAATTCCGCATGTAGCAGTGAAATGCGTAGATATGCGGAGGAACACCGATGGCGAAGGCAGCCTCCTGGGATAACACTGACGCTCATGCACGAAAGCGTGGGG |
| P-asv3941 | GCAAGCGTTGTCCGGAATTATTGGGCGTAAAGAGCTCGTAGGCGGCTTGTCGCGTCTGCTGTGAAAACCCGAGGCTCAACCTCGGGCCTGCAGTGGGTACGGGCAAGCTAGAGTGCGGTAGGGGAGATTGGAATTCCTGGTGTAGCGGTGGAATGCGCAGATATCAGGAGGAACACCGATGGCGAAGGCAGATCTCTGGGCCATAACTGACGCTGAGGAGCGAAAGCATAGGG |
| P-asv3942 | GCAAGCGTTAATCGGAATTACTGGGCGTAAAGCGCACGCAGGCGGTTGGATAAGTTAGATGTGAAAGCCCCGGGCTCAACCTGGGAATTGCATTTAAAACTGTCCAGCTAGAGTCTTGTAGAGGGGGGTAGAATTCCAGGTGTAGCAGTGAAATGCATAGAGATCTGGAGGAATACCGGTGGCGAAGGCGGCCCCCTGGACAAAGACTGACGCTCAGGTGCGAAAGCGTGGGG |
| P-asv3943 | GCAAGCGTTGTCCGGAATTATTGGGCGTAAAGCGCGCGCAGGCGGTCATTTAAGTCTGGTGTTTAAACCTTGGGCTCAACCTAAGGTCGCACTGGAAACTGGGTGACTTGAGTACAGAAGAGGAAAGTGGAATTCCACGTGTAGCGGTGAAATGCGTAGATATGTGGAGGAACACCAGTGGCGAAGGCGACTTTCTGGGCTGTAACTGACGCTGAGGCGCGAAAGCGTGGTG |

| | |
|---|---|
| P-asv3944 | GCAAGCGTTAATCGGAATTACTGGGCGTAAAGCGCGCGTAGGTGGTTCGTTAAGTTGGATGTGAAATCCCCGGGCTCAACCTGGGAACTGCATCCAAAACTGGCGAGCTAGAGTAGGGCAGAGGGTGGTGGAATTTCCTGTGTAGCGGTGAAATGCGTAGATATAGGAAAGAACACCAGTGGCGAAGGCGACCACCTGGACTCATACTGACACTGAGGTGCGAAAGCGTGGGG |
| P-asv3945 | CCAAGCGTTAATCGGAATTACTGGGCGTAAAGCGTGCGCAGGCGGTCTTGTAAGACAGGTGTGAAATCCCCGGGCTTAACCTGGGAACTGCACTTGTGACTGGAAGGCTGGAGTACGGCAGAGGGGGATGGAATTCCGCGTGTAGCAGTGAAATGCGTAGATATGCGGAGGAACACCGATGGCGAAGGCAATCCCCTGGGCCTGTACTGACGCTCATGCACGAAAGCGTGGGG |
| P-asv3946 | GCAAGCGTTGTCCGGAATTATTGGGCGTAAAGAGCTCGTAGGCGGTTTGTCGCGTCTGCTGTGAAAACGCAAGGCTTAACCTTGCGCCTGCAGTGGGTACGGGCAGACTAGAGTGCGGTAGGGGAGACTGGAATTCCTGGTGTAGCGGTGGAATGCGCAGATAACAGGAGGAACGCCGATGGCGAAGGCAGGTCTCTGGGCCGTTACTGACGCTGAGGAGCGAAAGCATGGGG |
| P-asv3947 | GCAAGCGTTACTCGGAATCACTGGGCGTAAAGAGCGTGTAGGCGGATTAATAAGTCAGATGTGAAAGCCAATGGCTCAACCATTGAACTGCATTTGAAACTGTTAATCTAGAATATGGGAGAGGTAGATGGAATTTCTGGTGTAGGGGTAAAATCCGTAGAGATCAGAAGGAATACCGATTGCGAAGGCGATCTACTGGAACATTATTGACGCTGAGACGCGAAAGCGTGGGG |
| P-asv3948 | TCAAGTGGTCAGGATGATTATTGGGCCTAAAGCATCCGTAGCTCGTTTTGTAAGTTTTCGGTTAAATCCATGCGCTCAACGTATGGTCTGCCGGGAATACTGCATAACTAGGAAGTGGGAGAGGTAGACGGTACTCGGTAGGAAGGGGTAAAATCCTTTGATCTATTGATGACCACCTGTGGCGAAGGCGGTCTACCAGAACACGTTCGACGGTGAGGGATGAAAGCTGGGGG |
| P-asv3949 | GTTAGCGTTGTTCGGAATTACTGGGCGTAAAGCGCACGTAGGCGGACCATCAAGTTGGGGGTGAAATCCCGGGGCTCAACCCCGGAACTGCCTTCAAAACTGGTGGTCTGGAGTTCGAGAGAGGTGAGTGGAATTCCGAGTGTAGAGGTGAAATTCGTAGATATTCGGAGGAACATCAGTGGCGAAGGCGGCTCACTGGCTCGATACTGACGCTGAGGTGCGAAAGTGTGGGG |
| P-asv3950 | GCGAGCGTTATTCGGAATTACTAGGCGTAAAGCGAGTGTAGGCGGACTCTTAAGTCTGACGTTAAATCTCCTGGCTTAACTAGGAGCTGTCGTCGGATACTGAGAGCCTCGAATGGGGTAGGGGGCAGCGGAATTCCCGGTGTAGCGGTGAAATGCGTAGATATCGGGAGGAACACCTATGGCGAAAGCAGCTGCCTGGGCCTCTATTGACGCTAAGACTCGAAAGCTGGGGG |
| P-asv3951 | CCGAGCGTTATCCGGATTTATTGGGCGTAAAGCGAGCGCAGGCGGTTAGATAAGTCTGAAGTTAAAGGCTGTGGCTTAACCATAGTACGCTTTGGAAACTGTTTAACTTGAGTGCAAGAGGGGAGAGTGGAATTCCATGTGTAGCGGTGAAATGCGTAGATATATGGAGGAACACCGGTGGCGAAAGCGGCTCTCTGGCTTGTAACTGACGCTGAGGCTCGAAAGCGTGGGG |
| P-asv3952 | GCAAGTGTTATCCGGAATCACTGGGCGTAAAGCGTCTGTAGGTGGTTTGATAAGTCAACTGTTAAATCTTGAAGCTCAACTTCAAAATCGCAGTCGAAACTATTAGACTTGAGTATAGTAGGGGTAAAGGGAATTTCCAGTGGAGCGGTGAAATGCGTAGAGATTGGAAAGAACACCGATGGCGAAGGCACTTTACTGGGCTATTACTGACACTCAGAGACGAAAGCTAGGGT |
| P-asv3953 | GCGAACGTTATTCGGAATCACTGGGCTTAAAGGGTGCGTAGGCGGTTATTTAAGTAGGATGTGAAATCTCCCGGCTCAACCGGGGAACTGCGTTCTAAACTGGATAACTTGAGTGAGATAGAGGTGTGCGGAACTTCCGGTGGAGCGGTGAAATGTGTTGATATCGGAGGGAACACCGTTGGCGAAAGCGGCACACTGGGTCTTAACTGACGCTGAGGCACGAAAGCCAGGGG |
| P-asv3954 | TCGAGTGATGGCCGATATTATTGGGCCTAAAGCGTTCGTAGCTGGCCAGTCAAGTCTGTTGGGAAATCCACGTGCCCAACACGTGGGCGTCCGACAGAAACTGACTGGCTTGGGACCGGAAGACCCGAAGAGTACGTCTGGGGTAGGAGTGAAATCCCGTAATCCTAGACGGACTACCGATGGCGAAAGCACTTCGGGAGTACGGATCCGACAGTGAGGAACGAAAGCTGGGGT |
| P-asv3955 | GCTAGCGTTGTTCGGAATTACTGGGCGTAAAGCGCACGTAGGCGGACATTTAAGTCAGGGGTGAAATCCCAGAGCTCAACTCTGGAACTGCCTTTGATACTGGGTGTCTAGAGTATGGAAGAGGTGAGTGGAATTCCGAGTGTAGAGGTGAAATTCGTAGATATTCGGAGGAACACCAGTGGCGAAGGCGGCTCACTGGTCCATTACTGACGCTGAGGTGCGAAAGCGTGGGG |
| P-asv3956 | CCAAGCGTTATCCGGAATCACTGGGTGTAAAGGGTACGCAGGCGGACATGTAGGTCCAATGTGAAAGTCCTGGGCTCAACCCAGGAATTGCATTGGAAACCCCATGTCTTGAGGATCAGAGAGGCAGGCGCAATTCGTGGTGTAGCGGTGAAATGCGTAGATATCACGAGGAACACCAGTGGTGTAGACGGCCTGCTGGATGAATCCTGACGCTGAGGTACGAAAGCGTGGGG |
| P-asv3957 | GCAAGCGTTAATCGGAATTACTGGGCGTAAAGTGCGCGTAGGCGGTCGGGTAAGTCGGATGTGAAAGCCCTGGGCTTAACCCAGGAATTGCATTCGATACTGCCTGACTCGAGTGCAGTAGAGGAGAGTGGAATTCCCGGTGTAGCGGTGAAATGCGTAGATATCGGGAGGAACACCAGTGGCGAAGGCGACTCTCTGGACTGACACTGACGCTGACAGCGCGAAAGCGTGGGG |
| P-asv3958 | GCAAGCGTTATCCGGAATCATTGGGTTTAAAGGGTCCGCAGGCGGACATATAAGTCAGTGGTGAAATCCCATCGCTTAACGATGGAACTGCCATTGATACTGTATGTCTTGAATTCGGTCGAAGTGGGCGGAATATGACATGTAGCGGTGAAATGCTTAGATATGTCATAGAACACCGATAGCGAAGGCAGCTCACTAGGCCTGGATTGACGCTCAGGGACGAAAGCGTGGGT |
| P-asv3959 | CCAAGTGATGGCCGATCTTATTGGGCCTAAAGCGTCCGTAGCCGGCCGGACAAGTCCGTTGGGAAATCGACGCGCTCAACGCGTCGGCGTCCAGCGGAAACTGTCCGGCTTGGGGCCGGAAGACCCGAGGGGTACGTCCGGGGTAGGAGTGAAATCCTGTAATCCTGGACGGACCACCGGTGGCGAAAGCGCCTCGGGAAGACGGACCCGACGGTGAGGGACGAAAGCTAGGGT |
| P-asv3960 | GCAAGCGTTGTTCGGATTTACTGGGCGTAAAGAGTGCGTAGGCGGTTTGGTAAGTTGGATGTGAAAGCCCCGGGCTTAACCTGGGAACTGCATTCAAAACTACCAGGCTCGAGTGTGGAAGAGGATGGCGGAATTCCCGGTGTAGAGGTGAAATTCGTAGATATCGGGAAGAACACCTGTGGCGAAGGCGGCCATCTGGTCCATTACTGACGCTGAGGCACGAAAGCGTGGGG |
| P-asv3961 | GCAAGCGTTATTCGGCATGACTGGGCGTAAAGGGTCCGTAGACGGTAGAATCAGTTCTTAGTTAAATATTAAAGCCTAACTTTAAAATAGCTTCGAATACTGTTCTACTTGAGTTTTATACAGAAGACTAGAATTCTATACGGAAAGGTAAAATTTGTAGATATATAGAGGAATATCAGTAGCGAAGGCGAGTCTTTGGTAAAAACTGACGTTGAGGGACGAAAGTATGGGG |
| P-asv3962 | GCAAGCGTTATCCGGAATCATTGGGTTTAAAGGGTCCGCAGGCGGATATATAAGTCAGTGGTGAAAGCCTACAGCTTAACTGTAGAACTGCCATTGAAACTGTATGTCTTGAATTCGATCGAAGTGGGCGGAATGTGTCATGTAGCGGTGAAATGCATAGATATGACACAGAACACCGATAGCGAAGGCAGCTCACTAGGTCTGAATTGACGCTCAGGGACGAAAGCGTGGGG |
| P-asv3963 | GCAAGCGTTAATCGGAATTACTGGGCGTAAAGCGCGCGTAGGTGGCTAAGTAAGATGGGTGTGAAATCCCCGGGCTCAACCTGGGAACTGCATCCATAACTGCTTGGCTAGAGTACGGTAGAGGGTAGTGGAATTTCCTGTGTAGCGGTGAAATGCGTAGATATAGGAAGGAACACCAGTGGCGAAGGCGACTACCTGGACTGATACTGACACTGAGGTGCGAAAGCGTGGGG |
| P-asv3964 | GCAAGCGTTGTCCGGAATTATTGGGAGTAAAGCGCGCGCAGGCGGTCTTTTAAGTCTGATGTGAAAACTTGCGGCTCAACCGCAAGCTGGCATTGGAAACTGGGAGACTTGAGTGTAGGAGAGGAAAGTGGAATTCCACGTGTAGCGGTGAAATGCGTAGATATGTGGAGGAACACCAGTGGCGAAGGCGACTTTCTGGCCTACAACTGACGCTGAGGCGCGAAAGCGTGGGG |
| P-asv3965 | CCAAGCGTTAATCGGAATTACTGGGCGTAAAGCGCGCGTAGGTGGTTAGTTAAGTTGGATGTGAAATCCCCGGGCTCAACCTGGGAACTGCTTTCAAAACTGCTGAGCTAGAGTACGGTAGAGGGTGGTGGAATTTCCTGTGTAGCGGTGAAATGCGTAGATATAGGAAGGAACACCAGTGGCGAAGGCGACCACCTGGACTGATACTGACACTGAGGTGCGAAAGCGTGGGG |
| P-asv3966 | CCAAGCGTTGTCCGGAATCACTGGGCGTAAAGGGTGTGCAGGTGGGGCAGCAAGTCAGGGGTGAAAGCCCGCGGCTTAACCGCGGAAGTGCCCTTGAAACTGCTGCTCTTGAGTCCCGGAGAGGCTGCCGGAATTCGTGGTGTAGCGGTGAAATGCGTAGATATCACGAGGAACACCAGAGGCGAAAGCGGGCAGCTGGACGGGTACTGACACTGAGGCACGAAAGCGTGGGG |
| P-asv3967 | ACGAGTGATGGCCGATATTATTGGGCCTAAAGCGTCCGTAGCCGGCCAGACAAGTCCGTTGGGAAATCGACGTGCCTAACACGTCGGCGTCCAGCGGAAACTGTCTGGCTTGGGGCCGGAAGATCCGAGGGGTACGTCCGGGGTAGGAGTGAAATCCCGTAATCCTGGACGGACCGCCGGTGGCGAAAGCGCCTCGGAAGGACGGACCCGACGGTCAGGGACGAAAGCTAGGGT |
| P-asv3968 | GCAAGCGTTGTCCGGATTTATTGGGTTTAAAGGGTGCGTAGGCGGCTTGTTAAGTCAGCGGTGAAAGTTTTCGGCTCAACCGGAAAAATGCCGTTGATACTGATGAGCTTGAGTCACGGAAGGGTACATGGAATTGGTGGTGTAGCGGTGAAATGCATAGATACCATCAGGAACACCGATAGCGAAGGCATTGTACTTGCCGCGAACTGACGCTGAGGCACGAAAGCGCGGGT |
| P-asv3969 | CCAAGCGTTGTTCGGATTTACTGGGCGTAAAGGGTGCGTAGGCGGTCGGATAAGTCTGATGTGAAATCTCCGAGCTTAACTCGGAAATGGCATTGGATACTATTCGGCTAGAGAGTTGGAGGGGGGACTGGAATACTTGGTGTAGCAGTGAAATGCGTAGATATCAAGTGGAACACCAGTGGCGAAGGCGAGTCCCTGGACAACTTCTGACGCTGAGGCACGAAAGCTAGGGG |
| P-asv3970 | GCGAGCGTTAATCGGAATTACTGGGCGTAAAGCGTGCGCAGGCGGTTCGCTAAGACAGGTGTGAAATCCCCGGGCTTAACCTGGGAACTGCGCTTGTGACTGGCGGGCTAGAGTACGGCAGAGGGGGGTGGAATTCCACGTGTAGCAGTGAAATGCGTAGATATGTGGAGGAACACCGATGGCGAAGGCAGCCCCCTGGGCCTGTACTGACGCTCATGCACGAAAGCGTGGGG |
| P-asv3971 | GCAAGCGTTAGTCGGAATTATTGGGCGTAAAGGGTGCGTAGGCGGAAAAGCAAGTCGGACGTGAAATTCTGGGGCTCAACCTCAGAGCTGCATCCGAAACTGCTTTTCTAGAGGATTGGCGGAGAAAACGGAATTCCATGTGTAGCGGTGAAATGCGTAGATATATGGAAGAACACCTGTGGCGAAAGCGGTTTTCTAGCCTCCTCCTGACGCTGAGGCACGAGAGCAAGGGG |
| P-asv3972 | CCGGGCGTTATCCGGATTTATTGGGTTTAAAGGGAGCGTAGGCTGTCTATTAAGCGTGTTGTGAAATATACCGGCTCAACCGGTGGCTTGCAGCGCGAACTGGTCGACTTGAGTATGCAGGAAGTAGGCGGAATTCATGGTGTAGCGGTGAAATGCTTAGATATCATGACGAACTCCGATTGCGCAGGCAGCTTACTGTAGCATAACTGACGCTGATGCTCGAAAGTGCGGGT |
| P-asv3973 | GCTAGCGTTGTTCGGAATTACTGGGCGTAAAGCGCATGTAGGCGGATATTTAAGTCAGGGGTGAAATCCCAGAGCTCAACTCTGGAACTGCCTTTGATACTGGGTATCTTGAGTATGGAAGAGGTAAGTGGAATTGCGAGTGTAGAGGTGAAATTCGTAGATATTCGCAGGAACACCAGTGGCGAAGGCGGCTTACTGGTCCATAACTGACGCTGAGGTGCGAAAGCGTGGGG |
| P-asv3974 | GCTAGCGTTGTTCGGAATTACTGGGCGTAAAGCGTACGTAGGCGGCTTTGTAAGTTAGAGGTGAAAGCCCGGGGCTCAACTCCGGAATTGCCTTTAAGACTGCATCGCTTGAATCATGGAGAGGTTAGTGGAATTCCGAGTGTAGAGGTGAAATTCGTAGATATTCGGAAGAACACCAGTGGCGAAGGCGACTAACTGGACATGTATTGACGCTGAGGTACGAAAGCGTGGGG |
| P-asv3975 | GCTAGCGTTGTTCGGAATTACTGGGCGTAAAGCGCACGTAGGCGGCCATTCAAGTCAGAGGTGAAAGCCCGGGGCTCAACCCCGGAACTGCCTTTGAAACTAGATGGCTTGAATCTTGGAGAGGCGAGTGGAATTCCGAGTGTAGAGGTGAAATTCGTAGATATTCGGAAGAACACCAGTGGCGAAGGCGACTCGCTGGGCAAGTATTGACGCTGAGGTGCGAAAGCATGGGG |
| P-asv3976 | TCCCTCTTTCCTCTTCCTTCCTGGGCGTAAAGCGCGCGTAGGTGGTTTGTTAAGTTGAATGTGAAAGCCCCGGGCTCAACCTGGGAACTGCATCCAAAACTGGCAAGCTAGAGTATGGCAGAGGGTGGTGGAATTTCCTGTGTAGCGGTGAAATGCGTAGATATAGGAAGGAACACCAGTGGCGAAGGCGACCACCTGGGCTAATACTGACACTGAGGTGCGAAAGCGTGGGG |
| P-asv3977 | GCGAGCGTTAATCGGAATTACTGGGCGTAAAGCGCGCGTAGGTGGCTTGATAAGCCGGTTGTGAAAGCCCCGGGCTCAACCTGGGAACGGCATCCGGAACTGTCAGGCTAGAGTGCAGGAGAGGAAGGTAGAATTCCCGGTGTAGCGGTGAAATGCGTAGAGATCGGGATCAATACCAGTGGCGAAGGCGGCCTTCTGGACTGACACTGACACTGAGGTGCGAAAGCGTGGGT |
| P-asv3978 | GCGAGCGTTAATCGGAATTACTGGGCGTAAAGCGCGCGTAGGTGGCTTGATGAGCCGGTTGTGAAGGCCCCGGGCTCAACCTGGGAACGGCATCCGGAACTGTCAGGCTAGAGTGCAGGAGAGGAAGGTAGAATTCCCGGTGTAGCGGTGAAATGCGTAGAGATCGGGAGGAATACCAGTGGCGAAGGCGGCCTTCTGGACTGACACTGACGCTGAGGTGCGAAAGCGTGGGT |
| P-asv3979 | GCGAGCGTTAATCGGAATTACTGGGCGTAGAGCGCGCGTAGGCGGCCTGATAAGCCGGTTGTGAAAGCCCCGGGCTCAACCTGGGAACGGCATCCGGAACTGTCAGGCTAGAGTGCAGGAGAGGAAGGTAGAATTCCCGGTGTAGCGGTGAAATGCGTAGAGATCGGGAGGAATACCAGTGGCGAAGGCGGCCTTCTGGACTGACACTGACGCTGAGGTGCGAAAGCGTGGGT |
| P-asv3980 | GCAAGCGTTGTCCGGAATTATTGGGCGTAAAGGGCTCGCAGGCGGTTTCTTAAGTCTGATGTGAAAGCCCCCGGCTCAACCGGGGAGGGTCATTGGAAGCTGGGAAACTTGAGTGCAGAAGAGGAGAGTGGAATTCCACGTGCAGCGGTGAAATGCGTAGAGATGTGGAGGAACACCAGTGGCGAAGGCGACTCTCTGGTCTGTAACTGACGCTGAGGAGCGAAAGCGTGGGG |
| P-asv3981 | GCAAGCGTTGCCCGGAATTATTGGGCGTAAAGAGCTCGTAGGCGGTTTGTCGCGTCTGCTGTGAAAACGCAAGGCTTAACCTTGCGCCTGCAGTGGGTACGGGCAGACTAGAGTGCGGTAGGGGAGACTGGAATTCCTGGTGTAGCGGTGGAATGCGCAGATATCAGGAGGAGCACCGATGGCGAAGGCAGGTCTCTGGGCCGTTTCTGACGCTGAGGAGCGAAAGCATGGGG |
| P-asv3982 | GCAAGCGTTACTCGGAATTACTGGGCGTAAAGGGTGCGTAGGTGGTCGTTTAAGTCCGTTGTGAAATCCCTGGGCTCAACCTGGGAATGGCAGTGGATACTGGACGGCTAGAATGCGGTAGAGGGTAGTGGAATTCCAGGTGTAGCAGTGAAATGCGTAGAGATCGGGAGGAACACCAGTGGCGAAGGCGGCTACCTGGACCAGCATTGACACTCAAGCACGAAAGCGTGGGG |

| | |
|---|---|
| P-asv3983 | CCGAGCGTTAGGCGGAATCACTGGGCTTAAAGCGTGTGTAGGCGGATGCCTAAGTACCTTGTGAAATCCCACGGCTCACCCGTGGAACTGCTTGGTATACTGGGTGTCTTGAGCCACTTAGGGGCTACCGGAACAAATGGTGGAGCGGTGAAATGCGTAGATATCATTTGGAACGCCAATGGTGAAACCAGGTAGCTGGGAGTGTGCTGACGCTGAGACACGAAAGCCAGGGG |
| P-asv3984 | ACAAGCGTTGTCCGGATTTACTGGGCGTAAAGAGCGCGCAGGCGGTCGTTCAAGTCGAGTGTGAAAGCCCCCGGCTCAACTGGGGAGGGTCATTCGATACTGATCGACTCGAAGGCAGGAGAGGGAAGTGGAATTCCCGGTGTAGTGGTGAAATGCGTAGATATCGGGAGGAACACCAGTGGCGAAGGCGACTTCCTGGCCTGTTCTTGACGCTGAGGCGCGAAAGCTAGGGG |
| P-asv3985 | GCGAGCGTTGTTCGGATTTAATGGGCGTAAAGGGCGCGTAGGCGGCCCTCCTTGTCGCGTGTGAAAGCCCCGGGCTCAACCTGGGAATTGCGCGCGAAACGGGGGGCTTGAGTTCGGGAGAGGAGCGTGGAATTCCCAGTGTAGAGGTGAAATTCGTAGATATTGGGAAGAACACCGGTGGCGAAGGCGGCGCTCTGGACCGAGACTGACGCTGAGGCGCGAAAGCGTGGGG |
| P-asv3986 | GCCAGCGTTGTTCGGAATTACTGGGTGTAAAGGGTTCGTAGGCGGTGCGGCAAGTTGGGAGTGAAATCTCTGGGCTTAACCCAGAGGCTGCTTCCAAAACTGCTGTGCTCGAGTGTGAGAGAGGCTCGTGGAATTGCAGGTGTAGCGGTGAAATGCGTAGATATCTGCAGGAACACCAGAGGCGAAGGCGGCTCACTGGATCGCAACTGACGCTGAGAGACGAAAGCTAGGGG |
| P-asv3987 | GCAAGCGTTGTTCGGAATTACTGGGCGTAAAGGGCGCGTAGGCGGCCTTCTAAGTCGAACGTGAAATCCCTGGGCTCAACCCAGGAACTGCGTCCGATACTGGGAGGCTTGAATCCGGGAGAGGGATGCGGAATTCCAGGTGTAGCGGTGAAATGCGTAGATATCTGGAGGAACACCGGTGGCGAAGGCGGCACCCTGGGCCGGCATTGACGCTGAGGCGCGAAAGCCAGGGG |
| P-asv3988 | CCAAGCGTTGTTCGGATTCACTGGGCGTAAAGGGTGCGTAGGCGGTCGGATAAGTCTGATGTGAAATCTTCGAGCTCAACTCGGAAACTGCATCGGAAACTATTCGGCTGGAGGATTGGAGGGGAGACTGGAATTCTCGGTGTAGCAGTGAAATGCGTAGATATCGAGAGGAACACCAGTGGCGAAGGCGAGTCTCTGGACAATTCCTGACGCTGAGGCACGAAAGCTAGGGG |
| P-asv3989 | GCAAGCGTTAATCGGATTTACTGGGCGTAAAGGGCGCGTAGGTGGTAAGGCAAGTAAGATGTGAAAGCCCTGGGCTTAACCTGGGAAGTGCATCTTAAACTGCCTGGCTAGAGTATGGTAGAGGGTAGCGGAATTTCCGGTGTAGCGGTGAAATGCGTAGAGATCGGAAGGAACGCCAGTGGTGAAGACGGCTACCTGGACCAATACTGACACTGAGGCGCGAAAGCGTGGGG |
| P-asv3990 | GCGAACGTTGTTCGGAATCACTGGGCTTAAAGGGCGCGTAGGCGGGTCTTCAAGTTTGGGGTGAAATCCTCCGGCTCAACCGGAGAACTGCCTCGAATACTGAAGGTCTCGAGGGGGATCGGGGCATGCGGAACTACGGGTGGAGCGGTGAAATGCGTTGATATCCGTAGGAACTCCGGTGGCGAAGGCGGCGTGCTGGATCCCTTCTGACGCTGAGGCGCGAAAGCCAGGGG |
| P-asv3991 | GCGAGCGTTGTTCGGATTTACTGGGCGTAAAGGGCGCGTAGGCGGCCTTGTTTGTCAGGCGTGAAAGCCCTGGGCTCAACCTGGGAACTGCGCTTGGGACGGCGAGGCTAGAGATCGGGAGAGGAGAGTGGAATTCCCAGTGTAGAGGTGAAATTCGTAGATATTGGGAAGAACACCGGTGGCGAAGGCGGCTCTCTGGACCGATACTGACGCTGAGGCGCGAAAGCGTGGGG |
| P-asv3992 | GCGAGCGTTGTCCGGAATTATTGGGCGTAAAGAGCTCGTAGGCGGCTTGTCACGTCGGTTGTGAAAGCCCGGGGCTTAACCCCGGGTCTGCAGTCGATACGGGCAGGCTAGAGTGTGGTAGGGGAGATCGGAATTCCTGGTGTAGCGGTGAAATGCGCAGATATCAGGAGGAACACCGGTGGCGAAGGCGGATCTCTGGGCCATTACTGACGCTGAGGAGCGAAAGCGTGGGG |
| P-asv3993 | GCAAGCGTTGTTCGGAATTACTGGGCGTAAAGGGCTCGTAGGCGGCCAACTAAGTCAGACGTGAAATCCCTCGGCTTAACCGGGGAACTGCGTCTGATACTGGATGGCTTGAGGTTGGGAGAGGGATGCGGAATTCCAGGTGTAGCGGTGAAATGCGTTGATATCTGGAGGAACACCGGTGGCGAAGGCGGCATCCTGGACCAATTCTGACGCTGAGGAGCGAAAGCCAGGGG |
| P-asv3994 | GCGAGCGTTGTCCGGAATTACTGGGCGTAAAGGGCTTGTAGGTGGTTTGTCGCGTTGTTCGTGAAATGCCACAGCTTAACTGTGGGCGTGCGGGCGATACGGGCAGACTGGAGTACTGCAGGGGAGACTGGAATTCCTGGTGTAGCGGTGGAATGCGCAGATATCAGGAGGAACACCGGTGGCGAAGGCGGGTCTCTGGGCAGTAACTGACGCTGAGGAGCGAAAGCGTGGGG |
| P-asv3995 | GCGAGCGTTGTTCGGATTTACTGGGCGTAAAGGGCGCGTAGGCGGCGGCCCAAGTCGAGTGTGAAAGCCCCGGGCTCAACCTGGGAGGTGCGCTCGAGACTGGTCTGCTTGAGTTCGGGAGAGGAGAGTGGAATTCCCAGTGTAGAGGTGAAATTCGTAGATATTGGGAAGAACACCGGTGGCGAAGGCGGCTCTCTGGACCGAGACTGACGCTGAGGCGCGAAAGCGTGGGG |
| P-asv3996 | GCTAGCGTTGTTCGGAATCACTGGGCGTAAAGGGCGCGTAGGCGGCCAATTAAGTCGGGGGTGAAAGCCTGTGGCTCAACCACAGAATTGCCTTCGATACTGATTGGCTTGAGACCGGAAGAGGTAAGTGGAACTGCGAGTGTAGAGGTGAAATTCGTAGATATTCGCAAGAACACCAGTGGCGAAGGCGGCTTACTGGTCCGGTTCTGACGCTGAGGCGCGAAAGCGTGGGG |
| P-asv3997 | GCAAGCGTTGTTCGGATTTACTGGGCGCACAGGGAGCGTAGGCGGTTCGGTAAGCCCTTTGTGAAATCTCCAGGCTTAACCTGGAACAGTCGGAGGGGACTGCTGGGCTAGAGGACGGGAGAGGAGCGCGGAATTCCCGGTGTAGCGGTGAAATGCGTAGAGATCGGGAGGAAGGCCGGTGGCGAAGGCGGCGCTCTGGAACGTTTCTGACGCTGAGGCTCGAAAGCGTGGGG |
| P-asv3998 | GCCAGCGTTGTTCGGAATTACTGGGTGTAAAGGGTTCGTAGGCGGTGCGGCAAGTTGGGAGTGAAATCTCTGGGCTTAACCCAGAGGCTGCTTCCAAAACTGCTGTGCTCGAGTGTGAGAGAGGCTCGTGGAATTGCAGGTGTAGCGGTGAAATGCGTAGATATCTGCAGGAACACCCGTGGCGAAGGCGGCTCACTGGATCGCAACTGACGCTGAGAGACGAAAGCTAGGGG |
| P-asv3999 | TCAAGCGTTGTCCGGATTTATTGGGCGTAAAGAGCTCGTAGGTGGTTCGATAAGTCGGGTGTGAAACCTCCAGGCTCAACCTGGAGACGCCACCCGATACTGTCGTTACTAGAGTCCAGTAGGGGAGCGTGGAATTCCTGGTGTAGCGGTGAAATGCGCAGATATCAGGAGGAACACCAGCGGCGAAGGCGGCGCTCTGGGCTGGCACTGACACTGAGGAGCGAAAGCGTGGGG |
| P-asv4000 | GCAAGCGTTGTTCGGAATCATTGGGCGTAAAGCGCGTGTAGGCGGCTTGGCAAGTCGGGTGTGAAATCCCGGGGCTCAACCCCGGAACTGCATTCGATACTGCCTCGCTAGAGTCCCGGAGAGGAGGGCGGAATTCCCGGTGTAGAGGTGAAATTCGTAGATATCGGGAGGAACACCAGTGGCGAAGGCGGCCCTCTGGACGGTGACTGACGCTGAGATGCGAAAGCGTGGGG |
| P-asv4001 | GCAAGCGTTGTTCGGAATTACTGGGCGTAAAGGGCTCGTAGGTGGCCAACTAAGTCGGACGTGAAATCCCTCGGCTTAACTGGGGAACTGCGTCTGATACTGGCGAGCTTGAGTGCAGGAGAGGAACGCGGAATTCCAGGTGTAGCGGTGAAATGCGTAGATATCTGGAGGAACACCGGTGGCGAAGGCGGCGTTCTGGACTGCAACTGACACTGAACGACGAAAGCCAGGGG |
| P-asv4002 | GCTAGCGTTGTCCGGAATCATTGGGCGTAAAGAGCGTGTAGGCGGTCCGGTAAGTCCGCTGTGAAAGTCGGGGGCTCAACCCTCGAATGCCGGTGGATACTGTCGGGCTAGAGTGCGGAAGAGGCGAGTGGAATTCCTGGTGTAGCGGTGAAATGCGCAGATATCAGGAGGAACACCAATTGCGAAGGCAGCTCGCTGGGACGTAACTGACGCTGAGACGCGAAAGCGTGGGG |
| P-asv4003 | ACGGGCGTTGTCCGGATTCATTGGGCGCAAAGCGCGCGTAGGCGGCTCGGAAAGTCGGTCGTGAAATGCCGGGGCTCAACCCCGGGACTGCGTCCGATACTTCCGGGCTGGAGGTAGGTAGGGGAGATCGGAATTCCTGGTGTAGCGGTGAAATGCGCAGATATCAGGAGGAACACCGGTGGCGAAGGCGGATCTCTGGGCCTTACCTGACGCTGAGGCGCGAAAGCTAGGGG |
| P-asv4004 | GCAAGCGTTGTTCGGAATTACTGGGCGTAAAGGGCGCGTAGGCGGCCTGTTAAGTCGAACGTGAAATCCCCGGGCTTAACCCGGGAACTGCGTCCGATACTGGCAGGCTTGAATCCGGGAGAGGGATGTGGAATTCCAGGTGTAGCGGTGAAATGCGTAGATATCGAGAGGAACACTTGCGGCGAAGGCGGGTTGCTGGGCCGACACTGACGCTGAGGCGCGAAAGCTAGGGG |
| P-asv4005 | GCAAGCGTTGTTCGGAATTATTGGGTGTAAAGGGCGCGTAGGCGGCGCGACAAGTCACCTGTGAAATCTCTGGGCTTAACTCAGAGCCTGCAGGCGAAACTGTCGTGCTGGAGTATGGGAGAGGTGCGTGGAATTCCCGGTGTAGCGGTGAAATGCGTAGATATCGGGAGGAACACCTGTGGCGAAAGCGGCGCACTGGACCATAACTGACGCTGAGGCGCGAAAGCTAGGGG |
| P-asv4006 | GCAAGCGTTATCCGGATTTATTGGGCGTAAAGGGCTCGTAGGCGGCTCGTCGCGTCCGGTGTGAAAGTCCATCGCTTAACGGTGGATCTGCGCCGGGTACGGGCGGGCTGGAGTGCGGTAGGGGAGACTGGAATTCCCGGTGTAACGGTGGAATGTGTAGATATCGGGAAGAACACCGATGGCGAAGGCAGGTCTCTGGGCCGTCACTGACGCTGAGGAGCGAAAGCGTGGGG |
| P-asv4007 | GCAAGCGTTGTTCGGAATTACTGGGCGTAAAGGGCGCGTAGGCGGCCTTCTAAGTCGAACGTGAAATCCCTGGGCTCAACCCAGGAACTGCGTCCGATACTGGAAGGCTTGAATCCGGGAGAGGGATGCGGAATTCCAGGTGTAGCGGTGAAATGCGTAGATATCTGGGGGAACACCGGTGGCGAAGGCGGCATCCTGGACCGGTATTGTCGCTGAGGCGCGAAAGCCAGGGG |
| P-asv4008 | GCAAGCGTTATTCGGAATTATTGGGCGTAAAGGGCGCGTAGGCGGTTTTTTAAGTCGGATGTGCAATCCCTGGGCTCAACCCAGGAACTGCATCCGAGACTGGAAGGCTAGAGTACTGGAGAGGGTGGTGGAATTCCTCGTGTAGCGGCGAAATGCGTAGAGATGAGGAGGAACACCAGTGGCGAAGGCGGCCACCTGGACAGTAACTGACGCTGAGGCGCGAAAGCGTGGGT |
| P-asv4009 | GCAAGCGTTGTTCGGAATTACTGGGCGTAAAGGGCTCGTAGGCGGCCAACTAAGTCAGACGTGAAATCCCTCGGCCTAACCGGGGAACTGCGTCTGATACTGGATGGCTAGAGTTTGGGAGAGGGATGCGGAATTCCAGGTGTAGCGGTGAAATGCGTAGATATCTGGAGGAACACCGGTGGCGAAGGCGGCATCCTGGACCAACACTGACGCTGAGGAGCGAAAGCCAGGGG |
| P-asv4010 | GCTAGCGTTGCTCGGAATCACTGGGCGTAAAGCGCACGTAGGCGGGTCGTTAAGTCGAGGGTGAAATCCTGGAGCTCAACTCCAGAACTGCCTTCGATACTGGCGGCCTTGAGTCCGGGAGAGGTGAGTGGAACTGCGAGTGTAGAGGTGAAATTCGTAGATATTCGCAAGAACACCAGTGGCGAAGGCGGCTCACTGGCCCGGTACTGACGCTGAGGTGCGAAAGCGTGGGG |
| P-asv4011 | GCAAGCGTTGCCCGGAATTACTGGGCGTAAAGGGCGCGTAGGCGGGATGGCAAGTCGGTCGTGAAATCCCTCGGCTCAACTGAGGAATGTCGATCGATACTGCTGTTCTTGAGGATTGCAGAGGAGACTGGAATTCCCGGTGTAGCGGTGAAATGTGTAGAGATCGGGAGGAACACCGGTGGCGAAGGCGGGTCTCTGGGCAATTCCTGACGCTGAGGCGCGAAAGCGTGGGG |
| P-asv4012 | GCTAGCGTTGCTCGGAATCACTGGGCGTAAAGCGCACGTAGGCGGCTTTTTAAGTCAGAGGTGAAATCCTGGAGCTCAACTCCAGAACTGCCTTTGATACTGAGGAGCTTGAGTCCGGGAGAGGTGAGTGGAACTGCGAGTGTAGAGGTGAAATTCGTAGATATTCGCAAGAACACCAGTGGCGAAGGCGGCTCACTGGCCCGGTACTGACGCTGAGGTGCGAAAGCGTGGGG |
| P-asv4013 | GCCAGCGTTGCTCGGAATTACTGGGTGTAAAGGGTTCGTAGGCGGTGCGGCAAGTCGGGAGTGAAATCTCTGGGCTCAACCCAGAGGCTGCTTCCGAAACTGCTGTGCTTGAGTGTGGGAGAGGCGCGTGGAATTGCAGGTGTAGCGGTGAAATGGGTAGATATCTGCAGGAACACCCGTGGCGAAAGCGGCGCGCTGGACCACAACTGACGCTGAGGAACGAAAGCTAGGGG |
| P-asv4014 | GCGAGCGTTAATCGGAATTACTGGGCGTAAAGTGTGCGCAGGCGGCCTCGCAAGTCGAGTGTGAAATCCCCGAGCTTAACTTGGGAATTGCGCTCGAAACTACGGGGCTTGAGTGTGGCAGAGGGGGGTGGAATTCCACGTGTAGCGGTGAAATGCGTAGAGATGTGGAGGAACACCAATGGCGAAGGCAGCCCCCTGGGCCAACACTGACGCTCATGCACGAAAGCGTGGGG |
| P-asv4015 | GCAAACGTTGTTCGGAATCACTGGGCGTAAAGGGCGTGTAGGCGGCTGAATGAGTGGGATGTTAAATCCCCCGGCTCAACCGGGGAACTGCATTTCATACTGTTCAGCTTGAGTACATCAGAGGAGACTGGAATTCCCGGTGTAGCGGTGAAATGCGTAGATATCGGGAGGAACACCAGTGGCGAAGGCGAGTCTCTGGGATGATACTGACGCTGAGGCGCGAAAGCCAGGGG |
| P-asv4016 | GCGAGCGTTGTTCGGATTTACTGGGCGTAAAGGGCGCGCAGGCGGCGGACCAAGTCAGGCGTGAAAGCCGCGGGCTTAACCTGGGAGGTGCGTCTGGGACTGGTATGCTAGAGTTCGGGAGAGGAGAGTGGAATTCCCAGTGTAGAGGTGAAATTCGTAGATATTGGGAAGAACACCGGTGGCGAAGGCGGCTCTCTGGACCGAGACTGACGCTGAGGCGCGAAAGCGTGGGG |
| P-asv4017 | GCAAGCGTTGTTCGGAGTTACTGGGCGTAAAGAGTGTGTAGGCTGTTCTCTAAGTTTGGTGTGAAATTTCCCGGCTTAACTGGGAGGGTGCGCCGAAAACTGGAGGGCTTGAGTATGGGAGAGGAAAGCGGAATTCCTGGTGTAGCGGTGAAATGCGTAGATATCAGGAGGAACACCTGCGGTGTAGACGGCTTTCTGGACCATCACTGACGCTGAGACACGAAAGCGTGGGT |
| P-asv4018 | GCTAGCGTTGTTCGGAATTACTGGGCGTAAAGCGCGCGCAGGCGGCCTTCCAAGTCAGGGGTGAAAGCCCGGAGCTCAACTCCGGAATTGCCTTTGAAACTGTCAGGCTAGAGTACGGGAGAGGTGAGTGGAATTCCCAGTGTAGAGGTGAAATTCGTAGATATTGGGAAGAACACCGGTGGCGAAGGCGGCTCACTGGCCCGTTTCTGACGCTCAGGCGCGACAGCGTGGGG |
| P-asv4019 | GCAAGCGTTAATCGGAATTACTGGGCGTAAAGCGTGCGCAGGCGGTTATGTAAGACAGGTGTGAAATCCCCGGGCTTAACCTGGGAACTGCATTTGTGACTGCATAGCTAGAGTACGGTAGAGGGGGATGGAATTCCGCGTGTAGCAGTGAAATGCGTAGATATGCGGAGGAACACCGATGGCGAAGGCAATCCCCTGGACCTGTACTGACGCTCATGCACGAAAGCGTGGGG |
| P-asv4020 | GCAAACGTTGCTCGGAATCACTGGGCGTAAAGCGCACGTAGGCGGCTCGAAAAGTCGGATGTGAAAGCCCTGGGCTCAACCTAGGAAGTGCATTCGAAACTGCCGAGCTTGAGTACGGGAGAGGGTTGCGGAATTCCCGGTGTAGAGGTGAAATTCGTAGATATCGGGAGGAACACCAGTGGCGAAGGCGGCAGCCTGGACCGATACTGACGCTGAGGTGCGAAAGCGTGGGG |
| P-asv4021 | CCAAGCGTTGTTCGGATTCACTGGGCGTAAAGGGTGCGTAGGTGGTGGGGTAAGTCGGATGTGAAATCTCCGAGCTCAACTCGGAAACCGCATTGGAAACTACCCTGCTAGAGGGTCGGTGGGGGGACTGGAATACTTGGTGTAGCAGTGAAATGCGTAGATATCAAGTGGAACACCAGTGGCGAAGGCGAGTCCCTGGACGACTCCTGACACTGAGGCACGAAAGCTAGGGG |

| | |
|---|---|
| P-asv4022 | GCGAGCGTTGTCCGGAGTTACTGGGCGTAAAGGGTGCGCAGGCGGCCGCGCGCGTTGGGGATGAAATCTCCCTGCTCAACGGGGAGGGTGTTTCCAAGACGGCGTGGCTTGAGGCTCGGGGGGGGGCGTGGAATTCCGGGTGGAGCGGTGAAATGCGTAGAGATCCGGAGGAACACCGACGGCGTAAGCAGCGCCCTAGACGAGTCCTGACGCTCAGGCACGAAAGCTGGGGG |
| P-asv4023 | GCCAGCGTTGTTCGGAATTACTGGGTGTAAAGGGTTCGTAGGCGGTGTGGCAAGTTGGGAGTGAAATCTCTGGGCTCAACCCAGAGGCTGCTTCCAAAACTGCCATGCTTGAGTGTGGGAGAGGCGCGTGGAATTGCAGGTGTAGCGGTGAAATGCGTAGATATCTGCAGGACACCCGTGGCGAAAGCGGCGCGCTGGACCACAACTGACGCTGAGGAACGAAAGCTAGGGG |
| P-asv4024 | GCTAGCGTTGCTCGGAATCATTGGGCGTAAAGGGCGCGTAGGCGGCGCAGCTAGTCAGGCGTGAAACCCCCGGGCTCAACCCGGGGACTGCGTTTGATACGGCTGTGCTAGAGGATGGAAGAGGGCCGCGGAATTCCCAGTGTAGAGGTGAAATTCGTAGATATTGGGAAGAACACCGGTGGCGAAGGCGGCGGCCTGGTCCATTACTGACGCTGAGGCGCGACAGCGTGGGG |
| P-asv4025 | GCTAGCGTTGTTCGGAATTACTGGGCGTAAAGCGCGCGTAGGCGGCGGGACAAGTTGGGCGTGAAAGCCCGGGGCTCAACCCCGGAGGTGCGCCCGAGACTGTCCTGCTGGAGGCCGAGAGAGGGTGGCGGAATTCCCAGTGTAGAGGTGAAATTCGTAGATATTGGGAAGAACACCGGTGGCGAAGGCGGCCACCTGGCTCGGAACTGACGCTGAGGCGCGAAAGCGTGGGG |
| P-asv4026 | GCGAGCGTTATTCGGAATGACTGGGCGTAAAGCGCACGTAGGCGGTTTGCTAAGTCCATTGTGAAAGCCCACGACTCAATCGTGGAAGTGCAGCGGATACTGGCAGACTGGAGTGCAGGAGAGGAGAGCGGAATTCCTGGTGTAGAGGTGAAATTCGTAGATATCAGGAGGAACAACAGCGGCGAAGGCGGCTCTCTGGACTGTAACTGACGCTGAGGTGCGAAAGTGTGGGT |
| P-asv4027 | GCAAGCGTTAATCGGAATTACTGGGCGTAAAGCATGCGCAGGCGATTTTGTAAGTCTGATGTGAAATCCCCGGGCTCAACCTGGGAACTGCATTGGAGACTGCAAGGCTAGAGTGTGTCAGAGGGGGGTAGAATTCCACGTGTAGCAGTGAAATGCGTAGAGATGTGGAGGAATACCGATGGCGAAGGCAGCCCCCTGGGATAACACTGACGCTCATGCACGAAAGCGTGGGG |
| P-asv4028 | GCGAGCGTTATCCGGAATTATTGGGTGTAAAGGGTGCGTAGACGGGAGAACAAGTTAGTTGTGAAATACCTCGGCTCAACTGAGGAACTGCAACTAAAACTGTACTTCTTGAGTGCAGGAGAGGTAAGTGGAATTCCTAGTGTAGCGGTGAAATGCGTAGATATTAGGAGGAACACCAGTGGCGAAGGCGACTTACTGGACTGTAACTGACGTTGAGGCACGAAAGTGTGGGG |
| P-asv4029 | GCAAGCGTTAATCGGAATTACTGGGCGTAAAGCGAGCGTAGGTGGCTTGATAAGTCAGATGTGAAATCCCCGGGCTTAACCTGGGAACTGCATCTGATACTGTTAAGCTAGAGTAGGTGAGAGGGAAGTAGAATTTCAGGTGTAGCGGTGAAATGCGTAGAGATCTGAAGGAATACCGATGGCGAAGGCAGCTTCCTGGCATCATACTGACACTGAGGCTCGAAAGCGTGGGT |
| P-asv4030 | GCAAGCGTTAATCGGAATTACTGGGCGTAAAGCGCGCGTAGGTGGTTCGTTAAGTTGGATGTGAAATCCCCGGGCTCAACCTGGGAACTGCATTCAAAACTGTCGAGCTAGAGTATGGTAGAGGGTGGTGGAATTTCCTGTGTAGCGGTGAAATGCGTAGATATAGGAAGGAACACCAGTGACGAAGGCGGCCACCTGGACTGATACTGACACTGAGGTGCGAAAACGTGGGG |
| P-asv4031 | GCTAGCGTTGTTCGGAATTACTGGGCGTAAAGCGCACGTAGGCGGATCGATCAGTCAGGGGTGAAATCCCAGGGCTCAACCCTGGAACTGCCTTTGATACTGTCGATCTGGAGTATGGAAGAGGTGAGTGGAATTCCGAGTGTAGAGGTGAAATTCGTAGATATTCGGAGGAACACCAGTGGCGAAGGCGGCTCACTGGTCCATTACTGACGCTGAGGTGCGAAAGCGTGGGG |
| P-asv4032 | GCAAGCGTTAATCGGAATTACTGGGCGTAAAGCGCACGCAGGCGGTCTGTCAAGTCGGATGTGAAATCCCCGGGCTCAACCTGGGAACTGCATCCGAAACTGGCAGGCTAGAGTCTTGTAGAGGGGGGTAGAATTCCAGGTGTAGCGGTGAAATGTGTAGAGATCTGGAGGAATACTGGTGGCGAAGGCGGCCCCTTGGACAAAGACTGACGCTCAGGTGCGAAAGCGTGGGG |
| P-asv4033 | GCAAGCGTTATCCGGAATTACTGGGTGTAAAGGGTGCGTAGGTGGTATGGCAAGTCAGAAGTGAAAACCCAGGGCTTAACTCTGGGACTGCTTTTGAAACTGTCAGACTAGAGTGCAGGAGAGGTAAGCGGAATTCCTAGTGTAGCGGTGAAATGCGTAGATATTAGGAGGAACATCAGTGGCGAAGGCGGCTTACTGGACTGAAACTGACACTGAGGCACGAAAGCGTGGGG |
| P-asv4034 | CCAAGCGTTATCCGGAATCATTGGGTTTAAAGGGTCCGTAGGCGGTTTTTTAAGTCAGAGGTGAAATCCTGCAGCTCAACTGTAGCATTGCCTTTGATACTGAAAGACTTGAGTTATTGTGAAGTAGTTAGAATGTGTAGTGTAGCGGTGAAATGCATAGATATTACACAGAATACCGATTGCGAAAGCAGATTACTAACAATATACTGACGCTGAGGGACGAAAGCGTGGGT |
| P-asv4035 | CCAAGTGATGGCCGCTATTATTGGGCCTAAAGCGTCCGTAGCTTGCCGAGCAAGTCCATCGGGAAATCGACGCGCTCAACGCGTCGGCTTCCGGTGGAAACTGCCCGGCTTGGGGCCGGAAGACTCGGCGGGTACGTCCGGGGTAGGAGTGAAATCCCGTAATCCTGGACGGACCACCAATGGCGAAAGCACGCCGAGAAGCCGGACCCGACAGTGAGGGACGAAAGCCAGGGT |
| P-asv4036 | ACAAGCGTTGTCCGGAATTACTGGGTGTAAAGGGAGCGTAGGCGGGAAGACAAGTTGGAAGTGAAATCTATGGGCTCAACCCATAAACTGCTTTCAAAACTGTTTTTCTTGAGTAGTGCAGAGGTAGGCGGAATTCCCGGTGTAGCGGTGGAATGCGTAGATATCGGGAGGAACACCAGTGGCGAAGGCGGCCTACTGGGCACCAACTGACGCTGAGGCTCGAAAGTGTGGGT |
| P-asv4037 | CCGAGCGTTATCCGGATTTACTGGGCGTAAAGAGTCCGTAGCTGGCATTGTAAGTCTTGATTCAAATACTCTGACTCACTCAGAGAAAGGGTCAAGATACTGCAAAGCTAGAGATCATAGGGGGTTAGTGGAATTTCCGGTGGAGGGGTGAAATCCGTTGATATCGGAAGGAATGCCGAAAGCGAAAGCAGCTAACTACTATATTCTGACAGTGAAGGACGAAAGCTTGGGG |
| P-asv4038 | GCGAACGTTACTCGGAATCACTGGGCTTACAGGGTGCGTAGGTGGCTTGTTAAGCAGGGTGTGAAATCCCTCGGCTCAACCGAGGAACTGCGCTCTGAACTGGCGAGCTAGAGTATGCTAGAGGTGAATGGAACATCCAGTGGAGCGGTGAAATGTGTTGATATTGGATGGAACACCAGAGGCGAAGGCGATTCACTGGGGCATAACTGACACTGAGGCACGAAAGCCAGGGG |
| P-asv4039 | GCAAACGTTATTCGGAATTATTGGGCGTAAAGGGTGTGTAGGCGGCTTATCAAGTCTATAGTTAAATCTCTCGGCTTAACCGAGGGGCCGCTATGGAAACTGATTTGCTAGAGTATGGAAGAGAGAAGTGGAATTCTCGGAGTAGCGGTAAAATGCGTGGATCTCGAGAGGAACACCGATGGCGAAGGCAGCTTCTTGGTCCATTTCTGACGCTGAAACACGAAAGCGTGGGG |
| P-asv4040 | GCAAGCGTTAATCGGAATTACTGGGCGTAAAGCGTGCGCAGGCGGTTATATAAGACAGTTGTGAAATCCCCGGGCTCAACCTGGGAACTGCATCTGTGACTGTATAGCTAGAGTACGGTAGAGGGGGATGGAATTCCGCGTGTAGCAGTGAAATGCGTAGATATGCGGAGGAACACCGATGGCGAAGGCAATCCCCTGGACTGTACTGACGCTCATGCACGAAAGCGTGGGG |
| P-asv4041 | GCAAGCGTTAATCGGAATTACTGGGCGTAAAGCGTGCGTAGGTGGTTCGCTAAGTCAGGTGTGAAAGCCCGGGGCTTAACCTCGGAATTGCATTTGATACTGGCGAGCTAGAGTGTGGTAGAGGGTGGTGGAATTCCCGGTGTAGCGGTGAAATGCGTAGATATCGGGAGGAACATCAGTGGCGAAGGCGGCCACCTGGACCAACACTGACACTGAGGCACGAAAGCGTGGGT |
| P-asv4042 | GCTAGCGTTGCTCGGAATCACTGGGCGTAAAGGGCGCGTAGGCGGACATTTAAGTCGGGGGTGAAAGCCCAGGGCTCAACCCTGGAATTGCCTCCGATACTGGGTGTCTAGAGACCGGAAGAGGTGAGTGGAACTGCGAGTGTAGAGGTGAAATTCGTAGATATTCGCAAGAACACCAGTGGCGAAGGCGGCTCACTGGTCCGGATCTGACGCTGAGGCGCGAAAGCGTGGGG |
| P-asv4043 | GCAAGCGTTGCTCGGATTTATTGGGCGTAAAGCGCGCGCAGGCGGATCGACAAGTCAGATGTGAAATCTCGGGGCTCAACCCCGAAACTGCGTCTGAAACTATCGATCTAGAATGTCATAGAGGGAAGAGGAATTTCACATGTAGGGGTAAAATCCGTAGAGATGTGAAGGAACACCAGAGGCGAAGGCGTCTTCCTGGATGATTATTGACGCTGAGGCGCGAAAGCGTGGGG |
| P-asv4044 | GCAAGCGTTGTCCGGATTTATTGGGCGTAAAGCGAGCGCAGGCGGTTTCTTAAGTCTGATGTGAAAGCCTTCGGCTCAACCGAAGAAGTGCATCGGAAACTGGGAAACTTGAGTGCAGAAGAGGACAGTGGAACTCCATGTGTAGCGGTGAAATGCGTAGATATATGGAAGAACACCAGTGGCGAAGGCGGCTGTCTGGTCTGTAACTGACGCTGAGGCTCGAAAGCATGGGT |
| P-asv4045 | CCGGGCGTTATCCGGATTTATTGGGTTTAAAGGGAGCGTAGGCTGTCTATTAAGCGTGTTGTGAAATTTACCGGCTCAACCGGTGGCTTGCAGCGCGAACTGGTCGACTTGAGTATGCAGGAAGTAGGCGGAATTCATGGTGTAGCGGTGAAATGCTTAGATATCATGACGAACTCCGATTGCGCAGGCAGCTTACTGTAGCATAACTGACGCTGATGCTCGAAAGTGCGGGT |
| P-asv4046 | GCTAGCGTTGTTCGGAATTACTGGGCGTAAAGCGCATGTAGGCGGATATTTAAGTCAGGGGTGAAATCCCGCAGCTCAACTGCGGAACTGCCTTTGATACTGGGTATCTTGAGTATGGAAGAGGTGAGTGGAATTCCGAGTGTAGAGGTGAAATTCGTAGATATTCGGAGGAACACCAGTGGCGAAGGCGGCTCACTGGTCCATTACTGACGCTGAGGTGCGAAAGCGTGGGG |
| P-asv4047 | GCAAGCGTTATCCGGAATCACTGGGTTTAAAGGGTGCGTAGGCGGCATGGTAAGTCAGGGGTGAAAGGCCGTCGCTTAACGATGGAACTGCCTTTGATACTGTCATGCTTGAATCAAGTTAAGGCCGGCGGAATGTGGCATGTAGCGGTGAAATGCATAGATATGCCATAGAACACCGATTGCGAAGGCAGCTGGCTAGGCTTAGATTGACGCTGAGGCACGAAAGCGTGGGG |
| P-asv4048 | CCAAGCGTTGTCCGGATTTATTGGGTTTAAAGGGTGCGTAGGCGGATTAATAAGTCAGTGGTGAAAGCTTTCGGCTCAACCGGAAAATTGCCATTGAAACTGTTAATCTTGAGTATAGTTGGGGTGAATGGAATGTTGCATGTAGCGGTGAAATGCTTAGATATGCAACAGAACACCGATCGCGAAGGCAGTTCACTAAACTATAACTGACGCTGAGGCACGAAAGCGTGGGT |
| P-asv4049 | CCGAGTGATGGCCGATATTATTGGGCCTAAAGCGTCCGTAGCTGGCCGGACAAGTCCGTTGGGAAATCTGTCCGCTTAACGGGCAGGCGTCCAGCGGAAACTGTTCGGCTTGGACCGGAAGACCTGAGGGGTACGTCCGGGGTAGGAGTGAAATCCTGTAATCCTGGACGGACCACCGGTGGCGAAAGCGCCTCAGGAGGACGGATCCGACAGTGAGGGACGAAAGCTAGGGT |
| P-asv4050 | CCGAGTGGTAGTCGTTATTATTGGGCTTAAAGTGTTCGTAGCCTGCCGTGTAAGTCCGCTGTGAAACCCACACGCTCAACGTGTGGATGTGCAGCGGATACTGCATGGCTAGGGACCGGACGGCGTCGAGGCTATTCCAGGAGTAGGAGTGAAATCCAAAAATTCCTGGAGGAGCACCTGTGGCGAAAGCGCTCGACGAGTACGGATCCGACGGTGAGGAACGAAAGCCGGGGG |
| P-asv4051 | GCGAGCGTTGTCCGGAATTATTGGGCGTAAAGAGCTCGTAGGCGGTCTGTCGCGTCTGCTGTGAAAACCCGAGGCTCAACCTCGGGCCTGCAGTGGGTACGGGCAGACTAGAGTGCGGTAGGGGAGAATGGAATTCCTGGTGTAGCGGTGGAATGCGCAGATATCAGGAGGAACACCGATGGCGAAGGCAGTTCTCTGGGCCGTAACTGACGCTGAGGAGCGAAAGCGTGGGG |
| P-asv4052 | GCAAGCGTTAATCGGAATTACTGGGCGTAAAGCGTGCGCAGGCGGTAATGTAAGACAGATGTGAAATCCCCGGGCTCAACCTGGGAACTGCATTTGTGACTGCATTGCTGGAGTACGGCAGAGGGGGATGGAATTCCGCGTGTAGCAGTGAAATGCGTAGATATGCGGAGGAACACCGATGGCGAAGGCAATCCCCTGGGCCTGTACTGACGCTCATGCACGAAAGCGTGGGG |
| P-asv4053 | GCTAGCGTTGTTCGGAATGACTGGGCGTAAAGCGCGCGTAGGCGGATTGGCAAGTTGGGGGTGAAATCCCGGGGCTCAACCTCGGAATGGCCTTCAAGACTGCTGGTCTTGAGTTCGAGAGAGGTGAGTGGAATTCCGAGTGTAGAGGTGAAATTCGTAGATATTCGGAGGAACACCAGTGGCGAAGGCGGCTCACTGGCTCGACACTGACGCTGAGGTGCGAAAGCGTGGGG |
| P-asv4054 | CCGAGCGTTGTTCGGAATCACTGGGCGTAAAGGGAGCGTAGGCGGCGTGGTAAGTCAGATGTGAAATCCCGGGGCTCAACCCCGGAACTGCATCCGATACTGCCGTGCTAGAGTAATGGAGGGGTAGCTGGAATTATCGGTGTAGCAGTGAAATGCGTGGATATCGATAGGAACACTCGTGGCGAAAGCGAGCTACTGGACATTTACTGACGCTGAGGCTCGAAGGCCAGGGG |
| P-asv4055 | CCAAACGTTATTCGGAATTACTGGGCTTAAAGAGTTCGTAGGCGGCTTGGAAAGTGGGGTGTGAAAGCCCTCGGCTCAACCGAGGAATTGCGCTCCAAACTCCCAAGCTTGAGGAGGCTAGAGGTGAGCAGAACTGATGGTGGAGCGGCGAAATGCGTTGATATCATCAGGAATACCGGAGGCGAAGGCGGCTCACTGGGACCTTTCTGACGCTGAGGAACGAAAGCTAGGGG |
| P-asv4056 | GCTAGCGTTGTTCGGAATTACTGGGCGTAAAGCGCACGTAGGCGGACCCGTAAGTCAGAGGTGAAAGCCTGGAGCTTAACTCCAGAACTGCCTTTGATACTGCGGGTCTTGAGGATGGGAGAGGTGAGTGGAACTGCGAGTGTAGAGGTGAAATTCGTAGATATTCGCAAGAACACCAGTGGCGAAGGCGGCTCACTGGCCCATTACTGACGCTGAGGTGCGAAAGCGTGGGG |
| P-asv4057 | GCTAGCGTTGTTCGGAATCACTGGGCGTAAAGCGCACGTAGGCGGACTGATCAGTCAGGGGTGAAATCCCGGGGCTTAACCTCGGAACTGCCTTTGATACTGTCAGTCTTGAGTCCGAGAGAGGTGAGTGGAACTCCGAGTGTAGAGGTGAAATTCGTAGATATCCGGAAGAACACCAGTGGCGAAGGCGGCTCACTGGCTCGGTACTGACGCTGAGGTGCGAAAGCGTGGGG |
| P-asv4058 | GCAAGCGTTGTCCGGATTTATTGGGTTTAAAGGGTGCGTAGGCGGCCCCTTAAGTCAGCGGTGAAAGCCCGGGGCTCAACCCCGGAACCGCCGTTGATACTGGGGGGCTTGAGTATGGACGAGGTTGGCGGAATTGAGGGTGTAGCGGTGAAATGCTTAGATACCCTCAAGAACACCGATTGCGTAGGCAGCTAACTAGGCCATAACTGACGCTGAGGCACGAAAGCGTGGGG |
| P-asv4059 | GCGAGCGTTGTCCGGATTTATTGGGCGTAAAGGGTTCGTAGGCGGTCTTTTAAGTCAGATGTGAAAGGTCACGGCTCAACCGTGGTAAGCATTTGAAACTGAAAGACTTGAGTTAAGGAGAGGAAAGTGGAATTCCTAGTGTAGCGGTGAAATGCGTAGATATTAGGAGGAATACCAGTGGCGAAGGCGACTTTCTGGACTTATACTGACGCTGAGGAACGAAAGCGTGGGG |
| P-asv4060 | GCAAGCGTTGTCCGGATTTACTGGGCGTAAAGAGCGCGCAGGCGGTCGAATAAGTCGAATGTGAAAGCCCCCGGCTCAACTGGGGAGGGTCATTCGATACTGTTCGACTCGAAGGCAGGAGAGGGAAGCGGAATTCCCGGTGTAGTGGTGAAATGCGTAGATATCGGGAGGAACACCAGTGGCGAAGGCGGCTTTCTGGCCTGCTCTTGACGCTGAGGCGCGAAAGCCAGGGG |

| | |
|---|---|
| P-asv4061 | GCGAGCGTTACCCGGATTTACTGGGTGTAAAGGGCGTGTAGGCGGTCTCTCAAGTCCGATGCTAAAGACCGAAGCTCAACTTCGGGGGTGCGTTGGATACTGTGAGGCTAGACGGTCGGAGAGGGTAGCGGAATTTCCGGAGTAGCGGTGAAATGCGCAGATACCGGAAGGAACGCCGATAGCGAAAGCAGCTACCTGGACGATTTGTGACGCTGAGGCGCGAAAGCGTGGGG |
| P-asv4062 | GCAAGCGTTAATCGGAATTACTGGGCGTAAAGCGCGCGTAGGTGGTTTGGTAAGTCGGATGTGAAAGCCCCGGGCTTAACCTGGGAATTGCATTCGATACTGCTTGACTAGAGTGTGGTAGAGGGAAGTGGAATTCCGCATGTAGTGGTGAAATGCGTAGAGATGCGGAGGAACACCAATGGCGAAGGCAACTTCCTGGACCAACACTGACACTGAGGTGCGAAAGCGTGGGT |
| P-asv4063 | GCTAGCGTTGTTCGGAATTACTGGGCGTAAAGCGCACGTAGGCGGACTGGAAAGTCAGAGGTGAAATCCCAGGGCTCAACCTTGGAACTGCCTTTGAAACTATCAGTCTGGAGTTCGAGAGAGGTGAGTGGAATTCCGAGTGTAGAGGTGAAATTCGTAGATATTCGGAGGAACACCAGTGGCGAAGGCGGCTCACTGGCTCGATACTGACGCTGGGGTGCGAAAGCGTGGGG |
| P-asv4064 | GCTAGCGTTGTTCGGAATTACTGGGCGTAAAGCGCACGTAGGCGGCTTTTTAAGTCAGGGGTGAAATCCCGGGGCTCAACCCCGGAACTGCCCTTGAAACTGGCAAGCTAGAATCTTGGAGAGGTCAGTGGAATTCCGAGTGTAGAGGTGAAATTCGTAGATATTCGGAAGAACACCAGTGGCGAAGGCGACTGACTGGACAAGTATTGACGCTGAGGTGCGAAAGCGTGGGG |
| P-asv4065 | GCAAGCGTTATCCGGAATTATTGGGCGTAAAGTGTCTGTAGGTGGCTTTTCAAGTCCGTCGTCAAATCCCAGGGCTCAACCCTGGACAGGCGGTGGAAACTATCAAGCTAGAGTACGGTAGAGGCAGAGGGAATTTCCGGTGGAGCGGTGAAATGCGTTGAGATCGGGAGGAACACCAAGGGCGAAAGCACTCTGCTGGGCCGTTACTGACACTCAGAGACGAAAGCTAGGGG |
| P-asv4066 | GCAAGCGTTGTTCGGAATTATTGGGCGTAAAGGGCGCGTAGGTGGTGCGGTAAGTCGCCTGTGAAACCTCTAGGCTCAACCTAGAGCCTGCAGGCGAAACTGCCGTGCTTGAGGGTGGGAGAGGTGCGTGGAATTCCCGGTGTAGCGGTGAAATGCGTAGATATCGGGAGGAACACCTGTGGCGAAAGCGGCGCACTGGACCACATCTGACGCTGAGGCGCGAAAGCTAGGGG |
| P-asv4067 | GCGAGCGTTGTCCGGAATTATTGGGCGTAAAGAGCTTGTAGGCGGTTTGTCGCGTCTGCTGTGAAAATCCGAGGCTCAACCTCGGACTTGCAGTGGGTACGGGCAGACTAGAGTGTGGTAGGGGAGACTGGAATTCCTGGTGTAGCGGTGGAATGCGCAGATATCAGGAGGAACACCGATGGCGAAGGCAGGTCTCTGGGCCACTACTGACGCTGAGAAGCGAAAGCTAGGGG |
| P-asv4068 | GCAAGCGTTGTCCGGATTTACTGGGCGTAAAGAGCGCGCAGGCGGTCGATTTAGTCGTGTGTGAAAGCCCCCGGCTCAACTGGGGAGGGTCACGCGATACTGATCGACTCGAAGGCAGGAGAGGGTAGTGGAATTCCCGGTGTAGTGGTGAAATGCGTAGATATCGGGAGGAACACCAGTGGCGAAGGCGACTACCTGGCCTGTTCTTGACGCTGAGGCGCGAAAGCTAGGGG |
| P-asv4069 | GCGAGCGTTGTTCGGAATTATTGGGCGTAAAGGGCGCGTAGGCGGTGCCACAAGTCGCCTGTGAAATCCCCGGGCTTAACCCGGGGCCTGCAGGCGAAACTGTGGTGCTGGAGTGTGGGAGAGGTGCGTGGAATTCCCGGTGTAGCGGTGAAATGCGTAGATATCGGGAGGAACACCTGTGGCGAAAGCGGCGCACTGGACCACAACTGACGCTGAGGCGCGAAAGCTAGGGG |
| P-asv4070 | GCAAGCGTTGCTCGGAATTACTGGGCGTAAAGGGCGCGTAGGCGGGATGGCAAGTCGGTCGTGAAATCCCTCGGCTCAACTGAGGATCGTCGACCGATACTGCCGTTCTTGAGGATTGCAGAGGAGACTGGAATTCCCGGTGTAGCGGTGAAATGTGTAGAGATCGGGAGGAACACCGGTGGCGAAGGCGGGTCTCTGGGCAATTCCTGACGCTGAGGCGCGAAAGCGTGGGG |
| P-asv4071 | GCCAGCGTTGTTCGGAATTACTGGGTGCAAAGGGCTCGTAGGCGGTGTGGCAAGTTGGGAGTGAAATCTCTGGGCTCAACCCAGAGACGGCTTCCAAAACTGCCGTGCTTGAGTGTGGGAGAGGCTCGTGGAATTGCAGGTGTAGCGGTGAAATGCGTAGATATCTGCAGGAACACCCGTGGCGAAAGCGGCGAGCTGGACCACTACTGACGCTGAAGAACGAAAGCTAGGGG |
| P-asv4072 | GCAAGCGTTGTTCGGATTGACTGGGCGTAAAGGGCGTGTAGGCGGTCGGTTCAGTCCGGTGTGAAAGCCCGGGGCTCAGCCCCGGAAGTGCATCGGAAACTAGCCGACTTGAGCGCGATAGAGGAGGGTGGAATTCCTGGTGTAGCGGTGAAATGCGTAGATATCAGGAGGAACACCGGCGGCGAAGGCGGCCCTCTGGATCGTTGCTGACGCTGAGACGCGAAAGCGTGGGG |
| P-asv4073 | GCCAGCGTTGTTCGGAATTATTGGGCGTAAAGGGCGCGTAGGCGGTGCGGTAAGTCACCTGTGAAATCCCCGGGCTTAACTCGGGGCCTGCAGGCGAAACTGCCGTGCTGGAGTATGGGAGAGGTGCGTGGAATTCCCGGTGTAGCGGTGAAATGCGTAGATATCGGGAGGAACACCTGTGGCGAAAGCGGCGCACTGGACCATAACTGACGCTGAGGCGCGAAAGCTAGGGG |
| P-asv4074 | GCGAGCGTTGTCCGGAGTTATTGGGCGTAAAGAGCGTGTAGGCGGTTCGGTAAGTCTGCCGTGAAAACCCAGGGCTCAACCCTGGGCGTGCGGTGGATACTGCCGGGCTAGAGGGTGGTAGAGGCGAGTGGAATTCCCGGTGTAGCGGTGAAATGCGCAGATATCGGGAGGAACACCAGTAGCGAAGGCGGCTCGCTGGGCCATACCTGACGCTGAGACGCGAAAGCGTGGGG |
| P-asv4075 | GCAAGCGTTGTTCGGAATCACTGGGCTTAAAGAGCATGTAGGTGGGGTGTCAGGCGTCGGGTGAAATCCCTCGGCTCAACCGAGGAACTGCCTGGCGAACCGGCATTCTTGAGGCAGGTAGAGGTGGCTGGAACGACGGGTGGAGCGGTGAAATGCGTAGATATCCGTTGGAACGCCGGCGGCGAAAGCGAGCCACTGGGCCTGTCCTGACACTGAGATGCGAAAGCGTGGGG |
| P-asv4076 | GCAAGCGTTGTCCGGAATTATTGGGCGTAAAGAGCTCGTAGGCGGTCTGTTGCGTCGGCTGTGAAAACCCGGGGCTCAACCCCGGGCCTGCAGCCGATACGGGCAGACTAGAGTTCGGCAGGGGAGACTGGAATTCCTGGTGTAGCGGTGAAATGCGCAGATATCAGGAGGAACACCGGTGGCGAAGGCGGGTCTCTGGGCCGATACTGACGCTGAGGAGCGAAAGCGTGGGG |
| P-asv4077 | GCGAGCGTTGTCCGGAGTTACTGGGCGTAAAGGGCACGCAGGCGGTTGTGTCAGTTTGCGGGGACAGTGGGCGGCTTAACTGCCCAAGGGTCGCAAAGACGGCACGACTGGAGAGCGAGAGAGGGACCTGGAACTCCGGGTGGAGCGGTGAAATGCGTAGAGATCCGGAAGAACACCACTGGCGAAGGCAAGGTCCTGGCTCGTATCTGACGCTGAGGTGCGAAAGCTAGGGG |
| P-asv4078 | GCGAGCGTTAATCGGAGTTACTGGGCGTAAAGCGTGCGCAGGCGGTTTGTTAAGTCTGATGTGAAAGCCCCGGGCTCAACCTGGGAACTGCATTGGAAACTGGCAAGCTCGAGTATGGCAGAGGGGGGTGGAATTCCGCGTGTAGCAGTGAAATGCGTAGAGACGCGGAGGAACACCGATGGCGAAGGCAGCCCCCTGGGCCAATACTGACGCTCAGGCACGAAAGCGTGGGG |
| P-asv4079 | GCTAGCGTTGCTCGGAATCACTGGGCGTAAAGCGCACGTAGGCGGCTTCTTAAGTCAGGGGTGAAATCCTGGAGCTCAACTCCAGAACTGCCTTTGATACTGAGGAGCTGGAGTCCGGGAGAGGTGAGTGGAACTGCGAGTGTGGAGGTGAAATTCGTAGATATTCGCAAGAACACCAGTGGCGAAGGCGGCTCACTGGCCCGGTACTGACGCTGAGGTGCGAAAGCGTGGGG |
| P-asv4080 | GCGAGCGTTAATCGGAATTACTGGGCGTAAAGGGTGCGCAGGCGGTTCTGCAAGTCAGGCGTGAAAGCCCCGGGCTTAACCTGGGAACGGCGCTTGAAACTACAGGACTGGAGTATGGCAGAGGGAGGTGGAATTCCACGTGTAGCGGTGAAATGCGTAGAGATGTGGAGGAACACCGATGGCGAAGGCAGCCTCCTGGGCCAATACTGACGCTCATGCACGAAAGCGTGGGG |
| P-asv4081 | GCGAGCGTTGTTCGGATTTACTGGGCGTAAAGGGCGCGTAGGCGGCCCCTCAAGTCGCGTGTGAAAGCCCCGGGCTCAACCTGGGAATGGCGCGCGATACTGGGTGGCTGGAGTTCGGGAGAGGAGCGTGGAATTCCCAGTGTAGAGGTGAAATTCGTAGATATTGGGAAGAACACCGGTGGCGAAGGCGGCGCTCTGGACCGAGACTGACGCTGAGGCGCGAAGGCGTGGGG |
| P-asv4082 | GCGAGCGTTGTTCGGAATTATTGGGCGTAAAGGGTGCGTAGGCGGTGCCACAAGTCACTTGTGAAATCCCCAGGCTTAACCTGGGGCCTGCAGGCGAAACTGTGGTGCTGGAGTATGGGAGAGGTGCGTGGAATTCCCGGTGTAGCGGTGAAATGCGTAGATATCGGGAGGAACACCTGTGGCGAAAGCGGCGCACTGGACCATAACTGACGCTGATGCACGAAAGCTAGGGG |
| P-asv4083 | GCGAACGTTATTCGGAATCACTGGGCTTAAAGCGCGTGTAGGCGGACGGCCGCGTCGCTTGCTGAAAGCCCCCGGCTCAACCGGGGGACCGGCGGCGATACGGGCCGTCTGGAGGGGCGTAGGGGGAGCTGGAACTCACGGTGGAGCGGTGAAATGCGTTGAGATCGTGAGGAACGCCCGCGACGAAAGCGAGCTCCTGGACGCCTGCTGACGCTGAGACGCGAAAGCCAGGGG |
| P-asv4084 | GCAAGCGTTGTCCGGATTCACTGGGCGTAAAGAGCGTGTAGGCGGCCAGGCAGGTCTGTTGTGAAAACTGGAGGCTCAACCTCCAGACGTCGACAGGAACCGTCTGGCTAGAGTCCGGAAGAGGAGAGTGGAATTCCTGGTGTAGCGGTGAAATGCGCAGATATCAGGAAGAACACCAATGGCGAAGGCAGCTCTCTGGGACGGTACTGACGCTGAGACGCGAAAGCGTGGGG |
| P-asv4085 | GCGAGCGTTAATCGGAATTACTGGGCGTAAAGCGTGCGCAGGCGGTTCGTTAAGACAGATGTGAAGTCCCCGGGCTTAACCTGGGAACTGCATTTGTGACTGGCGAGCTAGAGTATGGCAGAGGGAGGTAGAATTCCACGTGTAGCAGTGAAATGCGTAGAGATGTGGAGGAATACCGATGGCGAAGGCAGCCTCCTGGGCCAATACTGACGCTCATGCACGAAAGCGTGGGG |
| P-asv4086 | GCTAGCGTTGTTCGGAATTACTGGGCGTAAAGCGTGTGTAGGCGGCTATGCAAGTCGGATGTGAAAGCCCCAGGCTTAACCTGGGAAGTGCACCCGAAACTGCATGGCTCGAGTATCGGAGAGGTTGGTGGAATTCTCGGTGTAGAGGTGAAATTCGTAGATATCGAGAGGAACACCGGCGGCGAAGGCGGCCAACTGGACGAATACTGACGCTGAGACACGAAAGCGTGGGG |
| P-asv4087 | GCAAGCGTTAATCGGAATTACTGGGCGTAAAGCGTACGTAGACGGTTGGATAAGTCGGATGTGAAAGCCCCGGGCTTAACCTGGGAATTGCATTCGAGACTGTGCAGCTAGGGTGCGGAAGAGGGAAGCGGAATTCCAGGTGTAGCGGTGAAATGCGTAGATATCTGGAGGAACACCAGTGGCGAAAGCGGCTTCCTGGTCCAGCACCGACGTTCAGGTACGAGAGCATGGGG |
| P-asv4088 | GCCAGCGTTGTTCGGAATTACTGGGTGTAAAGGGTTCGTAGGCGGTGTGGCAAGTTGGGAGTGAAATCTCTGGGCTCAACCCAGAGGCTGCTTCCAAAACTGCTGTGCTGGAGTGTGGGAGAGGCGCGTGGAATTGCAGGTGTAGCGGTGAAATGCGTAGATATCTGCAGGAACACCCGTGGCGAAGGCGGCGCGCTGGACCACTACTGACGCTGAGGAACGAAAGCTAGGGG |
| P-asv4089 | GCGAGCGTTGTCCGGAATTATTGGGCGTAAAGAGCTCGTAGGCGGTTTGTCGCGTCGGCCGTGAAAACTTGGGGCTTAACCCCGAGCTTGCGGTCGATACGGGCAGACTTGAGTTCGGCAGGGGAGACTGGAATTCCTGGTGTAGCGGTGAAATGCGCAGATATCAGGAGGAACACCGGTGGCGAAGGCGGGTCTCTGGGCCGATACTGACGCTGAGGAGCGAAAGCGTGGGG |
| P-asv4090 | GCAAGCGTTGTCCGGAATCACTGGGCGTAAAGCGCACGTAGGCGGATTTTTAAGTCAGGGGTGAAATCCCGAGGCTCAACCTCGGAACTGCCTTTGATACTGGGGATCTGGAGTCCGGGAGAGGTGAGTGGAACTGCGAGTGTAGAGGTGAAATTCGTAGATATTCGCAAGAACACCAGTGGCGAAGGCGGCTCACTGGCCCGGAACTGACGCTGAGGTGCGAAAGCGTGGGG |
| P-asv4091 | GCGAGCGTTGTCCGGATTTACTGGGCGTAAAGCGCACGCAGGCGGTTCCGAAAGTCGAATGTGAAAGCTCCCGGCCCAACTGGGAGAGGCCGTTCGAAACTTCGGAACTTGAAGGCGGGAGAGGCAGATGGAATTTCCGGTGTAGCGGTAAAATGCGTAGATACCGGAAGGAACACCAGTGGCGAAAGCGATCTGCTGGCCCGCCCTTGACGCTCAGGTGCGAAAGCTAGGGG |
| P-asv4092 | GCAAGCGTTGTTCGGAATTACTGGGCGTAAAGGGTTTGTAGGTGGCCGACTAAGTCAGACGTGAAATCCCTCAGCTTAACTGGGGAACTGCGTCTGGGACTGGTCGGCTTGAGGGCCGGAGAGGAACGCGGAATTCCAGGTGTAGCGGTGAAATGCGTAGATATCTGGAGGAACACCGGTGGCGAAGGCGGCGTTCTGGACGGCATCTGACACTGAGAAACGAAAGCCAGGGG |
| P-asv4093 | GCTAGCGTTGTTCGGAATTACTGGGCGTAAAGCGCACGTAGGCGGATCTTTAAGTCAGAGGTGAAATCCCAAGGCTCAACCTTGGAACTGCCTTTGATACTGGGGATCTTGAGTCCGGAAGAGGTGAGTGGAACTGCGAGTGTAGAGGTGAAATTCGTAGATATTCGCAAGAACACCAGTGGCGAAGGCGGCTCACTGGTCCGGTACTGACGCTGAGGTGCGAAAGCGTGGGG |
| P-asv4094 | GCGAACGTTGTTCGGATTTACTGGGCGTAAAGGGCGCGCAGGCGGCGGATTTTGTCAGGCGTGAAAGCCCCGGGCTTGACCTGGGAACTGCGCTTGAGACGGGTTCGCTAGAGATCGGGAGAGGAGAGTGGAATTCCCAGTGTAGAGGTGAAATTCGTAGATATTGGGAAGAACACCGGTGGCGAAGGCGGCTCTCTGGACCGATACTGACGCTGAGGCGCGAAAGCGTGGGG |
| P-asv4095 | GCGAGCGTCGTCCGGATTCATGGGGCGTAAAGAGCTCGTAGGCGGCTTGATAAGTCGGATGTGAAACCTCCAGGCTCAACCTGGAGTCGCCATCCGATACTGTCATGGCTAGAGTCCGGTAGGGGCCCACGGAACTCCTGGTGTAGCGGTGAAATGCGCAGATATCAGGAAGAACACCGGTGGCGAAGGCGGTGGGCTGGGCCGGTACTGACGCTGAGGAGCGAAAGCGTGGGG |
| P-asv4096 | GCGAGCGTTGTCCGGAATCACTGGGCGTAAAGGGCGCGTAGGCGGCTTTGTAAGGGTGCGGTGAAAGCCCGGGGCTTACCCCCGGGTCGGCCGTGCCGACTGCAAGGCTTGGGCACTGTAGAGGCAGGCGGAATTCCGGGTGTAGCGGTGGAATGCGTAGAGATCCGGAAGAACACCGGTGGCGAAGGCGGCCTGCTGGGCAGTTGCTGACGCTGAGGCGCGACAGCGTGGGG |
| P-asv4097 | GCAAGCGTTAATCGGAATTACTGGGCGTAAAGCGCACGCAGGTGGCTGAGTCAGTCGGATGTGAAAGCCCTGGGCTTAACCTGGGAATTGCATTCGATACTACTCAGCTAGAGTATGGGAGAGGGAAGTGGAATTCCCGGTGTAGCGGTGAAATGCGTAGATATCGGGAGGAACATCAGTGGCGAAGGCGGCTTCCTGGCCCAATACTGACACTCAGGTGCGACAGCGTGGGG |
| P-asv4098 | GCAAGCGTTATCCGGATTCACTGGGTTTAAAGGGTGCGTAGGCGGGTTGGTAAGTCAGTGGTGAAATCTTCAAGCTTAACTTGGAAACTGCCGTTGATACTATCAATCTTGAATATTGTGGAGGTAAGCGGAATATGTCATGTAGCGGTGAAATGCTTAGATATGACATAGAACACCGATAGCGAAGGCAGCTTGCTACACAATAATTGACGCTGAGGCACGAAAGCGTGGGG |
| P-asv4099 | TCAAGCGTTGTTCGGATTCATTGGGCGTAAAGGGTGCGTAGGTGGCGCCATAAGTCGGGTGTGAAATTTCGGAGCTTAACTCCGAAACTGCATTCGATACTGTGGTGCTTGAGGACTGGAGAGGAGACTGGAATTTACGGTGTAGCAGTGAAATGCGTAGAGATCGTAAGGAAGACCAGTGGCGAAGGCGGGTCTCTGGACAGTTCCTGACACTGAGGCACGAAGGCCAGGGG |

| | |
|---|---|
| P-asv4100 | GCTAGCGTTGCTCGGAATCACTGGGCGTAAAGCGCACGTAGGTGGGTCGTTAAGTCGGGGGTGAAATCCTGGAGCTCAACTCCAGAACTGCCCTTGATACTGAGGATCTCGAGTTCGGGAGAGGTGAGTGGAACTGCGAGTGTAGAGGTGAAATTCGTAGATATTCGCAAGAACACCAGTGGCGAAGGCGGCTCACTGGCCCGATACTGACGCTGAGGTGCGAAAGCGTGGGG |
| P-asv4101 | GCAAGCGTTGTTCGGAATCACTGGGCGTAAAGGGCGTGTAGGCGGTCCGCTAAGTCTGATGTGAAAGCCCGGGGCTCACCCCCGGAAGGGCATTGGAAACTGGCGGACTCGAGTCCCGGAGAGGAAGGTGGAATTCCTGGTGTAGCGGTGAAATGCGTAGATATCAGGAGGAATACCGGTGGCGAAGGCGGCCTTCTGGACGGTGACTGACGCTGAGACGCGAAAGCGTGGGG |
| P-asv4102 | GCAAGCGTTGTTCGGAATTATTGGGCGTAAAGGGTGCGTAGGCGGTGCGGTAAGTCTATGGTGAAATCTCCGGGCTCAACTCGGAGCCTGCCACGGAAACTGCCGTGCTTGAGTATGGGAGAGGTGAGTGGAATTCCCGGTGTAGCGGTGAAATGCGTAGATATCGGGAGGAACACCTGTGGCGAAAGCGGCTCACTGGACCATAACTGACGCTGATGCACGAAAGCTAGGGG |
| P-asv4103 | GCAAGCGTTGTCCGGAATCATTGGGCGTAAAGAGCGCGTAGGCGGCCAGATAAGTCCGGTGTGAAAGTCACAGGCTCAACCTGTGAATGCCGCTGGAAACTGTCTGGCTAGAGTCCGGTAGGGGCGAGTGGAATTCCTGGTGTAGCGGTGAAATGCGCAGATATCAGGAGGAACACCAATGGCGAAGGCAGCTCGCTGGAACGGTACTGACGCTGAGGCGCGAAAGCGTGGGG |
| P-asv4104 | GCGAGCGTTGTTCGGATTTATTGGGCGTAAAGCGAACGCAGGTGGTTTGGTAAGTCAGGGGTGAAAGACCACAGCTTAACTGTGGCATTGCCTTTGATACTGCCTTTCTTGAGTAGAGTAGGGGGAAGCGGAATTCCGAGTGTAGAGGTGAAATTCGTAGATATTCGGAGGAACACCGGTGGCGAAGGCGGCTTCCTGGACTCTAACTGACACTCATGTTCGAAAGCGTGGGG |
| P-asv4105 | GCCAGCGTTGTTCGGAATTACTGGGCGTAAAGGGTTCGTAGGCGGTGCGGCAAGTTGGGAGTGAAATCTCTGGGCTCAACCCAGAGTCAGCTTCCGAAACTGCTGTGCTTGAGTGTGGGAGAGGCGCGTGGAATTGCAGGTGTAGCGGTGAAATGCGTAGATATCTGCAGGAACACCCGTGGCGAAAGCGGCGCGCTGGACCACTACTGACGCTGAGGAACGAAAGCTAGGGG |
| P-asv4106 | CCAAGCGTTGTTCGGATTCACTGGGCGTAAAGGGTGCGTAGGTGGTGAGGTAAGTCGGATGTGAAAGCTCGGAGCTCAACTCCGAAATGGCATTGGAAACTACCTTGCTCGAGGGTTGGAGGGGGGACTGGAATTCTCGGTGTAGCAGTGAAATGCGTAGATATCGAGAGGAACACCAGTGGCGAAGGCGAGTCCCTGGACAACTCCTGACACTGAGGCACGAAAGCTAGGGG |
| P-asv4107 | GCGAGCGTTAATCGGAATTACTGGGCGTAAAGCGTGCGCAGGCGGTTGTGCAAGACAGATGTGAAATCCCCGGGCTTAACCTGGGAATGGCATTTGTGACTGCACGGCTAGAGTGTGTCAGAGGGGGGTAGAATTCCACGTGTAGCAGTGAAATGCGTAGATATGTGGAGGAACACCGATGGCGAAGGCAGCCCCCTAGGATAACACTGACGCTCATGCACGAAAGCGTGGGG |
| P-asv4108 | GCAAGCGTTAATCGGAATTACTGGGCGTAAAGCGCGCGTAGGTGGTTTTTTAAGTTGGATGTGAAAGCCCCGGGCTCAACCTGGGAACTGCATCCAAAACTGGCAAGCTAGAGTACAGTAGAGGGTGGTGGAATTTCCTGTGTAGCGGTGAAATGCGTAGATATAGGAAGGAACACCAGTGGCGAAGGCGACCACCTGGACTGATACTGACACTGAGGTGCGAAAGCGTGGGG |
| P-asv4109 | GCGAGCGTTGTCCGGATTTACTGGGCGTAAAGGGTGAGTAGGCGGTTGTATATGTCAGATGTAAAAGGCACAGGCTTAACTTGTGTAAGCATTTGAAACTGTACAACTTGAGTGCAGGAGAGGTAAGTGGAATTCCTAGTGTAGCGGTGAAATGCGTAGATATTAGGAGGAACACCAGTGGCGAAGGCGACTTACTGGACTGTAACTGACGCTGAGTCACGAAAGCGTGGGT |
| P-asv4110 | GCAAGCGTTGTCCGGAATTATTGGGCGTAAAGAGCTCGTAGGCGGTTTGTCGCGTCTGCTGTGAAATCCCGAGGCTCAACCTCGGGCTTGCAGTGGGTACGGGCAGACTAGAGTGCGGTAGGGGAGATTGGAATTCCTGGTGTAGCGGTGGAATGCGCAGATATCAGGAGGAACACCGATGGCGAAGGCAGATCTCTGGGCCGTAACTGACGCTGAGGAGCGAAAGCGTGGGG |
| P-asv4111 | GCAAGCGTTAATCGGAATTACTGGGCGTAAAGCGTGCGCAGGCGGTTCGGAAAGAAAGATGTGAAATCCCAGGGCTCAACCTTGGAACTGCATTTTTAACTACCGGACTAGAGTATGTCAGAGGGAGGTGGAATTCCACGTGTAGCAGTGAAATGCGTAGATATGTGGAGGAACACCGATGGCGAAGGCAGCCTCCTGGGATAATACTGACGCTCATGCACGAAAGCGTGGGG |
| P-asv4112 | GCAAGCGTTGTCCGGAATTATTGGGCGTAAAGCGCGCGCAGGCGGTCCTTTAAGTCTGATGTGAAAGCCCACGGCTCAACCGTGGAGGGTCATTGGAAACTGGAGGACTTGAGTACAGAAGAGGAAAGCGGAATTCCACGTGTAGCGGTGAAATGCGTAGAGATGTGGAGGAACACCAGTGGCGAAGGCGGCTTTCTGGTCTGTAACTGACGCTGAGGCGCGAAAGCGTGGGG |
| P-asv4113 | GCGAGCGTTATCCGGAATTACTGGGCGTAAAGGGTGCGTAGGCTGCGAGACAAGTCGGGTGTAAAAGGCACGGGCTCAACCTGTGTGTGCACTCGAAACTGTTTTGCTGGAGGGATAGAGAGGCAAGCGGAATTCCCGGTGTAGCGGTGAAATGCGTAGATATCGGGAAGAACACCAGTGGCGAAGGCGGCTTGCTAGCTATACCCTGACGCTGAGGCACGAAAGCCAGGGG |
| P-asv4114 | GCGAGCGTTAATCGGAATTACTGGGCGTAAAGCGAGCGCAGACGGTTACTTAAGCAGGATGTGAAATCCCCGGGCTCAACCTGGGAACTGCGTTCTGAACTGGGTGACTAGAGTGTGTCAGAGGGAGGTAGAATTCCACGTGTAGCAGTGAAATGCGTAGAGATGTGGAGGAATACCGATGGCGAAGGTAGCCTCCTGGGATAACACTGACGTTCATGCTCGAAAGCGTGGGT |
| P-asv4115 | TCAAGTGGTCGTCGCTTTTATTGGGCCTAAAACGTCCGTAGCCTGTTTGGTAAATCTGTGGGTAAATCAACCAGCTTAACTGGTTGAATTCTGCAGAGACTGCCAGACTAGGGACCGGGAGAGGTGTGGGGTACTCTCGGGGTAGGGGTAAAATCCTGTCATCCTGGGAGGACCACCAGTTGCGAAGGCGCCACACTGGAACGGATCCGACGGTCAGGGACGAAGCCTAGGGG |
| P-asv4116 | GCCAGCGTTGTTCGGAATTACTGGGTGTAAAGGGTTCGTAGGCGGTGCGGCAAGTTGGGAGTGAAATCTATGGGCTTAACCCAGAGGCTGCTTCCAAAACTGCTGTGCTTGAGTGTGAGAGAGGCTCGTGGAATTGCAGGTGTAGCGGTGAAATGCGTAGATATCTGCAGGAACACCCGTGGCGAAAGCGGCGAGCTGGATCACAACTGACGCTGAGGAACGAAAGCTAGGGG |
| P-asv4117 | GCAAGCGTTAATCGGAATTACTGGGCGTAAAGCATGCGCAGGCGGTTTTGTAAGACAGTGGTGAAATCCCCAGGCTCAACCTGGGAACTGCCATTGTGACTGCAAAGCTAGAGTGCGGCAGAGGGGGATGGAATTCCGCGTGTAGCAGTGAAATGCGTAGATATGCGGAGGAACACCGATGGCGAAGGCAATCCCCTGGGCCTGCACTGACGCTCATGCACGAAAGCGTGGGG |
| P-asv4118 | GCGAGCGTTGCCCGGAATTACTGGGCGTAAAGGGGGCGTAGGCGGTCAAATAAGTCAGATGTGAAATACTACTGCTCAACGGTAGAAATGCGTTTGAAACTGTAAGACTTGAGGGTACCAAAGGTAGACGGAATTACCTGAGTAGGGGTGAAATCCGTAGATACAGGTAAGAACGCCAGTGGGGAAGCCGGTCTACTGGGGTAACCCTGACGCTGAGGCCCGAAAGCTAGGGG |
| P-asv4119 | GCGAGCATTAATCGGACTTACTAGGCGTAAAGCGTGCGTAGGCGGCTTCTTAAGTCGGATGTGAAATCCCTGAGCTTAACTTAGGAATTGCATTCGATACTGGGAAGCTAGAGTATGGGAGAGGATGGTAGAATTCCAGGTGTAGCGGTGAAATGCGTAGAGATCTGGAGGAATACCGATGGCGAAGGCAGCCATCTGGCCTAATACTGACGCTGAGGTACGAAAGCATGGGG |
| P-asv4120 | GCAAGCGTTAATCGGATTTACTGGGCGTAAAGCGCGCGTAGGCGGCCAATTAAGTCAAATGTGAAATCCCCGAGCTTAACTTGGGAATTGCATTCGATACTGGTTGGCTAGAGTATGGGAGAGGATGGTAGAATTCCAGGTGTAGCGGTGAAATGCGTAGAGATCTGGAGGAATACCGATGGCAAAGGCAACCATCTGGCCTAACACTGACGCTGAGGTGCGAAAGCATGGGG |
| P-asv4121 | GCGAGCGTTAATCAGATTTACTGAGCGTAAAGCGTGCGTAGGCGGCTTTTTAAGTCGGATGTGAAATCCCTGAGCTTAACTTAGGAATTGCATTCGATACTGGGAAGCTAGAGTATGGGAGAGGATGGTAGAATTCCAGGTGTAGCGGTGAAATGCGTAGAGATCTGGAGGAATACCGATGGCGAAGGCAGCCATCTGGCCTAATACTGACGCTGAGGTACGAAAGCATGGGG |
| P-asv4122 | GCTAGCGTTGTTCGGAATCACTGGGCGTAAAGGGCGCGTAGGCGGCCATTTAAGTCGGGGGTGAAATCCCGGGGCTCAACCCCGGAATGGCCTTCGAGACTGGGTGGCTGGAGACCGGAAGAGGAGGGTGGAATACCCAGTGTAGAGGTGAAATTCGTAGATATTGGGTGGAACACCAGTGGCGAAAGCGGCTCTCTGGTCCGGGTCTGACGCTGAGGCGCGAAAGCGTGGGG |
| P-asv4123 | GCTAGCGTTGCTCGGAATCACTGGGCGTAAAGGGCGCGTAGGCGGCCATTCAAGTCGGGGGTGAAAGCCTGTGGCTCAACCACAGAATTGCCTTCGATACTGTTTGGCTTGAGTTTGGTAGAGGTTGGTGGAACTGCGAGTGTAGAGGTGAAATTCGTAGATATTCGCAAGAACACCAGTGGCGAAGGCGGCCAACTGGACCAATACTGACGCTGAGGCGCGAAAGCGTGGGG |
| P-asv4124 | GCAAGCGTTATCCGGAATTATTGGGCGTAAAGCGCGTGTAGGCGGCCCGTTAAGTCGACTGTGAAATCCCAGGGCTCAACCCTGGAACTGCAGTCGATACTGGCGGGCTTGAGGAAGGTAGAGGAGAGTGGAATTCCCGGTGTAGCGGTGGAATGCGCAGATATCGGGAGGAACACCGATGGCGAAGGCAGCTCTCTGGGCCTTACCTGACGCTGAGACGCGAAAGCATGGGT |
| P-asv4125 | GCGAGCGTTAATCGGAATTACTGGGCGTAAAGCGGGCGCAGACGGTTACTTAAGCGGGATGTGAAATCCCCGGGCTCAACCCGGGAATTGCGTTCCGAACTGGGTGGCTAGAGTGTGTCAGAGGGGGGTAGAATTCCACGTGTAGCAGTGAAATGCGTAGAGATGTGGAGGAATACCGATGGCGAAGGCAGCCCCCTGGGATAACACTGACGTTCATGCCCGAAAGCGTGGGT |
| P-asv4126 | GCAAGCGTTGTCCGGAATTATTGGGCGTAAAGAGCTCGTAGGCGGTTTGTCGCGTCTGCTGTGAAAACGCAAGGCTTAACCTTGCGCCTGCAGTGGGTACGGGCAGACTAGAGTGCGGTAGGGGAGACTAGAACTCCTGGTGTAGCGGTGGAATGCGCAGATATCAGGAGGAACACCGATGGCGAAGACAGGTCTCTGGGCCGTTACTGACGCTGAGGAGCGAAAGCATGGGG |
| P-asv4127 | GCTAGCGTTGTTCGGAATTACTGGGCGTAAAGCGCACGTAGGCGGCTATTTAAGTCAGGGGTGAAATCCCGGGGCTCAACCCCGGAACTGCCTCTGAAACTGGATAGCTAGAATCTTGGAGAGGTCAGTGGAATTCCGAGTGTAGAGGTGAAATTCGTAGATATTCGGAAGAACACCAGTGGCGAAGGCGACTGACTGGACAAGTATTGACGCTGAGGTGCGAAAGCGTGGGG |
| P-asv4128 | GCGAGCGTTAATCGGAATTACTGGGCGTAAAGCGCACGTAGGCGGCCAAGTAAGTTGGATGTGAAAGCCCCGGGCTTAACCTGGGAATTGCATTCGATACTGCTTGGCTGGAGTGTGGTAGAGGGAAGCGGAATTCCGGGTGTAGCGGTGAAATGCGTAGATATCCGGAGGAACACCAGTGGCGAAGGCGGCTTCCTGGACCAACACTGACGCTGAGGTGCGAAAGCGTGGGG |
| P-asv4129 | CCAAGCGTTGTCCGGATTTATTGGGTTTAAAGGGTGCGCAGGCGGCCCGATAAGTCAGTGGTGAAAGCCGACAGCTTAACTGTCGAAATGCCATTGATACTGTTGGGCTTGAGTATAGTTGAGGTTGGCGGAATGTAACATGTAGCGGTGAAATGCTTAGATATGTTACAGAACACCGATTGCGAAGGCAGCTGACTAAGCTATTACTGACGCTCATGCACGAAAGCGTGGGG |
| P-asv4130 | GCTAGCGTTATTCGGAATTACTGGGCGTAAAGCGCGCGTCGGCGGCTTGCCAAGTCAGGTGTGAAAGCCCGGGGCTTAACCCCGGAATTGCATTTGAAACTGGCAGGCTAGAGTCTGGTAGAGGGTGGTGGAATTCCACGTGTAGAGGTGAAATTCGTAGATATGTGGAGGAACACCGGTGGCGAAGGCGACCACCTGGACCAGTACTGACGCTATAGCGCGAAAGCGTGGGG |
| P-asv4131 | GCAAGCGTTGTTCGGATTTATTGGGCGTAAAGCGCGCGCAGGCGGATCAGCAAGTCAGATGTGAAATCTCAGGGCTCAACCCTGAAACTGCGTCTGAAACTGCTAGTCTAGAATGTCGGAGGGGGCAGGGGAATTTCACGTGTAGGGGTAAAATCCGTAGAGATGTGAAGGAACACCGGAGGCGAAGGCGCCTGCCTGGACGATACTTGACGCTGAGGCGCGAAAGCGTGGGG |
| P-asv4132 | GCAAGCGTTAATCGGAATTACTGGGCGTAAAGCGCGCGTAGGTGGTCTGATAAGTCAAATGTGAAAGCCCCGGGCTCAACCTGGGAATTGCATTCGATACTGTCGGACTAGAGTCTGGTAGAGGGTGGGGGAATTCCCGGTGTAGCGGTGAAATGCGTAGATATCGGGAGGAACACCAGTGGCGAAGGCGCCCACCTGGACCAAGACTGACACTGAGGTGCGAAAGCGTGGGG |
| P-asv4133 | GCAAGCGTTATCCGGAATTATTGGGCGTAAAGAGCTCGTAGGCGGTTTGTCGCGTCTGCTGTGAAATCCCGAGGCTCAACCTCGGGTCTGCAGTGGGTACGGGCAGACTAGAGTGCGGTAGGGGAGATTGGAATTCCTGGTGTAGCGGTGGAATGCGCAGATATCAGGAGGAACACCGATGGCGAAGGCAGATCTCTGGGCCGTAACTGACGCTGAGGAGCGAAAGCGTGGGG |
| P-asv4134 | GCGAGCGTTGTCCGGAATTACTGGGCGTAAAGGGTGCGTAGGCTGCGAGACAAGTCGGGTGTAAAAGGCACGGGCTCAACCTGTGTGTGCACTCGAAACTGTTTTGCTGGAGGGGTAGAGAGGCAAGCGGAATTCCCGGTGTAGCGGTGAAATGCGTAGATATCGGGAAGAACACCAGTGGCGAAGGCGGCTTGCTAGCTACACCCTGACGCTGAGGCACGAAAGCCAGGGG |
| P-asv4135 | GCAAGCGTTGTCCGGAATTATTGGGCGTAAAGCGCGCGCAGGCGGTCTTTAAAGTCTGATGTGAAATCTTGCGGCTTAACCGTAAGGGGTCATTGGAAACTGGAGGACTTGAGTGCAGAAGAGGAGAGTGGAATTCCACGTGTAGCGGTGAAATGCGTAGAGATGTGGAGGAACACCAGTGGCGAAGGCGACTCTCTGGTCTGTAACTGACGCTGAGGTGCGAAAGCGTGGGT |
| P-asv4136 | GCGAGCGTTAATCGGAATTACTGGGCGTAAAGCGTGCGCAGGCGGTTATGTAAGACAGATGTGAAATCCCCGGGCTCAACCTGGGAACTGCGTTTGTGACTGCATAACTAGAGTACGGCAGAGGGAGGTGGAATTCCACGTGTAGCAGTGAAATGCGTAGAGATGTGGAGGAACACCGATGGCGAAGGCAGCCTCCTGGGCCAGTACTGACGCTCATGCACGAAAGCGTGGGG |
| P-asv4137 | GCAAGCGTTAATCGGAATTACTGGGCGTAAAGCGTGCGCAGGCGGTTGTGCAAGACAGATGTGAAATCCCCGGGCTTAACCTGGGAATGGCATTTGTGACTGCACGGCTAGAGTGTGTCAGAGGGGGGTAGAATTCCACGTGTAGCAGTGAAATGCGTAGAGATGTGGAGGAATACCGATGGCGAAGGCAGCCCCCTGGGATAACACTGACGCTCATGCACGAAAGCGTGGGG |
| P-asv4138 | GCAGGCGTTAATCGGAATTACTGGGCGTAAAGCGTGCGCAGGCGGTTTGTTAAGACAGGTGTGAAATCCCCGGGCTTAACCTGGGAATTGCGCTTGTAACTGGCAAGCTAGAGTGTGGCAGAGGGGGGTGGAATTCCACGTGTAGCAGTGAAATGCGTAGATATGTGGAGGAACACCGATGGCGAAGGCAGCCCCCTGGGTTAACACTGACGCTCATGCACGAAAGCGTGGGG |

| | |
|---|---|
| P-asv4139 | GCAAGCGTTACTCGGAATTACTGGGCGTAAAGCGTGCGTAGGTGGTGATTTAAGTCCGTTGTGAAAGCCCTGGGCTCAACCTGGGAATTGCAGTGGATACTGGGTCACTAGAGTGTGGTAGAGGGTGGCGGAATTCCCGGTGTAGCAGTGAAATGCGTAGAGATCGGGAGGAACATCCGTGGCGAAGGCGGCCACCTGGGCCAACACTGACACTGAGGCACGAAAGCGTGGGG |
| P-asv4140 | GCAAGCGTTAATCGGAATTACTGGGCGTAAAGCGTGCGTAGGTGGTTTGTTAAGTCTGATGTGAAAGCCCTGGGCTCAACCTGGGAATGGCATTGGATACTGGCTTACTAGAGTGCGGTAGAGGGTAGCGGAATTTCCGGTGTAGCGGTGAAATGCGTAGAGATCGGAAGGAACATCCGTGGCGAAGGCGGCTACCTGGACCAGCACTGACACTGAGGCACGAAAGCGTGGGG |
| P-asv4141 | GCAAGCGTTAATCGGATTTACTGGGCGTAAAGCGTGCGTAGGCGGCTATTTAAGTCGGATGTGAAAGCCCTGGGCTTAACCTGGGAACTGCATTCGATACTGGGTAGCTGGAGATCGGTAGAGGGCGGCGGAATTCCGGGTGTAGCGGTGAAATGCGTAGATATCCGGAGGAACACCGATGGCGAAGGCAACCGCCTGGGCCTGATCTGACGCTGAGGCACGAAAGCGTGGGG |
| P-asv4142 | GCTAGCGTTGTTCGGAATCACTGGGCGTAAAGGGCGCGTAGGCGGCTTTATAAGTCGGGGTGAAAGCCTGTGGCTCAACCACAGAATTGCCTTCGATACTGTATGGCTTGAGACCGGAAGAGGTAAGTGGAACTGCGAGTGTAGAGGTGAAATTCGTAGATATTCGCAAGAACACCAGTGGCGAAGGCGGCTTACTGGTCCGGATCTGACGCTGAGGCGCGAAAGCGTGGGG |
| P-asv4143 | GCAAGCGTTGTCCGGATTTATTGGGTTTAAAGGGTGCGTAGGCGGCCCTGTAAGTCAGTGGTGAAAGCCCATAGCTCAACTATGGAACTGCCATTGATACTGCAGGGCTTGAGTACAGACGAGGTTGGCGGAATTGATAGTGTAGCGGTGAAATGCATAGATACTATCAAGAACACCGATTGCGAAGGCAGCTAACTAGGCTGTAACTGACGCTGAGGCACGAAAGCGTGGGG |
| P-asv4144 | GCAAGCGTTATCCGGATTTACTGGGTTTAAAGGGTGTGTAGGCGGATATTTAAGTCAGAGGTGAAATCTCCGAGCTCAACTCGGAAACTGCCTTTGATACTATTTATCTTGAATTTCGTTGAGGTTGGCGGAATATATCATGTAGCGGTGAAATGCATAGATATGATATAGAACACCGATTGCGTAGGCAGCTGGCTGAACGAACATTGACGCTGAGGCACGAAAGCGTGGGT |
| P-asv4145 | GCAAGCGTTATCCGGAATTATTGGGCGTAAAGAGCTCGTAGGCGGTTTGTCGCGTCTGCCGTGAAAGTCCGGGGCTCAACCCCGGATCTGCGGTGGGTACGGGCAGACTAGAGTGATGTAGGGGAGACTGGAATTCCTGGTGTAGCGGTGAAATGCGCAGATATCAGGAGGAACACCGATGGCGAAGGCAGGTCTCTGGGCATTAACTGACGCTGAGGAGCGAAAGCATGGGG |
| P-asv4146 | GCAAGCGTTGTCCGGAATTATTGGGCGTAAAGCGCGCGCAGGCGGTTCCTTAAGTCTGATGTGAAAGCCCCCGGCTCAACCGGGGAGGGTCATTGGAAACTGGGGAACTTGAGTACTGAAGAGGAAAGCGGAATTCCACGTGTAGCGGTGAAATGCGTAGAGATGTGGAGGAACACCAGTGGCGAAGGCGGCTTTCTGGTCAGTAACTGACGCTGAGGCGCGAAAGCGTGGGG |
| P-asv4147 | GCGAGCGTTGTCCGGAATTATTGGGCGTAAAGGGCTCGTAGGCGGTTTGTCGCGTCGGGAGTGAAAACTCGGGGCTTAACCCTGAGCCTGCTTCCGATACGGGCAGACTAGAGGTATGCAGGGGAGAACGGAATTCCTGGTGTAGCGGTGAAATGCGCAGATATCAGGAGGAACACCGGTGGCGAAGGCGGTTCTCTGGGCATTACCTGACGCTGAGGAGCGAAAGTGTGGGG |
| P-asv4148 | GCGAGCGTTAATCGGAATTACTGGGCGTAAAGCGCGCGTAGGTGGCTTGATAAGCCGGTTGTGAAAGCCCCGGGCTCAACCTGGGAACGGCATCCGGAACTGTCAGGCTAGAGTGCAGGAGAGGAAGGTAGAATTCCCGGTGTAGCGGTGAAATGCGTAGAGATCGGGAGGAATACCAGTGGCGAAGGCGGCCTTCTGGACTGGCACTGACACTGAGGTGCGTAAGCGTGGGT |
| P-asv4149 | GCAAGCGTTGTCCGGAATTATTGGGCGTAAAGCGCGCGCAGGCGGTCCTTTAAGTCTGATGCGAAAGCCCACGGCTCAACCGTGAAGGGTCATTGGAAACTGGGGACTTGAGTGCAGAAGAGGAAAGTGGAATTCCATGTGTAGCGGTGAAATGCGTAGAGATGTGGAGGAACACCAGTGGCGAAGGCGACTTTCTGGTCTGTAACTGACGCTGAGGCGCGAAAGCGTGGGG |
| P-asv4150 | GCTAGCGTTGCTCGGAATCACTGGGCGTAAAGCGCACGTAGGCGGCTTTCTAAGTCAGGGGTGAAATCCTGGAGCTCAACTCCAGAACTGCCTTTGATACTGGAAAGCTTGAGTTCGAGAGAGGTGAGTGGAACTGCGAGTGTAGAGGTGAAATTCGTAGATATTCGCAAGAACACCAGTGGCGAAGGCGGCTCACTGGCTCGATACTGACGCTGAGGTGCGAAAGCGTGGGG |
| P-asv4151 | GCAAGCGTTGTCCGGAATTATTGGGCGTAAAGAGCTCGTAGGCGGTTTGTCGCGTCTGCTGTGAAAACGCAAGGCTTAACCTTGCGCCTGCAGTGGGTACGGGCGGACTAGAGTGCGGTAGGGGAGACTGGAATTCCTGGTGTAGCGGTGGAATGCGCAGATATCAGGAGGAACACCGATGGCGAAGGCAGGTCTCTGGGCCGTTACTGATGCTGAGGAGCGAAAGCATGGGG |
| P-asv4152 | GCGAGCGTTAATCGGAATTACTGGGCGTAAAGCGTGCGCAGGCGGTTATGTAAGACAGATGTGGAAGCCCCGGGCTCAACCTGGGAACTGCGTTTGTGACTGCATAACTAGAGTACGGCAGAGGGGGGTGGAATTCCGCGTGTAGCAGTGAAATGCGTAGAGATGCGGAGGAACACCGATGGCGAAGGCAGCCCCCTGGGCCAGTACTGACGCTCATGCACGAAAGCGTGGGG |
| P-asv4153 | GCTAGCGTTGTTCGGAATTACTGGGCGTAAAGCGCACGTAGGCGGCTTCCTAAGTCAGGGGTGAAAGCCTGGAGCTCAACTCCAGAACTGCCCTTGAGACTGGGTCGCTCGAATCCGGGAGAGGTGAGTGGAATTCCGAGTGTAGAGGTGAAATTCGTAGATATTCGGAAGAACACCAGTGGCGAAGGCGGCTCACTGGACCGGTATTGACGCTGAGGTGCGAAAGCGTGGGG |
| P-asv4154 | GCCAGCGTTGCTCGGAATTACTGGGCGTAAAGGGTTCGTAGGCGGTGCGGCAAGTCGGGAGTGAAATCTCTGGGCTTAACCCAGAGGCTGCTTCCGAAACTGCCGTGCTAGAGTGCGGGAGAGGCGAGTGGAATTACAGGTGTAGCGGTGAAATGCGTAGATATCTGTAGGAACACCCGTGGCGAAGGCGGCTCGCTGGACCGCAACTGACGCTGAGGAACGAAAGCTAGGGG |
| P-asv4155 | GCAAGCGTTGTTCGGATTTACTAGGCGTACAGGGAGCGTAGGCGGTTCGGTAAGCCCTTTGTGAAATCTCCAGGCTTAACCTGGAACAGTCGGAGGGGACTGCTGGGCTAGAGGACGGGAGAGGAGCGCGGAATTCCCGGTGTAGCGGTGAAATGCGTAGAGATCGGGAGGAAGGCCGGTGGCGAAGGCGGCGCTCTGGAACGTTTCTGACGCTGAGGCTCGAAAGCGTGGGG |
| P-asv4156 | GCAAGCGTTGTTCGGAATCACTGGGCGTAAAGGGCGTGTAGGCGGTCAGGTAAGTGGGATGTGAAATGCCCCGGCTCACCCGGGGATCTGCATTCCAAACTGCTTGGCTTGAGTATGGGAGAGGATGAGGGAATTCCTGGTGTAGCGGTGAAATGCGTAGATATCAGGAGGAACACCGGTGGCGAAGGCGCTCATCTGGCCCAATACTGACGCTGAGGCGCGAAAGCTAGGGG |
| P-asv4157 | GCAAGCGTTGTTCGGAATTACTGGGCGTAAAGGGCGCGTAGGCGGCCTCCTAAGTCGAACGTGAAATCCCCGGGCTCAACCCGGGAACTGCGTTCGAGACTGGTTGGCTTGAATCCGGGAGAGGGATGCGGAATTCCAGGTGTAGCGGTGAAATGCGTAGATATCTGGAGGAACACCGGTGGCGAAGGCGGCATCCTGGACCGGCATTGACGCTGAGGCGCGAAAGCCAGGGG |
| P-asv4158 | TCAAGCGTTGTTCGGATTCACTGGGCGTAAAGGGTGCGTAGGTGGTGGGGTAAGTCGGATGTGAAATCTCCGAGCTCAACTCGGAAATGGCATTGGAAACTACCCTGCTCGAGGGTCGGAGGGGGGACTGGAATTCTCGGTGTAGCAGTGAAATGCGTAGATATCGAGAGGAACACCAGTGGCGAAGGCGAGTCCCTGGACGACTCCTGACACTGAGGCACGAAAGCTAGGGG |
| P-asv4159 | CCCAGCGTTGTTCGGAATTACTGGGTGTAAAGGGTCAGTAGGCGGTGTGGCAAGTTGGGAGTGAAATCTATGGGCTCAACCCATAGGCTGCTTCCAAAACTGCTGTGCTTGAGTGTGGGAGAGGCGCGTGGAATTGCAGGTGTAGCGGTGAAATGCGTAGATATCTGCAGGAACACCCGTGGCGAAAGCGGCGCGCTGGACCACAACTGACGCTGAATGACGAAAGCTAGGGG |
| P-asv4160 | GCAAGCGTTGTTCGGAATTATTGGGCGTAAAGGGTGCGTAGGTGGTACGGCAAGTCTTGTGTGAAATCTTCAGGCTCAACTTGAAGTCTGCACAAGAAACTGCCGGGCTTGAGTATGGGAGAGGTGGGTGGAATTTCCGGTGTAGCGGTGAAATGCGTAGATATCGGAAGGAACACCTGTGGCGAAAGCGGCTCACCGGACCATTACCGACACTGAGGCACGAAAGCTAGGGG |
| P-asv4161 | GCTAGCGTTGTTCGGAATCACTGGGCGTAAAGCGCGTGTAGGGGCCCATTAAGTCTGATGTGAAAGCCCGGGGCTCACCCCCGGAAGTGCATTGGAAACTGATGGGCTAGAGTCCCGGAGAGGAGGGTGGAATTCCTGGTGTAGCGGTGAAATGCGTAGATATCAGGAGGAACACCGGTGGCGAAGGCGGCCCTCTGGACGGTGACTGACGCTGAGACGCGAAAGCGTGGGG |
| P-asv4162 | GCCAGCGTTGTTCGGAATCATTGGGCGTAAAGGGCGCGTAGGCGGCCACGCAAGTCGGATGTGAAATCCCTCGGCTCAACCGAGGAACTGCACCCGAAACTGCGAGGCTCGAGTCCTGGAGAGGGTAGTGGAATTCCCAGTGTAGCGGTGAAACGCGTAGATACTGGGAGGAACACCGGTGGCGAAGGCGGCTACCTGGACAGGTACTGACGCTGAGGCGCGAAAGCCAGGGG |
| P-asv4163 | GCCAGCGTTGTTCGGAATCATTGGGCGTAAAGGGCGCGTAGGCGGCCGATCAAGTCGGAGGTGAAATCCCTCGGCTCAACTGAGGAATTGCCTCCGATACTGTTCGGCTTGAGTCCTGGAGAGGGTAGTGGAATTCCGAGTGTAGCGGTGAAGTGCGTAGAGACTCGGAAGAACACCAGTGGCGAAGGCGGCTACCTGGACAGGTACTGACGCTGAAGCGCGAAAGCTAGGGG |
| P-asv4164 | GCAAGCGTTGCCCGGATTTATTGGGCGTAAAGAGCGTGTAGGCGGCCAGACAGGTCCGTTGTGAAAACTAGAGGCCTAACCTCTAGACGTCGATGGAAACCGTCTGGCTAGAGTCCGGAAGAGGAGAATGGAATTCCTGGTGTAGCGGTGAAATGCGCAGATATCAGGAAGAACACCCGTGGCTAAGGCGGTTCTCTAGTACGGTACTGACGCTGAGACGCGAAAGCGTGGGG |
| P-asv4165 | GCGAGCGTTGTCCGGATTTATTGGGCGTAAAGAGCTCGTAGGCGGCTTGTCGCGTCGACTGTGAAAACCCGCGGCTCAACCGCGGGCTTGCAGCCGATACGGGCAGGCTAGAATTCGGTAGGGGAGACTGGAATTCCTGGTGTAGCGGTGAAATGCGCAGATATCAGGAGGAACACCGGTGGCGAAGGCGGGTCTCTGGGCCGATATTGACGCTGAGGAGCGAAAGCGTGGGG |
| P-asv4166 | GCAAGCGTTGTCCGGATTTATTGGGCGTAAAGAGCGTGTAGGCGGCCAGGTAGGTGGGTTGTGAAAACTGGAGGCTCAACCTTCAGACGTCGACCCAAACCCCCTGGCTAGAGTCTGGAAGAGGAGAGTGGAATTCCTGGTGTAGCGGTGAAATGCGCAGATATCAGGAAGAACACCCGTGGCGGAGGCGGCTCTCTGGGACGGTACTGACGCTGAGACGCGAAAGCGTGGGG |
| P-asv4167 | CCAAGCGTTGTTCGGAATCATTGGGCGTAAAGCGTGTGCAGGTGGTTTGGTAAGTTGGATGTGAAAGCCCCGGGCTTAACCCGGGAAGTGCATTCAAAACTACCAAACTTGAGTATGGAAGAGGGTCGTGGAATTCCAGGTGTAGTGGTGAAATACGTAGATATCTGGAGGAACACCGGTGGCGAAGGCGGCGACCTGGTCCTATACTGACGCTGAGGTGCGAAAGCGTGGGG |
| P-asv4168 | GCAAACGTTGTTCGGAATCACTGGGCGTAAAGGGCGTGTAGGCGGCTGAATAAGTGGGATGTTAAATCCCCCGGCTCAACCGGGGACCTGCATCTCATACTGTTCAGCTCGAGCACATCAGAGGAGACTGGAATTCCCGGTGTAGCGGTGAAATGCGTAGATATGTGGCGGAACACCGATGGCGAAGGCAGCCCCCTGGGCCAAGATTGACGCTCAGGCACGAAAGCGTGGGG |
| P-asv4169 | GCAAGCGTTGTTCGGAATTACTGGGCGTAAAGGGCGCGTAGGCGGCCACCTAAGTCAGACGTGAAATCCCTCGGCTCAACCGGGGAACTGCGTCTGATACTGGGCGGCTTGAATTCGGGAGAGGAATGCGGAATTCCAGGTGTAGCGGTGAAATGCGTAGATATCTGGAGGAACACCGGTGGCGAAGGCGGCATTCTGGACCGACATTGACGCTGAGGCGCGAAAGCCAGGGG |
| P-asv4170 | GCAAACGTTGTTCGGATTCACTGGGCTTAAAGGGCGCGTAGGCGGGGGTTCAAGTCGGTGGTGAAATCTTTCGGCCCAACCGGAAAATCGCCTCCGATACTGGATCTCTTGAGGGGGGTAGGGGCATGCGGAACTTCGGGTGGAGCGGTGAAATGCGTAGATATCCGAAGGAACGCCGGTGGCGAAAGCGGCGTGCTGGACCCTTTCTGACGCTGAGACGCGAAAGCGTGGGG |
| P-asv4171 | GCAAGCGTTGTTCGGAATTATTGGGCGTAAAGGGCGCGTAGGCGGTGCGGTAAGTCTTCTGTGAAATCTCCGGGCTCAACTCGGAGCCTGCAGAGGAAACTGCCGTGCTGGAGTGTGGGAGAGGAGAGTGGAATTCCCGGTGTAGCGGTGAAATGCGTAGATATCTGCAGGAACACCCGTGGCGAAAGCGGCGCGCTGGACCACTACTGACGCTGAGGAACGAAAGCTAGGGG |
| P-asv4172 | GCGAGCGTTAATCGGATTTACTGGGCGTAAAGCGTGTGTAGGCGGCTTGGCAAGTTGGATGTGAAAGCCCACGGCTCAACCGTGGAAGCGCATCCAAAACTGCTTCGCTCGAGTCTCGGAGAGGAAGGTGGAATGCTTGGTGTAGAGGTGAAATTCGTAGATATCAAGCAGAACACCTGCGGCGAAGGCGGCCTTCTGGACGATGACTGACGCTGAGACACGAAAGCGTGGGG |
| P-asv4173 | GCTAGCGTTGTTCGGAATTACTGGGCGTAAAGCGCACGTAGGCGGATTGTTAAGTCAGAGGTGAAATCCCGGAGCTCAACTTCGGAACTGCCTTTGATACTGGCAATCTAGAGTCCGGAAGAGGTTAGTGGAATTCCCAGTGTAGCGGTGAAATGCGTAGATATTGGGAGGAACACCGGTGGCGAAGGCGGCCATCTGGACCGGTCCTGACGCTGAGGCGCGAAAGCGTGGGG |
| P-asv4174 | CCAAGCGTTAGGCGGAATCACTGGGCTTAAAGCGTGTGTAGGTGGATGCCTAAGTACCTTGTGAAATCCCACGGCTCAACCGTGGAACTGCTGGGTATACTGGGCGTCTTGGGTCACCTAGGGGTTGCCGGAACAAATGGTGGAGCGGTGAAATGCGTAGATATCATTTGGAACGCCAATGGCGAAAGCAGGCAACTGGGGGTGCACCGACACTGAGACACGAAAGCCAGGGG |
| P-asv4175 | GCAAACGTTGTTCGGAACTACTGGGCGTAAAGCGCGTGTAGGCGGCAGGGTGTGTCGGATGTGAAATCCCGTGGCTCAACCACGGAAGTGCACCCGAAACTACCCAGCTCGAGTCTCGGAGAGGAAGGTGGAATGCTTGGTGTAGAGGTGAAATTCGTAGATATCAAGCGGAACACCGGTGGCGAAGGCGGCCTTCTGGACGATGACTGACGCTGAGACGCGAAAGCGTGGGG |
| P-asv4176 | GCAAGCGTTGTCCGGATTTACTGGGCGTAAAGGGCGTGCAGGCGGTCTGTTAAGTCTAGGGTGAAGGCTCCCGGCTCAACTGGGAGAGGTCCTTGGATACTGGCAGACTCGAGGAGGGTAGAGGAGAGTGGAATTCCCGGTGTAGTGGTGATATGCGTAGATATCGGGAGGAACACCAGTGGCGAAGGCGGCTCTCTGGGCCCTACCTGACGCTGAGACGCGAAAGCGTGGGG |
| P-asv4177 | GCCAGCGTTGCTCGGAATTACTGGGCGTAAAGCGTGCGTAGGTGGCTTGTTAAGTCTGCTGTGAAAGCCCCGGGCTCAACCTGGGAATGGCAGTGGATACTGGCAAGCTAGAGTGCGGTAGAGGGTAGTGGAATTCCCGGTGTAGCGGTGAAATGCGTAGAGATCGGGAGGAACACCAGTGGCGAAGGCGGCTACCTGGACCAGCACTGACACTGAGGCACGAAAGCGTGGGG |

| | |
|---|---|
| P-asv4178 | GCTAGCGTTGCTCGGAATCACTGGGCGTAAAGCGCACGTAGGCGGACTCTTAAGTCAGGGGTGAAATCCTGGAGCTCAACTCCAGAACTGCCCTTGATACTGGGAGCCTTGAGTTCGGGAGAGGTGAGTGGAACTGCGAGTGTAGAGGTGAAATTCGTAGATATTCGCAAGAACACCAGTGGCGAAGGCGGCTCACTGGCCCGATACTGACGCTGAGGTGCGAAAGCGTGGGG |
| P-asv4179 | GCAAGCGTTGCTCGGAATCACTGGGCGTAAAGGGCGCGTAGGCGGGATGGCAAGTCGGTCGTGAAATCCCTCGGCTCAACTGAGGAGAGTCGGCCGATACTGCTGTTCTTGAGGGATGCAGAGGGGACTGGAATTCCCGGTGTAGCGGTGAAATGTGTAGAGATCGGGAGGAACACCGGTGGCGAAGGCGGGTCTCTGGGCATCTCCTGACGCTGAGGCGCGAAAGCGTGGGG |
| P-asv4180 | GCCAGCGTTGTTCGGAATTATTGGGCGTAAAGGGCGCGTAGGCGGTCGAGTAAGTCTGACGTGAAATCCCTTGGCTCAACCAAGGAACTGCGTCTTAAACTGCTCGGCTTGAGTCCGGGAGAGGGTAGTGGAATTCCCAGTGTAGCGGTGAAATGCGTAGATATTGGGAGGAACACCGGTGGCGAAGGCGGCTACCTGGACCGGTACTGACGCTGAGGCGCGAAAGCTAGGGT |
| P-asv4181 | GCGAACGTTGTTCGGAATCACTGGGCTTAAAGGGCGCGTAGGCGGGTTTTCAAGTCTGTGGTGAAATCCTCCAACTAAACTGGAGAAGTGCCGTGGATACTGGAGACCTCGAGGAGGGTAGGGGCATCTGGAACAGCCGGTGGAGCGGTGAAATGCGTTGATATCGGCTGGAACTCCGATGGCGAAGGCAAGGTGCTGGACCCTCTCTGACGCTGAGGCGCGAAAGCCAGGGG |
| P-asv4182 | GCTAGCGTTGCTCGGAATTACTGGGCGTAAAGCGCACGTAGGCGGATTGTTAAGTCGGGGGTGAAATCCTGGAGCTCAACTCCAGAACTGCCTTCGATACTGGCGATCTCGAGTTCGGGAGAGGTGAGTGGAACTGCGAGTGTAGAGGTGAAATTCGTAGATATTCGCAAGAACACCAGTGGCGAAGGCGGCTCACTGGCCCGATACTGACGCTGAGGTGCGAAAGCGTGGGG |
| P-asv4183 | GCTAGCGTTGTTCGGAATCACTGGGCGTAAAGGGCGCGTAGGCGGCCAGGCAAGTCGGCTTGTGAAAGCCCGGGGCTCAACCCCGGAGGTGCGGCCGGGACTGCTTGGCTGGAGGCCGGAAGAGGAGGGTGGAATACCCAGTGTAGAGGTGAAATTCGTAGATATTGGGTGGAACACCGGTGGCGAAAGCGGCTCTCTGGTCCGGTCCTGACGCTGAGGCGCGAAAGCGTGGGG |
| P-asv4184 | GCAAGCGTTGTTCGGAATTACTGGGCGTAAAGGGCGCGTAGGCGGTCTTGTCAGTCCCTTGTGAAATCCCCAGGCTCAACTGGGGAACTGCACGGAAACAGCAGGGCTTGAGTTCGGGAGAGGGAAGCGGAATTCCGGGTGTAGCGGTGAAATGCGTAGATATCCGGAGGAACACCGGTGGCGAAGGCGGCTTCCTGGACCGATACTGACGCTGAGGCGCGAAAGCTAGGGG |
| P-asv4185 | CCGAGTTGTCGGGACGATTATTGGGCCTAAAGCATCCGTAGCCTGTTCTGCAAGTCCTCCGTTAAATCCACCTGCTCAACGGGTGGGCTGCGGAGGATACCGCAGAGCTAGGAGGCGGGAGAGGCAAACGGTACTCAGTGGGTAGGGGTAAAATCCATTGATCTACTGAAGACCACCAGTGGCGAAGGCGGTTTGCCAGAACGCGCTCGACGGTGAGGGATGAAAGCTGGGGG |
| P-asv4186 | ACAAGCGTTGTCCGGATTTACTGGGCGTAAAGAGCGCGCAGGCGGTCGATTTAGTCGAGTGTGAAAGCCCCCGGCTCAACTGGGGAGGGTCACTCGATACTGATCGACTCGAAGGCGGGAGAGGGAAGCGGAATTCCCGGTGTAGTGGTGAAATGCGTAGATATCGGGAGGAACACCAGTGGCGAAGGCGGCTTCCTGGCCCGTTCTTGACGCTGAGGCGCGAAAGCTAGGGG |
| P-asv4187 | GCGAGCGTTAATCGGAATTACTGGGCGTAAAGCGTGCGCAGGCGGTTCGCTGTGTCCTGTGTGAAAGCCCCGGGCTCAACCTGGGAATGGCATGGGAAACTGGCGAGCTGGAGTGTGGCAGAGGGGGGTGGAATTCCGCGTGTAGCAGTGAAATGCGTAGAGATGCGGAGGAACACCGATGGCGAAGGCAGCCCCCTGGGCCAACACTGACGCTCAGGCACGAAAGCGTGGGG |
| P-asv4188 | GCGAGCGTTGTCCGGATTTATTGGGCGTAAAGAGCTCGTAGGCGGCTTGTCGCGTCGACTGTGAAAACCTGCGGCTCAACCGTGTGGTCTGCAGTCGATACGGGCAGGCTAGAGTTCGGTAGGGGAGACTGGAATTCCTGGCGTAGCGGTGGAATGCGCAGATATCAGGAGGAACACCGGTGGCGAAGGCGGGTCTCTGGGCAGTAACTGACGCTGAGGAGCGAAAGCGTGGGG |
| P-asv4189 | GCAAGCGTTGTCCGGAATTATTGGGCGTAAAGAGCTCGTAGGCGGTTCGTCGCGTCGGCTGTGAAAACCCGGAGCTCAACTCCGGGCCTGCAGTCGATACGGGCGGACTGGAGTTCGGCAGGGGAGACTGGAACTCCTGGTGTAGCGGTGAAATGCGCAGATATCAGGAGGAACACCGGTGGCGAAGGCGGGTCTCTGGGCCGATACTGACGCTGAGGAGCGAAAGCGTGGGG |
| P-asv4190 | GCAAGCGTTGTTCGGGATTATTGGGCGTAAAGGGTGCGTAGGCGGTGCGGTAAGTCTGTAGTGAAATCTCTGGGCTCAACTCAGAGTCTGCTATAGAAACTGCCGTGCTTGAGTGTGGGAGAGGTGAGTGGAATTCCCGGTGTAGCGGTGAAATGCGTAGATATCGGGAGGAACACCTGTGGCGAAAGCGGCTCACTGGACCACAACTGACGCTGATGCGCGAAAGCTAGGGG |
| P-asv4191 | GCGAGCGTTGTCCGGAGTTACTGGGCGTAAAGGGCATGCAGGCGGTCGTACGCGTTTCGGGTGACAGCGGGTGGCTTAACTACCCGAGTGTTCGAAAGACGGTACGACTTGAGGGCTCGAGAGGGACACGGAATTCCGGGTGGAGTGGTGAAATGCGTAGATATCAGGAGGAACACCGGTGGCGAAGGCGGCCCTCTGGACGGTGACTGACGCTGAGACGCGACAGCGTGGGG |
| P-asv4192 | GCAAGCGTTACTCGGAATTACTGGGCGTAAAGCGTGCGTAGGCGGTTTGTTAAGTCTGATGTGAAAGCCCCGGGCTCAACCTGGGAATGGCATTGGATACTGGCAGGCTAGAATGTGGTAGAGGGTGGTGGAAACCCCGGTGTAGCGGTGAAATGCGTAGAGACCGGGAGGAACATCCGTGGCGAAGGCGGCCACCTGGACCAACATTGACGCTGAGGCACGAAAGCGTGGGG |
| P-asv4193 | GCAAGCGTTGTCCGGATTTATTGGGCGTAAAGCGAGCGCAGGCGGTTCTTTAAGTCTGATGTGAAAGCCCCCGGCTCAACCGGGGAGGGTCATTGGAAACTGGAGGACTTGAGTGCAGAAGAGGAGAGTAGAATTCCATGTGTAGCGGTGAAATGCGTAGATATATGGAGGAACACCAGTGGCGAAGGCGGCTCTCTGGTCTGTAACTGACGCTGAGGCTCGAAAGCGTGGGG |
| P-asv4194 | GCAAGCGTTGTCCGGAATTATTGGGCGTAAAGAGCGTGTAGGCGGCTCGGTGAGTCCGCCGTGAAAATCCGAGGCTCAACCTCGGGCTTGCGGTGGATACTGTCGGGCTAGAGGATGGTAGAGGGGAGTGGAATTCCCGGTGTAGCGGTGAAATGCGCAGATATCGGGAGGAACACCAGTAGCGAAGGCGGCTCCCTGGGCCATTCCTGACGCTGAGACGCGAAAGCTAGGGG |
| P-asv4195 | GCAAGCGTTATTCGGATTTACTGGGCGTACAGGGAGCGTAGGCGGTTCGGTAAGCCCTCTGTGAAATCTTCAGGCTTAACCTGGAAAAGTCGGAGGGGACTGCTGGGCTAGAGGACGGGAGAGGAGCGCGGAATTCCCGGTGTAGCGGTGAAATGCGTAGAGATCGGGAGGAAGGCCGGTGGCGAAGGCGGCGCTCTGGAACGTTTCTGACGCTGAGGCTCGAAAGCGTGGGG |
| P-asv4196 | GCAAGCGTTAATCGGATTTACTGGGCGTAAAGCGCGCGTAGGCGGCTAATTAAGTCAAATGTGAAATCCCCGAGCTTAACTTGGGAATTGCATTCGATACTGGTTAGCTAGAGTGTGGGAGAGGATGGTAGAATTCCAGGTGTAGCGGTGAAATGCGTAGAGATCTGGAGGAATACCGATGGCGAAGGCAGCCATCTGGCCTAACACTGACGCTGAGGTACGAAAGCATGGGG |
| P-asv4197 | GCAAGCGTTGTCCGGAATTATTGGGCGTAAAGCGCGCGCAGGCGGTCTTTTAAGTCTGATGTGAAAGCCCACGGCTCAACCGTGGAAGGTCATTGGAAACTGAAAGACTTGAGTACAGAAGAGGAAAGCGGAATTCCACGTGTAGCGGTGAAATGCGTAGAGATGTGGAGGAACACCAGTGGCGAAGGCGGCTTTCTGGTCTGTAACTGACGCTGAGGCGCGAAAGCGTGGGG |
| P-asv4198 | GCAAGCGTTATCCGGAATTATTGGGCGTAAAGAGCTTGTAGGCGGTTCGTCGCGTCTGGTGTGAAAGCCCATCGCTTAACGGTGGGTCTGCGCCGGGTACGGGCGGGCTAGAGTGCAGTAGGGGAGACTGGAATTCCCGGTGTAACGGTGGAATGTGTAGATATCGGGAAGAACACCAATGGCGAAGGCAGGTCTCTGGGCTGTTACTGACGCTGAGAAGCGAAAGCGTGGGG |
| P-asv4199 | GCAAGCGTTGTTCGGAATTACTGGGCGTAAAGCGTGCGTAGTCGGTATTGAGAGTCACGGGTGAAATCCCAGGGCTTAACCCTGGAACTGCCTGTGAGACCTCAGTACTAGAGTGTGAGAGGGGATAGTGGAATACCCAGTGTAGCGGTGAAATGCGTGGAGATTGGGTAGAACACCGGTGGCGAAGGCGACTATCTGGCTCACAACTGACGATCAGGCACGAAAGCGTGGGG |
| P-asv4200 | GCTAGCGTTGTTCGGAATTACTGGGCGTAAAGCGCACGTAGGCGGCCATTCAAGTCAGAGGTGAAAGCCCGGGGCTCAACCCCGGAACTGCCTTTGAATCTAGATGGCTTGAATCTTGGAGAGGCGAGTGGAATTCCGAGTGTAGAGGTGAAATTCGTAGATATTCGGAAGAACACCAGTGGCGAAGGCGACTCGCTGGGCAAGTATTGACGCTGAGGTGCGAAAGCGTGGGG |
| P-asv4201 | GCAAGCGTTGTCCGGATTTACTGGGCGTAAAGAGCGTGTAGGCGGCCTGACAGGTCTGTTGTGAAAACTGGAGGCTCAACCTCCAGACGTCGACAGAAACCGTTTGGCTAGAGTCCGGAAGAGGAGAGTGGAATTCCTGGTGTAGCGGTGAAATGCGCAGATATCAGGAAGAACACCGATGGCGAAGGCAGCTCTCTGGGACGGTACTGACGCTGAGACGCGAAAGCGTGGGG |
| P-asv4202 | GCAAGCGTTGTTCGGAATCATTGGGCGTAAAGGGCGTGTAGGCGGTCTGTTAAGTCATGTGTGAAATCCCTCGGCTCAACCGGGGAAGGACGCATGAAACTGACAAGCTAGAGTACCAAAGAGGGGGGTGGAATTCCCGGTGTAGCGGTGAAATGCGTAGATATCGGGAGGAACACCGGTGGCGAAGGCGGCCCCCTGGTTGGATACTGACGCTGAGACGCGAAAGCGTGGGG |
| P-asv4203 | GCAAGCGTTAATCGGAATTACTGGGCGTAAAGCGCGCGTAGGTGGTTTGTTAAGTTGGATGTGAAAGCCCCGGGCTCAACCTGGGGACTGCATCCAAAACTGGCAAGCTAGAGTACGGTAGAGGGTGGTGGAATTTCCTGTGTAGCGGTGAAATGCGTAGATATAGGAAGGAACACCAGTGGCGAAGGCGACCACCTGGACTGATACTGACACTGAGGTGCGAAAGCGTGGGG |
| P-asv4204 | GCAAGCGTTGTCCGGATTCATTGGGCGTAAAGAGTTCGTAAGCGGTTTGTTAAGTCTGATGTTAAATATTAGGGCCCAACCCTGAAACTGCATTGGATACTGGCAGACTGGAGTGCAGTAGAGGCAAGTGGAATTCCCAGTGTAGCGGTGAAATGCGTAGATATTGGGAAGAACACCGGTGGCGCAGGCGACTTGCTGGGCTGTAACTGACGCTGAGGAACGAAAGCTAGGGG |
| P-asv4205 | GCTAGCGTTGTTCGGAATTACTGGGCGTAAAGGGCGCGTAGGCGGCCTGTTTAGTCAGAAGTGAAAGCCCCGGGCTCAACCTGGGAACAGCTTTTGATACTGGCAGGCTCGAGTTCCGGAGAGGATGGTGGAATTCCCAGTGTAGAGGTGAAATTCGTAGATATTGGGAAGAACACCGGTGGCGAAGGCGGCCATCTGGACGGACACTGACGCTGAGGCGCGAAAGCGTGGGG |
| P-asv4206 | GCTAGCGTTGCTCGGAATTACTGGGCGTAAAGGGCGCGTAGGCGGACATTTAAGTCAGGGGTGAAATCCCAGAGCTCAACTCTGGAACTACCTTTGATACTGGGTGTCTTGAGTGTGAGAGAGGTATGTGGAACTCCGAGTGTAGAGGTGAAATTCGTAAATATTCGGAAGAACACCAGTGGCGAAGGCGACATACTGGCTCATTACTGACGCTGAGGCGCGAAAGCGTGGGG |
| P-asv4207 | GCTAGCGTTGTTCGGAATTACTGGGCGTAAAGCGCGCGTAGGCGGCTACTCAAGTCAGAGGTGAAAGCCCGGGGCTCAACCCCGGGACTGCCTTTGAAACTAGGTAGCTAGAATCTTGGAGAGGTCAGTGGAATTCCGAGTGTAGAGGTGAAATTCGTAGATATTCGGAAGAACACCAGTGGCGAAGGCGACTGACTGGACAAGTATTGACGCTGAGGTGCGAAAGCGTGGGG |
| P-asv4208 | GCAAGCGTTAATCGGAATTACTGGGCGTAAAGCGTGCGCAGGCGGTTTTGTAAGTCTGATGTGAAATCCCCGGGCTCAACCTGGGAATTGCATTGGAGACTGCAAGGCTAGAATCTGGCAGAGGGGGGTAGAATTCCACGTGTAGCAGTGAAATGCGTAGATATGTGGGGGAACACCGATGGCGAAGGCGGCTCTCTGGTCTGTAACTGACGCTGAGGCGCGAAAGCGTGGGG |
| P-asv4209 | GCAAGCGTTAATCGGAATTACTGGGCGTAAAGCGCACGTAGGCGGTTGGTTAAGTTGGATGTGAAAGCCCCGGGCTCAACCTGGGAATTGCATTCAAAACTGGCCAACTAGAATACGGGAGAGGGGGGTAGAATTCCAAGTGTAGCGGTGAAATGCATAGAGATTTGGAGGAATACCAGTGGCGAAGGCGGCCCCCTGGCCCGATATTGACGCTGAGGTGCGAAAGCGTGGGG |
| P-asv4210 | GCTAGCGTTGCTCGGAATCACTGGGCGTAAAGGGCGCGTAGGCGGACTTTTAAGTCGGAGGTGAAAGCCCAGGGCTCAACCCTGGAATTGCCTTCGATACTGGGAGTCTTGAGTTCGGAAGAGGTTGGTGGAACTGCGAGTGTAGAGGTGAAATTCGTAGATATTCGCAAGAACACCGGTGGCGAAGGCGGCCAACTGGTCCGATACTGACGCTGAGGCGCGAAAGCGTGGGG |
| P-asv4211 | GCAAGCGTTGTCCGGAATTACTGGGCGTAAAGGGCGTGTGGGCGGTGCGATACGTAGCGGTTTTAAGCCTCCGGCTCACCCGGAGGAGGGCGGCTAAACGGTCGCGCTGGAGGGCAGGAGAGGTGCGTGGAATTCCTGGTGGAGCGGTGAAATGCGTAGAGATCAGGAAGAACACCCGTGGCGAAGGCGGCGCACTGGCCTGGCCCTGACGCTGAGGCGCGACAGCGTGGGG |
| P-asv4212 | GCAAGCGTTATCCGGATTTACTGGGTGTAAAGGGAGCGTAGACGGCTTTGCAAGTCTGATGTGAAAGGCGGGGGCTCAACCCCTGGACTGCATTGGAAACTGTGAGGCTTGAGTGCCGGAGAGGTAAGCGGAATTCCTAGTGTAGCGGTGAAATGCGTAGATATTAGGAGGAACACCAGTGGCGAAGGCGGCTTACTGGACGGTAACTGACGTTGAGGCTCGAAAGCGTGGGG |
| P-asv4213 | GCAAGCGTTACTCGGAATTACTGGGCGTAAAGCGTGCGTAGGTGGTTCGTTAAGTCTGATGTGAAAGCCCTGGGCTCAACCTGGGAATTGCATTGGATACTGGCGGGCTAGAGTACGGTAGAGGGTGGCGGAATTCCTGGTGTAGCAGTGAAATGCGTAGATATCAGGAGGAACATCCGTGGCGAAGGCGGCCACCTGGACCAGTACTGACACTGAGGCACGAAAGCGTGGGG |
| P-asv4214 | GCAAGCGTTATCCGGATTTATTGGGCGTAAAGCGCGTCTAGGCGGTTTGGTAAGTCTGATGTGAAAATGCGGGGCTCAACTCCGTATTGCGTTGGAAACTGCCAAACTAGAGTACTGGAGAGGTGGGCGGAACTACAAGTGTAGAGGTGAAATTCGTAGATATTTGTAGGAATGCCGATGGGGAAGCCAGCCCACTGGACAGATACTGACGCTAAAGCGCGAAAGCGTGGGT |
| P-asv4215 | TCCCTCTTTCTCCTTCCTCCCTGGGCGTAAAGGGTGCGTAGGCGGCCTAACAAGTCAGAGGTGAAAGGCATTGGCTCAACCAATGTAAGCCTTTGAAACTGTTGGGCTTGAGTGCAGGAGAGGTAAGTGGAATTCCTAGTGTAGAGGTGAAATTCGTAGATATTAGGAGGAACACCAGTGGCGAAGGCGACTTACTGGACTGTAACTGACGCTGAGGCACGAAAGCGTGGGG |
| P-asv4216 | GCTAGCGTTGTTCGGAATTACTGGGCGTAAAGCGCACGTAGGCGGCTTTGTAAGTCAGGGGTGAAAGCCTGGAGCTCAACTCCAGAACTGCCTTTGAGACTGCATCGCTTGAATCCGGGAGAGGTGAGTGGAATTCCGAGTGTAGAGGTGAAATTCGTAGATATTCGGAAGAACACCAGTGGCGAAGGCGGCTCACTGGACCGGTATTGACGCTGAGGTGCGAAAGCGTGGGG |

| | |
|---|---|
| P-asv4217 | GCAAGCGTTGTTCGGATTTACTGGGCGTAAAGCGCGCGCAGGCGGATTAACAAGTCAGATGTGAAATCTCGGGGCTCAACCTCGAAACTGCGTCTGAAACTGTCAATCTAGAGTCTCACAGGGGGCAGGGGAATTTCACGTGTAGGGGTAAAATCCGTAGAGATGTGAAGGAACACCGGTGGCGAAAGCGCCTGCCTGGATGAGTACTGACGCTGAGGCGCGAAAGCGTGGGG |
| P-asv4218 | CCGAGTGGTAGTCGTTTTTATTGGGCTTAAAGTGTTCGTAGCCTGCCATGTAAGTCCGCTGTGAAATCCAAGCGCTCAACGTTTGGACGTGCAGCGGATACTGCATGGCTAGGGACCGGACGGCGTCGAGGCTATTCCAGGAGTAGGAGTGAAATCCAAAAATTCCTGGAGGAGCACCTGTGGCGAAAGCGCTCGACGAGTACGGATCCGACGGTGAGGAACGAAAGCCGGGGG |
| P-asv4219 | GCAAGCGTTAATCGGAATTACTGGGCGTAAAGCGCACGTAGGTGGATATTTAAGTCGGATGTGAAATCCCCGGGCTCAACCTGGGAATTGCACCCGATACTGGGTATCTAGAGTATGATAGAGGGAAGTGGAATTTCCGGTGTAGCGGTGAAATGCGTAGATATCGGAAAGAACACCAGTGGCGAAGGCGGCTTCCTGGATCAATACTGACACTGAGGTGCGAAAGCGTGGGG |
| P-asv4220 | GCTAGCGTTGTTCGGAATCACTGGGCGTAAAGCGCGCGTAGGCGGCGATTCAAGTCAGGGGTGAAATCCTGGGGCTCAACCCCAGAATTGCCTTTGAAACTGTATTGCTAGAGTACTGGAGGGGATAGCGGAATTCCTAATGTAGAGGTGAAATTCGTAGATATTAGGAGGAACACCGGTGGCGAAGGCGGCTATCTGGACAGTAACTGACGCTGAGGCGCGAAAGCGTGGGG |
| P-asv4221 | GCAAACGTTGTTCGGAATTACTGGGCTTAAAGCGCGCGTAGGCGGCATCGTTAGTGTCTGGTGAAATCCCACGGCTCACCCGTGGAACTGCCGGGCATACTGCGGTGCTAGAGTGTGGTAGGGGAGAGCGGAACTCCTGGTGGAGCGGTGAAATGCGTAGATATCAGGAGGAACACCGGTGGCGAAGGCGGCTCTCTGGTCCACAACTGACGCTGAGGTGCGAAAGCTAGGGG |
| P-asv4222 | TCAAGCGTTACCCGGAATCATTGGGTGTAAAGGGTTTGTAGGCTGATAATTAAGTCAGGTATGAAAGACCGGAGCTCAACTCCGTGTTTGTATTTGAAACTGGTTGTCTAGAATCAGGGAGAGGTAAGCGGAATTCTAAGTGTAGGGGTGCAATCCGTAGATACTTAGAGGAACACCAAAAGCGAAGGCAGCTTACTGGAACTGTATTGACGCTGATAAACGAAAGCGTGGGG |
| P-asv4223 | GCGAGCGTTGTTCGGAATCATTGGGCGTAAAGCGTATGTAGGCGGACTAGAAAGTCAGAAGTGAAATCCCGGGGCTCAACCTCGGAATTGCTTTTGAAACTCCTAGTCTAGTGTCCCGGAGGGGTTGACGGAATTCCTAGTGTAGAGGTGAAATTCGCAGATATTAGGAGGAACACCAGTGGCGTAGGCGGTCAACTGGACGGGTACAGACGCTGAGATACGAAAGCATGGGT |
| P-asv4224 | CCAAGCGTTATCCGGATTTATTGGGTTTAAAGGGTGCGTAGGCGGCTTATTAAGTCAGGGGTGAAAGACGGTGGCTCAACCATCGCAGTGCCTTTGATACTGATGAGCTTGAATATACTAGAGGTAGGCGGAATGTGACAAGTAGCGGTGAAATGCATAGATATGTCACAGAACACCGATTGCGAAGGCAGCTTACTATGGTATTATTGACGCTGAGGCACGAAAGCGTGGGG |
| P-asv4225 | CCGAGCGTTGTCCGGAATTACTGGGCGTAAAGAGCTCGTAGGTGGTTTGTCGCGTTGTTCGTGAAAACTCACAGCTTAACTGTGGGCGTGCGGGCGATACGGGCAGACTAGAGTACTGCAGGGGAGACTGGAATTCCTGGTGTAGCGGTGGAATGCGCAGATATCAGGAGGAACACCGGTGGCGAAGGCGGGTCTCTGGGCAGTAACTGACGCTGAGGAGCGAAAGCGTGGGG |
| P-asv4226 | GCAAGCGTTGTTCGGAATCATTGGGCGTAAAGCGCGCGCAGGCGGATTGATAAGTCAGGTGTGAAATCTCGAGGCTCAACCTCGAAACTGCGCCTGAAACTATTAGTCTAGAATGTCGGAGGGGACAGGGGAATTTCACGTGTAGGGGTAAAATCCGTAGAGATGTGAAGGAACACCGGAGGCGAAGGCGCCTGTCTGGACGACTATTGACGCTGAGGCGCGAAAGCGTGGGG |
| P-asv4227 | GCAAGCGTTATTCGGATTCACTGGGCGTAAAGGGTGCGCAGGCGGTCGTGTGTGTCAGGTGTGAAATCCCGGAGCTTAACTCCGGAACTGCGCCTGAAACTACACGACTAGAGCATTGGAGAGGGAAGTGGAATTCATGGTGTAGCAGTGAAATGCGTAGATATCATGAGGAACACCAGAGGCGAAGGCGACTTCCTGGACAATTGCTGACGCTCAGGCACGAAAGCATGGGG |
| P-asv4228 | GCAAGCGTTAATCGGAATTACTGGGCGTAAAGCGCGCGTAGGTGGTTTGTTAAGTTGGATGTGAAAGCCCTGGGCTCAACCTAGGAACTGCATCCAAAACTAACTCACTAGAGTACGATAGAGGGAGGTAGAATTCATAGTGTAGCGGTGGAATGCGTAGATATTATGAAGAATACCAGTGGCGAAGGCGGCCTCCTGGATCTGTACTGACACTGAGGTGCGAAAGCGTGGGT |
| P-asv4229 | CCGAGTGATGGCCGATATTATTGGGCCTAAAGCGTCCGTAGCCTGCTGTGTAAGTCCATTGGGAAATCGACCAGCTCAACTGGTCGGCGTCCGGTGGAAACTACACAGCTTGGGGCCGAGAGACTCAACGGGTACGTCCAGGGTAGGAGTGAAATCCTGTAATCCTGAACGGACCACCAATGGGGAAACCACGTTGAGAGACCGGACCCGACAGTGAGGGACGAAAGCCAAGGT |
| P-asv4230 | GCTAGCGTTGTTCGGAATCACTGGGCGTAAAGCGCACGTAGGTGGATTAGTAAGTCAGGGGTGAAAGCCCAGAGCTCAACTCTGGAATTGCCCTTGATACTGCTAGTCTAGAGTTTGGGAGAGGTAAGCGGAATTCCGAGTGTAGAGGTGAAATTCGTAGATATTCGGAGGAACACCAGAGGCGAAGGCGGCTTACTGGCCCAATACTGACACTGAGGTGCGAAAGCGTGGGG |
| P-asv4231 | GCAAGCGTTGTCCGGAATCATTGGGCGTAAAGCGCGTGTAGGCGGCTCGGTAAGTCCGCTTTGAAAGTCCGGGGCTCAACCCCGGGAGGCGGGTGGATACTGTCGAGCTCGAGTCCGGAAGAGGCGAGTGGAATTCCTGGTGTAGCGGTGAAATGCGCAGATATCAGGAGGAACACCGATGGCGAAGGCAGCTCGCTGGGACGTGACTGACGCTGAGACGCGAAAGCGTGGGG |
| P-asv4232 | GCTAGCGTTGTTCGGAATTACTGGGCGTAAAGCGCACGTAGGCGGCCTTGTAAGATGGGGGTGAAATCCCCGGGCTCAACCCGGGAATTGCCTTCATGACTGCAAAGCTCGAGGATGTGAGAGGATAGTGGAATTCCGAGTGTAGAGGTGAAATTCGTAGATATTCGGAAGAACACCAGTGGCGAAGGCGACTGTCTGGCACATTTCTGACGCTGAGGTGCGAAAGCGTGGGG |
| P-asv4233 | GCGAGCGTTGTCCGGAATTATTGGGCGTAAAGCGCGCGTAGGCGGCTGGTTAAGTCCTGTGTGAAAGACCACGGCTCAACCGTGGGGGTGCACGGGAAACTGGCCGGCTTGAGTGCAGGAGAGGGGAGCGGAATTCCCGGTGTAGCGGTGGAATGCGTAGAGATCGGGAGGAACACCAGTGGCGAAGGCGGCTCCCTGGCCTGTAACTGACGCTGAGGCGCGAAAGCGTGGGG |
| P-asv4234 | GCGAGCGTTGTCCGGAATTATTGGGCGTAAAGAGCTCGTAGGCGGTCTGTCGCGTCGAATGTGAAAACCCGGGGCTCAACTCCGGGCTTGCATTCGATACGGGCAGGCTAGAGTTCGGTAGGGGAGTCTGGAATTCCTGGTGTAGCGGTGAAATGCGCAGATATCAGGAGGAACACCGGTGGCGAAGGCGGGACTCTGGGCCGATACTGACGCTGAGGAGCGAAAGCGTGGGG |
| P-asv4235 | ACAAGCGTTGTCCGGAATTATTGGGCGTAAAGGGAGCGCAGGCGGTTGGAATAGTCTGATGTGAAAGCCCACGGCTTAACCGTGGAAGGTCATTGGAAACTGTCCAACTTGAATGCAGAAGAGGTAAGTGGAATTCCAAGTGTAGCGGTGAAATGCGTAGATATTTGGAGGAACACCAGTGGCGAAGGCGACTTACTGGTCTGTGATTGACGCTGAGGCTCGAAAGCGTGGGT |
| P-asv4236 | GCTAGCGTTGTTCGGAATTACTGGGCGTAAAGCGCACGTAGGCGGCTACTCAAGTCAGAGGTGAAAGCCCGGGGCTCAACCCCGGAACTGCCTTTGAAACTAGGTAGCTAGAATCTTGGAGAGGTCAGTGGAATTCCGAGTGTAGAGGTGAAATTCGTAGATATTCGGAAGAACACCAGTGGCGAAGGCGACTGACTGGACAAGTATTGACGCTGAGGTACGAAAGCGTGGGG |
| P-asv4237 | GCGAGCGTTGTCCGGATTCATTGGGCGTAAAGCGCGCGCAGGCGGTGGCGTGAGCCCGGCGTGAAAGCCCCCAGCTCAACTGGGGGAGGCCGTCGGAGACCGCGCCACTTGAGGCCGGTAGGGGAGAGCGGAATTCCCGGTGTAGTGGTGGAATGCGTAGATATCGGGAGGAACACCAGTGGCGAAGGCGGCTCTCTGGACCGGTGCTGACGCTCAGGCGCGAAAGCGTGGGG |
| P-asv4238 | TCGAGTGGTGGTCACTTTTATTGAGCCTAAAGCGTTCGTAGCCGGTCCTGTAAATCTTCAGGAAAATTCTTCCGCTCAACGGAAGATCCTCTGAAGAGACTGCAGGACTAGGGACCGGGTGGGGTTGGATGTACTTCTGGGGTAGGGGTAAAATCCTGTAATCCTGGAAGGACAACCGGCGGCGAAAGCGTCCAACTAGAACGGATCCGACGGTGAGGAACGAGGGCTAGGGG |
| P-asv4239 | TCTAGTGGTGGTCACTTTTATTGAGCCTAAAGCGTTCGTAGCCGGTCCTGTAAATCTCCAGGAAAATTCTCCCGCTCAACGGGAGATCCTCTGGAGAGACTGCGGGACTAGGGACTGGGTGAGGTTGAATGTACTTCCAGGGTAGGGGTAAAATCCTGTAATCCCGGAAGGACAACCGGTGGCGAAGGCGTTCAACTAGAACAGATCCGACGGTGAGGAACGAAGGCTAGGGG |
| P-asv4240 | GCAAGCGTTATCCGGATTTACTGGGTGTAAAGGGAGCGTAGACGGCGATGCAAGTCTGGAGTGAAAGCCCGGGGCTCAACCCCGGGACTGCTTTGGAAACTGTGTTGCTAGAGTGCAGGAGAGGTAAGTGGAATTCCTAGTGTAGCGGTGAAATGCGTAGATATTAGGAGGAACACCAGTGGCGAAGGCGGCTTACTGGACTGTAACTGACGTTGAGGCTCGAAAGCGTGGGG |
| P-asv4241 | GCAAGCGTTAATCGGAATTACTGGGCGTAAAGCGTACGTAGGCGGTTTTATAAGCAAGATGTGAAAGCCCAGGGCTCAACCTTGGAACTGCATTTTGAACTGTAAGACTAGAGTACTGTAGAGGGTGGTGGAATTTCCAGTGTAGCGGTGAAATGCGTAGAGATTGGAAGGAACATCAGTGGCGAAGGCGGCCACCTGGACAGATACTGACGCTGAGGTACGAAAGCGTGGGG |
| P-asv4242 | GCAAGCGTTGTTCGGGATTACTGGGCGTAAAGGGCGCGTAGGCGGTCCCGTAAGTTCCGCGTGAAATCCCCCGGCTCAACCGGGGAATGGCGCGGAAGACTGCGGGGCTTGAGTTCGGGAGAGGGAAGCGGAATTCCGGGTGTAGCGGTGAAATGCGTAGATATCCGGAGGAACACCGGTGGCGAAGGCGGCTTCCTGGACCGACACTGACGCTGAGGCGCGAAAGCTAGGGG |
| P-asv4243 | GCGAACGTTGTTCGGAATCACTGGGCTTAAAGGGCGCGTAGGCGGGCCTCCCAGTCCGGGGTGAAAGCTTTCGGCTTAACCGGAAAAGTGCCCTGGATACTGGAGGCCTTGAGGGAGGTAGGGGCGGCCGGAACTTCCGGTGGAGCGGTGAAATGCGTTGATATCGGAAGGAACGCCGGTGGCGAAAGCGGCCGCTGGATCTCTACTGACGCTGAGGCGCGAAAGCCAGGGG |
| P-asv4244 | GCAAGCGTTGTCCGGAATCATTGGGCGTAAAGCGCGTGTAGGCGGTTTGGTAAGTCCGCTCTGAAAGTCCGGGGCTCAACCCCGGAAGGCGGGTGGATACTGTCAGACTAGAGTGCGGAAGAGGCGAGTGGAATTCCCGGTGTAGCGGTGAAATGCGCAGATATCGGGAGGAACACCAATTGCGAAGGCAGCTCGCTGGGACGTAACTGACGCTGAGACGCGAAAGCGTGGGG |
| P-asv4245 | GCGAGCGTTGTCCGGATTTACTGGGCGTAAAGAGCGCGCAGGCGGTCGTGCAAGTCGCGTGTGAAAGCCCCCGGCTCAACTGGGGAGGGTCACGCGATACTGTTCGACTTGAAGGCAGGAGAGGGAAGTGGAATTCCCGGTGTAGTGGTGAAATTCGTAGATATCAGGAGGAACACCGGTGGCGAAGGCGGCCCTCTGGACGGTGACTGACGCTGAGACGCGAAAGCGTGGGG |
| P-asv4246 | GCTAGCGTTATTCGGAATTACTGGGCGTAAAGGGCTCGTAGGTGGCCAACTAAGTCGGACGTGAAATCCCTCAGCTTAACTGGGGAACTGCGTCCGATACTGGATGGCTTGAGTTTGGGAGAGGGATGCGGAATTCCAGGTGTAGCGGTGAAATGCGTAGATATCTGGAGGAACACCGGTGGCGAAGGCGGCATCCTGGACCAACACTGACACTGAGGAGCGAAAGCCAGGGG |
| P-asv4247 | GCGAACGTTGTTCGGAATCACTGGGCTCAAAGGGCGCGTAGGCGGCTTGCCAAGTCAGTGGTGAAATCCCGCAGCTTAACTGCGGAAGTGCCTTTGATACTGGCGAGCTCGAGGGAGGTAGGGGTATGTGGAACTTCTGGTGAAGCGGTGAAATGCGTTGATATCAGAAGGAGCGCCGGTGGCGAAAGCGACGTACTGGACCTCTTCTGACGCTGAGGCGCGAAAGCTAGGGG |
| P-asv4248 | GCGAGCGTTGTCCGGATTTATTGGGCGTAAAGCGCGTGTAGGCGGTTTGGTAAGTCCGCTCTGAAAGTCCGGGGCTCAACCCCGGAAGGCGGGTGGATACTGTCAGACTAGAGTGCGGAAGAGGCGAGTGGAATTCCCGGTGTAGCGGTGAAATGCGCAGATATCGGGAGGAACACCAATTGCGAAGGCAGCTCGCTGGGACGTAACTGACGCTGAGACGCGAAAGCGTGGGG |
| P-asv4249 | GCGAGCGTTGTCCGGAATCACTGGGCGTAAAGGGCGCGTAGGCGGTTCGTTAAGTTGCTGGTGAAATCCCGGGGCTCAACTCCGGGGCTGCCGGCAAGACTGGCGGGCTAGAGCACGGTAGAGGCAGATGGAATTCCCGGTGTAGCGGTGGAATGCGTAGAGATCGGGAAGAATACCGGTGGCGAAGGCGTTCTGCTGGGCCGTTGCTGACGCTGAGGCGCGACAGCGTGGGG |
| P-asv4250 | GCAAGCGTTGTTCGGAATTACTGGGCGTAAAGAGCGTGTAGGCGGCCAGGTAGGTGGGTTGTGAAAACTGGAGGCTCAACCTTCAGACGTCGACCCAAACCCCCTGGCTAGAGTCCGGAAGAGGAGAGTGGAATTCCTGGCGTAGCGGTGAAATGCGCAGATATCAGGAAGAACACCCGTGGCGAAGGCGGCTCTCTGAGACGGTACTGACGCTGAGGCGCGAAAGCGTGGGG |
| P-asv4251 | CCAAGCGTTATCCGGATTTATTGGGTTTAAAGGGTGCGTAGGTGGCTTTTTAAGTCAGGGGTGAAAGACATCGGCTCAACCGATGCAGTGCCTTTGATACTGAAGAGCTTGAATTATGTTGAAGATGGCGGAATGAGACAAGTAGCGGTGAAATGCATAGATATGTCTCAGAACACCGATTGCGAAGGCAGCTGTCTAAGCATTGATTGACACTGATGCACGAAAGCGTGGGG |
| P-asv4252 | GCGAGCGTTGTCCGGATTCATTGGGCGTAAAGCGCGCGTAGGCGGCCCGGCAAGTCGGCCGTGAAATCTCCCAGCTCAACTGGGAGGGGTCGGGCGAGACTGCGGGGCTCGAGGTGGGCAGAGGGCAGTGGAATTCCGGGTGTAGTGGTGGAATGCGTAGAGATCCGGAGGAACACCAGTGGCGAAGGCGGCTGCCTGGGCCTGGCCTGACGCTGAGGCGCGAAAGCGTGGGG |
| P-asv4253 | GCGAGCGTTGTCCGGAATTATTGGGCGTAAAGGGCTCGTAGGCGGTCTGTCACGTCGGGAGTGAAAACTCGGGGCTCAACCCCGAGCCTGCTTCCGATACGGGCAGACTAGAGGTATGCAGGGGAGAATGGAATTCCTGGTGTAGCGGTGAAATGCGCAGATATCAGGAGGAACACCGATGGCGAAGGCAGGTCTCTGGGCTGTAACTGACGCTGAGGAGCGAAAGCATGGGG |
| P-asv4254 | GCAAGCGTTACTCGGAATCACTGGGCGTAAAGCGTACGTAGGCGGTTGGTTAAGTCTGCTGTGAAAGCCCTGGGCTCAACCTGGGAACTGCAGTGGATACTGGCCAGCTAGAGTGTGATAGAGGATGGTGGAATTCCCGGTGTAGCGGTGAAATGCGTAGAGATCGGGAGGAACACCAGTGGCGAAGGCGGCCATCTGGATCAACACTGACGCTGAGGCACGAAAGCGTGGGG |
| P-asv4255 | GCAAGCGTTGTCCGGAATTATTGGGCGTAAAGTGCGCGCAGGCGGTCCTTTAAGTCTGATGTGAAAGCCCACGGCTCAACCGTGGAGGGTCATTGGAAACTGGGGGACTTGAGTACAGAAGAGGAAAGCGGAATTCCACGTGTAGCGGTGAAATGCGTAGAGATGTGGAGGAACACCAGTGGCGAAGGCGGCTTTCTGGTCTGTAACTGACGCTGAGGCGCGAAAGCGTGGGG |

| | |
|---|---|
| P-asv4256 | GCAAGCGTTGTTCGGAATTACTGGGCGTAAAGGGCGCGTAGGCGGCCTTCTAAGTCGAACGTGAAATCCCCGGGCTCAACCCGGGAACTGCGTCCGATACTGGGAGGCTTGAATCCGGGAGAGGGATGTGGAATTCCAGGTGTAGCGGTGAAATGCGTAGATATCTGGAGGAACACCGGTGGCGAAGGCGGCATCCTGGACCGGTATTGACGCTGAGGCGCGAAAGCTAGGGG |
| P-asv4257 | GCGAGCGTTAATCGGAATTACTGGGCGTAAAGCGTGCGTAGACGGTTACGTAAGTCGGGTGTGAAAGCCCCGGGCTCAACCTGGGAATGCCATTCGAGACTGCATAGCTAGGGTGCGGAAGAGGGAAGCGGAATTTCAGGTGTAGCGGTGAAATGCGTAGATATCTGAAGGAACACCAGTGGCGAAAGCGGCTTCCTGGTCCAGCACCGACGTTCAGGCACGAAAGCGTGGGG |
| P-asv4258 | GCAAGCGTTGTCCGGAATCATTGGGCGTAAAGAGCGCGTAGGCGGCCCTGTAAGTCCGTTCTGAAAGTCGGAGGCTCAACCTCCGAAAGCGGATGGATACTGCAGGGCTAGAGTCCGGAAGAGGCGAGTGGAATTCCTGGTGTAGCGGTGGAATGCGCAGATATCAGGAGGAACACCAACGGCGAAGGCAGCTCGCTGGGACGGTACTGACGCTGAGGCGCGAAAGCGTGGGG |
| P-asv4259 | GCTAGCGTTGCTCGGAATCACTGGGCGTAAAGCGCACGTAGGCGGGTCGTTAAGTCAGGGGTGAAATCCTGGAGCTCAACTCCAGAACTGCCCTTGATACTGGCGATCTCGAGTTCGGGAGAGGTGAGTGGAACTGCGAGTGTAGAGGTGAAATTCGTAGATATTCGCAAGAACACCAGTGGCGAAGGCGGCTCACTGGCCCGATACTGACGCTGAGGTGCGAAAGCGTGGGG |
| P-asv4260 | GCAAACGTTGTTCGGAATTACTGGGCGTAAAGCGCGTGTAGGCGGCAAGGCAGGTCGGATGTGAAAGCCCGCGGCTCAACCGTGGAAGTGCACTCGAAACCGCCTAGCTAGAGTCTCGGAGAGGAAGGTGGAATGCTTGGTGTAGAGGTGAAATTCGTAGATATCAAGCAGAACACCTGTGGCGAAGGCGGCCTTCTGGACGATGACTGACGCTGAGACGCGAAAGCGTGGGG |
| P-asv4261 | GCGAGCGTTAATCGGAATTATTGGGCGTAAAGAGCTCGTAGGCGGTTTGTCGCGTCTGCTGTGAAAACTGGAGGCTCAACCTCCAGCTTGCAGTGGGTACGGGCAGACTAGAGTGCGGTAGGGGAGATTGGAATTCCTGGTGTAGCGGTGGAATGCGCAGATATCAGGAGGAACACCGATGGCGAAGGCAGATCTCTGGGCCGTAACTGACGCTGAGGAGCGAAAGCATGGGG |
| P-asv4262 | GCAAGCGTTGCTCGGAATCACTGGGCGTAAAGGGCGCGTAGGCGGGATGGCAAGTCAGTCGTGAAATCTCTCGGCTCAACCGAGAAACGTCGGCTGATACTGCCGTTCTTGAGGGATGCAGAGGAGACTGGAATTCCCGGTGTAGCGGTGAAATGTGTAGAGATCGGGAGGAACACCGGTGGCGAAGGCGGGTCTCTGGGCATCTCCTGACGCTGAGGCGCGAAAGCGTGGGG |
| P-asv4263 | GCAAGCGTTGTTCGGAATTACTGGGCGTAAAGGGTTCGTAGGTGGCCAACTAAGTCAAACGTGAAATCCCTCAGCTTAACTGGGGAGCTGCGTCTGAGACTGGATGGCTTGAGTGCAGGAGAGGAACGCGGAATTCCAGGTGTAGCGGTGAAATGCGTAGATATCTGGAGGAACACCAGTGGCGAAGGCGGCTTCCTGGCCTGCAACTGACGCTGAGACGCGAAAGCGTGGGG |
| P-asv4264 | GCAAGCGTTGTCCGGATTTATTGGGCGTAAAGAGCGTGTAGGCGGCCAGACAGGTCCGTTGTGAAAACTCGAGGCTCAACCTCCAGACGTCGATGGAAACCGCTTGGCTAGAGTCCGGAAGAGGAGAGTGGAATTCCTGGTGTAGCGGTGAAATGCGCAGATATCAGGAAGAACACCGATGGCGAAGGCAGCTCTCTGGGACGGTACTGACGCTGAGACGCGAAAGCGTGGGG |
| P-asv4265 | GCGAGCGTTGTTCGGATTTACTGGGCGTAAAGGGCGCGTAGGCGGCGGGTCAAGTCGAGTGTGAAAGCCCCGGGCTCAACCTGGGAGGTGCGCTCGAGACTGGTCTGCTAGAGTCCGGGAGAGGAGCGTGGAATTCCCAGTGTAGAGGTGAAATTCGTAGATATTGGGAAGAACACCGGTGGCGAAGGCGGCGATCTGGACGATGACTGACGCTGAGACGCGAAAGCGTGGGG |
| P-asv4266 | GCGAGCGTTGTCCGGAATTATTGGGCGTAAAGAGCGTGTAGGCGGTTCGGTAAGTCTGCTGTGAAATCCTGGGGCTCAACCCTGGGCGTGCAGCGGATACTGCCGGGCTAGAGGGTGGTAGAGGCGAGTGGAATTCCCAGTGCAGCGGTGAAATGCGCAGATATCGGGAGGAACACCAGTAGCGAAGGCGGCTCGCTGGGCCATTCCTGACGCTGAGACGCGAAAGCTAGGGG |
| P-asv4267 | GCGAGCGTTGTTCGGAATTACTGGGCGTAAAGGGCGCGCAGGCGGCTCTGAAAGTCAGATGTGAAAGCCCCGGGCTTAACCTGGGAATTGCATTTGATACTCTAGAGCTTGAGTACGGGAGAGGAGAGTGGAATTCCCAGTGTAGAGGTGAAATTCGTAGATATTGGGAAGAACACCAGTGGCGAAGGCGGCTCTCTGGACCGTAACTGACGCTGAGGCGCGAAAGCGTGGGG |
| P-asv4268 | GCAAGCGTTGTCCGGATTTATTGGGTTTAAAGGGTGCGTAGGCGGCCTGTTAAGTCAGTGCTGAAATATCCCGGCTTAACCGGGAGGGTGGCATTGATACTGATGGGCTTGAGTGGAGTCGAGGTAGGCGGAATTGACGGTGTAGCGGTGAAATGCTTAGATATCGTCAAGAACACCGATAGTGTAGACAGCTTACTAGGCTCAAACTGACGCTGAGGCACGAAAGTGTGGGG |
| P-asv4269 | GCAAGCGTTATCCGGATTCACTGGGTTTAAAGGGTGCGTAGGCGGGTTGGTAAGTCCGTGGTGAAATCTCCAAGCTTAACTTGGAAACTGCCGTGGATACTATCAATCTTGAATATCGTGGAGGTGAGCGGAATATGTCATGTAGCGGTGAAATGCTTAGATATGACATAGAACACCCATTGCGAAGGCAGCTCGCTACACGGTTATTGACGCTGAGGCACGAAAGCGTGGGG |
| P-asv4270 | GCAAGCGTTGTTCGGAATTATTGGGCGTAAAGGGCGCGTAGGCGGTTTGGCAAGTTTGGTGTGAAATCTATGGGCTCAACCCATAGCCTGCATCGAAAACTGTCGGGCTTGAGTATGGGAGAGGTGAGTGGAATTTCCGGTGTAGCGGTGAAATGCGTAGATATCGGAAGGAACACCTGTGGCGAAAGCGGCTCACTGGACCATAACTGACGCTGATGCACGAAAGCTAGGGG |
| P-asv4271 | GCTAGCGTTGTTCGGAATTACTGGGCGTAAAGCGCGTGTAGGCGGTTGTCCAAGTTGGATGTGAAAGCCTTGAGCTTAACTCAAGAAATGCATTCAGGACTGGATGACTAGAGGACCGGAGAGGATAGTGGAATTCCCAGTGTAGTGGTGAAATACGTAGAGATTGGGAAGAACACCAGTGGCGAAGGCGGCTATCTGGACGGTTTCTGACGCTAAGACGCGAAAGCGTGGGG |
| P-asv4272 | GCTAGCGTTGTTCGGAATTACTGGGCGTAAAGCGCGCGCAGGCGGTCCTTTAAGTCAGGGGTGAAAGCCCAGAGCTCAACTCTGGAATTGCCCTTGAAACTAGAGGACTTGGGTACGGGAGAGGTGAGTGGAATTCCGAGTGTAGAGGTGAAATTCGTAGATATTCGGAAGAACACCAGTGGCGAAGGCGGCTCACTGGCCCGTTACCGACGCTCAGGCGCGACAGCGTGGGG |
| P-asv4273 | GCTAGTGTTGTTCGGAATTACTGGGCGTAAAGGGCGCGCAGGCGGCATGGCAAGTCGAGTGTGAAGTCCCTTGGCTTAACTGAGGAATGGCGCTCGAAACTACCTCGCTGGAGGGCGGGAGAGGGAAGTGGAACTCTCGGTGTAGGGGTGAAATCTGTAGATATCGAGAGGAACACCGGTGGCGAAGGCGGCTTCCTGGCCCGTACCTGACGCTGAGGCGCGAAAGCGTGGGG |
| P-asv4274 | GCAAACGTTGTTCGGAATCACTGGGCGTAAAGGGCGTGTAGGCGGTTCGGTAAGTGGAATGTGAAATCCCCCGGCTTACCCGGGGAACTGCATCCCATACTGCTGGACTTGAGTGCGGGAGAGGATGGAGGAATTCCTGGTGTAGCGGTGAAATGCGTAGATATGCGGAGGAACACCGATGGCGAAGGCAGTCCCCTGGGCCTGCACTGACGCTCATGCACGAAAGCGTGGGG |
| P-asv4275 | GCAAGCGTTAATCGGAATTACTGGGCGTAAAGCGTGCGCAGGCGGTTTTGCAAGACAGATGTGAAATCCCCGGGCTTAACCTGGGAACTGCATTTGTGACTGCAAGGCTAGAGTGTGTCAGAGGGAGGTGGAATTCCGCGTGTAGCAGTGAAATGCGTAGAGATGCGGAGGAACACCGATGGCGAAGGCAGCCTCCTGGGATAACACTGACGCTCATGCACGAAAGCGTGGGG |
| P-asv4276 | GCAAGCGTTGTCCGGAATCATTGGGCGTAAAGCGCGTGTAGGCGGCCCGGAAAGTCTGGTGTGAAAACTTTGGGCTCAACCCTAAGCCTGCACTGGATACTATCGGGCTTGAATCCGGAAGAGGAGAGTGGAATTCCTGGTGTAGCGGTGGAATGCGCAGATATCAGGAGGAACACCAATGGCGAAGGCAGCTCTCTGGTACGTGATTGACGCTGAGACGCGAAAGCGTGGGG |
| P-asv4277 | GCAAGCGTTGTTCGGAGTCACTGGGCGTAAAGGGCGCGTAGGCGGTGCATAAAGTCAGGCGTGAAAGCCCCGGGCTTAACCTGGGAATTGCGCTTGATACTGATGCGCTTGAGAACGGGAGAGGATGGCGGAATTCCCAGTGTAGAGGTGAAATTCGTAGATATTGGGAAGAACACCGGTGGCGAAGGCGGCCATCTGGACCGTTTCTGACGCTGAGGCGCGAAAGCGTGGGG |
| P-asv4278 | GCGAGCGTTGTCCGGAATTATTGGGCGTAAAGGGCTCGTAGGCGGTTTGTCGCGTCGGGAGTGTAATCCGCCGGCTTAACTGGTGGGGTGCTTTCGGTACGGGCAGACTGGAGGTATGCAGGGGAGAACGGAATTCCTGGTGTAGCGGTGAAATGCGCAGATATCAGGAGGAACACCGGTGGCGAAGGCGGTTCTCTGGGCATGTCCTGACGCTGAGGAGCGAAAGTGTGGGG |
| P-asv4279 | GCAAGCGTTGTTCGGAATTACTGGGCGTAAAGGGTTTGTAGGTGGCTTGTTAAGTCAGACGTGAAATCCCTCAGCTTAACTGGGGAACTGCGTCTGATACTGGTGAGCTTGAGTGCAGGAGAGGAACGCGGAATTCCAGGTGTAGCGGTGAAATGCGTAGATATCTGGAGGAACACCGGTGGCGAAGGCGGCGTTCTGGACTGCAACTGGCACTGATAAACGAAAACCAGGGT |
| P-asv4280 | GCAAGCGTTGTCCGGAATTATTGGGCGTAAAGAGCTCGTAGGTGGTCCGCTACGTCCGCTGTGAAATTCCAGGGCTTAACCCTGGACTTGCAGTGGATACGGGTGGACTAGAGGTAGGTAGGGGAGAATGGAATTCCCGGTGTAGCGGTGAAATGCGCAGATATCGGGAGGAACACCGGTGGCGAAGGCGGTTCTCTGGGCCTTACCTGACACTGAGGAGCGAAAGCGTGGGG |
| P-asv4281 | CCGAGCGTTGTCCGGAATTATTGGGCGTAAAGGGCTCGTAGGCGGTTCGTCACGTCGGGAGTGAAAACTCGGAGCTCAACTCCGAGCCTGCTTCCGATACGGGCAGACTAGAGGTAGGCAGGGGAGAGCGGAACTCCTGGTGTAGCGGTGGAATGCGCAGATATCAGGAAGAACACCGGTGGCGAAGGCGGCTCTCTGGGCCTTACCTGACGCTGAGGAGCGAAAGCGTGGGT |
| P-asv4282 | GCGAGCGTTAATCGGAATTACTGGGCGTAAAGCGTGCGCAGGCGGCGCCGCAAGTCAGGCGTGAAATCCCCGAGCTTAACTTGGGAATGGCGCTTGAAACTACGGTGCTGGAGTGTGGCAGAGGGAGGTGGAATTCCACGTGTAGCGGTGAAATGCGTAGAGATGTGGAGGAACACCGATGGCGAAGGCAGCCTCCTGGGCCAACACTGACGCTCATGCACGAAAGCGTGGGG |
| P-asv4283 | GCTAGCGTTGCTCGGAATGACTGGGCGTAAAGGGCGCGTAGGCGGTTTGCATAGTCGGGCGTGAAATTCCCGGGCTCAACCCGGGGGCTGCGTTCGAGACGTGCAGACTGGAGTGGGGAAGAGGGTCGTGGAATTCCCAGTGTAGAGGTGAAATTCGTAGATATTGGGAAGAACACCGGTGGCGAAGGCGGCGACCTGGTCCTTGACTGACGCTGAGGCGCGAAAGCGTGGGG |
| P-asv4284 | GCGAGCGTTGTCCGGAATCACTGGGCGTAAAGGGCGCGTAGGTGGCGCGACAAGTCCGCGGTGAAAGCTCGACGCTTAACGTCGAGTCGGCCACGGATACTGTTGGGCTTGAGCACTGTAGAGGCTAATGGAATTCCGGGTGTAGCGGTGGAATGCGTAGAGATCCGGAAGAACACCGGTGGCGAAGGCGGTTAGCTGGGCAGTTGCTGACACTGAGGCGCGACAGCGTGGGG |
| P-asv4285 | GCAAGCGTTGTCCGGAATTATTGGGCGTAAAGGGCGCGCAGGCGGCTTCTTAAGTCTGTCTTAAAAGTGCGGGGCTTAACCCCGTGATGGGATGGAAACTGGGAAGCTCAGAGTATCGGAGAGGAAAGCGGAATTCCTAGTGTAGCGGTGAAATGCGTAGATATTAGGAGGAACACCAGTGGCGAAAGCGGCTTTCTGGACGAAAACTGACGCTGAGGCGCGAAAGCCAGGGG |
| P-asv4286 | GCTAGCGTTGCTCGGAATTACTGGGCGTAAAGGGCGCGTAGGCGGGTATTTAAGTCAGTGGTGAAATCCTAGAGCTTAACTCTAGAACTGCCATTTATACTATTTATCTTGAATATTGTGGAGGTAAGCGGAATATGTCATGTAGCGGTGAAATGCTTAGATATGACATAGAACACCTATTGCGAAGGCAGCTTACTACGCATATATTGACGCTGAGGCACGAAAGCGTGGGG |
| P-asv4287 | ACGAGTAGTGGTCACTTTTATTGAGCCTAAAGCGTTCGTAGCCGGTTTTGTAAATCTTCAGATAAAGCCTGAAGCTTAACCCCAGAAAGTCTGAAGAGGCTGCAGGACTTGAGATCGGGTGAGGTTAAACGTACTTTCAGGGTAAGGGTAAAATCCTGTAATCCCGGAAGGACGACCAGTGGCGAAAGCGTTTAACTAGAACGAATCTGACGGTAAGGAACGAAGGCTAGGGT |
| P-asv4288 | GCGAGCGTTGTCCGGATTTATTGGGCGTAAAGAGTTCGTAGGTGGTTTGTTAAGTTTGGTGTAAAGATCGGGGCTCAACCCTGGGACTGCACTGAATACTGGCAGACTCGAGTGTGGTAGAGGCTAGTGGAATTCCCAGTGTAGCGGTGAAATGCGTAGATATTGGGAAGAACACCGGTGGCGTAGGCGACTAGCTGGGCCATAACTGACGCTGAGGAACGAAAGCCAGGGG |
| P-asv4289 | GCAAGCGTTGTCCGGAATTATTGGGCGTAAAGAGCTCGTAGGCGGTCTGTCACGTCGGCTGTGAAAACCCGAGGCTCAACCTCGGGCCTGCAGTCGATACGGGCAAACTAGAGTACTGCAGGGGAGACTGGAATTCCTGGTGTAGCGGTGAAATGCGCAGATATCAGGAGGAACACCGGTGGCGAAGGCGGGTCTCTGGGCAGTAACTGACGCTGAGGAGCGAAAGCGTGGGG |
| P-asv4290 | GCAAGCGTTGTTCGGAATTACTGGGCGTAAAGGGCTCGTAGGCTGCCAACTAAGTCAGACGTGAAATCCCTCGGCTTAACCGGGGAACTGCGTCTGATACTGGATGGCTTGAGGTTGGGAGAGGGATGCGGAATTCCAGGTGTAGCGGTGAAATGCGTAGATATCTGGAGGAACACCGGTGGCGAAGGCGGCATCCTGGACCAATTCTGACGCTGAGGAGCGAAAGCCAGGGG |
| P-asv4291 | GCAAACGTTGTTCGGAATTACTGGGCGTAAAGCGTGTGTAGGCGGTCTCATAAGTCGGGTGTGAAAGCCCCGGGCTCAACCCGGGAAGTGCACTCGAAACTGCGAGGCTAGAGTACTGGAGAGGTTGGTGGAATTCTCGGTGTAGAGGTGAAATTCGTAGATATCGAGAGGAACACCGGTGGCGAAGGCGGCCAACTGGACAGATACTGACGCTGAGACACGAAAGCGTGGGG |
| P-asv4292 | GCAAGCGTTGTTCGGAATTATTGGGCGTAAAGGGTGCGTAGGCGGTGCGGTAAGTCTTCTGTGAAATCCCCGGGCTCAACTCGGAGCCTGCAGGAGAAACTGCCGTGCTTGAGTATGGGAGAGGTGAGTGGAATTCCTGGTGTAGCGGTGAAATGCGTAGATATCAGGAGGAACACCTGTGGCGAAAGCGGCTCACTGGACCATAACTGACGCTGAGGCACGAAAGCTAGGGG |
| P-asv4293 | GCGAGCGTTATCCGGATTTATTGGGTTTAAAGGGTGCGTAGGCGGCTTTTTAAGTCAGCGGTGAAAGGTTCGGGCTCAACCCGGACATTGCCGTTGAAACTGGGGAGCTTGAGTATGGATGAGGTAGGCGGAATGCGTGGTGTAGCGGTGAAATGCATAGATATCACGCAGAACTCCGATTGCGAAGGCAGCTTACTAAGGCATTACTGACGCTGAAGCACGAAAGCGTGGGT |
| P-asv4294 | GCGAGCGTTAATCGGAATTACTGGGCGTAAAGTGTGCGCAGGTGGCTGAGCAAGTCGAGTGTGAAATCCCCGGGCTTAACTTGGGAATTGCGCTCGAAACTACTTGGCTCGAGTGTGGCAGAGGAAGGTGGAATTCCAGGTGTAGCGGTGAAATGCGTAGAGATCTGGAGGAACACCGATGGCGAAGGCAGCCTTCTGGGCCAACACTGACGCTCATGCACGAAAGCGTGGGG |

| | |
|---|---|
| P-asv4295 | TCGAGCGTTGTCCGGAGTTACTGGGCGTAAAGCGCATGCAGGCGGCTCATTACGCATGGCGTGAAAGCCCCCGGCTTAACCGGGGAGGGTCGTCGTGAACGGGTGAGCTTGAGGGTATCAGGGGCTGGTGGAATTCCCGGTGTAGTGGTGAAATGCGTAGAGATCGGGAGGAACACCTGTGGCGAAGGCGGCCAGCTGGGATACACCTGACGCTGAGATGCGAAGGCGTGGGG |
| P-asv4296 | GCAAGCGTTAATCGGAATTACTGGGCGTAAAGCGCACGTAGGCGGTTTGTTAAGTTGGATGTGAAAGCCCCGGGCTCAACCTGGGAATTGCATTCAAAACTGGCCAACTAGAGTATGTGAGAGGGGGGTAGAATTCCAAGTGTAGCGGTGAAATGCGTAGAGATTTGGAGGAATACCAGTGGCGAAGGCGGCCCCCTGGCACAATACTGACGCTGAGGTGCGAAAGCGTGGGG |
| P-asv4297 | GCAAGCGTTAATCGGAATTACTGGGCGTAAAGCGTTCGTAGGCGGTTTGTTAAGCAAGATGTGAAAGCCCAGGGCTCAACCTTGGAACTGCATTTTGAACTGGCAAACTAGAGTACTGTAGAGGGTGGTGGAATTTCCAGTGTAGCGGTGAAATGCGTAGAGATTGGAAGGAACATCAGTGGCGAAGGCGGCCACCTGGACAGATACTGACGCTGAGGAACGAAAGCGTGGGG |
| P-asv4298 | GCAAGCGTTGTCAGGAATTATTGGGCGTAAAGCGCGCGCAGGCGGTCATTTAAGTCTGGTGTTTAAACCTTGGGCTCAACCTGGGGTCGCACTGGAAACTGGGTGACTTGAGTACAGAAGAGGAAAGTGGAATTCCACGTGTAGCGGTGAAATGCGTAGATATGTGGAGGAACACCAGTGGCGAAGGCGACTTTCTGGGCTGTAACTGACGCTGAGGCGCGAAAGCGTGGTG |
| P-asv4299 | GCGAGCGTTAATCGGAATTACTGGGCGTAAAGCGTGCGCAGGCGGTTTTGTAAGACAGGCGTGAAATCCCCGGGCTCAACCTGGGAATTGCGCTTGTGACTGCAAGGCTAGAGTGTGTCAGAGGGGGGTAGAATTCCACGTGTAGCAGTGAAATGCGTAGAGATGTGGAGGAATACCGATGGCGAAGGCAGCCCCCTGGGACATGACTGACGCTCATGCACGAAAGCGTGGGG |
| P-asv4300 | GCAAGCGTTGTCCGGAATTATTGGGCGTAAAGCGCGCGCAGGCGGCCTCTTAAGTCTGATGTGAAAGCCCCCGGCTCAACCGGGGAGGGCCATTGGAAACTGGGAGGCTTGAGTATAGGAGAGAAGAGTGGAATTCCACGTGTAGCGGTGAAATGCGTAGAGATGTGGAGGAACACCAGTGGCGAAGGCGACTCTTTGGCCTATAGCTGACGCTGAGGCGCGAAAGCGTGGGG |
| P-asv4301 | CCAAGCGTTATCCGGATTTATTAGGTTTAAAGGGTCCGCAGGCGGGATTTTAAGTCAGTGGTGAAAGCCTACAGCTCAACTGTAGAACTGCCATTGATACTGAAATTCTTGAGTGTGGTTGAAGTGGGCGGAACATGTCATGTAGCGGTGAAATGCATAGATATGACATAGAACACCAATTGCGAAGGCAGCTCACTAAGCCATTACTGACGCTGAGGGACGAAAGCGTGGGG |
| P-asv4302 | CCAGGCGTTATCCGGATTTATTGGGTTTAAAGGGAGCGTAGGCGGATTGTTAAGTCAGCGGTTAAAGGGTGTGGCTCAACCATACATTGCCGTTGAAACTGGCGATCTTGAGTGCAGACAGGGATGCCGGAATTCGTGGTGTAGCGGTGAAATGCTTAGATATCACGAAGAACTCCGACCGCGAAGGCAGGTGTCCGGGCTGCAACTGACGCTGAGGCTCGAAAGTGTGGGT |
| P-asv4303 | GCGAGCGTTAATCGGGATTACTGGGCGTAAAGCGCGCGTAGGTGGTTCGTTAAGTTGGATGTGAAAGCCCCGGGCTCAACCTGGGAATTGCATCCAAAACTGGCGAGCTAGAGTACGGTAGAGGGTGGTGGAATTTCCTGTGTAGCGGTGAAATGCGTAGATATAGGAAGGAACACCAGTGGCGAAGGCGACCACCTGGACTGATACTGACACTGAGGTGCGAAAGCGTGGGG |
| P-asv4304 | GCGAGCGTTGTCCGGATTTATTGGGTTTAAAGGGTGCGTAGGCGGCTCTTTAAGTCAGTGGTGAAAGCCCGCAGCTTAACTGTGGAACTGCCATTGAAACTGGAGAGCTTGAGTACGGTTGAAGTAGGCGGAATTTATGGTGTAGCGGTGAAATGCATAGATACCATAAAGAACACCGATAGCGTAGGCAGCTTACTAAGCCGTAACTGACGCTGAGGCACGAAAGCATGGGT |
| P-asv4305 | CCAAGCGTTAATCGGAATCACTGGGCTTAAAGGGTGCGCAGGCGGACCCGTGTGTGTGCCGTGAAATCCCTCAGCTCAACTGAGGAATTGCGGTGCAAACTGCGGGTCTTGAGGCAGGTAGGGGCTGTTGGAACGATCGGTGGAGCGGTGAAATGCGTTGATATCGATCGGAACGCCGAAGGTGAAGACAAGCAGCTGGGCCTGTCCTGACGCTCAGGCACGAAAGCGTGGGG |
| P-asv4306 | GCAAGCGTTATCCGGAATTATTGGGCGTAAAGCGCGCGCAGGCGGTCTCTTAAGTCTGATGTGAAAGCCCACGGCTCAACCGTGGAGGGTCATTGGAAACTGGGAGACTTGAGTGCAGGAGAGAAAAGTGGAATTCCACGTGTAGCGGTGAAATGCGTAGAGATGTGGAGGAACACCAGTGGCGAAGGCGGCTTTTTGGCCTGTAACTGACGCTGAGGCGCGAAAGCGTGGGG |
| P-asv4307 | GCAAGCGTTAATCGGAATCACTGGGCGTAAAGGGTGCGTAGGCGGTGCATTAGGTCTGTCGTGAAATCCCCGGGCTCAACCTGGGAATGGCGGTGGAAACCGGTGTACTAGAGTATGGGAGAGGGTGGTGGAATTCCAGGTGTAGCGGTGAAATTCGTAGATATTCGGAAGAACACCAGTGGCGAAGGCGGCTCACTGGACTGGTATTGACGCTGAGGCACGAAAGCGTGGGG |
| P-asv4308 | CCTCTTCTTTTTCACTTTTATTGAGCCTAAAGCGTTCGTAGCCGGTTTTGTAAATCTTCAGATAAAGCCTGAAGCTTAACTCCAGAAAGTCTGAAGAGACTGCAAGACTTGAGATCGGGTGAGGTTAAACGTACTTTCAGGGTAGGGGTAAAATCCTGTAATCCCGGAAGGACGACCAGTGGCGAAAGCGTTTAACTAGAACGAATCTGACGGTAAGGAACGAAGGCTAGGGT |
| P-asv4309 | GCAAGCGTTATCCGGATTCACTGGGTTTAAAGGGTGCGTAGGCGGGTATTTAAGTCAGTGGTGAAATCCTAGAGCTTAACTCTAGAACTGCCATTGATACTATTTATCTTGAATATTGTGGAGGTAAGCGGAATATGTCATGTAGTGGTGAAATTCGTAGATATTCGGAAGAACACCAGTGGCGAAGGCGGCTCACTGGACTGGTATTGACGCTGAGGTGCGAAAGCGTGGGG |
| P-asv4310 | GCAAGCGTTAATCGGAATGACTGGGCGTAAAGGGCGAGTAGGCGGTTGTATAGGTGTGGTGTGAAAGACACGGGCTTAACCTGTGGGTGTCACCGCAAACGGTATGACTAGAGTGCAGTAGAGGGGAGTGGAATTTCCGGAGGAGCGGTGAAATGCGTAGATCTCGGAAGGAACACCGGTGGCGAAGGCGACTCTCTGGACTGTAACTGACGCTGAGGCGCGAAGGCGTGGGG |
| P-asv4311 | TCAAGTGATGGCCGATCTTATTGGGCCTAAAGCGTCCGTAGCCTGCCGCACGGGTCCGTCGGGAAATCCGCCCGCTCAACGGGCGGACGTCCGGCGGAAACCATACGGCTTGGGACCAGAGGACCCGAGGGGTACGTCGGGGGTAGGAGTGAAATCCCGAGATCCTCGACGGACCGCCGGTGGCGAAAGCGCCTCGGGAAGATGGATCCGACGGTCAGGGACGAAAGCTTGGGT |
| P-asv4312 | CCGAGTGATGGCCGATATTATTGGGCCTAAAGCGTCCGTAGCTGGCCGGACAAGTCCGTTGGGAAATCTGCTCGCTTAACGAGCAGGCGTCCAGCGGAAACTGTTCGGCTTGGGACCGGAAGACCTGAGGGGTACGTCCGGGGTAGGAGTGAAATCCCGTAATCCTGGACGGACCACCGGTGGCGAAAGCGCCTCAGGAGGACGGATCCGACAGTGAGGAACGAAAGCTAGGGT |
| P-asv4313 | GCGAGCGTTGTCCGGAATCATTGGGTGTAAAGGGTGCGTAGGCGGATTGCTAAGTCTGGGGTGAAATCCTGCCGCTTAACGGTAGAATGGCCCCCGATACTGGCAATCTTGAGTACAGAAGAGGTCGGTGGAATTCGTAGTGTAGCGGTGAAATGCATAGATATTACGAAGAACACCAGTGGCGAAGGCGGCCGGCTGGTCTGTAACTGACGCTGAGGCACGAAAGCGTGGGG |
| P-asv4314 | GCTAGCGTTGTTCGGAATTACTGGGCGTAAAGGGCGTGCAGGCGGATCCGCCAGTCAGAGGTGAAAGCCCCGGGCTTAACCTGGGAATTGCCTTTGATACTGTGGATCTAGAGTTCGGGAGAGGGTCGTGGAATTCCCAGTGTAGAGGTGAAATTCGTAGATATTGGGAAGAACACCAGTGGCGAAGGCGGCGACCTGGACCGATACTGACGCTGAGGCGCGAAAGCGTGGGG |
| P-asv4315 | GCAAGCGTTGTCCGGAATTATTGGGCGTAAAGAGCTCGTAGGCGGTTTGTCGCGTCTGCTGTGAAAACCCGAGGCTCAACCTCGGGCCTGCAGTGGGTACGGGCAGACTAGAGTGCGGTAGGGGAGACTGGAATTCCTGGTGTAGCGGTGGAATGCGCAGATATCAGGAGGAACACCGATGGCGAAGGCAGGTCTCTGGGCCGCAACTGACGCTGAGGAGCGAAAGCATGGGG |
| P-asv4316 | GCAAGCGTTAATCGGAATTACTGGGCGTAAAGCGTGCGTAGGCGGTCTTTTAAGTTGGATGTGAAATCCCCGGGCTTAACCTGGGAACTGCATTCAAGACTGGGAGACTGGAGTCTGGGAGAGGATGGTGGAATTTCCGGTGTAGCGGTGAAATGCGTAGATATCGGAAGGAACATCAGTGGCGAAGGCGACCATCTGGCCTAAGACTGACGCTGAGGCACGAAAGCGTGGGG |
| P-asv4317 | GCAAGCGTTATCCGGAATCACTGGGCGTAAAGGGTGCGTAGGTGGTTTTGTAAGTCTGAGGTAAAAGGTCACGGCTCAACCGTGGTAAGCCTTGGAAACTGCAAAACTTGAATGCAGGAGAGGAAAGTGGAATTCCTAGTGTAGCGGTGAAATGCGTAGATATTAGGAGGAACATCCATGGCGAAGGCAGCTTTCTGGACTGTTATTGACACTGAGGCACGAAAGCGTGGGG |
| P-asv4318 | GCGAGCGTTAATCGGATTTACTGGGCGTAAAGGGCGCGTAGGCGGTGAGATATGTGTGATGTGAAAGCCCCGGGCTTAACCTGGGAAGTGCATTGCAAACGGTCTGACTGGAGTATATGAGAGGGTGGCGGAATTTCCGGTGTAGCGGTGAAATGCGTAGATATCGGAAGGAACGTCAATGGCGAAGGCAGCCACCTGGCATAATACTGACGCTGAGGCGCGAAAGCGTGGGG |
| P-asv4319 | GCAAGCGTTGTTCGGAATTACTGGGCGTAAAGGGTGCGTAGGCGGTGCTCTAAGTTTGGTGTGAAATCTCCCGGCTCAACTGGGAGGGTGCGCCGGAAACTGGAGCGCTTGAGTGTGGGAGAGGAAAGCGGAATTCCTGGTGTAGCGGTGAAATGCGTAGAGATCAGGAGGAACACCTGCGGTGTAGACGGCTTTCTGGACCATAACTGACGCTGAGGCACGAAAGCGTGGGT |
| P-asv4320 | GCTAGCGTTGTTCGGAATTACTGGGCGTAAAGCGCGCGTAGGCGGATCTTTAAGTCAGAGGTGAAATCCTGGAGCTCAACTCCAGAACTGCCTTTGATACTGGAGATCTTGAGTCCGGAAGAGGTTAGTGGAATTCCCAGTGTAGAGGTGAAATTCGTAGATATTGGGAAGAACACCAGTGGCGAAGGCGGCTAACTGGTCCGGTACTGACGCTGAGGTGCGAAAGCGTGGGG |
| P-asv4321 | GCGTGCGTTATTCATACGAACTTGGCGTAAAGCGTCTTTAGGTTGTTTAGCGTAGTTAGCATCTAAAGCTCAGAACGTAATTCTGGAATGTGTGCTAAGACGGTTATTCTTGAGTTGATTTAGGGTTAGTAGTATTTCTTCTCGAGGGTTAAAATTCATAGAAAGAAGAAAGAGGGTCTAGGGTGAAGACAACTAACCGATTTCAACTGACACCAAAGGACTACAGCGTAGGG |
| P-asv4322 | GCTAGCGTTGTTCGGAATTACTGGGCGTAAAGCGCACGTAGGCGGACTATTAAGTCAGAGGTGAAATCCCACGGCTCAACCGTGGAACGGCCTTTGATACTGGTAGTCTTGAGTTCGAGAGAGGTGAGTGGAATTCCGAGTGTAGAGGTGAAATTCGTAGATATTCGGAGGAACACCAGTGGCGAAGGCGGCTCACTGGCTCGATACTGACGCTGAGGTGCGAAAGCGTGGGG |
| P-asv4323 | GCAAGCGTTAATCGGAATTACTGGGCGTAAAGCGTACGTAGGCGGCGTGTTAAGCAAGATGTGAAAGCCCCGGGCTCAACCTGGGAATGGCATTTTGAACTGGCACGCTAGAGTCCTGAAGAGGGTGGTAGAATTTCCAGTGTAGCGGTGAAATGCGTAGATATTGGAAGGAATACCGGTGGCGAAGGCGGCCACCTGGTCAGAGACTGACGCTGAGGTACGAAAGCGTGGGG |
| P-asv4324 | GTTAGCGTTGTTCGGAATTACTGGGCGTAAAGCGCACGTAGGCGGACCATCAAGTTGGGGGTGAAATCCCGGGGCTCAACCCCGGAACTGCCTTCAAAACTGGTGGTCTGGAGTTCGAGAGAGGTGAGTGGAATTCCGAGTGTAGAGGTGAAATTCGTAGATATTCGGAGGAACATCAGTGGCGAAGGCGGCTCACTGGCTCGATACTGACGCTGAGGTGCGAAAGTGTGGGT |
| P-asv4325 | GTTAGCGTTGTTCGGAATTACTGGGCGTAAAGCGCACGTAGGCGGATCAGAAAGTTGGGGGTGAAATCCCGGGGCTCAACCCCGGAACTGCCTTCAAAACTTCTGGTCTGGAGTTCGAGAGAGGTGAGTGGAATTCCGAGTGTAGAGGTGAAATTCGTAGATATTCGGAGGAACATCAGTGGCGAAGGCGGCTCACTGGCTCGATACTGACGCTGAGGTGCGAAAGTGTGGGT |
| P-asv4326 | CCGAGCGTTGTTCGGAATCACTGGGCGTAAAGGGAGCGTAGGCGGCGTGGTAAGTCAGATGTGAAATCCCGGGGCTCAACCCCGGAACTGCATCCGATACTGCCGTGCTAGAGTAATGGAGGGGTAGCTGGAATTCTCGGTGTAGCAGTGAAATGCGTGGATATCGAGAGGAACACTCGTGGCGAAAGCGAGCTACTGGACATTTACTGACGCTGAGGCTCGAAGGCCAGGGG |
| P-asv4327 | GCAAGCGTTGTCCGGAATTATTGGGCGTAAAGAGCTCGTAGGCGGTTTGTCGCGTCTGCTGTGAAAGCCCGGGGCTTAACCCCGGGTGTGCAGTGGGTACGGGCAGACTTGAGTGCAGTAGGGGAGACTGGAACTCCTGGTGTAGCGGTGAAATGCGCAGATATCAGGAAGAACACCGATGGCGAAGGCAGGTCTCTGGGCTGTTACTGACGCTGAGGAGCGAAAGCATGGGG |
| P-asv4328 | GCAAGCGTTAATCGGAATTACTGGGCGTAAAGCGCGCGTAGGTGGCTTGATAAGTTGGATGTGAAATCCCCGGGCTCAACCTGGGAACTGCATCCAAAACTGTCTGGCTAGAGTGTGGTAGAGGGTAGTGGAATTTCCAGTGTAGCGGTGAAATGCGTAGATATTGGAAGGAACACCAGTGGCGAAGGCGACTACCTGGACTAACACTGACACTGAGGTGCGAAAGCGTGGGG |
| P-asv4329 | GCAAGCGTTACTCGGAATTACTAGGCGTAAAGGGCAGGTAGGTGGTTTGATAAGTCTGTTGTGTCAGCTCCTGGCTTAACTGGGAGAGGTCAACGGAAACTATCAGGCTAGAGTGTGGGAGAGGATACTGGAATTCCTGGTGTAGCGGTGAAATGCGCAGATATCAGGAGGAACACCAATGGCGAAAGCAAGTATCTGGACCACTACTGACACTAAGCTGCGAAAGCTAGGGG |
| P-asv4330 | GCTAGCGTTGTTCGGAATTACTGGGCGTAAAGCGCACGTAGGCGGACATTTAAGTCAGGGGTGAAATCCCAGAGCTCAACTCTGGAACTGCCTTTGATACTGGGTGTCTTGAGTGTGGAAGAGGTCAGTGGAATTGCGAGTGTAGAGGTGAAATTCGTAGATATTCGCAGGAACACCAGTGGCGAAGGCGGCTGACTGGTCCACAACTGACGCTGAGGTGCGAAAGCGTGGGG |
| P-asv4331 | CCAAGCGTTATCCGGAATTACTGGGCGTAAAGAGTTGCGTAGGCGGCATAGTAAGCAGATAGTGAAAGCGTGTGGCTCAACCATACTCACATTATCTGAACTGCTAAGCTAGAGGATGAGAGAGGTCACTGGAATTCCTAGTGTAGGAGTGAAATCCGTAGATATTAGGAGGAACACCAATGGCGTAGGCAGGTGACTGGCTCATTCCTGACGCTCAGGCACGAAAGCGTGGGT |
| P-asv4332 | GCAAGCGTTGTCCGGAATTATTGGGCGTAAAGCGCTCGCAGGCGGTCTTTTAAGTCTGATGTGAAAGCCCACGGCTTAGCCGTGGAGGGTCATTGGAAACTGGAGGACTTGAGTACAGAAGAGGAGAGTGGAATTCCACGTGTAGCGGTGAAATGCGTAGATATGTGGAGGAACACCAGTGGCGAAGGCGACTCTCTGGTCTGTAACTGACGCTGAGGAGCGAAAGCGTGGGG |
| P-asv4333 | GCGAGCGTTAATCGGATTTACTGGGCGTAAAGCGTGCGTAGGCGGCTTCGTAAGTTGGATGTGAAATCCCCGAGCTTAACTTGGGAATTGCATTCAAAACTGCGAGGCTAGAGTATGGGAGAGGATGGTAGAATTCCAGGTGTAGCGGTGAAATGCGTAGAGATCTGGAGGAATACCGATGGCGAAGGCAGCCATCTGGCCTAATACTGACGCTGAGGTACGAAAGCATGGGG |

| | |
|---|---|
| P-asv4334 | GCAAGCGTTATCCGGAATCACTGGGTTTAAAGGGTGCGTAGGCGGTCATGTAAGTCAGGGGTGAAAGCCCGGCGCTCAACGTCGGAATTGCCTTTGATACTGCATGACTTGAATCAGGTTGAGGTGGGCGGAATGTGACATGTAGCGGTGAAATGCATAGATATGTCATAGAACACCAATTGCGAAGGCAGCTCACTGGCCCTGAATTGACGCTGAGGCACGAAAGCGTGGGG |
| P-asv4335 | GCAAGCGTTGTCCGGATTTATTGGGCGTAAAGCGAGCGCAGGCGGTTCTTTAAGTCTGATGTGAAAGCCCTCGGCTCAACCGAGGAAGGTCATTGGAAACTGGAGAACTTGAGTGCAGAAGAGGAGAGTGGAATTCCATGTGTAGCGGTGAAATGCGTAGATATATGGAGGAACACCAGTGGCGAAGGCGACTCTCTGGTCTGTAACTGACGCTGAGGCTCGAAAGCGTGGGG |
| P-asv4336 | GCTAGCGTTGTTCGGATTTACTGGGCGTAAAGCGCACGTAGGCGGATTGTTAAGTGAGGGGTGAAATCCCGGGGCTCAACCTCGGAACTGCCTTTCATACTGGCAATCTAGAGTCCGGAAGAGGTGAGTGGAACTCCTAGTGTAGAGGTGGAATTCGTAGATATTAGGAAGAACACCAGTGGCGAAGGCGGCTCACTGGTCCGGTACTGACGCTGAGGTGCGAAAGCGTGGGG |
| P-asv4337 | GCAAACGTTGCTCGGAATCATTGGGCGTAAAGCGCACGTAGGCGGCCTGATAAGTCGGATGTGAAATCCCTCGGCTTAACCGAGGAAGTGCATTCGAAACTGTTGGGCTCGAGTATGGAAGAGGGTCGCAGAATTCCCGGTGTAGAGGTGAAATTCGTAGATATCGGGAGGAATACCGGTGGCGAAGGCGGCGACCTGGGCCAATACTGACGCTGAGGTGCGAAAGCGTGGGG |
| P-asv4338 | GCTAGCGTTGTTCGGAATTACTGGGCGTAAAGCGCACGTAGGCGGATTTTTAAGTCAGGGGTGAAATCCCAAGGCTCAACCTTGGAACTGCCTTTGATACTGGAAGTCTCGAGATCGGGAGAGGTGAGTGGAATTCCTAGTGTAGAGGTGAAATTCGTAGATATTAGGAAGAACACCAGTGGCGAAGGCGGCTCACTGGCCCGATACTGACGCTGAGGTGCGAAAGCGTGGGG |
| P-asv4339 | GCGAGCGTTAATCGGAATTACTGGGCGTAAAGGGTGCGTAGGCGGTTTATTAAGTTATCTGTGAAAGACCTGGGCTTAACCTTGGGATGTCAGATGATACTGATGAACTAGAGTATAGGAGAGGGCAGTGGAATTTCCGGTGTAGCGGTGAAATGCGTAGATATCGGAAGGAACACCGGTGGCGAAGGCGGCTGCCTGGCCTGATACTGACGCTGAGGCACGAAAGCGTGGGT |
| P-asv4340 | GCGAGCGTTGTTCGGAATTACTGGGCGTAAAGCGCGCGCAGGCGGCTTCTTAAGTCTATGGGTGAAAGACCGGGGCTCAACCCCGTGTCAGCCCGGGATACTGGGAAGCTAGGGACATGTAGGGGCGAGCGGAATTCCGGGTGTAGCGGTGGAATGCGTAGAGATCCGGAAGAACACCGGGGGCGAAGGCGGCTCGCTGGGCATGGTCCGACGCTGAGGCGCGAAAGCGTGGGG |
| P-asv4341 | GCAAGCGTTGTCCGGATTTACTGGGTGTAAAGGGCGTGTAGGCGGAGATGCAAGTTGGGAGTGAAATCCCGGGGCTCAACCCCGGAACTGCTTTCAAAACTGTATCCCTTGAGTGTCGGAGAGGCAAGCGGAATTCCTAGTGTAGCGGTGAAATGCGTAGATATTAGGAGGAACACCAGTGGCGAAGGCGGCTTGCTGGACGATAACTGACGCTGAGGCGCGAAAGCGTGGGG |
| P-asv4342 | CCAAGCGTTAGGCGGAATCACTGGGCTTAAGGCGCGTGTAGGTGGGTCGTTAAGTGTCCGCGTGAAATCCCACGGCTCAACCGTGGAACTGCCGGGCAGACTGGCGATCTTGAGGCATTCAGGGGCGAGCGGAACAAATGGTGGAGCGGTGAAATGCGTAGATATCATTTGGAACGCCAATGGTGAAAACAGCTCGCTGGGAATGATCTGACACTGAGACGCGAAAGCCAGGGG |
| P-asv4343 | CCGAGCGTTGTCCGGATTTATTGGGCGTAAAGCGAGCGCAGGTGGTTTAATAAGTCTGATGTAAAAGGCAGTGGCTCAACCATTGTGTGCATTGGAAACTGTTAGACTTGAGTGCAGTAGAGGAGAGTGGAATTCCATGTGTAGCGGTGAAATGCGTAGATATATGGAGGAACACCGGTGGCGAAAGCGGCTCTCTGGACTGTAACTGACACTGAGGCTCGAAAGCGTGGGT |
| P-asv4344 | GCAAGCGTTAATCGGAATTACTGGGCGTAAAGCGCGCGTAGGTGGTTCGTTAAGTTGGATGTGAAATCCCCGGGCTCAACCTGGGAACTGCATCCAAAACTGGCGAGCTAGAGTAGGGCAGAGGGTGGTGGAATTTCCTGTGTAGCGGTGAAATGCGTAGATATAGGAAGGAACACCAGTGGCGAAGGCGACCACCTGGACTGATACTGACACTGAGGTGCGAAAGCGTGGGG |
| P-asv4345 | GCAAGCGTTGTCCGGAATTATTGGGCGTAAAGAGCTCGTAGGCGGTTTGTCGCGTTTGCTGTGAAAACGCGAGGCTCAACCTCGCGCCTGCAGAGGGTACGCGCAAACTAGAGTGCGGCAGGGGAGATTGGAATTCCTGGTGTAGCGGTGGAATGCGCAGATATCAGGAGGAACACCGATGGCGAAGGCAGATCTCTGGGCCGTAGCTGACGCTGAGGAGCGAAAGCATGGGG |
| P-asv4346 | GCAAGCGTTGTCCGGAATTATTGGGCGTAAAGAGCTCGTAGGCGGTGTGTCGCGTCTGCTGTGAAAACTCAGGGCTCAACTCTGAGCTTGCAGTGGGTACGGGCACACTAGAGTGCTGTAGGGGAGACTGGAATTCCTGGTGTAGCGGTGAAATGCGCAGATATCAGGAGGAACACCGGTGGCGAAGGCGGGTCTCTGGGCAGTTACTGACGCTGAGGAGCGAAAGCATGGGG |
| P-asv4347 | GCAAGCGTTATCCGGATTTATTGGGCGTAAAGCGCTCGCAGGCGGTCTTTTAAGTCTGATGTGAAATCTCGAGGCTCAACCTCGAGCGGTCATTGGAAACTGGGAGACTTGAGTGCAGAAGAGGAGAGTGGAATTCCACGTGTAGCGGTGAAATGCGTAGAGATGTGGAGGAACACCAGTGGCTAAGGCGACTCTCTGGTCTGTAACTGACGCTGAGGAGCGAAAGCGTGGGG |
| P-asv4348 | GCAAGCGTTGTTCGGATTTACTGGGCGTAAAGGGCGCGTAGGCGGCCTGTGCAAGTCAGTTGTGAAATCTCCGAGCTTAACTCGGAAAGGTCAACTGATACTGCTCGGCTAGAGTGCGGAAGGGGCAACTGGAATTCTTGGTGTAGCGGTGAAATGCGTAGATATCAAGAGGAACACCTGCGGCGAAGGCGGGTTGCTGGGCCGACACTGACGCTGAGGCGCGAAAGCTAGGGG |
| P-asv4349 | GCAAGCGTTGTTCGGAATTACTGGGCGTAAAGGGCGCGTAGGCGGCCTGTTAAGTCGAACGTGAAATCCCCGGGCTCAACCCGGGAACTGCGTCCGATACTGGCAGGCTTGAGTTCGGGAGAGGGATGCGGAATTCCAGGTGTAGCGGTGAAATGCGTAGATATCTGGAGGAACACCGGTGGCGAAGGCGGCATCCTGGACCGAGATTGACGCTGAGGCGCGAAAGCCAGGGG |
| P-asv4350 | GCAAGCGTCGTCCGGAATCATTGGGCGTAAAGCGCGCGTAGGCGGCCCGGTAAGTCCGCTCTGAAAGTCAAAGGCTCAACCTTTGAAAGCGGGTGGATACTGCCGGGCTAGAGTCCGGAAGAGGCGAGTGGAATTCCTGGTGTAGCGGTGGAATGCGCAGATATCAGGAGGAACACCAACGGCGAAGGCAGCTCGCTGGGACGGTACTGACGCTAAGGCGCGAAAGCGTGGGG |
| P-asv4351 | GCGAGCGTTAATCGGAATTACTGGGCGTAAAGGGTGCGTAGGTGGTTGAATAAGTTAGGTGTGAAATTCCCGGGCTTAACCTGGGGACTGCGCCTGATACTGTTGAGCTAGAGTACTGTAGAGGGGAGTGGAATTTCCGGAGTAGCGGTGAAATGCGTAGATATCGGAAGGAACACCAGTGGCGAAAGCGACTTTCTGGACAGATACTGACACTGAGGCACGAAAGCGTGGGG |
| P-asv4352 | GCAAGCGTTGTCCGGAATTATTGGGCGTAAAGAGCTCGTAGGCGGTTTGTCGCGTCTGCTGTGAAAACTCAGGGCTTAACCCTGAGCCTGCAGTGGGTACGGGCAGACTAGAGTGTGGTAGGGGAGATCGGAATTCCTGGTGTAGCGGTGGAATGCGCAGATATCAGGAGGAACACCGGTGGCGAAGGCGGATCTCTGGGCCATAACTGACGCTGAGGAGCGAAAGCATGGGG |
| P-asv4353 | GCTAGCGTTGCTCGGAATCACTGGGCGTAAAGCGCACGTAGGCGGCTTTTTAAGTCAGGGGTGAAATCCTGGAGCTCAACTCCAGAACTGCCTTTGATACTGAGAGGCTAGAGTCCGGAAGAGGTAAGTGGAACTGCGAGTGTAGAGGTGAAATTCGTAGATATTCGCAAGAACACCAGTGGCGAAGGCGGCTTACTGGTCCGGTACTGACGCTGAGGTGCGAAAGCGTGGGG |
| P-asv4354 | GCCAGCGTTGCTCGGAATTACTGGGCGTAAAGGGTTCGTAGGCGGTGCGGCAAGTTGGGAGTGAAATCTCTGGGCTTAACCCAGAGGCTGCTTTCAAAACTGCCGTGCTCGAGTGCGGGAGAGGTGAGTGGAACTGCGAGTGTAGAGGTGAAATTCGTAGATATTCGCAAGAACACCAGTGGCGAAGGCGGCTCACTGGCCCGATACTGACGCTGAGGTGCGAAAGCGTGGGG |
| P-asv4355 | GCAAGCGTTGTCCGGAATCATTGGGCGTAAAGAGCGTGTAGGCGGCCTGGTAAGTCCGCTGTGAAAGTCCAGGGCTCAACCCTGGAATGCCGGTGGAAACTGTCAGGCTCGAGTCCGGAAGAGGCGAGTGGAATTCCTGGTGTAGCGGTGAAATGCGCAGATATCAGGAGGAACACCAATGGCGAAGGCAGCTCGCTGGGACGGTACTGACGCTGAGACGCGAAAGCGTGGGG |
| P-asv4356 | GCGAGCGTTGTTCGGAATTACTGGGCGTAAAGGGCGCGCAGGCGGCCGATCAAGTCAGGCGTGAAAGCCCCGGGCTTAACCTGGGAATTGCGCTTGGGACTGGTTGGCTTGAGTTCGGGAGAGGAGAGTGGAATTCCCAGTGTAGAGGTGAAATTCGTAGATATTGGGAAGAACACCGGTGGCGAAGGCGGCTCTCTGGACCGATACTGACGCTCAGGCGCGACAGCGTGGGG |
| P-asv4357 | GCAAGCGTTGTTCGGAATTACTGGGCGTAAAGGGCGCGTAGGCGGCAGGGCAAGTCAGGTGTGAAAGCCCCGGGCTTAACTCAGGAACTGCATCTGAAACTGCTTTGCTTGAGTCCTGGAGGGGGTAGTGGAATTCCCAGTGTAGCGGTGAAATGCGTAGATATTGGGAGGAACACCGGTGGCGAAGGCGGCTACCTGGACAGTGACTGACGCTGAGGCGCGAAAGCTAGGGG |
| P-asv4358 | GCAAGCGTTGTTCGGAATTACTGGGCGTAAAGGGCGCGTAGGCGGCCTGTTAAGTCGAACGTGAAACCCCCGGGCTCAACCCGGGAACTGCGTCCGATACTGGCAGGCTTGAATCCGGGAGAGGGATGTGGAATTCCAGGTGTAGCGGTGAAATGCGTAGATATCTGGAGGAACACCGGTGGCGAAGGCGGCATCCTGGACCGGTATTGACGCTGAGGCGCGAAAGCCAGGGG |
| P-asv4359 | GCGAGCGTTGTCCGGATTTATTGGGCGTAAAGAGCTCGTAGGCGGCTTGTCGCGTCGACTGTGAAAACCCGCGGCCCAACCGTGCGGTCTGCAGTCGATACGGGCAGGCTAGAGTTCGGTAGGGGAGACTGGAATTCCTGGTGTAGCGGTGAAATGCGTAGATATCGGGAGGAACACCAGTGGCGAAAGCGGCTACCTGGCCCACTCTTGACGCTGAGGCGCGAAAGCTAGGGG |
| P-asv4360 | GCGAGCGTTGTCCGGAATCACTGGGCGTAAAGGGCGCGTAGGTGGCGAGATAAGCGTGTGGTGAAAGCCCGGGGCTCAACCCCGTGTCTGCCATGCGGACTGTTTCGCTCGAGCGTAGTAGAGGCAGGTGGAATTCCGGGTGTAGCGGTGGAATGCGTAGAGATCCGGAAGAACACCGGTGGCGAAGGCGGCCTGCTGGGCTACTGCTGACACTGAGGCGCGACAGCGTGGGG |
| P-asv4361 | GCGAGCGTTAATCGGAATTACTGGGCGTAAAGCGTGCGCAGGCGGTTCGCTGTGTCCGATGTGAAAGCCCCGGGCTCAACCTGGGAATGGCATGGGAAACTGGCGGACTGGAGTGTGGCAGAGGGGGGTGGAATTCCACGTGTAGCAGTGAAATGCGTAGAGATGTGGAGGAACACCGATGGCGAAGGCAGCCCCCTGGGCTAACACTGACGCTCAGGCACGAAAGCGTGGGG |
| P-asv4362 | GCAAGCGTTGTTCGGAATTACTGGGCGTAAAGCGCGATTAGGCGGTTTGAAAAGTCAGATGTGAAAGCCCCGGGCTCAACTTGGGAACTGCATTTGATACTGTCAGACTAGAGTACAGAAGAGGTGACTGGAATTCCCAGTGTAGCGGTGAAATGCGTAGATATTGGGAGGAACACCAGATGCGAAGGCGGGTCACTGGGCTGATACTGACGCTGATATTGCGAAAGCCAGGGG |
| P-asv4363 | GCAAACGTTGTCCGGATTTATTGGGTGTAAAGGGCATGTAGGTGGTTTTGTAAGTCAAAGGTGAAATGGTTTGGCTCAACCAAATCATTGCCTTTGAAACTGCAGAACTTGAGTACGGAAGAGGAGAGCGGAATTCCCAGTGTAGCGGTGAAATGCGTAGATATTGGGAGGAACACCGGTGGCGAAAGCGGCTCTCTGGTCCGAAACTGACACTGAGATGCGAAAGCCAGGGG |
| P-asv4364 | GCAAGCGTTATCCGGATTCACTGGGTTTAAAGGGTGCGTAGGTGGGTTGGTAAGTCAGTGGTGAAATCTCCAGGCTTAACTTGGAAACTGCCATTGATACTATCAATCTTGAATACCGTGGAGGTCAGCGGAATATGTCATGTAGCGGTGAAATGCTTAGATATGCGGAGGAACACCGATGGCGAAGGCAATCCCCTGGGCCTGCACTGACGCCCATGCACGAAAGCGTGGGG |
| P-asv4365 | GCGAGCGTTGTCCGGAATCACTGGGCGTAAAGGGTGAGTAGGTGGCCTGTTAAGCGTGTGGTGAAAGCCTGGGGCTCAACCCCAGGTCTGCCGTGCGAACTGACAGGCTTGAGCAGAGTAGAGGCGAGTGGAATTTCGGGTGTAGCGGTGGAATGCGTAGATATCCGAAAGAACACCGGTGGCGAAGGCGACTCGCTGGGCTTTTGCTGACACTGAATCACGACAGCGTGGGG |
| P-asv4366 | GCAAGCGTTGTCCGGAATTATTGGGCGTAAAGCGCGCGCAGGCGGTTGATTAAGTCAGATGTGAAAGGCTACGGCTCAACCGTAGAGTTGCATTTGAAACTGGTCAGCCTGAGTGCAGGAGAGGAGAGTGGAATTCCCGGTGTAGCGGTGGAATGCGTAGATATCGGGAGGAACACCAGTGGCGAAGGCGGCTCTCTGGCCTGCAACTGACGCTGAGGCGCGAAAGCGTGGGG |
| P-asv4367 | GCGAGCGTTGTCCGGATTTATTGGGCGTAAAACGCTTGTAGGCGGCCACGCTGCGTCGTGCGTGAAATCTCCCGGCTTAACTGGGAGGGGTCGTGCGATACGAGCGGGCTCGAGGCAGGTAGGGGAAGCGGAATTCCCGGTGTAGTGGTGGAATGCGTAGATATCGGGAGGAACACCAGTGGCGAAGGCGGCTTCCTGGACCTGATCTGACGCTGAGGAGCGAAAGCGTGGGG |
| P-asv4368 | GCAAGCGTTGTTCGGAATTACTGGGCGTAAAGGGCTCGTAGGTGGCCAACTAAGTCAGACGTGAAATCCCTCGGCTTAACCGGGGAACTGCGTCTGATACTGGATGGCTTGAGTTTGGGAGAGGGACGCGGAATTCCAGGTGTAGCGGTGAAATGCGTAGATATCAGGAGGAACACCGGTGGCGAAGGCGGGTCTCTGGGCCGTAACAGACGCTGAGGAGCGAAAGCGTGGGG |
| P-asv4369 | GCTAGCGTTGCTCGGAATCACTGGGCGTAAAGGGCGCGTAGGCGGGGCGGCAAGTCGGTCGTGAAATCCTCCGGCTCAACCGGAGAACGTCGGCTGAAACTGCCGTTCTTGAGGGATGCAGAGGAGACTGGAATTCCCGGTGTAGCGGTGAAATGTGTAGAGATCGGGAGGAACACCGGTGGCGAAGGCGGGTCTCTGGGCATCTCCTGACGCTGAGGCGCGAAAGCGTGGGG |
| P-asv4370 | GCAAGCGTTGCTCGGAATCATTGGGCGTAAAGCGCGTGTAGGCGGGACAGTATGTCGGGTGTGAAAGCCCAGGGCTCAACCCTGGAAGTGCGCCCGAAACTGCTGTTCTTGAGGTCGGGAGGGGGTCGTGGAATTCCCGGTGTAGCGGTGAAATGCGTAGATATCGGGAGGAACACCTGTGGCGAAGGCGGCGACCTGGACCGAATCTGACGCTGAGACGCGAAAGCGTGGGG |
| P-asv4371 | GCAAACGTTGCTCGGAATCATTGGGCGTAAAGCGCACGTAGGCGGTCTGTTATGTCGGATGTGAAAGCCCTTGGCTTAACTAAGGAAGTGCATCCGAAACTGGCAGGCTAGAGTACTAAAGAGGGTCGCGGAATTCCCGGTGTAGAGGTGAAATTCGTAGATATCGGGAGGAACACCGGTGGCGAAGGCGGCGACCTGGGTAGATACTGACGCTGAGGTGCGAAAGCGTGGGG |
| P-asv4372 | GCGAGCGTTGTTCGGAGTTACTGGGCGTAAAGGGTGTGTAGGCGGTTGTTTAAGTTTGGTGTGAAATCTCCCGGCTCAACCGGGAGGGTGCGCCGAATACTGAATGACTTGGAGTGCGGGAGAGGAGAGTGGAATTCCTGGTGTAGCGGTGAAATGCGTAGATATCAGGAGGAACACCGGTGGTGTAGACGGCTCTCTGGACCGTAACTGACGCTGAGACACGAAAGCGTGGGT |

| | |
|---|---|
| P-asv4373 | CCAAGCGTTGTTCGGATTTACTGGGCGTAAAGGGTGCGTAGGTGGCGAGGTAAGTTCGATGTGAAATCTCCGAGCTCAACTCGGAAATGGCATTGAATACTATCCCGCTTGAGGGTCGGAGGGGAGACTGGAATTCTCGGTGTAGCAGTGAAATGCGTAGATATCGAGAGGAACACCAGTGGCGAAGGCGAGTCTCTGGACGACTCCTGACACTGAGGCACGAAAGCTGGGGG |
| P-asv4374 | GCAAACGTTGTTCGGAATTACTGGGCGTAAAGCGCGTGTAGGCGGCTATGTAAGTCGGGCGTGAAATCCCACGGCTCAACCGTGGAAGTGCACCCGAAACTGCGTAGCTAGAGTCCTGGAGAGGAAGGTGGAATGCTTGGTGTAGAGGTGAAATTCGTAGATATCAAGCGGAACACCGGTGGCGAAGGCGGCCTTCTGGACAGTGACTGACGCTGAGACGCGAAAGCGTGGGG |
| P-asv4375 | GCGAGCGTTGTCCGGAATCACTGGGCGTAAAGGGCGCGTAGGCGGCCGTGTAAGTCACTTGTGAAATCCCGGGGCTCAACTCCGGGGCTGCAGGTGAGACTGCATGGCTAGAGCACGGTAGAGGCGAGTGGAATTCCCGGTGTAGCGGTGGAATGCGTAGATATCGGGAAGAACACCGGTGGCGAAGGCGGCTCGCTGGGCCGTTGCTGACGCTGAGGCGCGAAAGCGTGGGG |
| P-asv4376 | GCAAGCGTTGTTCGGAATTACTGGGCGTAAAGAGTGTGTAGGCGGTGTTCTAAGTCTGTCGTGAAATCTCCCGGCTTAACCGGGAGGGTGCGGCGGAAACTGGAATGCTTGAGGGTGGGAGAGGTAAGCGGAATTCCTGGTGTAGCGGTGAAATGCGTAGATATCAGGAGGAACACCTGTGGTGTAGACAGCTTACTGGACCACTTCTGACGCTGAGACACGAAAGCGTGGGT |
| P-asv4377 | GCGAGCGTTGTTCGGATTTACTGGGCGTAAGGGGCGCGTAGGCGGCGATGAAAGTCAGGCGTGAAAGCCCCGGGCTCAACCTGGGAGGTGCGCTTGGGACTACATTGCTGGAGTTCGGGAGAGGAGCGTGGAATTCCCAGTGTAGAGGTGAAATTCGTAGATATTGGGAAGAACACCGGTGGCGAAGGCGGCGCTCTGGACCGATACTGACGCTGAGGCGCGAAAGCGTGGGG |
| P-asv4378 | GCAAGCGTTGTTCGGATTTACTGGGCGTACAGGGAGCGTAGGCGGTTTGATAAGCCCTCTGGGAAATCTTCGGGCTTAACCCGGAAAGTTCAGAGGGGACTGCCAAGCTAGAGGACGGGAGAGGAGCGCGGAATTCCCGGTGTAGCGGTGAAATGCGTAGAGATCGGGAGGAAGGCCGGTGGCGAAGGCGGCGCTCTGGAACGTATCTGACGCTGAGGCTCGAAAGCGTGGGG |
| P-asv4379 | GCCAGCGTTGCTCGGAATTACTGGGCGTAAAGGGTCTGTAGGCGGTGCGGCAAGTCGGAAGTGAAATCCCAGGGCTTAACTCTGGGGCTGCTCCCGAAACTGCTGTGCTAGAGTGTGAGATAGGCGAGTGGAATTGCAGGTGTAGCGGTGAAATGCGTAGATATCTGCAGGAACATCCGAGGCGAAAGCGGCTCGCTAGATCACAACTGACGCTGAGAGACGAAAGCTAGGGG |
| P-asv4380 | GCAAGCGTTATTCGGAATTATTGGGCGTAAAGCGCGCGTAGGCGGCTGCGCAAGTCGGGTGTGAAACCCCGGGGCTCAACCCCGGAACTGCATCCGATACTGCGTGGCTAGAGGTAGGTAGGGGAGATCGGAATTCCTGGTGTAGCGGTGAAATGCGCAGATATCAGGAGGAACACCGGTGGCGAAGGCGGATCTCTGGGCCTTACCTGACGCTCAGGCGCGAAAGCTAGGGG |
| P-asv4381 | GCAAGCGTTGTCCGGATTTACTGGGCGTAAAGAGCGCGCAGGCGGTCGTACAAGTCGCGTGTGAATGCCCCCGGCTCAACTGGGGAGGGTCACGCGATACTGTTCGACTCGAAGGCAGGAGAGGGTAGTGGAATTCCCGGTGTAGTGGTGAAATGCGTAGATATCGGGAGGAACACCAGTGGCGAAGGCGACTACCTGGCCTGTTCTTGACGCTGAGGCGCGAAAGCTAGGGT |
| P-asv4382 | GCAAGCGTTGTCCGGATTTACTGGGCGTAAAGGGCTTGTAGGCGGTTCGTTAAGTCCGGTGTGAAATCTCCCGGCTCAACTGGGAGGAGCCATCGGAAACTGGCGAGCTAGAGGTGGGCAGAGGAAAGCGGAATTCCCGGTGTAGCGGTGAAATGCGTAGATATCGGGAGGAACACCAGTGGCGAAGGCGGCTTTCTGGGCCTATCCTGACGCTGAGGAGCGAAAGCGTGGGG |
| P-asv4383 | GCGAGCGTTAATCGGAATTACTGGGCGTAAAGCGTGCGCAGGCGGTTCTCCAAGTCCGTTGTGAGAGCCCCGGGCTCAACCTGGGAATGGCAGTGGAAACTAGGGGACTTGAATCTGGCAGAGGGGGGTGGAATGCCGCGTGTAGCAGTGAAATGCGTAGAGATGCGGCGGAACACCGATGGCGAAGGCAGCCCCCTGGGCTGAGATTGACGCTCAGGCACGAAAGCGTGGGG |
| P-asv4384 | GCAAGCGTTAATCGGAATTACTGGGCGTAAAGCGTGCGCAGGCGGTTCGGAAAGAAAGATGTGAAATCCCAGGGCTTAACCTTGGAACTGCATTTTTAACTACCGGGCTAGAGTGTGTCAGAGGGGGGTGGAATTCCACGTGTAGCAGTGAAATGCGTAGAGATGTGGAGGAACACCGATGGCGAAGGCAGCCCCCTGGGATGACACTGACGCTCATGCACGAAAGCGTGGGG |
| P-asv4385 | GCAAGCGTTGTTCGGAATTACTGGGCGTAAAGGGCGCGTAGGCGGCGCCGCAAGTCACTTGTGAAATCTCCGGGCTTAACTCGGAACGGCCAAGTGAAACTGCGGTGCTAGAGTGCGGGAGGGGCAATCGGAATTCTTGGTGTAGCGGTGAAATGCGTAGATATCAAGAGGAACACCTGAGGTGAAGACGGGTTGCTGGGCCGACACTGACGCTGAGGCGCGAAAGCCAGGGG |
| P-asv4386 | GCAGGCGTTAATCGGAATTACTGGGCGTAAAGCGTGCGCAGGCGGTCTACTAAGACAGGTGTGAAATCGCTCGGCTCAACTGGGCGGGGTCAGCGGATACGGCGTGGCTGGAGCAAGCTAGGGGGCAATGGAATTCCCGGTGTAGTGGTGGAATGCGTAGATATCGGGAGGAACACCAGTGGCGAAGGCGGTTGCCTGGAGCTTTGCTGACGCTGATGCGCGAAAGCGTGGGG |
| P-asv4387 | GCCAGCGTTGCTCGGAATTACTGGGCGTAAAGGGTCCGTAGGCGGTGTGGCAAGTCGGGAGTGAAATCTCTGGGCTTAACCCAGAGGCTGCTTCCGAAACTGCCGTGCTAGAGTGTGAGAGAGGCGAGTGGAATTGCGGGTGTAGCGGTGAAATGCGTAGATATCCGCAGGAACATCCGAGGCGAAAGCGGCTCGCTGGATCACAACTGACGCTGAGGGACGAAAGCTAGGGG |
| P-asv4388 | GCAAGCGTTGTCCGGATTTATTGGGCGTAAAGAGCTCGTAGGCGGTTTGATAAGTCGGGTGTGAAATCTCCAGGCTCAACCTGGAGCTGCCACTCGATACTGTCATGACTAGAGTTCGGTAGGGGACCACGGAATTCCTGGTGTAGCGGTGAAATGCGCAGATATCAGGAGGAACACCAGCAGCGAAGGCGGTGGTCTGGGCCGACACTGACGCTGAGGAGCGAAAGCGTGGGG |
| P-asv4389 | GCGAGCGTTGTCCGGAATTATTGGGCGTAAAGAGCGTGTAGGCGGTTCGGTAAGTCTGCTGTGAAATCCTGGGGCTCAACCCTGGGCGTGCAGCGGATACTGCCGGGCTAGAGGGTGGTAGAGGCGAGTGGAATTCCGAGTGTAGCGGTGAAATGCGCAGATATTCGGAGGAACACCAGTAGCGAAGGCGGCTCGCTGGGCCACACCTGACGCTGAGACGCGAAAGCGTGGGG |
| P-asv4390 | GCAAGCGTTATCCGGAATTATTGGGCGTAAAGCGTCCGTAGGCGGCTAGTCAAGTCTGCTGTCAAAGGTCAGGGCTTAACCCTGGATAGGCAGTGGAAACTGATTAGCTAGAGAGCGGCAGGGGTAGAGGGAATTCCCAGTGTAGCGGTGAAATGCGTAGATATTGGGAAGAACACCAGTGGCGAAGGCGCTCTACTGGGCCGGTACTGACGCTGAGGCGCGAAAGCGTGGGG |
| P-asv4391 | GCGAGCGTTAATCGGAATTACTGGGCGTAAAGGGCGCGTAGGCGGATGTATGTGAGTGATGTGAAAGCCCCGGGCTTAACCTGGGAAGCGCATTGCTGACTGTACGACTAGAGTATATGAGAGGGTGGCGGAATTTCCGGTGTAGCGGTGAAATGCGTAGAGATCGGAAGGAACGTCAATGGCGAAGGCAGCCACCTGGCATGATACTGACGCTGAGGCGCGAAAGCGTGGGG |
| P-asv4392 | GCAAGCGTTGTCCGGATTTATTGGGCGTAAAGCGAGTGCAGGCGGCTCGATAAGTCTGATGTGAAAGCCTTCGGCTAGACCGGAGAATTGCATCAGAAACTGTCGAGCTTGAGTACAGAAGAGGAGAGTGGAACTCCATGTGTAGCGGTGAAATGCGTAGATATATGGAAGAACACCGGTGGCGAAGGCGGCTCTCTGGTCTGTTACTGACGCTGAGGCTCGAAAGCATGGGT |
| P-asv4393 | GCAAGCGTTGTCCGGAATTATTGGGCGTAAAGCGCGCGCAGGCGGTTCCTTAAGTCTGATGTGAAAGCCCACGGCTCAACCGTGGAGGGTCATTGGAAACTGGGGAACTTGAGTGCAGGAGAGAAGAGTGGAATTCCACGTGTAGCGGTGAAATGCGTAGAGATGTGGAGGAACACCAGTGGCGAAGGCGACTCTTTGGCCTGTAACTGACGCTGAGGCACGAAAGCGTGGGG |
| P-asv4394 | GCAAGCGTTGTCCGGAATTATTGGGTTTAAAGGGTGCGTAGGCGGGTTAATAAGTCAGAGGTGAAATCCTACAGCTTAACTGTAGAACTGCCTTTGATACTGTTAACCTTGAGTACGGTAGAGGTAGGCGGAATTGACGGTGTAGCGGTGAAATGCATAGATATCGTCAAGAACACCAATTGCGTAGGCAGCTTACTGGACCGTAACTGACGCTGAGGCACGAAAGCGTGGGT |
| P-asv4395 | GCAAGCGTTAATCGGAATTACTGGGCGTAAAGCGCGCGTAGGTGGTTCGTTAAGTTGGATGTGAAAGCCCCGGGCCCAACCTGGGAACTGCATCCAAAACTGGCGAGCTAGAGTATGGTAGAGGGTGGTGGAATTTCCTGTGTAGTGGTGAAATGCGTAGATATAGGAAGGAACACCAGTGGCGAAGGCGACCACCTGGACTGATACTGACACTGAGGTGCGAAAGCGTGGGG |
| P-asv4396 | GCCAGCGTTATTCGGAATTACTGGGCGTAAAGCGCACGTAGGCGGCTTTGTAAGTTAGAGGTGAAAGCCTGGAGCTCAACTCCAGAATTGCCTTTAAGACTGCATCGCTTGAATCCAGGAGAGGTGAGTGGAATTCCGAGTGTAGAGGTGAAATTCGTAGATATTCGGAAGAACACCAGTGGCGAAGGCGGCTCACTGGACTGGTATTGACGCTGAGGTGCGAAAGCGTGGGG |
| P-asv4397 | GCAAACGTTGTTCGGAATTACTGGGCGTAAAGCGCGTGTAGGCTGCTCTGCAAGTCGGGTGTGAAAGCCCTGGGCTCAACCTAGGAAGTGCACTCGAAACTGCAGAGCTTGAGTCCTGGAGAGGATCGCGGAATTCTCGGTGTAGAGGTGAAATTCGTAGATATCGAGAGGAACACCGGTGGCGAAGGCGGCGATCTGGACAGTGACTGACGCTGAGACGCGAAAGCGTGGGG |
| P-asv4398 | GCAAGCGTTAATCGGAATTACTGGGCGTAAAGCGCGCGTAGGCGGCTTTTTAAGTCGGATGTGAAAGCCCCGGGCTCAACCTGGGAACTGCATCCGATACTGGAAGGCTAGAGTGCGAGAGAGGGAGGTAGAATTCCATGTGTAGCGGTGAAATGCGTAGATATATGGAGGAATACCAGTGGCGAAGGCGGCCTCCTGGCTCGACACTGACGCTGAGGTGCGAAAGCGTGGGG |
| P-asv4399 | GCTAGCGTTGTTCGGAATTACTGGGCGTAAAGCGCACGTAGGCGGATACTTAAGTTAGGGGTGAAATCCCAGGGCTCAACCCTGGAACTGCCTTTAATACTGGGTATCTCGAGTCCGAGAGAGGTGAGTGGAATTCCGAGTGTAGAGGTGAAATTCGTAGATATTCGGAGGAACACCAGTGGCGAAGGCGGCTCACTGGCTCGGTACTGACGCTGAGGTGCGAAAGCGTGGGG |
| P-asv4400 | GCAAGCGTTATCCGGAATTATTGGGCGTAAAGCGCGCGTAGGCGGTTTCTTAAGTCTGATGTGAAAGCCCCCGGCTCAACCGGGGAGGGTCATTGGAAACTGGGGAACTTGAGTACAGAAGAGGAGAGTGGAATTCCATGTGTAGCGGTGAAATGCGCAGAGATATGGAGGAACACCAGTGGCGAAGGCGGCTCTCTGGTCTGAAACTGACGCTGAGGTGCGAAAGCGTGGGG |
| P-asv4401 | GCGAGCGTTATTCGGAATTACTGGGCGTAAAGCGCACGCAGGCGGTTGCCCAAGTCAGATGTGAAAGCCCCGGGCTTAACCTGGGAACTGCATTTGAAACTGGGCGACTAGAGTATGAAAGAGGAAAGCGGAATTTCCAGTGTAGCAGTGAAATGCGTAGATATTGGAAGGAACACCGATGGCGAAGGCAGCTTTCTGGGTCGATACTGACGCTCATGTGCGAAAGCGTGGGG |
| P-asv4402 | GCAAGCGTTAATCGGAATTACTGGGCGTAAAGCGTGCGCAGGCGGCTGATTAAGTCAAATGTGAAAGCCCCGGGCTTAACCTGGGAATGGCATTCGATACTGGTCAGCTAGAGTTTGGTAGAGGTAAGCGGAATTCCGGGTGTAGCGGTGAAATGCGTAGATATCCGGAGGAACATCAGTGGCGAAGGCGGCTTACTGGACCAAAACTGACGCTCAGGCACGAAAGCGTGGAT |
| P-asv4403 | GCGAGCGTTAATCGGAATTACTGGGCGTAAAGCGCGCGTAGGTGGCTTGATAAGCCGGTTGTGAAAGCCCCGGGCTCAACCTGGGAACGGCATCCGGAACTGTCAGGCTAGAGTGCAGGAGAGGAAGGTAGAATTCCCGGTGTAGCGGTGAAATGCGTAGAGATCGGGAGGAATACCAGTGGCGAAGGCGGCCTTCTGGACTGGCACTGACACTGAGGCGCGAAAGCGTGGGT |
| P-asv4404 | TCGAGTGATGGCCGATCTTATTGGGCCTAAAGCGTTCGTAGCTGGCTAGTCAAGTCCGTTGGGAAATCGACGTGCCTAACACGTCGGCGTCCAGCGGAAACTGACTTGCTTGGGACCGGAAGACTTGAGGGGTACGTCCGGGGTAGGAGTGAAATCCCGTAATCCTGGACGGACCGCCAATGGCGAAAGCACCTCAAGAGCACGGATCCGACAGTGAGGAACGAAAGCTGGGGT |
| P-asv4405 | GCAAGCGTTACCCGGAATCACTGGGCGTAAAGGGCGTGTAGGCGGGAAGCTAAGTCCGACTTTAAAGACCGGGGCTCAACCCCGGGCCTGGGTTGGAGACTGGCTTTCTGGACCTCTGGAGAGGCAACTGGAATTCCTGGTGTAGCGGTGGAATGCGTAGATACCAGGAGGAACACCGATGGCGAAGGCAGGTTGCTGGACAGAAGGTGACGCTGAGGCGCGAAAGTGTGGGG |
| P-asv4406 | GCTAGCGTTGTTCGGAATTACTGGGCGTAAAGCGCACGTAGGCGGGTATTTAAGTCAGGGGTGAAATCCCAGAGCTCAACTCTGGAACTGCCTTTGATACTGGGTACCTAGAGTATGGAAGAGGTGAGTGGAATTCCGAGTGTAGAGGTGAAATTCGTAGATATTCGGAGGAACACCAGTGGCGAAGGCGGCTCACTGGTCCATTACTGACGCTGAGGTGCGAAAGCGTGGGG |
| P-asv4407 | GCAAGCGTTATCCGGATTTACTGGGTGTAAAGGGAGCGCAGGCGGTTTGGCAAGTTGAGAGTGGAAGCAGGGGGCTCAACCCCTTGACTGCTCCCAAAACTGTTAAACTTGAGTATGGGAGAGGCAGGCGGAATTCCTAGTGTAGCGGTGAAATGCTTAGATATTAGGAAGAACACCGGTGGCGAAGGCGGCCTGCTGGACCAAAACTGACGCTGAGGCTCGAAAGCGTGGGT |
| P-asv4408 | GCAAGCGTTAATCGGAATCACTGGGCGTAAAGGGCACGCAGGCGGTTGGATAAGTCGGGTGTGAAAGCCCCGGGCTCAACCTGGGAATGGCATTCGATACTGTCCAGCTAGAGTCTGGTAGAGGGAAGTGGAATTTCCGGTGTAGCGGTGAAATGCGTAGATATCGGAAAGAACACCAGTGGCGAAGGCGACTTCCTGGACCAAGACTGACGCTGAGGTGCGAAAGCGTGGGT |
| P-asv4409 | TCGAGTGATGGCCGATATTATTGGGCCTAAAGCGTTCGTAGCTGGCCGAGCAAGTCTGTCGGGAAATCTGTTCGCTCAACGAGCAGGCGTCCGGCAGAAACTGCTCGGCTTGGGACCGGAAGACTCGAGGGGTACGTCCGGGGTAGGAGTGAAATCCCGTAATCCTGGACGGACCACCGATGGCGAAAGCACCTCGAGAGCACGGATCCGACAGTGAGGAACGAAAGCTGGGGT |
| P-asv4410 | GCAAGCGTTGTCCGGAATCATTGGGTGTAAAGGGTGCGCAGGCTGGCGCCTAAGTCGGAGGTGAAATCTTGCCGCTTAACGGTAAAATGGCCTCCGATACTGGGTGTCTTGAATCTGACAGAGGTCGGTGGAATTCGTGGTGTAGCGGTGAAATGCATAGATATCACGAAGAACACCCGTGGCGAAGGCGGCCGGCTGGGTCAGTATTGACGCTGAGGCACGAGAGCGTGGGG |
| P-asv4411 | GCAAGCGTTATCCGGAATTATTGGGCGTAAAGCGTCCGCAGGGGGCGCATCAAGTCTGCTGTTAAAGGCCCGAGCTTAACTCCGGTAGAGCAGTGGAAACTGGTGCGCTAGAGGGCGACAGGGGTAGAGGGAATTCCCAGTGTAGCGGTGAAATGCGTAGAGATTGGGAAGAACACCGGTGGCGAAAGCGCTCTACTGGGTCGCACCTGACCCTCAGGGACGAAAGCTAAGGT |

| | |
|---|---|
| P-asv4412 | GCGAGCGTTGTTCGGAATTACTGGGCGTAAAGCGTGCGCAGGCGGATAATCAAGTCAGAAGTGAAAGCCCAGGGCTCAACCTTGGAATTGCTTTTGAAACTGGTTGTCTGGAGTCTCGGAGAGGTTGGCGGAAT TCCTAGTGTAGAGGTGAAATTCGTAGATATTAGGAGGAACACCAGTGGCGTAAGCGGCCAACTGGACGAGTACTGACGCTCATGCACGAAAGCGTGGGG |
| P-asv4413 | CCAAGCGTTACTCGGAATTACTGGGCGTAAAGAATGCGTAGGCTGTTTTGCAAGTTGGTGATGAAATGCTGGGGCTCAACTCCAGTGCTGTTACCAAAACTGCTTAACTTGAGTGCGGTAGGGGGCAATGGAATT CTTGGTGTAGGGGTTAAATCCTTTGATATCAAGAGGAACGCCGGAGGCGAAAGCGATTGCCTGGGCCGTTACTGACGCTAATGCATGACAGTGTGGGG |
| P-asv4414 | ACAAGTGGAAATCATGTTTACTGGGTCTAAAGTGACCGTAGCCGGCTGAGTAAGTCTTTTGTGAAAGTTCCGTAGCTTAACTCGGAATTTGCAAAAGATACTGCTTAGCTTGAGGCCGGGAGGGGTATGAGGTATT GTTTGGGTAGCGGTAAAATGTTATAATCCTTACAAGACCAACTAAGGCGAAGGCATCATACTTGAACGGACCTGACGGTGAGGGTCGAGAGCTGGGGG |
| P-asv4415 | GCGAGCGTTAATCGGAATTACTGGGCGTAAAGCGTGCGCAGGCGGTTTCTTAAGCAAGATGTGAAATCCCCGGGCTTAACCTGGGAACTGCATTTTGAACTGGGAGACTAGAGTGTGTCAGAGGGGGGTGGAAT TCCACGTGTAGCAGTGAAATGCGTAGAGATGTGGAGGAACACCGATGGCGAAGGCAGCCCCCTGGGATAACACTGACGCTCATGCACGAAAGCGTGGGG |
| P-asv4416 | GCGAGCGTTATTCGGATTCACTGGGCGTAAAGGGTGCGCAGGCGGTCGTGTGTGTCAGGTGTGAAATCTCGAGGCTCAACCTCGAAATGGCGCCTGAAACTACACGACTAGAGTACTGGAGAGGGTAACGGAAT TCACGGTGTAGCAGTGAAATGCGTAGATATCGTGAGGAACACCAGAGGCGAAGGCGGTTACCTGGACAGTTACTGACGCTCAGGCACGAAAGCGTGGGG |
| P-asv4417 | GCAAACGTTGTTCGGAATCACTGGGCATAAAGCGCACGTAGGCGGTTTCGTAAGTGTGGAGTGAAATCCCACGGCTCACCCGTGGAACTGCTCCGCATACTGCGGGACTTGAGGACGGTAGGGGAGAGCAGAA CTCCTGGTGGAGCGGTGAAATGCGTAGATATCAGGAGGAATACCGGTGGCGAAGGCGGCTCTCTGGTCCGTTTCTGACGCTGAGGTGCGAAAGCCAGGGT |
| P-asv4418 | GCGAGCGTTGTCCGGAATTACTGGGCGTAAAGAGCACGTAGGCGGCTATATGAGTTGGGTGTGAAATCCCGGGGCTTAACCCCGGAACTGCATTCAATACTGTATAGCTAGAGACCGAGAGAGGAAAATGGAAT TACGAGTGTAGAGGTGAAATTCGTAGATATTCGTAGGAACACCAGAGGCGAAAGCGATTTTCTGGCTCGGTACTGACGCTCAGGTGCGAAAGCGTGGGG |
| P-asv4419 | GCAAACGTTGTTCGGAATCACTGGGCATAAAGCGCACGTAGGCGGTCGATGAAGTGCGGGGTGAAAGCCCACGGCTCACCCGTGGAATTGCTTCGCATACTGATCGACTGGAGGTCGGGAGAGGAGAGCGGAA CTCTTGGAGGAGCGGTGAAATGCGTAGATATCAAGAGGAACACCAGAGGCGAAGGCGGCTCTCTGGCCCGATACTGACGCTGAGGTGCGAAAGCCAGGGT |
| P-asv4420 | GCAAGCGTTGTTCGGATTCATTGGGTGTAAAGGGTCCGTAGGTGGCCCGTTAAGTCCGGCGTGAAATCCCAGAGCTCAACTCTGGAACTGCGTCGGAAACTATCGGGCTTGAGTGCCGGAGAGGTGAAGGGAAT TCCCGGTGTAAGGGTGAAATCTGTAGATATCGGGAGGAACACCAGTGGCGAAGGCGCTTCACTGGCCGGTTGCTGACGCTGAGGGACGAAAGCAAGGGG |
| P-asv4421 | GCGAACGTTATTCGGAATCACTGGGCTTAAAGCGCGCGTAGGCGGGCCGTCGCGTCGGTCGCTGAAAGCCCCCGGCTCAACCGGGGAACGGGCACCGATACGGGCGGCCTGGAGTGGAGGAGGGGGATCCG GAACTCCCGGTGGAGCGGTGAAATGCGTTGAGATCGGGAGGAACGCCGGCGGCGAAAGCGGGATCCTGGACTTCGACTGACGCTGAGGCGCGAAAGCCAGGGG |
| P-asv4422 | GCAAGCGTTACTCGGAATTACTGGGCGTAAAGCGTGCGCAGGCGGTTCGTTAAGTCTGCTGTGAAAGCCCTGGGCTCAACCTGGGAATGGCAGTGGATACTGGCGAGCTAGAGTACGGTAGAGGAAGGTGGAA TTCCCGGTGTAGCGGTGAAATGCGTAGAGATCGGGAGGAACACCAGTGGCGAAGGCGACCTTCTGGACCAGTACTGACGCTCATGCACGAAAGCGTGGGG |
| P-asv4423 | CCAAGCGTTGTTCGGAATAACTGGGCGTAAAGCGTGTGTAGGCTGCGCGGAAAGTCAAATGTGAAAGCCAAGGGCTCAACCCTTGAACTGCATTCGATACTCCCGTGCTAGAGTAATGGAGAGGTAAGTGGAATT CCTGGTGTAGCAGTGAAATGCGTGGATATCAGGAGGAACATCAATGGCGAAGGCAACTTACTGGACATTTACTGACGCTCAGACACGAAGGCTAGGGG |
| P-asv4424 | CCGAGCGTTAATCGGAATCACTGGGCTTAAAGCGTACGCAGGCGGACTTGTAGGTATCTTGTGAAATCCCACGGCTTAACCGTGGAACTGCAGGGTAAACCACAAGTCTTGAGACATGTAGGGGTCAGTGGAACC ATGAGTGGAGCGGTGAAATGCGTAGATATTCATGGGAACGCCAATGGCGAAGGCAGCTGACTGGGCATGTTCTGACGCTCAGGTACGAAAGCGTGGGT |
| P-asv4425 | ACGAGCGTTGTCCGGAATTATTGGGCGTAAAGAGCTCGTAGGTGGTTGGTCACGTCTGCTGTGGAAACGCAACGCTTAACGTTGCGCGTGCAGTGGGTACGGGCTGACTAGAGTGCAGTAGGGGAGTCTGGAA TTCCTGGTGTAGCGGTGAAATGCGCAGATATCAGGAGGAACACCGGTGGCGAAGGCGGGACTCTGGGCTGTAACTGACACTGAGGAGCGAAAGCATGGGG |
| P-asv4426 | GCGAGCGTTGTCCGGAATCACTAGGCGTAAAGGGCAAGTAGGCGGAGCTATAAGTCAGATGTTAAAGGCATCGGCTCAACCGATGTACGGCATTCGATACTGTAGTTCTAGAGTGGGGTAGAGGAAGACGGAAT TCCTGGTGTAGCGGTGGAATGCGCAGATATCAGGAAGAACACCGATGGCGAAGGCAGTCTTCTGGGCCTCAACTGACGCTAAATTGCGAAGGTGTGGGT |
| P-asv4427 | CCGAGCGTTGTCCGGAATCATTGGGCGTAAAGGGTACGTAGGCGGATAAGCAAGTTAGAAGTTAAATCCTATAGCTCAACTATAGCAAGCTTTTAAAACTGCTCATCTTGAGGTATGGAAGGGAAAGTGGAATTCC TAGTGTAGCGGTGAAATGCGCAGATATTAGGAGGAATACCGGTGGCGAAGGCGACTTTCTGGCCATAAACTGACGCTGAGGTACGAAAGCGTGGGT |
| P-asv4428 | GCAAGCGTTGTCCGGAATCATTGGGCGTAAAGAGTTCGTAGGTGGTATGTTAAGTCTGGTGTTAAAGCCCGAAGCTCAACTTCGGTTCGGCACTGGATACTGGCAAACTTGAATGTGGTAGAGGTAAAGGGAATT CCTGGTGTAGCGGTGAAATGCGTAGATATCAGGAGGAACATCGGTGGCGAAAGCGCTTTACTGGACCATTATTGACACTGAGGAACGAAAGCCGGGGT |
| P-asv4429 | GCAAGCGTTGTTCGGAATCACTGGGCGTAAAGCGCGTGTAGGCGGCCCGATTAGTCGAGTGTGAAAGCCCGAGGCTCAACCTCGGAAGCGCACTCGAAACTGTCAGGCTTGAGTACGGGAGAGGGAAGTGGAA TTCCTGGTGTAGAGGTGAAATTCGTAGATATCAGGAGGAACACCGGTGGCGAAGGCGGCTTCCTGGACCGATACTGACGCTGAGACGCGAAAGCGTGGGG |
| P-asv4430 | GCAAGCGTTATCCGGAATGATTGGGCGTAAAGAGTCCGTAGGTAGTCATCCAAGTCTGCTGTTAAAGAGCGAGGCTTAACCTCGTAAAGGCAGTGGAAACTGGAAGACTAGAGTGTAGTAGGGGCAGAGGGAAT TCCTGGTGTAGCGGTGAAATGCGTAGAGATCAGGAAGAACACCGGTGGCGAAAGCGCTCTGCTGGGCTATAACTGACACTGAGGGACGAAAGCTAGGGG |
| P-asv4431 | GCAAGCGTTGTTCGGAATTATTGGGCGTAAAGGGCGCGTAGGCGGTTTGACAAGTCTGATGTGAAATCTCTGGGCTCAACCCAGAGTCTGCATTAGAAACTGTCGGGCTTGAGTATGGGAGAGGTGAGTGGAAT TTCCGGTGTAGCGGTGAAATGCGTAGATATCGGAAGGAACACCTGTGGCGAAAGCGGCTCACTGGACCATAACTGACGCTGAGGCGCGAAAGCTAGGGG |
| P-asv4432 | GCGAGCGTTGTCCGGAATTATTGGGCGTAAAGGGCTCGTAGGCGGTTTGTCGCGTCGGGAGTGAAAACACTGGGCTTAACTCAGTGCTTGCTTCCGATACGGGCAGACTAGAGGTATTCAGGGGAGAACGGAAT TCCTGGTGTAGCGGTGAAATGCGCAGATATCAGGAGGAACACCGGTGGCGAAGGCGGTTCTCTGGGAATTACCTGACGCTGAGGAGCGAAAGTGTGGGG |
| P-asv4433 | GCTAGCGTTGTCCGGAGTTACTGGGCGTAAAGCGCGCGTAGGCGGCTTTTTAAGTCGGGTGTGAAAGCCCGGGGCTCAACCCCGGATGCGCATCCGAGACTGGAAGGCTTGAGTCCGGGAGAGGGCGGCGGA ATTCCCAGTGTAGAGGTGAAATTCGTAGATATTGGGAAGAACACCGGTGGCGAAGGCGGCCGCCTGGCCCGGAACTGACGCTGAGGCGCGAAAGCGTGGGG |
| P-asv4434 | GCAAGCGTTGTCCGGAATTATTGGGCGTAAAGCGCGCGCAGGCGGTCCTTTAAGTCTGATGTGAAATCCCGCGGCTCAACCGCGGGCGGCCATTGGAAACTGGGGGACTTGAGTGCAGAAGAGGAAAGTGGAA TTCCACGTGTAGCGGTGAAATGCGTAGAGATGTGGAGGAACACCAGTGGCGAAGGCGACTTTCTGGTCTGTAACTGACGCTGAGGCGCGAAAGCGTGGGG |
| P-asv4435 | GCGAGCGTTAATCGGAATTACTGGGCGTAAAGCGTACGCAGGCGGTTTGTTAAGCGAGATGTGAAAGCCCCGGGCTCAACCTGGGAACTGCATTTCGAACTGGCAAACTAGAGTGTGATAGAGGGTGGTAGAAT TTCAGGTGTAGCGGTGAAATACGTAGAGATCTGAAGGAATACCGATGGCGAAGGCAGCCACCTGGGTCAACACTGACGCTCATGTACGAAAGCGTGGGG |
| P-asv4436 | GCAAGCGTTACCCGGAATCACTGGGCGTAAAGGGCGTGTAGGCGGGAAGCTAAGTCCGACTTTAAAGACCGGGGCTCAACCCCGGGAGTGGGTTGGAGACTGGCTTTCTAGACCTCTGGAGAGGCAACTGGAA TTCCTGGTGTAGCGGTGGAATGCGTAGATACCAGGAGGAACACCAATGGCGAAGGCAGGTTGCTGGACAGAAGGTGACGCTGAGGCGCGAAAGTGTGGGG |
| P-asv4437 | GCAAGCGTTGTCCGGAATTACTGGGCGTAAAGAGCTCGTAGGTGGTTTGTCGCGTCGTCTGTGAAATCCCGGGGCTTAACTTCGGGCGTGCAGGCGATACGGGCATAACTTGAGTGCTGTAGGGGAGACTGGA ATTCCTGGTGTAGCGGTGAAATGCGCAGATATCAGGAGGAACACCGATGGCGAAGGCAGGTCTCTGGGCAGTTACTGACGCTGAGGAGCGAAAGCATGGGT |
| P-asv4438 | GCGAGCGTTAATCGGAATTACTGGGCGTAAAGCGCGCGTAGGTGGCTTGATAAGCCGGTTGTGAAAGCCCCGGGCTCAACCTGGGAACGGCATCCGGAACTGTCAGGCTAGAGTGCAGGAGAGGAAGGTAGAA TTCCCGGTGTAGCGGTGAAATGCGTAGAGATCGGGAGGAATACCAGTGGCGAAGGCGGCCTTCTGGACTGTCACTGACACTGAGGTGCGAAAGCGTGGGT |
| P-asv4439 | GCAAGCGTTGTTCGGAGTTACTGGGCGTAAAGAGTCTGTAGGTGGTCTGTCAAGTCTTTGGTGAACGGCCGTGGCTTAACCATGGGAATGCCAAAGAGACTGGCAGACTGGAGGCTGGGAGAGGGAAGCGGAA TTTCTGGTGTAGCGGTGAAGTGCGTAGATATCAGAAGGAAGGCCGGTGGCGAAGGCGGCTTCCTGGAACAGACCTGACACTGAGAGACGAAAGCGTGGGG |
| P-asv4440 | GCGAGCGTTAATCGGAATTACTGAGCGTAAAGCGCGCGTAGGCGGTCTGATAAGCCGGTTGTGAAAGCCCCGGGCTCAACCTGGGAACGGCATCCGGAACTGGCGAGCTAGAGTACGGTAGAGGGTGGTGGAA TTTCCTGTGTAGCGGTGAAATGCGTAGATATAGGAAGGAACACCAGTGGCGAAGGCGACCACCTGGACTGATACTGACACTGAGGTGCGAAAGCGTGGGG |
| P-asv4441 | GCAAGCGTTATCCGGAATCACTGGGCGTAAAGGGCGCGTAGGCGGTTTGTTAAGTCCGATGTTAAAGACCGAGGCTCAACCTCGACACAGCGTTGGATACTGGCAAGCTGGACGGTTGGAGAGGAAGGTAGAAT TACCAGTGTAGCGGTGGAATGCGTAGATACTGGTAGGAATACCCATTGCGAAGGCAGCCTTCTGGACAACACGTGACGCTGAGGCGCGAAAGTGTGGGG |
| P-asv4442 | GCAAGCGTTGTCCGGATTTATTGGGCGTAAAGAGTTCGTAGGCGGTTTGTTAAGTCTGACGTTAAAGATCGGGGCTCAACCTCGGGAGTGCATTGGATACTGGCAGACTGGAGTGTGGTAGAGGTTAGTGGAATT CCCAGTGTAGCGGTGAAATGCGTAGATATTGGGAAGAACACCAGTGGCGTAGGCGACTAACTGGGCCATAACTGACGCTGAGGAACGAAAGCCAGGGG |
| P-asv4443 | GCTAGCGTTGCTCGGAATTACTGGGCGTAAAGCGCACGTAGGCGGCTTCTTAAGTCGGAGGTGAAATCCTGGAGCTCAACTCCAGAACTGCCTTCGATACTGGGAAGCTTGAGTTCGGGAGAGGTGAGTGGAAC TGCGAGTGTAGAGGTGAAATTCGTAGATATTCGCAAGAACACCAGTGGCGAAGGCGGCTCACTGGCCCGATACTGACGCTGAGGTGCGAAAGCGTGGGG |
| P-asv4444 | GCGAGCGTTGTTCGGAATTACTGGGCGTAAAGGGCGTGTAGGCGGTGCGGTAAGTCGGGCGTGAAATCCCCGGGCTTAACCCGGGGCGGTCGTTCGAAACTGCCGTGCTAGAGTGCAGGAGAGGGAAGTGGA ATTCCCGGTGTAGCGGTGAAATGCGTAGATATCGGGAGGAACACCTGCGGCGAAGGCGGCTTCCTGGACTGCAACTGACGCTGAGGCGCGAAAGCTAGGGG |
| P-asv4445 | GCGAGCGTTAATCGGAATTACTGGGCGTAAAGCGTGCGTAGACGGTTACGTAAGTCGGGTGTGAAAGCCCCGGGCTCAACCTGGGAATTGCATTCGAGACTGCATAGCTAGGGTGCGGAAGAGGGAAGCGGAA TTTCAGGTGTAGCGGTGAAATGCGTAGATATCTGAAGGAACACCAGTGGCGAAGGCGGCTTCCTGGCCCAGCACCGACGTTCAGGCACGAAAGCGTGGGG |
| P-asv4446 | GCAAGCGTTGTCCGGAATTATTGGGCGTAAAGAGCTCGTAGGCGGTCTGTCGCGTCGGCTGTGAAAACCTGGGGCTCAACCCCGGGCCTGCAGCCGATACGGGCAGACTAGAATTCGGTAGGGGAGACTGGAA TTCCTGGTGTAGCGGTGAAATGCGCAGATATCAGGAGGAACACCGGTGGCGAAGGCGGGTCTCTGGGCCGATATTGACGCTGAGGAGCGAAAGCGTGGGG |
| P-asv4447 | GCAAGCGTTGTTCGGAATCACTGGGCGTAAAGGGCGTGTAGGCGGTCAGGTAAGTGGGACGTGAAATGCCTCGGCTCACCCGAGGAGCGGCGTCCCATACTGCTTGGCTTGAGTACGGGAGAGGATGTGGGA ATTCCTGGTGTAGCGGTGAAATGCGTAGATATCAGGAGGAACACCGGTGGCGAAGGCGCACATCTGGCCCGATACTGACGCTGAGGCGCGAAAGCTAGGGG |
| P-asv4448 | GCGAGCGTTGTTCGGAATTACTGGGCGTAAAGGGCGCGTAGGCGGCCATTCAAGTCAGGCGTGAAAGCCCCGGGCTCAACCCGGGAATGGCGCTTGGGACTGGGTGGCTGGAGTTCGGGAGAGGAGCGTGG AATTCCCAGTGTAGAGGTGAAATTCGTAGATATTGGGAAGAACACCGGTGGCGAAGGCGGCGCTCTGGACCGATACTGACGCTGAGGCGCGAAAGCGTGGGG |
| P-asv4449 | CCGAGTGGTCGGGACGATTATTGGGCCTAAAGCATCCGTAGCCGGTCCTGCAAGTCCTCCGTTAAATCCACCCGCTCAACGGTTGGGCCGCGGGGGATACTACAGGGCTAGGAGGCGGGAGAGGCAAGCGGTA CTCGATGGGTAGGGGTAAAATCCACTGATCCATTGAAGACCACCAGTGGCGAAGGCGGCTTGCCAGAACGCGCTCGACGGTGAGGGATGAAAGCTGGGGG |
| P-asv4450 | GCAAGCGTTGCTCGGAATCACTGGGCGTAAAGGGCGCGTAGGCGGGATGGCAAGTCGGTCGTGAAATCCTTCGGCTCAACCGAAGAACGTCGGGCGATACTGCCGTTCTTGAGGGATGCAGAGGAGATTGGAA TTCCCGGTGTAGCGGTGAAATGTGTAGAGATCGGGAGGAACACCGGTGGCGAAGGCGGGTCTCTGGGCATCTCCTGACGCTGAGGCGCGAAAGCGTGGGG |

| | |
|---|---|
| P-asv4451 | GCGAGCGTTAATCGGAATTACTGGGCGTAAAGAGTGCGTAGGCGGTTCTTTGTGTCCGCTGTGAAAGCCCCGGGCTTAACCTGGGAGTGGCAGTGGAAACTAGAGGACTTGAATCCGGCAGAGGGGGGTGGAA<br>TGCCACGTGTAGCAGTGAAATGCGTAGATATGTGGCGGAACACCGATGGCGAAGGCAGCCCCCTGGGCTGAGATTGACGCTCAGGCACGAAAGCGTGGGG |
| P-asv4452 | GCAAGCGCTGTTCGGAATTATTGGGCGTAAAGCGCGTGTAGGCGGCTTTGTAAGTTGGGTGTGAAAGCCCTCGGCTTAACCGAGGAAGTGCGCCCAAGACTGCATGGCTTGAGTACCGGAGAGGGTGGCGGAA<br>TTCCCGGTGTAGAGGTGAAATTCGTAGATATCGGGAGGAACACCAGCGGCGAAGGCGGCCACCTGGACGGATACTGACGCTGAGACGCGAAAGCGTGGGG |
| P-asv4453 | GCAAGCGTTGTCCGGATTTATTGGGCGTAAAGAGCGTGTAGGCGGCCAGGTAGGTCCGTTGTGAAAACTGGAGGCTCAACCTCCAGAGGTCGATGGAAACCATCTGGCTAGAGTCCGGAAGAGGAGAGCGGAA<br>TTCCCGGTGTAGCGGTGAAATGCGCAGATATCGGGAAGAACACCCGTGGCGAAGGCGGCTCTCTGGGACGGTACTGACGCTGAGACGCGAAAGCGTGGGG |
| P-asv4454 | CCAAGCGTTGTTCGGATTCACTGGGCGTAAAGGGTGCGTAGGTGGCGGGGTAAGTCGGATGTGAAAGCTCGGAGCTCAACTCCGAAATGGCATTGGAAACTACCTTGCTCGAGGGTTGGAGGGGGGACTGGAA<br>TTCTCGGTGTAGCAGTGAAATGCGTAGATATCGAGAGGAACACCAGTGGCGAAGGCGAGTCCCTGGACAACTCCTGACACTGAGGCGCGAAAGCTAGGGT |
| P-asv4455 | GCAAGCGTTACTCGGAATTACTGGGCGTAAAGCGTGCGTAGGTGGTTTCTTAAGTCTGCTGTGAAAGCCCCGGGCTCAACCTGGGAATTGCAGTGGATACTGGGGAGCTAGAGTGCGGTAGAGGGTAGTGGAAT<br>TCCCGGTGTAGCGGTGAAATGCGTAGAGATCGGGAGGAACACCAGTGGCGAAGGCGGCTACCTGGACCAGCACTGACACTGAGGCACGAAAGCGTGGGG |
| P-asv4456 | CCAAGCGTTATCCGGATTTATTGGGTTTAAAGGGTGCGTAGGCGGCCTGTTAAGTCAGGGGTGAAAGACGGTAGCTTAACTATCGCAGTGCCCTTGATACTGATGGGCTTGAATACACTAGAGGTAGGCGGAATG<br>TGACAAGTAGCGGTGAAATGCATAGATATGTCACAGAACACCAATTGCGAAGGCAGCTTACTATGGTGTCATTGACGCTGAGGCACGAAAGCGTGGGG |
| P-asv4457 | GCAAGCGTTAATCGGAATTACTGGGCGTAAAGGGCGCGTAGGTGGTTATTTAAGTCAGATGTGAAATCCCTGGGCTCAACCTAGGAATTGCATTTGAAACTAGATGGCTAGAGTATGGTAGAGGGAAGTGGAATT<br>TCCGGTGTAGCGGTGAAATGCGCAGATATCGGGAGGAACACCGGTGGCGAAGGCGGTTCTCTGGGCCTTACCTGACACTGAGGAGCGAAAGCGTGGGG |
| P-asv4458 | CCTAGCGTTGTCCGGAATTATTGGGCGTAAAGAGCTCGTAGGCGGTTCGGCAAGTCGGATGTGAAATCTCCACGCTCAACGTGGAGGGGCCATTCGAAACTGCCGTGACTCGAGTCCGGTAGGGGAGTGTGGA<br>ATTCCCGGTGTAGCGGTGAAATGCGCAGATATCGGGAGGAACACCAGTAGCGAAGGCGGCACTCTGGGCCGGTACTGACGCTGAGGAGCGAAAGCGTGGGG |
| P-asv4459 | CCAAGCGTTGTTCGGATTCACTGGGCGTAAAGGGTGCGTAGGTGGTGGGGTAAGTCGGATGTGAAAGCTCGGAGCTCAACTCCGAAATGGCATTGGAAACTACCCTGCTAGAGGATTGGAGGGGGGACTGGAA<br>TACTTGGTGTAGCAGTGAAATGCGTAGATATCAAGTGGAACACCAGTGGCGAAGGCGAGTCCCTGGACAATACCTGACACTGAGGCACGAAAGCTAGGGG |
| P-asv4460 | GCAAGCGTTGTCCGGAATCATTGGGCGTAAAGAGCGCGTAGGCGGCCCAGTCAGTCCGCTGTGAAAGTCAGAGGCTCAACCTCTGGATGCCGGTGGATACTGCTGGGCTAGAGTCCGGAAGAGGCGAGTGGAA<br>TTCCTGGTGTAGCGGTGAAATGCGCAGATATCAGGAGGAACACCGATGGCGAAGGCAGCTCGCTGGGACGGTACTGACGCTGAGGCGCGAAAGCGTGGGG |
| P-asv4461 | GCGAGCGTTGTTCGGAATCACTGGGCTTAAAGCGCGTGTAGGCGGAGTGATGGGTGCCTTGTGAAATCCCTCGGCTCAACCGGGGAATGGCTCGGCAAACCGGCACTCCTGAGGCAGGTAGAGGTTACTGGAA<br>CCATTGGTGGAGCGGTGAAATGCGTAGAGATCAATGGGAACGCCGGTGGCGAGAGCGAGTAACTGGGCCTGATCTGACGCTGAGACGCGAAAGCGTGGGT |
| P-asv4462 | GCGAGCGTTGTCCGGAATCACTGGGCGTAAAGGGCGCGTAGGCGGTTTGATAAGGCTGTGGTGAAAGCCCGGGGCTCAACCCCGGGTCTGCCGTGGCGACTGTCAGACTGGAGGGCTCGAGAGGCAGACGGA<br>ATTCCGGGTGTAGCGGTGGAATGCGTAGAGATCCGGAAGAAGACCGGTGGCGAAGGCGGTCTGCTGGCGAGTACCTGACGCTGAGGCGCGACAGCGTGGGG |
| P-asv4463 | CCGAGTGGTCGGGACGATTATTGGGCCTAAAGCATCCGTAGCCGGTTCTGCAAGTCTTCCGTTAAATCCAACTGCTTAACAGTTGGGCTGCGGAGGATACTATAGAGCTAGGAGGCGGGAGAGGCAAGCGGTAC<br>TTAGTGGGTAGGGGTAAAATCCGTTGATCCACTGAAGACCACCAGTGGCGAAGGCGGCTTGCCAGAACGCGCTCGACGGTGAGGGATGAAAGCTGGGGG |
| P-asv4464 | GCAAGCGTTGTTCGGATTTACTGGGCGTAAAGGGCGCGTAGGCGGCCTCCGCAAGTCAATTGTGAAATCTCCGGGCTTAACTCGGAAAGGTCAACTGATACTGCGGGGCTAGAGTGCGGAAGGGGCAACTGGA<br>ATTCTCGGTGTAGCGGTGAAATGCGTAGATATCGAGAGGAACACCTGCGGCGAAGGCGGGTTGCTGGGCCGACACTGACGCTGAGGCGCGAAAGCTAGGGG |
| P-asv4465 | CCAAGCGTTGTTCGGATTCACTGGGCGTAAAGGGTGCGTAGGCGGTCGGGTAAGTTTGATGTGAAATCTCCGGGCCTAACCCGGAAATTGCATTGAATACTGCCTGGCTAGAGGATTGGAGGGGAGACTGGAAT<br>GCTCGGTGTAGCAGTGAAATGCGTAGATATCGAGCGGAACACCAGTGGCGAAGGCGAGTCTCTGGACAATTCCTGACGCTGAGGCACGAAAGCCAGGGG |
| P-asv4466 | GCGAGCGTTGTCCGGATTTATTGGGCGTAAAGAGCTCGTAGGCGGCTTGTCGCGTCGACTGTGAAATCCCGTGGCTCAACTGCGGGCTTGCAGCCGATACGGGCAGGCTGGAGTTCGGTAGGGGAGACTGGAA<br>TTCCTGGTGTAGCGGTGAAATGCGCAGATATCAGGAGGAACACCAGCGGCGAAGGCGGCGCTCTGGGCCGGAACTGACGCTGAGGAGCGAAAGCGTGGGT |
| P-asv4467 | GCAAGCGTTGTTCGGAATTACTGGGCGTAAAGCGCGGGCAGGCGGTCTGCTAAGTCGCTTGTGAAAATCCGGGGCTCAACCCCGGAAATGCATGCGATACTGGCAGGCTGGAGACCGGTAGAGGCTAGCGGAA<br>TTCCTGGCGTAGCGGTGGAATGCGTAGATATCAGGAAGAACACCTGTGGCGAAGGCGGTTAGCTGGGCCGGTTCTGACGCTCATCCGCGAAAGCGTGGGG |
| P-asv4468 | GCAAGCGTTGTTCGGATTTACTGGGCGTAGAGGGCGCGTAGGCGGCGCAACAAGTCACTTGTGAAATCTCCGGGCTTAACCCGGAGCGGCCAAGTGATACTGTCGTGCTAGAGTGCGGAAGGGGCTACTGGAA<br>TTCTCGGTGTAGTGGTGAAATGCGCAGATATCAGGAGGAACACCGGTGGCGAAGGCGGGTCTCTGGGCAACAACTGACGCTGAGGAGCGAAAGCGTGGGG |
| P-asv4469 | GCAAGCGTTGTCCGGAATTACTGGGCGTAAAGCGCACGTAGGCGGATCAGTTAGTCGGGGGTGAAATCCCGGGGCTCAACCCCGGAACTGCCTTCGATACTGCTGGTCTGGAGGTCGAGAGAGGCGAGTGGAA<br>TTCCGAGTGTAGAGGTGAAATTCGTAGATATTCGGAGGAACACCAGTGGCGAAGGCGGCTCGCTGGCTCGATACTGACGCTGAGGTGCGAAAGCGTGGGG |
| P-asv4470 | GCAAGCGTTGTTCGGAATTACTGGGCGTAAAGGGCTCGTAGGTGGCCAACTAAGTCAGACGTGAAATCCCTCGGCTTAACCGGGGAACTGCGTCTGATACTGGATGGCTTGAGTTTGGGAGAGGGACGCGGAAT<br>TCCAGGTGTAGCGGTGAAATGCGTAGAGATCGGAAAGAACATCGGTGGCGAAGGCGGCCGCCTGGGCCCAGACTGACACTGAGGTGCGAAAGCGTGGGG |
| P-asv4471 | GCTAGCGTTGTTCGGAATTACTGGGCGTAAAGCGCGTGTAGGCGGCCTGCTAAGTCTGATGTGAAAGCCCGGGGCTCACCCCCGGAAGTGCATTGGAAACTGGTAGGCTCGAGTCCCGGAGAGGAGGGTGGAA<br>TTCCTGGTGTAGCGGTGAAATGCGTAGATATCAGGAGGAACACCGGTGGCGAAGGCGGCCCTCTGGACGGTGACTGACGCTGAGACGCGAAAGCGTGGGG |
| P-asv4472 | GCAAGCGTTGTCCGGATTTATTGGGCGTAAAGAGCGTGTAGGCGGCCATACAGGTCAGCTGTGAAAACTCGAGGCTCAACCTCGAGACGTCGGTTGAAACCGTATGGCTAGAGTCCGGAAGAGGAGAGTGGAAT<br>TCCCGGTGTAGCGGTGAAATGCGCAGATATCGGGAAGAACACCTATGGCGAAGGCAGCTCTCTGGGACGGTACTGACGCTGAGACGCGAAAGCGTGGGG |
| P-asv4473 | GCTAGCGTTGTTCGGAATTACTGGGCGTAAAGCGTATGCAGGCGGATCGTTAAGTCGGGGGTGAAATCCCGAGGCTCAACCTCGGAACTGCCTTCGATACTGGCGACCTTCGAGTCCGGGGGAGGTGAGTGGA<br>ACTCCGAGTGTAGAGGTGAAATTCGTAGATATTCGGAAGAACACCAGTGGCGAAGGCGGCTCACTGGCCCGGTACTGACGCTGAGATACGAAAGCGTGGGG |
| P-asv4474 | GCTAGCGTTGTTCGGAATTACTGGGCGTAAGGCGAGTGTAGGTGGTTGTCCAAGTTGGATGTGAAAGCCTTGAGCTCAACTCAAGAAATGCATTCAGGACTGGGCGGCTAGAGGACCGGAGAGGATAGTGGAAT<br>TCCCAGTGTAGTGGTGAAATACGTAGAGATTGGGAAGAACACCAGTGGCGAAGGCGGCTATCTGGACGGTTTCTGACACTAAGACTCGAAAGCGTGGGG |
| P-asv4475 | GCGAGCGTTGTTCGGAATCACTGGGCATAAAGCGCACGTAGGCGGTGCCGTAAGTCGGATGTGAAATCCCTCGGCCTAACCGAGGAACTGCGTCCGATACTGCGGTGCTTGAGGATCAGAGGGGTGCATGGAA<br>TTTCCGGTGGAGCGGTGAAATGCGTTGATATCGGAAGGAACTCCGGCGGCGAAAGCGATGCACTGGCTGATTTCTGACGCTGAGGTGCGAAAGCCAGGGG |
| P-asv4476 | GCGAGCGTTGTCCGGAATTACTGGGCGTAAAGAGCTCGTAGGTGGTTTGTCGCGTTGTTCGTGAAAACCGAAGGCTTAACCTTTGGCGTGCGGGCGATACGGGCAGACTGGAGTACTGCAGGGGAGACTGGAA<br>TTCCTGGTGTAGCGGTGGAATGCGCAGATATCAGGAGGAACACCGGTGGCGAAGGCGGGTCTCTGGGCAGTAACTGACGCTGAGGAGCGAAAGCGTGGGG |
| P-asv4477 | GCAAGCGTTGTCCGGAATTATTGGGCGTAAAGAGCTCGTAGGCGGCCTGTTGCGTCCGCTGTGAAATCCCAGGGCTTAACCCTGGGCGTGCAGTGGATACGGGCTGGCTGGAGGCAGGCAGGGGAGAACGGA<br>ATTCCCGGTGTAGCGGTGAAATGCGCAGATATCGGGAGGAACACCGGTGGCGAAGGCGGTTCTCTGGGCCTGTTCTGACGCTGAGGAGCGAAAGCGTGGGG |
| P-asv4478 | GCTAGCGTTGTTCGGAATCACTGGGCGTAAAGCGCACGTAGGCGGATGTGTCAGTCAGGGGTGAAATCCCGGAGCTCAACTTCGGAACTGCCTTTGATACAGCACGTCTTGAGTCCGATAGAGGTGGGTGGAAT<br>TCCTAGTGTAGAGGTGGAATTCGTAGATATTAGGAAGAACACCGGTGGCGAAGGCGGCCCACTGGATCGGTACTGACGCTGAGGCGCGAAAGCCAGGGG |
| P-asv4479 | CCAAGCGTTATCCGGATTTACTGGGCGTAAAGCGCGTGTAGGTGGTTTGTTAAGTTGGGTGTTAAATCTCTCGGCTCAACTGAGAGGTGTCGCTCAAAACTAGCAGACTGGAGGACGGTAGAGGAAGGTGGAATT<br>CCGGGTGTAGTGGTGAAATGCGTAGATATCCGGAGGAACACCAGTGGCGAAGGCGGCCTTCTGGACCGCTCCTGACACTCAAACGCGAAAGCTAGGGG |
| P-asv4480 | CCGAGCGTTGTCCGGAGTTACTGGGCGTAAAGCGCGCGCAGGCGGCGGTGCTGGCCCGGCGTGAAAGCCCCCGGCTCAACCGGGGAGGGTCGTCGGGGACCGCACCGCTTGAGGGCGGTAGGGGCTGGTG<br>GAATGCCTGGTGTAGTGGTGAAATGTGTAGAGATCAGGCGGAACACCGGTGGCGAAGGCGGGATCCTGGGCCAGTACTGACGCTGAGGAGCGAAAGCGTGGGG |
| P-asv4481 | GCTAGCGTTGTTCGGAATTACTGGGCGTAAAGCGCACGTAGGCGGATCGATAAGTAAGAGGTGAAATCCCAGAGCTCAACTCTGGAACTGCCTTTTAAACTGTCGATCTGGAGCAGAGTAGAGGTGAGTGGAATT<br>CCGAGTGTAGAGGTGAAATTCGTAGATATTCGGAGGAACACCAGTGGCGAAGGCGGCTCACTGGACTCTTGCTGACGCTGAGGTGCGAAAGCGTGGGG |
| P-asv4482 | GCGAGCGTTGTCCGGAATTATTGGGCGTAAAGGGCTCGTAGGCGGCTTGTTGCGTCGGCTGTGAAAACCCGCGGCTCAACCGTGGGCTTGCAGCCGATACGGGCAGGCTAGAGTTCGGTAGGGGAGACTGGA<br>ATTCCTGGTGTAGCGGTGAAATGCGCAGATATCAGGAGGAACACCGGTGGCGAAGGCGGGTCTCTGGGCCGATACTGACGCTGAGGAGCGAAAGCGTGGGG |
| P-asv4483 | CCAAGCGTTGTTCGGATTCACTGGGCGTAAAGGGTGCGTAGGTGGCCGGGTAAGTCTGACGTGAAATCTCCGAGCCTAACTCGGAAACTGCATCGGAAACTACACGGCTCGAGGGTTGGAGGGGGGACTGGAA<br>TACTTGGTGTAGCAGTAAAATGCGTAGATATCAAGTGGAACACCAGTGGCGAAGGCGAGTCCCTGAACAACTCCTGACACTGAGGCACGAAAGCTAGGGG |
| P-asv4484 | GCGAACGTTATTCGGAATCACTGGGCTTACAGGGTGCGTAGGTGGCTTGTCAAGTAGGGTGTGAAAGCCCCCGGCTCAACCGGGGAATTGCGCCCTAAACTGGCGAGCTGGAGTGAGATAGGGGTGTGTGGAA<br>CTTCCGGTGGAGCGGTGAAATGTGTTGATATCGGAAGGAACGCCGGTGGGGAAAGCGACACACTGGATCTTGTCTGACGCTGAGGCACGAAAGCCAGGGG |
| P-asv4485 | GCGAGCGTTGTCCGGAATTACTGGGCGTAAAGGGTGCGTAGGTGGTTTGTTAAGTCCGTTGTGAAATCCCTGGGCTCAACCTGGGAATGGCAGTGGATACTGGCAAGCTAGAGTGCGGTAGAGGGTAGTGGAAT<br>TCCCGGTGTAGCGGTGAAATGCGTAGAGATCGGGAGGAACACCAGTGGCGAAGGCGGCTACCTGGACCAGCACTGACACTGAGGCACGAAAGCGTGGGG |
| P-asv4486 | GCAAGCGTTGTTCGGAATCACTGGGCGTAAAGCGCGTGTAGGCGGCCTCTTAAGTCTGATGTGAAAGCCCGGGGCTTACCCCCGGAAGTGCATTGGAAACTGGGAGACTAGAGTACCGGAGAGGAAGGTGGAA<br>TTCCTGGTGTAGTGGTGAAATGCGTAGATATCAGGAGGAACACCGGTGGCGAAGGCGGCCTTCTGGACGGATACTGACGCTGAGACGCGAAAGCGTGGGG |
| P-asv4487 | GCAAGCGTTGTTCGGAATTACTGGGCGTAAAGGGTTCGTAGGTGGCCAACTAAGTCAGACGTGAAATCCCTCAGCTTAACTGGGGAACTGCGTCTGAGACTGGATGGCTTGAGTGCAGGAGAGGAACGCGGAAT<br>TCCAGGTGTAGCGGTGAAACGCGTAGATATCTGGAGGAATACCGGTGGCGAAGGCGGCATCCTGGACCATTACTGACGCTGAGGCGCGAAAGCTAGGGG |
| P-asv4488 | GCAAGCGTTGTCCGGAATCATTGGGCGTAAAGCGCGCGTAGGTGGCTTCGTAAGTCCGCTGTGAAAGTCAAAGGCTCAACCTTTGAAAGCCGGTGGATACTGTGAAGCTAGAGTCCGGTAGAGGCGAGTGGAAT<br>TCCTGGTGTAGCGGTGGAATGCGCAGATATCAGGAGGAACACCAATAGCGAAGGCAGCTCGCTGGGACGGTACTGACACTAAGGCGCGAAAGCGTGGGG |
| P-asv4489 | GCTAGCGTTGTTCGGAATTACTGGGCGTAAAGCGCACGTAGGCGGCTTTGTAAGTCAGGGGTGAAATCCCGGGGCTCAACCCCGGAACTGCCCTTGAGACTGCAAGGCTCGAATCCAGGAGAGGTGAGTGGAA<br>TTCCGAGTGTAGAGGTGAAATTCGTAGATATTCGGAAGAACACCAGTGGCGAAGGCGGCTCACTGGACTGGTATTGACGCTGAGGTGCGAAAGCGTGGGG |

| | |
|---|---|
| P-asv4490 | GCAAGCGTTGTCCGGAATTATTGGGCGTAAAGCGCGCGCAGGCGGTCATTTAAGTCTGGTGTTTAATCCCGGGGCTCAACCCCGGATCGCACTGGAAACTGGGTGACTTGAGTGCAGAAGAGGAGAGTGGAATT<br>CCACGTGTAGCGGTGAAATGCGTAGATATGTGGAGGAACACCAGTGGCGAAGGCGACTCTCTGGGCTGTAACTGACGCTGAGGCGCGAAAGCGTGGGG |
| P-asv4491 | GCGAGCGTTAATCGGAATTACTGGGCGTAAAGCGTGCGCAGGCGGTTGTGTAAGACAGATGTGAAATCCCCGGGCTCAACCTGGGAACTGCATTTGTGACTGCACGGCTGGAGTGCGGCAGAGGGGGATGGAA<br>TTCCGCGTGTAGCAGTGAAATGCGTAGATATGCGGAGGAACACCGATGGCGAAGGCAATCCCCTGGGCCTGCACTGACGCTCATGCACGAAAGCGTGGGG |
| P-asv4492 | GCGAGCGTTAATCGGAATTACTGGGCGTAAAGCGTGCGCAGGCGGTTCGTTGTGTCGGTGGTGAAAGCCCCGGGCTCAACCTGGGAATGGCTATGGAAACTGGCGAACTGGAGTGTGGCAGAGGGGGGTGGA<br>ATTCCGCGTGTAGCAGTGAAATGCGTAGATATCAGGAGGAACACCGATGGCGAAGGCAGGTCTCTGGGCTGACACTGACGCTCATGCACGAAAGCGTGGGG |
| P-asv4493 | GCTAGCGTTGTTCGGAATTACTGGGCGTAAAGCGCACGTAGGCGGATTGTTAAGTCGGGGGTGAAATCCCAGGGCTCAACCCTGGAACTGCCTCCGATACTGGCATTCTTGAGTCCGAGAGAGGTGAGTGGAAT<br>TCCGAGTGTAGAGGTGAAATTCGTAGATATTCGGAAGAACACCAGTGGCGAAGGCGGCTCACTGGCTCGGTACTGACGCTGAGGTGCGAAAGCGTGGGG |
| P-asv4494 | GCTAGCGTTGTTCGGAATTACTGGGCGTAAAGCGCACGTAGGCGGCTACTCAAGTCAGAGGTGAAAGCCCGGGGCTCAACCCCGGAACTGCCTTTGAAACTAGGTAGCTAGAATCTTGGAGAGGCGAGTGGAAT<br>TCCGAGTGTAGAGGTGAAATTCGTAGATATTCGGAAGAACACCAGTGGCGAAGGCGACTCGCTGGACAAGTATTGACGCTGAGGTGCGAAAGCGTGGGG |
| P-asv4495 | GCAAGCGTTAATCGGAATTACTGGGCGTAAAGCGCGCGTAGGTGGTTCAGTAAGATGGGTGTGAAATCCCCGGGCTTAACCTGGGAACTGCTTTCATAACTGCTGAGCTAGAGTACGGTAGAGGGTAGTGGAAT<br>TTCCTGTGTAGCGGTGAAATGCGTAGATATAGGAAGGAACACCAGTGGCGAAGGCGACTACCTGGACTGATACTGACACTGAGGTGCGAAAGCGTGGGG |
| P-asv4496 | ACGAGCGTTATCCGGAATAATTGGGCGTAAAGGGTGCGTAGGCGGTGTGTTAAGTTCAAGGTTTAATTGTAGGGCTCAACCCTATATCGCTTTGAAAACTGGCACACTAGAGTAAGATAGAGGCAAGTGGAATTCC<br>ATGTGTAGCGGTAAAATGCGTAAATATATGGCAGAACACCAGAGGCGAAGGCGGCTTGCTGGGTCTTCACTGACGCTGAGGCACGAAAGCGTGGGG |
| P-asv4497 | GCAAGCGTTATCCGGAATGATTGGGCGTAAAGCGTCTGTAGGTGGCTTTTCAAGTCCGCCGTCAAATCCCAGGGCTCAACCCTGGACAGGCGGTGGAAACTACCAAGCTGGAGTACGGTAGGGGCAGAGGGAA<br>TTTCCGGTGGAGCGGTGAAATGCGTAGAGATCGGAAAGAACACCAACGGCGAAAGCACCCTGCTGGGCCGACACTGACACTGAGAGACGAAAGCTAGGGG |
| P-asv4498 | GCTAGCGTTGTTCGGAATTACTGGGCGTAAAGCGCGCGTAGGCGGTTTTTTAAGTCAGAGGTGAAAGCCCGGGGCTCAACCCCGGAATAGCCTTTGAAACTGGAAAACTTGAATCTTGGAGAGGTCAGTGGAATT<br>CCGAGTGTAGAGGTGAAATTCGTAGATATTCGGAAGAACACCAGTGGCGAAGGCGACTGACTGGACAAGTATTGACGCTGAGGTGCGAAAGCGTGGGG |
| P-asv4499 | CCGGGCGTTATCCGGATTTATTGGGTTTAAAGGGAGCGTAGGCCGGGGATTAAGTGTGTTGTGAAATGTAGGCGCCCAACGTCTGACTTGCAGCGCATACTGGTTCCCTTGAGTACGCGCAACGCCGGCGGAAT<br>TCGTCGTGTAGCGGTGAAATGCTTAGATATGACGAAGAACCCCGATTGCGAAGGCAGCCGGCGGGAGCGCAACTGACGCTGAAGCTCGAAGGTGCGGGT |
| P-asv4500 | GCTAGCGTTGCTCGGAATCACTGGGCGTAAAGGGCGCGTAGGCGGCCATTTAAGTCGGGGGTGAAAGCCTGTGGCTCAACCACAGAATTGCCTTCGATACTGGGTGGCTTGAGTCCGGAAGAGGTTGGTGGAA<br>CTGCGAGTGTAGAGGTGAAATTCGTAGATATTCGCAAGAACACCGGTGGCGAAGGCGGCCAACTGGTCCGGAACTGACGCTGAGGCGCGAAAGCGTGGGG |
| P-asv4501 | GCAAGCATTAATCGGAATTACTGGACGTAAAGCGCGCGTAGGTGGTTTGTTAAGTTGGATGTGAAAGCCCCGGGCTCAACCTGGGAACTGCATTCAAAACTGACAAGCTAGAGTATGGTAGAGGGTGGTGGAATT<br>TCCTGTGTAGCGGTGAAATGCGTAGATATAGGAAGGAACACCAGTGGCGAAGGCGACCACCTGGACTGATACTGACACTGAGGTGCGAAAGCGTGGGG |
| P-asv4502 | GCAAGCGTTAATCGGAATTACTGGGCGTAAAGCGCACGCAGGCGGTTTGTTAAGTCAGATGTGAAATCCCCGGGCTCAACCTGGGAACTGCATCTGATACTGGCAAGCTTGAGTCTCGTAGAGGGGGGTAGAAT<br>TCCAGGTGTAGCGGTGAAATGCGTAGAGATCTGGAGGAATACCGGTGGCGAAGGCGGCCCCCTGGACGAAGACTGACGCTCAGGTGCGAAAGCGTGGGT |
| P-asv4503 | GCTAGCGTTGTTCGGAATTACTGGGCGTAAAGCGCACGTAGGCGGACCGGAAAGTTGGGGGTGAAATCCCGGGGCTCAACCTCGGAACTGCCTTCAAAACTACTGGTCTGGAGTTCGAGAGAGGTGAGTGGAA<br>TTCCGAGTGTAGAGGTGAAATTCGTAGATATTCGGAGGAACACCAGTGGCGAAGGCGGCTCACTGGCTCGATACTGACGCTGAGGTGCGAAAGCGTGGGG |
| P-asv4504 | GCAAGCGTTGTCCGGAATTATTGGGCGTAAAGAGCTCGTAGGCGGTTTGTCGCGTCTGCTGTGAAGTCCCGGGGCTCAACCCCGAGTCTGCAGTGGGTACGGGCAGACTAGAGTGCAGTAGGGGAGACTGGAG<br>TTCCTGGTGTAGCGGTGAAATGCGCGGATATCAGGAGGAACACCGATGGCGAAGGCAGGTCTCTGGGCTGTAACTGACGCTGAGGAGCGAAAGCATGGGG |
| P-asv4505 | GCAAGCGTTAATCGGAATTACTGGGCGTAAAGCGTGCGTAGGCGGTTTGTTAAGTCCGATGTGAAAGCCCTGGGCTCAACCTGGGAATGGCATTGGATACTGGCGAGCTAGAGTCAGGTAGAGGGTGGTGGAAT<br>TCCCGGTGTAGCGGTGAAATGCGTAGAGATCGGGAGGAACATCTGTGGCGAAGGCGGCCACCTGGACCATGACTGACGCTGAGGCACGAAAGCGTGGGG |
| P-asv4506 | GCAAGCGTTAATCGGAATTACTGGGCGTAAAGCGCGCGTAGGCGGTTTGATAAGCAGGATGTGAAAGCCCTGGGCTTAACCTAGGAATGGCATTCTGAACTGTCTGACTGGAGTATGGTAGAGGGTAGTGGAAT<br>TTCCTGTGTAGCGGTGAAATGCGTAGATATAGGAAGGAACACCAGTGGCGAAGGCGACTACCTGGACCAATACTGACGCTGAGGTGCGAAAGCGTGGGG |
| P-asv4507 | TCGAGTGGTCAGGAGGTTTATTGGGCCTAAAGCATCCGTAGCCGGTTGCATAAGTTTTCGGTTAAATCTATGCGCTTAGCGTATAGGCCGCCGAAAATACTGTGCAGCTAGGGAGTGGGAGAGGTACACGGTACT<br>CGATAGGAAGGGGTAAAATCCTTTGATCTATTGATGACCACCTGTGGCGAAGGCGGTGTACCAGAACACGTTCGACGGTGAGGGATGAAAGCTGGGGG |
| P-asv4508 | GCTAGCGTTATCCGGATTTACTGGGCGTAAAGGGTGCGTAGGTGGTTTCTTAAGTCAGGAGTGAAAGGCTACGGCTTAACCGTAGTAAGCTCTTGAAACTGGGAAACTTGAGTGCAGGAGAGGAAAGTGGAATTC<br>CTAGTGTAGCGGTGAAATGCGTAGATATTAGGAGGAACACCAGTAGCGAAGGCGGCTTTCTGGACTGTAACTGACACTGAGGCACGAAAGCGTGGGG |
| P-asv4509 | GCAAGCGTTGTCCGGAATTATTGGGCGTAAAGCGCGCGCAGGCGGTCCTTTAAGTCTGATGTGAAAGCCCACGGCTCAACCGTGGAGAGTCATTGGAAACTGGGGGACTTGAGTGCAGAAGAGGAGAGCGGAA<br>TTCCACGTGTAGCGGTGAAATGCGTAGAGATGTGGAGGAACACCAGTGGCGAAGGCGGCTCTCTGGTCTGTAACTGACGCTGAGGCGCGAAAGCGTGGGG |
| P-asv4510 | ACTAGCGTTGTTCGGAATTACTGGGCGTAAAGCGCGCGTAGGCGGACTATTAAGTCAGGGGTGAAATCCCGGGGCTCAACCCCGGAACTGCCCTTGATACTGGTAGTCTGGAGTTCTGGAGAGGTGAGTGGAAT<br>TCCGAGTGTAGAGGTGAAATTCGTAGATATTCGGAGGAACACCAGTGGCGAAGGCGGCTCACTGGCCAGATACTGACGCTGAGGTGCGAAAGTGTGGGG |
| P-asv4511 | GCGAGCGTTGTCCGGAATTATTGGGCGTAAAGAGCTTGTAGGCGGTTTGTCGCGTCTGCTGTGAAAGGCCGGGGCTTAACTCCGTGTATTGCAGTGGGTACGGGCAGACTAGAGTGCAGTAGGGGAGACTGGA<br>ATTCCTGGTGTAGCGGTGAAATGCGCAGATATCAGGAGGAACACCGATGGCGAAGGCAGGTCTCTGGGCTGTAACTGACGCTGAGGAGCGAAAGCATGGGG |
| P-asv4512 | GCTAGCGTTGTTCGGAATTACTGGGCGTAAAGCGCACGTAGGCGGATATTTAAGTCAGGGGTGAAATCCCAGAGCTCAACTCTGGAACTGCCTTTGATACTGGGTATCTTGAGTATGGAAGAGGTAAGTGGAATT<br>GCGAGTGTAGAGGTGAAATTCGTAGATATTCGCAGGAACACCAGTGGCGAAGGCGGCTTACTGGTCCATTACTGACGCTGAGGTGCGAAAGCGTGGGG |
| P-asv4513 | GCAAGCGTTATCCGGATTTACTGGGTTTAAAGGGTGCGCAGGCGGTAATATAAGTCAGTGGTGAAAGCTTCCAGCTCAACTGGGAAACTGCCATTGATACTGTAATACTTGAATTTGGGAGAGGTAGGCGGAATA<br>CATCATGTAGCGGTGAAATGCTTAGATATGATGTAGAACACCGATTGCGAAGGCAGCTTACTAGACCAACATTGACGCTCATGCACGAAAGCGTGGGG |
| P-asv4514 | TCGAGTGATGGCCGATCTTATTGGGCCTAAAGCGTCCGTAGCCGGTCGCGCAAGTCCATCGGGAAATCCGCCAGCCCAACTGGCGGGCGTCCGGTGGAACCTGCACGACTTGGGACCGGAAGACCCAACGGGT<br>ACGTCCGGGGTAGGAGTGAAATCCCGTAATCCTGGACGGACCGCCGATGGCGAAAGCACGTTGGGAAGACGGATCCGACGGTGAGGGACGAAAGCCAGGGT |
| P-asv4515 | GCTAGCGTTGTTCGGAATTACTGGGCGTAAAGCGCACGTAGGCTGTGCCGCAAGTCAGGGGTGAAATCCCAGAGCTCAACTCTGGAACTGCCTTTGAAACTGCGGTGCTAGAATCTCAGAGGGGGTGGCGGAAT<br>TCCAAATGTAGGGGTGAAATCCGTAGATATTTGGAGGAACACCGGTGGCGAAGGCGGCTACCTGGATGAGTATTGACGCTGAGGTGCGAAAGCGTGGGG |
| P-asv4516 | GCAAGCGTTATCCGGAATCATTGGGTTTAAAGGGTCCGCAGGCGGGCGTATAAGTCAGTGGTGAAAGCCCATCGCTTAACGATGGAACTGCCATTGATACTGTATGCCTTGAATTCGGTCGAAGTGGGCGGAATA<br>TGACATGTAGCGGTGAAATGCTTAGATATGTCATAGAACACCGATAGCGAAGGCAGCTCACTAGGCCTGGATTGACGCTCAGGGACGAAAGCGTGGGT |
| P-asv4517 | GCAAGCGTTGTCCGGATTTATTGGGCGTAAAGGGAACGCAGGCGGTCCTTTAAGTCTGATGTGAAAGCCTTCGGCTTAACCGAAGAATTGCATTGGAAACTGGAGGACTTGAGTGCAGAAGAGGAGAGTGGAAC<br>TCCATGTGTAGCGGTGAAATGCGTAGATATATGGAAGAACACCAGTGGCGAAAGCGGCTCTCTGGTCTGTAACTGACGCTGAGGTTCGAAAGTGTGGGT |
| P-asv4518 | GCTAGCGTTGTTCGGAATTACTGGGCGTAAAGGGCGCGTAGGCGGCGCTGTAAGTGAGATGTGAAAGACCTGGGCTCAACCTGGGAAGTGCATTTCATACTGCAGTGCTTGAGTGTAAGAGAGGAAAGTGGAAT<br>GATGCATGTAGAGGTGAAATTCGTAGATATGCATTGGAACACCAGTGGCGAAGGCGACTTTCTGGCTTACAACTGACGCTAAGGCGCGAAAGCGTGGGG |
| P-asv4519 | GCTAGCGTTGTTCGGAATCACTGGGCGTAAAGCGCACGTAGGCGGCCTGCTAAGTCAGGTGTGAAATCCCAGGGCTTAACCCTGGAATTGCACTTGAAACTGGTTGGCTTGAGATTGACAGAGGATAGCGGAAT<br>TCCGAGTGTAGAGGTGAAATTCGTAGATATTCGGAGGAACACCAGTGGCGAAGGCGGCTATCTGGGTCAATACTGACGCTAAGGTGCGAAAGCGTGGGG |
| P-asv4520 | GCAAGCGTTATTCGGAATTACTGGGCGTAAAGCGCGCGTAGGCGGATTGTTAAGTCAGTTGTGAAAGCCCTGGGCTCAACCTAGGAATGGCGATTGAAACTGGCAGTCTAGAGTTTAGTAGAGGGAAGGGGAAT<br>TTCTGGAGTAGCAGTGAAATGCGTAGATATCAGAAGGAACATCAGTGGCGAAGGCGCCTTCCTGGACTAAAACTGACGCTGAGGTGCGAAAGCGTGGGG |
| P-asv4521 | ACTAGCGTTATTCGGAATTACTGGGCGTAAAGCGCACGTAGGCGGCTTTGTAAGTAAGGGGTGAAAGCCTGGAGCTCAACTCCAGAACTGCCTTTTAGACTGCATCGCTTGAATCCGGGAGAGGTGAGTGGAATT<br>CCGAGTGTAGAGGTGAAATTCGTAGATATTCGGAAGAACACCAGTGGCGAAGGCGGCTCACTGGACCGGTATTGACGCTGAGGTGCGAAAGCGTGGGG |
| P-asv4522 | GCTAGCGTTGCTCGGAATTACTGGGCGTAAAGGGAGCGTAGGCGGACATTTAAGTCAGGGGTGAAATCCCAGAGCTCAACTCTGGAACTGCCTTTGATACTGGGTGTCTTGAGTGTGATAGAGGTAAGTGGAACT<br>CCGAGTGTAGAGGTGAAATTCGTAGATATTCGGAAGAACACCAGTGGCGAAGGCGACTTACTGGATCATTACTGACGCTGAGGCTCGAAAGCGTGGGG |
| P-asv4523 | GCAAGCGTTAATCGGAATTACTGGGCGTAAAGCGCGCGTAGGTGGTTTGTTAAGTTGAATGTGAAAGCCCCGGGCTCAACCTGGGAACTGCATCCAAAACTGGCAAGCTAGAGTATGGCAGAGGGTGGTGGAAT<br>TTCCTGTGTAGCGGTGAAATGCGTAGAGATCTGAAGGAATACCGGTGGCGAAGGCGGCCCCCTGGACAGATACTGACACTCAGATGTGAAAGCGTGGGG |
| P-asv4524 | GCGAGCGTTAATCGGAATTACTGGGCGTAAAGCGTGCGCAGGCGGCTATACAAGACAGGCGTGAAATCCCCGGGCTTAACCTGGGAATGGCGCCTGTGACTGTATAGCTAGAGTGTGTCAGAGGGGGGTAGAA<br>TTCCACGTGTAGCAGTGAAATGCGTAGAGATGTGGAGGAATACCGATGGCGAAGGCAGCCCCCTGGGATAACACTGACGCTCATGCACGAAAGCGTGGGG |
| P-asv4525 | GCAAGCGTTGTCCGGAATTATTGGGCGTAAAGAGCTCGTAGGCGGTTTGTCGCGTCTGGTGTGAAAACTCAAGGCTCAACCTTGAGCTTGCATCGGGTACGGGCAGACTAGAGTGCGGTAGGGGTGACTGGAAT<br>TCCTGGTGTAGCGGTGGAATGCGCAGATATCAGGAGGAACACCGATGGCGAAGGCAGGTCACTGGGCCGCAACTGACGCTGAGGAGCGAAAGCATGGGG |
| P-asv4526 | CCAAGCGTTGTTCGGAATTACTGGGCGTAAAGCGGGTGTAGGCGGCTATATAAGTCGGATGTGAAAGCCCTGGGCTCAACCCAGGAAGTGCATCCGATACTGCATAGCTTGAGTACTAGAGAGGGAAGTGGAAT<br>TCCTGGTGTAGCGGTGAAATGCGTAGATATCAGGAGGAACATCGGTGGCGAAGGCGGCTTCCTGGCTATGTACTGACGCTCAGACCCGAAAGCGTGGGT |
| P-asv4527 | GCGAGCGTTATTCGGAATTACTAGGCGTAAAGCGAGTGTAGGCGGACTCTTAAGTCCGACGTTAAATCTCCCGGCTTAACTGGGAGCTGTCGTCGGATACTGAGAGCCTCGAATGGGGTAGGGGGCAGCGGAAT<br>TCCCGGTGTAGCGGTGAAATGCGTAGATATCGGGAGGAACACCTATGGCGAAAGCAGCTGCCTGGGCCTCTATTGACGCTAAGACTCGAAAGCTGGGGG |
| P-asv4528 | GCGAGCGTTGTCCGGAATTACTGGGCGTAAAGAGCTCGTAGGTGGTTTGTCGCGTCGTTTGTGTAAGTCCACAGCTTAACTGTGGGACTGCAGGCGATACGGGCATAACTTGAGTGCTGTAGGGGAGACTGGAA<br>TTCCTGGTGTAGCGGTGAAATGCGCAGATATCAGGAGGAACACCGATGGCGAAGGCAGGTCTCTGGGCAGTAACTGACGCTGAGGAGCGAAAGCATGGGT |

| | |
|---|---|
| P-asv4529 | GCTAGCGTTATCCGGAATTACTGGGCGTAAAGGGTGCGTAGGTGGTTTCTTAAGTCAGAGGTGAAAGGCTACGGCTCAACCGTAGTAAGCCTTTGAAACTGAGAAACTTGAGTGCAGGAGAGGAGAGTAGAATTCCTAGTGTAGCGGTGAAATGCGTAGATATTAGGAGGAATACCAGTTGCGAAGGCGGCTCTCTGGACTGTAACTGACACTGAGGCACGAAAGCGTGGGG |
| P-asv4530 | GCAAGCGTTACTCGGAATCACTGGGCGTAAAGGGCGCGTAGGCGGGTTCGTAAGTCAGATGTGAAATCCAATAGCTCAACTATTGAACTGCATTTGAAACTACTTACCTAGAGCGTGGGAGAGGCAGATGGAATTAGTGGTGTAGGGGTAAAATCCGTAGATATCACTAGGAATACTCATTGCGAAGGCGATCTGCTGGAACTCTGCTGACGCTGAGGCGCGAAAGCGTGGGG |
| P-asv4531 | TCCCTTTTTATCCGGAATCACTGGGCGTAAAGCGTCTGTAGGTGGTTTAATAAGTCAACTGTTAAATCTTGAAGCTCAACTTCAAAATCGCAGTCGAAACTATTAGACTAGAGTATAGTAGGGGTAAAGGGAATTTCCAGTGGAGCGGTGAAATGCGTAGAGATTGGAAAGAACACCGATGGCGAAGGCACTTTACTGGGCTATTACTGACACTCAGAGACGAAAGCTAGGGT |
| P-asv4532 | GCGAGCGTTGTCCGGAATTATTGGGCGTAAAGCGCGCGCAGGCGGCTCTTTAAGTCAGGTGTGAAAGGCTGCGGCTCAACCGCAGAGCGGCACCTGAAACTGGAGAGCTTGAGTGCAGGAGAGGGGAGCGGAATTCCCGGTGTAGCGGTGGAATGCGTAGAGATCGGGAGGAACACCAGTGGCGAAGGCGGCTCTCTGGCCTGTAACTGACGCTGAGGCGCGAAAGCGTGGGG |
| P-asv4533 | GCAAGCGTTGTCCGGAATGATTGGGCGTAAAGCATCTGTAGGTGGTTTTTAAAGTCGACTGTTAAATCCCAGAGCTCAACTTTGGCAGGGCAGTCGAGCACTTAGAAACTTGAGTACGGTAGGGGTAGAGGGAATTCTTAGTGGAGCGGTAAAATGCACAGAGATTAGGAAGAACACCAGTGGCGAAAGCGCTCTACTGGGCCGTGACTGACACTGAGAGATGAAAGTTGGGGG |
| P-asv4534 | GCGAGCGTTGTCCGGAATTACTGGGCGTAAAGGGCGTGTAGGCGGCCTGTTAAGTCAGGGGTGAAAGACCGGGGCTCAACCCCGGGGGTGCCTCTGAAACTGGCGGGCTTGAGTGCTCTAGAGGGCAGTGGAATTCCCGGTGTAGCAGTGAAATGCGTAGAGATCGGGAGGAACACCAGTGGCGAAGGCGGCTGCCTGGGGAGTCACTGACGCTGAGGCGCGAAAGCTAGGGG |
| P-asv4535 | GCGAGCGTTGTCCGGAATTATTGGGCGTAAAGAGCTTGTAGGCGGCTTGTCGCGTCTGCTGTGAAAGCCCGGGGCTTAACCCCGGGTCTGCAGTGGGTACGGGCAGGCTAGAGTATGGTAGGGGAGACTGGAATTCCTGGTGTAGCGGTGGAATGCGCAGATATCAGGAGGAACACCGATGGCGAAGGCAGGTCTCTGGGCCATTACTGACGCTGAGAAGCGAAAGCGTGGGG |
| P-asv4536 | GCGAGCGTTGTCCGGAATTACTGGGCGTAAAGCGCGCGCAGGCGGTCCTTTAAGTCTGATGTGAAATTCCGCGGCTCAACCGCGGGTGTGCATTGGAAACTGGGGGACTTGAGTGCGGAAGAGGGGAGCGGAATTCCCGGTGTAGCGGTGAAATGCGTAGAGATCGGGAGGAACACCAGTGGCGAAGGCGGCTCTCTGGTCCGCAACTGACGCTGAGGCGCGAAAGCGTGGGG |
| P-asv4537 | CCGAGTGGTCGGGACGTTTATTGGGCCTAAAGCATCCGTAGCCGGTTCTACAAGTCTTCCGTTAAATCCACCTGCTTAACAGATGGGCTGCGGAAGATACTATAGAGCTAGGAGGCGGGAGAGGCAAGCGGTACTTAGTGGGTAGGGGTAAAATCCGTTGATCCATTGAAGACCACCAGTGGCGAAGGCGGCTTGCCAGAACGCACTCGACGGTGAGGGATGAAAGCTGGGGG |
| P-asv4538 | GCTAGCGTTGCTCGGAATCACTGGGCGTAAAGGGCGCGTAGGCGGCGTTTTAAGTCGGGGGTGAAAGCCTGTGGCTCAACCACAGAATGGCCTTCGATACTGGGACGCTTGAGTATGGTAGAGGTTGGTGGAACTGCGAGTGTAGAGGTGAAATTCGTAGATATTCGCAAGAACACTGGTGGCGAAGGCAATCCCCTGGGCCTGCACTGACGCTCATGCACGAAAGCGTGGGG |
| P-asv4539 | GCAAGCGTTATTCGGAATTATTGGGCGTAAAGGGCGCGTAGGCGGCCTGGTCAGTGGAAAGTGAAAGCCCTCGGCTTAACCGAGGAACTGCTTGCCATACTGCCTGGCTAGAGGATGGGAGAGGAGAGTGGAATTCCCGGTGTAGCGGTGAAATGCGTAGAGATCGGGAGGAACACCAGTGGCGAAGGCGGCTCTCTGGACCATTACTGACGCTGAGGCGCGAAAGCGTGGGT |
| P-asv4540 | GCAAGCGTTGTTCGGAATCATTGGGCGTAAAGGGCGCGTAGGCGGTCTCGTATGTCCGCTGTAAAAGCACACGGCTCAACCGTGTTGGGTCGGTGGAAACTGCGAGACTTGAGGCTGGAAGGGGCTAGTGGAATTCCCCATGTAGCGGTGAAATGCGTAGAGATGGGGGAGAACACTCGTGGCGAAGGCGGCTAGCTGGACCAGTTCTGACGCTGAGGCGCGAAAGCGTGGGG |
| P-asv4541 | GCTAGCGTTGTTCGGAATCACTGGGCGTAAAGGGCGCGTAGGCGGCCTTGGCAGTCGGTTTGTGAAAGCCCGGGGCTCAACCCCGGAGGTGCGGCCGAAACACCAAGGCTGGAGGCCGGAAGAGGAGGGTGGAATACCCAGTGTAGAGGTGAAATTCGTAGATATTGGGTGGAACACCGGTGGCGAAAGCGGCTCTCTGGTCCGGACCTGACGCTGAGGCGCGAAAGCGTGGGG |
| P-asv4542 | GCAAGCGTTGTCCGGAATTATTGGGCGTAAAGAGCTCGTAGGCGGTTTGTCGCGTCTGCTGTGAAAACGCAAGGCTTAACCTTGCGCCTGCAGTGAGCACGGGCAGACTAGAGTGCGGTAGGGGAGACTGGAATTCCTGGTGTAGCGGTGGAATGCGCAGATATCAGGAGGAACACCGATGGCGAAGGCAGGTCTCTGGGCCGTTACTGACGCTGAGGAGCGAAAGCATGGGG |
| P-asv4543 | GCAAGCGTTATCCGGAGTAACTGGGCGTAAAGCGCGCGCAGGCGGCGGTGTTGGCCCGGCGTGAAAGCCCCCGGCTCAACCGGGGAGGGTCGTCGGGGACCGCACCGCTTGAGGGCGGTAGGGGCTGGTGGAATGCCTGGTGTAGTGGTGAAATGCGTAGAGATCAGGCGGAACACCCGTGGCGAAGGCGGCCAGCTGGGCCGTCCCTGACGCTGAGGCGCGAAGGCGTGGGG |
| P-asv4544 | CCAAGCGTTAATCGGAATTACAGGGCGTAAAGCGTGCGCAGGCGGTTGTGCAAGACCGATGTGAAATCCCCGAGCTTAACTTGGGAATTGCATTGGTGACTGCACGGCTAGAGTGAGGCAGAGGGGGGTAGAATTCCACGTGTAGCAGTGAAATGCGTAGATATGTGGAGGAATACCGATGGCGAAGGCAGCCCCCTGGGATAACACTGACGCTCATGCACGAAAGCGTGGGG |
| P-asv4545 | GCTAGCGTTGTCCGGAATCACTGGGCGTAAAGGGCGCGTAGGCGGCCCGATAAGTAGGGGGTGAAATCTTGCGGCTCAACCGCAAGGCTGCCTTCTAAACTGTCGGGCTGGAGCACGGTAGAGGCAGATGGAATTCCCGGTGTAGCGGTGGAATGCGTAGAGATCGGGAAGAACACCAGTGGCGAAGGCGGTCTGCTGGGCCGTTGCTGACGCTGAGGCGCGACAGCGTGGGG |
| P-asv4546 | GCAAGCGTTAATCGGAATTACTGGGCGTAAAGCGCGCGTAGGCGGCTTGATAAGATGGGTGTGAAATCCCCGGGCTTAACCTGGGAACTGCATCCATAACTGTCTGGCTAGAGTACAGTAGAGGGTGGTGGAATTTCCTGTGTAGCGGTGAAATGCGTAGATATATGAAGGAACACCAGTGGCGAAGGCGACCACCTGGACTGATACTGACGCTGAGGTGCGAAAGCGTGGGG |
| P-asv4547 | GCAAGCGTTAATCGGAATTACTGGGCGTAAAGCGCGCGTAGGTGGCTTGATAAGTTGGATGTGAAATCCCCGGGCTCAACCTGGGAACTGCACCCAAAACTGTCTGGCTAGAGTGCGGTAGAGGGTAGTGGAATTTCCAGTGTAGCGGTGAAATGCGTAGATATTGGAAGGAACACCAGTGGCGAAGGCGACTACCTGGACTGACACTGACACTGAGGTGCGAAAGCGTGGGG |
| P-asv4548 | GCTAGCGTTGTTCGGAATTACTGGGCGTAAAGCGCGCGTAGGCGGCCATTCAAGTCAGAGGTGAAAGCCCGGGGCTCAACCCCGGAACTGCCTTTGAAACTAGATGGCTTGAATCTTGGAGAGGCGAGTGGAATTCCGAGTGTAGTGGTGAAATTCGTAGATATTCGGAAGAACACCAGCGGCGAAGGCGACTCGCTGGACAAGTATTGACGCTGAGGTGCGAAAGCGTGGGG |
| P-asv4549 | GCAAGCGTTGTCCGGAATTATTGGGCGTAAAGCGCTCGCAGGCGGTCTTTTAAGTCTGATGTGAAAGCCCACGGCTTAACCGTGGAGGGTCATTGGAAACTGGAGGACTTGAGTACAGAAGAGGAGAGTGGAATTCCACGTGTAGCGGTGAAATGCGTAGAGATGTGGAGGAACACCAGTGGCGAAGGCGACTCTCTGGTCTGTAACTGACGCTGAGGAGCGAAAGCGTGGGT |
| P-asv4550 | GCAAGCGTTGTCCGGAATTATTGGGCGTAAAGAGCTCGTAGGCGGTCTGTCGCGTCTGCTGTGAAAACGCAAGGCTTAACCTTGCGCCTGCAGTGGGTACGGGCAGACTAGAGTGCGGTAGGGGAGACTGGAATTCCTGGTGTAGCGGTGGAATGTGCAGATATCAGGAGGAACACCGATGGCGAAGGCAGGTCTCTGGGCCGTTACTGACGCTGAGGAGCGAAAGCATGGGG |
| P-asv4551 | GCAAGCGTTAATCGGAATTACTGGGCGTAAAGCGTGCGTAGGTGGTTTCTTAAGTCTGTCGTGAAAGCCCTGGGCTCAACCTGGGAATGGCGATGGATACTGGGAGGCTGGAGTGCGGTAGAGGATGGCGGAATTCCCGGTGTAGCGGTGAAATGCGTAGAGATCGGGAGGAACATCCGTGGCGAAGGCGGCCATCTGGACCAGCACTGACACTGAGGCACGAAAGCGTGGGG |
| P-asv4552 | GCGAGCGTTGTCCGGAGTTACTGGGCGTAAAGCGCACGTAGGCGGTCTGCTAAGTTCGGAGTGACAGTCACCAGCTTAACTGGTGAAGTACTTCGAAAACTGGCGGACTTGAGAGCGAAAGAGGGACACGGAATTCCGGGTGGAGCAGTGAAATGCGTAGAGATCCGGAGGAACACCAAAGGCGAAGGCAGTGTCCTGGCTCGTTTCTGACGCTGAGGTGCGAAAGCTAGGGG |
| P-asv4553 | GCAAGCGTTGTCCGGATTTATTGGGCGTAAAGCGCGTGTAGGCGGCCAGAAAAGTCCGCTGTGAAAACTCGAGGCTCAACATCGAGCTGTCGGTGGAAACTTTCTGGCTAGAGTCCGGAAGAGGAGAATGGAATTCCTGGTGTAGCGGTGAAATGCGCAGATATCAGGAAGAACACCCGTGGCGAAGGCGGTTCTCTGGGACGGTACTGACGCTGAGACGCGAAAGCGTGGGG |
| P-asv4554 | GCAAACGTTGTTCGGAATCACTGGGCGTAAAGGGCGTGTAGGCGGTTCGGTAAGTGGAATGTGAAATCCCCCGGCTTACCCGGGGAACTGCATCCCATACTGCTGGACTTGAGTGCGGGAGAGGATGGAGGAATTCCTGGTGCAGCGGTGAAATGCGTAGATATCAGGAGGAACACCTGTGGCGAAGGCGTCCATCTGGCCCGTTACTGACGCTGATGCGCGAAAGCTAGGGG |
| P-asv4555 | GCGAGCGTTAATCGGAATTACTGGGCGTAAAGCATGCGCAGGTGGTTCGTTGTGTCGGTGGTGAAAGCCCCGGGCTCAACCTGGGAATGGCCATGGAAACTGGCGAACTGGAGTGCGGCAGAGGGGGGTGGAATTCCGCGTGTAGCAGTGAAATGCGTAGAGATGCGGAGGAACACCGATGGCGAAGGCAGCCCCCTGGGCTAGCACTGACGCTCAGGCACGAAAGCGTGGGG |
| P-asv4556 | GCAAGCGTTATCCGGATTTACTGGGTGTAAAGGGAGCGCAGGCGGTACGGCAAGTCTGATGTGAAAGCCCGGGGCTCAACCCCGGTACTGCATTGGAAACTGTCGGACTAGAGTGTCGGAGGGGTAAGTGGAATTCCTAGTGTAGCGGTGAAATGCGTAGATATTAGGAGGAACACCAGTGGCGAAGGCGGCTTACTGGACGATTACTGACGCTGAGGCTCGAAAGCGTGGGG |
| P-asv4557 | GCAAGCGTTGTTCGGAATTACTGGGCGTAAAGGGTCTGTAGGTGGCCAACTAAGTCAGACGTGAAATCCCTCAGCTTAACTGGGGAACTGCGTCTGATACTGGATGGCTTGAGTGCAGGAGAGGAATGCGGAATTCCAGGTGTAGCGGTGAAATGCGTAGATATCTGGAGGAACACCGGTGGCGAAGGCGGCATCCTGGACCGACACTGACACTGAGGAGCGAAAGCCAGGGG |
| P-asv4558 | GCAAGCGTTGTCCGGAATCATTGGGCGTAAAGAGCGTGTAGGCGGCTCGGTAAGTCCGCTGTAAAAGTCCAGGGCTCAACCCTGGAAAGCCGGTGGAAACTGTCGAGCTAGAGTCCGGAAGAGGCGAGTGGAATTCCTGGTGTAGCGGTGAAATGCGCAGATATCAGGAGGAACACCAATGGCGAAGGCAGCTCGCTGGGACGGTACTGACGCTGAGACGCGAAAGCGTGGGG |
| P-asv4559 | GCGAGCGTTGTCCGGAATCACTGGGCGTAAAGGGCGCGTAGGCGGTCTGCTAAGCGTGTGGTGAAAGCTCGGGGCTCAACCCCGAGTCTGCCATGCGAACTGGTGGACTGGAGCACTGTAGAGGCAGGTGGAATTCCGGGTGTAGCGGTGGAATGCGTAGAGATCCGGAAGAACACCAGTGGCGAAGGCGGCCTGCTGGGCAGTTGCTGACGCTGAGGCGCGACAGCGTGGGG |
| P-asv4560 | GCGAGCGTTGTCCGGAATCACTGGGCGTAAAGGGCGCGTAGGCGGCTCGGTAAGGTTGCGGTGAAAGCCCGGGGCTCAACCCCGGGTCGGCCGTGGCAACTGCTGGGCTCGAGCACTGTAGAGGCAGGTGGAATTCCGGGTGTAGCGGTGGAATGCGTAGAGATCCGGAAGAACACCGGTGGCGAAGGCGGCCTGCTGGGCAGTAGCTGACGCTGATGCGCGACAGCGTGGGG |
| P-asv4561 | GCAAGCGTTGTCCGGATTTATTGGGCGTATAGAGCGTGTAGGCGGCCAGGTAGGTCTGCTGTGAAAACTCGAGGCTTAACCTCGAGATGTCGGCGGAAACCATCTGGCTAGAGTCCGGAAGAGGAGAGTGGAATTCCTGGTGTAGCGGTGAAATGCGCAGATATCAGGAAGAACACCCGTGGCGAAGGCGGCTCTCTGGGACGGTACTGACGCTGAGACGCGAAAGCGTGGGG |
| P-asv4562 | CCAAGCGTTGTCCGGATTTATTGGGCGTAAAGAGCTCGTAGGCGGTTCGGTAAGTCGGGTGTGAAAACTCTGGGCTCAACCCAGAGCCGCCACCCGATACTGCTGTGACTTGAGTTCGGTAGGGGAGCGGGGAATTCCTGGTGTAGCGGTGAAATGCGCAGATATCAGGAGGAACACCAGTGGCGAAGGCGCCGCTCTGGGCCGAGACTGACGCTGAGGAGCGAAAGCGTGGGT |
| P-asv4563 | GCAAGCGTTGTTCGGAATTACTGGGCGTAAAGGGCGCGTAGGCGGCCTGTTAAGTCGAACGTGAAATCCCCGGGCTCAACCCGGGAACTGCGTCCGATACTGGCAGGCTTGAATCCGGGAGAGGGATGCGGAATTCCAGGTGTAGCGGTGAAATGCGTAGATATCTGGAGGAACACCGGTGGCGAAGGCGGCATCCTGGACCGGCATTGACGCTGAGGTGCGAAAGCGTGGGG |
| P-asv4564 | GCGAGCGTTGTTCGGATTTACTGGGCGTAAAGGGCGCGTAGGCGGTGCCGCAAGTCAGGCGTGAAAGCCCCGGGCTCAACCTGGGAGGTGCACTTGGGACTGCGGCACTGGAGTTCGGGAGAGGAGCGTGGAATTCCCAGTGTAGAGGTGAAATTCGTAGATATTGGGAAGAACACCGGTGGCGAAGGCGGCGCTCTGGACCGAGACTGACGCTGAGGCGCGAAAGCGTGGGG |
| P-asv4565 | GCAAGCGTTATCCGGATTCACTGGGTTTAAAGGGTGCGTAGGTGGGTCTGTAAGTCAGTGGTGAAATCTCCAAGCTTAACTTGGAAACTGCCGTTGATACTATAGATCTTGAATGCTGTGGAGGTGAGCGGAATATGTCATGTGGCGGTGAAATGCTTAGATATGACATAGAACACCAATTGCGAAGGCAGCTCACTACCCAGATATTGACACTGAGGCACGAAAGCGTGGGG |
| P-asv4566 | GCAAGCGTTGTTCGGAATCACTGGGCTTAAAGCGCGTGTAGGCGGCCCGTCAAGTACCTTGTGAAATCCCTCGGCTCAACAGGGGAATGGCTGGGTATACTGACGGGCTTGAGGTTGGTAGGGGTGCCTGGAACTCTAGGTGGAGCGGTGAAATGCGTAGATATCTAGAGGAACGCCCGTGGCGAAAGCGGGGCACTGGGCCACACCTGACGCTAAGACGCGAAAGCGTGGGG |
| P-asv4567 | TCAAGCGTTGTCCGGATTTATTGGGCGTAAAGCGCCCGCAGGCGGCTTGCCAAGTTCGATGTGAAATCTCCCGGCTCAACTGGGAGGGTGCATCGAAAACTGGCAGGCTTGAGGCAGGGAGAGGGATGTGGAATTCCAGGTGTAGTGGTGAAATGCGTAGATATCTGGAGGAACACCAGTGGCGAAGGCGGCATCCTGGCCCTGTCCTGACGCTCAGGGGCGAAAGCGTGGGG |

| | |
|---|---|
| P-asv4568 | CCAAGCGTTGTTCGGATTCACTGGGCGTAAAGGGTGCGCAGGTGGGTTGGCAAGTTTGATGTGAAAGCCCCGGGCTTAACCCGGGAATGGCATTGAATACTGCTGATCTGGAGGATTGGAGGGGGGACTGGAATTCTTGGTGTAGCAGTGAAATGCGTAGATATCAAGAGGAACACCAGTGGCGAAGGCGAGTCCCTGGACAATACCTGACACTGAGGCACGAAAGCCAAGGG |
| P-asv4569 | GCAAGCGTTGTCCGGATTTACTGGGCGTAAAGAGCGCGCAGGCGGCTGGTCAAGTCCGATGTGAAATCTCCCGGCTCAACTGGGAAACTGCATCGGAAACTGATCGGCTTGAAGGTGGGAGAGGGTAGTGGAATTCCCGGTGTAGTGGTGAAATGCGTAGATATCGGGAGGAACACCAGTGGCGAAGGCACCTTTCTGGTTCATTTCTGACACTGAGATTCGAAAGCATGGGT |
| P-asv4570 | GCAAACGTTGTTCGGAATTACTGGGCATAAAGCGCGCGTAGGCCGTTCGCTAAGTCAGGTGTGAAAGCCCCCGGCTCAACCGGGGAATTGCATTTGATACTGGCGAGCTTGAGGATGAGATGGGTGGGCGGAACTCCAGGTGGAGCGGTGAAATGCGTAGATATCTGGAGGAACACCGGTGGCGAAAGCGGCCCACTGGACCATCACTGACGCTGAGGCGCGAAAGCTAGGGT |
| P-asv4571 | GCAAGCGTTGTTCGGATTTACTGGGCGTAGAGGGCGCGTAGGCGGCCACCGCAAGTCGGTTGTGAAATCTCTGGGCTTAACCCAGAAAGGTCAACCGATACTGCGGGGCTAGAGTGCAGAAGGGGCAACTGGAATTCTCGGTGTAGCGGTGAAATGCGTAGATATCGAGAGGAACACCTGCGGCGAAGGCGGGTTGCTGGGCTGACACTGACGCTGAAGCGCGAAAGCTAGGGG |
| P-asv4572 | GCGAGCGTTGTCCGGAATTATTGGGCGTAAAGAGCTCGTAGGCGGTTTGGATAGTCGGGTGTGAAAGCTCCGAGCTCAACTCGGAGACGCCACCCGATACTGCCATGACTAGAGTCCGGTAGGGGAGCAGGGAATTCCTGGTGTAGCGGTGAAATGCGCAGATATCAGGAGGAACACCGGTGGCGAAGGCGGGTCTCTGGGCCGATACTGACGCTGAGGAGCGAAAGCGTGGGG |
| P-asv4573 | GCTAGCGTTGCTCGGAATGACTGGGCGTAAAGGGCGCGTAGGCGGTTTGGACAGTCAGATGTGAAATTCCTGGGCTCAACCTGGGGGCTGCATTTGATACGTACAGACTGGAGTGAGGAAGAGGGTCGTGGAATTCCCAGTGTAGAGGTGAAATTCGTAGATATTGGGAAGAACACCGGTGGCGAAGGCGGCGACCTGGTCCTTTACTGACGCTGAGGCGCGAAAGCGTGGGG |
| P-asv4574 | GCAAGCGTTGTTCGGAATTACTGGGCGTAAAGGGCGCGTAGGCGGCCTTCTAAGTCAGACGTGAAATCCCTCGGCTCAACCGAGGAACTGCGTCTGATACTGGAAGGCTTGAATCCGGGAGAGGGATGTGGAATTCCAGGTGTAGCGGTGAAATGCGTAGATATCTGGAGGAACACCGGTGGCGAGGGCGGCATCCTGGACCGGTATTGACGCTGAGGCGCGAAAGCTAGGGG |
| P-asv4575 | GCAAGCGTTGTTCGGAGTTACTGGGCGTAAAGGGTGTGTAGGCGGTCTACTAAGTCTGGTGTGAAATCTCCCGGCTCAACTGGGAGGGTGCGCCGGAAACTGGAAGGCTAGAGTGCGGGATGGGAGAGTGGAATTCCTGGTGTAGCGGTGAAATGCGTAGATATCAGGAGGAACACCTGTGGTGTAGACGGCTCTCTGGACCGCAACTGACGCTGAGACACGAAAGCGTGGGT |
| P-asv4576 | GCTAGCGTTGTTCGGAATTACTGGGCGTAAAGCGCACGTAGGCGGACCGGAAAGTCAGAGGTGAAATCCCAGGGCTCAACCTTGGAACTGCCTTTGAAACTCCTGGTCTTGAGGTCGAGAGAGGTGAGTGGAATTCCGAGTGTAGAGGTGAAATGCGTAGATATCGGGAGGAACACCAGTGGCGAAGGCGACTACCTGGCCTGTTCTTGACGCTGAGGCGCGAAAGCTAGGGG |
| P-asv4577 | CTAGCGTTGCTCGGAATGACTGGGCGTAAAGGGCGCGTAGGCGGAAGTGACAGTCAGACGTGAAATTCCTGGGCTCAACCTGGGGACTGCGTTTGAGACGTTACATCTAGAGTGAGGAAGAGGGTTGTGGAATTCCCAGTGTAGAGGTGAAATTCGTAGATATTGGGAAGAACACCGGTGGCGAAGGCGGCAACCTGGTCCTTTACTGACGCTGAGGCGCGAAAGCGTGGGG |
| P-asv4578 | GCAAGCGTTGTTCGGAATTACTGGGCGTAAAGGGCTCGTAGGCGGCCAACTAAGTCAGACGTGAAATCCCTCGGCTTAACCGGGGAACTGCGTCTGATACTGGATGGCTTGAATCCGGGAGAGGGATGCGGAATTCCAGGTGTAGCGGTGAAATGCATAGATATGTCACAGAACACCAATTGCGAAGGCAGCTTACTATGGTTTTATTGACACTGAGGCACGAAAGCGTGGGG |
| P-asv4579 | CCGAGCGTTGTCCGGAGTTACTGGGCGTAAAGCGCGCGCAGGCGGCCCGCTGCGCCCGGCGTGAAAGCCCCCGGCACAACCGGGGAACGGCGTCGGGGACGGGCGGGCTTGAGGGCGGCAGGGGCTGGTGGAATGCCCGGTGTAGTGGTGAAATGCGTAGAGATCGGGCGGAACACCCGTGGCGAAGGCGGCCAGCTGGGCCGACCCTGACGCTGAGGCGCGAAGGCGTGGGG |
| P-asv4580 | CAAGCGTTGCTCGGAATTACTGGGCGTAAAGGGCGCGTAGGCGGAATGGCAAGTCGGTCGTGAAATCCTTCGGCTCAACCGAAGAACGTCGATCGAAACTGCCGTTCTTGAGGATTGCAGAGGAGACTGGAATTCCCGGTGTAGCGGTGAAATGTGTAGAGATCGGGAGGAACACCGGTGGCGAAGGCGGGTCTCTGGGCAATTCCTGACGCTGAGGCGCGAAAGCGTGGGG |
| P-asv4581 | GCGAGCGTTAATCGGAATTACTGGGCGTAAAGCGTGCGCAGGCGGTTACTTAAGTCAGGTGTGAAAGCCCTGGGCTTAACCTAGGAATTGCGCTTGAGACTGGGTAGCTTTGAGTGTGGCAGAGGGAGGTGGAATTCCACGTGTAGCGGTGAAATGCGTAGATATCGGGAGGAACACCTGTGGCGAAAGCGGCGCACTGGACCACAACTGACGCTGAGGCGCGAAAGCTAGGGG |
| P-asv4582 | GCAAGCGTTGTCCGGAATCATTGGGCGTAAAGAGCGTGTAGGCGGTCCGGTCAGTCGGTTGTGAAAGTCAAGGGCTCAACCCTTGAATGCCGGCCGATACTGCCGGGCTAGAGTCCGGAAGAGGTGAGTGGAATTCCTGGTGTAGCGGTGAAATGCGCAGATATCAGGAGGAACACCGATGGCGAAGGCAGCTCACTGGGACGGTACTGACGCTGGGACGCGAAAGCGTGGGG |
| P-asv4583 | GCAAGCGTTGTTCGGATTTATTGGGCGTAAAGGGTGCGTAGGCGGTTTGGCAAGTCTTGTGTGAAATCTTTGGGCTCAACCCAAAGTCTGCATAAGAAGCTGCCGGGCTTGGGTATGGGAGGGGTGAGTGGAATTTCCGGTGTAGCGGTGAAATGCGTAGATATCGGAAGGAACACCTGTGGCGAAAGCGGCTCACTGGACCATTACTGACGCTGAGGCACGAAAGCTAGGGG |
| P-asv4584 | GCCAGCGTTGTTCGGAATAACTGGGCGTAAAGGGTCCGTAGGCGGTGTGGCAAGTCGGGAGTGAAATCTCTGGGCTTAACTCAGGGGCTGCTTCCGAAACTGCTGTGCTAGAGTGCGAGAGAGGTGAGTGGAATTGCAGGCGTAGCGGTGAAATGCGTAGATATCTGCAGGAACACCCGTGGCGAAGGCGGCTCACTGGATCGCAACTGACGCTGAGAGACGAAAGCTAGGGG |
| P-asv4585 | GCGAGCGTTGTTCGGAATCACTGGGCGTAAAGCGCGTGTAGGCGGCCCACTAAGTCTGATGTGAAAGCCCGGGGCTCACCCCCGGAAGTGCATTGGAAACTGGTGGACTCGAGTCCCGGAGAGGAGAGTGGAATTCCCGGTGTAGCGGTGAAATGCGTAGATATCAGGAGGAACACCGGTGGCGAAGGCGGCTCTCTGGACGGTGACTGACGCTGAGACGCGAAAGCGTGGGG |
| P-asv4586 | GCAAGCGTTGTCCGGATTTACTGGGCGTAAAGAGCGTGTAGGCGGCCAGACAGGTCTGTTGTGAAAACTGGAGGCTTAACCTCCAGACGTCGACAGAAACCGTCTGGCTAGAGTCCGGAAGAGGAGAGTGGAATTCCTGGTGTAGCGGTGAAATGCGCAGATATCAGGAAGAACACCAATGGCGAAGGCAGCTCTCTGGGACGGTACTGACGCTGAGACGCGAAAGCGTGGGG |
| P-asv4587 | GCGAGCGTTAATCGGAATTACTGGGCGTAAAGCGTGCGCAGGCGGTAAGTTAAGTCTGCTGTGAAAGCCCCGGGCTCAACCTGGGAATGGCAGTGGATACTGGACAGCTAGAGTGCGGTAGAGGGTAGTGGAATTCCCGGTGTAGCAGTGAAATGCGTAGAGATCGGGAGGAACACCAGTGGCGAAGGCGGCTACCTGGACCAGCACTGACACTGAAGCACGAAAGCGTGGGG |
| P-asv4588 | GCGAGCGTTAATCGGAATTACTGGGCGTAAAGCGTGCGCAGGCGGTGAGTTAAGTCTGATGTGAAAGCCCCGGGCTCAACCTGGGAATGGCATTGGAAACTGGCTCGCTGGAGTATGGCAGAGGGGGGTGGAATTCCGCGTGTAGCAGTGAAATGCGTAGAGATGCGGAGGAACACCGATGGCGAAGGCAGCCCCCTGGGCCAATACTGACGCTCAGGCACGAAAGCGTGGGG |
| P-asv4589 | GCGAACGTTGTTCGGAATCACTGGGCTTAAAGGGCGCGTAGGCGGGTTGCCAAGTCCGGGGTGAAAGCCCCCAGCTCAACTGGGGAAGTGCCTTGGATACTGGTAATCTCGAGGGAGGTAGGGGCATGCGGAACTTCCGGTGGAGCGGTGAAATGCGTAGATATCGGAAGGAACGCCGGTGGCGAAAGCGGCGTGCTGGACCTCTTCTGACGCTGAGGCGCGAAAGCCAGGGG |
| P-asv4590 | GCAAGCGTTGTCCGGAATCATTGGGCGTAAAGCGCGTGTAGGCGGTTTGGTAAGTCCGCTCTGAAAGTCCGGGGCTCAACCCCGGAAGGCGGGTGGATACTGTCAGACTAGAGTGCAGAAGAGGCGAGTGGAATTCCCGGTGTAGCGGTGAAATGCGCAGATATCGGGAGGAACACCAATTGCGAAGGCAGCTCGCTGGGACGTAACTGACGCTGAGACGCGAAAGCGTGGGG |
| P-asv4591 | TCAAGCGTTGTCCGGATTTATTGGGCGTAAAGCGCCCGCAGGCGGCTTGCCAAGTTCGATGTGAAATCTCCCGGCTCAACTGGGAGGGTGCATCGAAAACTGGCAGGCTTGAGACAGGGAGAGGGATGTGGAATTCCAGGTGTAGTGGTGAAATGCGTAGATATCTGGAGGAACACCAGTGGCGAAGGCGGCATCCTGGCCCTGTCCTGACGCTCAGGGGCGAAAGCGTGGGG |
| P-asv4592 | GCAGGCGTTGTCCGGATTTATTGGGCGTAAAGAGCGTGTAGGCGGCCAGACAAGTCCGTTGTGAAAACTAGAGGCTTAACCTCTAGACGTCGATGGAAACTGTTTGGCTAGAGTCCGGAAGAGGAGAATGGAATTCCTGGTGTAGCGGTGAAATGCGCAGATATCAGGAAGAACACCCGTGGCTAAGGCGGTTCTCTAGTACGGTACTGACGCTGAGACGCGAAAGCGTGGGG |
| P-asv4593 | GCGAGCGTTGTCCGGATTTATTGGGCGTAAAGGGCTCGTAGGCGGTTGATCGCGTCGGAAGTGGAAACTTGATGCTTAACGTTGAGCATGCTTTCGATACGGGTTGACTTGAGGAAGGTAGGGGAGAATGGAATTCCTGGTGGATCGGTGGAATGCGCAGATATCAGGAGGAACACCAGTGGCGAAGGCGGTTCTCTGGGCCTTTCCTGACGCTGAGGAGCGAAAGCGTGGGG |
| P-asv4594 | GCAAGCGTTGTCCGGAATTATTGGGCGTAAAGAGCTTGTAGGCGGTTTGTCGCGTCTGCTGTGAAAACGCAAGGCTTAACCTTGCGCCTGCAGTGGGTACGGGCAGACTAGAGTGCGGTAGGGGAGACTGGAATTCCTGGTGTAGCGGTGGAATGCGCAGATATCAGGAGGAACACCGATGGCGAAGGCAGGTCTCTGGGCCGTTACTGACGCTGAGGAGCGAAAGCAAGGGG |
| P-asv4595 | ACGAGTAGTGGTCACTTTTATTGAGCCTAAAGCGTTCGTAGCCGGTTTTGTGAATCTTCAGATAAAGCCTGAAGCTTAACTCCAGAAAGTCTGAAGAGACTGCAAGACTTGAGATCGGGTGAGGTTAAACGTACTTTCAGGGTAGGGGTAAAATCCTGTAATCCCGGAAGGACGACCAGTGGCGAAAGCGTTTAACTAGAACGAATCTGACGGTAAGGAACGAAGGCTAGGGT |
| P-asv4596 | GCAAGCGTTGTCCGGAATTATTGGGCGTAAAGAGCTCGTAGGCGGTTTGTCGCGTCTGCTGTGAAAACGCAAGGCTTAACCTTGCACCTGCAGTGGGTACGGGCAGACTAGAGTGCGGTAGGGGAGACTGGAATTCCTGGTGTAGCGGTGGAATGCGCAGATATCAGGAGGAACACCGATGGCGAAGGCAGGTCTCTGGGCCGTTACTGACGCTGAGGAGCGAAAGCATGGGG |
| P-asv4597 | CCGAGCGTTGTCCGGAATCATTGGGCGTAAAGGGTACGTAGGCGGCCTAGTAAGTTAGAAGTGAAATAATATAGCTCAACTATATAAAGCTTTTAAAACTGCTAAGCTTGAGAGATGAAAGGGAAAGTGGAATTCCTAGTGTAGCGGTGAAATGCGCAGATATTAGGAAGAATACCAGTGGCGAAGGCGACTTTCTGGTCATCATCTGACGCTGAGGTACGAAAGCGTGGGT |
| P-asv4598 | GCAAGCGTTGTCCGGAATCATTGGGCGTAAAGCGCGCGCAGGCGGTCCCTTAAGTCTGATGTGAAAGCCCACGGCTCAACCGTGGAGGGTCATTGGAAACTGGGGGACTTGAGTGCAGAAGAGGAAAGCGGAATTCCAAGTGTAGCGGTGAAATGCGTAGAGATTTGGAGGAACACCAGTGGCGAAGGCGGCTTTCTGGTCTGTAACTGACGCTGAGGCGCGAAAGCGTGGGG |
| P-asv4599 | GCTAGCGTTGCTCGGAATTACTGGGCGTAAAGGGCGCGTAGGCGGACATTTAAGTCAGGGGTGAAATCCCAGAGCTCAACTCTGGAACTGCCTTTGATACTGGGTGTCTTGAGTGTGAGAGAGGTATGTGGAACTCCGAGTGTAGAGATGAAATTCGTAGATATTCGGAAGAACACCAGTGGCGAAGGCGACATACTGGCTCATTACTGACGCTGAGGCGCGAAAGCGTGGGG |
| P-asv4600 | GCTAGCGTTGTTCGGAATTACTGGGCGTAAAGCGCACGTAGGCGGACTTTTAAGTCAGGGGTGAAATCCCGGGGCTCAACCTCGGAACTGCCTTTGATACTGGAAGTCTGGAGTATGGAAGAGGTAAGTGGAATTCCGAGTGTAGAGGTGAAATTCGTAGATATTCGGAGGAACACCAGTGGCGAAGGCGGCTTACTGGTCCATTACTGACGCTGAGGTGCGAAAGCGTGGGG |
| P-asv4601 | TCCCTCTTTTTCCTTCCTTCTTTTTCTTAAAGGTCTCGCAGGCGGTTTCTTAAGTCTGATGTGAAAGCCCCCGGCTCAACCGGGGAGGGTCATTGGAAACTGGGAAACTTGAGTGCAGAAGAGGAGAGTGGAATTCCACGTGTAGCGGTGAAATGCGTAGAGATGTGGAGGAACACCAGTGGCGAAGGCGACTCTCTGGTCTGTAACTGACGCTGAGGAGCGAAAGCGTGGGG |
| P-asv4602 | GCTAGCGTTGTTCGGAATCACTGGGCGTAAAGCGCACGTAGGTGGATTGTTAAGTCAGGGGTGAAATGCCGAGGCTCAACCTCGGAACTGCCTTTGATACTGGCAATCTCGAGTCCGGGAGAGGTGAGTGGAACTGCGAGTGTAGAGGTGAAATTCGTAGATATTCGCAAGAACACCAGTGGCGAAGGCGGCTCACTGGCCCGGTACTGACACTGAGGTGCGAAAGCGTGGGG |
| P-asv4603 | CCAAGCGTTGTTCGGAATTACTGGGCGTAAAGGGTGCGTAGGCGGCTCTCTAAGTCAGGTGTGAAAGCCCCGAGCTTAACTCGGGAACTGCACTTGATACTGGAGGGCTAGAGATTGGAAGAGGTGAGTGGAATTCCTAGTGTAGAGGTGAAATTCGTAGATATTAGGAAGAATATCAGTGGCGAAGGCGGCTCACTGGTCCAATACTGACGCTAAGGCACGAAAGCGTGGGG |
| P-asv4604 | GCTAGCGTTGTTCGGATTTACTGGGCGTAAAGCGCACGTAGGCGGATTGGTCAGTTAGAGGTGAAATCCCGGAGCTCAACTCCGGAACTGCCTTTAATACTGCCAGTCTAGAGTCCGGAAGAGGTGAGTGGAACTCCTAGTGTAGAGGTGGAATTCGTAGATATTAGGAAGAACACCAGTGGCGAAGGCGGCTCACTGGTCCGGTACTGACGCTGAGGTGCGAAAGCGTGGGG |
| P-asv4605 | GCAAGCGTTATCCGGAATCACTGGGCGTAAAGGGCGCGTAGGCGGTTTGTTAAGTCCGATGTTAAAGACCGAGGCTCAACCTCGACACGGCGTTGGATACTGACAAGCTGGACGGTTGGAGAGGAAGGTAGAATTACCAGTGTAGCGGTGGAATGCGTAGATACTGGTAGGAATACCCATTGCGAAGGCAGCCTTCTGGACAACACGTGACGCTGAGGCGCGAAAGTGTGGGG |
| P-asv4606 | CCGAGTGGTGACCAGTATTATTGGGCTTAAAGTGTTCGTAGCCGGACCTGCAAGTCTGCTGTGAAATCCAGACGCTCAACGTTTGGAAGTGCAGTGGATACTGCGGGTCTAGGGACCGGACGGCGTCGAGGCTATTCCATGGGTAGGAGTGAAATCCTAAAATCCATGGAGGAGCACCTGTGGCGAAGCGCTCGGCGAGAACGGATCCGACAGTGAGGGACGAAGGCTAGGGT |

| | |
|---|---|
| P-asv4607 | GCTAGCGTTGTTCGGAATTACTGGGCGTAAAGCGCACGTAGGCGGATATTTAAGTCAGGGGTGAAATCCCGGGGCTCAACCCCGGAACTGCCTCTGATACTGGGTATCTTGAGTTCGGAAGAGGTGAGTGGAAT<br>TGCGAGTGTAGAGGTGAAATTCGTAGATATTCGCAGGAACACCAGTGGCGAAGGCGGCTCACTGGTCCGATACTGACGCTGAGGTGCGAAAGCGTGGGG |
| P-asv4608 | GCTAGCGTTGCTCGGAATTACTGGGCGTAAAGGGCGCGTAGGCGGATAGTTTAGTCAGAGGTGAAAGCCCAGGGCTCAACCTTGGAATTGCCTTTGATACTGGCTATCTTGAGTATGGAAGAGGTATGTGGAACT<br>CCGAGTGTAGAGGTGAAATTCGTAGATATTCGGAAGAACACCAGTGGCGAAGGCGACATACTGGTCCATTACTGACGCTGAGGCGCGAAAGCGTGGGG |
| P-asv4609 | GTTAGCGTTGTTCGGAATTACTGGGCGTAAAGCGCACGTAGGCGGACCATTAAGTCAGGGGTGAAATCCCGGGGCTCAACCCCGGAACTGCCTTTGATACTGGTGGTCTTGAGTTCGAGAGAGGTGAGTGGAAT<br>TCCGAGTGTAGAGGTGAAATTCGTAGATATTCGGAGGAACACCAGTGGCGAAGGCGGCTCACTGGCTCGATACTGACGCTGAGGTGCGAAAGTGTGGGG |
| P-asv4610 | GCGAACGTTATTCGGAATCACTGGGCTTAAAGCGCGTGTAGGCGGGCGACCACGTCGGACGTTGAAAGCCCCCGGCTCAACCGGGGGATCGGCGCCGATACGGGCCGCCTGGAGGGGCGTAGGGGGAGCTG<br>GAACTTCCGGTGGAGCGGTGAAATGCGTTGAGATCGGAAGGAACGCCCGCGGCGAAAGCGAGCTCCTGGACGCCTACTGACGCTGAGACGCGAAAGCCAGGGG |
| P-asv4611 | GCAAGCGTTGTTCGGAATTATTGGGCGTAAAGCGAGTGTAGGCGGTTTGGCAAGTCGGATGTGAAAGCCCAGGGCCCAACCCTGGAATTGCATCCGATACTGTCAGGCTAGAGTTCCGGAGAGGGTGGCGGAA<br>TTCCCGGTGTAGGGGTGAAATCCGTAGATATCGGGAGGAACACCAGTGGCGAAGGCGGCCACCTGGACGGTGACTGACGCTGAGACTCGAAAGCGTGGGT |
| P-asv4612 | GCAAGCGTTAATCGGAATTACTGGGCGTAAAGGGCTCGCAGGCGGTTAGTTTAGTCGGATGTGAAATCCCCGGGCTTAACCTGGGGTGGTCATTCGATACTGGCTAACTAGAGTAAGGTAGAGGATAGTGGAATT<br>TCCGGTGTAGCGGTGAAATGCGTAGATATCGGAAGGAACACCAGTGGCGAAGGCGGCTATCTGGACCTTAACTGACGCTCAGGAGCGAAAGCGTGGGG |
| P-asv4613 | GCAAGCGTTACTCGGAATTACTGGGCGTAAAGCGTGCGTAGGTGGTTGTTTAAGTCTGTTGTGAAAGCCCTGGGCTCAACCTGGGAACTGCAGTGGAAACTGGACGACTAGAGTGTGGTAGAGGGTAGCGGAAT<br>TCCCGGTGTAGCAGTGAAATGCGTAGAGATCGGGAGGAACATCCATGGCGAAGGCAGCTACCTGGACCAACACTGACACTGAGGCACGAAAGCGTGGGG |
| P-asv4614 | GCAAGCGTTGTCCGGAATCATTGGGCGTAAAGAGTTCGTAGGTGGTATGTTAAGTCTGGTGTTAAAGCCCGAAGCTCAACTTCGGTTCGGCACTGGATACTGGCAAACTTGAATGTGGTAGAGGTAAAGGGAATT<br>CCTGGTGTAGCGGTGAAATGCGTAGATATCAGGAGGAACATCGGTGGCGTAAGCGCTTTACTGGGCCATAATTGACACTGAGGAACGAAAGCCAGGGT |
| P-asv4615 | GCGAGCGTTAATCGGAATTACTGGGCGTAAAGGGTGCGCAGGCGGATGTTTAAGTTAGAAGTGAAATCCCCGGGCTTAACCTGGGAATTGCTTTTAAGACTGGGCATCTAGAGTTTGTCAGAGGGGGGTGGAAT<br>TCCAAGTGTAGCAGTGAAATGCGTAGAGATTTGGAGGAACACCAATGGCGAAGGCAGCCCCCTGGGATAAAACTGACGCTCATGCACGAAAGCGTGGGT |
| P-asv4616 | GCGAGCGTTAATCGGAATTACTGGGCGTAAAGCGTGCGCAGGCGGTTTTTTAAGATAGGCGTGAAATCCCCGGGCTCAACCTGGGAACTGCGCTTATGACTGGAAGACTCGAGTATGGCAGAGGGGGGTGGAAT<br>TCCACGTGTAGCAGTGAAATGCGTAGAGATGTGGAAGAACACCGATGGCGAAGGCGGCTCACTGGCTCGATACTGACGCTGAGGTGCGAAAGCGTGGGG |
| P-asv4617 | GCAAGCGTTATCCGGAATTATTGGGCGTAAAGGGTGCGTAGGTGGTTAGTTAAGTCTTTGGTGTAAGTGCAGTGCTTAACGCTGTGATGCTAAAGAAACTGGCTGACTTGAGTGTGGTAGAGGCAAGTGGAATTC<br>CATGTGTAGCGGTAAAATGCGTAAATATATGGAGGAACACCAGTGGCGAAGGCGGCTTGCTGGGCCACAACTGACACTGAGGCACGAAAGCGTGGGG |
| P-asv4618 | GCTAGCGTTGTTCGGAATCACTGGGCGTAAAGCGCGCGTAGGCGGCTTGTCAAGTCAGGGGTGAAATCCCAGAGCTCAACTCTGGAATTGCCTTTGAAACTGGCAAGCTTGAGTGTCGGAGGGGATAGCGGAAT<br>TCCTAATGTAGAGGTGAAATTCGTAGATATTAGGAGGAACACCGGTGGCGAAGGCGGCTATCTGGACGACAACTGACGCTGAGGCGCGAAAGCGTGGGG |
| P-asv4619 | GCAAGCGTTGTTCGGAATTACTGGGCATAAAGGGTGCGTAGGTGGCGCGGTGCGTCTGTGGTGAAAGTCTTCGGCCCAACCGGAGAATAGCCGCGGATACGGCCGTGCTTGAGTGCATTAGGGGTAGGGGGAA<br>CGAGCAGTGTAGCGGTGAAATGCGTAGATATTGCTCGGAAGGCCAGTGCCGAAGGGGCCCTACTGGGATGTAACTGACACTGAGGCACGAAAGCGTGGGG |
| P-asv4620 | GCAAGCGTTAATCGGAATCACTGGGCGTAAAGCGTGCGTAGGCGGCTTTGCAAGTCAGGGGTGAAAGCCCTCGGCTCAACCGAGGAATTGCCTTTGAAACTGCTTGGCTTGAGTACTGGAGAGGCAGGCGGAAT<br>TCCGGGTGTAGGGGTGAAATCCGTAGATATCCGGAGGAACACCAGTGGCGAAGGCGGCCTGCTGGACAGATACTGACGCTGAGGTACGAAAGCGTGGGG |
| P-asv4621 | GCAAGCGTTATTCGGAATAACTGGGCGTAAAGAGCGCGTAGGCGGTTCGTTAAGTCTGTTGTTAAATTGCCTGGCTCAACCAGGTGTCGGCAATAGAAACTGGCAAACTAGAGGACAAGAGAGAGGGGTGGAATT<br>CTCGGAGTAGCGGTAAAATGCGTAGATCTCGAGAGGAACACCGATGGCGAAGGCAGCCCCTTGGCTTGTATCTGACGCTCAAGTGCGAAAGCGTGGGG |
| P-asv4622 | GCGAACGTTAATCGGAATTACTGGGCGTAAAGCGCGCGTAGGCGGTCCGGCAAGTGGGATGTGAAAGCCCTGGGCTTAACCTGGGAACTGCATTCCAAACTATCGGGCTAGAGTATGGTAGAGGGAGGTGGAA<br>TTTCCTGTGTAGCGGTGAAATGCGTAGATATAGGAAGGAACACCAGTGGCGAAGGCGGCCTCCTGGACCAATACTGACGCTGAGGTGCGAAAGCGTGGGG |
| P-asv4623 | GCTAGCGTTGTTCGGAATTACTGGGCGTAAAGCGCGCGTAGGCGGGCTGGAAAGTTGGGGGTGAAATCCCGGGGCTCAACCCCGGAACTGCCTTCAAAACTGACGGTCTGGAGTTCGAGAGAGGTGAGTGGAA<br>TTGCGAGTGTAGAGGTGAAATTCGTAGATATTCGCAGGAACACCGGTGGCGAAGGCGGCTCACTGGCTCGATACTGACGCTGAGGCGCGAAAGCGTGGGG |
| P-asv4624 | GCTAGCGTTGCTCGGAATTACTGGGCGTAAAGGGAGCGTAGGCGGACATTTAAGTCAGAGGTGAAATCCCGGAGCTTAACTTCGGAACTGCCTTTGATACTGGATGTCTTGAGTGTGAGAGAGGTATGTGGAACT<br>CCGAGTGTAGAGGTGAAATTCGTAGATATTCGGAAGAACACCAGTGGCGAAGGCGACATACTGGCTCATTACTGACGCTGAGGCGCGAAAGCGTGGGG |
| P-asv4625 | GCTAGCGTTGCTCGGAATCACTGGGCGTAAAGGGCGCGTAGGCGGACATTTAAGTCGGGGGTGAAAGCCCAGGGCTCAACCCTGGAATTGCCTCCGATACTGGGTGTCTCGAGACCGGAAGAGGTGAGTGGAA<br>CTGCGAGTGTAGAGGTGAAATTCGTAGATATTCGCAAGAACACCAGTGGCGAAGGCGGCTCACTGGTCCGGTTCTGACGCTGAGGCGCGAAAGCGTGGGG |
| P-asv4626 | GCAAGCGTTAATCAGAATTACTGGGCGTAAAGCGCGCGTAGGTGGATATTTAAGTCGGATGTGAAAGCCCTGGGCTTAACCTGGGAATTGCACCCGATACTGAGTATCTAGAGTATGGTAGAGGGAAGTGGAATT<br>TCCGGTGTAGCGGTGAAATGCGTAGATATCGGAAAGAACACCAGTGGCGAAGGCGACTTCCTGGACCAATACTGACACTGAGGTGCGAAAGCGTGGGG |
| P-asv4627 | GCGAGCGTTATCCGGAATTACTGGGTTTAAAGGGTGCGTAGGCGGGTTATTAAGTCAGTGGTGAAATGGTGTCGCTTAACGATAGAATTGCCATTGATACTGATAATCTTGAATTGGGTTGAGGTTAGCGGAATGT<br>GACATGTAGCGGTGAAATGCATAGATATGTCATGGAACACCAATTGCGAAGGCAGCTAACTAGGCCTTGATTGACGCTGAGGCACGAAAGCGTGGGT |
| P-asv4628 | GCTAGCGTTGTTCGGAATTACTGGGCGTAAAGCGCGCGTAGGCGGATCGATCAGTCAGAGGTGAAATCCCGGAGCTCAACTTCGGAACTGCCTTTGATACTGTCGATCTGGAGTCCGGGAGAGGTGCATGGAAT<br>TCCGAGTGTAGAGGTGAAATTCGTAGATATTCGGAGGAACACCAGAGGCGAAGGCGATGCACTGGACCGGTACTGACGCTGAGGTGCGAAAGCGTGGGG |
| P-asv4629 | GCAAGCGTTGTTCGGAATTACTGGGCGTAAAGCGTGCGTAGGCGGCCAAGCAAGTTGGAAGTGAAATCCCAGGGCTCAACCCTGGAACTGCTTTCAAAACTGCTAGGCTAGAATCTCGGAGAGGATAGCGGAAT<br>TTCCAGTGTAGAGGTGAAATTCGTAGATATTGGAAGGAACACCAGTGGCGTAGGCGGCTATCTGGACGAGTATTGACGCTAAGGCACGAAAGTGTGGGG |
| P-asv4630 | GCAAGCGTTAATCGGAATTACTGGGCGTAAAGCGCGCGTAGGTTGTTTGCTAAGCGAGATGTGAAAGCCCCGGGCTCAACCTGGGAACGGCATTTCGAACTGGCAAGCTAGAGTACAGTAGAGGGTGGCGGAAT<br>TTCCTGTGTAGCGGTGAAATGCGTAGAGATGGGAAGGAACATCAGTGGCGAAGGCGGCCACCTGGACTGATACTGACACTGAGGTGCGAAAGCGTGGGG |
| P-asv4631 | GCAAGCGTTATCCGGATTTATTGGGCGTAAAGGGCTCGTAGGCGGTTCGTCGCGTCCGGTGTGAAAGTCCATCGCTTAACGGTGGATCCGCGCCGGGTACGGGCGGGCTTGAGTGCGGTAGGGGAGACTGGA<br>ATTCCCGGTGTAACGGTGGAATGTGTAGATATCGGGAAGAACACCAATGGCGAAGGCAGGTCTCTGGGCCGTTACTGACGCTGAGGAGCGAAAGCGTGGGG |
| P-asv4632 | GCGAGCGTTGTCCGGAATTACTGGGCGTAAAGAGCTCGTAGGCGGTTTGTCGCGTCGTCTGTGAAAACCCGCAGCTCAACTGCGGGCTTGCAGGCGATACGGGCAGACTTGAGTACTGCAGGGGAGACTGGAA<br>TTCCTGGTGTAGCGGTGAAATGCGCAGATATCAGGAGGAACACCGGTGGCGAAGGCGGGTCTCTGGGCAGTAACTGACGCTGAGGAGCGAAAGCGTGGGT |
| P-asv4633 | CCAAGCGTTATCCGGAATCATTGGGTTTAAAGGGTCCGTAGGCGGTCTAGTAAGTCAGTGGTGAAAGCCCATCGCTCAACGGTGGAACGGCCATTGATACTGCTGGACTTGAATTATTAGGAAGTAACTAGAATAT<br>GTAGTGTAGCGGTGAAATGCTTAGAGATTACATGGAATACCAATTGCGAAGGCAGGTTACTACTAATGGATTGACGCTGATGGACGAAAGCGTGGGT |
| P-asv4634 | GCAAGCGTTGTTCGGAATCATTGGGCGTAAAGCGCGCGCAGGCGGCTTGCCAAGTCTGATGTGAAATCCCGGGGCTCAACCTCGGAACTGCATTGGAAACTGGTAGGCTTGAGTATGTGAGGGGTTAGTGGAAT<br>TCCTGGTGTAGGGGTGAAATCCGTAGAGATCAGGAGGAACACCGGTGGCGAAGGCGACTGACTGGAACATAACTGACGCTGAGGCGCGAAAGCGTGGGT |
| P-asv4635 | GCTAGCGTTGCTCGGAATTACTGGGCGTAAAGGGCGCGTAGGCGGCACGATTAGTCAGGCGTGAAATTCCTGGGCTCAACCTGGGGACTGCGCTTGATACGGTCGAGCTAGAGGATGGAAGAGGGTCGCGGAA<br>TTCCCAGTGTAGAGGTGAAATTCGTAGATATTGGGAAGAACACCGGTGGCGAAGGCGGCGACCTGGTCCATTACTGACGCTGAGGCGCGACAGCGTGGGG |
| P-asv4636 | CCAAACGTTATTCGGAATCACTGGGCTTAAAGCGTGCGTAGGCGGTCCGGAAAGTAGGGTGTGAAATCCCACAGCTCAACTGTGGAACTGCGCTCTAAACTACCGGGCTAGAGGGATATAGAGGTGAGCAGAAC<br>AGATGGTGGAGCGGTGAAATGCGTTGATATCATCTGGAATACCGGAGGCGAAGGCGGCTCACTGGGTATCTTCTGACGCTGAGGTACGAAAGCTAGGGG |
| P-asv4637 | GCAAGCGTTATCCGGAATCATTGGGCGTAAAGCGTCCGCAGGTGGCACTTCAAGTCTGCTGTCAAAGGTCGGGGCTCAACTCCGAATCGGCGGTGGAAACTGAAAGCTAGAGTGCGGTAGGGGCAGAGGGAA<br>TTCCTGGTGTAGCGGTGAAATGCGTAGAGATCAGGAAGAACACCGGTGGCGAAAGCGCTCTGCTAGGCCGCAACTGACACTCAGGGACGAAAGCTAGGGG |
| P-asv4638 | GCTAGCGTTGTTCGGAATCACTGGGCGTAAAGCGCACGTAGGCGGATTTGCCAGTCAGGGGTGAAATCCCGGGGCTCAACCTCGGAACTGCCTTTGATACAGCAAGTCTAGAGTCCGGAAGAGGTGAGTGGAAT<br>TCCTAGTGTAGAGGTGAAATTCGTAGATATTAGGAGGAACACCAGTGGCGAAGGCGGCTCACTGGTCCGGCACTGACGCTGAGGTGCGAAAGCGTGGGG |
| P-asv4639 | GCGAACGTTATTCGGAATCACTGGGCTTAAAGCGCGTGTAGGCGGATGACCACGTCGACGGCTGAAAGCCCCCGGCTCAACCGGGGGATCGGCCTCGATACGGGCCATCTGGAGGGGCGTAGGGGGAGCTGG<br>AACTCACGGTGGAGCGGTGAAATGCGTTGAGATCGTGAGGAACGCCCGCGGCGAAAGCGAGCTCCTGGACGCCTGCTGACGCTGAGACGCGAAAGCCAGGGG |
| P-asv4640 | GCAAGCGTTGTCCGGATTTATTGGGTTTAAAGGGTGCGTAGGCGGCCTTTTAAGTCAGCGGTGAAAGACCCCGGCTCAACCGGGGCAGTGCCGTTGATACTGGAAGGCTTGAGTGTTGGAGGGGTACATGGAAT<br>TGATGGTGTAGCGGTGAAATGCATAGATACCATCAGGAACACCGATAGCGAAGGCATTGTACTGGCCAACAACTGACGCTGAGGCACGAAAGCATGGGT |
| P-asv4641 | GCGAGCGTTAATCGGAATTACTGGGCGTAAAGCGTGCGCAGGCGGTTTTGTAAGACAGGTGTGAAATCCCCGGGCTTAACCTGGGAACTGCGCTTGTGACTGCAAGACTTGAGTACGGCAGAGGGGGGTAGAAT<br>TCCACGTGTAGCAGTGAAATGCGTAGAGATGTGGAGGAATACCGATGGCGAAGGCAGCCCCCTGGGTCGATACTGACGCTCATGCACGAAAGCGTGGGG |
| P-asv4642 | GCGAACGTTATTCGGAATCACTGGGCTTAAAGCGCGTGTAGGCGGATCGGAACGTCGGAGCCTGAAATCCCCCGGCTCAACCGGGGAACGCGGCCCGAAACGACCGGTCTCGAGGGGTGTAGGGGGGACTGG<br>AACTTCCGGTGGAGCGGTGAAATGCGTTGAGATCGGAAGGAACGCCCGTGGCGAAAGCGAGTCCCTGGACGCCTGCTGACGCTGAGACGCGAAAGCTAGGGG |
| P-asv4643 | GCAAGCGTTATCCGGATTTACTGGGTGTAAAGGGAGCGCAGGCGGCATGATAAGTCTGATGTGAAAACCCAAGGCTCAACCATGGGACTGCATTGGAAACTGTCGTGCTGGAGTGTCGGAGAGGTAAGCGGAAT<br>TCCTAGTGTAGCGGTGAAATGCGTAGATATTAGGAGGAACACCAGTGGCGAAGGCGGCTTACTGGACGATGACTGACGCTGAGGCTCGAAAGCGTGGGG |
| P-asv4644 | CCGGGCGTTATCCGGATTTATTGGGTTTAAAGGGAGCGTAGGCTGCCGGTTAAGTGTGTTGTGAAATGCAGGCGCTCAACGTCTGCCTTGCAGCGCATACTGTCCGGCTTGAGTACGCACAACGTTGGCGGAAT<br>TCGTCGTGTAGCGGTGAAATGCTTAGATATGACGAAGAACTCCGATTGCGAAGGCAGCTGACGGGAGCGTTACTGACGCTGAAGCTCGAAAGTGCGGGT |
| P-asv4645 | GCGAGCGTTGTCCGGATTTATTGGGCGTAAAGGGCTCGTAGCTGGTTGATCGCGTCGGAAGTGTAATCTTGGGGCTTAACCCTGAGCGTGCTTTCGATACGGGTTGACTTGAGGAAGGTAGGGGAGAATGGAAT<br>TCCTGGTGGAGCGGTGGAATGCGCAGATATCAGGAGGAACACCAGTGGCGAAGGCGGTTCTCTGGGCCTTTCCTGACGCTGAGGAGCGAAAGCGTGGGG |

| | |
|---|---|
| P-asv4646 | GCTAGCGTTGCTCGGAATTACTGGGCGTAAAGGGCGCGTAGGCGGACAGTTAAGTTGGGGGTGAAAGCCCGGGGCTCAACCTCGGAAATGCCTTCAATACTGGCTGTCTTGAGTACGGGAGAGGTGAGTGGAACTCCGAGTGTAGAGGTGAAATTCGTAGATATTCGGAAGAACACCAGTGGCGAAGGCGACTCACTGGCCCGTTACTGACGCAGAGGCGCGAAAGCGTGTGG |
| P-asv4647 | GCAAGCGTTGTCCGGAATCATTGGGCGTAAAGAGCGTGTAGGCGGTCCGATAAGTCCGCTCTGAAAGCCCGGGGCTCAACCCCGGGAGGCGGGCGGATACTGTCGGGCTAGAGTCCGGAAGAGGCAAGTGGAATTCCTGGTGTAGCGGTGAAATGCGCAGATATCAGGAGGAACACCGATGGCGAAGGCAGCTTGCTGGGACGGTACTGACGCTGAGACGCGAAAGCGTGGGG |
| P-asv4648 | GCGAGCGTTGTCCGGAATTATTGGGCGTAAAGGGCTCGTAGGCGGTTTGTCGCGTCGGGAGTGAAAACCAGGTGCTTAACACCTGGCTTGCTTTCGATACGGGCAGACTAGAGGTATTCAGGGGAGAACGGAATTCCTGGTGTAGCGGTGAAATGCGCAGATATCAGGAGGAACACCGGTGGCGAAGGCGGTTCTCTGGGAATGACCTGACGCTGAGGAGCGAAAGTGTGGGG |
| P-asv4649 | ACGAGCGTTATCCGGATTTATTGGGTTTAAAGGGTGCGCAGGTGGTCCTGCAAGTCAGTGGTGAAAAGCTGAGGCTCAACCTCAGCCTTGCCGTTGAAACTGCAAGACTTGAGAGTACATGATGTGGGCGGAATGCGTAGTGTAGCGGTGAAATGCATAGATATTACGCAGAACTCCGATTGCGAAGGCAGCTCACAAAGGTATATCTGACACTGAGGCACGAAAGCGTGGGG |
| P-asv4650 | GCGAGTGGTCGGGACGTTTACTGGGCCTAAAGCGCCTGTAGCCGGCTCACTAAGTTGCTCCTCAAAGCCCCGGGCTCAACCCGGGGACTGGGAGCAATACTAGTGAGCTAGGGGGCGGGAGAGGCCAGGGGTACTCCCGGATAAGGGGCGAAATCCGTAGATCCCGGGAGGACCACCAGTGGCGGAGGCGCCTGGCTAGAACGCGCCCGACGGTGAGAGGCGAAAGCCGGGGC |
| P-asv4651 | GCAAGCGTTATCCGGAATTATTGGGCGTAAAGCGCGCGTAGGCGGTTTTTTAAGTCTGATGTGAAAGCCCACGGCTCAACCGTGGAGGGTCATTGGAAACTGTAAAACTTGAGTGCGGAAGAGGAAAGTGGAATTCCATGTGTAGCGGTGAAATGCGCAGAGATATGGAGGAACACCAGTGGCGAAGGCGACTTTCTGGTCTGTAACTGACGCTGATGTGCGAAAGCGTGGGG |
| P-asv4652 | GCAAGCGTTGTCCGGAATTATTGGGCGTAAAGAGCTCGTAGGCGGTTTGTCGCGTCTGCTGTGAAAGCCCGGGGCTCGACCCCGGGTCTGCAGTGGGTACGGGCAGACTAGAGTGCAGTAGGGGAGACTGGAATTCCTGGTGTAGCGGTGAAATGCGCAGATATCAGGAGGAACACCGATGGCGAAGGCAGGTCTCTGGGCTGTTACTGACGCTGAGGAGCGAAAGCATGGGG |
| P-asv4653 | GCTAGCGTTGCTCGGAATCACTGGGCGTAAAGGGTGCGCAGGTGGATCTTTAAGTCAGGGGTGAAATCCCAGGGCTTAACCCCGGAACTGCCTGTGAGACCTCAGTACTAGAGTGTGAGAGGGGATAGTGGAATACCCAGTGTAGCGGTGAAATGCGTAGAGATTGGGTGGAACACCGGTGGCGAAGGCGGCTATCTGGCTCACAACTGACGATCAGGCACGAAAGCGTGGGG |
| P-asv4654 | GCGAGCGTTAATCGGAATTACTGGGCGTAAAGCGAGCGCAGACGGTTACTTAAGCAAGATGTGAAATCCCCGGGCTTAACCTGGGAACTGCGTTTTGAACTGAGTGACTAGAGTATGTCAGAGGGGGGTAGAATTCCACGTGTAGCAGTGAAATGCGTAGAGATGTGGAGGAATACCGATGGCGAAGGCAGCCCCCTGGGATAATACTGACGTTCATGCTCGAAAGCGTGGGT |
| P-asv4655 | CCAAGCGTTATCCGGATTTATTGGGCGTAAAGCGAGCGCAGACGGTTATTTAAGTCTGAAGTGAAAGCCCTCAGCTCAACTGAGGAATTGCTTTGGAAACTGGATGACTTGAGTGCAGTAGAGGAAAGTGGAACTCCATGTGTAGCGGTGAAATGCGTAGATATATGGAAGAACACCAGTGGCGAAGGCGGCTTTCTGGACTGTAACTGACGTTGAGGCTCGAAAGTGTGGGT |
| P-asv4656 | GCAAGCGTTGTTCGGAATTACTGGGCGTAAAGGGCGCGTAGGCGGTTCGCTAAGTCGGATGTGAAAACTCTGGGCTCAACCCAGAGACTGCATCCGAAACTGACGGGCTAGAGTTCTGGAGGGGGTAGCGGAATTCCTGGTGTAGCGGTGAAATGCGTAGATATCAGGAGGAACACCGGTGGCGAAGGCGGCTACCTGGACAGAAACTGACGCTGAGGCGCGAAAGCTAGGGG |
| P-asv4657 | GCAAGCGTTGTCCGGATTTACTGGGCGTAAAGAGCGCGCAGGCGGTCGGATTAGTCGCGTGTGAAAGCCCCCGGCTCAACTGGGGAGGGTCACGCGATACTGTTCGACTCGAAGGCAGGAGAGGGAAGTGGAATTCCCGGTGTAGTGGTGAAATGCGTAGATATCGGGAGGAACACCAGTGGCGAAGGCGACTTCCTGGCCTGTTCTTGACGCTGAGGCGCGAAAGCTAGGGT |
| P-asv4658 | ACAAACGTTATTCGGAATCACTGGGCTTAAAGGGTGCGTAGGCGGCCACGTAAGTGAGATGTGAAATCCCTCGGCTCAACCGAGGAAGTGCATTTCATACTGCATGGCTTGAGGGAGACAGGGGTAAGCGGAACTGATGGTGGAGCGGTGAAATGCGTTGATATCATCAGGAACACCGGTGGCGAAAGCGGCTTACTGGGTCTCTTCTGACGCTGAGGTACGAAAGCTAGGGT |
| P-asv4659 | GCAAGCGTTGTTCGGATTTATTGGGCGTAAAGGGTGCGTAGGCGGTTTGGCAAGTCTTGTGTGAAATCTTCGGGCTCAACCCGAAGTCTGCATAAGAAACTGCCGGGCTTGAGTATGGGAGAGGTGAGTGGAATTTCCGGTGTAGCGGTGAAATGCGTAGATATCGGAAGGAACACCTGTGGCGAAAGCGGCTCACTGGACCATAACTGACGCTGATGCACGAAAGCTAGGGG |
| P-asv4660 | GCAAGCGTTATCCGGATTTACTGGGTGTAAAGGGAGCGCAGGCGGCTTGGCAAGTCTGATGTGAAAGGCTGGGGCTCAACCCCAGGACTGCATTGGAAACTGTTAGGCTGGAGTGTCGGAGAGGTAAGCGGAATTCCTAGTGTAGCGGTGAAATGCGTAGATATTAGGAGGAACACCAGTGGCGAAGGCGGCTTACTGGACGACAACTGACGCTGAGGCTCGAAAGCGTGGGG |
| P-asv4661 | ACAAGCGTTGTCCGGATTTACTGGGCGTAAAGAGCGCGCAGGCGGTCGTTCAAGTCGAGTGTGAAAGCCCCCGGCTCAACTGGGGAGGGTCACTCGATACTGATCGACTCGAAGGCAGGAGAGGGAAGTGGAATTCCCGGTGTAGTGGTGAAATGCGTAGATATCGGGAGGAATACCGGTGGCGAAGGCGGCATCCTGGACCGAGATTGACACTGAGTGAGCGAAAGCCAGGGG |
| P-asv4662 | GCAAGCGTTGCTCGGAATCACTGGGCGTAAAGGGCGCGTAGGCGGGATGGCAAGTCAGTCGTGAAATCTCTCGGCTCAACCGAGAAACGTCGGCTGATACTGCCGTTCTTGAGGGATGCAGAGGAGACTGGAATTCCCGGTGTAGCGGTGAAATGTGTAGAGATCGGGAGGAACACCGGTGGCGAAGGCGGGTCTCTGGGCATTTCCTGACGCTGAGGCGCGAAAGCGTGGGG |
| P-asv4663 | GCGAGCGTTGTCCGGATTTACTGGGCGTAAAGAGCGCGTAGGCGGAGACTTAAGTGTGGCCTGAAAGCCCCCGGCTCAACCGGGGAGGGTGGCTGCAAACTGGGTTTCTAGGGGGGGATAGAGGAAAGTGGAATTCCGGGTGTAGCGGTGAAATGCGTAGAGATCCGGAGGAACACCAGTGGCGAAGGCGGCTTTCTGGGTCCGCCCTGACGCTGAGGCGCGAAAGCGTGGGG |
| P-asv4664 | GCTAGCGTTGTTCGGAATTACTGGGCGTAAAGGGCGCGTAGGCGGCTGTGCAAGTCAGGTGTGAAAGCCCTGGGCTCAACCCAGGATGTGCACTTGATACTGCGCGGCTAGAGTGTCGGAGAGGATGGTGGAATTCCCAGTGTAGAGGTGAAATTCGTAGATATTGGGAAGAACACCGGTGGCGAAGGCGACCATCTGGACGACAACTGACGCTGAGGCGCGAAAGCGTGGGG |
| P-asv4665 | GCTGGCGTTGTCCGGAATCATTGGGCGTAAAGCGCGTGTAGGCGGTCCGGTAAGTCCGCTGTGAAAGTCGGGGGCTCAACCCTCGAATGCCGGTGGATACTGTCGGGCTAGAGTGCGGAAGAGGCGAGTGGAATTCCTGGTGTAGCGGTGAAATGCGCAGATATCAGGAGGAACACCAATTGCGAAGGCAGCTCGCTGGGACGTTACTGACGCTGAGACGCGAAAGCGTGGGG |
| P-asv4666 | GCAAGCGTTGTTCGGAATCACTGGGCGTAAAGGGCGTGTAGGCGGTCTATCAAGTCTGGTGTGAAATCCCCTCGCTCAACGAGGGAACTGCGTCGGAAACTGATAGACTAGAGTACCGGAGAGGAAGGTGGAATTCCCGGTGTAGCGGTGAAATGCGTAGATATCGGGAGGAACACCGGTGGCGAAGGCGGCCTTCTGGACGGATACTGACGCTGAGACGCGAAAGCGTGGGG |
| P-asv4667 | GCGAGCGTTGTCCGGATTTATTGGGCGTAAAGAGCTCGTAGGCGGTTTGGCAAGTTGGGTGTGAAAACTCCGGGCTCAACTCGGAGACGCCACTCAATACTGTCATGACTCGAGTCCGGTAGGGGAGTGCGGAATTCCTGGTGTAGCGGTGAAATGCGCAGATATCAGGAGGAACACCGGTGGCGAAGGCGGCACTCTGGGCCGGTACTGACGCTGAGGAGCGAAAGCGTGGGG |
| P-asv4668 | GCAAGCGTTATCCGGATTTATTGGGCGTAAAGGGAACGCAGGCGGTCTTTTAAGTCTGATGTGAAAGCCTTCGGCTTAACCGGAGTAGTGCATTGGAAACTGGGAGACTTGAGTGCAGAAGAGGAGAGTGGAACTCCATGTGTAGCGGTGAAATGCGTAGATATATGGAAGAACACCAGTGGCGAAAGCGGCTCTCTGGTCTGTAACTGACGCTGAGGTTCGAGAGCGTGGGT |
| P-asv4669 | GCTAGCGTTGTTCGGAATCACTGGGCGTAAAGGGCGCGTAGGCGGCCGTTCAAGTCGGGGGTGAAAGCCTGTGGCTCAACCACAGAATGGCCTTCGATACTGGACGGCTCGAGACCGGAAGAGGTCAGTGGAACTGCGAGTGTAGAGGTGAAATTCGTAGATATTCGCAAGAACACCAGTGGCGAAGGCGGCTGACTGGTCCGGTTCTGACGCTGAGGCGCGAAAGCGTGGGG |
| P-asv4670 | GCAAGCGTTGTCCGGATTTATTGGGTTTAAAGGGTGCGTAGGCGGTTGATTAAGTCAGTTGTGAAATACGGCAGCTTAACTGTCGAGGTGCAATTGATACTGGTTGACTTGAGTACAGACGAGGTAGGCGGAATTGACGGTGTAGCGGTGAAATGCATAGATATCGTCAAGAACACCGATAGCGAAGGCAGCTTACTAGACTGTAACTGACGCTGAGGCACGAAAGTGTGGGG |
| P-asv4671 | GCGAACGTTATTCGGAATCACTGGGCTTAAAGCGCGTGTAGGCGGGACGGCACGTCGGTCGCTGAAAGCCCCCGGCTCAACCGGGGAACGGCGCCGATACGACCGGCCTGGAGGGACGTAGGGGGACCTGGAACTTCCGGTGGAGCGGTGAAATGCGTTGAGATCGGAAGGAACGCCCGTGGCGAAAGCGAGGTCCTGGACGTTTTCTGACGCTGGGACGCGAAAGCCAGGGG |
| P-asv4672 | GCGAGCGTTGTTCGGAATTACTGGGCGTAAAGGGCGTGTAGGCGGTGCGATAAGTCGGGTGTGAAATCCCCGGGCTTAACCCGGGGCGGTCGCTCGAAACTGTCGTGCTTGAGGGTGGGAAAGGGAAGTGGAATTCCCGGTGTAGCGGTGAAATGCGTAGATATCGGGAGGAACACCTGCGGCGAAGGCGGCTTCCTGGACCATTACCGACGCTGAGGCGCGAAAGCTAGGGG |
| P-asv4673 | GCTAGCGTTGTTCGGAATTACTGGGCGTAAAGGGCGCGTAGGCGGCCTGGTCAGTCGGATGTGAAAGCCCGGGGCTCAACCCCGGAACGGCATCCGAGACGGCCAGGCTGGAGACCGGGAGAGGAGAGTGGAATAGCCAGTGTAGAGGTGAAATTCGTAGATATTGGCTGGAACACCGGTGGCGAAAGCGGCTCTCTGGCCCGGTTCTGACGCTGAGGCGCGAAAGCGTGGGG |
| P-asv4674 | GCAAGCGTTGTTCGGAATTACTGGGCGTAAAGCGCGTGTAGGCGGCCTCCTCAGTCTGGTGTGAAAGCCCGGGGCTCAACCCCGGAAGTGCATTGGATACTGGGAGGCTGGAGTACCGGAGAGGAAGGTGGAATTCCTGGTGTAGCGGTGAAATGCGTAGATATCAGGAGGAACACCGGTGGCGAAGGCAGCCCCCTGGGTCAAGATTGACGCTCATGCACGAAAGCGTGGGG |
| P-asv4675 | GCAAGCGTTGTTCGGAATCACTGGGCGTAAAGGGCGCGCAGGCGGTTCGTTAAGTTCGAGGTGAAATCTCCCGGCTCAACTGGGAGGGGTCTTCGGATACTGGCGGACTTGAGGAGAGCAGAGGTGAGCGGAATTCTGGGTGTAGTGGTGAAATGCGTAGATATCCAGAGGAACACCAGAGGCGAAGGCGGCTCACTGGGCTCCACCTGACGCTGAGGCGCGAAAGCGTGGGG |
| P-asv4676 | GCAAACGTTGTTCGGAATTACTGGGCGTAAAGAGTGTGTAGGCGGCTTGGCGTGTTTGGTGTGAAATCTCCCGGCTTAACTGGGAGGGTGCGCCAAAACTGCCAGACTAGAGTACTGGAGAGGAAAGCGGAATTCCTGGTGTAGCGGTGAAATGCGTAGATATCAGGAGGAACACCGGTGGTGTAGACGGCTTTCTGGACAGTCACTGACGCTGAGACACGAAAGCGTGGGG |
| P-asv4677 | GCGAGCGTTGTCCGGATTCATTGGGCGTAAAGCGCGCGCAGGCGGCTCAGAGAGCCCGGCGTGAAAGCCCTCGGCTCAACCGGGGGAGGTCGTCGGGGACCTGTGGGCTAGAGGCCGGTAGGGGCGAGTGGAATTCCCGGTGTAGTGGTGGAATGCGTAGAGATCGGGAGGAACACCAGTGGCGAAGGCGACTCGCTGGACCGGTCCTGACGCTCAGGCGCGAAAGCGTGGGG |
| P-asv4678 | GCGAGCGTTGTCCGGATTTATTGGGCGTAAAGAGCGCGCAGGCGGCCCGTCAAGTCACCACGTGAAATCCCCCGGCTCAACCGGGGAGGGTCGGGTGAAACTGGCGGGCTCGAGGATGGGAGAGGGGTGCGGAATTCCCGGTGTAGTGGTGAAATGCGTAGATATCGGGAGGAACACCAGTGGCGAAGGCGGCACCCTGGCCCATACCTGACGCTGAGGCGCGAAGGCCGGGGG |
| P-asv4679 | GCAAGCGTTGTTCGGAATTATTGGGCGTAAAGGGTGCGTAGGTGGTTCGGCAAGTCTTGTGTGAAATCTTCAGGCTCAACTTGAAGTCTGCACAAGAAACTGCCGGGCTTGAGTATGGGAGAGGTGAGTGGAATTTCCGGTGTAGCGGTGAAATGCGTAGATATCGGAAGGAACACCTGTGGCGAAAGCGGCTCACTGGACCATTACTGACGCTGAGGCGCGAAAGCTAGGGG |
| P-asv4680 | GCTAGCGTTGTTCGGAATTACTGGGCGTAGAGCGCGTGTAGGCGGGTGTTTAAGTCAGGGGTGAAATGCCAAGGCTCAACCTTGGAACTGCCTTTGATACTGGATACCTTGAGTCCGGGAGAGGTGAGTGGAACTGCGAGTGTAGAGGTGAAATTCGTAGATATTCGCAAGAACACCAGTGGCGAAGGCGGCTCACTGGCCCGGAACTGACGCTGAGACGCGAAAGCGTGGGG |
| P-asv4681 | GCAAGCGTTGTTCGGATTTACTGGGTGTAAAGGGCGCGCAGGCGGCTAGTTTTGTCGAGTGTGAAAGCCCCGGGCTTAACCTGGGAACTGCGCTCGAAACGGGCTGGCTAGAGATCGGGAGAGGAGAGTGGAATTCCCAGTGTAGAGGTGAAATTCGTAGATATTGGGAAGAACACCGGTGGCGAAGGCGGCTCTCTGGACCGATACTGACGCTGAGGCGCGAAAGCGTGGGG |
| P-asv4682 | GCTAGCGTTGTTCGGAATTACTGGGCGTAAAGGGCGCGTAGGCGGCCTCCTAAGTCGGACGTGAAATCCCCGAGCTCAACTCGGGAACTGCGTCCGATACTGGTTGGCTTGAATCCGGGAGAGGGATGCGGAATTCCAGGTGTAGCGGTGAAATGCGTAGATATCTGGAGGAACACCGGTGGCGAAGGCGGCATCCTGGACCGGCATTGACGCTGAGGCGCGAAAGCCAGGGG |
| P-asv4683 | GCCAGCGTTGTTCGGAATTACTGGGTGTAAAGGGTTCGTAGGCGGTGTGGCAAGTCGGGAGTGAAAGCCTGGGGCTCACCCCCAGAAGTGCATCCGAAACTAGCCAGCTAGAGCGCGATAGAGGAGGGTGGAATTCCTGGTGTAGCGGTGAAATGCGTAGATATCAGGAGGAACACCGGCGGCGAAGGCGGCCCTCTGGATCGTTGCTGACGCTGAGACGCGAAAGCGTGGGG |
| P-asv4684 | GCGAGCGTTATCCGGATTTACTGGGTGTAAGGGGAGCGCAGGCGGCGTGGCAAGTCAGATGTGAAAACCCGGGGCCCAACCCCGGGACTGCATTTGAAACTGCCATGCTGGAGTGCCGGAGAGGTAAGTGGAATTCCTAGTGTAGCGGTGAAATGCGTAGATATTAGGAGGAACACCAGTGGCGAAGGCGGCTTACTGGACGGTAACTGACGCTGAGGCTCGAAAGCGTGGGG |

| | |
|---|---|
| P-asv4685 | ACAAGCGTTGTCCGGATTTACTGGGCGTAAAGAGCGCGCAGGCGGTCGTTCAAGTCGAGTGTGAAAGCCCCCGGCTCAACTGGGGAGGGTCACTCGATACTGATCGACTCGAAGGCAGGAGAGGGAAGTGGAATTCCCGGTGTAGTGGTGAAATGCGTAGATATCGGGAGGAACACCAGTGGCGAAGGCGACTTCCTGGCCTGTTCTTGACGCTGAGGCGCGAAAGCTAGGGG |
| P-asv4686 | GCGAGCGTTGTTCGGATTTACTGGGCGTAAAGGGCGCGTAGGCGGTGCCGAAGGTCAGGCGTGAAAGCCCCGGGCTCAACCTGGGAGGTGCGCTTGGGACTGCGGCACTGGAGTTCGGGAGAGGAGCGTGGAATTCCCAGTGTAGAGGTGAAATTCGTAGATATTGGGAAGAACACCGGTGGCGAAGGCGGCGCTCTGGACCGAGACTGACGCTGAGGCGCGAAAGCGTGGGG |
| P-asv4687 | GCAAGCGTTGTTCGGATTTACTGGGCGTAAAGGGCGCGTAGGCGGTCAGCACAAGTCAATTGTGAAATCTCCGAGCTTAACTCGGAAAGGTCAACTGAAACTGTGCGACTAGAGTGCGGAAGGGGCAACTGGAATTCTTGGTGTAGCGGTGAAATGCGTAGATATCAAGAGGAACACCTGCGGCGAAGGCGGGTTGCTGGGCCGACACTGACGCTGAAGCGCGAAAGCCAGGGG |
| P-asv4688 | GCGAGCGTTAATCGGAATTACTGGGCGTAAAGCGCACGTAGGTGGCTTTGCAAGTCGATTGTGAAATCCCCGGGCTTAACCTGGGAACTGCAGTCGAGACTGCTTTGCTTGAGTATGGGAGAGGGAGGTGGAATTTCCGGTGTAGCGGTGAAATGCGTAGATATCGGAAGGAACATCAGTGGCGAAGGCGACCTCCTGGACCAATACTGACACTCAAGTGCGAAAGCGTGGGG |
| P-asv4689 | GCTAGCGTTGTTCGGAATGACTGGGTGTAAAGGGCGCGTAGGCGGTATTTTAAGTGAGGCGTGAAAGCCCTGGGCTTAACCCAGGAGGTGCGTTTCATACTGAAATGCTAGAGTGTGAGAGAGGAAAGTGGAATTCCTAGTGTAGAGGTGAAATTCGTAGATATTAGGAAGAACACCAGAGGCGAAGGCGGCTTTCTGGCTCGCAACTGACGCTGAGGCGCGAAAGCGTGGGG |
| P-asv4690 | GCAAGCGTTAATCGGAATTACTGGGCGTAAAGCGTGCGCAGGCGGTCTTGCAAGACAGATGTGAAATCCCCGGGCTTAACCTGGGAACTGCGTTTGTGACTGCAAGGCTGGAGAGCGGCAGAGGGGGATGGAATTCCGCGTGTAGCAGTGAAATGCGTAGACATGCGGAGGAACACCGATGGCGAAGGCAATCCCCTGGGCCTGCACTGACGCTCATGCACGAAAGCGTGGGG |
| P-asv4691 | CCAAGCGTTATCCGGAATCATTGGGTTTAAAGGGTCCGTAGGCGGGCTATTAAGTCAGAGGTGAAAGTTTGCAGCTCAACTGTAAAATTGCCTTTGAAACTGGTAGTCTTGAATTATTATGAAGTGGTTAGAATAAGTAGTGTAGCGGTGAAATGCATAGATATTACTTAGAATACCAATTGCGAAGGCAGATCACTAATAATATATTGACGCTGAGGGACGAAAGCGTGGGG |
| P-asv4692 | GCGAGCGTTATCCGGATTCACTGGGTTTAAAGGGTGCGTAGGCGGGTATTTAAGTCAGTGGTGAAATCCTAGAGCTTAACTCTAGAACTGCCATTGATACTATTTATCTTGAATATTGTGGAGGTAAGCGGAATATGTCATGTAGCGGTGAAATGCTTAGATATGACATAGAACACCCATTGCGAAGGCAGCTTACTACGCATATATTGACGCTGAGGCACGAAAGCGTGGGG |
| P-asv4693 | ACTAGCGTTGTTCGGAATCACTGGGCGTAAAGCGCACGTAGGCGGATTTGTAAGTCAGGGGTGAAATCCCGGGGCTCAACCTCGGAACTGCCTTTGATACTGCGAGTCTTGAGTCCGATAGAGGTGGGTGGAATTCCTAGTGTAGAGGTGAAATTCGTAGATATTAGGAAGAACACCGGTGGCGAAGGCGGCCCACTGGATCGGTACTGACGCTGAGGTGCGAAAGCGTGGGG |
| P-asv4694 | GCAAGCGTTATCCGGAATTATTGGGCGTAAAGGGAGCGCAGGCGGTTTATCAAGTTTATGGTTAAAGTTCGGGGCTTAACCCCGTGATGCCATAGAAACTGGTAGACTAGAGTGCAGGAGAGGTTAGTGGAATTCCATGTGTAGCGGTAAAATGCGTAGATATATGGAGGAACACCAGTGGCGAAGGCGGCTAACTGGCCTGTAACTGACGCTGAGGCTCGAAAGCGTGGGG |
| P-asv4695 | GCTAGCGTTGCTCGGAATCACTGGGCGTAAAGGGCGCGTAGGCGGCAATTCAAGTCGGGGGTGAAAGCCTGTGGCTCAACCACAGAATTGCCTTCGATACTGTTTGGCTTGAGTATGGCAGAGGTCAGTGGAACTGCGAGTGTAGAGGTGAAATTCGTAGATATTCGCAAGAACACCAGTGGCGAAGGCGGCTGACTGGGCCATTACTGACGCTGAGGCGCGAAAGCGTGGGG |
| P-asv4696 | GCAAGCGTTGTCCGGATTTATTGGGTTTAAAGGGTGCGTAGGTGGCCCGTTAAGTCCGGGGTGAAAGCCCACAGCTCAACTGTGGAACTGCCCTGGATACTGGCGGGCTTGAGTCCAGACGAGGTTGGCGGAATGGAAGCTGTAGCGGTGAAATGCATAGATAGCTTCCAGAACCCCGATTGCGTAGGCAGCTGACTAGGCTGGTACTGACACTGAGGCACGAAAGCGTGGGG |
| P-asv4697 | GCAAGCGTTAATCGGAATTACTAGGCGTAAAGCGCGCGTAGGTGGTTCGTTAAGTTGGATGTGAAATCCCCGGGCTCAACCTGAGAACTGCATTCAAAACTGACGAGCTAGAGTATGGTAGAGGGTGGTGGAATTTCCTGTGTAGCGGTGAAATGCGTAGATATAGGAAGGAACACCAGTGGCGAAGGCGACCACCTGGACTGATACTGACACTGAGGTGCGAAAGCGTGGGG |
| P-asv4698 | GCAAGCGTTAATCGGAATTACTGGGCGTAAAGCGTGCGCAGGCGGTTTTGTAAGTCTGATATGAAATCCCCGGGCTCAACCTGGGAATTGCATTAAAGACTGCAAGGCTAGAATCTGGCAGAGGGGGGTAGAATTCCACGTGTAGCAGTGAAATGCGTAGATATGTGGAGGAACACCGATGGCGAAGGCAGCCCCCTGGGTCAAGATTGACGCTCATGCACGAAAGCGTGGGG |
| P-asv4699 | GCAAGCGTTGTCCAGAATTATTGGGCGTAAAGCACGCACAGGCAGTCCTTTAAGTCTGATATGAAAGCCCACAGCTCAACCGTAGAGGGTCATTGGAAACTGGGGGACTTGAGTGCAGAAGAGAAGAGTGGAATTCCACGTGTAGCGGTGAAATGCGTAGAGATGTGGAGGAACACCAGTGGCGAAGGCGACTCTTTGGTCTGTAACTGACGCTGAGGCGCGAAAGCGTGGGG |
| P-asv4700 | GCAAGCGTTATCCGGAATTATTGGGCGTAAAGGGTGCGTAGGCGGTCATTTAAGTCTTTGGTGTAAGTGCAAGGCTCAACCTTGTGATGCTAAGGAAACTGGATGACTTGAGTGTGATAGAGGTAAGTGGAATTCCATGTGTAGCGGTAAAATGCGTAAATATATGGAGGAACACCAGTGGCGAAGGCGGCTTACTGGGTCACAACTGACGCTGAGGCACGAAAGCGTGGGG |
| P-asv4701 | GCTAGCGTTGTTCGGAATTACTGGGCGTAAAGCGTACGCAGGCGGTTGATCAAGTCAGGTGTGAAAGCCCGGGGCTCAACCTCGGAATTGCGCCTGAGACTGGTCGGCTAGAGTTCGGAAGAGGAGAGTGGAATTCCCAGTGTAGAGGTGAAATTCGTAGATATTGGGAAGAACACCAGTGGCGAAGGCGACTCTCTGGTCCGATACTGACGCTCAAGTACGAAAGCGTGGGG |
| P-asv4702 | GCAAGCGTTGTCCGGAATCATTGGGCGTAAAGCGCGTGTAGGCGGCTCGGTAAGTCCGCTCTGAAAGCCCGGGGCTCAACCCCGGGAGGCGGGTGGATACTGTCGAGCTCGAGTCCGGAAGAGGCAAGTGGAATTCCCGGTGTAGCGGTGAAATGCGCAGATATCGGGAGGAACACCAATGGCGAAGGCAGCTTGCTGGGACGGTACTGACGCTGAGACGCGAAAGCGTGGGG |
| P-asv4703 | GCAAGCGTTGTCCGGAATTATTGGGCGTAAAGCGCGCGCAGGCGGCTTACTAAGTCTGATGTGAAAGCCCACGGCTCAACCGTGGAGGGCCATTGGAAACTGGTAGGCTTGAGTGCAGGAGAGGAGAGCGGAATTCCCGGTGTAGCGGTGAAATGCGTAGATATCGGGAGGAACACCCGTGGCGAAGGCGGCTCTCTGGCCTGTAACTGACGCTGAGGCGCGAAAGCGTGGGG |
| P-asv4704 | GCTAGCGTTGTTCGGAATTACTGGGCGTAAAGCGCATGTAGGCGGATATTTAAGTCAGGGGTGAAATCCCGCAGCTCAACTGCGGAACTGCCTTTGATACTGGGTATCTTGAGTATGGAAGAGGTGAGTGAAATTCCGAGTGTAGAGGTGAAATTCGTAGATATTCGGAGGAACACCAGTGGCGAAGGCGGCTCACTGGTCCATTACTGACGCTGAGGTGCGAAAGCGTGGGG |
| P-asv4705 | GCAAGCGTTGTCCGGATTTACTGGGCGTAAAGGATGCGTAGGCGGATATTTAAGTCAGATGTGAAAACCCCGAGCTTAACTTGGGGACTGCATTTGAAACTGGATATCTAGAGTGTGGGAGAGGAGAATGGAATTCCTAGTGTAGCGGTGAAATGCGTAGAGATTAGGAAGAACACCAGTGGCGAAGGCGATTCTCTGGACCATAACTGACGCTGAGGCATGAAAGCGTGGGT |
| P-asv4706 | GCGAGCGTTATCCGGATTTATTGGGTTTAAAGGGTGCGTAGGTGGGCTATTAAGTCAGTGGTGAAAGTTTGTCGCTCAACGATAAAATTGCCGTTGAAACTGGTGGTCTTGAGTATGGATGAAGTAGGCGGAATGCGTGGTGTAGCGGTGAAATGCATAGAGATCACGCAGAACTCCGATTGCGAAGGCAGCTTACTAAGGCATAACTGACACTGAAGCACGAAAGCGTGGGT |
| P-asv4707 | GCAAGTGTTATCCGGAATTATTGGGCGTAAAGCGTCTGCAGGTGGATTTTTAAGTCTTTTGTTAAATCTTTGAGCTCAACTCGAAATCTGCAAAAGAAACTATTTATCTAGAGTATGGTAGAGGTAAAGGGAATTTTCAGTGGAGCGGTGAAATGCGTAGATATTGGAAAGAACACCAAAGGCGAAGGCACTTTACTGGGCTATTACTGACACTCAGAGACGAAAGCTAGGGG |
| P-asv4708 | GCAAGCGTTGTCCGGATTTACTGGGCGTAAAGGGAGCGTAGGTGGATATTTAAGTGGGATGTGAAATACTCGGGCTTAACCTGGGTGCTGCATTCCAAACTGAATATCTAGAGTGCAGGAGAGGAAAGTAGAATTCCTAGTGTAGCGGTGAAATGCGTAGAGATTAGGAAGAATACCAGTGGCGAAGGCGACTTTCTGGACTGTAACTGACACTGAGGCTCGAAAGCGTGGGG |
| P-asv4709 | GCTAGCGTTGTTCGGAATTACTGGGCGTAAAGCGCGTGTAGGCGGTTATTCAAGTCAGGTGTGAAAGCCTTGAGCTCAACTCAAGAAATGCACTTGATACTGGATGACTAGAGGACCGGAGAGGATAGTGGAATTCCCAGTGTAGTGGTGAAATACGTAGAGATTGGGAAGAACACCAGTGGCGAAGGCGGCTATCTGGACGGTTTCTGACGCTAAGACGCGAAAGCGTGGGG |
| P-asv4710 | GCGAGCGTTGTTCGGAATTACTGGGCGTAAAGCGCGCGTAGGCGGTCGTTTAAGTCTCACTGTGAAAGACCGGGGCTCAACCCCGGGGCGGCAGGGGATACTGGACGACTGGAGACAGGTAGAGGCGAGCGGAATTCCGGGTGTAGCGGTGGAATGCGTAGAGATCCGGAAGAACACCAGAGGCGAAGGCGGCTAGCTGGGCCTGGTCTGACGCTGAGGCGCGAAAGCGTGGGG |
| P-asv4711 | GCGAGCGTTGTCCGGAATCATTGGGCGTAAAGAGCTCGTAGGCGGCTTGGTAAGTCGGATGTGAAACCTCCAGGCTCAACCTGGAGTCGCCATCCGATACTGCCATGGCTAGAGTCCGGTAGGGGCCCACGGAACTCCTGGTGTAGCGGTGAAATGCGCAGATATCAGGAAGAACACCGGTGGCGAAGGCGGTGGGCTGGGCCGGCACTGACGCTGAGGAGCGAAAGCGTGGGG |
| P-asv4712 | ACGAGTGATGGCCGATTTTATTGGGCCTAAAGCGTCCGTAGCTGGCCGACCAAGTCCGTCGGGAAATCCACCAGCCCAACTGGTGGGCGTCCGGCGGAAACTGCTCGGCTTGGAGCCGGAAGACCCAAGGGGTACGTCCGGGGTAGGAGTGAAATCCCGTAATCCTGGACGGACCACCGATGGGGAAACCACTTCAAGAGAACGGATCCGACAGTGAGGGACGAAAGCTAGGGT |
| P-asv4713 | GCGAGCGTTGTTCGGAATCACTGGGCGTAAAGCGTGCGCAGGCGGTTATGAAAGTCAGATGTGAAAGCCCAGGGCTTAACCCTGGAATTGCATTTGAAACTCCATGACTAGAGTACTGGAGGGGTTGGCGGAATTTCGAGTGTAGCAGTGAAATGCGTAGATATTCGAAGGAACACCGATGGCGTAGGCAGCCAACTGGACAGTAACTGACGCTCATGCACGAAAGCGTGGGG |
| P-asv4714 | CCAAGCGTTACCCAGATTCACTGGGCGTAAAGGGTGCGTAGGCGGTCATATTAGTCGGGTGTTAAATCCTGGGGCTCAACCTCAGGCTCGCATCCGAAACGGTATGACTAGAGGAAGTGAGAGGTAAGCAGAACTCATGGTGTAGGGGTGAAATCCGTTGATATCATGGGGAATACCAAATGCGAAGGCAGCTTACTGGCACTTTCCTGACGCTGAGGCACGAAAGCATGGGT |
| P-asv4715 | GCAAGCGTTGTCCGGATTTATTGGGCGTAAAGCGAGCGCAGGCGGTCAATTAAGTCTGATGTGAAAGCCCCCGGCTCAACCGGGGAGGGTCATTGGAAACTGGTTGACTTGAGTGCAGAAGAGGAGAGTGGAATTCCATGTGTAGCGGTGAAATGCGTAGATATATGGAGGAACACCAGTGGCGAAGGCGACTCTCTGGTCTGTAACTGACGCTGAGGCTCGAAAGCGTGGGT |
| P-asv4716 | GCGAGCGTTGTTCGGAATTACTGGGCGTAAAGCGTGCGCAGGCGGATAATCAAGTCAGAAGTGAAAGCCCAGGGCTCAACCTTGGAATTGCTTTTGAAACTGGTTGTCTGGAGTCTCGGAGAGGTTGGCGGAATTCCTAGTGTAGAGGTGAAATTCGTAAATATTAGGAGGAACACCAGTGGCGTAAGCGGCCAACTGGACGAGTACTGACGCTCATGCACGAAAGCGTGGGG |
| P-asv4717 | GCAAGCGTTGTTCGGATTTATTGGGCGTAAAGCGCGCGCAGGCGGATGTGCAAGTCAGATGTGAAATCTCGGGGCTCAACCCCGAAACTGCGTCTGAAACTGTTCATCTAGAATCATATAGGGGGTAGGGGAATTTCACATGTAGGGGTAAAATCCGTAGAGATGTGAAGGAACACCGGAGGCGAAGGCGCCTACCTGGATATGTATTGACGCTGAGGCGCGAAAGCGTGGGG |
| P-asv4718 | GTTAGCGTTGTTCAGAATTACTGGGCGTAAAGCGCACGTAGGCGGATCAGAAAGTTGGGGGTGAAATCCCAGGGCTCAACCCCGGAACGGCCCTCGATACTGGCAGCCTAGAGTTCGAGAGAGGTGAGTGGAATTCCGAGTGTAGAGGTGAAATTCGTAGATATTCGGAGGAACACCAGTGGCGAAGGCGGCTCACTGGCTCGATACTGACGCTGAGGTGCGAAAGCGTGGGG |
| P-asv4719 | GCGAGCGTTGTCCGGATTTATTGGGCGTAAAGAGCTCGTAGGCGGTTCGGTAAGTCGGGTGTGAAAATCCAGGGCTCAACCCTGGGACGCCACCTGATACTGCTGTGACTAGAGTCCGGTAGAGGAGCGTGGAATTCCTGGTGTAGCGGTGAAATGCGCAGATATCAGGAGGAACACCAGCGGCGAAGGCGGCGCTCTGGGCCGGTACTGACGCTGAGGAGCGAAAGCGTGGGG |
| P-asv4720 | GCTAGCGTTGTTCGGATTTACTGGGCGTAAAGCGCACGTAGGCGGATTGTTAAGTCAGAGGTGAAATCCCGGAGCTCAACTCCGGAACTGCCTTTGATACTGGCAATCTAGAGTCCGGAAGAGGTGAGTGGAACTCCTAGTGTAGAGGTGGAATTCGTAGATATTAGGAAGAACACCAGTGGCGAAGGCGGCTCACTGGTCCGGTACTGACGCTGAGGTGCGAAAGCGTGGGG |
| P-asv4721 | GCTAGCGTTGTTCGGATTTACTGGGCGTAAAGCGCACGTAGGCGGCGGATCAAGTGAGGCGTGAAAGGCCTGGGCTCAACCTGGGAACGGCGCTTCAGACTGATTGGCTCGAGTTCGGGAGAGGGTGGTGGAATTTCCAGTGTAGAGGTGAAATTCGTAGATATTGGAAGGAACACCGGTGGCGAAGGCGGCCACCTGGCCCGAGACTGACGCTCAAGTGCGAAAGCGTGGGG |
| P-asv4722 | GCAAGCGTTAATCGGATTTACTGGGCGTAAAGCGTGCGTAGGCGGTTCGTTAAGTCGATTGTGAAAGCCCTGGGCTTAACCTGGGAATTGCAGTCGATACTGGCGGACTAGAGAACGGTAGAGGGAGGCGGAACTCCAGGTGTAGCGGTGAAATGCGTAGATATTCGGAAGAACGCCAGTGGCGAAGGCGACTCACTGGCCCGTTACTGACGCTGAGGCGCGAAAGCGTGGGG |
| P-asv4723 | GCAAGCGTTACTCGGAATTACTGGGCGTAAAGCGTGCGTAGGCGGTTTGTTAAGTCAGATGTGAAAGCCCCGGGCTCAACCTGGGAATTGCATTTGATACTGGCAAGCTAGAGTGTGGTAGAGGGTGGTGGAATTCCCGGTGTAGCGGTGAAATGCGTAGAGATCGGGAGGAACATCCGTGGCGAAGGCGGCCACCTGGACCAACACTGACGCTGAGGCACGAAAGCGTGGGG |

| | |
|---|---|
| P-asv4724 | CCAAGCGTTATCCGGATTTATTGGGTTTAAAGGGTTCGTAGGCGGGATATTAAGTCAGTGGTGAAAGCCCACAGCTCAACTGTGGAACTGCCATTGATACTGATATTCTTGAGTTTATTTGGAGTTAGCGGAATATGTCATGTAGCGGTGAAATGCTTAGATATGACATAGAACACCAATCGCGAAAGCAGCTAACTAAGATAATACTGACGCTGAGGAACGAAAGCGTGGGG |
| P-asv4725 | GCAAGCGTTATCCGGAATCACTGGGCGTAAAGGGCGCGTAGGCGGTCTGTTAAGTCCGATGTTAAAGACCGGGGCTCAACCCCGACACGGCGTTGGATACTGGCAGGCTTGACGGCTGGAGAGGAAAGTGGAATTACCAGTGTAGCGGTGAAATGCGTAGATACTGGTAGGAACACCCATTGCGAAGGCAGCTTTCTGGACAGTTTGTGACGCTGAGGCGCGAAAGTGTGGGG |
| P-asv4726 | GTTAGCGTTGTTCGGAATTACTGGGCGTAAAGCGCACGTAGGCGGACTATTAAGTCAGGGGTGAAATCCCGGGGCTCAACCCCGGAACTGCCTTTGATACTGGTAGTCTAGAGTTCGAGAGAGGTGAATGGAATTCCGAGTGTAGAGGTGAAATTCGTAGATATTCGGAAGAACACCAGTGGCGAAGGCGGCTCACTGGCTCGATACTGACGCTGAGGTGCGAAAGTGTGGGT |
| P-asv4727 | GCGAGCGTTGTCCGGATTCATTGGGCGTAAAGAGTTCGTAGGCGGTTTGTTAAGTCTGATGTTAAAGATCGGGGCTCAACCCTGGGAGTGCATTGGATACTGGCAGACTGGAGTGCAGTAGAGGCGAGTGGAATTCCCAGTGTAGCGGTGAAATGCGTAGATATTGGGAAGAACACCGGTGGCGCAGGCGACTCGCTGGGCTGTAACTGACGCTGAGGAACGAAAGCTAGGGG |
| P-asv4728 | GCAAGCGTTATTCGGAATTATTGGGCGTAAAGAGTGCGTAGACGGCTTATCAAGTCAACTGTTAAATTCTCCGGCCTAACTGGAGGTGAGCGGTAGAAACTGATAAGATTGAGGTTGGAAGAGAGAAGTGGAATTCTCGGAGTAGCGGTAAAATGCGTAGATCTCGAGAGGAACACCGATGGCGAAGGCAGCTTCTTGGTCCATACCTGACGTTGAGGCACGAAAGCGTGGGT |
| P-asv4729 | GCTAGCGTTGTTCGGAATTACTGGGCGTAAAGCGCACGTAGGCGGACCGCTAAGTCAGGGGTGAAATCCCGGGGCTCAACCTCGGAATTGCCTTTGATACTGGTGGTCTAGAGTCCGAGAGAGGTGAGTGGAACTCCGAGTGTAGAGGTGAAATTCGTAGATATTCGGAAGAACACCAGTGGCGAAGGCGGCTCACTGGCTCGGCACTGACGCTGAGGTGCGAAAGCGTGGGG |
| P-asv4730 | GCAAGCGTTGTTCGGAATTACTGGGCGTAAAGCGTTTGTAGGCGGTGCTGAGAGTCCTAGGTGAAATCCCCGAGCTCAACTCGGGAACTGCCTGGGAGACCTCAGCACTAGAGTACTGAAGGGGTTGCTGGAATTCCATGTGTAGCGGTGAAATGCGTAGATATATGGAGGAACACCTGTGGCGAAGGCGGGCAACTGGTCAGAGACTGACGCTGAGAAACGAAAGCGTGGGG |
| P-asv4731 | ACGAGTGATGGCCAATCTTATTGGGCCTAAAGCGTCCGTAGCCGGCCAGACAAGTCCGTTGGGAAATCGACGCGCCTAACGCGTCGGCGTCCAGCGGAAACTGTCTGGCTTGGGGCCGGAAGACCCGAGGGGTACGTCCGGGGTAGGAGTGAAATCCCGTAATCCTGGACGGACCACCGGTGGCGAAAGCGCCTCGGGAAGACGGACCCGACGGTGAGGGACGAAAGCTAGGGT |
| P-asv4732 | CCGAGTGATGGCCGATATTATTGGGCCTAAAGCGTCCGTAGCTGGCCGCGGAGGTCCGTCGGGAAATCTACTGGCCTAACCAGTAGGCGTCCGGCGGAAACCCCGTGGCTTGGGACCGGGAGACCTGAGGGGTACGTCCGGGGTAGGAGTGAAATCCCGTAATCCTGGACGGACCACCGGTGGCGAAAGCGCCTCAGGAGAACGGATCCGACAGTGAGGGACGAAAGCTAGGGT |
| P-asv4733 | GCAAGCGTTGTCCGGAATTATTGGGCGTAAAGAGCTCGTAGGCGGTTTGTCGCGTCTGCTGTGAAAACCTGAGGCTCAACCTCGGGCTTGCAGTGGGTACGGGCAGACTAGAGTGCGGTAGGGGTGACTGGAATTCCTGGTGTAGCGGTGGAATGCGCAGATATCAGGAGGAACACCGATGGCGAAGGCAGGTCACTGGGCCGCAACTGACGCTGAGGAGCGAAAGCATGGGG |
| P-asv4734 | CCAAGCGTTGTCCGGATTTATTGGGTTTAAAGGGTGCGTAGGCGGATTATTAAGTCAGAGGTGAAAGCCGGTAGCTCAACTATCGAATAGCCTTTGAAACTGTTAGTCTTGAGTATTGTTGAGGTAGGCGGAATGTATCATGTAGCGGTGAAATGCATAGATATGATACAGAACACCGATTGCGAAGGCAGCTTACTAAGCAATAACTGACGCTGAGGCACGAAAGCGTGGGG |
| P-asv4735 | GCTAGCGTTATTCGGATTTACTGGGCGTAAAGCGTGCGTAGGCGGCTTTTTAAGTCGGGTGTGAAATCCCCGGGCTCAACCTGGGCACTGCAGTTAGAACTGGCAAGCTAGAGTAGGGTAGAGGGGTGTGGAATTCCAGGTGTAGCGGTGAAATGCGTAGATATCTGGAGGAACATCAGTGGCGAAGGCGACACCCTGGACTCATACTGACACTGAGGTGCGAAAGCGTGGGG |
| P-asv4736 | GCGAGCGTTATCCGGATTCATTGGGTTTAAAGGGTACGTAGGCGGCCCGATAAGTCAGTGGTGAAATCCTGCAGCTTAACTGTAGAACTGCCATTGATACTGTTGGACTTGAATGTAGTTGAGGTAGGCGGAATGTGTTGTGTAGCGGTGAAATGCTTAGATATAACACAGAACACCGATTGCGAAGGCAGCTTACTAGACTATGATTGACGCTGAGGTACGAAAGCGTGGGT |
| P-asv4737 | GCGAGCGTTGTCCGGAATTATTGGGCGTAAAGGGCTCGTAGGCGGTCTGTTGCGTCAGGAGTGAAAACTCGGGGCTTAACCCCGAGCCTGCTTCTGATACGGGCAGACTAGAGGTATGCAGGGGGGAACGGAATTCCTGGTGTAGCGGTGAAATGCGCAGATATCAGGAGGAACACCGGTGGCGAAGGCGGTTCTCTGGGCATTACCTGACGCTGAGGAGCGAAAGTGTGGGG |
| P-asv4738 | GCAAGCGTTGTTCGGATTTACTGGGCGTAAAGCGTGCGTAGGCGGTATTGAGAGTCGCGGGTGAAATCCCAGGGCTTAACCCTGGAACTGCCTGCGAGACCTCAGTACTAGAGTATGAGAGGGGATAGTGGAATTCCTGGTGTAGCGGTGAAATGCGTAGAGATCAGGAGGAACACCGGTGGCGAAGGCGGCTATCTGGCTCATAACTGACGCTCAGGCACGAAAGCGTGGGT |
| P-asv4739 | GCAAGCGTTATCCGGATTTACTGGGTGTAAAGGGAGCGTAGGTGGCATGGCAAGTCCGATGTGAAAGCCCGGGGCTCAACCCCGGGATTGCATTAGAAACTGTCAGGCTAGAGTGCAGGAGAGGTAAGTGGAATTCCTAGTGTAGCGGTGAAATGCGTAGATATTAGGAGGAACACCAGTGGCGAAGGCGGCTTACTGGACTGTAACTGACACTGAGGCTCGAAAGCGTGGGG |
| P-asv4740 | GCAAGCGTTGTCCGGAATCATTGGGCGTAAAGAGCGTGTAGGCGGCCCGGTAAGTCCGCTGTGAAAGTCGAGGGCTCAACCCTCGGATGCCGGCGGATACTGTCGGGCTAGAGTCCGGAAGAGGCGAGTGGAATTCCTGGTGTAGCGGTGAAATGCGCAGATATCAGGAGGAACACCTACGGCGAAGGCAGCTCGCTGGGACGGTACTGACGCTGAGACGCGAAAGCGTGGGG |
| P-asv4741 | GCAAACGTTATTCGGAATCACTGGGCTTAAAGGGTGCGTAGGCGGCCATGAGGGTGGGGTGTGAAATCCCTCGGCTTAACCGAGGAATCGCGCTCCAAACCGCATGGCTGGAGTGAGACAGAGGTGAGCGGAACTGATGGTGGAGCGGTGAAATGCGTTGATATCATCAGGAACACCAGAGGCGAAGGCGGCTCACTGGGTCTCAACTGACGCTGATGCACGAAAGCTAGGGT |
| P-asv4742 | GCAAGCGTTAATCGGAATTACTTGGCGTAAAGCGTGCGTAGTCGGCTTTTCAAGTCGGGTGTGAAAGCCCCGGGCTCAACCTGGGAATTGCATTCGATACTGGATCGCTAGAGTTCGGCAGAGGGAAGTGGAATTTCTGGTGTAGCGGTGAAATGCGTAGATATCAGAAGGAACATCAGTGGCGAAAGCGACTTCCTGGACCAGAACTGACGATCAGGCACGAAAGCGTGGGG |
| P-asv4743 | GCAAACGTTGCTCGGAATCACTGGGCGTAAAGGGCGTGTAGGCGGGCAGGAAAGTCAGGTGTGAAATCCCCGAGCTCAACTCGGGAACTGCATTTGAAACTCCCTGCCTTGAGTACCGGAGAGGAAGGTGGAATTCCTGGTGTAGAGGTGAAATTCGTAGATATCAGGAGGAACACCAGTGGCGAAGGCGGCCTTCTGGACGGTAACTGACGCTGAGACGCGAAAGCGTGGGG |
| P-asv4744 | GCGAGCGTTGTCCGGAATTATTGGGCGTAAAGGGCTCGTAGGCGGTCTGTCGCGTCAGGAGTGAAAACTCGGGGCTCAACCCCGAGCCTGCTTCTGATACGGGCAGACTAGAGGTATGCAGGGGAGAACGGAATTCCTGGTGTAGCGGTGAAATGCGCAGATATCAGGAGGAACACCGGTGGCGAAGGCGGTTCTCTGGGCATTACCTGACGCTGAGGAGCGAAAGTGTGGGG |
| P-asv4745 | GCAAGCGTTACTCGGAATTACTGGGCGTAAAGCGTGCGTAGGTGGTTGTTTAAGCCTGTTGTGAAAGCCCTGGGCTCAACCTGGGAATTGCAGTGGATACTGGGCGACTAGAGTGTGGTAGAGGGTAGTGGAATTCCCGGTGTAGCAGTGAAATGCGTAGAGATCGGGAGGAACATCCATAGCGAAGGCAGCTACCTGGACCAACACTGACACTGAGGCACGAAAGCGTGGGG |
| P-asv4746 | GCAAGCGTTGTCCGGAATTATTGGGCGTAAAGAGCTCGTAGGCGGTTTGTCGCGTCTGCTGTGTAAACGCAAGGCTTAACCTTGCGCCTGCAGTGGGTACGGGCAGACTAGAGTGCGGTAGGGGAGACTGGAATTCCTGGTGTAGCGGTGGAATGCGCAGATATCAGGAAGAAAACCGATGGCGAAGGCAGGTCTCTGGGCCGTTACTGACGCTGAGGAGCGAAAGCATGGTG |
| P-asv4747 | GCTAGCGTTGCTCGGAATTACTGGGCGTAAAGGGAGCGTAGGCGGATAGTTTAGTCAGGGGTGAAAGCCCAGGGCTCAACCTTGGAATTGCCTTTGATACTGGCTATCTTGAGTATGGAAGAGGTATGTGGAACTCCGAGTGTAGAGGTGAAATTCGTAGATATTCGGAAGAACACCAGTGGCGAAGGCGACATACTGGTCCATTACTGACGCTGAGGCTCGAAAGCGTGGGG |
| P-asv4748 | GCAAGCGTTACTCGGAATCACTGGGCGTAAAGGACGCGTAGGCGGGTTGTCAAGTCAGGTGTGAAATCCTACAGCTTAACTGTAGAACTGCACTTGAAACTGGCAACCTAGAGTATGGGAGGGGGAGATGGAATTAGTGGTGTAGGGGTAAAATCCGTAGATATCACTAGGAATACCTAAAGCGAAGGCGATCTCCTGGAACATTACTGACGCTAAGGCGTGAAAGCGTGGGG |
| P-asv4749 | ACAAGCGTTGTCCGGATTTATTGGGCGTAAAGCGAGCGCAGGCGGTCTTTTAAGTCTGATGTGAAAGCCCACGGCTCAACCGTGGATGGTCTTTGGAGACTGGAAGACTTGAATGCAGAAGAGGAAAGTGGAATTCCATGTGTAGCGGTGAAATGCGTAGATATATGGAGGAACACCAGTGGCGAAGGCGACTTTCTGGTCTGTAATTGACGCTGAGGCTCGAAAGCGTGGGG |
| P-asv4750 | GCTAGCGTTGCTCGGAATTACTGGGCGTAAAGGGCGCGTAGGCGGATAGTTTAGTCAGAGGTGAAAGCCCAGGGCTCAACCTTGGAACTGCCTTTGATACTGGCTATCTTGAGTTCGGGAGAGGTGAGTGGAATGCCGAGTGTAGAGGTGAAATTCGTAGATATTCGGCGGAACACCAGTGGCGAAGGCGACTCACTGGCCCGATACTGACGCTGAGGCGCGAAAGCGTGGGG |
| P-asv4751 | TCAAGTGATGGCAGATCTTATTGGGCCTAAAGCGTCCGTAGCTTGCCGTGCAAGTCCGTCGGGAAATCCACCCGCTAAACGGGTGGGCGTCCGGCGGAAACTGGATGGCTTGGGACCGGGAGGCTTGAAGGGTACGTCCGGGGTAGGAGTGAAATCCCGTAATCCTGGACGGACCACCGATGGGGAAACCACTTCAAGAGAACGGATCCGACAGTGAGGGACGAAAGCTAGGGT |
| P-asv4752 | GCAAGCGTTGTCCGGAATTATTGGGCGTAAAGAGCTCGTAGGCGGTCTGTTACGTCGGCTGTGAAAACCTGGGGCTCAACCCCGGGCCTGCAGCCGATACGGGCAGACTAGAATTCGGTAGGGGAGACTGGAATTCCTGGTGTAGCGGTGAAATGCGCAGATATCAGGAGGAACACCGGTGGCGAAGGCGGGTCTCTGGGCCGATATTGACGCTGAGGAGCGAAAGCGTGGGG |
| P-asv4753 | GCCAGCGTTGCTCGGAATTACTGGGCGTAAAGGGTTCGTAGGTGGCGCGGCAAGTTGGGAGTGAAATCTCTGGGCTTAACCCAGAGGCTGCTTTCAAAACTGCCGTGCTCGAGTGCGAGAGAGGTCAGTGGAATTGCAGGTGTAGCGGTGAAATGCGTAGATATCTGCAGGAACACCCGTGGCGAAGGCGGCTGACTGGATCGCAACTGACGCTGATGCACGAAAGCTGGGGG |
| P-asv4754 | GCGAGCGTTAATCGGAATTACTGGGCGTAAAGCGTGCGCAGGCGGTGGCTCAAGCCAGGCGTGAAATCCCCGGGCTTAACCTGGGAATGGCGTTTGGGACTGGGTCGCTAGAGTGTGGCAGAGGGAGGTGGAATTCCACGTGTAGCAGTGAAATGCGTAGAGATGTGGAGGAACACCGATGGCGAAGGCAGCCTCCTGGGCCAGCACTGACGCTCATGCACGAAAGCGTGGGG |
| P-asv4755 | GCTAGCGTTGTTCGGAATTACTGGGCGTAAAGCGCGCGCAGGCGGTCCTTCAAGTCAGAGGTGAAAGCCCAGAGCTCAACTCTGGAACGGCCTTTGAGACTAGAGGACTTGGGTACGGGAGAGGTGAGTGGAATTCCGAGTGTAGAGGTGAAATTCGTAGATATTCGGAAGAACACCAGTGGCGAAGGCGGCTCACTGGCCCGTTACCGACGCTCAGGCGCGACAGCGTGGGG |
| P-asv4756 | GCAAGCGTTGTCCAGATTTACTGGGCGTAAAGAGCGTGTAGGCGGCCAGGCAGGTCTGTTGTGAAAACTGGAGGCTCAACCTCCAGACGTCGACAGAAACCGTCTGGCTAGAGTCCGGAAGAGGAGAGTGGAATTCCTGGTGTAGCGGTGAAATGCGCAGATATCAGGAAGAACACCAATGGCGAAGGCAGCTCTCTGGGACGGTACTGACGCTGAGACGCGAAAGCGTGGGG |
| P-asv4757 | GCAAGCGTTACTCGGAATTACTGGGCGTAAAGCGTGCGTAGGTGGTTTGTTAAGTCTGATGTGAAAGCCCTGGGCTCAACCTGGGAACTGCATTGGAAACTGGCTGACTAGAGTGCGGTAGAGGGGTGTGGAATTCCCGGTGTAGCAGTGAAATGCGTAGATATCGGGAGGAACATCAGTTGCGAAGGCGGCTCTCTGGACCAGCACTGACACTGAGGCACGAAAGCGTGGGG |
| P-asv4758 | GCGAGCGTTAATCGGAACTACTGGGCGTAAAGCGTGCGTAGGCGGTTCGTTAAGTCCGTTGTGAAAGCCCTGGGCTCAACCTGGGAATGGCGCTGGATACTGACGAGCTCGAGTGCGGTAGAGGAGGGTGGAATTCCCGGTGTAGCGGTGAAATGCGTAGAGATCGGGAGGAACATCAGTGGCGAAGGCGGCCCTCTGGACCAGCACTGACGCTGAGGCACGAAAGCGTGGGG |
| P-asv4759 | GCTAGCGTTGTTCGGATTTACTGGGCGTAAAGCGCACGTAGGCGGATACTTAAGTCAGGGGTGAGATCCCGGGGCTCAACCCCGGAACTGCCTTTGATACTGGGTATCTCGAGTTCGAGAGAGGTGAGTGGAATTCCGAGTGTAGAGGTGAAATTCGTAGATATTCGGAGGAACACCAGTGGCGAAGGCGGCTCACTGGCTCGATACTGACGCTGAGGTGCGAAAGCGTGGGG |
| P-asv4760 | GCAAGCGTTGTTCGGAATCACTGGGCGTAAAGGGCGTGTAGGCGGTTCGGTAAGTGGGATGTGAAATGCCCGGGCTCACCCCGGGACCTGCATCCCATACTGCTGAGCTTGAGTACAGGAGAGGGTGGGGAATTCCTGGTGTAGCGGTGAAATGCGTAGATATCAGGAGGAACACCGGTGGCGAAGGCGCCCACCTGGCCTGCTACTGACGCTGATGCGCGAAAGCTAGGGG |
| P-asv4761 | GCTAGCGTTGTCCGGAATCATTGGGCGTAAAGCGCGTGTAGGCGGCCCGGTAAGTCCGCTGTGAAAGTCGGGGGCTCAACCCTCGGATGCCGGTGGATACTGTCGGGCTCGAGTGCGGGAGAGGCGAGTGGAATTCCTGGTGTAGCGGTGAAATGCGCAGATATCAGGAGGAACACCGATTGCGAAGGCAGCTCGCTGGAACGTTACTGACGCTGAGACGCGAAAGCGTGGGG |
| P-asv4762 | GCGAGCGTTGTTCGGATTTACTGGGCGTAAAGGGCGCACAGGCGGCTTTGTTTGTCAGGCGTGAAAGCCCCGGGCTTAACCTGGGAACTGCGCCTGGGACGGCGAGGCTAGAGATCGGGAGAGGAGAGTGGAATTCCCAGTGTAGAGGTGAAATTCGTAGATATTGGGAAGAACACCGGTGGCGAAGGCGGCTCTCTGGACCGATACTGACGCTGAGGCGCGAAAGCGTGGGG |

| | |
|---|---|
| P-asv4763 | GCGAGCGTTAATCGGAATTACTGGGCGTAAAGTGTGCGCAGGCGGTCTCGCCAGTCAAGTGTGAAAGCCCCGGGCTTAACCTGGGAACTGCGCTTGAAACTACGAGACTCGAGTGTGGCAGAGGAAGGTGGAATTCCACGTGTAGCGGTGAAATGCGTAGAGATGTGGAGGAACACCGATGGCGAAGGCATCCTCCTGGGCCAGCACTGACGCTCATGCACGAAAGCGTGGGG |
| P-asv4764 | GCGAACGTTATTCGGAATCACTGGGCTTAAAGCGCGTGTAGGCGGGACGGCACGTCGGTCGTTGAAAGCCCCCGGCTTAACCGGGGAAGTGGCGCCGATACGACCGTCCTGGAGGGACGTAGGGGGACCTGGAACTTCCGGTGGAGCGGTGAAATGCGTTGAGATCGGAAGGAACGCCCGTGGCGAAAGCGAGGTCCTGGACGTTTTCTGACGCTGAGACGCGAAAGCCAGGGG |
| P-asv4765 | GCGAGCGTTATTCGGAATTATTGGGCGTAAAGGGCGCGTAGGTGGCCCTGTAAGTCAAAGGTGAAATCCCTCGGCTCAACCGAGGAATTGCCCTTGAAACTGCAGGGCTTGAGGCCGGGAGGGGGTAGCGGAATTCCCAGTGTAGCGGTGAAATGCGTAGATACTGGGAGGAACACCGGTGGCGAAGGCGGCTACCTGGACCGGTTCTGACACTGATGCGCGAAAGCGTGGGT |
| P-asv4766 | ACAAGCGTTGTCCGGATTTACTGGGCGTAAAGAGCGCGCAGGCGGTCGTTCAAGTCGAGTGTGAAAGCCCCCGGCTCAACTGGGGAGGGTCATTCGATACTGATCGACTCGAAGGCAGGAGAGGGTAGTGGAATTCCAGGTGTAGCGGTGGAATGCGTAGAGATCGGGAAGAACATCGGTGGCGAAGGCGGCCTGCTGGGCTCTTGCTGACACTGAGGCGCGACAGCGTGGGG |
| P-asv4767 | GCGAGCGTTGTCCGGAATCACTGGGCGTAAAGGGCGCGTAGGTGGCCTGATAAGTAGGGGGTGAAATCTCGAGGCTCAACTTCGAGGCTGCCTCCTAAACTGTCAGGCTCGAGCGGAGTAGAGGCAGGTGGAATTCCGGGTGTAGCGGCGGAATGCGTAGAGATCCGGAAGGACACCGGTGGCGAAGGCGGCCTGCTGGGCAGTAGCTGACGCTGAGGCGCGACAGCGTGGGG |
| P-asv4768 | GCGAGCGTTGTCCGGAATTATTGGGCGTAAAGAGCTCGTAGGCGGCCTGTTGCGTCCGCTGTGAAATCCTGGGGCTTAACCCCGGGCGTGCAGTGGATACGGGCAGGCTGGAGGCAGGCAGGGGAGAACGGAATTCCCGGTGTAGCGGTGAAATGCGCAGATATCGGGAGGAACACCGGTGGCGAAGGCGGTTCTCTGGGCCTGTACTGACGCTGAGGAGCGAAAGCGTGGGG |
| P-asv4769 | GCAAGCGTTGTTCGGAATTATTGGGCGTAAAGGGCGCGTAGGCGGCGCGACAAGTCACCTGTGAAATCTCCGGGCTTAACTCGGAGCCTGCAGGCGAAACTGTCGTGCTGGAGTATGGGAGAGGTGCGTGGAATTCCCGGTGTAGCGGTGAAATGCGTAGATATCGGGAGGAACACCAGTGGCGAAGGCGATCATCTGGGGTTGTTCTGACGCTGATGAGCGAAAGCTAGGGG |
| P-asv4770 | GCTAGCGTTGTTCGGAATTATTGGGCGTAAAGGGCGCGTAGGAGGGCGAACAAGTCTGACGTGAAATCCCTCGGCCTAACCGGGGAACTGCGTTTGAAACTGTTCGGCTCGAGTCCGGGAGAGGGTAGTGGAATTCCCAGTGTAGCGGTGAAATGCGTAGATATTGGGAGGAACACCGGTGGCGAAGGCGGCTACCTGGACCGGCACTGACGCTGAGGCGCGAAAGCCAGGGG |
| P-asv4771 | GCAAACGTTGCTCGGAATTATTGGGCGTAAAGCGCACGTAGGCGGCTTTGTAAGTCGGATGTGAAATCCCTCGGCTTAACCGAGGACGTGCATCCGAAACTGCGAGGCTCGAGTATGGAAGAGGGTCGCAGAATTCCCGGTGTAGAGGTGAAATTCGTAGATATCGGGAGGAATACCAGTGGCGAAGGCGGCGACCTGGGCCAATACTGACGCTGAGGTGCGAAAGCGTGGGG |
| P-asv4772 | ACAAGCGTTGTCCGGAATTACTGGGTGTAAAGGGAGCGCAGGCGGGAGGACAAGTTGGAAGTGAAATCCATGGGCTCAACCCATGAACTGCTTTCAAAACTGTTTTTCTTGAGTAGTGCAGAGGTAGGCGGAATTCCCGGTGTAGCGGTGGAATGCGTAGATATCGGGAGGAACACCAGTGGCGAAGGCGGCCTACTGGGCACCAACTGACGCTGAGGCTCGAAAGTGTGGGT |
| P-asv4773 | GCAAGCGTTGTCCGGAATTACTGGGCGTAAAGAGCGCGTAGGTGGCTAGATAAGTCCGGTGTGAAATCCCACGGCTTAACCGGGGAATGGCACCGGAGACTGTATAGCTCGAGGACGGGAGAGGAGAGCGGAATTCCCGGTGTAGCGGTGAAATGCGTAGAGATCGGGAGGAAGGCCGGTGGCGAAGGCGGCTCTCTGGAACGTCCCTGACGCTGAGGCGCGAAAGCGTGGGG |
| P-asv4774 | CCAAGCGTTGTTCGGATTCACTGGGCGTAAAGGGTGCGTAGGTGGCAGGGTAAGTTTGATGTGAAATCTCCGGGCTTAACCCGGAAACTGCATTGAATACTATCTGGCTAGAGGGTTGGAGGGGGGACTGGAATACTTGGTGTAGCAGTGAAATGCGTAGATATCAAGTGGAACACCAGTGGCGAAGGCGAGTCCCTGGACAACTCCTGACACTGAGGCACGAAAGCTAGGGG |
| P-asv4775 | GCCAGCGTTGTTCGGAATTACTGGGTGTAAAGGGTCGGTAGGCGGTGTGGCAAGTTGGGAGTGAAATCTATGGGCTCAACCCATTGGCTGCTTCCAAGACTGCTGTGCTTGAGTGTGGGAGAGGCGCGTGGAATTGCAGGTGTAGCGGTGAAATGCGTAGATATCTGCAGGAACACCCGTGGCGAAAGCGGCGCGCTGGACCACAACTGACGCTGAATGACGAAAGCTAGGGG |
| P-asv4776 | GCAAGCGTTGTCCGGAATTACTGGGCGTAAAGGGCGCGTAGGCGGCCCGATAAGTAGGAGGTGAAATTCCGTGGCTCAACCACGGGGCGGCCTTCTAGACTGTCAGGCTTGAGCACGGTAGAGGCAGATGGAATTCTCGGTGTAGCGGTGGAATGCGTAGAGATCGGGAAGAACACCAGTGGCGAAGGCGGTCAGCTGGGCCGTTGCTGACGCTGAGGCGCGACAGCGTGGGG |
| P-asv4777 | GCAAGCGTTGTTCGGATTTACTGGGCGTAAAGGGCGCGTAGGCGGCGCCCCGAGTCAATTGTGAAATCTCCGGGCTTAACTCGGAAAGGTCAACTGATACTGCGGGGCTAGAGTGCGGAAGGGGCAACTGGAATTCTCGGTGTAGCGGTGAAATGCGTAGATATCGAGAGGAACACCTGCGGCGAAGGCGGGTTGCTGGGCCGACACTGACGCTGAGGCGCGAAAGCTAGGGG |
| P-asv4778 | GCGAGCGTTAATCGGAATAACTGGGCGTAAAGGGCACGCAGGCGGCTATTTAAGTGAGGTGTGAAATCCCCGGGCTTAACCTGGGAATTGCATTTCATACTGGGTAGCTAGAGTACTTTAGGGAGGGGTAGAATTCCACGTGTAGCGGTGAAATGCGTAGAGATGTGGAGGAATACCGAAGGCGAAGGCAGCCCCTTGGGAATGTACTGACGCTCATGTGCGAAAGCGTGGGG |
| P-asv4779 | GCAAGCGTTGTTCGGAACTACTGGGCGTAAAGCGCGTGTAGGTGGTCGAGTAAGTCGAATGTGAAAGCCCCGGGCTCAACCCGGGAAGTGCATCCGAAACTGCTCGACTAGAGTCCCGGAGAGGAAGGCAGAATTCCCAATGTAGGGGTGAAATCCGTAGATATTGGGAGGAATACCGGTGGCGAAGGCGGCCTTCTGGACGGTGACTGACACTCAGACGCGAAAGCGTGGGG |
| P-asv4780 | GCGAGCGTTATTCGGAATTATTGGGCGTAAAGGGCGCGTAGGCGGCCCGGTAAGTCAAAGGTGAAATCCCTCGGCTCAACTGAGGAACTGCCTTTGAAACTGTCGGGCTTGAGGCCGGGAGAGGGTAGTGGAATTCCCAGTGTAGCGGTGAAATGCGTAGATATTGGGAGGAACACCAGTGGCGAAGGCGGCTACCTGGACCGGTTCTGACGCTGAGGCGCGAAAGCGTGGGT |
| P-asv4781 | GCAAGCGTTGTCCGGAATCATTGGGCGTAAAGCGCGCGTAGGCGGCCCGGTAAGTCCGTTGTGAAAGTCAAAGGCTCAACCTTTGAATGCCGATGGATACTGCCGGGCTAGAGTCCGGAAGAGGCGAGTAGAATTCCTGGTGTAGCGGTGGAATGCGCAGATATCAGGAGGAATACCCATAGCGAAGGCAGCTCGCTGGGACGGTACTGACGCTAAGGCGCGAAAGCGTGGGG |
| P-asv4782 | GCTAGCGTTGTTCGGAATTACTGGGCGTAAAGCGCACGTAGGCGGATTTTTAAGTCAGGGGTGAAATCCCGAGGCTCAACCTCGGAACTGCCTTTGATACTGGAGATCTTGAGTTCGGAAGAGGTTGGTGGAACAGCTAGTGTAGAGGTGAAATTCGTAGATATTAGCTAGAACACCAGTGGCGAAGGCGGCCAACTGGTCCGATACTGACGCTGAGGTGCGAAAGCGTGGGG |
| P-asv4783 | GCTAGCGTTGCTCGGAATCACTGGGCGTAAAGGGCGCGTAGGCGGCGCCGCAAGTCGGGCGTGAAATCCCAGGGCTCAACCCGGGGGCTGCGCCCGAGACTGCGGGGCTGGAGGACGGGAGAGGCTCGCGGAATTCCCAGTGTAGAGGTGAAATTCGTAGATATTGGGAAGAACACCGGTGGCGAAGGCGGCGAGCTGGCCCGTTTCTGACGCTGAGGCGCGACAGCGTGGGG |
| P-asv4784 | GCAAGCGTTGTCAGGAATTATTGGGCGTAAAGCGCGCGCAGGCGGCCTCTTAAGTCTGATGTGAAAGCCCCCGGCTCAACCGGGGAGGGCCATTGGAAACTGGGAGGCTTGAGTATAGGAGAGAAGAGTGGGATTCCACGTGTAGCGGTGAAATGCGTAGAGATGTGGAGGAACACCAGTGGCGAAGGCGACTCTTTGGCCTATAACTGACGCTGAGGCGCGAAAGCGTGGGG |
| P-asv4785 | GCAAGCGTTATCCGGAATCATTGGGTTTAAAGGGTCCGCAGGCGGTCAATTAAGTCAGAGGTGAAATCCAGTCGCTTAACGATTGAACTGCCTTTGATACTGGTTGACTTGAGTCATATGGAAGTAGATAGAATGTGTAGTGTAGCGGTGAAATGCTTAGATATTACACAGAATACCGATTGCGAAGGCAGTCTACTACGTAGGTACTGACGCTCATGGACGAAAGCGTGGGG |
| P-asv4786 | GCGAGCGTTAATCGGAATTACTGGGCGTAAAGCGCGCGTAGGTGGTTCGGTAAGTCGGTTGTGAAATCCCTGGGCTTAACCTGGGAATTGCATTCGATACTGCCGGACTGGAGTGTGGTAGAGGTAAGCGGAATTCCCGGTGTAGCGGTGAAATGCGTAGATATCGGGAGGAACACCAGTGGCGAAGGCGGCTTACTGGACCAACACTGACACTGAGGTGCGAAAGCGTGGGT |
| P-asv4787 | GCGAGCGTTGTCCGGAATTACTGGGCGTAAAGAGCTTGTAGGCGGTTCGTCGCGTCGTTCGTGAAAACTTGGGGCTCAACCCCAAGCTTGCGGGCGATACGGGCGGACTAGAGTACTGCAGGGGAGACTGGAATTCCTGGTGTAGCGGTGAAATGCGCAGATATCAGGAGGAACACCGGTGGCGAAGGCGGGTCTCTGGGCAGTAACTGACGCTGAGAAGCGAAAGCGTGGGT |
| P-asv4788 | CCAAGCGTTGTCCGGATTTATTGGGCGTAAAGAGCTCGTAGGCGGTTCAGTTAGTCGGGTGTGAAAACTCTGGGCTCAACTCAGAGACGCCACCCGATACTGCTGTGACTTGAGTCCGGTAGGGGAGCGGGGAATTCCAGGTGTAGCGGTGAAATGCGTAGAGATTGGGAAGAACACCAGTGGCGAAGGCGGCTATCTGGACGGTTTCTGACGCTAAGACGCGAAAGCGTGGGG |
| P-asv4789 | GCTAGCGTTGCTCGGAATCGCTGGGCGTAAAGGGCGCGTAGGCGGCGTTTTAAGTCGGGGGTGAAAGCCTGTGGCTCAACCACAGAATGGCCTTCGATACTGGGACGCTTGAGTATGGTAGAGGTCGGTGGAACTGCGAGTGTAGAGGTGAAATTCGTAGATATTCGCAAGAACACCGGTGGCGAAGGCGGCCGACTGGACCATTACTGACGCTGAGGCGTGAAAGCGTGGGG |
| P-asv4790 | CCAAGCGTTAATCGGAATTACTGGGCGTAAAGCGTGCGCAGGCGGTTTGTTAAGCGAGATGTGAAAGCCCCGGGCTCAACCTGGGAATTGCATTTCGAACTGGCAAACTAGAGTCTTGTAGAGGGGGGTAGAATTCCAGGTGTAGCGGTGAAATGCGTAGAGATCTGGAGGAATACCGGTGGCGAAGGCGGCCCCCTGGACAAAGACTGACGCTAAGGCACGAAAGCGTGGGG |
| P-asv4791 | GCAAGCGTTATCCGGATTTATTGGGTTTAAAGGGTCCGTAGGCGGATGTGTAAGTCAGTGGTGAAATCTCGCAGCTTAACTGCGAAACTGCCATTGATACTGCATGTCTTGAGTAAATTTGAAGTGGCTGGAATAAGTAGTGTAGCGGTGAAATGCATAGATATTACTTAGAACACCAATTGCGAAGGCAGGTCACTAAGATTTAACTGACGCTGATGGACGAAAGCGTGGGG |
| P-asv4792 | GCAAGCGTTAATCGGAATTACTGGGCGTAAAGCGCGCGTAGGCGGTTAAATAAGTCAGATGTGAAATCCCTGGGCTCAACCTAGGAACTGCATTTGAAACTATTTGGCTAGAGTATGGTAGAGGTGAGTGGAATTTCAGGTGTAGCGGTGAAATGCGTAGAGATCTGAAGGAACATCAGTGGCGAAGGCGACTCACTGGGCCATTACTGACGCTGAGGTGCGAAAGCGTGGGT |
| P-asv4793 | GCAAGCGTTAATCGGAATTACTGGGCGTAAAGCGCGCGTAGGTGGTTATTTAAGTCAGATGTGAAAGCCCCGGGCTTAACCTGGGAACTGCATCTGATACTGGATAACTAGAGTAGGTGAGAGGGGAGTAGAATTCCAGGTGTAGCGGTGAAATGCGTAGAGATCTGGAGGAATACCGATGGCGAAGGCAGCTCCCTGGCATCATACTGACACTGAGGTGCGAAAGCGTGGGT |
| P-asv4794 | GCAAGCGTTAATCGGAATTACTGGGCGTAAAGCGCGCGTAGGTGGTTTATTAAGTTGGATGTGAAATCCCCGGGCTCAACCTGGGAACTGCATCCAAAACTGGCAAGCTAGAGTATGGTAGAGGGTGGTGGAATTTCCTGTGTAGCGGTGAAATGCGTAGATATAGGAAGGAACACCAGTGGCGAAGGCGACCACCTGGACTGATACTGACACTGAGGTGCGAAAGCGTGGGG |
| P-asv4795 | GCGAGCGTTAATCGGAATTACTGGGTGTAAAGCGCGCGTAGGCGGTTTGTTAAGCCGGTTGTGAAATCCCTAGGCTCAACCTAGGAACTGCACCCGGAACTGGCAAGCTAGAGTATAGTAGAGGAAGGTGGAATTCCACGTGTAGCGGTGAAATGCGTAGATATGTGGAGGAACATCAGTGGCGAAGGCGGCCTTCTGGACTAATACTGACGCTGAGGTGCGAAAGCGTGGGG |
| P-asv4796 | GCTAGCGTTGTTCGGAATTACTGGGCGTAAAGCGCACGTAGGCGGATCAGTCAGTTAGAGGTGAAATCCCAGGGCTCAACCCTGGAACTGCCTTTAATACTGCTGGTCTGGAGGTCGAGAGAGGCGAGTGGAATTCCGAGTGTAGAGGTGAAATTCGTAGATATTCGGAGGAACACCAGTGGCGAAGGCGGCTCGCTGGCTCGATACTGACGCTGAGGTGCGAAAGCGTGGGG |
| P-asv4797 | GCAAGCGTTGTCCGGAATTATTGGGCGTAAAGCGCGCGCAGGCGGTCCTTTAAGTCTGATGTGAAAGCCCACGGCTCAACCGTGGAGGGTCATTGGAAACTGGGGACTTGAGTGCAGAAGAGAAGAGTGGAATTCCACGTGTAGCGGTGAAATGCGTAGATATTCGGAAGAACACCAGTGGCGAAGGCGGCTCGCTGGACTGGTATTGACGCTGAGGTGCGAAAGCGTGGGG |
| P-asv4798 | GCGAGCGTTGTCCGGAATTACTGGGTGTAAAGGGAGCGTAGGCGGGACGGCAAGTCAGATGTGAAAACTATGGGCTCAACTCATAGACTGCATTTGAAACTGTTGTTCTTGAGTGAGGTAGAGGTAAGCGGAATTCCTGGTGTAGCGGTGAAATGCGTAGAGATCAGGAGGAACATCGGTGGCGAAGGCGGCTTACTGGGCCTTTACTGACGCTGAGGCTCGAAAGCGTGGGG |
| P-asv4799 | GCAAGCGTTAATCGGAATTACTGGGCGTAAAGCGTACGTAGGTGGTTTGTTAAGTTGGATGTGAAAGCCCAGGGCTCAACCTTGGAACTGCATTCAAAACTGATGAACTAGAGTATGGTAGAGGTAAGTGGAATTTCCTGTGTAGCGGTGAAATGCGTAGAGATGGGAAGGAACATCAGTGGCGAAGGCGACTTACTGGACCAATACTGACACTGAGGTACGAAAGCGTGGGT |
| P-asv4800 | GCAAGCGTTATCCGGATTTACTGGGTGTAAAGGGAGCGCAGGCGGTGCGGCAAGTCTGATGTGAAAGCCCGGGGCTCAACCCCGGTACTGCATTGGAAACTGTCGTACTAGAGTGTCGGAGGGGTAAGCGGAATTCCTAGTGTAGCGGTGAAATGCGTAGATATTAGGAGGAACACCAGTGGCGAAGGCGGCTTACTGGACGATAACTGACGCTGAGGCTCGAAAGCGTGGGG |
| P-asv4801 | GCGAGCGTTGTTCGGAATCACTGGGCGTAAAGCGCGCGCAGGCGGGACTTGCAAGTCGGAAGTGAAATCTCGGGGCTCAACTTCGAGGCTGCTTCCGATACTGCGGACCTGGAGACCAGTAGGGGTTGGTGGAATTCCCGGTGTAGCGGTGGAATGCGTAGAGATCGGGAAGAACACCTGTGGCGAAGGCGGCCAACTGGGCTGGATCTGACGCTGAGGCGCGAAAGCGTGGGG |

| | |
|---|---|
| P-asv4802 | GCTAGCGTTGTTCGGAATTACTGGGCGTAAAGCGTGCGTAGGCGGATTATTAAGTTAGGGGTGAAATCCCGGGGCTCAACCTCGGAACTGCCTCTAAAACTGGTAATCTAGAGATATGGAGAGGTAAGTGGAATT<br>CCTAGTGTAGAGGTGAAATTCGTAGATATTAGGAGGAACACCAGAGGCGAAGGCGGCTTACTGGACATATACTGACGCTGAGGCACGAAAGTGTGGGT |
| P-asv4803 | GCAAGCGTTGTCCGGAATTATTGGGCGTAAAGAGCTCGTAGGCGGCTTGTCGCGTCGGATGTGAAAGCCCGGGGCTTAACCCCGGGTCTGCATTCGATACGGGCAGGCTAGAGTTCGGTAGGGGAGATCGGAA<br>TTCCTGGTGTAGCGGTGAAATGCGCAGATATCAGGAGGAACACCGGTGGCGAAGGCGGATCTCTGGGCCGATACTGACGCTGAGGAGCGAAAGCGTGGGG |
| P-asv4804 | GCGAGCGTTGTCCGGATTTACTGGGCGTAAAGGGTGAGTAGGCGGTTGTATATGTCAGGTGTAAAAGGCTATGGCTTAACCATGGTTAGCACTTGAAACTGTACAACTTGAGTGCAGGAGAGGTAAGTGGAATTC<br>CTAGTGTAGCGGTGAAATGCGTAGATATTAGGAGGAACACCAGTGGCGAAGGCGACTTACTGGACTGTAACTGACGCTGAGTCACGAAAGCGTGGGT |
| P-asv4805 | GCGAGCGTTAATCGGAATCACTGGGCGTAAAGCGTACGTAGGCGGCGTGTCAAGTCAGGTGTGAAAGCCCTCGGCTCAACCGGGGAATTGCACTTGAAACTGATATGCTAGAGTTCTGGAGGGGGTAGCGGAAT<br>TCCCGGTGTAGGAGTGAAATCCGTAGATATCGGGAGGAACACCAGTGGCGAAGGCGGCTACCTGGACAGTAACTGACGCTGAGGTACGAAAGTGTGGGG |
| P-asv4806 | GCGAGCGTTGTCCGGAATCACTGGGCGTAAAGCGCGCGCAGGCGGCGATCTACGTCTGGGGTGAAAGCCCGGAGCTCAACTCCGGGACTGCCTTGGATACGGGATCGCTTGAGGATTGGAGAGGCAAGGGGA<br>ATTCCACGTGTAGCGGTGAAATGCGTAGAGATGTGGAGGAACACCTGTGGCGAAGGCGCCTTGCTGGCCAATTACTGACGCTGAGGCGCGAAAGCGTGGGG |
| P-asv4807 | GCGAACGTTAATCGGAATTACTGGGCGTAAAGCGCGTGTAGGCGGTTCGTTAAGTGGGATGTGAAAGCCCTGGGCTCAACCCAGGAATTGCATTCCAGACTGGCGGACTCGAGTGCAGTAGAGGGAGGTGGAA<br>TTCCAGGTGTAGCGGTGAAATGCGTAGATATCTGGAGGAACATCGGTGGCGAAGGCGGCCTCCTGGACTGACACTGACGCTGAGACGCGAAAGCGTGGGG |
| P-asv4808 | GCAAGCGTTAATCGGAATTACTGGGCGTAAAGCGAGCGTAGGTGGCTTGATAAGTCAGATGTGAAAGCCCCGGGCTTAACCTGGGAACGGCATCTGATACTGTTAGGCTAGAGTAGGTGAGAGGAAGGTAGAAT<br>TCCAGGTGTAGCGGTGAAATGCGTAGAGATCTGGAGGAATACCGATGGCGAAGGCAGCCTTCTGGCATCATACTGACACTGAGGTTCGAAAGCGTGGGG |
| P-asv4809 | GCGAGCGTTGTTCGGAATCACTGGGCGTAAAGCGTGCGTAGGCGGTTTGATAAGTCAGGAGTGAAATCCCGGAGCTCAACTCCGGAACTGCTTCTGAAACTATCTTACTAGAGTATGGTAGAGGTTGGGAGAATT<br>CCTAGTGTAGAGGTGAAATTCGTAGATATTAGGAGGAATACCAGTGGCGTAGGCGCCCAACTGGACCATTACTGACGCTCAGGCACGAAAGCGTGGGG |
| P-asv4810 | TCAAGTGGTCAGGATGATTATTGGGCCTAAAGCATCCGTAGCTCGTTTTGTAAGTTTTCGGTTAAATCCATACGCTTAACGTATGGGCTGCCGGGAATACTGCATAACTAGGAAGTGGGAGAGGTAGACGGTACTC<br>GGTAGGAAGGGGTAAAATCCTTTGATCTATTGATGACCACCTGTGGCGAAGGCGGTCTACCAGAACACGTTCGACGGTGAGGGATGAAAGCTGGGGG |
| P-asv4811 | GCAAGCGTTATTCGGAATGACTGGGCGTAAAGAGCGTGAAGGCGGTTCTTTAAGTCATCTGTTAAAGCGCTCGGCTTAACCGAGCATATGCGGGTGATACTGGAGGGCTAGAGTACAGGAGAGAAAGGTAGAAT<br>TCTCGGAGTAGCGGTAAAATGCGTAGATCTCGAGAGGAATACCGGTTGCGAAGGCGGCCTTTTGGCCTGTTACTGACGCTTAAGCGCGAAAGCGTGGGG |
| P-asv4812 | GCGAGCGTTGTCCGGAGTTACTGGGCGTAAAGAGCGTGTAGGCGGTTGAAAAAGTCAGGTGTGAAATTCTTCGGCTCAACCGGGGGACTGCGCCTGAAACTTTTCGACTGGAGTTCGGGAGGGGCAACTGGAA<br>TTCCTGGTGTAACGGTGAAATGTGTAGATATCAGGAGGAACACCGGTGGCGAAGGCGGGTTGCTGGACCGAAACTGACGCTGAGACGCGAAAGCGTGGGG |
| P-asv4813 | GCTAGCGTTGTTCGGAATTACTGGGCGTAAAGCGCACGTAGGCGGACTGGAAAGTCAGAGGTGAAATCCCAGGGCTCAACCTTGGAACTGCCTTTGATACTGGTTGACTTGAGTCATATGGAAGTAGATAGAATG<br>TGTAGTGTAGCGGTGAAATGCTTAGATATTACACAGAATACCGATTGCGAAGGCAGTCTACTACGTATGTACTGACGCTCATGGACGAAAGCGTGGGG |
| P-asv4814 | GCGAGCGTTGTTCGGAATTACTGGGCGTAAAGCGTGCGTAGGCGGCTATGAAAGTTAGAAGTGAAAGCCCCGGGCTCAACCCGGGAATTGCTTTTAAAACTCCATAGCTTGAATTCGGGAGAGGATAGCGGAATT<br>CTCAGTGTAGCAGTGAAATGCGTAGATATTGAGAGGAACACCAGTGGCGTAGGCGGCTATCTGGCCCGACATTGACGCTGAGGCACGAAAGCGTGGGG |
| P-asv4815 | GCAAGCGTTATCCGGATTCACTGGGTTTAAAGGGTGCGTAGGCGGGTATGTAAGTCAGTGGTGAAATCTCGGAGCTTAACTCCGAAACTGTCATTGATACTATATATCTTGAATATTGTGGAGGTTTGCGGAATAT<br>GTCATGTAGCGGTGAAATGCTTAGATATGACATAGAACACCTATTGCGAAGGCAGCAGGCTACACATATATTGACGCTGAGGCACGAAAGCGTGGGG |
| P-asv4816 | GCTAGCGTTGTTCGGAATTACTGGGCGTAAAGGGCGCGTAGGCGGCTTTTTGCGTGAGATGTGAAAGACCTGGGCTCAACCTGGGAAGTGCATTTCATACGGGAAAGCTAGAGTGCGAGAGGGGAAAGTGGAAT<br>GATGCGTGTAGAGGTGAAATTCGTAGATATGCATCGGAACACCAGTGGCGAAGGCGACTTTCTGGCTCGTAACTGACGCTAAGGCGCGAAAGCGTGGGG |
| P-asv4817 | GCTAGCGTTGTTCGGAATGACTGGGCGTAAAGGGCGCGTAGGCGGCGATCTAAGTCAGGCGTGAAAGCCCCGGGCTTAACCTGGGAATTGCGCATGATACTGGGTTGCTTGAGTGCGGGAGAGGATGGTGGAA<br>TTCCCAGTGTAGAGGTGAAATTCGTAGATATTGGGAAGAACACCGGTGGCGAAGGCGGCCATCTGGACCGCAACTGACGCTGAGGCGCGAAAGCGTGGGG |
| P-asv4818 | GCTAGCGTTGTTCGGAATTACTGGGCGTAAAGCGCACGTAGGCGGCTTTGTAAGTTAGAGGTGAAAGCCCGGGGCTCAACTCCGGAACTGCCTTTAAGACTGCATCGCTTGAATCATGGAGAGGTTAGTGGAATT<br>CCGAGTGTAGAGGTGAAATTCGTAGATATTCGGAAGAACACCAGTGGCGAAGGCGACTAACTGGACATGTATTGACGCTGAGGTGCGAAAGCGTGGGG |
| P-asv4819 | GTTAGCGTTGTTTGGAATTACTGGGCGTAAAGCGCGCGTAGGCGGGCTGGAAAGTTGGGGGTGAAATCCCGGGGCTCAACCCCGGAACTGCCTTCAAAACTCCCAGTCTGGAGGTCGAGAGAGGTGAGTGGAA<br>TTCCGAGTGTAGAGGTGAAATTCGTAGATATTCGGAGGAACACCAGTGGCGAAGGCGGCTCACTGGCTCGCACCTGACGCTGAGGCGCGAAAGTGTGGGG |
| P-asv4820 | GCAAACGTTATTCGGAATCACTGGGCTTAAAGGGCGCGTAGGCGGGCTGTCAAGTCTGGGGTGAAATCCCGCGGCTCAACCGTGGAACTGCCTCAGATACTGACGGCCTCGAGGGAGGTAGGGGCATGCGGAA<br>CTGTAGGTGGAGCGGTGAAATGCGTTGATATCTACAGGAACTCCGGTGGCGAAAGCGGCGTGCTGGACCTCTTCTGACGCTGAGGCGCGAAAGCTAGGGG |
| P-asv4821 | TCGAGTGATGACCGATTTTATTGGGCCTAAAGCGTCCGTAGCTGGCCACGAAAGCCCGTCGGGAAATCTCTCGGCTCAACCGAGAGGCGTCCGGCGGAACCTCCGTGGCTTGGAACCGGAAGGCTCAGAGGGT<br>ACGTCCAGGGTAGGAGTGAAATCCTGTAATCCTGGACGGACCACCGATGGCGAAAGCACTCTGAGAGGACGGATTCGACAGTGAGGGACGAAAGCTAGGGT |
| P-asv4822 | GCGAGCGTTGTCCGGATTTATTGGGTTTAAAGGGTGCGTAGGTGGCCTAGTAAGTCAGTGGTGAAATACGGTTGCTCAACAATCGAGGTGCCATTGATACTGCAAGGCTTGAAATAATTGGAGGCTGCCGGAATG<br>GATGGTGTAGCGGTGAAATGCATAGATATCATCCAGAACACCGATTGCGAAGGCAGGTGGCTACGATTTGTTTGACACTGAGGCACGAAAGCATGGGG |
| P-asv4823 | GCGAACGTTGTTCGGAATTACTGGGCGTAAAGCGCGTGTAGGCGGTGAGGCAAGTCGGATGTGAAAGCCCAGGGCTCAACCCTGGAAGTGCACTCGATACTGCCTTGCTGGAGTCCCGGAGAGGATGGTGGAA<br>CTCTCGGTGTAGAGGTGAAATTCGTAGATATCGAGAAGAACACCGGTGGCGAAGGCGGCCATCTGGACGGTGACTGACGCTGAGACGCGAAAGCGTGGGG |
| P-asv4824 | CCAAGCGTTATCCGGATTTATTGGGTTTAAAGGGTCCGTAGGCGGGTCATTAAGTCAGTGGTGAAATCCTACAGCTCAACTGTAGAACTGCCATTGAAACTGATGACCTTGAATATAATTGAAGTGGGCGGAATGT<br>GTCATGTAGCGGTGAAATGCTTAGATATGACACAGAACACCGATTGCGAAGGCAGCTCACTAAGTTATTATTGACGCTGAGGGACGAAAGCCTGGGT |
| P-asv4825 | CCGGGCGTTATCCGGATTTATTGGGTTTAAAGGGAGTGTAGGCGGCCTGTTAAGCGTGTTGTGAAATGTAGATGCTCAACATCTGAACTGCAGCGCGAACTGGCTGGCTTGAGTACACGCAACGTGGGCGGAAT<br>TCATGGTGTAGCGGTGAAATGCTTAGATATCATGAAGAACTCCTATTGCGAAGGCAGCTCACGGGAGTGTCACCGACGCTTAAGCTCGAAGGTGCTGGT |
| P-asv4826 | ACGAGTGATGGCCGATGTTATTGGGCCTAAAGCGTCCGTAGCCGGCCAAACAAGTCCGTTGGGAAATCGACGCGCTCAACGCGTCGGCGTCCGGCGGAAACTGTTTGGCTTGGGGCCGGAAGACCTGAGGGGT<br>ACGTCCGGGGTAGGAGTGAAATCCCGTAATCCTGGACGGACCACCGTTGGCGAAAGCGCCTCAGGAAGACGGACCCGACGGTGAGGGACGAAAGCTAGGGT |
| P-asv4827 | GCAAGCGTTGTCCGGAATTATTGGGCGTAAAGCGCGCGCAGGCGGCCATTTAAGTCTGGTGTTTAATCCCGGGGCTCAACTCCGGGTCGCATCGGAAACTGGGTGGCTGGAGTGCAGAAGAGGAAAGTGGAAT<br>TCCACGTGTAGCGGTGAAATGCGTAGAGATGTGGAGGAACACCAGTGGCGAAGGCGACTTTCTGGGCTGTAACTGACGCTGAGGCGCGAAAGCGTGGGG |
| P-asv4828 | GCAAGCGTTACTCGGAATTACTGGGCGTAAAGCGTGCGTAGGTGGTGGCTTAAGTCCGTTGTGAAAGCCCTGGGCTCAACCTGGGAATTGCAGTGGATACTGGGTCACTAGAGTGTGGTAGAGGGTGGCGGAAT<br>TCCCGGTGTAGCAGTGAAATGCGTAGAGATCGGGAGGAACATCCGTGGCGAAGGCGGCCACCTGGGCCAACACTGACACTGAGGCACGAAAGCGTGGGG |
| P-asv4829 | GCAAGCGTTAATCGGAATTACTGGGCGTAAAGCGCACGTAGGCGGCCCTGCAAGTCAGATGTGAAAGCCGACGGCTCAACCGTTGAATTGCATTTGAAACTGCAGGGCTCGAGTCCTGGAGAGGGTGGCGGAAT<br>TCCCGGTGTAGGAGTGAAATCCGTAGATATCGGGAGGAACACCAGTGGCGAAGGCGGCCACCTGGACAGGTACTGACGCTGAGGTGCGAAAGTGTGGGG |
| P-asv4830 | TCAAGCGTTGTTCGGAATTACTGGGCGTAAAGCGAGTGTAGGTGGCGTATTAAGTCAGATGTGAAAGCCCTAGGCTTAACCTAGGAAGAGCATTTGATACTGATATGCTTGAGTGCGAGAGAGGAAAGTGGAATT<br>ACAGGTGTAGGGGTGAAATCCGTAGATATCTGTAGGAATACCAATTGTGAAGACAGCTTTCTGGCTCGTAACTGACACTCAGACTCGAAAGCGTGGGG |
| P-asv4831 | GCGAGCGTTGTCCGGAATTACTGGGCGTAAAGGGTGCGTAGGCGGTTTTTTAAGTGGGATGTGAAATCCCTGGGCTCAACTCAGGAATTGCATTCCAAACTGGAAGACTAGAGTGCAGGAGAGGAAAGTGGAAT<br>TCCCAGTGTAGCGGTGAAATGCGTAGAGATTGGGAGGAACACCAGTGGCGAAGGCGACTTTCTGGACTGTAACTGACGCTGATGCACGAAAGCGTGGGG |
| P-asv4832 | TCAAGCGTTACCCGGAATCATTGGGTGTAAAGGGTTTGTAGGTGGATAATTAAGTCAGGTATGAAAGACCGGAGCTCAACTCCGTGTTTGTATTTGAAACTGGTTGTCTAGAATCAGGGAGAGGTAAGCGGAATTC<br>TAAGTGTAGGGGTGCAATCCGTAGATACTTAGAGGAACACCAAAAGCGAAGGCAGCTTACTGGAACTGTATTGACACTGATAAACGAAAGCGTGGGG |
| P-asv4833 | GCGAGCGTTAATCGGAATTACTGGGCGTAAAGCGCATGCAGGCGGTTTATTAAGCCAGATGTGAAAGCCCCGGGCTTAACCTGGGAATTGCATTTGGAACTGGTAAACTAGAGTCTTGTAGAGGGGGGTAGAATT<br>TCAGGTGTAGCGGTGAAATGCGTAGAGATCTGAAGGAATACCGGTGGCGAAGGCGGCCCCCTGGACAAAGACTGACGCTCAGATGCGAAAGCGTGGGG |
| P-asv4834 | GCAAGTGTTATCCGGAATTATTGGGCGTAAAGCGTCTGTAGGTTGATTTATAAGTCTTTTGTTAAATATTTAAGCTTAACTTAAAAAATGCAAAGGAAACTATTTATCTAGAGTTTGGTAGAGGTAAAGGGAATTTCCA<br>GTGAAATGGTGAAATATTTAGATATTGGAAAGAACACCAAGGGCGAAGGCATTTTACTGGGCCATAACTGACACTGAGAGACGAAAGCTAGGGG |
| P-asv4835 | GCGAGCGTTAATCGGAATTACTGGGCGTAAAGCGCGCGTAGGTGGTTAGGTAAGCGAGATGTGAAAGCCCCGGGCTCAACCTGGGAATGGCATTTCGAACTGCTTGGCTAGAGTGTGGTAGAGGGTAGTGGAA<br>TTTCCTGTGTAGCGGTGAAATGCGTAGATATAGGAAGGAACACCAGTGGCGAAGGCGGCTACCTGGACCAACACTGACACTGAGGTGCGAAAGCGTGGTT |
| P-asv4836 | GCAAGCGTTACTCGGAATTACTGGGCGTAAAGCGTGCGTAGGCGGTTCGTTAAGTCTGCTGTGAAAGCCCTGGGCTCAACCTGGGAACTGCAGTGGATACTGGCGAGCTAGAGTGTGTCAGAGGATGGTGGAAT<br>TCCCGGTGTAGCGGTGAAATGCGTAGAGATCGGGAGGAACACCAGTGGCGAAGGCGGCCATCTGGGACAACACTGACGCTGAGGCACGAAAGCGTGGGG |
| P-asv4837 | TCTATCTTTAATCTTATTTACTTTTCTTAAATCTTTCTTAGTCTGCTTTTTAATTCGTATGTGAAATCCCTGAGCTTAACTTAGGAATTGCATTCGATACTGGGAAGCTAGAGTATGGGAGAGGATGGTAGAATTCCAG<br>GTGTAGCGGTGAAATGCGTAGAGATCTGGAGGAATACCGATGGCGAAGGCAGCCATCTGGCCTAATACTGACGCTGAGGTACGAAAGCATGGGG |
| P-asv4838 | GCTAGCGTTGTTCGGATTTACTGGGCGTAAAGCGCACGTAGGCGGACTTTTAAGTCAGGGGTGAAATCCCGGGGCTCAACCCCGGAACTGCCTTTGATACTGGAAGTCTTGAGTATGGTAGAGGTGAGTGGAAT<br>TCCGAGTGTAGAGGTGAAATTCGTAGATATTCGGAGGAACATCAGTGGCGAAGGCGGCTCACTGGACCATTACTGACGCTGAGGTGCGAAAGCGTGGGG |
| P-asv4839 | GCGAGCGTTAATCGGATTTACTGGGCGTAAAGCGTGCGTAGGCGGCTTTTTAAGTCGGATGTGAAATCCCTGAGCTTAACTTAGGAATTGCATTCGATACTGGGAAGCTAGAGTATGGGAGAGGATGGTAGAATT<br>CCAGGTGTAGCGGTGAAATGCGTAGAGATCTGGAGGAATACCGATGGCGAAGGCAGCCTTCTGGCATCATATTGACACTGAGGTTCGAAAGCGTGGGT |
| P-asv4840 | GCAAGCGTTATCCGGAATCATTGGGTTTAAAGGGTCCGCAGGCGGTCAATTAAGTCAGAGGTGAAATCCCATAGCTTAACTATGGAACTGCCTTTGATACTGGTTGACTTGAGTTATACGGAAGTAGATAGAATAA<br>GTAGTGTAGCGGTGAAATGCATAGATATTACTTAGAATACCGATTGCGAAGGCAGTCTACTACGTATATACTGACGCTCATGGACGAAAGCGTGGGG |

| | |
|---|---|
| P-asv4841 | GCTAGCGTTGCTCGGAATCACTGGGCGTAAAGGGTGCGTAGGAGGATCTTTAAGTCAGGGGTGAAATCCTGGAGCTCAACTCCAGTACTGCCTTTGATACTGAGGATCTTGAGTTCGGGAGAGGTAAGTGGAACTGCGAGTGTAGAGGTGAAATTCGTAGATATTCGCAAGAACACCAGTGGCGAAGGCGGCTTACTGGCCCGATACTGACACTGAGGCACGAAAGCGTGGGG |
| P-asv4842 | GCTAGCGTTGTTCGGAATTACTGGGCGTAAAGCGCACGTAGGCGGCTGTTCAAGTCAGAGGTGAAAGCCCGGAGCTCAACTTCGGAACTGCCTTTGAAACTAGACAGCTAGAATCCTGGAGAGGTGAGTGGAATTCCGAGTGTAGAGGTGAAATTCGTAGATATTCGGAAGAACACCAGTGGCGAAGGCGGCTCACTGGACAGGTATTGACGCTGAGGTGCGAAAGCGTGGGG |
| P-asv4843 | GCTAGCGTTGTTCGGATTTACTGGGCGTAAAACGCACGTAGGCGGACTATTAAGTCAGGGGTGAAATCCCGGGGCTCAACCCCGGAACTGCCTTTGATACTGGTAGTCTTGAGTTCGAGAGAGGTGAGTGGAATTCCGAGTGTAGAGGTGAAATTCGTAGATATTCGGAGGAACACCAGTGGCGAAGGCGGCTCACTGGCTCGATACTGACGCTGAGGTGCGAAAGCGTGGGG |
| P-asv4844 | GCAAGCGTTAATCGGAATTACTGGGCGTAAAGCGAGCGTAGGTGGCTTAATAAGTCAGATGTGAAATCCCCGGGCTTAACCTGGGAACTGCATCTGATACTGTTGGGCTAGAGTAGGTGAGAGGGAGGTAGAATTTCAGGTGTAGCGGTGAAATGCGTAGAGATCTGAAGGAATACCGATGGCGAAGGCAGCCTCCTGGCATCATACTGACACTGAGGTTCGAAAGCGTGGGT |
| P-asv4845 | GCGAGCGTTATCCGGAATTATTGGGCATAAAGGGCATCTAGGCGGCCTTTCAAGTCAGGGGTGAAAACCTGCGGCTCAACCGCAGGCCTGCCTTTGAAACTGATAGGCTGGAGTACCGGAGAGGTGGACGGAACTGCACGAGTAGAGGTGAAATTCGTAGATATGTGCAGGAATGCCGATGATGAAGATAGTTCACTGGACGGTAACTGACGCTGAAGTGCGAAAGCCGGGGG |
| P-asv4846 | GCAAGCGTTGTTCGGAATTACTGGGCGTAAAGGGCGAGTAGGTGGCCATCTAAGTCAGACGTGAAATCCCCCGGCTTAACCTGGGAACTGCGTCTGATACTGGACGGCTTGAGTATGGGAGAGGAATGTAGAATTCCAGGTGTAGCGGTGAAATGCGTAGATATCTGGAGGAATACCGGTGGCGAAGGCGGCATTCTGGACCATTACTGACACTGAATCGCGAAAGCTAGGGG |
| P-asv4847 | TCCCTCTTTCCTCGGAATCACTGGGCGTAAAGGGTGCGTAGGCGGTACGTTAGGTCTGTCGTGAAATCCCCGGGCTCAACCTGGGAATGGCGGTAGAAACCGGCGCACTAGAGTATGGGAGAGGGTGGTGGAATTCCAGGTGTAGCGGTGAAATGCGTAGAGATCTGGAGGAACATCAGTGGCGAAGGCGGCCACCTGGCCCAATACTGACGCTGAGGCACGAAAGCGTGGGG |
| P-asv4848 | GCGAGCGTTAATCGGAATTACTGGGCGTAAAGCGTGCGCAGGCGGTTGTGTAAGACAGGTGTGAAATCCCCGGGCTTAACCTGGGAATGGCGCTTGTGACTGCACGGCTAGAGTACGGCAGAGGGGGGTGGAATTCCACGTGTAGCAGTGAAATGCGTAGAGATGTGGAGGAACACCGATGGCGAAGGCAGCCCCCTGGGCCGATACTGACGCTCATGCACGAAAGCGTGGGG |
| P-asv4849 | TCCCTCTTTCCTCGGAATTACTGGGCGTAAAGCGTGCGCAGGCGGTTGTGTAAGACAGGTGTGAAATCCCCGGGCTTAACCTGGGAATTGCATTTGTGACTGCACGACTAGAGTGTGTCAGAGGGGGGTGGAATTCCACGTGTAGCAGTGAAATGCGTAGATATGTGGAGGAACACCAATGGCGAAGGCAGCCCCCTGGGATAACACTGACGCTCATGCACGAAAGCGTGGGG |
| P-asv4850 | GCAAGCGTTGTCCGGAATTATTGGGCGTAAAGAGCTCGTAGGCGGTTTGTCGCGTCTGCTGTGAAAACCCGAGACTCAACCTCGGGCCTGCAGTGGGTACGGGCAGACTAGAGCGCGGTAGGGGAGACTGGACTTCCTGGTGTAGCGGTGGAATGCGCAGACATCAGGAGGAACACCGATGGCGAAGGCAGGTCTCTGGCCCGCAACTGACGCTGAGGAGCGAAAGCATGGGG |
| P-asv4851 | GCAAGCGTTACTCGGAATTACTGGGCGTAAAGCGTGCGTAGGTGGTTTGTTAAGTCAGATGTGAAAGCCCTGGGCTCAACCTGGGAACTGCATTTGATACTGGCAAGCTAGAGTGCGGTAGAGGATAGCGGAATTCCTGGTGTAGCAGTGAAATGCGTAGAGATCAGGAGGAACATCCGTGGCGAAGGCGGCTATCTGGGCCAGCACTGACACTGAGGCACGAAAGCGTGGGG |
| P-asv4852 | GCAAGCGTTAATCGGAATCACTGGGCGTAAAGCGCGCGTAGGTGGATATTTTAGTCGGATGTGAAAGCCCCGAGCTTAACTTGGGAATTGCACCCGATACTGGATATCTAGAGTATAGTAGAGGGAAGTGGAATTTCCGGTGTAGCGGTGAAATGCGTAGATATCGGAAAGAACATCAGTGGCGAAGGCGGCTTCCTGGACTAATACTGACACTGAGGCGCGAAAGCGTGGGG |
| P-asv4853 | GCAAGCGTTAATCGGAATTACTGGGCGTAAAGCGCACGCAGGCGGTCTGTCAAGTCGGATGTGAAATCCCCGGGCTCAACCTGGGAACTGCATCCGAAACTGGCAGGCTAGAGTCTTGTAGAGGGGGGTAGAATTCCAGGTGTAGCGGTAAAATGCGTAGAGATCTGGAGGAATACCGGTGGCGAAGGCGGCTATCTGGCTCACAACTGACGATCAGGCACGAAAGCGTGGGG |
| P-asv4854 | GCAAGCGTTAATCGGAATTACTGGGCGTAAAGGGTGTGTAGGTGGTTTGATAAGTTAGGTGTGAAAGTCCTGGGCTTAACCTGGGGATTGCGCCTAATACTGTCAAGCTAGAGTACTGTAGAGGAAGGTGGAATTTCCGGTGTAGCGGTGAAATGCGTAGATATCGGAAGGAACATCAGTGGCGAAGGCGACCTTCTGGACAGATACTGACACTGAGACACGAAAGCATGGGT |
| P-asv4855 | GCCAGCGTTGCTCGGAATTACTGGGCGTAAAGCGTGTGTAGGCGGCTATGCAAGTCGGATGTGAAAGCCCCAGGCTTAACCTGGGAAGTGCACCCGAAACTGCAGAGCTTGAGTACCGGAGAGGATGGCGGAATTCCCCAAGTAGAGGTGAAATTCGTAGATATGGGGAGGAACACCGGTGGCGAAGGCGGCCATCTGGACGGATACTGACGCTGAGACGCGAAAGCGTGGGG |
| P-asv4856 | GCGAGCGTTGTCCGGAATCACTGGGCGTAAAGGGCGCGTAGGTGGTCTTATAAGGGTGTGGTGAAAGCCCGGGGCTCAACCCCGGGTCGGCCGTGCCGACTGTAAGACTCGAGTGCTGTAGGGGCAGGCGGAATTCCGGGTGTAGCGGTGGAATGCGTAGAGATCCGGAGGAAGACCGGTGGCGAAGGCGGCCTGCTGGGCAGTTACTGACACTGAGGCGCGACAGCGTGGGG |
| P-asv4857 | GCAAGCGTTGTCCGGAATTACTGGGCGTAAAGCGCGCGTAGGCGGCTAGATAAGTCCGGTGTGAAATCCCACGGCTTAACCGGGGAACGGCATCGGGGACTGTTTAGCTTGAGGACGGGAGAGGAGAGCGGAATTCCCGGTGTAGCGGTGAAATGCGTAGAGATCGGGAGGAAGGCCGGTGGCGAAGGCGGCTCTCTAGAACGTCCCTGACGCTGAGGCGCGAAAGCGTGGGG |
| P-asv4858 | GCAGGCGTTAATCGGAATTACTGGGCGTAAAGCGTGCGCAGGCGGCTCTTTTAGTCAGATGTGAAAGCCCCGGGCTTAACCTGGGAATTGCGTTTGATACTGGAGAGCTGGAGTGTGGAAGAGGGGGGTGGAATTCCAGGTGTAGCAGTGAAATGCGTAGATATCTGGAGGAACACCGGTGGCGAAGGCGGCATCCTGGACCGGCATTGACGCTGAGGCGCGAAAGCCAGGGG |
| P-asv4859 | GCAAGCGTTGTCCGGAATTATTGGGCGTAAAGCGCGCGCAGGTGGTGGTTTAAGTCTGATGTGAAAGCCCACGGCTCAACCGTGGAAGGTCATTGGAAACTGGATCACTTGAGTGCAGGAGAGGAAAGTGGAATTCCACGTGTAGCGGTGAAATGCGTAGAGATGTGGAGGAACACCAGTGGCGAAGGCGACTTTCTGGCCTGTAACTGACACTGAGGCGCGAAAGCGTGGGG |
| P-asv4860 | GCGAGCGTTGTCCGGAATCACTGGGCGTAAAGGGCGCGTAGGCGGCGCGGGAAGTCAACGGTGAAAGCTTGGGGCTCAACTCCGAGTCGGCCGTCGATACTCCCGTGCTCGAGCACTGTAGAGGCAGATGGAATTCCGGGTGTAGCGGTGGAATGCGTAGAGATCCGGAAGAACACCGGTGGCGAAGGCGGTCTGCTGGGCAGTTGCTGACGCTGATGCGCGACAGCGTGGGG |
| P-asv4861 | GCAAGCGTTGTCCGGATTTATTGGGCGTAAAGCGCGCGCAGGCGGCTTACTAAGTCTGATGTGAAATCCCACGGCTCAACCGTGGAGGGCCATTGGAAACTGGTAAGCTTGAGTGTAGGAGAGGAGAGCGGAATTCCTGGTGTAGCGGTGAAATGCGTAGATATCAGGAGGAACACCCGTGGCGAAGGCGGCTCTCTGGCCTATAACTGACGCTGAGGTGCGAAAGCGTGGGG |
| P-asv4862 | GCAAGCGTTGTCCGGATTTATTGGGTTTAAAGGGTGCGTAGGCGGCCCTGTAAGTCAGTGCTGAAATATCCCGGCTTAACCGGGAGGGTGGCATTGATACTGCGGGGCTAGAGTACAGATGAGGTAGGCGGAATTGACGGTGTAGCGGTGAAATGCTTAGGTATCGTCAAGAACACCTATAGCGAAGGCAGCTTACTAGGCTGTAACTGACGCTGAGGCACGAAAGTGTGGGG |
| P-asv4863 | GCAAGCGTTGTTCGGAATTACTGGGCGTAAAGCGCGCGCAGGCGGTCTGTTAAGTCTGGTGTGAAATCCCCCCGCTCAACGAGGGAACGGCGCCGGAAACTGGCAAACTAGAGTACCGGAGAGGAAGGTGGAATTCCCGGTGTAGCGGTGAAATGCGTAGATATCGGGAGGAACACCGGTGGCGAAGGCGGCCTTCTGGACGGATACTGACGCTGATACGCGAAAGCGTGGGG |
| P-asv4864 | GCTAGCGTTGTTCGGAATTACTGGGCGTAAAGCGCACGTAGGCGGCTTTGTAAGTTAGAGGTGAAAGCCCGGGGCTCAACCCCGGAATAGCCTTTAAGACTGCATCGCTTGAACGTCGGAGAGGTGAGTGGAATTCCGAGTGTAGAGGTGAAATTCGTAGATATTCGGAAGAACACCAGTGGCGAAGGCAGCTCTCTGGGACGGTACTGACGCTGAGACGCGAAAGCGTGGGG |
| P-asv4865 | GCCAGCGTTGCTCGGAATTACTGGGCGTAAAGGGTCCGTAGGCGGTGCGGCAAGTCGGATGTGAAATCTCTGGGCTTAACCCAGAGGCTGCTTCCGAAACTGCCGCGCTAGAGGATGAGAGAGGCGAGTGGAATTGCGGGTGTAGCGGTGAAATGCGTAGATATCCGCAGGAACATCCGAGGCGAAGGCGGCTCGCTGGATCATTTCTGACGCTGAGGGACGAAAGCTAGGGG |
| P-asv4866 | GCTAGCGTTGTTCGGAATTACTGGGCGTAAAGCGCGTGTAGGCGGCCCGCTAAGTCTGATGTGAAAGCCCGGGGCTCACCCCCGGAAGTGCATTGGAGACTGGCGGGCTAGAGTCCCGGAGAGGAGGGTGGAATTCCTGGTGTAGCGGTGAAATGCGTAGATATCAGGAGGAACATCAGTGGCGAAGGCGGCCCTCTGGTCCAGTACTGACACTCAAGTGCGAAAGCGTGGGG |
| P-asv4867 | GCGAGCGTTGTCCGGAATTATTGGGCGTAAAGGGCTCGTAGGCGGTCTGTCGCGTCAGGAGTGAAAACTCGGGGCTCAACCCCGAGCCTGCTTCTGATACGGGCAGACTAGAGGTATGCAGGGGAGAACGGAATTCCTGGTGTAGCGGTGAAATGCGCAGATATCAGGAGGAACACCGGTGGCGAAGGCGGTTCTCTGGGCATGATCTGACGCTGAGGAGCGAAAGTGTGGGG |
| P-asv4868 | GCGAGCGTTAATCGGAATTACTGGGCGTAAAGCGTGCGTAGACGGTTCCGTAAGTCGGGTGTGAAAGCCCCGGGCTCAACCTGGGAATTGCATTCGAGACTGCGAAGCTAGGGTGCGGAAGAGGGAAACGGAATTTCTGGTGTAGCGGTGAAATGCGTAGATATCAGAAGGAGCATCAGTGGCGAAAGCGGCTTCCTGGTCCAGCGCCGACGTTCAGGCACGAAAGCGTGGGG |
| P-asv4869 | GCGAACGTTATTCGGAATCACTGGGCTTAAAGGGCGCGTAGACGGGCGACCGCGTCGATGGCTGAAAGCCCCCGGCCCAACCGGGGAACGGGCCTCGATACGGGTCGCCTCGAGTGGAGGAGGGGGACCTGGAACTGCCGGTGGAGCGGTGAAATGCGTTGAGATCGGCAGGAACGCCTGTGGCGAAAGCGAGGTCCTGGACTCCCACTGACGTTGAGGCGCGAAAGCCAGGGG |
| P-asv4870 | GCGAGCGTTAATCGGAATTACTGGGCGTAAAGCGTGCGTAGACGGTTTCGTAAGTCGGGTGTGAAATCCCCGGGCTCAACCTGGGAATTGCATTCGAAACTGCGAGGCTAGGGTACGGAAGAGGGAAGCGGAATTTCTGGTGTAGCGGTGAAATGCGTAGATATCAGAAGGAACATCAGTGGCGAAAGCGGCTTCCTGGTCCAGTACCGACGTTCAGGCACGAAAGCGTGGGG |
| P-asv4871 | GCGAGCGTTGTCCGGAATCACTGGGCGTAAAGGGCGCGTAGGCGGCTTGGTAAGCGTGCGGTGAAAGCTCGAGGCTCAACCTCGAGTCTGCCGTGCGAACTGCCGAGCTGGAGCACGGTAGAGGCACATGGAATTCCGGGTGTAGCGGTGGAATGCGTAGAGATCCGGAAGAACACCGGTGGCGAAGGCGGTGTGCTGGGCCGTAGCTGACGCTGAGGCGCGACAGCGTGGGG |
| P-asv4872 | GCGAGCGTTGTTCGGATTTATTGGGCGTAAAGAGCGTGTAGGCGGCCGTACAGGTCAGCTGTGAAAACTCGAGGCTCAACCTCGAGACGTCGGTTGAAACCGTACGGCTAGAGTCCGGAAGAGGAGAGCGGAATTCCCGGTGTAGCGGTGAAATGCGCAGATATCGGGAAGAACACCCGTGGCGAAGGCGGCTCTCTGGGACGGTACTGACGCTGAGACGCGAAAGCGTGGGG |
| P-asv4873 | GCTAGCGTTGTTCGGAATTACTGGGCGTAAAGCGTACGTAGGCGGCTTTGTAAGTTAGAGGTGAAAGCCCGGGGCTCAACCCCGGAATTGCCTTTAAGACTGCATCGCTCGAATCATGGAGAGGTGAGTGGAATTCCGAGTGTAGAGGTGAAATTCGTAGATATTCGGAAGAACACCAGTGGCGAAGGCGACTCACTGGACATGTATTGACGCTGAGGTACGAAAGCGTGGGG |
| P-asv4874 | GCTAGCGTTGCTCGGAATCACTGGGCGTAAAGGGTGCGTAGGCGGGTCTTTAAGTCAGGGGTGAAATCCTGGAGCTCAACTCCAGAACTGCCTTTGATACTGAAGATCTTGAGTTCGGGAGAGGTGAGTGGAACTGCGAGTGTAGAGGTGAAATTCGTAGATATTCGCAAGAACACCAGTGGCGAAGGCGGCCCTCTGGCCCGATACTGACGCTGAGGTGCGAAAGCGTGGGG |
| P-asv4875 | GCTAGCGTTGCTCGGAATCACTGGGCGTAAAGCGCACGTAGGCGGCTTCTTAAGTCAGGGGTGAAATCCTGGAGCTCAACTCCAGAACTGCCTTTGATACTGAGGAGCTTGGAGTCCGGGAGAGGTGAGTGGAACTGCGAGTGTAGAGGTGAAATTCGTAGATATTCGCAAGAACACCAGTGGCGAAGGCGGCTCACTGGCCCGGTACTGACGCTGAGGTGCGAAAGCGTGGGG |
| P-asv4876 | GCGAGCGTTGTTCGGAATTATTGGGCGTAAAGGGCGCGTAGGCGGTACGGTAAGTCACCTGTGAAACCTCTGGGCTTAACTCAGAGCCTGCAGGCGAAACTGCCGTGCTGGAGTGTGGGAGAGGTGCGTGGAATTCCCGGTGTAGCGGTGAAATGCGTAGATATCGGGAGGAACACCTGTGGCGAAAGCGGCGCACTGGACCACAACTGACGCTGAGGCGCGAAAGCTAGGGG |
| P-asv4877 | GCGAGCGTTGTCCGGAATCACTGGGCGTAAAGGGCGCGTAGGCGGCCTGATAAGTAGGGGGTGAAATCCTGCGGCTTAACCGCAGGGCTGCCTTCTAGACTGTCAGGCTAGAGCACAGTAGAGGCAGGTGGAATTCCCGGTGTAGCGGTGGAATGCGTAGAGATCGGGAAGAACGTCAGTGGCGAAGGCGGCCTGCTGGGCTGTTGCTGACGCTGAGGCGCGACAGCGTGGGG |
| P-asv4878 | GCTAGCGTTGCTCGGAATTACTGGGCGTAAAGGGCGCGTAGGCGGCTTGGCCAGTCAGGCGTGAAATTCCTGGGCTTAACCTGGGGACTGCGCTTGATACTGCCAGGCTTGAGGCCGAGAGAGGGTGGTGGAATTCCCAGTGTAGAGGTGAAATTCGTAGATATTGGGAAGAACACCGGTGGCGAAGGCGACCACCTGGCTCGGTACTGACGCTGAGGCGCGACAGCGTGGGG |
| P-asv4879 | GCTAGCGTTATTCGGAATTATTGGGCGTAAAGGGCGCGTAGGTGGCATGTTAAGTTTGCTGTTAAATCTTCCAGCCTAACTGGAAAACTGCAGCAAAAACTGGCAAGCTAGAGTGTAAGAGAGAGAAGTGGAATTCTCGGAGTAGCGATAAAATGCGTAGATCTCGAGAGGAACACCGATGGCGAAGGCAGCTTCTTGGCTTACTACTGACACTGAGGCGCGAAAGCGTGGGG |

| | |
|---|---|
| P-asv4880 | GCAAGCGTTATCCGGATTTATTGGGTTTAAAGGGTCCGTAGGCGGACTTATAAGTCAGTGGTGAAATCCTGTCGCTTAACGATAGAACTGCCATTGATACTGTAAGTCTTGAGTATATTTGAGGTAGCTGGAATAAGTAGTGTAGCGGTGAAATGCATAGATATTACTTAGAACACCAATTGCGAAGGCAGGTTACCAAGATATAACTGACGCTGAGGGACGAAAGCGTGGGG |
| P-asv4881 | GCGAGCGTTGTCCGGAATTATTGGGCGTAAAGCGCGCGCAGGCGGTTTCTTAAGTCTGATGTGAAATCTCGCGGCTCAACCGCGAGCGGTCATTGGAAACTGGGGAACTTGAGTACAGAAGAGGAGAGTGGAATTCCACGTGTAGCGGTGAAATGCGTAGATATGTGGAGGAACACCAGTGGCGAAGGCGACTCTCTGGTCTGTAACTGACGCTGAGGCGCGAAAGCGTGGGG |
| P-asv4882 | GCAAGCGTTGTCCAGAATTATTGGGCGTAAAGCGCGCACAGGCGGTCCTTTAAGTCTGATGTGAAAGCCCACGGCTCAACCGTGGAGGGTCATTGGAAACTGGGGGACTTGAGTGCAGAAGAGAAGAGTAGAATTCCACATGTAGCGGTGAAATGCATAGAGATGTGGAGGAACACCAGTGGCGAAGGCGACTCTTTGGTCTGTAACTGACGCTGAGGCGCGAAAGCGTGGAA |
| P-asv4883 | GCGAGCGTTAATCGGAATTACTGGGCGTAAAGCGCGCGTAGGCGGTCTGATAAGCCGGTTGTGAAAGCCCCGGGCTCAACCTGGGAACGGCATCCGGAACTGTCAGGCTAGAGTGCAGGAGAGGAAGGTAGAATTCCCGGTGTAGCTGTGAAATGCGTAGAGATCGGGAGGAATACCAGTGGCGAAGGCGGCCTTCTGGACTGACACTGACGCTGAGGTGCGAAAGCGTGGGT |
| P-asv4884 | GCTAGCGTTGTTCGGAATTACTGGGCGTAAAGCGCACGTAGGCGGCTTTGCAAGTCAGAGGTGAAAGCCCGGGGCTCAACCCCGGAATTGCCTTTGAGACTGCATGGCTAGAATTGTGGAGAGGTAAGTGGAATTCCGAGTGTAGAGGTGAAATTCGTAGATATTCGGAAGAACACCAGTGGCGAAGGCGACTTACTGGACACATATTGACGCTGAGGTGCGAAAGCGTGGGG |
| P-asv4885 | GCTAGCGTTGTTCGGAATCATTGGGCGTAAAGGGCGTGTAGGCGGTTTGCTAAGTCATGTGTGAAATCCCTCGGCTCAACCGGGGAACGACGCATGAAACTGACAAGCTAGAGTACCAAAGAGGGGGGTGGAATTCCCGGTGTAGCGGTGAAATGCGTAGATATCGGGAGGAACACCGGTGGCGAAGGCGGCCCCCTGGTTGGATACTGACGCTGAGACGCGAAAGCGTGGGG |
| P-asv4886 | GCAAGCGTTGTTCGGAATTACTGGGCGTAAAGGGCGCGTAGGCGGCCTTGTGAGTCAGACGTGAGATCCCCAGGCTTAACTTGGGAACTGCGTCTGATACTGCAAGGCTCGAGTGCGGGAGAGGGATGTGGAATTCCAGGTGTAGCGGTGAAATGCGTAGATATCTGGAGGAACACCGGTGGCGAAGGCGGCATCCTGGACCGACACTGACGCTGAGGCGCGAAAGCTAGGGG |
| P-asv4887 | GCAAGCGTTATCCGGATTTATTGGGTTTAAAGGGTGCGTAGGCGGTATTACAAGTCAGGGGTGAAATCTTGGTGCTTAACATTAAAATTGCCTTTGAAACTGTGGTACTTGAGTGTAAAAGAGGTAGGCGGAATGTGTTGTGTAGCGGTGAAATGCATAGATATAACACAGAACCCCGATTGCGAAGGCAGCTTACTATCATACAACTGACGCTGATGCACGAAAGCGTGGGG |
| P-asv4888 | GCTAGCGTTGCTCGGAATCACTGGGCGTAAAGGGTGCGTAGGCGGGTCTTTAAGTCAGAGGTGAAATCCTGGAGCTCAACTCCAGAACTGCCTTTGATACTGAGGATCTTGAGTCCGGAAGAGGTGAGTGGAACTGCGAGTGTAGAGGTGAAATTCGTAGATATTCGCAAGAACACCAGTGGCGAAGGCGGCTCACTGGTCCGGTACTGACGCTGAGGCACGAAAGCGTGGGG |
| P-asv4889 | GCTAGCGTTGTTCGGAATTACTGGGCGTAAAGCGCACGTAGGCGGCTATTCAAGTCAGAGGTGAAAGCCTGGAGCTTAACTCCAGAACTGCCTTTGAAACTAGATAGCTAGAATCATGGAGAGGTGAGTGGAATTCCGAGTGTAGAGGTGAAATTCGTAGATATTCGGAAGAACACCAGTGGCGAAGGCGGCTCACTGGACATGTATTGACGCTGAGGTGCGAAAGCGTGGGG |
| P-asv4890 | CCAAGCGTTATCCGGAATCATTGGGTTTAAAGGGTCCGTAGGCGGTTTTGTAAGTCAGTGGTGAAATCTGGTCGCTTAACGATCAAACGGCCATTGATACTGCAAGACTTGAATTATTGGGAAGTAACTAGAATATGTAGTGTAGCGGTAAAATGCTTAGATATTACATGGAATACCAATTGCGAAGGCAGGTTACTATCAATATATTGACGCTGATGGACGAAAGCGTGAAG |
| P-asv4891 | TCCCTCTTTTTCCTTCCTTCTTGGGCGTAAAGGGCTCGCAGGCGGTTTCTTAAGTCTGATGTGAAAGCCCCCGGCTCAACCGGGGAGGGTCATTGGAAACTGGGAAACTTGAGTGCAGAAGAGGAGAGTGGAATTCCACGTGTAGCGGTGAAATGCGTAGAGATGTGGAGGAACACCAGTGGCGAAGGCGACTCTCTGGTCTGTAACTGACGCTGAGGAGCGAAAGCGTGGGG |
| P-asv4892 | TCAAGTGATGTCCCATATTATTGGGCCTAAAGCGTCCGTAGCTGGCCAACCAAGTCTATCGGGAAATCCACCTGCCCAACAGGTGGGCGTCCGGTAGAAACTGGCTGGCTTGGAACCGGAAGGCTCAGAGAGTACGTCCGCGGTAGGAGTGAAATCCCGTAATCCCGGACGGACTACCGATGGCGAAAGCACTCTGAGAAGACGGATTCGACAGTGAGGGACGAAAGCTCGGGT |
| P-asv4893 | GCTAGCGTTGTTCGGAATCACTGGGCGTAAAGCGCACGTAGGCGGACCATTAAGTCAGGGGTGAAAGCATGGAGCTCAACTCCAGAACTGCCTTTGATACTGATGGTCTCGAGTTCGGAAGAGGTTGGTGGAACTGCGAGTGTAGAGGTGAAATTCGTAGATATTCGCAAGAACACCAGTGGCGAAGGCGGCCAACTGGTCCGATACTGACGCTGAGGTGCGAAAGCGTGGGG |
| P-asv4894 | GCAAGCGTTAATCGGAATTACTGGGCGTAAAGCGCACGTAGGCGGTTTGTTAAGTTGGATGTGAAAGCCCCGGGCTCAACCTGGGAATTGCATTCAAGACTGGCAAGCTAGAGTATGTGAGAGGGGGGTAGAATTCCAAGTGTAGCGGTGAAATGCGTAGAGATTTGGAGGAATACCAGTGGCGAAGGCGGCCCCCTGGCACAATACTGACGCTCAGGTGCGAAAGCGTGGGG |
| P-asv4895 | CCGAGCGTTGTCCGGATTTATTGGGCGTAAAGCGAGCGCAGGCGGTTTGATAAGTCTGAAGTAAAAGGCTGTGGCTTAACCATAGTATGCTTTGGAAACTGTTAAACTTGAGTGCAGAAGGGGAGAGTGGAATTCCATGTGTAGCGGTGAAATGCGTAGATATATGGAGGAACACCGGTGGCGAAAGCGGCTCTCTGGTCTGTAACTGACGCTGAGGCTCGAAAGCGTGGGT |
| P-asv4896 | GCAAGCGTTATCCGGAATTATTGGGCATAAAGGGCATCTAGGCGGCCCTGTAAGTCTAGGGTGAAAACCTGCGGCTCAACCGCAGGCCTGCCCCGGAAACTACAGGGCTAGAGTATCGGAGAGGTGGACGGAACTGCACGAGTAGAGGTGAAATTCGTAGATATGTGCAGGAATGCCGATGATGAAGATAGTTCACTGGACGGTAACTGACGCTGAAGTGCGAAAGCTAGGGG |
| P-asv4897 | ACGAGCGTTATCCGGATTTATTGGGTTTAAAGGGTGCGTAGGTGGCTTTATAAGTCAGTGGTGAAAAGCTGTGGCTCAACCATAGTCTTGCCGTTGAAACTGTGGAGCTTGAGTGCGGATGAGGTAGGCGGAACGCGTAGTGTAGCGGTGAAATGCATAGATATTACGCAGAACTCCGATTGCGAAGGCAGCTTACCAAACCGCAACTGACACTGAAGCACGAAAGCGTGGGT |
| P-asv4898 | TCAAGTGATGTCCAATATTATTGGGCCTAAAGCGTCCGTAGCTGGCCAGGCAAGTCTATCGGGAAATCCACCTGCCCAACAGGTGGGCGTCCGGTAGAAACTGCCCGGCTTGGAACCGGAAGGCTCAGAGAGTACGTACGGGGTAGGAGTGAAATCCCGTAATCCTGAACGGACCACCGATGGCGAAAGCACCTCGAGAGGACTGATCCGACAGTGAGGGACGAAAGCTAGGGT |
| P-asv4899 | GCAAGCGTTGTCCGGAATTATTGGGCGTAAAGAACTCGTAGGCGGTTTGTCGCGTCTGCTGTGAAAACGCGAGGCTCAACCCCGCGCCTGCAGTGGGTACGGGCAGACTAGAGTGCGGTAGGGGAGATTGGAATTCCTGGTGTAGCGGTGGAATGCGCAGATATCAGGAGGAACACCGATGGCGAAGGCAGATCTCTGGGCCGTAACTGACGCTGAGGAGCGAAAGCGTGAGG |
| P-asv4900 | ACAAGCGTTGTCCGGAATTATTGGGCGTAAAGAGCTCGTAGGTGGTCTGTCACGTCTGCTGTGGAAACGCAAGGCTCAACCTTGCGCGTGCAGTGGGTACGGGCGGACTAGAGTGCAGTAGGGGAGTCTGGAATTCCTGGTGTAGCGGTGAAATGCGCAGATATCAGGAGGAACACCGGTGGCGAAGGCGGGACTCTGGGCTGTAACTGACGCTGAGGAGCGAAAGCATGGGG |
| P-asv4901 | TCGAGTGATAGCCACTTTTATTGGGCCTAAAGCGTCCGTAGCCGGCCGAACGGGTCCGTCGGGAAATCCACTCGCCCAACGAGTGGACGTCCGGCGGAAACCAGTCGGCTTGGGGCCGGGAGACCAGAGAGGTACGTCCGGGGTAGGAGTGAAATCCTGTAATCCTGGACGGACCGCCGGTAGCGAAAGCGTCTCTGGAGAACGGACCCGACGGTGAGGGACGAAAGCTTGGGT |
| P-asv4902 | GCAAGCGTTAATCGGAATTACTGGGCGTAAAGCGCGCGTAGGCGGTTTGTTAAGTCGGATGTGAAAGCCCCGGGCTCAACCTGGGAATTGCATTCGATACTGGTCGACTAGAGTACGAGAGAGGGAGGTAGAATTCCACGTGTAGCGGTGAAATGCGTAGATATGTGGAGGAATACCGGTGGCGAAGGCGGCCTCCTGGCTCGATACTGACGCTGAGGTGCGAAAGCGTGGGG |
| P-asv4903 | GCAAGCGTTATCCGGATTCACTGGGTTTAAAGGGAGCGTAGGCGGGCAGGTAAGTCAGTGGTGAAATCTCCGGGCTTAACCCGGAAACTGCCATTGATACTATCTGTCTTGAATATCCTGGAGGTAAGCGGAATATGTCATGTAGCGGTGAAATGCTTAGATATGACATAGAACACCCATTGCGAAGGCAGCTTACTACGGGATGATTGACGCTGAGGCTCGAAAGCGTGGGG |
| P-asv4904 | GCTAGCGTTGTTCGGAATTACTGGGCGTAAAGCGCGCGTAGGCGGGCTGGTCAGTCAGGGGTGAAAGCCCGAGGCTTAACCTCGGAACTGCCTTTGATACTGCCAGTCTAGAGTCCGGCAGAGGTGAGTGGAATTCCTAGTGTAGAGGTGAAATTCGTAGATATTAGGAAGAACACCAGTGGCGAAGGCGGCTCACTGGGCCGAGACTGACGCTGAGGCGCGAAAGCGTGGGG |
| P-asv4905 | GCAAACGTTGTTCGGAATTACTGGGCTTAAAGCGCACGTAGGCGGCGCCGCAAGTGTGGGGTGAAAGCCCACGGCTTACCCGTGGAATTGCCTGGCAGACTGCGGTGCTAGAGTGTGGTAGGGGAGAGCGGAACTCCTGGTGGAGCGGTGAAATGCGTAGATATCAGGAGGAACACCGGTGGCGAAGGCGGCTCTCTGGTCCACAACTGACGCTGAGGTGCGAAAGCTAGGAG |
| P-asv4906 | GCTAGCGTTGCTCGGAATTACTGGGCGTAAAGGGAGCGTAGGCGGACTGTTTAGTCAGAGGTGAAAGCCCAGGGCTCAACCTTGGAATTGCCTTTGATACTGGCAGTCTTGAGTACGGAAGAGGTATGTGGAACTCCGAGTGTAGAGGTGAAATTCGTAGATATTCGGAAGAACACCAGTGGCGAAGACGACATACTGGTCCGTTACTGACGCTGAGGCTCGAAAGCGTGGGG |
| P-asv4907 | GCAAGCGTTAATCGGAATTACTGGGCGTAAAGCGTGCGCAGGCGGCTTTGTAAGACGGATGTGAAAGCCCCGGGCTCAACCTGGGAAGTGCATTCGTGACTGCAAGGCTAGAGTGTGTCAGAGGGAGGTGGAATTCCGCGTGTAGCAGTGAAATGCGTAGAGATGCGGAGGAACACCGATGGCGAAGGCAGCCTCCTGGGATAACACTGACGCTCAGGCACGAAAGCGTGGGG |
| P-asv4908 | GCAAGCGTTGTCCGGAATTATTGGGCGTAAAGAGCTCGTAGGCGGTTTGTCGCGTCTGGTGTGAAAACTCGAGGCTCAACCTCGAGCTTGCATCGGGTACGGGCAGACTAGAGTGCGGTAGGGGAGACTGGAATTCCTGGTGTAGCGGTGGAATGCGCAGATATCAGGAGGAACACCGATGGCGAAGGCAGGTCTCTGGGCCGCAACTGACGCTGAGGAGCGAAAGCATGGGG |
| P-asv4909 | GCAAGCGTTACCCGGAATTACTGGGCGTAAAGGGAGCGTAGGCGGGAGGTCAAGTCCGGTGTGAAAGACCCTGGCTCAACCAGGGATGAGGGCTGGAAACTGACAATCTTGAGTACAGCAGAGGAAGAGGGAATTACCGGTGTAGGGGTGAAATCCGTAGAGACCGGTAAGAAGGCCAGTGGCGAAGGCGCTTTTCTGGGCTGTTACTGACGCTGAGGCTCGAAAGCCAGGGG |
| P-asv4910 | GCAAGCGTTGTTCGGAATCACTGGGCTTAAAGCGTGCGTAGGCGGACTTACAAGTCAGAAGTGAAATCCCCGGGCTTAACCCGGGAACTGCTTTTGAAACTGTAAGTCTAGAATCCGGCAGGGGTTGGGAGAATTCCTAGTGTAGAGGTGAAATTCGCAGATATTAGGAGGAATACCAGTGGCGTAGGCGCCCAACTGGACCGGTATTGACGCTGAGGTACGAAAGCGTGGGG |
| P-asv4911 | CCGAGTGGTCGGGACGATTATTGGGCCTAAAGCATCCGTAGCTTGTTCTGCAAGTCCTCCGTTAAATCCACCTGCTCAACAGATGGGCTGCGGAGGATACTACAGGGCTAGGAGGCGGGAGAGGCAAGCGGTACTCAGTGGGTAGGGGTAAAATCCTCTGATCCACTGAAGACCACCAGTGGCGAAGGCGGCTTGCCAGAACGCGCTCGACGGTGAGGGATGAAAGCTGGGGG |
| P-asv4912 | GCGAGCGTTGTTCGGAATCACTGGGCGTAAAGGGTGCGCAGGCGGTCATGAAAGTTAGGAGTGAAATCCCCGGGCTTAACCCGGGAATTGCTCTTAAAACTCCATGACTGGAGTACTGGAGAGGTTAGCGGAATTCCGAGTGTAGCAGTGAAATGCGTAGATATTCGGAGGAACACCAATGGCGTAGGCAGCTAACTGGACAGTAACTGACGCTCATGCACGAAAGCGTGGGG |
| P-asv4913 | GCTAGCGTTATTCGGAATCACTGGGCGTAAAGAGCACGTAGGCGGCTTGGAAAGCCAGGGGTGAAAGCCCGGAGCTTAACTCCGGAATTGCCTTTGGAACTATCAAGCTAGCATCTGGCAGAGGGTGGTGGAATTCCTAGTGTAGAGGTGAAATTCGTAGATATTAGGAGGAACACCGGTGGCGAAGGCGACCACCTGGGCCAGTATGGACGCTGAGGTGCGAAAGCGTGGGG |
| P-asv4914 | GCTAGCGTTGTTCGGAATCACTGGGCGTAAAGCGCACGTAGGCGGATTTGTTAGTCAGGGGTGAAATCCCGGGGCTCAACCTCGGAACTGCCTCTGATACTGCAAGTCTCGAGTCCGAGAGAGGTGAGTGGAATTCCTAGTGTAGAGGTGAAATTCGTAGATATTAGGAAGAACACCAGTGGCGAAGGCGGCTCACTGGCTCGGTACTGACGCTGAGGTGCGAAAGCGTGGGG |
| P-asv4915 | GCAAGCGTTGTTCGGAATTACTGGGCGTAAAGAGCACGTAGGCGGGTTCGTCAGCCATTTGTGAAAGCCCAAGGCTCAACCTTGGAATTGCAGGTGGAACTGCGAATCTTGAATACCTGAGAGGATGAGGGAATTCCTGGTGTAGAAGTGAAATCCGTAGATATCAGGAGGAACACCCGAGGCGAAGGCGCTCGTCTGGCAGTGTATTGACGCTGGGGTGCGAAAGCGTGGGG |
| P-asv4916 | GCAAGCGTTGTCCGGAATTATTGGGCGTAAAGCGTTCGTAGGCGGCATGTCAAGTCTGATGTTAAATCCCAGAGCTCAACTCTGGATCGGCATTGGATACTGGCAAGCTAGAGTGCAGAAGAGGCAAGGGGAATTCCTAGTGTAGCGGTGAAATGCGTAGATATTAGGAGGAACACCGGTGGCGTAAGCGCCTTGCTGGGCTGTAACTGACGCTGAGGGACGAAAGCCAGGGG |
| P-asv4917 | GCTAGCGTTGTTCGGAAATACTGGGCGTAAAGCGAACGTAGGCGGTCTATTAAGTCAGGGGTGAAATCCCAGAGCTCAACTCTGGAACTGCCCTTGAAACTGCTAGACTAGAATCCTGGAGAGGTCAGTGGAATTCCGAGTGTAGAGGTGAAATTCGTAGATATTCGGAAGAACACCAGTGGCGAAGGCGACTGACTGGACAGGTATTGACGCTGAGGTTCGAAAGCGTGGGG |
| P-asv4918 | CCAAGCATTATCCGGATTCACTGGGCGTAAAGGGTGCGTAGGCGGTGTGATTAGTCAGGTGTTAAATCCTGGGGCTTAACCTCAGAATCGCATTTGAAACGGTCATACTAGAAGAAGTCAGAGGTAAACAGAACTCTCGGTGTAGGGGTGAAATCCGTTGATATCGAGGGGAATACCAAATGCGAAGGCAGTTTACTGGGACTTTCTTGACGCTGAGGCACGAAAGCATGGGT |

| | |
|---|---|
| P-asv4919 | GCAAGCGTTGTTCGGAATTACTGGGCGTAAAGAGCGCGTAGGCGGTCTACCAAGTCAGACGTGAAAGCCCTGGGCTCAACCCGGGAACTGCGTTTGAAACTGGTAAACTTGAGTGTCAGAGGGGTGAGTGGAATTCCTAGTGTAGAGGTGAAATTCGTAGATATTAGGAGGAACACCAGTGGCGAAGGCGACTCACTGGCTGAATACTGACGCTAAGGCGCGAAAGCGTGGGG |
| P-asv4920 | GCGAGCGTTGTCCGGATTTATTGGGTTTAAAGGGTGCGTAGGCGGCCTATTAAGTCAGTGCTGAAATACCCCGGCTTAACCGGGGGGGTGGCATTGATACTGATAGGCTTGAGTACAGATGAGGTAGGCGGAATTGACGGTGTAGCGGTGAAATGCATAGATATCGTCAAGAACACCGATAGCGAAGGCAGCTTACTAAGCTGTAACTGACGCTGATGCACGAAAGTGTGGGG |
| P-asv4921 | CCAAGCGTTGTCCGGATTTATTGGGTTTAAAGGGTGCGTAGGCGGGATATTAAGTCAGTGGTGAAAGCCTACAGCTTAACTGTAGAATTGCCTCTGAAACTGGTATCCTTGAGTGTGGTTGAGGTGGATGGAATGTATCATGTAGCGGTGAAATGCTTAGATATGATACAGAACACCGATTGCGAAGGCAGTCCACTAAGCCATTACTGACGCTGAGGCACGAAAGCATGGGG |
| P-asv4922 | GCAAGCGTTATCCGGATTCATTGGGTTTAAAGGGTGCGCAGGTGGGAATGTAAGTCAGTGGTGAAATACTGCCGCTTAACGGTAGAGCTGCCATTGATACTGCGTTTCTTGAGTATGGTTGAGGTAGGTGGAATGTGTTGTGTAGCGGTGAAATGCATAGATATGACACAGAACGCCGATTGCGAAGGCAACTTACTAAGCCATTACTGACGCTGAGGCACGAAAGCGTGGGT |
| P-asv4923 | GCAAGCGTTAATCGGAATTACTGGGCGTAAAGCGTGCGCAGGCGGCCTATTAAGTCAGATGTGAAATCCCCGAGCTCAACTTGGGAACTGCATTTGAAACTGATAGGCTAGAGTGTGTCAGAGGGAGGTAGAATTCCACGTGTAGCAGTGAAATGCGTAGAGATGTGGAGGAATACCAGTGGCGAAGGCGGCCTCCTGGGATAACACTGACGCTCATGTACGAAAGCGTGGGG |
| P-asv4924 | GCAAGCGTTATTCAGATTCACTGGGCGTAAAGGGTGCGCAGGTGGCCAAGTGTGTGAGGCGTGAAAGCCTGGGGCTTAACTCCAGAATTGCACCTCAAACTACATGGCTAGAGCATTGGAGAGGGTAGCAGAATTCATGGTGTAGCAGTGAAATGCGTAGATATCATGAGGAATACCAGAGGCGAAGGCGGCTACCTGGACAATTGCTGACACTCAGGCACGAAAGCGTGGGG |
| P-asv4925 | TCAAGCGTTATCCGGAATCATTGGGTTTAAAGGGTTCGTAGGCGGGACAATAAGTCAGTGGTGAAAGTTTGTGGCTCAACCATAAAATTGCCATTGATACTGTTGTTCTTGAGTTATTGTGAAGTGGTTAGAATGTGTAGTGTAGCGGTGAAATGCATAGATATTACACAGAACACCAATTGCGAAAGCATATCACTAACAATATACTGACGCTGATGAACGAAAGCGTAGGT |
| P-asv4926 | GCTAGCGTTGTTCGGAATTACTGGGCGTAAAGCGCACGTAGGCGGACGGTCAAGTTGGGGGTGAAAGCCCGAGGCTCAACCTCGGAATTGCCTTCAAAACTGATCGTCTCGAGGCCGGGAGAGGGCAGTGGAATTCCCAGTGTAGAGGTGAAATTCGTAGATATTGGGAAGAACACCGGTGGCGAAGGCGGCTGCCTGGCCCGGACCTGACGCTGAGGTGCGAAAGCGTGGGG |
| P-asv4927 | CCGAGTGATGGCCGATCTTATTGGGCCTAAAGCGTCCGTAGCCAGCCGAACAGGTCCGCCGGGAAATCTGCTCGCTCAACGAGCAGGCGTCCGGCGGAAACCAGTCAGCTTGGGACCGAGAGGACCGAAGGGTACGTCTGGGGTAGGAGTGAAATCCCGTAATCCTAGACGGACCACCGATGGCGAAAGCACTTCGGTAGATCGGATCCGACGGTGAGGGACGAAAGCTAGGGT |
| P-asv4928 | GCAAGCGTTACTCGGAATCACTGGGCGTAAAGGACGCGTAGGCGGGATGTCAAGTCTGATGTGAAATCCTATGGCTCAACCATAGAACTGCATTGGAAACTGGCAACCTAGAGTATGGGAGGGGGAGATGGAATTAGTGGTGTAGGGGTAAAATCCGTAGATATCACTAGGAATACCGAAAGCGAAGGCGATCTCCTGGAACATAACTGACGCTAAGGCGTGAAAGCGTGGGG |
| P-asv4929 | GCGAGTGTTACTCATAATAACTGGGCGTAAAGGGTACGTAGATGGTTTATATGGTTACTTTGTGAAAGAATAATGTAATGGTTATTTTGCTAGTAATACGTGTAAACTAGAGTATGACAGAGGAAAGTGGAATTTTTAATGTAAAGGTGGTATTTATCGATATTAAAAAGACGATCTAAGGCGAAGGCAACTTTCTGGGTTATTACTGACATTGAGGTACTGAAGCATGGGG |
| P-asv4930 | ACGAGTGATGGCCGATCTTATTGGGCCTAAAGCGTCCGTAGCCGGCCGGGCAAGTCCGTTGGGAAATCCACGCGCTCAACGCGTGGGCGTCCGGCGGAAACTGTTCGGCTTGGGGCCAGAGGACTCGAGGGGTACGTCCGGGGTAGGAGTGAAATCCCGTAATCCTGGACGGACCACCGGTGGCGAAAGCGCCTCGAGAAGATGGACCCGACGGTGAGGGACGAAAGCTTGGGT |
| P-asv4931 | GCAAGCGTTAATCGGAATTACTGGGCGTAAAGGGCTCGTAGGCGGCTGACTAAGTCGGATGTGAAATCCCTGGGCTTAACCTAGGACTGTCATTCGGGACTGGTCGGCTAGAGTTGGATAGAGGAAAGCGGAATTTCCGGTGTAGCGGTGAAATGCGTAGATATCGGAAGGAACACCAGTGGCGAAGGCGGCTTTCTGGATCCAAACTGACGCTGAGGAGCGAGAGCGTGGGT |
| P-asv4932 | GCAAGCGTTGTTCGGATTTATTGGGCGTAAAGCGCGCGCAGGCGGATTAATCAGTCAGGTGTGAAATCTCGGGGCCTAACCTCGAAACTGCGCCTGAAACTATTAATCTAGAATTGTCTAGAGGGAAGGGGAATTTCACGTGTAGGGGTAAAATCCGTAGAGATGTGAAGGAACACCAGAGGCGAAGGCGCCTTCCTGGAGACCAATTGACGCTGAGGCGCGAAAGCGTGGGG |
| P-asv4933 | GCAAGCGTTGTTCGGAATTACTGGGCGTAAAGCGTACGTAGGCGGCGGGTTAAGTCGAATGTTAAAGCCTGGAGCTCAACTCCAGAGATGCATTCGATACTGGCTTGCTAGAGGTTGATAGAGGAAAGCGGAATTCCGAGTGTAGCGGTGAAATGCGTAGATATTCGGAGGAACATCAGTAGCGAAGGCGGCTTTCTGGGTCACACCTGACGCTGAGGTACGAAAGCGTGGGG |
| P-asv4934 | GCTAGCGTTGCTCGGAATTACTGGGCGTAAAGGGCGCGTAGGCGGACAGTTAAGTCAGAGGTGAAAGCCCAGGGCTCAACCTTGGAATTGCCTTTGATACTGGCTGTCTTGAGTTCGGGAGAGGTGAGTGGAATGCCGAGTGTAGAGGTGAAATTCGTAGATATTCGGCGGAACACCAGTGGCGAAGGCGACTCACTGGCCCGATACTGACGCTGAGGCGCGAAAGCGTGGGG |
| P-asv4935 | GCAAGCGTTATTCGGAATCATTGGGCGTAAAGGGTGCGTAGACGGCATATTAAGTCAACTGTTAAATCCTCCGGCCTAACTGGAGTTCAGCGGTAGAAACTGATAAGCTTGAGGTTGGAAGAGAGAAGTGGAATTCTCGGAGTAGCGGTAAAATGCGTAGATCTCGAGAGGAACACCAATGGCGAAGGCAGCTTCTTGGTCCAAATCTGACGTTGAGGCACGAAAGCGTGGGG |
| P-asv4936 | CCAAGCGTTATCCGGAATCATTGGGTTTAAAGGGTCCGTAGGCGGTCTTATAAGTCAGTGGTGAAATCTCCCCGCTCAACGGGGAAACGGCCATTGATACTGTAGGACTTGAATTATTAGGAAGTAACTAGAATATGTAGTGTAGCGGTGAAATGCTTAGAGATTACATGGAATACCAATTGCGAAGGCAGGTTACTACTAATGGATTGACGCTGATGGACGAAAGCGTGGGT |
| P-asv4937 | GCTAGCGTTGCTCGGAATCACTGGGCGTAAAGGGCGCGTAGGCGGCCGATCAAGTCAGGGGTGAAAGCCTGTGGCTCAACCACAGAATTGCCTTTGATACTGTTCGGCTCGAGACCGGAAGAGGTTGGTGGAACTGCGAGTGTAGAGGTGAAATTCGTAGATATTCGCAAGAACACCAGTGGCGAAGGGGCCAACTGGTCCGGTTCTGACGCTGAGGCGCGAAAGCGTGGGG |
| P-asv4938 | GCAAGCGTTATCCGGAATTACTGGGTTTAAAGGGTGCGTAGGCGGCTATATAAGTCAGAGGTGAAAGCCCGGAGCTTAACTTCGGAATTGCCTTTGATACTGTATAGCTTGAAAGAGGTTGAGGTCAGCGGAATGTGACATGTAGCGGTGAAATGCATAGATATGTCATAGAACACCAATTGCGAAGGCAGCTGGCTGGGCCTACTTTGACGCTGAGGCACGAAAGCGTGGGG |
| P-asv4939 | CCAAGCGTTGTTCGGATTTATTGGGCGTAAAGCGCGCGCAGGCGGGTCAGCAAGTCAGGTGTGAAATCTCGGAGCTCAACTCCGAAACTGCGCCTGAAACTGCTAATCTAGAATGTCGGAGGGGACAGGGGAATTTCACGTGTAGGGGTAAAATCCGTAGAGATGTGAAGGAACACCGGAGGCGAAGGCGCCTGTCTGGACGACTATTGACGCTGAGGCGCGAAAGCGTGGGG |
| P-asv4940 | GCGAGCGTTGTCCGGATTTATTGGGCGTAAAGAGCTCGTAGGCGGCTCGGTAAGTCGGATGTGAAATCTCCAGGCTTAACCTGGAGAGGCCATTCGATACTGCTGTGGCTTGAGTCCGGTAGAGGAGCATGGAATTCCCGGTGTAGCGGTGAAATGCGCAGATATCGGGAGGAACACCAGTAGCGAAGGCGGTGCTCTGGGCCGGTACTGACGCTGAGGAGCGAAAGCGTGGGG |
| P-asv4941 | GCAAGCGTTACTCGGAATCACTGGGCGTAAAGAGCGTGTAGGCGGACTAATAAGTCAGAAGTGAAATCCAATAGCTTAACTATTGAACTGCTTTTGAAACTGTTGGTCTAGAGTGTGGGAGAGGTAGATGGAATTTCTGGTGTAGGGGTAAAATCCGTAGATATCAGAAGGAATACCGATTGCGAAGGCGATCTACTGGAACATAACTGACGCTGAGACGCGAAAGCGTGGGG |
| P-asv4942 | GCGAGCGTTAATCGGATTTACTGGGCGTAAAGCGCGCGTAGGTGGATACTTAAGTCGGATGTGAAAGCCCTGGGCTTAACCTAGGAACTGCATCCGATACTGGGTGTCTAGAGTATGGTAGAGGGAAGTGGAATTTCCGGTGTAGCGGTGAAATGCGTAGATATCGGAAAGAACACCAGTGGCGAAGGCGGCTTCCTGGACTAATACTGACACTGAGGTGCGAAAGCGTGGGG |
| P-asv4943 | GCGAGCGTTGTTCGGAATTACTGGGCGTAAAGGGCGCGTAGGCGGTGACGTTAGTCGGGCGTGAAAGCCCCGGGCTCAACCTGGGATGTGCGTTCGATACTGCGTTGCTTGAGAGCGGAAGAGGGTAGTGGAATTCCCAGTGTAGAGGTGAAATTCGTAGATATTGGGAAGAACACCAGTGGCGAAGGCGGCTACCTGGTCCGTTTCTGACGCTGAGGCGCGAAAGCGTGGGG |
| P-asv4944 | GCGAGCGTTGTCCGGAATTACTGGGCGTAAAGAGCTCGTAGGCGGTTTGTCGCGTCGTCTGTGAAATTCTGCAACTCAATTGCAGGCGTGCAGGCGATACGGGCAGACTTGAGTACTACAGGGGAGACTGGAATTCCTGGTGTAGCGGTGAAATGCGCAGATATCAGGAGGAACACCGGTGGCGAAGGCGGGTCTCTGGGTAGTAACTGACGCTGAGGAGCGAAAGCGTGGGT |
| P-asv4945 | GCGAGCGTTCCTCTTCTTTCCTTTTCTTAAAGCGTGCGTAGGCGGCTTTTTAAGTCGGATGTGAAATCCCTGAGCTTAACTTAGGAATTGCATTCGATACTGGGAAGCTAGAGTATGGGAGAGGATGGTAGAATTCCAGGTGTAGCGGTGAAATGCGTAGAGATCTGGAGGAATACCGATGGCGAAGGCAGCCATCTGGCCTAATACTGACGCTGAGGTACGAAAGCATGGGG |
| P-asv4946 | GCAAGCGTTAATCGGAATTACTGGGCGTAAAGCGAGCGTAGGTGGCTTGATAAGTCAGATGTGAAATCCCCGGGCTTAACCTGGGAACTGCATCTGAAACTGTTAGGCTAGAGTAGGTGAGAGGGAAGTAGAATTTCAGGTGTAGCGGTGAAATGCGTAGAGATCTGAAGGAATACCGATGGCGAAGGCAGCTTCCTGGCATCATACTGACACTGAGGCTCGAAAGCGTGGGT |
| P-asv4947 | GCGAGCGTTAATCGGAATTACTGGGCGTAAAGCGTGCGCAGGCGGTTTTGTAAGCCAGATGTGAAATCCCCGGGCTTAACCTGGGAATGGCATTTGGGACTGCAAGGCTTGAGTGCGGCAGAGGAGACTGGAATTCCTGGTGTAGCAGTGAAATGCGTAGATATCAGGAGGAACACCGATGGCGAAGGCAGGTCTCTGGGCTGACACTGACGCTCATGCACGAAAGCGTGGGG |
| P-asv4948 | GCGAGCGTTATCCGGAATTACTGGGTGTAAAGGGCGTGTAGGCGGCACTGTAAGTCAGATGTGAAATCTCCCGGCTCAACCGGGAGCGTGCATCTGATACTGCAATGCTTGAGTGATAGAGGGGAAAGCGGAATTCCTAGTGTAGCGGTAAAATGCGTAGATATTAGGAGGAACACCAGTGGCGAAGGCGGCTTTCTGGCTATTAACTGACGCTGAGGCGCGAAAGCGTGGGG |
| P-asv4949 | GCGAGCGTTATCCGGAATTATTGGGCGTAAAGAGCTCGTAGGCGGTTTGTCGCGTCTGTCGTGAAAGTCCGGGGCTTAACCCCGGATCTGCGGTGGGTACGGGCAGACTAGAGTGCAGTAGGGGAGACTGGAATTCCTGGTGTAGCGGTGAAATGCGCAGATATCAGGAGGAACACCGATGGCGAAGGCAGGTCTCTGGGCTGTAACTGACGCTGAGGAGCGAAAGCATGGGG |
| P-asv4950 | GCAAGCGTTACTCGGAATCACTGGGCGTAAAGGACGCGTAGGCGGATTATCAAGTCTTTTGTGAAATCTAGTGGCTTAACCACTAAACTGCTTAGGAAACTGATAGTCTAGAGTAGGGGAGAGGTAGATGGAATTCTTGGTGTAGGGGTAAAATCCGTAGAGATCAAGAAGAATACCCATTGCGAAAGCGATCTGCTGGAACCTAACTGACGCTGAGGCGTGAAAGCGTGGGT |
| P-asv4951 | GCGAGCGTTGTCCGGAATTATTGGGCGTAAAGCGCGCGCAGGCGGTCTCTTAAGTCTGATGTGAAAGCCCCCGGCTCAACCGGGGAGGGTCATTGGAAACTGGGAGACTTGAGTGCAGGAGAGGGAAGCGGAATTCCATGTGTAGCGGTGAAATGCGTAGAGATATGGAGGAACACCAGTGGCGAAGGCGGCTTCCTGGCCTGTAACTGACGCTGAGGCGCGAAAGCGTGGGG |
| P-asv4952 | GCGAGCGTTAATCGGATTTACTGGGCGTAAAGCGTGCGTAGGTGGCCAATTAAGTCAAATGTGAAATCCCCGAGCTTAACTTGGGAATTGCATTCGATACTGGTTGGCTAGAGTATGGGAGAGGATGGTAGAATTCCAGGTGTAGCGGTGAAATGCGTAGAGATCTGGAGGAATACCGATGGCGAAGGCAGCCATCTGGCCTAATACTGACACTGAGGTACGAAAGCATGGGG |
| P-asv4953 | GCAAGCGTTGTCCGGATTTATTGGGCGTAAAGAGCTCGTAGGCGGTTTGTCGCGTCGGCCGTGAAAACCTGGGGCTTAACTCTGGGCGTGCGGTCGATACGGGCAGACTTGAGTGTGGCAGGGGAGACTGGAATTCCTGGTGTAGCGGTGAAATGCGCAGATATCAGGAGGAACACCGGTGGCGAAGGCGGGTCTCTGGGCCGATACTGACGCTGAGGAGCGAAAGCGTGGGG |
| P-asv4954 | GCTAGCGTTGTTCGGAATTACTGGGCGTAAAGGGCGCGTAGGCGGTTTTTCAAGTCAGGCGTGAAAGCCCCGGGCTTAACCTGGGAACTGCGCTTGAGACTGTGAGACTAGAGTACCGGAGAGGTAGGTGGAATTCCCAGTGTAGAGGTGAAATTCGTAGATATTGGGAAGAACACCAGTGGCGAAGGCGGCCTACTGGACGGTAACTGACGCTGAGGCGCGAAAGCGTGGGG |
| P-asv4955 | TCGAGTGATGTCCCATATTATTGGGCCTAAAGCGTCCGTAGCTGGCCAGGCAAGTCTATCGGGAAATCCACCTGCTCAACAGGTGGGCGTCCGGTAGAAACTGCCTGGCTTGGAACCGGAAGGCTCAGAGAGTACGTCCGGGGTAGGAGTTAAATCCCGTAATCCCGGACGGACTACCGATGGCGAAAGCACTCTGAGAAGACGGCTTCGACAGTGAGGGACGAAAGCTAGGGT |
| P-asv4956 | TCTAGCGTTATCCGGAATTACTGGGCGTAAAGGGTGCGTAGGTGGTTTTTTAAGTCAGAAGTGAAAGGCTACGGCTCAACCGTAGTAGGCTTTTGAAACTAGAGAACTTGAGTGCAGGAGAGGAGAGTAGAATTCCTAGTGTAGCGGTGAAATGCGTAGATATTAGGAGGAATACCAGTAGCGAAGGCGGCTCTCTGGACTGTAACTGACACTGAGGCACGAAAGCGTGGGG |
| P-asv4957 | GCAAGCGTTGTCCGGAATTATTGGGCGTAAAGAGCTCGTAGGCGGTTTGTCGCGTCCGCTGTGAAAGCCCGGGGCTCAACCCCGGGTCGGCAGTGGGTACGGGCAGACTAGAGTGCAGTAGGGGAGACTGGAATTCCTGGTGTAGCGGTGAAATGCGCAGATATCAGGAGGAACACCGATGGCGAAGGCAGGTCTCTGGGCTGTAACTGACGCTGAGGAGCGAAAGCATGGGG |

| | |
|---|---|
| P-asv4958 | GCGAGCGTTAATCGGAATTACTGGGCGTAAAGCGCGCGTAGGCGGTCTGATAAGCCGGTTGTGAGAGCCCCGGGCTCAACCTGGGAACGGCATCCGGAACTGTCAGGCTAGAGTGCAGGAGAGGAAGGTAGAATTCCCGGTGTAGCGGTGAAATGCGTAGAGATCGGGAGGAATACCAGTGGCGAAGGCGGCCTTCTGGACTGACACTGACACTGAGGTGCGAAAGCGTGGGT |
| P-asv4959 | GCTAGCGTTGTTCGGAATCACTGGGCGTAAAGCGCACGTAGGCGGATGTGTCAGTCAGGGGTGAAATCCCGGAGCTCAACTTCGGAACTGCCTTTGATACAGCACGTCTTGAGTCCGATAGAGGTGGGTGGAATTCCTAGTGTAGAGGTGAAATTCGTAGATATTAGGAAGAACACCGATGGCGAAAGCAGCTCTCTGGGACGGTACTGACGCTGAGGCGCGAAAGCGTGGGG |
| P-asv4960 | GCTAGCGTTGCTCGGAATTACTGGGCGTAAAGGGAGCGTAGGCGGATAGTTAAGTCAGAGGTGAAAGCCCAGGGCTCAACCTTGGAATTGCCTTTGATACTGGCTATCTTGAGTATGGGAGAGGTTAGCGGAACTCCGAGTGTAGAGGTGAAATTCGTAGATATTCGGAAGAACACCAGTGGCGAAGGCGGCTAACTGGCCCATTACTGACGCTGAGGCTCGAAAGCGTGGGG |
| P-asv4961 | GCTAGCGTTGTTCGGAATTACTGGGCGTAAAGCGTGCGCAGGCGGCTTTTCAAGTCAGAGGTGAAAGCCCCGGGCTCAACCCGGGAATGGCCTTTGAGACTGGATGGCTAGAGTGCGAGAGAGGTGAGTGGAATTCCCAGTGTAGAGGTGAAATTCGTAGATATTGGGAAGAACACCCGTGGCGAAGGCGGCTCACTGGCTCGCAACTGACGCTCAGGCACGACAGCGTGGGG |
| P-asv4962 | GCGAGCGTTGTCCGGAATTACTGGGCGTAAAGAGCTCGTAGGTGGTTTGTCGCGTTGTCCGTGAAAACCGGAGGCTTAACCTCTGGCGGGCGGGCGATACGGGCAGACTGGAGTACTGCAGGGGAGACTGGAATTCCTGGTGTAGCGGTGGAATGCGCAGATATCAGGAGGAACACCGGTGGCGAAGGCGGGTCTCTGGGCAGTAACTGACGCTGAGGAGCGAAAGCGTGGGG |
| P-asv4963 | GCAAGCGTTGTTCGGATTGACTGGGCGTAAAGGGCGTGTAGGCGGTCGGCCAAGTCTGATGTGAAAGCCCGGGGCTTAGCTCCGGAAGTGCGTCGGAAACTAGTCGGCTGGAGCGCGATAGAGGAGGGTGGAATTCCTGGTGTAGCGGTGAAATGCGTAGATATCAGGAGGAACACCGGCGGCGAAGGCGGCCCTCTGGATCGTTGCTGACGCTGAGACGCGAAAGCGTGGGG |
| P-asv4964 | GCTAGCGTTGCTCGGAATCACTGGGCGTAAAGGGCGCGTAGGCGGATCTTTAAGTCGGGGGTGAAAGCCCAGGGCTCAACCCTGGAATTGCCTTCGATACTGGAGATCTTGGGACCGGAAGAGGTTAGTGGAACTGCGAGTGTAGAGGTGAAATTCGTAGATATTCGCAAGAACACCAGTGGCGAAGGCGGCTGACTGGTCCGGTTCTGACGCTGAGGCGCGAAAGCGTGGGG |
| P-asv4965 | GCAAGCGTTGTTCGGAATTACTGGGCGTAAAGGGCTCGTAGGTGGCCAACTAAGTCAGACGTGAAATCCCTCAGCTTAACTGGGGAACTGCGTCTGATACTGGATGGCTTGAGTTTGGGAGAGGGATGCGGAATTCCAGGTGTAGCGGTGAAATGCGTAGATATCTGGAGGAACACCGGTGGCGAAGGCGGCGTTCTGGACTGCAACTGACACTGAGGAACGAAAGCCAGGGG |
| P-asv4966 | CCGAGCGTTGTCCGGATTTATTGGGCGTAAAGAGCTCGTAGGCGGCTCACTAAGTCGGATGTGAAAACTCGGGGCTCAACCCCGAGACGCCATCCGATACTGGTGTGGCTTGAATCTGGTAGGGGAGATCGGAATTCCTGGTGTAGCGGTGAAATGCGCAGATATCAGGAGGAACACCGGTGGCGAAGGCGGATCTCTGGGCCAGTATTGACGCTGAGGAGCGAAAGCGTGGGT |
| P-asv4967 | GCTAGCGTTGCTCGGAATCACTGGGCGTAAAGCGCACGTAGGCGGCTTTTTAAGTCGGGGGTGAAATCCTGGAGCTCAACTCCAGAACTGCCTTCGATACTGAGAAGCTCGAGTCCGGGAGAGGTGAGTGGAACTGCGAGTGTAGAGGTGAAATTCGTAGATATTCGCAAGAACACCAGTGGCGAAGGCGGCTCACTGGCCCGGTACTGACGCTGAGGTGCGAAAGCGTGGGG |
| P-asv4968 | GCTAGCGTTGTTCGGAATTACTGGGCGTAAAGCGCACGTAGGCGGCTTTGTAAGTTAGAGGTGAAAGCCCGGGGCTCAACTCCGGAATTGCCTTTAAGACTGCATCGCTAGAACGTCGGAGAGGTGAGTGGAATTCCGAGTGTAGAGGTGAAATTCGTAGATATTCGGAAGAACACCAGTGGCGAAGGCGACTCACTGGACGACCGTTGACGCTGAGGTGCGAAAGCGTGGGG |
| P-asv4969 | GCAAACGTTGTTCGGAATCACTGGGCATAAAGCGCATGTAGGCGGCTCCGTAAGTGTCGGATGAAATCCCACGGCCTACCCGTGGAACTGTTCGGCATACTGCGGTGCTAGAGGATGGTAGAGGAGAGCGGAACTCTTGGTGGAGCGGTGAAATGCGTAGATATCAAGAGGAACACCGGTGGCGAAGGCGGCTCTCTGGTCCATTTCTGACGCTGAGGTGCGAAAGCTAGGGG |
| P-asv4970 | CCGAGTGGTCGGGACGATTATTGGGCCTAAAGCATCCGTAGCCGGTTCTGCAAGTCTTCCGTTAAATCCAGCCGCTCAACGGATGGGCCGCGGAGGATACTACAGGACTAGGAGGCGGGAGAGGCAAGCGGTACTCAGTGGGTAGGGGTAAAATCCTTTGATCCATTGAAGACCACCAGTGGCGAAGGCGGCTTGCCAGAACGCGCTCGACGGTGAGGGATGAAAGCTGGGGG |
| P-asv4971 | GCGAGCGTTGTCCGGAATCATTGGGCGTAAAGAGCGCGCAGGCGGCTTGGCAAGTCGAGTGTGAAATCCCGGGGCTCAACCCCGGGACTGCATCCGGTACTTCCGGGCTGGAGGTAGGTAGGGGAGATCGGAATTCCTGGTGTAGCGGTGAAATGCGCAGATATCAGGAGGAACACCGGTGGCGAAGGCGGATCTCTGGGCCTTACCTGACGCTGAGGCGCGAAAGCTAGGGG |
| P-asv4972 | GCAAGCGTTGTTCGGAATTACTGGGCGTAAAGGGCTCGTAGGTGGCCAACTAAGACACACGTGAAATCCCTCGGCTTAACCGGGGAACTGCGTGTGATACTGGATGGCTTGAGTTTGGGAGAGGGATGCGGAATTCCAGGTGTAGCGGTGAAATGCGTAGATATCTGGAGGAACACCGGTGGCGAAGGCGGCATCCTGGACCAATACTGACACTGAGGAGCGAAAGCCAGGGG |
| P-asv4973 | GCAAGCGTTGTCCGGATTTATTGGGCGTAAAGAGCGTGTAGGCGGCCAGGTAGGTCTGCTGTGAAAACTCGAGGCTTAACCCCGAGCTGTCGGCGGAAACCATCTGGCTAGAGTCCGGAAGAGGAGAATGGAATTCCCGGTGTAGCGGTGAAATGCGCAGATATCGGGAAGAACACCCGTGGCGAAGGCGGTTCTCTGGGACGGTACTGACGCTGAGACGCGAAAGCGTGGGG |
| P-asv4974 | GCTAGCGTTGTTCGGAATTACTGGGCGTAAAGCGCGCGCGGGCGGTCTTCCAAGTCAGTGGTGAAAGCCCGGAGCTCAACTCCGGAAATGCCATTGAAACTGTGAGACTGGAGTACGAGAGAGGTGAGTGGAATTCCCAGTGTAGAGGTGAAATTCGTAGATATTGGGAAGAACACCGGTGGCGAAGGCGGCTCACTGGCTCGTAACTGACGCTCAGGCGCGACAGCGTGGGG |
| P-asv4975 | GCAAGCGTTGTCCGGATTTATTGGGCGTAAAAAGCGTGTAGGCGGCTAGGTAGGTACGTTGTGAAAACTCGAGGCTCAACCTCGAGACGTCGATGGAAACCATCTAGATAGAGTCCGGAAGAGGAGGGTGGAATTCCTGGTGTAGCGGTGAAATGCGCAGATATCAGGAAGAACACCCGTGGCGAAGGCGGCTCTCTGGTACGTGACTGACGCTGAGACGCGAAAGCGTGGGG |
| P-asv4976 | GCTAGCGTTGCTCGGAATCACTGGGCGTAAAGCGCACGTAGGCGGCTTTTTAAGTCGGGGGTGAAATCCTGGAGCTCAACTCCAGAACTGCCTTCGATACTGAGAAGCTCGAGTCCGGGAGAGGTGAGTGGAACTGCGAGTGTAGAGGTGAAATTCGTAGATATTCGCAAGAACACCAGTGGCGAAGGCGGCTCACTGGCCCGATACTGACGCTGAGGCACGAAAGCGTGGGG |
| P-asv4977 | GCCAGCGTTGCTCGGAATTACTGGGCGTAAAGGGTCCGTAGGCGGTGCGGCAAGTCGGAAGTGAAATCTCCAGGCTTAACTTGGAGGCTGCTCCCGAAACTGCCGTGCTGGAGTGCGGGAGAGGCGAGTGGAATTGCAGGTGTAGCGGTGAAATGCGTAGATATCTGCAGGAACACCAGAGGCGAAGGCGGCTCGCTGGACCGCAACTGACGCTGAGGGACGAAAGCTAGGGG |
| P-asv4978 | GCTAGCGTTGCTCGGAATTACTGGGCGTAAAGGGCGCGTAGGCGGCTCGCTTAGTCAGGCGTGAAATTCCTGGGCTCAACCTGGGGACTGCGCTTGAGACGGGTGGGCTAGAGGACGGAAGAGGCTCGCGGAATTCCCAGTGTAGAGGTGAAATTCGTAGATATTGGGAAGAACACCGGTGGCGAAGGCGGCGAGCTGGTCCGTTACTGACGCTGAGGCGCGACAGCGTGGGG |
| P-asv4979 | GCGAGCGTTGTCCGGAATTATTGGGCGTAAAGAGCTTGTAGGCGGTCTGTCGCGTCTGCTGTGAAAATCCGGGGCTCAACCCCGGACTTGCAGTGGGTACGGGCAGACTAGAGTGTGGTAGGGGAGACTGGAATTCCTGGTGTAGCGGTGAAATGCGCAGATATCAGGAGGAACACCGATGGCGAAGGCAGGTCTCTGGGCCACTACTGACGCTGAGAAGCGAAAGCATGGGG |
| P-asv4980 | GCTAGCGTTGTTCGGATTTACTGGGCGTAAAGCGCGTGTAGGCGGACTTGTAAGTTGGGGGTGAAATCCCGAGGCTCAACCTCGGAACTGCCTCCAAAACTACAAGTCTGGAGCAAGATAGAGGCAAGTGGAATTGCGAGTGTAGAGGTGAAATTCGTAGATATTCGCAGGAACACCAGTGGCGAAGGCGACTTGCTGGATCTTTGCTGACGCTGAGACGCGAAAGCGTGGGG |
| P-asv4981 | CCGAGCGTTAGGCGGAATCACTGGGCTTAAAGCGTGTGTAGGCGGGCGGGCAAGTACCTTGTGAAATCCCACGGCTCAACCGTGGAACGGCTGGGTATACTGCCTGTCTTGAGCCAGCTAGGGGCGACCGGAACAGACGGTGGAGCGGTGAAATGCGTAGATATCGTCTGGAACGCCAAAGGTGAAAACGGGTCGCTGGGGCTGTGCTGACGCTGAGACACGAAAGCCAGGGG |
| P-asv4982 | GCAAGCGTTGTTCGGAATTATTGGGCGTAAAGCGCGTGTAGGCGGTCCTGCAAGTTGGGTGTGAAAGCCCTCGGCTTAACCGAGGAAGTGCGCCCAAAACTACAGAACTTGAGTACCGGAGAGGGTGGCGGAATTCCCGGTGTAGAGGTGAAATTCGTAGATATCGGGAGGAACACCAGTGGCGAAGGCGGCCACCTGGACGGATACTGACGCTGAGACGCGAAAGCGTGGGT |
| P-asv4983 | GCAAACGTTGCTCGGAATTACTGGGCGTAAAGCGCGTGTAGGCGGACCTGTAAGTTGGACGTGAAAGCCCTGGGCTCAACCCAGGAATTGCATCCAAAACTACAGGCCTAGAGTGATGGAGAGGGCAGCAGAATTCCTGGTGTAGAGGTGAAATTCGTAGATATCAGGAGGAACACCGGTGGCGAAGGCGGCTGCCTGGACATCAACTGACGCTGAGACGCGAAAGCGTGGGG |
| P-asv4984 | GCGAGCGTTAATCGGAATTACTGGGCGTAAAGCGTGCGCAGGCGGTACGTTGTGTCGGCGGTGAAATCCCCGGGCTTAACCTGGGAATTGCCGTTGAAACTGGTGTACTTGAGTGTGGCAGAGGGGGGTGGAATTCCGCGTGTAGCAGTGAAATGCGTAGAGATGCGGAGGAACACCGATGGCGAAGGCAGCCCCCTGGGCTAACACTGACGCTCAGGCACGAAAGCGTGGGG |
| P-asv4985 | GCGAGCGTTGTCCGGATTTATTGGGCGTAAAGAGCTCGTAGGCGGCTCGTTGCGTCAATCGTGAAAACCTACTGCTTAACGGTGGGCTTGCGGTTGATACGGGCGGGCTAGAGTTCGGTAGGGGAGACTGGAATTCCTGGTGTAGCGGTGGAATGCGCAGATATCAGGAGGAACACCGGTGGCGAAGGCGGGTCTCTGGGCCGACATTGACGCTGAGGAGCGAAAGCGTGGGG |
| P-asv4986 | GCTAGCGTTGTTCGGAATTACTGGGCGTAAAGCGCACGTAGGCGGCTTTGTAAGTTAGAGGTGGAAGCCTGGAGCTCAACTCCAGAATTGCCTTTAAGACTGCAGAGCTAGAATCGCGGAGAGGTGAGTGGAATTCCGAGTGTAGAGGTGAAATTCGTAGATATTCGGAAGAACACCAGTGGCGAAGGCGGCTCACTGGACGCGTATTGACGCTGAGGTGCGAAAGCGTGGGG |
| P-asv4987 | GCAAGCGTTGTTCGGAATTATTGGGCGTAAAGGGCGCGTAGGCGGCGCGACAAGTCACCTGTGAAATCCCCAGGCTTAACCTGGGGCCTGCAGGCGAAACTGTCGTGCTGGAGTATGGGAGAGGTGCGTGGAATTCCCAGTGTAGCGGTGAAATGCGTAGATATCGGGAGGAACACCTGTGGCGAAAGCGGCGCACTGGACCATAACTGACGCTGAGGCGCGAAAGCTAGGGG |
| P-asv4988 | GCAAGCGTTGTCCGGAATTATTGGGCGTAAAGCGCGCGCAGGCGGTCTTTTAAGTTTGGTGTTTAAGCCCGGGGCTCAACCCCGGTTCGCACTGAAAACTGGGAGACTTGAGTGCAGGAGAGGAAAGCGGAATTCCACGTGTAGCGGTGAAATGCGTAGAGATGTGGAGGAACACCAGTGGCGAAGGCGGCTTTCTGGACTGTAACTGACGCTGAGGCGCGAAAGCGTGGGG |
| P-asv4989 | GCTAGCGTTATTCGGAATTATTGGGCGTAAAGGGCGCGTAGGCGGCCATGCGTGTCTTCCGTGAAATCCCTCGGCTTAACTGAGGACGTGCGGAAGAAACTGCAAGGCTTGAGTCCGGGAGAGGGGGGTGGAATTCCCAGTGTAGCGGTGAAATGCGTAGATATTGGGAGGAACACCGGTGGCGAAGGCGGCCCCCTGGACCGGTACTGACGCTGAGGCGCGAAAGCGTGGGT |
| P-asv4990 | CCGAGCGTTGTCCGGAGTTACTGGGCGTAAAGCGCGCGCAGGCGGTTTGCCACGTCGGGTGTGAAAGCCCCCCGCTCAACGGGGGAGGGTCATTCGAAACGGGCAGACTGGAGGCAGGGAGAGGTCGGTGGAATTCCCGGTGTAGTGGTGAAATGCGTAGATATCGGGAGGAACACCAGTGGCGAAGGCGGCCGGCTGGACCTGACCTGACGCTGAGGCGCGAAGGCGTGGGG |
| P-asv4991 | GCAAGCGTTGTCCGGATTTATTGGGCGTAAAGCGAGCGCAGGCGGTTTCTTAAGTCTGATGTGAAAGCCCTCGGCTCAACCGAGGAAGGTCATTGGAAACTGGGGAACTTGAATGCAGAAGAGGAGAGTGGAATTCCATGTGTAGCGGTGAAATGCGTAGATATATGGAGGAACACCAGTGGCGAAGGCGACTCTCTGGTCTGTAATTGACGCTGAGGCTCGAAAGCGTGGGG |
| P-asv4992 | GCGAGCGTTGTTCGGAATTATTGGGCGTAAAGAGCGTGTAGGCGGTTTGGTAAGTCTGCCGTGAAAACCCGGGGCTCAACCCCGGGCGTGCGGTGGATACTGCCAGGCTAGAGGATGGTAGAGGCGAGTGGAATTCCCGGTGTAGCGGTGAAATGCGGGGATATCGGGAGGAACACCAGTAGCGAAGGCGGCTCGCTGGGCCATTCCTGACGCTGAGACGCGAAAGCTAGGGG |
| P-asv4993 | CCAGGCGTTATCCGGATTTATTGGGTTTAAAGGGAGCGTAGGCGGACTGTTAAGTCAGCGGTTAAAGAGTGTGGCTCAACCATACGATGCCGTTGAAACTGGTGGTCTTGAGTGCACACAGGGATGCTGGAACTCGTGGTGTAGCGGTGAAATGCTTAGATATCACGATGAACTCCGATCGCGAAGGCAGGTGTCCGGGGTGTAACTGACGCTGAGGCTCGAAAGTGTGGGT |
| P-asv4994 | GCGAGTGATCGGGACTCTTATTGGGCCTAAAGCGCCCGTAGCCGGCTCTGTAAGTCACTGCTTAAAGACCCGAGCTCAACTCGGGGAAGGGTAGTGATACTACAGAGCTAGGGGGCGGGAGAGGTCGGGGGTACTCCTAGAGTAGGGGCGAAATCCGTAGATCCTAGGAGGACCACCAGTGGCGGAGGCGCCCGGCTAGAACGCGCCCGACGGTGAGGGGCGAAAGCTGGGGT |
| P-asv4995 | CCGAGCGTTGATCGGAATCATTGGGCTTAAAGGGTGCGCAGGCGGATGGGCAAGTGCGTTGTGAAATCCCATGGCTTAACCATGGAATGGCTTCGCAGACTGCCCGTCTTGAACCCGGCAGGGGCTGTCAGAACGTTAGGTGGAGCGGTGAAATGCGTAGATATCTAACGGAATGCCGATGGAGAAGTCAGACAGCTGGGCCGGCGTTGACGCTCAGGCACGAAAGCGTGGGT |
| P-asv4996 | CCGAGCGTTGTCCGGAATCATTGGGCGTAAAGGGTACGTAGGCGGTTATCTAAGTTAGAAGTCAAAGGCTATAGCTCAACTATAGTAAGCTTTTAAAACTGGATAACTTGAGAGATGGAAGGGAAAGTGGAATTCCTAGTGTAGCGGTGAAATGCGCAGATATTAGGAGGAATACCGGTGGCGAAGGCGACTTTCTGGCCATTATCTGACGCTGAGGTACGAAAGCGTGGGT |

| | |
|---|---|
| P-asv4997 | GCAAGCGTTATCCGGAATCACTGGGTTTAAAGGGTGCGTAGGCGGATATTTAAGTCAGAGGTGAAAGCTCGCAGCTCAACTGCGGAATTGCCTTTGATACTGATTATCTTGAATTATGTTGAGGTTAGCGGAATGAGTCATGTAGCGGTGAAATGCATAGATATGACTTAGAACACCAATTGCGAAGGCAGCTAGCTAGGCATATATTGACGCTGAGGCACGAAAGCGTGGGG |
| P-asv4998 | CCGAGCGTTATCCGGATTTATTGGGTTTAAAGGGAGCGTAGGCGGAAGCTTAAGTCAGTTGTGAAAGTTTGCGGCTCAACCGTAAAATTGCAGTTGATACTGGGTTTCTTGAGTGCAGTAGAGGTAGGCGGAATTCGTGGTTTAGCGGTGAAATGCTTAGATATCACGAAGAACTCCGATTGCGAAGGCAGCTTACTGGACTGTAACTGACGCTGATGCTCGAAAGTGTGGGT |
| P-asv4999 | GCAAGCGTTAATCGGAATTACTGGGCGTAAAGCGCGCGTAGGTGGTTTGTTAAGTTGAATGTGAAAGCCCCGGGCTCAACCTGGGAACTGCATCCAAAACTGGCAAGCTAGAGTATGGCAGAGGGTGGTGGAATTTCCTGTGTAGCGGTGAAATGCGTAGATATAGGAAGGAACACCAGTGGCGAAGGCGACCACCTGGGCTAATTCTGACACTGAGGTGCGAAAGCGTGGGG |
| P-asv5000 | GCAAGCGTTGTCCGGAATTATTGGGCGTAAAGCGCGCGCAGGTGGTTCCTTAAGTCTGACGTGAAAGCCCACGGCTCAACCGTGGAGGGTCATTGGAAACTGGGGAACTTGAGTGCAGAAGAGGAAAGTGGAATTCCACGTGTAGCGGTGAAATGCGTAGAGATGTGGAGGAACACCAGTGGCGAAGGCGACTTTCTGGTCTGTAACTGACACTGAGGCGCGAAAGCGTGGGG |
| P-asv5001 | GCAAGCGTTAATCGGAATTACTGGGCGTAAAGCGTGCGCAGGCGGTTATGCAAGACAGAGGTGAAATCCCCGGGCTCAACCTGGGAACTGCCTTTGTGACTGCATGGCTAGAGTACGGTAGAGGGGGATGGAATTCCGCGTGTAGCAGTGAAATGCGTAGATATGCGGAGGAACACCGACGGCGAAGGCAGCCCCCTGGGATAACACTGACGCTCATGCACGAAAGCGTGGGG |
| P-asv5002 | GCAAGCGTTGTCCGGATTTATTGGGTTTAAAGGGTGCGTAGGCGGCCCCTTAAGTCAGCGGTGAAAGCCCGGGGCTCAACCCCGGAACTGCCGTTGATACTGGGGGGCTTGAGTACAGACGAGGTAGGCGGAATTGAGGGTGTAGCGGTGAAATGCTTAGATACCCTCAAGAACACCGATTGCGTAGGCAGCTTACTAGGCTGTGACTGACGCTGAGGCACGAAAGCGTGGGG |
| P-asv5003 | GCGAGCGTTGTCCGGATTTATTGGGTTTAAAGGGTGCGTAGGTGGCTTTATAAGTCAGTGGTGAAATACAGCCGCTCAACGGTTGAGGTGCCATTGATACTGTAGAGCTTGAAACAATTGGAGGCTGCCGGAATGGATGGTGTAGCGGTGAAATGCATAGATATCATCCAGAACACCGATTGCGAAGGCAGGTGGCTACGTTTGATTTGACACTGAGGCACGAAAGCATGGGG |
| P-asv5004 | GCGAGCGTTGTCCGGAATTATTGGGCGTAAAGAGTGCGTAGGCGGCAAATTAAGTCAGATGTGAAAACTAAGGGCTCAACCCATAGATTGCATCTGAAACTGATATGCTTGAGTCAAGGAGAGGAAAGTGGAATTCCTAGTGTAGCGGTGGAATGCGTAGATATTAGGAGGAATACCGGTGGCGAAGGCGACTTTCTGGACCTGAACTGACGCTGAGGCACGAAAGCGTGGGG |
| P-asv5005 | CCAAGCGTTATCCGGAATCATTGGGTTTAAAGGGTCCGTAGGCGGGATGATAAGTCAGTGGTGAAAGTCTGCAGCTCAACTGTAGAATTGCCATTGATACTGTTGTTCTTGAGTGCTTGTGAAGTGGTTAGAATGAGTAGTGTAGCGGTGAAATGCATAGATATTACTCAGAATACCGATTGCGAAGGCAGATCACTAACAATACACTGACGCTGATGGACGAAAGCGTAGGT |
| P-asv5006 | CCGAGCGTTATCCGGATTTATTGGGTTTAAAGGGTTCGTAGGCGGTAATTTAAGTCAGTGGTGAAATCCTGCAGCTCAACTGTAGAACTGCCATTGAAACTGAATTACTTGAGTACACTTGAGGTAGGCGGAATATGTAGTGTAGCGGTGAAATGCTTAGATATTACATAGAACACCAATTGCGAAGGCAGCTTACTAAGGTGTAACTGACGCTGATGAACGAAAGCGTGGGG |
| P-asv5007 | ACAAGTGGTCATCACGTTTATTGGGCCTAAAGCGTTCGTAGCCGGTTTGGTAAATCTTTTGTGAAATTGTTCGGCTTAACCGGACAGCGCGCAGGGGACACTGCCAAACTCGAGACCGGGAGGCGCAAGAAGTATGCCGTGGGGACTGGTAAAATGGGATAATCCACGGTAGACTACCGATGGCGAAGGCATCTTGCGAGAACGGATCTGACGGTGAGGAACGAAAGCTAGGGG |
| P-asv5008 | GCAAGCGTTATCCGGATTTATTGGGTTTAAAGGGTCCGTAGGCGGATCTGTAAGTCAGTGGTGAAATCTCACAGCTTAACTGTGAAACTGCCATTGATACTGCAGGTCTTGAGTAAATTTGAAGTGGCTGGAATAAGTAGTGTAGCGGTGAAATGCATAGATATTACTTAGAACACCAATTGCGAAGGCAGGTCACTAAGATTTAACTGACGCTGATGTACGAAAGCGTGGGG |
| P-asv5009 | ACAAGTGGGGATCACATTTATTTGGCCTAAAGCGTTCGTAGCCGGCTCGTTAAATCCTTTGTGAAATCGTTTCGTTCAATCAAACGACGTGCGGGGGAGACTGGCGAGCTCGGGACTGGGAGGAGTCAGGAGTATGCAGGGGGGAGCAGTAAAATGCTATAATCCCCTGGAGACTACCGGTGGCGAAGGCGCCTGACCAGAACAGATCCGACGGTGAGGAACGAAAGCTAGGGG |
| P-asv5010 | ACGAGTGATGGCCGCTCTTATTGGGCCTAAAGCGTTCGTAGCCGGCCGAGCAAGTTCGTTGGGAAATCTGCCCGCTTAACGGGCAGGCGTCCGGCGAAAACTGCTCGGCTTGGGGCCGGAAGATCTGAGGGGTACGTCCGGGGTAGGAGTGAAATCCCGTAATCCTGGACGGACCACCGGTGGCGAAAGCGCCTCAGAAAAACGGACCCGACGGTGAGGGACGAAAGCTAGGGT |
| P-asv5011 | CCGAGTGATGGCCGATATTATTGGGCCTAAAGCGTCCGTAGCTGGCTAGACAAGTCCGTTGGGAAATCTGCTCGCTTAACGACCAGGCGTCCAGCGGAAACTGTGTAGCTTGGGACCGGAAGACCTGAGGGGTACGTCTGGGGTAGGAGTGAAATCCCGTAATCCTGGACGGACCGCCGGTGGCGAAAGCGCCTCAGGAGAACGGATCCGACAGTGAGGGACGAAAGCTAGGGT |
| P-asv5012 | GCTAGCGTTGTTCGGAACTACTGGGCGTAAAGGGCGCGTAGGCGGCTTCTTGCGTCGGATGTGAAAGCCCGGGGCTCAACCTCGGAACGGCATTCGATACGGGGGAGCTGGAGACCGGGAGAGGAGAGTGGAATAGCCAGTGTAGAGGTGAAATTCGTAGATATTGGCTGGAACACCGGTGGCGAAAGCGGCTCTCTGGCCCGGTTCTGACGCTGAGGCGCGAAAGCGTGGGG |
| P-asv5013 | GCAAGCGTTATCCGGATTTATTGGGTTTAAAGGGTGCGTAGGCGGAGTCATAAGTCAGTGGTGAAAGTATGCCGCTCAACGGTATAATTGCCATTGAAACTGTGATTCTTGAGTATGGACGAGGCAGGCGGAATGTGTAATGTAGCGGTGAAATGCATAGATATTACACAGAACCCCGATTGCGTAGGCAGCTTGCCAGACCATTACTGACGCTGATGCACGAAAGCGTGGGG |
| P-asv5014 | GCAAGTGATGACCGATCTTATTGGGCCTAAAGCGTCCGTAGCCGGCCGCGCAAGTCCATCGGGAAATCCAGTCGCTTAACGACTGGGCGTCCGGTGGAAACTGCACGGCTTGGGGCCGGAAGACCTCAGGGGTACGTCCGGGGTAGGAGTGAAATCCCGTAATCCTGGACGGACCACCGATGGGGAAACCACCTCAAGAGAACGGATCCGACAGTGAGGGACGAAAGCTAGGGT |
| P-asv5015 | CCAAGTGATGGCCGATATTATTGGGCCTAAAGCGTCCGTAGCTGGCCGAGTAAGTTCGTTGGGAAATCCACTCGCCTAACGAGTGGGCGTCCGGCGAAAACTGCTCGGCTTGGGACCGGAAGACCCAGGGGGTACGTCCGGGGTAGGAGTGAAATCCCGTAATCCTGGACGGACCACCGGTGGCGAAAGCGCCCTGGGAAGACGGATCCGACAGTGAGGGACGAAAGCTAGGGT |
| P-asv5016 | GCTAGCGTTGCTCGGAATTATTGGGCGTAAAGCGCGCGTAGGCGGCCGCATAAGTCTGATGTGAAAGCCCAGGGCTCAACCCTGGAAAGCCATTGGAAACTGTGCGGCTTGAGAACAGGAGGGGTAGGTGGAATTCCTGGTGTAGGAGTGAAATCCGTAGAGATCAGGAGGAACACCGGTGGCGAAGGCGACCTACTGGACTGGATCTGACGCTGAGGCGCGAAAGCGTGGGG |
| P-asv5017 | GCAAGCGTTAATCGGAATTACTGGGCGTAAAGCGCACGTAGGCGGACTGGAAAGTTGGGGGTGAAATCCCGGGGCTCAACCTCGGAACTGCCTTCAAAACTATCAGTCTGGAGTTCGAGAGAGGTGAGTGGAATTCCGAGTGTAGAGGTGAAATTCGTAGATATTCGGAGGAACACCAGTGGCGAAGGCGGCTCACTGGCTCGATACTGACGCTGAGGTGCGAAAGCGTGGGG |
| P-asv5018 | GCTAGCGTTGTTCGGATTTACTGGGCGTAAAGCGCACGTAGGTGGATTGTTAAGTGAGGGGTGAAATCCCAGGGCTCAACCCTGGAACTGCCTTTCATACTGGCAATCTTGAGTTCGGAAGAGGTGAGTGGAATTCCAAGTGTAGAGGTGAAATTCGTAGATATTTGGAAGAACACCGGTGGCGAAGGCGGCTCACTGGTCCGATACTGACACTGAGGTGCGAAAGCGTGGGG |
| P-asv5019 | CCTAGCGTTGTTCGGAATTATTGGGCGTAAAGCGGGTGTAGGCGGACATATAAGTCAGGTGTGAAATCCCAGGGCTCAACCCTGGAAGTGCACTTGATACTGTATGTCTTGAGTGTGGTAGAGGCTATTAGAATTCTTGGTGTAGTGGTGAAATACGTAGATATCAAGAGGAATACCGGAGGCGAAGGCGGATAGCTGGGCCAACACTGACGCTGAGACCCGAAAGCGTGGGG |
| P-asv5020 | GCAAGCGTTAATCGGAATTACTGGGCGTAAAGCGTGCGCAGGCGGTTTTGTAAGACAGGCGTGAAATCCCCGGGCTCAACCTGGGAATTGCGCTTGTGACTGCAAGGCTGGAGTGCGGCAGAGGGGGATGGAATTCCGCGTGTAGCAGTGAAGTGCGTAGATATGCGGAGGAACACCGATTGCGAAGGCAGTCTACTACGTATGTACTGACGCTCATGGACGAAAGCGTGGGG |
| P-asv5021 | GCTAGCGTTGTTCGGAATTACTGGGCGTAAAGCGCGCGTAGGCGGACTGTTAAGTCGGGGGTGAAATCCCGGGGCTCAACCCCGGAACTGCCTTCGATACTGGCAGTCTAGAGGACGGAAGAGGTCAGTGGAATGGCGAGTGTAGAGGTGAAATTCGTAGATATTCGCCGGAACACCAGTGGCGAAGGCGACTGACTGGTCCGTTTCTGACGCTGAGGTGCGAAAGCGTGGGG |
| P-asv5022 | GCAAGCGTTACTCGGAATTACTGGGCGTAAAGCGCGCGTAGGCGGTTTGATAAGTCTGATGTGAAATCCCAGGGCTTAACCCTGGAAGTGCATTGGATACTGTCAGACTTGAGTATAGGAGAGGAGAGTGGAATTCCTGGAGTAGAGGTGAAATTCGTAGATATCAGGAGGAACATCAGTGGCGAAGGCGACTCTCTGGACTATAACTGACGCTGAGGTGCGAAAGCGTGGGT |
| P-asv5023 | GCAAGCGTTATTCGGAATAACTGGGCGTAAAGGGTGCGTAGACGGCTCATTAAGTCCGCTGTTAAATACCCCGGCCTAACCGGGGAATCGCGGTGGAAACTGATAGGCTTGAGGATGGAAGAGAGAAGTGGAATTCTCGGAGTAGCGGTAAAATGCGTAGATCTCGAGAGGAACACCGATGGCGAAGGCAGCTTCTTGGTCCATTTCTGACGTTGATGCACGAAAGCGTGGGG |
| P-asv5024 | GCGAGCGTTAATCGGAATTACTGGGCGTAAAGCGCGCGTAGGCGGTTTGTTAAGTCTGATGTGAAATCCCCGGGCTCAACCTGGGAATTGCATCGGATACTGGCAAGCTAGAATGCGGTAGAGGGGGGTGGAATTCCAGGTGTAGCGGTGAAATGCGTAGATATCTGGAGGAACACCAGTGGCGAAGGCGGCCCCCTGGACTGACATTGACGCTGAGGTGCGAAAGCGTGGGG |
| P-asv5025 | CCAAGCGTTGCTCGGAATCATTGGGCGTAAAGCGGGTGTAGGCGGCCTTGCAAGTCGGAAGTGAAAGCCCTGGGCTCAACCCAGGAAGTGCTTTCGATACTGCTTGGCTTGAGTGTGGGAGAGGATCGTGGAATTCCAGGTGTAGTGGTGAAATACGTAGATATCTGGAGGAACATCGGTGGCGAAGGCGGCGATCTGGCCCAACACTGACGCTCAGACCCGAAAGCGTGGGG |
| P-asv5026 | TCAAGCGTTATCCGGATTTACTGGGCGTAAAGTGTCCGTAGACTGATATATAAGTCCGTTTTCAAATATCCCGGCTCAACCGGGAAAAGGGGATGGATACTGTGTATCTAGAAGTTATTAGAGGTTAGTGGAATTTCCGGTGGAGCGGTGAAATGCGTTGATATCGGAAAGAACGCCGAAAGCGAAAGCAGCTAACTATAATAAACTTGACGTTGAGGGACGACAGTGTGGGG |
| P-asv5027 | GCAAGCGTTAATCGGAATTACTGGGCGTAAAGAGCGCGTAGGTGGTTTGGTAAGTCGGATGTGAAATCCCTGGGCTTAACCTAGGAACTGCATTCGATACTACCAGGCTAGAGTAGAGTAGAGGGAAGTGGAATTTCCGGTGTAGCAGTGAAATGCGTAGATATCGGAAAGAACATCAGTGGCGAAGGCGACTTCCTGGACTCATACTGACACTGAGGTGCGAAAGCATGGGG |
| P-asv5028 | GCAAGCGTTAATCGGAATTATTGGGCGTAAAGCGTGCGTAGGCGGTTCGGAAAGTTAGGTGTGAAATCCCAGGGCTTAACCTTGGAATTGCGCTTAAAACTCCCGAGCTAGAGTACTGTAGAGGATAGTGGAATTTCCGGTGTAGCGGTGAAATGCGTAGAGATCGGAAGGAACACCAGTGGCGAAGGCGACTATCTGGACAGTAACTGACGCTGAGGTACGAAAGCGTGGGT |
| P-asv5029 | GCGAGCGTTACTCGGATTTACTGGGCGTAAAGGGTCCGTAGGCGGCCTTGTGTGTCGGGTGTGAAATCCCACGGCTTAACCGTGGAACTGCGCCCGAAACTACAAGGCTAGAGTATTGGAGAGGTGAGTGGAATTCCAGGTGTAGCGGTGAAATGCGTAGATATCTGGAGGAACACCAACGGCGAAGGCAGCTCACTGGACAATTACTGACGCTGAGGGACGAAAGTGTGGGG |
| P-asv5030 | GCAAGCGTTATCCGGAATCATTGGGTTTAAAGGGTCCGCAGGCGGTCAATTAAGTCAGAGGTGAAATCCAGTCGCTTAACGATTGAACTGCCTTTGATACTGGTTGACTTGAGTCATATGGAAGTAGATAGAATGTGTAGTGTAGCGGTGAAATGCTTAGATATTACACAGAATACCGATTGCGAAGGCAGTCTACCACGTATGTACTGACGCTCATGGACGAAAGCGTGGGG |
| P-asv5031 | GCGAGCGTTGTCCGGATTTATTGGGTTTAAAGGGTGCGTAGGCGGCCCGTTAAGTCCGGGGTGAAAGCCCACTGCTCAACAGTGGAACTGCCCTGGATACTGGCGGGCTTGAGTCCAGACGAGGTTGGCGGAATAGATGCTGTAGCGGTGAAATGCATAGATAGCATCTAGAACCCCGATTGCGTAGGCAGCTGACTAGGCTGGCACTGACGCTGAGGCACGAAAGCGTGGGG |
| P-asv5032 | GCGAGCGTTGTCCGGAATTATTGGGCGTAAAGGGCTCCTAGGCGGTTCAGTAAGTCGACTGTGAAAATCCAAGGCTCAACCTTGGGACGCCAGTCGATACTGCTGTGACTCGAGTTCGGTAGAGGAGTGTGGAATTCCTGGTGTAGCGGTGAAATGCGCAGATATCAGGAGGAACACCAACGGCGAAGGCAGCACTCTGGGCCGATACTGACGCTGAAGAGCGAAAGCGTGGGG |
| P-asv5033 | GCTAGCGTTGTTCGGAATTACTGGGCGTAAAGCGCGCGTAGGCGGTTTTTTAAGTCAGAGGTGAAAGCCCAGTGCTCAACACTGGAACTGCCTTTGAAACTGGAAAACTTGAATCTTGGAGAGGTCAGTGGAATTCCGAGTGTAGAGGTGAAATTCGTAGATATTCGGAAGAACACCAGTGGCGAAGGCGACTGACTGGACAAGTATCGACGCTGAGGTGCGAAAGCGTGGGG |
| P-asv5034 | TCCCTCTTTTTTCTTCCTTACTGGGCGTAAAGCGTGCGTAGTCGGTATTGAGAGTCACGGGTGAAATCCCAGGGCTTAACCCTGGAACTGCCTGTGAGACCTCAGTACTAGAGTGTGAGAGGGGATAGTGGAATACCCAGTGTAGCGGTGAAATGCGTAGAGATTGGGTGGAACACCGGTGGCGAAGGCGGCTATCTGGCTCACAACTGACGATCAGGCACGAAAGCGTGGGG |
| P-asv5035 | GCAAACGTTGTTCGGAATTACTGGGCGTAAAGCGCGTGTAGGCGGCACGTTAAGTCGACTGTGAAAGCCCATGGCTCAACCATGGAAGTGCAGTCGATACTGGCATGCTCGAGTCCCGGAGAGGGTGGTGGAATTCCCAGTGTAGAGGTGAAATTCGTAGATATTGGGAGGAACACCGGTGGCGAAGGCGACCACCTGGACGGGTACTGACGCTGAGACGCGAAAGCGTGGGG |

| | |
|---|---|
| P-asv5036 | GCAAGCGTTACTCGGAATCACTGGGCGTAAAGAGCATGTAGGCGGATAGATAAGTTGGAAGTGAAATCCACTGGCTTAACCAGTGAACTGCTCCCAAAACTGTCTATCTAGAATATGGGAGAGGTAGATGGAATTTCTGGTGTAGGGGTAAAATCCGTAGAGATCAGAAGGAATACCGATTGCGAAGGCGATCTACTGGAACATTATTGACGCTCAGATGCGAAAGCGTGGGG |
| P-asv5037 | GCGAGCGTTGTCCGGAATTATTGGGCGTAAAGAGCGTGTAGGCGGCTTGGTAAGTCTGCTGTGAAACCTTGGGGCTCAACCCCAAGCGTGCAGTGGATACTGTCAAGCTAGAGGATGGTAGAGGCCAGTGGAATTCCCAGTGTAGCGGTGAAATGCGCAGATATTGGGAGGAACACCAGTAGCGAAGGCGGCTGGCTGGGCCATTCCTGACGCTGAGACGCGAAAGCTAGGGG |
| P-asv5038 | GTTAGCGTTGTTCGGAATTACTGGGCGTAAAGCGTACGTAGGCGGATTAATAAGTTAGAGGTGAAATCCCAGGGCTCAACCCTGGAACTGCCTTTAAAACTGTTAGTCTTGAGATCGAGAGAGGTGAGTGGAATTCCAAGTGTAGAGGTGAAATTCGTAGATATTTGGAGGAACACCAGTGGCGAAGGCGGCTCACTGGCTCGATACTGACGCTGAGGTACGAAAGTGTGGGG |
| P-asv5039 | GCAAGCGTTATCCGGAATTATTGGGCGTAAAGCGAGTCTAGGCGGTTTGTTAAGTCAGATGTGAAAATGCGGGGCTCAACTCCGTATGGCCTTTGATACTGGCAAACTAGAGTACTGGAGAGGTGGGCGGAACTACAAGTGTAGAGGTGAAATTCGTAGATATTTGTAGGAATGCCGATAGTGAAGACAGCTCACTGGACAGATACTGACGCTAAAGCTCGAAAGCGTGGGG |
| P-asv5040 | GCAAGCGTTATCCGGATTTATTGGGTTTAAAGGGTCCGTAGGCGGATTAATAAGTCAGTGGTGAAAGCCCGCAGCTTAACTGTGGAACTGCCATTGATACTGTTAGTCTTGAGTGTAGTTGAAGTGGCTGGAATGAGTAGTGTAGCGGTGAAATGCATAGATATTACTCAGAACACCAATTGCGAAGGCAGGTCACTAAGTTACAACTGACGCTGATGGACGAAAGCGTGGGG |
| P-asv5041 | TCGAGTGGTCAGGACGTTTATTGGGCCTAAAGCATCCGTAGCCGGCTTTGCAAGTCTTCGGTTAAATCCACCTGCTCAACAGATGGGCCGCTGGAGATACTACAAAGCTAGGGAGTGGGAGAGGCAGACGGTATTCAGTGGGTAGGGGTAAAATCCTCTGATCCATTGAGGACCACCAGTGGCGAAGGCGGTCTGCCAGAACACGTTCGACGGTGAGGGATGAAAGCTGGGGG |
| P-asv5042 | GCAAGCGTTATTCGGATTCACTGGGCGTAAAGGGTGCGCAGGCGGCCATGTGTGTCAGGTGTGAAAGCCCGAGGCTCAACCTCGGAATTGCGCCTGAAACTACATGGCTAGAGTATTGGAGAGGGTAGCGGAATTCACGGTGTAGCAGTGAAATGCGTAGATATCGTGAGGAACACCAGAGGCGAAGGCGGCTACCTGGACAATTACTGACGCTCAGGCACGAAAGCGGGGGG |
| P-asv5043 | GCAAGCGTTGTTCGGAATTACTGGGCGTAAAGCGTGCGTAGTCGGTATTGAGAGTCGCGGGTGAAATCCCAGGGCTTAACCCTGGAACTGCCTGCGAGACCTCAGTACTAGAGTGTGAGAGGGGATATTGGAATACCCAGTGTAGCGGTGAAATGCGTAGAGATTGGGTAGAACACCGGTGGCGAAGGCGGATATCTGGCTCACAACTGACGATCAGGCACGAAAGCGTGGGG |
| P-asv5044 | GCTAGCGTTGTTCGGAATTACTGGGCGTAAAGCGCACGTAGGCGGACTGTCAAGTCAGGGGTGAAATCCCGGGGCTCAACCTCGGAACTGCCTTTGATACTGTCAGTCTAGAGTCCGGAAGAGGTGGGTGGAATTCCTAGTGTAGAGGTGAAATTCGCAGATATTAGGAAGAACACCGGTGGCGAAGGCGGCCCACTGGTCCGGAACTGACGCTGAGGTGCGAAAGCGTGGGG |
| P-asv5045 | GCGAACGTTATTCGGAATCACTGGGCTTAAAGCGCGTGTAGGCGGACGACCACGTCGACGGCTGAAAGCCCCCGGCTCAACCGGGGGACCGGCCTCGATACGGGCCGTCTGGAGGGGCGTAGGGGGAGCTGGAACTCACGGTGGAGCGGTGAAATGCGTTGAGATCGTGAGGAACGCCCGCGGCGAAAGCGAGCTCCTGGACGCCTGCTGACGCTGAGACGCGAAAGCCAGGGG |
| P-asv5046 | ACGAGCGTTAATCGGAATCACTGGGCTTAAAGGGTGCGCAGGCGGCTCGAAAAGCGTTCCGTGAAATCCCCCGGCTCAACCGGGGAACTGCGGAGCGAACTGCCGAGCTTGAGGCAAGTAGGGGCGGGCAGAACTGTAGGTGGAGCGGTGAAATGCGTAGATATCTACAGGAATGCCGATGGTGAAGACGGCCCGCTGGGCTTGACCTGACGCTCATGCACGAAAGCGTGGGG |
| P-asv5047 | CCGAGCGTTATCCGGATTTATTGGGTTTAAAGGGTGCGTAGGCGGCTTATTAAGTCAGGGGTGAAATACGGCAGCTCAACTGTCGCAGTGCCTTTGATACTGATAAGCTTGAATGTGGTTGAAGATGGCGGAATGAGACAAGTAGCGGTGAAATGCATAGATATGTCTCAGAACTCCGATTGCGAAGGCAGCTATCTAAGCCATTATTGACGCTGATGCACGAAAGCGTGGGG |
| P-asv5048 | CCGAGTGGTAGTCGTTATTATTGGGCTTAAAGTGTTCGTAGCCTGCCGTGTAAGTCCGCTGTGAAACCCACACGCTCAACGTGTGGATGTGCAGCGGATACTGCATGGCTAGGGACCGGACGGCGTCGAGGCTATTCCAGGGGTAGGAGTGAAATCCAAAAATCCCTGGAGGAGCACCTGTGGCGAAAGCGCTCGACGAGTACGGATCCGACGGTGAGGAACGAAAGCTGGGGG |
| P-asv5049 | GCAAGCGTTGTCCGGATTTACTGGGCGTAAAGGATGCGTAGGTGGACATTTAAGTCAGATGTGAAATACCCGAGCTCAACTTGGGTGCTGCATTTGAAACTGGATGTCTAGAGTGCAGGAGAGGAAAGCGGAATTCCTAGTGTAGCGGTGAAATGCGTAGAGATTAGGAAGAACACCAGTGGCGAAGGCGGCTTTCTGGACTGTAACTGACACTGAGGCATGAAAGCGTGGGG |
| P-asv5050 | GCAAGCGTTATTCGGAATTATTGGGCGTAAAGGGCGCGCAGGCGGTCTTGTCCGTCAGGTGTGAAAGCTCGGGGCTCAACCCCGGAAGTGCACTTGAAACAGCAAGACTTGAATACGGGAGAGGAAAGCGGAATTCCTGGTGTAGAGGTGAAATTCGTAGATATCAGGAGGAACACCGATGGCGAAGGCAGCTTTCTGGACCGATATTGACGCTGAGGCGCGAAGGCGTGGGT |
| P-asv5051 | GCAAGCGTTGTCCGGATTTACTGGGCGTAAAGGGAGCGTAGGTGGATATTTAAGTGGGATGTGAAATACTCGGGCTTAACCTGAGTGCTGCATTCCAAACTGGATATCTAGAGTGCAGGAGATGAAAGTAGAATTCCTAGTGTAGCGGTGAAATGCGTAGAGATTAGGAAGAATACCAGTGGCGAAGGCGACTTTCTGGACTGTAACTGACACTGAGGCTCGAAAGCGTGGGG |
| P-asv5052 | GCAAGCGTTGTTCGGAATTACTGGGCGTAAAGAGCACGTAGGCGGGTTCGTCAGCCATTTGTGAAAGCCCAAGGCTCAACCTTGGAATTGCAGGTGGAACTGCGAATCTTGAATACCTGAGAGGATGAGGGAATTCCTGGTGTAGAAGTGAAATTCGTAGATATCAGGAGGAACACCCGAGGCGAAGGCGCTCGTCTGGCAGTGTATTGACGCTGAGGTGCGAAAGCGTGGGG |
| P-asv5053 | GCAAGCGTTGTTCGGAATTACTGGGCGTAAAGCGCGTGTAGGCGGCGCTATAAGTCGGATGTGAAAGCCCTCGGCTCAACCGAGGAAGTGCATCCGATACTGTAGTGCTTGAGTGTCGGAGGGGGTGGCGGAATTCCTGGTGTAGAGGTGAAATTCGTAGATATCAGGAGGAACACCTGTGGCGAAGGCGGCCACCTGGACGACAACTGACGCTGAGACGCGAAAGCGTGGGT |
| P-asv5054 | CCGAGTGATGGCCGATATTATTGGGCCTAAAGCGTCCGTAGCTGGCCGGACAAGTCCGTTGGGAAATCTGTTCGCTTAACGAGCAGGCGTCCAGCGGAAACTGTTCGGCTTGGGACCGGAAGACCTGAGGGGTACGTCCGGGGTAGGAGTGAAATCCTGTAATCCTGGACGGACCACCGGTGGCGAAAGCGCCTCAGGAGGACGGATCCGACAGTGAGGGACGAAAGCTAGGGT |
| P-asv5055 | GCAAGCGTTGTTCGGAATTATTGGGCGTAAAGAGCATGTAGGCGGACTGTTAAGTCTGGTGTGAAAGCCCGGGGCTCAACCCCGGAAGTGCATTGGATACTGGCAGTCTTGAGTATGGGAGAGGAAAGTGGAATTCCGAGTGTAGGAGTGAAATCCGTAGATATTCGGAGGAACACCAGTGGCGAAGGCGGCTTTCTGGACCAATACTGACGCTGAGATGCGAAAGCGTGGGG |
| P-asv5056 | GCGAGCGTTATCCGGATTCATTGGGCGTAAAGCGCGCGTAGGCGGCCCGGCAGGCAGGGGGTCAAATGGCGGGGCTCAACCCCGTCCCGCCCCCTGAACCGCCGGGCTCGGGTCCGGTAGGGGAGGGTGGAACACCCGGTGTAGCGGTGGAATGCGCAGATATCGGGTGGAACACCGGTGGCGAAGGCGGCCCTCTGGGCCGAGACCGACGCTGAGGCGCGAAAGCTGGGGG |
| P-asv5057 | GCAAGCGTTACTCGGAATTACTGGGCGTAAAGCGTGCGTAGGTGGTGATTTAAGTCTGGTGTGAAAGCCCTGGGCTCAACCTGGGAATGGCATTGGAAACTGGGTCACTGGAGTGCGGTAGAGGATGGCGGAATTCCCGGTGTAGCAGTGAAATGCGTAGAGATCGGGAGGAACATCCGTGGCGAAGGCGGCCATCTGGACCAGCACTGACACTGAGGCACGAAAGCGTGGGG |
| P-asv5058 | GCAAGCGTTGTCCGGAATTATTGGGCGTAAAGCGCGCGCAGGCGGTTTCTTAAGTCTGATGTGAAAGCCCACGGCTCAACCGTGGAGGGTCATTGGAAACTGGGGAACTTGAGTACAGAAGAGGAGAGTGGAATTCCACGTGTAGCGGTGAAATGCGTAGAGATGTGGAGGAACACCAGTGGCGAAGGCGACTCTCTGGTCTGTCACTGACGCTGAGGCGCGAAAGCGTGGGG |
| P-asv5059 | GCAAGCGTTAATCGGAATTACTGGGCGTAAAGCGCGCGTAGGTGGTCTGTTAAGTCGGATGTGAAAGCCCCGGGCTCAACCTGGGAACTGCATTCGATACTGGCAGACTAGAGTGCGGGAGAGGGAGGTAGAATTCCATGTGTAGCGGTGAAATGCGTAGATATGTGGAGGAATACCAGTGGCGAAGGCGGCCCCCTGGCTCGACACTGACACTGAGGTGCGAAAGCGTGGGG |
| P-asv5060 | CCAAGCGTTGTTCGGAATTACTGGGCGTAAAGCGAGTGTAGGCGGTCCGTCAAGTTGGTTGTGAAATCTCCTGGCTCAACTGGGAGGGTGCGACCAAAACTGACGGACTAGAGTCTGGGAGAGGAGAGTGGAATTCCTGGTGTAGCGGTGAAATGCGTAGATATCAGGAGGAACACCGGTGGTGAAGACGGCTCTCTGGACCGGTACTGACGCTGAGACTCGAAAGCGTGGGG |
| P-asv5061 | GCAAGCGTTGTTCGGAATCACTGGGCTTAAAGCGTATGTAGGCGGTCATGCAAGTCAGGAGTGAAATCCCGGGGCTCAACCCCGGAATTGCTTTTGAAACTGCACGACTAGAATACCGGAGAGGTTGGGAGAATTCCTAGTGTAGAGGTGAAATTCGCAGATATTAGGAGGAATACCAGTGGCGTAGGCGCCCAACTGGACGGTTATTGACGCTCAGATACGAAAGCGTGGGG |
| P-asv5062 | GCTAGCGTTGTTCGGAATTACTGGGCGTAAAGCGCGCGTAGGCGGCCTTGCAAGTTGGGGGTGAAATCCCGGAGCTCAACTCCGGAACTGCCTTCAAAACTGCAAAGCTGGAGGATGCGAGAGGTGAGCGGAATTCCGAGTGTAGAGGTGAAATTCGTAGATATTCGGAAGAACACCAGTGGCGAAGGCGGCTCACTGGCGCATTTCTGACGCTGAGGCGCGAAAGCGTGGGG |
| P-asv5063 | GCAAGCGTTGTCCGGATTTATTGGGCGTAAAGAGCTCGTAGGCGGCCTGTCGCGTCGACTGTGAAAACCCGCGGCTTAACTGCGGGAGTGCAGTCGATACGGGCAGGCTAGAGTTCGGTAGGGGAGACTGGAATTCCTGGTGTAGCGGTGAAATGCGCAGATATCAGGAGGAACACCGGTGGCGAAGGCGGGTCTCTGGGCCGATACTGACGCTGAGGAGCGAAAGCGTGGGG |
| P-asv5064 | TCAAGCGTTATCCGGATTTATTGGGTTTAAAGGGTGCGTAGGCGGTTTGATAAGTTAGAGGTGAAATCCCGGGGCTTAACTCCGGAACTGCCTCTAATACTGTTAGACTAGAGAGTAGTTGCGGTAGGCGGAATGTATGGTGTAGCGGTGAAATGCTTAGAGATCATACAGAACACCGATTGCGAAGGCAGCTTACCAAACTATATCTGACGTTGAGGCACGAAAGCGTGGGG |
| P-asv5065 | GCAAGCGTTGTCCGAAATTATTGGGCGTAAAGAGCTCGTAGGCGGTTTGTCGCGTCTGCTGTGAAAACGCAAGGCTTAACCTTGCGCCAGCAGTGGGTACGGGCAGACTATAGTGCGGTAGGGGAGACTGGATTTCCTGGTGTAGCGGTGGAATGCGCAGATATCAGGAGGAACGCCGGTGGCGAAGGCAGGTCTCTGGGCCGTTACTGACGCTGAGGAGCGAAAGCATGGGG |
| P-asv5066 | GCAAGCGTTAATCGGAATTACTGGGCGTAAAGCGCGCGTAGGTGGTTCAGTAAGTTGGAAGTGAAATCCCCGGCCTCAACCTGGGAACTGCTTTCAAAACTGCTGAGCTAGAGTACGGTAGAGGGTGGTGGAATTTCCTGTGTAGCGGTGAAATGCGTAGATATAGGAAGGAACACCAGTGGCGAAGTCGACCACCTGGACTGATACTGACACTGAGGTGCGAAAGCGTGGGG |
| P-asv5067 | GCGAGCGTTAATCGGAATTACTGGGCGTAAAGCGTGCGTAGACGGTCGTGCAAGTCGGGTGTGAAAGCCCCAGGCTCAACCTGGGAATTGCATTCGAAACTGCTCGGCTAGGGTGCGGAAGAGGGAAGCGGAATTTCCGGTGTAGCGGTGAAATGCGTAGATATCGGAAGGAGCACCAGTGGCGAAAGCGGCTTCCTGGTCCGACACCGACGTTCAGGCACGAAAGCGTGGGG |
| P-asv5068 | GCGAGCGTTATCCGGAATCATTGGGTTTAAAGGGTCCGTAGGCGGGCGTATAAGTCAGAGGTGAAAGCACTGAGCTCAACTGAGTAACTGCCTTTGAAACTGTATGTCTTGAATGTTTGTGAAGTAGCTGGAATGTGTAGTGTAGCGGTGAAATGCATAGATATTACACAGAACACCGATTGCGAAGGCATGTTACTAACAGATTATTGACGCTGATGGACGAAAGCGTGGGG |
| P-asv5069 | CCGAGCGTTATCCGGAATCATTGGGTTTAAAGGGTCCGTAGGCGGGAGAATCAGTCAGTGGTGAAAGTCTGTGGCTCAACCATAGAATTGCCATTGAAACTGTTTTCCTTGAGTCATTATGAAGTGGTTAGAACATGTAGTGTAGCGGTGAAATGCATAGATATTACATAGAATACCGATTGCGAAGGCAGATCACTAATAATGTACTGACGCTGAGGGACGAAAGCGTAGGT |
| P-asv5070 | GCGAGCGTTATTCGGAATTATTGGGCGTAAAGGGCGCGTAGGCGGCCGGATAAGTCAAAGGTGAAATCCCTCGGCTCAACTGAGGAACTGCCTTTGATACTGTCAGGCTTGAGTCCGGGAGGGGGTAGCGGAATTCCCAGTGTAGCGGTGAAATGCGTAGATATTGGGAGGAACACCAGTGGCGAAGGCGGCTACCTGGACCGGAACTGACGCTGAGGCGCGAAAGCGTGGGG |
| P-asv5071 | GCTAGCGTCGTTCGGAATTATTGGGCGTAAAGGGTGCGTAGGCGGTGCGGTAAGTCTGTGGTGAAATCTCCGGGCTCAACTCGGAGCCTGCCACGGAAACTGCCGTGCTTGAGTATGGGAGAGGTGAGTGGAATTCCCGGTGTAGCGGTGAAATGCGTAGATATCGGGAGGAACACCTGTGGCGAAAGCGGCTCACTGGACCATAACTGACGCTGATGCACGAAAGCTAGGGG |
| P-asv5072 | GCCAGCGTTGTTCGGAATTATTGGGCGTAAAGGGTGCGTAGGCGGTGCGGTAAGTCTTCTGTGAAATCTCCGGGCTCAACCCGGAGCCTGCAGAGGAAACTGCCGTGCTGGAGTATGGGAGAGGTGAGTGGAATTCCCGGTGTAGCGGTGAAATGCGTAGATATCGGGAGGAACACCTGTGGCGAAAGCGGCTCACTGGACCATTACTGACGCTGATGCACGAAAGCTAGGGG |
| P-asv5073 | GCTAGCGTTGTTCGGAATTACTGGGCGTAAAGGGCGCGTAGGCGGTCTGCTAAGTTGGATGTGAAAACTCTGGGCTTAACCCAGAGCCTGCATTCAATACTGACAGGCTAGAGTTCTGGAGGGGATAGCGGAATTCCTGGTGTAGCGGTGAAATGCGTAGATATCAGGAGGAACACCGGTGGCGAAGGCGGCTATCTGGACAGAGTCTGACGCTGAGGCGCGAAAGCTAGGGG |
| P-asv5074 | GCAAGCGTTGTCCGGATTTACTGGGCGTAAAGAGCGCGCAGGCGGTCGTGCAAGTCGCGTGTGAAAGCCCCCGGCTCAACTGGGGAGGGTCACGCGATACTGCTCGACTCGAAGGCAGGAGAGGGAAGTGGAATTCCCGGTGTAGTGGTGAAATGCGTAGATATCGTAAGGAAGACCAGTGGCGAAGGCGGGTCTCTGGACAGTTCCTGACGCTGTGGCACGAAGGCCAGGGG |

| | |
|---|---|
| P-asv5075 | GCGAGCGTTGTCCGGAGTTACTGGGCGTAAAGGGCACGCAGGCGGTCTGCCACGTTCGCCATGACAGTGTCGAGCTTCACTCGACAAGGGGGGCGAAAACGAGCGGACTTGAGACTCAGAGAGGGACCTGGAATTCCGGGTGGAGTGGTGAAATGCGTAGATATCCGGAGGAAGACCAAAGGCGAAGGCAGGGTCCTGGCTGAGGGCTGACGCTGAGGTGCGAAAGCCAGGGG |
| P-asv5076 | GCAAGCGTTGTTCGGAATTACTGGGCGTAAAGGGCGCGTAGGCGGCCTTCTAAGTCAGACGTGAAATCCCCGGGCTCAACCCGGGAACTGCGTCTGATACTGGGAGGCTTGAATCCGGGAGAGGGATGTGGAATTCCAGGTGTAGCGGTGAAATGCGTAGATATCTGGAGGAACATCGGTGGCGAAGGCGGCATCCTGGACCGGTATTGACGCTGAGGCGCGAAAGCCAGGGG |
| P-asv5077 | GCGAGCGTTGTCCGGATTTATTGGGCGTAAAGAGCTCGCAGGCGGCTTGTTGCGTCGACTGTGAAAACCCGCGGCTCAACCGTGTGGTCTGCAGTCGATACGGGCAGGCTAGAATTCGGTAGGGGAGACTGGAATTCCTGGTGTAGCGGTGAAATGCGCAGATATCAGGAGGAACACCGGTGGCGAAGGCGGGTCTCTGGGCCGATATTGACGCTGAGGAGCGAAAGCGTGGGG |
| P-asv5078 | GCGAGCGTTAATCGGAATTACTGGGCGTAAAGCGCGCGTAGGCGGCCAGGCAAGTCGGGTGTGAAGGGCCTGGGCTCAACCTGGGAATTGCATTCGAAACTATCTGACTCGAGTATGGGAGAGGGAAGCGGAATTTCCGGTGTAGCGGTGAAATGCGTAGATATCGGAAGGAACACCAGTGGCGAAGGCGGCTTCCTGGACTAATACTGACGCTGAGGTGCGAAAGCGTGGGG |
| P-asv5079 | GCAAGCGTTGTCCGGAATTATTGGGCGTAAAGAGCTCGTAGGCGGTCTGTCGCGTCGGCTGTGAAAACTCGGGGCTCAACCCCGAGCCTGCAGTCGATACGGGCAGACTAGAGTGCGGTAGGGGAGACTGGAATTCCTGGTGTAGCGGTGAAATGCGCAGATACCAGGAGGAACACCGGTGGCGAAGGCGGGTCTCTGGGCCGTAACTGACGCTGAGGAGCGAAAGCGTGGGG |
| P-asv5080 | GCGAGCGTTGTCCGGAATCATTGGGCGTAAAGGGCGCGTAGGTGGCCCGCTCAGCGTGCGGTGAAAGCTCGGGGCTCAACCCCGAGTCGGCCGTGCGAACTGGTGGGCTCGAGCACTGTAGAGGCAGGCGGAATTCCGGGTGTAGCGGTGGAATGCGTAGAGATCCGGAAGAACACCGGTGGCGAAGGCGGCCTGCTGGGCAGTAGCTGACACTGAGGCGCGACAGCGTGGGG |
| P-asv5081 | GCAAGCGTTGTCCGGAATTATTGGGCGTAAAGAGCTCGTAGGCGGCTTGTCGCGTCGACTGTGAAAACCCGCGGCTCAACTGCGGGCCTGCAGCCGATACGGGCAGGCTAGAGTTCGGTAGGGGAGACTGGAATTCCTGGTGTAGCGGTGAAATGCGCAGATATCAGGAGGAACACCGGTGGCGAAGGCGGGTCTCTGGGCCGATACTGACGCTGAGGAGCGAAAGCGTGGGG |
| P-asv5082 | GCGAGCGTTGTCCGGAATCACTGGGCGTAAAGGGCGCGTAGGCGGCCGTGTAAGTCACTTGTGAAATCCCGGGGCTCAACTCCGGGGCTGCAGGTGAGACTGCATGGCTAGAGCACGGTAGAGGCGAGTGGAATTCCCGGTGTAGCGGTGGAATGCGTAGATATCGGGAAGAACACCGGTGGCGAAGGCGGCTCGCCGGGCCGTTGCTGACGCTGAGGCGCGAAAGCGTGGGG |
| P-asv5083 | GCGGGCGTTAATCGGAATTACTGGGCGTAAAGCGCGCGCAGGCGGTCCTTCAAGTCAGAGGTGAAAGCCCAGAGCTCAACTCTGGAACGGCCTTTGAGACTAGAGGACTTGGGTACGGGAGAGGTGAGTGGAATTCCGAGTGTAGAGGTGAAATTCGTAGATATTCGGAAGAACACCAGTGGCGAAGGCGGCTCACTGGCCCGTTACCGACGCTCAGGCGCGACAGCGTGGGG |
| P-asv5084 | GCTAGCGTTGTTCGGATTTACTGGGCGTAAAGCGCGTGTAGGCGGGCCGTTAAGTTGGGGGTGAAAGCCCGAGGCTCAACCTCGGAACTGCCTTCAAAACTGGCGGCCTGGAGCAGATGAGAGGCGAGTGGAATTCCGAGTGTAGAGGTGAAATTCGTAGATATTCGGAGGAACACCAGTGGCGAAGGCGACTCGCTGGCATCTTGCTGACGCTGAGACGCGAAAGCGTGGGG |
| P-asv5085 | GCAAGCGTTACTCGGAATTACTGGGCGTAAAGCGTGCGTAGGTGGTGAGTTAAGTCTGTCGTGAAAGCCCCGGGCTCAACCTGGGAATGGCGATGGATACTGGCTCGCTAGAGTGCGGTAGAGGGTGGTGGAATTCCCGGTGTAGCAGTGAAATGCGTAGAGATCGGGAGGAACATCAGTTGCGAAGGCGGCCACCTGGACCAGCACTGACACTGAGGCACGAAAGCGTGGGG |
| P-asv5086 | GCAAGCGTTGTCCGGATTTATTGGGCGTAAAGAGCTCGTAGGCGGTTTGTCGCGTCGGATGTGAAAACCCAGGGCTCAACCCTGGGCCTGCATTCGATACGGGCAAACTAGAGTATGGCAGGGGAGACTGGAATTCCTGGTGTAGCGGTGAAATGCGCAGATATCAGGAGGAACACCGGTGGCGAAAGCGGCTCACTGGATCTCTTCTGACGCTGAGGCACGAAAGCTAGGGT |
| P-asv5087 | GCAAACGTTGCTCGGAATTATTGGGCGTAAAGCGCACGTAGGCGGCCTTTCAAGTCGAATGTGAAAGCCCTGGGCTCAACCCAGGAAGTGCATCCGAAACTGATTGGCTTGAATGCCGGAGGGGCCCAGAGAATTCCCGGTGTAGAGGTGAAATTCGTAGATATCGGGAGGAATACCAGTGGCGAAGGCGCTGGGCTGGACGGTCACTGACGCTGAGGTGCGAAAGCGTGGGG |
| P-asv5088 | CCGAGCGTTGTCCGGAATTACTGGGCGTGAAGCGCGCGCAGGCGGCCCTCTGCGCCCGGCGTGAAAGCCCCCGGCTCAACCGGGGAGCGTCGTCGGGGACGGAGGGCTTGAGGGCGAGAGGGGCCGGTGGAATCCCTGGTGTAGTGGTGAAATGCGTAGAGATCAGGGGGAACACCCGTGGCGAAGGCGGCCGGCTGGCTCGACCCTGACGCTGAGGCGCGAAGGCGTGGGG |
| P-asv5089 | GCAAGCGTTGTCCGGAATTATTGGGCGTAAAGCGCGCGCAGGCGGTCTTTTAAGTCTGATGTGAAATCCCGCGGCTTAACCGCGGGCGGTCATTGGAAACTGGAGGACTTGAGTGCAGAAGAGGAGAGTGGAATTCCACGTGTAGCGGTGAAATGCGTAGAGATGTGGAGGAACACCAGTGGCGAAGGCGACTCTCTGGTCTGTAACTGACGCTGAGGCGCGAAAGCGTGGGT |
| P-asv5090 | GCAAACGTTGCTCGGAATTACTGGGCGTAAAGCGCGTGTAGGCGGGTCGGCAAGTCGGGCGTGAAATCCCTAGGCTCAACCTAGGAACTGCGCTCGAAACTGCCGGTCTTGAGTGGCGGAGAGGAAAGCGGAATTCCTGGTGTAGAGGTGAAATTCGTAGATATCAGGAGGAACACCGGTGGCGAAGGCGGCTTTCTGGACGTCTACTGACGCTGAGACGCGAAAGCGTGGGG |
| P-asv5091 | GCAAACGTTATTCGGAATTACTGGGCGTAAAGAGTGCGTAGGCGGTTCTCCAAGTCCGTTGTGAAAGCCCCGGGCTCAACCTGGGAATGGCAGTGGAAACTAGGGGACTTGAATCTGGCAGAGGGGGGTGGAATGCCGCGTGTAGCAGTGAAATGCGTAGAGATGCGGCGGAACACCGATGGCGAAGGCAGCCCCCTGGGCTGAGATTGACGCTCAGGCACGAAAGCGTGGGG |
| P-asv5092 | GCTAGCGTTGTTCGGAATTACTGGGCGTAAAGCGCACGTAGGCGGCGATTCAAGTCAGAGGTGAAAGCCCGGGGCTCAACCCCGGAACTGCCTTTGAAACTAGATTGCTAGAATCCTGGAGAGGTTAGTGGAATTCCGAGTGTAGAGGTGAAATTCGTAGATATTCGGAAGAACACCAGTGGCGAAGGCGACTAACTGGACAGGTATTGACGCTGAGGTGCGAAAGAGTGGGG |
| P-asv5093 | GCAAGCGTTATTCGGAATTATTGGGCGTAAAGCGCTCGTAGGCGGGACTGCAAGTCCGTGTTGAAAGACCTCGGCTCAACTGAGGGACCGGCTCGGATACTGTGGTTCTTGAGGCAATCAGAGGGTGATGGAATTCCCGGTGTAGCGGTGAAATGCGTAGATATCGGGAGGAACACCAGTGGCGAAGGCGATCACCTGGGGTTGTTCTGACGCTGAGGAGCGAAAGCTAGGGG |
| P-asv5094 | GCAAGCGTTACTCGGAATTACTGGGCGTAAAGCGTGCGTAGGCGGTTTGTTAAGTCCGCTGTGAAAGCCCTGGGCTCAACCTGGGAATTGCAGTGGATACTGGCTAGCTAGAGTGCGGTAGAGGATGGTGGAATTCCCGGTGTAGCGGTGAAATGCGTAGAGATCGGGAGGAACACCCGTGGCGAAGGCGGCCATCTGGACCAGCACTGACGCTGAGGCACGAAAGCGTGGGG |
| P-asv5095 | GCGAGCGTTGTCCGGATTTATTTGGCGTAAAGGGCTCGTAGGTGGTTGATCGCGTCGGAAGTGTAATCTTGGGGCTTAACCCTGAGCGTGCTTTCGATACGGGTTGACTTGAGGAAGGTAGGGGAGAATGGAATTCCTGGTGGAGCGGTGGAATGCGCAGATATCAGGAGGAACACCAGTGGCGAAGGCGGTTCTCTGGGCCTTTCCTGACGCTGAGGAGCGAAAGCGTGGGG |
| P-asv5096 | GCAAGCGTTATCCGGATTCACTGGGTTTAAAGGGTGCGTAGGTGGGCTGGTAAGTCAGTGGTGAAATCTCCGGGCTTAACTCGGAAACTGCCATTGATACTATCAGTCTTGAATACCGTGGAGGTCAGTAGAATATGTCATGTAGCGGTGAAATGCTTAGATATGACATAGAATACCAATTGCGAAGGCAGCTGGCTACACGAATATTGACACTCAGGCACGAAAGCGTGGGG |
| P-asv5097 | GCAAACGTTGCTCGGAATTATTGGGCGTAAAGCGCACGTAGGCGGCTGTCTAAGTCGGATGTGAAATCCCTCGGCTTAACCAAGGAAGTGCACCCGAAACTGGATAGCTTGAGTACAGAAGAGGATCGCGGAATTCCCGGTGTGGAGGTGAAATTCGTAGATATCGGGAGGAACACCAGTGGCGAAGGCGGCGATCTGGGCTGATACTGACGCTGAGGTGCGAAAGCGTGGGG |
| P-asv5098 | GCAAGCGTTAATCGGAATTACTGGGCGTAAAGCGTGCGTAGGCGGTTCGTTAAGTCTGCTGTGAAAGCCCCGGGCTCAACCTGGGAATGGCAATGGATACTGGCGAGCTAGAGTGTGTCAGAGGATGGTGGAATTCCCGGTGTAGCGGTGAAATGCGTAGAGATCGGGAGGAACATCAGTGGCGAAGGCGGCCATCTGGGACAACACTGACGCTGAAGCACGAAAGCGTGGGG |
| P-asv5099 | GCAAGCGTTAATCGGAGTTACTGGGCGTAGAGGGCGCGTAGGCGGTTAAGTGAGTGTGATGTGAAAGCCCTGGGCTTAACCTAGGAAGTGCATCGCAAACGACTTAACTGGAGTATAAGAGAGGGTGGCGGAATTTCCGGTGTAGCGGTGAAATGCGTAGAGATCGGAAGGAACGTCAATGGCGAAGGCAGCCACCTGGCTTAATACTGACGCTGAGGCGCGAAAGCGTGGGG |
| P-asv5100 | GCGAGCGTTGTCCGGAATCATTGGGCGTAAAGCGCGTGTAGGCGGTTCGGTAAGTCTGCTGTGAAAGTCCAGGGCTCAACCCTGGGATGCCGGTGGATACTGCCGAACTAGAGTGCGGAAGAGGCGAGTGGAATTCCCGGTGTAGCGGTGAAATGCGCAGATATCGGGAGGAACACCAATTGCGAAGGCAGCTCGCTGGGACGTTACTGACGCTGAGACGCGAAAGCGTGGGG |
| P-asv5101 | GCGAGCGTTGTCCGGAATTATTGGGCGTAAAGGGCTCGTAGGCGGTTTGTCGCGTCGGGAGTGAAAACACTGGGCTTAACCCAGTGCTTGCTTTCGATACGGGCAGACTAGAGGTATGCAGGGGAGAACGGAATTCCTGGTGTAGCGGTGAAATGCGCAGATATCAGGAGGAACACCGGTGGCGAAGGCGGTTCTCTGGGCATGATCTGACGCTGAGGAGCGAAAGTGTGGGG |
| P-asv5102 | GCAAGCGTTAATCGGAATTACTGGGCGTAAAGCGTGCGTAGACGGTTACATAAGTCGGGTGTGAAAGCCCCGGGCTCAACCTGGGAATTGCATTCGAGACTGCGTAGCTAGGGTGCGGAAGAGGGAAGCGGAATTTCGGGTGTAGCGGTGAAATGCGTAGATATCCGAAGGAACATCAGTGGCGAAAGCGGCTTCCTGGTCCAGCACCGACGTTCAGGCACGAAAGCGTGGGG |
| P-asv5103 | GCAAACGTTGCTCGGAATCATTGGGCGTAAAGCGCACGTAGGCGGTTCGTTATGTCGGATGTGAAAGCCCTTGGCTTAACCGAGGAAGTGCATCCGAAACTGGCGAGCTAGAGTACAAAAGAGGATCGCGGAATTCCCGGTGTAGAGGTGAAATTCGTAGATATCGGGAGGAACACCGGTGGCGAAGGCGGCGATCTGGGTTGATACTGACGCTGAGGTGCGAAAGCGTGGGG |
| P-asv5104 | GCTAGCGTTGCTCGGAATGACTGGGCGTAAAGGGCGCGTAGGCGGCTCGTACAGTCGGATGTGAAATTCCCGGGCTCAACCTGGGGGCTGCATTCGATACGTACGGGCTTGAGTTCGGAAGAGGGTCGTGGAATTCCCAGTGTAGAGGTGAAATTCGTAGATATTGGGAAGAACACCGGTGGCGAAGGCGGCGACCTGGTCCGATACTGACGCTGAGGCGCGAAAGCGTGGGG |
| P-asv5105 | GCAAGCGTTGTCCGGAATTACTGGGTGTAAAGGGTTCATAGGCGGTGTGGCAAGTCGGGAATGAAATTTCTGGGCTCAACTCAGAGACTGCTTCCGAAACTGCTGTGCTGGAGTGTGGGAGAGGCGCGTGGAATTGCAGGTGTAGCGGTGAAATGCGTAGATATCTGCAGGAACACCCGTGGCGAAAGCGGCGCGCTGGACCACTACTGACGCTGAGGAACGAAAGCTAGGGG |
| P-asv5106 | GCGAGCGTTGTCCGGATTTATTGGGCGTAAAGAGCTCGTAGGCGGCTTGTCGCGTCGACTGTGAAAACCCGCGGCTCAACTGCGGGCTTGCAGCCGATACGGGCAGGCTAGAGTTCGGTAGGGGAGACTGGAATTCCTGGTGTAGCGGTGAAATGCGCAGATATCAGGAGGAACACCGGTGGCGAAGGCGGGTCTCTGGGCCGATACTGACGCTGAGGAGCGAAAGCGTGGGG |
| P-asv5107 | GCTAGCGTTATTCGGAATTATTGGGCGTAAAGGGCGCGTAGGCGGCCTTGCGTGTCTTTCGTGAAATCCCTCGGCTTAACTGAGGACGTGCGGAAGAAACTGCAGGGCTCGAGTCCGGGAGAGGGGGGTGGAATTCCCAGTGTAGCGGTGAAATGCGTAGATATTGGGAGGAACACCGGTGGCGAAGGCGGCCCCCTGGACCGGTACTGACGCTGAGGCGCGAAAGCGTGGGT |
| P-asv5108 | GCAAGCGTTGTTCGGAATTACTGGGCGTAAAGGGCGCGTAGGCGGCCTTCCAAGTCCTGTGTGAAATCCCTCGGCTCAACTGAGGAATCGCACGGGAAACTGTTTGGCTTGAGTTCGGGAGAGGGAAGCGGAATTCCGGGTGTAGCGGTGAAATGCGTAGATATCCGGAGGAACACCGGTGGCGAAGGCGGCTTCCTGGACCGACACTGACGCTGAGGCGCGAAAGCTAGGGG |
| P-asv5109 | GCAAGCGTTATCCGGAATTATTGGGCGTAAAGCGTCCGCAGGTGGCAATTCAAGTCTGCTGTTAAAGGTTCTGGCTCAACCAGAAACAGGCAGTGGAAACTGAATAGCTAGAGTGCGGTAGGGGCAGAGGGAATTCCCAGTGTAGCGGTGAAATGCGTAGAGATTGGGAAGAACACCGGTGGCGAAAGCGCTCTGCTAGGCCGTAACTGACACTCATGGACGAAAGCTAGGGG |
| P-asv5110 | GCAAGCGTTGTCCGGAATTATTGGGCGTAAAGCGCGCGCAGGCGGTTTCTTAAGTCTGATGTGAAAGCCCACGGCTCAACCGTGGAGGGTCATTGGAAACTGGGGAACTTGAGGACAGAAGAGGAGAGTGGAATTCCACGTGTAGCGGTGAAATGCGTAGATATGTGGAGGAACACCAGTGGCGAAGGCGACTCTCTGGTCTGTTTCTGACGCTGAGGTGCGAAAGCGTGGGT |
| P-asv5111 | GCAAGCGTTATCCGGAATTATTGGGCGTAAAGCGCGCGTAGGCGGTTTCTTAAGTCTGATGTGAAAGCCCACGGCTCAACCGTGGAGGGTCATTGGAAACTGGGAAACTTGAGTGCAGAAGAGGAGAGTGGAATTCCATGTGTAGCGGTGAAATGCGCAGAGATATGGAGGAACACCAGTGGCGAAGGCGGCTCTCTGGTCTGTAACTGACGCTGATGTGCGAAAGCGTGGGG |
| P-asv5112 | GCAAGCGTTGTCCGGAATCATTGGGCGTAAAGAGCTCGTAGGCGGTTTGTCGCGTCTGCTGTGAAAACGCGAGGCTCAACCTCGCGCCCGCAGTGGGTACGGGCAAACTAGAGTGCGGTAGGGGAGATTGGAATTCCTGGTGTAGCGGTGGAATGCGCAGATATCAGGAGGAACACCGATGGCGAAGGCAGATCTCTGGGCCGTAACTGACGCTGAGGAGCGAAAGCATGGGG |
| P-asv5113 | GCGAGCGTTGTCCGGAATTATTGGGCGTAAAGGGCTCGTAGGCGGTTTGTCGCGTCGGGAGTGAAAACACTGGGCTTAACCGAGTGCTTGCTTTCGATACGGGCAGACTAGAGGTATGCAGGGGAGAACGGAATTCCTGGTGTAGCGGTGGAATGCGCAGATATCAGGAGGAACACCGGTGGCGAAGGCGGTTCTCTGGGCATTACCTGACGCTGAGGAGCGAAAGCATGGGG |

| | |
|---|---|
| P-asv5114 | GCAAGCGTTAATCGGAATTACTGGGCGTAAAGCGCACGTAGGCGGTTTGTTAAGTTGGATGTGAAAGCCCCGGGCTCAACCTGGGAATTGCATTCAAAACTGGCCAACTAGAGTATGTGAGAGGGGGGTAGAATTCCAAGTGTAGCGGTGAAATGCATAGAGATTTGGAGGAATACCAGTGGCGAAGGCGGCCCCCTGGCACAATACTGACGCTGAGGTGCGAAAGCGTGGGG |
| P-asv5115 | GCAAGCGTTAATCGGAATTACTGGGCGTAAAGCGCACGCAGGCGGTCTGTTAAGTCAGGGGTGAAATCCTGGAGCTCAACTCCGGAACTGCCTTTGATACTGAGGATCTTGAGTTCGGGAGAGGTAAGTGGAACTGCGAGTGTAGAGGTGAAATTCGTAGATATTCGCAAGAACACCAGTGGCGAAGGCGGCTTACTGGCCCGATACTGACACTGAGGCACGAAAGCGTGGGG |
| P-asv5116 | GCAAGCGTTATCCGGAATTATTGGGCGTAAAGAGCTCGTAGGCGGTTTGTCGCGTCTGCTGTGAAAGCCCGGGGCTCAACCCCGGGTCTGCAGTGGGTACGGGCAGACTAGAGTGCAGTAGGGGAGACTGGAATTCCTGGTGTAGCGGTGAAATGCGCAGATATCAGGAGGAACACCGATGGCGAAGGCAGGTCTCTGGGCTGTAACTGACGCTGAGGAGCGAAAGCATGGGG |
| P-asv5117 | GTTAGCGTTGTTCGGAATTACTGGGCGTAAAGCGCACGTAGGCGGATCAGAAAGTTGGGGGTGAAATCCCGGGGCTCAACCCCGGAACTGCCTCCAAAACTTCTGGTCTGGAGTTCGAGAGAGGTGAGTGGAATTCCGAGTGTAGAGGTGAAATTCGTAGATATTCAGAGGAACACCAGTGGCGAAGGCGGCTCACTGGCTCGATACTGACGCTGAGGTGCGAAAGTGTGGGG |
| P-asv5118 | GCGAGCGTTAATCGGAATCACTGGGCGTAAAGGGCGCGTAGACGGTTAGTTACGTCTGCCGTGAAATCCCCGGGCTCAACCTGGGAATGGCGGTGGAAACGGGCTGACTAGAGTATGGGAGAGGGTGATGGAATTCCAGGTGTAGCGGTGAAATGCGTAGAGATCTGGAGGAACATCAGTGGCGAAGGCGGTCACCTGGCCCAATACTGACGCTGAGGCGCGAAAGCGTGGGG |
| P-asv5119 | GCTAGCGTTGTTCAGAATTACTGGGCGTAAAGCGCACGTAGGCGGATTGGAAAGTCAGAGGTGAAATCCCAGGGCTCAACCCTGGAACTGCCTTTGAAACTCCCAGTCTTGAGGTCGAGAGAGGTGAGTGGAATTCCGAGTGTAGAGGTGAAATTCGTAGATATTCGGAGGAACACCAGTGGCGAAGGCGGCTCACTGGCTCGATACTGACGCTGAGGTGCGAAAGCGTGGGG |
| P-asv5120 | GCGAGCGTTAATCGGAATTACTGGGCGTAAAGCGTGCGCAGGCGGTTTCGCAAGACAGATGTGAAATCCCCGGGCTTAACCTGGGAACTGCATTTGTGACTGCGAGGCTAGAGTGCGTCAGAGGGAGGTGGAATTCCGCGTGTAGCAGTGAAATGCGTAGAGATGCGGAGGAACACCGATGGCGAAGGCAGCCTCCTGGGATGACACTGACGCTCATGCACGAAAGCGTGGGG |
| P-asv5121 | GCAAGCGTTACTCGGAATTACTGGGCGTAAAGCGTGCGTAGGTGGTTCGTTAAGTCTGATGTGAAAGCCCTGGGCTCAACCTGGGAACTGCATTGGATACTGGCGATCTGGAGTGCGGTAGAGGGTGGCGGAATTCCCGGTGTAGCAGTGAAATGCGTAGATATCGGGAGGAACATCCGTGGCGAAGGCGGCCACCTGGACCAGCACTGACACTGAGGCACGAAAGCGTGGGG |
| P-asv5122 | GCAAGCGTTATCCGGAATGATTGGGCGTAAAGCGTCTGTAGGTGGCTTTTTAAGTCCGCCGTCAAATCCCAGGGCTCAACCCCGGACAGGCGGTGGAAACTACCAAGCTGGAGTACGGTAGGGGCAGAGGGAATTTCCGGTGGAGCGGTGAAATGCGTAGAGATCGGAAAGAACACCAACGGCGAAAGCACCCTGCTGGGCCGACACTGACACTGAGAGACGAAAGCTAGGGG |
| P-asv5123 | GCAAGCGTTATCCGGAATGATTGGGCGTAAAGCGTCCGCAGGTGGCAGTGTAAGTCTGCTGTCAAAGGTCAGGGCTCAACCCTGAACAGGCAGTGGAAACTACACAGCTAGAGTGCGGTGGGGGCAGAGGGAATTCCCAGTGTAGCGGTGAAATGCGTAGAGATTGGGAAGAACACCGGTGGCGAAAGCGCTCTGCTAGGCCGCAACTGACACTCATGGACGAAAGCTAGGGG |
| P-asv5124 | CCAAGCGTTATCCGGATTTATTGGGTTTAAAGGGTGCGTAGGCGGCCTGTTAAGTCAGGGGTGAAAGACGGTAGCTCAACTATCGCAGTGCCCTTGATACTGATGGGCTTGAATGGACTAGAGGTAGGCGGAATGTGACAAGTAGCGGTGAAATGCATAGATATGTCACAGAACACCGATTGCGAAGGCAGCTTACTATGGTCTTATTGACGCTGAGGCACGAAAGCGTGGGG |
| P-asv5125 | CCGAGCGTTGTCCGGATTTATTGGGCGTAAAGCGAGCGCAGGCGGTTCTTTAAGTCTGAAGTTAAAGGCAGTGGCTTAACCATTGTACGCTTTGGAAACTGGAGGACTTGAGTGCAGAAGGGGAGAGTGGAATTCCATGTGTAGCGGTGAAATGCGTAGATATATGGAGGAACACCGGTGGCGAAAGCGGCTCTCTGGTCTGTAACTGACGCTGAGGCTCGAAAGCGTGGGG |
| P-asv5126 | GCGAGCGTTGTCCGGAATCATTGGGCGTAAAGCGCGTGTAGGCGGCTTGGTCAGTCTGGTGTGAAAACTCGGGGCTTAACCCCTAGCCTGCACTGGATACTGCCAAGCTTGAGTCCGGGAGGGGAGAGCGGAATTCCTAGTGTAGCGGTGAAATGCGCAGATATTAGGAAGAACACCGGTGGCGAAGGCGGCTCTCTGGAACGGTACTGACGCTGAGACGCGAAAGCGTGGGG |
| P-asv5127 | TCGAGCGTTGTCCGGAGTTACTGGGCGTAAAGCGTGCGCAGGCGGCTCATTGCGCCCGGCGTGAAAGCCCCCGGCTCAACCGGGGAGGGTCGTCGGGGACGGGTGAGCTTGAGGGTATCAGGGGTTGGTGGAATTCCCGGTGTAGTGGTGAAATGCGTAGAGATCGGGAGGAACACCCGTGGCGAAGGCGGCCAACTGGGATACACCTGACGCTGAGGCACGAAGGCGTGGGG |
| P-asv5128 | GCGAGCGTTAATCGGAATTACTGGGCGTAAAGAGTGCGCAGGCGGTTTTGCAAGACCGATGTGAAATCCCCGGGCTTAACCTGGGAACTGCATTGGTGACTGCAAGGCTAGAGTGTGTCAGAGGGAGGTGGAACTCCGCATGTAGCAGTGAAATGCGTAGATATGCGGAAGAACACCGATGGCGAAGGCAGCCTCCTGGGATAACACTGACGCTCATGCACGAAAGCGTGGGG |
| P-asv5129 | CCGGGCGTTATCCGGATTTATTGGGTTTAAAGGGAGCGTAGGCCGGAGATTAAGCGTGTTGTGAAATGTAGATGCTCAACATCTGAACTGCAGCGCGAACTGGTTTCCTTGAGTACGCACAAAGTGGGCGGAATTCGTGGTGTAGCGGTGAAATGCTTAGATATCACGAAGAACTCCGATTGCGAAGGCAGCTCACTGGAGCGCAACTGACGCTGAAGCTCGAAAGTGCGGGT |
| P-asv5130 | GCGAGCGTTAATCGGAATTACTGGGCGTAAAGCGTGCGCAGGCGGTTATGTAAGACAGGTGTGAAATCCCCGGGCTTAACCTGGGAACTGCATTTGTGACTGCATAGCTAGAGTACGGTAGAGGGGGATGGAATTCCGCGTGTAGCAGTGAAATGCGTAGATATGCGGAGGAACACCGATGGCGAAGGCAATCCCCTGGACCTGTACTGACGCTCATGCACGAAAGCGTGGGG |
| P-asv5131 | GCAAGCGTTGTCCGGAATTACTGGGCGTAAAGAGCTCGTAGGTGGTTTGTCGCGTCGTCTGTGAAAGCCCGGGGCTTAACTCCGGGTCTGCAGGCGATACGGGCATAACTAGAGTGCTGTAGGGGAGACTGGAATTCCTGGTGTAGCGGTGGAATGCGCAGATATCAGGAGGAACACCGATGGCGAAGGCAGGTCTCTGGGCAGTAACTGACGCTGAGGAGCGAAAGCATGGGG |
| P-asv5132 | GCAAGCGTTATTCGGAATTATTGGGCGTAAAGGGTGCGTAGGCTGCGTACTAAGTCAACTGTTAAATTTCTCGGCCTAACTGAGAATCTGCAGTAGATACTGGTATGCTAGAGGGTGGAAGAGAGAAGTGGAATTCTCGGAGTAGCGGTAAAATGCGTAGATCTCGAGAAGAACACCAATGGCGAAAGCAGCTTCTTGGTCCACACCTGACGCTGAGGCACGAAAGCGTGGGG |
| P-asv5133 | GCGAGCGTTACCCGGATTTACTGGGTGTAAAGGGCGTGTAGGCGGTCTTTCAAGTCCGATGCTAAAGACCGAAGCTCAACTTCGGGGGTGCGTTGGATACTGTGAGGCTAGACGGTCGGAGAGGGTAGCGGAATTTCCGGAGTAGCGGTGAAATGCGCAGATACCGGAAGGAACGCCGATAGCGAAAGCAGCTACCTGGACGATTTGTGACGCTGAGGCGCGAAAGCGTGGGG |
| P-asv5134 | CCAGGCGTTATCCGGATTTATTGGGTTTAAAGGGAGCGTAGGCTGGAGATTAAGTGTGTTGTGAAATGTAGACGCTCAACGTCTGAATTGCAGCGCATACTGGTTTCCTTGAGTACGCACAACGTTGGCGGAATTCGTTGTGTAGCGGTGAAATGCTTAGATATGACGAAGAACTCCGATTGCGAAGGCAGCTGACGGGAGCGCAACTGACGCTTAAGCTCGAAGGTGCGGGT |
| P-asv5135 | GTTAGCGTTGTTCGGAATTACTGGGCGTAAAGCGCGCGTAGGCGGATTGGAAAGTTGGGGGTGAAATCCCGGGGCTCAACCCCGGAACTGCCTCCAAAACTCCCAGTCTTGAGTTCGAGAGAGGTGAGTGGAACTCCGAGTGTAGAGGTGAAATTCGTAGATATTCGGAAGAACACCAGTGGCGAAGGCGGCTCACTGGCTCGATACTGACGCTGAGGTGCGAAAGTGTGGGG |
| P-asv5136 | GCAAGCGTTGTCCGGATTTATTGGGTTTAAAGGGTGCGTAGACGGCTTAGTAAGTCCATGGTGAAATCCGACAGCTCAACTGTTGAACTGCCATGGATACTGCTAGGCTTGAGTACAGACGAGGTAGGCGGAATGGATAGTGTAGCGGTGAAATGCATAGATACTATCCAGAACCCCGATTGCGAAGGCAGCTTACTAGGCTGTAACTGACGCTGAGGCACGAAAGCGTGGGG |
| P-asv5137 | GCTAGCGTTGTTCGGATATACTGGGCGTAAAGCGCACGTAGGCGGCGCCGTAAGTCAGGGGTGAAATCCCGGAGCTCAACTCCGGAACTGCCCTTGAAACTGCAGTGCTAGAATCCTGGAGAGGCGAGTGGAATTCCGAGTGTAGAGGTGAAATTCGTAGATATTCGGAAGAACACCAGTGGCGAAGGCGACTCGCTGGACAGGTATTGACGCTGAGGTGCGAAAGCGTGGGG |
| P-asv5138 | GCAAGCGTTAATCGGAATTACTGGGCGTAAAGTGCGCGTAGGCGGCCGGGTAAGTCGGATGTGAAAGCCCTGGGCTCAACCTAGGAATTGCATTCGATACTGCTTGGCTAGAGTGCAGTAGAGGGGAGTGGAATTCCCGGTGTAGCGGTGAAATGCGTAGATATCGGGAGGAACACCAGTGGCGAAGGCGACTCCCTGGACTGACACTGACGCTGACAGCGCGAAAGCGTGGGG |
| P-asv5139 | CCGAGCGTTAAGCGGAGTAACTGGGCTTAAAGCGTGCGTAGGCGGACCAGTAAGTATCTTGTGAAATCCCATCGCTCAACGGTGGAATTGCAGGGTATACTGCTGGTCTTGAGGTCGGTAGGGGCTGCCGGAACTCTAGGTGTAGCGGTGAAATGCGTAGATATCTAGAGGAACGCCAAAGACGAAGGTAGGCAGCTGGGCCGATTCTGACGCTGAGGTACGAAAGCGTGGGG |
| P-asv5140 | GCAAGCGTTATTCGGAATTACTGGGCGTAAAGCGCGCGTAGGCGGCGAGGCAAGTCAGATGTGAAAGCCCTCGGCTCAACCGAGGAATTGCATTTGAAACTGCCTGGCTAGAGTTCGGGAGAGGGTGGCGGAATTCCCGGTGTAGGAGTGAAATCCGTAGATATCGGGAGGAACACCAGAGGCGAAGGCGGCCACCTGGACCGAAACTGACGCTGAGGTGCGAAAGCGTGGGT |
| P-asv5141 | GCAAGCATTAATCGGATTTACTGGGCGTAAAGGGTGCGTAGGCGGATAGGTCAGTCAGGTGTGAAATTCCGGTACTCAATATCGGAACAGCATTTGATACTACCTGTCTAGAGGTCAGACGGAGAGAACGGAATTCCATGTGTAGCGGTGAAATGCGTAGATATATGGAAGAACACCTGTGGCGAAAGCGGTTCTCTAGTTTGATACTGACGCTGAGGCACGAGAGCGTGGGG |
| P-asv5142 | CCAAGCGTTATCCGGATTTATTGGGTTTAAAGGGTCCGTAGGCGGATGATTAAGTCAGAGGTGAAATCCTGCAGCTCAACTGTAGCATTGCCTTTGATACTGGTTGTCTTGAGTATATCTGAAGTGGGCGGAATGTGTCATGTAGCGGTGAAATGCATAGATATGACACAGAACACCGATTGCGAAGGCAGCTCACTAAGATACAACTGACGCTGATGGACGAAAGCGTGGGT |
| P-asv5143 | CCAAGCGTTAATCGGAATTACTGGGCGTAAAGCGTGCGCAGGCGGTTGTGCAAGACCGATGTGAAATCCCCGAGCTTAACTTGGGAATTGCATTGGTGACTGCACGGCTAGAGTGTGTCAGAGGGGGGTAGAATTCCACGTGTAGCAGTGAAATGCGTAGAGATGTGGAGGAACACCGGTGGCGAAGGCGACCTACTGGACTGGATCTGACGCTGAGGCGCGAAAGCGTGGGG |
| P-asv5144 | GCGAGCGTTATCCGGAATTATTGGGCGTAAAGAGCTCGTAGGCGGTTTGTCGCGTCTGCCGTGAAAGTCCGGGGCTTAACCCCGGATCTGCGGTGGGTACGGGCAGACTAGAGTGCAGTAGGGGAGACTGGAATTCCTGGTGTAGCGGTGAAATGCGCAGATATCAGGAGGAACACCGATGGCGAAGGCAGGTCTCTGGGCTGTTACTGACGCTGAGGAGCGAAAGCATGGGG |
| P-asv5145 | CCGAGCGTTGACCGGATTTATTGGGCGTAAAGCGTGCGCAGGCTGTTTTGTAAGTCATCTGTTAAATCTTCAGGCTCAACCTGGAGGCTGCGGGTGATACTGCAAAACTAGAGTATGGGAGAGGTCAGTGGAATGCCTAGTGTAGGGGTAAAATCCGTTAAGATTAGGTAGAACACCAAATGCGAAAGCAGCTGACTGGAACATTACTGACGCTGATGCACGAAAGCGTGGGT |
| P-asv5146 | ACAAACGTTGTTCGGAATCACTGGGCTTAAAGGGCGTGTAGGCGGAGCAATAGGTGTCTGGTGAAAACCCCGGTCTCAACCCGGGAATTGCTGGGCAAACCGGTGCTCTTGAGGCAGGTAGAGGTGACTGGAACAACTGGTGGAGCGGTGAAATGCGTTGATATCAGTTGGAACGCCGGCGGCGAAAGCGGGTCACTGGGCCTGTCCTGACGCTGAGGCGCGAAAGCGTGGGG |
| P-asv5147 | GCGAGCGTTAATCGGAATTACTGGGCGTAAAGCGCACGCAGGCGGTTTGTTAAGCTAGATGTGAAAGCCCCGGGCTCAACCTGGGATGGTCATTTAGAACTGGCAGACTAGAGTCTTGGAGAGGGGAGTGGAATTCCAGGTGTAGCGGTGAAATGCGTAGATATCTGGAGGAACATCAGTGGCGAAGGCGACTCCCTGGCCAAAGACTGACGCTCATGTGCGAAAGTGTGGGT |
| P-asv5148 | GCTAGCGTTGCTCGGAATTACTGGGCGTAAAGGGAGCGTAGGCGGACATTTAAGTCAGGGGTGAAATCCCGAGGCTCAACCTCGGAACTGCCTTTGATACTGGGTGTCTTGAGTGTGAGAGAGGTATGTGGAACTCCGAGTGTAGAGGTGAAATTCGTAGATATTCGGAAGAACACCAGTGGCGAAGGCGACATACTGGCTCATTACTGACGCTGAGGCTCGAAAGCGTGGGG |
| P-asv5149 | GCGAGCGTTAATCGGAATTACTGGGCGTAAAGCGTGCGCAGGCGGATTGTTAAGCAAGATGTGAAATCCCCGGGCTTAACCTGGGAATGGCATTTTGAACTGGCAGTCTAGAGTGTGTCAGAGGGGGGTGGAATTCCACGTGTAGCAGTGAAATGCGTAGAGATGTGGAGGAACACCGATGGCGAAGGCAGCCCCCTGGGATAACACTGACGCTCATGTACGAAAGCGTGGGT |
| P-asv5150 | CCAAGCATTATCCGGATTCACTGGGCGTAAAGGGTGCGTAGGCGGTGTGATTAGTCAGGTGTTAAATCCTGGGGCTTAACCCCAGAATCGCATTTGAAACGGTCATACTAGAAGTGGTGAGAGGTAAGCAGAACTCACGGTGTAGGGGTGAAATCCGTTGATATCGTGGGGAATACCAAATGCGAAGGCAGCTTACTGGCACGTACTTGACGCTGAGGCACGAAAGCGTGGGT |
| P-asv5151 | GCAAGCGTTGTTCGGAATTACTGGGCGTAAAGAGCGCGTAGGCGGTCTATCAAGTTGGGAGTGAAATCCCGGGGCTCAACCTCGGAATTGCTTTCAAAACTGGTAGACTCGAGAATGGTAGGGGAAAGTGGAATTCCTAGTGTAGAGGTGAAATTCGTAGATATTAGGAGGAACACCAGAGGCGAAGGCGGCTTTCTGGACCATTTCTGACGCTGAGGCGCGAAAGCGTGGGG |
| P-asv5152 | CCGAGCGTTACCCGGAATTATTGGGTGTAAAGGGTTCGTAGGTGGATTTTTAAGTCAGGTATGAAAGACCGAAGCTCAACTTCGTGTTTGTATTTGAAACTGAAAGTCTAGAATCAGGGAGAGGTAAGCGGAATTCTCAGTGTAGGGGTGCAATCCGTAGATACTGAGAGGAACACCAAAAGCGAAGGCAGCTTACTGGAACTGTATTGACACTGAGGAACGAAAGCGTGGGG |

| | |
|---|---|
| P-asv5153 | GCGAGCGTTGTCCGGAATTACTGGGCGTAAAGGGTTCGCAGGCGGCATGGCAAGTCCGATGTAAAAGGCGAAGGCTCAACCTTCGTAAGCATCGGAAACTGTCAAGCTTGAGTGAAGGAGAGGCAAGTGGAATTCCTAGTGTAGCGGTGGAATGCGTAGATATTAGGAGGAATACCGGTGGCGAAGGCGACTTGCTGGACTTCAACTGACGCTGAGGAACGAAAGCGTGGGT |
| P-asv5154 | GCAAGCGTTACTCGGAATCACTGGGCGTGAAGAGCGTGTAGGCGGGTAGTTAAGTCAGAAGTGAAATCCAATAGCTCAACTATTGAACTGCTTTTGAAACTGATTACCTAGAATATGGGAGAGGTAGATGGAATTTCTGGTGTAGGGGTAAAATCCGTAGAGATCAGAAGGAATACCGATTGCGAAGGCGATCTACTGGAACATTATTGACGCTGAGACGCGAAAGCGTGGGG |
| P-asv5155 | GCAAGCGTTGTTCGGATTTACTGGGCGTAAAGCGCGCGCAGGCGGATTTGTAAGTCAGATGTGAAATCTCGGGGCTCAACCTCGAAACTGCGTCTGAAACTACTGATCTAGAGTAATGGAGGGGGAAGTGGAATTTCGCATGTAGGGGTAAAATCCGTAGAGATGCGAAGGAACACCAGTGCCGAAGGGGACTTCCTGGCCATTTACTGACGCTGAGGCGCGAAAGCGTGGGG |
| P-asv5156 | CCGAGCGTTGTCCGGAATTATTGGGCGTAAAGGGCTCGTAGGCGGTTTGTCGCGTCGGGAGTGAAAACTCAGGGCTCAACCCTGAGCGTGCTTACGATACGGGCAAACTAGAGGTATGCAGGGGAGAACGGAATTCCTGGTGTAGCGGTGGAATGCGCAGATATCAGGAGGAACACCGGTGGCGAAGGCGGTTCTCTGGGCATTACCTGACGCTGAGGAGCGAAAGCATGGGG |
| P-asv5157 | GCAAGCGTTATCCGGAATCACTGGGTTTAAAGGGTACGTAGGCTGTAATATAAGTCAGACGTCAAAGGCTGTGGCTAAACCATAGTACGCGTTTGAAACTATATCACTTGAATATGTTCGAGGTGCACGGAATGCGTCATGTAGCGGTGAAATGCATAGATATGACGTAGAACACCAAATGCGAAGGCAGTGCACTAGGGCATCATTGACGCTGAGGTACGAAAGCGTGGGT |
| P-asv5158 | CCAAGCGTTATCCGGAATCATTGGGTTTAAAGGGTCCGTAGGCGGTCTTGTAAGTCAGTGGTGAAAGCCCATCGCTCAACGATGGAACGGCCATTGATACTGCAGGACTTGAATTATTGGGAAGTAACTAGAATATGTAGTGTAGCGGTGAAATGCTTAGAGATTACATGGAATACCAATTGCGAAGGCAGGTTACTACCAATGGATTGACGCTGATGGACGAAAGCGTGGGT |
| P-asv5159 | CCAAGCGTTATCCGGAATCATTGGGTTTAAAGGGTCCGTAGGCGGTTTTATAAGTCAGTGGTGAAATCCGGCAGCTCAACTGTCGAACTGCCATTGATACTGTAGAACTTGAATTATTGTGAAGTAACTAGAATATGTAGTGTAGCGGTGAAATGCTTAGATATTACATGGAATACCCATTGCGAAGGCAGGTTACTAACAATTGATTGACGCTGATGGACGAAAGCGTGGGG |
| P-asv5160 | GCAAGCGTTATTCGGAATTATTGGGCGTAAAGGGTGCGCAGACGGCTTGTTAAGTCACTTGTTAAATTCTCCGGCCTAACCGGAGGCGAGCATGTGAAACTAGCAGGCTAGAGGGTGAGAGAGAGAAGTGGAATTCTCGGAGTAGCGGTAAAATGCGTAGATCTCGAGAGGAACACCAATGGCGAAGGCAGCTTCTTGGCTCACTCCTGACGTTCAGGCACGAAAGCGTGGGG |
| P-asv5161 | GCAAGCGTTATCCGGAATCATTGGGTTTAAAGGGTCCGCAGGCGGACGTATAAGTCAGTGGTGAAATCCTGCAGCTTAACTGCAGAACTGCCATTGATACTGTACGTCTTGAATTCGGTCGAAGTGGGCGGAATGTGTAGTGTAGCGGTGAAATGCATAGATATTACACAGAACACCGATAGCGAAGGCAGCTCACTAGGCCTGGATTGACGCTCAGGGACGAAAGCGTGGGG |
| P-asv5162 | GCGAGCGTTATCCGGATTCATTGGGCGTAAAGAGCGCGTAGGCGGCCTCTCAAGCGGGATCTCTAATCCGAGGGCTCAACCCCCGGCCGGATCCCGAACTGGGAGGCTCGAGTTCGGTAGAGGCAGGCGGAATTCCCGGTGTAGCGGTGGAATGCGCAGATATCGGGAAGAACACCGATGGCGAAGGCAGCCTGCTGGGCCGCAACTGACGCTGAGGCGCGAAAGCTAGGGG |
| P-asv5163 | GCTAGCGTTGTTCGGAATTACTGGGCGTAAAGGGCGTGTAGGCGGACGATCCAGTCAGGGGTGAAAGGCCAGGGCTCAACCTTGGAACGGCCCTTGAAACTGATTGTCTAGAGTCCGGGAGAGGATGGCGGAATTCCCAGTGTAGAGGTGAAATTCGTAGATATTGGGAAGAACACCAGTGGCGAAGGCGGCCATCTGGACCGGTACTGACGCTGAGGCGCGAAAGCGTGGGG |
| P-asv5164 | CCCCTCTTTCCTCGTCCTTACTGGGCGTAAAGCGTGCGCAGGCGGTTGTGCAAGACCGATGTGAAATCCCCGAGCTTAACTTGGGAATTGCATTGGTGACTGCACGGCTAGAGTGTGTCAGAGGGGGGTAGAATTCCACGTGTAGCAGTGAAATGCGTAGAGATGTGGAGGAATACCGATGGCGAAGGCAGCCCCCTGGGATAACACTGACGCTCATGCACGAAAGCGTGGGG |
| P-asv5165 | TCGAGTGATGGCCGATCTTATTGGGCCTAAAGCGTCCGTAGCTGGCCGTGCAAGTCCGTCGGGAAATCCACCCGCTCAACGGGTGGGCGGCCGACGGAAACTGGACGGCTTGGGACCGGGAGGCTTGAAGGGTACGTCCGGGGTAGGAGTGAAATCCCGTAATCCTGGACGGACCACCGATGGGGAAGCCACTTCAAGAGAACGGATCCGACAGTGAGGGACGAAAGCTAGGGT |
| P-asv5166 | CCAAGCGTTAATCGGAATTACTGGGCGTAAAGCGTCCGCAGGTGGCTTGATAAGATAGAGGTGAAATCCCTGGGCTCAACCTAGGAATTGCCTTTATGACTGTCTCGCTAGAGTATGGGAGAGGGGGGTGGAATTCCACGTGTAGCAGTGAAATGCGTAGAGATGTGGAGGAACACCGATGGCGAAGGCAACCCCCTGGCCTAATACTGACACTCATGGACGAAAGCGTGGGG |
| P-asv5167 | GCGAGCGTTAATCGGAATTACTGGGCGTAAAGAGAGCGCAGGCGGTTTTGCAAGACCGATGTGAAATCCCCGGGCTTAACCTGGGAACTGCATTGGTGACTGCAAGGCTAGAGTGTGTCAGAGGGAGGTGGAACTCCGCATGTAGCAGTGAAATGCGTAGATATGCGGAAGAACACCGATGGCGAAGGCAGCCTCCTGGGATAACACTGACGCTCATGCACGAAAGCGTGGGG |
| P-asv5168 | GCTAGCGTTGTTCGGAATTACTGGGCGTAAAGCGCACGTAGGCGGCTTTGTAAGTTAGGGGTGAAAGCCCGGTGCTCAACACCGGAATTGCCTTTAAGACTGCATCGCTTGAATCCAGGAGAGGTGAGTGGAATTCCGAGTGTAGAGGTGAAATTCGTAGATATTCGGAAGAACACCAGTGGCGAAGGCGGCTCACTGGACTGGTATTGACGCTGAGGTGCGAAAGCGTGGGG |
| P-asv5169 | GCGAGCGTTAATCGGAATTACTGGGCGTAAAGCGTGCGCAGGCGGCTTCGCAAGTCAGACGTGAAATCCCCGGGCTTAACCTGGGAATGGCGTTTGAAACTACGGAGCTGGAGTGTGGCAGAGGGGGGTAGAATTCCACGTGTAGCGGTGAAATGCGTAGATATGTGGAGGAATACCGATGGCGAAGGCAGCCCCCTGGGCCAACACTGACGCTCATGCACGAAAGCGTGGGG |
| P-asv5170 | GCAAGCGTTAATCGGAATTACTGGGCGTAAAGGGTGTGTAGGTGGTTAGATAAGTTATCTGTGAAAGCCCTGGGCTCAACCTGGGAAGGTCAGATAAGACTGTTTGGCTCGAGTATAAGAGAGGGTAGTGGAATTTCCGGTGTAGCGGTGAAATGCGTAGAGATCGGAAGGAACACCAGTGGCGAAGGCGGCTACCTGGCTTAATACTGACACTGAAACACGAAAGCGTGGGG |
| P-asv5171 | GCTAGCGTTGTTCGGAATTACTGGGCGTAAAGCGCACGTAGGCGGACTGGAAAGTTGGGGGTGAAATCCCAGGGCTCAACCCTGGAACTGCCTTCAAAACTATCAGTCTGGAGTTCGAGAGAGGTGAGTGGAATTCCGAGTGTAGAGGTGAAATTCGTAGATATTCGGAGGAACACCAGTGGCGAAGGCGGCTCACTGGCTCGATACTGACGCTGAGGTGCGAAAGCGTGGGG |
| P-asv5172 | TCTCTCTTTTCTCTTCCTCCCTTTTCTTAAAGGTTGCGTAGGCGGGTCTTTAAGTCAGGGGTGAAATCCTGGAGCTCAACTCCAGAACTGCCTTTGATACTGAAGATCTTGAGTTCGGGAGAGGTGAGTGGAACTGCGAGTGTAGAGGTGAAATTCGTAGATATTCGCAAGAACACCAGTGGCGAAGGCGGCTCACTGGCCCGATACTGACGCTGAGGCACGAAAGCGTGGGG |
| P-asv5173 | GCAAGCGTTGTCCGGAATTATTGGGCGTAAAGCGCTCGCAGGCGGTTCTTTAAGTCTGATGTGAAATCTTACGGCTCAACCGTAAACGTGCATTGGAAACTGGGGAACTTGAGTGCAGAAGAGGAGAGTGGAATTCCACGTGTAGCGGTGAAATGCGTAGAGATGTGGAGGAACACCAGTGGCGAAGGCGACTCTCTGGTCTGTAACTGACGCTGAGGAGCGAAAGCGTGGGG |
| P-asv5174 | GCAAGCGTTATCCGGAATTATTGGGCGTAAAGCGCGCGTAGGCGGTAAGATAAGTCTGATGTGAAAGCCCCCGGCTCAACCGGGGAGGGTCATTAGAAACTGTTTTACTTGAGTACAGAAGAGGAGAGTGGAATTCCATGTGTAGCGGTGAAATGCGCAGAGATATGGAGGAACACCAGTGGCGAAGGCGGCTCTCTGGTCTGTAACTGACGCTGAGGTGCGAAAGCGTGGGG |
| P-asv5175 | GCAAGCATTAATCGGATTTACTGGGTGTAAAGGGCACGTAGGCGGGTAAGCAAGTCAGATGTGAAATTCCGGAGCTCAACTCCGGAGCTGCATTTGAAACTACTTATCTTGAGGGTGGACGGAGAAAACGGAATTCCACGTGTAGCGGTGAAATGCGTAGATATGTGGAAGAACACCGGTGGCGAAGGCGGTTTTCTAGTTCATTCCTGACGCTGAGGCGCGAGAGCAAGGGG |
| P-asv5176 | GCAAACGTTGTTCGGAATTACTGGGCGTAAAGCGCGTGTAGGCGGTCTTGCAGGTCGGATGTGAAATCCCTCGGCTCAACCAAGGAAGTGCATTCGAAACCGCAAGACTTGAGTGTCGGAGAGGAAGGCGGAATTGTCGGTGTAGAGGTGAAATTCGTAGATATCGACAGGAACACCTGTGGCGAAAGCGGCCTTCTGGACGAACACTGACGCTGAGACGCGAAAGCGTGGGG |
| P-asv5177 | GCGAGCGTTAATCGGATTTACTGGGCGTAAAGCGTGCGTAGGCGGCTTTTTAAGTCGGATGTGAAATCCCTGAGCTTAACTTAGGAATTGCATTCGATACTGGGAAGCTAGAGTATGGGAGAGGATGGTAGAATTCCAGGTGTAGCGGTGGAATGCGCAGATATCAGGAGGAACACCGGTGGCGAAGGCGGGTCTCTGGGCAGTAACTGACGCTGAGGAGCGAAAGCGTGGGG |
| P-asv5178 | GCGAACGTTGTTCGGAATCACTGGGCTTAAAGGGCGCGTAGGCGGTCTGCCAAGTCGGTGGTGAAATCTTTCGGCTTAACCGGAAAAGTGCCAACGATACTGGCGGACTTGAGGGAGGTAGGGGCAGATGGAACTTCCGGTGGAGCGGTGAAATGTGTTGATATCGGAAGGAACGCCGGTGGCGAAAGCGATCTGCTGGACCTCTACTGACGCTGAGGCGCGAAAGCCAGGGG |
| P-asv5179 | GCAAGCGTTGTCCGGATTCATTGGGCGTAAAGAGCGTGTAGGCGGCCAGGTAGGTCAGCTGTGAAAACTGGAGGCTCAACCTTCAGACGTCGGCTGAAACCATCTGGCTAGAGTCCGGGAGAGGAGAGTGGAATTCCTGGTGTAGCGGTGAAATGCGCAGATATCAGGAAGAACACCCGTGGCGAAAGCGGCTCTCTGGAACGGTACTGACGCTGAGACGCGAAAGCGTGGGG |
| P-asv5180 | GCAAGCGTTGTCCGGATTTACTGGGCGTAAAGAATGCGTAGGTGGATATTTAAGTGGGATGTGAAATCCCCGGGCTCAACCCGGGAACTGCATTCCAAACTGGATATCTGGAGTGCAGGAGAGGAAAGCGGAATTCCTAGTGTAGCGGTGAAATGCGTAGATATTAGGAGGAACACCAGTGGCGAAGGCGGCTTTCTGGACTGTAACTGACACTGAGGCATGAAAGCGTGGGG |
| P-asv5181 | GCGAGCGTTGTTCGGAATTACTGGGCGTAAAGCGTATGTAGGCGGTTTTGTAAGACAGATGTGAAATCCCAGGGCTTAACCTTGGAACTGCATTTGTGACTGCAAGACTTGAGGATCGGAGAGGTTGGCGGAATTCCAAGTGTAGAGGTGAAATTCGCAGATATTTGGAGGAACACCGATGGCGTAAGCAGCCAACTGGACGATTACTGACGCTGAGATACGAAAGCGTGGGG |
| P-asv5182 | ACAAACGTTATTCGGAATCACTGGGCTTAAAGGGTGCGTAGGCGGCGCGGAAAGTTGGGTGTGAAATCCCTCGGCTCAACCGAGGAACTGCGCCCAAAACTACCGTGCTCGAGGGAGACAGAGGTGAGCGGAACTTAGGGTGGAGCGGTGAAATGCGTTGATATCCTAAGGAACACCAGTAGCGAAAGCGGCTCACTGGGTCTCTTCTGACGCTGAGGCACGAAAGCTAGGGT |
| P-asv5183 | GCAAGCGTTGTCCGGAATGATTGGGCGTAAAGCATCTGTAGGTTGTTTCTAAAGTCAACTGTTAAATCCCAGAGCTCAACTTTGGCAGGGCAGTTGAGTACTTAGGGACTGGAGAGTGGTAGGGGTAGAGGGAATTCCTTGTGAAGCGGTGAAATGCACAGAGATAAGGAAGAACACCAGCGGCGAAAGCGCTCTACTGGGCCATTACTGACACTGAGAGATGAAAGTTAGGGG |
| P-asv5184 | GCAAGCGTTGTCCGGAATTATTGGGCGTAAAGAGCTCGTAGGCGGTTTGTCGCGTCTGCTGTGAAATCCCGAGGCTCAACCTCGGGACTGCAGTGGGTACGGGCAGACTAGAGTGTGGTAGGGGAGACTGGAATTCCTGGTGTAGCGGTGGAATGCGCAGATATCAGGAGGAACACCGATGGCGAAGGCAGGTCTCTGGGCCACTACTGACGCTGAGGAGCGAAAGCATGGGT |
| P-asv5185 | CCAAGCGTTGTTCGGATTTACTGGGCGTAAAGGGTGCGTAGGCGGCGTGTCAAGTCACTTGTGAAATCTCCGAGCTTAACTCGGAACAGCCAAGTGATACTGATGTGCTAGAGTACAGAAAGGGTAACTGGAATTCTTGGTGTAGCGGTGAAATGCGTAGATATCAAGGGGAACACCTGAGGCGAAGGCGAGTTACTAGGCTGATACTGACGCTGAGGCACGAAAGCTAGGGG |
| P-asv5186 | GCAAGCGTTAATCGGAATTACTGGGCGTAAAGCGTGCGTAGGTGGTTTGTTAAGTCTGATGTGAAAGCCCTGGGCTCAACCTGGGAATGGCATTGGATACTGGCTCACTAGAGTGCGGTAGAGGGATGCGGAATTTCCGGTGTAGCGGTGAAATGCGTAGAGATCGGAAGGAACATCTGTGGCGAAGGCGGCATCCTGGACCAGCACTGACACTGAGGCACGAAAGCGTGGGG |
| P-asv5187 | GCTAGCGTTGTTCGGAATTACTGGGCGTAAAGCGCACGTAGGCGGGTTTGTAAGTAGGGGGTGAAATCCCAGGGCTCAACCCTGGAACTGCCTTCTAGACTGCAAGCCTGGAGGTCAGGAGAGGCGAGTGGAATACCGAGTGTAGAGGTGAAATTCGTAGATATTCGGTGGAACACCAGTGGCGAAGGCGACTCGCTGGACTGATACTGACGCTGAGGTGCGAAAGCGTGGGG |
| P-asv5188 | GCGAGCGTTAATCGGAATTACTGGGCGTAAAGCGCGCGTAGGCGGTTTGTTAAGTCGGATGTGAAATCCCCGGGCTCAACCTGGGAATTGCATTCGATACTGGCAAGCTAGAGTATGGGAGAGGAAGGTAGAATTCCAGGTGTAGCGGTGAAATGCGTAGAGATCTGGAGGAATACCGATGGCGAAGGCAGCCTTCTGGCCTAATACTGACGCTGAGGTGCGAAAGCATGGGG |
| P-asv5189 | GCTAGCGTTGTTCGGAATTACTGGGCGTAAAGCGCACGTAGGCGGCTTTGTAAGTTAGAGGTGAAAGCCCGGGGCTCAACTCCGGAACTGCCTTTAAGACTGCATCGCTAGAATTGTGGAGAGGTGAGTGGAATTCCGAGTGTAGAGGTGAAATTCGTAGATATTCGGAAGAACACCAGTGGCGAAGGCGACTCACTGGACACATATTGACGCTGAGGTGCGAAAGCGCGGGG |
| P-asv5190 | GCAAGCGTTGTCCGGAATTATTGGGCGTAAAGCGCGCGCAGGCGGTCTCTTAAGTCTGATGTGAAAGCCCACGGCTTAACCGTGGAGGGTCATTGGAAACTGAGGGACTTGAGTACAGAAGAGGAGAGTGGAATTCCACGTGTAGCGGTGAAATGCGTAGAGATGTGGAGGAACACCAGTGGCGAAGGCGACTCTCTGGTCTGTAACTGACGCTGAGGTGCGAAAGCGTGGGT |
| P-asv5191 | CCGAGCGTTGTCCGGAATTATTGGGCGTAAAGAGCTCGTAGGCGGTTTGTCGCGTCTGCTGTGAAATCCCGAGGCTCAACCTCGGGCCTGCAGTGGGTACGGGCAGACTGGAGTGCGGTAGGGGAGAATGGAATTCCTGGTGTAGCGGTGGAATGCGCAGATATCAGGAGGAACACCAATTGCGAAGGCAGCTCGCTGGGACGTTACTGACGCTGAGACGCGAAAGCGTGGGG |

| | |
|---|---|
| P-asv5192 | GCAAGCGTTGTCCGGAATTATTGGGCGTAAAGCGCGCGCAGGCGACCTCTTAAGTCTGATGTGAAAGCCCCCGGCTCAACCGGGGAGGGCCATTGGAAACTGGGAGGCTTGAGTATAGGAGAGAAGAGTGGAATTCCACGTGTAGCGGTGAAATGCGTAGAGATGTGGAGGAACACCAGTGGCGAAGGCGACTCTTTGGCCTATAACTGACGCTGAGGCGCGAAAGCGTGGGG |
| P-asv5193 | GCTAGCGTTGCTCGGAATTACTGGGCGTAAAGGGCGCGTAGGCGGATCGTTAAGTCAGAGGTGAAATCCCAGGGCTCAACCCTAGAACTGCCTTTGATACTGGCGATCTTGAGTACGAGAGAGGTATGTGGGACTCCGAGTGTAGAGGTGAAATTCGTAGATATTCGGAAGAACACCAGTGGCGAAGGCGACATACTGGCTCATTACTGACGCTGAGGCGCGAAAGCGTGGGG |
| P-asv5194 | GCGAACGTTATTCGGAATCACTGGGCTTAAAGGGTGCGTAGGCGGCTAATCAAGCCCGAGGTGAAATGTCCCGGCCTAACCGGGGACGTGCTTCGGGGACTGATTGGCTTGGGCGATCTAGGGGTCTGTGGAACTCCCGGTGGAGCGGTGAAATGTGTAGAGATCGGGAGGAACGCCGGTGGCGAAAGCGACAGACTGGGGATTGGCCGACGCTGAGGCACGAAAGCTAGGGG |
| P-asv5195 | GCGAGCGTTAATCGGAATCACTGGGCGTAAAGCGTACGTAGGCGGCGTGTCAAGTCAGGTGTGAAAGCCCTCGGCTCAACCGGGGAATTGCACTTGAAACTGATATGCTAGAGTTCTGGAGGAGGTAGCGGAATTCCCGGTGTAGGAGTGAAACCCGTAGATATCGGGAGGAACACCAGTAGCGAAAGCGGCTACCTGGACAGTAACTGACGCTGAGGTACGAAAGTGTGGGG |
| P-asv5196 | GCTAGCGTTGCTCGGAATCACTGGGCGTAAAGGGCGCGTAGGCGGCCGATCAAGTCAGAGGTGAAAGCCCAGGGCTCAACCCTGGAATTGCCTTTGATACTGTTCGGCTAGAGACCGGAAGAGGTTAGTGGAACTGCGAGTGTAGAGGTGAAATTCGTAGATATTCGCAAGAACACCAGTGGCGAAGGCGGCTAACTGGTCCGGTTCTGACGCTGAGGCGCGAAAGCGTGGGG |
| P-asv5197 | GCGAGCGTTGTCCGGATTTATTGGGCGTAAAGAGCTCGTAGGTGGTTTGGTAAGTCGGATGTGAAATCTCCAGGCTCAACCTGGAGGGGTCATTCGAGACTGCTATGACTAGAGTTCGGTAGAGGAGTGTGGAATTCCTGGTGGAGCGGTGAAATGCGCAGATATCAGGAGGAACACCCGTAGCGAAGGCGGCACTCTGGGCCGATACTGACACTGAGGAGCGAAAGCGTGGGG |
| P-asv5198 | GCAAGCGTTGTCCGGAATCATTGGGCGTAAAGCGCGTGTAGGCGGCTCGATAAGTCGGTTGTGAAAGTCCAGGGCTCAACCCTGGAATGCCGGCCGATACTGTCGAGCTAGAGTCCGGAAGAGGCGAGTGGAATTCCTGGTGTAGCGGTGAAATGCGCAGATATCAGGAGGAACACCAATGGCGAAGGCAGCTCGCTGGGACGGTACTGACGCTGAGACGCGAAAGCGTGGGG |
| P-asv5199 | GCAAGCGTTATCCGGATTTACTGGGTTTAAAGGGTGCGTAGGTGGCAGGAAAGTCAGTGGTGAAATCTTCAAGCTTAACTTGGAAACTGCCATTGATACTTTTTGTCTTGAATATCCCGGAGGTAAGCGGAATATGTCATGTAGCGGTGAAATGCTTAGATATGACATAGAACACCCATTGCGAAGGCAGCTTACTACAGGATTATTGACACTGAGGCACGAAAGCGTGGGG |
| P-asv5200 | GCGAGCGTTGTTCGGATTTACTGGGCGTAAAGGGCGCGTAGGCGGCCATTCAAGTCAGGCGTGAAAGCCCCGGGCTCAACCTGGGAATTGCGCTTGGGACTGGGTGGCTGGAGTTCGGGAGAGGAGCGTGGAATTCCCAGTGTAGAGGTGAAATTCGTAGATATTGGGAAGAACACCGGTGGCGAAGGCGGCGCTCTGGACCGATACTGACGCTGAGGCGCGAAAGCGTGGGG |
| P-asv5201 | GCGAGCGTTGTTCGGAATTATTGGGCGTAAAGGGCGCGTAGGCGGCCATGCGTGTCTCCCGTGAAATCCCTCGGCTTAACTGAGGACGTGCGGAAGAAACTGCAAGGCTCGAGTCCGGGAGAGGGGGGTGGAATTCCCAGTGTAGCGGTGAAATGCGTAGGTATCTGGAGGAACATCAGTGGCGAAGGCGGCTTCCTGGTCCAACACTGACACTGAGGTGCGAAAGCGTGGGG |
| P-asv5202 | GCGAGCGTTGTCCGGAGTTACTGGGCGTAAAGGGCGCGTAGGCGGTGCGGTAAGTCACCTGTGAAATCTCCGAGCTCAACTCGGAGCCTGCAGGCGAAACTGCCGTGCTGGAGTATGGGAGAGGTGCGTGGAATTCCCGGTGTAGCGGTGAAATGCGTAGATATCGGGAGGAACACCTGTGGCGAAAGCGGCGCACTGGACCATAACTGACGCTGAGGCGCGAAAGCTAGGGG |
| P-asv5203 | GCAAGCGTTGTTCGGAATTACTGGGCGTAAAGGGCGCGTAGGCGGCCTGCTAAGTCGAACGTGAAATCCCTGGGCTCAACCCAGGAACTGCGTCCGATACTGGTTGGCTTGAATCCGGGAGAGGGATGTGGAATTCCAGGTGTAGCGGTGAAATGCGTAGATATCTGGAGGAACACCGGTGGCGAAGGCGGCGTTCTGGACTGCAACTGACACTGATAAGCGAAAGCCAGGGT |
| P-asv5204 | GCAAGCGTTGTTCGGAGTTACTGGGCGTAAAGGGTGCGTAGGCGGCCTTTTAAGTTTGGTGTGAAATCTCCCGGCTCAACTGGGAGGGTGCGCCGAATACTGAGAGGCTAGAGTGTGGGAGAGGAAAGTGGAATTCCTGGTGTAGCGGTGAAATGCGTAGATATCAGGAGGAACACCGGTGGTGTAAACGGCTTTCTGGACCATAACTGACGCTGAGGCACGAAAGCGTGGGT |
| P-asv5205 | GCGAGCGTTAATCGGAATTACTGGGCGTAAAGGGCGCGTAGGTGGTTTATTAAGATTGTTGTGAAATCCCTGGGCTTAACCTAGGAACTGCAACGATGACTGGTAGGCTGGAGTACTGTAGAGGGTAGTGGAATTTCCGGTGTAGCGGTGAAATGCGTAGAGATCGGAAGGAACACCAGTGGCGAAGGCGGCTACCTGGACAAATACTGACACTGAGGCGCGAGAGCGTGGGG |
| P-asv5206 | GCAAGCGTTGTCCGGAATCATTGGGCGTAAAGAGCTCGTAGGCGGCCTGTTGCGTCTGCTGTGAAAGCCCGGGGCTTAACCCCGGGTCTGCAGTGGATACGGGTAGGCTAGAGTCCGGAAGAGGAGAGTGGAATTCCTGGTGTAGCGGTGAAATGCGCAGATATCAGGAGGAACACCGGTGGCGAAGGCGGCACTCTGGGCCGGTACTGACGCTGAGGAGCGAAAGCGTGGGG |
| P-asv5207 | GCAAGCGTTGTTCGGAATTACTGGGCGTAAAGGGCTCGTAGGTGGCCAGCTAAGTCAGACGTGAAATCCCCAGGCTCAACTTGGGAACTGCGTCTGATACTGGATGGCTTGAATTCGGGAGAGGGATGCAGAATTCCAGGTGTAGCGGTGAAATGCGTAGATATCTGGAGGAATACCGGTGGCGAAGGCGGCATCCTGGACCGAGATTGACACTGAGGAGCGAAAGCTAGGGG |
| P-asv5208 | GCGAGTGTTGTTCGGAATTACTGGGCGTAAAGGGCGCGCAGGCGGCCTGGCAAGTCGAGTGTGAAATCCCTTGGCTTAACTGAGGAATGGCGCTCGAAACTACCTCGCTGGAGGGCGGGAGAGGGAAGTGGAACTCTCGGTGTAGGGGTGAAATCTGTAGATATCGAGAGGAACACCGGTGGCGAAGGCGGCTTCCTGGCCCGTACCTGACGCTGAGGCGCGAAAGCGTGGGG |
| P-asv5209 | GCAAGCGTTGTCCGGATTTACTGGGCGTAAAGGGCTTGTAGGCGGTTCGTTAAGTCCGGTGTGAAATCTCCCGGCTCAACTGGGAGCGGCCATCGGATACTGGCGAGCTAGAGGTGGGCAGAGGAAAGCGGAATTCCCGGTGTAGCGGTGAAATGCGTAGATATCGGGAGGAACACCAGTGGCGAAGGCGGCTTTCTGGGCCTATCCTGACGCTGAGGAGCGAAAGCGTGGGG |
| P-asv5210 | GCGAGCGTTAATCGGAATTACTGGGCGTAAAGCGCGCGTAGGCGGTCTTGTAAGTCGGATGTGAAAGCCCCGGGCTTAACCTGGGAACTGCATTTGGGACTCTAGAGCTAGAGATCGGGAGAGGTGGGTGGAATTCCCAGTGTAGAGGTGAAATTCGTAGATATTGGGAAGAACACCGGTGGCGAAGGCGGCTCACTGGACCGATACTGACGCTGAGGCGCGAAAGCGTGGGG |
| P-asv5211 | GCAAGCGTTGTTCGGAATCACTGGGCGTAAAGGGCGTGTAGGCGGTCGAGTAAGTGGAATGTGAAATACTCCGGCTCACCCGGAGCACTGCATCCCAAACTGCTCGGCTTGAGTATGGGAGAGGACGAGGGAATTCCTGGTGTAGCGGTGAAATGCGTAGATATCAGGAGGAACACCGGTGGCGAAAGCGCTCGTCTGGCCCAATACTGACGCTGATGCGCGAAAGCTAGGGG |
| P-asv5212 | GCGAGCGTTGTCCGGAGTTACTGGACGTAAAGGGTGCGTAGGCGGTGCGGTAAGTCTGTTGTGAAATCTCCGGGCTCAACTCGGAGCCTGCAACGGAAACTGCCGTGCTTGAGTGTGGGAGAGGTGAGTGGAATTCCCGGTGTAGCGGTGAAATGCGTAGATATCGGGAGGAACACCTGTGGCGAAAGCGGCTCACTGGACCACAACTGACGCTGATGCGCGAAAGCTAGGGG |
| P-asv5213 | GCAAGCGTTATCCGGGTTTATTGGGCGTAAAGAGAGTGCAGGCGGTTTTCTAAGTCTGATGTGAAAGCCTTCGGCTTAACCGGAGAGGTGCATCGGAAACTGGATAACTTGAGTGCAGAAGAGGGTAGTGGAACTCCATGTGTAGCGGTGGAATGCGTAGATATATGGAAGAACACCAGTGGCGAAGGCGGCTACCTGGTCTGCAACTGACGCTGAGACTCGAAAGCATGGGT |
| P-asv5214 | CCAAGCGTTGTTCGGATTCATTGGGCGTAAAGCGCGCGCAGGCGGTTGGCAGAGCCCGGCGTGAAAGCCCCCAGCTCAACTGGGGGAGGCCGTCGGAGACCTGTCGACTTGAGGCCGGTAGGGGAGAGCGGAATTCCCGGTGTAGTGGTGGAATGCGTAGATATCGGGAGGAACACCAGTGGCGAAGGCGGCTCTCTGGACCGGTACTGACGCTCAGGCGCGAAAGCGTGGGG |
| P-asv5215 | GCGAGCGTTATTCGGAATTATTGGGCGTAAAGCGCTCGTAGGCGGGATCGATAGTCCGTGAAGAAAGACCTGGGCTCAACTCAGGGAACGGCACGGATACTTCGATTCTTGAGGCAATCAGAGGGTGATGGAATTCCCGGTGTAGCGGTGAAATGCGTAGATATCGGGAGGAACACCAGTGGCGAAGGCGATCACCTGGGGTTGTTCTGACGCTGAGGAGCGAAAGCTAGGGG |
| P-asv5216 | GCTAGCGTTGTTCGGAATTACTGGGCGTAAAGCGCGCGTAGGCGGCTTTTCAAGTCAGAAGTGAAAGCCCCGGGCTCAACCCGGGAATAGCTTTTGATACTGTTGAGCTCGAGTCCGGGAGAGGATAGTGGAATTCCCAGTGTAGAGGTGAAATTCGTAGATATTGGGAAGAACACCAGTGGCGAAGGCGGCTATCTGGACCGGTACTGACGCTGAGGCGCGAAAGCGTGGGG |
| P-asv5217 | GCGAGCGTTGTCCGGAATCATTGGGCGTAAAGCGCGCGCAGGCGGTTGGCAGAGCCCGGCGTGAAAGCCCCCAGCTCAACTGGGGGAGGCCGTCGGAGACCTGTCGACTTGAGGCCGGTAGGGGAGAGCGGAATTCCCGGTGTAGTGGTGGAATGCGTAGATATCGGGAGGAACACCAGTGGCGAAGGCGGCTCTCTGGACCGGTACTGACGCTCAGGCGCGAAAGCGTGGGG |
| P-asv5218 | GCTAGCGTTGCTCGGAATCACTGGGCGTAAAGCGCACGTAGGCGGCGAAGCAAGTCAGGGGTGAAATCCCGGGGCTCAACCCCGGAACTGCCCTTGAAACTGCTTTGCTAGAATCCTGGAGAGGCGAGTGGAATTCCGAGTGTAGAGGTGAAATTCGTAGATATTCGGAAGAACACCAGTGGCGAAGGCGACTCGCTGGACAGGTATTGACGCTGAGGTGCGAAAGCGTGGGG |
| P-asv5219 | ACAAGCGTTGTCCGGATTTATTGGGCGTAAAGCGAGCGCAGGCGGTCCGGTAAGTCTGATGTGAAAGCCCACGGCTCAACCGTGGAACGGCATTGGAAACTGGCGGACTTGAATGTAGCAGAGGAAAGTGGAATTCCATGTGTAGCGGTGGAATGCGTAGATATATGGAGGAACACCAGTGGCGAAAGCGACTTTCTGGGCTATGATTGACGCTGAGGCTCGAAAGCGTGGGG |
| P-asv5220 | GCAAGCGTTGTCCGGAATTATTGGGCGTAAAGGGCTCGCAGGCGGTTCCTTAAGTCTGATGTGAAAGCCCCCGGCTCAACCGGGGAGGGTCATTGGAAACTGGGGAACTTGAGTGCAGAAGAGGAGAGTGGAATTCCAGGTGTAGCGGTGAAATGCGTAGAGATCTGGAGGAATACCGGTGGCGAAGGCGGCCCCCTGGACGAAGACTGACGCTCAGGTGCGAAAGCGTGGGG |
| P-asv5221 | GCTAGCGTTGTTCGGAATTACTGGGCGTAAAGCGCGCGCAGGCGGTCTTCCAAGTCAGTGGTGAAAGCCTGGAGCTCAACTCCAGAACTGCCATTGAAACTGTTAGACTTGAGTACGAGAGAGGTGAGTGGAATTCCCAGTGTAGGGGTGAAATTCGTAGATATTGGGAAGAACACCAGTGGCGAAGGCGGCTCACTGGCTCGTAACTGACGCTCAGGCGCGACAGCGTGGGG |
| P-asv5222 | CCAAGCGTTGTTCGGATTCATTGGGCGTAAAGGGTGCGTAGGTGGCGGGGCAAGTCAGGTGTGAAATCCCGGGGCTCAACCCCGGAACTGCATTTGATACTGCCTTGCTGGAGTCCTGGAGAGGAGACTGGAATTTACGGTGTAGCAGTGAAATGCGTAGAGATCGTAAGGAAGACCAGTGGCGAAGGCGAGTCTCTGGACAGGTACTGACACTGAGGCACGAAGGCCAGGGG |
| P-asv5223 | GCGAGCGTTAATCGGAATTACTGGGCGTAAAGCGCACGTAGGTGGCGACGTAAGTCGGGTGTGAAAGCCCCGGGCTCAACCTGGGAATTGCATTCGAGACTGCATTGCTGGAGTACGGAAGAGGAGGGCGGAATTCCCGGTGTAGCGGTGGAATGCGTAGAGATCGGGAAGAACACCAGTGGCGAAGGCGGTCTGCTGGGCCGTTGCTGACGCTGAGGCGCGACAGCGTGGGG |
| P-asv5224 | TCGAGTGGTCGGGGCGATTATTGGGCCTAAAGCATCCGTAGCCGGTTCCGTAGGTCCTCTGTTAAATCCAACCGCTTAACGGTTGACCTGCAGAGGATACCGCGGGACTAGGAGGCGGGAGAGGCGGACGGTACTCCGCGTGTAGGGGTAAAATCCTTTGATCCGCGGAAGACCACCAGTGGCGAAGGCGGTCCGCCAGAACGCGCTCGACGGTGAGGGATGAAAGCTGGGGG |
| P-asv5225 | GCAAGCGTTGTTCGGATTGACTGGGCGTAAAGGGCGTGTAGGCGGTCGGTTCAGACCGGTGTGAAAGCCCGGGGCTCAGCCCCGGAAGTGCATCGGAAACTAGCCGACTTGAGCGCGATAGAGGAGGGTGGAATTCCTGGTGTAGCGGTGAAATGCGTAGATATCAGGAGGAACACCGGCGGCGAAGGCGGCCCTCTGGATCGTTGCTGACGCTGAGACGCGAAAGCGTGGGG |
| P-asv5226 | GCAAGCGTTGTTCGGAATTATTGGGCGTAAAGGGTGCGTAGGCGGTTTGGTAAGTCTTATGTGAAATCTCCGGGCTCAACTCGGAGCCTGCATGGGAAACTGCCGGGCTTGAGTGTGGGAGAGGTGAGTGGAATTTCCGGTGTAGCGGTGAAATGCGTAGATATCGGAAGGAACACCTGTGGCGAAAGCGGCTCACTGGACCACAACTGACGCTGATGCACGAAAGCTAGGGG |
| P-asv5227 | GCAAGCGTTAATCGGAATTATTGGGCGTAAAGCGAGTGCAGACGGTTACTTAAGCCAGATGTGAAATCCCCAAGCTTAACTTGGGACGTGCATTTGGAACTGGGTGACTAGAGTGTGTCAGAGGGAGGTAGAATTCACATGTAGCGGTGGAATGCGTAGAGATGTGGAGGAATACCGGTGGCGAAGGCGGCCCCCTGGACAAAGACTGACGCTCAGGTGCGAAAGCGTGGGG |
| P-asv5228 | GCGAGCGTTGTCCGGAATTATTGGGCGTAAAGAGCTCGTAGGCGGTTCGTCGCGTCTGCTGTGAAATCCCGAGGCTCAACCTCTGGCCTGCAGTGGGTACGGGCGAACTAGAGTGCAGTAGGGGAGACTGGAATTCCTGGTGTAGCGGTGGAATGCGCAGATATCAGGAGGAACACCGATGGCGAAGGCAGGTCTCTGGGCTGTAACTGACGCTGAGGAGCGAAAGCGTGGGG |
| P-asv5229 | GCGAGCGTTGTTCGGAATTATTGGGCGTAAAGGGCATGCAGGCGGGTCGCCAAGCTTGGTAAGAAATACCGGGGCTCAACTCCGGAGCTATATTGAGAACTGGCGAGCTAGAGTTGCCGAAGGGTATCCGGAATTCCGCGTGAAGGGGTGAAATCTGTAGATATGCGGAAGAACACCGATGGCGAAGGCAGGATACCGGCGGACGACTGACGCTGAGGTGCGAAGGTGCGGGG |
| P-asv5230 | GCAAGCGTTATCCGGAATTATTGGGCGTAAAGAGTGCGTAGGTGGTTACCTAAGCGCAGGGTTTAATTTAGAGGCTCAACCTCTACTCGCCTTGCGAACTGGGCTACTTGAGTGCAGGAGGGGAAAGCGGAATTCCTAGTGTAGCGGTGAAATGCGTAGATATTAGGAGGAACACCAGCGGCGAAGGCGGCTTTCTGGACTGTAACTGACACTGAGGCACGAAAGCGTGGGT |

| | |
|---|---|
| P-asv5231 | GCAAGCGTTGTCCGGAATTATTGGGCGTAAAGCGCGCGCAGGCGGTCCTTTAAGTCTGATGTGAAATCCCACGGCTCAACCGTGGAAGGTCATTGGAAACTGGGGGACTTGAGTACAGAAGAGGAAAGCGGAAT<br>TCCAAGTGTAGCGGTGAAATGCGTAGAGATTTGGAGGAACACCAGTGGCGAAGGCGGCTTTCTGGTCTGTAACTGACGCTGAGGCGCGAAAGCGTGGGG |
| P-asv5232 | GCTAGCGTTGTTCGGAATTACTGGGCGTAAAGCGCACGTAGGCGGCTTTATAAGTTAGGGGTGAAAGCCTGGAGCTCAACTTCAGAATTGCCCTTAAGACTGCATCGCTTGAATCCAGGAGAGGTGAGTGGAATT<br>CCGAGTGTAGAGGTGAAATTCGTAGATATTCGGAAGAACACCAGTGGCGAAGGCGGCTCACTGGACTGGTATTGACGCTGAGGTGCGAAAGCGTGGGG |
| P-asv5233 | GCAAGCGTTAATCGGAATCACTGGGCGTAAAGGGTGCGTAGGCGGTACGTTAGGTCTGTCGTGAAATCCCCGGGCTCAACCTGGGAATGGCGGTAGAAACCGGCGCACTAGAGTATGGGAGAGGGTGGTGGAA<br>TTCCAGGTGTAGCGGTGAAATGCGTAGAGATCTGGAGGAACATCAGCGGCGAAGGCGGCTCACTGGACTGGTATTGACGCTGAAGTGCGAAAGCGTGGGG |
| P-asv5234 | CCAAGCGTTGTTCGGATTCATTGGGCGTAAAGGGTGCGTAGGTGGCGTGGTAAGTCGGGTGTGAAATTTCGGAGCTTAACTCCGAAACTGCATTTGATACTGCCCTGCTTGAGGACTGGAGAGGAGACTGGAATT<br>TACGGTGTAGCAGTGAAATGCGTAGAGATCGTAAGGAAGACCAGTGGCGAAGGCGGGTCTCTGGACAGTTCCTGACACTGAGGCACGAAGGCCAGGGG |
| P-asv5235 | GCTAGCGTTGTTCGGAATTACTGGGCGTAAAGCGCACGTAGGCGGATATTTAAGTCAGGGGTGAAATCCCGGGGCTCAACCCCGGAACTGCCTTTGATACTGGGTATCTTGAGTTCGGAAGAGGTAAGTGGAATT<br>GCGAGTGTAGAGGTGAAATTCGTAGATATTCGCAGGAACACCAGTGGCGAAGGCGGCTTACTGGTCCGATACTGACGCTGAGGTACGAAAGCGTGGGG |
| P-asv5236 | GCGAGCGTTGTCCGGAATTATTGGGCGTAAAGAGCTTGTAGGCGGTTTGTCGCGTCTGCTGTGAAAGGCCGGGGCTTAACCCCGTGTATTGCAGTGGGTACGGGCAGACTAGAGTGCAGTAGGGGAGACTGGA<br>ATTCCTGGTGTAGCGGTGGAATGCGCAGATATCAGGAGGAACACCGATGGCGAAGGCAGGTCTCTGGGCTGTAACTGACGCTGAGGAGCGAAAGCATGGGG |
| P-asv5237 | GCGAGCGTTAATCGGAATTACTGGGCGTAAAGGGTGCGTAGGTGGTTTTGTAAGTTAGGTGTGAAAGTCCCGGGCTTAACCTGGGGATTGCGCCTAATACTGCAAAGCTAGAGTACTGTAGAGGAAAGTGGAATT<br>TCCAGTGTAGCAGTGAAACGCGTAGATATTGGAAGGAACATCAGTGGCGAAGGCGACTTTCTGGACAGATACTGACACTGAGGCACGAAAGCGTGGGG |
| P-asv5238 | GCGAGCGTTAATCAGATTTACTGGGAGTAAAGCGTGCGTAGGCGGCTTTTTAAGTCGGATGTGAAATCCCCGAGCTTAACTTGGGAATTGCATTCGATACTGGGAAGCTAGAGTATGGGAGAGGATGGTAGAATT<br>CCAGGTGTAGCGGTGAAATGCGTAGAGATCTGGAGGAATACCGATGGCGAAGGCAGCCATCTGGCCTAATACTGACGCTGAGGTACGAAAGCATGGGG |
| P-asv5239 | GCAAGCGTTGTTCGGAATTATTGGGCGTAAAGGGCTCGCAGGTGGTTTGTTAAGTTGGTGGTTTAATCTCTGGGCTCAACCCAGAGTCAGCCTCCAAAACTGGCGAACTTGAGTACGATAGGGGATAGCGGAATT<br>CTCGGTGTAGCGGTGGAATGCGTAGATATCGAGAGGAACACCAATGGCGAAGGCAGCTATCTGGATCGTAACTGACACTCATGAGCGAAAGTGCGGGG |
| P-asv5240 | TCCTTCTTTCTCCTTCCTTCTTGGGCGTAAAGCGCGCGTAGGCGGTTTTTTAAGTCTGATGTGAAAGCCCACGGCTCAACCGTGGAGGGTCATTGGAAACTGGAAAACTTGAGTGCAGAAGAGGAAAGTGGAATT<br>CCATGTGTAGCGGTGAAATGCGCAGAGATATGGAGGAACACCAGTGGCGAAGGCGACTTTCTGGTCTGTAACTGACGCTGATGTGCGAAAGCGTGGGG |
| P-asv5241 | GCAAGCGTTAATCGGAATTACTGGGCGTAAAGCGCGCGTAGGCGGTTGTGCAAGTTGGATGTGAAATCCCCGGGCTTAACCTGGGCACTGCATTCAAAACTGCACGGCTAGAGTATGGGAGAGGAAGGTAGAAT<br>TCCAGGTGTAGCGGTGAAATGCGTAGAGATCTGGAGGAATACCGATGGCGAAGGCAGCCTTCTGGCCTAATACTGACGCTGAGGTGCGAAAGCATGGGG |
| P-asv5242 | GCGAGCGTTATCCGGAATTATTGGGCGTAAAGGGTGCGTAGGCGGCCTGTTAAGTTTATGGTGAAAGCGTGGGGCTCAACCCCATAAAGCCATAGATACTGGCAGGCTAGAGTACTGGTGAGGGTAGTGGAATT<br>CCATGCGTAGCGGTAAAATGCGTAGATATATGGAGGAACACCGGTGGCGAAGGCGGCTACCTAGACAGAGACAGACGCTGAGGCACGAAAGCGTGGGG |
| P-asv5243 | GCAAGCGTTGTCCGGAATTACTGGGCGTAAAGCGCGCGCAGGCGGCCATTTGCGTCCGGGGTGAAAGCCCAAGGCTCAACCTTGGGATTGCCTTGGATACGGGATGGCTTGAGGATCGGAGAGGCAAGGGGA<br>ATTCCACGTGTAGCGGTGAAATGCGTAGAGATGTGGAGGAACACCTGTGGCGAAGGCGCCTTGCTGGCCGATTTCTGACGCTGAGGCGCGAAAGCGTGGGG |
| P-asv5244 | GCGAGCGTTGTCCGGAATTATTGGGCGTAAAGAGCTTGTAGGCGGTTTGTCGCGTCTGCTGTGAAAGCCCGGGGCTTAACCCTGGGTCTGCAGTGGGTACGGGCAGGCTAGAGTGTGGTAGGGGAGACTGGAA<br>TTCCTGGTGTAGCGGTGGAATGCGCAGATATCAGGAGGAACACCGATGGCGAAGGCAGGTCTCTGGGCCATTACTGACGCTGAGAAGCGAAAGCGTGGGG |
| P-asv5245 | GCAAGCGTTAATCGGAATTACTGGGCGTAAAGCGCGCGTAGGTGGTTCGTTAAGTTGGATGTGAAAGCCCCGGGCTCAACCTGGGAACTGCATCCAAAACTGGCGAGCTAGAGTACGGTAGAGGGTGGTGGAAT<br>TTCCTGTGTAGCGGTGAAATGCGTAGAAATAGGAAGGAACACCAGTGGCGAAGGCGGCCTTCTGGACTGACACTGACGCTGAGGTGCGAAAGCGTGGGT |
| P-asv5246 | GCAAGCGTTGCTCGGAATTATTGGGCGTAAAAAGCTCGTAGGCGGTCTGTCGCGTCTGCTGTGAAAACCCGAGGCTCAACCTCGGGCCTGCAGTGGGTACGGGCAGACTAGAGTGCGGTAGGGGAGACTGGAA<br>TCCCTGGTGTAGCGGTGGAATGCGCAGATATCAGGAGGAACACCGATGGCGAAGGCAGGTCTCTGGGCCGCAACTGACGCTGAGGAGCGAAAGCATGGGG |
| P-asv5247 | TCAAGTGATGTCCAATATTATTGGGCCTAAAGCGTCCGTAGCTGGCCGCGCAAGTCCGTCGGGAAATCCACCTGCCCAACAGGTGGGCGTCCGGCGGAAACTGTGTGGCTTGGAACCGGAAGGCTCAGAGGGT<br>ACGTCCGGGGTAGGAGTGAAATCCCGTAATCTTGAACGGACCACCGATGGCGAAAGCACCTCGGAAGCACGGATCCGACAGTGAGGGACGAAAGCTTGGGT |
| P-asv5248 | TCAAGTGATGACCGATTTTATTGGGCCTAAAGCGTCCGTAGCTGGCCGCGCAAGTCCGTCGGGAAATCCACCCGCTTAACGGGTGGGCGTCCGGCGGAAACTGTGCGGCTTGGGACAGGAAGGCTCGAGGGGT<br>ACGTTCGGGGTAGGAGAGAAATCCCGTAATCCCGGACGGACTACCGATGGCGAAAGCACTCTGAGAAGACGGCTTCGACAGTGAGGGACGAAAGCTAGGGT |
| P-asv5249 | GCGTGCGTTGTCCGGAATTATTGGGCGTAAAGGGCTCGTAGGCGGTTTGTCGCGTCGGGAGTGAAAACACTGGGCTTAACTCAGTGCTTGCTTCCGATACGGGCAGACTAGAGGTATTCAGGGGAGAACGGAAT<br>TCCTGGTGTAGCGGTGAAATGCGCAGATATCAGGAGGAACACCGGTGGCGAAGGCGGTTCTCTGGGAATTACCTGACGCTGAGGAGCGAAAGTGTGGGG |
| P-asv5250 | GCGAGCGTTGTCCGGAATTACTGGGCGTAAAGAGCTCGTAGGTGGTTTGTCGCGTCGTCTGTGAAATCCCGGGGCTTAACTTCGGGCGTGCAGGCGATACGGGCAAGCTGGAGTGCTGTAGGGGAGACTGGAA<br>TTCCTGGTGTAGCGGTGGAATGCGCAGATATTAGGAAGAACACCGATGGCGAAGGCAGGTCTCTGGGCAGTAACTGACACTGAGGAGCGAAAGCATGGGT |
| P-asv5251 | GCGAGCGTTGTTCGGAATTACTGGGCGTAAAGCGTGCGCAGGCGGCTCTTTAAGTCTGTCGGTCAAACGCCGGGGCTCAACCCCGTGCTCGCCGGAGATACTGGAGAGCTGGAGACATGTAGAGGCAAGCGGA<br>ATTCCGGGTGTAGCGGTGGAATGCGTAGAGATCCGGAAGAACACCGGAGGCGAAGGCGGCTTGCTGGGCATGGTCTGACGCTGAGGCGCGAAAGCGTGGGG |
| P-asv5252 | CCGAGCGTTGTCCGGATTTATTGGGCGTAAAGCGAGCGCAGGTGGTTACATAAGTCTGAAGTAAAAGGCAGTGGCTCAACCATTGTATGCTTTGGAAACTGTGTAACTTGAGTGCAGAAGGGGAGAGTGGAATTC<br>CATGTGTAGCGGTGAAATGCGTAGATATATGGAAGAACACCGGTGGCGAAAGCGGCTCTCTGGTCTGTAACTGACACTGAGGCTCGAAAGCGTGGGG |
| P-asv5253 | GCGAGCGTTATCCGGAATTATTGGGCATAAAGGGCATCTAGGCGGCCTAACAAGTCAGGGGTGAAAACCTGCGGCTCAACCGCAGGCTTGCCTTTGAAACTGTGAGGCTGGAGTATCGGAGAGGTGGACGGAA<br>CTGCACGAGTAGAGGTGAAATTCGTAGATATGTGCAGGAATGCCGATGATGAAGATAGTTCACTGGACGATAACTGACGCTGAAGTGCGAAAGCCGGGGG |
| P-asv5254 | GCAAGCGTTATCCGGAATCACTGGGCGTAAAGGGCGCGTAGGCGGTCTGTTAAGTCCGATGTTAAAGACCGGGGCTCAACCCCGACACGGCGTTGGATACTGGCAGGCTTGACGACTGGAGAGGCAAGTGGAA<br>TTACCAGTGTAGCGGTGGAATGCGTAGATACTGGTAGGAACACCCATTGCGAAGGCAGCTTGCTGGACAGTACGTGACGCTGAGGCGCGAAAGTGTGGGG |
| P-asv5255 | GCAAGCGTTGTCCGGAATTATTGGGCGTAAAGGGTACGCAGGCGGATGATCAAGTCAAGCGTGAAAGGTGTCGGCTTAACCGACAGACTGCGTTTGAAACTGGTTATCTTGAGTGTAACAGAGGAGAGTGGAATT<br>CCTAGTGTAGTGGTGAAATACGTAGATATTAGGAAGAACACCAGTGGCGAAGGCGACTCTCTGGGTTAACACTGACGCTGAGGTACGAAAGCTGGGGG |
| P-asv5256 | GCTAGCGTTGTTCGGAATTACTGGGCGTAAAGCGCACGTAGGCGGGCTGGAAAGTTGGGGGTGAAATCCCGGGGCTCAACCCCGGAACTGCCTTCAAAACTATCAGCCTAGAGGTCGAGAGAGGTGAGTGGAA<br>CTCCGAGTGTAGAGGTGAAATTCGTAGATATTCGGAAGAACACCAGTGGCGAAGGCGGCTCACTGGCTCGATACTGACGCTGAGGTGCGAAAGCGTGGGG |
| P-asv5257 | GCAAGCGTTACTCGGAATTACTGGGCGTAAAGCGTGCGTAGGTGGTCGTTTAAGTCTGTTGTGAAAGCCCTGGGCTCAACCTGGGAACTGCAGCGGAAACTGGACGACTAGAGTGTGGTAGAGGGTAGCGGAAT<br>TCCTGGTGTAGCAGTGAGATGCGTAGAGATCAGGAGGAACATCCATGGCGAAGGCAGCTACCTGGACCAACACTGACACTGAGGCACGAAAGCGTGGGG |
| P-asv5258 | GCTAGCGTTGTTCGGAATCACTGGGCGTAAAGCGCGCGTAGGCGGCTTGGAAAGTTGGGTGTGAAATCCCGGGGCTTAACCCCGGAACTGCACTCAAAACTTCCTGGCTAGAGACTTAGAGAGGTGGGCGGAAT<br>TCCGAGTGTAGAGGTGAAATTCGTAGATATTCGGAAGAACACCGGTTGCGAAGGCGGCTCACTGGCTAAGTTCTGACGCTGAGGTGCGAAAGCGTGGGG |
| P-asv5259 | GCTAGCGTTGTTCGGAATTACTGGGCGTAAAGCGCGCGTAGGCGGATTGGCCAGTCAGGGGTGAAATCCCGGGGCTTAACCTCGGAACTGCCTTTGATACTGCCAGTCTCGAGTCCGGAAGAGGTGAGTGGAA<br>TTCCTAGTGTAGAGGTGAAATTCGTAGATATTAGGAAGAACACCGGTGGCGAAGGCGGCTCACTGGTCCGGTACTGACGCTGAGGCGCGAAAGCGTGGGG |
| P-asv5260 | GCTAGCGTTATTCGGAATTACTGGGCGTAAAGCGCACGTAGGCGGCTTTGTAAGTTAGAGGTGAAAGCCTGGAGCTCAACTCCAGAATTGCCTTTAAGACTGCATCGCTTGAATCCAGGAGAGGTGAGTGGAATT<br>CCGAGTGTAGGGGTGAAATCCTAAAATCCTGAGAGGAGCACCTGTGGCGAAAGCGCTCGACGAGTACGGATCCGACGGTGAGGAACGAAAGCTAGGGG |
| P-asv5261 | CCAAGCGTTATCCGGATTTACTGGGCGTAAAGCGTCTGTAGCCGGTAATATAAGTCTGCATTCAAATCCCGAAGCTCAACTTCGGAAAGGGTGCAGATACTGTATAACTTGAATGGAATAGGGGGTAGTAGAATTT<br>CCAGTGGAGGAGTGAAATCCGTTGATATTGGAAGGAATACCAAAAGCGAAGGCAGCTACCTATATTCTTATTGACGGTCAGAGACGAAAGCTTGGGT |
| P-asv5262 | GCAAGCGTTACTCGGAATCACTGGGCGTAAAGCGCACGTAGGCGGACTTACAGGTCAATTGTGAAATACCTGGGCTCAACCCGGGAACTGCAGTTGAGACCGTTTGTCTTGAATACCTGAGAGGGTGGGGGAAT<br>ACCTGGTGGAGAAGTGAAATTCGTAGAGATCAGGTGGAACACCGGAGGCGAAGGCGCCTGCCTGGCAGTGTATTGACGCTGAGGTGCGAAAGTGCGGGG |
| P-asv5263 | GCAAGCGTTATCCGGAATTATTGGGCGTAAAGAGCTCGTAGGCGGTTTGTCGCGTCTATCGTGAAAGTCCGAGGCTCAACCTCGGATCTGCGGTGGGTACGGGCAGACTAGAGTGATGTAGGGGAGACTGGAAT<br>TCCTGGTGTAGCGGTGAAATGCGCAGATATCAGGAGGAACACCGATGGCGAAGGCAGGTCTCTGGGCATTAACTGACGCTGAGGAGCGAAAGCATGGGG |
| P-asv5264 | GCTAGCGTTGCTCGGAATCACTGGGCGTAAAGGGCGCGTAGGCGGGCTATCAAGCCCGGGGTGAAATCCTCCCGCTCAACGGGAGAACTGCCTCGGATACTGATGGCCTCGAGGGGGATAGGGGCACGCGGA<br>ACTGTGGGTGGAGCGGTGAAATGCGTTGATATCCATAGGAACTCCGGTGGCGAAAGCGGCGTGCTGGATCCCTTCTGACGCTGAGGCGCGAAAGCCAGGGG |
| P-asv5265 | GCAAGCGTTACCCGGAATCACTGGGCGTAAAGGGCGTGTAGGCGGAAGGTTAAGTCCGACTTTAAAGACCGGGGCTCAACCCCGGGCCTGGGTTGGAGACTGGCTTTCTGGACCTCTGGAGAGGCAACTGGAA<br>TTCCTGGTGTAGCGGTGGAATGCGTAGATACCAGGAGGAACACCGATGGCGAAGGCAGGTTGCTGGACAGAAGGTGACGCTGAGGCGCGAAAGTGTGGGG |
| P-asv5266 | GCGAGCGTTAATCAGAATTACTGGGCGTAAAGGGCGCGTAGGTGGTTTTTTAAGTCAGATGTGAAATCCCCCAGCTTAACTGGGGTCGGTCATCTGAGACTGGAAAACTAGAGTTAGTGAGAGGAAAGTAGAATT<br>TCCGGTGTAGCGGTGAAATGCGTAGATATCGGAAAGAATACCGATGGCGAAGGCAGCTTTCTGGCACCAAACTGACACTGAGGCGCGAAAGCGTGGGG |
| P-asv5267 | GCTAGCGTTGTTCGGAATTACTGGGCGTAAAGCGCACGTAGGCGGCTTCTTAAGTCAGAGGTGAAATCCAGTCGCTTAACGATTGAACTGCCTTTGATACTGGTTGACTTGAGTCATATGGAAGTAGATAGAATGT<br>GTAGTGTAGCGGTGAAATGCTTAGATATTACACAGAATACCGATTGCGAAGGCAGTCTACTACGTATGTACTGACGCTCATGGACGAAAGCGTGGGG |
| P-asv5268 | CCAGGCGTTATCCGGATTTATTGGGTTTAAAGGGAGCGTAGGCTGGAGATTAAGTGTGTTGTGAAATGTAGACGCTCAACGTCTGAATTGCAGCGCATACTGGTTTCCTTGAGTACGCACAACGTTGGCGGAATTC<br>GTCGTGTAGCGGCGAAATGCTTAGATATGACGAAGAACTCCGATTGCGAAGGCAGCTGACGGGAGCGCAACTGACGCTTAAGCTCGAAGGTGCGGGT |
| P-asv5269 | GCAAGCGTTAATCGGAATTACTGGGCGTAAAGGGTGCGTAGGCGGTTTAACAAGTCAGGTGTGAAAGCCCCGAGCTCAACTCGGGAACTGCGCTTGAGACTGTTAGGCTGGAATTCGGAAGAGGTGAGTGGAAT<br>TCCCAGTGTAGAGGTGAAATTCGTAGATATTGGGAAGAACACCGGTGGCGAAGGCGGCTCACTGGTCCGACATTGACGCTGAGGCACGAAAGCGTGGGG |

| | |
|---|---|
| P-asv5270 | CCAAGCGTTATCCGGAATCATTGGGTTTAAAGGGTCCGTAGGCGGGCAGTTAAGTCAGTGGTGAAAGTCTTCCGCTCAACGGGAGAACTGCCATTGATACTGATTGTCTTGAGTTATTATGAAGTGGTTAGAATGAGTAGTGTAGCGGTGAAATGCATAGATATTACTCAGAATACCGATTGCGAAGGCAGATCACTAATAATATACTGACGCTGATGGACGAAAGCGTAGGT |
| P-asv5271 | GCAAGCGTTGTTCGGAATTACTGGGCGTAAAGGGAGCGTAGGCGGGATTACAAGACAGGGGTTAAATCTCCGGGCTCAACCTGGAACTTGCCTTTGTGACTGTAGTCCTTGAATATGGTAGAGGTCAGTGGAATTTCCGGTGTAGCGGTGGAATGCGTAGAGATCGGAAAGAACACCAGAGGCGAAGGCGACTGACTGGACCAATATTGACGCTGAGGCTCGAAAGCGTGGGG |
| P-asv5272 | CCAAGCGTTGTTCGGATTCACTGGGCGTAAAGGGTGCGTAGGCGGTGGGATAAGTCTGGTGTGAAAGCTCGGAGCTTAACTCCGAAATTGCATTGGAAACTATCTCACTTGAGGTTTGGAGGGGGGACTGGAATTCTCGGTGTAGCAGTGAAATGCGTAGATATCGAGAGGAACACCAGTGGCGAAGGCGAGTCCCTGGACAAATCCTGACGCTGAGGCACGAAAGCTAGGGG |
| P-asv5273 | GCAAACGTTGTTCGGAATCACTGGGCTTAAAGAGCACGTAGGCGGCGCTACAAGTGTCGGGTGAAATCCCACGGCTCACCCGTGGAACTGCCCGGCAGACTGTGGTGCTCGAGTGTGGTAGGGGAGAGCGGAACTCCAGGTGGAGCGGTGAAATGCGTAGATATCTGGAGGAACACCGGTGGCGAAGGCGGCTCTCTGGTCCACAACTGACGCTGAGGTGCGAAAGCTAGGGG |
| P-asv5274 | GCAAGCGTTGTTCGGATTCACTGGGCGTACAGGGAGCGTAGGCGGTTGGGTAAGCCCTCCGTGAAATCTCCGGGCCTAACCCGGAAAGTGCAGAGGGGACTGCTCGGCTAGAGGATGGGAGAGGAGCGCGGAATTCCCGGTGTAGCGGTGAAATGCGTAGAGATCGGGAGGAAGGCCGGTGGCGAAGGCGGCGCTCTGGAACATTTCTGACGCTGAGGCTCGAAAGCGTGGGG |
| P-asv5275 | GCAAGCGTTGTTCGGAATTACTGGGCGTAAAGGGCGTGTAGGTGGTAAATTTAGTTAGAGGTGAAATCCCAGGGCTCAACCTTGGAATTGCCTTTAAAACGGGTTTACTAGAGGACAATAAAGGATAACGGAATACCCAGTGTAGAGGTGGAATTCGTAGATATTGGGTAGAACACCAGTGGCGAAAGCGGTTATCTAGGTTGTGACTGACACTGAGACGCGAAAGCGTGGGG |
| P-asv5276 | CCGAGCGTTATCCGGTATTATTGGGCGTAAAGAGTACGTAGGCGGTTATGTAAGACAATGGTTAAATCTCCGGGCTCAACTCGGAACCAGCCGTTGTGACTGCGTAACTTGTCGGTAGCAGGGGCAAGCGGAATTCTCGGTGTAGGAGTGAAATCCGTTGAGATCGAGAAGAACACCAATGGCGAAGGCAGCTTGCTAGGTTACTTGAGACGCTGAGGTACGAAAGCGTGGGG |
| P-asv5277 | GCAAGCGTTATCCGGAATTACTGGGTGTAAAGGGCGTGTAGGCGGGGATGTAAGTCAAATGTGAAATTCTGAAGCTCAACTTCAGCGCTGCATTTGAAACTATATCTCTTGAGTACTGGAGAGGAAAGTGGAATTCCTAGTGTAGCGGTGAAATGCGTAGATATTAGGAGGAACACCAGTGGCGAAGGCGACTTTCTGGACAGTAACTGACGCTGAGGCGCGAAAGCGTGGGG |
| P-asv5278 | GCAAACGTTAATCGGAATTACTGGGCGTAAAGCGCGCGTAGGCGGTTTGCTAAGTGGGATGTGAAAGCCCCGGGCTCAACCTGGGAACTGCATTCCAAACTGGCAGGCTAGAGTTTGGTAGAGGGAGGTGGAATTTCCTGTGTAGCGGTGAAATGCGTAGATATAGGAAGGAACACCAGTGGCGAAGGCGGCCTCCTGGACCAAAACTGACGCTGAGGTGCGAAAGCGTGGGG |
| P-asv5279 | ACGAGTGGCAATCGCATATTGGGCCTAAAGCGTCCGTAGTCGGCTTAGTAAATCTCTTGTGAAATTGTTTGGCTTAACCAAACAGCGTGCGGGAGACACTGCTTTGCTTGGGACCGGGAGGAGTTAGAAGTATTCTGAGGGGAGCGGTAAAATGTTATAATCCTTGGAGGACTACCTGTGGCAAAGGCGTCTAACTAGAACGGGTCCGACGATGAGGGACGAAGGCTAGGGG |
| P-asv5280 | GCTAGCGTTGTTCGGAATCACTGGGCGTAAAGGGCGCGTAGGCGGCGCGGTAAGTTGGGTGTGAAAGACCCGGGCTTAACCCGGGAAATGCGCTTGATACTGCCGTGCTTGAGACCGGGAGAGGTGAGTGGAATTCCCAGTGTAGAGGTGAAATTCGTAGATATTGGGAAGAACACCAGTGGCGAAGGCGGCTCACTGGCCCGGTTCTGACGCTGAGGCGCGAAAGCGTGGGG |
| P-asv5281 | GCGAGCGTTAATCGGAATTACTGGGCGTAAAGCGCACGTAGGCGGCCTGTTAAGCAAGATGTGAAAGCCCAGGGCTCAACCTTGGAATTGCATTTTGAACTGGCAGGCTAGAGTTTTGAAGAGGGTGGTAGAATTTCCAGTGTAGCGGTGAAATGCGTAGATATTGGAAGGAATACCAGTGGCGAAGGCGGCCACCTGGTCAAAAACTGACGCTGAGGAGCGAAAGCGTGGGG |
| P-asv5282 | GCGAACGTTATTCGGAATCACTGGGCTTAAAGCGCGTGTAGGCGGGTTGGCACGTCGACGTTGAAAGACCCCGGCTCAACCGGGGGACCGGCGCCGATACGGCCAGCCTCGAGGGGCGTAGGGGGACCTGGAACTTCCGGTGGAGCGGTGAAATGCGTTGAGATCGGAAGGAACGCCGGTGGCGAAAGCGAGGTCCTGGCCGCCTGCTGACGCTGAGACGCGAAAGCCAGGGG |
| P-asv5283 | GCTAGCGTTGCTCGGAATCACTGGGCGTAAGGGGTGCGTAGGCGGGTCTTTAAGTCAGGGGTGAAATCCTGGAGCTCAACTCCAGAATTGCCTTTAAGACTGCATCGCTTGAATCCAGGAGAGGTGAGTGGAATTCCGAGTGTAGAGGTGAAATTCGTAGATATTCGGAAGAACACCAGTGGCGAAGGCGGCTCACTGGACTGGTATTGACGCTGAAGTGCGAAAGCGTGGGG |
| P-asv5284 | GCAAGCGTTACCCGGAATCACTGGGCGTAAAGGGCGTGTAGGCGGAAGGTTAAGTCCGACTTTAAAGACCGGGGCTCAACCCCGGGCCTGGGTTGGAGACTGGCTTTCTGGACCTCTGGAGAGGCAACTGGAATTCCTGGTGTAGCGGTGGAATGCGTAGATACCAGGAGGAACACCAATGGCGAAGGCAGGTTGCTGGACAGAAGGTGACGCTGAGGCGCGAAAGTGTGGGG |
| P-asv5285 | GCAAGCGTTAATCGGAATTACTGGGCGTAAAGAGCGCGTAGGTGGTTTATTAAGTCGCATGTGAAAGCCCTGAGCTCAACTCAGGAACTGCATGTGATACTGATAAGCTAGAGTATAGTAGAGGGAAGTGGAATTTCCGGTGTAGCGGTGAAATGCATAGATATCGGAAGGAACATCAGTGGCGAAGGCGACTTCTTGGGCTATAACTGACACTGAGGCGCGAAAGCGTGGGT |
| P-asv5286 | GCGAGCGTTGTTCGGAATCATTGGGCGTAAAGGGCGCGCAGGCGGCTGCGCAAGCCTGGTGTGAAAGACCGCAGCTCAACTGCGGGTACGCGCTGGGAACTGTGTGGCTTGAGTTCCGGAGGGGAAGCTGGAATTCCTGGTGTAGGGGTGAAATCTGTAGATATCAGGAAGAACATCAGTGGCGAAGGCGAGCTTCTGGCCGTGAACTGACGCTGAGGCGCGAAAGCGTGGGG |
| P-asv5287 | GCGAGCGTTAATCGGAATTACTGGGCGTAAAGCGCGCGTAGGCGGTTTGGTAGGTCGGATGTGAAATCCCCGGGCTCAACCCGGGAACTGCATTCGATACTGTCAGGCTAGAGTGTGTAAGAGGATAGTGGAATTCCAGGTGTAGCGGTGAAATGCGTAGATATCTGGAGGAACATCAGTGGCGAAGGCGACTGTCTGGTACAACACTGACGCTGAGGTGCGAAAGCGTGGGG |
| P-asv5288 | GCTAGCGTTGCTCGGAATTACTGGGCGTAAAGGGAGCGTAGGCGGGTTATCAAGTTGGAGGTGAAAGCCCAGGGCTCAACCTTGGAATTGCCTTCAAAACTGATAACCTAGAGGATAGAAGAGGTAAGTGGAACTCCGAGTGTAGAGGTGAAATTCGTAGATATTCGGAAGAACACCAGTGGCGAAGGCGACTTACTGGTCTATTACTGACGCTGAGGCTCGAAAGCGTGGGG |
| P-asv5289 | CCAAGCGTTACCCGGAATCACTGGGTGTAAAGGGTTCGTAGGTGGTTTGATAAGTTGGATATGAAAGACCGGAGCTCAACTCCGTGTTTGTGTTCAAAACTGTTAAACTAGAATTAGGGAGAGGTAAGTGGAATTCTGTGTGTAGGGGTGCAATCCGTAGATACACAGAGGAACACCAAAAGCGAAGGCAACTTACTGGAACTATATTGACACTGATGAACGAAAGCGTGGGG |
| P-asv5290 | CCGAGTGATGGCCGCTATTATTGGGCCTAAAGCGTCCGTAGCTTGCCGTGCAAGTCTATCGGGAAATCGACGCGCTCAACGCGTCGGCTTCCGGTGGAAACTGCACGGCTTGGGGCCGGAAGACTCAGCGGGTACGTCCGGGGTAGGAGTGAAATCCTGTAATCCTGGACGGACCACCAATGGCGAAAGCACGCTGAGAAGCCGGACCCGACAGTGAGGGACGAAAGCCAGGGT |
| P-asv5291 | GCTAGCGTTGTTCGGAATCACTGGGCGTAAAGCGCACGTAGGCGGACTTTTAAGTCAGTGGTGAAATCCTAGTGCTTAACTCTAGAACTGCCATTGATACTATTTATCTTGAATATTGTGGAGGTAAGCGGAATATGTCATGTAGCGGTGAAATGCTTAGATATGACATAGAACACCTATTGCGAAGGCAGCTTACTACGCATATATTGACGCTGAGGCACGAAAGCGTGGGG |
| P-asv5292 | GCAAGCGTTGTCCGGAATTATTGGGCGTAAAGGGCTCGTAGGCGGTTTGTCGCGTCGGGAGTGAAAACACTGGGCTTAACCGAGTGCTTGCTTTCGATACGGGCAGACTAGAGGTATGCAGGGGAGAACGGAATTCCTGGTGTAGCGGTGAAATGCGCAGATATCAGGAGGAACACCGGTGGCGAAGGCGGTTCTCTGGGCATTACCTGACGCTGAGGAGCGAAAGTGTGGGG |
| P-asv5293 | CCGAGCGTTATCCGGATTTATTGGGTTTAAAGGGTGCGTAGGTGGTTTGATAAGTCAGCGGTGAAAGTTTGCAGCTTAACTGTAAAAATGCCGTTGAAACTGTCGGACTTGAGTGTAAATGAGGTAGGCGGAATGCGTGGTGTAGCGGTGAAATGCATAGATATCACGCAGAACTCCGATTGCGAAGGCAGCTTACTAAGCTACAACTGACACTGAAGCACGAAAGCGTGGGG |
| P-asv5294 | GCTAGCGTTATTCGGAATTATTGGGCGTAAAGAGTTCGTAGGCGGTTACCTAAGTCTGGTGTTAAAGACGGAAGCTCAACTTCCGCATGGCATTGGATACTGGGTGGCTAGAGTGTGAAAGAGGTAACTGGAATTCCAGGTGTAGCGGTGAAATGCGTAGATATCTGGAGGAACACCGGTGGCGAAAGCGAGTTACTGGTTCACAACTGACGCTGAGGAACGAAAGCCAGGGT |
| P-asv5295 | GCAAGCGTTATCCGGATTCATTGGGTTTAAAGGGTGCGCAGGCGGGCTGGTAAGTCAGTGGTGAAATACTGTCGCTTAACGATAGGGTTGCCATTGATACTGCTGGTCTTGAGTTTGGTTGAGGTAGGCGGAATGTGCAGTGTAGCGGTGAAATGCATAGAGATTGCACAGAACACCAATTGCGAAGGCAGCTTACTAAACCAAAACTGACGCTGAGGCACGAAAGCGTGGGG |
| P-asv5296 | GCAAGCGTTAATCGGAATTACTGGGCGTAAAGCGTGCGTAGGTGGTTTGGTAAGTTATCTGTGAAATCCCTGGGCTCAACCTGGGCAGGTCAGATAATACTGCTGAACTCGAGTATGGGAGAGGGTAGTGGAATTTCCGGTGTAGCGGTGAAATGCGTAGAGATCGGAAGGAACACCAGTGGCGAAGGCGGCTACCTGGCCTAATACTGACACTGAGGCACGAAAGCGTGGGG |
| P-asv5297 | GCTAGCGTTGTTCGGAATTACTGGGCGTAAAGCGCGCGTAGGCGGCTTGGCAAGTGAGGCGTGAAAGTCCTGGGCTCAACCTGGGGACTGCGCTTCATACTGCTTGGCTTGAGTCCGGAAGAGGGTGGTGGAATTCCGAGTGTAGAGGTGAAATTCGTAGATATTCGGAGGAACACCGGTGGCGAAGGCGGCCACCTGGTCCGGTACTGACGCTCAAGTGCGAAAGCGTGGGG |
| P-asv5298 | GCGAGCGTTGTCCGGAATCACTGGGCGTAAAGCGCGCGCAGGCGGCCGTCTGCGTCCGGGGTGAAAGCCCGGAGCTCAACTCCGGGACTGCCTTGGATACGGGACGGCTTGAGGATCGGAGAGGCAAGGGGAATTCCACGTGTAGCGGTGAAATGTGTAGAGATGTGGAGGAACACCTGTGGCGAAGGCGCCTTGCTGGCCGATTTCTGACGCTGAGGCGCGAAAGCGTGGGG |
| P-asv5299 | GCTAGCGTTACTCGGAATTACTGGGCGTAAAGGGCGCGTAGGCGGCTTGGCCAGTCAGGCGTGAAATTCCTGGGCTTAACCTGGGGACTGCGCTTGATACTGCCAGGCTTGAGGCCGAGAGAGGGTGGTGGAATTCCCAGTGTAGAGGTGAAATTCGTAGATATTGGGAAGAACACCGGTGGCGAAGGCGACCACCTGGCTCGGTACTGACGCTGAGGCGCGACAGCGTGGGG |
| P-asv5300 | GCTAGCGTTATTCGGAATTACTGGGCGTAAAGCGCGCGTAGGCGGCTTTGTAAGTTAGAGGTGAAAGCCCGGAGCTCAACTCCGGAACTGCCTTTAAGACTGCATCGCTAGAATTGTGGAGAGGTGAGTGGAACTCCGAGTGTAGAGGTGAAATTCGTAGATATTCGGAAGAACACCAGTGGCGAAGGCGACTCACTGGACACATATTGACGCTGAGGTGCGAAAGCGTGGGG |
| P-asv5301 | CCAAGCGTTATCCGGATTCATTGGGTTTAAAGGGTGCGTAGGCGGGACGTTAAGTCAGTGGTGAAATCTCTCAGCTCAACTGAGAAACTGCCTCTGATACTGATGTTCTTGAGTTTGGCTGAAGTGGGCGGAATGAGGCATGTAGCGGTGAAATGCATAGATATGCCTCAGAACACCAATTGCGAAGGCAGCTCACTAAACCAATACTGACGCTGAGGCACGAAAGCGTGGGG |
| P-asv5302 | GCTAGCGTTGTTCGGAATTACTGGGCGTAAAGCGCACGCAGGCGGTGCGATCAGTCAGAAGTGAAAGCCCCGGGCTTAACCTGGGAACTGCTTTTGATACTGTCGCGCTTGAATCACGGAGAGGGTAGTGGAATTCCGAGTGTAGAGGTGAAATTCGTAGATATTCGGAAGAACACCAGTGGCGAAGGCGACTACCTGGCCGTCGATTGACGCTCATGTGCGAAAGCGTGGGG |
| P-asv5303 | GCAAGCGTTGTTCGGAATGACTGGGCGTAAAGGGCGTGTAGGCGGCCATGTGCGTCAGATGTGAAAGTCCTGGGCTTAACCTGGGGATTGCATTTGATACGGCAAGGCTAGAGGTTATGAGGGGAGAGCGGAATTGCGCATGTAGAAGTGAAATTCGTAGAGATGCGCAGGAACACCGATGTCGAAGGCAGCTCTCTGGTATTAACCTGACGCTAAGGCGCGAAAGCGTGGGG |
| P-asv5304 | ACAAGTGTTATCCGGAATTATTGGGCGTAAAGCGTTTGTAGGTGGATAAAAAAGTCTTTTGTTAAATCTTTAAGCTTAACTTAAAACAAGCAAAAGATACTCTTTTTCTAGAGTATGGGAGAGGTAAAGGGAATTTTCAGAGAAGCGGTGAAATGCGTAGATATTGAAAGGAACACCAATGGCGAAAGCACTTTACTGGATCAATACTGACACTAAGAAACGAAAGCTAGGGG |
| P-asv5305 | ATAAGCGTTACTCAGCATTATTGGGCGTAAAGGGTGCGTAGGTTGTTTAATAAGTTAATACAAAAATCCAAGACAATTAATTCTTGTAAAGTGATTAATACTATTAAACTAGAGTGTAAAAGAGGATAGTAGTATTCTTAGTGTAGAGGTAAAATTCGTAAATATTAAGAGGACTATCAACAGGCGAAGGCAACTATCTGGTTTATTCTGACACTGAGGCACGAAAGCATGGGG |
| P-asv5306 | GCAAGCGTTGCCCGGAATTACTGGGCGTAAAGAGCTCGTAGGCGGGGGCGTGCGTCCGAAAGGAAATCTCAAGGCTCAACCTTGGGGCTTTTTCGGATACGGCGCTTCTTGAGGGTGAGAGAGGAAAGCGGAATTCCAGGTGTAGCGGTGAAATGCGTAGATATCTGGAGGAACACCAGCGGCGAAGGCGGCTTTCTGGCTCATTCCTGACGCTGAGGCACGAAAGCGTGGGG |
| P-asv5307 | GCGAGCGTTGTCCGGAATTATTGGGCGTAAAGGGCTCGTAGGCGGTTTGTCGCGTCAGGAGTGAAAACTCGGGGCTTAACTCCGAGCCTGCTTCTGATACGGGCAGACTAGAGGTATGCAGGGGAGAACGGAATTCCTGGTGTAGCGGTGAGATGCGCAGATATCAGGAGGAACACCGGTGGCGAAGGCGGTTCTCTGGGCATTACCTGACGCTGAGGAGCGAAAGTGTGGGG |
| P-asv5308 | GCGAGCGTTGTTCGGAATCATTGGGCGTAAAGCGTATGTAGGCGGACTAGAAAGTCAGAAGTGAAATCCCAGGGCTCAACCCTGGAATTGCTTTTGAAACTCCTAGTCTAGTGTCCCGGAGGGGTTGACGGAATTCCTAGTGTAGAGGTGAAATTCGCAGATATTAGGAGGAACACCAGTGGCGTAGGCGGTAACTGGACGGGTACAGACGCTGAGATACGAAAGCGTGGGG |

| | |
|---|---|
| P-asv5309 | CCGAGTGATGGCCGAAATTATTGGGCCTAAAGCGTTCGTAGCTGGCCGGACAAGTCCGTTGGGAAATCCGCGCGCTTAACGCGCGGGCGTCCAGCGGAAACTGTTCGGCTTGGGACCGGAAGACCTGAGGGGTACGTCCGGGGTAGGAGTGAAATCCTGTAATCCTGGACGGACCACCGGTGGCGAAAGCGCCTCAGGAGGACGGATCCGACAGTGAGGGACGAAAGCTAGGGT |
| P-asv5310 | GCTAGCGTTCTTCTTCCTTCCTGGGCGTAAAGCGCACGTAGGCGGCTTTGTAAGTTAGAGGTGAAAGCCTGGAGCTCAACTCCAGAATTGCCTTTAAGACTGCATCGCTTGAATCCAGGAGAGGTGAGTGGAATTCCGAGTGTAGAGGTGAAATTCGTAGATATTCGGAAGAACACCAGTGGCGAAGGCGGCTCACTGGACTGGTATTGACGCTGAGGTGCGAAAGCGTGGGG |
| P-asv5311 | GCAAGCGTTGTTCGGAATTACTGGGCTTAAAGCGTGTGTAGGCGGAATGGAAGGTGTCGTGTGAAAGCCCCCGGCTCAACCGGGGAACGGCTCGGCAAACCACCATTCTTGAGGCCAGCAGGGGTACCCGGAACCTTGGGTGGAGCGGTGAAATGCGTTGATATTCAAGGGAACACCGATGGCGAAAGCGGGGTACTGGGCTGGTTCTGACGCTGAGACACGAAAGCGTGGGG |
| P-asv5312 | GCAAGCGTTGTTCGGAATCATTGGGCGTAAAGCGCGTGTAGGTGGTTATGCAAGTTGGATGTGAAAGCCCTCGGCTCAACCGGGGAAGTGCATCCAAAACTACATAGCTAGAGGACGGCAGAGGGAGGTGGAATTCCAGGTGTAGAGGTGAAATTCGTAGATATCTGGAGGAATACCTGAGGCGAAGGCGGCCTCCTGGACCGTTCCTGACACTGAGACGCGAAAGCGTGGGG |
| P-asv5313 | GCAAGCGTTGTCCGGAATTATTGGGCGTAAAGAGCTCGTAGGCGGTTTGTCGCGTCTGGTGTGAAAACTCAAGGCTCAACCTTGAGCTTGCATCGGGTACGGGCAGACTAGAGTGTGGTAGGGGAGACTGGAATTCCTGGTGTAGCGGTGGAATGCGCAGATATCAGGAGGAACACCGATGGCGAAGGCAGGTCTCTGGGCCACTACTGACGCTGAGGAGCGAAAGCATGGGT |
| P-asv5314 | GCTAGCGTTGCTCGGAATTACTGGGCGTAAAGGGCGCGTAGGCGGCTATGTTAGTCAGGCGTGAAATTCCTGGGCTCAACCTGGGGACTGCGCTTGATACGGCATGGCTTGAGGATGGAAGAGGCTCGCGGAATTCCCAGTGTAGAGGTGAAATTCGTAGATATTGGGAAGAACACCGGTGGCGAAGGCGGCGAGCTGGTCCATTACTGACGCTGAGGCGCGACAGCGTGGGG |
| P-asv5315 | GCGAGCGTTGTCCGGATTTACTGGGCGTAAAGGGCGCGCAGGCGGTCTGTTAAGTTCAGGGTGAAAGCTCCCGGCTCAACTGGGAGAGGTCCTTGGATACTGGCAGACTCGAGGGAGGCAGAGGAAAGTGGAATTCCCGGTGTAGTGGTGATATGCGTAGATATCGGGAGGAACACCAGTGGCGAAGGCGGCTTTCTGGGCCTTACCTGACGCTGAGGCGCGAAAGCGTGGGG |
| P-asv5316 | GCAAGTGTTATCCGGAATCACTGGGCGTAAAGCGTCTGTAGGTGGTTTAATAAGTCAACTGTTAAATCTTGAAGCTCAACTTCAAAATCGCAGTCGAAACTATTAGACTAGAGTATAGTAGGGGTAAAGGGAATTTCCAGTGGAGCGGTGAAATGCGTAGAGATTGGAAGGAACACCGATGGCGAAGACACTTTACTGGGCTATTACTGACACTCAGAGACGAAAGCTAGGGT |
| P-asv5317 | GCGAGCGTTAATCGGAATTACTGGGCGTAAAGCGTGCGCAGGCGGTTGTGTAAGACAGGTGTGAAATCCCCGGGCTTAACCTGGGAACTGCGCTTGTGACTGCACGGCTAGAGTACGGCAGAGGGGGGTGGAATTCCACGTGTAGCAGTGAAATGCGTAGATATGTGGAGGAACACCGATGGCGAAGGCAGCCCCCTGGGCCGATACTGACGCTCATGCACGAAAGCGTGGGG |
| P-asv5318 | GCAACCGTTGTTCGGAATCACTGGGCATAAAGAGCACGTAGGCGGGTATGTAAGTCGGGTGTGAAATCCCTCGGCCTAACCGAGGAACTGCGCCCGAAACTGCATGTCTTGAGGACAGGAGGGGAGCGCGGAATTCCTGGTGGAGCGGTGAAATGCGTTGATATCAGGAGGAACACCGGCGGCGAAGGCGGCGCTCTGGCCTGTTTCTGACGCTGAGGTGCGAAAGCCAGGGG |
| P-asv5319 | GCGAGCGTTAATCGGAATTATTGGGCGTAAAGGGTGCGTAGGTGGTTAGATAAGTCAGATGTGAAAGCCCTGGGCTTAACCTAGGAATGGCGTTTGATACTGTTTAGCTAGAGTACGAGAGAGGGAAGTGGAATTTCCGGTGTAGCGGTGAAATGCGTAGATATCGGAAGGAACATCAGTGGCGAAAGCGACTTCCTGGCTCGAAACTGACACTGAGGCACGAAAGCGTGGGG |
| P-asv5320 | GCGAACGTTATTCGGAATCACTGGGCTTAAAGGGTGCGTAGGCGGCCAGTTAAGTAGGGTGTGAAATCCCTCGGCTCAACCGAGGAATTGCGCTCTAAACTGTCTGGCTTGAGTGAGATAGGGGTATACGGAACTTCCGGTGGAGCGGTGAAATGTGTTGAGATCGGAAGGAACGCCGGTGGCGAAAGCGGTATACTGGGTCTTAACTGACGCTGAGGCACGAAAGCCAGGGT |
| P-asv5321 | TCTCTCTTTTTCTTCCTTCCTGGGCTTAAAGCGCACGTAGGCGGCGATTTAAGTCAGAGGTGAAAGCCCGGGGCTCAACCCCGGAACTGCCTTTGAGACTGGATTGCTTGAATCCTGGAGAGGTGAGTGGAATTCCGAGTGTAGAGGTGAAATTCGTAGATATTCGGAAGAACACCAGTGGCGAAGGCGGCTCACTGGACAGGTATTGACGCTGAGGTGCGAAAGCGTGGGG |
| P-asv5322 | ACGAGCGTTATCCGGAATTATTGGGTTTAAAGGGTGCGTAGGTGGCGTATTAAGTCAGTGGTGAAAAGCTGCAGCTCAACTGTAGTCTTGCCGTTGAAACTGATATGCTAGAGAGGAGACGAGGTATGCGGAATGTGTGGTGTAGCGGTGAAATGCATAGATATCACACAGAACGCCGATTGCGAAGGCAGCGTACCAGGCTCCGTCTGACACTGAAGCACGAAAGCGTGGGG |
| P-asv5323 | TCTCTCTTTCCTCGGAATTACTGGGCGTAAAGGGCGCGTAGGCGGTGGGTTACGTCTGCCGTGAAATCCCCGGGCTCAACCTGGGAATGGCGGTGGAAACGGGCTGACTGGAGTATGGGAGAGGGTGATGGAATTCCAGGTGTAGCGGTGAAATGCGTAGAGATCTGGAGGAACACCAGTGGCGAAGGCGGTCACCTGGCCCAATACTGACGCTGAGGCGCGAAAGCGTGGGG |
| P-asv5324 | GCAAGCGTTATCCGGATTCACTGGGTTTAAAGGGTGCGTAGGCGGGTATGCAAGTCAGAGGTGAAAGCCTGGAGCTCAACTCCAGAACTGCCTTTGAGACTGCATCGCTTGAATCCAGGAGAGGTGAGTGGAATTCCGAGTGTAGAGGTGAAATTCGTAGATATTCGGAAGAACACCAGTGGCGAAGGCGGCTCACTGGACTGGTATTGACGCTGAAGTGCGAAAGCGTGGGG |
| P-asv5325 | GCTAGCGTTGCTCGGAATGACTGGGCGTAAAGGGCGCGTAGGCGGACGGCACAGTCAGGCGTGAAATTCCTGGGCTCAACCTGGGGATTGCGTCTGAGACGTGTTGTCTTGAGTATGGAAGAGGGTTGTGGAATTTCCAGTGTAGAGGTGAAATTCGTAGATATTGGAAAGAACACCGGTGGCGAAGGCGGCAACCTGGTCCATTACTGACGCTGAGGCGCGAAAGCGTGGGG |
| P-asv5326 | GCGAGCGTTGTCCGGAATTACTGGGCGTAAAGGGTTCGCAGGCGGCATGACAAGTCCGATGTAAAAGGCGAAAGCTCAACTTTCGTAAGCATCGGAAACTGTCAAGCTTGAGTGAAGGAGAGGCAAGTGGAATTCCTAGTGTAGCGGTGGAATGCGTAGATATTAGGAGGAATACCGGTGGCGAAGGCGACTTGCTGGACTTCAACTGACGCTGAGGAACGAAAGCGTGGGT |
| P-asv5327 | GCAAGCGTTGTCCGGAATTATTGGGCGTAAAGCGCTCGTAGGTGGATCTACAAGTCTGCGAAGAAAGACCCGGGCTCAACTCGGGGAACGGCGTAGATACTGTTGATCTTGAGACAATCAGAGGATGATGGAATTCCCGGTGTAGCGGTGAAATGCGTAGATATCGGGAGGAACACCAGTGGCGAAGGCGATCATCTGGGGTTGTTCTGACACTGAGGAGCGAAAGCTAGGGG |
| P-asv5328 | GGCAAGCGTTGTTCGGAATTACTGGGCGCAAAGGGCTCGTAGGCGGCTAACTAAGTCAGACGTGAAATCCCTCGGCTTAACCGGGGAACTGCTTCTGATACTGGATAGCTTGAGTTTGGGAGAGGGATGCGGAATTCCAGGTGTAGCGGTGAAATGCGTAGATATCTGGAGGAACACCGGTGGCGAAGGCGGCATCCTGGACCAACACTGACGCTGATGAGCGAAAGCCAGGGG |
| P-asv5329 | GCAAGCGTTGTCCGGATTTATTGGGCGTAAAGCGCACGCAGGCGGTCTGCCAAGTTCGGAGTGACAGTTTCCGGCTTAACTGGGAAAGGTCTTCGAAAACTGGTGGACTTGAGAACTTGAGGGGGAGACGGAATTCCGGGTGGAGCGGTGAAATGCGTAGAGATCCGGAGGAACACCGAAAGCGAAGGCAGTCTCCTGGCAAGTTTCTGACGCTCAGGTGCGAAAGCTAGGGG |
| P-asv5330 | GCGAACGTTGTTCGGAATCACTGGGCTTAAAGGGCGCGTAGGCGGGCTGTCAAGTCCGGGGTGAAAGCCTCCCGCTTAACGGGAGAAGAGCCTCGGATACTGACGGCCTCGAGGAGGGTAGGGGCAAGCGGAACTATGGGTGGAGCGGTGAAATGCATTGATATCCATAGGAACTCCGGTGGCGAAGGCGGCTTGCTGGACCCTTTCTGACGCTGAGGCGCGAAAGCCAGGGG |
| P-asv5331 | GCCAGCGTTGTTCGGAATTACTGGGTGTAAAGGGTTCGTAGGCGGTGTGGCAAGTTGGGAGTGAAATCTCTGGGCTCAACCCAGAGGCTGCTTCCAAAACTGCCGTGCTTGAGTGTGGGAGAGGCGCGTGGAATTGCAGGTGTAGCGGTGAAATGCGTAGATATCTGCAGGAACACCCGTGGCGAAAGCGGCGCGCTGGACCACTACTGACGCTGAGGCACGAAAGCGTGGGG |
| P-asv5332 | CCAAGCGTTGTTCGGATTCACTGGGCGTAAAGCGCTCGTAGGCGGTCTGTTACGTCGGCTGTGAAAACCTGGGGCTCAACCCCGGGCCTGCAGCCGATACGGGCAGACTAGAATTCGGTAGGGGAGACTGGAATTCCTGGTGTAGCGGTGAAATGCGCAGATATCAGGAGGAACACCGGTGGCGAAGGCGGGTCTCTGGGCCGATATTGACGCTGAGGAGCGAAAGCGTGGGG |
| P-asv5333 | GCAAGCGTTATTCGGAATTATTGGGCGTAAAGGGCGCGTAGGCGGCCTGTCAAGTCTCTCGTGAAATCCCTCGGCTTAACTGAGGACGTGCGGGAGATACTGTCAGGCTGGAGTCCGGAAGAGGGTGGTGGAATTCCCAGTGTAGCGGTGAAATGCGTAGATATTGGGAGGAACACCGGTGGCGAAGGCGGCCACCTGGGCCGGAACTGACGCTGAGGCGCGAAAGCGTGGGG |
| P-asv5334 | GCGAGCGTTGTTCGGAATTATTGGGCGTAAAGGGTGCGTAGGCGGTTTGATAAGTCTTATGTGAAATCTCTGGGCTCAACCCAGAGTCTGCATAGGAAACTGTCGGGCTTGAGTATGGGAGAGGTGAGTGGAATTTCCGGTGTAGCGGTGAAATGCGTAGATATCGGAAGGAACACCTGTGGCGAAAGCGGCTCACTGGACCATAACTGACGCTGATGCACGAAAGCTAGGGG |
| P-asv5335 | CCAAGCGTTGTTCGGATTCACTGGGCGTAAAGGGTGCGTAGGCGGTCGGGTAAGTCTGACGTGAAATCTTCAAGCTCAACTTGGAAACTGCGTCGGATACTATTCGGCTAGAGGAATGGAGGGGAGACTGGAATACTTGGTGTAGCAGTGAAATGCGTAGATATTAGGAGGAACACCGGTGGCGAAGGCGGGTCTCTGGGCCGATACTGACGCTGAGGAGCGAAAGCGTGGGG |
| P-asv5336 | GCCAGCGTTGTTCGGAATTACTGGGTGTAAAGGGTTCGTAGGCGGTGTGGCAAGTCGGGAGTGAAATCTCTGGGCTCAACCCAGAGACTGCTTCCGAAACTGCTGTGCTTGAGTGTGAGAGAGGCTCGTGGAATTGCAGGTGTAGCGGTGAAATTCGTAGATATCAGGAGGAACACCGGCGGCGAAGGCGGCCCTCTGGATCGTTGCTGACGCTGAGACGCGACAGCGTGGGG |
| P-asv5337 | GCAAGCGTTGTCCGGAATTATTGGGCGTAAAGCGCTCGTAGGTGGATCGGCAAGTCCGCAAGGAAAGACCCGGGCTCAACTCGGGAACCCGTGTGGATACTGCCGACCTTGAGGTAGTCAGAGGATGATGGAATTCCCGGTGTAGCGGTGAAATGCGTAGATATCGGGAGGAACACCAGTGGCGAAGGCGATCGTCTGGGGCTATTCTGACACTGAGGAGCGAAAGCTAGGGG |
| P-asv5338 | GCGAGCGTTGTCCGGAATCATTGGGCGTAAAGCGCGTGTAGGCGGCTCGATAAGTCTGGTGTGAAATCTTGGGGCCCAACCCCAAGCCTGCACTGGATACTGTCGAGCTTGAGTCCGGGAGGGGAGAGCGGAATTCCTAGTGTAGCGGTGAAATGCGCAGATATTAGGAAGAACACCGGTGGCGAAGGCGGCTCTCTGGAACGGTACTGACGCTGAGACGCGAAAGCGTGGGG |
| P-asv5339 | GCAAGCGTTGTTCGGAGTTACTGGGCGTAAAGGGTGTGTAGGCGGCTTTCTAAGTTTGTTGTGAAATCTCCCGGCTTAACCGGGAGGGTGCGGCGAAGACTGGAAGGCTAGAGTATGGGATGGGAGAGTGGAATTCCTGGTGTAGCGGTGAAATGCGTAGATATCAGGAGGAACACCTGCGGTGTAGACGGCTCTCTGGACCATTACTGACGCTGAGACACGAAAGCGTGGGT |
| P-asv5340 | GCTAGCGTTGTTCGGAATTACTGGGCGTAAAGCGCACGTGGGCGGGTTGTTAAGTCAGGGGTGAAATCCCGGAGCTCAACTCCGAAACTGCCTTTGATACTGGCAATCTAGAGACTGGAAGAGGTTAGTGGAATTCCCAGTGTAGAGGTGAAATTCGTAGATATTGGGAAGAACACCAGTGGCGAAGGCGGCTAACTGGTCCAGTACTGACGCTGAGGTGCGAAAGCGTGGGG |
| P-asv5341 | GCAAGCGTTGTTCGGAATTATTGGGCGTAAAGGGTGCTTAGGCGGTTTGATAAGTCATCTGTGAAATCTCTGGGCTCAACCCAGAGTCTGCAGGCGAAACTGTCGGGCTTGAGTATGGGAGAGGTGAGTGGAATTTCCGGTGTAGCGGTGAAATGCGTAGATATCGGAAGGAACACCTGTGGCGAAAGCGGCTCACTGGACCATAACTGACGCTGATGCACGAAAGCTAGGGG |
| P-asv5342 | GCTAGCGTTGTTCGGAATTACTGGGCGTAAAGCGCACGTAGGCGGATTGTTAAGTCAGAGGTGAAATCCCGGAGCTCAACTTCGGAACTGCCTTTGATACTGGCAATCTCGAGTCCGGAAGAGGTTAGTGGAATTCCCAGTGTAGAGGTGAAATTCGTAGATATTGGGAAGAACACCGGTGGCGAAGGCGGCCATCTGGACCGTATCTGACGCTGAGGCGCGAAAGCGTGGGG |
| P-asv5343 | GCGAGCGTTAATCGGAATTACTGGGCGTAAAGCGTGCGTAGACGGTCGTGCAAGTCGGGTGTGAAAGCCCCAGGCTCAACCTGGGAATTGCATTCGAAACTGCTCGGCTAGGGTGCGGAAGAGGGAAGCGGAATTTCCGGTGTAGCGGTGAAATCCGTAGATATCGGAAGGAACATCAGTGGTGAAAGCGACTTCCTGGTCCGACACCGACGTTCAGGCACGAAAGCGTGGGG |
| P-asv5344 | GCAAGCGTTGTTCGGATTTACTGGGCATAAAGCGCGCGTAGGTGGTCTTGTAAGCGGGGGATGAAAGCCCATGGCTCACCCATGGAACTGTCCCTCGGACTGCAAGGCTGGAGGGCGGGAGGGGAGAATGGAATTCCCGGTGTAGCGGTGAAATGCGTAGATATCGGGAGGAACGCCGGTGGCGAAAGCGATTCTCTGGCCCGTCCCTGACACTGAGGCGCGAGAGCTAGGGG |
| P-asv5345 | GCTAGCGTTGTCCGGATTTATTGGGCGTAAAGAGCGTGTAGGCGGCTTGGTAGGTCTGCTGTGAAAACTCGGGGCTCAACCCCGAGAGGTCGGTGGAAACCACCTGGCTAGAGTCCGGAAGAGGAGAGTGGAATTCCCGGTGTAGCGGTGAAATGCGCAGATATCGGGAAGAACACCCGTGGCGAAGGCGGCTCTCTGGGACGGTACTGACGCTGAGACGCGAAAGCGTGGGG |
| P-asv5346 | GCAAGCGTTGTCCGGATTTACTGGGCGTACAGCGCGCGCAGGCGGCGAGGTAGGTACCTGGTGACATTCTCGTGCTCAACACGAGACGGTCTGGGTAGACCGCCTGGCTGGAGGCAGGGAGAGGGATGTGGAATACCCGGTGGAGCGGTGAAATGCGTAGAGATCGGGTGGAACACCAGTGGCGAAGGCGGCATCCTGGCCCTGACCTGACGCTGAGGCGCGATAGCGTGGGG |
| P-asv5347 | ACGAGCGTTGTCCGGATTCATTGGGCGTAAAGCGCGCGTAGGCGGCTCGGAAAGTCGGTCGTGAAATGCCGGGGCTCAACCCCGGGACTGCGTCCGATACTTCCGGGCTGGAGGTAGGTAGGGGAGATCGGAATTCCTGGTGTAGCGGTGAAATGCGCAGATATCAGGAGGAACACCGGTGGCGAAGGCGGATCTCTGGTACGTGACTGACGCTGAGACGCGAAAGCGTGGGG |

| | |
|---|---|
| P-asv5348 | GCAAACGTTGTTCGGAATCACTGGGCTTAAAGGGCGCGTAGGCGGGTCTCCCAGTCTGTGGTGAAATCCTCCAGCTTAACTGGAGAAGTGCCGTGGATACTGGAGACCTCGAGGAGGGTAGGGGCATCTGGAA CGGTCGGTGGAGCGGTGAAATGCGTTGATATCGACCGGAACTCCGATGGCGAAGGCAAGGTGCTGGCCCCTTTCTGACGCTGAGGCGCGAAAGCCAGGGG |
| P-asv5349 | GCTAGCGTTGTTCGGAATTACTGGGCGTAAAGGGCGCGTAGGCGGTGCTGTAAGTCAGGCGTGAAATCCCCGGGCTCAACCTGGGAATTGCGTTTGATACTGCGGCGCTGGAATGCGGGAGAGGATGGCGGAA TTCCCAGTGTAGAGGTGAAATTCGTAGATATTGGAAGAACACCGGTGGCGAAGGCGGCCATCTGGCCCGTAATTGATGCTGAGGCGCGAAAGCGTGGGG |
| P-asv5350 | GTGAGCGTTGTCCGGATTTATTGGGCGTAAAGAGCTCGTAGGCGGCTTGTCGCGTCGACCGTGAAAACTTGGGGCTCAACCCTAAGCCTGCGGTCGATACGGGCAGGCTAGAGTTCGGTAGGGGAGACTGGAA TTCCTGGTGTAGCGGTGAAATGCGCAGATATCAGGAGGAACACCGGTGGCGAAGGCGGGTCTCTGGGCCGATACTGACGCTGAGGAGCGAAAGCGTGGGG |
| P-asv5351 | GCGAACGTTGCTCGGAATTACTGGGCGTAAAGCGTGTGTAGGCGGCCACGCAAGTCGGATGTGAAAGCCCTGGGCTCAACCCAGGAAGTGCACCCGATACTGCGACGGCTTGAGTACCGGAGAGGTTGGTGGA ATTCCTGGTGTAGCGGTGAAATGCGCAGATATCAGGAAGAACACCTATGGCGAAGGCAGCTCTCTGGGACGGTACTGACGCTGAGACGCGAAAGCGTGGGG |
| P-asv5352 | GCAAACGTTGTTCGGAATTACTGGGCATAAAGCGCACGTAGGCGGTTCGTTAAGTCATGTGTGAAATCCCCCGGCTCAACCGGGGAATTGCGCATGATACTGGCGAGCTAGAGATCGATAGGGGTGATCGGAAC TCCAGGTGGAGCGGTGAAATTCGTAGATATTCGCAAGAACACCAGTGGCGAAGGCGGCTCACTGGCCCGATACTGACGCTGAGGCGCGAAAGCGTGGGG |
| P-asv5353 | GCGAGCGTTGTCCGGAATTATTGGGCGTAAAGAGCGTGTAGGCGGTTCGGTAAGTCTGCCGTGAAAACCCAGGGCTCAACCCTGGGCGTGCGGTGGATACTGCCGGGCTAGAGGGTGGTAGAGGCGAGTGGA ATTCCCGGTGTAGCGGTGAAATGCGCAGATATCGGGAGGAACACCAGTAGCGAAGGCGGCTCGCTGGGCCATACCTGACGCTGGGACGCGAAAGCGTGGGG |
| P-asv5354 | GCAAGCGTTAATCGGAATTACTGGGCGTAAAGCGAGCGTAGGTGGCTTGATAAGTCAGATATGAAAGCCACGGGCTTAACCTGGGAACGGCATCTGATACTGTTAGGCTAGAGTAGGTGAGAGGAAGGTAGAAT TCCAGGTGTAGCGGTGAAATGCGTAGAGATCTGGAGGAATACCGATGGCGAAGGCAGCCTTCTGGCATCATACTGACACTGAGGTTCGAAAGCGTGGGT |
| P-asv5355 | GCTAGCGTTGTTCGGAATTACTGGGCGTAAAGCGCACGTAGGCGGCTTTGTGAGTCAGAGGTGAAAGGCATTGGCTCAACCAATGTAAGCCTTTGAAACTGTTGGGCTTGAGTGCAGGAGAGGTAAGTGGAATT CCTAGTGTAGAGGTGAAATTCGTAGATATTAGGAGGAACACCAGTGGCGAAGGCGACTTACTGGACTGTAACTGACGCTGAGGCACGAAAGCGTGGGG |
| P-asv5356 | GCAAGCGTTGTCCGGAATTATTGGGCGTAAAGCGCGCGCAGGCGGTCCTTTAAGTCTGATGTGAAAGCCCACGGCTCAACCGTGGAGGGTCATTGGAAACTGGGGGACTTGAGTGCAGAAGAGAAGAGTGGAA TTCCACATATAGCGGTGAAATGCGTAGAGATGTGGAGGAACACCAGTAGCGAAGGCAACTCTTTGGTCTGTAACTGACGCTGAGGCGCGAAAGCGTGGGG |
| P-asv5357 | GCAAGCGTTAATCGGAATTATTGGGCGTAGAGAGCTCGTAGGCGGCCTGTTGCGTCTGCTGTGAAAGCCCGGGGCTTAACCTGGGAATGGCGATGGATACTGGCGAGCTAGAGTGTGATAGAGGATGGCGGAA TTCCCGGTGTAGCGGTGAAATGCGTAGAGATCGGGAGGAACATCTGTGGCGAAGGCGGCCATCTGGATCAACACTGACGCTGAGGCACGAAAGCGTGGGG |
| P-asv5358 | GCAAGCGTTATCCGGAATTATTGGGCGTAAAGCATCCGCAGGGGGCGCATCAAGTCGGCTGTTAAAGGCTGGAGCTTAACTCCGGTAAGGCAGTGGAAACGGGTGCGCTAGAGGGCGACAGGGGTAGAGGGAA TTCCCAGTGTAGCGGTGAAATGCGCAGAGATTGGGAAGAACACCGGTGGCGAAAGCGCTCTACTGGGTCGCGCCTGACCCTCAGGGACGAAAGCTAAGGT |
| P-asv5359 | GCAAGCGTTAATCGGAATTACTGGGCGTAAAGCGTTCGTAGGCGGTCTATTAAGCAAGATGTGAAAGCCCAGGGCTCAACCTTGGAACTGCATTTTGAACTGGTAGACTAGAGTACTGTAGAGGGTGGTGGAATT TCCAGTGTAGCGGTGAAATGCGTAGAGATTGGAAGGAACATCAGTGGCGAAGGCGGCCACCTGGACAGATACTGACGCTGAGGAACGAAAGCGTGGGG |
| P-asv5360 | GCAAGCGTTACTCAGAATCACTGGGCGTAAAGAGCGTGTAGGCGGATAGATAAGTCAGAAGTGAAATCCAATAGCTTAACTATTGAACTGCTTTTGAGACTGTCTATCTAGAGTATGGGGGAGGTAGATGGAATTT CTGGTGTAGGGGTAAAATCCGTAGAGATCAGAAGGAATACCGATTGCGAAGGCGATCTACTGGAACATAACTGACGCTGAGACGCGAAAGCGTGGGG |
| P-asv5361 | ACAAGCGTTGTCCGGATTTATTGGGCGTAAAGGGAGCGCAGGCGGTCTGTTTAGTCTAATGTGAAAGCCCACGGCTTAACCGTGGAACGGCATTGGAAACTAACAGACTTGAATGTAGAAGAGGAAAATGGAATT CCAAGTGTAGCGGTGGAATGCGTAGATATTTGGAGGAACACCAGTGGCGAAGGCGATTTTCTGGTCTAACATTGACGCTGAGGCTCGAAAGCGTGGGG |
| P-asv5362 | GCTAGCGTTGTTCGGAATTACTGGGCGTAAAGGGCGCGTAGGCGGCCGTTTGCGTCGGATGTGAAAGCCCGGGGCTCAACCTCGGAACGGCATTCGATACGGGACGGCTGGAGACCGGGAGAGGAGAGTGGA ATAGCCAGTGTAGAGGTGAAATTCGTAGATATTGGCTGGAACACCGGTGGCGAAAGCGGCTCTCTGGCCCGGTTCTGACGCTGAGGCGCGAAAGCGTGGGG |
| P-asv5363 | CCTAGCGTTGTTCGGAATTACTGGGCGTAAAGGGAGCGCAGGCGGATTGATAAGTCTAAAGTGGAATCCCGGGGCTCAACCTCGGACGTGCTTTGGATACTGTCAGTCCTGAGTTCGTGAGAGGAAAGCGGAAT TCAGGGTGTAGCGGTGAAATGCTTAGATATCCTGAGGAACACCAGTGGCGAAAGCGGCTTTCTGGCACGAAACTGACGCTGAGGCTCGAAAGCGTGGGT |
| P-asv5364 | ACGAGCGTTATCCGGATTTATTGGGTTTAAAGGGTGCGCAGGTGGTCTTGCAAGTCAGTGGTGAAAAGCTGAGGCTCAACCTCAGCCTTGCCGTTGAAACTGTAAGACTTGAGAGTACATGATGTGGGCGGAATG CGTAGTGTAGCGGTGAAATGCATAGATATTACGCAGAACTCCGATTGCGAAGGCAGCTCACAAAGGTATATCTGACACTGAGGCACGAAAGCGTGGGG |
| P-asv5365 | CCAAGCGTTGTTCGGATTCACTGGGCGTAAAGGGTGCGTAGGTGGTGAGGTAAGTCGGATGTGAAAGCTCGGAGCTCAACTCCGAAATGGCATTGGAAACTACCTTGCTCGAGGGTTGGAGGGGGGACTGGAA TACTTGGTGTAGCAGTGAAATGCGTAGATATCAAGTGGAACACCAGTGGCGAAGGCGAGTCCCTGGACAACTCCTGACACTGAGGCGCGAAAGCTAGGGG |
| P-asv5366 | GCAAGCGTTAATCGGAATTACTGGGCGTAAAGCGCACGCAAGCGGTCTGTCAAGTCGGATGTGAAATCCCCGGGCTCAACCTAAGAACTGCATCCGAAACTGGCAGGCTAGAGTCTTGTAGAGGGGGGTAGAAT TCCAGGTGTAGCGGTGAAATGCGTAGAGATCTGGAAGAATACCGGTGGCGAAGGCGGCCCCCTGGACAAAGACTGACGCTCAGGTGCGAAAGCGTGGGG |
| P-asv5367 | GCAAGCGTTATCCGGATTTATTGGGTTTAAAGGGTCCGTAGGCGGGCCCGTAAGTCAGTGGTGAAATCTCGCAGCTTAACTGTGAAACTGCCATTGATACTGCGGGTCTTGAGTAAATTTGAAGTGGCTGGAATAA GTAGTGTAGCGGTGAAATGCATAGATATTACTTAGAACACCAATTGCGAAGGCAGGTCACTAAGATTTAACTGACGCTGATGGACGAAAGCGTGGGG |
| P-asv5368 | GCAAGCGTTAATCGGAATTACTGGGCGTAAAGCGTGCGCAGGCGGTTCGGAAAGAAAGGTGTGAAATCCCAGAGCTTAACTTTGGAACTACACTTTTAACTACCGGGCTAGAGTATGTCAGAGGGAGGTGGAATT CCGCGTGTAGCAGTGAAATGCGTAGATATGCGGAGGAACACCGATGGCGAAGGCAGCCTCCTGGGATAATACTGACGCTCATGCACGAAAGCGTGGGG |
| P-asv5369 | GCAAGCGTTACCCGGAATCACTGGGCGTAAAGGGCGTGTAGGCGGGAAGCTAAGTCCGACTTTAAAGACCGGGGCTCAACCCCGGGAGTGGGTTGGAGACTGGCTTTCTGGACCTCTGGAGAGGCAACTGGAA TTCCTGGTGTAGCGGTGGAATGCGTAGATACCAGGAGGAACGCCGATGGCGAAGGCAGGTTGCTGGACAGAAGGTGACGCTGAGGCGCGAAAGTGTGGGG |
| P-asv5370 | GCAAGCGTTATCGGAATTACTGGGCGTAAAGCGCGCGTAGGTGGTTCGTTAAGTTGGATGTGAAAGCCCCGGGCTCAACCTGGGAACGGCATCCGGAACTGTCAGTCTAGGGTGCAGGAGAGGAAGGTAGAAC TCCGGGTGTAGCGGTGAAATGCGTAGAGATCGGGAGGAATACCAGTGGCGAAGGCGGCCTTCTGGACTGACACTGACGCTGAGGTGCGAAAGCGTGGGT |
| P-asv5371 | GCTAGCGTTGCTCGGAATTACTGGGCGTAAAGGGAGCGTAGGCGGACTGTTAAGTTAGAGGTGAAAGCCCAGGGCTCAACCTTGGAATTGCCTTTGATACTGGCAGTCTTGAGTACGGAAGAGGTATGTGGAAC TCCGAGTGTAGAGGTGAAATTCGTAGATATTCGGAAGAACACCAGTGGCGAAGGCGACATACTGGTCCGTTACCGACGCTGAGGCTCGAAAGCGTGGGG |
| P-asv5372 | GCAAGCGTTGTCCGGATTTATTGGGCGTAAAGAGTTCGTAGGCTGTTTGTTAAGTCTGATGTTAAAGACTGGGGCTCAACCTCGGAAATGCATTGGATACTGGCAGACTGGAGTGCAGTAGAGGCTAGTGGAATT CCCAGTGTAGCGGTGAAATGCGTAGATATTGGGAAGAACACCGGTGGCGTAGGCGACTAGCTGGGCTGTAACTGACGCTGAGGAACGAAAGCCAGGGG |
| P-asv5373 | GCTAGCGTTGTTCGGATTTACTGGGCGTAAAGCGCACGTAGGCGGATTGTTAGGTTAGGGGTGAAATCCCAGGGCTCAACTCCAGAATTGCCTTTAAGACTGCATCGCTTGAATCCAGGAGAGGTGAGTGGAATT CCGAGTGTAGAGGTGAAATTCGTAGATATTCGGAAGAACACCAGTGGCGAAGGCGGCTCACTGGACTGGTATTGACGCTGAGGTGCGAAAGCGTGGGG |
| P-asv5374 | GCGAGCGTTAATCGGAATTACTGGGCGTAAAGCGTGCGCAGGCGGTTTCGTAAGCTGGAGGTGAAATCCCCGGGCTTAACCTGGGAATGGCCTTCAGGACTGCGAGGCTAGAGTGCGGCAGAGGGAGGTGGA ATTCCACGTGTAGCAGTGGAATGCGTAGAGATGTGGAGGAACACCGATGGCGAAGGCAGCCTCCTGGGCCAGCACTGACGCTCATGCACGAAAGCGTGGGG |
| P-asv5375 | CCAAGCGTTAATCGGAATTACTGGGCGTAAAGCGCACGTAGGCGGTTTGTTAAGCAAGATGTGAAAGCCCAGGGCTCAACCTTGGAATAGCATTTTGAACTGGCAGACTAGAGTTTTGAAGAGGGTGGTAGAATT TCCAGTGTAGCGGTGAAATGCGTAGATATTGGAAGGAATACCAGTGGCGAAGGCGGCCACCTGGTCAAAAACTGACGCTGAGGAGCGAAAGCGTGGGG |
| P-asv5376 | ACGAACGTTATTCGGAATTACTGGGCTTAAAGGGTTTGTAGGCGGCCTTGTAAGTCAGGTGTGAAAGCCCTCGGCTCAACCGAGGAACAGCGCTTGATACTGCAAGGCTTGAGGGAGACAGGGGTAAGCGGAAC TGATGGTGGAGCGGTGAAATGCGTTGATATCATCAGGAACACCGGTGGCGAAAGCGGCTTACTGGGTCTCTTCTGACGCTGAGGAACGAAAGCTAGGGT |
| P-asv5377 | CCGAGCGTTGTCCGGAATCATTGGGCGTAAAGGGTACGTAGGCGGCCTAGTAAGTTAGAAGTGAAATAATATAGCTCAACTATATAAAGCTTTTAAAACTGCAAGGCTTGAGAGATGGAAGGGAAAGTGGAATTCC TAGTGTAGCGGTGAGATGCGCAGATATTAGGAAGAATACCAGTGGCGAAGGCGACTTTCTGGTCATCCTCTGACGCTGAGGTACGAAAGCGTGGGT |
| P-asv5378 | CCAGGCGTTATCCGGATTTATTGGGTTTAAAGGGAGCGTAGGCGGATTGTTAAGTCAGCGGTTAAAGGGTGTGGCTCAACCATGCATTGCCGTTGAAACTGGCGATCTTGAGTGCAGACAGGGATGCCGGAATTC GTGGTGTAGCGGTGAAATGCTTAGATATCACGAAGAACTCCGATCGCGAAGGCAGGTGTCCGGGCTGCAACTGACGCTGAGGCTCGAAAGTGTGGGT |
| P-asv5379 | GCAAGCGTTGCTCGGAATTACTGGGCGTAAAGCGCACGCAGGCGGTTTATCAGGTCGGATGTGAAAGCCCGGGGCTCAACCCCGGAACTGCATTCGAAACCGGTAGACTTGAGTTCGAGAGAGGGTGGTGGAA TTCCTGGTGTAGAGGTGAAATTCGTTGATATCAGGAGGAACACCGGTGGCGAAGGCGACCACCTGGCTCGATACTGACGCTCATGTGCGAAAGCGTGGGG |
| P-asv5380 | ACAAGCGTTATCCGAATTTATTGGGCGTAAAGCGCGTCTAGGTGGTTATGTAAGTCTGATGTGAAAATGCAGGGCTCAACTCTGTATTGCGTTGGAAACTGCATGACTAGAGTACTGGAGAGGTAAGCGGAACTAC AAGTGTAGAGGTGAAATTCGTAGATATTTGTAGGAATGCCGATGGGGAAGCCAGCTTACTGGACAGATACTGACGCTAAAGCGCGAAAGCGTGGGT |
| P-asv5381 | GCAAGCGTTAATCGGAATTACTGGGCGTAAAGGGTGCGTAGGTGGTTATGTAAGTTAGCTGTGAAATTCCTGGGCTTAACCTAGGGGGGTCAGCTAAGACTGTGTGACTAGAGTATGGGAGAGGGTAGTGGAAT TTCCGGTGTAGCGGTGAAATGCGTAGAGATCGGAAGGAACACCAGTGGCGAAGGCGGCTACCTGGCCTAATACTGACACTGAAGCACGAAAGCGTGGGG |
| P-asv5382 | CCGAGCGTTGTCCGGAATTATTGGGCGTAAAGGGCTCGTAGGCGGTTTGTCACGTCGGGAGTGAAATCTTACAGCTTAACTGTGAACTTGCTTCCGATACGGGCAGACTTGAGGCATTCAGGGGAGAACGGAATT CCTGGTGTAGCGGTGAAATGCGCAGATATCAGGAGGAACACCGGTGGCGAAGGCGGTTCTCTGGGAATGTACTGACGCTGAGGAGCGAAAGTGTGGGG |
| P-asv5383 | GCAAGCGTTATCCGGATTTACTGGGTGTAAAGGGAGCGCAGACGGCCAAGCAAGTCTGAAGTGAAATGCATGGGCTCAACCCATGAATTGCTTTGGAAACTGTTTGGCTTGAGTGTCGGAGGGGTAAGCGGAAT TCCTAGTGTAGCGGTGAAATGCGTAGATATTAGGAGGAACACCGGAGGCGAAGGCGGCTTACTGGACGACAACTGACGTTGAGGCTCGAAGGCGTGGGG |
| P-asv5384 | GCAAGCGTTGTCCGGAATTATTGGGCGTAAAGAGCTCGTAGGCGATTTGTCGCGTCTGCTGTGAAAACGCGAGGCTCAACCTCGCGCCTGCAGTGGGTACGGGCAAACTTGAGTGCGGTAGGGGAGATTGGAA TTCCTGGTGTAGCGGTGGAATGCGCAGATATCGGGAGGAGCACCGATGGCGAAGGCAGATCTCTGGGCCGTAACTGACGCTGAGGAGCGAAAGCATGGGG |
| P-asv5385 | GCGAGCGTTAATCGGAATTACTGGGCGTAAAGCGCATGCAGGCGGCTTGTTAAGCAAGATGTGAAAGCCCGGGGCTTAACCTCGGAACCGCATTTTGAACTGGCAGGCTAGAGTCTTGTAGAGGGGGGTAGAAT TCCAGGTGTAGCGGTGAAATGCGTAGAGATCTGGAGGAATACCGGTGGCGAAGGCGGCCCCCTGGACAAAGACTGACGCTCAGGTGCGAAAGCGTGGGG |
| P-asv5386 | GCAAGCGTTATCCGGATTTATTGGGCGTAAAGCGCTCGCAGGCGGTCTTTTAAGTCTGATGTGAAAGCCCACGGCTCAACCGTGGAGGGTCATTGGAAACTGGAAGACTTGAGTGCAGAAGAGGAGAGTGGAAT TCCACGTGTAGCGGTGAAATGCGTAGAGATGTGGAGGAACACCAGTGGCGAAGGCGACTCTCTGGTCTGTAACTGACGCTGAGGAGCGAAAGCGTGGGG |

| | |
|---|---|
| P-asv5387 | GCAAGCGTTATTCGGAATGACTGGGCGTAAAGAGCACGTAGGCGGCTTATTAAGTCTGATGTGAAAGCCCCGAACTCAATTTGGGAAGTGCATTGGATACTGGTAAGCTAGAGTATAGGAGAGGAGAGTGGAATTCCAGGTGTAGAGGTGAAATTCGTAGATATCTGGAGGAACAACGGAGGCGAAGGCGACTCTCTGGACTATTACTGACGCTGAGGTGCGAAAGCGTGGGG |
| P-asv5388 | GCTAGCGTTGCTCGGAATTACTGGGCGTAAAGGGAGCGTAGGCGGGTTGACAAGTTGGAGGTGAAAGCCCAGGGCTCAACCCTGGAATTGCCTTCAAAACTGTCAGCCTTGAGTATGGAAGAGGTAAGTGGAACTCCGAGTGTAGAGGTGAAATTCGTAGATATTCGGAAGAACACCAGTGGCGAAGGCGACTTACTGGTCCATGACTGACGCTGAGGCTCGAAAGCGTGGGG |
| P-asv5389 | GCGAGCGTTATTCGGAATCATTGGGTTTAAAGGGTCTGTAGGCGGGCTATTAAGTCAGGGGTGAAAGGTTTCAGCTTAACTGAGAAATTGCCTTTGATACTGGTAGTCTTGAATATCTGTGAAGTTCTTGGAATGTGTAGTGTAGCGGTGAAATGCTTAGATATTACACAGAACACCGATTGCGGAGGCAGGGGACTAACAGACGATTGACGCTGAGAGACGAAAGCGTGGGG |
| P-asv5390 | ACAAGCGTTGTCCGGATTTATTGGGCGTAAAGCGAGCGCAGGCGGTCTTTTAAGTCTGATGTGAAAGCCCACGGCTCAACCGTGGATGGTCATTGGAAACTGGAAGACTTGAATGCAGAAGAGGAAAGTGGAATTCCATGTGTAGCGGTGAAATGCGTAGATATATGGAAGAACACCAGTGGTGAAGGCGACTTTCTGGTCTGTAATTGACGCTGAGGCTCGAAAGCGTGGGG |
| P-asv5391 | GCGAGCGTTAATCGGAATTACTGGGCGTAAAGGGTACGTAGGCGGTCCGTTAAGTCGGATGTGAAAGCCCTGGGCTTAACCTGGGAATTGCATTCGATACTGGCGGACTTGAGTCTGACAGAGGGAAGTGGAATTCCGGGTGTAGCGGTGAAATGCGTAGATATCCGGAAGAACACCAGTAGCGAAGGCGGCTTCCTGGGTCAAGACTGACGCTGAGGTACGAAAGCGTGGGG |
| P-asv5392 | GCAAGCGTTAATCGGAATCACTGGGCGTAAAGGGTGCGTAGGCGGTACGTTAGGTCTGTCGTGAAATCCCCGGGCTCAACCTGGGAATGGCGGTAGAAACCGGCGCACTAGAGTATGGGAGAGGGTGGTGGAATTCCAGGTGTAGCGGGGAAATGCGTAGAGATCTGGAAGAACATCAGTGGCGAAGGCGGCCACCTGGCCCAATACTGACGCTGAGGCACGAAAGCGTGGGG |
| P-asv5393 | GCAAGCGTTGTCCGGAATCATTGGGTGTAGAGGGTGCGCAGGCGGGCGTCTAAGTCTGGGGTGGAATCTTACCGCTTAACGGTAAAATGGCCCCCGATACTGGGCGTCTTGAGCACGACAGAGGTTGGTGGAATTCGTAGTGTAGCGGTGAAATGCATAGATATTACGAAGAACACCGGTGGCGAAAGCGGCCAACTGGGTCGGAGCTGACGCTGAGGCACGAGAGCGTGGGG |
| P-asv5394 | GCTAGCGTTGTTCGGAATTACTGGGCGTAAAGGGCGCGTAGGCGGCCCGAGAAGTCGGATGTGAAAGCCCGGGGCTCAACCTCGGAACGGCATTCGAAACTTTCGGGCTCGAGAACGGAAGAGGAGAGTGGAATACCCAGTGTAGAGGTGAAATTCGTAGATATTGGGTGGAACACCGGTGGCGAAAGCGGCTCTCTGGTCCGGGTCTGACGCTGAGGCGCGAAAGCGTGGGG |
| P-asv5395 | GCAAGCGTTAATCGGAATTACTGGGCGTAAAGGGTGCGTAGGCGGCATGACAAGCTAGATGTGAAAGCCCTGGGCTTAACCTGGGAAGTGCATCTGGGACTGTTGAGCTAGAGTATGGTAGAGGAAAGTGGAATTTCTAGTGTAGCGGTGAAATGCGTAGATATTAGAAAGAACACCAGAGGCGAAGGCGACTTTCTGGACCGATACTGACGCTGAGGCACGAAAGCGTGGGG |
| P-asv5396 | CCAAGCGTTGTTCGGAATCACTGGGCTTAAAGGGTGCGTAGGTGGCCAGGCAAGTCAGTGGTGAAAGTCGTCCGCCCAACGGGCGAATTGCCATTGATACTGCCTGGCTCGAGTGTGCTAGGGGGATGCGGAACGAGTGGTGTAGCGGTGAAATGCGTAGATATCACTCGGAAGGCCAGAGGCGAAGGCGGCATCCTGGGGCACAACTGACACTGAGGCACGAGAGCGTGGGG |
| P-asv5397 | ACGAGTGATGGCCGATATTATTGGGCCTAAAGCGTTCGTAGCCGGCCGGTCAAGTCTGTTGGGAAATCCACGTGCCCAACACGTGGGCGTCCAGCAGAAACTGACTGGCTTGGGGCCGGAAGACTCGAGGGGTACGTCCGGGGTAGGAGTGAAATCCCGTAATCCTGGACGGACCACCGGTGGCGAAAGCGCCTCGAGAAGACGGACCCGACGGTGAGGAACGAAAGCTGGGGT |
| P-asv5398 | GCAACCGTTGTTCGGAATCACTGGGCATAAAGCGCGCGTAGGCGGCGTGGTAAGTCAGGTGTGGAAGCCCTCGGCCCAACCGAGGAATCGCGCTTGAAACTACCTCGCTTGAGGACAGGAGGGGAGCGTGGAATTCCTGGTGGAGCGGTGAAATGCGTTGATATCAGGAGGAACACCGGCGGCGAAGGCGACGCTCTGGCCTGTTTCTGACGCTGAGGTGCGAAAGCCAGGGG |
| P-asv5399 | GCGAGTGATGACCGATCTTATTGGGCCTAAAGCGTCCGTAGCCGGCCGCGCAAGTCCATCGGGAAATCCAGTCGCTTAACGACTGGGCGTCCGGTGGAAACTGCACGGCTTGGGGCCGGAAGACCTCAGGGGTACGTCCGGGGTAGGAGTGAAATCCCGTAATCCTGGACGGACCACCGATGGGGAAACCACCTCAAGAGAACGGATCCGACAGTGAGGGACGAAAGCTAGGGT |
| P-asv5400 | CCAAGCGTTAATCAGAATTATTGGGCGTAAAGCGCGCGCAGGCGGCTTTGTAAGTCTTGTGTTTAAACGCGGGGCTCAACCCCGTGTCGCATGGGAAACTGCAAGGCTTGAGTGCAGAAGAGGAAAGTGGAATTCCACGTGTAGCGGTGAAATGCGTAGAGATGTGGAGGAACACCAGTGGCGAAGGCGACTTTCTGGGCTGTAACTGACGCTGAGGCGCGAAAGCGTGGGG |
| P-asv5401 | ACGAACGTTATTCGGAATCACTGGGCTTAAAGGGTGCGCAGGCTGTGCAGAAAGTCGGGTGTGAAATCCCTCGGCTCAACCGAGGAACTGCGCCCGAAACTACTGTGCTCGAGGGAGATAGAGGTGAGCGGAACTTAGGGTGGAGCGGTGAAATGCGTTGATATCCTAAGGAACACCGGTGGCGAAAGCGGCTCACTGGATCTCTTCTGACGCTCATGCACGAAAGCCAGGGG |
| P-asv5402 | GCTAGCGTTATTCGGAATTATTGGGCGTAAAGAGCGCGTAGGCTGGTTAGTAAGCTAAAAGTGAAATCCCAAAGTTCAACTTTGGAATTGCTTTTAAAACTGCTAACCTAGAGATTGAAAGAGGATAGAGGAATTCCTAGTGTAGAGGTGAAATTCGTAAATATTAGGAGGAGCACCAGTGGCGAAGGCATCTATCTGGTTCAAATCTGACGCTGAGGCGCGAAGGCGTGGGG |
| P-asv5403 | CCAAGCATTATCCGGATTTACTGGGCGTAAAGGGTGCGTAGGCGGTTATATTAGTCGGGTGTTAAATCCTGGGGCTCAACCTCAGGTTCGCATTCGAAACGGTATAACTAGAAGGTGTGAGAGGTAAGCAGAACTCATGGTGTAGGGGTGAAATCCGTTGATATCATGGGGAATACCAAATGCGAAGGCAGCTTACTGGCACATTCTTGACGCTGAGGCACGAAAGCGTGGGT |
| P-asv5404 | GCAAGCGTTACTCGGAATTACTGGGCGTAAAGCGTGCGTAGGCGGTTCGTTAAGTCCGATGTGAAAGCCCTGGGCTCAACCTGGGAACTGCATTGGAAACTGGCGGGCTGGAGTGCGGTAGAGGGTGGCGGAATTCCCGGTGTAGCAGTGAAATGCGTAGATATCGGGAGGAACACCCGTGGCGAAGGCGGCCACCTGGACCAGCACTGACGCTGAGGCGCGAAAGCGTGGGG |
| P-asv5405 | GCAAGCGTTGTTCGGAATTATTGGGCGTAAAGGGCATGTAGGCGGTTTGGTAAGCCTGGTGTGAAATCCTGCAGCTTAACTGTAGAATTGCATTGGGTACTGCCAGACTTGAATCACAGAGGGGAAACTAGAATTCCAGGTGTAGGGGTGGAATCTGTAGATATCTGGAAGAATACCGGTGGCGAAGGCGGGTTTCTAGCTGATGATTGACGCTGAGGTGCGAAAGTGCGGGG |
| P-asv5406 | GCTAGCGTTGTTCGGAATTACTGGGCGTAAAGGGCGTGTAGGCGGTACCATAAGTCAGATGTGAAAGCCCTAGGCTTAACCTAGGAGGTGCATTTGATACTGTGGAACTAGAGATCGAGAGAGGAAAGTGGAATTACGAGTGTAGAGGTGAAATTCGTAGATATTCGTAAGAACACCAGTGGCGAAGGCGACTTTCTGGCTCGATACTGACGCTAAGGCGCGAAAGCGTGGGG |
| P-asv5407 | GCAAGCGTTAATCGGAATTACTGGGCGTAAAGCGCGCGTAGGTGGTTTGTTAAGTCGGATGTGAAATCCCTGGGCTCAACCTAGGAATGGCATTCGAAACTGACGAACTAGAGTATGGTAGAGGGAAGGGGAATTCCAGGTGTAGCGGTGAAATGCGTAGATATCTGGAGGAACATCAGTGGCGAAGGCGCCTTCCTGGACTAATACTGACACTGAGGTGCGAAAGCGTGGGT |
| P-asv5408 | GCAAGCGTTATCCGGAATTATTGGGCGTAAAGGGCGTGTAGGTGGTCTTTTAAGTCAGGAGTTAAATTTTTGAGCTCAACTCAAAAACTGCTTTTGAAACTAATTGACTAGAGTTTAGGAGAAGAAGGTGGAATTATCAGAGTAGCGGTGAAATGCGTAGATCCTGATAGGAAGGCCAAAGACGAAGGTAGCCTTCTGGACTAATACTGACACTGAGGCGCGAAAGCTAGGGG |
| P-asv5409 | GCAAGCGTTGTTCGGAATCACTGGGCTTAAAGCGTGCGTAGGCGGATTTGCAAGTCAGAAGTGAAATCCCCGGGCTCAACCCGGGAATTGCTTTTGAAACTGCAAGTCTAGAAGCCGGGAGAGGTTAGCAGAATTCCGAGTGTAGAGGTGAAATTCGCAGATATTCGGAGGAATACCAGTGGCGTAGGCGGCTAACTGGACCGGTTTTGACGCTGAGGCACGAAAGCGTGGGG |
| P-asv5410 | GCGAGCGTTGTTCGGAATTATTGGGCGTAAAGGGCGTGTAGGCGGTTAAATAAGTCTTGTTTTAAAGACCGGTGCTCAACACCGGGTATGGACAGGAAACTGTTTGACTAGAATCCGGTAGGGAAATGTGGAATTCCTAGTGTAGGGGTGAAATCTGTTGATATTAGGAAGAACACCAATGGCGAAGGCAACATTTTGGGCCTGGATTGACGCTGAGGCGCGAAAGCATAGGG |
| P-asv5411 | GCAAGTGTTATCCGGAATCACTGGGCGTAAAGCGTCTGTAGGTGGTTTAATAAGTCAACTGTTAAATCTTGAAGCTCAACTTCAAAATCGCAGTCGAAACTATTAGACTAGAGTATAGTAGGGGTAAAGGGAATTTCCAGTGGAGCGGTGAAATTCGTAGATATTCGGAAGAACACCAGTGGCGAAGGCGACTGACTGGACCGGTATTGACGCTGAGGTACGAAAGCGTGGGT |
| P-asv5412 | GCAAGCGTTAACCGGATTTACTGGGCGTAAAGAGCGCGTAGGTTGGGGAATAAGTCAGATGTGAAATACCGGTACTCAATACCGGTGCTGCATTTGAAACTGTTCTTCTAGAGGGTAGATGGAGAAAGCGGAATTCCGTATGTAGCGGTGGAATGCGTAGATATACGGAAGAACACCCGTGGCGAAAGCGGCTTTCTAATTTATTTCTGACACTGAGGCGCGAAAGCGTGGGG |
| P-asv5413 | GCAAGCGTTATCCGGATTTATTGGGTTTAAAGGGTCCGTAGGCGGCTAAATAAGTCAGTGGTGAAAGCCGGGCAGCTTAACTGTCGAACTGCCATTGATACTGTTTAGCTTGAATTGATTTGAAGTAGCTGGAATGTGTAGTGTAGCGGTGAAATGCATAGATATTACACAGAACACCGATAGCGAAGGCAGGTTACTAAGATCATATTGACGCTGAGGGACGAAAGCGTGGGG |
| P-asv5414 | GCAGGTGTTATTCATCCTGACTGGGCGTAAAGGGCACGTAGGCGATTTACTGTATTTTTAGTAAAAGTCTCCGGCTTAACCGGAGAACGACTTTTTAGATTAGTAGATTTGAGTTTGTTTGAATAGGATGGTACGTTTAGTGTAGAAGTGAAATTTGCAGAAATTCGACGGACCACCGTCTGGCGAAGGCGATCCTATAATAGTTAACTGACGCTAAGGTGCAAAAGCATAGGG |
| P-asv5415 | CCGAACGTTATTCGGTATCACTGGGCTTAAAGCGTTCGTAGGCGGCCTAGTAGGTGAGATGTGAAAGCCCTCGGCTCAACCGAGGAACAGCGTTTCAAACCACTAGGCTAGAGGGTGGCAGGGGTAATGGGAACTGATGGTGGAGCGGTGAAATGCGTTGATATCATCAGGAACACCGGTGGCGAAAGCGCATTACTGGGCCATTTCTGACGCTGAGGAACGAAAGCTAGGGT |
| P-asv5416 | GCTAGCGTTATTCGGAATTACTGGGCGTAAAGCGCACGTAGGCGGATCAGTCAGTTAGGGGTGAAATCCCGGGGCTCAACCCCGGAACTGCCTTTAATACTGCTGATCTAGAGTCCGAGAGAGGTGAGTGGAATTCCGAGTGTAGAGGTGAAATTCGTAGATATTCGGAGGAACACCAGTGGCGAAGGCGGCTCACTGGCTCGGTACTGACGCTGAGGTGCGAAAGCGTGGGG |
| P-asv5417 | GCAAGCGTTGTTCGGAATCACTGGGCGTAAAGCGCGCGCAGGCGGGACCTGCAAGTCGGAAGTGAAATCCCGGGGCTCAACTCCGGGGCTGCTTCCGAAACTGCGGACCTGGAGACCGGTAGAGGTGGGTGGAATTCCCGGTGTAGCGGTGGAATGCGTAGATATCGGGAAGAACACCTGTGGCGAAGGCGGCCCACTGGGCCGGATCTGACGCTGAGGCGCGAAAGCGTGGGG |
| P-asv5418 | GCGAGCGTTAATCGGAATTACTGGGCGTAAAGCGTACGTAGGCGGATTATTAAGCGAGATGTGAAATCCCGGGGCTCAACCTCGGAACTGCATTTCGAACTGGTAGTCTAGAGTATTGTAGAGGGTGGTGGAATTTCCAGTGTAGCGGTGAAATGCGTAGAGATTGGAAGGAACATCAGTGGCGAAGGCGGCCACCTGGACAAATACTGACGCTGAGGTACGAAAGCGTGGGG |
| P-asv5419 | ACAAGCGTTGTCCGGATTTATTGGGCGTAAAGCGAGCGCAGGCGGTCTTTTAAGTCTGATGTGAAAGCCCACGGCTCAACCGTGGATGGTCATTGGAGACTGGAAGACTTGAATGCAGAAGAGGAAAGTGGAATTCCATGTGTAGCGGTGAAATGCGTAGATATATGGAGGAACACCAGTGGCGAAGGCGACTTTCTGGTCTGTAATTGACGCTGAGGCTCGAAAGCGTGGGG |
| P-asv5420 | ACAAGTGTTGAGCGTCTTGACTGGGCCTAAAGGGTGGGCAGGTGGATTTCAGAGTCTGTTTAGTGATTTCCCCACTCCCGAGTGGGCCTCCTAAAGGAACCTGATCTCTTTGGGTCACAATCAGGATCAGAGAATAGTCCAAGCACCGGTAAAATGGGACCATATGGATTGGAATGCCAATGGCGAAGGCACTGGTCTGGTTGTGCACCGACACTCAGCCACGGAAGCGTGGGG |
| P-asv5421 | GCTAGCGTTGCTCGGAATGACTGGGCGTAAAGGGCGCGTAGGCGGACGGCACAGTCAGGCGTGAAATTCCTGGGCTCAACCTGGGGACTGCGTCTGAGACGTGTTGTCTTGAGTATGGAAGAGGGTTGTGGAATTTCCAGTGTAGAGGTGAAATTCGTAGATATTGGAAAGAACACCGGTGGCGAAGGCGGCTTACTGGCCCGATACTGTCACTGAGGCACGAAAGCGTGGGG |
| P-asv5422 | GCAAGCGTTAATCGGAATTACTGGGCGTAAAGCGTGCGCAGGCGGTTCGGAAAGAAAGATGTGAAATCCCAGGGCTCAACCTTGGAACTGCATTTTTAACTCCCGAACTAGAGTATGTCAGAGGGGGGTGGAATTCCACGTGTAGCAGTGAAATGCGTAGATATGTGGAGGAACACCGATGGCGAAGGCAGCCCCCTGGGATAATACTGACGCTCATGCACGAAAGCGTGGGG |
| P-asv5423 | GCAAGCGTTAATCGGAATTACTGGGCGTAAAGCGTGCGCAGGCGGTTTGTTAAGACAGATGTGAAATCCCCGGGCTCAACCTGGGAACTGCATTTGTGACTGGCAAGCTAGAGTCTGGCAGAGGGGGGTGGAATTCCGCGTGTAGCAGTGAAATGCGTAGATATGCGGAGGAACACCGATGGCGAAGGCAGCCCCCTGGGCCAACACTGACGCTCATGCACGAAAGCGTGGGG |
| P-asv5424 | GCTAGCGTTGCTCGGAATTACTGGGCGTAAAGGGCGCGTAGGCGGCTTAGTTAGTCAGGCGTGAAATTCCTGGGCTCAACCTGGGGACTGCGCTTGATACGGCTAGGCTAGAGGATGGAAGAGGGTCGCGGAATTCCCAGTGTAGAGGTGAAATTCGTAGATATTGGGAAGAACACCGGTGGCGAAGGCGGCGACCTGGTCCATTACTGACGCTGAGGCGCGACAGCGTGGGG |
| P-asv5425 | TCGAGTGATGGCCGATTTTATTGGGCCTAAAGCGTTCGTAGCCGGCCGAGCAAGTCCATCGGGAAATCCGCCAGCCCAACTGGCGGGCGTCCGGTGGAAACTGCACGGCTCGGGACCGGAAGACCCGAAGGGTACGTCTTGGGTAGGAGTGAAATCCCGTAATCCTGGACGGACCACCGATGGCGAAAGCACTTCGGGAAGACGGATCCGACGGTGAGGGACGAAAGCCAGGGT |

| | |
|---|---|
| P-asv5426 | GCAAGCGTTACTCGGAATCACTGGGCGTAAAGGACGCGTAGGCTGTTTAGCAAGTCAGATGTGAAATCCAATGGCTCAACCATTGAACTGCATTTGAAACTGCTAGACTAGAGTATGGGAGGGGGAGATGGAATTAGTGGTGTAGGGGTAAAATCCGTAGATATCACTAGGAATACCTAAAGCGAAGGCGATCTCCTGGAACATTACTGACGCTAAGGCGTGAAAGTGTGGGG |
| P-asv5427 | GCAAGCGTTAATCGGAATTACTGGGCGTAAAGCGCATGCAGGTGGTTTGTTAAGTCAGATGTGAAAGCCCTGGGCTCAACCTAGGAATCGCATTTGAAACTGACAAGCTAGAGTACTGTAGAGGGGGGTAGAATTTCAGGTGTAGCGGTGAAATGCGTAGAGATCTGAAAGAATACCGGTGGCGAAGGCGGCCCCCTGGACAGATACTGACACTCAGATGCGAAAGCGTGGGG |
| P-asv5428 | GCAAGCGTTAATCGGAATTACTGGGCGTAAAGCGTGCGTAGGTGGCCAGGCAAGTCTATCGGGAAATCCACCTGCCCAACAGGTGGGCGTCCGGTAGAAACTGCCCGGCTTGGAACCGGAAGGCTCAGAGAGTACGTCCGGGGTAGGAGTGAAATCCCGTAATCCCGGACGGACTACCGATGGCGAAAGCACTCTGAGAAGACGGCTTCGACAGTGAGGGACGAAAGCTAGGGT |
| P-asv5429 | GCGAGCGTTATCCGGATTTATTGGGTTTAAAGGGTTCGTAGGCGGGAGAATTAGTCAGTGGTGAAAGCCTGCAGCTTAACTGTAGAACTGCCATTGATACTGTTCTTCTTGAATATATATGAAGTGGGCGGAATGCGTCATGTAGCGGTGAAATGCACAGATATGACGCAGAACACCGATAGCGAAGGCAGCTCACTAATATATTATTGACGCTCAGGAACGAAAGCATGGGG |
| P-asv5430 | CCAAGCGTTATCCGGATTTATTGGGTTTAAAGGGTGCGTAGGCGGCCTGTTAAGTCAGGGGTGAAAGACGGCAGCTCAACTATCGCAGTGCCCTTGATACTGATGGGCTTGAATGGACTAGAGGTAGGCGGAATGAGACAAGTAGCGGTGAAATGCATAGATATGTCTCAGAACACCGATTGCGAAGGCAGCTTACTATGGTCTTATTGACGCTGAGGCACGAAAGCGTGGGG |
| P-asv5431 | GCAAACGTTATTCGGAATCACTGGGCTTAAAGGGTGCGTAGGCGGCTCATTAAGTAGGGTGTGAAAGCCCCCGGCTCAACCGGGGAATTGCGCTCTAAACTGATGGGCTGGAGTGAGATAGGGGTGTGTGGAACTTCCGGTGGAGCGGTGAAATGTGTTGATATCGGAAGGAACGCCGGTGGCGAAAGCGACACACTGGGTCTCAACTGACGCTGAGGCACGAAAGCCAGGGG |
| P-asv5432 | GCTAGCGTTGTTCGGAATTACTGGGCGTAAAGCGCGTGTAGGCGGTTGTTCAAGTCGGGTGTGAAAGCCTTGAGCTCAACTCAAGAAATGCACTCGGTACTGGATGACTAGAGGACCGGAGAGGATAGTGGAATTCCCAGTGTAGTGGTGAAATACGTAGAGATTGGGAAGAACACCAGTGGCGAAGGCGGCTATCTGGACGGTTACTGACGCTAAGACGCGAAAGCGTGGGG |
| P-asv5433 | GCGAGCGTTATTCGGATTTATTGGGCGTAAAGGGTTCGTAGGAGGTTGGATAAGTTTGATGTGAAAGCCCGTGGCTCAACCACGGAACTGCATTGGAAACTGTCCAACTTGAGGTTCGGAGAGGAAAGCAGAATTCCCGGTGTAGCGGTGGAATGCGTAGATATCGGGAGGAATACCGACAGCGAAGGCAGCTTTCTGGACGACACCTGACTCTGAGGAACGAAAGCCTGGGG |
| P-asv5434 | GCAAGCGTTGTCCGGATTTACTGGGCTTAAAGAATGCGTAGGCGGATATTTAAGTGAGATGTGAAATTCCCGGGCTTAACTTGGGAGCTGCATTTCAAACTGGATATCTAGAGTGCGGGAGAGGAAAGTGGAATTCCTAGTGTAGCGGTGAAATGCGTAGAGATTAGGAAGAACACCAGTGGCGAAGGCGACTTTCTGGACCGTAACTGACGCTGAGGCATGAAAGCGTGGGG |
| P-asv5435 | GCAAGCGTTAATCGGAATTACTGGGCGTAAAGGGCGCGTAGGTGGTATCTTAAGTTGGGTGTGAAATCCCCGGGCTCAACCTGGGAATTGCATCCAAAACTGGGGTGCTAGAGTGTGGTAGAGGGTAGCGGAATTTCTGGTGTAGCGGTGAAATGCGTAGATACCAGAAGGAACATCAATGGCGAAGGCAGCTACCTGGGCCAACACTGACGCTGAGGCGCGAAAGCGTGGGG |
| P-asv5436 | TCTCTCTTTCCTCTTCTTTCCTTTTCTTAAAGCGTGCGTAGGCGGCTTTTTAAGTCGGATGTGAAATCCCTGAGCTTAACTTAGGAGTTGCATTCGATACTGGGAAGCTAGAGTATGGGAGAGGATGGTAGAATTCCAGGTGTAGCGGTGAAATGCGTAGAGATCTGGAGGAATACCGATGGCGAAGGCAGCCATCTGGCCTAATACTGACGCTGAGGTACGAAAGCATGGGG |
| P-asv5437 | GCGAGCGTTAATCGGAATAATTGGGCGTAAAGGGTGCGTAGGCGGATTCTTAAGTCAGATGTGAAAGCCCCGGGCTCAACCTGGGAATGGCATTTGATACTGGGAGTCTAGAGTAGAGGAGAGGAGAGTGGAATTTCAGGTGTAGCGGTGAAATGCGTAGATATCTGAAAGAACACCAGTGGCGAAGGCGGCTCTCTGGCCTCATACTGACGCTGAGGCACGAAAGCGTGGGG |
| P-asv5438 | GCTAGCGTTGTTCGGAATCACTGGGCGTAAAGCGCACGTAGGCGGACCTTTAAGTCAGGGGTGAAAGCCTGGAGCTCAACTCCAGAACTGCCTTTGATACTGGAGGTCTCGAGTGTGGAAGAGGTTGGTGGAACTGCGAGTGTAGAGGTGAAATTCGTAGATATTCGCAAGAACACCAGTGGCGAAGGCGGCCAACTGGTCCATTACTGACGCTGAGGTGCGAAAGCGTGGGG |
| P-asv5439 | GCGAGCGTTGTCCGGAATTATTGGGCGTAAAGGGCTCGTAGGCGGTTTGTCGCGTCGGAAGTGAAAACTCAGGGCTTAACCCTGTTCGTGCTTCCGATACGGGCAGACTAGAGGTAGGCAGGGGAGAACGGAATTCCTGGTGTAGCGGTGGAATGCGCAGATATCAGGAGGAACACCGGTGGCGAAGGCGGTTCTCTGGGCCTATTCTGACGCTGAGGAGCGAAAGCATGGGG |
| P-asv5440 | CCAAGCGTTAATCGGAATTACTGGGCGTAAAGGGAGCGTAGGCGGACTGTTTAGTCAGAGGTGAAAGCCCAGGGCTCAACCTTGGAATTGCCTTTGATACTGAAGATCTTGAGTTCGGGAGAGGTGAGTGGAACTGCGAGTGTAGAGGTGAAATTCGTAGATATTCGCAAGAACACCAGTGGCGAAGGCGGCTCACTGGCCCGATACTGACGCTGAGGCACGAAAGCGTGGGG |
| P-asv5441 | CCAAGCGTTGTTCGGAATTATTGGGCGTAAAGCGGGTGTAGGTGGCTCTATAAGTCAGGAGTGAAATCCCCGAGCTTAACTTGGGAAGTGCTTTTGATACTGCAGAGCTTGAGTGTGGGAGAGGATCGTAGAATTCCAGGTGTAGTGGTGAAATACGTAGATATCTGGAGGAATACCGGTGGCGAAGGCGGCGATCTGGCCCAACACTGACACTGAGACCCGAAAGCGTGGGG |
| P-asv5442 | GCAAGCGTTGTTCGGAATTATTGGGCGTAAAGGGCGTGTAGGTGGCTTACCAAGTCTGGTGTGAAATCCCACGGCTCAACCGTGGAACTGCATTGGAAACTGGTGAGCTCGAGTATTGGAGGGGGTCGCGGAATTCCCGGTGTAGAGGTGAAATTCGTAGATATCGGGAGGAACACCAGTGGCGAAGGCGGCGACCTGGCCATTTACTGACACTGAGACGCGAAAGCGTGGGG |
| P-asv5443 | GCAAGCGTTGTCCGAAATTATTGGGCGTAAAGAGCTCGTAGGCGGTTTGTCGCGTCTGCTGTGAAAGCCCGGGGCTCAACCCCGGGTCGGCAGTGGGTACGGGCAGACTGGAGTGCAGTAGGGGAGACTGGAATTCCTGGTGTAGCGGTGAAATGCGCAGATATCAGGAGGAACACCGATGGCGAAGGCAGGTCTCTGGGCTGTTACTGACGCTGAGGAGCGAAAGCATGGGG |
| P-asv5444 | GCAAGCGTTGTTCGGAATTACTGGGCGTAAAGCGTACGTAGGCGGTGAGTTAAGTCGAATGTTAAAGCCCGAGGCTCAACCTCGTGGATGCATTCGATACTGGCTCGCTAGAGGGTGATAGAGGAAAGTGGAATTCCGAGTGTAGCGGTGAAATGTGTAGATATTCGGAGGAACATCAGTAGCGAAGGCGACTTTCTGGGTCACACCTGACGCTGAGGTACGAAAGCGTGGGT |
| P-asv5445 | GCGAGCGTTATCCGGAATCACTGGGTTTAAAGGGTGCGCAGGCGGCGTGCTAAGTCAGGGGTGAAAGGCCACCGCTCAACGGTGGGACTGCCTCTGATACTGGCATGCTCGAACGGGCGCGAGGCGGGCGGAATGTGGCATGTAGCGGTGAAATGCTTAGATATGCCATAGAACACCGATTGCGAAGGCAGCCCGCCAGAGCCTAGTTGACGCTCATGCACGAAAGCGTGGGG |
| P-asv5446 | GCAAGCGTTATCCGGAATTATTGGGCGTAAAGCGCGCGTAGGCGGTAAGATAAGTCTGATGTGAAAGCCCCCGGCTCAACCGGGGAGGGTCATTGGAAACTGTTTTACTTGAGTACAGAAGAGGAGAGTGGAATTCCATGTGTAGCGGTGAAATGCGCAGAGATATGGAGGAACACCAGTGGCGAAGGCGGCTCTCTGGTCTGTAACTGACGCTGAGGTGCGAAAGCGTGGGG |
| P-asv5447 | GCAAGCGTTGTTCGGAATTACTGGGCGTAAAGGGCGCGTAGGCGGCTCATCAAGTCAGGCGTGAAATTCCCGGGCTCAACCTGGGGGCTGCGCTTGATACTGATGGGCTTGAATGCGGGAGAGGATAGTGGAATTCCCAGTGTAGAGGTGAAATTCGTAGATATTGGGAAGAACACCGGTGGCGAAGGCGGCTATCTGGCCCGTGATTGACGCTGAGGCGCGAAAGCGTGGGG |
| P-asv5448 | GCGAGCGTTGTCCGGAATTACTGGGCGTAAAGAGCGCGTAGGCGGCGTCGTACGACCGGTGTGAAAGCCCCCGGCTCAACTGGGGAGGGACACCGGTAACGGCGATGCTACGAGTGCAGGAGAGGGACGTGGAACTCCGGGAGTAGCGGTGAAATGCGTAGAGACCCGGAGGAACACCAGTGGCGAAGGCGGCGTCCTGGCCTGCGACTGACGCTGTGGCGCGAAAGCTAGGGG |
| P-asv5449 | GCAAGCGTTATCCGGATTTATTGGGTTTAAAGGGTTCGTAGGCGGAAAAATAAGTCAGTGGTGAAAGCCTGCAGCTCAACTGTAGAACTGCCATTGAAACTATTTTTCTTGAATATATATGAAGTAGGCGGAATGCGTCATGTAGCGGTGAAATGCACAGATATGACGCAGAACACCGATAGCGAAGGCAGCTTACTAATGTATTATTGACGCTGATGAACGAAAGCGTGGGT |
| P-asv5450 | GCTAGCGCTGTTCGGAATTACTGGGCGTAAAGCGCGTGTAGGCGGTTGCCAAAGTCGGGTGTGAAAGCCTTGAGCTCAACTCAAGAAATGCACTCGATACTTGGTGACTAGAGGACCGGAGAGGATAGTGGAATTCCCAGTGTAGTGGTGAAATACGTAGAGATTGGGAAGAACACCAGTGGCGAAGGCGGCTATCTGGACGGTTTCTGACGCTAAGACGCGAAAGCGTGGGG |
| P-asv5451 | GCAAGCGTTATCCGGATTTATTGGGTTTAAAGGGTCCGTAGGCGGATTAGTAAGTCAGTGGTGAAAGCCCGCAGCTCAACTGTGGAACTGCCATTGATACTGCTAGTCTTGAGTGTAGTTGAAGTAGCTGGAATGAGTAGTGTAGCGGTGAAATGCATAGATATTACTCAGAACACCAATTGCGAAGGCAGGTTACTAAGTTACAACTGACGCTGATGGACGAAAGCGTGGGG |
| P-asv5452 | GCGAGCGTTGTCCGGAATTACTGGGCGTAAAGAGCTCGTAGGTGGTTTGTCGCGTCGTCTGTGAAAGTCCAGCGCTTAACGCTGGGATTGCAGGCGATACGGGCTTACTTGAGTGCTGTAGGGGAGACTGGAATTCCTGGTGTAGCGGTGAAATGCGCAGATATCAGGAGGAACACCAATGGCGAAGGCAGGTCTCTGGGCAGTCACTGACACTGAGGAGCGAAAGCATGGGT |
| P-asv5453 | GCAAGCGTTACTCGGAATTACTGGGCGTAAAGCGTGCGTAGGTGGTTCGTTAAGTCTGCCGTGAAAGCCCCGGGCTCAACCTGGGAATGGCGGTGGATACTGGCGGACTAGAGTGCGGTAGAGGGTGGTGGAATTCCCGGTGTAGCAGTGAAATGCGTAGAGATCGGGAGGAACATCTGTGGCGAAGGCGGCCACCTGGACCAGCACTGACACTGAGGCACGAAAGCGTGGGG |
| P-asv5454 | TCAAGCGTTATCCGGATTTACTGGGTGTAAAGGGTCCGCAGGCGGATTAGCGGGTCAGAAGTTAAAACTTCGAGCTCAACTCGAAGACTGCTTTTGAAACTACTAATCTAGAGTTCGGAAGAGGCAATCGGAATTGCTGGTGTAGTCGTAATAAGCGTTGATATCAGCAAGAACACCAAATGCGAAGGCAGATTGCTAGTACGTAACTGACGCTCAGGGACGAAAGCGTGGGG |
| P-asv5455 | GCAAGCGTTAATCGGAATTACTGGGCGTAAAGGGCGCGTAGGCGGCCCTGTCAGTCGGGTGTGAAAGCCCGGGGCTCAACCCCGGAACGGCACCCGAGACGGCAGGGCTGGAGACCGGGAGAGGAGGGTGGAATAGCCAGTGTAGAGGTGAAATTCGTAGATATTGGCTGGAACACCGGTGGCGAAAGCGGCTCTCTGGCCCGGTTCTGACGCTGAGGCGCGAAAGCGTGGGG |
| P-asv5456 | CCGAGCGTTAATCGGAATTACTGGGCGTAAAGCGTGCGCAGGCGGTTTGTTAAGCAAGATGTGATAGCCCCGGGCTCAACCTGGGAATTGCATTTTGAACTGGCAAACTAGAGTCTTGTAGAGGGGGGTAGAATTCCAGGTGTAGCGGTGAAATGCGTAGAGATCTGGAGGAATACCGGTGGCGAAGGCGGCCCCCTGGACAAAGACTGACGCTCAGGCACGAAAGCGTGGGG |
| P-asv5457 | GCAAGCGTTATTCGGAATTACTGGGCGTAAAGGGTGCGTAGACGGTTCATTAAGTCTTTTGTTTAAGACCCCGGCCTAACTGGGGACCAGCAGGAGATACTGATAGACTAGAGGATGGAAGAGAGAAGCGGAATTCTCGGAGTAGCGGTTAAATGCGTAGATCTCGAGAGGAACACCGATGGCGAAAGCAGCTTCTTGGTCCATATCTGACGTTGAGGCACGAAAGCGTGGGG |
| P-asv5458 | GCAAGCGTTACTCGGATTTACTGGGCGTAAAGCGCGCGCAGGCGGATCGGCAAGTCAGGTGTGAAATCTCGGGGCTCAACTCCGAAACTGCGCCTGAAACTATCGATCTTGAGTCTTGGCGGGGGGGGTGGAATATCGCATGTAGGGGTAAAATCCGTAGAGATGCGATGGAACACCAGAGGCGAAGGCGACCTCCTGGACAAGTACTGACGCTGAGGCGCGAAAGCGTGGGG |
| P-asv5459 | GCAAGCGTTATCCAGATTCACTGGGTTTAAAGGGAGCGTAGGCGGAATGTTAAGTCCGTGGTGAAAGCCCTCAGCTCAACTGAGGAACTGCCATGGATACTATCATTCTTGAATGTCGTTGAGGTTTGCGGAATGGGTCATGTAGCTGTGAAATGCTTAGATATGACCCGGAACACCAATTGCGAAGGCAGCAGGCTAAACGAATATTGACGCTGAGGCTCGAAAGCGTGGGG |
| P-asv5460 | GCAAGCGTTAATCGGAATTACTGGACGTAAAGCGAGCGTAGGTGGCTTGATAAGTCAGATGTGAAATCCCCGGGCTCAACCTGGGAATGGCGGTAGAAACCGGCGCACTAGAGTATGGGAGAGGGTGGTGGAATTCCAGGTGTAGCGGTGAAATGCGTAGAGATCTGGAGGAACATCAGTGGCGAAGGCGGCCACCTGGCCCAATACTGACGCTGAGGCACGAAAGCGTGGGG |
| P-asv5461 | GCAAGCGTTGCCCGGAATTATTGGGCGTAAAGAGCTCGTAGGCGGTTTGTCGCGTCTGCTGTGAAAACGCAAGGCTTAACCTTGCGCCTGCAGTGGGTACGGGCAGACTAGAGTGCGGTAGGGGAGACTGGAATTCCTGGTGTAGCGGTGGAATGCGCAGATATCAGGGGGAACACCGATGGCGAAGGCAGGTCTCTGGGCCGTTACTGACGCTGAGGAGCGAAAGCATGGGG |
| P-asv5462 | GCAAGCGTTAATCGGAATTACTGGGCGTAAAGCGCACGTAGGCGGTTGGTTAAGTCAGATGTGAAAGCCCCGGGCTCAACCTGGGAATTGCATTTGAAACTGGCCAACTAGAGTATGTGAGAGGGGGGTAGAATTCCAAGTGTAGCGGTGAAATGCGTAGAGATTTGGAGGAATACCAGTGGCGAAGGCGGCCCCCTGGCACAATACTGACGCTCAGGTGCGAAAGCGTGGGG |
| P-asv5463 | GCAAGCGTTGTCCGGAATTATTGGGCGTAAAGCGCGCGCAGGCGGTCTCTTAAGTCTGATGTGAAAGCCCACGGCTCAACCGTGGAGGGTCATTGGAAACTGGGAGACTTGAGTGCAGAAGAGAAGAGTGGAATTCCATGTGTAGCGGTGAAATGCGTAGAGATATGGAGGAACACCAGTGGCGAAGGCGACTCTTTGGTCTGTAACTGACGCTGAGGCGCGAAAGCGTGGGG |
| P-asv5464 | GCGAGCGTTAATCGGAATTACTGGGCGTAAAGCGCACGCAGGCGGTTTGTTAAGCTAGATGTGAAAGCCCCGGGCTCAACCTGGGACGGTCATTTAGAACTGGCAGACTAGAGTCTTGGAGAGGGGAGTGGAATTCCAGGTGTAGCGGTGAAATGCGTAGATATCTGGAGGAACATCAGTGGCGAAGGCGACTCCCTGGCCAAAGACTGACGCTCATGTGCGAAAGTGTGGGT |

| | |
|---|---|
| P-asv5465 | GCAAACGTTGTTCGGAATTACTGGGCGTAAAGCGCGTGTAGGCTGCTCCGCAAGTCGGATGTGAAAGCCCTGGGCTCAACCTAGGAAGTGCATTCGATACTGCAGAGCATGAGTCCTGGAGAGGAAGGCGGAATTCTCGGTGTAGAGGTGAAATTCGTAGATATCGAGAGGAACACCGGTGGCGAAGGCGGCCTTCTGGACAGTGACTGACGCTGAGACGCGAAAGCGTGGGG |
| P-asv5466 | CCAAGCGTTGTTCGGATTCATTGGGCGTAAAGGGCGCGTAGGTGGCGCGAGTAGTCAACGGTGAAAGCCCGGGGCTCAACCCCGGGTCGGCCGTCGATACTCTCGTGCTTGAGCACTGTAGAGGCAGGTGGAATTCCGGGTGTAGCGGTGGAATGCGTAGAGATCCGGAAGGACACCGGTGGCGAAGGCGGCCAGCTGGGCAGTAGCTGACACTGAGGCGCGACAGCGTGGGG |
| P-asv5467 | GCGAGCGTTAATCGGAATTACTGGGCGTAAAGCGCACGTAGGTGGCGACGTAAGTCGGGTGTGAAATCCCTGGGCTCAACCTGGGGACGCCACTCGATACTGCTGTGGCCAGAGTCCAGTAGGGGAGTGTGGAATTCCCGGTGTAGCGGTGAAATGCGCAGATATCGGGAGGAACACCAGCGGCGAAGGCGGCACTCTGGGCTGGAACTGACGCTGAGGAGCGAAAGCGTGGGT |
| P-asv5468 | GCAAGCGTTGTCCGGATTTATTGGGCGTAAAGAGCGTGTAGGCGGCCAGATAGGTCCGTTGTGAAAACTGGAGGCTTAACCTCCAGACGTCGATGGAAACCATCTGGCTAGAGTCCGGAAGAGGAGAATGGAATTCCTGGTGTAGCGGTGAAATGCGCAGATATCAGGAAGAACACCCGTGGCGAAGGCGGTTCTCTGGGACGGTACTGACGCTGAGACGCGAAAGCGTGGGG |
| P-asv5469 | GCAAGCGTTGTCCGGAATTACTGGGCGTACAGCGCGCGCAGGCGGTCCACTACGTGCCGTGTGACATCCGTCGGCTTAACCGAGGGAGGCCAGGGTACACGGGTGAACTTGAGGCAGAGAGAGGGATGTGGAATACCCGGTGGAGCGGTGAAATGCGTAGAGATCGGGTGGAACACCAGGGGCGAAGGCGGCATCCTGGCTCTGTACTGACGCTGAGGCGCGAGAGCGTGGGG |
| P-asv5470 | GCTAGCGTTGTTCGGAATTACTGGGCGTAAAGGGCGCGTAGGCGGCCTTGTTAGTCGAGTGTGAAAGCCCTGGGCTTAACCCGGGAAGCGCGCTTGATACTGCAGGGCTTGAGTGTGGGAGAGGATGATGGAATTCCCAGTGTAGAGGTGAAATTCGTAGATATTGGGAAGAACACCGGTGGCGAAGGCGATCATCTGGACCACAACTGACGCTGAGGCGCGAAAGCGTGGGG |
| P-asv5471 | GCAAACGTTGTTCGGAATTACTGGGCGTAAAGCGCGTGTAGGCTGCTTCGAAAGTCGGGTGTGAAAGCCCAGGGCTCAGCCCTGGAAGTGCGCCGGAAACTGGGAGGCTCGAGTGCCGGAGGGGAGAGTGGAATTCCCGGTGTAGCGGTGAAATGCGTAGATATCGGGAGGAATACCGGTGGCGAAAGCGACCCTCTGGACGGTAACTGACGCTGAGGCACGAAAGCGTGGGG |
| P-asv5472 | GCAAGCGTTGTCCGGAATCATTGGGCGTAAAGAGCGTGTAGGCGGCTTGGTAAGTCGGCTGTGAAAGTCGGGGGCTCAACCCTCGAATGCCGGCCGATACTGTCAAGCTAGAGTCCGGAAGAGGCGAGTGGAATTCCTGGTGTAGCGGTGAAATGCGCAGATATCAGGAGGAACACCGATGGCGAAGGCAGCTCGCTGGGACGGTACTGACGCTGAGGCGCGAAAGCCAGGGG |
| P-asv5473 | GCGAGCGTTAATCGGAATTACTGGGTGTAAAGGGCGCGTAGGCGGTAAGATGTGTGTGTTGTGAAAGCCCCGGGCTTAACCTGGGAAGTGCAATGCAAACTATCTAACTAGAGTATATGAGAGGGTGGCGGAATTTCCGGTGTAGCGGTGAAATGCGTAGAGATCGGAAGGAACGTCAATGGCGAAGGCAGCCACCTGGCATAATACTGACGCTGAGGCGCGAAAGCGTGGGG |
| P-asv5474 | GCAAGCGTTGTCCGGATTCATTGGGCGTAAAGAGCGTGTAGGCGGCCAGGTAGGTCGGTTGTGAAAACTGGAGGCTCAACCTTCAGAGGTCGACCGAAACCATCTGGCTAGAGTCCGGGAGAGGAGAGTGGAATTCCTGGTGTAGCGGTGAAATGCGCAGATATCAGGAAGAACACCCGTGGCGAAAGCGGCTCTCTGGAACGGTACTGACGCTGAGACGCGAAAGCGTGGGG |
| P-asv5475 | GCAAGCGTTATTCGGAATTACTGGGCGTAAAGCGCGTGCAGGCGGGCCGGCAAGTCTGATGTGAAAGCCCCGGGCTTAACCTGGGAAGTGCATTGGAAACTGCTGGTCTTGAGTTCTGGAGAGGAAGGGGAATTCCCGGTGTAGAGGTGAAATTCGTAGAGATCGGGAGGAATACCAGTGGCGAAGGCGCCCTTCTGGACAGTAACTGACGCTGAGACGCGAAAGCGTGGGG |
| P-asv5476 | GCTAGCGTTGTCCGGAATCATTGGGCGTAAAGCGCGTGTAGGCGGCCCGGTAAGTCCGCTGTGAAAGTCAAGGGCTCAACCCTTGAAAGCCGGTGGATACTGTCGGACTAGAGTGCGGGAGAGGCGAGTGGAATTCCTGGTGTAGCGGTGAAATGCGCAGATATCAGGAGGAACACCAATTGCGAAGGCAGCTCGCTGGAACGTTACTGACGCTGAGACGCGAAAGCGTGGGG |
| P-asv5477 | GCGAACGTTGTTCGGATTCACTGGGCTTAAAGGGCGCGTAGGCGGCTGCTCAAGTCGGGGGTGAAATCTTTCCGCTCAACGGGAAAAGGGCCTCCGATACTGGGCGGCTGGAGGGAGGTAGGGGCATGTGGAACTTCCGGTGGAGCGGTGAAATGCGTAGATATCGGAAGGAACGCCGGTGGCGAAAGCGACGTGCTGGACCTCCACTGACGCTGAGGCGCGAAAGCCAGGGG |
| P-asv5478 | GCTAGCATTGCTCGGGATCACTGGGCGTAAAGCGCACGTAGGCGGCTTTTTAAGTCAGAGGTGAAATCCTGGAGCTCAACTCCAGAACTGCCTTTGATACTGAGAGGCTCGAGTCCGGGAGAGGTGAGTGGAACTGCGAGTGTAGAGGTGAAATTCGTAGATATTCGCAAGAACACCAGTGGCGAAGGCGGCTCACTGGCCCGGTACTGACGCTGAGGTGCGAAAGCGTGGGG |
| P-asv5479 | GCTAGCGTTGTTCGGAATTACTGGGCGTAAAGCGTGCGCAGGCGGCTATCCAAGTCAGTGGTGAAAGCCCGGAGCTCAACTCCGGAATTGCCATTGAAACTGTTTAGCTTGAGGACGAGAGAGGTGAGTGGAATTCCCAGTGTAGAGGTGAAATTCGTAGATATTGGGAAGAACACCGGTGGCGAAGGTGGCTCACTGGCTCGTTTCTGACGCTCATGCACGACAGCGTGGGG |
| P-asv5480 | GCGAGCGTTGTTCGGATTTACTGGGCGTAAAGGGCGCGCAGGCGGCCTTGTTTGTCAGGCGTGAAAGCCCCGGGCTTAACCTGGGAACGGCGCTTGGGACGGCGAGGCTAGAGATCGGGAGAGGAGAGTGGAATTCCCAGTGTAGAGGTGAAATTCGTAGATATTGGGAAGAACACCGGTGGCGAAGGCGGCTCTCTGGACCGATACTGACGCTGAGGCGCGAAAGCGTGGGG |
| P-asv5481 | CCAAGCGTTGTCCGGATTTATTGGGCGTAAAGAGCTCGTAGGCGGCTCGGTAAGTCGGGTGTGAAACCTCCAGGCTCAACCTGGAGACGCCACCCGATACTGCTGTGGCTAGAGTCCGGTAGGGGAGCGTGGAATTCCTGGTGTAGCGGTGAAATGCGCAGATATCAGGAGGAACACCAGCGGCGAAGGCGGCGCTCTGGGCCGGTACTGACGCTGAGGAGCGAAAGCGTGGGG |
| P-asv5482 | GCTAGCGTTGTTCGGAATTACTGGGCGTAAAGGGCGCGTAGGCGGCCTTGTGCGTCGTGCGTGAAAGGCCTGGGCTCAACCCAGGAATGGCGCTCGATACGGCATGGCTAGAGTGCAGGAGAGGATGGTGGAATTCCCAGTGTAGAGGTGAAATTCGTAGATATTGGGAAGAACACCGGTGGCGAAGGCGGCCATCTGGACTGCAACTGACGCTGAGGCGCGAAAGCGTGGGG |
| P-asv5483 | GCAAGCGTTAATCGGAATTACTGGGCGTAAAGCGCACGCAGGCGGTCTGTCGAGTCGGATGTGAAATCCCCGGGCTCAACCTGGGAACTGCATTCGAGACTGGCAGGCTTGAGTCTTGTAGAGGGGGGTAGAATTCCAGGTGTAGCGGTGAAATGCGTAGAGATCTGGAGGGATACCGGTGGCGAAGGCGGCCCCCTGGACAAAGACTGACGCTCAGGTGCGAAAGCGTGGGG |
| P-asv5484 | GCGAACGTTATTCGGAATCACTGGGCTTAAAGGGCGCGTAGGCGGGCGACCAAGTCCGGGGTGAAAGCCCACGGCCCAACCGTGGAAGGGCCCCGGATACTGGTCGCCTCGAGGGAGGTAGGGGCGTGTGGAACTTCCGGTGGAGCGGTGAAATGCGTAGAGATCGGAAGGAACGCCGGTGGCGAAAGCGACACGCTGGACCTCTGCTGACGCTGAGGCGCGAAAGCCAGGGG |
| P-asv5485 | GCTAGCGTTGCTCGGAATCACTGGGCGTAAAGGGCGCGTAGGCGGCGCCTCTAGTCGGGCGTGAAATCCCCGGGCTCAACCCGGGGGCTGCGCCCGATACGGGGGCGCTGGAGGACGGGAGAGGCTCGCGGAATTCCCAGTGTAGAGGTGAAATTCGTAGATATTGGGAAGAACGCCGGTGGCGAAGGCGGCGAGCTGGCCCGTTTCTGACGCTGAGGCGCGACAGCGTGGGG |
| P-asv5486 | GCGAGCGTTAATCGGAATTACCGGGCGTAAAGCGCGCGTAGGTGGTTAATTAAGTCAGATGTGAAATCCCAGGGCTCAACCTTGGAACTGCATTTGAAACTGGTTAACTGGAGTTTTGTAGAGGGTGGTAGAATTTCAGGTGTAGCGGTGAAATGCGTAGAGATCTGAAGGAATACCAGTGGCGAAGGCGGCCACCTGGACAAAGACTGACACTGAGGCGCGAAGGCGTGGGG |
| P-asv5487 | GCGAGCGTTATCCGGACTTACTGGGCGTAAAGGGTGTGCAGGCGGGGGTTCAAGTGGTGTATGAAAGGTCCCAGCTAAACTGGGATAGGTTATGCCAGACTGGACTTCTAGAGTGCGAGAGAGGGGCGTGGAATTCCGGGTGTAGTGGTGAAATGCGTAGAGATCCGGAGGAACCCCAGAGGCGAAGGCGGCGCCCTGGCTCGCAACTGACGCTCAGACACGACAGCATGGGG |
| P-asv5488 | GCCAGCGTTGTTCGGAATTACTGGGCGTAAAGCGTGCGCAGGCGGTTCGTTAAGTTCGATGTGAAAGCCCCGGGCTCAACCTGGGAATTGCATTGGAAACTGGCGGGCTTGAGTGTGGCAGAGGGGGGTGGAATTCCACGTGTAGCAGTGAAATGCGTAGAGATGTGGAGGAACACCGATGGCGAAGGCAGCCCCCTGGGCTAACACTGACGCTCAGGCACGAAAGCGTGGGG |
| P-asv5489 | GCAAACGTTATTCGGAATCACTGGGCGTAAAGGGTGCGTAGACGGTATGTTAAGTTCGTTGTTAAAGGCTCCAGCCTAACTGGAGTACCGCAATGAAAACTGGCAAGCTAGAGGGTGAGAGAGAGAAGTGGAATTCCCGGAGTAGCGGTAAAATGCGTGGATCTCGGGAGGAACACCAATGGCGAAGGCAGCTTCTTGGCTCATACCTGACGTTGAGGCACGAAAGCGTGGGG |
| P-asv5490 | GCGAGCGTTGTTCGGAATTATTGGGCGTAAAGGGTGCGTAGGCGGTTCGGTAAGTCTTGTGTGAAATCTTCGGGCTCAACTCGAAGTCTGCACGAGAAACTGCCGGGCTTGAGTGTGGGAGAGGTGAGTGGAATTTCCGGTGTAGCGGTGAAATGCGTAGATATCGGAAGGAACACCTGTGGCGAAAGCGGCTCACTGGACCACAACTGACGCTGATGCACGAAAGCTAGGGG |
| P-asv5491 | GCTAGCGTTGCTCGGAATCACTGGGCGTAAAGGGCGCGTAGGCGGCTCTGTGAGTCAGGCGTGAAATTCCTGGGCTTAACCTGGGGACTGCGCTTGATACTGCGGGGCTTGAGGCCGAGAGAGGGTGGTGGAATTCCCAGTGTAGAGGTGAAATTCGTAGATATTGGGAAGAACACCGGTGGCGAAGGCGACCACCTGGCTCGGTACTGACGCTGAGGCGCGACAGCGTGGGG |
| P-asv5492 | GCGAGCGTTAATCGGAATTACTGGGCGTAAAGCGTGCGCAGGCGGTTCGTCGTGTCCGATGTGAAAGCCCCGGGCTCAACCTGGGAATGGCATTGGAAACTGGCGGGCTTGAGTGTGGCAGAGGGGGGTGGAATTCCGCGTGTAGCAGTGAAATGCGTAGAGATGCGGAGGAACACCGATGGCGAAGGCAGCCCCCTGGGCCAACACTGACGCTCAGGCACGAAAGCGTGGGG |
| P-asv5493 | GCGAACGTTGTTCGGAATCACTGGGCGTAAAGGGCGTGTAGGCGGCTGAATAAGTGGAGTGTTAAATCCCCCGGCTCAACCGGGGAACTGCACTTCATACTGTTCAGCTTGGGTACAGCAGAGGAGACTGGAATTCCCGGTGTAGCGGTGAAACGCGTAGATATCGGGAGGAACACCAGTGGCGAAGGCGAGTCTCTGGGCTGATACCGACGCTGAGGCGCGAAAGCCAGGGG |
| P-asv5494 | GCGAGCGTTGTTCGGAATTATTGGGCGTAAAGGGCGCGTAGGCGGTGCCACAAGTCACTTGTGAAATCTCCGGGCTTAACTCGGAGCCTGCAGGCGAAACTGTGGTGCTGGAGTGTGGGAGAGGTGCGTGGAATTCCCGGTGTAGCGGTGAAATGCGTAGGTATCGGGAGGAACACCCGTGGCGAAGGCGGCTCTCTGGGACGGTACTGACGCTGAGACGCGAAAGCGTGGGG |
| P-asv5495 | GCTAGCGTTGTTCGGAATCACTGGGCGTAAAGCGCACGTAGGCGGATATGTCAGTCAGGGGTGAAATCCCGGAGCTCAACCTCGGAAATGCCTTCGATACTGGCTGTCTTGAGTACGGGAGAGGTGAGTGGAACTCCGAGTGTAGAGGTGAAATTCGTAGATATTCGGAAGAACACCAGTGGCGAAGGCGACTCACTGGCCCGTTACTGACGCTGAGGCGCGAAAGCGTGGGG |
| P-asv5496 | GCAAGCGTTGTCCGGAATTATTGGTCGTAAAGCGCGCGCAGGCGGTCCTTTAAGTCAGGGGTGAAATCCTGGAGCTCAACTCCAGAACTGCCTTTGATACTGAGGATCTTGAGTTCGGGAGAGGTAAGTGGAACTGCGAGTGTAGAGGTGAAATTCGTAGATATTCGCAAGAACACCAGTGGCGAAGGCGGCTTACTGGCCCGATACTGACACTGAGGCACGAAAGCGTGGGG |
| P-asv5497 | GCGAGCGTTGTTCGGAATTATTGGGCGTAAAGCGCGCGCAGGCGGCGTGTTAAGTCGACGGGTGAAATGCCGGGGCTCAACCCCGTGCCCGCCCGCGATACTGGCAGGCTGGAGACAGGTAGAGGCGAGCGGAATTCCGGGTGTAGCGGTGGAATGCGTAGAGATCCGGAAGAACACCAGAGGCGAAGGCGGCTCGCTGGGCCTGGTCTGACGCTGAGGCGCGAAAGCGTGGGG |
| P-asv5498 | GCAAGCGTTGTTCGGAATTACTGGGCGTAAAGCGCGGGCAGGCGGTCTGTTAAGTCGTTTGTGAAACTCCGGGGCTCAACCCCGGACATGCAGACGATACTGGCGGACTAGAGAATGGTAGAGGCAAGCGGAATTCCTAGTGTAGCGGTGGAATGCGTAGATATCAGGAAGAACACCTGTGGCGAAGGCGGCTTGCTGGGCCATTTCTGACGCTCATCCGCGAAAGCGTGGGG |
| P-asv5499 | GCGAGCGTTGTCCGGAATTACTGGGCGTAAAGAGCTCGTAGGTGGTTTGTCGCGTCGTTTGTGTAAGTCCACAGCTTAACTGTGGGACTGCAGGCGATACGGGCATAACTTGAGTGCTGTAGGGGAGACTGGAATTCCTGGTGTAGCGGTGGAATGCGCAGATATCAGGAGGAACACCGATGGCGAAGGCAGGTCTCTGGGCAGTAACTGACGCTGAGGAGCGAAAGCATGGGT |
| P-asv5500 | GCGAGCGTTAATCGGAATTACTGGGCGTAAAGCGCGCGTAGGTGGTTAATTAAGTCAGATGTGAAATCCCAGGGCTCAACCTTGGAACTGCATTTGAAACTGGTTAACTAGAGTTTTGTAGAGGGTGGTAGAATTTCAGGTGTAGCGGTGAAATGCGTAGAGATCTGAAGGAATACCAGTGGCGAAGGCGGCCACCTGGACAAAGACTGACACTGAGGCGCGAAGGCGTGGGG |
| P-asv5501 | GCAAGCGTTATCCGGAATTATTGGGCGTAAAGAGCTTGTAGGCGGTTCGTCGCGTCTGGTGTGAAAGCCCATCGCTTGACGGTGGGTTTGCGCCGGGTACGGGCGGGCTAGAGTGCAGTAGGGGAGACTGGAATTCTCGGTGTAACGGTGGAATGTGTAGATATCGGGAAGAACACCAATGGCGAAGGCAGGTCTCTGGGCTGTTACTGACGCTGAGAAGCGAAAGCGTGGGG |
| P-asv5502 | GCAAGTGTTATCCGGAATTACTGGGCGTAAAGCGTCTGTAGGTTGTTAAACAAGTCAACTGTTAAATCTTGAAGCTCAACTTCAAAATCGCAGTCGAAACTGTTTGACTAGAGTATAGTAGAGGTAAAGGGAATTTCCAGTGGAGCGGTGAAATGCGTAGATATTGGAAAGAACACCGATGGCGAAAGCACTTTACTGGGCTATTACTGACACTCAGAGACGATAGCTAGGGT |
| P-asv5503 | GCAAGCGTTGTCCGGAATTATTGGGCGTAAAGAGCTCGTAGGCGGTCTGTCACGTCGGCTGTGAAAACTCGGGGCTCAACCCTGAGCCTGCAGTCGATACGGGCAGACTAGAGTTCGGCAGGGGAGACTGGAATTCTTGGTGTAGCGGTGAAATGCGCAGATATCAGGAGGAACACCGGTGGCGAAGGCGGGTCTCTGGGCCGATACTGACGCTGAGGAGCGAAAGCGTGGGG |

| | |
|---|---|
| P-asv5504 | TCAAGTAATGTCCAATATTATTGGCCTAAAGCGTCCGTAGCTGGCCGCGCAAGTCCGTCGGGAAATCCACCTGCCCAACAGGTGGGCGTCCGGCGGAAACTGTGTGGCTTGGAACCGGAAGGCTCAGAGGGTACGTCCGGGGTAGGAGTGAAATCCCGTAATCCCGGACGGACTACCGATGGCGAAAGCACTCTGAGAAGACGGCTTCGACAGTGAGGGACGAAAGCTAGGGT |
| P-asv5505 | TCTCTCTTTTTCTTCCTTCCTTTTCTTAAAGCTCACGTAGGCGGCTTTGTAAGTCAGAGGTGAAAGCCTGGAGCTCAACTCCAGAACTGCCTTTGAGACTGCATCGCTTGAATCCAGGAGAGGTGAGTGGAATTCCGAGTGTAGAGGTGAAATTCGTAGATATTCGGAAGAACACCAGTGGCGAAGGCGGCTCACTGGACTGGTATTGACGCTGAAGTGCGAAAGCGTGGGG |
| P-asv5506 | CCGAGCGTTATCCGGATTTATTGGGTTTAAAGGGAGCGTAGATGGATGTTTAAGTCAGTTGTGAAAGTTTGCGGCTCAACCGTAAAATTGCAGTTGATACTGGATGTCTTGAGTGCAGTTGAGGCAGGCGGAATTCGTGGTGTAGCGGTGAAATGCTTAGATATCACGAAGAACTCCGATTGCGAAGGCAGCCTGCTAAGCTGCAACTGACATTGAGGCTCGAAAGTGTGGGT |
| P-asv5507 | GCAAGCGTTGTCCGGATTTATTGGGCGTAAAGCGCGCGCAGGTGGTTTCTTAAGTCTGATGTGAAAGCCCCCGGCTCAACCGGGGAGGGTCATTGGAAACTGGGAGACTTGAGTGCAGAAGAGGATAGTGGAATTCCAAGTGTAGCGGTGAAATGCGTAGAGATTTGGAGGAACACCAGTGGCGAAGGCGACTGTCTGGTCTGTAACTGACACTGAGGCGCGAAAGCGTGGGG |
| P-asv5508 | GCGAGCGTTGTCCGGATTTATTAGGCGTAAAGAGTTCGTAGGTGGTTTGTTAAGTTTGGTGTTAAAGATCGGGGCTCAACCCTGGGACTGCACTGAATACTGGCAGACTCGAGTGTGGTAGAGGCTAGTGGAATTCCCAGTGTAGCGGTGAAATGCGTAGATATTGGGAAGAACACCGGTGGCGTAGGCGACTAGCTGGGCCATAACTGACGCTGAGGAACGAAAGCCAGGGG |
| P-asv5509 | CCAAGCGTTAATCGGAATTACTGGGCGTAAAGCGTGCGCAGGCGGTTTTGTAAGACAGAGGTGAAATCCCCGGGCTCAACCTGGGAATTGCCTTTGTGACTGCAAGGCTAGAGTGCGGCAGAGGGGGGTAGAATTCCACGTGTAGCAGTGAAATGCGTAGAGATGTGGAGGAATACCGATGGCGAAGGCAGCCCCCTGGGCCGATACTGACGCTCATGCACGAAAGCGTGGGG |
| P-asv5510 | GCTAGCGTTGTTCGGAATGACTGGGCGTAAAGGGCGCGTAGGTGGTTACGCAAGTCTTGTGTGAAAGCCCCGGGCTTAACCTGGGAACTGCACAGGAGACTGCGTGGCTAGAGTACAGAAGGGGAAACTGGAATTGCGAGTGGAGAGGTGAAATTCGTAGAGATTCGCAAGAACACCAATGGCGAAGGCATGTTTCTGGTCTGTTACTGACACTGAGGCGCGAAAGCGTGGGG |
| P-asv5511 | GCAAGCGTTGTCCAGAATTATTAGGCATAAAGCGCACACAGGCAGTCCTTTAAGTCTGATATGAAAGCCCACGGCTCAACCGTGGAGGGTCATTAGAAACTGGGGGACTTGAGTACAGAAGAGAAGAGTAGAATTCCACATGTAGCGGTAAAATGCATAGAGATGTAGAAAAACACCAGTGACGAAGGCAACTCTTTGGTCTATAACTGACACTGAGGCGCAAAAGCGTAGGA |
| P-asv5512 | GCGAGCGTTAATCGGAATTACTGGGCGTAAAGCGTGCGCAGGCGGCTTCGTAAGCTGGTGGTGAAATCCCCGGGCTTAACCTGGGAATGGCCTTCAGGACAGCGAGGCTAGAGTGCGGCAGAGGGAGGTGGAATTCCGCGTGTAGCAGTGAAATGCGTAGAGATGTGGAGGAACACCGATGGCGAAGGCAGCCTCCTGGGCCAGCACTGACGCTCATGCACGAAAGCGTGGGG |
| P-asv5513 | GCAAGCGTTGTCCGGAGTTATTGGGCGTGAAGAGCTCGTAGGCGGTTTGTCGCGACTGCTGTGAAAACGCAAGGCTTAACCTTGCGCCTGCAGTGGGTACGGGCAGACTAGAGTGCGGTAGGGGAGACTGGAATTCCTGGTGTAGCGGTGGAATGCGCAGATATCAGGAGGAACACCGATGGCGAAGGCAGGTCTCTGGGCCGTTACTGACGCTGAGGAGCGAAAGCATGGGG |
| P-asv5514 | GCAAGCGTTGTTCGGAATTACTGGGCGTAAAGCGTGCGTAGTCGGTATTGAGAGTCACGGGTGAAATCCCAGGGCTTAACCCTGGAACTGCCTGTGAGACCTCAGTACTAGAGTGTGAGAGGGGATAGTGGAATACCCAGTGTAGCGGTGAAATGCGTAGAGATTGGGCGGAACACCGGTGGCGAAGGCGGCTCACTGGACTGGTATTGACGCTGAAGTGCGAAAGCGTGGGG |
| P-asv5515 | GCGAGCGTTGTCCGGATTTATTGGGCGTAAAGAGTTCGTAGGCGGTTTGTTAAGTCTGATGTTAAAGATCGGGGCTCAACCCTGGGAGTGCATTGGATACTGGCAGACTGGAGTGTGGTAGAGGCTAGTGGAATTCCTAGTGTAGCGGTGAAATGCGTAGATATTAGGAAGAACACCAGTGGCGTAGGCGACTAGCTGGGCCATAACTGACGCTGAGGAACGAAAGCCAGGGG |
| P-asv5516 | GCAAGCGTTGTCCGGATTTATTGGGCGTAAAGCGAGCGCAGGCGGTTTCTTAAGTCTGATGTGAAAGCCTTCGGCTCAACCGAAGAAGTGCATCGGAAACTGGGGAACTTGAGTGCAGAAGAGGACAGTGGAACTCCATGTGTAGCGGTGAAATGCGTAGATATATGGAAGAACACCAGTGGCGAAGGCGGCTGTCTGGTCTGTAACTGACGCTGAGGCTCGAAAGCATGGGT |
| P-asv5517 | ACGAACGTTATTCGGAATTACTGGGCTTAAAGGGTGCGTAGGCTGCACGGAAAGTTGGGTGTGAAAGCCCTCAGCTCAACTGAGGAATTGCATCCAAAACTGCCGTGCTTGAGGGAGACAGAGGTGAGCGGAACTCAAGGTGGGGCGGTGAAATGCGTTGATATCTTGAGGAACACCGGTGGCGAAGGCGGCTCACTGGGTCTCTTCTGACGCTGAGGCACGAAAGCTAAGGT |
| P-asv5518 | GCAAGCGTTATCCGGAATCACTGGGCATAAAGCGTCTGTAGGTTGCCTACCAAGTCTGCTGTTAAAGATCAGGGCCTAACCCTGGGAAAGCAGTGGAAACTAGTAGGCTTGAGTGTGGTAGAGGTAGAGGGAATTCCTGGTGTAGCGGTGAAATGCGTAGATATTAGGAAGAACACCAATGGCGAAAGCACTCTACTGGGCCATAACTGACACTGAGAGACGACAGCTAGGGG |
| P-asv5519 | CCAAGCGTTGTCCGGAATCACTGGGTGTAAAGGGTGCGTAGGCGGATTCGTAAGTCAGAGGTGAAAGCCCACGGCTCAACCGTGGAACTGCCTTTGATACTGCGAGTCTTGAGTCCTGGAGGGGCTGGCGGAATTCGTGGTGTAGCGGTGAAATGCGTAGATATCACGAGGAACACCGGAGGTGTAGACGGCCAGCTGGACAGGTACTGACGCTGAGGCACGAAAGCGTGGGG |
| P-asv5520 | GCAAGCGTTATCCGGATTCACTGGGTTTAAAGGGTGCGTAGGCGGTCTGGTAAGTCAGATGTGAAAGCTCACAGCTTAACTGTGGAATTGCATTTGATACTGTCAGACTTGAATCAAGATGAGGTAGGCGGAATGTGTCATGTAGCGGTGAAATGCTTAGATATGACATAGAACACCGATTGCGAAGGCAGCTTACTAGACTTGTATTGACGCTGAGGCACGAAAGCGTGGGG |
| P-asv5521 | GCAAGCGTTGTCCGGAATCATTGGGCGTAAAGGGTGCGTAGGCGGCCATGTAAGTCAGATGTGAAAGACTGCGGCTTAACCGTAGGGTTGCATTTGAAACTGTATGGCTTGAGTGCTGGAGAGGGTAGTGGAATTCCCGGTGTAGCGGTGAAATGCGTAGATATCGGGAGGAACACCAGTGGCGAAGGCGACTACCTGGACAGTAACTGGCGCTGAGGCACGAAAGCTAGGGG |
| P-asv5522 | GCAAGCGTTGTCCGGATTTATGGGCGTAAAGAGCTCGTAGGCGGTCTGTCGCGTCTGCTGTGAAAACCCGAGGCTCAACCTCGGGCCTGCAGTGGGTACGGGCAGACTAGAGTGCGGTAGGGGAGACTGGAATTCCTGGTGTAGCGGTGGAATGCGCAGATATCAGGAGGAACACCGATGGCGAAGGCAGGTCTCTGGGCCGTTACTGACGCTGAGGAGCGAAAGCATGGGG |
| P-asv5523 | GCGAGCGTTGTCCGGAATTATTGGGCGTAAAGGGCTTGTAGGTGGCTGGTTGCGTCTGTCGTGAAAGCTCATGGCTTAACTGTGGGTTTGCGGTGGGTACGGGCTGGCTTGAGTGCAGTAGGGGAGGCTGGAATTCCTGGTGTAGCGGTGGAATGCGCAGATATCAGGAGGAATACCGGTGGCGAAGGCGGGTCTCTGGGCTGTTACTGACACTGAGGAGCGAAAGCATGGGG |
| P-asv5524 | GCTAGCGTTGTTCGGAATTACTGGGCGTAAAGCGCACGTAGGCGGATATTTAAGTCAGGGGTGAAATCCCAGAGCTCAACTCTGGAACTGCCTTTGATACTGGGTATCTCGAGTCCGAGAGAGGTGAGTGGAATTCCGAGTGTAGAGGTGAAATTCGTAGATATTCGGAGGAACACCAGTGGCGAAGGCGGCTCACTGGCTCGGTACTGACGCTGAGGTGCGAAAGCGTGGGG |
| P-asv5525 | GCAAGCGTTGTCCGGAGTTACTGGGCGTAAAGAGTGCGTAGGCGGTGCCCTAAGTCTGTTGTGAAATCTCCTGGCTCAACTGGGAGGGTGCGGTGGAAACTGGGGTGCTTGAGAGTGGGAGAGGTTAGCGGAATTCCTGGTGTAGCGGTGAAATGCGTAGATATCAGGAGGAACACCTGTGGTGTAGACGGCTGACTGGACCACGACTGACGCTGAGGCACGAAAGCGTGGGT |
| P-asv5526 | GCGAGCGTTAATCGGAATTACTGGGCGTAAAGCGTGCGTAGGCGGATTATTAAGTGAGATGTGAAAGCCCCGGGCTCAACCTGGGAACTGCATTTCAAACTGGTAGTCTAGAGTTTTGTAGAGGGTGGTGGAATTTCCAGTGTAGCGGTGAAATGCGTAGAGATTGGAAGGAACATCAGTGGCGAAGGCGGCCACCTGGACAAAAACTGACGCTGAGGCACGAAAGCGTGGGG |
| P-asv5527 | GCGAGCGTTAATCGGAATTACTGGGCGTAAAGCGCATGCAGGTGGATGATTAAGTCAGATGTGAAAGCCCGGGGCTCAACCTCGGAATAGCATTTGAAACTGGTCATCTAGAGTACTGTAGAGGGGGGTAGAATTTCAGGTGTAGCGGTGAAATGCGTAGAGATCTGAAGGAATACCAGTGGCGAAGGCGGCCCCCTGGACAGATACTGACACTCAGATGCGAAAGCGTGGGG |
| P-asv5528 | CCGGGCGTTATCCGGATTTATTGGGTTTAAAGGGAGCGTAGGCCGTGGATTAAGCGTGTTGTGAAATGTAGACGCTCAACGTCTGACTTGCAGCGCGAACTGGTCCACTTGAGTGTGCACAACGCAGGCGGAATTCGTCGTGTAGCGGTGAAATGCTTAGATATGACGAAGAACCCCGATTGCGAAGGCAGCTTGCGGGAGCACGACTGACGCTGAAGCTCGAAAGTGCGGGT |
| P-asv5529 | GCGAGCGTTAATCGGAATTACTGGGCGTAAAGCGCACGTAGGCGGCTTGGCATGTCGGGTGTGAAATCCCCGGGCTCAACCTGGGAACTGCATCCGAAACTGCCTTGCTAGAGTATGGGAGAGGGAAGTGGAATTTCCGGTGTAGCGGTGAAATGCGTAGATATCGGAAGGAACATCAGTGGCGAAGGCGACTTCCTGGACCAATACTGACGCTCAGGTGCGAAAGCGTGGGG |
| P-asv5530 | GCAAGCGTTGTCCGGATTTATTGGGTTTAAAGGGTGCGTAGGCGGCCTATTAAGTCAGGGGTGAAAGACTGCAGCTTAACTGTAGCAGTGCCTTTGATACTGATAGGCTTGAGTACAGATGAGGCCGGCGGAATGGGTGGTGTAGCGGTGAAATGCATAGATACCATCCAGAACACCGATTGCGAAGGCAGCCGGCTAAACTGTCACTGACGCTGAGGCACGAAAGCGTGGGG |
| P-asv5531 | GCGAGCGTTATCCGGATTTATTGGGTTTAAAGGGAGCGTAGGCGGTCTTATTAGTCAGTGGTGAAATCTCGTTGCTTAACAACGACCGTGCCATTGATACTGTAGGACTTGAGTACAGTTGCCGTTGGCGGAATGTGTCATGTAGCGGTGAAATGCATAGATATGACACAGAACACCGATTGCGAAGGCAGCTGACGAAACTGTAACTGACGCTGAGGCTCGAAGGCGTGGGG |
| P-asv5532 | GCGAGCGTTAATCGGAATTACTGGGCGTAAAGCGCGCGTAGGTGGTCTGTTAAGTCGGATGTGAAAGCCCCGGGCTCAACCTGGGAACTGCATTCGATACTGGCAGACTAGAGTGCGAGAGAGGGAGGTAGAATTCCATGTGTAGCGGTGAAATGCGTAGATATGTGGAGGAATACCAGTGGCGAAGGCGGCCTCCTGGCTCGACACTGACACTGAGGTGCGAAAGCGTGGGG |
| P-asv5533 | GCGAGCGTTATCCGGATTTATTGGGTTTAAAGGGTGCGTAGGTGGCTTGTTAAGTCAGCGGTGAAAGGTTCGAGCTCAACTTGGACATTGCCGTTGAAACTGGCGAGCTTGAGTGTAGATGAGGTAGGCGGAATGCATGGTGTAGCGGTGAAATGCTTAGATATCATGCAGAACTCCGATTGCGAAGGCAGCTTACTAAGGTACAACTGACACTGAAGCACGAAAGCGTGGGT |
| P-asv5534 | GCGAGCGTTGTCCGGAATTACTGGGCGTAAAGAGCTTGTAGGCGGTCTGTCACGTCGATTGTGAAAATTCACGGCTCAACCGTGGACCTGCAGTCGATACGGGCAGACTAGAGTACTTCAGGGGAGACTGGAATTCCTGGTGTAGCGGTGAAATGCGCAGATATCAGGAGGAACACCGGTGGCGAAGGCGGGTCTCTGGGAAGTAACTGACGCTGAGAAGCGAAAGCGTGGGT |
| P-asv5535 | GCAAGCGTTGTTCGGAATTACTGGGCGTAAAGGGCGCGTAGGCGGCCGATCAAGTCAGGCGTGAAATTCCCGGGCTCAACCCGGGGGCTGCGCTTGATACTGATTGGCTTGAATGCGGGAGAGGATAGTGGAATTCCCAGTGTAGAGGTGAAATTCGTAGATATTGGGAAGAACACCAGTGGCGAAGGCGGCTATCTGGCCCGTTATTGACGCTGAGGCGCGAAAGCGTGGGG |
| P-asv5536 | GCAAGCGTTGTCCGGATTTATTGGGTTTAAAGGGTGCGTAGGTGGTTTATTAAGTCAGTGGTGAAAGACGGTCGCTTAACGATTGCAGTGCCATTGATACTGATAGACTTGAGTTTTGTTGAGGTAGGCGGAATTGATAGTGTAGCGGTGAAATGCATAGATATTATCAAGAACTCCAATTGCGTAGGCAGCTTACTGAGCAAAAACTGACACTGAGGCACGAAAGTGTGGGT |
| P-asv5537 | GCGAGCGTTGTCCGGAATTACTGGGCGTAAAGAGCACGTAGGCGGCTATATAAGTCAGATGTAAAAAACCTGGGCTTAACCGAGGGGATGCAACTGAAACTATATAGCTTGAGTCAAGGAGAGGAGAGCGGAATTCCTGGTGTAGCGGTGAAATGCGTAGAGATCAGGAAGAATACCAGTGGCGAAAGCGGCTCTCTGGACTTGAACTGACGCTGAGGTGCGAAAGCGTGGGG |
| P-asv5538 | GCGAACGTTGTTCGGAATCACTGGGCTTAAAGGGCGCGTAGGCGGGTCATCAAGTCCGGGGTGAAATCCTCCAGCTTAACTGGAGAATAGCCTTGGATACTGGTGGTCTCGAGTGAGGTAGGGGCAGCTGGAACTTCCGGTGGAGCGGTGAAATGCGTTGATATCGGAAAGAACGCCGGTGGCGAAAGCGAGCTGCTGGACCTCTTCTGACGCTGAGGCGCGAAAGCCAGGGG |
| P-asv5539 | GCAAGCGTTACCCGGAATCACTGGGCGTAAAGGGCGTGTAGGCGGGAAGCTAAGTCCGACTTTAAAGACCGGGGCTCAACCCCGGGAGTGGGTTGGAGACTGGCTTCCTGGACCTCTGGAGAGGCAACTGGAATTCCTGGTGTAGCGGTGGAATGCGTAGATACCAGGAGGAACACCAATGGCGAAGGCAGGTTGCTGGACAGAAGGTGACGCTGAGGCGCGAAAGTGTGGGG |
| P-asv5540 | GCGAGCGTTGTCCGGATTTATTGGGCGTAAAGAGTTCGTAGGCGGTTTGTTAAGTCTGATGTTAAAGACCGGGGCTCAACCTCGGAAATGCATTGGATACTGGCAGACTGGAGTGCAGTAGAGGCTAGTGGAATTCCCAGTGTAGCGGTGAAATGCGTAGATATTGGGAAGAACACCGGTGGCGTAGGCGACTAGCTGGGCTGTAACTGACGCTGAGGAACGAAAGCCAGGGG |
| P-asv5541 | GCAAGCGTTATTCGGATTTACTGGGCATAAAGAGCGCGTAGGCGGCTCGGTCAGTCGGAGGTGAAAGCCCGCAGCTCAACTGCGGAACTGCCCCCGATACTGCCGAGCTTGAGTGCAGGAGGGGTGACTGGAACTCAGGGTGTAGCGGTGAAATGCGTAGATATCCTGAGGAACACCGGAGGCGAAGGCGAGTCACTGGACTGCTACTGACGCTGAGGCGCGAAAGCTAGGGG |
| P-asv5542 | GCAAGCGTTGTTCGGAATCACTGGGCGTAAAGCGCGCGCAGGCGGGTGTGAAAGTCGGAGGTGAAAGGCCGGGGCTCAACCCCGGACCGTTCTTTCGATACTTCATGCCTAGGGACCGGTAGAGGCCAGTAGAATTCCCGGAGTAGCGGTGGAATGCGTAGAGATCGGGAAGAACACCAGAGGCGAAGGCGACTGGCTGGGCCGGATCCGACGCTCAGGCGCGAAAGCGTGGGG |

| | |
|---|---|
| P-asv5543 | CCAAGTGATGGCCGATCTTATTGGGCCTAAAGCGTCCGTAGCCGGCCTCGCAAGTCCGTCGGGAAATCCAGTAGCTCAACTACTGGGCGTCCGGCGGAAACTCCGAGGCTTGGGACCGGAAGGCTCGAGGGGTACGTCCGGGGTAGGAGTGAAATCCCGTAATCCCGAACGGACCACCGATGGCGAAAGCACCTCGAGAAGACGGCTCCGACAGTGAGGGACGAAAGCTCGGGT |
| P-asv5544 | TCCCTTTCTTTCCAATATTATTGGGCCTAAAGCGTCCGTAGCTGGCCGCGCAAGTCCGTCGGGAAATCCACCTGCCCAACAGGTGGGCGTCCGGCGGAAACTGTGTGGCTTGGAACCGGAAGGCTCAGAGGGTACGTCCGGGGTAGGAGTGAAATCCCGTAATCCCGGACGGACCACCGATGGCGAAAGCACTCTGAGAAGACGGCTTCGACAGTGAGGGACGAAAGCTAGGGT |
| P-asv5545 | CCAAGCGTTAATCGGAATCACTGGGCTTAAAGCGTGCGCAGGCGGACCGGTAAGTACCTTGTGAAATCCCACGGCTCAACCGTGGAACTGCAGGGTATACTGCCGGTCTTGAGGCCAGTAGGGGTCGTCGGAACTGTAGGTGTAGCGGTGAAATGCGTAGATATCTACAGGAACGCCAAAGGTGAAGACAGACGACTGGGCTGGTTCTGACGCTCATGCACGAAAGCGTGGGG |
| P-asv5546 | GCAAGCGTTATCCGGATTTACTGGGCGTATAGAGCGTTGAGGCGGCTCTGTAAGTTGGGCGTGAAAGTCCCGGGCCTAACCCGGGGAAGCCGTTCAAGACTGCAGAGCTTGAGGGCAGCAGAGGGAGGTGGAATTCCCGGTGTAGCGGTGAAATGCGTAGATATCGGGAGGAACACCAGTGGCGAAGGCGGCCTCCTGGGCTGTACCTGACGCTGAATGCGAAAGCTAGGGG |
| P-asv5547 | CCGAGCGTTACTCGGAGTCACTGGGCTTAAAGGGTGCGCAGGCGGACCGCGAGGTGTCCTGTGAAAGCCCACCGCTCAACGGTGGAATTGCAGGGCAAACCAACGGTCTTGAGGCAGGCAGGGGTCGACGGAACTGTGGGTGGAGCGGTGAAATGCGTAGATATCCACAGGAACGCCAGTGGCGAAGGCGGTCGACTGGGCCTGTTCTGACGCTGAGGCACGAAAGCGTGGGG |
| P-asv5548 | GCAAGCGTTAATCGGAATTACTGGGCGTAAAGCGTGCGTAGGCGGCCACGTAAGTCGGATGTGAAAGCCCCGGGCTTAACCTGGGAATTGCATTCGATACTGCGTGGCTAGAGTTTGGTAGAGGTAAGTGGAATTCCAGGTGTAGCGGTGAAATGCGTAGATATCTGGAGGAACACCAGTGGCGAAGGCGGCTTACTGGACCAAAACTGACGCTGAGGTACGAAAGCGTGGGT |
| P-asv5549 | TCGAGCGTTATCCGGATTTATTGGGCGTAAAGCGTCCGCAGTCGGATTGAAAAGTCTTTGCCTAAATCCTTGGGCTCAACCCAAGGCATGGCATTGAAACTTTCGATCTAGAGTATGTCAGAGGTAAGCGGAACTCATGGTGTAGCAGTGATAAGCGTTGATATCATGGGGAACACCAAAGGCGAAGGCAGCTTACTGGGACATTACTGACGATCAGGGACGAAAGTGTAGGT |
| P-asv5550 | GCTAGCGTTATTCGGAATTACTGGGCGTAAAGCGCACGTAGGCGGCTTCGTAAGTTAGAGGTGAAAGCCCCGAGCTCAACCTGGGAACTGCATCCAAAACTGGCAAGCTAGAGTATGGCAGAGGGTGGTGGAATTTCCTGTGTAGCGGTGAAATGCGTAGATATAGGAAGGAACACCAGTGGCGAAGGCGACCACCTGGGCTAATACTGACACTGAGGTGCGAAAGCGTGGGG |
| P-asv5551 | GCTAGCGTTGTTCGGAATTACTGGGCGTAAAGCGCACGTAGGCGGCTTTGTAAGTTAGAGGTGAAAGCCCAGAGCTCAACTCTGGAACTGCCTTTAAGACTGCATCGCTTGAATCATGGAGAGGTTAGTGGAATTCCGAGTGTAGAGGTGAAATTCGTAGATATTCGGAAGAACACCAGTGGCGAAGGCGACTAACTGGACATGTATTGACGCTGAGGTGCGAAAGCGTGGGG |
| P-asv5552 | GCGAGCGTTGTCCGGAATCATTGGGCGTAAAGGGTGCGCAGGCGGTATGGAAGGTCAGTTGTGAAAGCCACAGGCTCAACTTGTGGAGTGCAATTGAAACCATCATACTAGAGAGCGGAAGAGGAAAGCGGAATTCCATGTGTAGCGGTGAAATGCTTAGATATGTGGAAGAACACCAATGGCGAAGGCAGCTTTCTGGTCCGTTACTGACGCTGAGGCACGAAAGCGTGGGG |
| P-asv5553 | GCAAGCGTTACTCGGAATTACTGGGCGTAAAGCGTCTGTAGGCGACGTCGTAAGTCTGGCGTTAAATCGCGGAGCTCAACTCCGCAACCGCGCTGGAAACTGCGATGATAGAGGGACTCAGAGGCATCTGGAATGCCGTGTGTAGGGGTAAAATCCGTTGATCCACGGTAGAACGCCAAAAGTGAAGACAGGATGCTGGGAGTCACCTGACGCTGAGAGACGAAAGCGTGGGG |
| P-asv5554 | GTTAGCGTTGTTCGGAATTACTGGGCGTAAAGCGCACGTAGGCGGATCAGAAAGTTGGGGGTGAAATCCCAGGGCTCAACCCTGGAACGGCCTCCAAAACTCCTGGTCTTGAGTTCGAGAGAGGTGAGTGGAATTCCTAGTGTAGCGGTGAAATGCTTGGATATTACACAGAATACCGATTGCGAAGGCAGTCTACTACGTATGTACTGACGCTCATGGACGAAAGCGTGGGG |
| P-asv5555 | GCAAGCGTTGCTCGGAATTATTGGGCGTAAAGCGCGTGTAGGCGGTCTTCTAAGTCTGGTGTGAAATCCCGGGGCTTAACCCCGGAAGTGCATTGGATACTGGAAGGCTAGAGGACGAGAGAGGATGGTGGAATTCCTGGTGTAGGGGTGAAATCCGTAGATATCAGGAGGAACATCAGTGGCGAAGGCGGCCATCTGGCTCGTACCTGACGCTGAGACGCGAAAGCGTGGGG |
| P-asv5556 | TCGAGCGTTGCGCGGAATCACTGGGCTTAAAGGGTGCGTAGGCGGATGTACAAGTGCCTTGTGAAATCCCCGGGCTCAACCCGGGAACTGCTTGGTAGACTGTATGTCTTGAGGTCGGTAGGGGTTGCTGGAACAGTTGGTGGAGCGGTGAAATGCGTAGATATCAACTGGAACGCCAAAGGCGAAGGCAAGCAACTGGGCCGATTCTGACGCTGAGGCACGAAAGCGTGGGG |
| P-asv5557 | GCAAACGTTGCTCGGAATCATTGGGCGTAAAGCGCGCGTAGGCGGCCGGACAAGTTGGATGTGAAAGCCCTGGGCTCAACCCGGGATGTGCATTCAAAACTGTCTGGCTTGAATCTCGGAGGGGGTCAGGGAATTCTCGGTGTAGAGGTGAAATTCGTAGATATCGGGAGGAACACCAGTGGCGAAGGCGCTGACCTGGACGATGATTGACGCTGAGGCGCGAAAGCGTGGGG |
| P-asv5558 | GCTAGCGTTGTTCGGAATTACTGGGCGTAAAGGGCGCGTAGGCGGACCTTTTAGTCAGAGGTGAAATCCCGGGGCTCAACCTCGGAACTGCCTTTGATACTGGAGGTCTTGAGTCTGTGAGAGGGTGGCGGAATACCCAGTGTAGAGGTGAAATTCGTAGATATTGGGTAGAACACCAGTGGCGAAGGCGGCCACCTGGCACAGTACTGACGCTGAGGCGCGAAAGCGTGGGG |
| P-asv5559 | GCAAGCGTTAATCGGAATTACTGGGCGTAAAGCGCGCGTAGGTGGATTGATATGTTGGCTGTGAAAGCCCCGGGCTCAACCCGGGAAGGCCAGCCAAAACTGTCAGCCTAGAGTATAGTAGAGGGAAGTGGAATTTCAGGTGTAGCGGTGAAATGCGTAGATATCTGAAGGAACATCAGTGGCGAAGGCGACTTTCTGGACTAATACTGACACTGAGGCGCGAAAGCGTGGGG |
| P-asv5560 | ACGAGTGGCTATCACGTTTATTGGGCCTAAAGCGTCCGTAGTCTGCCAGGTGCATCCTCTGTGAAATCGCAGCGCTTAACGTTGCGGCATGCGGAGGACACGACCTGGCTCGAGAGCGGGAGGCGTCGGGGTATTCCAAGGGGACCGGTAAAATGGTATAATCCTTGGAGGACCACCTGTGGCGAAGGCGCCCGACGAGAACGCATCTGACGATGAGGGACGAAAGCCAGGGG |
| P-asv5561 | GCGAGCGTTGTCCGGAATCATTGGGCGTAAAGAGCACGTAGGTGGATCAATTAGTCTGCTGTGAAAGTCGAAGGCTCAACCTTCGAAAGCCGGTGGATACTGTTGATCTAGAGTACGGAAGAGGCGAGTGGAATTCCTGGTGTAGCGGTGGAATGCGCAGATATCAGGAGGAACACCAATAGCGAAGGCAGCTCGCTGGGACGTAACTGACACTGAGGTGCGAAAGCGTGGGG |
| P-asv5562 | CCGAGCGTTGTCCGGAATCATTGGGCGTAAAGGGTACGTAGGCGGCCCAGTAAGTTAGAAGTGAAATAATATAGCTCAACTATATAAAGCTTTTAAAACTGCTAGGCTTGAGAGATGGAAGGGAAAGTGGAATTCCTAGTGTAGCGGTGAAATGCGCAGATATTAGGAAGAATACCGGTGGCGAAGGCGACTTTCTGGTCATCATCTGACGCTGAGGTACGAAAGCGTGGGT |
| P-asv5563 | TCTCTCTTTTTTCTTCCTTCCTGGGCGTAAAGCGCACGTAGGCGGCTTTGTAAGTCAGAGGTGAAAGCCTGGAGCTCAACTCCAGAACTGCCTTTGAGACTGCATCGCTTGAATCCAGGAGAGGTGAGTGGAATTCCGAGTGTAGAGGTGAAATTCGTAGATATTCGGAAGAACACCAGTGGCGAAGGCGGCTCACTGGACTGGTATTGACGCTGAAGTGCGAAAGCGTGGGG |
| P-asv5564 | GCAAGCGTTAATCGGATTTATTGGGCGTAAAGAGCGCGTAGGCGGGGTGGACAGTTAGATGTGAAAGCCCGGAGCTCAACTCCGGACCCGCATCTAATACTACCATCCTAGAGGATGGACGGAGAAAAGGGAATTCCACGTGTAGCGGTGAAATGCGTAGATATGTGGAAGAACACCAGTGGCGAAGGCGCTTTTCTAGTTTATTCCTGACGTTGAGGTGCGAAAGTAAGGGG |
| P-asv5565 | ACAAGTGGAGATCATGATTACTGGGTCTAAAGTGTTCGTAGCTGGTTGTGCAAGTTTCTTGTTAAATATTTGTAGCTTAACTCAAATCTGCAAGAAATACTGTTCAACTTGAGACCGGAAGGGGTATGAAGTATGAATTGGGTAGCGGTAAAATGCTATAATCCTTTTCAGACTAACAAAGGCGAAGGCATCATACTAAGACGGATCTGACAGTGAGGAACGAAAGCTAGGGG |
| P-asv5566 | TCAAGCGTTGTCCGGATTTATTGGGCGTAAAGAGCTCGTAGGCGGTTCAACAAGTCGGTCGTAAAAGTTTAGGGCTCAACCCTAAAATGTCGATCGATACTGTTGTGACTAGGATACGGTAGAGGTGAATGGAATTCCGAGTGTAGCGGTGAAATGCGTAGATATTCGGAGGAACACCAATTGCGAAGGCAGTTCACTGGACCGTGATCGACGCTGAGGAGCGAAAGCTAGGGG |
| P-asv5567 | GCAAGCGTTGTCCGGAATCACTGGGCGTAAAGCGCGCGCAGGCGGCCATTTGCGTCCGGGGTGAAAGCCCAGAGCTCAACTCTGGGATTGCCTTGGATACGGGATGGCTTGAGGATCGGAGAGGCAAGGGGAATTCCACGTGTAGCGGTGAAATGCGTAGAGATGTGGAGGAACACCTGTGGCGAAGGCGCCTTGCTGGCCGATTTCTGACGCTGAGGCGCGAAAGCGTGGGG |
| P-asv5568 | GCAAGCGTTAATCGGAATTACTGGGCGTAAAGCGTGCGCAGGCGGTTATGCAAGACAGAGGTGAAATCCCCGGGCTCAACCTGGGAACTGCCTTTGTGACTGCATAGCTAGAGTACGGTAGAGGGGGATGGAATTCCGCGTGTAGCAGTGAAATTCGTAGATATTCGCAAGAACACCAGTGGCGAAGGCGGCTTACTGGCCCGATACTGACACTGAGGCACGAAAGCGTGGAG |
| P-asv5569 | GCAAGCGTTGTCCGGAATTATTGGGCGTAAAGCGCGCGCAGGCGGCTACTTAAGTCCATCTTAAAAGTGCGGGGCTTAACCCCGTGATGGGATGGAAACTGAGAAGCTGGAGTGTCGGAGAGGAAAGTGGAATTCCTAGTGTAGCGGTGAAATGCGTAGAGATTAGGAAGAACACCGGTGGCGAAGGCGACTTTCTGGACGACAACTGACGCTTAGGCGCGAAAGCGTGGGG |
| P-asv5570 | GCGAACGTTATTCGGAATCACTGGGCTTAAAGCGCGTGTAGGCGGGCGACCACGTCGGACGTTGAAAGCCCCCGGCTCAACCGGGGGATCGGCGCCGATACGGGCCGCCTGGAGGGGCGTAGGGGGAGCTGGAACTTCCGGTGGAGCGGTGAAATGCGTTGAGATCGGAAGGAACGCCCGCGGCGAAAGCGAGCTCCTGGACGCCTACCGACACTGAGGCGCGAAAGCCAGGGG |
| P-asv5571 | GCTAGCGTTGTTCGGAATTACTGGGCGTAAAGCGTACGTAGGCGGCTTTGTAAGTTAGAGGTGAAAGCCTGGTGCTCAACACCAGAACTGCCTTTAAGACTGCATCGCTTGAACGTCGGAGAGGTGAGTGGAATTCCGAGTGTAGAGGTGAAATTCGTAGATATTCGGAAGAACACCAGTGGCGAAGGCGGCTCACTGGACGACTGTTGACGCTGAGGTGCGAAAGCGTGGGG |
| P-asv5572 | GCAAGCGTTGTTCGGAATCATTGGGCGTAAAGCGTGCGCAGGCGGCCTGTTAAGTCTGGTGTGAAAACCCAGGGCTCAACCCTGGAGGTGCATTGGAAACTGGCAGGCTTGAATCCGAGAGGGGTCAGTGGAATTCGTGGTGTAGGAGTGAGATCCGTAGAGATCACGAGGAACACCGGCGTCGAAGGCGACTGACTGGATCGGTATTGACGCTGAGGCACGAAAGCGTGGGG |
| P-asv5573 | GCTAGCGTTGTTCGGAATTACTGGGCGTAAAGCGCGCGTAGGCGGATCGAAAAGTTGGGGGTGAAATCCCGGAGCTCAACTCCGGAACTGCCCTCAAAACTAGCGGTCTCGAGTTCTGGAGAGGTGAGTGGAATTCCGAGTGTAGAGGTGAAATTCGTAGATATTCGGAGGAACACCAGTGGCGAAGGCGGCTCACTGGCCAGATACTGACGCTGAGGCGCGAAAGCGTGGGG |
| P-asv5574 | TCGAGTGATGACCGATCTTATTGGGCCTAAAGCGTCCGTAGCCGGCCGGGCAAATCCGTCGGGAAATCTACCAGCTCAACTGGTAGGCGTCCGGCGGAAACTGCTCGGCTTGGGACAGGGAGACTCAAGAGGTACGTCCGGGGTAGGAGTGAAATCCCGTAATCCTGGACGGACCACCGATGGCGAAAGCATCTTGAGAGAACTGCTCCGACGGTCAGGGACGAAAGCTGGGGT |
| P-asv5575 | TCAAGCGTTGTTCGGAATCACTGGGCGTAAAGGGTGCGTAGGTGGCGTGGTAAGTCAGATGTGAAAGCCCGGGGCTCAACCTCGGAACTGCATCCGATACTGCCATGCTAGAGTACTGGAGAGGTGACTAGAATTCTCGGTGTAGCAGTGAAATGCGTAGATATCGAGAGGAATACCAAAGGCGTAGGCAGGTCACTGGACAGTTACTGACACTGAGGCACGAAGGCCAGGGG |
| P-asv5576 | GCGAGCGTTAATCGGAATTACTGGGCGTAAAGCGTGCGCAGGCGGTTTTGAAAGTCAGATGTGAAATCCCCGAGCTCAACTTGGGAACTGCGTTTGAAACTCCAAAGCTAGAGTATAGGAGAGGGGGGTAGAATTCCACGTGTAGCAGTGGAATGCGTAGAGATGTGGAGGAATACCAGTGGCGAAGGCGGCCTTCTGGACTGACACTGACACTGAGGTGCGAAAGCGTGGGT |
| P-asv5577 | GCGAGCGTTATCCGGAATCACTGGGCGTAAAGGGCGCGTAGGCGGTCTAGTAAGTCCGATGTTAAAGAGCGGGGCTCAACCCCGTCACGGCGTTGGATACTGCTAGGCTTGACGACTGGAGAGGTGAGTGGAATTACCAGTGTAGCGGTGGAATGCGTAGATACTGGTAGGAACACCCATTGCGAAGGCAGCTCACTGGACAGTACGTGACGCTGAGGCGCGAAAGTGTGGGG |
| P-asv5578 | GCAAGCGTTAATCGGAATTACTGGGCGTAAAGCGTGCGCAGGCGGTTTTGTAAGTCTGATGTGAAAGCCCCGGGCTCAACCTGGGAATTGCATTGGAGACTGCAAGGCTTGAATCTGGCAGAGGGGGGTAGAATTCCACGTGTAGCAGTGAAATGCGTAGAGATGTGGAGGAACACCGGTGGCGAAGGCAGCCCCCTGGGTCAAGATTGACGCTCATGCACGAAAGCGTGGGG |
| P-asv5579 | CCAAGCGTTAATCGGAATTACTGGGCGTAAAGCGTGCGCAGGCGGTTGCGCAAGACCGATGTGAAATCCCCGAGCTTAACTTCGGAACTGCCTTTGATACAGCATATCTCGAGTCCGAGAGAGGTGAGTGGAATTCCTAGTGTAGAGGTGAAATTCGTAGATATTAGGAAGAACACCGGTGGCGAAGGCGGCTCACTGGCTCGGTACTGACGCTGAGGTGCGAAAGCGTGGGG |
| P-asv5580 | GCAAGCGTTAATCGGATTTACTGGGCGTAAAGCGTGCGTAGGCGGTTCGTTAGGTCGATTGTGAAAGCCCTGGGCTTAACCTGGGAATTGCAGTCGATACTGGCGGACTAGAGAACGGTAGAGGGAGGCGGAACTCCAGGTGTAGCGGTGAAATGCGTAGATATCTGGAAGAACACCGATGGCGAAGGCAACCTCCTGGGCCTGTTCTGACGCTGAGGCACGAAAGCGTGGGT |
| P-asv5581 | TCGAGTGTTGTCCCATATTATTGGGCCTAAAGCGTCCGTAGCTGGCCAGGCAAGTCTATCGGGAAATCGACCAGCTCAACTGGTCGGCGTCCGGTAGAAACTACGCGGCTTGGGGCCGAGAGACTTGACGGGTACGTCCAGGGTAGGAGTGAAATCCTGTAATCCTGAACGGACCACCAATGGGGAAACCACGTCAAGAGACCGGACCCGACAGTGAGGGACGAAAGCCAGGGT |

| | |
|---|---|
| P-asv5582 | GCAAGCGTTAATCGGAATGACTGGGCGTAAAGCGTGCGTAGGCGGTGATATTAGTCAGCCGTGAAAGCCCCGGGCTCAACCTGGGAATCGCGGTTGAAACGGGATGACTCGAGTGGGCTAGAGGACTGTGGAACTCCCGGTGTAGCGGTGAAATGCGTAGAGATCGGGAAGAACACCGATGGCGAAGGCAGGTCTCTGGGCCACTACTGACGCTGAGAAGCGAAAGCATGGGG |
| P-asv5583 | GCAAGCGTTAATCGGAATTACTGGGCGTAAAGCGTGCGCAGGCGGACTTTTAAGCCAGATGTGAAAGCCCCGAGCTTAACTTGGGAATTGCGTTTGGAACTGGGAGTCTAGAGTCTGTCAGAGGGGGATGGAATTCCACGTGTAGCAGTGAAATGCGTAGAGATGTGGAGGAACACCGATGGCGAAGGCAGTCCCCTGGGATAAGACTGACGCTCATGCACGAAAGCGTGGGG |
| P-asv5584 | ACAAACGTTATTCGGAATCACTGGGCTTAAAGAGTTCGTAGGCGGCCGTGTAGGTGAGATGTGAAATCCCTCGTCTCAACCGAGGAATTGCGTTTCAAACCACATGGCTTGAGGAAGATAGGGGTGAGCGGAACTGTTGGTGGAGCGGTGAAATGCGTTGATATCAACAGGAACACCAGAGGCGAAGGCGGCTCACTGGGTCTTTTCTGACGCTGAGGAACGAAAGCTAGGGG |
| P-asv5585 | GCGAACGTTGTTCGGAATCACTGGGCTTAAAGGGCGCGTAGGCGGGCTATCAAGTCCGGGGTGAAATCCTCCAGCTCAACTGGAGAACTGCCTCGGATACTGATGGTCTCGAGGAGGATAGGGGCACACGGAACTGTGGGTGGAGCGGTGAAATGCGTTGATATCCATAGGAACTCCGGTGGCGAAAGCGGTGTGCTGGATCCTTTCTGACGCTGAGGCGCGAAAGCCAGGGG |
| P-asv5586 | GCAAGCGTTATCCGGAATCATTGGGTTTAAAGGGTCCGCAGGCGGAATAATAAGTCAGTGGTGAAATCTTACAGCTTAACTGTGAAATTGCCATTGATACTGTTGTTCTTGAATTTGGTCGAAGTGGGCGGAATATGACATGTAGCGGTGAAATGCATAGATATGTCATAGAACACCGATAGCGAAGGCAGCTCACTAGGCCTGAATTGACGCTCATGGACGAAAGCGTGGGG |
| P-asv5587 | GCAAGCGTTAATCGGAATTACTGGGCGTAAAGCGTGCGCAGGCGGTTTTGTAAGTCTGTTGTGAAATCCCCGGGCTTAACCTGGGAATGGCAATGGAGACTGCAAGGCTAGAGTTTGGCAGAGGGGGGTAGAATTCCACGTGTAGCAGTGAAATGCGTAGAGATGTGGAGGAACACCGATGGCGAAGGCAGCCCCCTGGGTCAAGACTGACGCTCATGCACGAAAGCGTGGGG |
| P-asv5588 | GCAAGCGTTGTCCGGATTTATTGGGCGTAAAGAGCTCGTAGGCGGTCTGTCGCGTCTGCTGTGAAAACTCGGGGCTTAACCCCGGGCCTGCAGTGGGTACGGGCAGACTAGAGTGCGGTAGGGGAGACTGGAATTCCTGGTGTAGCGGTGGAATGCGCAGATATCAGGAGGAACACCGGTGGCGAAGGCGGGTCTCTGGGCCGTAACTGACGCTGAGGAGCGAAAGCATGGGG |
| P-asv5589 | GCAAGCGTTATCCGGATTTATTGGGTTTAAAGGGTCCGTAGGCGGATCTGTAAGTCAGTGGTGAAATCTCACAGCTTAACTGTGAAACTGCCATTGATACTGCAGGTCTTGAGTGTTATTGAAGTAGCTGGAATAAGTAGTGTAGCGGTGAAATGCATAGATATTACTTAGAACACCAATTGCGAAGGCAGGTTACTAAGCAACAACTGACGCTGATGGACGAAAGCGTGGGG |
| P-asv5590 | GCAAACGTTGCTCGGAATCATTGGGCGTAAAGCGCGCGTAGGCGGTTGAACAAGTTGGATGTGAAAGCCCTGGGCTCAACCCAGGATGTGCATTCAAAACTGTTCGACTTGAATCTCGGAGGGGGTCAGAGAATTCCCGGTGTAGAGGTGAAATTCGTAGATATCGGGAGGAACACCAGTGGCGAAGGCGCTGACCTGGGCGATGATTGACGCTGAGGCGCGAAAGCGTGGGG |
| P-asv5591 | GCAAGCGTTGTTCGGATTTATTGGGCGTAAAGAGTATGTAGGCGGTTCTACAAGTCAGGTGTGAAATCTTGGGGCTCAACCCCGAAACTGCGCCTGAAACTATAGAACTAGAATCTTGAAGGGGAAGGTAGAATTTCGCATGTAGGGGTAAAATCCGTAGAGATGCGAAGGAACATCAGAGGCGAAGGCGACCTTCTGGGCAAGTATTGACGCTGAGATACGAAAGCGTGGGT |
| P-asv5592 | GCAAGCGTTAATCGGAATTACTGGGCGTAAAGGGAGCGTAGGCGGGAGGTCAAGTCCGGTGAGAAAGGCCCTGGCTCAACCAGGGACGAGCGCTGGAAACTGACAATCTTGAGTACAGCAGAGGAAGAGGGAATTACCGGTGTAGGGGTGAAATCCCTAGAGACCGGTAAGAAGGCCAGTGGCGAAGGCGCTTTTCTGGGCTGTAACTGACGCTGAGGCTCGAAAGCCAGGGG |
| P-asv5593 | GCAAGCGTTAATCGGAATTACTGGGCGTAAAGCGCACGTAGGCGGTGATCCATGTCAGAGGTGAAATCCCGGAGCTCAACTTCGGAATTGCCTTTGAAACGGGATCGCTTGAGTCCGAGAGAGGATGGCGGAATTCCTAGTGTAGAGGTGAAATTCGTAGATATTAGGAAGAACACCAGTGGCGAAGGCGGCCATCTGGCTCGGTACTGACGCTCAGGTGCGAAAGCGTGGGG |
| P-asv5594 | ACGAGTGATGGCCGATATTATTGGGCCTAAAGCGTCCGTAGCCGGCCGCGCAAGTCCGTTGGGAAATCCACGCGCTCAACGCGTGGGCGTCCAGCGGAAACTGTGCGGCTTGGGGCCGGAAGACTCGAGGGGTACGTCCGGGGTAGGAGTGAAATCCCGTAATCCTGGACGGACCACCGGTGGCGAAAGCGCCTCGAGAAGACGGACCCGACGGTGAGGGACGAAAGCTTGGGT |
| P-asv5595 | TCGAGCGATGACCGATCTTATTGGGCCTAAAGCGTCCGTAGCCGGCCGGGCAAGTTCATCGGGAAATCTACCAGCTCAACTGGTAGGCGTCCGGTGAAAACTGCTCGGCTTGGGACAGGGAGACTCAAGAGGTACGTCCGGGGTAGGAGTGAAATCCCGTAATCCTGGACGGACCACCGATGGCGAAAGCATCTTGAGAGAACTGCTCCGACGGTCAGGGACGAAAGCTGGGGT |
| P-asv5596 | GCAAGCATTAATCGGATTTATTGGGCGTAAAGGGTGCGTAGGCGGCGAGGAAAGTCAGATGTGAAATCTCGGAGCTCAACTCCGGAATAGCATTTGAAACTGTCTTGCTAGAGGGTAGACGGAGAAAATGGAATTCCACAAGTAGCGGTGAAATGCGTAGATATGTGGAGGAACACCTGTGGCGAAAGCGATTTTCTAGTTTATACCTGACGCTGAGGCGCGAAAGCAAGGGG |
| P-asv5597 | GCGAGCGTTAATCGGATTTACTGGGCGTAAAGCGTGCGTAGGCGGCTTTTTAAGTCGGATGTGAAATCCCTGAGCTTAACTTAGGAATTGCATTCGATACTGGGAAGCTAGAGTATGGGAGAGGATGGTAGAATTCCAGGTGTGGCGGTGAAATGCGTAGAGATCTGGAGGAACATCAGTGGCGAAGGCGGCCACCTGGCCCAATACTGACGCTGAGGCACGAAAGCGTGGGG |
| P-asv5598 | GCAAGCGTTAATCGGAATTACTGGGCGTAAAGGGCTTGTAGGTGGTTATCTAAGTCGGATGTTAAATCCCCGGGCTTAACCTGGGACTGTCATTCGATACTGGATAACTTTGAGTTGGGTAGAGGTAAGTGGAATTTCCGGTGTAGCGGTGAAATGCGTAGATATCGGAAGGAACATCAGTGGCGAAGGCGGCTTACTGGACCCAAACTGACACTGAGGAGCGAAAGCGTGGGG |
| P-asv5599 | GCGAGCGTTGTCCGGAATTATTGGGCGTAAAGGGCTCGTAGGCGGTTCGTCGCGTCGGGAGTGAAAACTCAGGGCTTAACCCTGAGCCTGCTTCCGATACGGGCGGACTAGAGGTAGGTAGGGGAGAACGGAATTCCTGGTGTAGCGGTGGAATGCGCAGATATCAGGAGGAACACCGATGGCGAAGGCAGTTCTCTGGGCCTTACCTGACGCTGAGGAGCGAAAGCGTGGGG |
| P-asv5600 | GCTAGCGTTGTTCGGAATCACTGGGCGTAAAGGGCGCGTAGGCGGCTGTGGCAGTCGGGTGTGAAAGCCCGGAGCTCAACTCCGGAACGGCACTCGAGACGCCACGGCTGGAGGCCGGAAGAGGAGGGTGGAATACCCAGTGTAGAGGTGAAATTCGTAGATATTGGGTGGAACACCGGTGGCGAAAGCGGCTCTCTGGTCCGGACCTGACGCTGAGGCGCGAAAGCGTGGGG |
| P-asv5601 | GCTAGCGTTGTTCGGAATTACTGGGCGTAAAGCGCACGTAGGCGGATCGATCAGTCAGGGGTGAAATCCCGCAGCTCAACTGCGGAACTGCCTTTGATACTGTCGGTCTAGAGTATGGAAGAGGTGAGTGGAATTCCGAGTGTAGAGGTGAAATTCGTAGATATTCGGAGGAACACCAGTGGCGAAGGCGGCTCACTGGTCCATTACTGACGCTGAGGTGCGAAAGCGTGGGG |
| P-asv5602 | TCGAGCGTTACCCAGATTCACTGGGCGTAAAGGGTGCGTAGGCGGATATATTAGTCAGATGTTAAATCTTTGGGCTCAACCCAAAGGCCGCATTTGAAACGGTATATCTAGAAGAAGTCAGAGGTGTGCGGAACTCAAGGTGTAGGGGTGAAATCCGTTGATATCTTGGGGAACACCAAATGCGAAGGCAGCACACTGGGACTTTCTTGACGCTGAGGCACGAAAGCGTGGGT |
| P-asv5603 | GCAAGCGTTATCCGGAATTATTGGGCGTAAAGCGCGCGCAGGTGGTTTCTTAAGTCTGATGTGAAAGCCTACGGCTCAACCGTGGAGGGTCATTGGAAACTGGGAGACTTGAGTGCAGAAGAGGAAAGTGGAATTCCATGTGTAGCGGTGAAATGCGTAGAGATATGGAGGAACACCAGTGGCGAAGGCGACTTTCTGGTCTGTAACTGACACTGAGGCGCGAAAGCGTGGGG |
| P-asv5604 | GCGAGCGTTGTCCGGAATCACTGGGCGTAAAGGGCGCGTAGGCGGTCCGCTAAGTGTGTGGTGAAAGCTCGGGGCTCAACCCCGAGTCGGCCGTGCAAACTGGTGGACTTGAGCATGTCAGAGGCAGGCGGAATTCCGGGTGTAGCGGTGGAATGCGTAGAGATCCGGAAGAACACCGGTGGCGAAGGCGGCCTGCTGGGACATGGCTGACGCTGAGGCGCGACAGCGTGGGG |
| P-asv5605 | GCGAGCGTTGTCCGGAATTATTGGGCGTAAAGCGCGCGCAGGCGGTCTCTTAAGTCTGATGTGAAAGCCCACGGCTCAACCGTGGAGGGTCATTGGAAACTGGGGGACTTGAGGGCAGGAGAGGAGAGCGGAATTCCACGTGTAGCGGTGAAATGCGTAGAGATGTGGAGGAACACCAGTGGCGAAGGCGGCTCTCTGGCCTGCACCTGACGCTGAGGCGCGAAAGCGTGGGG |
| P-asv5606 | GCTAGCGTTATTCGGAATTACTGGGCGTAAAGCGCACGTAGGCGGCTTTGTAAGTTAGAGGTGAAAGCCTGGAGCTCAACTCCAGAATGGCCTTCAATACTGGCTGTCTTGAGTACGGGAGAGGTGAGTGGAACTCCGAGTGTAGAGGTGAAATTCGTAGATATTCGGAAGAACACCAGTGGCGAAGGCGACTCACTGGCCCGTTACTGACGCTGAGGCGCGAAAGCGTGGGG |
| P-asv5607 | GCAAGCGTTGTCCGGATTTATTGGGCGTAAAGCGCTCGCAGGCGGTCTTTTAAGTCTGATGTGAAAGCCCACGGCTTAACCGTGGAGGGGCATTGGAAACTGGAGGACTTGAGTGCAGAAGAGGAGAGTGGAATTCCATGTGTAGCGGTGAAATGCGTAGAGATATGGAGGAACACCAGTGGCGAAGGCGACTCTCTGGTCTGTAACTGACGCTGAGGAGCGAAAGCGTGGGT |
| P-asv5608 | GCGAGCGTTAATCGGAATTACTGGGCGTAAAGCGTGCGCAGGCGGCTTTTTAAGCCAGATGTGAAATCCCCGGGCTTAACCTGGGAACTGCATTTGGAACTGGAAGGCTAGAGTGTAGCAGAGGGGGGTAGAATTCCACGTGTAGCAGTGAAATGCGTAGAGATGTGGAGGAATACCGATGGCGAAGGCAGCCCCCTGGGCTAACACTGACGCTCATGCACGAAAGCGTGGGG |
| P-asv5609 | GCAAGCGTTGTCCGGAATTATTGGGCGTAAAGAGCTCGTAGGCGGTTCGTCGCGTCGGCTGTGAAATCCCGAGGCTCAACCTCGGGCCTGCAGTCGATACGGGCGGACTTGAGTTCGGCAGGGGAGACTGGAATTCCTGGTGTAGCGGTGAAATGCGCAGATATCAGGAGGAACACCGGTGGCGAAGGCGGGTCTCTGGGCCGAAACTGACGCTGAGGAGCGAAAGCGTGGGG |
| P-asv5610 | GCGAGCGTTATCCGGAATTATTGGGCGTAAAGAGCTCGTAGGCGGTTTGTCACGTCTGCTGTGAAAGCCCGGGGCTCAACCCCGGGTGTGCAGTGGGTACGGGCAGACTAGAGTGCAGTAGGGGAGACTGGAATTCCTGGTGTAGCGGTGAAATGCGCAGATATCAGGAGGAACACCGATGGCGAAGGCAGGTCTCTGGGCTGTTACTGACGCTGAGGAGCGAAAGCATGGGG |
| P-asv5611 | GCTAGCGTTATTCGGAATTACTGGGCGTAAAGCGCACGTAGGCGGACTGGAAAGTCAGAGGTGAAATCCCAGGGCTCAACCTTGGAACTGCCTTTGAAACTCCCAGTCTTGAGGTCGAGAGAGGTGAGTGGAATTCCGGGTGTAGAGGTGAAATTCGTAGAGATGTGGAGGAATACCGATGGCGAAGGCAGCCCCCTGGGATAACACTGACGCTCATGCACGAAAGCGTGGGG |
| P-asv5612 | GCTAGCGTTGTTCGGAATGACTGGGCGTAAAGGGCGCGTAGGCGGCTTTATTAGTCAGGCGTGAAAGCCCCGGGCTTAACCTGGGAACTGCGTTTGATACTGTAGAGCTTGAGGATGGAAGAGGAAAGTGGAATTCCTAGTGTAGAGGTGAAATTCGTAGATATTAGGAGGAACATCAGAGGCGAAGGCGGCTTTCTGGTCCATCACTGACGCTGAGGCGCGAAAGCGTGGGG |
| P-asv5613 | GCGAGCGTTAATCGGATTTACTGGGCGTAAAGCGTGCGTAGGCGGCTTCTTAAGTCGGATGTGAAATCCCTGAGCTTAACTTAGGAATTGCATTCGATACTGGGAAGCTAGAGTATGGGAGAGGATGGTAGAATTCCAGGTGTAGCAGTGAAATGCGTAGATATCGGGAGGAACACCCGTGGCGAAGGCGGCCACCTGGACCAGCACTGACGCTGAGGCACGAAAGCGTGGGG |
| P-asv5614 | GCTAGCGTTATTCGGAATTACTGGGCGTAAAGCGCACGTAGGCGGACTGGAAAGTCAGAGGTGAAATCCCAGGGCTCAACCTTGGAACTGCCTTTGAAACTCCCAGTCTTGAGGTCGAGAGAGGTGAGTGGAATTCCGAGTGTAGAGGTGAAATTCGCAGATATTAGGAGGAACACCAGTGGCGAAGGCGACTGACTGGACCGGTATTGACGCTGAGGTACGAAAGCGTGGGT |
| P-asv5615 | GCTAGCGTTGTTCGGAATCACCGGGCGTAAAGCGCACGTAGGCGGCTTTTTAAGACAGATGTGAAATCCCCGGGCTCAACCTGGGAACTGCATTTGTGACTGTATAGCTAGAGTACGGCAGAGGGGGATGGAATTCCGCGTGTAGCAGTGAAATGCGTAGATATGCGGAGGAACACCGATGGCGAAGGCAATCCCCTGGGCCTGTACTGACGCTCATGCACGAAAGCGTGGGG |
| P-asv5616 | GCTAGCGTTGTTCGGAATTACTGGGCGTAAAGCGCACGTAGGCGGTATGTTAAGTTGGATGTGAAAGCCCCGGGCTCAACCTGGGAATTGCATTCGATACTGGGAAGCTAGAGTATGGGAGAGGATGGTAGAATTCCAGGTGTAGCGGTGAAATGCGTAGAGATCTGGAGGAATACCGATGGCGAAGGCAGCCATCTGGCCTAATACTGACGCTGAGGTACGAAAGCATGGGG |
| P-asv5617 | GCAAGCGTTAATCGGAATTACTGGGCGTAAAGCGCACGCAGGCGGCTTTTTAAGTCGGATGTGAAAGCCCCGGGCTCAACCTGGGAATTGCATTCGATACTGTAAAGCTAGAGTATGTGAGAGGGGGGTAGAATTCCAAGTGTAGCGGTGAAACTCGTAGATATTCGGAAGAACACCAGTGGCGAAGGCGACTCACTGGCCCATTACTGACGCTGAGGCTCGAAAGCGTGGGG |
| P-asv5618 | GCGAGCGTTGTCCGGAATTATTGGGCGTAAAGAGCTTGTAGGTGGTTTGTCGCGTCTGCGGTGAAATCCCAACGCTTAACGTTGGGCTTGCCGTGGGTACGGGCAGACTAGAGTGTGGTAGGGGAGACTGGAATTCCTGGTGTAGCGGTGAAATGCGCAGATATCAGGAGGAACACCGATGGCGAAGGCAGGTCTCTGGGCCATAACTGACACTGAGAAGCGAAAGCATGGGG |
| P-asv5619 | GCAAGCGTTGTTCGGAATCACTGGGCGTAAAGCGCGCGTAGGCGGCGATGCAAGTCAGAGGTGAAATCCCAAGGCTTAACCTTGGAATTGCCTTTGAAACTGCATTGCTAGAGTACTGGAGGGGATGGCGGAATTCCTAATGTAGCGGTGAAATGCGTAGATATTAGGAGGAACACCGGTGGCGAAGGCGGCCATCTGGACAGTCACTGACGCTGAGGCGCGAAAGCGTGGGG |
| P-asv5620 | GCGAACGTTATTCGGAATCACTGGGCTTAAAGCGCGTGTAGGCGGATCGCCGCGTCGGACGCTGAAATCCCCCGGCTTAACCGGGGAACGGGCGCCGAAACGGGCGGTCTGGAGGGGAGTAGGGGGACCTGGAACTTCCGGTGGAGCGGTGAAATGCGTTGAGATCGGAAGGAACGCCCGTGGCGAAAGCGAGGTCCTGGGCTCCTGCTGACGCTGAGACGCGAAAGCCAGGGG |

| | |
|---|---|
| P-asv5621 | GCAAGCGTTGTCCGGAATTATTGGGCGTAAAGGGCGCGCAGGCGGCCTGTTAAGTCTGATGTGAAAGACCGCGGCTCAACCGCGGGGTGGCATCAGAAACTGGCAGGCTTGAGTGCAGGAGAGGAGAGTGGAATTCCCGGTGTAGCGGTGGAATGCGTAGAGATCGGGAGGAACACCAGTGGCGAAGGCGGCTCTCTGGCCTGTTACTGACGCTGAGGCGCGAAAGCGTGGGG |
| P-asv5622 | GCAAACGTTGTTCGGAATCATTGGGCGTAAAGGGCGCGTAGGTGGCGATGTATGTCTGATGTGAAAGCCCGAAGCTTAACTTCGGAAGTGCGTCGGAAACTGCATCACTTGAGTCTCGGAGAGGATAACGGAATTCCTGATGTAGAGGTGAAATTCGTAGATATCAGGAGGAACACCGGTGGCGAAGGCGGTTATCTGGACGAAGACTGACGCTGAGGCGCGAAAGCGTGGGG |
| P-asv5623 | CCGAGCGTTAATCGGAATCACTGGGCTTAAAGCGTACGCAGGCGGACTTGTAGGTATCTTGTGAAATCCCACGGCTTAACCGTGGAACTGCAGGGTAAACCACAAGTCTTGAGACATGTAGGGGTCAATGGAACCATGAGTGGAGCGGTGAAATGCGTAGATATTCATGGGAACGCCAATGGCGAAGGCAGCTGACTGGGCATGTTCTGACGCTCAGGTACGAAAGCGTGGGT |
| P-asv5624 | GCAAGCGTTAATCGGAATTACTGGGCGTAAAGCGCGCGTAGGCGGCTTGTTAAGTCGGATGTGAAATCCCCGGGCTCAACCTGGGAACTGCATCCGATACTGGCTGGCTAGAGTGTGGTAGAGGGAAGTGGAATTCCAGGTGTAGCGGTGAAATGCGTAGATATCTGGAGGAACATCAGTGGCGAAGGCGGCTTCCTGGACCAACACTGACGCTGAGGTGCGAAAGCGTGGGG |
| P-asv5625 | GCGAGCGTTGTCCGGAATTATTGGGCGTAAAGGGCTTGTAGGCGGTTGGTCGCGTCTGCCGTGAAATTCTCTGGCTTAACTGGGGGCGTGCGGTGGGTACGGGCTGACTTGAGTGCGGTAGGGGAGACTGGAACTCCTGGTGTAGCGGTGGAATGCGCAGATATCAGGAAGAACACCGGTGGCGAAGGCGGGTCTCTGGGCCGTTACTGACGCTGAGGAGCGAAAGCGTGGGG |
| P-asv5626 | GCTAGCGTTGTTCGGAATCACTGGGCGTAAAGCGCACGTAGGCGGACTTTTAAGTCAGGGGTGAAATCCCAAGGCTCAACCTTGGAACTGCCTTTGATACTGGAAGTCTTGAGTTCGGGAGAGGTGAGTGGAACTGCGAGTGTAGAGGTGAAATTCGTAGATATTCGCAAGAACACCAGTGGCGAAGGCGGCTCACTGGCCCGATACTGACGCTGAGGTGCGAAAGCGTGGGG |
| P-asv5627 | GCTAGCGTTGTTCGGAATTACTGGGCGTAAAGCGCGCGTAGGCGGCTTTGTCAGTCAGGGGTGAAATCCCGGGGCTTAACCTCGGAACTGCCTTTGATACTGCAAGGCTTGAGTCTGTGAGAGGATGGTGGAATACCCAGTGTAGAGGTGAAATTCGTAGATATTGGGTGGAACACCAGTGGCGAAGGCGGCCATCTGGCACAGTACTGACGCTGAGGTGCGAAAGCGTGGGG |
| P-asv5628 | GCAAGCGTTGTCCGGATTTATTGGGTTTAAAGGGTGCGTAGGCGGCCTTGTAAGTCAGTGCTGAAATATCCCGGCTTAACCGGGAGGGTGGCATTGATACTGCAGGGCTTGAGTGGAGTCGAGGTAGGCGGAATTGACGGTGTAGCGGTGAAATGCTTAGATATCGTCAAGAACACCGATAGTGTAGACAGCTTACTAGGCTTCAACTGACGCTGAGGCACGAAAGTGTGGGG |
| P-asv5629 | GCTAGCGTTATCCGGATTTACTGGGCGTAAAGGGTGCGTAGGCGGTCTTTTAAGTCAGGAGTGAAAGGCCACGGCTCAACCGTAGTAAGCTCTTGAAACTGGAGGACTTGAGTGCAGGAGAGGAGAGTGGAATTCCTAGTGTAGCGGTGAAATGCGTAGATATTAGGAGGAACACCAGTAGCGAAGGCGGCTCTCTGGACTGTAACTGACGCTGAGGCACGAAAGCGTGGGG |
| P-asv5630 | GCTGGCGTTGTTCGGAATTACTGGGCGTAAAGGGCGCGTAGGCGGCGCTTCAAGTCAGATGTGAAAGCCCCGGGCTCAACCTGGGAACCGCATTTGAGACTGTTGTGCTTGAGGTCCGGAGAGGTGGGTGGAATTCCCAGTGTAGAGGTGAAATTCGTAGATATTGGGAAGAACACCGGTGGCGAAGGCGGCCCACTGGACGGAACCTGACGCTGAGGCGCGAAAGCGTGGGG |
| P-asv5631 | GCAAGCGTTGTCCAGAATCACTGGGCGTAAAGCGCGCGCAGGCGGCCATTTGCGTCCGAGGTGAAAGCCCAAGGCTCAACCTTGGGATTGCCTTGGATACGGGATGGCTTGAGGATCGGAGAGGCAAGGGGAATTCCACGTGTAGCGGTGAAATGCGTAGAGATGTGGAGGAACACCTGTGGCGAAGGCGCCTTGCTGGCCGATTTCTGACGCTGAGGCGCGAAAGCGTGGGG |
| P-asv5632 | GCGAACGTTGTTCGGAATCACTGGGCTTAAAGGGCGCGTAGGCGGCCTGGTCAGTCAGGGGTGAAATCTTTCGGCTCAACCGGAAAAGTGCCTTTGATACTACCAGGCTTGAGGGAGATAGGGGCGTGTGGAACTTCCAGTGGAGCGGTGAAATGCGTTGATATTGGAAGGAACGCCGGTGGCGAAAGCGACACGCTGGGTCTCTTCTGACGCTGAGGCGCGAAAGCTAGGGG |
| P-asv5633 | GCGAGCGTTATCCGGAATTATTGGGCGTAAAGCGCGCGCAGGCGGCCGATTAAGTCAGGTGTGAAAGGCTGCGGCTCAACCGCAGAGCGGCATCTGAAACTGGTCGGCTTGAGTGCAGGAGAGGGGAGCGGAATTCCCGGTGTAGCGGTGGAATGCGTAGAGATCGGGAGGAACACCAGTGGCGAAGGCGGCTCTCTGGCCTGTAACTGACGCTGAGGCGCGAAAGCGTGGGG |
| P-asv5634 | GCGAGCGTTGTCCGGAATTATTGGGCGTAAAGGGCTCGTAGGCGGTTTGTCGCGTCGGGAGTGAAAACACCGGGCTTAACTCGGTGCTTGCTTTCGATACGGGCAGACTAGAGGTATGCAGGGGAGAACGGAATTCCTGGTGCAGCGGTGAAATGCGCAGATACCAGGAGGAACACCGGTGGCGAAGGCGGTTCTCTGGGCATGTCCTGACGCTGAGGAGCGAAAGCATGGGG |
| P-asv5635 | GCTAGCGTTGTTCGGAATTACTGGGCGTAAAGCGCACGTAGGCGGCTATTCAAGTCAGAGGTGAAAGCCCGGGGCTCAACCCCGGAACTGCCTTTGAAACTAGATAGCTTGAATCCAGGAGAGGTGAGTGGAATTCCGAGTGTAGAGGTGAAATGCGTAGAGATCTGAGGAACATCAGTGGCGAAGGCGGCCACCTGGCCCAATACTGACGCTGAGGCACGAAAGCGTGGGG |
| P-asv5636 | GCAAGCGTTGTTCGGAGTTACTGGGCGTAAAGAGTCTGTAGGTGGTCTGTCAAGTCTTTGGTGAAAGGCCGTGGCTTAACCATGGGAATGCCAAAGAGACTGGCAGACTGGAGGCTGGGAGAGGGAAGCGGAATTTCTGGTGTAGCTGTGAAATGCGTAGATATCAGAAGGAAGGCCGGTGGCGAAGGCGGCTTCCTGGAACAGACCTGACACTGAGAGACGAAAGCGTGGGG |
| P-asv5637 | GCTAGCGTTGTTCGGAATTATTGGGCGTAAAGCGAGTGCAGACGGTTACTTAAGCCAGATGTGAAATCCCTAAGCTTAACTTGGGACGTGCATTTGTGACTGCACGACTAGAGTGTGTCAGAGGGGGGTGGAATTCCACGTGTAGCAGTGAAATGCGTAGATATGTGGAGGAACACCAATGGCGAAGGCAGCCCCCTGGGATAACACTGACGCTCATGCACGAAAGCGTGGGG |
| P-asv5638 | CCAAGCGTTATCCGGATTTATTGGGTTTAAAGGGTGCGTAGGCGGCAGATTAAGTTAGAGGTGAAATTCCGAGGCTTAACCTTGGCACTGCCTCTGATACTGGTTTGCTAGCGAGTAGATGCTGTAGGCGGAATGTGTGGTGTAGCGGTGAAATGCATAGAGATCACACAGAACACCGATTGCGAAGGCAGCTTACAAATCTATATCGGACGCTGAGGCACGAAAGCGTGGGG |
| P-asv5639 | CCGAGCGTTAGGCGGAATCACTGGGCTTAAAGCGTGTGTAGGCGGACCCCTAAGTGTCTTGTGAAATCCCACGGCTCACCCGTGGAACTGCTGGGCATACTGGGGGTCTTGGGCCGTTCAGGGGCGACTGGAACAAGCGGTGGAGCGGTGAAATGCGTAGATATCGTGAGGAACACCAGAGGCGAAGGCGGTTACCTGGACAATTACTGACGCTCAGGCACGAAAGCATGGGG |
| P-asv5640 | GCTAGCGTTGTTCGGAATTACTGGGCGTAAAGCGCGCGCAGGCGGTTCTTCAAGTCAGGGGTGAAAGCCCAGAGCTCAACTCCGGAACTGCCATTGAAACTGTGAAACTTGAGGACGAGAGAGGTGAGTGGAATTCCCAGTGTAGAGGTGAAATTCGTAGATATTGGGAAGAACACCGGTGGCGAAGGCGGCTCACTGGCTCGTTTCTGACGCTCAGGCGCGACAGCGTGGGG |
| P-asv5641 | GCGAGCGTTAATCGGAATTACTGGGCGTAAAGCGTGCGCAGGCGGTTCGCCGTGTCCGATGTGAAAGCCCCGGGCTTAACCTGGGACTGGCATTGGAAACTGGCGGGCTGGAGTATGGCAGAGGGGGGTGGAATTCCGCGTGTAGCAGTGAAATGCGTAGAGATGCGGAGGAACACCGATGGCGAAGGCAGCCCCCTGGGCCAATACTGACGCTCAGGCACGAAAGCGTGGGG |
| P-asv5642 | GCAAGCGTTGTTCGGAATTACTGGGCGTAAAGGGTTCGTAGGTGGCCTGCTAAGTCAGACGTGAAATCCCTCAGCTTAACTGGGGAACTGCGTCTGAGACTGGTGGGCTTGAGGGCAGGAGAGGAACGTGGAATTCCAGGTGTAGCGGTGAAATGCGTAGATATCTGGAGGAACACCGGTGGCGAAGGCGGCGTTCTGGACTGCATCTGACACTGAGGAACGAAAGCCAGGGG |
| P-asv5643 | GCAAGCGTTGTCCGGATTTATTGGGCGTAAAGAGCTCGTAGGCGGCCTGTCGCGTCCGCTGCGAAATCCCAGGGCTTAACCCTGGGCCTGCAGTGGATACGGGCTGGCTGGAGGCAGGCAGGGGAGAACGGAATTCCCGGTGTAGCGGTGAAATGCGCAGATATCGGGAGGAACACCGGTGGCGAAGGCGGTTCTCTGGGCCTGTTCTGACGCTGAGGAGCGAAAGCGTGGGG |
| P-asv5644 | GCGAGCGTTACTCGGAATTACTGGGCGTAAAGCGTGCGTAGGTGGTGCGTTAAGTCTGTCGTGAAAGCCCCGGGCTCAACCTGGGAATGGCGATGGATACTGGCGCGCTAGAGTGCGGTAGAGGAGAGTGGAATTCCCGGTGTAGCAGTGAAATGCGTAGAGATCGGGAGGAACATCAGTTGCGAAGGCGGCTCTCTGGACCAGCACTGACACTGAGGCACGAAAGCGTGGGG |
| P-asv5645 | GCGAACGTTGTTCGGAATCACTGGGCTTAAAGGGCGCGTAGGCGGCCGCTCAAGTCGGGGGTGAAATCGTTCCGCTTAACGGGACAAGAGCTTTCGATACTGGGCGGCTGGAGGGAGGTAGGGGTGCGCGGAACTTCCGGTGGAGCGGTGAAATGCGTAGATATCGGAAGGAACGCCGGTGGCGAAAGCGGCGCACTGGACTTCCACTGACGCTGAGGCGCGAAAGCCAGGGG |
| P-asv5646 | GCAAGCGTTGTTCGGAATTACTGGGCGTAAAGGGCTCGTAGGCGGCCAACTAAGTCAGACGTGAAATCCCTCGGCTTAACCGGGGAACTGCGTCTGATACTGGATGGCTTGAATCCGGGAGAGGGATGCGGAATTCCAGGTGTAGCGGTGAAATGCGTAGATATCTGGAGGAACACCGGTGGCGAAGGCGGCATCCTGGACCGGTATTGACACTGAGGAGCGAAAGCCAGGGG |
| P-asv5647 | GCGAGCGTTGTCCGGAATCACTGGGCGTAAAGGGCGCGTAGGTGGCTTTTCAAGCGCGCGGTGAAAGCCCGGGGCTCAACCCCGGGTCGGCCGTGCGGACTGGGAGGCTTGAGCACTGTAGGGGCGGGTGGAATTCCGGGTGTAGCGGTGGAATGCGTAGAGATCCGGGAGAACACCGGTGGCGAAGGCGGTCGACTGGGCCAGAACTGACGCTGAGGCGCGACAGCGTGGGG |
| P-asv5648 | GCAAGCGTTATCCGGATTTACTGGGTGTAAAGGGAGCGCAGACGGCAGGGCAAGTCTGATGTGAAAGTCCGGGGTTCAACCCCGGGACTGCATTGGAAACTGTCCGGCTAGAGTGCAGGAGAGGTAAGTGGAATTCCTAGTGTAGCGGTGAAATGCGTAGATATTAGGAGGAACACCAGTGGCGAAGGCGGCTTACTGGACTGTAACTGACGTTGAGGCTCGAAAGCGTGGGG |
| P-asv5649 | GCAAGCGTTGTTCGGAATTACTGGGCGTAAAGGGCCCGTAGGCGGCCAACTAAGTCCGACGTGAAATCCCCAGGCTTAACCTGGGAACTGCGTCGGATACTGGATGGCTTGAATCCGGGAGAGGGATGCAGAATTCCAGGTGTAGCGGTGAAATGCGTAGATATCTGGAGGAATACCGGTGGCGAAGGCGGCATCCTGGACCGGCATTGACGCTGAACGGCGAAAGCTAGGGG |
| P-asv5650 | GCAAGCGTTGTTCGGAATTACTGGGCGTAAAGAGTGTGTAGGCGGTGTTCTAAGTCTGTCGTGAAATCTCCCGGCTTAACTGGGAGGGTGCGGCGGAAACTGGAACGCTAGAGGGTGGGAGAGGAAAGCGGAATTCCTGGTGTAGCGGTGAAATGCGTAGATATCAGGAGGAACACCTGCGGTGTAGACAGCTTTCTGGACCACTTCTGACGCTGAGACACGAAAGCGTGGGT |
| P-asv5651 | GCAAGCGTTGTTCGGAATTACTGGGCGTAAAGGGCGCGTAGGCGGTTCGTTAAGTCCTGTGTGAAATCCCCCGGCTCAACTGGGGAACTGCACAGGAAACTAGCGGGCTTGAGTTCGGGAGAGGGAAGCGGAATTCCAGGTGTAGCGGTGAAATGCGTAGATATCCGGAGGAACACCGGTGGCGAAGGCGGCTTCCTGGACCGACACTGACGCTGAGGCGCGAAAGCTAGGGG |
| P-asv5652 | GCTAGCGTTGTTCGGATTTACTGGGCGTAAAGCGCGCGTAGGCGGATTGTTAAGTCAGGGGTGAAATGCCGAGGCTCAACCTCGGAACTGCCTTTGATACTGGCAATCTGGAGTTCGGGAGAGGTGAGTGGAATTCCGAGTGTAGAGGTGAAATTCGTAGATATTCGGAGGAACACCAGAAGCGAAGGCGGCTCACTGGCCCGATACTGACGCTGAGGCGCGAAAGCGTGGGG |
| P-asv5653 | GCAAGCGTTAATCGGAATTACTGGGCGTAAAGCGTGCGCAGGCGGTTCTGTAAGATAGATGTGAAATCCCCGGGCTCAACCTGGGAATTGCATATATGACTGCAGGACTTGAGTTTGTCAGAGGAGGGTGGAATTCCACGTGTAGCAGTGAAATGCGTAGATATGTGGAAGAACACCGATGGCGAAGGCAGCCCTCTGGGACATGACTGACGCTCATGCACGAAAGCGTGGGG |
| P-asv5654 | GCGAGCGTTGTCCGGATTCATTGGGCGTAAAGCGCTCGCAGGCGGCCGGCCGCGTCTTGCGTGAAGGCCCCCGGCTCAACCGGGGGTGGTCGTAAGATACGGGTCGGCTGGAGGTAGGTAGGGGGACGCGGAATTCCCGGTGTAGTGGTGGAATGCGTAGAGATCGGGAGGAACACCGGCGGCGAAGGCGGCGTCCTGGACCTGGCCTGACGCTCAGGAGCGAAAGCGTGGGG |
| P-asv5655 | GCTAGCGTTGCTCGGAATCACTGGGCGTAAAGGGCGCGTAGGCGGGTCTTTAAGTCAGGGGTGAAATCCTGGAGCTCAACTCCAGAACTGCCTTTGGTACTGAGAAGCTCGAGTCCGGGAGAGGTGAGTGGAACTGCGAGTGTAGAGGTGAAATTCGTAGATATTCGCAAGAACACCAGTGGCGAAGGCGGCTCACTGGCCCGGTACTGACGCTGAGGTGCGAAAGCGTGGGG |
| P-asv5656 | GCAAGCGTTGTTCGGAATCACTGGGCGTAAAGGGCGTGTAGGCGGTCTGCTAAGTCCGGTGTGAAATCCCCCCGCTTAACGGGGGAACGGCGCCGGAAACTGGCAGACTAGAGTACCGGAGAGGAAGGTGGAATTCCAGGTGTAGCGGTGAAATGCGTAGATATCTGGAGGAACACCGGTGCGAAGGCGGCATCCTGGACCAACACTGACGCTGAGGAGCGAAAGCCAGGGG |
| P-asv5657 | GCGAGCGTTAATCGGAATCACTGGGCGTAAAGCGTGCGCAGGCGGTTCGTCGTGTCCGATGTGAAAGCCCCGGGCTCAACCTGGGAATGGCATTGGAAACTGGCGGGCTTGAGTGTGGCAGAGGGGGGTGGAATTCCGCGTGTAGCAGTGAAATGCGTAGAGATGCGGAGGAACACCGATGGCGAAGGCAGCCCCCTGGGCCAACACTGACGCTCAGGCACGAAAGCGTGGGG |
| P-asv5658 | GCGAGCGTTGTCCGGATTCATTGGGCGTAAAGCGCGCGCAGGCGGCTTCGCAAGTCGGTCGTGAAATCTCCCGGCTCAACCGGGAGGGGTCGGGCGAGACTGCGGGGCTCGAGGTGGGCAGAGGGCAGTGGAATTCCGGGTGTAGTGGTGGAATGCGTAGAGATCCGGAGGAACACCAGTGGCGAAGGCGGCTGCCTGGGCCTGGCCTGACGCTGAGGCGCGAAAGCGTGGGG |
| P-asv5659 | CCAAGCGTTGTTCGGATTCATTGGGCGTAAAGCGCGCGCAGGCGGTTGCGAGAGCCCGGCGTGAAAGCCCCCAGCTCAACTGGGGGAGGCCGTCGGAGACCTTGCGACTGGAGGCCGGTAGGGGAGAGCGGAATTCCCGGTGTAGTGGTGGAATGCGTAGATATCGGGAGGAACACCAGTGGCGAAGGCGGCTCTCTGGACCGGTTCTGACGCTCAGGCGCGAAAGCGTGGGG |

| | |
|---|---|
| P-asv5660 | GCTAGCGTTATTCGGATTCACTGGGCATAAAGCGCGCGTAGGCGGCTCTTCAAGTCGGTGGTGAAAGCCCGCAGCTCAACTGCGGAACAGCCTTCGATACTGTCGAGCTTGAGTGCAGGAGGGGTGATCGGAACTCCAGGTGTAGCGGTGAAATGCGTAGATATCTGGAGGAACACCGGAGGCGAAGGCGGATCACTGGACTGCCACTGACGCTGAGGCGCGAAAGCTAGGGG |
| P-asv5661 | GCGAGCGTTGTCCGGAATTATTGGGCGTAAAGAGCTCGTAGGCGGTGTGTCGCGTCGGCCGTGAAAACTTTCAGCTTAACTGGGAGCTTGCGGTCGATACGGGCATCACTGGAGTTCGGCAGGGGAGACTGGAATTCCCGGTGTAGTGGTGAAATGCGTAGAGATCGGGAGGAACACCCGCGGCGAAGGCGGCCACCTGGACCAGACCTGACGCTGAGGCGCGAAGGCCGGGGG |
| P-asv5662 | GCCAGCGTTGCTCGGAATTACTGGGCGTAAAGGGTCCGTAGGCGGTGCGGTAAGTTGGGAGTGAAATCTCTGGGCTTAACCTAGAGGCTGCTTCCAAAACTACCGTGCTAGAGTGCGAGAGAGGCAAGTGGAATTGCGGGTGTAGCGGTGAAATGCGTAGATATCCGCAGGAACATCCGAGGCGAAAGCGGGTTGCTGGATCGCAACTGACGCTGAGGGACGAAAGCTAGGGG |
| P-asv5663 | GCAAGCGTTACCCGGAATCACTGGGCGTAAAGGGCGTGTAGGCGGACCGCTAAGTCTGGCTTTGAAGACTGGGGCTCAACCCCAGGCATGGGCTGGGTACTGGAGGTCTGGACCTCTGGAGAGGGAACCGGAATTCCTGGTGTAGCGGTGGAATGCGTAGATACCAGGAGGAACACCGATGGCGAAGGCAGGTTCCTGGACAGCAGGTGACGCTGAGGCGCGAAAGTGTGGGG |
| P-asv5664 | GCGAACGTTAATCGGAATTACTGGGCGTAAAGCGCGCGTAGGCGGCTTGTCAAGTCGGATGTGAAAGCCCCGGGCTCAACCCGGGAACTGCATTCGAAACTGGCAGGCTCGAGTGCAGTAGAGGGAGGTGGAATTCCAGGTGTAGCGGTGAAATGCGTAGATATCTGGAGGAACACCAGTGGCGAAGGCGGCCTCCTGGACTGACACTGACGCTGAGGTGCGAAAGCGTGGGG |
| P-asv5665 | GCTAGCGTTGCTCGGAATTACTGGGCGTAAAGGGCGCGTAGGCGGATCGTTAAGTCAGAGGTGAAATCCCAGGGCTCAACCCTGGAACTGCCTTTGATACTGGCGATCTTGAGTATGAGAGAGGTATGTGGAACTCCGAGTATAGAGGTGAAATTCGTAGATATTCAGAAGAACACCAGTGGCGAAGGCGACATACTGGCTCATTACTGACGCTGAGGCGCGAAAGCGTGGGG |
| P-asv5666 | GCAAGCGTTGTCCGGAATTATTGGGCGTAAAGAGCTCGTAGGCGGTTTGTCGCGTCTGCTGTGAAAGCCCGGGGCTTAACCCCGGGTGTGCAGTGGGTACGGGCAGACTAGAGTGCAGTAGGGGAGACTGGAATTCCTGGTGTAGCGGTGGAATGCGCAGATATCAGGAGGAACACCGATGGCGAAGGCAGGTCTCTGGGCTGTAACTGACGCTGAGGAGCGAAAGCATGGGG |
| P-asv5667 | GCGAACGTTATTCGGAATCACTGGGCTTAAAGAGCGCGTAGGCGGATTGCTACGTCGGTGTCTGAAATCCCCCGGCTCAACCGGGGAAGTGGATCCGATACGGGCAGTCTTGAGGGACGTAGGGGGGGCTGGAACTTCCGGTGGAGCGGTGAAATGCGTTGAGATCGGAAGGAACGCCCGTGGCGAAAGCGAGCCCCTGGACGTCTGCTGACGCTGAGGCGCGAAAGCTAGGGG |
| P-asv5668 | GCAAGCGTTAATCGGAATTACTGGGCGTAAAGGGTTCGTAGGCGGTTTATTAAGTCAGATGTGAAAGCCCAGGGCTCAACCTTGGAACTGCATTTGATACTGGTAGACTAGAGTATAGTAGAGGAGTGGGGAATTTCTGGTGTAGCGGTGAAATGCGTAGAGATCAGAAGGAACACCAATGGCGAAGGCAACACTCTGGACTAATACTGACGCTGAGGGACGAAAGCGTGGGG |
| P-asv5669 | GCGAGCGTTAATCGGATTTACTGGGCGTAAAGCGTGTGCAGGTGGTTCACTGCGTCGGTTGTGAAATCCCCGGGCTCAACCTGGGAATGGCTTCCGAAACGGGTGGACTCGAGTACGGTAGAGGGTGGTGGAATTCCCGGTGTAGCGGTGAAATGCGTAGAGATCGGGAGGAACATCAATGGCGAAGGCAGCCGCCTGGGCCTGTACTGACACTGAGACACGAAAGCGTGGGG |
| P-asv5670 | GCAAGCGTTAATCGGAATTACTGGGCGTAAAGCGCACGCAGGCGGTCTGTCAAGTCGGATGTGAAATCCCCGGGCTCAACCTGGGGATTGCATTTAAAACTGTCCAGCTAGAGTCTTGTAGAGGGGGGTAGAATTCCAGGTGTAGCGGTGAAATGCGTAGAGATCTGGAGGAATACCGGTGGCGAAGGCGGCCCCCTGGACAAAGACTGACGCTCAGGTGCGAAAGCGTGGGG |
| P-asv5671 | GCGAGCGTTACCCGGATTTACTGGGCGTAAAGGGCGTGTAGGCGGCCTGGGGCGTCCCATGTGAAAGGCCACGGCTCAACCGTGGAGGAGCGTGGGATACGCTCAGGCTAGAGGGTGGGAGAGGGTGGTGGAATTCCCGGAGTAGCGGTGAAATGCGCAGATACCGGGAGGAACGCCGATGGCGAAGGCAGCCACCTGGTCCACTTCTGACGCTGAGGCGCGAAAGCGTGGGG |
| P-asv5672 | GCGAGCGTTGTCCGGATTCACTGGGCGTAAAGCGCGCGCAGACGGCTTTCTGCGTCCGGGGTGAAAACCCGGGGCTCAACCCCGGGAGGGCCTTGGATACGGGAGAGCTTGAGGGCCGGAGAGGCAAGGGGAATTCCACGTGTAGCGGTGAAATGCGTAGAGATGTGGAGGAACACCTGTGGCGAAGGCGCCTTGCTGGCCGGTTCCTGACGTTGAGGCGCGAAAGCGTGGGG |
| P-asv5673 | GCGAGCGTTAATCGGAATTACTGGGCGTAAAGGGCGCGTAGGCGGTGGGTTACGTCTGCCGTGAAATCCCCGGGCTCAACCTGGGAATGGCGGTGGAAACGGGCTGACTGGAGTATGGGAGAGGGTGATGGAATTCCAGGTGTAGCGGTGAAATGCGTAGAGATCTGGAGGAACACCAGTGGCGAAGGCGGTCGCCTGGCCCAATACTGACGCTGAGGCGCGAAAGCGTGGGG |
| P-asv5674 | TCCTTCTTTCTCCTTCCTCCCTTTTCTTAAAGGGTGCGTAGGCGGCCTAACAAGTCAGAGGCGAAAGGCATTGGCTCAACCAATGTAAGCCTTTGAAACTGTTGGGCTTGAGTGCAGGAGAGGTAAGTGGAATTCCTAGTGTAGAGGTGAAATTCGTAGATATTAGGAGGAACACCAGTGGCGAAGGCGACTTACTGGACTGTAACTGACGCTGAGGCACGAAAGCGTGGGG |
| P-asv5675 | GCAAGCGTTGTTCGGAATTACTGGGCGTAAAGAGTGTGTAGGCGGTTGGGTAAGTTTGGTGTGAAATCTCCCGGCTTAACTGGGAGGGTGCGCCGAAAACTGCCTGGCTAGAGTGCCGGAGAGGAAAGCGGAATTCCTGGTGTAGCGGTGAAATGCGTAGATATCAGGAGGAACACCGGTGGTGTAGACGGCTTTCTGGACGGCCACTGACGCTGAGACACGAAAGCGTGGGT |
| P-asv5676 | GCAAGTGTTATCCGGAATCACTGGGCGTAAAGCGTCTGTAGGTGGTTGAATAAGTCAACTGTTAAATCTTGAGGCTCAACTTCAAAATCGCAGTCGAAACTATTCGACTAGAGTATAGTAGGGGTAAAGGGAATTTCCAGTGGAGCGGTGAAATGCGTAGAGATTGGAAAGAACACCGATGGCGAAGGCACTTTACTGGGCTATTACTAACACTGAGAGACGAAAGCTAGGGT |
| P-asv5677 | GCGAGCGTTGTCCAGATTTATTGGGCGTAAAGGGCTCGTAGGTGGTTGATCGCGTCGGAAGTGTAATCTTGGGGCTTAACCCTGAGCGTGCTTTCGATACGGGTTGACTTGAGGAAGGTAGGGGAGAATGGAATTCCTGGTGGAGCGGTGGAATGCGCAGATATCAGGAGGAACACCAGTGGCGAAGGCGGTTCTCTGGGCCTTTCCTGACGCTGAGGAGCGAAAGCGTGGGG |
| P-asv5678 | CCAAGCGTTAATCGGAATTACTGGGCGTAAAGCGTACGCAGGCGGTTCATTAAGCCAGATGTGAAATCCCCGGGCTCAACCTGGGAATTGCATTTGGAACTGGTGAACTAGAGTCTTGTAGAGGGGGGTAGAATTTCAGGTGTAGCGGTGAAATGCGTAGAGATCTGAAGGAATACCGGTGGCGAAGGCGGCCCCCTGGACAAAGACTGACGCTCATGTACGAAAGCGTGGGG |
| P-asv5679 | GCAAGCGTTGTCCGGAATTATTGGGCGTAAAGCGCGCGCAGGCGGCATTGTAAGTCTTGTGTTTAAACGCGGGGCTCAACCCCGTGTCGCATGGGAAACTGCAAGGCTTGAGTGCAGAAGAGGAAAGTGGAATTCCACGTGTAGCAGTGAAATGCGCAGATATGCGGAGGAACACCGATGGCGAAGGCAATCCCCTGGGCCTGTACTGACGCTCATGCACGAAAGCGTGGGG |
| P-asv5680 | GCAAGCGTTATCCGGATTTATTAGGTTTAAAGGGTCCGTAGGCGGAAATTTAAGTCAGTGGTGAAAGCCCATAGCTCAACTATGGAACTGCCATTGATACTGAATTTCTTGAATATGATTGAAGTGGGCGGAATATGTCATGTAGCGGTGAAATGCTTAGATATGACATAGAACACCGATAGCGAAGGCAGCTCACTAAGTCATTATTGACGCTGAGGGACGAAAGCGTGGGG |
| P-asv5681 | GCGAGCGTTATCCGGAATCATTGGGTTTAAAGGGTCCGTAGGCGGGCTAATAAGTCAGAGGTGAAAGCGCTCAGCTCAACTGAGCAACTGCCTTTGAAACTGTTAGTCTTGAATGGTTGTGAAGTAGTAGGAATGTGTAGTGTAGCGGTGAAATGCTTAGATATTACACAGAACACCGATAGCGAAGGCATATTACTAACAATTAATTGACGCTGATGGACGAAAGCGTGGGG |
| P-asv5682 | GCTAGCGTTGTTCGGAATTACTGGGCGTAAAGCGCACGTAGGCGGCTTTGTAAGTTAGGGGTGAAAGCCTGGTGCTCAACACCAGAACTGCCTTTGATACTGCATGGCTTGAATCCGAGAGAGGTGAGTGGAATTCCGAGTGTAGAGGTGAAATTCGTAGATATTCGGAAGAACACCAGTGGCGAAGGCGGCTCACTGGCTCGGTATTGACGCTGAGGTGCGAAAGCGTGGGG |
| P-asv5683 | GCAAGCGTTAATCGGATTTACTGGGCGTAAAGCGTGCGTAGGCGGCATGTTAAGTCGGATGTGAAATCCCAGGGCTTAACCTTGGAACTGCATCCGATACTGGCAAGCTAGAGATCGGTAGAGGTTGGTGGAATTTCCAGTGTAGCGGTGAAATGCGTAGATATTGGAAGGAACATCGATGGCGAAGGCAGCCAACTGGGCCTGATCTGACGCTGAGGTACGAAAGCGTGGGG |
| P-asv5684 | GCTAGCGTTATTCGGAATTACTGGGCGTAAAGCGCACGTAGGCGGACTGGAAAGTCAGAGGTGAAAGCCCAGGGCTCAACCCTGGAACTGCCTTTGAAACTCCCAGTCTTGAGGTCGAGAGAGGTGAGTGGAATTCCGAGTGTAGAGGTGAAATTCGTAGATATTCGGAGGAACACCAGTGGCGAAGGCGGCTCACTGGCTCGATACTGACGCTGAGGTGCGAAAGCGTGGGG |
| P-asv5685 | CCGAGCGTTATCCGGAATCATTGGGTTTAAAGGGTCCGCAGGCGGTCAATTAAGTCAGAGGTGAAATCCCATAGCTCAACTATGGAACTGCCTTTGATACTGGTTGACTTGAGTCATATGGAAGTAGATAGAATGTGTAGTGTAGCGGTGAAATGCATAGATATTACACAGAATACCGATTGCGAAGGCAGTCTACTACGTATGTACTGACGCTGAGGGACGAAAGCGTGGGG |
| P-asv5686 | CCGAGCGTTATCCGGATTTATTGGGTTTAAAGGGAGCGTAGGCGGACGCTTAAGTCAGTTGTGAAAGTTTGCGGCTCAACCGTAAAATTGCAGTTGATACTGGGTGTCTTGAGTACAGTAGAGGCAGGCGGAATTCCTGGTGTAGCGGTGAAATGCTTAGATATCACGAAGAACTCCGATTGCGAAGGCAGCCTGCTGGACTGTAACTGACGCTGATGCTCGAAAGTGTGGGT |
| P-asv5687 | GCAAGCGTTATTCGGAATCATTGGGCGTACAGGGTGTGTAGGCGGCATGTTAAGTCTGTTGTTAAAGACTCCGGCCTAACCGGAGATGGGCAGCGGAAACTGGCATACTAGAGGTTGGAAGAGAGAAGCGGAATTCTCGGTGTAGCGGTAAAATGCGTAGATATCGAGAGGAACACCGATGGCGAAGGCAGCTTCTTGGTCCATACCTGACGCTGAAACACGAAAGCGTGGGG |
| P-asv5688 | GCTAGCGTTGCTCGGAATTACTGGGCGTAAAGGGCGCGTAGGCGGACGTTTAAGTCGGGGGTGAAAGCCCAGGGCTCAACCCTGGAATTGCCTTCGATACTGGACGTCTCGAGACCGGTAGAGGTAAGCGGAACTGCGAGTGTAGAGGTGAAATTCGTAGATATTCGCAAGAACACCAGTGGCGAAGGCGGCTTACTGGACCGGTTCTGACGCTGAGGCGCGAAAGCGTGGGG |
| P-asv5689 | TCAAGTGATGACCGATTTTATTGGGCCTAAAGCGTCCGTAGCTGGCCGCGCAAGTCCGTCGGGAAATCCACCCGCTTAACGGGTGGGCGTCCGGCGGAAACTGTGCGGCTTGGGACAGGAAGGCTCGAGGGGTACGTTCGGGGTAGGAGTGAAAACCCGTAATCCTGGACGGACCACCGATGGGGAAACCACCTCAAGAGAACGGATCCGACAGTGAGGGACGAAAGCTAGGGT |
| P-asv5690 | GCTAGCGTTGCTCGGAATCACTGGGCGTAAAGGGCGCGTAGGCGGCTGATTTAGTCGAGGGTGAAAGCCCGTGGCTCAACCACGGAATGGCCTTCGATACTGGTTGGCTTGAGACCGGAAGAGGACAGCGGAACTGCGAGTGTAGAGGTGAAATTCGTAGATATTCGCAAGGACACCAGTGGCGAAGGCGGCTGTCTGGTCCGGTTCTGACGCTGAGGCGCGAAAGCGTGGGG |
| P-asv5691 | GCAAGCGTTAATCGGAATTACTGGGCGTAAAGCGCACGCGGGCGGTTTGTTAAGTCAGATGTGAAATCCCCGGGCTCAACCTGGGAACTGCATCTGATACTGGCAAGCTTGAGTCTCGTAGAGGGGGGTAGAATTCCAGGTGTAGCGGTGAAATGCGTAGAGATCTGGAGGAATACCGGTGGCGAAGGCGGCCCCCTGGACGAAGACTGACGCTCAGGTGCGAAAGCGTGGGG |
| P-asv5692 | GCGAGCGTTGTCCGGAATCATTGGGCGTAAAGAGCGTGTAGGCGGCCCGGTAAGTCTGCTGTGAAAACCCGGGGCTCAACCCCGGGCGTGCAGTGGAAACTGCCGGGCTAGAGGGCGGCAGAGGCGAGTGGAATTCCCGGTGTAGCGGTGAAATGCGCAGATATCGGGAGGAACACCAGTAGCGAAGGCGGCTCGCTGGGCCGTTCCTGACGCTGAGACGCGAAAGCTAGGGG |
| P-asv5693 | GCGAGCGTTGTCCGGATTTATTGGGCGTAAAGAGTTCGTAGGTGGTTTGTTAAGTTTGATGTTAAAGATCGGGGCTCAACCTCGGGACAGCATTGAATACTGGCAGACTTGAGTATGGTAGAGGCTAGTGGAATTCCCAGTGTAGCGGTGAAATGCGTAGATATTGGGAAGAACACCGGTGGCGTAGGCGACTAGCTGGGCCATAACTGACGCTGAGGAACGAAAGCCAGGGG |
| P-asv5694 | GCTAACGTTATTCGGATTTACTGGGCGTAAAGGGTGCGTAGACGGCAGGATAAGTCAACTGTTAAATTTTCCGGCCTAACCGGAAGTCGGCGGTAGAAACTATTTTGCTAGAGGGTGGAAGAGAGAAGTGGAATTCTCGGAGTAGCGGTAAAATGCGTAGATATCGAGAGGAACACCGATGGCGAAGGCAGCTTCTTGGTCCACACCTGACGTTGAGGCACGAAAGCGTGGGT |
| P-asv5695 | GCAAGCGTTATCCGGAATTATTGGGCGTAAAGAGCGCGCAGGTGGTTGATTAAGTCTGATGTGAAAGCCCACGGCTTAACCGTGGAGGGTCATTGGAAACTGGTCGACTTGAGTGCAGAAGAGGGAAGTGGAATTCCATGTGTAGCGGTGAAATGCGTAGAGATATGGAGGAACACCAGTGGCGAAGGCGGCTTCCTGGTCTGTAACTGACACTGAGGCGCGAAAGCGTGGGG |
| P-asv5696 | GCAAGCATTGGTCGGAATTACTGGGCGTAAAGGGCGCGTAGGCGGCACGATAAGTCGGATGTGAAATTCCGGGGCTCAACCCCGGAGCTGCATTTGAAACTATCGAGCTAGAGTTCAGTTAGGGTAAACGGAATTCCACGTGTAGCGGTGAAATGCGTAGATATGTGGAAGAACACCGGTGGTGAAAACGGTTTACTGGGCTGAAACTGACGCTGAGGCGCGAAAGCATGGGG |
| P-asv5697 | GCTAGCGTTGCTCGGAATTACTGGGCGTAAAGGGAGCGTAGGCGGACTGTTTAGTCAGAGGTGAAAGCCCAGGGCTCAACCTTGGAATTGCCTTTGATACTGGCAGTCTTGAGTATGGGAGAGGTGAGTGGAACTCCGAGTGTGGAGGTGAAATTCGTAGATATTAGGAGGAACACCAGTGGCGAAGGCGACTTACTGGACTGTAACTGACGCTGAGGCACGAAAGCGTGGGG |
| P-asv5698 | GCGAGCGTTGTTCGGAATTACTGGGCGTAAAGCGCGCGCAGGCGGCGGGGCAAGTCGGACGTGAAAGCCCTCGGCTCAACCGAGGAACTGCGTACGATACTGCTTCGCTTGAGTCCGGTAGGGGGTAGCGGAATTCCCGGTGTAGCGGTGGAATGCGTAGATATCGGGAGGAACACCGATGGCGAAGGCGGCTACCTGGGCCGGTTCTGACGCTGAGGCGCGAAAGCTAGGGG |

| | |
|---|---|
| P-asv5699 | GCAAGCGTTACTCGGAATCACTGGGCGTAAAGGGTGCGTAGGCTGGCTTCTAAGTCAGATGTGAAATCCAATGGCTTAACCATTGAACTGCATTTGAAACTGGGAGCCTAGAGTTCAGAAGGGGCAGATGGAATTAGTGGTGTAGGGGTAAAATCCGTAGATATCACTAGGAATATCAAAAGCGAAGGCGATCTGCTGGGATGATACTGACGCCGAGGCACGAAAGCGTGGGG |
| P-asv5700 | GCGAACGTTAATCGGAATTACTGGGCGTAAAGCGCGCGTAGGTGGTCTTTTAAGTGGGATGTGAAAGCCCAGGGCTCAACCCTGGAATTGCATTCCAAACTGAAAGACTCGAGTACGATAGAGGGAGGTAGAATTCAGAGTGTAGCGGTGAAATGCGTAGATATTCTGAGGAATACCAGTGGCGAAGGCGGCCTCCTGGATTGATACTGACACTGAGGCGCTAAAGCGTGGGG |
| P-asv5701 | CCGAGCGTTAATCGGAATCACTGGGCTTAAAGCGTGCGCAGGCGGACCCGTTAGTGTCTTGTGAAATCCCACTGCTCAACGGTGGAATTGCAGGGCATACTGCGGGTCTTGAGGTCGGTAGAGGTTGCCGGAACTGTAGGTGTAGCGGTGAAATGCGTAGATATCTACAGGAACACCCAAGGCGAAGGCAGGCAACTGGGCCGATTCTGACGCTCAGGCACGAAAGCGTGGGG |
| P-asv5702 | GCTAGCGTTGTTCGGAATCACTGGGCGTAAAGGGTCCGTAGGCGGCCATGCAGGTCCTGGGTGAAATCCCTCGGCTTAACCGAGGAATTGCCTGGGAAACCGTTTGGCTTGAGTCCGGTAGAGGGAGATGGAATTCCCGGTGTAGCGGTGAAATGCGTAGATATCGGGAGGAACACCAGTGGCGAAGGCGATCTCCTGGACCGGTTACTGACGCTGAGGGACGAAAGCTAGGGG |
| P-asv5703 | GCAAGCGTTAATCGGAATTACTGGGCGTAAAGCGCACGTAGGCGGCTGTTTAAGCCAGTTGTGAAAGCCCCGGGCTCAACCTGGGAATGGCAGTTGGAACTGGGCAGCTGGAGTACAGTAGAGGGGTGTGGAATTTCAGGTGTAGCGGTGAAATGCGTAGATATCTGAAGGAACATCAGTGGCGAAGGCGACACCCTGGACTGATACTGACGCTCAGGTGCGAAAGCGTGGGG |
| P-asv5704 | GCTAGCGTTGTTCGGAATCACTGGGCGTAAAGGGCGTGTAGGCGGCGGATTTAGTTAGGCGTGAAAGCCCCGGGCTCAACCTGGGAACTGCGCTTAATACTGGTCTGCTAGAGTCCGGGAGAGGCGAGTGGAATATCCAGTGTAGAGGTGAAATTCGTAGATATTGGATGGAACACCGATGGCGAAGGCAGCTCGCTGGACCGGAACTGACGCTGAGGCGCGAAAGCGTGGGG |
| P-asv5705 | GCAAGCGTTGTTCGGAATTACTGGGCGTAAAGCGTGTGTAGGCGGCTTGGTAAGTCGGATGTGAAAGCCCTTGGCTCAACTGAGGAAGTGCATCTGATACTGTCAAGCTTGAGTCCTGGAGAGGGGTGTGGAATTCCCGGTGTAGAGGTGAAATTCGTAGATATCGGGAGGAACACCAGTGGCGAAGGCGACACCCTGGATGGAGACTGACGCTGAGACACGAAAGCGTGGGG |
| P-asv5706 | GCTAGCGTTGTTCGGAATTACTGGGCGTAAAGCGCGCGTAGGCGGCCGGCCAAGTCGGGGGTGAAAGCCCGGGGCTCAACCCCGGAACGGCCTCCGAAACTGGTCGGCTCGAGGACGGGAGAGGAGAGTGGAATTCCCAGTGTAGAGGTGAAATTCGTAGATATTGGGAAGAACACCGGTGGCGAAGGCGGCTCTCTGGACCGTATCTGACGCTGAGGCGCGAAAGCGTGGGG |
| P-asv5707 | CCAAGCGTTGTCCGGAATTACTGGGTGTAAAGGGTGCGCAGGCGGGCTTGTGCGTCAGGGGTTAAAGACCACAGCTTAACTGTGGGAGTGCCTCTGATACGGCGAGTCTTGAGTGTGAGAGAGGAAAGTGGAATTCCTGGTGTAGCAGTGAAATGCGTAGATATCAGGAGGAACACCAGTGGCGAAGGCGGCTTTCTGGCTCATTACTGACGCTCAGGCACGAAAGCGTGGGG |
| P-asv5708 | GCAAGCGTTAATCGGAATTACTGGGCGTAAAGGGTGCGTAGGCGGCATGACAAGCTAGATGTGAAAGCCCTGGGCTTAACCTGGGAAGTGCGTCTGGGACTGTTGAGCTAGAGTATGGTAGAGGAAAGTGGAATTTCTAGTGTAGCGGTGAAATGCGTAGATATTAGAAGGAACACCAGAGGCGAAGGCGACTTTCTGGACCGATACTGACGCTGAGGCACGAAAGCGTGGGG |
| P-asv5709 | GCAAGCGTTACTCGGAATTACTGGGCGTAAAGCGTGCGTAGGTGGTTTGTTAAGTCTGATGTGAAAGCCCTGGGCTCAACCTGGGAATGGCACTGGATACTGGCAAGCTAGAGTGTGGCAGAGGGTAGTGGAATTCCTGGTGTAGCAGTGAAATGCGTAGAGATCAGGAGGAACATCCGTGGCGAAGGCGACTGCCTGGGCCAACACTGACACTGAGGCACGAAAGCGTGGGG |
| P-asv5710 | GCTAGCGTTGTTCGGAATTACTGGGCGTAAAGCGCACGTAGGCGGGCGATTAAGTCAGGGGTGAAATCCCGGAGCTCAACTCCGGAACTGCCTTTGATACTGGTCGTCTAGAGATCGGAAGAGGTGAGTGGAATTCCCAGTGTAGAGGTGAAATTCGTAGATATTGGGAAGAACACCGGTGGCGAAGGCGGCTCACTGGTCCGATACTGACGCTGAGGTGCGAAAGCGTGGGG |
| P-asv5711 | CCAAGCGTTATCCGGATTTATTGGGCGTAAAGCGTCTGTAGGTTGTTAATCAAGTCTGAAGTGAAAGAATGGGGCTCAACCCCATATACGCTTTGGAAACTGGTTGACTAGAGAATGTTAGAGGTAAGTGGAATTCCTAGTGTAGCGGTGAAATGCGTAGATATTAGGAAGAACACCAGAGGCGAAGGCGACTTACTGGGACATTTCTGACACTGAGAGACGAAAGCATGGGG |
| P-asv5712 | GCAAGCGTTATCCGGATTCATTGGGCGTAAAGCGCGCGTAGGCGGTCTGTTAGGTCAGGAGTTAAATCTGGGGGCTCAACCCCTATCCGCTCCTGATACCGGCAGGCTTGAGTCTGGTAGGGGAAGATGGAATTCCAAGTGTAGCGGTGAAGTGCGCAGATATTTGGAAGAACACCGGTGGCGAAGGCGGTCTTCTGGGCCATGACTGACGCTGAGGCGCGAAAGCTAGGGG |
| P-asv5713 | CCAAGCGTTATTCGGAATCACTGGGCTTAAAGCGTGCGCAGGCGGACCCGTAGGCATCTCGTGAAATCCCACGGCTCAACCGTGGAACTGCGGGGTGAACCACGGGTCTCGAGGCAGGTAGGGGTCAGCGGAACTATGGGTGGAGCGGTGAAATGCGTAGATATCCATGGGAACGCCGATGGCGAAGGCAGCTGACTGGGCCTGTCCTGACGCTCAGGCACGAAAGCGTGGGT |
| P-asv5714 | CCAGGCGTTATCCGGATTTATTGGGTTTAAAGGGAGTGTAGGCGGTTTGTTAAGCGTGTTGTGAAATTTAGGTGCTCAACATTTAACTTGCAGCGCGAACTGTCAGACTTGAGTACACGCAACGTATGCGGAATTCATGGTGTAGCGGTGAAATGCTTAGATATCATGAAGAACTCCGATTGCGAAGGCAGCATACGGGAGTGTAACTGACGCTTAAGCTCGAAGGTGCGGGT |
| P-asv5715 | GCAAGCGTTGTTCGGATTTACTGGGCGTAAAGCGCGCGCAGGCGGATAAAAAAGTCAGGTGTGAAATCTCGGGGCTCAACCCCGAAACTGCGCCTGAAACTCTTTGTCTAGAATCTTGGAGGGGGTAGGGGAATTTCTCATGTAGGGGTAAAATCCGTAGAGATGAGAAGGAACACCGGTGGCGAAAGCGCCTACCTGGACAAGAATTGACGCTGAGGCGCGAAAGCGTGGGG |
| P-asv5716 | GCAAGCGTTATTCGGAATGATTGGGCGTAAAGAGCGTGTAGGCGGTTGAATAAGTTTGATGTTTAATTCTCTGGCTTAACTAGAGGCATGCATTGAATACTGTTCGGCTAGAGTGCGATAGAGAAAAGTGGAATTCTTGGAGTAGCGGTAAAATGCGTAGATCTCAAGAGGAACGCTAATGGCGAAGGCAGCTTTTTGGAGCGCAACTGACGCTGAGACGCGAAGGCGTGGGG |
| P-asv5717 | CCGAGCGTTGTCCGGATTTATTGGGCGTAAAGAGCTCGTAGGCGGTTCCGTAAGTCGGATGTGAAAACTCGGGGCTCAACCCCGAGACGCCATCTGATACTGCGGTGACTAGAGTCCGGTAGGGGAGCATGGAATTCCTGGTGTAGCGGTGGAATGCGCAGATATCAGGAGGAACACCAATGGCGAAGGCAGTGCTCTGGGCCGGTACTGACGCTGAGGAGCGAAAGCATGGGG |
| P-asv5718 | CCTAGCGTTGTTCGGAATCATTGGGCGTAAAGCGGGTGTAGGCGGGTTATTAAGTCAGAAGTGAAAGCCCTGGGCTCAACTCAGGAAGTGCTTTTGAAACTGATGATCTTGAGTGTGGGAGAGGACAGTAGAATTCCTGGTGTAGTGGTGAAATACGTAGATATCAGGAGGAACCCCGGTGGGGAAGCCGGCTGTCTGGCCCAACACTGACGCTCAGACCCGAAAGCGTGGGT |
| P-asv5719 | GCAGGCGTTGTTCAGATTTATTGGGCGTAAAGGGTCCGTAGGCGGTTCGTAAAGTCGGATGTGAAATCCTACAGCTTAACTGTGGAACTGCATTCGAAACTCACGGACTAGAGTGTGAGAGGGGTAAAGGGAATTCCCGGTGTAAGGGTGAAATCTGCAGAGATCGGGAGGAACACCAGTCGCGAAGGCGCTTTACTGGCTCACTACTGACGCTCAGGGACGAAAGCTAGGGG |
| P-asv5720 | GCTAGCGTTGTTCGGAATTACTGGGCGTAAAGCGCGTGTAGGCGGTTGCCCAAGTCGGGTGTGAAAGCCTTGAGCTCAACTCAAGAAATGCACTCGATACTGGGTGACTAGAGGACCGGAGAGGATAGTGGAATTCCCAGTGTAGTGGTGAAATACGTAGAGATTGGGAAGAACACCAGTGGCGAAGGCGGCTATCTGGACGGTTTCTGACGCTAAGACGCGAAAGCGTGGGG |
| P-asv5721 | GCTAGCGTTGTTCGGAATTACTGGGCGTAAAGCGCGCGTAGGCTGTGCCGCAAGTCAGAGGTGAAATCCCGGAGCTTAACTCCGGAACTGCCTTTGAAACTGCGGTGCTGGAATCTCAGAGGGGGTAGCGGAATTCCAAATGTAGGGGTGAAATCCGTAGATATTTGGAGGAACACCGGTGGCGAAGGCGGCTACCTGGATGAGTATTGACACTGAGGCTCGAAAGCGTGGGG |
| P-asv5722 | GCAAGCGTTGCTCGGAATCACTGGGCGTAAAGCGTACGCAGGCGGTCGTTCAAGTCGGGTGTGAAAGCCCGGGGCTCAACCCCGGAATTGCACTCGAAACTGGATGGCTTGAGTACGGGAGAGGGAAGCAGAATTCCTGGTGTAGAGGTGAAATTCGTAGATATCAGGAGGAATACCTGTGGCGAAAGCGGCTTCCTGGCCCGATACTGACGCTCATGTACGAAAGCGTGGGG |
| P-asv5723 | GTTAGCGTTGTTCGGAATTATTGGGCGTAAAGCGTGTCTAGGTGGATTGGTAAGTCAGGTGTGAAAGACCCGAGCTTAACTTGGGAATTGCGCTTGAAACTGCTGGTCTTGAGTAACAAAGAGGTGAGTGGAATTCCTGGTGTAGCAGTGAAATGCGTAGAGATCAGGAAGAACATCGGAGGCGAAGGCGGCTCACTGGTTGTTTACTGACACTGAAGCACGAAAGCGTGGGG |
| P-asv5724 | GTTAGCGTTGTTCGGAATTACTGGGCGTAAAGCGCGCGTAGGCGGGCTGTTAAGTCGGGGGTGAAATCCCGGGGCTCAACCCCGGAACGGCCCTCGATACTGGCAGCCTAGAGTTCGAGAGAGGTGAGTGGAATTCCGAGTGTAGAGGTGAAATTCGTAGATATTCGGAGGAATACCGGTGGCGAAGGCGGCCAACTGGACAGATACTGACGCTGAGACACGAAAGCGTGGGG |
| P-asv5725 | GCAAACGTTGCTCGGAATCATTGGGCGTAAAGCGCACGTAGGCGGCACAGCAAGTTGGATGTGAAAGCCCAGGGCTCAACCTTGGATGTGCATTCAAAACTGCTGAGCTTGAGTTCCGGAGGGGCCCAGAGAATTCCCGGTGTAGAGGTGAAATTCGTAGATATCGGGAGGAATACCAGTGGCGAAGGCGCTGGGCTGGACGGATACTGACGCTGAGGTGCGAAAGCGTGGGG |
| P-asv5726 | GCGAGCGTTGTTCGGAATTATTGGGCGTAAAGGGCGCGTAGGCGGATTGTCAAGTCTGGTGTGAAAGACCGCAGCTTAACTGCGGGGGCGCGCTGGAAACTGACGATCTTGAATCCTGGAGGGGGAGCTGGAATTCCTGGTGTAGGGGTGAAATCTGTAGATATCAGGAAGAACACCGGTGGCGAAGGCGAGCTCCTGGCCGTGGATTGACGCTGAGGCGCGAAAGCGTGGGG |
| P-asv5727 | GCGAGCGTTACTCGGAATCACTAGGCGTAAAGCGCGTGTAGGCGGGTGCTTAAGTCCGCTGTGAAATCTCCCGGCTCAACTGGGAGCGGTCAGCGGATACTGGGCGCCTAGAGTGGGGTAGGGGGTGCTGGAATTCCCGGTGTAGCGGTGAAATGCGTAGATATCGGGAGGAACACCGATGGCGAAAGCAGGCACCTGGGCCCTCACTGACGCTAAGACGCGAAAGCTAGGGG |
| P-asv5728 | GCTAGCGTTGTTCGGAATTACTGGGCGTAAAGGGCGCGTAGGCGGCCTGGCAAGTCGGATGTGAAAGCCCGGGGCTCAACCTCGGAACGGCATTCGAAACTGCTGGGCTGGAGAACGGAAGAGGAGAGTGGAATACCCAGTGTAGAGGTGAAATTCGTAGATATTGGGTGGAACACCGGTGGCGAAAGCGGCTCTCTGGTCCGGATCTGACGCTGAGGCGCGAAAGCGTGGGG |
| P-asv5729 | GCTAGCGTTGTTCGGAATGACTGGGCGTAAAGGGCGCGTAGGCGGCTTACCAAGTCAGGTGTGAAATCCTTGGGCTTAACCCAAGAACTGCATTTGATACTGGTTTGCTAGAGGAAATGAGAGGTAAGTGGAATTCCTAGTGTAGAGGTGAAATTCGTAGATATTAGGAGGAACACCAGAGGCGAAGGCGGCTTACTGGCATTTACCTGACGCTGAGGCGCGAAAGCGTGGGG |
| P-asv5730 | GCAAGCGTTAATCGGATTTACTGGGCGTAAAGGGTGCGTAGGTGGTTGTATAAGTCGGATGTGAAAGCCCGGGGCTTAACCTCGGAACTGCATTCGATACTGTATGACTAGAGTTCGGTAGAGGGCGGCGGAATTTCCGGTGTAGCGGTGAAACGCGTAGATATCGGAAGGAACATCAATGGCGAAGGCAGCCGCCTGGGCCTGAACTGACACTGAGGCACGAAAGCGTGGGT |
| P-asv5731 | GCGAGCGTTAATCGGAATTACTGGGCGTAAAGCGTACGCAGGCGGTTTGTTAAGCGAGATGTGAAAGCCCCGGGCTCAACCTGGGAACAGCATTTCGAACTGGCAAGCTAGAGTGTGATAGAGGGTGGTAGAATTTCAGGTGTAGCGGTGAAATGCGTAGAGATCTGAAGGAATACCGATGGCGAAGGCAGCCACCTGGGTCAACACTGACGCTCATGTACGAAAGCGTGGGG |
| P-asv5732 | GCGAGCGTTAATCGGAATTACTGGGCGTAAAGCGCATGCAGGTGGTTCATTAAGTCAGATGTGAAAGCCCGGGGCTCAACCTCGGAACTGCATTTGAAACTGGTGAACTAGAGTACTGTAGAGGGGGGTAGAATTTCAGGTGTAGCGGTGAAATGCGTAGAGATCTGAAGGAATACCAGTGGCGAAGGCGGCCCCCTGGACAGATACTGACACTCAGATGCGAAAGCGTGGGG |
| P-asv5733 | GCGAGCGTTGTCCGGATTTATTGGGTTTAAAGGGTGCGTAGGTGGCTGTTTAAGTCGTGGGTGAAATGCTGTGGCTCAACCATAGAAGTGCCTGCGATACTGGATAGCTTGAGTAATTTGGAGGTAGCTGGAATTGATGGTGTAGCGGTGAAATGCATAGATACCATCAGGAACACCGATAGCGAAGGCAGGCTACTACGGATTAACTGACACTGAGGCACGAAAGCGTGGGT |
| P-asv5734 | GCAAGCGTTAATCGGAATTACTGGGCGTAAAGCGCGCGTAGGTGGTTTGATAAGCGAGATGTGAAAGCCCCGGGCTTAACCTGGGAACGGCATTTCGAACTGTCAGGCTAGAGTGTGGTAGAGGGTAGTGGAATTTCCTGTGTAGCGGTGAAATGCGTAGATATAGGAAGGAACACCAGTGGCGAAGGCGACTACCTGGACCAACACTGACACTGAGGTGCGAAAGCGTGGGG |
| P-asv5735 | GCTAGCGTTGCTCGGAATCACTGGGCGTAAGGGGTGCGTAGGCGGGTCTTTAAGTCAGGGGTGAAATCCTGGAGCTCAACTCCAGAATTGCCTTTAAGACTGCATCGCTTGAATCCAGGAGAGGTGAGTGGAATTCCGAGTGTAGAGGTGAAATTCGTAGATATTCGGAAGAACACCAGTGGCGAAGGCGGCTCACTGGACTGGTATTGACGCTGAGGTGCGAAAGCGTGGGG |
| P-asv5736 | GCGAGCGTTGTCCGGAATCATTGGGCGTAAAGCGCGTGTAGGCGGCCATCCAAGTCTGATGTGAAAGCTCGGGGCTCAACCCCGAGTCTGCATTGGAAACTGTATGGCTTGAGTCCGGAAGAGGAGACTGGAATTCCTGGTGTAGCGGTGAAATGCGCAGATATCAGGAAGAACACCGGTGGCGAAGGCGGGTCTCTGGGACGGTACTGACGCTGAGACGCGAAAGCGTGGGG |
| P-asv5737 | TCAAGTGGTCAGGATGATTATTGGGCCTAAAGCATCCGTAGCCGGCTCTGTAAGTTTTCGGTTAAATCTGTACGCTCAACGTACAGGCTGCCGGGAATACTGCAGAGCTAGGGAGTGGGAGAGGTAGACGGTACTCGGTAGGAAGGGGTAAAATCCTTTGATCTATTGATGACCACCTGTGGCGAAGGCGGTCTACCAGAACACGTCCGACGGTGAGGGATGAAAGCTGGGGG |

| | |
|---|---|
| P-asv5738 | GCGAGCGTTAATCGGAATTACTGGGCGTAAAGCGTGCGCAGGCGGTTTTGAAAGTCAGATGTGAAATCCCCGAGCTCAACTTGGGAACTGCGTTTGAAACTCCAAAGCTAGAGTATAGGAGAGGGGGGTAGAATTCCACGTGTAGCAGTGAAGTGCGTAGAGATGTGGAGGAATACCGATGGCGAAGGCAGCCTCCTGGGATAACACTGACGTTGAGGCTCGAAAGCGTGGGG |
| P-asv5739 | GCAAGCGTTATCCGGAATCATTGGGTTTAAAGGGTCCGTAGGCGGACAATTAAGTCAGGGGTGAAAGTCTGTAGCTCAACTATAGAATTGCCTTTGATACTGGTTGTCTTGAATTACTGTGAAGTGGTTAGAATAAGTAGTGTAGCGGTGAAATGCATAGATATTACTTAGAATACCAATTGCGAAGGCAGATCACTAACAGTATATTGACGCTGATGGACGAAAGCGTGGGG |
| P-asv5740 | CCAAGCGTTATCCGGATTCATTGGGTTTAAAGGGTCCGCAGGCGGGGCATTAAGTCAGTGGTGAAAGCCTACCGCTCAACGGTAGGACTGCCATAGATACTGGTGCTCTTGAGTGTAGTTGAAGTGGGCGGAATGAGTCATGTAGCGGTGAAATGCATAGATATGACTCAGAACACCGATTGCGAAGGCAGCTCACTAAGCTAACACTGACGCTCAGGGACGAAAGCGTGGGT |
| P-asv5741 | GCGAGCGTTATTCGGATTTACTGGGCGTAAAGGGTCCGTAGGGGGACATTTAAGTCAGATGTGAAATTTCAGGGCTCAACCCTGAACCTGCATTTGAAACTGAGTGTCTAGAGTTTGGTAGAGGTAAGTGGAATTTGTGGTGTAGCGGTGGAATGCGTAGATATCACAAGGAACATCAAAGGCGAAAGCAGCTTACTGGGCCTAAACTGACCCTGAGGGACGAAAGCTAAGGT |
| P-asv5742 | GCAAGCGTTGTCCGGAATCACTGGGCGTAAAGGGTGTGCAGGTGGGGCAGCAAGTCAGCGGTGAAATCCTACGGCTTAACCGTAGAAGTGCCGTTGAAACTGTTGCTCTTGAGTCCTAGAGAGGTCGCTGGAATTCGTGGTGTAGCGGTGAAATGCGTAGATATCACGAGGAACACCAGAGGCGTAAGCGAGCGACTAGATAGGTACTGACGCTGAGGCACGAAAGCGTGGGG |
| P-asv5743 | GCAAGCGTTGTCCGGAATCATTGGGCGTAAGGCGCGTGTAGGCGGTTACGTAAGTCTGCTGTGAAAGTCGGAGGCTCAACCTTCGAATGCCAGTGGATACTGCGTGACTAGAGTTCGGAAGAGGCAAGTGGAATTCCTGGTGTAGCGGTGAAATGCGCAGATATCAGGAGGAACACCAACGGCGAAGGCAGCTTGCTGGTACGACACTGACGCTGAGACGCGAAAGCGTGGGG |
| P-asv5744 | CCTAGCGTAGTTCGGAATTACTGGGCTTAAAGAGTTCGTAGGTGGTTGAAAAAGTTGGTGGTGAAATCCCAGAGCTTAACTCTGGAACTGCCATCAAAACTTTTCAGCTAGAGTATGATAGAGGAAAGCAGAATTTCTAGTGTAGAGGTGAAATTCGTAGATATTAGAAAGAATACCAATTGCGAAGGCAGCTTTCTGGATCATTACTGACACTGAGGAACGAAAGCATGGGT |
| P-asv5745 | CCAAGTGATGGCCGATATTATTGGGCCTAAAGCGTCCGTAGCTTGCTGTGTAAGTCCATTGGGAAATCGACCAGCTCAACTGGTCGGCGTCCGGTGGAAACTACACAGCTTGGGGCCGAGAGACTCAACGGGTACGTCCGGGGTAGGAGTGAAATCCTGTAATCCTGGACGGACCACCAATGGGGAAACCACGTTGAGAGACCGGACCCGACAGTGAGGGACGAAAGCCAGGGT |
| P-asv5746 | CCGAGTGATGGCCAATATTATTGGGCCTAAAGCGTCCGTAGCTGGCTAGACAAGTCCGTTGGGAAATCTGTTCGCTTAACGAGCAGGCGTCCAGCGGAAACTGTGTAGCTTGGGACCGGAAGACCTGAGGGGTACGTCTGGGGTAGGAGTGAAATCCCGTAATCCTGGACGGACCACCGGTGGCGAAAGCGCCTCAGGAGAACGGATCCGACAGTGAGGGACGAAAGCTAGGGT |
| P-asv5747 | GCAAGCGTTGTCCGGAATTACTGGGCGTAAAGAGCTCGTAGGCGGTTTGTCACGTCGTCTGTGAAATCCTCCAGCTCAACTGGGGGCGTGCAGGCGATACGGGCAGACTTGAGTACTACAGGGGAGACTGGAATTCCTGGTGTAGCGGTGAAATGCGCAGATATCAGGAGGAACACCGGTGGCGAAGGCGGGTCTCTGGGTAGTAACTGACGCTGAGGAGCGAAAGCATGGGT |
| P-asv5748 | GCAAGCGTTATCCGGAATCACTGGGCATAAAGCGTCTGTAGGTGGTTTGGTAAGTCTGCTGTTAAAGACTGGGGCTCAACCCCAGAAAAGCAGTGGAAACTGCTAGACTTGAGTGTGGTAGAGGTAGAGGGAATTCCTAGTGTAGCGGTGAAATGCGTAGATATTAGGAAGAACACCAATGGCGAAGGCACTCTACTGGGCCATAACTGACACTGAGAGACGACAGCTAGGGG |
| P-asv5749 | GCAAGCGTTGTCCGGAATTACTGGGCGTAAAGAGTTCGTAGGCAGTCAAATAAGTCAGAAGTGAAAGGCTACGGCTCAACCGTAGTAAGCTTTTGAAACTGTTTGACTTGAGTACTGGAGAGGAAAGTGGAATTCCTAGTGTAGCGGTGAAATGCGTAGATATTAGGAGGAACACCAGTGGCGAAGGCGGCTTTCTGGACAGTAACTGACGCTGAGGAACGAAAGCGTGGGG |
| P-asv5750 | GCGAGCGTTATCCGGAATCACTGGGCGTAAAGGGCGTGTAGGCGGTTTGTCAAGTCCGATGTTAAAGCGTGAGGCTCAACCTCATATGAGCGTTGGATACTGGCGAGCTGGACGGTTGGAGAGGAAGGTAGAATTACCAGTGTAGCGGTGGAATGCGTAGATACTGGTAGGAATACCCATTGCGAAGGCAGCCTTCTGGACAACACGTGACGCTGAGGCGCGAAAGTGTGGGG |
| P-asv5751 | GCAAGCGTTGTTCGGATTTACTGGGCGTAAAGCGCGCGCAGGCGGATTATCAAGTCAGATGTGAAATCTCGGGGCTCAACCCCGAAACTGCGTCTGAAACTGATAGTCTAGAGTCTCAGAGGGGGTAGGGGGATTTCACGTGTAGGGGTAAAATCCGTAGAGATGTGAAGGAACACCAGTGGCGAAGGCGCCTACCTGGATGAGTACTGACACTGAGGCGCGAAAGCGTGGGG |
| P-asv5752 | GCTAGCGTTGCTCGGAATTACTGGGCGTAAAGGGTGCGTAGGCGGGTCTTTAAGTCAGTGGTGAAAGCCTGCAGCTTAACTGTAGAACTGCCATTGATACTGGAGACCTTGAGTGTACTAGAGGTAGGCGGAATTTATGGTGTAGCGGTGAAATGCATAGATACCATAAAGAACACCGATAGCGTAGGCAGCTTACTGGAGTACAACTGACGCTGAGGCACGAAAGCATGGGG |
| P-asv5753 | GCTAGCGTTGCTCGGAATCACTGGGCGTAAAGGGTGCGTAGGCGGGTCTTTAAGTCAGGGGTGAAATCCTGGAGCTTAACTCCAAAACTGCCATTGATACTATATATCTTGAATATTGTGGAGGTTTGCGGAATATGTCATGTAGCGGTGAAATGCTTAGATATGACATAGAACACCTATTGCGAAGGCAGCAGGCTACACATATATTGACGCTGAGGCACGAAAGCGTGGGG |
| P-asv5754 | GCAAGCGTTAATCGGAATTACTGGGCGTAAAGGGTGCGTAGGTGGTTTAATAAGTTGGCTGTGAAATTCCTGGGCTCAACCTGGGGCGGTCAGCCAAGACTGTTAGACTCGAGTTTGGGAGAGGGTAGTGGAATTTCCGGTGTAGCGGTGAAATGCGTAGAGATCGGAAGGAACACCAGTGGCGAAGGCGGCTACCTGGCCTAAAACTGACACTGAAGCACGAAAGCGTGGGG |
| P-asv5755 | GCAAGCGTTGTCCGGAATCACTGGGCGTACAGGGTGCGCAGGCGGACGTGCAAGTCTGACGTGAAAGCCCTGAGCTCAACTCAGGAAGTGCGTCGGAAACTGCATGTCTTGAGGGCCAGAGAGGCCGGTGGAATTCGCGGTGGAGCGGTGAAATGCCTAGATATCGCGAGGAACACCCGTGGCGAAAGCGACCGGCTGGATGGACCCTGACGCTGAGGCACGAAAGCGTGGGT |
| P-asv5756 | GCAAGCGTTAATCGGAATTACTGGGCGTAAAGCGTGCGCAGGCGGCCTATTAAGTCAGATGTGAAATGCCCGAGCTCAACTTGGGAACTGCATTTGAAACTGATAGGCTAGAGTGTGTCAGAGGGAGGTAGAATTCCACGTGTAGCAGTGAAATGCGTAGAGATGTGGAGGAATACCAGTGGCGAAGGCGGCCTCCTGGGATAACACTGACGCTCATGTACGAAAGCGTGGGG |
| P-asv5757 | GCTAGCGTTGTTCGGAATCACTGGGCGTAAAGCGCGCGTAGGCGGCTTTTCAAGTCAGGGGTGAAATCCTGAGGCTTAACCTCAGAATTGCCTTTGAAACTGTAAAGCTAGAGTGTCGGAGGGGATAGCGGAATTTCCAATGTAGCGGTGAAATGCGTAGATATTGGAAGGAACACCGGTGGCGAAGGCGGCTATCTGGACGACAACTGACGCTGAGGCGCGAAAGCGCGGGG |
| P-asv5758 | GCAAGCGTTATCCGGAATCACTGGGCGTAAAGGGTGCGTAGGCGGCTTGTCAAGCCAGAGGTGAAAGGCATTGGCTCAACCAATGTAAGCCTTTGGAACTGACGAGCTTGAGTACAGGAGAGGTAAGTGGAATTCCTAGTGTAGAGGTGAAATTCGTAGATATTAGGAGGAACACCAGTGGCGAAGGCGACTTACTGGACTGTAACTGACGCTGAGGCACGAAAGCGTGGGT |
| P-asv5759 | GCGAGCGTTATCCGGATTTATTGGGTTTAAAGGGTGCGTAGGCGGTTTCTTAAGTCAGCGGTAAAATTTCAGGGCTTAACCCTGTCGTGCCGTTGAAACTGGGGGACTTGAGTCTGGATGAAGTGGGCGGAATTTGTAGTGTAGCGGTGAAATGCATAGATATTACAAGGAACTCCAATTGCGCAGGCAGCTCACTAAACCATCACTGACGCTGAAGCACGAAAGCGTGGGT |
| P-asv5760 | GCAAGCGTTGTTCGGAATCACTGGGCTTAAAGCGTGCGTAGGCGGATTTGCAAGTCAGGAGTGAAATCCAGGGGCTCAACCCCTGAATTGCTCTTGAAACTGCAAATCTAGAATCCGGCAGGGGTTGGGAGAATTCCTAGTGTAGAGGTGAAATTCGCAGATATTAGGAGGAATACCAGTGGCGTAGGCGCCCAACTGGACCGGTATTGACGCTGAGGTACGAAAGCGTGGGG |
| P-asv5761 | CCTAGCGTTGTTCGGAATCATTGGGCGTAAAGGGTGTGTAGGTGGCCATGTAAGTCAGGTGTGAAAGCCCCGGGCTCAACCCGGGAAGTGCATTTGATACTGCGTGGCTTGAGTACCGAAGAGGGTAGTGGAATTCCAGGTGTAGTGGTGAAATACGTAGATATCTGGAGGAACACCGGTGGCGAAGGCGGCTACCTGGTCGAGTACTGACACTGAGACACGAAAGCGTGGGG |
| P-asv5762 | GCAAGCGTTGTCCGGATTTATTGGGTTTAAAGGGTGCGTAGGCGGCCCATTAAGTCAGCGGTGAAAGCCCGGGGCTCAACCCCGGAACCGCCGTTGATACTGGTGGGCTTGAGTATGGACGAGGTTGGCGGAATTGAGGGTGTAGCGGTGAAATGCTTAGATACCCTCAAGAACACCGATTGCGTAGGCAGCTAACTAGGCCATAACTGACGCTGAGGCACGAAAGCGTGGGG |
| P-asv5763 | GCAAGCGTTGTCCAGATTTATTGGGTTTAAAGGGTGCGTAGGCGGCTGTTTAAGTCAGTGGTGAAAGACTCCGGCTCAACCGGAGCAGTGCCATTGATACTGGACAGCTTGAGTATTGGAGGGGTACATGGAATTGATGGTGTAGCGGTGAAATGCATAGATACCATCAGGAACACCGATAGCGAAGGCATTGTACTGGCCAATAACTGACGCTGATGCACGAAAGCATGGGT |
| P-asv5764 | GCAAGCGTTATCCGGATTTACTGGGTGTAAAGGGAGCGTAGGTGGTATGGCAAGTCAGATGTGAAAGCCCGGGGCTCAACCCCGGGACTGCATTTGAAACTGTCAAACTAGAGTGCAGGAGAGGTAAGTGGAATTCCTAGTGTAGCGGTGAAATGCGTAGATATTAGGAGGAACACCAGTGGCGAAGGCGGCTTACTGGACTGTAACTGACACTGAGGCTCGAAAGCGTGGGG |
| P-asv5765 | GCAAGCGTTATCCGGAATTATTGGGCGTAAAGCGTCCGCAGGCGGCCTTGTAAGTCTGCTGTCAAAGAGTGGAGCTCAACTCCATAGAGGCAGTGGAAACTGTAAGGCTTGAGTGTGGTAGGGGCAGAGGGAATTCCCGGTGTAGCGGTGAAATGCGTAGATATCGGGAAGAACACCAGTGGCGAAGGCGCTCTGCTGGGCCATAACTGACGCTCATGGACGAAAGCCAGGGG |
| P-asv5766 | GCAAGCGTTATCCGGATTCACTGGGTTTAAAGGGTGCGTAGGAGGGCAGATAAGTCAGTGGTGAAATCTTCGAGCTTAACTCGGAAACTGCCGTTGATACTATTTGTCTTGAATATCGTGGAGGTGAGCGGAATATGTCATGTAGCGGTGAAATGCTTAGATATGACATAGAACACCAATTGCGAAGGCAGCTCGCTACACGAATATTGACTCTGAGGCACGAAAGCGTGGGG |
| P-asv5767 | GCAAGCGTTATCCGGATTTATTGGGCGTAAAGCGAGCGCAGGCGGTTGCTTAGGTCTGATGTGAAAGCCTTCGGCTTAACCGAAGAAGGGCATCGGGAACCGGGCGACTTGAGTGCAGAAGAGGACAGTGGAACTCCATGTGTAGCGGTGGAATGCGTAGATATATGGAAGAACACCAGTGGCGAAGGCGGCTGTCTGGTCTGCAACTGACGCTGAGGCTCGAAAGCATGGGT |
| P-asv5768 | GCAAGCGTTAATCGGAATTACTGGGCGTAAAGCGCACGTAGGCGGTATGTTAAGTTGGATGTGAAAGCCCCGGGCTCAACCTGGGAATTGCATTCAAAACTGGCATGCTAGAGTATGTGAGAGGGGGGTAGAATTCCAAGTGTAGAGGTGAAATTCGTAGATATTCGGAGGAACACCAGTGGCGAAGGCGGCTCACTGGCCCGATACTGACGCTGAGGCACGAAAGCGTGGGG |
| P-asv5769 | GCAAGCGTTGTCCGGATTTACTGGGCGTAAAGAGCGCGCAGGCGGTCGTGCAAGTCGCGTGTGAAAGCCCCCGGCTCAACTGGGGAGGGTCACGCGATACTGCTCGACTCGAAGGCAGGAGAGGGAAGTGGAATTCCCGGTGTAGTGGTGAAATGCGTAGATATCGGGAGGAACACCAGTGGCGAAGGCGACTTCCTGGCCTGTTCTTGACGCTGAGGCGCGAAAGCTAGGGG |
| P-asv5770 | GCGAGCGTTGTTCGGATTTACTGGGCGTAAAGCGTGTGTAGGCGGTTTTTCAAGTCAGAAGTGAAAGCCCTGGGCTTAACCCAGGAATTGCTTTTGAAACTGGATCACTAGAATGTGGGAGAGGCAAGCAGAGTTCCGAGTGTAGAGGTGAAATTCGTAGATATTCGGAGGAATACCAAGGGCGTAGGCAGCTTGCTGGAACACTATTGACGCTGAGACACGAAAGCATGGGG |
| P-asv5771 | GCAAGCGATACTCGGAATCACTAGGCGTAAAGCGCAGGTAGGCGGTTCGGTAAGTCTTTTGTGAAAGCTCCCGGCTTAACTGGGAGAGGTCAAGGGAAACTGCTGGGCTAGAGTGTGGTAGGGGATACTGGAATTCCTGGTGTAGCGGTGAAATGCGTAGATATCAGGAGGAATACCTATGGCGAAAGCAGGTATCTGGGCCATCACTGACGCTGAGCTGCGAAAGCTAGGGG |
| P-asv5772 | GCTAGCGTTGTTCGGAATTACTGGGCGTAAAGGGCGCGTAGGCGGCCGTTTAAGTTAGATGTGAAAGTCCTGGGCTCAACCTGGGGACTGCATTTAATACTGGATGGCTTGAATTCGGGAGAGGATAGCGGAATTCCCAGTGTAGAGGTGAAATTCGTAGATATTGGGAAGAACACCGGTGGCGAAGGCGGCTATCTGGACCGACATTGACGCTGAGGCGCGAAAGCGTGGGG |
| P-asv5773 | GCAAGCGTTGTCCGGAATCACTGGGCGTAAAGGGTACGCAGGCGGATATACAAGTCAAATGTGAAATATACCGGCTCAACCGGTAAGGTGCATTTGAAACTGTATATCTTGAGTCTAGGAGAGGAAAGCGGAATTCCTGGTGGAGCGGTGGAATGCGTAGATATCAGGAAGAACACCAGTGGCGAAGGCGGCTTTCTGGCCTAAGACTGACGCTCATGTACGAAAGCTAGGGG |
| P-asv5774 | GCAAGCGTTATCCGGAATCACTGGGTTTAAAGGGTGCGTAGGCGGACCAGTAAGTCAGAGGTGAAAGCCCGTCGCTTAACGACGGAATTGCCTTTGATACTGTTGGTCTTGAATCAGGTTGAGGTTAGCGGAATGTGGCATGTAGCGGTGAAATGCGTAGATATGCCATAGAACACCGATTGCGAAGGCAGCTAACTATCCCTGTATTGACGCTGAGGCACGAAAGCGTGGGG |
| P-asv5775 | GCGAGCATTATTCGGAATGATTAGGCGTAAAGGGTTTGTAGGTGGTTTTTTAAGTTGAAAAAAACGAATTAAAGCTCAACTTTATAAATTTTTTCAAAACTGAAAAACTTGAGTATACATAGAGGATAATGGAATTTCTATTGGAGGGATAGAATATGTTGATAATAGAAGGAAGGTCGATAGCGAAGGCAATTATCTGGGTATATACTGACACTGAAAAACGAAAGCTTGGGT |
| P-asv5776 | GCGAGCGTTAATCGGAATTACTGGGCGTAAAGCGTGCGCAGGCGGTTTTGTAAGACAGATGTGAAATCCCCGGGCTTAACCTGGGAACTGCGTTTGTGACTGCAAGGCTAGAGTACGGCAGAGGGGGGTGGAATTCCTGGTGTAGCAGTGAAATGCGTAGAGATGTGGAGGAATACCGGTGGCGAAGGCGGCCCCCTGGACAAAGACTGACGCTGAGGTGCGAAAGCGTGGGG |

| | |
|---|---|
| P-asv5777 | GCAAGCGTTGTCCGGAATTATTGGGCGTAAAGGGCTCGCAGGCGGTTTCTTAAGTCTGATGTGAAATCTTGTGGCTTAACCACAAGCGGTCATTGGAAACTGGGAGACTTGAGTGCAGAAGAGGAGAGTGGAATT CCACGTGTAGCGGTGAAATGCGTAGAGATGTGGAGGAACACCAGTGGCGAAGGCGACTCTCTGGTCTGTAACTGACGCTGAGGAGCGAAAGCGTGGGG |
| P-asv5778 | GCAAGCGTTGTCCGGAATCATTGGGTGTAAAGGGTGCGTAGACGGGCTAGTAAGTCTGGGGTGAAATCTTACCGCTTAACGGTAAAACTGCCTTGGATACTGCTGGTCTTGAGTGTGAGAGAGGTCGGTGGAAT TCGTGGTGTAGCGGTGAAATGCTTAGATATCACGAAGAACACCTGTGGTGAAGACGGCCGGCTGGATCACAACTGACGCTGAGGCACGAGAGCGTGGGG |
| P-asv5779 | GCAAGCGTTGTCCGGAATTATTGGGCGTAAAGGGCGCGTAGGCGGATTGTTAAGTCCGGTGTGAAAGATCAGGGCTCAACTCTGAGAGTGCATCGGAAACTGGGAATCTTGAGGACAGGAGAGGAAAGTGGAAT TCCACGTGTAGCGGTGAAATGCGTAGATATGTGGAGGAACACCGGTGGCGAAGGCGACTTTCTGGACTGTAACTGACGCTGAGGCGCGAAAGCGTGGGG |
| P-asv5780 | GCAAGCGTTAATCGGAATTACTGGGCGTAAAGCGCACGCAGGCGGTCTGTTAAGTCAGATGTGAAATCCCCGGGCTCAACCCGGGAACTGCATTTGAAACTGGCAGGCTTGAGTCTCGTAGAGGGGGGTAGAAT TCCAGGTGTAGCGGTGAAATGCGTAGAGATCTGGAGGAATACCGGTGGCGAAGGCGGCCCCCTGGACGAAGACTGACGCTCAGGTGCGAAAGCGTGGGG |
| P-asv5781 | GCTAGCGTTGTTCGGAATTATTGGGCGTAAAGGGCATGTAGGCGGTTTGGTAGGCCTGATGTGAAATCCTGCAGCTTAACTGTAGAATTGCATTGGGTACTGCCGAACTTGAATCACTGAGGGGAAACTAGAATT CCAGGTGTAGGGGTGGAATCTGTAGATATCTGGAAGAATACCGGTGGCGAAGGCGGGTTTCTAGCAGATGATTGACGCTGAGGTGCGAAAGTGCGGGG |
| P-asv5782 | GCTAGCGTTGTTCGGAATTACTGGGCGTAAAGCGCACGTAGGCGGACTTTTAAGTCTGGGGTGAAAGCCCGGAGCTCAACTCCGGAACTGCCTTGGATACTGGGAGTCTAGAGGATGGAAGAGGTAAGTGGAAT ACCGAGTGTAGAGGTGAAATTCGTAGATATTCGGTGGAACACCAGTGGCGAAGGCGACTTACTGGTCCATTTCTGACGCTGAGGTGCGAAAGTGTGGGG |
| P-asv5783 | GCAAGCGTTATCCGGATTTACTGGGTGTAAAGGGAGCGTAGACGGTGAGGTAAGTCAGATGTGAAAGCCCAGGGCTCAACCCTGGGACTGCATTTGAAACTACCTGGCTGGAGTGCAGGAGAGGTAAGTGGAAT TCCTAGTGTAGCGGTGAAATGCGTAGATATTAGGAGGAACACCAGTGGCGAAGGCGGCTTACTGGACTGTAACTGACGTTGAGGCTCGAAAGCGTGGGG |
| P-asv5784 | CCAAACGTTATTCGGAATTATTGGGCTTAAAGGGTGCGTAGGCGGCGCTTTAAGTCGGGTGTGAAATCCTTCGGCTCAACCGAAGAATTGCGCCCGATACTGGAGTGCTAGAGGGAGACAGGGGTGAGCGGAAC TGATGGTGGAGCGGTGAAATGCGTTGATATCATCAGGAACACCGGTGGCGAAAGCGGCTCACTGGGTCTCTTCTGACGCTGAGGCACGAAAGCCAGGGT |
| P-asv5785 | CCAAGCGTTGTTCGGATTTATTGGGCGTAAAGCGCGCGCAGGTGGATTAGCAAGTCAGGTGTGAAATCTCGGAGCTTAACTCCGAAACTGCGCCTGAAACTGCTAGTCTAGAATGTCGGAGGGGGCAGGGGAAT TTCACGTGTAGGGGTAAAATCCGTAGAGATGTGAAGGAACACCGGAGGCGAAGGCGCCTGCCTAGACGATTATTGACACTGAGGCGCGAAAGCGTGGGG |
| P-asv5786 | GCGAGCGTTGTTCGGAATCACTGGGCGTAAAGCGTGCGTAGGCCGTCTTGTAAGTCAGAAGTGAAAGCCCGGGGCTTAACCCCGGAATTGCTTTTGAAACTGCAAGACTAGAATTCGGGAGAGGATAGCGGAAT TCCCAGTGTAGTAGTGAAATACGTAGATATTGGGAGGAACACCAGTGGCGTAAGCGGCTATCTGGACCGACATTGACGCTGAGGCACGAAAGCGTGGGG |
| P-asv5787 | GCAAGCGTTGTCCGGAATTATTGGGCGTAAAGAGCTCGTAGGCGGTCTGTCGCGTCTGCTGTGAAAACTCGAGGCTTAACCTCGGGCTTGCAGTGGGTACGGGCAGACTAGAGTGCAGTAGGGGAGACTGGAA TTCCTGGTGTAGCGGTGAAATGCGCAGATATCAGGAGGAACACCGATGGCGAAGGCAGGTCTCTGGGCTGTTACTGACGCTGAGGAGCGAAAGCATGGGG |
| P-asv5788 | GCTAGCGTTGTTCGGAATCATTGGGCGTAAAGCGCGTGTAGGTGGCTTGCTAAGTCGATTGTGAAATCCCTAGGCTCAACCAAGGAAGTGCATTCGAAACTAGCTCGCTCGAGGATGGCAGAGGAAGGTGGAAT TCCAAGTGTAGAGGTGAAATTCGTAGATATTTGGAGGAATACCGGCGGCGAAGGCGGCCTTCTGGACCATTCCTGACACTGAGACGCGAAAGCGTGGGG |
| P-asv5789 | GCTAGCGTTGTTCGGATTTACTGGGCGTAAAGCGCGCGTAGGCGGATAAATTAGTTAGAGGTGAAATCCCAGGGCTCAACCCTGGAACTGCCTTTAATACTGTTTATCTAGAGTATGAAAGAGGTGAGTGGAATTC CGAGTGTAGAGGTGAAATTCGTAGATATTCGGAGGAACATCAGTGGCGAAGGCGGCTCACTGGTTCATTACTGACGCTGAGGTGCGAAAGCGTGGGG |
| P-asv5790 | GCGAGCGTTAATCGGATTTACTGGGCGTAAAGCGTGCGTAGGCGGCTTTTTAAGTCGGATGTGAAATCCCTGAGCTTAACTTAGGAATTGCATTCGATACTGGGAAGCTAGAGTATGGGAGAGGATGGTAGAATT CCAGGTGTAGCGGTGAAATGCGTAGAGATCTGGAGGAATACCAGTGGCGAAGGCGACTTTCTGGTCTGTAACTGACGCTGAGGCTCGAAAGCATGGGG |
| P-asv5791 | GCGAGCGTTATTCGGAATCATTGGGCGTAAAGGGTGCGTAGACGGTGTGCTAAGTCTGTTGTTAAATTTTCCGGCCTAACCGGAAACCTGCGGTGGAAACTGGCGCACTAGAGGGTGGAAGAGAGAAGTGGAAT TCTCGGAGTAGCGGTAAAATGCGTGGATCTCGAGAGGAACACCGATGGCGAAGGCAGCTTCTTGGTCCATACCTGACGTTGAGGCACGAAAGCGTGGGG |
| P-asv5792 | GCTAGCGTTGCTCAGAATTACTGGGCGTAAAGGGCGCGTAGGCGGACAGTTAAGTTGGGGGTGAAAGCCCCGGGCTCAACCTGGGAATTGCATTCGATACTGTAAAGCTAGAGTATGTGAGAGGGGGGTAGAAT TCCAAGTGTAGCGGTGAAATGCGTAGAGATTTGGAGGAATACCAGTGGCGAAGGCGGCCCCCTGGCACAATACTGACGCTCAGGTGCGAAAGCGTGGGG |
| P-asv5793 | TCAAGTGGTCGTCGCTTTTATTGGGCCTAAAACGTCCGTAGCCTGTTTGGTAAATCTGTGGGTAAATCAACCAGCTTAACTGGTTGAATTCTGCAGAGACTGCCAGACTAGGGACCGGGAGAGGTGTGGGGTACC CTAGGGGTAGGGGTAAAATCCTGTCATCCTTAGAGGACCACCAGTTGCGAAGGCGCCACACTGGAACGGATCCGACGGTCAGGGACGAAGCCTAGGGG |
| P-asv5794 | GCACGCGTTGTCCGGAATTATTGGGCGTAAAGCGCGCGCAGGCGGTCTCTTAAGTCTGATGTGAAAGCCCACGGCTCAACCGTGGAGGGTCATTGGAAACTGGGAGACTTGAGTGTAGGAGAGGAAAGTGGAA TTCCACGTGTAGCGGTGAAATGCGTAGATGTGTGGAGGAACACCAGTGGCGAAGGCGACTTTCTGGCCTATAACTGACGCTGAGGCGCGAAAGCGTGGGG |
| P-asv5795 | GCAAGCGTTATCCGGATTCACTGGGTTTAAAGGGTGCGTAGGCGGGAATGTAAGTCAGTGGTGAAATCTCGGAGCTTAACTCCGAAACTGCCATTGATACTATGTTTCTTGAATATTGTGGAGGTGAGCGGAATAT GTCATGTAGCGGTGAAATGCTTAGATATGACATAGAACACCGATTGCGAAGGCAGCTCGCTACACAAATATTGACGCTGAGGCACGAAAGCGTGGGG |
| P-asv5796 | GCGAGCGTTGTCCGGAATTATTGGGCGTAAAGAGCTTGTAGGCGGTTTGTCGCGTCTGCTGTGAAAGGCCGGAGCTTAACTCCGTGTATTGCAGTGGGTACGGGCAGACTAGAGTGCAGTAGGGGAGACTGGA ATTCCTGGTGTAGCGGTGGAATGCGCAGATATCAGGAGGAACACCGATGGCGAAGGCAGGTCTCTGGGCTGTAACTGACGCTGAGGAGCGAAAGCATGGGG |
| P-asv5797 | GCGAGCGTTAATCGGAATTACTGGGCGTAAAGCGCGCGTAGGCGGCCAGGTCAGTTGGATGTGAAAGCCCAGGGCTTAACCCTGGAACTGCATTCAATACTGCCGGGCTAGAGTACGGTAGAGGGGGGTAGAA TTCCACGTGTAGCGGTGAAATGCGTAGAGATGTGGAGGAATATCAGTGGCGAAGGCGGCCCCCTGGACTGATACTGACGCTGAGGTGCGAAAGCGTGGGG |
| P-asv5798 | CCAAGCGTTATCCGGAATCATTGGGTTTAAAGGGTCCGTAGGCGGTTTTATAAGTCAGTGGTGAAATCCGGCAGCTCAACTGTCGAACTGCCATTGATACTGTAGAACTTGAATTATTGTGAAGTAACTAGAATATG TAGTGTAGCGGTGAAATGCTTAGAGATTACATGGAATACCAATTGCGAAGGCAGGTTACTAACAATCGATTGACGCTGATGTACGAAAGCGTGGGT |
| P-asv5799 | GCAAGCGTTGTCCGGAATTATTGAGCGTAAAGAGCTCGTAGGCGGTTTGTCGCGTCTGCTGTGAAAACGCGAGGCTCAACCTCGCGCCTGCAGTGGGTACGGGCAAACTAGAGTGCGGTAGGGGAGATTGGAA ATCCTGGTGTAGCGGTGGAATGCGCAGATATCAGGAGGAACACCGATGGCGAAGGCAGATCTCTGGGCCGTAACTGACGCTGAGGAGCGAAAGCATGGGG |
| P-asv5800 | GCAAGCGTTATCCGGATTCACTGGGTTTAAAGGGTGCGTAGGCGGACAGGTAAGTCCGTGGTGAAATCTCCGAGCTTAACTCGGAAACTGCCATGGATACTCTCTGTCTTGAATACTGTGGAGGTGAGCGGAATA TGTCATGTAGCGGTGAAATGCTTAGATATGACATAGAACACCAATTGCGAAGGCAGCTCACTGCGCAGTGATTGACGCTGAGGCACGAAAGCGTGGGG |
| P-asv5801 | GCTAGCGTTGTTCGGAATTACTGGGCGTAAAGCGAGTGTAGGTGGTTGCCATAGTCAGGCGTGAAAGCCTTGAGCTCAACTCAAGAAATGCGCTTGATACTGGGCAGCTAGAGGACCGGAGAGGATAGTGGAAT TCCCAGTGTAGTGGTGAAATACGTAGAGATTGGGAAGAACACCAGTGGCGAAGGCGGCTATCTGGACGGTTTCTGACACTAAGACTCGAAAGCGTGGGG |
| P-asv5802 | GCGAGCGTTAATCGGAATAACTGGGCGTAAAGGGCACGCAGGCGGTGACTTATGTGAGGTGTGAAAGCCCCGGGCTTAACCTGGGAATTGCATTTCATACTGGGTCGCTAGAGTACTTTAGGGAGGGGTAGAAT TCCACGTGTAGCGGTGAAATGCGTAGAGATGTGGAGGAATACCGAAGGCGAAGGCAGCCCCTTGGGAATGTACTGACGCTCATGTGCGAAAGCGTGGGG |
| P-asv5803 | GCTAGCGTTGTTCGGAATTACTGGGCGTAAAGCGCACGTAGGCGGACATTTAAGTCAGGGGTGAAATCCAGGGGCTCAACCCCGGAACTGCCTTTGATACTGGGTGTCTTGAGTTCGAGAGAGGTGAGTGGAAT TGCGAGTGTAGAGGTGAAATTCGTAGATATTCGCAGGAACACCAGTGGCGAAGGCGGCTCACTGGCTCGATACTGACGCTGAGGTGCGAAAGCGTGGGG |
| P-asv5804 | GCTAGCGTTGTTCGGAATTACTGGGCGTAAAGCGCACGTAGGCGGCTATGTAAGTCAGAGGTGAAAGCCTGGAGCTCAACTCCAGAACTGCCTTTGATACTGGCAGTCTTGAGTACGGAAGAGGTATGTGGAACT CCGAGTGTAGAGGTGAAATTCGTAGATATTCGGAAGAACACCAGTGGCGAAGGCGACATACTGGTCCGTTACTGACGCTGAGGCTCGAAAGCGTGGGG |
| P-asv5805 | GCAAGCGTTGTTCGGAGTTACTGGGCGTAAAGGGCGCGTAGGCGGTGCAACAAGTCAGGCGTGAAAGCCCCGGGCTTAACCTGGGAATTGCGCTTGATACTGTTGCGCTAGAGATCGGGAGAGGATGGCGGAA TTCCCAGTGTAGAGGTGAAATTCGTAGATATTGGGAAGAACACCGGTGGCGAAGGCGGCCATCTGGACCGCATCTGACGCTGAGGCGCGAAAGCGTGGGG |
| P-asv5806 | GCTAGCGTTGTTCGGAATTACTGGGCGTAAAGGGCGTGTAGGCGGCCCGTAAAGTCAGGTGTGAAATTCCTGGGCTCAACCCAGGACGTGCGCCTGATACTGGCGGGCTTGAGTCCGGGAGAGGATGGTGGAA CTCCCAGTGTAGAGGTGAAATTCGTAGATATTGGGGAGAACACCCGTGGCGAAGGCGGCCATCTGGACCGGTACTGACGCTAAGGCGCGAAAGCGTGGGG |
| P-asv5807 | GCAAGCGTTATCCAGATTTACTGGGCGTAAAGCGCACGCAGGCGGATGACTAAGTCAGCTGTGAAAGCCCCCGGCTCAACTGGGGAGGGTCGGCTGAAACTGGTTGTCTTGAGGGCAGCAGAGGAGTGTGGAG TTGCGGGTGTAGTGGTGAAATGCGTAGATATCCGCAGGAACACCGGTGGCGAAGGCGGCACTCTGGGCTGTACCTGACGCTGATGTGCGAAGGCTGGGGG |
| P-asv5808 | GCTAGCGTTGCTCGGAATCACTGGGCTTAAAGGGCGCGTAGGCGGGTGCTCAAGTCCGGGGTGAAATCCTCCAGCTCAACTGGAGAACTGCCTCGGATACTGGGTGCCTCGAGGGTGGTAGGGGTATGCGGAA CTATCGGTGGAGCGGTGAAATGCGTTGATATCGATAGGAACTCCGGTGGCGAAGGCGGCATACTGGACCACATCTGACGCTGAGGCGCGAAAGCCAGGGG |
| P-asv5809 | GCGAGCGTTGTCCGGAATTATTGGGCGTAAAGAGCTCGTAGGCGGTGTGTCGCGTCGGCCGTGAAAACTTGCGGCTCAACTGTGGGCGTGCGGTCGATACGGGCATCACTGGAGTTCGGCAGGGGAGACTGG AATTCCTGGTGTAGCGGTGAAATGCGCAGATATCAGGAGGAACACCGGTGGCGAAGGCGGGTCTCTGGGCCGATATTGACGCTGAGGAGCGAAAGCGTGGGG |
| P-asv5810 | GCAAGCGTTGTCCGGAGTTACTGGGCGTAAAGCGCACGCAGGCGGTCCCATCAGTCTGTAGGGACAGGGCACAGCTTAACTGTGTCAGGCTTACGGAAACTGTGGGACTTGAGGGCTTCAGAGGGATGGGGAA TTCCGGGTGGAGTGGTGAAATGCGTAGAGATCCGGAAGAACACCAAAGGCGAAGGCACCATCCTGGGGAGTCACTGACGCTCGGGTGCGAAAGCTAGGGG |
| P-asv5811 | GCAAGCGTTGTCCGGAATTATTGGGCGTAAAGAGCTCGTAGGCGGTTTGTTACGTCCGCTGTGAAAACCCGGGGCTTAACCCTGGGCTTGCAGTGGATACGGGCAGACTAGAGGCAGGTAGGGGAGAATGGAA TTCCCGGTGTAGCGGTGAAATGCGCAGATATCGGGAGGAACACCGGTGGCGAAGGCGGTTCTCTGGGCCTGACCTGACGCTGAGGAGCGAAAGCGTGGGG |
| P-asv5812 | GCAAGCGTTGTTCGGAATTACTGGGCGTAAAGGGCTCGTAGGTGGCCAACTAAGTCAGACGTGAAATCCCTCAGCTTAACTGGGGAACTGCGTCTGATACTGGATGGCTTGAGTTTGGGAGAGGGATGCGGAAT TCCAGGTGTAGCGGTGAAATGCGTAGATATCGGGAGGAACACCGGTGGCGAAGGCGGCTTTCTGGACAGTAACTGACGCTGAGGCGCGAAAGCTAGGGG |
| P-asv5813 | GCGAGCGTTATCCGGAATTACTGGGTTTAAAGGGTGCGTAGGCGGCTATGTAAGTCAGGAGTGAAAGTTTGCGGCTCAACCGTAAAATTGCTTTTGATACTGCATGGCTGGAATCAGGATGAGGTCAGCGGAATG TGGCATGTAGCGGTGAAATGCATAGATATGCCATAGAACACCAATTGCGAAGGCAGCTGGCTAGACCTGTATTGACGCTGAGGCACGAAAGCGTGGGG |
| P-asv5814 | GCAAGCGTTGTTCGGAATTATTGGGCGTAAAGGGAGTGTAGGCGGTTTGGTAAGTCGTGTGTGAAATCTTCGGGCTCAACCCGAAGTCTGCACACGAAACTGCCGGGCTTGAGTGTGGGAGAGGTGAGTGGAAT TCCAGGTGTAGCGGTGAAATGCGTAGATATCTGGAGGAACACCGACGGCGAAGGCAGCCCGCTGGACAAATCCTGACACTCAGGCTCGAAAGTATGGGG |
| P-asv5815 | GCAAGCGTTATTCGGAATTATTGGGCGTAAAGGGCGCGTAGGCGGCTTTTCAAGTGACGGGTGAAATCCCTCGGCTTAACCGAGGAACTGCCTGTCAGACTGGAAGGCTTGAGACCGGGAGAGGTGAGTGGAAT TCCCAGTGTAGCGGTGAAATGCGTAGATATTGGGAGGAACACCAGTGGCGAAGGCGGCTCACTGGACCGGAACTGACGCTGAGGCGCGAAAGCGTGGGG |

| | |
|---|---|
| P-asv5816 | GCAAGCGTTGTTCGGAATTATTGGGCGTAAAGGGAGAGTAGGCGGTTTTGTAAGTCGTGTGTGAAATCTTCGGGCTCAACCCGAAGTCTGCACACGAAACTGCAGGGCTTGAGTATGGGAGAGGTGAGTGGAATTCCTGGTGTAGCGGTGAAATGCGTAGATATCAGGAGGAACACCTGTGGCGAAAGCGGCTCACTGGACCATTACTGACGCTGAGACTCGGAAGCTAGGGG |
| P-asv5817 | GCGAGCGTTGTTCGGAATTACTGGGCGTAAAGCGCGTGTAGGCGGCCGATTAAGTCTGGTGTGAAAGCCCGGGGCTCACCCCCGGAAGTGCGCTGGAAACTGGTCGGCTAGAGTATGGGAGAGGAAAGTGGAATTCCCGGTGTAGCGGTGAAATGCGTAGATATCGGGAGGAACACCAGCGGCGAAGGCGGCTTTCTGGACCAATACTGACGCTGAGACGCGAAAGCGTGGGG |
| P-asv5818 | GCAAGCGTTGTTCGGAATTACTGGGCGTAAAGGGTGCGTAGGCGGCTTTCTAAGTTGGATGTGAAAACTCTGGGCTCAACCCAGAGACTGCATCCAAAACTGGAGAGCTTGGGTTCTGGAGGGGAAAGCGGAATTCCTGGTGTAGCGGTGAAATGCGTAGATATCAGGAGGAACACCGGTGGCGAAGGCGGCTTTCTGGACAGATACCGACGCTGAGGCACGAAAGCTAGGGG |
| P-asv5819 | GCAAGCGTTGTTCGGAATCATTGGGCGTAAAGGGTATGTAGGCGGTGCGGTAAGTCTGGTGTGAAATCCCCAAGCTTAACTTGGGTTCCGCGCCAGAGACTGCCGTGCTCGAGTGTGGGAGAGGAAAGTGGAATTCCCGGTGTAGCGGTGAAATGCGTAGATATCGGGAGGAACACCCGTGGCGAAGGCGGCTTTCTGGACCACTACTGACGCTGAGATACGAAAGCCGGGGG |
| P-asv5820 | GCAAGCGTTATCCGGAATCACTGGGCGTAAGGGGTGCGTAGGCGGCCTAACAAGTCAGAGGTGAAAGGCATTGGCTCAACCTCGGAAATGCCTTCAATACTGGCTGTCTTGAGTACGGGAGAGGTGAGTGGAACTCCGAGTGTAGAGGTGAAATTCGTAGATATTCGGAAGAACACCAGTGGCGAAGGCGACTCACTGGCCCGTTACTGACGCTGAGGCGCGAAAGCGTGGGG |
| P-asv5821 | GCAAGCGTTGTCCGGATTTATTGGGTTTAAAGGGTGCGTAGGCGGCTTATTAAGTCAGTGGTGAAATACTCTAGCTCAACTAGAGGGGTGCCATTGATACTGATGAGCTTGAGTACAGATGAGGTAGGCGGAATTGACGGTGTAGCGGTGAAATGCTTAGATATCGTCAAGAACACCTATAGCGAAGGCAGCTTACTAGGCTGTAACTGACGCTGAGGCACGAAAGTGTGGGG |
| P-asv5822 | GCTAGCGTTGTTCGGAATTACTGGGCGTAAAGCGCACGCAGGCGGATTGATAAGTCGGGGGTGAAATCCCGGGGCTCAACCTCGGAATTGCCTTCGATACTGTTAGTCTTGAGTCCGGGAGAGGTGAGTGGAATTCCTAGTGTAGAGGTGGAATTCGTAGATATTAGGAAGAACACCAGTGGCGAAGGCGGCTCACTGGCCCGGTACTGACGCTCATGTGCGAAAGCGTGGGG |
| P-asv5823 | GCAAGCGTTGTCCGGATTTATTGGGTTTAAAGGGTGCGTAGGCGGTCCTATAAGTCAGTGGTGAAATACGGCAGCTTGACTGTCGAGGTGCCATTGATACTGCAGGACTAGAGTACTGATGAGGTAGGCGGAATGGACGGTGTAGCGGTGAAATGCTTAGATATCGTCCAGAACACCGATAGCGAAGGCAGCTTACTAAGGAGTAACTGACGCTGAGGCACGAAAGTGTGGGG |
| P-asv5824 | GCAAGCGTTGTCCGGAATTATTGGGCGTAAAGGGAGCGCAGGTGGACGGTAAGTCCGTCTTAAAAGGCAGGGGCTCAGCCCCTGTAAGGGATGGAAACTATCGATCTTGAGTGCCGGAGAGGAAAGCGGAATTCCCAGTGTAGCGGTGAAATGCGTAGATATTGGGAAGAACACCAGTGGCGAAGGCGGCTTTCTGGACGGCAACTGACACTGAGGCTCGAAAGCCAGGGG |
| P-asv5825 | GCAAGTGTTATCCGGAATTATTGGGCGTAAAGCGTCTGTAGGTGGTTTAATAAGTCAACTGTTAAATCTTGAGGCTTAACTTCAAAACAGCAGTCGAAACTATTAAACTAGAGTATAGTAGGGGTAAAGGGAATTTCCAGTGGAGCGGTGAAATGCGTAGATATTGGAAAGAACACCGATGGCGAAAGCACTTTACTGGGCTATTACTAACACTCAGAGACGAAAGCTAGGGT |
| P-asv5826 | GCCAGCGTTGTTCGGAGTCATTGGGCGTAAAGGGCGCGTAGGCGGCCGACCAAGTCGGAGGTGAAATCCCTCGGCTCAACTGAGGAACTGCCTCCGATACTGTTCGGCTTGAGTCCCGGAGAGGGTAGTGGAATTCCCGGTGTAGCGGTGAAATGCGTAGAGACCGGAAAGAACACCAGTGGCGAAGGCGGCTACCTGGACGGGTACTGACGCTGAAGCGCGAAAGCTAGGGG |
| P-asv5827 | GCGAGCGTTATCCGGAATCATTGGGTTTAAAGGGTCCGTAGGCGGGCTGATAAGTCAGAGGTGAAAGCGCTTAGCTCAACTAAGCAACTGCCTTTGAAACTGTCAGTCTTGAATGATTGTGAAGTAGTTGGAATGTGTAGTGTAGCGGTGAAATGCTTAGATATTACACAGAACACCGATAGCGAAGGCATATTACTAACAATTTATTGACGCTGATGGACGAAAGCGTGGGG |
| P-asv5828 | ACTAGCGTTGTCCGGAATTATTGGGCGTAAAGAGCTCGTAGGTGGTTTGTCGCGTCTGCTGTGGAAACGTGCCGCTTAACGGTGCGCGTGCAGTGGGTACGGGCGGACTAGAGTGCAGTAGGGGAGTCTGGAATTCCTGGTGTAGCGGTGAAATGCGCAGATATCAGGAGGAACACCGGTGGCGAAGGCGGGACTCTGGGCTGTTACTGACGCTGAGGAGCGAAAGCATGGGG |
| P-asv5829 | ACTAGTGTTATTCATCTTTATTAGGTTTAAAGGGTACCTAGACGGTAAATCAAGCCCTTGCAGAGCAAGATGGGACTAATTTACTAGAGTTACTTACAAGGAGGCTATAAAGTACTAATGGTGTAGAGATAAAATTTTGTAATACCAATTCTCTCTAGAGAAGTTATGGCACTGATAAAAGCGAAAGCATCTTCTTATGTGATAACTGACGTTGAGGGACGAAGGCTATGGAT |
| P-asv5830 | GCGAGCGTTGTTCGGAATTATTGGGCGTAAAGGGCATGTAGGCGGATAGGCAAGCTTGGTGTGAAATGTTACAGCTTAACTGTGAGACAGCATTGAGAACTGCAGATCTTGAATTACTGAAGGGTAACCAGAATTCCACGTGTAGGGGTGAAATCTGTAGATATGTGGAAGAATACCAATGGCGAAGGCAGGTTACCGGCAGATAATTGACGCTGAGGTGCGAAAGTGCGGGG |
| P-asv5831 | GCAAGCGTTAATCGGAATTACTGGGCGTAAAGCGCGCGTAGGCGGTCTGTTAAGTCGGATGTGAAAGCCCCGGGCTTAACCTGGGAATGGCATTCGATACTGGCGGACTGGAATACGGGAGAGGGAGGTAGAATTCCATGTGTAGCGGTGAAATGCGTAGATATATGGAAGAATACCAGTGGCGAAGGCGGCCTCCTGGCCTGATATTGACGCTGAGGTGCGAAAGCGTGGGG |
| P-asv5832 | GCAAGCGTTGTCCGGAATTATTGGGCGTAAAGAGCTCGTAGGCGGTTTGTCGCGTCTGTTGTGAAAACGCGAGGCTCAACCTCGCGCCTGCAGTGGGTACGGGCAAACTAGAGTGCGGTAGGGGAGATTGGAATTCCTGGTGTAGCGGTGGAATGCGCAGATATCAGGAGGAACACCGATGGCGAAGGCAGATCTCTGGGCCGTAACTGACGCTGAGGAGCGAAAGCATGGGG |
| P-asv5833 | GCTAGCGTTGTTCGGAATCACTGGGCGTAAATCGCACGTAGGCGGATATGTCAGTCAGGGGTGAAATCCCGGAGCTCAACTTCGGAACTGCCTTCAATACTGGCTGTCTTGAGTACGGGAGAGGTGAGTGGAACTCCGAGTGTAGAGGTGAAATTCGTAGATATTCGGAAGAACACCAGTGGCGAAGGCGACTCACTGGCCCGTTACTGACGCTGAGGCGCGAAAGCGTGGGG |
| P-asv5834 | GCGAGCGTTGTCCGGATTTATTGGGTTTAAAGGGTGCGTAGGCGGCCCGTTAAGTCCGGGGTGAAAGCCCGTTGCTCAACAACGGAACTGCCCTGGATACTGACGGGCTTGAGTACAGACGAGGTTGGCGGAATGGACCGAGTAGCGGTGAAATGCATAGATACGGTCCAGAACCCCGATTGCGAAGGCAGCTGACTAGGCTGATACTGACGCTGAGGCACGACAGCGTGGGG |
| P-asv5835 | GCTAGCGTTATCCGGATTTACTGGGCGTAAAGGGTGCGTAGGCGGTCTTTCAAGTCAGGAGTTAAAGGCTACGGCTCAACCGTAGTAAGCTCCTGATACTGTCTGACTTGAGTGCAGGAGAGGAAAGCGGAATTCCCAGTGTAGCGGTGAAATGCGTAGATATTGGGAGGAACACCAGTAGCGAAGGCGGCTTTCTGGACTGTAACTGACGCTGAGGCACGGAAGCGTGGGG |
| P-asv5836 | GCAAGCGTTATTCGGAATGACTGGGCGTAAAGGGTGCGTAGACGGTAGACTAAGTCCGTTGTTAAATTCTTCGGCCTAACCGGAGGTCAGCGATGGAAACTAGTCTGCTAGAGGGTGAGAGAGAGAAGTGGAATTCTCGGAGTAGCGGTAAAATGCGTAGATCTCGAGAGGAACACCAATGGCGAAGGCAGCTTCTTGGCTCATTCCTGACGTTGAGGCACGAAAGCGTGGGG |
| P-asv5837 | GCGAGCGTTGTCCGGAATTACTGGGCGTAAAGGGTGCGTAGGCGGTTTGGTAAGTTGGATGTGAAATACCCGGGCTTAACTTGGGGGCTGCATCCAATACTGTCGGACTTGAGTGCAGGAGAGGAAAGCGGAATTCCTAGTGTAGCGGTGAAATGCGTAGATATTAGGAGGAACACCGGTGGCGAAGGCGGCTTTCTGGACTGTAACTGACGCTGAGGCACGAAAGCGTGGGG |
| P-asv5838 | GCAAGCGTTGTCCGGAATTATTGGGCGTAAAGAGCTCGTAGGCGGTTGGTCGCGTCGGCTGTGAAATCCCGGAGCTCAACTCCGGGTCTGCAGTCGATACGGGCCGACTTGAGTGTTGCAGGGGAGACTGGAATTCCTGGTGTAGCGGTGAAATGCGCAGATATCAGGAGGAACACCGGTGGCGAAGGCGGGTCTCTGGGCAACAACTGACGCTGAGGAGCGAAAGCGTGGGG |
| P-asv5839 | GCAAGCGTTAATCAGAATTACTGGGCGTAAAGCGTACGTAGGCGGCGTGTTAAGCAAGATGTGAAAGCCCCGGGCTCAACCTGGGAATTGCATTTTGAACTGGCATGCTAGAGTCCTGAAGAGGGTGGTAGAATTTCCAGTGTAGCGGTGAAATGCGTAGATATTGGAAGGAATACCGGTGGCGAAGGCGGCCACCTGGTCAGAGACTGACGCTGAGGTACGAAAGCGTGGGG |
| P-asv5840 | GCGAGCGTTAATCGGAATTACTGGGCGTAAAGCGCACGCAGGCGGTTTGTTAAGCCAGATGTGAAAGCCCCGGGCTCAACCTGGGAGGGTCATTTGGAACTGGCAAGCTAGAGTCTTGGAGAGGGGAGTGGAATTCCAGGTGTAGCGGTGAAATGCGTAGAGATCTGGAGGAACATCAGTGGCGAAGGCGGCTCCCTGGCCAAAGACTGACGCTCATGTGCGAAAGTGTGGGT |
| P-asv5841 | GCGAGCGTTGTCCGGAATTATTGGGCGTAAAGAGCTCGTAGGCGGTGTGTCGCGTCGGCCGTGAAAACCTTCGGCTTAACCGGGGGCTTGCGGTCGATACGGGCATCACTTGAGTTCGGCAGGGGAGACTGGAATTCCTGGTGTAGCGGTGAAATGCGCAGATATCAGGAGGAACACCGGTGGCGAAGGCGGGTCTCTGGGCCGATACTGACGCTGAGGAGCGAAAGCGTGGGG |
| P-asv5842 | GCAAGCGTTATCCGGATTCACTGGGTTTAAAGGGTGCGTAGGCGGGTATGTAAGTCAGTGGTGAAATCTCGGAGCTTAACTCCGAAACTGCCATTGATACTATATATCTTGAATATTGTGGAGGTTTGCGGAATATGTCATGTAGCGGTGAAATGCTTAGATATGACATAGAACATCTATTGCGAAGGCAGCAGGCTACACATATATTGACGCTGAGGCACGAAAGCGTGGGG |
| P-asv5843 | GCAAGCGTTATCCGGAATCACTGGGCGTAAAGGGTGCGTAGGCGGCCTAACAAGTCAGAGGTGAAAGGCATAGGCTCAACCAATGTAAGCCTTTGAAACTGGGGAACTTGAGTGCAGAAGAGGAGAGCGGAATTCCACGTGTAGCGGTGAAATGCGTAGAGATGTGGAGGAACACCAGTGGCGAAGGCGGCTCTCTGGTCTGTAACTGACGCTGAGGCGCGAAAGCGTGGGG |
| P-asv5844 | CCGAGCGTTATCCGGATTTATTGGGTTTAAAGGGTGCGTAGGCGGCCTATTAAGTCAGGGGTGAAATACGGTGGCTCAACCATCGCAGTGCCTTTGATACTGATGGGCTTGAATCCATTTGAAGTGGGCGGAATAAGACAAGTAGCGGTGAAATGCATAAATATGTCTTAGAACTCCGATTGCGAAGGCAGCTCACTAAACTGATATTGACGCTGATGCACGAAAGCGTGGGG |
| P-asv5845 | CCGGGCGTTATCCGGATTTATTGGGTTTAAAGGGAGCGTAGGCCGTGGATTAAGCGTGTTGTGAAATGCAGGTGCTCAACGTCTGCACTGCAGCGCGAACTGGTCCACTTGAGTGTGCACAACGCAGGCGGAATTCGTCGTGTAGCGGTGAAATGCTTAGATATGACGAAGAACTCCGATTGCGAAGGCAGCTTGCGGGAGCACAACTGACGCTGAAGCTCGAAAGTGCGGGT |
| P-asv5846 | CCAAGCGTTAATCGGAATTACTGGGCGTAAAGCGTGCGCAGGCGGTTGTGCAAGACCGATGTGAAATCCCCGAGCTTAACTTGGGAATTGCATTGGTGACTGCACGGCTAGAGTGTGTCAGAGGGGGGTAGAATTCCACGTGTAGCAGTGAAATGCGTAGAGATGTGGAGGAATACCGATGGCGAAGGGGACTCTTTGGTCTGTAACTGACGCTGAGGCGCGAAAGCGTGGGG |
| P-asv5847 | CCAGGCGTTATCCAGATTTATTGGGTTTAAAGGGAGCGTAGGCGGATTGTTAAGTCAGCGGTTAAAGGGTGTGGCTCAACCATACATTGCCGTTGAAACTGGCGATCTTGAGTGCAGACAGGGATGCCGGAATTCGTGGTGTAGCGGTGAAATGCTTAGATATCACGAAGAACTCCGATCGCGAAGGCAGGTGTCCGGGCTGCAACTGACGCTGAGGCTCGAAAGTGTGGGT |
| P-asv5848 | GCAAGCGTTAATCGGATTTACTGGGCGTAAAGCGTGCGTAGGCGGTTCGTTAAGTCGATTGTGAAAGCCCTGGGCTTAACCTGGGAATTGCGGTCGATACTGGCGGACTAGAGAACGGTAGAGGGAGGCGGAACTCCAGGTGTAGCGGTGAAATGCGTAGATATCTGGAAGAACACCGATGGCGAAGGCAACCTCCTGGGCCTGTTCTGACGCTGAGGCACGAAAGCGCGGGT |
| P-asv5849 | GCGAGCGTTAATCGGAATTACTGGGCGTAAAGCGCGCGTAGGCGGTCTGATAAGCCGGTTGTGAAAGCCCCGGGCTCAACCTGGGAACGGCATCCGGAACTGTCAGGCTAGAGTGCAGGAGAGGAAGGTAGGATTTCCGGTGTAGCGGTGAAATGCGTAGAGATCGGGAGGAATACCAGTGGCGAAGGCGGCCTTCTGGACTGACACTGACGCAGAGGTGCGAAAGCGTGGGT |
| P-asv5850 | GCTAGCGTTGTTCGGAATTACTGGGCGTAAAGCGTGCGCAGGCGGCTTCTCAAGTCAGTGGTGAAAGCCCGGAGCTCAACTCCGGAATTGCCATTGAAACTGTGAGGCTTGAGGACGAGAGAGGTGAGTGGAATTCCCAGTGTAGAGGTGAAATTCGTAGATATTGGGAAGAACACCGGTGGCGAAGGCGGCTCACTGGCTCGTTTCTGACGCTCATGCACGATAGCGTGGGG |
| P-asv5851 | GCGAACGTTGTTCGGAATCACTGGGCTTAAAGGGCGCGTAGGCGGGCTGTCAAGTTTGGGGTGAAATCCCACGGCTCAACCGTGGAACTGCCTCAAATACTGACGGTCTCGAGGAAGGTAGGGGCGAGCGGAACTGATGGTGGAGCGGTGAAATGCGTTGATATCATCAGGAACTCCGGTGGCGAAGGCGGCTCGCTGGACCTTTTCTGACGCTGAGGCGCGAAAGCTAGGGG |
| P-asv5852 | TCTCTCTTTTCCCTTCCTCCCTTTTCTTAAAGGTTGCGTAGGCGGGTCTTTAAGTCAGGGGTGAAATCCTGGAGCTCAACTCCAGAACTGCCTTTGATACTGAAGATCTTGAGTTCGGGAGAGGTGAGTGGAACTGCGAGTGTAGAGGTGAAATTCGTAGATATTCGCAAGAACACCAGTGGCGAAGGCGGCTCACTGGCCCGATACTGACGCTGAGGCACGAAAGCGTGGGG |
| P-asv5853 | GCAAGCGTTATCCGGATTTATTGGGTTTAAAGGGTCCGTAGGCGGATCTGTAAGTCAGTGGTGAAATCTCACAGCTTAACTGTGAAACTGCCATTGATACTGCAGGTCTTGAGTAAGGTAGAAGTGGCTGGAATAAGTAGTGTAGCGGTGAAATGCATAGATATTACTTAGAACACCAATTGCGAAGGCAGGTCACTATGTCTTAACTGACGCTGATGGACGAAAGCGTGGGG |
| P-asv5854 | GCGAACGTTATTCGGAATCACTGGGCTTACAGGGTGCGTAGGCGGCTTCTCAGGTCCGATGTGAAATCCCCCAGCTCAACTGGGGAATGGCATTGGAAACCGGGAGGCTTGAGTGAGCTAGAGGTACACGGAACTTCCGGTGGAGCGGTGAAATGTGTTGAGATCGGAAGGAACACCAGAGGCGAAGGCGGTGTACTGGGGCTTGTCTGACGCTGAGGCACGAAAGCCAGGGG |

| | |
|---|---|
| P-asv5855 | GCAAGCGTTGTTCGGAATTACTGGGCGTAAAGGGCGCGTAGGCGGCCACCTAAGTCAGACGTGAAATCCCTCGGCTCAACCGGGGAACTGCGTCTGATACTGGGCGGCTTGAATTCGGGAGAGGGATGCGGAA<br>TTCCAGGTGTAGCGGTGAAATGCGTAGATATCTGGAGGAACACCGGTGGCGAAGGCGGCATCCTGGACCGAGATTGACGCTGAGGCGCGAAAGCCAGGGG |
| P-asv5856 | GCGAGCGTTATCCAGATTTATTGGGTTTAAAGGGAGCGTAGGCGGTCTTATAAGTCAGTGGTGAAATCTCGTTGCTTAACAACGACCGTGCCATTGATACTGTAGGACTTGAGTACAGTTGCCGTTGGCGGAATGT<br>GTCATGTAGCGGTGAAATGCATAGATATGACACAGAACACCGATTGCGAAGGCAGCTGACGAAACTGTAACTGACGCTGAGGCTCGAAAGCGTGGGG |
| P-asv5857 | GCAAGCGTTACTCGGAATTACTGGGCGTAAAGCGTGCGTAGGTGGTCGTTTAAGTCTGTTGTGAAAGCCCTGGGCTCAACCTGGGAACTGCAGTGGAAACTGGACGACTAGAGTGTGGTGGAGGGTAGCGGAAT<br>TCCTGGTGTAGCAGTGAAATGCGTAGAGATCAGGAAGAACATCCATGGCGAAGGCAGCTACCTGGACCAACACTGACACTGAGGCACGAAAGCGTGGGG |
| P-asv5858 | GCGAGCGTTGTCCGGATTTATTGGGCGTAAAGAGCTCGTAGGCGGCTTGTCGCGTCGACTGTGAAATCCCGCGGCTCAACTGCGGGTCTGCAGTCGATACGGGCAGGCTAGAGTTCGGTAGGGGAGACTGGAA<br>TTCCTGGTGTAGCGGTGAAATGCGCAGATATCAGGAGGAACACCGGTGGCGAAGGCGGGTCTCTGGGCCGATACTGACGCTGAGGAGCGAAAGCGTGGGG |
| P-asv5859 | GCAAGCGTTATCCGGATTCACTGGGTTTAAAGGGTGCGTAGGCGGGTAGGTAAGTCAGTGGTGAAATCCCAGGGCTTAGCCCTGGGACTGCCTGTGAGACCTCAGTACTAGAGTGTGAGAGGGGATAGTGGAAT<br>ACCCAGTGTAGCGGTGAAATGCGTAGAGATTGGGTGGAACACCGGTGGCGAAGGCGGCTATCTGGCTCACAACTGACGATCAGGCACGAAAGCGTGGGG |
| P-asv5860 | GCAAGCGTTATCCGGAATCATTGGGTTTAAAGGGTCCGTAGGCGGTTTAATAAGTCAGTGGTGAAAGCCCATCGCTCAACGATGGAACGGCCATTGATACTGTTAAACTTGAATTATTAGGAAGTAACTAGAATAT<br>GTAGTGTAGCGGTGAAATGCTTAGAGATTACATGGAATACCAATTGCGAAGGCAGGTTACTACTAATATATTGACGCTGATGGACGAAAGCGTGGGG |
| P-asv5861 | GCTAGCGTTGTTCGGAATTACTGGGCGTAAAGCGCACGTAGGCGGATCGATCAGTCAGGGGTGAAATCCCAGGGCTTAACCCTGGAACTGCCTTTGATACTGTCGATCTAGAGTATGGAAGAGGTGAGTGGAATT<br>CCGAGTGTAGAGGTGAAATTCGTAGATATTCGGAGGAACACCAGTGGCGAAGGCGGCTCACTGGTCCATTACTGACGCTGAGGTGCGAAAGCGTGGGG |
| P-asv5862 | GCAAGCGTTAATCGGAATTACTGGGCGTAAAGCGCGCGTAGGCGGTTAGTTAAGTCAGATGTGAAATCCCCGGGCTCAACCTGGGAATGGCCTCTGATACTGGCTAGCTAGAGTTTGGAAGAGGGAAGTAGAAT<br>TTCAGGTGTAGCGGTGAAATGCATAGAGATCTGAAGGAATACCAGTGGCGAAGGCGACTTCCTGGTCCAGAACTGACGCTGAGGTGCGAAAGCGTGGGT |
| P-asv5863 | GCAAGCGTTGTCCGGAATTACTGGGTGTAAAGGGAGCGCAGGCGGGACAGCAAGTTGGATGTGAAATACCGAGGCTTAACCACGGAGCTGCATCCAAAACTGTTGTTCTTGAGTGAAGTAGAGGTAGGCGGAAT<br>TCCGAGTGTAGCGGTGAAATGCGTAGATATTCGGAGGAACACCAGTGGCGAAGGCGGCCTACTGGGCTTTAACTGACGCTGAGGCTCGAAAGTGTGGGG |
| P-asv5864 | TCAAGTGATGTCCAATATTATTGGGCCTAAAGCGTCCGTAGCTGGCCGCGCAAGTCCGTCGGGAAATCCACCTGCCCAACAGGTGGGCGTCCGGCGGAAACTGTGTGGCTTGGAACCGGAAGGCTCAGAGGGT<br>ACGTCCGGGGTAGGAGTGAAATCCCGTAATCCCGGACGGACCACCGATGGCAAAGCACCTCGAGAGGACTGATCCGACAGTGAGGGACGAAAGCTGGGGT |
| P-asv5865 | GCAAGCGTTGTCCGGAATTATTGGGCGTAAAGCGCGCGCAGGCGGTTTTTTAAGTCTGATGTATAAGCCCAAGGCTCAACTTTGGGAGGCATTGGAAACTGGGAGACTGGAGTGTAGGAGAGGAAAGTGGAATT<br>CCACGTGTAGCGGTGAAATGCGTAGAGATGTGGAGGAACACCAGTGGCGAAGGCGACTTTCTGGCCTATAACTGACGCTGAGGCGCGAAAGCGTGGGG |
| P-asv5866 | ACGAGTGGCAATCGCGTTTATTGGGCCTAAAGCGTCCGTAGCAGGTTTGTTAAATTTCTTGTGAAATTGCCAGGCTCAACTTGGCAGTGTGCAGGGAACACTGGCAGACTAGGGACCGGGAAGGGTCAGAGGTAT<br>TCCTGAGGGAGCGGTAAAATGCTATAATCTCAGGAGGACCACCTGTGGCGAAGGCGTCTGACTAGAACGGATCCGACTGTGAGGGACGAAAGCCAGGGG |
| P-asv5867 | GCAAGCGTTAATCGGAATTACTGGGCGTAAAGCGCGCGCAGGCGGTTGTGTAAGTCAGATGTGAAATCCCCGGGCTCAACCTGGGAATTGCATTTGAGACTGCACGGCTAGAGTGTGTCAGAGGGGGGTAGAAT<br>TCCACGTGTAGCAGTGAAATGCGTAGAGATGTGGAGGAATACCGATGGCGAAGGCAGCCCCCTGGGATAACACTGACGCTCATGCACGAAAGCGTGGGG |
| P-asv5868 | GCAAGCGTTGTTCGGAATTACTGGGCGTAAAGGGCGCGTAGGCGGCTTAGGAAGTCATGGGTGAAAGGCCTCGGCTCAACCGAGGAATTGCCTGTGAAACCACGGAGCTAGAGTGCTGGAGAGGAAAGCGGAA<br>TTCCCAGTGTAGCGGTGAAATGCGTAGATATTGGGAGGAACATCTGAGGCGAAGGCGGCTTTCTGGACAGACACTGACGCTGAGGCGCGAAAGCCAGGGG |
| P-asv5869 | GCAAGCGTTATCCGGAATTATTGGGCGTAAAGAGCTCGTAGGCGGTTTGTCGCGTCTGCTGTGAAATCTGGGGGCTCAACCCCCAGCCTGCAGTGGGTACGGGCAGACTAGAGTGCGGTAGGGGAGATTGGAA<br>TTCCTGGTGTAGCGGTGGAATGCGCAGATATCAGGAGGAACACCGATGGCGAAGGCAGATCTCTGGGCCGTAACTGACGCTGAGGAGCGAAAGCGTGAGG |
| P-asv5870 | GCAAGCGTTAATCGGAATTACTGGGCGTAAAGCGCGCGTAGGTGGCTGAGTAAGATGGGTGTGAAATCCCCGGGCTTAACCTGGGAACTGCATCCATAACTGCTCGGCTAGAGTACGATAGAGGGTAGTGGAAT<br>TTCCTGTGTAGCGGTGAAATGCGTAGATATAGGAAGGAACACCAGTGGCGAAGGCGACTACCTGGACTGATACTGACACTGAGGTGCGAAAGCGTGGGG |
| P-asv5871 | GCAAGCGTTAATCAGAATTATTGGGCGTAAAGCGAGTGCAGACGGTTACTTAAGCCAGATGTGAAATCCCCGAGCTTAACTTGGGAATTGCATTGGTGACTGCACGGCTAGAGTGTGTCAGAGGGGGGTAGAATT<br>CCACGTGTAGCAGTGAAATGCGTAGAGATGTGGAGGAATACCGATGGCGAAGGCAGCCCCCTGGGATAACACTGACGCTCATGCACGAAAGCGTGGGG |
| P-asv5872 | GCTAGCGTTGTCCGGAATCACTGGGCGTAAAGGGTTCGCAGGCGGAAATGCAAGTCAGGTGTAAAAGGCAGTAGCTTAACTACTGTAAGCATTTGAAACTGCATATCTTGAGAAGAGTAGAGGTAAGTGGAATTT<br>TTAGTGTAGCGGTGAAATGCGTAGATATTAAAAAGAATACCGGTGGCGAAGGCGACTTACTGGGCTCATTCTGACACTGAGGAACGAAAGCGTGGGT |
| P-asv5873 | GCGAGCGTTAATCGGAATTACTGGGCGTAAAGCGCGCGTAGGTGACTTGATAAGCCGGTTGTGAAAGCCCCGAGCTTAACTTGGGAATTGCATTGGTGACTGCACGGCTAGAGTGTGTCAGAGGGGGGTAGAAT<br>TCCACGTGTAGCAGTGAAATGCGTAGAGATGTGGAGGAATACCGATGGCGAAGGCAGCCCCCTGGGATAACACTGACGCTCATGCACGAAAGCGTGGGG |
| P-asv5874 | GCGAGCGTTATTCGGATTTATTGGGCGTAAAGGGTTCGTAGGAGGTTGGATAAGTTTGATGTGAAAGCCCGTGGCTCAACCACGGAACTGCATTGGAAACTGTCCAACTTGAGGTTCGGAGAGGAAAGCAGAATT<br>CCCGGTGTAGCGGTGGAATGCGTAGATATCGGGAGGAATACCGACAGCGAAGGCAGTTTTCTGGACGACACCTGACTCTGAGGAACGAAAGCCTGGGG |
| P-asv5875 | GCAAGCGTTATCCGGAATTATTGGGCGTAAAGCGTCCGTAGGCGGCTAGTCAAGTTTGCTGTCAAAGGTCGGGGCTCAACCCTGGATAGGCAGTGGAAACTGATTAGCTAGAGAGCGGCAGGGGTAGAGGGAAT<br>TCCCAGTGTAGCGGTGAAATGCGTAGATATTGGGAAGAACACCAGTGGCGAAGGCGCTCTACTGGGCCGTTACTGACGCTGAGGGACGAAAGCTAGGGG |
| P-asv5876 | GCGAGCGTTGTCCGGAATTATTGGGCGTAAAGAGCGTGTAGGCGGTTTGGTAAGTCTGCCGTGAAAACCCGGGGCTCAACCCCGGGCGTGCGGTGGATACTGCCAGGCTAGAGGATGGTAGAGGCGAGTGGA<br>ATTCCCGGTGTAGTGGTGAAATGCGCAGATATCGGGAGGAACACCAGTAGCGAAGGCGGCTCGCTGGGCCACTCCTGACGCTGAGACGCGAAAGCTAGGGG |
| P-asv5877 | GCGAACGTTATTCGGAATCACTGGGCTTAAAGCGCGTGTAGGCGGGTCGGCACGTCCGTGGCTGAAAGCCCCCGGCTCAACCGGGGAAGTGGCGCGGATACGGCCGATCTGGAGGGGGGTAGGGGGACCTG<br>GAACTTCCGGTGGAGCGGTGAAATGCGTTGAGATCGGAAGGAACGCCCGTGGCGAAAGCGAGGTCCTGGACCCTTTCTGACGCTGAGACGCGAAAGCCAGGGG |
| P-asv5878 | GCTAGCGTTGTTCGGATTTACTGGGCGTAAAGCGCACGTAGGCGGGTCGTTAAGTCAGGGGTGAAATCCTGGAGCTCAACTCCAGAACTGCCTTTGATACTGGCGACCTAGAGTCCGGAAGAGGTGAGTGGAAC<br>TCCTAGTGTAGAGGTGGAATTCGTAGATATTAGGAAGAACACCAGTGGCGAAGGCGGCTCACTGGTCCGGTACTGACGCTGAGGTGCGAAAGCGTGGGG |
| P-asv5879 | GCGAGCGTTAATCGGAATTACTGGGCGTAAAGGGTGCGTAGGTGGTTTATTAAGTTAGGTGTGAAAGTCCTGGGCTCAACCTGGGGATTGCGCCTGATACTGGTAGGCTAGAGTATTGTAGAGGGAAGTGGAAT<br>TTCCGGTGTAGCGGTGAAATGCGTAGATATCGGAAGGAACATCAGTGGCGAAGGCGACTTCCTGGACAACTACTGACACTGAGGCACGAAAGCGTGGGG |
| P-asv5880 | GCTAGCGTTATTCGGAATTACTGGGCGTAAAGCGCACGTAGGCGGCTTTGTAAGTTAGAGGTGAAAGCCTGGAGCTCAACTCCAGAATTGCCTTTAAGACTGCATCGCTTGAATCCAGGAGAGGTGAGTGGAATT<br>CCGAGTGTAGAGGTGAAATTCGTAGATATCGGGAGGAACATCAGTGGCGAAGGCGACCACCTGGACCAAAACTGACGCTGAGGTGCGAAAGCGTGGGG |
| P-asv5881 | GCAAGCGTTGTCCGGAATTATTGGGTTTAAAGGGTGCGCAGGCGGGTCTTTAAGTCAGAGCTGAAATCCCACAGCTTAACTGTGGAACTGGCTTTGATACTGGAGGTCTTGAGTACACTAGAGGTATGCGGAATT<br>TATGGTGTAGCGGTGAAATGCATAGATACCATAAAGAACACCGATAGCGAAGGCAGCATACTTGGGTGTAACTGACGCTGAGGCACGAAAGCGTGGGT |
| P-asv5882 | GCAAGTGGTGGCCACGATTATTGGGCTTAAAGTGTTCGTAGCCGGTCTGACGAGTCTACTGTGAAATCTAGCGGCTCAACCGTTAGGAGCGCAGTGGATACCACCAGACTAGGGGCCGGAGGGCGTCGAGGCT<br>ATTCCTGGAGTAGGGGTGAAATCCCAAAATTCCAGGAGGAGCACCAGTGGCGAAAGCGCTCGGCGACTACGGGCCCGACGGTGAGGAACGAAAGCCAGGGG |
| P-asv5883 | GCTAGCGTTGTTCGGAATTACTGGGCGTAAAGCGCGCGTAGGCGGACTGGTCAGTCAGGGGTGAAAGCCCGGGGCTCAACCTCGGAAATGCCTTTGATACTGCCAGTCTAGAGTCCGGCAGAGGTGAGTGGAA<br>TTCCTAGTGTAGAGGTGAAATTCGTAGATATTAGGAAGAACACCAGTGGCGAAGGCGGCTCACTGGGCCGAGACTGACGCTGAGGCGCGAAAGCGTGGGG |
| P-asv5884 | GCGAGCGTTAATCGGATTTACTGGGCGTAAAGCGTGCGTAGGCGGCTTTTTAAGTCGGATGTGAAATCCCTGAGCTTAACTTAGGAATTGCATTCGATACTGGGAAGCTAGAGTATGGGAGAGGATGGTAGAATT<br>CCAAGTGTAGCGGTGAAATGCGTAGAGATTTGGAGGAATACCAGTGGCGAAGGCGGCCCCCTGGCACAATACTGACGCTCAAGTGCGAAAGCGTGGGG |
| P-asv5885 | GCGAGCGTTGTCCGGATTCACTGGGCGTAAAGGGCTCGTAGGCGGCCTGTCGCGTCGGCTGTGAAAGTCCGGAGCTCAACTCCGGGATTGCAGCCGATACGGGCAGGCTAGAGGGCGGTAGAGGGGAGTGG<br>AATTCCCGGTGTAGCGGTGAAATGCGCAGATATCGGGAGGAACACCAGTGGCGAAGGCGGCTCCCTGGGCCGTCCCTGACGCTGAGGAGCGAAAGCGTGGGG |
| P-asv5886 | GCAAGCGTTGTTCGGAATCACTGGGCTTAAAGCGTCCGCAGGCGGATTTGCAAGTCAGAAGTGAAATCCCCGGGCTTAACCCGGGAACTGCTTTTGAAACTGCAAATCTAGAATATTGGAGAGGTTGGGAGAATT<br>CCGAGTGTAGAGGTGAAATTCGCAGATATTCGGAGGAATACCAGTGGCGAAGGCGCCCAACTGGACAATCATTGACGCTCAGGGACGAAAGCGTGGGG |
| P-asv5887 | GCTAGCGTTGTTCGGAATTACTGGGCGTAAAGGGCGTGTAGGCGGCGGATTTAGTTAGGCGTGAAAGCCCCGGGCTCAACCCGGGAACTGCGCTTAATACTGATTTGCTAGAGTCCGGGAGAGGCGAGTGGAA<br>TATCCAGTGTAGAGGTGAAATTCGTAGATATTGGATGGAACACCAGTGGCGAAGGCGGCTCGCTGGACCGGAACTGACGCTGAGGCGCGAAAGCGTGGGG |
| P-asv5888 | GCTAGCGTTGTTCGGAATCACTGGGCGTAAAGGGCGCGTGGGCGGCCGATCAAGTCGGGGGTGAAAGCCTGTGGCTCAACCACAGAATTGCCTTCGATACTGTTCGGCTCGAGACCGGAAGAGGTCAGTGGAA<br>CTGCGAGTGTAGAGGTGAAATTCGTAGATATTCGCAAGAACACCAGTGGCGAAGGCGGCTGACTGGTCCGGATCTGACGCTGGGGCGCGAAAGCGCGGGG |
| P-asv5889 | GCAAGCGTTGTTCGGAATCACTGGGCTTAAAGCGTGCGTAGGCGGACTTGCAAGTCAGGAGTGAAATCCCGGGGCTCAACCTCGGAATTGCTTTTGAAACTATGAGTCTAGAAACCGGTAGGGGCTAGCGGAAT<br>TCCGAGTGTAGAGGTGAAATTCGCAGATATTCGGAGGAACACCAGTGGCGTAGGCGGCTGGCTGGACCGGTATTGACGCTGAGGCACGAAAGCGTGGGG |
| P-asv5890 | CCTAGCGTTGTTCGGAATTATTGGGCGTAAAGCGGGTGTAGGTGGCTTAGCAAGTCAGATGTGAAAGCCCTGGGCTCAACCCAGGAAGTGCGTCTGATACTGCTTTGCTAGAGCACGGTAGAGGGAACTGGAAT<br>TCCAGGTGTAGGGGTGAAATCCGTAGATATCTGGAGGAATACCAATGGCGAAGGCAGGTTCCTGGTCCGTTGCTGACACTCAGACCCGAAAGCGTGGGG |
| P-asv5891 | CCAAGCGTTAATCGGAATTACTGGGCGTAAAGCGTGCGCAGGCGGCTCAGAAAGTTGGGTGTGAAATCCCTCGGCTCAACCGAGGAATTGCGCCCAAAACTACTGAGCTTGAGGGAGTTAGAGGTGAGCGGAAC<br>TCACGGTGGAGCGGTGAAATGCGTTGATATCGTGAGGAACACCGGAGGCGAAAGCGGCTCACTGGGGCTTCACTGACGCTGAGGCACGAAAGCTAGGGG |
| P-asv5892 | GCAAGCGTTGTTCGGAATTATTGGGCGTAAAGCGCGTGTAGGCGGCCATCTAAGTCTGATGTGAAATCCCGGGGCTTAACCCCGGAAGTGCATTGGATACTGGATGGCTAGAGGGCGAGAGAGGGAAACAGAAT<br>TCCTGGTGTAGGGGTGAAATCCGTAGATATCAGGAGGAATACCGGCGGCGAAGGCGGTTTCCTGGCTCGTTACTGACGCTGAGACGCGAAAGCGTGGGG |
| P-asv5893 | GCAAGCGTTGTTCGGAGTTACTGGGCGTAAAGAGTGCGTAGGCGGTTGACCAAGTCTGTCGTGAAATCTCCTGGCTTAACTGGGAGGGTGCGGTGGAAACTGGTTGGCTTGAGAGTGGGAGAGGTAAGCGAAA<br>TTCCTGGTGTAGCGGTGAAATGCGTAGATATCAGGAGGAACACCGGTGGTGTAGACGGCTTACTGGACCACGACTGACGCTGAGGCACGAAAGCGTGGGT |

| | |
|---|---|
| P-asv5894 | GCGAGCGTTGTCCGGAATTACTGGGCGTAAAGGATGCGTAGGTGGATATTTAAGTGGGATGTGAAATCCCCGGGCTCAACCCGGGAACTGCATTCCAAACTGGATATCTAGAGTGCAGGAGAGGAAAGCGGAATTCCTAGTGTAGCGGTGAAATGCGTAGATATTAGGAGGAACACCAGTGGCGAAGGCGGCTTTCTGGACTGTAACTGACACTGAGGCATGAAAGCGTGGGT |
| P-asv5895 | GCAAGCGTTATCCGGAATCATTGGGTTTAAAGGGTCCGCAGGCGGTCTTATAAGTCAGTGGTGAAAGCCTACAGCTTAACTGTAGAACTGCCATTGATACTGTAGGACTTGAATTCGGTCGAAGTGGGCGGAATATGTCATGTAGCGGTGAAATGCTTAGATATGACATAGAACACCGATAGCGAAGGCAGCTCACTAGGCCTGGATTGACGCTCAGGGACGAAAGCGTGGGT |
| P-asv5896 | GCGAGCGTTGTCCGGAATTATTGGGCGTAAAGGGTGCGTAGGCGGTTTTTTAAGTCTGCATTGTAAGTTCAGTGCTTAACGCTGAAATTGGTGCGGAAACTGGAAGACTAGAGTACTGTAGGGGAAAGCGGAATGTCCGGTGCAGCAGTGAAATGCGTAGAGATCGGATGGAACACCGAAAGCGAAGGCAGCTTTCTGGGCAGTAACTGACGCTGAGGCACGAAAGCGTGGGG |
| P-asv5897 | GTAAGCGTTAATCGGAATTACTGGGCGTAAAGCGCGCGTAGGCGGTTTGATAAGTTGGATGTGAAAGCCCCGGGCTCAACCCGGGAAGTGCATTTGAAACTGGTTATCTTGAGTGCAGTAGAGGAAGGTGGAATTCCAAGTGTAGTGGTGAAATACGTAGATATTTGGAGGAACATCGGTGGCGAAGGCGACCTTCTGGACTGTTACTGACGCTGAGACCCGAAAGCGTGGGG |
| P-asv5898 | GCTAGCGTTGTTCGGAATTACTGGGCGTAAAGGGTGCGTAGGTTGTTTAATAAGTTAATACAAAAATCCAAGACAATTAATTCTTGTAAAGTGATTAATACTATTAAACTAGAGTGTAAAAGAGGATAGTAGTATTCTTAGTGTAGAGGTAAAATTCGTAAATATTAAGAGGACTATCAACAGGCGAAGGCAACTATCTGGTTTATTCTGACACTGAGGCACGAAAGCATGGGG |
| P-asv5899 | GCTAGCGTTGTTCGGAATTACTGGGCGTAAAGGGCGCGTAGGCGGCCTTGTCAGTCAGAAGTGAAAGCCCCGGGCTCAACCTGGGAATTGCTTTTGATACTGCAAGGCTAGAGTTTCGGAGAGGATGGTGGAATTCCCAGTGTAGAGGTGAAATTCGTAGATATTGGGAAGAACACCGGTGGCGAAGGCGGCCATCTGGACGAACACTGACGCTGAGGCGCGAAAGCGTGGGG |
| P-asv5900 | GCAAGCGTTAATCGGAATTACTGGGCGTAAAGGGCGCGTAGGCGGTCTGATAAGCGGGTTGTGAAAGCCCCGGGCTCAACCTGGGAACGGCAATCCGAACTGTCAGGCTAGAGTGCAGCAGAGGGCAGTGGAATTCCCGGTGTAGCGGTGAAATGCGTAGATATCGGGAGGAACACCAGTGGCGAAGGCGACTGCCTGGGCTGACACTGACGCTGAGGTGCGAAAGCGTGGGT |
| P-asv5901 | GCAAGCGTTATCCGGAATCATTGGGCGTAAAGCGTTCGTAGGTGGTTTGGAAAGTTTGCGGTTAAATCTGGTCGCTCAACGATCAAATCGCTGCAAAAACTCCCAAGCTAGAGATTGGCAGAGGCAACTGGAATTCCACATGTAGGGGTAAAATCCGTTGATATGTGGAGGAACACCAAAAGCGAAGGCAGGTTGCTGGGCCATTTCTGACACTGAGGAACGAAAGCGTGGGG |
| P-asv5902 | GCAAGCGTTGTTCGGAATCACTGGGCTTAAAGCGTGCGTAGGCGGATTTGCAAGTCAGGAGTGAAATCCCGGGGCTCAACCCCGGAATTGCTCTTGAAACTGCAAATCTAGAATCCTGGAGAGGTTGGGAGAATTCCTAGTGTAGAGGTGAAATTCGCAGATATTAGGAGGAATACCAGTGGCGTAGGCGCCCAACTGGACAGGTATTGACGCTCAGGCACGAAAGCGTGGGG |
| P-asv5903 | GCAAGCGTTAATCGGAATTACTGGGCGTAAAGCGTGCGCAGGCGGTTTTGTAAGACAGGCGTGAAATCCCCGGGCTCAACCTAGGAACCGCCTTTGAAACTGGAGAGCTAGAATCTTGGAGAGGCGAGTGGAATTCCGAGTGTAGAGGTGAAATTCGTAGATATTCGGAAGAACACCAGTGGCGAAGGCGACTCGCTGGACAAGTATTGACGCTGAGGTGCGAAAGCGTGGGG |
| P-asv5904 | GCGAACGTTATTCGGAATCACTGGGCTTAAAGGGTGCGTAGGCGGCGAACTAAGTAGGGTGTGAAATCCCATCGCTCAACGATGGAACTGCGCTCTAAACTGGTTTGCTTGAGTAAGATAGGGGTGTGCGGAACTTCAAGTGGAGCGGTGAAATGTGTAGATATTTGAAGGAACACCGGTGGCGAAAGCGGCACACTAGGTCTTAACTGACGCTGAGGCACGAAAGCCAGGGT |
| P-asv5905 | GCAAGCGTTAATCAGAATTACTGGGCGTAAAGCGTGCGCAGGCGGTGATGTAAGACAGGCGTGAAATCCCGGGGCTCAACCCCGGAACTGCCCTTGAAACTGGGAAGCTAGAATCTTGGAGAGGCGAGTGGAATTCCGAGTGTAGAGGTGAAATTCGTAGATATTCGGAAGAACACCAGTGGCGAAGGCGACTCGCTGGACAAGTATTGACGCTGAGGTGCGAAAGCGTGGGG |
| P-asv5906 | GCTAGCGTTGTTCGGAATTACTGGGCGTAAAGCGCGCGTAGGCGGCGCTGTAAGCCAGGGGTGAAATCCCGGAGCTCAACTCCGGAATTGCCTTTGGAACTACAGTGCTAGAGTATTGGAGGGGAGTGTGGAATTCTTAATGTAGGGGTGAAATCCGTAGATATTAAGAGGAACACCGGTGGCGAAGGCGACACTCTGGACAATTACTGACGCTGAGGTGCGAAAGCGTGGGG |
| P-asv5907 | GCTAGCGTTAATCGGAATTACTGGGCGTAAAGGGCCTGTAGGTGGTTTCTTAAGTTAGATGTGAAAGCCCTGGGCTTAACCTGGGAACTGCATCTAAAACTGGGTGACTCGAGTATGGTAGAGGGAAGTGGAATTTCTAGTGTAGCGGTGAAATGCGTAGATATTAGAAAGAACACCAGTGGCGAAGGCGACTTCCTGGACCAATACTGACACTGAGGGGCGAAAGCGTGGGG |
| P-asv5908 | CCGAGCGTTAGGCGGAATCACTGGGCTTAAAGGGTGCGTAGGCGGATCAGTAAGCATTTTGTGAAATCCCTCGGCTCAACCGAGGAATTGCAGAGTGAACTGCTGATCTTGAGGTAGGCAGAGGTCATCGGAACGTACGGTGGAGCGGTGAAATGCGTAGATATCGTACGGAACGCCGAAGGGGTAGCCAGGTGACTGGGCCTTACCTGACGCTGAGGCACGAAAGCGTGGGG |
| P-asv5909 | GCAAGCGTTATCCGGATTCATTGGGTTCAAAGGGTGCGCAGGCGGATTATTAAGTCAGTGGTGAAATCCTACAGCTCAACTGTAGAACTGCCATTGATACTGATAATCTTGAGTATATTTGAAGTGGGCGGAATGTAGCATGTAGCGGTGAAATGCATAGATATGCTACAGAACACCAATTGCGAAGGCAGCTCACTAAACTATAACTGACGCTCATGCACGAAAGCGTGGGG |
| P-asv5910 | CCGAGCGTTATCCGGAATTACTGGGCGTAAAGCGTCTGTAGCCGGATGTATAAGTCTGTCTTTAAATACTCTGATTCAATCAGAGAAAAGGGATGGATACTGTATGTCTAGAGGATTTTAGAGGTTAGTGGAATTTCCGGTGGAGCGGTGAAATGCGTTGATATCGGAAGGAATGCCGAAAGCGAAAGCAGCTAACTATAAAATATCTGACGGTCAGAGACGACAGCTTGGGG |
| P-asv5911 | GCAAGCGTTGTCCGGATTTATTGGGCGTAAAGAGTTCGTAGGCGGTTATTTAAGTCTGGTGTTAAATACTGTGGCCCAACCGCAGAACTGCATTGGATACTGGATGACTTGAGTGCGGTAGAGGCGAGTGGAATTCCCAGTGTAGCGGTGAAATGCGTAGATATTGGGAAGAACACCAGTGGCGTAAGCGACTCGCTGGCCCGTAACTGACGCTGAGGAACGAAAGCCAGGGG |
| P-asv5912 | GCAAGCGTTATCCGGAATTATTGGGCGTAAAGCGTCCGTAGGCGGTTTTTTAAGTCTGTGGTTAAATCCCGGGGCTCAACTCCGGAATCGTCACGGAAACTAGAAAACTTGAGATGTGCAGAGGCAGTTGGAATTTCGGAAGTAGGGGTAAAATCCGTAGATCTCCGAAGGAACACCAAACGCGAAGGCATTCTGCTGGGCACATTCTGACGCTGAGGGACGAAAGCGTGGGG |
| P-asv5913 | GCTAGCGTTGTTCGGAATTACTGGGCGTAAAGGGTGCGTAGGCGGCTTCGTAAGTTTGGTGTGAAAGACCTGGGCTTAACCTGGGAATTGCATCAAAAACTGCGTTGCTTGAGAACGAGAGAGGAAAACGGAATTGCGAGTGTAGAGGTGGAATTCGTAGATATTCGCAAGAACACCAATGGCGAAGGCAGTTTTCTGGCTCGATATCTGACGCTGAGGCACGAAAGCGTGGGG |
| P-asv5914 | GCAAGCGTTATCCGGATTTACTGGGTTTAAAGGGTGCGTAGGCGGTCTATTATGTCAGTGGTGAAAGCCCGGGGCTCAACTCCGGAACTGCCATTGAAACTGATAGACTTGAGTTTGGGAGAGGTAGGCGGAATACAGCATGTAGCGGTGAAATGCATAGATATGCTGTAGAACACCAATTGCGAAGGCAGCTTACTATACCAATACTGACGTTCAGGCACGAAAGCGTGGGG |
| P-asv5915 | GCGAACGTTAATCGGAATTACTGGGCGTAAAGCGCGCGTAGGCGGCTCGTTAAGTGGGATGTGAAAGCCCCGGGCTCAACCCGGGAATTGCATTTCAAACTGGCGAGCTCGAGTGCAGTAGAGGGAGGTGGAATTCCAGGTGTAGCGGTGAAATGCGTAGATATCTGGAGGAACATCGGTGGCGAAGGCGGCCTCCTGGACTGACACTGACGCTGAGGCGCGAAAGCGTGGGG |
| P-asv5916 | CCAAGCGTTGTCCGGAATCACTGGGCGTAAAGGATGCGCAGGCGGGGCAGCAAGTCAGAGGTGAAATCCTGCGGCTTAACCGCAGACGTGCCTTTGAAACTGCTGCTCTTGAGTCCTGGAGAGGTTGCCGGAATTCGTGGTGTAGCGGTGAAATGCGTAGATATCACGAGGAACACCAGAGGCGAAAGCGGGCAACTGGACAGGTACTGACGCTGAGGCATGAGAGCGTGGGG |
| P-asv5917 | GCAAGCGTTAATCGGAATTACTGGGCGTAAAACGTGCGCAGGCGGTTTTGTAAGTCTGACGTGAAATCCCCGGGCTCAACCTGGGAATTGCAGTCGATACTGGCGGACTAGAGAACGGTAGAGGGAGGCGGAACTCCAGGTGTAGCGGTGAAATGCGTAGATATCTGGAAGAACACCGATGGCGAAGGCAACCTCCTGGGCCTGTTCTGACGCTGAGGCACGAAAGCGTGGGT |
| P-asv5918 | GCGAGCGTTAATCGGAATTACTGGGCGTAAAGAGCGCGTAGGTGGTCTACTAAGTCATGTGTGAAAGCCCCGAGCTTAACTTGGGAATTGCACGTGATACTGGTAGTCTAGAGTATGGTAGAGGGAAGTGGAATTTCCGGTGTAGCGGTGAAATGCATAGATATCGGAAGGAACACCAGTGGCGAAGGCGACTTCCTGGCCAATACTGACACTGAGGCGCGAAAGCGTGGGG |
| P-asv5919 | CCAAGCGTTGTTCGGATTTACTGGGCGTAAAGCGTGCGTAGGAGGCTGGGAAAGTCTGATGTGAAATGTCACCGCTCAACGGTGAAAGTGCATTGGAAACTATTCAGCTCGAGGATCGGAGAGGAAAGCGGAATTCCAGGTGTAGCGGTGAAATGCGTAGATATCTGGAGGAACACCAACGGCGAAGGCAGCTTTCTGGACGGTTCCTGACTCTGAAGCACGAAGGCCAGGGG |
| P-asv5920 | GCGAGCGTTGTCCGGAATTACTGGGCGTAAAGGGCGTGTAGGCGGCCTTTTAAGTCAGACGTGAAATCCCTGGGCTTAACCCAGGACCTGCGTTTGAAACTGGAGGGCTTGAGGACCGGAGAGGGTAGTGGAATTCCCAGTGTAGCGGTGAAATGCGTAGATATTGGGAGGAACACCAGTGGCGAAGGCGACTGCCTGGACGGCACCTGACGCTGAGGCGCGAAAGCCAGGGG |
| P-asv5921 | ACGAGCGTTATCCGGATTTATTGGGCGTACAGCGCGTCTAGGTGGTTATATAAGTCTGATGTGAAAATGCAGGGCTCAACTCTGTATTGCGTTGGAAACTGTATAACTAGAGTACTGGAGAGGTAAGCGGAACTACAAGTGTAGAGGTGAAATTCGTAGATATTTGTAGGAATGCCGATGGGGAAGCCAGCTTACTGGACAGATACTGACGCTGAAGCGCGAAAGCGTGGGT |
| P-asv5922 | GCTAGCGTTGTTCGGAATTACTGGGCGTAAAGCGCGCGTAGGCGGCGCATTAAGTCAGAGGTGAAAGCCCGGGGCTCAACCCCGGAAATGCCTTTGAGACTGGTGTGCTTGAGTGCGGGAGAGGTGAGTGGAATTCCCAGTGTAGAGGTGAAATTCGTAGATATTGGGAAGAACACCAGTGGCGAAGGCGGCTCACTGGCCCGTTACTGACGCTGAGGTGCGAAAGCGTGGGG |
| P-asv5923 | GCAAGCGTTATCCGGAATTACTGGGTTTAAAGGGTGCGTAGGCGGAAATATAAGTCAGAGGTGAAATCACACAGCTTAACTGGGTAATTGCCTTTGATACTGTAATTCTTGAATCAGATTGAGGTTAGCGGAATGAGTCATGTAGCGGTGAAATGCATAGATATGACTTAGAACACCAATTGCGAAGGCAGCTAGCTAGGTCTGTATTGACGCTGAGGCACGAAAGCGTGGGG |
| P-asv5924 | GCAAGCGTTGTTCGGAATTACTGGGCGTAAAGCGTGCGTAGTCGGTATTGAGAGTCACGAGTGAAATCCCAGGGCTCAACCCTGGAACGGCCTCCAAAACTCCTGGTCTTGAGTTCGAGAGAGGTGAGTGGAATTCCGAGTGTAGAGGTGAAATTCGTAGATATTCGGAGGAACACCAGTGGCGAAGGCGGCTCACTGGCTCGATACTGACGCTGAGGTGCGAAAGTGTGGGG |
| P-asv5925 | GCAAGCGTTGTCCGGAATTACTGGGCGTAAAGGGCGTGTAGGCGGTGTCCCACGTAGCGGTTTTCAGCCGCCGGCTCACCCGGCGGAGGGCGGCTAAACGAGGAGACTTGAGGGCAGGAGAGGTGCACGGAATTCCTGGTGGAGCGGTGAAATGCGTAGAGATCAGGAAGAACACCCGTGGCGAAGGCGGTGCACTGGCCTGGCCCTGACGCTGAGGCGCGACAGCGTGGGG |
| P-asv5926 | GCGAGCGTTATTCGGAATCATTGGGTTTAAAGGGTCTGTAGGCGGGCTATTAAGTCAGGGGTGAAAGGTTTCAGCTTAACTGAGAAATTGCCTTTGATACTGGTAGTCTTGAATATCTGTGAAGTTCTTGGAATGCGTAGTGTAGCGGTGAAATGCTTAGATATTACACAGAACACCGATTGCGAAGGCAGGGGACTAACAGACAATTGACGCTGAGAGACGAAAGCGTGGGG |
| P-asv5927 | GCAAGCGTTGTCCGGAATTATTGGGCGTAAAGCGCTCGCAGGCGGCCTTTTAAGTCTGATGTGAAATCTTACGGCTTAACCGTAAGTGGCCATTGGAAACTGAAGGGCTTGAGGACAGAAGAGGAGAGTGGAATTCCACGTGTAGCGGTGAAATGCGTAGAGATGTGGAGGAACACCAGTGGCGAAGGCGACTCTCTGGTCTGTATCTGACGCTGAGGAGCGAAAGCGTGGGG |
| P-asv5928 | GCAAGCGTTGTCCGGATTTACTGGGTGTAAAGGGAGCGCAGGCGGTTATCTAAGTCAGAGGTGAAATCCTACAGCTTAACTGTAGAACTGCCTTTGATACTGGATAACTTGAGTACTAGAGAGAGAGACGGAATTCCAGGTGTAGTGGTGAAATACGTAGATATCTGGAAGAACACCAGAGGCGAAGGCGGTCTCTTGGCTAGTAACTGACGCTGAGGCTCGAAAGCGTGGGG |
| P-asv5929 | GCAAGCGTTGTCCGGATTTATTGGGCGTAAAGCGAGCGCAGGCGGTTTCTTAAGTCTGATGTGAAAGCCTTCGGCTCAACCGAAGAAGTGCATCGGAAACTGGGAAACTTGAGTGCAGAAGAGGACAGTGGAACTCCATGTGTAGCGGTGAAATGCGTAGATATATGGAAGAACACCAGTGGCGAAGGCGGCTGTCTGGCCTGTAACTGACGCTGAGGCTCGAAAGCATGGGT |
| P-asv5930 | GCAAGCGTTGTTCGGAGTTACTGGGCGTAAAGAGTCTGTAGGTGGTCTGTCGAGTCTTTGGTGAAATCCCCGAGCTTAACTTGGGAATTGCATTGGTGACTGCACGGCTAGAGTGTGTCAGAGGGGGGTAGAATTCCACGTGTAGCAGTGAAATGCGTAGAGATGTGGAGGAATACCGATGGCGAAGGCAGCCCCCTGGGATAACACTGACGCTCATGCACGAAAGCGTGGGG |
| P-asv5931 | GCTAGCGTTGCTCGGAATTACTGGGCGTAAAGGGAGCGTAGGCGGACAGTTAAGTTAGAGGTGAAAGCCCAGGGCTCAACCTTGGAATTGCCTTTGATACTGGCTGTCTTGAGTATGGGAGAGGTGTGTGGAACTCCGAGTGTAGAGGTGGAATTCGTAGATATTCGGAAGAACACCAGTGGCGAAGGCGGCTCACTGGACTGGTATTGACGCTGAAGTGCGAAAGCGTGGGG |
| P-asv5932 | GCGAGCGTTGTCCGGAATTACTGGGCGTAAAGAGCTCGTAGGTGGTTTGTCGCGTTGTTCGTGAAATCTCACGGCTTAACTGTGAGCGTGCGGGCGATACGGGCAGACTTGAGTACTGCAGGGGAGACTGGAATTCCTGGTGTAGCGGTGGAATGCGCAGATATCAGGAGGAACACCGGTGGCGAAGGCGGGTCTCTGGGCAGTAACTGACGCTGAGGAGCGAAAGCGTGGGG |

| | |
|---|---|
| P-asv5933 | GCTAGCGTTGTTCGGAATTACTGGGCGTAAAGCGCACGTAGGCGGATATTTAAGTCAGGGGTGAAATCCCGGGGCTCAACCCCGGAACTGCCTTTGATACTGGGTATCTTGAGTGTGGTAGAGGTGAGTGGAATTGCGGGTGTAGAGGTGAAATTCGTAGATATTCGCAGGAACACCAGTGGCGAAGGCGGCTCACTGGACCACAACTGACGCTGAGGTGCGAAAGCGTGGGG |
| P-asv5934 | GCGAGCGTTAATCGGAATTACTGGGCGTAAAGCGTGCGCAGGCGGTTATGTAAGACAGATGTGAAATCCCCGGGCTCAACCTGGGAACTGCATTAGTGACTGCATAGCTGGAGTACGGCAGAGGGGGATGGAATTCCGCGTGTAGCAGTGAAATGCGTAGATATGCGGAAGAACACCGATGGCGAAGGCGACTTTCTGGTCTGTAACTGACACTGAGGCGCGAAAGCGTGGGG |
| P-asv5935 | TCGAGTGATGGCCAATCTTATTGGGCCTAAAGCGTTCGTAGCCGGCCACGCAAGTCCATCGGGAAATCCGCCAGCCCAACTGGCGGGCGTCCGGTGGAACCTGTGTGGCTTGGGACCGGAAGACCCGACAGGTACGTCTTGGGTAGGAGTGAAATCCCGTAATCCTGGACGGACCACCGATAGCGAAAGCATGTCGGGAAGACGGATCCGACGGTGAGGGACGAAAGCCAGGGT |
| P-asv5936 | CCAAGCGTTATTCGGAATCACTGGGCTTAAAGCGTGCGCAGGCGCACCCGTAGGCATCTCGTGAAATCCCACGGCTCAACCGTGGAACTGCGGGGTGAACCACGGGTCTCGAGGCAGGTAGGGGTCAGCGGAACTATGGGTGGAGCGGTGAAATGCGTAGATATCCATGAGAACGCCGATGGCGAAGGCAGCTGACTGGGCCTGTCCTGACGCTCAGGCACGAAAGCGTGGGT |
| P-asv5937 | GCAAGCGTTGTTCGGAATTACTGGGCGTAAAGGGCGCGCAGGCGGTCGGCCACGTTAGATGTGAAAGCCCCGGGCTCAACCTGGGAACTGCATTTAATACGGGCTGGCTTGAGAGCGGGAGAGGAGTGTGGAATTCCCAGTGTAGAGGTGAAATTCGTAGATATTGGGAAGAACACCAGTGGCGAAGGCGGCACTCTGGCCCGTTTCTGACGCTGAGGCGCGAAAGCGTGGGG |
| P-asv5938 | GCGAGCGTTGTCCGGAATTATTGGGCGTAAAGAGCTTGTAGGCGGTTTGTCGCGTCTGCTGTGAAAACTCGAGGCTTAACCTCGAGCCTGCAGTGGGTACGGGCAGACTAGAGTATGGTAGGGGAGACTGGAATTCCTGGTGTAGCGGTGGAATGCGCAGATATCAGGAGGAACACCGATGGCGAAGGCAGGTCTCTGGGCCATAACTGACGCTGAGAAGCGAAAGCATGGGT |
| P-asv5939 | CCAAGCGTTATTCGGAATCACTGGGCGTAAAGGGTGCGTAGGCGGCGTGGAAAGTCAGATGTGAAAGCCCCGGGCTCAACCCGGGAATTGCATCCGATACTCCCGCGCTAGAGGATTGGAGAGGAGTCTGGAATTCACGGTGTAGCAGTGAAATGCGTGGATATCGTGAGGAACACTTGTGGCGAAAGCGAGACTCTGGACAATTCCTGACGCTGATGCACGAAGGCCGGGGG |
| P-asv5940 | GCGAGCGTTAATCGGAATTACTGGGCGTAAAGCGCGCGTAGGCGGCTATTTAAGTCTGATGTGAAATCCCCGGGCTCAACCTGGGAATGGCATCGGATACTGGGTAGCTAGAGTGCAGTAGAGGAAGGTGGAATTCCAGGTGTAGCGGTGAAATGCGTAGAGATCTGGAGGAACACCAGTGGCGAAGGCGGCCTTCTGGACTGACACTGACGCTGAGGTGCGAAAGCGTGGGG |
| P-asv5941 | GCAAGCGTTATCCGGAATCACTGGGTTTAAAGGGTGCGTAGGTGGCGATGTAAGTCAGAGGTGAAAGCCCGGCGCTCAACGTCGGAATTGCCTTTGATACTGCGTTGCTTGAATGAGGTTGAGGTAGGCGGAATGTGGCATGTAGCGGTGAAATGCATAGATATGCCATAGAACACCAATTGCGAAGGCAGCTTACTGGACCTTTATTGACGCTGAGGCACGAAAGCGTGGGT |
| P-asv5942 | GCAAACGTTATTCGGATTTACTGGGTGTAAAGGGTGCGTAGACGGCATAGTAAGTCAACTGTTAAATTTTCCGGCCTAACCGGAAGCCAGCGGTAGAAACTACTAAGCTAGAGGATGGAAGAGAGAAGTGGAATTCTCGGAGTAGCGGTAAAATGCGTAGATCTCGAGAGGAACACCGATGGCGAAAGCAGCTTCTTGGTCCATATCTGACGTTGAGGCACGAAAGCGTGGGG |
| P-asv5943 | GCGAACGTTAATCGGAATTACTGGGCGTAAAGCGCGCGTAGGTGGTCTTTTAAGTGGGATGTGAAAGCCCAGGGCTCAACCCTGGAATTGCATTCCAAACTGAAAGACTCGAGTACGATAGAGGGAGGTAGAATTCAGAGTGTAGCGGTGAAATGCGTAGATATTCTGAGGAATACCAGTGGCGAAGGCGGCCTCCTGGATTGATACTGACACTGAGGCGCGAAAGCGTGGGG |
| P-asv5944 | GCTAGCGTTGTTCGGATTTACTGGGCGTAAAGCGCACGTAGGCGGATTGTTAAGTTAGGGGTGAAATCCCAGGGCTCAACCCTGGAACTGCCTTTAATACTGGCAATCTCGAGTCCGGAAGAGGTGAGTGGAATTCCGAGTGTAGAGATGAAATTCGTAGATATTCGCAAGAACACCAGTGGCGAAGGCGGCTCACTGGCCCATAACTGACGCTGAGGCACGAAAGCGTGGGG |
| P-asv5945 | GCTAGCGTTAATCGGATTTACTGGGCGTAAAGCGTGTGTAGGTGGTTGTTCAAGTCAGTTGTGAAATCCCTGGGCTTAACCTGGGAACTGCTTCTGAGACTGAACGACTAGAGTACGGTAGAGGGTGGTGGAATTTCCGGTGTAGCGGTGAAATGCGTAGATATCGGAAGGAACATCAATGGCGAAGGCAGCCACCTGGGCCTGTACTGACACTGAGACACGAAAGCGTGGGG |
| P-asv5946 | GCAAGCGTTAATCGGAATTACTGGGCGTAAAGAATGCGTAGGCTGTTTTGCAAGTTGGTGATGAAATGCTGGGGCTCAACTCCAGTGCTGTTACCAAAACTGCTTAACTTGAGTGCGGTAGGGGGCAATGGAATTCTTGGTGTAGGGGTTAAATCCTTTGATATCAAGAAGAACGCCGGAGGCGAAAGCGATTGCCTGGGCCGTTACTGACGCTAATGCATGACAGTGTGGGG |
| P-asv5947 | CCGAGCGTTGTCCGGAATCATTGGGCGTAAAGGGTACGTAGGCGGGTCATTAAGTTAGAAGTCAAAGGCTATAGCTCAACTATAGTAAGCTTCTAAAACTGGAGACCTTGAGTAATGGAAGGGAAAGTGGAATTCCTAGTGTAGCGGTGGAATGCGCAGATATTAGGAGGAATACCGGTGGCGAAGGCGACTTTCTGGCCATTAACTGACGCTGAGGTACGAAAGCGTGGGT |
| P-asv5948 | GCAAGCGTTAATCGGAATTACTGGGCGTAAAGTGCACGTAGGCGGCCTTGTAAGTTGGGGGTGAAATCCCGGAGCTCAACTCCGGAACTGCCTTCAAAACTACAAAGCTAGAGGATGTGAGAGGATAGTGGAATTCCGAGTGTAGAGGTGAAATTCGTAGATATTCGGAAGAACACCAGTGGCGAAGGCGACTGTCTGGCACATTTCTGACGCTGAGGTGCGAAAGCGTGGGG |
| P-asv5949 | GCGAGCGTTACTCGGATTTACTGGGCGTAAAGGGCGCGTAGGCGGTTTGTTAAGTCTGATGTGAAAGCCCACGACTCAATCGTGGGAGTGCATTGGATACTGGCAGACTAGAGTTCAAGAGGGGTGAGCGGAATTCCAGGTGTAGAGGTGAAATTCGTAGATATCTGGAGGAACAACGGTGGCGAAGGCGGCTCACTGGATTGATACTGACGCTGAGGCGCGAAAGCGTGGGG |
| P-asv5950 | GCTAGCGTTGTTCGGAATTACTGGGCGTAAAGCGCGCGCAGGCGGCCATTTAAGTCAGATGTGAAAGCCCGGGGCTCAACCTTGGAACTGCATTTGAAACTGAGTGGCTAGAGTATGGTAGGGGAGAGGGGAATTCCTAGTGTAGCGGTGAAATGCGTAGAGATTAGGAGGAACACCAGTGGCGAAGGCGCCTCTCTGGGCCATTACTGACGCTGAGGCGCGAAAGCGTGGGG |
| P-asv5951 | GCAAGCGTTGTTCGGATTCATTGGGTGTAAAGGGTGCGTAGGCGGCCTGGTAAGTCAGATGTGAAATCCTACAGCTCAACTGTAGAACTGCATTTGAAACTACCAAGCTTGAGTGCGGAAGAGGTGAAGGGAATTCCCGGTGTAAGGGTGAAATCTGCAGAGATCGGGAAGAACACCAGCAGCGAAGGCGCTTCACTGGTCCGTAACTGACGCTGAGTGCACGAAGGCGTGGGG |
| P-asv5952 | CCAAGCGTTGTTCGGATTTACTGGGCGTAAAGGGCGCGTAGGCGGCGCGGTAAGTCACTTGTGAAATCTCTGAGCTTAACTCAGAACGGCCAAGTGATACTGCAGTGCTAGAGTGTGGAAGGGGCAATCGGAATTCTTGGTGTAGCGGTGAAATGCGTAGATATCAAGAGGAACACCTGAGGTGAAGACGGGTTGCTGGGCCAACACTGACGCTGATGCGCGAAAGCCAGGGG |
| P-asv5953 | GCAAGCGTTGTTCGGAATCATTGGGCTTAAAGGGCGTGCAGGCGGCCGCGCAAGTGTCCTGTGAAAGCCCCCGGCTCAACCGGGGAAGTGCGGGGCAAACTGCGCGGCTTGAGGTCGGTAGAGGCGCCTGGAACTCTAGGTGGAGCGGTGAAATGCGTAGATATCTAGAGGAACGCCAGGAGCGAAAGCGGGGCGCTGGGCCGATTCTGACGCTGAGACGCGAAAGCGTGGGG |
| P-asv5954 | CCGAGCGTTATCCGGAATTATTGGGCGTAAAGGGTCCGTAGGTGGCCATATTAGTCTTTTGTCGAAGACCCGTGCTCAACACGGGGGAGGCAAAGGAAACGGTATGGCTTGAGTCTGTGAGAGGTGTGCGGAACTCATGGTGTAGGGGTGAAATCCGTTGATATCATGGGGAACACCAAAAGCGAAGGCAGCACACTGGCACACGACTGACACTGAGGGACGAAACCCTGGGT |
| P-asv5955 | GCGAGCGTTGTCCGGATTTATTGGGTTTAAAGGGTGCGTAGGCGGCTTTTTAAGTCTGGGGTGAAAGCCCGCTGCTCAACAGCGGAACTGCCCTGGATACTGGAGAGCTTGAGTACAGACGAGGTTGGCGGAATGGACTGAGTAGTGGTGAAGTGCATAGATACAGTCCAGAACCCCGATTGCGAAGGCAGCTGACTAGGCTGTTACTGACGCTGAGGCACGAAAGCGTGGGG |
| P-asv5956 | GCAAGCGTTATCCGGATTTATTGGGCGTAAAGCGCTCGTAGGCGGTCTGTTAGGTCGGGAGTTAAATCCGGGGGCTCAACCCCCGTCCGCTCCCGATACCGGCAGACTTGAGTTTGGTAGGGGAAGGTGGAATTCCTAGTGTAGCGGTGGAATGCGCAGATATTAGGAAGAACACCAGTGGCGAAGGCGGCCTTCTGGACCATAACTGACGCTGAGGAGCGAAAGCTAGGGG |
| P-asv5957 | GCAAGCGTTATCCGGAATTATTGGGCGTAAAGCGCGCGCAGGCGGCCGGTTAAGTCAGGTGTGAAAGGCTGCGGCTCAACCGCAGAGCGGCATCTGAAACTGGTCGGCTTGAGTGCAGGAGAGGGGAGCGGAATTCCCGGTGTAGCGGTGGAATGCGTAGAGATCGGGAGGAACACCAGTGGCGAAGGCGGCTCTCTGGCCTGTAACTGACGCTGAGGCGCGAAAGCGTGGGG |
| P-asv5958 | GCAAGCGTTACTCAGAATTACTGGGCGTAAAGCGTGCGTAGGTGGTGGTTTAAGTCTGCTGTGAAAGCCCTGGGCTCAACCTGGGAATTGCAGTGGATACTGGATCACTAGAGTGTGGTAGAGGGATGCGGAATTTCTGGTGTAGCAGTGAAATGCGTAGAGATCAGAAGGAACATCCGTGGCGAAGGCGGCATCCTGGGCCAACACTGACACTGAGGCACGAAAGCGTGGGG |
| P-asv5959 | CCTAGCGTTGTTCGGAATCATTGGGCGTAAAGCGGGTGCAGGTGGCTATGTAAGTCAGATGTGGAAGCCCAGGGCTCAACCCTGGAAGTGCATTTGATACTGCGTAGCTTGAGTGCTGGAGAGGTTACTAGAATTCTTGGTGTAGTGGTGAAATACGTAGATATCAAGAGGAATACCGGCGGCGAAGGCGGGTAACTGGCCAGACACTGACACTCAGACCCGAAAGCGTGGGG |
| P-asv5960 | GCTAGCGTTGCTCGGAATTACTGGGCGTAAAGGGCGCGTAGGCGGACCTGCAAGTCAGGGGTGAAATCCCGGGGCTCAGCCTCGGAACTGCCCTTGAGACTGTTGGTCTAGAGTCTGTGAGAGGATGACGGAATACCCAGTGTAGAGGTGAAATTCGTAGATATTGGGTAGAACACCGGTGGCGAAGGCGGTCATCTGGCGCAGCACTGACGCTGAGGCGCGAAAGCGTGGGG |
| P-asv5961 | TCAAGTGATGTCCCATATTATTGGGCCTAAAGCGTCCGTAGCTGGCCGGGCAAGTCCATCGGGAAATCCACCTGCCCAACAGGTGGGCGACCGGTGGAAACTGCCTGGCTTGGGATCGGAAGGCTCAGAGAGTACGACCGGGGTAGGAGTGAAATCCCGTAATCCAGGACGGACTACCGATGGCGAAAGCACTCTGAGAAGACGACTCCGACAGTGAGGGACGAAAGCTCGGGT |
| P-asv5962 | GCAAGCGTTGTTCGGAATTACTGGGCGTAAAGGGTTCGTAGGTGGCCATTTAAGTCAGACGTGAAATCCCTCAGCTCAACTGGGGAATTGCGTCTGAAACTGGATGGCTTGAGTGCAGGAGAGGAATGCGGAATTCCAGGTGTAGCGGTGAAATGCGTAGATATCTGGAAGAACACCGGTGGCGAAGGCGGCATTCTGGACTGTAACTGACACTGAGGAACGAAAGCCAGGGG |
| P-asv5963 | GCGAGCGTTGTCCGGATTCATTGGGCGTAAAGAGCTCGTAGGCGGTTCAGTAAGTCGGGTGTGAAAGCTCCGAGCTCAACTCGGAGAGGCCACCCGATACTGCTGTGACTAGAATCCAGTAGGGGAGTGTGGAACTCCTGGTGTAGCGGTGAAATGCGCAGATATCAGGAGGAACACCGACGGCGAAGGCAGCACTCTGGGCTGGTATTGACGCTGAGGAGCGAAAGTGTGGGT |
| P-asv5964 | GCAAGCGTTATCCAGTTTTATTGGGTTTAAAGGGTGCGTAGGCGGCTTATTAAGTCAGTTGTGAAATCCTAGCGCTCAACGTTAGAACTGCGATTGATACTGATAGGCTTGAGTCAAGAAGAGGTAGGTAGAATTTATGGTGTAGTAGTGAAATGCTTAGATATCATAAGGAATACCAATAGCGTAGGCAGCTTACTGGTCTTGAACTGACGCTGAGGCACGAAAGCGTGGGG |
| P-asv5965 | GCAAGCGTTGTTCGGAATTACTGGGCGTAAAGCGTGCGTAGTCGGTATTGAGAGTCACGGGTGAAATCCCCAAGCTTAACTTGGGAACTGCCTGTGAGACCTCAGTACTAGAGTGTGAGAGGGGATAGTGGAATACCAAGTGTAGCGGTGAAATGCGTAGAGATTTGGTGGAACACCGGTGGCGAAGGCGGCTATCTGGCTCACAACTGACGATCAGGCACGAAAGCGTGGGG |
| P-asv5966 | GCGAGCGTTATCCGGAATTATTGGGCGTAAAGAGTACGTAGGTGGTTTTGTAAGCGTGGGGTGAAAGACCGAAGCTCAACTTCGGTTAGCCTTGCGAACTGCAAGACTTGAGTGCAGGAGAGGAAAGTGGAATTCCTAGTGTAGCGGTGAAATGCGTAGATATTAGGAGGAACATCAGTGGCGAAGGCGACTTTCTGGACTGTAACTGACACTGAGGTACGAAAGCGTGGGT |
| P-asv5967 | ACGAGTGGTGACCACGTTTATTGGGCCTAAAGCGTCCGTAGCTGGCTTGTTAAATCCTTTGTGAAATCGTCCGGCTTAACCGTACGACGCGCAAGGGAGACTGGCAGGCTCGAGACCGGGAGGAGTCAGAAGTATTCCTGAGGGAGCGGTAAAATGTTATAATCTCAGGAGGACTACCTGTGGCGAAGGCGTCTGACTAGAACGGATCTGACAGTGAGGGACGAAAGCCAGGGG |
| P-asv5968 | GCAAGCGTTAATCGGAATCACTGGGCGTAAAGCGTGCGTAGGCGGCCTGTCAAGTCAGGTGTGAAAGCCCTCGGCTCAACCGGGGAATTGCACTTGAAACTGTCAGGCTGGAGTCTCGGAGAGGGTGGCGGAATTCCCGGTGTAGGAGTGAAATCCGTAGATATCGGGAGGAACACCAGTGGCGAAGGCGGCCACCTGGACGATGACTGACGCTGAGGTACGAAAGCGTGGGT |
| P-asv5969 | GCAAGCGTTGTCCGGATTTATTGGGTTTAAAGGGTGCGTAGGCGGTCATATAAGTCAGTGGTGAAATACTTCAGCTCAACTGGAGAGGTGCCATTGATACTGTATGACTTGAGTACAGACGAGGTAGGCGGAATTGACGGTGTAGCGGTGAAATGCTTAGATATCGTCAAGAACACCGATAGCGAAGGCAGCTTACTAGACTGTAACTGACGCTGATGCACGAAAGTGTGGGG |
| P-asv5970 | GCAAGCGTTGTCCGGAATCATTGGGCGTAAAGGGCGTGTAGGTGGTTTCGTAAGTCGAATATGAAATCTCACGGCTCAACCGTGAAATTGCGTTCGAAACTACGAGACTTGAGTTAGGGAGAGGTGTACGGAATTCCTGGTGTAGCGGTGGAATGCGTAGATATCAGGAAGAACACCAATGGCGAAGGCAGTACACTGGCCTGTAACTGACACTGAGGCGCGAAAGCGTGGGG |
| P-asv5971 | GCAAACGTTATTCAGAATCACTGGGCTTAAAGGGTGCGTAGGCGGCTCATTAAGTAGGGTGTGAAAGCCCCCGGCTCAACCGGGGAATTGCGCTCTAAACTGATGGGCTGGAGTGAGATAGGGGTGTGTGGAACTTCCGGTGGAGCGGTGAAATGTGTTGATATCGGAAGGAACGCCGGTGGCGAAAGCGACACACTGGGTCTCAACTGACGCTGAGGCACGAAAGCCAGGGG |

| | |
|---|---|
| P-asv5972 | GCTAGCGTTGCTCGGAATTACTGGGCGTAAAGGGCGCGTAGGTGGTTTAGTAAGTTATCTGTGAAATTCCTGGGCTTAACCTAGGGTGGTCAGATAAGACTGTTGGACTGAGAGTATAGGAGAGGGTAGTGGAAT<br>TTCCGGTGTAGCGGTGAAATGCGTAGAGATCGGAAGGAACACCAGTGGCGAAGGCGGCTACCTGGCCTAATACTGACACTGAGGCACGAAAGCGTGGGG |
| P-asv5973 | GCAAGCGTTGTCCGGAATTATTGGGCGTAAAGAGCATGTAGGCGGTATATTAAGTTGGATGTAAAAGGCAATGGCTCAACCATTGTAAGCGTCCAAAACTGGTATACTTGAGTGCAGGAGAGGTGAGTGGAATTC<br>CCGGTGTAGCGGTGGAATGCGTAGAGATCGGGAGGAACACCAGTGGCGAAGGCGACTCACTGGCCTGTAACTGACGCTGAGATGCGAAAGCGTGGGG |
| P-asv5974 | GCGAGCGTTGTCCGGAATTATTGGGCGTAAAGGGCTCCTAGGTTGTTCAGTAAGTCTACTGTGAAAATCCAAGGCTTAACCTTGGGACGCCAGTGGATACTGCTGTGACTCGAGTTCGGTAGAGGAGTGTGGAGT<br>TCCTGGTGTAGCGGTGAAATGCGCAGATATCAGGAGGAACACCAACGGCGAAGGCAGCACTCTGGGCCGATACTGACACTGAAGAGCGAAAGCATGGGG |
| P-asv5975 | GCGAGCGTTAATCGGAATCACTGGGCGTAAAGCGCACGTAGGCGGCGACGTAAGTCAGACGTGAAAGCCCTCGGCTCAACCGAGGAATTGCGTTTGATACTGCGTTGCTTGAGTCCCGGAGAGGGTGGCGGAA<br>TTCCGGGTGTAGGAGTGAAATCCGTAGATATCCGGAAGAACACCGGTGGCGAAGGCGGCCACCTGGACAGGTACTGACGCTGAGGTGCGAAAGCGTGGGG |
| P-asv5976 | GCAAGCGTTATTCAGATTCACTGGGCGTAAAGGGTGCGCAGGCGGTCGTGCGTGTCAGGTGTGAAATCCCGGAGCTTAACTCCGGAACTGCGCCTGAAACTACACGACTAGAGCATTGGAGAGGGAAGTGGAAT<br>TCATGGTGTAGCAGTGAAATGCGTAGATATCATGAGGAACACCAGAGGCGAAGGCGACTTCCTGGACAATTGCTGACGCTCAGGCACGAAAGCATGGGG |
| P-asv5977 | CCAAGTGGTGGTCGATTTTATTGAGTCTAAAGCGTTCGTAGCCGGTTTGATAAATCCTTGGGTAAATCGGAAAGCTTAACTTCCCGAATTCCGAGGAGACTGTCAAACTTGGGACCGGGAGAGGCTAGAGGTACTT<br>CTGGGGTAGGGGTAAAATCCTGTAATCCTAGAAGGACCACCGGTGGCGAAGGCGTCTAGCTAGAACGGATCCGACGGTGAGGGACGAAGCCCTGGGT |
| P-asv5978 | ACAAACGTTATTCGGAATTACTGGGCTTAAAGGGTGCGTAGGCTGTGCGGAAAGTTAGGTGTGAAATCCCCCGGCTCAACCGGGGAATTGCGCTTAAAACTACCGTGCTAGAGGAATACAGAGGTGAGCGGAACT<br>TCGGGTGGAGCGGTGAAATGCGTTGATATCCGAAGGAACACCGGTGGCGAAAGCGGCTCACTGGGTATTTTCTGACGCTGATGCACGAAAGCTAGGGG |
| P-asv5979 | CCAAGCATTATCCGGATTTACTGGGCGTAAAGGGTGCGTAGGCGGTGTGATTAGTCAGGTGTTAAATCCTGGGGCTTAACCCCAGAATCGCATTTGAAACGGTCATACTAGAAGAGGTGAGAGGTAAGCAGAACT<br>CACGGTGTAGGGGTGAAATCCGTTGATATCGTGGGGAATACCAAATGCGAAGGCAGCTTACTGGCACATTCTTGACGCTGAGGCACGAAAGCGTGGGT |
| P-asv5980 | GCTAGCGTTGTTCGGAATTACTGGGCGTAAAGGGCGCGTAGGCGGCCATTTAAGTTAGGTGTGAAAGTCCTGGGCTCAACCTGGGGACTGCACTTAATACTGGGTGGCTTGAATTCGGGAGAGGATAGCGGAAT<br>TCCCAGTGTAGAGGTGAAATTCGTAGATATTGGGAAGAACACCGGTGGCGAAGGCGGCTATCTGGACCGACATTGACGCTGAGGCGCGAAAGCGTGGGG |
| P-asv5981 | TCAAGCGTTGTTCGGATTCATTGGGCGTAAAGGGTGCGTAGGCGGCGCTGTAAGTCGGATGTGAAAGCCCGGGGCTCAACTCCGGAATTGCATTCGATACTGCAGCGCTAGAGGACTGGAGAGGAGACTGGAA<br>TTCACAGTGTAGCAGTGAAATGCGTAGATATTGTGAGGAAGGCCAGTGGCGAAGGCGGGTCTCTGGACAGTTCCTGACGCTGAGGCACGAAGGCTAGGGT |
| P-asv5982 | GCGAGCGTTAATCGGAATTATTGGGCGTAAAGCGAGTGCAGACGGTTGTTTAAGCCAGATGTGAAATCCCTGAGCTTAACTTAGGAGTTGCATTCGATACTGGGAAGCTAGAGTATGGGAGAGGATGGTAGAATT<br>CCAGGTGTAGCGGTGAAATGCGTAGAGATCTGGAGGAATACCGATGGCGAAGGCAGCCATCTGGCCTAATACTGACGCTGAGGTACGAAAGCATGGGG |
| P-asv5983 | GCAAGCGTTGTTCGGAATCACTGGGCGTAAAGCGCGTGTAGGCGGCCCGATTAGTCGAGTGTGAAAGTCCGAGGCTCAACCTCGGAAGCGCACTCGAAACTGTCAGGCTTGAGTACGGGAGAGGGAAGTGGAA<br>TTCCTGGTGTAGAGGTGAAATTCGTAGATATCAGGAGGAACACCGGTGGCGAAGGCGGCTTCCTGGACCGATACTGACGCTGAGACGCGAAAGCGTGGGG |
| P-asv5984 | CCGAGCGTTATCCGGAATCATTGGGTTTAAAGGGTCCGTAGGCGGGCGCATAAGTCAGGGGTGAAAGTTTGCCGCTCAACGGTAAAATTGCCTTTGATACTGTGTGCCTTGAATAGCGGTGAAGTGGTTAGAATG<br>TGCAGTGTAGCGGTGAAATGCTTAGATATTGCACAGAATACCGATTGCGAAGGCAGGCCACTAACCGCGTATTGACGCTGAGGGACGAAAGCGTGGGG |
| P-asv5985 | GCAAGCGTTATCCGGATTTATTGGGCGTAAAGCGAGCGCAGGCGGTTGTTTAGGTCTGATGTGAAAGCCTTCGGCTTAACCGAAGAAGTGCATCGGAAACCGGGCGACTTGAGTGCAGAAGAGGACAGTGGAAC<br>TCCATGTGTAGCGGTGGAATGCGTAGATATATGGAAGAACACCAGTGGCGAAGGCGGCTGTCTGGTCTGCAACTGACGCTGAGGCTCGAAAGCATGGGT |
| P-asv5986 | GCAAGCGTTATTCGGATTCACTGGGCGTAAAGGGTGCGCAGGCGGTTGCGTGTGTTGGGTGTGAAAGCCCGGAGCTCAACTCCGGAACTGCGCCCAAAACTACGCGACTAGAGTACCGGAGAGGATAGCGGAA<br>TTCACGGTGTAGCAGTGAAATGCGTAGATATCGTGAAGAACACCAGCGGCGAAGGCGGCTATCTGGAAGGTAACTGACGCTCAGGCACGAAAGCGTGGGG |
| P-asv5987 | GCAAACGTTATTCGGAATCACTGGGCGTAAAGGGTGCGTAGACGGCAATATAAGTCAATTATTAAATCTCCTAGCTTAACTAGGAATCTGTGATTGAAACTATATGGCTTGAGGATGGAAGAGAGAAGTGGAATTCT<br>CGGAGTAGCGGTAAAATGCGTAGATCTCGAGAGGAACACCGATGGCGAAGGCAGCTTCTTGGTCCATTTCTGACGTTGAGGCACGAAAGCGTGGGG |
| P-asv5988 | CCGAGCGTTAATCGGAATTACTGGGCGTAAAGCGTGCGCAGGCGGTTTGTTAAGCGAGATGTGAAAGCCCTGGGCTCAACCTAGGAATAGCATTTCGAACTGGCGAACTAGAGTCTTGTAGAGGGGGGTAGAAT<br>TCCAGGTGTAGCGGTGAAATGCGTAGAGATCTGGAGGAATACCGGTGGCGAAGGCGGCCCCCTGGACAGAGACTGACGCTCAGGTGCGAAAGCGTGGGG |
| P-asv5989 | GCAAGCGTTGTCCGGAATTATTGGGCGTAAAGGGAGCGCAGGCGGGAAACTAAGCGGATATTAAAAGTGCGGGGCTCAACCCCGTGATGGGGTCCGAACTGGTTTTCTTGAGTGCAGGAGAGGAAAGCGGAAT<br>TCCCAGTGTAGCGGTGAAATGCGTAGATATTGGGAAGAACACCAGTGGCGAAGGCGGCTTTCTGGACTGTAACTGACGCTGAGGCTCGAAAGCTAGGGT |
| P-asv5990 | GCAAGCGTTGTCCGGAATTATTGGGCGTAAAGCGCGCGCAGGCGGTTCCTTAAGTCTGATGTGAAAGCCCCCGGCTCAACCGGGGAGGGTCATTGGAAACTGGGGAACTTGAGTGCAGAAGAGGAGAGCGGAA<br>TTCCACGTGTAGCGGTGAAATGCGTAGAGATGTGGAGGAACACCAGTGGCGAAGGCGGCTCTCTGGACTGTAACTGACGCTGAGGCGCGAAAGCGTGGGG |
| P-asv5991 | GCGAGCGTTAATCGGATTTACTGGGCGTAAAGCGCGCGTAGGTGGCCAATTAAGTCAAATGTGAAATCCCCGAGCTTAACTTGGGAATTGCATTCGATACTGGTTGGCTAGAGTATGGGAGAGGATGGTAGAATT<br>CCAGGTGTAGCGGTGAAATGCGTAGAGATCTGGAGGAATACCGATGGCGAAGGCAGCCATCTGGCCTAATACTGACACTGAGGTGCGAAAGCATGGGG |
| P-asv5992 | GCAAGCGTTAATCGGAATCACTGGGCGTAAAGGGTGCGTAGGCGGTACGTTAGGTCTGTCGTGAAATCCCCGGGCTCAACCTGGGAATGGCGGTAGAAACCGGCGCACTAGAGTATGGGAGAGGGTGGTGGAA<br>TTCCAGGTGTCGCGGTGAAATTCGTAGATATTCGGAAGAACACCAGTGGCGAAGGCGGCTCGCTGGACAAGTATTGACGCTGAGGTGCGAAAGCGTGGGG |
| P-asv5993 | GCAAGCGTTATCCGGAATTATTGGGCGTAAAGCGTCCGCAGGCGGCCCTACAAGTCTGCTGTTAAAAAGTGGAGCTTAACTCCATCATGGCAGTGGAAACTGTTGGGCTTGAGTGTGGTAGGGGCAGAGGGAAT<br>TCCCGGTGTAGCGGTGAAATGCTTAGATATCGGGAAGAACACCAGTGGCGAAGGCGCTCTGCTGGGCCATCACTGACGCTCATGGACGAAAGCCAGGGG |
| P-asv5994 | GCAAGCGTTGTCCGGAATTATTGGGCGTAAAGAGCTCGTAGGCGGTTTGTCGCGTCTGCTGTGAAATCTGGGGGCTCAACCCCCAGCCTGCAGTGGGTACGGGCAGACTAGAGTGCGGTAGGGGAGATTGGAA<br>TTCCTGGTGTAGCGGTGGAATGCGCAGATATCAGGAGGAACACCGATGGCGAAGGCAGATCTCTGGGCCGTAACTGACGCTGAGGAGCGAAAGCATGGGG |
| P-asv5995 | CCGAGCGTTGTCCGGAATTATTGGGCGTAAAGGGCTCGTAGGCGGTTTGTCGCGTCGGGAGTGAAAACTCAGGGCTCAACCCTGAGCGTGCTTCCGATACGGGCAGACTAGAGGTATGCAGGGGAGAACGGAA<br>TTCCTGGTGTAGCGGTGGAATGCGCAGATATCAGGAGGAACACCGGTGGCGAAGGCGGTTCTCTGGGCATTACCTGACGTTGAGGAGCGAAAGCATGGGG |
| P-asv5996 | GCAAGCGTTATCCGGATTCACTGGGTCTAAAGGGTGCGTAGGCGGACTTGTAAGTCCGGGGTGAAAGCCCCCAGCTCAACTGGGGAAGCGCCTTGGATACTTCAAGTCTGGAGGGAGGTAGGGGCGTGCGGAA<br>CTTCCGGTGGAGCGGTGAAATGCGTAGATATCGGAAGGAGCACCGGTGGCGAAAGCGGCGCGCTGGACCTCTTCTGACGCTGAGGCGCGAAAGCCAGGGG |
| P-asv5997 | GCGAGCGTTAATCGGAATTACTGGGCGTAAAGGGCGAGTAGGTGGCCATCTAAGTCAGACGTGAAATCCCTCGGCTTAACCGGGGAACTGCGTCTGATACTGGATGGCTTGAGTTCGGGAGAGGGATGCGGAAT<br>TCCAGGTGTAGCGGTGAAATGCGTAGATATCTGGAGGAACACCGGTGGCGAAGGCGGCATCCTGGACCGACACTGACACTGAGGAGCGAAAGCCAGGGG |
| P-asv5998 | GCAAACGTTGTTCGGAATTACTGGGCATAAAGCGCACGTAGGCGGTCCACTAAGTCAGGTGTGAAATTCCCCGGCTCAACCGGGGAATTGCGCCTGATACTGGTGGGCTCGAGGTCGATAGGGGTGAGCGGAA<br>CTCCAGGTGGAGCGGTGAAATGCGTAGATATCTGGAGGAACACCGGTGGCGAAGGCGGCTCACTGGATCGAATCTGACGCTGAGGCGCGAAAGCTAGGGT |
| P-asv5999 | GCAAGCGTTGTCCGGATTTATTGGGTTTAAAGGGTGCGTAGGCGGTTTTTTAAGTCAGTTGTGAAATACGGCAGCTTAACTGTCGAGGTGCAATTGATACTGGGAAACTTGAGTACAGACGAGGTAGGCGGAATT<br>GACGGTGTAGCGGTGAAATGCATAGATATCGTCAAGAACACCGATAGCGAAGGCAGCTTACTAGACTGTAACTGACGCTGATGCACGAAAGTGTGGGG |
| P-asv6000 | GCGAGCGTTGTCCGGAATCACTGGGCGTAAGGGCGCGTAGGCGGCTTCTCACGCGTGCGGTGAAAGCCCGGGGCTTAACTCCGGGTCTGCCGTGCGAACGGGGAGGCTTGAGTCTGGTAGAGGTCGATGGAA<br>TTCCGGGTGTAGCGGTGGAATGCGTAGAGATCCGGAAGAACACCGGTGGCGAAGGCGGTCGACTGGGCCAGAACTGACGCTGAGGCGCGACAGCGTGGGG |
| P-asv6001 | GCAAGCGTTGTTCGGATTTATTGGGCGTAAAGGGTGCGTAGGCGGTTTGGCAAGTCTTGTGTGAAATCTTCGGGCTCAACCCGAAGTCTGCATAAGAAACTGCCGGGCTTGAGTATGGGAGAGGTGAGTGGAAT<br>TTCCGGTGTAGCGGTGAAATGCGTAGATATCGGGAGGAACACCTGTGGCGAAGGCGAGTCTCTGGACATTTCCTGACGCTGAGGCACGAAAGCCAGGGG |
| P-asv6002 | GCGGGCGTTATCCGGATTTACTGGGTGTAAAGCGCGTGCAGGCGGTTGGTTAAGTTGGGTGTGAAAGCTCCTGGCTCAACTGGGAGAGGTCGCTCAAGACTGGCCAACTGGAGAGCAGTAGGGGAAGGTGGAA<br>TTCCGGGTGTAGTGGTGAAATGCGTAGATATCCGGAGGAACACCAGTGGCGAAAGCGGCCTTCTGGACTGCATCTGACGCTCAGACGCGAAAGCTAGGGG |
| P-asv6003 | GCAAGCGTTGTCCGGAATTATTGGGCGTAAAGCGCGCGCAGGCGGATAGGTCAGTCTGTCTTAAAAGTTCGGGGCTTAATCCCGTGATGGGATGGAAACTGCCAATCTAGAGTATCGGAGAGGAAAGTGGAATT<br>CCTAGTGTAGCGGTGAAATGCGTAGATATTAGGAAGAACACCAGTGGCGAAGGCGACTTTCTGGACGAAAACTGACGCTGAGGCGCGAAAGCCAGGGG |
| P-asv6004 | TCGAGTGGTGATCACTTTTATTGAGTCTAAAGCGTCCGTAACCGGTCTTATAAATCTTCAGATAAATTCTTCCGCTTAATGGAAGAACTTCTGAAGAGACTGTAAGACTTGGGACCGGGTGAGGTTGAATGTACTTT<br>CAGGGTAGGGGTAATATCCTGTAATCCTGAAAGGACGACCGGTGGCGAAAGCGTTCAACTAGAACGGATCCGACGGTGAGGGACGAAGGCTAGGGG |
| P-asv6005 | GCTAGCGTTGTCCGGAATCACTGGGCGTAAAGGGTTCGCAGGCGGGATAACAAGTCAGATGTGAAAGGCATGGGCTCAACCCATGTAAGCATTTGAAACTGTAATTCTTGAGAAGTGGAGAGGTAAGTGGAATTC<br>CTAGTGTAGCGGTGAAATGCGTAGATATTAGGAGGAATACCTGTGGCGAAGGCGACTTACTGGACACAAATCTGACGCTGAGGAACGAAAGCGTGGGT |
| P-asv6006 | GCAAGCGTTATTCGGAATTATTGGACGTAAAGGGCGCGCAGGCGGTCGTTCAAGTCGAGTGTGAAAGCCCCCGGCTCAACTGGGGAGGGTCATTCGATACTGATCGACTCGAAGGCAGGAGAGGGTAGTGGAA<br>TTCCCGGTGTAGTGGTGAAATGCGTAGATATCGGGAGGAACACCAGTGGCGAAGGCGACTACCTGGCCTGTTCTTGACGCTGAGGCGCGAAAGCTAGGGG |
| P-asv6007 | GCAAGCGTTGTCCGGAATTATTGGGCGTAAAGCGCTCGTAGGTGGATCGAAAAGTCTGCAAGCAAAGCCCCGAGCTCAACTCGGGAATCCGTGTGGAAACTGTCGATCTTGAGGGAGTCAGAGGATGATGGAAT<br>TCCCGGTGTAGCGGTGAAATGCGTAGATATCGGGAGGAACACCAGTGGCGAAGGCGATCATCTGGGGCTAACCTGACACTGAGGAGCGAAAGCTAGGGG |
| P-asv6008 | GCTAGCGTTGCTCGGAATTACTGGGCGTAAAGCGCACGTAGGCGGCTTTCTAAGTCGGGGGTGAAATCCTGGAGCTCAACTCCAGAACTGCCTTCGATACTGGAGGGCTCGAGTTCGGGAGAGGTGAGTGGAA<br>CTGCGAGTGTAGAGGTGAAATTCGTAGATATTCGCAAGAACACCAGTGGCGAAGGCGGCTCACTGGCCCGATACTGGCGCTGAGGTAGGAAAGCGTGGGG |
| P-asv6009 | CCAAGCGTTATCCGGATTTATTGGGTTTAAAGGGTGCGTAGGTGGCTTTTTAAGTCAGGGGTGAAAGACGGTAGCTTAACTATCGCAGTGCCTTTGATACTGAAGAGCTTGAATGAACTAGAGGTAGGCGGAATGT<br>GACAAGTAGCGGTGAAATGCATAGATATGTCACAGAACACCAATTGCGAAGGCAGCTTACTATGGTTTTATTGACACTGAGGCACGAAAGCGTGGGG |
| P-asv6010 | GCGAACGTTGCTCGGAATTACTGGGCGTAAAGCGCGTGTAGGCGGGTGCGCAAGTCCGTTGTGAAATCCCTGGGCTCAACCCAGGAACTGCATCGGAAACTGCGCATCTTGAGTAATGGAGAGGGAAGTGGAAT<br>TCCCGGTGTAGAGGTGAAATTCGTAGATATCGGGAGGAACACCAGTGGCGAAGGCGGCTTCCTGGACATTTACTGACGCTGAGACGCGAAAGCGTGGGG |

| | |
|---|---|
| P-asv6011 | GCGAGCGTTGTTCGGAATTACTGGGCGTAAAGCGCGTGTAGGTGGTTTGTCATGTCAGGTGTGAAAGCCCTCGGCCTAACCGAGGAAGTGCGCCTGAAACTGCCAAACTTGAGTACGAGAGAGGAGGACGGAATTCCCAGTGTAGAGGTGAAATTCGTAGATATTGGGAAGAACACCGGTGGCGAAAGCGGTTCTCTGGATCGATACTGACACTAAGACGCGAAAGCTAGGGG |
| P-asv6012 | CCAAACGTTATTCGGATTTACTGGGCTTAAAGAGTTCGTAGGCGGCTTGGAAGGTGGGGTGTGAAAGCCCTCGGCTCAACCGAGGAATTGCGCTCCAAACTACCGAGCTTGAGGAGGCTAGAGGTGAGCAGAACAGATGGTGGAGCGGTGAAATGCGTTGATATCATCTGGAATACCGGAGGCGAAGGCGGCTCACTGGGACCTTTCTGACGCTGAGGAACGAAAGCTAGGGG |
| P-asv6013 | GCGAGCGTTGTCCGGAATCACTGGGCGTAAAGGGCGCGTAGGTGGCGCGAGTAGTCAACGGTGAAAGCCCGGGGCTCAACCCCGGGTCGGCCGTCGATACTCTCGTGCTTGAGCACTGTAGAGGCAGGTGGAATTCCGGGTGTAGCGGTGGAATGCGTAGAGATCCGGAAGGACACCGGTGGCGAAGGCGGCCAGCTGGGCAGTAGCTGACACTGAGGCGCGACAGCGTGGGG |
| P-asv6014 | GCAAGCGTTGCTCGGAATTATTGGGCGTAAAGGGCGTGTAGGCGGTTTGTTAAGTCATGTGTAAAAGCCCTCGGCTCAACCGGGGAAGGACGCATGAAACTGGCAAGCTAGAGTACCAAAGAGGGGGGTGGAATTCCCGGTGTAGCGGTGAAATGCGTAGATATCGGGAGGAACACCTGTGGCGAAGGCGGCCCCCTGGTTGGATACTGACGCTGAGACGCGAAAGCGTGGGG |
| P-asv6015 | GCGAACGTTGTTCGGAATCACTGGGCTTAAAGGGCGCGTAGGCGGCAGGGTAGGTCGGGGGTGAAATCCCACAGCTTAACTGTGGAAGAGCCTCCGATACCGTCTTGCTGGAGGAGGGTAGGGGTGCGCGGAACTTCTGGTGGAGCGGTGAAATGCGTTGATATCAGAAGGAACGCCGGCGGCGAAAGCGGCGCACTGGACCCTTTCTGACGCTGAGGCGCGAAAGCCAGGGG |
| P-asv6016 | GCAAGCATTAATCGGATTTATTGGGCGTAAAGGGCGCGTAGGCGGGAAAGAAAGTTAGATGTGAAATCCCAGAGCTCAACTCTGGGGCTGCATCTAATACTTCTTTTCTTGAGGATAGGCGGAGAAAAGGAATTCCACGTGTAGCGGTGGAATGCGTAGATATGTGGAAGAACACCGGTGGCGAAGGCGCTTTTCTAGCTTATTCCTGACGCTGAGGCGCGAAAGCCAGGGG |
| P-asv6017 | CCAAGCGTTATCCGGAATCATTGGGTTTAAAGGGTCCGTAGGCGGGAGAATCAGTCAGTGGTGAAAGTCTGTGGCTTAACCATAGAATTGCCATTGAAACTGTTTTCCTTGAGTCATTATGAAGTGGTTAGAACATGTAGTGTAGCGGCGAAATGCATAGATATTACATAGAATACCGATTGCGAAGGCAGATCACTAATAATGTACTGACGCTGAGGGACGAAAGCGTAGGT |
| P-asv6018 | TCAAGTGGTCGTCGCTTTTATTGGGCCTAAAACGTCCGTAGCCTGTTTGGTAAATCTGTGGGTAAATCAATCCGCTCAACGGATTGAATTCTGCAGAGACTGCCAGACTAGGGACCGGGAGAGGTGTGGGGTACTCGAGGGGTAGGGGTAAAATCCTGTCATCCTTTGAGGACCACCAGTTGCGAAGGCGCCACACTGGAACGGATCCGACGGTCAGGGACGAAGCCTAGGGG |
| P-asv6019 | CCGAGCGTTGTCCGGAATCATTGGGCGTAAAGGGTACGTAGGCGGGTCATTAAGTTAGAAGTTAAAGGCTATAGCTCAACTATAGTAAGCTTCTAAAACTGGAGACCTTGAGTAATGGAAGGGAAAGTGGAATTCCTAGTGTAGCGGTGGAATGCGCAGATATTAGGAGGAATACCGGTGGCGAAGGCGACTTTCTGGCCATTAACTGACGCTGAGGTACGAAAGCGTGGGT |
| P-asv6020 | GCGAGCGTTAATCGGAATTACTGGGCGTAAAGCGTGCGCAGGCGGTTTTGAAAGTCAGATGTGAAATCCCCGAGCTCAACTTGGGAACTGCGTTTGAAACTCCAAAGCTAGAGTATAGGAGAGGGGGGTAGAATTCCACGTGTAGCGGTGAAATGCGTAGAGGTTGGGTGGAACACCGGTGGCGAAGGCGGCTATCTGGCTCACAACTGACGATCAGGCACGAAAGCGTGGGG |
| P-asv6021 | GCTAGCGTTATCCGGAATTACTGGGCGTAAAGGGTGCGTAGGTGGTTTCTTAAGTCAGAGGTGAAAGGCTACGGCTCAACCGTAGTAAGTCTTTGAAACTGGGAAACTTGAGTGCAGGAGAGGAGAGTGGAATTCCTAGTGTAGCGGTGAAATGCGTAGATATTAGGAGGAATACCAGTTGCGAAGGCGGCTCTCTGGACTGTAACTGACACTGAGGCACGAAAGCGTGGGG |
| P-asv6022 | GCGAGCGTTGTCCGGAATTACTGGGCGTAAAGAGCACGTAGGCGGGAATTTAAGTCAGGTGTGAAAGACTGCAGCTTAACTGCGGGGTTGCACTTGAAACTGGATTTCTTGAGGGCAGGAGAGGAAAGCAGAATTCCTGGTGTAGCGGTGAAATGCGTAGAAATCAGGAGGAATACCAGTGGCGAAGGCGGCTTTCTGGACTGACCCTGACGCTGAGGTGCGAAAGCGTGGGT |
| P-asv6023 | GCCAGCGTTGTTCGGAGTTACTGGGCGTAAAGCGCACGTAGGCGGCTTTGTAAGTCAGAGGTGAAAGCCTGGAGCTCAACTCCGAAACTGCCATTGATACTATATATCTTGAATATTGTGGAGGTTTGCGGAATATGTCATGTAGCGGTGAAATGCTTAGATATGACATAGAACACCTATTGCGAAGGCAGCAGGCTACACATATATTGACGCTGAGGCACGAAAGCGTGGGG |
| P-asv6024 | CCGAGCGTTGTCCGGAATTACTGGGCGTAAAGAGCTCGTAGGTGGTTTGTCGCGTTGTTCGTGAAAACTCACAGCTTAACTGTGGGCGTGCGGGCGATACGGGCAGACTGGAGTACTGCAGGGGAGACTGGAATTCCCGGTGTAGCGGTGGAATGCGCAGATATCAGGAGGAACACCGGTGGCGAAGGCGGGTCTCTGGGCAGTAACTGACGCTGAGGAGCGAAAGCGTGGGG |
| P-asv6025 | GCGAACGTTATTCGGAATCACTGGGCTTACAGGGTGCGTAGGCGGCTTATTAAGTAGGGTGTGAAAGCCCCCGGCTCAACCGGGGAATTGCGCCCTAAACTGGCGAGCTGGAGTGAGATAGGGGTGTGTGGAACTTCCGGTGGAGCGGTGAAATGTGTTGATATCGGAAGGAACGCCGGTGGCGAAAGCGACACACTGGATCTTGTCTGACGCTGAGGCACGAAAGCCAGGGG |
| P-asv6026 | GCGAGCGTTGTCCGGAATTATTGGGCGTAAAGAGCTTGTAGGCGGTTTGTCGCGTCTGCTGTGAAAGGCCGGGGCTTAACCCCGTGTATTGCAGTGGGTACGGGCAGACTAGAGTGCAGTAGGGGAGACTGGAATTCCTGGTGTAGCGGTGAAATGCGCAGATATCAGGAGGAACACCGATGGCGAAGGCAGGTCTCTGGGCTGTAACTGACGCTGAGGAGCGAAAGCATGGGG |
| P-asv6027 | GCGAGCGTTATCCGGAATTATTGGGCATAAAGGGCATCTAGGCGGATATACAAGTCAGGGGTGAAAACTTAGGGCTCAACTCAAAGCTTGCCTTTGAAACTGTATATCTAGAGTGCTGGAGAGGTGGACGGAACTACACGAGTAGAGGTGAAATTCGTAGATATGTGTAGGAATGCCGATGATGAAGATAGTCCACTGGACAGCAACTGACGCTGAAGTGCGAAAGCTAGGGG |
| P-asv6028 | GCTAGCGTTGCTCGGAATCACTGGGCGTAAAGGGCGCGTAGGCGGCTTGCCAAGTCGGGGGTGAAAGCCCGTGGCTCAACCACGGAATGGCCTTCGATACTGGCAGGCTTGAGACCGGAAGAGGACAGCGGAACTGCGAGTGTAGAGGTGAAATTCGTAGATATTCACAAGAACACCAGTGGCGAAGGCGGCTGTCTGGTCCGGTTCTGACGCTGAGGCGCGAAAGCGTGGGG |
| P-asv6029 | GCAAGCGTTGTCCGGATTTATTGGGCGTAAAGCGAGCGCAGGCGGTTCTTTAAGTCTGATGTGAAAGCCCCCAGCTCAACTGGGGAAGGTCATTGGAAACTGGGGAACTTGAGTGCAGAAGAGGAGAGTGGAATTCCATGTGTAGCGGTGAAATGCGTAGATATATGGAGGAACACCAGTGGCGAAGGCGACTCTCTGGTCTGTAACTGACGCTGAGGCTCGAAAGCGTGGGT |
| P-asv6030 | CCAAGCGTTATCCGGATTTATTGGGTTTAAAGGGTCCGTAGGCGGGTTATTAAGTCAGTGGTGAAATCCTACAGCTCAACTGTAGAACTGCCATTGAAACTGGTAACCTTGAGTATAATTGAAGTGGGCGGAATGTGTCATGTAGCGGTGAAATGCTTAGATATGACACAGAACACCGATTGCGAAGGCAGCTCACTAAGTTATAACTGACGCTGAGGGACGAAAGCGTGGGT |
| P-asv6031 | GCAAGCGTTAATCGGAATTACTGGGCGTAAAGCGCACGTAGGTGGTTTTCTAAGCCAGCTGTGAAATCCCCGGGCTCAACCTGGGCACTGCAGTTGGAACTGGAAAGCTAGAGTAGGGTAGAGGGGTGTGGAATTCCAGGTGTAGCGGTGAAATGCGTAGATATCTGGAGGAACATCAGTGGCGAAGGCGACACCCTGGACTCATACTGACACTGAGGTGCGAAAGCGTGGGG |
| P-asv6032 | ACAAGCGTTGTCCGGATTTATTGGGCGTAAAGGGAGCGCAGGCGGTCTGTTTAGTCTAATGTGAATGCCCACGGCTTAACCGTGGAACGGCATTGGGAACTGACAGACTTGAATGTAGAAGAGGAAAATGGAATTCCAAGTGTAGCGGTGGAATGCGTAGATATTTGGAGGAACACCAGTGGCGAAGGCGATTTTCTGGTCTAACATTGACGCTGAGGCTCGAAAGCGTGGGG |
| P-asv6033 | GCAAGCGTTAATCGGAATTACTGGGCGTAAAGCGCGCGTAGGCGGTCAGATAAGTCAGATGTGAAATCCCCGGGCTCAACCTGGGAACTGCACCTGATACTGTCTGACTAGAGTTTAGGAGAGGGAAGTAGAATTTCAGGTGTAGCGGTGAAATGCATAGAGATCTGAAGGAATACCAGTGGCGAAGGCGACTTCCTGGCCTAAAACTGACGCTGAGGTGCGAAAGCGTGGGT |
| P-asv6034 | GCGAGCGTTAATCGGAATAACTGGGCGTAAAGGGCACGCAGGCGGTGACTTAAGTGAGGTGTGAAAGCCCCGGGTTTAACCTGGGAATTGCATTTCATACTGGGTCGCTAGAGTACTTTAGGGAGGGGTAGAATTCCACGTGTAGCGGTGAAATGCGTAGAGATGTGGAGGAATACCGAAGGCGAAGGCAGCCCCTTGGGAATGTACTGACGCTCATGTGCGAAAGCGTGGGG |
| P-asv6035 | GCAAGCGTTAATCGGAATTACTGGGCGTAAAGCGCGCGTAGGTGGTTTGTTAAGTTGGATGTGAAATCCCCGGGCTCAACCTGGGAACTGCATTCAAAACTGACTGACTAGAGTATGGTAGAGGGTGGTGGAATTTCCTGTGTAGCGGTGAAATGCGTAGATATAGGAAGGAACACCAGTGGCGAAGGCGACCACCTGGACTGATACTGACACTGAGGTGCGAATGCGTGGGG |
| P-asv6036 | GCGAGTGTTATCCGGAATTATCGGGCGTAAAGCGTCTGTAGGTTGAGTTCAAAGTCTTTTGTAAAATCTTTAAGCTCAACTTAAAAACCGCAAAAGTTACTAGTTTTCTTGAGTATGGTAGAGGTAAAGGGAATTTCCAGTGGAGCGGCGAAATGCGTAGAGATTGGAAGGAACACCAAAAGCGAAGGCACTTTACTGGGCCATTACTGACACTGAGAGACGAAAGCTAAGGG |
| P-asv6037 | GCAAGCGTTAATCGGAATTACTGGGCGTAAGGCGTGCGCAGGCGGTTCGGAAAGAAAGATGTGAAATCCCAGAGCTTAACTTTGGAACTGCATTTTTAACTACCGGGCTAGAGTGTGTCAGAGGGAGGTGGAATTCCGCGTGTAGCAGTGAAATGCGTAGATATGCGGAGGAACACCGATGGCGAAGGCAGCCTCCTGGGATAACACTGACGCTCATGCACGAAAGCGTGGGG |
| P-asv6038 | GCGAGCGTTGTCCGGATTTATTGGGCGTAAAGGGCTCGTAGGTGGTTGATCGCGTCGGAAGTGTAATCTTGGGGCTTAACCCTGAGCGTGCTTTCGATACGGGTTGACTTGAGGAAGGTAGGGGAGAATGGAATTCCTGGCGGAGCGGTGGAATGCGCAGATATCAGGAGGAACACCAGTGGCGAAGGCGGTTCTCTGGGCCTTTCCTGACGCTGAGGAGCGAAAGCGTGGGG |
| P-asv6039 | GCTAGCGTTGTTCAGAATTACTGGGCGTAAAGCGCACGTAGGCGGCTTTGTAAGTCAGAGGTGAAAGCCTGGAGCTCAGCTCCAGAAATGCCTTCAATACTGGCTGTCTTGAGTACGGGAGAGGTGAGTGGAACTCCGAGTGTAGAGGTGAAATTCGTAGATATTCGGAAGAACACCAGTGGCGAAGGCGACTCACTGGCCCGTTACTGACGCTGAGGCGCGAAAGCGTGGGG |
| P-asv6040 | CCAAGCGTTATCCGGAATCATTGGGTTTAAAGGGTCCGTAGGCGGGACAGTAAGTCAGTGGTGAAAGTCTGCAGCTCAACTGTAGAACTGCCATTGAAACTGTTGTCCTTGAGTCATTATGAAGTGGTTAGAATATGTAGTGTAGCGGTGAAATGCATAGATATTACATAGAATACCGATTGCGAAGGCAGATCACTAATAATGTACTGACGCTGAGGGACGAAAGCGTAGGT |
| P-asv6041 | TCCCTCTTTCTCCGGAATCACTGGGCGTAAAGGGTGCGTAGGCGGCCTAACAAGTCAGAGGTGAAAGGCATTGGCTCAACCAATGTAAGCCTTTGAAACTGTTAGGCTTGAGTGCAGGAGAGGTAAGTGGAATTCCTAGTGTAGAGGTGAAATTCGTAGATATTAGGAGGAACACCAGTGGCGAAGGCGACTTACTGGACTGTAACTGACGCTGAGGCACGAAAGCGTGGGG |
| P-asv6042 | GCAAGCGTTATCCGGAATTATTGGGCGTAAAGCGTCCGTAGGTGGCTTTCTCAAGTCTGCGGTCAAAGGCTGGGGCTTAACTCCAGAAAGGCCGTGGAAACTGAAAGGCTAGAGAGCCGTAGGGGTAGAGGGAATTCCCGGTGTAGCGGTGAAATGCGTAGAGCTCGGGAAGAACATCGGTGGCGAAAGCGCTCTACTGGACGGTATCTGACACTGAGGGACGAAAGCTAGGGG |
| P-asv6043 | GCTAGCGTTGTTCGGAATTACTGGGCGTAAAGCGCACGTAGGCGGATCGATCAGTCAGGGGTGAAATCCCAGAGCTCAACTCTGGAACTGCCTTTGATACTGTCGATCTAGAGTATGGAAGAGGTGAGTGGAATTCCGAGTGTAGAGGCGAAATTCGTAGATATTCGGAGGAACACCAGTGGCGAAGGCGGCTCACTGGTCCATTACTGACGCTGAGGTGCGAAAGCGTGGGG |
| P-asv6044 | CCAAGCGTTGTTCGGATTTACTGGGCGTAAAGGGCGCGTAGGCGGCCTGGCAAGTCACTTGTGAAATCTCCGAGCTTAACTCGGAACGGTCAAGTGAAACTGCCATGCTAGAGTGCAGAAGGGGCAATCGGAATTCTTGGTGTAGCGGTGAAATGCGTAGATATCAAGAAGAACACCTGAGGTGAAGACGGGTTGCTGGGCTGACACTGACGCTGAGGCGCGAAAGCCAGGGG |
| P-asv6045 | GCAAGCGTTATCCGGAATCATTGGGCTTAAAGGGTCCGCAGGCGGACGTATAAGTCAGTGGTGAAATCCTGCAGCTTAACTGCAGAACTGCCATTGATACTGTACGTCTTGAATTCGGTCGAAGTGGGCGGAATGTGTAGTGTAGCGGTGAAATGCATAGATATTACACAGAACACCGATAGCGAAGGCAGCTCACTAGGCCTGGATTGACGCTCAGGGACGAAAGCGTGGGG |
| P-asv6046 | GCAAGCGTTGTCCGGATTTACTGGGCGTAAAGGGAGCGTAGGCGGATCTTTAAGTGGGATGTGAAATACTCGGGCTCAACCTGGGGGCTGCATTCCAAACTGGGGATCTAGAGTACAGGAGGGGAGAGTGGAATTCCTAGTGTAGCGGTGAAATGCGTAGAGATTAGGAAGAACACCAGTGGCGAAGGCGACTCTCTGGACTGTAACTGACGCTGAGGCTCGAAAGCGTGGGG |
| P-asv6047 | GCGAGCGTTATCCAGATTTATTGGGTTTAAAGGGTGCGCAGGCGGGATTTTAAGTCAGCGGTGAAATTTTCAGGCTCAACCTGAACACTGCCGTTGAAACTGGGATTCTTGAGTATGGATGAAGTAGGCGGAATTCGTTGTGTAGCGGTGACATGCTCAGATATAACGAGGAACTCCGATTGCGTAGGCAGCTTACTGAGCCATAACTGACGCTCAAGCACGAAAGCGTGGGG |
| P-asv6048 | GCAAGCGTTGTCCGGAATTATTGGGCGTAAAGCGCGCGCAGGCGGCTATGTAAGTCTGGTGTTAAAGCCCGGGGCTCAACCCCGGTACGCATCGGAAACTGTGTAGCTTGAGTGCAGAAGAGGAAAGCGGTATTCCACGTGTAGCGGTGAAATGCGTAGAGATGTGGAGGAACACCAGTGGCGAAGGCGGCTTTCTGGTCTGTAACTGACGCTGAGGCGCGAAAGCGTGGGG |
| P-asv6049 | GTAAACGTTGCCCGGATTTATTGGGCGTAAAGTGTTTCGTAGGCGACTCATCGCATCTTGTTTTAAAGCTTCTGGCTCAACCAGAAAAAGGGACAAGAGATGGGTGAGATTGAAGGTTGTCGGGGTCACTGGAACTGTCGGTGTAGGGGTGAAATCCGTTGATATCGATAGGAACTCCAAAGGCGTAGGCGGGTGACTAGGCAAATCTTGACGCTGATGAACGAGAGCTAGGGG |

| | |
|---|---|
| P-asv6050 | GCAAGCGTTATCCGGATTTACTGGGTGTAAAGGGAGCGCAGACGGTTGCGCAAGTCTGAAGTGAAATCCCGAGGCTTAACCACGGGACTGCTTTGGAAACTGTGCGACTTGAGTATCGGAGGGGCAGGCGGAATTCCTAGTGTAGCGGTGAAATGCGTAGATATTAGGAAGAACACCGGTGGCGAAGGCGGCCTGCTGGACGAAAACTGACGTTGAGGCTCGAAGGCGTGGGG |
| P-asv6051 | GCAAGCGTTATCCGGACTTACTGGGTTTAAAGGGTGCGCAGGCGGTAATTTAAGTCAGTGGTGAAAGCCTACAGCTCAACTGTAGAACTGCCATTGAAACTGAAATACTTGAGTGTGGTAGAGGTAGGCGGAATGAGTCATGTAGCGGTGAAATGCATAGATATGACTCAGAACACCGATTGCGAAGGCAGCTTACTAAACCATTACTGACGCTCAGGCACGAAAGCGTGGGG |
| P-asv6052 | GCAAGCGTTGTCCGGAATTACTGGGCGTAAAGAGCTCGTAGGTGGTTTGTCGCGTCGTCTGTGAAATCCCGGGGCTTAACTCCGGGTCTGCAGGCGATACGGGCATAACTAGAGTGCTGTAGGGGAGACTGGAATTCCTGGTGTAGCGGTGGAATGCGCAGATATCAGGAGGAACACCGATGGCGAAGGCAGGTCTCTGGGCAGTAACTGACGCTGAGGAGCGAAAGCATGGGG |
| P-asv6053 | GCAAGCATTAATCGGATTTATTGGGCGTAAAGGGCGCGTAGGCGGGAACATAAGTCAGATGTGAAATCCTGGAGCTCAACTCCAGAACAGCATTTGAAACTATATTTCTTGAGGGTAGGCGGAGAAAACGGAATTCCACATGTAGCGGTGAAATGCGTAGATATGTGGAAGAACATCGGTGGCGAAGGCGGTTTTCTAGCTTATTCCTGACGCTGAGGCGCGAAAGCAAGGGA |
| P-asv6054 | GCAAGCGTTACTCGGAATTACTGGGCGTAAAGCGTGCGTAGGTGGTTTGTTAAGTCTGATGTGAAAGCCCTGGGCTCAACCTGGGAATGGCATTGGAAACTGGCGGACTCGAGTGCGGTAGAGGGGTGTGGAATTCCCGGTGTAGCAGTGAAATGCGTAGAGATCGGGAGGAACACCAGTGGCGAAGGCGACACCCTGGACCAGCACTGACACTGAGGCACGAAAGCGTGGGG |
| P-asv6055 | GCAAGCGTTATCCGGAATCACTGGGTTTAAAGGGTGCGTAGGCGGTTAGATAAGTCAGAGGTGAAAGTTTGCAGCTTAACTGTAAAATTGCCTTTGAAACTGTATAACTTGAATAGCGTTGAGGTCAGCGGAATGTGACATGTAGCGGTGAAATGCATAGATATGTCATAGAACACCAATTGCGAAGGCAGCTGGCTGGGCGTTTATTGACGCTGAGGCACGAAAGCGTGGGG |
| P-asv6056 | GCGAGCGTTGTCCGGAATTATTGGGCGTAAAGGGTGCGCAGGCGGCTTTACAAGTTGGATGTGAAATATTGTGGCTCAACCACAAACGTGCATCCAAAACTGCAAAGCTTGAGTTAAGGAGAGGTAAGTGGAATTCCTGGTGTAGCGGTGGAATGCGTAGATATCAGGAGGAATACCGGTGGCGAAGGCGACTTACTGGACTTAAACTGACGCTCAGGCACGAAAGCGTGGGG |
| P-asv6057 | GCTAGCGTTGTTCGGAATTACTGGGCGTAAAGCGCACGTAGGCGGCCTTGTAAGTTGGGGGTGAAATCCCGGAGCTCAACTCCGGAACTGCCTCCAAAACTACAAAGCTCGAGAATGTGAGAGGTGAGTGGAATTCCGAGTGTAGAGGTGAAATTCGTAGATATTCGGAAGAACACCAGTGGCGAAGGCGACTCACTGGCACATTTCTGACGCTGAGGTGCGAAAGCGTGGGG |
| P-asv6058 | GCAAGCGTTGTCCGGAATTATTGGGCGTAAAGGGCTCGCAGGCGGTTTCTTAAGTCTGATGTGAAAGCCCCCGGCTCAACCGGGGAGGGTCATTGGAAACTGGGAAACTTGAGTGCAGAAGAGGAGAGTGGAATTCCATGTGTAGCGGTGAAATGCGTAGATATATGGAGGAACACCAGTGGCGAAGGCGGCTCTCTGGTCTGTAACTGACGCTGAGGCTCGAAAGCGTGGGG |
| P-asv6059 | GCAAGCGTTATCCGGAATCATTGGGCGTAAAGGGTCCTGAGGTGGCCTGCCCAGTCTACTGTGAAAGATCAGGGCTCAACCCTGAAGATGCGGTAGAAACTAGCAGGCTAGGATACGGTAGGGGCAGGGGGAATTCCCGTTGTAGCGGTGAAATGCATAGATATCGGGAGGAACACCAAAGGCGAAGGCACCCTGCTGGGCCGATATTGACACTGACGGACGAGAGCTAGGGG |
| P-asv6060 | GCGAGCGTTGTTCGGAATCACTGGGCGTAAAGCGTGCGTAGGCGGTTATACAAGTCAGAAGTGAAAGCCAGGGCTTAACCCTGGAATTGCTTTTGAGACTGTATGGCTAGAGATCTAGAGAGGTTGGGGGAATTCCGAGTGTAGAGGTGAAATTCGCAGATATTCGGAGGAACACCAGTGGCGTAGGCGCCCAACTGGCTAGATACTGACGCTGAGGCACGAAAGCGTGGGG |
| P-asv6061 | GCGAGCGTTATTCGGAATTATTGGGCGTAAAGGGTGCGTAGACGGCACGTTAAGTCAACTGTTAAATTCCTCGGCCTAACTGAGGATCAGCAGTAGAAACTGGCGCGCTTGAGGATGAGAGAGAGAAGTGGAATTCTCGGAGTAGCGGTAAAATGCGTAGATCTCGAGAGGAACACCGATGGCGAAGGCAGCTTCTTGGCTCATTCCTGACGTTGAGGCACGAAAGCGTGGGG |
| P-asv6062 | GCAAGCGTTGTCCGGATGTACTGGGTGTAAAGGGCGTGTAGGCGGGTCAGCAAGTCAGATGTGAAATGCCGGGGCTTAACTCCGGAACTGCAACTGAAACTGTTGATCTTGAGTACTGGAGAGGAAAGTGGAATTCCTAGTGTAGCGGTGAAATGCATAGAGATTAGGAGGAACACCAGTGGCGAAGGCGGCTTTCTGGACAGTAACTGACGCTGAGGCGCGAAAGCGTGGGG |
| P-asv6063 | GCTAGCGTTGTTCGGATTTACTGGGCGTAAAGGGAGCGTAGGCGGTTTTTCAAGTCAGGCGTGAAAGCCCGAGGCTTAACCTCGGAATTGCGCTTGAAACTGATTGACTGGAGTATGGGAGAGGATAGCGGAATTCCTAGTGTAGAGGTGAAATTCGTAGATATTAGGAAGAACACCGGCGGCGAAGGCGGCTATCTGGACCATTACTGACGCTGAAGCTCGAAAGCATGGGG |
| P-asv6064 | CCAAGTGGTAGTCATTATTATTGGGTTTAAAGTGTTCGTAGCCGGCCGTACAAGTCTGCTGTGAAATCTATGCGCTCAACGGGTGGGCGGCCGACGGAAACTGGACGGCTTGGGACCGGGAGGCTTGAAGGGTACGTCCGGGGTAGGAGTGAAATCCCGTAATCCTGGACGGACCACCGATGGGGAAACCACTTCAAGAGAACGGATCCGACAGTGAGGGACGAAAGCTAGGGT |
| P-asv6065 | GCAAGCGTTGTTCGGAGTTACTGGGCGTAAAGAGTCTGTAGGTGGTCTGTCAAGTCTTTGGTGAAAGGCCGTGGCTTAACCATGGGAATGCCAAAGAGACTGGCAGACTGGAGGCTGGGAGAGGGAAGCGGAATTTCTGGTGTAGCGGTGAAATGCGTAGATATCAGAAGGAGGCCGGTGGCGAAGGCGGCTTCCTGGAACAGACCTGACACTGAGAGACGAAAGCGTGGGG |
| P-asv6066 | CCAAGCGTTAGTCGGAATCACTGGGCTTAAAGCGTGCGTAGGCGGATCGGTAGGCGTCTTGTGAAAGCCCACGGCTCAACCGTGGAATTGCAGGGCGAACCATCGATCTTGAGGCAGGTAGGGGTCGATAGAACTGTGGGTGGAGCGGTGAAATGTGTAGATATCCACAGGAATGCCGGTGGCGAAGGCGATCGACTGGGCCTGTCCTGACGCTGAGGCACGAAAGCGTGGGG |
| P-asv6067 | GCGAACGTTGTCCGGATTCACTGGGCGTAAAGCGCGCGCAGGCGGGTTGGTAAGTCTGAGGTGAAAGTTCCCGGCTCAACTGGGAGAAGCCCTTGGATACTGCCAGCCTTGAGACACTTAGAGGAGAGTGGAATTCCCGGTGTAGTGGTGGAATGCGTAGATATCGGGAGGAACACCAGTGGCGAAAGCGGCTCTCTGGGAGTGTTCTGACGCTGAGGCGCGAAAGCGTGGGG |
| P-asv6068 | GCAAGCATTAATCGGATTTACTGGGTGCAAAGGGCGCGTAGGCGGGTAAGCAAGTCAGATGTGAAATTCCGGAGCTCAACTCCGGAGCTGCATTTGAAACTACTTATCTTGAGGGTGGACGGAGAAAACGGAATTCCACGTGTAGCGGTGAAATGCGTAGATATGTGGAAGAACACCGGTGGCGAAGGCGGTTTTCTAGTTCATTCCTGACGCTGAGGCGCGAGAGCAAGGGG |
| P-asv6069 | GCTAGCGTTATTCGGAATTACTGGGCGTAAAGCGCACGTAGGCGGCTCAATGAGTCAGTGGTGAAAGCCGGCAGCTTAACTGTCGGATTGCCATTGATACTGTTGAGCTTGAGTATAGTTGAGGTTGGCGGAATGTATCATGTAGCGGTGAAATGCTTAGATATGATACAGAACACCGATTGCGAAGGCAGCTGACTAAGCTATTACTGACGCTCATGCACGAAAGCGTGGGG |
| P-asv6070 | ACGAGTGATGGCCGATCTTATTGGGCCTAAAGCGTCCGTAGCTGGCCGTACAAGTCCGTCGGGAAATCCACCAGCTCAACTGGTGGGCGTCCGGCGGAAACTGTGTGGCTTGGAACCGGAAGGCTCAGAGGGTACGTCCGGGGTAGGAGTGAAATCCCGTAATCCCGGACGGACCACCGATGGCGAAAGCACTCTGAGAAGACGGCTTCGACAGTGAGGGACGAAAGCTAGGGT |
| P-asv6071 | GCGAGCGTTAATCGGAGTTACTGGGCGTAAAGGGCGCGTAGGTGGTGATACGAGTCAGAGGTGAAAGCCCTAGGCTTAACCTAGGAATTGCCTATGATACGGTATGACTAGAGTCGCGGAGAGGATGGTGGAATTTCCGGAGTAGCGGTGAAATGCGTAGATATCGGAAGGAACACTGGTGGCGAAGGCGACCATCTGGAAGCAGACTGACACTGAGGCGCGAAGGCGTGGGG |
| P-asv6072 | GCAAGCGTTGTCCGGAATTATTGGGCGTAAAGCGCGCGCAGGTGGTTTAATAAGTCTGATGTGAAAGCCCACGGCTCAACCGTGGAGGGTCATTGGAAACTGTTAAACTTGAGTGCAGGAGAGAAAAGTGGAATTCCTAGTGTAGCGGTGAAATGCGTAGAGATTAGGAGGAACACCAGTGGCGAAGGCGGCTCACTGGCCCGATACTGACGCTGAGGCACGAAAGCGTGGGG |
| P-asv6073 | GCTAGCGTTGTTCGGAATTACTGGGCGTAAAGGGCGCGTAGGCGGCTTTTTGCGTGAGATGTGAAAGACCTGGGCTCAACCTGGGAAGTGCATTTCATACGGGAAAGCTAGAGTGCGAGAGGGGAAAGTGGAATGATGCGTGTAGAGGTGAAATTCGTAGATATGCATCGGAGCACCAGTGGCGAAGGCGACTTTCTGGCTCGTAACTGACGCTAAGGCGCGAAAGCGTGGGG |
| P-asv6074 | GCAAGCGTTGTCCGGAATTATTGGGCGTAAAGAGCTCGTAGGTGGTTCGATACGTCGGATGTGAAAATCAGGGGCTCAACCCCTGACCTGCATCCGATACGGTCGAGCTAGAGTTTGGTAGGGGAGACTGGAATTCCTGGTGTAGCGGTGGAATGCGCAGATATCAGGAGGAACACCGATGGCGAAGGCAGGTCTCTGGGCCAATACTGACACTGAGGAGCGAAAGCGTGGGG |
| P-asv6075 | GCAAGCGTTATTCGGAATTACTGGGCGTAAAGCGCGCGTAGGCGGATTGTTAAGTCAGTTGTGAAAGCCCTGGGCTCAACCTAGGAGTTGCATCTGATACTGGCAGTCTAGAGTTTAAGAGAGGGAAGGGGAATTCCAGGTGTAGCAGTGAAATGCGTAGATATCTGGAGGAACATCAGTGGCGAAGGCGCCTTCCTGGCTTAAAACTGACGCTGAGGTGCGAAAGCGTGGGT |
| P-asv6076 | GCAGACGTTGTTCGGAATTACTGGGCGTAAAGCGTGTGTAGGCGGCTATGTAAGTCGGATGTGAAAGCCCTGGGCTCAACCCAGGAGGTGCACTCGATACTGCGTAGCTTGAGTCTCGGAGAGGTTGGTGGAATTCTCGGTGTAGAGGTGAAATTCGTAGATATCGAGAGGAACACCGGTGGCGAAGGCGGCCAACTGGACGAAGACTGACGCTGAGACACGAAAGCGTGGGG |
| P-asv6077 | GCAAGCGTTACTCGGAATTACTAGGCGTAAAGGGCAGGTAGGTGGTTTGATAAGTCTGTTGCGAAAGCTCCTGGCTTAACTGGGAGAGGTCAACAGATACTATTGAACTTGAGTATAGGAGAGGTTACTGGAATTCCCGGTGTAGCGGTGAAATGCGCAGAGATCGGGAGGAACACCAATGGCGAAAGCAGGTAACTGGACTATTACTGACACTCAGCTGCGAAAGCTAGGGG |
| P-asv6078 | TCTCTCTTTTCTCTTCCTCCCTTTTCTTAAAGGGTGCGTAGGCGGGTCTTTAAGTCAGGGGTGAAATCCTGGAGCTCAACTCCAGAACTGCCTTTGATACTGAAGATCTTGAGTATGGGAGAGGTGAGTGGAACTGCGAGTGTAGAGGTGAAATTCGTAGATATTCGCAAGAACACCAGTGGCGAAGGCGGCTCACTGGCCCATAACTGACGCTGAGGCACGAAAGCGTGGGG |
| P-asv6079 | CCGAGCGTTATCCGGATTTATTGGGCGTAAAGGGCGCGTAGGCGGCTTGGTGCGTCTCATGTTAAGTCTCACTGCTCAACGGTGAAATCGCATGAGGTACGATCAAGCTTGAGAGTGTTAGAGGTAGATGGAACTCATGGTGTAGGGGTGACATCCGTTGATATCATGGGGAACACCAAAGGCGAAGGCATTCTACTGGGACATTTCTGACGCTGATGCGCGAAAGCGTGGGG |
| P-asv6080 | GCAAGCGTTGTCCGGAATTATTGGGCGTAAAGCGCGCGCAGGCGGTCATTTAAGTCTGGTGTTTAAACCTTGGGCTCAACCTGGGGTCGCACTGGAAACTGGGTGACTTGAGTACAGAAGAGGAAAGTGGAATTCCACGTGTAGCGGTGAAATGCGTAGATATGTGGAGGAACACCAGTGGCGAAGGCGACTTTCTGGGCTGTAACTGACGCTGAGGCGCGAAAGCGTGGGG |
| P-asv6081 | GCGAGCGTTAATCGGATTTACTGGGCGTAAAGGGCGCGTAGGCGGTGAGATGTGTGTGATGTGAAAGCCCCGGGCTTAACCTGGGAAGTGCATCGCAAACTGTCTGACTGGAGTATATGAGAGGGTGGCGGAATTTCCGGTGTAGCGGTGAAATGCGTAGATATCGGAAGGAACGTCGATGGCGAAGGCAGCCACCTGGCATAATACTGACGCTGAGGCGCGAAAGCGTGGGG |
| P-asv6082 | GCGAGCGTTGTTCGGAGTCACTGGGCGTAAAGAGTACGTAGGTGGACTGGCAAGTCAGAAGTGAAATCCCGGGGCTCAACCCCGGAATGGCTTTTGAAACTGCAAGTCTAGAGTCCCGGAGGGGCCAGCGGAATTCCGAGTGTAGAGGTGAAATTCGCAGATATTCGGAAGAACACCAGTGGCGAAGGCGGCTGGCTGGACGGGTACTGACACTGAGGTACGAAAGCGTGGGG |
| P-asv6083 | GCAAGCGTTGTCCGGATTTACTGGGCGTAAAGGGAGCGTAGGCGGACACTTAAGTGGGACGTGAAATCCCCGGGCTTAACCCGGGAGCTGCATTCCAAACTGGGTGTCTAGAGTGCAGGAGAGGAAAGTGGAATTCCTAGTGTAGCGGTGAAATGCGTAGAGATTAGGAAGAACACCAGTGGCGAAGGCGACTTTCTGGACTGTAACTGACGCTGAGGCTCGAAAGCGTGGGG |
| P-asv6084 | TCAAGTGGTCAGGAGGATTATTGGGCCTAAAGCATCCGTAGCCGGTTCTGCAAGTTTTCGGTTAAATCCATTCGCTCAACGAATGGGCTGCCGAAAATACTACAGACCTAGGGAGTGGGAGAGGTAGACGGTATTCGGTAGGAAGGGGTAAAATCCTTTGATCTATTGAGGACCACCTATGGCGAAGGCAGTCTACCAGAACACGTTCGACGGTGAGGGATGAAAGCTGGGGG |
| P-asv6085 | GCAAGCGTTAATCGGAATTACTGGGCGTAAAGCGTGCGTAGGCGGATGCTTAAGTCAGATGTGAAAGCCCTGGGCTTAACCTAGGAATTGCATCTGATACTGGGCATCTAGAGTATGGTAGAGGATAGTGGAATTTCAGGTGTAGCGGTGAAATGCGTAGATATCTGAAGGAACACCAGTGGCGAAGGCGACTGTCTGGGCCAATACTGACGCTGAGGTACGAAAGCATGGGT |
| P-asv6086 | GCTAGCGTTACTCGGAATTATTGGGCGTAAAGCGTGCGTAGGCGGCTTTGCAAGTTGGAAGTGAAAGCCCAGGGCTTAACCCTGGAATTGCTTTCAAAACTACAAGGCTAGAATGTCAGAGGGGATAGCGGAATTCCTAGTGTAGAGGTGAAATTCGTAGATATTAGGAGGAACACCGGAGGCGAAAGCGGCTATCTGGCTGACAATTGACGCTGTGGCACGAAAGCGTGGGG |
| P-asv6087 | GCAAGCGTTGTCCGGAATGATTGGGCGTAAAGCATCTGTAGGTGGTTTTTAAAGTCGACTGTTAAAGCCCAGGGCTTAACTTTGGAGAAGCGGTCGAGTACTTAAGGACTTGAGTTTGGTAGGGGTAGAGGGAATTCCTAGTGGAGCGGTGAAATGCACAGAGATTAGGAAGAACACCGAGGGCGAAAGCACTCTACTGGGCCACAACTGACACTGAGAGATGAGAGTCGGGGT |
| P-asv6088 | GCTAGCGTTGTTCGGAATTACTGGGCGTAAAGGGCGCGTAGGCTGCCATGTGTGTCAGGTGTGAAATCCCAAGGCTTAACCTTGGAACTGCATCTGAAACTGCATGGCTTGAGTGAGTAAGAGGCGGACGGAATTCCGAGTGTAGCAGTGAAATGCGTAGATATTCGGAGGAACATCGGTGGCGAAGGCGGTTCGCTGGTACTCTACTGACGCTGAGGCGCGAAAGCGTGGGG |

| | |
|---|---|
| P-asv6089 | CCGAGCGTTAATCGGAATCACTGGGCTTAAAGGGTGCGTAGGCGGATGTGTAGGTGTCTTGTGAAATCCCTCCGCTCAACGGAGGAACTGCACGGCAAACCACACATCTTGAGGCAGGTAGAGGCCGTCGGAAC<br>GATAGGTGGAGCGGTGAAATGCGTAGATATCTATCGGAACGCCGAAGGTGAAGACAGACGGCTGGGCCTGTCCTGACGCTGAGGCACGAAAGCGTGGGG |
| P-asv6090 | GCAAGCGTTGTTCGGAATGACTGGGCGTAAAGAGCGCGTAGGCGGGTTCGTATGTCATCTGTGAAATCCCCGGGCTTAACCTGGGAATTGCAGGTGAAACTGCGAATCTTGAATACTGGAGGGGATGGTGGAAT<br>TCCTGGTGTAGAAGTGAAATTCATAGATATCAGGAGGAACACCTGAGGCGAAGGCGGCCATCTGGCCATGTATTGACGCTGTGGCGCGAAAGCGTGGGT |
| P-asv6091 | CCAAGCGTTGTCCGGATTTATTGGGTTTAAAGGGTGCGTAGGCGGACCATTAAGTCAGAGGTGAAAGTTTGCAGCTTAACTGTAAAATTGCCTTTGAAACTGGTGGTCTTGAGTATAGTTGAGGTGGGCGGAATGT<br>GTCATGTAGCGGTGAAATGCTTAGATATGATACAGAACACCGATTGCGAAGGCAGCTCACTAAACTGTAACTGACGCTGAGGCACGAAAGCGTGGGG |
| P-asv6092 | ACAAGTGTTACTCGTCATAATTGGGCGTAAAGGGTGCGCAGGCGGTTCTTTAAGTTATAGGTGAAAGGTTAATTTTAAGAGTTAAAAAGACCTATAAAACTAAAGGAGTAGAGTATACAAGAGGAAAATGGAACTTT<br>TACAGTAGAGGTGGAATTCTTGAATAGTAAAGGGACGACCAAAGGCGAAGGCAATTTTCTGGGGTATTACTGACGCTGAAGCACGAAAGCATGGGG |
| P-asv6093 | GCAAGCGTTGTCCGGATTTACTGGGTTTAAAGGGTACGTAGGCGGTTGTGCCAAGTCTGGGGTGAAAGTTTGCAGCTCAACTGTAAAATTGCCTTGGATACTGGTCGGCTAGAGTAGTGGAGCGGTGGCTGGAA<br>TTTATAGTGTAGCGGTGAAATGCTTAGATACTATGAGGAACACCAATTGCGAAGGCAGGTCACTGGTCACTAACTGACGCTGAGGTACGAAAGTGTGGGG |
| P-asv6094 | GCGAGCGTTAATCGGAATTACTGGGCGTAAAGGGTGCGCAGGTGGTTTTTTAAGTTATCTATGAAAGACACGGGCTTAACCTGTGTTAGTTGGATGATACTGGGAGACTTGAGTATAGGAGAGGGTAGTGGAATT<br>TCCGGTGTAGCGGTGAAATGCGTAGATATCGGAAGGAACACCGGTGGCGAAGGCGGCTACCTGGCCTGATACTGACACTGAGGCACGAAAGCGTGGGT |
| P-asv6095 | GCGAGCGTTGTTCGGAATCACTGGGCGTAAAGGGCGTGTAGGCGGTCCGCTAAGTCTGATGTGAAAGCCCGGGGCTCACCCCCGGAAGTGCATTGGAAACTGGTGGACTCGAGTCCCGGAGAGGAAGGTGGA<br>ATTCCTGGTGTAGCGGTGAAATGCGTAGATATCAGGAGGAATACCGGTGGCGAAGGCGGCCTTCTGGACGGTGACTGACGCTGAGACGCGAAAGCGTGGGG |
| P-asv6096 | ACGAGCGTTGTCCGGAATTACTGGGCGTAAAGGGCGCGCAGGCTGTGCTGCAAGTCAGCTGTTAAAGACTGGGGCTTAACCCTGGTATGCGGTTGAGACTGCAGTACTGGAGTGCTGTAGAGGCAAGTGGAATT<br>CCCAGTGTAGCGGTGAAATGCGTAGATATTGGGAAGAACACCGGTGGCGAAGGCGACTTGCTGGGCAGCAACTGACGCTGAGGCGCGAAAGCCAGGGG |
| P-asv6097 | GCGAGCGTTATCCGGAATTATTGGGTTTAAAGGGTACGCAGGCGGCCTAATAAGTCAGTGGTGAAATCTCTCGGCTCAACCGAGAAACTGCCATTGATACTGTTGGGCTTGAATATAGTTGAAGTAGGCGGAATG<br>TGTAGTGTAGCGGTGAAATGCTTAGATATTACACAGAACACCGATAGCGAAGGCAGCTTACTAAACTATTATTGACGCTCATGTACGAAAGCGTGGGG |
| P-asv6098 | GCAAGCGTTGTTCGGAATCATTGGGCTTAAAGCGCGTGTAGGCGGCCTCGCAAGTACTTTGTGAAATCCCCGGGCTCAACCTGGGAATGGCTGGGTATACTGCGGGGCTTGAGGCTGGTAGGGGTGCCTGGAA<br>CTCTAGGTGGAGCGGTGAAATGCGTAGATATCTAGGGGAACGCCCGTGGCGAAAGCGGGGCACTGGGCCAGACCTGACGCTGAGACGCGAAAGCGTGGGG |
| P-asv6099 | GCGAGCGTTGTCCGGAGTCACTGGGCGTAAAGGGCGCGTAGGTGGCGGGGTAAGCGTGTGGTGAAAGTCCGGGGCTCAACCCCGGATCGGCCATGCGAACTGCCTCGCTGGAGCACGGTAGAGGCACATGG<br>AATTCCGGGTGTAGCGGTGGAATGCGTAGAGATCCGGAAGAACACCGGTGGCGAAGGCGGTGTGCTGGGCCGTTGCTGACACTGAGGCGCGACAGCGTGGGG |
| P-asv6100 | GCTAGCGTTGTTCGGAATCACTGGGCGTAAAGCGCGCGCAGGCGGCTATTCAAGTCAGGGGTGAAATCCCTGGGCTCAACCCAGGAATTGCCTTTGAAACTGTATGGCTGGAGTGTCGGAGGGGATAGCGGAA<br>TTCCTAATGTAGCGGTGAAATGCGTAGATATTAGGAGGAACACCGGTGGCGAAGGCGGCTATCTGGACGACAACTGACGCTGAGGCGCGAAAGCGTGGGG |
| P-asv6101 | GCAAGCGTTGTCCGGAATTATTGGGCGTAAAGCGCGCGCAGGCGGCTAATTAAGTCAGATGTGAAAGGCCACGGCTCAACCGTGGAGTTGCATTTGAAACTGGTTAGCTTGAGTGCAGGAGAGGTGAGTGGAAT<br>TCCCGGTGTAGCGGTGGAATGCGTAGATATCGGGAGGAACACCAGTGGCGAAGGCGGCTCACTGGCCTGCAACTGACGCTGAGGCGCGAAAGCGTGGGG |
| P-asv6102 | GCTAGCGTTGTTCGGAATTACTGGGCGTAAAGCGCGCGTAGGCGGATCGTTAAGTCGGGGGTGAAATCCCGGGGCTCAACCCCGGAACTGCCTTCGATACTGGCGATCTAGAGGTCGAGAGAGGTGAGTGGAA<br>TTCCGAGTGTAGAGGTGAAATTCGTAGATATTCGGAGGAACACCAGTGGCGAAGGCGGCTCACTGGCTCGATACTGACGCTGAGGTGCGAAAGTGTGGGG |
| P-asv6103 | GCAAGCGTTGTTCGGAATCACTGGGCGTAAAGAATGCGTAGGCGGCTAGACAAGTCAGGAGTGAAATCCCAGGGCTCAACCCTGGAACTGCTTTTGAAACTGTCTTGCTAGAGTCCCGGAGGGGTTGGCGGAAT<br>TCCGAGTGTAGAGGTGAAATTCGCAGATATTCGGAAGAACACCAGTGGCGAAGGCGACTCACTGGCCCGTTACTGACGCTGAGGCGCGAAAGCGTGGGG |
| P-asv6104 | ACGAGTGATGGCCGATATTATTGGGCCTAAAGCGTTCGTAGCCGGCCAGTCAAGTCTGTTGGGAAATCGACGCGCTCAACGCGTCGGCGTCCAGCGGAAACTGATTGGCTTGGGGCCGGAAGATCTGAGGGGT<br>ACGTCCGGGGTAGGAGTGAAATCCCGTAATCCTGGACGGACCACCGGTGGCGAAAGCGCCTCAGAAAGACGGACCCGACGGTGAGGAACGAAAGCTAGGGT |
| P-asv6105 | CCAAGCGTTGTTCGGATTCACTGGGCGTAAAGGGTGCGTAGGCGGCGCGGAAAGTCGGATGTGAAAGCCCCGGGCTCAACCCGGGAACTGCATCCGATACTCCCGTGCTAGAGGATTGGAGAGGAGTCTGGAA<br>TTCACGGTGTAGCAGTGAAATGCGTAGGTATCGTGAGGAACACTCGTGGCGAAGGCGAGACTCTGGACAATTCCTGACGCTGAGGCACGAAGGCCGGGGG |
| P-asv6106 | CCAAGCGTTATCCGGATTTATTGGGTTTAAAGGGTCCGTAGGCGGGTCATTAAGTCAGTGGTGAAATCCTACAGCTCAACTGTAGAACTGCCATTGAAACTGATGACCTTGAATATAATTGAAGTGGGCGGAATGT<br>GTCATGTAGCGGTGAAATGCTTAGATATGACACAGAACACCGATTGCGAAGGCAGCTCACTAAGTTATTATTGACGCTGAGGGACGAAAGCGTGGGT |
| P-asv6107 | GCAAGCGTTAACCGGATTTACTGGGCGTAAAGGGTGCGTGGGCGGATTGGTAAGTCAGATGTGAAATTTCGGCACTCAATACCGAAGCTGCATTTGAAACTGCCTATCTAGAGGGTAGATGGAGAAAGCGGAATT<br>CCATGTGTAGCGGTGAAATGCGTAGATATATGGAAGAACACCTGTGGCGTAAGCGGCTTTCTAATTTATTCCTGACGCTGAGGCACGAGAGTGTGGGG |
| P-asv6108 | GCTAGCGTTGTTCGGAATTACTGGGCGTAAAGGGCGCGTAGGCGGACCTGTAAGTCAGGGGTGAAATCCCGGGGCTCAACCTCGGAACTGCCCTTGAGACTGTTGGTCTAGAGTCTGTGAGAGGATGACGGAA<br>TACCCAGTGTAGAGGTGAAATTCGTAGATATTGGGTAGAACACCGGTGGCGAAGGCGGTCATCTGGCGCAGCACTGACGCTGAGGCGCGAAAGCGTGGGG |
| P-asv6109 | CCAAGCGTTATCCGGATTTATTGGGTTTAAAGGGTGCGTAGGCGGCTTTTTAAGTCAGGGGTGAAAACTTGCGGCTCAACCGTAAGCTTGCCTTTGATACTGAGGAGCTTGAATACACTTGAGGTAGGCGGAATG<br>TGACAAGTAGCGGTGAAATGCATAGATATGTCACAGAACACCAATTGCGAAGGCAGCTTACTAAAGTGTGATTGACGCTGAGGCACGAAAGCGTGGGG |
| P-asv6110 | GCAAGCGTTATCCGGAATCATTGGGCGTAAAGCGCATGTAGGCGGCCATGTAAGTCGGTTGTGAAATCTCGGGGCTCAACCCCGAAACTGCATCCGATACTGCATGGCTTGAGGAAGGTAGAGGAGAGTGGAAT<br>TCCCGGTGTAGCGGTGGAATGCGCAGATATCGGGAGGAACACCGGCGGCGAAGGCGGCTCTCTGGGCCTCTCCTGACGCTGAGATGCGAAAGTGTGGGT |
| P-asv6111 | GCAAGCGTTAATCGGAATTACTGGGCGTAAAGGGTGCGTAGGTGGTTTGGTAAGTTATGTGTGAAATCCCCGGGCTTAACCTGGGAGGGTCAGATAATACTGCCGAACTAGAGTATGAGAGAGGGTAGTGGAATT<br>TCCGGTGTAGCGGTGAAATGCGTAGAGATCGGAAGGAACACCAGTGGCGAAGGCGGCTGCCTGGCTTAATACTGACACTGAGGCACGAAAGCGTGGGG |
| P-asv6112 | CCAAGCGTTATCCGGAATCATTGGGTTTAAAGGGTCCGTAGGTGGTCCATTAAGTCAGAGGTGAAAGCCTATCGCTCAACGATAGAACTGCCTTTGATACTGATAGACTTGAGTTATTGTGAAGTAGTTAGAATATG<br>TAGTGTAGCGGTGAAATGCATAGATATTACATAGAATACCGATTGCGAAGGCAGATTACTAACAATATACTGACACTGAGGGACGAAAGCGTGGGT |
| P-asv6113 | GCGAGCGTTAATCGGAATTACTGGGCGTAAAGCGTGCGCAGGCGGCTTTTTAAGCCAGATGTGAAATCCCCGGGCTTAACCTGGGAACTGCATTTGGAACTGGAAAGCTAGAGTGTGTCAGAGGGGGGTGGAAT<br>TCCGCGTGTAGCAGTGAAATGCGTAGATATGCGGAGGAACACCGATGGCGAAGGCAGCCCCCTGGGATAACACTGACGCTCATGCACGAAAGCGTGGGG |
| P-asv6114 | GCAAGCGTTACTCGGAATCACTGGGCGTAAAGGATGTGTAGGCTGTACTATAAGTCAGGAGTGAAATCCAACGGCTCAACCGTTGAACTGCTTTTGAAACTGTTGTACTAGAATATGGGAGAGGTAGATGGAATTG<br>GTGGTGTAGGGGTAAAATCCGTAGATATCACCAGGAATACCGATTGCGAAGGCGATCTACTGGAACATTATTGACGCTGAGACATGAAAGCGTGGGG |
| P-asv6115 | GCGAGCGTTGTCCGGATTTATTGGGCGTAAAGAGCTCGTAGGCGGCTTAGTAAGTCGGGTGTGAAAACTCCAGGCTCAACCTGGAGACGCCACCTGATACTGCTATGGCTAGAGTCCGGTAGAGGAGTGTGGAA<br>TTCCCAGTGTAGCGGTGAAATGCATAGATATTGGGAGGAACACCAATGGCGAAGGCAGCACTCTGGGCCGGTACTGACGCTGAGGAGCGAGAGCGTGGGG |
| P-asv6116 | GCGAGCGTTGTCCGGATTTATTGGGCGTAAAGAGCTCGTAGGCGGTTTCGTATGTCGGGTGTGAAAACTCCAGGCTCAACCTGGAGACGCCACCCGAAACTGCGATGACTAGAGTCCGGTAGGGGAGTGTGGA<br>ATTCCTGGTGTAGCGGTGAAATGCGCAGATATCAGGAGGAACACCAACAGCGAAGGCAGCACTCTGGGCCGGAACTGACGCTGAGGAGCGAAAGCGTGGGT |
| P-asv6117 | GCGAGCGTTAATCGGAATTACTGGGCGTAAAGCGTGCGCAGGCGGTTTTGAAAGTCAGATGTGAAATCCCCGAGCTCAACTTGGGAACTGCGTTTGAAACTCCAAAGCTAGAGTATAGGAGAGGGGGGTAGAAT<br>TCCACGTGTAGCAGTGAAATGCTTAGATATTCGCAAGAACACCAGTGGCGAAGGCGGCTTACTGGCCCGATACTGACACTGAGGCACGAAAGCGTGGGG |
| P-asv6118 | GCTAGCGTTATTCGGAATTACTGGGCGTAAAGCGCACGTAGGCGGCTTTGTAAGTTAGAGGTGAAAGCCTGGAGCTCAACTCCAGAATTGCCTTTAAGACTGCATCGCTTGAATCCAGGAGAGGTGAGTGGAATT<br>CCTGGTGTAGGGGTGGAATGCGCAGATATCAGGAGGAACACCGATGGCGAAGGCAGGTCTCTGGGCCGTTACTGACGCTGAGGAGCGAAAGCATGGGG |
| P-asv6119 | CCAAGCGTTAATCGGAATCACTGGGCTTAAAGCGTGCGCAGGCGGACCCGTCAGTGTCTTGTGAAATCCCACTGCTCAACGGTGGAATTGCAGGGCATACTGCGGGTCTTGAGGTCGGTAGAGGTTGCCGGAAC<br>TGTAGGTGTAGCGGTGAAATGCGTAGATATCTACAGGAACACCCAAGGCGAAGGCAGGCAACTGGGCCGATTCTGACGCTCAGGCACGAAAGCGTGGGG |
| P-asv6120 | GCAAGCGTTAATCGGAATTACTGGGCGTAAAGCGTGCGCAGGCGGTTATACAAGACAGATGTGAAATCCCCGGGCTCAACCTGGGAATTGCATTTGTGACTGTATAGCTAGAGTGTGTCAGAGGGGGGTAGAATT<br>CCACGTGTAGCAGTGAAATGCGTAGATATTCGGAAGAACACCAGTGGCGAAGGCGGCTCACTGGACTGGTATTGACGCTGAGGTGCGAAAGCGTGAGG |
| P-asv6121 | GCTAGCGTTGTTCGGAATTACTGGGCGTAAAGCGCGCGTAGGCGGATAGATTAGTCAGATGTGAAATCCCAAGGCTCAACCTTGGAACTGCATTTGAAACTGTCTATCTAGAGACATGGAGAGGTTAGTGGAATA<br>CCGAGTGGAGAGGTGAAATTCGTAGATATTCGGTGGAACACCAGTGGCGAAGGCGACTAACTGGACATGTACTGACGCTGAGGCGCGAAAGTGTGGGG |
| P-asv6122 | GCGAACGTTATTCGGAATCACTGGGCTTAAAGGGTGCGTAGGCGGCCTGTTAAGTAGGGTGTGAAATTTGTCGGCTCAACCGGCAAACGGCATTCTAAACTGGCAGGCTTGAGTGAGTCAGGGGTGTGCGGAAC<br>TTCCGGTGGAGCGGTGAAATGTGTTGATATCGGAAGGAACGCCGGTGGCGAAAGCGGCACACTGGGGCTTTTCTGACGCTGAGGCACGAAAGCCGGGGG |
| P-asv6123 | GCTAGCGTTGCTCGGAATTACTGGGCGTAAAGGGCGTGTAGGCGGCGTTTTAAGTCGGGGGTGAAAGCCTGGAGCTCAACTCCAGAATTGCCTTTAAGACTGCATCGCTTGAATCCAGGAGAGGTGAGTGGAAT<br>TCCGAGTGTAGAGGTGAAATTCGTAGATATTCGGAAGAACACCAGTGGCGAAGGCGGCTCACTGGACTGGTATTGACGCTGAGGTGCGAAAGCGTGGGG |
| P-asv6124 | GCTAGCGTTGTTCGGAATTACTGGGCGTAAAGCGCACGTAGGCGGCTTTGTAAGTCAGAGGTGAAAGCCTGGAGCTCAACTCCAGAACTGCCTTTGAGACTGCATCGCTTGAATCCAGGAGAGGTGAGTGGAAT<br>TCCTGGTGTAGGGGTGGAATGCGCAGATATCAGGAGGAACACCGATGGCGAAGGCAGGTCTCTGGGCCGTTACTGACGCTGAGGAGCGAAAGCATGGGG |
| P-asv6125 | GCAAGCGTTGTCCGGATTTACTGGGCGTAAAGGGAGCGTAGGTGGATATTTAAGTGGGATGTGAAATACCCGGGCTTAACCTGGGTGCTGCATTCCAAACTGGATATCTAGAGTGCAGGAGAGGAAAGGAGAAT<br>TCCTAGTGTAGCGGTGAAATGCGTAGAGATTAGGAAGAATACCAGTGGCGAAGGCGCCTTTCTGGACTGTAACTGACACTGAGGCTCGAAAGCGTGGGG |
| P-asv6126 | GCAAGCGTTGTTCGGAATTATTGGGCGTAAAGGGCATGCAGGCGGTCGTGCAAGCCCGGCGTGAAAGACGGGGGCTCAACTCCCGAAGTGCGCTGGGAACTGCAGGGCTTGAGTTTCTGAAGGGCAGCCGGA<br>ATTCCACGTGTAGGGGTGAAATCTGTAGATATGTGGAAGAACACCAATGGCGAAGGCAGGCTGCCGGCAGAAGACTGACGCTGAGGTGCGAAGGTGCGGGG |
| P-asv6127 | GCAAGCGTTGTCCGGAATCATTGGGCGTAAAGCGCGCGTAGGCGGCCTTGTAAGTCCGTTGTGAAAGTCAAAGGCTCAACCTTTGAAAGCCGATGGATACTGCAGGGCTAGAGTCCGGAAGAGGCGAGTGGAAT<br>TCCTGGTGTAGCGGTGGAATGCGCAGATATCAGGAGGAACACCCATAGCGAAGGCAGCTCGCTGGGACGGTACTGACGCTAAGGCGCGAAAGCGTGGGG |

| | |
|---|---|
| P-asv6128 | GCAAGCGTTGTCCGGATTTATTGGGTTTAAAGGGTGCGTACGCGGCCTGTTAAGTCAGAGGTGAAATCCCGGGGCTCAACCCCGGAACTGCCTTTGAGACCTCAGTACTAGAGTGTGAGAGGGGATAGTGGAATACCCAGTGTAGCGGTGAAATGCGTAGAGATTGGGTGGAACACCGGTGGCGAAGGCGGCTATCTGGCTCACAACTGACGATCAGGCACGAAAGCGTGGGG |
| P-asv6129 | GCGAGCGTTATCCGGATTCATTGGGTTTAAAGGGTGCGTAGGCGGTTTCTTAAGTCAGTGGTGAAATCCTGCGGCTCAACCGTAGAACTGCCATTGATACTGAGGAACTTGAATTTAGTTGAGGTAGGCGGAATGAGTAGTGTAGCGGTGAAATGCTCAGATATTACTCAGAACACCGATTGCGAAGGCAGCTCACCAAGCTACGATTGACGCTGATGCACGAAAGCGTGGGG |
| P-asv6130 | GCAAGCGTTGTCCGGAATTATTGGGCGTAAAGCGCGCGCAGGCGGCCTGTTAAGTCTGGTGTGAAAGACCACGGCTCAACCGAGGGGTGGCATCAGAAACTGGCGGGCTTGAGTGCAGGAGAGGAGAGTGGAATTCCCGGTGTAGCGGTGGAATGCGTAGAGATCGGGAGGAACACCAGTGGCGAAGGCGACTCTCTGGCCTGTAACTGACGCTGAGGCGCGAAAGCGTGGGG |
| P-asv6131 | CCGAGTGGTCGGGACGATTATTGGGCCTAAAGCATCCGTAGCCGGTACTGCAAGTCTTCCGTTAAATCCACCTGCTTAACAGTTGGGCTGCGGGGGATACTACAGCGCTAGGAGGCGGGAGAGGCAAGCGGTACTCAATGGGTAGGGGTAAGATCCGCTGATCCATTGAAGACCACCAGTGGCGAAGGCGGCTTGCCAGAACGCGCTCGACGGTGAGGGATGAAAGCTGGGGG |
| P-asv6132 | GCAAGCGTCAATCGGAATCACTGGGCGTAAAGGGTGCGTAGGCGGTGCATTAGGTCTGTCGTGAAATCCCCGGGCTCAACCAATGTAAGCCTTTGAAACTGTTGGGCTTGAGTGCAGGAGAGGTAAGTGGAATTCCTAGTGTAGAGGTGAAATTCGTAGATATTAGGAGGAACACCAGTGGCGAAGGCGACTTACTGGACTGTAACTGACGCTGAGGCACGAAAGCGTGGGG |
| P-asv6133 | GCTAGCGTTGCTCGGAATCACTGGGCGTAAAGGGCGCGTAGGCGGCGTTTTAAGTCGGGGGTGAAAGCCTGTGGCTCAACCACAGAATGGCCTTCGATACTGGGACGCTTGAGTATGGTAGAGGTTGGTGGAACTGCGAGTGTAAAGGTGAAATTCGTAGATATTCGGAAGAACACCAGTGGCGAAGGCGGTTCACTGGACTGGTATTGACGCTGAAGTGCGAAAGCGTGGGG |
| P-asv6134 | TCAAGTGATGTCCCATATTATTGGGCCTAAAGCGTCCGTAGCTGGCCGACCAAGTCTATCGGGAAATCCACCCGCTCAACGGGTGGGCGTCCGGTAGAAACTGGCCGGCTTGGAACCGGAAGGCTCAGAGAGTACGTCCGGGGTAGGAGTGAAATCCCGTAATCCCGGACGGACTACCGATGGCGAAAGCACTCTGAGAAGACGGATTCGACAGTGAGGGACGAAAGCTCGGGT |
| P-asv6135 | GCTGGCGTTGTTCGGAATTACTGGGCATAAAGCGCACGTAGGCGGCTTTTTAAGTCAGGGGTGAAATCCCGGGGCTCAACCCCGGAACTGCCCTTGAAACTGGGAAGCTAGAATCTTGGAGAGGCGAGTGGAATTCCGAGTGTAGAGGTGAAATTCGTAGATATTCGGAAGAACACCAGTGGCGAAGGCGACTCGCTGGACAAGTATTGACGCTGAGGTGCGAAAGCGTGGGG |
| P-asv6136 | ACGAGTGATGGCCGATTTTATTGGGCCTAAAGCGTCCGTAGCTGGCCGACCAAGTCCGTCGGGAAATCCACCCGCCCAACGGGTGGGCGTCCGGTAGAAACTGGCCGGCTTGGAACCGGAAGGCTCAGAGAGTACGTCCGGGGTAGGAGTGAAATCCCGTAATCCCGGACGGACTACCGATGGCGAAAGCACTCTGAGAAGACGGCTTCGACAGTGAGGGACGAAAGCTCGGGT |
| P-asv6137 | GCGAGCGTTAATCGGAATTACTGGGCGTAAAGGGTGCGTAGGTGGTTATTTAAGTCAGATGTGAAATCCCTAGGCTTAACCTAGGAATTGCGTTTGAAACTGAATAGCTAGAGTGCAGGAGAGGGAAGTGGAATTTCCGGTGTAGCGGTGAAATGCGTAGATATCGGAAGGAACATCAGTGGCGAAGGCGACTTCCTGGACTGTTACTGACACTGAGGCACGAAAGCGTGGGG |
| P-asv6138 | GCAAGCGTTGTTCGGAATTATTGGGCGTAAAGCGCGCGCAGGCGGGTTGCTAAGTCTGATGTGAAATCCCTCGGCTCAACCGAGGACGTGCATTGGATACTGGCAATCTTGAATCCCGGAGAGGGTGGCGGAATTCCCAGTGTAGAGGTGAAATTCGTAGATATTGGGAGGAACACCAGTGGCGAAGGCGGCCACCTGTACGGTGATTTACGCTGAGGCGCGAAAGCGTGGGT |
| P-asv6139 | GCAAGCGTTGCTCGGAATCACTGGGCGTAAAGCGAACGCAGGCGGACCATTAAGTCGGGTGTGAAATCCCGGGGCTCAACCCCGGAACTGCATTCGAAACTGCTGGTCTCGAGTATGGGAGAGGGAAGCGGAATTCCTGGTGTAGAGGTGAAATTCGTAGATATCAGGAGGAACACCTGTGGCGAAAGCGGCTTCCTGGCCCAATACTGACGCTCATGTTCGAAAGCGTGGGG |
| P-asv6140 | GCAAGCGTTGTCCGGAATTACTGGGCGTAAAGAGTTCGTAGGCGGGAATGTAAGTCAGAAGTGAAAGGCAATGGCTCAACCATTGTAAGCTTTTGAAACTGCATTTCTTGAGTATCGGAGAGGAAAGTGGAATTCCTAGTGTAGCGGTGAAATGGGTAGATATTAGGAGGAACACCAGTGGCGAAGGCGACTTTCTGGACGAAAACTGACGCTGAGGAACGAAAGCGTGGGG |
| P-asv6141 | GCAAACGTTGTTCGGAATTACTGGGCGTAAAGCGCGTGTAGGCGGTCTTGCAGGTCGGATGTGAAATCCCTCGGCTCAACCAAGGAAGTGCATTCGAAACCGCAAGACTTGAGTATCGGAGAGGAAGGCGGAATTGTCGGTGTAGAGGTGAAATTCGTAGATATCGACAGGAACACCTGTGGCGAAAGCGGCCTTCTGGACGAATACTGACGCTGAGACGCGAAAGCGTGGGG |
| P-asv6142 | GCAAGCGTTGTTCGGAATTATTGGGCGTAAAGCGCGCGCAGGCGGGTCGCAAAGTCCGATGTGAAATCCCTCGGCTCAACCGAGGACGTGCATTGGATACTGGCGACCTTGAGTCCTGGAGAGGGGAGCGGAATTTCCAGTGTAGAGGTGAAATTCGTAGATATTGGAAGGAACACCAGTGGCGAAGGCGGCTCCCTGGACAGTGACTGACGCTGAGGCGCGAAAGCGTGGGT |
| P-asv6143 | GCTAGCGTTGTTCGGAATTACTGGGCGTAAAGGGCGCGTAGGCGGTCTTGTAAGTCAGGCGTGAAAGGCCCGGGCTCAACCTGGGGACTGCGCTTGATACTGCGAGGCTTGAATCCGGGAGAGGATGGGGGAATTCCCAGTGTAGAGGTGAAATTCGTAGATATTGGGAAGAACACCGGTGGCGAAGGCGGCCATCTGGACCGGTATTGACGCTGAGGCGCGAAAGCGTGGGG |
| P-asv6144 | GCGAGCGTTGTTCGGAATCACTGGGCGTAAAGCGTATGTAGGCGGTCATGAAAGTCAGAAGTGAAATCCCAGGGCTTAACCCTGGAATTGCTTTTGAAACTTCATGACTAGAGTTCTGGAGAGGCAAGGGGAATTCCGAGTGTAGCAGTGAAATGCGCAGATATTCGGAGGAACACCAGTGGCGTAGGCGCCTTGCTGGACAGATACTGACGCTGAGATACGAAAGCGTGGGG |
| P-asv6145 | CCAAGCGTTGTCCGGATTTATTGGGCGTAAAGAGCTCGTAGGCGGCTCAGTAAGTCGGATGTGAAAACTCCAGGCTCAACCTGGAGACGCCATTCGATACTGCTGTGGCTTGAGTCTGGTAGGGGAGCACGGAATTCCTGGTGTAGCGGTGAAATGCGCAGATATCAGGAGGAACACCGGTGGCGAAGGCGGTGCTCTGGGCCAGTACTGACGCTGAGGAGCGAAAGCGTGGGG |
| P-asv6146 | GCAAGCGTTAATCGGAATTACTGGGCGTAAAGCGTTCGTAGGCGGCCTATCAAGTCTGATGTTAAAGCCCGGAGCTCAACTCCGGTTCAGCATTGGATACTGATAAGCTAGAGTACAGGAGAGGCAAGGGGAATTCCCAGTGTAGCGGTGAAATGCGTAGATATTGGGAGGAACACCAGTGGCGTAGGCGCCTTGCTGGACTGTAACTGACGCTGAGGGACGAAAGCCAGGGG |
| P-asv6147 | GCAAGCGTTATCCGGATTCATTGGGCGTAAAGCGCTCGTAGGCGGTCTGTTAGGTCGGGAGTTAAATCCGGGGGCTCAACCCCCGTTCGCTCCCGATACCGGCAGACTTGAGTTTGGTAGGGGAAGGTGGAATTCCTAGTGTAGCGGTGGAATGCGCAGATATTAGGAAGAACACCAGTGGCGAAGGCGGCCTTCTGGGCCATAACTGACGCTGAGGAGCGAAAGCTAGGGG |
| P-asv6148 | CCAAGTGGTAGCCACTTTTATTGGGCTTAAAGTGTTCGTAGCCGGCCGTGTAAGTCTGCTGTGAAATCCACGCGCTCAACGTGTGGAAGTGCAGCGGATACTGCATGGCTAGGAACCGGATGGCGTCGAGGCTATTCCAGGGGTAGGAGTGAAATCCAAAAATCCCTGGAGGAGCACCTGTGGCGAAAGCGCTCGACGAATACGGATTCGACGGTGAGGAACGAAAGCCAGGGG |
| P-asv6149 | GCAAGCGTTGTTCGGAATCACTGGGCGTAAAGGGCGCGCAGGCGGTTTGGTAAGTCAGATGTGAAAGCCCACGGCTTAACCGTGGAAGTGCATTTGAAACTGTCAGACTTGAGTACCAGAGGGGAAAGTGGAATTCCCGGTGTAGAGGTGAAATTCGTAGATATCGGGAGGAATACCGGTGGCGAAGGCGACTTTCTGGCTGAGTACTGACGCTGAGGCGCGAAAGCGTGGGG |
| P-asv6150 | GCTAGCGTTGTTCGGAATTACTGGGCGTAAAGCGCGCGCAGGCGGTTCAATAAGTTAGGGGTGAAATCCCCGGGCTCAACCCGGGAATGGCCTTTAAGACTGTCGAGCTAGAGGACGAGAGAGGTGAGTGGAATTCCCAGTGTAGAGGTGAAATTCGTAGATATTGGGAAGAACACCGGTGGCGAAGGCGGCTCACTGGCTCGTTTCTGACGCTGAGGCGCGAAAGCGTGGGG |
| P-asv6151 | GCAAGCGTTGTCCGGAATTATTGGGCGTAAAGAGCTCGTAGGCGGTCTGTCGCGTCTGCTGTGAAAACTCAAGGCTCAACCTTGGGCTTGCAGTGGGTACGGGCAGACTAGAGTGCGGTAGGGGTGACTGGAATTCCTGGTGTAGCGGTGGAATGCGCAGATATCAGGAGGAACACCGATGGCGAAGGCAGGTCACTGGGCCGCAACTGACGCTGAGGAGCGAAAGCATGGGG |
| P-asv6152 | GCAAGCGTTAATCGGAATCACTGGGCGTAAAGCGTGCGTAGGCTGCTTGATAAGTCAGACGTGAAATCCCTCGGCTCAACCGGGGAACTGCGTTTGATACTGTCGAGCTAGAGTCCTGGAGAGGGTGGCGGAATTCCTGGTGTAGGAGTGAAATCCGTAGATATCAGGAGGAACACCGGTGGCGAAGGCGGCCACCTGGACAGGTACTGACGCTGAGGTACGAAAGCGTGGGG |
| P-asv6153 | GCAAGCGTTAATCGGAATTACTGGGCGTAAAGCGCGCGTAGGTGGTTTGATAAGTTGAATGTGAAAGCCCCGGGCTCAACCCTGGAACTGCGTCTGGAACTGGTAATCTAGAGTTATGGAGGGGAGAGCGGAATTTCACATGTAGGGGTAAAATCCGTAGAGATGTGAAGGAACACCAGTGCCGAAGGGGGCTCTCTGGCCATTAACTGACGCTGAGGCGCGAAAGCGTGGGT |
| P-asv6154 | GCAAGCGTTGTTCGGAATCATTGGGCGTAAAGGGCTCGTAGGCGGACTAGTAAGTCTGGCGTGAAATCTCCACCGCTCACCGGGGAGAGGTCGCTGGATACTGCTAGTCTTGAGTTCGGTAGAGGGTCGTGGAATTTCGGGTGTAGCGGTGGAATGCGTAGATATCCGAAAGAACACCGGTGGCGAAGGCGGCGACCTGGGCCTGAACTGACGCTGAGGAGCGAAAGCTAGGGG |
| P-asv6155 | CCGAGCGTTATCCGGATTTATTGGGTTTAAAGGGAGCGTAGGCGGACGCTTAAGTCAGTTGTGAAAGTTTGCGGCTCAACCGTAAAATTGCAGTTGATACTGGGTGTCTTGAGTGCAGCAGAGGTAGGCGGAATTCGTGGTGTAGCGGTGAAATGCTTAGATATCACGAAGAACTCCGATTGCGAAGGCAGCTTACTGGACTGTAACTGACGCTGATGCTCGAAAGTGTGGGT |
| P-asv6156 | GCAAGCGTTATCCGGAATCACTGGGCGTAAAGGGCGCGTAGGCGGTCTGTTAAGTCCGATGTTAAAGCCCGGGGCTCAACCCCGGCACGGCGTTGGATACTGGCAGGCTTGACGGCTGGAGAGGCGAGTGGAATTACCAGTGTAGCGGTGGAATGCGTAGATACTGGTAGGAACACCTATGGCGAAGGCAGCTCGCTGGACAGCACGTGACGCTGAGGCGCGAAAGTGTGGGG |
| P-asv6157 | GCAAGCGTTGTTCGGATTCATTGGGTGTAAAGGGCAGGTAGGTGGCTTAGCAAGTCTGGTGTGAAATCCCAGGGCTCAACCCTGGAACTGCACTGGAAACTACTGAGCTTGAGTACCGGAGAGGTAAGGGGAATTCTCGGTGTAAAGGTGAAATCTGTAGATATCGAGAGGAATACCAGTGGCGAAGGCGCCTTACTGGCCGGTTACTGACACTGAGCTGCGAAAGCAAGGGG |
| P-asv6158 | CCGAGCGTTAATCGGAATTACTGGGCGTAAAGCGCGCGTAGGTGGTTCAGTCAGTCAGATGTGAAAGCCCCGGGCTCAACCTGGGAACTGCACCTGATACTGCTGAGCTAGAGTACGGTAGAGGGTAGTGGAATTTCCGGTGTAGCGGTGAAATGCGTAGAGATCGGAAGGAACACCAGTGGCGAAGGCGACTACCTGGACCGATACTGACACTGAGGTGCGAAAGCGTGGGT |
| P-asv6159 | GCAAACGTTGTTCGGATTCACTGGGCTTAAAGGGCGCGTAGGCGGCTCGCCAAGTCAGGGGTGAAATCCCCCGGCTCAACCGGGGAACAGCTTCTGATACTGGCGGGCTCGAGGGGGGTAGGGGCAGGCGGAACTTCCGGTGGAGCGGTGAAATGCGTAGATATCGGAAGGAACGCCGGTGGCGAAAGCGGCCTGCTGGACCCCTTCTGACGCTGAGGCGCGAAAGCCAGGGG |
| P-asv6160 | GCAAGCGTTACTCGGAATTATTGGGCGTAAAGCGTGCGTAGGCGGCTTTATAAGTTGGAAGTGAAAGCCTTTGGCTCAACCAAAGAATTGCTTACAAAACTGTAAAGCTAGAGTATTAGAGAGGATAGAAGAATTTCTGATGTAGGGGTGAAATCCGTAGATATCAGAAAGAATATCGAAGGCGAAAGCATCTGTCTGGCTAAATACTGACGCTGTTGCACGAAAGCGTGGGG |
| P-asv6161 | GCAAGCGTTGTCCGGAATTATTGGGCGTAAAGAGCTCGTAGGCGGCTTGTCGCGTCGGTTGTGAAAGCCCGGGGCTTAACTCCGGGTCTGCAGTCGATACGGGCAGGCTAGAGTTCGGTAGGGGAGATCGGAATTCCTGGTGTAGCGGTGAAATGCGCAGATATCAGGAGGAACACCGGTGGCGAAGGCGGATCTCTGGGCCGATACTGACGCTGAGGAGCGAAAGCGTGGGG |
| P-asv6162 | GCAAGCGTTATCCGGATTCATTGGGTTTAAAGGGAGCGTAGGTGGACTTGTAAGTCAGTGGTGAAATCTCTCAGCTTAACTGAGAAACTGCCATTGATACTGCAGGTCTTGAGTACAGATGATGTGGGCGGAATATGACATGTAGTGGTGAAATACTTAGATATGTCATAGAACACCGATTGCGAAGGCAGCTCACAAAACTGTAACTGACACTGAGGCTCGAAAGAGCGGGG |
| P-asv6163 | GCAAGCGTTGTCCGGAATCACTGGGCGTAAAGCGCACGCAGGCGGCACGGTAAGTGGGGTGTGACAGCGGACGGCTTAACCGTCCGAGGCCATCTCAGACTGCCGCGCTTGAGCCGTGCAGAGGTCAGCGGAATTGCCGGTGTAGGAGTGACATCCGTAGAGACCGGCAGGAACACCGAAGGGGAAGCCAGCTGACTGGGCACGAGCTGACGCTCAGGTGCGATAGCGTGGGG |
| P-asv6164 | GCTAGCGTTATTCGGAATTACTGGGCGTAAAGCGCACGTAGGCGGATATGTCAGTCAGGAGTGAAATCCCGGAGCTCAACTTCGGAACTGCCTTTGATACTGAAGATCTTGAGTTCGGGAGAGGTGAGTGGAACTGCGAGTGTAGAGGTGAAATTCGTAGATATTCGCAAGAACACCAGTGGCGAAGGCGGCTCACTGGCCCGATACTGACGCTGAGGCACGAAAGCGTGGGG |
| P-asv6165 | GCAAACGTTGCTCAGAATCACTGGGCGTAAAGGGCGTGTAGGCGGGTAGGCAAGTCGGGCGTGAAATCCCTGAGCTCAACTCAGGAACTGCACTCGAAACTGCTTATCTTGAGTGCCGGAGAGGAAAGCGGAATTCCTGGTGTAGAGGTGAAATTCGTAGATATCAGGAGGAACACCAGTGGCGAAGGCGGCTTTCTGGACGGTAACTGACGCTGAGACGCGAAAGCGTGGGG |
| P-asv6166 | GCAAGCGTTATCCAGATTCACTGGGTTTAAAGGGTGCGTAGGCGGGTTGGTAAGTCCGTGGTGAAATCTCCAAGCTTAACTTGGAAACTGCCGTGGATACTATCAATCTTGAATGTCGTGGAGGTAAGCGGAATATGTCATGTAGCGGTGGAATGCTTAGAGATGACATAGAACACCAATTGCGTAGGCAGCTTACTACACGAATATTGACGCTGAGGCACGAAAGCGTGGGG |

| | |
|---|---|
| P-asv6167 | TCAAGCGTTGTTCGGAATCACTGGGCGTAAAGGGTGCGTAGGTGGCGTGGTAAGTCAGATGTGAAAGCCCGGGGCTCAACCCCGGAATTGCATCCGATACTGCCGCGCTAGAGTACTGAAGAGGTGACTAGAAT<br>TCTAGGTGTAGCAGTGAAATGCGTAGATATCTAGAGGAATACCAAAGGCGAAGGCAGGTCACTGGGCAGTTACTGACACTGAGGCACGAAGGCCAGGGT |
| P-asv6168 | GCAAGCGTTATCCGGAATCACTGGGTTTAAAGGGTGCGTAGGCGGTTTGATAAGTCAGAGGTGAAAGCTAGCAGCTTAACTGTTAAATTGCCTTTGATACTGTCAGACTTGAATCAAGTTGAGGTAGGCGGAATGT<br>GACATGTAGCGGTGAAATGCATAGATATGTCATAGAACACCAATTGCGTAGGCAGCTTGCTGGGCTTGAATTGACGCTGAGGCACGAAAGCGTGGGG |
| P-asv6169 | CCAAGCGTTGTTCGGAATCACTGGGCGTAAAGGGAGCGTAGGCGGCGCGGTAAGTCAGATGTGAAATCCCGGGGCTCAACCCCGGAACTGCATCCGATACTGCCGTGCTTGAGCAATGGAGAGGTAGCTGGAA<br>TTATCGGTGTAGCAGTGAAATGCGTAGATATCGATAGGAACACTCGTGGCGAAAGCGAGCTACTGGACATTTGCTGACGCTGAGGCTCGAAGGCTAGGGG |
| P-asv6170 | GCAAGCGTTAATCGGAATCACTGGGCGTAAAGCGCACGTAGGCGGCGAGGTCAGTCAGATGTGAAAGCCCCGGGCTTAACCTGGGAACTGCATTTGATACTGCCTGGCTAGAGTTTGATAGAGGGGGGTGGAAT<br>TCCAGGTGTAGCGGTGAAATGCGTAGATATCTGGAGGAACACCAGTGGCGAAGGCGGCCCCCTGGATCAAAACTGACGCTGAGGTGCGAAAGCGTGGGG |
| P-asv6171 | CCAAGCGTTGTTCGGATTCACTGGGCGTAAAGGGTGCGTAGGTGGTCGGGTAAGTCTGATGTGAAATCTCCGAGCTCAACTCGGAAACTGCATCGGATACTATTCGGCTCGAGGATTGGAGGGGGGACTGGAAT<br>ACTTGGTGTAGCAGTGAAATGCGTAGATATCAAGTGGAACACCAGTGGCGAAGGCGAGTCCCTGGACAATTCCTGACACTGAGGCACGAAAGCTAGGGG |
| P-asv6172 | CCAAGCGTTGTTCGGAATCACTGGGCGTAAAGCGTGTGTAGGCGGCGCGGAAAGTCAAATGTGAAAGCCCGAGGCTCAACCTCGGAACTGCATTCGATACTCCCGTGCTGGAGTCATGGAGAGGTAAGTGGAAT<br>TCCTGGTGTAGCAGTGAAATGCGTGGATATCAGGAGGAACATCAATGGCGAAGGCAACTTACTGGACATGTACTGACGCTGAGACACGAAGGCTAGGGG |
| P-asv6173 | GCTAGCGTTGTTCGGAATTACTGGGCGTAAAGAGAGCGTAGGCGGCTCTATAAGTTGGATGTGAAAGCCCCGAGCTTAACTTGGGAATAGCATTCAATACTGTAGGGCTAGAGAGTGATAGAGGATGATGGAATT<br>CCTAGTGTAGAGGTGAAATTCGTAGATATTAGGAGGAACACCAGTGGCGAAGGCGGTCATCTGGGTCACAACTGACGCTAAGGCTCGAAAGCGTGGGG |
| P-asv6174 | GCGAGCGTTGTCCGGAATTACTGGGCGTAAAGGACGCGTAGGCTGTCCATTAAGTTGGTAGTGAAATACCGGGGCTCAACCTCGGGGCTGCTATCAAAACTGATGGGCTAAGAGTATCGAAGGGGAGAACAGAA<br>TTCCAGGTGTAGCGGTGAAATGCGTAGATATCTGGGAGAATACCGGTGGCGAAGGCGGTTCTCTGGTCGATTACTGACGCTGATGCGTGAAAGCGTGGGG |
| P-asv6175 | CCAAGCGTTATCCGGAATCATTGGGTTTAAAGGGTCCGTAGGCGGCCTTATAAGTCAGTGGTGAAATCTCCTAGCTCAACTAGGAAACTGCCATTGATACTGTAGGGCTTGAATTTTTGTGAAGTAACTAGAATATG<br>TAGTGTAGCGGTGAAATGCTTAGATATGCTACGGAACACCGATTGCGAAGGCAGCTTACTATTTCATGATTGACGCTGAGGCACGAAAGCGTGGGG |
| P-asv6176 | GCAAGCGTTGTCCGGATTTATTGGGCGTAAAGAGTTCGTAGGCGGTTAGTTAAGTCTGGTGTTAAATCCCGTGGCCCAACCACGGAGCTGCATCAGATACTGGCTGACTTGAGTGCGGTAGAGGCGAGTGGAAT<br>TCCCAGTGTAGCGGTGAAATGCGTAGATATTGGGAAGAACACCAGTGGCGTAAGCGACTCGCTGGGCCGTAACTGACGCTGAGGAACGAAAGCCAGGGG |
| P-asv6177 | CCGGGCGTTATCCGGATTTATTGGGTTTAAAGGGAGCGTAGGCCGGAGATTAAGCGTGTTGTGAAATGTAGACGCTCAACGTCTGCACTGCAGCGCGAACTGGTTTCCTTGAGTACGCACAAAGTGGGCGGAATT<br>CGTGGTGTAGCGGTGAAATGCTTAGATATCACGAAGAACTCCGATTGCGAAGGCAGCTCACTGGAGCGCAACTGACGCTGAAGCTCGAAAGTGCGGGT |
| P-asv6178 | GCTAGCGTTGTTCAGAATTACTGGGCGTAAAGCGCACGTAGGCGGCTTTGTAAGTCAGAGGTGAAATCCCCGGGCTCAACCTGGGAACTGCATCTGTGACTGCATAGCTAGAGTACGGTAGAGGGGGATGGAAT<br>TCCGCGTGTAGCAGTGAAATGCGTAGATATGCGGAGGAACACCGATGGCGAAGGCAATCCCCTGGACCTGTACTGACGCTCATGCACGAAAGCGTGGGG |
| P-asv6179 | GCAAGCGTTATCCGGAATTATTGGGCGTAAAGGGTGCGTAGGTGGTTAGTTAAGTCTTTGGTGTAAGTGCAGTGCTTAACGCTGTGATGCTAAAGAAACTGGCTGACTTGAGTGTGGTAGAGGCAAGTGGAATTC<br>CATGTGTAGCGGTAAAATGCGTAAATATATGGAGGAACACCAGTGGCGAAGGCGGCTTGCTGGGCCATAACTGACACTGAGGCACGAAAGCGTGGGG |
| P-asv6180 | GCAAACGTTGTTCGGAATCACTGGGCATAAAGCGCACGTAGGCGGTTTCGTAAGTGTGGGGTGAAATCCCACGGCTCACCCGTGGAACTGCTCCGCATACTGCGGGACTTGAGGACGGTAGGGGAGAGCAGAA<br>CTCCTGGTGGAGCGGTGAAATGCGTAGATATCAGGAGGAATACCGGTGGCGAAGGCGGCTCTCTGGTCCGTTTCTGACGCTGAGGTGCGAAAGCCAGGGT |
| P-asv6181 | GCGAGCGTTAATCGGATTTACTGGGCGTAAAGCGTGCGTAGGCGGCCGCTTAAGTCGAACGTGAAATCCCCGGGCTCAACCCGGGAACTGCGTCCGATACTGGGCGGCTTGAATTCGGGAGAGGGATGCGGAA<br>TTCCAGGTGTAGCGGTGAAATGCGTAGATATCTGGAGGAACACCGGTGGCGAAGGCGGCATCCTGGACCGACATTGACGCTGAGGCGCGAAAGCCAGGGG |
| P-asv6182 | GCGAACGTTGTTCGGAATCACTGGGCTTAAAGGGCGCGTAGGCGGGCTGCCGCGTCTGGGGTGAAATCCTCCGGCTCAACCGGAGAACGGCCTCGGGTACGGGCGGCCTCGAGGGAGATAGGGGCGTGCGG<br>AACTGTGGGTGGAGCGGTGAAATGCGTTGATATCCACAGGAACTCCGGTGGCGAAGGCGGCACGCTGGATCTTCACTGACGCTGAGGCGCGAAAGCCAGGGG |
| P-asv6183 | CCAAGCGTTATCCGGAATCATTGGGTTTAAAGGGTCCGTAGGTGGATAATTAAGTCAGAGGTGAAATCCTGCAGCTCAACTGTAGAATTGCCTTTGATACTGGTTATCTTGAATCAATGTGAAGTGGTTAGAATATG<br>TAGTGTAGCGGTGAAATGCATAGATATTACATAGAATACCAATTGCGAAGGCAGATCACTAACATTGTATTGACACTGATGGACGAAAGCGTGGGG |
| P-asv6184 | GCAAGCGTTAATCAGAATTATTGGGCGTAAAGCGCATGTAGGTGGTTTAATAAGTTTGATGTGAAATCCCCGGGCTTAACCTGGGAATGGCGTTGAATACTGTTAGACTAGAGTCAGACAGAGGGTAGTGGAATTT<br>CCGGTGTAGCGGTGAAATGCGTAGAGATCGGAAAGAACACCAGTGGCGAAGGCGGCTACCTGGGTCCAGACTGACACTGAGGTGCGAAAGCGTGGGG |
| P-asv6185 | ACGAGTGGCATTCACATTTATTGGGTCTAAAGCGTTCGTAGCTGGATATATACATCTTTAGTGAAATCGGTCAACTTAATTGCCCGGATTGCTTGAGACACGGTATATCTAGAGACCGGGAGGAGTCAGAGGTATG<br>CCTGGGGGAGTGGTAAAATATGATAATCCCAGGTAGACCACCTGCGGCGAAGGCGTCTGACTAGAACGGATCTGACAGTGAGGAACGAAGGCTAGGGG |
| P-asv6186 | GCGAGCGTTGTCCGGAATCATTGGGCGTAAAGGGCTCCTAGGCGGTTCAGTAAGTCGACTGTGAAAATCCAAGGCTCAACCTTGGGACGCCAGTCGATACTGCTGTGACTCGAGTTCGGTAGAGGAGTGTGGAA<br>TTCCTGGTGTAGCGGTGAAATGCGCAGATATCAGGAGGAACACCAACGGCGAAGGCAGCACTCTGGGCCGATACTGACGCTGAAGAGCGAAAGCGTGGGG |
| P-asv6187 | GCTAGCGTTATCCGGATTTACTGGGCGTAAAGGGTGCGTAGGCGGTCTTTCAAGTCAGGAGTTAAAGGCTACGGCTCAACCGTAGTAAGCTCCTGATACTGTCTGACTTGAGTGCAGGAGAGGAAAGCGGAATTC<br>CCAGTGTAGCGGTGAAATGCGTAGATATTGGGAGGAACACCAGTAGCGAAGGCGGCTTTCTGGACTGTAACTGACGCTGAGGCACGAAAGCGTGGGG |
| P-asv6188 | CCAAGCGTTATCCGGAATCATTGGGCGTAAAGGGTGTGTAGGTGGTCTCGTTAGTCTCCTGTTAAAACTTTTGGCTTAACCGGAAGATTGCAGGGGAAACGGCGAGACTGGAGGGTGCGAGAGGTGAATGGAAC<br>TCATGGAGTAGGGGTGAAATCCGTTGATATCATGGGGAACACCAAATGCGAAGGCAATTCACTGGCGCATTTCTGACACTGAAACACGAAGGCGTGGGT |
| P-asv6189 | GCGAGCGTTGTCCGGATTTATTGGGTTTAAAGGGTGCGTAGGTGGCCTTGTAAGTCAGTGGTGAAATGCGGTCGCTCAACGATCGAACTGCCATTGATACTGCGGGGCTTGAGACAGATGGAGGTCGCCGGAAC<br>GGATGGTGTAGCGGTGAAATGCATAGATATCATCCAGAACACCGATTGCGAAGGCAGGTGACTACGCCTGTTCTGACACTGAGGCACGAAAGCATGGGG |
| P-asv6190 | GCAAGCGTTGTTCGGATTTACTGGGCGTAAAGAGCACGTAGGCGGTTGGGAAAGCCTTTTGGGAAATCTCCCGGCTTAACCGGGAAAGGTCGAGAGGAACTATTCAGCTAGAGGACGGGAGAGGAGCGCGGAA<br>TTCCCGGTGTAGCGGTGAAATGCGTAGATATCGGGAAGAAGGCCGGTGGCGAAGGCGGCGCTCTGGAACGTACCTGACGCTGAGGTGCGAAAGCGTGGGG |
| P-asv6191 | CCAAACGTTATTCGGTATCACTGGGCTTAAAGCGTGCGTAGGCGGCCTGGTAGGTGAGATGTGAAAGCCCACGGCTCAACCGTGGAATTGCGTTTCAAACCACCAGGCTTGAGGATGGCAGGGGTGATGGGAAC<br>TGATGGTGGAGCGGTGAAATGCGTTGATATCATCAGGAACACCGGTGGCGAAGGCGCATCACTGGGCCATTTCTGACGCTGAGGCACGAAAGCTAGGGT |
| P-asv6192 | GCGAGCGTTGTCCGGGTTTATTGGGCGTAAAGCGCGCGTAGGCGGCCCGGAAAGTCGGATGTGAAAACCCACGGCTCAACCGTGGGCCTGCATCCGATACTGCCGGGCTCGAGGCAGGCAGGGGAGATCGGA<br>ATTCCTGGTGTAGCGGTGAAATGCGCAGATATCAGGAGGAACACCGGTGGCGAAGGCGGATCTCTGGGCCTGTCCTGACGCTCAGGCGCGAAAGCTGGGGG |
| P-asv6193 | GCAAGCATTAATCGGATTTATTGGGCGTAAAGGGCGCGTAGGCGGATAGTCAAGTCGTGTGTGAAATTTCGGGGCTCAACCTCGAAGCGGCACAGGAAACTAACTATCTAGAGGGTAGATGGAGAGAATGGAATT<br>CCACAAGTAGCGGTGAAATGCGTAGATATGTGGAAGAACACCAGTGGCGAAGGCGATTCTCTAATTTATTCCTGACGCTGAGGCGCGAAAGCAAGGGG |
| P-asv6194 | CCAAGCGTTAATCGGAATTACTGGGCGTAAAGCGTGCGCAGGCGGCGTTTTAAGTCGGGGGCGAAAGCCTGTGGCTCAACCACAGAATTGCCTTCGATACTGGGACGCTTGAGTCTGGTAGAGGTTGGTGGAAC<br>TGCGAGTGTAGAGGTGAAATTCGTAGATATTCGCAAGAACACCGGTGGCGAAGGCGGCCAACTGGACCAGTACTGACGCTGAGGCGCGAAAGCGTGGGG |
| P-asv6195 | GCTAGCGTTGTTTGGAATTACTGGGCGTAAAGGGCGCGTAGGCGGCGCTTCAAGTCAGATGTGAAAGCCCCGGGCTCAACCTGGGAACCGCATTTGAGACTGTTGTGCTTGAGGTCCGGAGAGGTGGGTGGAA<br>TTCCCAGTGTAGAGGTGAAATTCGTAGATATTGGGAAGAACACCGGTGGCGAAGGCGGCCCACTGGACGGAACCTGACGCTGAGGCGCGAAAGCGTGGGG |
| P-asv6196 | GCAAGCGTTATCCGGAATCATTGGGTTTAAAGGGTCCGCAGGCGGGCGTATAAGTCAGTGGTGAAAGCCCATCGCTTAACGATGGAACTGCCATTGATACTGTATGCCTTGAATTCGGTCGAAGTGGGCGGAATA<br>TGACATGTAGCGGTGAAATGCTTAGATATGTCATAGAACACCGATAGCAAAGGCAGCTCACTAGGCCTGGATTGACGCTCAGGGACGAAAGCGTGGGT |
| P-asv6197 | GCTAGCGTTGTTCGGAATTACTGGGCGTAAAGCGCACGTAGGCGGACCTGTCAGTCAGAGGTGAAATCCCGGAGCTTAACTTCGGAACTGCCTTTGATACTGCAGGTCTTGAGGATGGGAGAGGTGAGTGGAAC<br>TGCGAGTGTAGAGGTGAAATTCGTAGATATTCGCAAGAACACCAGTGGCGAAGGCGGCTCACTGGCCCATTACTGACGCTGAGGTGCGAAAGCGTGGGG |
| P-asv6198 | GCGAACGTTGTTCGGAATCACTGGGCTTAAAGGGCGCGTAGGCGGGTTGCCAAGTCTGTGGTGAAATCCTCCAGCTTAACTGGAGAACGGCCGTGGATACTGGCGATCTCGAGGGAGGTAGGGGCGTGCGGAA<br>CTGTGCGTGGAGCGGTGAAATGCATTGATATGCACAGGAACTCCGGTGGCGAAGGCGGCACGCTGGACCTCTTCTGACGCTGAGGCGCGAAAGCCAGGGG |
| P-asv6199 | GCGAGCGTTAATCGGAATTACTGGGCGTAAAGCGCGCGTAGGCGGTTCGGTAAGTCAGGTGTGAAAGCCCCGGGCTTAACCTGGGAATTGCATTTGATACTGCCGGGCTAGAGTCTGGGAGAGGGAAGCGGAA<br>TTCCGGGTGTAGCGGTGAAATGCGTAGATATCCGGAGGAACACCGGTGGCGAAGGCGGCTTCCTGGCCCAAGACTGACGCTGAGGTGCGAAAGCGTGGGG |
| P-asv6200 | GCTAGCGTTATTCGGATTTACTGGGCATAAAGCGCACGTAGGCGGCTCGGTAAGTCGGGGGTGAAAGCCCGCAGCTCAACTGCGGAATTGCCTTCGATACTGCCGAGCTTGAGTGCAGGAGGGGTGACTGGAA<br>CTCCAGGTGTAGCGGTGAAATGCGTAGATATCTGGAGGAACACCAGAGGCGAAGGCGGGTCACTGGACTGCAACTGACGCTGAGGTGCGAAAGCTAGGGG |
| P-asv6201 | CCGAACGTTATTCGGAATCACTGGGCTTAAAGCGTGCGTAGGCGGTCTTAAAAGTTAGGTGTGAAATCCCACAGCTCAACTGTGGAACTGCGCTTAAAACTATAAGGCTAGAGGGATATAGAGGTGAGCGGAACT<br>GATGGTGGAGCGGTGAAATGCGTTGATATCATCAGGAACACCGGCGGCGAAGGCGGCTCACTGGGTATCTTCTGACGCTGAGGTACGAAAGCTAGGGG |
| P-asv6202 | GCGAGCGTTAATCGGAATTACTGGGCGTAAAGCGCACGTAGGTGGCTCCTTAAGTCGGATGTGAAAGCCCTGGGCTTAACCTGGGAATTGCATCCGATACTGGGTGGCTAGAGTACGGTAGAGGGAGGTGGAAT<br>TCCAGGTGTAGCGGTGAAATGCGTAGAGATCTGGAGGAACACCGGTGGCGAAGGCGGCCTCCTGGACTGATACTGACACTGGGGTGCGAAAGCGTGGGG |
| P-asv6203 | GCAAGCGTTGTCCGGATTTACTGGGCGTAAAGAGTATGTAGGTGGGCACTTAAGTCAGATGTGAAATTCCCGGGCTTAACCTGGGAGCTGCATTTGATACTGGGTGTCTAGAGTGCAGGAGAGGAAAGTGGAATT<br>CCTAGTGTAGCGGTGAAATGCGTAGAGATTAGGAAGAACACCAGTGGCGAAGGCGACTTTCTGGACTGTAACTGACACTGAGATACGAAAGCGTGGGT |
| P-asv6204 | GCAAGCGTTGTCCGGAATCATTGGGCGTAAAGCGCGTGTAGGCGGCTCGGTAAGTCCGCTCTGAAAGCCCGGGGCTCAACCCTGGGAGGCGGGCGGATACTGCCGAGCTCGAGTCCGGAAGAGGCGAGTGGA<br>ATTCCTGGTGTAGCGGTGAAATGCGCAGATATCAGGAAGAACACCAATGGCGAAGGCAGCTCGCTGGGACGGTACTGACGCTGAGGTACGAAAGCATGGGG |
| P-asv6205 | CCTAGCGTTGTTCGGAATCATTGGGCGTAAAGCGGGTGTAGGTGGCTTTGTAAGTCGGAAGTGAAAGCCCTGGGCTCAACCCAGGAAGTGCTTTCGATACTGCTTAGCTTGAGTGTGGGAGAGGATGCTGGAAT<br>TCCAGGTGTAGTGGTGAAATACGTAGATATCTGGAGGAACACCGGTGGCGAAGGCGGGCATCTGGCCCAACACTGACACTCAGACCCGAAAGTGCGGGG |

| | |
|---|---|
| P-asv6206 | GCGAGCGTTAATCGGAATTACTGGGCGTAAAGCGAGCGCAGACGGTTACTTAAGCAGGATGTGAAATCCCCGGGCTTAACCTGGGAACTGCGTTCTGAACTGGGTGACTAGAGTGTGTCAGAGGGAGGTAGAATTCCACGTGTAGCAGTGAAATGCGTAGAGATGTGGAGGAATACCGATGGCGAAGGCAGCCTCCTGGGATAACACTGACGTTCATGCTCGAAAGCGTGGGT |
| P-asv6207 | GCGAGCGTTGTTCGGAATCACTGGGCGTAAAGCGTGCGCAGGCGGTTATGAAAGTCAGATGTGAAAGCCCGGGGCTTAACCCTGGAATTGCATTTGAAACTCCATGACTAGAGTACTGGAGGGGTTGGCGGAATTTCGAGTGTAGCAGTGAAATGCGTAGATATTCGAAGGAACACCGGTGGCGTAGGCAGCCAACTGGACAGTAACTGACGCTCATGCACGAAAGCGTGGGG |
| P-asv6208 | GCAAGCGTTATCCGGAATTACTGGGTGTAAAGGGCGTGTAGGCGGGAATGTAAGTCAGATGTGAAATTCCAAGGCTCAACTTTGGAGCTGCATCTGAAACTATGTTTCTTGAGTGCTGGAGAGGAAAGCGGAATTCCTAGTGTAGCGGTGAAATGCGTAGATATTAGGAGGAACACCAGTGGCGAAGGCGGCTTTCTGGACAGTAACTGACGCTGAGGCGCGAAAGCGTGGGG |
| P-asv6209 | CCGGGCGTTATCCAGATTTATTGGGTTTAAAGGGAGCGTAGGCCGGAGATTAAGTGTGTTGTGAAATGTAGACGCTCAACGTCTGACTTGCAGCGCATACTGGTTTCCTTGAGTACGCACAACGTTGGCGGAATTCGTCGTGTAGCGGTGAAATGCTTAGATATGACGAAGAACTCCGATTGCGAAGGCAGCTGACGGGAGCGCAACTGACGCTTAAGCTCGAAGGTGCGGGT |
| P-asv6210 | GCAAGTGTTATCCGGAATCACTGGGCGTAAAGCGTCTGTAGGTGGTTTGATAAGTCAACTGTTAAATCTTGAGGCTCAACCTCAAAATCGCGGTCGAAACTATTAGACTAGAGTATAGTAGGGGTAAAGGAATTTCCAGTGGAGCGCTGAAATGCGTAGATATTGGAAGGAACACCGATGGCGAAGGCACTTTACTGGGCTATTACTAACACTCAGAGACGAAAGCTAGGGT |
| P-asv6211 | GCAAGCGTTGTTAGGAGTCACTGGGCTTAAAGCGTGCGTAGGCGGACTGCCAAGTCAGGAGTGAAATCCAGGGGCTCAACCCCTGAACTGCTCTTGAAACTGGCAGTCTAGAATCCGGGAGAGGTTGGGAGAATTCCTAGTGTAGAGGTGAAATTCGCAGATATTAGGAGGAATACCAGTGGCGTAGGCGCCCAACTGGACCGGTATTGACGCCCAGGCACGAAAGCGTGGGG |
| P-asv6212 | GCAAGCGTTGTTCGGATTTATTGGGCGTAAAGCGCGCGCAGGCGGGTTTTTAAGCCAGATGTGAAATCTCGGGGCTCAACCCCGAAGCTGCGTCTGGAACTGGAGACCTAGAATCTCGGAGGGGGAAGGAGAATTTCACATGTAGGGGTAAAATCCGTAGAGATGTGAAGGAACACCAGAGGCGAAGGCGCCTTCCTGGACGAGTATTGACGCTGAGGCGCGAAAGCGTGGGG |
| P-asv6213 | GCGAACGTTATCCGGATTTACTGGGCGTAAAGCGCGTGCAGGCGGCTCGTTAAGTCGTGCGTGAAAGCTCCCAGGCTCAACCGGGAGAGGTCGTGCGATACTGGCGGGCTAGAGACAGAGAGAGGCAAGTAGAATTCCGGGTGTAGTGGTGAAATGCGTAGATATCCGGAGGAACACCAGTGGCGAAGGCGGCTTGCTGGCTCTGTTCTGACGCTCAGACGCGAAGGCATGGGG |
| P-asv6214 | GCGAGCGTTAATCGGAATTACTGGGCGTAAAGCGCGCGTAGGTGGTTAATCAAGTCAGATGTGAAAGCCCAGGGCTCAACCTTGGAACTGCATTTGAAACTGGTTGACTAGAGTTTTGTAGAGGGCGGTAGAATTTCAGGTGTAGCGGTGAAATGCGTAGAGATCTGAAGGAATACCAGTGGCGAAGGCGGCCACCTGGACATAGACTGACACTGAGGCGCGAAGGCGTGGGG |
| P-asv6215 | GCAAGCGTTGTCCGGAATTATTGGGCGTAAAGAGCTTGTAGGTGGCTTGTCGCGTCTGCCGTGAAAACCCGAGGCTCAACCTCGGGCGTGCGGTGGGTACGGGCAGGCTAGAGTGTGGTAGGGGAGACTGGAACTCCTGGTGTAGCGGTGATATGCGCAGATATCAGGAAGAACACCGATGGCGAAGGCAGGTCTCTGGGCCATTACTGACACTGAGAAGCGAAAGCATGGGG |
| P-asv6216 | GCAAATGTTGTTCGGAATCACTGGGCGTAAAGGGAGCGTAGGCGGGCTTTTAAGTTAGGTGTTTAATCCCGGGGCCCAACCCCGGACAGGCACCTAAAACTGGGAGTCTTGAATAGTGGAGAGGACGGCGGAATTGGGGGTGTAGCGGTGGAATGCGTAGAGATCCCCAAGAACACCAGTTGCGAAGGCGGCCGTCTGGACACATATTGACGCTGAGGCTCGAAGGCATGGGG |
| P-asv6217 | GCAAGCGTTGTCCGGATTTATTGGGTTTAAAGGGTGCGTAGGTGGCTTGTTAAGTCTGGGGTGAAAGGCCGTGGCTCAACTACGAGCAGGGCCCTGGATACTGGCAGGCTTGAGTGCGCGTGAGGCCGGTGGAACGGGTGGTGTAGCGGTGAAATGCATAGATACCATCCAGAACACCGATTGCGGGGCAGCCGGCTAACGCGCAACTGACACTGAGGCACGAAAGCGTGGGG |
| P-asv6218 | GCGAGCGTTAATCGGATTTACTGGGCGTAAAGGGCGCGTAGGCGGTAAGATGGGTGTGATGTGAAAGCCCCGGGCTTAACCTGGGAAGCGCATCGCAAACGATCTAACTAGAGTAGATGAGAGGGTGATGGAATTTCCGGTGTAGCGGTGAAATGCGTAGAGATCGGAAGGAACGTCAATGGCGAAGGCAGTCACCTGGCATCATACTGACGCTGAGGCGCGAAAGCGTGGGG |
| P-asv6219 | GCCAGCGTTGCTCGGAATAACTGGGCGCAAAGGGTTCGTAGGCGGTGCGGCAAGTTGGGAGTGAAATCTCTGGGCTTAACTCAGAGGCTGCTTCCAAAACTGCTGTGCTCGAGTGTGAGAGAGGCTCGTGGAATTGCAGGTGTAGCGGTGAAATGCGTAGATATCTGCAGGAACACCCGTGGCGAAAGCGGCGAGCTGGATCACTACTGACGCTGAGGAACGAAAGCTAGGGG |
| P-asv6220 | GCGAGCGTTGTTCGGAATTATTGGGCGTAAAGTGCGTGTAGGCGGTTTTTTAAGTCTGATGTGAAAGCCCTGGGCTCAACCTGGGAAGTGCATTGGAAACCGGAAGACTTGAGTACGGGAGAGGGCAGTGGAATTCCTAGTGTAGGGGTGAAATCCGTAGATATTAGGAGGAACACCGGTGGCGAAGGCGGCTGCCTGGACCGATACTGACGCTGAGACGCGAAAGCGTGGGG |
| P-asv6221 | GCGAGCGTTGTCCGGAATCACTGGGCGTAAAGGGTGAGTAGGTGGCTAGGTAAGCGTGTGGTGAAAGTCCGGGGCTCAACCCCGGATCTGCCGTGCGAACTGCTTAGCTCGAGCAATGTAGAGGCAGGTGGAATTTCGGGTGTAGCGGTGGAATGCGTAGATATCCGAAAGAACACCAGTGGCGAAGGCGGCCTGCTGGGCATTTGCTGACACTGAATCACGACAGCGTGGGG |
| P-asv6222 | GCGAGCGTTATCCGGATTTATTGGGTTTAAAGGGTGCGTAGGCGGAATATCAAGTCAGCGGTAAAAATTCGGGGCTCAACCCCGTCGTGCCGTTGAAACTGATGTTCTTGAGTGGGCGAGAAGTATGCGGAATGCGTGGTGTAGCGGTGAAATGCATAGATATCACGCAGAACTCCGATTGCGAAGGCAGCATACCGGCGCCCAACTGACGCTGAAGCACGAAAGCGTGGGT |
| P-asv6223 | GCTAGCGTTGCTCGGAATGACTGGGCGTAAAGGGCGCGTAGGCGGCACACGTAGTCAGGCGTGAAATTCCTGGGCTTAACCTGGGGGCTGCGTTTGATACATGTGAGCTAGAGGATGGAAGAGGCTCGTGGAATTCCCAGTGTAGAGGTGAAATTCGTAGATATTGGGAAGAACACCGGTGGCGAAGGCGGCGAGCTGGTCCATTACTGACGCTGAGGCGCGAAAGCGTGGGG |
| P-asv6224 | GCCAGCGTTGTTCGGAATTACTGGGTGTAAAGAGTGCGTAGGCGGTTTCCTAAGTTTGGTGTGAAATCTCCCGGCTTAACTGGGAGGGTGCGCCGAAGACTGGGGAGCTAGAGGATGGGAAAGGCAAGCGGAATTCCTGGTGTAGCGGTGAAATGCGTAGATATCAGGAGGAACACCGGCGGTGTAGACGGCTTGCTAGACCACCCCTGACGCTGAGGCACGAAAGCGTGGGT |
| P-asv6225 | CAAACGTTGTTCGGAATTACTGGGCTTGAAGCGCACGTAGGCGGCGTTGTAAGTGTCGGGTGAAATCCCACGGCTCACCCGTGGAACTGCCCGGCATACTGCGATGCTAGAGTGTGGTAGGGGAGAGCGGAACTCCTGGTGGAGCGGTGAAATGCGTAGATATCAGGAGGAACACCGGTGGCGAAGGCGGCTCTCTGGTCCACAACTGACGCTGAGGTGCGAAAGCTAGGGG |
| P-asv6226 | ACGAGTAGTGGTCACTTTTATTGAGCCTAAAGCGTTCCTAGCCGGTTTTGTAAATCTTCAGATAAAGCCTGGAGCTCAACTCCAGAATTGCCTTTAAGACTGCATCGCTTGAATCCAGGAGAGGTGAGTGGAATTCCGAGTGTAGAGGTGAAATTCGTAGATATTCGGAAGAACACCAGTGGCGAAGGCGGCTCACTGGACTGGTATTGACGCTGAGGTGCGAAAGCGTGGGG |
| P-asv6227 | GCAAGCGTTGTTCGGAATTACTGGGCGTAAAGGGCTCGTAGGTGGCCAACTAAGTCAGACGTGAAATCCCCAGGCTTAACCTGGGAACTGCGTCTGATACTGGATGGCTTGAATTCGGGAGAGGGATGCAGAATTCCAGGTGTAGCGGTGAAATGCGTAGATATCTGGAGGAATACCGGTGGCGAAGGCGGCATCCTGGACCGAGATTGGCACTGAGGAGCGAAAGCCAGGGG |
| P-asv6228 | CCAAGCGTTGTTCGGATTTACTGGGCGTAAAGGGCGCGTAGGCGGCCCCGCAAGTCACTTGTGAAATCTCCGGGCTTAACTCGGAACGGCCAAGTGAAACTGCGAGGCTAGAGTGCAGAAGGGGCAATCGGAATTCTTGGTGTAGCGGTGAAATGCGTAGATATCAAGAGGAACACCTGAGGTGAATACGGGTTGCTGGGCTGACACTGACGCTGAGGCGCGAAAGCCAGGGG |
| P-asv6229 | GCAAGCGTTAATCGGAATTACTGGGCGTAAAGCGTGCGCAGGCGGTTACGTATGTCGGACGTGAAAGCCCCGGGCTCAACCTGGGAATTGCATTCGAAACTGCGTGACTAGAGTGTGGTAGAGGAAGATGGAATTCCACGTGTAGCGGTGAAATGCGTAGATATGTGGAGGAACACCAATGGCGAAGGCAGTCTTCTGGGCCTACACTGACGCTGAGGTACGAAAGCGTGGGT |
| P-asv6230 | GTTAGCGTTGTTCGGAATTACTGGGCGTAAAGGGGGCGCAGGCGGACAGGTCAGTGAAAAGTGGAAGCCCGGGGCTCAACCCCGGAAGGCCTTTTCATACTGCCTGTCTTGAGTACGTGAGAGGAGAGCGGAATTCAGGGTGTAGCGGTGAAATGCTTAGATATCCTGAGGAACACCGATGGCGAAAGCAGCTCTCTGGCACGCTACTGACGCTGAGGCCCGAAAGCGTGGGG |
| P-asv6231 | GCTAGCGTTATTCGGAATCACTGGGCGTAAAGCGCACGTAGGCGGATATGTCAGTCAGGGGTGAAATCCCGGAGCTCAACTCCAGAACTGCCTTTGATACTGAGGATCTTGAGTTCGGGAGAGGTAAGTGGAACTGCGAGTGTAGAGGTGAAATTCGTAGATATTCGCAAGAACACCAGTGGCGAAGGCGGCTTACTGGCCCGATACTGACACTGAGGCACGAAAGCGTGGGG |
| P-asv6232 | CCGAGCGTTGTCCGGAATTATTGGGCGTAAAGAGCTCGTAGGCGGCATGTCGCGTCTGCTGTGAAAGACCGGGGCTTAACTCCGGTTCTGCAGTGGATACGGGCATGCTAGAGGTAGGTAGGGGAAACTGGAATTCCTGGTGTAGCGGTGAAATGCGCAGATATCAGGAGGAACACCGGTGGCGAAGGCGGGTTTCTGGGCCTTACCTGACGCTGAGGAGCGAAAGCATGGGG |
| P-asv6233 | GCAAGCGTTACTCGGAATTACTGGGCGTAAAGCGTGCGTAGGTGGTCGTTTAAGTCTGTTGTGAAAGCCCTGGGCTCAACCTGGGAACTGCAGTAGAAACTGGACGACTAGAGTGTGGTAGAGGGTAGCGGAATTCCTGGTGTAGCAGTGAAATGCGTAGAAATCAGGAGGAACATCCATGGCGAAGGCAGCTACCTGGACCAACACTGACACTGAGGCACGAAAGCGTGGGG |
| P-asv6234 | GCTAGCGTTGTTCGGAATTACTGGGCGTAAAGCGCACGTAGGCGGCTTTGTAAGTTGGGGGTGAAATCCCGGGGCTCAACTCCGGAACTGCCTCCAAAACTACATGGCTAGAGGATGTGAGAGGATAGTGGAATTCCGAGTGTAGAGGTGAAATTCGTAGATATTCGGAAGAACACCAGTGGCGAAGGCGACTGTCTGGCACATTTCTGACGCTGAGGTGCGAAAGCGTGGGG |
| P-asv6235 | CCGAGTGTTAATCGGAATTGCTGGGCGTAAAGCGTGCGCAGGCGGTTTGTTAAGCGAGATGTGAAAGCCCTGGGCTCAACCTAGGAATAGCATTTCGAACTGGCGAACTAGAGTCTTGTAGAGGGGGGTAGAATTCCAGGTGTAGCGGTGAAATGCGTAGAGATCTGGAGGAATACCGGTGGCGAAGGCGGCCCCCTGGACAAAGACTGACGCTCATGCACGAAAGCGTGGGG |
| P-asv6236 | GCAAGCGTTATTCGGATTTACTGGGCGTACAGGGAGCGTAGGCGGTTCGGTAAGCCCTCTGTGAAATCTTCAGGCTTAACCTGGAAAAGTCGGAGGGGACTGCCGGGCTAGAGGACGGGAGAGGAGCGCGGAATTCCCGGTGTAGCGGTGAAATGCGTAGAGATCGGGAGGAAGGCCGGTGGCGAAGGCGGCGCTCTGGAACGTTTCTGACGCTGAGGCTCGAAAGCGTGGGG |
| P-asv6237 | ACAAGCGTTGTCCGGATTTATTGGGCGTAAAGCGAGCGCAGGCGGCCTTTTAAGTCTGATGTGAAAGCCCACGGCTCAACCGTGGAGGGTCATTGGAAACTGGAAGACTTGAATGCAGAAGAGGAAAGTGGAATTCCATGTGTAGCGGTGAAATGCGTAGATATATGGAGGAACACCAGTGGCGAAGGCGACTTTCTGGTCTGTAATTGACGCTGAGGCTCGAAAGCGTGGGG |
| P-asv6238 | CCAGGCGTTATCCGGATTTATTGGGTTTAAAGGGAGCGTAGGCTGGAGATTAAGTGTGTTGTGAAATGTAGACGCTCAACGTCTGAATTGCAGCGCATACTGGTTTCCTTGAGTACGCACAACGTTGGCGGAATTCGTCGTGTAGCGGTGAAATGCTTAGATATGACGAAGAACTCCGATTGCGAAGGCAGCTAACGGGAGCGCAACTGACGCTTAAGCGCGAAGGTGCGGGT |
| P-asv6239 | GCAAGCGTTATCCAGATTTATTGGGTTTAAAGGGTCCGTAGGCGGGCCCGTAAGTCAGTGGTGAAATCTCACAGCTTAACTGTGAAACTGCCGTTGATACTGCGGGTCTTGAGTGTAGTTGAGGTAGCTGGAATGAGTAGTGTAGCGGTGAAATGCATAGATATTACTCAGAACACCAATTGCGAAGGCAGGTTACCAAGTTACAACTGACGCTGAGGGACGAAAGCGTGGGG |
| P-asv6240 | GCTAGCGTTGTTCGGAATCACTGGGCGTAAAGGGCGCGCAGGCGGCCTCTGTCGTCGGCCCGTGAAAGCCCGGGGCTCAACCCCGGAACGGCGGCCGATACGCAGGGGCTGGAGGGCGGAAGAGGAGGGTGGAATACCCAGTGTAGAGGTGAAATTCGTAGATATTGGGTGGAACACCGGTGGCGAGAGCGGCTCTCTGGTCCGGACCTGACGCTGAGGCGCGAAAGCGTGGGG |
| P-asv6241 | GCAAGCGTTGTCCGGAATTACTGGGCGTAAAGGGAGTGTAGGCGGTTTAATAAGTTAGAAGTGTAATACCCAGGGCTTAACTCGGGTGCTGCTTCTAAAACTATTAGACTTGAGTGCAGGAGAGGAGAGTGGAATTCCTAGTGTAGCGGTGGAATGCGTAGATATTAGGAAGAACATCAGTGGCGAAGGCGACTCTCTGGACTGTAACTGACGCTGAGGCTCGAAAGCGTGGGT |
| P-asv6242 | GCGAGCGTTGTCCGGAATTATTGGGCGTAAAGGGCTCGTAGGCGGTTTGTCGCGTCGGGAGTGGAAACCAGGTGCTTAACACCTGGCTTGCTTTCGATACGGGCAGACTAGAGGTATTCAGGGGAGAACGGAATTCCTGGTGTAGCGGTGAAATGCGCAGATATCAGGAGGAACACCGGTGGCGAAGGCGGTTCTCTGGGAATATCCTGACGCTGAGGAGCGAAAGTGTGGGG |
| P-asv6243 | GCAAGCGTTATCCGGATTCATTGGGTTTAAAGGGTGCGTAGGCGGACCTGTAAGTCAGTGGTGAAATCTCTCAGCTTAACTGAGAAACTGCCATTGATACTGCAAGTCTAGAGTATAGATGACGTTGGCGGAATATGACATGTAGTGGTGAAATACTTAGATATGTCATAGAACACCGATTGCGAAGGCAGCTAACGAAACTATAACTGACGCTGAGGCACGAAAGTGCGGGG |
| P-asv6244 | GCTAGCGTTGTTCGGAATCACTGGGCGTAAAGGGCGCGTAGGCGGCATGGCGTGTCGGCTTGTGAAAGCCCGGGGCTCAACCCCGGAGGTGCGGCCGAAACTGCCGCGCTGGAGGCCGGAAGAGGAGGGTGGAATACCCAGTGTAGAGGTGAAATTCGTAGATATTGGGTGGAACACCGGTGGCGAAAGCGGCTCCCTGGTCCGGACCTGACGCTGAGGCGCGAAAGCGTGGGG |

| | |
|---|---|
| P-asv6245 | GCAAGCGTTAATCGGAATTATTGGGCGTAAAGGGTGTGCAGGTTGTTTTTAAAAATTAATTTTTAAATATATTAAAGTAATTTAATATGAAAATTAATACTTATAAACTTGAGTATAAAAGAAGAAATTAGAATTTCAAGAGTAGCAGTGAAATGCTTAGATATTTGAAAGAATACTAGTGGCGAAGGCGTATTTCTATTTTATTACTGACACTAATACACGAAAGCATGGGG |
| P-asv6246 | GCGAGCGTTGTCCGGAATTACTGGGCGTAAAGGGCTCGTAGGTGGTGTGTCGCGTCGTCTGTGAAATTCCGGGGCTTAACTCCGGGCGTGCAGGCGATACGGGCATGCTTGAGTGCTGTAGGGGTAGCTGGAATGCCTGGTGTAGCGGTGAAATGCGCAGATATCAGGCAGAACGCCGATGGCGAAGGCAGGTTACTGGGCAGTTACTGACGCTGAGGAGCGAAAGCATGGGT |
| P-asv6247 | GCGAGCGTTGTCCGGAATTACTGGGCGTAAAGGGAGCGTAGGCGGTCGATTAAGTTAGATGTGAAACCCCCGGGCTTAACTTGGGGACTGCATCTAATACTGGTTGACTTAGAGTACAGGAGAGGGAAGCGGAATTCCTAGTGTAGCGGTGAAATGCGTAGATATTAGGAGGAACACCAGTGGCGAAGGCGGCTTTCTGGACTGACACTGACGCTGAGGCTCGAAAGCGTGGGG |
| P-asv6248 | CCGAGCGTTATTCAGAATCACTGGGCGTAAAGGGAGCGTAGGCGGCGCGGTAAGTCAGATGTGAAAGCCCAGGGCTCAACCCTGGAACTGCATTCGATACTGCCGTGCTAGAGTAATGGAGAGGTAATCGGAATTTTGGGTGTAGCAGTGAAATGCGTAGATATCCAAAGGAAGACCAATGGCGAAGGCAGATTACTGGACATTCACTGACGCTGAGGCTCGAAGGCTAGGGG |
| P-asv6249 | GCGAGCGTTATCCGGATTTATTGGGTTTAAAGGGTGCGTAGGCGGAAGTTCAAGTCAGCGGTAAAATCGAGGGGCTCAACCCCTTTAAGCCGTTGAAACTGGGTTTCTTGAGTGGGCGAGAAGTGTGCGGAATGTGTGGTGTAGCGGTGAAATGCATAGATATCACACAGAACTCCGATTGCGAAGGCAGCACACCGGCGCCCAACTGACGCTGAGGCACGAAAGCGTGGGG |
| P-asv6250 | GCTAGCGTTGTTCGGAACCACTGGGCGTAAAGCGCGCGTAGGCGGACTGATTAGTTGGGGGTGAAAGCCCGGAGCTCAACTCCGGAACTGCCTTCAATACTGTCAGTCTTGAGGTCGAGAGAGGTAAGTGGAATTCCGAGTGTAGAGGTGAAATTCGTAGATATTCGGAAGAACACCAGTGGCGAAGGCGACTTACTGGCTCGATTCTGACGCTGAGGTGCGAAAGCGTGGGG |
| P-asv6251 | GCTAGCGTTGCTCAGAATGACTGGGCGTAAAGGGCGCGTAGGCGGACGGCACAGTCAGGCGTGAAATTCCTGGGCTCAACCTGGGGACTGCGTCTGGGACGTGTTGCCTTGAGTATGGAAGAGGGTTGTGGAATTTCCAGTGTAGAGGTGGAACTCGTAGATATTGGAAAGAACACCGGTGGCGAAGGCGGCAACCTGGTCCATTACTGACGCTGAGGCGCGAAAGCGTGGGG |
| P-asv6252 | GCAAGCGTTAATCGGAATGACTGGGCGTGAAGGGCGAGTAGGCGGTTGTATAGGTGTGGTGTGTAAGACACGGGCTTAACCTGTGGGTGTCACCGCAAACGGTATGACTAGAGTGCAGTAGAGGGGAGTGGAATTTCCGGAGGAGCGGTGAAATGCGTAGATCTCGGAAGGAACACCGGTGGCGAAGGCGACTCTCTGGACTGTAACTGACGCTGAGGCGCGAAGGCGTGGGG |
| P-asv6253 | TCAAGCGTTGCCCGGATTTATTGGGCGTAAAGAGCTCGTAGGCGGTTCAACAAGTCGGTCGTAAAAGTTTAGGGCTCAACCCTAAAATGTCGATCGATACTGTTGTGACTAGGATACGGTAGAGGTGAATGGAATTCCGAGTGTAGCGGTGAAATGCGTAGATATTCGGAGGAACACCAATTGCGAAGGCAGTTCACTGGACCGTGATCGACGCTGAGGAGCGAAAGCTAGGGG |
| P-asv6254 | GCAAGCATTAATCGGATTTATTGGGCGTAAAGGGCGCGTAGGTGGGCCATTAAGTGAGTCGTGAAATTTCTGGGCTCAACCTAGAAGCTGCGGCTTAAACTGGTGGTCTAGAGGCTAGGCGGAGAAAACGGAATTCCAAGTGTAGCGGTGAAATGCGTAGATATTTGGAAGAACACCCGTGGCGAAAGCGGTTTTCTAGCTTATGCCTGACACTTAGGCGCGAAAGCAGGGGT |
| P-asv6255 | GCGAGCGTTGTCCGGAATCATTGGGCGTAAAGAGCTCGTAGGCGGCTTGGTAAGTCGGATGTGAAACCTCCAGGCTCAACCTGGAGTCGCCATCCGATACTGCCATAGCTAGAGTCCGGTAGGGGCCCACGGAACTCCTGGTGTAGCGGTGAAATGCGCAGATATCAGGAAGAACACCGGTGGCGAAGGCGGTGGGCTGGGCCGGCACTGACGCTGAGGAGCGAAAGCGTGGGG |
| P-asv6256 | GCGAGCGTTAATCGGAGTTACTGGGCGTAAAGCGCGCGTAGGCGGTTAGATGCGTCCGGTGTGAAAGCCCTGGGCTTAACCTGGGAATGGCATCGGAGACGGTTTAGCTAGAGTGGGACAGAGGATGGCGGAATTCCCGGTGTAGCGGTGAAATGCGTAGAGATCGGGAGGAACACCAGTGGCGAAGGCGGCCATCTGGGTCCACACTGACGCTGAGGTGCGAAAGCGTGGGG |
| P-asv6257 | GCAAGCGTTACTCGGAATTACTGGGCGTAAAGCGCGCGTAGGCGGTCTTGTATGTCGATTGTGAAAGCCCAGGGCTCAACCTTGGAACCGCAGCCGAAACTGCAAGACTTGAATGTTGGAGAGGGTGGGGGAATACCTAGTGGAGAAGTGAAATTCGTAGAGATTAGGTGGAACACCAGAGGCGAAGGCGCCTGCCTGGCCAAATATTGACGCTGAGGCGCGAAAGTGCGGGG |
| P-asv6258 | GCGAGCGTTATCCGGAATTATTGGGCGTAAAGCGCGCGCAGGCGGTCATTTAAGTCAGGTGTGAAAGGCTGCGGCTCAACCGCAGAGCGGCACCTGAAACTGGATGACTTGAGTGCAGGAGAGGAGAGTGGAATTCCCGGTGTAGCGGTGGAATGCGTAGAGATCGGGAGGAACACCAGTGGCGAAGGCGGCTCTCTGGCCTGTTACTGACGCTGAGGCGCGAAAGCGTGGGG |
| P-asv6259 | GCAAGCGTTAATCGGAATAACTGGGCGTAAAGGGCATGCAGGCGGTTCATCAAGTAGGATGTGAAATCCCCGGGCTCAACCTGGGAACAGCATACTAAACCGGTGGACTAGAGTATTGCAGGGGGAGACGGAATTCCAGGTGTAGCGGTGGAATGCGTAGATATCTGGAAGAACACCAAAGGCGAAGGCAGTCTCCTGGGCAAATACTGACGCTCATATGCGAAAGCGTGGGT |
| P-asv6260 | TCAAGTGATGTCCAATATTATTGGGCCTAAAGCGTCCGTAGCTGGCCGCGCAAGTCCGTCGGGAAATCCACCTGCCCAACAGGTGGGCGTCCGGCGGAAACTGTGTGGCTTGGAACCGGAAGGCTCAGAGGGTACGCCCGGGGTAGGAGTGAGATCCCGTAATCGTGGACGGACCACCGATGGGGAAACCACTCTGAGAAGACGGCTTCGACAGTGAGGGACGAAAGCTAGGGT |
| P-asv6261 | CCAGGCGTTATCCGGATTTATTGGGTTTAAAGGGAGTGTAGGCGGTTTGTTAAGCGTGTTGTGAAATTTAGATGCTCAACATTTAACTTGCAGCGCGAACTGGCGAACTTGAGTGCACACAACGTATGCGGAATTCATGGTGTAGCGGTGAAATGCTTAGATATCATGAAGAGCTCCGATTGCGAAGGCAGCATACGGGAGTGTAACTGACGCTTAAGCTCGAAGGTGCGGGT |
| P-asv6262 | CCAAGCGTTATCCGGATTTATTGGGTTTAAAGGGTGCGTAGGCGGCTCTATAAGTCAGTAGTGAAATCTCGAAGCTCAACTTCGAACGTGCTATTGAAACTGTAGAGCTTGAGTACAGACGAGGCAGGCGGAATGTGTTGTGTAGCGGTGAAATGCTTAGATATAACACAGAACACCGATAGCGAAGGCAGCTTGCCAGACTGTAACTGACGCTGAGGCACGAAAGCGTGGGG |
| P-asv6263 | GCTAGTGTTATTCGTATTGACTGGGTGTAAAGGGTAAGTAGGCGGTCTATTAAGTTTTAAATAAAATACTAAATTTTAATTTAGAAAGTTTAAAATACTATTAGACTAGAGTTAAACCGGAGATAACTGTACATTAAGTGTAAAAGTGAAATTTGTTAATATTTAAAGGACATGCGATAAGCGAAAGCTGTTATCTAGGTAAATACTGACGCTGAGGTACAAAAGCATAGGG |
| P-asv6264 | GCAAGCGTTAATCGGAATTATTGGGCGTAAAGCGAGTGCAGACGGTTACTTAAGCCAGTCGTGAAATCCCCGGGCTCAGCCTGGGAATGGCGGTGGAAACCGGCGCACTAGAGTATGGGAGAGGGTGGTGGAATTCCAGGTGTAGCGGTGAAATGCGTAGAGATCTGGAGGAACATCAGTGGCGAAGGCGGCCACCTGGCCCAATACTGACGCTGAGGCACGAAAGCGTGGGG |
| P-asv6265 | GCAAGCGTTAATCGGAATTACTGGGCGTAAAGCGCGCGTAGGCGGCTTGTTAAGTCAGATGTGAAATCCCCGAGCTCAACTTGGGAACTGCATTTGATACTGGCAGGCTAGAGTATGTTAGAGGAAAGCGGAATTCCGGGTGTAGCGGTGAAATGCGTAGATATCCGGAGGAACATCAGTGGCGAAGGCGGCTTTCTGGAACAATACTGACGCTGAGGTGCGAAAGCGTGGGG |
| P-asv6266 | GCAAGCGTTATCCGGAATCATTGGGTTTAAAGGGTCCGCAGGCGGTCAATTAAGTCAGAGGTGAAATCCAGTCGCTTAACGATTGAACTGCCTTTGATACTGGTTGACTTGAGTCATATGGAAGTAGATAGAATGTGTAGTGTAGCGGTGAAATGCATAGATATTACACAGAATACCGATTGCGAAGGCAGTCTACTACGTATGTACTGACGCTGAGGGACGAAAGCGTGGGG |
| P-asv6267 | CCGAGCGTTATCCGGATTTATTGGGTTTAAAGGGTGCGTAGGCGGCCTATTAAGTCAGGGGTGAAATACGGTGGCTCAACCATCGCAGTGCCTTTGATACTGATGGGCTTGAATCCATTTGAAGTGGGCGGAATAAGACAAGTAGCGGTGAAATGCATAGATATGTCTTAGAACTCCGATTGCGAAGGCAGCTCACTAAGCTGGTATTGACGCTGATGCACGAAAGCGCGGGG |
| P-asv6268 | GCAAGCGTTAATCGGAATTATTGGGCGTAAAGCGAGTGCAGACGGTTACTTAAGCCAGATGTGAAATCCCCAAGCTTAACTTGGGACGTGCATTTGGAACTGGGTGACTAGAGTGTGTCAGAGGGAGGTAGAATTCCACATGTAGCGGTGGAATGCGTAGAGATGTGGAGGAATACCAATGGCGAAGGTAGCCTCCTGGCCTAATACTGACGCTCATGCACGAAAGCGTGGGG |
| P-asv6269 | GCAAGCGTTATCCGGAATGATTGGGCGTAAAGGGTCCGCAGGTGGCATTGTAAGTCTGCTGTTAAAGAGTTTGGCTCAACCGAATAAAAGCAGTGGAAACTACAAAGCTAGAGTGTGGTCGGGGCAGAGGGAATTCCTGGTGTAGCGGTGAAATGCGTAGATATCAGGAAGAACATCGGTGGCGAAGGCGCTCTGCTAGGCCAAGACTGACACTGAGGGACGAAAGCTAGGGG |
| P-asv6270 | GCGAGCGTTGTCCGGAATGACTGGGCGTAAAGCGCGCGCAGGCGGTTTTCTACGTCTGGGGTGAAAGCCCGGGGCTCAACTCCGGGACGGCCTTGGAAACGGGGAGACTTGAGGGTCGGAGAGGCAAGGGGAATTCCACGTGTAGCGGTGAAATGCGTAGAGATGTGGAGGAACACCAGTGGCGAAGGCGCCTTGCTGGACGAAACCTGACGCTGAGGCGCGAAGGCGTGGGG |
| P-asv6271 | GCGAGCGTTAATCGGAATTACTGGGCGTAAAGGGTGCGTAGGTGGTTAAATTAGTTAGGTGTGAAATTCCTGAGCTCAACTTGGGGACTGCGCCTAATACTGTTCAACTAGAGTACTGTAGAGGGAAGTGGAATTTCCGGTGTAGCGGTGAAATGCGTAGATATCGGAAAGAACATCAGTGGCGAAGGCGACTTCCTGGACAGATACTGACACTGAGGCACGAAAGCGTGGGG |
| P-asv6272 | GCAAGCGTTGTTCGGATTTATTGGGCGTAAAGCGCGCGCAGGCGGATTTGTAAGTCAGATGTGAAATCCCGGGGCTCAACCCCGGAACTGCGTCTGAAACTATGAGTCTAGAATCTCAGAGGGGGTAGGGGAATTTCACGTGTAGGGGTAAAATCCGTAGAGATGTGAAGGAACACCGGTGGCGAAGGCGCCTACCTGGATGAGTATTGACGCTGAGGCGCGAAAGCGTGGGG |
| P-asv6273 | GCTAGCGTTACTCGGAATTACTGGGCGTAAAGGGCGCGTAGGCGGCGTTCCAAGTTAGGCGTGAAAGTCCTGGGCTCAACCTGGGAACTGCGCTTAAGACTGGAATGCTTGAGGATGGAAGAGGCTCGTGGAATTCCCAGTGTAGAGGTGAAATTCGTAGATATTGGGAAGAACACCGGTGGCGAAGGCGGCGAGCTGGTCCATTACTGACGCTGAGGCGCGATAGCGTGGGG |
| P-asv6274 | GCTAGCGTTATTCGGAATTACTGGGCGTAAAGCGCACGTAGGCGGCTTTGTAAGTTAGAGGTGTAAGCCTGGAGCTCAACTCCAGAATTGCCTTTAAGACTGCATCGCTTGAATCCAGGAGAGGTGAGTGGAATTCCGGGTGTAGAGGTGAAATTCGTAGATATTCGGAAGAACACCAGTGGCGAAGGCGGCTCACTGGACTGGTATTGACGCTGAGGTGCGAAAGCGTGGGG |
| P-asv6275 | ACAAACGTTGCTCGGAATTACTGGGCGTAAAGCGTACGTAGGCGGATTCGCAAGTCGGGCGTGAAATCCCCGGGCTCAACCCGGGAACTGCGTTCGAAACTGCGTATCTCGAGTGATGGAGAGGGAATCGGAATTCTCGGTGTAGAGGTGAAATTCGTAGATATCGAGAGGAACACCAGTGGCGAAAGCGGATTCCTGGACATTTACTGACGCTGAGGTACGAAAGCGTGGGG |
| P-asv6276 | GCGAGCGTTATTCGGTATTACTAGGCGTAAAGCGAGTGTAGGCGGACTCTTAAGTCTGACGTTAAATCTCCTGGCTTAACTAGGAGCCGTCGTCGGATACTGAGAGCCTCGAATGGGGTAGGGGGCAGCGGAATTCCCGGTGTAGCGGTGAAATGCGTAGATATCGGGAGGAACACCTATGGCGAAAGCAGCTGCCTGGGCCTCTATTGACGCTAAGACTCGAAAGCTGGGGG |
| P-asv6277 | GCAAGCGTTACTCGGAATTACTGGGCGTAAAGCGTGCGTAGGTGGTTCGTTAAGTCTGATGTGAAAGCCCTGGGCTCAACCTGGGAATTGCATTGGAAACTGGCGAGCTAGAGTACGGTAGAGGGTAGTGGAATTCCTGGTGTAGCAGTGAAATGCGTAGAGATCAGGAGGAACACCCGTGGCGAAGGCGACTACCTGGACCAGTACTGACACTGAGGCACGAAAGCGTGGGG |
| P-asv6278 | CCGAACGTTATTCGGTATCACTGGGCTTAAAGAGTTCGTAGGCGGCCATGCAGGTCAGGTGTGAAATCCCACAGCTCAACTGTGGAACTGCGCTTGAAACCGCATGGCTTGAGGGAGATAGAGGTGAGCGGAACAGATGGTGGAGCGGTGAAATGCGTTGATATCATCTGGAACACCGGTGGCGAAAGCGGCTCACTGGATCTCTACTGACGCTGAGGAACGAAAGCTAGGGT |
| P-asv6279 | TCGAGCGTTATCCGGAATTATTGGGCGTAAAGGGTGTGTAGGTGGACGTGTTAGTCCTGTGTGAAAGCCTGTCGCTCAACGACAGATCCGCATGGGAAACGGCACGACTGCGAGGGTGCGAGAGGTGAACGGAACTCATGGTGTAGGGGTGAAATCCGTTGATATCATGGGGAACGCCAAAAGCGAAGGCAGTTCACTGGCGCATGCCTGACACTGAGACACGAAAGCCAGGGT |
| P-asv6280 | CCAAGTGGTAGTCATTATTATTGGGCTTAAAGTGTTCGTAGCCGGACCGGCAAGTCCGCTGTGAAATCCACGCGCTTAACGTGTGGGCGTGCAGCGGAGACTGCAGGTCTAGGGACCGGACGGCGTCGAGGCTATTCCAGGGGTAGGGGTGAAATCCTAAAATCCTTGGAGGAGCACCTGTGGCGAAAGCGCTCGACGAGTACGGATCCGACGGTGAGGAACGAAAGCCAGGGG |
| P-asv6281 | CCAAGCGTTATTCGGAATCACTGGGCTTAAAGGGTGCGCAGGCGGATTAGCAAGCATCTCGTGAAATCCCTCCGCTCAACGGAGGAATTGCGGGGTGAACTGCTGGTCTTGAGGCAGGTAGAGGCCGTCGGAACGCTAGGTGGAGCGGTGAAATGCGTAGATATCTAGCGGAACGCCAAAGGTGAAGACAGGCGGCTGGGCCTGACCTGACGCTGAGGCACGAAAGCGTGGGG |
| P-asv6282 | GCGAGCGTTAATCGGAATTACTGGGCGTAAAGCGCACGTAGGCGGTTTGTTAAGCAGGATGTGAAATCCCCGGGCTCAACCTGGGAACTGCATCCTGAACTGGCAGGCTAGAATGCAGTAGAGGATAGTGGAATTCCTGGTGTAGCAGTGAAATGCGTAGATATCAGGAGGAACATCGATGGCGAAGGCAGCTATCTGGACTGACATTGACGCTGAGGTGCGAAAGCGTGGGG |
| P-asv6283 | TCGAGTGGTGATCACTTTTATTGAGTCTAAAGCGTCCGTAACCGGTCTTATAAATCTTCAGATAAATTCTTCCGCTTAATGGAAGAACTTCTGAAGAGACTGTAAGACTTGGGACCGGGTGAGGTTGAATGTACTTTCAGGGTAGGGGTAAAATCCTGTAATCCTGAAAGGACAACCGGTGGCGAAAGCGTTCAACTAGAACGGATCCGACGGTGAGGGACGAAGGCTAGGGG |

| | |
|---|---|
| P-asv6284 | GCTAGCGTTGTTCGGAATTACTGGGCGTAAGCGAGCGTAGGTTGCGCCGCAAGTCAGATGTGAAATCCCAGGGCTTAACCCTGGAACTGCATTTGAAACTGCGGTGCTGGAATCTCAGAGGGGGTAGCGGAATTCCAAATGTAGGGGTGAAATCCGTAGATATTTGGAGGAACACCGGTGGCGAAGGCGGCTACCTGGATGAGTATTGACACTGAGGCTCGAAAGCGTGGGG |
| P-asv6285 | GCTAGCGTTGTTCGGAATTACTGGGCGTAAAGCGCACGTAGGCGGATACTTAAGTCAGGGGTGAAATCCCGGGGCTCAACCCCGGAACTGCCTTTGATACTGGGTATCTGGAGTCCGGAAGAGGTGAGTGGAATTCCGAGTGCAGAGGTGAAATTCGTAGATATTCGCAAGAACACCAGTGGCGAAGGCGGCTCACTGGCCCGATACTGACGCTGAGGCACGAAAGCGTGGGG |
| P-asv6286 | CCGAGCGTTGTTCGGAATCACTGGGCGTAAAGGGAGCGTAGGCGGCGAGGCAAGTCAGATGTGAAATCCCAGGGCTCAACCCTGGAACTGCATTTGATACTGCTTTGCTAGAGTAATGGAGGGGTAGCTGGAATTATCGGTGTAGCAGTGAAATGCGTAGATATCGATAGGAACACTCGTGGCGAAAGCGAGCTACTGGACATTTACTGACGCTGAGGCTCGAAGGCCAGGGT |
| P-asv6287 | TCAAGCGTTAATCGGAATCACTGGGCTTAAAGGGTCCGCAGGCGGATTGGTAAGTATCTTGTGAAATACCATGGCTCAACCATGGAATTGCTCGGTAAACTGCTAATCTTGAACACGGTAGGGGCCATCGGAACTCTAGGTGGAGCGGTGAAATGCGTAGATATCTAGAGGAACGCCAGAGGCGAAGGCGGATGGCTGGGCCGTTGTTGACGCTCAGGGACGAAGGCATGGGT |
| P-asv6288 | TCAAGCGCTACCCGGAGTTACTGGGCGTAAAGGGCGTGTAGGGGGCTTTGTAAGTTCATTGTCAAATCTCAGGGCTTAACCCTGAGGCTGCAATGAAAACTGCAAGGCTAGAGGACGTCAGAGGAGAGTGGAACGCACGGTGTAGAGGTGAAATTCGTTGATATCGTGCGGAACATCAAAAGCGAAGGCATCTCTCTGGGGCGTTCCTGACCCTGAGGCGCTAAAGCTTGGGT |
| P-asv6289 | CCGAGCGTTGTTCGGAATCACTGGGCGTAAAGGGAGCGTAGGCGGCGTGGCAAGTCAGACGTGAAATCCCGGGGCTCAACCCCGGAACTGCGTCCGATACTGCCGTGCTAGAGGATTGGAGAGGTAGCTGGAATTCACGGTGTAGCAGTGAAATGCGTGGATATCGTGAGGAACACTCGTGGCGAAAGCGAGCTACTGGACAACTCCTGACGCTGAGGCTCGAAGGCCGGGGG |
| P-asv6290 | GCAAGCGTTATCCGGAATCACTGGGTTTAAAGGGTGCGTAGGCGGGACATTAAGTCTAAGGTGAAAGTTTGTCGCTTAACGATAAAATTGCCTTGGATACTGGTGTTCTTGAATTAGATTGAGGATAGCGGAATGTGGCATGTAGCGGTGAAATGCATAGATATGCCATAGAACACCAATTGCGAAGGCAGCTATCTGGGTCTGGACTGACGCTGAGGCACGAAAGCGTGGGG |
| P-asv6291 | CCAAGCGTTATCCGGATTCATTGGGTTTAAAGGGTGCGTAGGCGGATTAGTAAGTCAGTGGTGAAATCTTCCAGCTTAACTGGGAAACTGCCATTGATACTGCTAGTCTTGAGTGTACTTGAAGTGGGCGGAATAAGTCATGTAGCGGTGAAATGCATAGATATGACTTAGAACACCAATTGCGTAGGCAGCTCACTAAATTACAACTGACGCTGAGGCACGAAAGCGTGGGT |
| P-asv6292 | CCAAGCGTTATCCGGATTTACTGGGCGTAAAGCGTCTGTAGCCGGTAATATAAGTCTGCATTCAAATCCCGAAGCTCAACTTCGGAAAGGGTGCAGATACTGTATAACTTGAATGGAATAGGGGGTAGTCGAATTTCCAGTGGAGGAGTGAAATCCGTTGATATTGGAAGGAATACCAAAAGCGAAGGCAGCTACCTATATTCTTATTGACGGTCAGAGACGAAAGCTTGGGT |
| P-asv6293 | GCTAGCGTTGTTCGGAATCACTGGGCGTAAAGCGCACGTAGGTGGATTAGTAAGTCAGGGGTGAAAACCCAGAGCTCAACTCTGGAATTGCCCTTGATACTGCTAGTCTAGAGTTTGGGAGAGGTAAGCGGAATTCCGAGTGTAGAGGTGAAATTCGTAGATATTCGGAGGAACACCAGAGGCGAAGGCGGCTTACTGGCCCAATACTGACACTGAGGTGCGAAAGCGTGGGG |
| P-asv6294 | CCGAGCGTTATCCGGATTTACTGGGCGTAAAGAGTCCGTAGCTGGCATTGTAAGTCTTGATTCAAATACTCTGACTCACTCAGAGAAAGGGTCAAGGTACTGCAAAGCTAGAGATCATAGGGGGTTAGTGGAATTTCCGGTGGAGGGGTGAAATCCGTTGATATCGGAAGGAATGCCGAAAGCGAAAGCAGCTAACTACTATATTCCGACAGTGAGGGACGAAAGCTAGGGT |
| P-asv6295 | GCTAGCGTTGTCCGGATTTATTGGGCGTAAAGAGCTCGTAGGCGGTTCGGTAAGTCAGGTGTGAAATCCCCACGCTCAACGTGGAGGCGCCACCTGAAACTGCCGTGACTCGAGTCCGGTAGGGGAGTGTGGAATTCCCGGTGTAGCGGTGAGATGCGCAGATATCGGGAGGAACACCCGTGGCGAAGGCGGCACTCTGGGCCGGTACTGACGCTGAGGAGCGAAAGCGTGGGG |
| P-asv6296 | GCAAGCGTTATCCAGATTTATTGGGTTTAAAGGGTCCGTAGGCGGATCGGTAAGTCAGTGGTGAAATCTCATAGCTTAACTATGAAACTGCCATTGATACTGTCGGTCTTGAGTGTATTTGAAGTAGCTGGAATGAGTAGTGTAGCGGTGAAATGCATAGATATTACTCAGAACACCAATTGCGAAGGCAGGTTACTAAGATACAACTGACGCTGATGGACGAAAGCGTGGGG |
| P-asv6297 | GCAAGCGTTAATCGGAATTACTGGGCGTAAAGCGTGCGCAGGCGGTTTTGTAAGACAGGCGTGAAATCCCCGGGCTCAACCTGGGAATTGCGCTTGTGACTGCAAGGCTGGAGTGCGGCAGAGGGGGATGGAATTCCGCGTGTAGCAGTGAAATGCGTAGAGATCGGAAGGAACACCAATGGCGAAGGCGGCTACCTGGCCTAATACTGACACTGAGGCACGAAAGCGTGGGG |
| P-asv6298 | GCGAGCGTTAATCGGATTTACTGGGCGTAAAGCGTGCGTAGGCGGCTTTTTAAGTCGGATGTGAAATCCCTGAGCTTAACTTAGGAGTTGCATTCGATACTGGGAAGCTAGAGTATGGGAGAGGATGGTAGAATTCCAAGTGTAGCGGTGAAATGCGTAGAGATTTGGAGGAATACCAGTGGCGAAGGCGGCCCCCTGGCACAATACTGACGCTCAAGTGCGAAAGCGTGGGG |
| P-asv6299 | GCAACCGTTGCTCGGATTCACTGGGCATAAAGCGCGCGTAGGTGGTACTGCAAGTCACTGGTGAAATCCCCCGGCCCAACCGGGGAACGGCCGGTGAGACTGCGGTGCTGGAGGACGGGAGGGGAGAGTGGAACTCTTGGTGTAGCGGTGAAATGCGTAGATATCAAGAGGAACGCCGGTGGCGAAAGCGGCTCTCTGGACCGTACCTGACACTGAGGCGCGAAAGCTAGGGG |
| P-asv6300 | GCTAGCGTTGTTCGGAATTACTGGGCGTAAAGAGTGCGTAGGCGGTTTAGTAAGTTGGAAGTGAAAGCCCGGGGCTTAACCTCGGAATTGCTTTCAAAACTACTAATCTAGAGTGTAGTAGGGGATGATGGAATTCCTAGTGTAGAGGTGAAATTCTTAGATATTAGGAGGAACACCGGTGGCGAAGGCGGTCATCTGGGCTACGACTGACGCTGATGCACGAAAGCGTGGGG |
| P-asv6301 | GCAAGCGTTATCCGGAATTACTGGGTGTAAAGAGTACGTAGGCGGTCTGATAAGTCAGATGTGAAAGCCCGAGGCTCAACCTCGGGACTGCATTTGAAACTGTCAGACTAGAGTACAGGAGAGGTAAGTGGAATTCCTAGTGTAGCGGTGAAATGCGTAGATATTAGGAAGAACACCAGTGGCGAAGGCGGCTTACTGGACTGTAACTGACGCTGAGGTACGAAAGCGTGGGG |
| P-asv6302 | TCGAGTGGTGATCACTTTTATTGAGTCTGAAGCGTCCGTAACCGGTCTTATAAATCTTCAGATAAATTCTTCCGCTTAATGGAAGAACTTCTGAAGAGACTGTAAGACTTGGGACCGGGTGAGGTTGAATGTACTTTCAGGGTAGGGGTAAAATCCTGTAATCCTGAAAGGACGACCGGTGGCGAAAGCGTTCAACTAGAACGGATCCGACGGTGAGGGACGAAGGCTAGGGG |
| P-asv6303 | CCAAGTGGCAACCACATTTATTGGGCCTAAAGCGTCCGTAGCCGGTTAAGTAAATCTTCTGTGAAATTGTATTGCTCAACAATACGGCGTGCAGGAGACACTGCTTAGCTTGGGACCGGGAGGAGTCAGAGGTATTTCAGGGGGAGCGGTAAAATGCTATAATCCTTGGAAGACCACCTGTGGCGAAGGCGTCTGACTAGAACGGATCCGACGGTGAGGGACGAAAGCCAGGGG |
| P-asv6304 | GCGAGCGTTAATCGGATTTACTGGGCGTAAAGCGTGCGTAGGCGGCTTTTTAAGTCGGATGTGAAATCCCTGAGCTTAACTTAGGAATTGCATTCGATATTGGGAAGCTAGAGTATGGGAGAGGATGGTAGAATTCCAGGTGTAGCGGTGAAATGCGCAGAGATCTGGAGGAATACCGATGGCGAAGGCAGCCATCTGGCCTAATACTGACGCTGAGGTACGAAAGCATGGGG |
| P-asv6305 | GCTAGCGTTGTTCAGAATTACTGGGCGTAAAGCGCATGTAGGCGGCTTTTCAAGTTAGGTGTGAAATCCCTGGGCTTAACCCAGGAACTGCACTTAATACTGTTAAGCTTGAGAATGGTAGAGGATGATGGAATTCCCAGTGTAGAGGTGAAATTCGTAGATATTGGGAGGAACACCGATTGCGAAGGCAGTCATCTGGGCCATTTCTGACGCTGAGATGCGAAAGCGTGGGT |
| P-asv6306 | CCAAACGTTATTCGGAATTACTGGGCTTAAAGGGTGCGTAGGCGGCGCGGAAAGTCAGATGTGAAATCCCTCGGCTCAACCGAGGAACTGCGTTTGAAACTACCGTGCTCGAGTGAGACAGAGGTAAGCGGAACTTCGGGTGGAGCGGTGAAATGCGTTGATATCCGAAGGAACACCGGTGGCGAAAGCGGCTTACTGGGTCTTTACTGACGCTGATGCACGAAAGCTAGGGG |
| P-asv6307 | GCTAGCGTTGTTCGGAATTACTGGGCGTAAAGCGCGTGTAGGCTGGCTGTCAAGCCAGAGGTGAAATCCCAAGGCTCAACTTTGGAACTGCCTTTGGAACTGTCAGCCTAGAGTATCGGAGGGGATGGTGGAATTCCTAGTGTAGAGGTGAAATTCGTAGATATTAGGAGGAACACCGGTGGCGAAGGCGACCATCTGGACGATTACTGACGCTGAGACGCGAAAGCGTGGGG |
| P-asv6308 | GCAAGCGTTAATCGGAATTACTGGGCGTAAAGCGCGCGTAGGTGGTTTGTTAAGTTGAATGTGAAAGCCCCGGGCTCAACCTGGGAACTGCATCCAAAACTGGCAAGCTAGAGTATGGCAGAGGGTGGTGGAATTTCCTGTGTAGCGGTGAAATGCTTAGATATTGCATGGAATACCAATTGCGAAGGCAGGTTACTAACAAACGATTGACGCTGATGGACGAAAGTGTGGGG |
| P-asv6309 | GCGAGCGTTAATCGGAATTACTGGGCGTAAAGGGTGCGTAGGTGGTCAATTAAGTCAGATGTGAAAGCCCTGGGCTTAACCTAGGAATGGCATTTGATACTGGTTGACTAGAGTTTGGAAGAGGGAAGTGGAATTTCCGGTGTAGCGGTGAAATGCGTAGATATCGGAAGGAACATCAGTGGCGAAGGCGACTTCCTGGTCCGATACTGACACTGAGGCACGAAAGCGTGGGG |
| P-asv6310 | GCGAGCGTTAATCGGAATAATTGGGCGTAAAGCGTGCGTAGGTGGTGTTTTAAGTGGTATGTGAAAGACCGGGGCTTAACCTTGGGGTGTCATATCATACTGGAACGCTAGAGGGCAGTAGAGGAAGGTGGAATTTCCGGTGTAGCGGTGAAATGCGTAGATATCGGGAGGAACATCAGTGGCGAAGGCGACCTTCTGGGCTGCACCTGACACTGAGGCACGAAAGCATGGGG |
| P-asv6311 | GCAAGCGTTGTTCGGAATTACTGGGCGTAAAGCGCGGGCAGGTGGCCTGTTAAGTCTTTTGTGAAAACCCGGGGCTCAACTCCGTGGAATGCAGAAGATACTGGCAGGCTAGAGACCAGTAGAGGCTAGCGGAATTCCTGGTGTAGCGGTGGAATGCGTAGATATCAGGAAGAACACCTGTGGCGAAGGCGGCTAGCTGGGCTGGATCTGACACTCATCCGCGAAAGCGTGGGG |
| P-asv6312 | GCAAGCGTTGTCCGGAATTATTGGGCGTAAAGAGTTCGTAGGCGGTTTACTAAGTCTGATGTTAAAGACCTGAGCTCAACTCAGACATGGCATTGGAAACTGGTAGGCTAGAGTATGGCAGAGGCAAGGGGAATTCCCGGTGTAGCGGTGAAATGCGTAGATATCGGGAGGAACACCAGTGGCGAAAGCGCCTTGCTGGGCCATTACTGACGCTGAGGAACGAAAGCCAGGGT |
| P-asv6313 | GCAAGCGTTGTTCGGAATTACTGGGCGTAAAGGGCGCGTAGGCGGCCTAGCAAGTCAGATGTGAAATCCCGAGGCTTAACTTCGGAAGTGCATTTGAAACTGCAAGGCTCGAGTTCAGTAGAGGTTGGCGGAATACCCAGTGGAGAGGTGAAATTCGTAGATATTGGGTGGAACACCGGTGGCGAAGGCGGCCAACTGGGCTGACACTGACGCTGAGGCGCGAGAGCATGGGT |
| P-asv6314 | GCAAACGTTGTTCGGAATTACTGGGCGTAAAGCGCATGTAGGCGGTCTTGCAGGTCGGATGTGAAATCCCTCGGCTCAACCAAGGAAGTGCATTCGAAACCGCAAGACTTGAGTATCGGAGAGGAAGGCGGAATTGTCGGTGTAGAGGTGAAATTCGTAGATATCGACAGGAACACCTGTGGCGAAAGCGGCCTTCTGGACGAATACTGACGCTGAGACGCGAAAGCGTGGGG |
| P-asv6315 | GCGAGCGTTAATCGGAGTTACTGGGCGTAAAGGGCGCGTAGGCGGTTCTAGTGAGTGTGATGTGAAAGCCCTGGGCTTAACCTAGGAAGTGCATCGCAAACGACTGAACTGGAGTAGAAGAGAGGGTGGCGGAATTTCCGGTGTAGCGGTGAAATGCGTAGAGATCGGAAGGAACGTCAATGGCGAAGGCAGCCACCTGGCTTCATACTGACGCTGAGGCGCGAAAGCGTGGGG |
| P-asv6316 | GCAAGCGTTAATCGGAATTACTGGGCGTAAAGGGCGCGTAGGCGGATAGGTAAGTCAGATGTGAAATTTCGGAGCTCAACTTCGAAGCTGCATTTGAAACTACCAATCTGGAGGATAGGCGGAGAAAACGGAATTCCACGTGTAGCGGTGAAATGCGTAGAGATGTGGAAGAACACCTGTGGCGAAAGCGGTTTTCTAGCTTATTCCTGACGCTGAGGCGCGAAAGCAAGGGG |
| P-asv6317 | GCGAGCGTTGTTCGGATTTACTGGGCGTAAAGGGTGCGTAGGCGGTTAAGTAAGTTGGATGTGAAATCCCATGGCCCAACCATGGACGTGCATTTGATACTGCTTGACTAGAGTACAGGAGGGGAGAGCGGAATTCTTGGTGTAGCGGTGAAATGCGTTGATATCAAGAAGAACACCGGTGGCGAAGGCGGCTCTCTGGAATGTTACTGACGCTGAGGCACGAAAGCTGGGGG |
| P-asv6318 | GCTAGCGTTGTTCGGAATTACTGGGCGTAAAGCGTGCGCAGGCGGCTATGCAAGTCAGGGGTGAAATGCCAAGGCTCAACCTTGGAACTGCCTTTGAAACTGCATAGCTAGAGACTCGGAGGGGTTAGGGGAATTCCTAATGTAGCGGTGAAATGCGTAGATATTAGGAGGAGCACCAGTGGCGAAGGCGCCTAACTGGACGAGTACTGACGCTCATGCACGAAAGCGTGGGG |
| P-asv6319 | GCAAGCGTTACTCGGAATTACTGGGCGTAAAGCGTCTGTAGGCGACGTCGTAAGTCTGGCGTTAAATCGCGGAGCTCAACTCCGCAACCGCGCTGGAAACTGCGATGATAGAGGGACTCAGAGGCATCTGGAATGCCGTGTGTAGGGGTAAAATCCGTTGATCCACGGTAGAACGCCAAAAGTGAAGACAGGATGCCGGGAGTCACCTGACGCTGAGAGACGAAAGCGTGGGG |
| P-asv6320 | GCAAGCGTTGTTCGGAATTACTGGGCGTAAAGCGTACGTAGGCGGTGAGTTAAGTCGAATGTTAAAGCCCGAGGCTCAACCTCGTGGATGCATTCGATACTGGCTCGCTAGAGGGTGATAGAGGAAAGTGGAATTCCGAGTGTAGCGGTGAAATGCGTAGATATTCGGAGGAACATCAGTAGCGAAGGCGACTTTCTGGGTCACACCTGACGCTGAGGTACGAAAGCGTGGGG |
| P-asv6321 | GCGAGCGTTGTTCGGAATCACTGGGCGTAAAGCGTGCGTAGGCGGTTTTGTAAGACAGAAGTGAAATCCTGGAGCTTAACTCCAGAATTGCTTTTGTGACTGCAAAGCTTGAATCCAGGAGAGGTTGGGGGAATTCCGAGTGTAGAGGTGAAATTCGCAGATATTCGGAGGAACACCAGTGGCGTAGGCGCCCAACTGGACTGGTATTGACGCTGAGGCACGAAAGCGTGGGG |
| P-asv6322 | GCAAGCGTTATCCGGATTTATTGGGTTTAAAGGGTGCGCAGGCGGAATTTTAAGTCAGTGGTGAAAGCCCAAGGCTCAACCTTGGAACTGCCATTGATACTGAAATTCTTGAGTGTGGTTGATGTGGGCGGAATGTGCCATGTAGCGGTGAAATGCTTAGATATGGCACAGAACACCGATCGTGAGGACAGCTCACAAAACCATTACTGACGCTCAGGCACGAAAGCGTGGGG |

| | |
|---|---|
| P-asv6323 | CCAAGCGTTGTTCGGAATTACTGGGCGTAAAGGGAGCGCAGGCGGACAGGTAAGTCTAAAGTGGAAGCCCGGGGCTCAACCTCGGAAGGTCTTTGGATACTGCCTGTCTTGAGTTTGTCAGAGGAAAGCGGAATTCAGGGTGTAGCGGTGAAATGCTTAGATATCCTGAGGAACACCAATGGCGAAAGCAGCTTTCTGGGGCAAAACTGACGCTGAGGCTCGAAAGCGTGGGT |
| P-asv6324 | GCGAGCGTTGTTCGGAATAACTGGGCTTAAAGCGTGCGTAGGCGGATATGCAAGTCAGAGGTGAAATCCAGTCGCTTAACGATTGAACTGCCTTTGATACTGGTTGACTTGAGTCATATGGAAGTAGATAGAATGTGTAGTGTAGCGGTGAAATGCTTAGATATTACACAGAATACCGATTGCGAAGGCAGTCTACTACGTATGTACTGACGCTCATGGACGAAAGCGTGGGG |
| P-asv6325 | CCAAGCGTTATCCGGAATTACTGGGTATAAAGGGTGCGTAGGCGGTCTTGTGCGTCGAAGGTTAAAGGCCATGGCTCAACCATGGAAATGCCCTCGATACGGCAAGACTTGAGTATGAGAGAGGAAGATGGAATTCCTGGTGTAGCAGTGAAATGCGTAGATATCAGGAGGAACACCGGTGGCGAAGGCGGTCTTCTGGCTCATTACTGACGCTGAGGCACGAAAGTGTGGGG |
| P-asv6326 | GCAAGCGTTGTTCGGAATCATTGGGCGTAAAGCGCGCGTAGGCGGTCTTGTAAGTCTGGAGTGAAAGCCCTCGGCCTAACCGAGGAAGTGCTTCGGATACTGCGAGACTTGAATCCCGGAGGGGCGGGTGGAATTCCCAGTGTAGAGGTGAAATTCGTAGATATTGGGAGGAACACCAGTGGCGAAGGCGGCCCGCTGGACGGAGATTGACGCTGAGGCGCGAAAGCGTGGGG |
| P-asv6327 | CCAAGCGTTGTCCGGATTGACTGGGCGTAAAGCGCACGCAGGCGGTGGGGCACGTGGGGTGTGAAAGCGGCCCGCTTAACGGGGCGAGGCCATCCCATACGGCCCGACTCGAGAGTGGCAGAGGTTCGTGGAATTGACGGTGTAGTGGTGAAATGCGTAGAGATCGGCAGGAACACCAAGGGGGAAACCAGCGAACTGGGCCATATCTGACGCTCAGGTGCGAAAGCGTGGGG |
| P-asv6328 | GCAAGCGTTACACGGAATCACTGGGCGTAAAGGGTGCGTAGGCTGGCTTTTAAGTCAGATGTGAAATCTCGGGGCTTAACCCCGAAACTGCGTCTGAAACTGGTTGTCTAGAGTTTCATAGGGGGCAGGGGAATTTCACGTGTAGGGGTAAAATCCGTAGAGATGTGAAGGAACACCGGAGGCGAAGGCGCCTGCCTGGATGAATACTGACGCTGAGGCGCGAAAGCGTGGGG |
| P-asv6329 | GCGAGCGTTGCTCGGATTTATTGGGTGTAAAGGGTCTGTAGGCGGTCCCGTAAGTGCCCGGTGAAATTCTGGGGCTCAACCCCAGAGCGCCCGGGTAGACTGCGGGACTTGAGGGCTGGAGAGGCAAGCGGAATTCCCGGTGTAAGAGTGAAATCTGTAGATATCGGGAGGAACACCAGTGGCGAAGGCGGCTTGCTGGACAGCACCTGACGCTGAGAGACGAAAGCGTGGGG |
| P-asv6330 | GCTAGCGTTGCTCGGAATTACTGGGCGTAAAGGGAGCGTAGGCGGACTGTTTAGTCAGAGGTGAAAGCCCAGGGCTCAACCTTGGGATTGCCTTTGTGACTGCATAGCTAGAGTACGGTAGAGGGGGATGGAATTCCGCGTGTAGCAGTGAAATGCGTAGATATGCGGAGGAACACCGATGGCGAAGGCAATCCCCTGGACCTGTACTGACGCTCATGCACGAAAGCGTGGGG |
| P-asv6331 | GCGAACGTTGTTCGGAATCACTGGGCTTAAAGGGCGCGTAGGCGGGGTCGTAAGTCTGTGGTGAAATCCTCCGGCTCAACCGGAGAACTGCCTCGGATACTGCGGCTCTCGAGGGTGATAGGGGTATGCGGAACTATCGGTGAAGCGGTGAGATGCGTTGATATCGATAGGAACTCCGGTGGCGAAGGCGGCATACTGGATCACTTCTGACGCTGAGGCGCGAAAGCCAGGGG |
| P-asv6332 | GCTAGCGTTGTTCGGAATTACTGGGCGTAAAGCGCGCGTAGGCGGGTTTTTAAGTCAGATGTGAAATCCCGGGGCTCAACCTCGGAACTGCATTTGAAACTGGATACCTAGAGTCCGGGAGAGGTTCGTGGAATACCGAGTGTAGAGGTGAAATTCGTAGATATTCGGTGGAACACCAGTGGCGAAGGCGACGAACTGGACCGGTACTGACGCTGAGGTGCGAAAGTGTGGGG |
| P-asv6333 | GCAAGCGTTATCCGGATTCACTGGGTTTAAAGGGTGTGTAGGCGGGCGTTTAAGTCAGTGGTGAAATCTCCAGGCTCAACCTGGAAACTGCCATTGATACTATTCGTCTTGAATATCGTTGAGGTAGGCGGAATATATCATGTAGCGGTGAAATGCTTAGATATGATATAGAACACCGATTGCGAAGGCAGCTTGCTAAACGATGATTGACGCTGAGGCACGAAAGCGTGGGG |
| P-asv6334 | GCAAGCGTTAATCGGAATCACTGGGCGTAAAGCGTACGTAGGCGGCCATGCAAGTCAGACGTGAAAGCCCTCGGCTCAACCGAGGAATTGCGTTTGAAACTGCATGGCTCTGAGTTCGGGAGAGGGTAGCGGAATTCCCGGTGTAGGAGTGAAATCCGTAGATATCGGGAGGAACACCAGTGGCGAAGGCGGCTACCTGGACCGATACTGACGCTGAGGTATGAAAGCGTGGGG |
| P-asv6335 | GCGAGCGTTAATCGGAATTACTGGGCGTAAAGCGCATGCAGGTGGTTTGTTAAGTCAGATGTGAAATCCCCGAGCTTAACTTGGGACGTGCATTTGGAACTGGGTGACTAGAGTGTGTCAGAGGGAGGTAGAATTCCACATGTAGCGGTGGAATGCGTAGAGATGTGGAGGAATACCGATGGCGAAGGCAGCCTCCTGGGATAACACTGACGTTGAGGCTCGAAAGCGTGGGG |
| P-asv6336 | GCTAGCGTTGCTCGGAATCACTGGGCGTAAAGGGTGCGTAGGTGGATCTTTAAGTCAGGGGTGAAATCCTGGAGCTCAACCTGGGAATCGCATTGGTGACTGCACGGCTAGAGTGTGTCAGAGGGGGGTAGAATTCCACGTGTAGCAGTGAAATGCGTAGAGATGTGGAGGAATACCGATGGCGAAGGCAGCCCCCTGGGATAACACTGACGCTCATGCACGAAAGCGTGGGG |
| P-asv6337 | GCTAGCGTTGTTCGGAATCACTGGGCGTAAAGCGCACGTAGGCGGACTTTTAAGTCTGAGGTGAAAGCCTGGAGCTCAACTTCAGAACTGCCTTTGATACAGCATATCTCGAGTCCGAGAGAGGTGAGTGGAATTCCTAGTGTAGAGGTGAAATTCGTAGATATTAGGAAGAACACCGGTGGCGAAGGCGGCTCACTGGCTCGGTACTGACGCTGAGGTGCGAAAGCGTGGGG |
| P-asv6338 | GCAAGCGTTGTCCGGATTTATTGGGTTTAAAGGGTGCGTAGGCGGCCCGTTAAGTCAGCGGTGAAATCCCAGGGCTTAACCCTGGAACTGCCTGTGAGACCTCAGTACTAGAGTGTGAGAGGGGATAGTGGAATACCCAGTGTAGCGGTGAAATGCGTAGAGATTGGGTGGAACACCGGTGGCGAAGGCGGCTATCTGGCTCACAACTGACGATCAGGCACGAAAGCGTGGGG |
| P-asv6339 | GCAAGCGTTGTCCGGAATGACTGGGCGTAAAGGGTACGCAGGCGGATGAGTAAGTCAAATGTGAAAGGCACCGGCTCAACCGGTGAGCTGCATTTGAAACTGCCTATCTTGGGTTCAGGAGAGGAGAGCGGAATTCCTGGTGTAGCGGTGGAATGCGTAGATATCAGGAAGAACACCAGTGGCGAAGGCGGCTCTCTGGCCTGAAACTGACGCTCATGTACGAAAGCTGGGGG |
| P-asv6340 | GCGAGCGTTGTTCGGAATCACTGGGCGTAAAGGGTCCGTAGGCGGCCATGCAGGTCCTGGGTGAAATCCCTCGGCTTAACTGAGGAATTGCCTGGGAAACCGCATGGCTTGAGTTCGGTAGAGGGAGATGGAATTCCGGGTGTAGCGGTGAAATGCGTAGATATCCGGAGGAACACCAGTGGCGAAGGCGATCTCCTGGGCCGATACTGACGCTGAGGGACGAAAGCTAGGGG |
| P-asv6341 | GCAAGCATTAATCGGATTTATTGGGCGTAAAGGGCGCGTAGGCGGGTCTTTAAGTCAGATGTGAAATCCCGAGGCTCAACCTCGGAACAGCATTTGAAACTGAAGATCTTGAGGGTAGAAGGAGAGAATGGAATTCCACAAGTAGCGGTGAAATGCGTAGATATGTGGAAGAACACCGGTGGCGAAGGCGATTCTCTATTTTATTCCTGACGCTGAGGCGCGAAAGCGTGGGG |
| P-asv6342 | GCAAGCGTTAATCGGAATCACTGGGCGTAAAGGGTGCGTAGGCGGTACGTTAGGTCTGTCGTGAAGTCCCCGGGCTCAACCTGGGAACGGCATCCGGAACTATCAGGCTAGAGTGCAGGAGAGGAAGGTAGAATTCCCGGTGTAGCGGTGAAATGCGTAGAGATCGGGAGGAATACCAGTGGCGAAGGCGGCCTTCTGGACTGACACTGACACTGAGGTGCGAAAGCGTGGGT |
| P-asv6343 | GCTAGCGTTGCTCGGAATTACTGGGCGTAAAGGGCGCGTAGGCGGACAGTTAAGTTGGGGGTGAAGGCCCGGGGCTCAACCTCGGAACTGCATTCGATACTGTATGACTAGAGTTCGGTAGAGGGCGGCGGAATTTCCGGTGTAGCGGTGAAATGCGTAGATATCGGAAGGAACATCAATGGCGAAGGCAGCCGCCTGGGCCTGAACTGACACTGAGGCACGAAAGCGTGGGT |
| P-asv6344 | GCAAGCGTTATCTGGAATTACTGGGCATAAAGAGTTCGTAGGCGGTTAATCAAGCCAGCTAGTGAAAGCCATCGGCCTAACCGATGTCAGCTAATGGAACTGGTTAACTTGAGTACAGGAGAGGAAAGTGGAATTCCTAGTGTAGTAGTGAAATGCGTAGATATTAGGAAGAACATCAGTAGCGAAGGCGACTTTCTGGACTGTAACTGACGCTGAGGAACGAAAGCGTGGGG |
| P-asv6345 | CCAAGCGTTGTTCGGATCCACTGGGCGTAAAGGGAGCGTAGGCTGTGCGGTAAGTCAGATGTGAAATCTCGAGGCTCAACCTCGAAACTGCATCCGATACTGCCGTACTAGAGGATTGGAGAGGTAACAGGAATTCACGGTGTAGCAGTGAAATGCGTAGATATCGTGAGGAAGACCAATGGCGAAAGCATGTTACTGGACAACTCCTGACGCTGAGGCTCGAAGGCTAGGGT |
| P-asv6346 | GCGAGCGTTGTTCGGAATTATTGGGCGTAAAGGGCTTGTAGGCGGCCCGCTAAGTCTGGTGTGAAATCCCGCAGCTTAACTGCGGATCCGCATTGGATACTGGCGAGCTGGAGTCCGTGAGAGGAAACTGGAATTCCTGGTGTAGGGGTGAAATCTGCTGATATCAGGAAGAACACCTGAGGCGAAGGCGAGTTTCTGGCACAGGACTGACGCTAATGAGCGAAAGCGTAGGG |
| P-asv6347 | GCGAGCGTTAATCGGAATTACTGGGCGTAAAGCGCGCGTAGACGGTTTGGTAAGTCAGATGTGAAATCCCCGGGCTCAACCTGGGAACTGCATTTGATACTGCCTGGCTAGAGTATCGAAGAGGGAAGTGGAATTTCCGGTGTAGCGGTGAAATGCGTAGATATCGGAAGGAACACCAGTGGCGAAAGCGACTTCCTGGTCGAATACTGACGTTCATGTGCGAAAGCGTGGGG |
| P-asv6348 | GCTAGCGTTGTTCGGATTTACTGGGCGTAAAGCGTATGTAGGTGGTTTGTTTGGTCAGATGTGAAATCCCAGGGCTCAACCCTGGAACTGCATTTGATACCGGCAAGCTAGAGTATAGTAGAGGACAATGGAATTCTAAGTGTAGAGGTGAAATTCGTAGATATTTAGAGGAACACCAGTGGCGAAAGCGATTGTCTGGACTATTACTGACACTGAGATACGAAAGCGTGGGG |
| P-asv6349 | GCAAGCGTTAATCGGAATTACTGGGCGTAAAGCGTACGTAGGCGGCTGCCTAAGTTGGATGTGAAAGCCCCGGGCTTAACCCGGGAACTGCATCCAAAACTGGGTAGCTAGAGTGCGGAAGAGGAGTGTGGAGTTTCCTGTGTAGCGGTGAAATGCGTAGATATAGGAAGGAACACCAGTGGCGAAGGCGACACTCTGGTCTGACACTGACGCTGAGGTACGAAAGCGTGGGG |
| P-asv6350 | TCGAGTGATGGCCATTCTTATTGGGCCTAAAGCGTCCGTAGCTGGCCACGCAAGTTCGTCGGGAAATCCACATGCTCAACATGTGGGCGTCCGGCGAAACCTGCGTGGCTTGGGACCAGACGACCCAGAGGGTACGGCCGGGGTAGGAGTGAAATCCTGTAATCCCGGCCGGACCACCAGTGGCGAAAGCGCTCTGGGAGGATGGATCCGACGGTGAGGGACGAAAGCCAGGGT |
| P-asv6351 | GCAAGCGTTAATCGGATTTACTGGGCGTAAAGGGCGCGTAGGCGGTCGTATGAGTGTGATGTGAAAGCCCCGGGCTTAACCTGGGAAGTGCATCGCAAACGATACGACTGGAGTACATGAGAGGGTGGCGGAATTTCCGGTGTAGCGGTGAAATGCGTAGAGATCGGAAGGAACGTCGATGGCGAAGGCAGCCACCTGGCATGATACTGACGCTGAGGCGCGAAAGCGTGGGG |
| P-asv6352 | GCAAGCGTTGTTCGGAATTATTGGGCGTAAAGGGTGCGTAGGCGGTTTGACAAGTCTTATGTGAAATCTTCGGGCTCAACCCGAAGTCTGCATGAGAAACTGTTGAGCTTGAGTATGGGAGAGGTGAGTGGAATTTCCGGTGTAGCGGTGAAATGCGTAGATATCGGAAGGAACACCTGTGGCGAAAGCGGCTCACTGGACCATAACTGACGCTGAGGCACGAAAGCTAGGGG |
| P-asv6353 | GCAAGCGTTAATCGGAATCACTGGGCGTAAAGGGTGCGTAGGCGGGACGTTAAGTCAGTGGTGAAATCTCTCAGCTCAACTGAGAAACTGCCTCTGATACTGATGTTCTTGAGTTTGGCTGAAGTGGGCGGAATGAGGCATGTAGCGGTGAAATGCATAGATATGCCTCAGAACACCAATTGCGAAGGCAGCTCACTAAACCAATACTGACGCTGAGGCACGAAAGCGTGGGG |
| P-asv6354 | GCAAGCGTTAATCGGAATTACTGGGCGTAAAGCGCGCGTAGGTGGCTTGATCAGTTGGATGTGAAATCCCCGGGCTCAACCTGGGAACTGCATCCAATACTGGCTGTCTTGAGTACGGGAGAGGTGAGTGGAACTCCGAGTGTAGAGGTGAAATTCGTAGATATTCGGAAGAACACCAGTGGCGAAGGCGACTCACTGGCCCGTTACTGACGCTGAGGCGCGAAAGCGTGGGG |
| P-asv6355 | GCAAGCGTTGTCCGGATTTATTGGGCGTAAAGCGCGTGTAGGCGGCCATGCAGGTCCGCTGTGAAAACTGGAGGCTTAACCTCCAGAGGTCGGTGGAAACCGCATGGCTAGAGTCCGGAAGAGGGGACTGGAATTCCCGGTGTAGCGGTGGAATGCGCAGATATCGGGAGGAACACCCGTGGCGAAGGCGGGTCCCTGGGACGGTACTGACGCTGAGACGCGAAAGCGTGGGG |
| P-asv6356 | GCAAGCGTTGTTCGGAATTACTGGGCGTAAAGGGCGCGTAGGCGGCTTATCTAGTCAGGCGTGAAATTCCCGGGCTCCACCTGGGGGCTGCGCTTGATACTGATGAGCTTGAATGCGGGAGAGGATAGTGGAATTCCCAGTGTAGAGGTGAAATTCGTAGATATTGGGAAGAACACCGGTGGCGAAGGCGGCTATCTGGCCCGTGATTGACGCTGAGGCGCGAAAGCGTGGGG |
| P-asv6357 | GCTAGCGTTGTTCGGAATCACTGGGCGTAAAGCGCACGTAGGTGGACTTTTAAGTCAGAGGTGAAAGCCTGGAGCTCAACCTCGGAAATGCCTTCAATACTGGCTGTCTTGAGTACGGGAGAGGTGAGTGGAACTCCGAGTGTAGAGGTGAAATTCGTAGATATTCGGAAGAACACCAGTGGCGAAGGCGACTCACTGGCCCGTTACTGACGCTGAGGCGCGAAAGCGTGGGG |
| P-asv6358 | GCAAGCGTTATCCGGAATCATTGGGTTTAAAGGGTCCGCAGGCGGACATATAAGTCAGTGGTGAAATCCCATCGCTTAACGATGGAACTGCCATTGATACTGTATGTCTTGAGTTCGGTCGAAGTGGGCGGAATATGACATGTAGCGGTGAAATGCTTAGATATGTCATAGAACACCGATAGCGAAGGCAGCTCACTAGGCCTGGATTGACGCTCAGGGACGAAAGCGTGGGT |
| P-asv6359 | GCAAGCGTTAATCGGAATTACTGGGCGTAAAGTGCGCGTAGGCGGCTAGGTAAGTCGGATGTGAAAGCCCTGGGCTCAACCTAGGAATTGCATTCGAAACTGCTTGGCTAGAGTGCAGTAGAGGGGAGTGGAATTCCCGGTGTAGCGGTGAAATGCGTAGATATCGGGAGGAACACCAGTGGCGAAGGCGACTCCCTGGACTGACACTGACGCTGACAGCGCGAAAGCGTGGGG |
| P-asv6360 | GCAAGCGTTACTCGGAATCACTGGGCGTAAAGGGTGCGTAGGCTGGCTTCTAGGTCAGATGTGAAATCCAATGGCTTAACCATTGAACTGCATTTGAAACTGGGAGCCTAGAGTTCAGAAGGGGCAGATGGAATTAGTGGTGTAGGGGTAAAATCCGTAGATATCACTAGGAATATCAAAAGCGAAGGCGATCTGCTGGGATGATACTGACGCTGAGGCACGAAAGCGTGGGG |
| P-asv6361 | GCGAGCGTTAATCGGAATTACTGGGCGTAAAGCGCACGTAGGTGGCGACGTATGTCGGGTGTGAAAGCCCCGGGCTCAACCTGGGAATTGCATCCGAAACTGCGTTGCTGGAGTACGGAAGAGGAAGGCGGAATTCCCGGTGTAGCGGCGAAATGCGTAGATATCGGGAGGAACATCAGTGGCGAAGGCGGCCTTCTGGTCCAGTACTGACGCTCAAGTGCGAAAGCGTGGGG |

| | |
|---|---|
| P-asv6362 | GCGAGCGTTGTCCGGATTTATTGGGCGTAAAGGGCTCGTAGGCGGTTGATCGCGTCGGAAGTGGAAACTTGATGCTTAACGTTGAGCGTGCTTTCGATACGGGTTGACTTGAGGAAGGTAGGGGAGAATGGAATTCCTGGTGGAGCGGTGGAATGCGCAGATATCAGAAGGAACACCAGTGGCGAAGGCGGCTCACTGGCTCGATACTGACGCTGAGGTGCGAAAGTGTGGGG |
| P-asv6363 | GCAAGCGTTATTCGGAATTATTGGGCGTAAAGGGCGCGTAGGCGGCCTGTTAAGTCATCTGTGAAATTTCGGGGCTTAACCCCGGAGCTGCATGTGATACTGGCAGGCTTGAGTATGGCAGAGGAAAGCGGAATTCCTGGTGTAGCGGTGAAATGCGTAGATATCAGGAGGAACACCGGTGGCGAAGGCGGCTTTCTGGACCAATACTGACGCTGATGCGCGAAGGCGTGGGT |
| P-asv6364 | GCAAGCGTTACTCGGATTTACTGGGCGTAAAGGGTCCGCAGGTGGCTATGTGTGTCAGGTGTGAAATCCCACGGCTCAACCGTGGAACTGCGCCTGAAACTACATGGCTAGAGTATTGGAGAGGGTAGTGGAATTCCTAGTGTAGCGGTGAAATGCGTAGATATTAGGAGGAACACCAACGGCGAAGGCAGCTACCTGGACAATAACTGACACTCATGGACGAAAGCGTGGGG |
| P-asv6365 | GCGAGCGTTGTTCGGAATTATTGGGCGTAAAGGGCGCGTAGGCGGATTGTCAAGTCTGGTGTGAAAGACCGCAGCTTAACTGCGGGGGCGCTCTGGAAACTGACGATCTTGAATCCTGGAGGGGGAGCTGGAATTCCTGGTGTAGGGGTGAAATCTGTAGATATCAGGAAGAACACCGGTGGCGAAGGCGAGCTCCTGGCCGTGGATTGACGCTGAGGCGCGAAAGCGTGGGG |
| P-asv6366 | GCAAGCGTTGTCCGGAATCATTGGGCGTAAAGCGCGTGTAGGCGGCCACGTTAGTCCGCTCTGAAAGCCCGGGGCTCAGCCCCGGGAGGCGGGTGGATACTGCGTGGCTCGAGTCCGGAAGAGGCGAGTGGAATTCCTGGTGTAGCGGTGAAATGCGCAGATATCAGGAGGAACACCAATGGCGAAGGCAGCTCGCTGGGACGTGACTGACGCTGAGACGCGAAAGCGTGGGG |
| P-asv6367 | CCGAGCGTTATTCGGAATCACTGGGCGTAAAGGGAGCGTAGGCTGTATGGTAAGTCAGATGTGAAATCTCAGAGCTCAACTCTGAAACTGCATCCGATACTGCCGTACTAGAGTAATGGAGGGGTAACTGGAATTCTCGGTGTAGCAGTGAAATGCGTAGATATCGAGAGGAAGACCAATGGCGAAGGCAGGTTACTGGACATTTACTGACGCTGATGCTCGAAGGCTAGGGT |
| P-asv6368 | GCGAGCGTTATCCGGATTCATTGGGTTTAAAGGGTGCGTAGGCGGTTTCTTAAGTCAGTGGTGAAATCCTGCGGCTCAACCGTAGAACTGCCATTGATACTGAGGAACTTGAATTTAGTTGAGGTAGGCGGAATGAGTAGTGTAGCGGTGAAATGCTTAGATATTACTCAGAACACCGATTGCGAAGGCAGCTTACCAAGCTACGATTGACGCTGATGCACGAAAGCGTGGGG |
| P-asv6369 | TCGAGTGATGGTCGATCTTATTGGGCATAAAGCGTTCGTAGCTGGCCAGTCAAGTCTGTTGGGAAATCCACGTACCCAACACGTGGGCGTCCAACGGAAACTGACTGGCTTGGGACCGGAAGACCCGAAGAGTACGTCTGGGGTAGGAGTGAAATCCCGTAATCCTAGACGGACTACCGATGGCGAAAGCACTTCGGGAGTACGGATCCGACAGTGAGGAACGAAAGCTGGGGT |
| P-asv6370 | CCAAGCGTTAATCGGAATCACTGGGCGTAAAGAGTACGTAGGCGGTTTGGTAAGTCAGGTGTGAAATCCCGGAGCTCAACTCCGGAACTGCATCCGATACTGCTAGACTAGAGTAATGGAGAGGCAGGTGGAATTCTCGGTGTAGCAGTGAAATGCGTGGATATCGAGAGGAAGACCAATGGCGAAGGCAGCCTGCTGGACATTTACTGACGCTGAGGTACGAAGGCCAGGGT |
| P-asv6371 | GCAAGCGTTATTCGGACTTATTGGGTTTAAAGGGTCCGTAGGCGGGTTATTAAGTCAGTGGTGAAATCTCATAGCTCAACTATGAAACTGCCATTGATACTGATAGTCTTGAGTATATTTGAAGTAGCTGGAATAAGTAGTGTAGCGGTGAAATGCATAGATATTACTTAGAACACCAATAGCGAAGGCAGGTTACTAAGATATTACTGACGCTGAGGGACGAGAGCGTAGGG |
| P-asv6372 | GCAAGCGTTACTCGGATTCACTGGGCGTAAAGGGCACGTAGGCTGTTCCATAAGCCCTGCCTGAAAGGCTCTGGCTCAACCAGAGCAGTGGGCAGGGGACTGCGGAACTTGAGGATCAGAGAGGCGAGCGGAATTCCCGGTGTAAGGGTGAAATCTGTAGAGATCGGGAGGAACACCAGTGGCGAAAGCGGCTCGCTGGCTGATTTCTGACGCTGAGGTGCGAAAGCTAGGGG |
| P-asv6373 | CCAAGCGTTATCCGGATTCACTGGGTTTAAAGGGTGCGCAGGCGGTCAGGTAAGTCAGTGGTGAAAGCCTTCGGCTCAACCGAAGAATTGCCATTGAAACTACTTGACTTGAGTGTAGTTGAGGTGGGCGGAATGTAGCATGTAGCGGTGAAATGCATAGATATGCTACAGAACACCGATAGCGAAAGCGGCTCACCAGACTACGACTGACGCTGAGGCACGAAAGCATGGGT |
| P-asv6374 | GCTAGCGTTGTTCGGAATTACTGGGCGTAAAGGGCGCGTAGGCGGCCGCTTGCGTCGGATGTGAAAGCCCGGGGCTCAACCTCGGAACGGCATCCGATACGGGGCGGCTGGAGACCGGGAGAGGAGGGTGGAATAGCCAGTGTAGAGGTGAAATTCGTAGATATTGGCTAGAACACCGGTGGCGAAAGCGGCTCTCTGGCCCGGTTCTGACGCTGAGGCGCGAAAGCGTGGGG |
| P-asv6375 | TCAAGCGTTGTTCGGATTCATTGGGCGTAAAGGGTGCGTAGGCGGCGAGGCAAGTCTGGTGTGAAATCCCGGGGCTCAACCCCGGAACTGCACTGGATACTGCTTTGCTAGAGTACTGGAGAGGGAACTGGAATTTACGGTGTAGCAGTGAAATGCGTAGATATCGTAAGGAAGACCAGCGGCGAAGGCGAGTTCCTGGACAGTTACTGACGCTGAGGCACGAAGGCCAGGGG |
| P-asv6376 | GCAAGCGTTAATCGGAATTACTGGGCGTAAAGCGCGTGTAGGTGGTTAGTTAAGTTGGAAGTGAAATCCTATAGCTTAACTATAGAACTGCTTCCAAAACTGTTTACCTAGAGTATGGGAGAGGTAGATGGAATTTCTGGTGTAGGGGTAAAATCCGTAGAGATCAGAAGGAATACCGATTGCGAAGGCGATCTACTGGAACATTACTGACGCTCAGATGCGAAAGCGTGGGG |
| P-asv6377 | GCAAGCGTTATTCGGAATTACTGGGCGTAAAGCGCACGTAGGCGGCTTTGTAAGTTAGAGGTGAAAGCCTGGAGCTCAACTCCAGAACTGCTTTTGAAACTGGATGGCTAGAGTCCCGGAGAGGTTGGCGGAATTCCGAGTGTAGAGGTGAAATTCGCAGATATTCGGAGGAACACCAGTGGCGTAGGCGGCCAACTGGACGGGAACTGACGCTGAGGTACGAAAGCGTGGGG |
| P-asv6378 | CCAAGCGTTATCCGGAATCATTGGGTTTAAAGGGTCCGTAGGCGGAATAATAAGTCAGTGGTGAAAGTCTGCAGCTTAACTGTAGAATTGCCATTGATACTGTTATTCTTGAATTAGTATGAAGTGGTTGGAATGAGTAGTGTAGCGGTGAAATGCATAGATATTACTCGGAACACCTATTGCGAAGGCAGATCACTAATACTATATTGACGCTGAGGGACGAAAGCGTAGGT |
| P-asv6379 | GCAAGCGTTGTCCGGAGTTACTGGGCGTAAAGAGCTCGTAGGCGGTTTGTCGCGTCGGCCGTGAAATCCTGCTGCTCAACGGCAGGCGTGCGGTCGATACGGGCAGACTTGAGTTCGGCAGGGGAGACTGGAATTCCTGGTGTAGCGGTGAAATGCGCAGATATCAGGAGGAACACCGGTGGCGAAGGCGGGTCTCTGGGCCGATACTGACGCTGAGGAGCGAAAGCGTGGGG |
| P-asv6380 | CCTAGCGTTGTTCGGAATTATTGGGCGTAAGGCGGATGTAGGTGGCTTCGTAAGTCAGGTGTGAAAGCCTGGGGCTCAACCCCAGAAGTGCATTTGATACTGCGAAGCTTGAGTGTCGGAGAGGTTACTAGAATTGTTGGTGTAGTGGTGAAATACGTAGATATCAACAGGAATACCGGAGGCGAAGGCGGGTAACTGGCCGAACACTGACACTGAGATCCGAAAGCGTGGGG |
| P-asv6381 | GCGAGCGTTAATCGGAATTACTGGGCGTAAAGCGCGCGTAGGTGGTTAATTAAGTCAGATGTGAAATCCCAGGGCTCAACCTTGGAACTGCATTTGAAACTGGTTAACTAGAGTTTTGTAGTGGGTGGTAGAATTTCAGGTGTAGCGGTGAAATGCGTAGAGATCTGAAGGAATACCAGTGGCGAAGGCGGCCACCTGGACAAAGACTGACACTGAGGCGCGAAGGCGTGGGG |
| P-asv6382 | GCGAGCGTTAATCGGAATTACTGGGCGTAAAGCGTGCGCAGGCGGTTTTGTAAGCCAGATGTGAAATCCCCGGGCTTAACCTGGGAATGGCATTTGGGACTGCAAGGCTTGAGTATGGCAGAGGGGACTGGAATTCCTGGTGTAGCAGTGAAATGCGTAGATATCAGGAGGAATACCGATGGCGAAGGCAGGTCCCTGGGCTGATACTGACGCTCATGCACGAAAGCGTGGGG |
| P-asv6383 | GCTAGCGTTGCTCGGAATGACTGGGCGTAAAGGGCGCGTAGGCGGCACACACAGTCAGGCGTGAAATTCCCGGGCTTAACCTGGGGACTGCGCTTGATACATGTGAGCTAGAGGATGGAAGAGGCTCGTGGAATTCCCAGTGTAGAGGTGAAATTCGTAGATATTGGGAAGAACACCGGTGGCGAAGGCGGCGAGCTGGTCCATTACTGACGCTGAGGCGCGAAAGCGTGGGG |
| P-asv6384 | GCAAGCGTTATCCGGAATGATTGGGCCTAAAGCGTCCGTAGCTGGCCGTGCAAGCCCGTCGGGAAATCCACCCGCTCAACGGGTGGGCGGCCGACGGAAACTGGACGGCTTGGGACCGGGAGGCTTGAAGGGTACGTCCGGGGTAGGAGTGAAATCCCGTAATCCTGGACGGACCACCGATGGGGAAACCACTTCAAGAGAACGGATCCGACAGTGAGGGACGAAAGCTAGGGT |
| P-asv6385 | GCGAGCGTTGTTCGGAATTATTGGGCGTAAAGGGTCCGCAGGAGGTTCGTTAAAATTGGGGTGAAATCCCGGAGCTCAACTCCGGAACTGCCCTGATGACTGACGGACTAGAGTGCTGGAGGGGTAAGCGGAATTTCAGGTGTAGCGGTGGAATGCGTAGATATCTGAAAGAACACCGATGGCGAAGGCAGCTTACTGGACAGCAACTGACTCTCAGGGACGAAAGCGTGGGG |
| P-asv6386 | CCAAGCGTTGTCCGGAGTTACTGGGCGTAAAGGGTGCGCAGGCGGTGGGTTACGTCGTGTGTGAAAGCCCCCGGCTTAACCGGGGAGGGTCGCGCGAAACGGGCTCACTGGAGGCAAGGCGAGGTCGGTGGAATTCCCGGTGTAGTGGTGAAATGCGTAGATATCGGGAGGAACACCAGAGGCGAAGGCGGCCGACTGGACTTGACCTGACGCTGAGGCACGAAGGCGTGGGG |
| P-asv6387 | GCGAGCGTTATCCGGAATCACTGGGTTTAAAGGGTGCGTAGGCGGCCCGTTAAGTCAGGGGTGAAAGCCCGCGGCTCAACCGCGGAACTGCCCTTGATACTGGGGGGCTCGAATTGGGTCGAGGCAGGCGGAATGTGGCATGTAGCGGTGAAATGCTTAGATATGCCATAGAACACCGATTGCGAAGGCAGCCTGCCAGGCCTTGATTGACGCTGAGGCACGAAAGCGTGGGG |
| P-asv6388 | GCAAGCGTTATTCGGATTTACTGGGCGTAAAGCGCGCGTAGGTGGTTTAGTAAGTCTGGTGTGAAATCCCACGACTCAATCGTGGAACTGCATTGGATACTGCTAGACTAGAGTACTGGAGGGGTGAGTGGAATTCCTGGTGTAGAGGTGAAATTCGTAGATATCAGGAGGAACAACGGAGGCGAAGGCGACTCACTGGACAGTAACCGACACTGAGGTGCGAAAGTGTGGGT |
| P-asv6389 | GCGAGCGTTGTCCGGAATTACTGGGCGTAAAGAGCACGTAGGCGGGACTTTAAGTCAGGTGTGAAAGACCGAGGCTCAACCTCGGGGTGGCACTTGAAACTGGAGTTCTTGAGGGCAGGAGAGGAAAGCAGAATTCCTGGTGTAGCGGTGAAATGCGTAGAAATCAGGAGGAATACCAGTGGCGAAGGCGGCTTTCTGGACTGACCCTGACGCTGAGGTGCGAAAGCGTGGGT |
| P-asv6390 | GCAAGCGTTGTTCGGAATTACTGGACGTAAAGAGCGCGTAGGCGGTCTACCAAGTCAGACGTGAAAGCCCTGGGCTCAACCCGGGAACTGCGTTTGAAACTGGTAAACTTGAGTGTCAGAGGGGTGAGTGGAATTCCTAGTGTAGAGGTGAAATTCGTAGATATTAGGAGGAACACCAGTGGCGAAGGCGACTCACTGGCTGAATACTGACGCTAAGGCGCGAAAGCGTGGGG |
| P-asv6391 | TCGAGTGATGTCCCATATTATTGGGCCTAAGGCGTCCGTAGCTGGCCAGGCAAGTCTATCGGGAAATCCACCTGCCCAACAGGTGGGCGTCCGGTGGAAACTACACGGCTTGGGGCCGAGAGACTCGACGGGTACGTCCGGGGTAGGAGTGAAATCCCGTAATCCTGGACGGACCACCAATGGGGAAACCACGTCGAGAGACCGGACCCGACAGTGAGGGACGAAAGCCAGGGT |
| P-asv6392 | ACAAGCGTTGTCCAGAATTATTGGGCGTAAAGGGAGCGCAGGCGGTTGGAATAGTCTGATGTGAAAGCCCACGGCTTAACCGTGGAAGGTCATTGGAAACTGTCCAACTTGAATGCAGAAGAGGTAAGTGGAATTCCAAGTGTAGCGGTGAAATGCGTAGATATTTGGAGGAACACCAGTGGCGAAGGCGACTTACTGGTCTGTGATTGACGCTGAGGCTCGAAAGCGTGGGT |
| P-asv6393 | GCTAGCGTTACTCGGAATTATTGGGCGTAAAGAGTGCGTAGGTGGTTTTGTAAGTTAGAAGTGAAAGCCTTGGGCTTAACCTGAGAAATGCTTTTAATACTGCAAAGCTTGAGTTCTGAAGGGGATAGTGGAATTCCTAGTGTAGAGGTGACATTCGTAGATATTAGGAGGAACACCGGTGGCGAAAGCGACTATCTGGTCAGATACTGACACTGTTGCACGAAGGCGTGGGG |
| P-asv6394 | GCGAGCGTTGTCCGGATTTACTGGGCGTAAAGGGTGAGTAGGCGGTAATATGTGTCAGATGTAAAAGGCTATGGCTTAACCATAGTTGGCATTTGAAACTGTATTACTTGAGTGCAGGAGAGGTAAGTGGAATTCCTAGTGTAGCGGTGAAATGCGTAGATATTAGGAGGAACACCAGTGGCGAAGGCGACTTACTGGACTGTAACTGACGCTGAGTCACGAAAGCGTGGGT |
| P-asv6395 | GCGAGCGTTAATCGGAATTATTGGGCGTAAAGCGAGTGCAGACGGTTGTTTAAGCCAGATGTGAAATCCCCGAGCTTAACTTGGGACGTGCATTTGGAACTGGATAACTAGAGTGTGTCAGAGGGAGGTAGAATTCCACATGTAGCGGTGGAATGTGTAGAGATGTGGAGGAATACCGATGGCGAAGGCAGCCATCTGGCCTAATACTGACGCTGAGGTACGAAAGCATGGGG |
| P-asv6396 | GCGAGCGTTGTCCGGATTTATTGGGTTTAAAGGGTGCGTAGGCGGCTTGTTAAGTCCGGGGTGAAAGCCCACAGCTCAACTGTGGAACTGCCCTGGATACTGGCAAGCTTGAGTCCAGACGAGGTTGGCGGAATGGATGCTGTAGCGGTGAAATGCATAGATAGCATCCAGAACCCCGATTGCGTAGGCAGCTGACTAGGCTGGCACTGACGCTGAGGCACGAAAGCGTGGGG |
| P-asv6397 | GCGAGCGTTATCCGGAATGACTGGGCGTAAAGGGTCCGTAGGCAGCATTTTTAGTTAGAGGTTAAATCAAAAAGCTTAACTTTTTTCCGCTTCTAAAACGTAAATGCTTTGAGTTTGACAGGAGATAGCAGAATTTCACATGGAGGGGTGATATCCATAGATATGTGAAGGAATACCAGAGGCGAAGGCGGCTATCTAGGTCAATACTGACGTTGAGGGACGAAAGCATGGGG |
| P-asv6398 | CCAAGCGTTAATCGGAATTACTGGGCGTAAAGCGTGCGCAGGCGGTTGTGCAAGACCGATGTGAAAGCCTGTAGCTCAACTCCAGAATTGCCTTTAAGACTGCATCGCTTGAATCCAGGAGAGGTGAGTGGAATTCCGAGTGTAGAGGTGAAATTCGTAGATATTCGGAAGAACACCAGTGGCGAAGGCGGCTCACTGGACTGGTATTGACGCTGAGGTGCGAAAGCGTGGGG |
| P-asv6399 | GCAAGCGTTGTTCGGATTTACTGGGCGTACAGGGAGCGTAGGCGGTTTGGTAAGCCCCTCGGAAAATCTTCGGGCTCAACCCGGAAAGGTCGGGGGGGACTGCCAGGCTAGAGGATGGGAGAGGAGCGCGGAATTCCCGGTGTAGCGGTGAAATGCGTAGAGATCGGGAGGAAGGCCGGTGGCGAAGGCGGCGCTCTGGAACATTTCTGACGCTGAGGCTCGAAAGCGTGGGG |
| P-asv6400 | CCAAGCGTTGTTCGGAATTACTGGGCGTAAAGGGAGCGCAGGCGGACAGGTAAGTCTAAAGTGGAAGCCCCGGGCTCAACCTCGGAAGGTCTTTGGATACTGCCTGTCTTGAGTTTGTCAGAGGAAAGCGGAATTCAGGGTGTAGCGGTGAAATGCTTAGATATCCTGAGGAACACCAATGGCGAAAGCAGCTTTCTGGGGCAAAACTGACGCTGAGGCTCGAAAGCGTGGGT |

| | |
|---|---|
| P-asv6401 | GCAAGCGTTATCCGGATTTACTGGGTTTAAAGGGTGCGCAGGCGGTAATGTAAGTCAGTGGTGAAAGCTTCCAGCTCAACTGGAAAACTGCCATTGATACTGCATTACTTGAGTTTGGGAGAGGTAGGCGGAATACATCATGTAGCGGTGAAATGCTTAGATATGATGTAGAACACCGATTGCGAAGGCAGCTTACTATACCAACACTGACGCTCATGCACGAAAGCGTGGGG |
| P-asv6402 | GCGAGCGTTGTTCGGAATTATTGGGCGTAAAGGGCACGCAGGCGGATTAGAAAGTCTGGTGTGGAAGACCGCAGCTCAACTGCGGGAACGCACTGGAAACTTTTAATCTAGAGTTCAAGAGGGGAAGCTGGAATTCCAGGTGTAGGGGTGAAATCTATAGATATCTGGAAGAACATCAGTGGCGAAGGCGAGCTTCTGGCTATGAACTGACGCTGAGGTGCGAAAGCGTGGGG |
| P-asv6403 | GCGAGCGTTAATCGGAATCACTGGGCGTAAAGCGTGCGTAGGCTGCGTGGCAAGTCAGGCGTGAAAGCCCTCGGCTCAACCGAGGAATTGCGTTTGAAACTGCCATGCTAGAGTCTCGGAGAGGTTGGCGGAATTCCAGGTGTAGGAGTGAAATCCGTAGATATCTGGAGGAACACCGACGGCGAAGGCGGCCAACTGGACGAGTACTGACGCTGAGGTACGAAAGCGTGGGT |
| P-asv6404 | GCGAGCGTTAATCGGAATTACTGGGCGTAAAGCGCGCGTAGGCGGTTTGTTAAGCCAGATGTGAAAGCCCCGGGCTCAACCTGGGAACTGCATCTGGAACTGGCAAGCTAGAGTACAGTAGAGGAAGGTGGAATTCCACGTGTAGCGGTGAAATGCGTAGATATGTGGAGGAACATCAGTGGCGAAGGCGGCCTTCTGGACTGATACTGACGCTGAGGTGCGAAAGCGTGGGG |
| P-asv6405 | GCAAGCGTTATCCGGATTTACTGGGTTTAAAGGGTGCGTAGGCGGGCCATTAAGTCAGATGTGAAAGCCTCAAGCTCAACTTGAGAACTGCGTTTGATACTGGTGGCCTTGAATTCGATAGAAGTGGGCGGAACAAATCGTGTAGCGGTGAAATGCTTAGATATGATTTGGAACACCAATTGCGAAGGCAGCTCACTATGCCGATATTGACGCTGAGGCACGAAAGCGTGGGG |
| P-asv6406 | GCAAGCGTTATCCGGAATTATTGGGCGTAAAGGGTGCGTAGGTGGTTAGTTAAGTCTTTGGTGTAAGTGCAGTGCTTAACGCTGTGATGCCAAAGAAACTGGCTGACTTGAGTGTGGTAGAGGCAAGTGGAATTCCATGTGTAGCGGTAAAATGCGTAAATATATGGAGGAACACCAGTGGCGAAGGCGGCTTGCTGGGCCACAACTGACACTGAGGCACGAAAGCGTGGGG |
| P-asv6407 | GCAAGCGTTAATCAGAATTACTGGGCGTAAAGCGCGCGTAGGCGGACTGTTAAGTCGGGGGTGAAATCCCGGGGCCCAACCCCGGAACTGCCTTCGATACTGGCAGTCTAGAGGACGGAAGAGGTCAGTGGAATGGCGAGTGTAGAGGTGAAATTCGTAGATATTCGCCGGAACACCAGTGGCGAAGGCGACTGACTGGTCCGTTTCTGACGCTGAGGTGCGAAAGCGTGGGG |
| P-asv6408 | GCGAGCGTTGTCCGGAATTATTGGGCGTAAAGGGTGCGTAGGCGGTTTTTTAAGTCTGCATTGTAAGTTCAGTGCTTAACGCTGAAATTGGTGCGGAAACTGGAAGACTAGAGTACTGTAGGGGAAAGCGGAATGTCCGGTGTAGCAGTGAAATGCGTAGAGATCGGATGGAACACCGAAAGCGAAGGCAGCTTTCTGGGCAGTAACTGACGCTGAGGCACGAAAGCGTGGGG |
| P-asv6409 | GCTAGCGTTGTCCGGAATCATTAGGCGTAAAGAGCGTGTAGGCGGCCCGGTAAGTCCGCTGTGAAAGTCGGGGGCTCAACCCTCGAATGCCGGTGGATACTGTCGGGCTAGAGTGCGGAAGAGGCGAGTGGAATTCCTGGTGTAGCGGTGAAATGCGCAGATATCAGGAGGAACACCGATTGCGAAGGCAGCTCGCTGGGACGTAACTGACGCTGAGACGCGAAAGCGTGGGG |
| P-asv6410 | GCAAACGTTGTTCGGACTCACTGGGCTTAAAGGGCGCGTAGGCGGCTCGCCAAGTCAGGGGTGAAATCCCCCGGCTCAACCGGGGAACAGCTTCTGATACTGGCGGGCTCGAGGGGGGTAGGGGCAGGCGGAACTTCCGGTGGAGCGGTGAAATGCGTAGATATCGGAAGGAACGCCGGTGGCGAAAGCGGCCTGCTGGACCCCTTCTGACGCTGAGGCGCGAAAGCCAGGGG |
| P-asv6411 | GCGAGCGTTGTCCGGAGTTACTGGGCGTAAAGAGCGTGTAGGCGGTTGAAAAAGTCAGGTGTGAAAATTTTCGGCTCAACCGAGAAACTGCGCCTGAAACTTTTCGACTGGAGTTTGGGAGGGGCAACTGGAATTCCTGGTGTAACGGTGAAATGTGTAGATATCAGGAGGAACACCCGTGGTGAAGGCGGGTTGCTGGACCAAAACTGACGCTGAGACGCGAAAGCGTGGGT |
| P-asv6412 | GCAAGCGTTGCCCGGAATTACTGGGCGTAAAGAGCTCGTAGGCGGGGGCGTGCGTCCGAAAGGAAATCTCAAGGCTCAACCTTGGGGCTTTTTCGGATACGGCGATTCTTGAGGGTGAGAGAGGAAAGCGGAATTCCAGGTGTAGCGGTGAAATGCGTAGATATCTGGAGGAACACCAGCGGCGAAGGCGGCTTTCTGGCTCATTCCTGACGCTGAGGAGCGAAAGCGTGGGT |
| P-asv6413 | GCTAGCGTTGTTCGGAATTACTGGGCGTAAAGGGCGCGTAGGCGGATGTCTAAGTTAGGGGTGAAATCCCGGGGCTCAACCTCGGAACTGCCCTTGATACTGGATGTCTTGAGTTCGTGAGAGGGTGGCGGAATACCCAGTGTAGAGGTGAAATTCGTAGATATTGGGTAGAACACCAGTGGCGAAGGCGGCCACCTGGCACGACACTGACGCTGAGGCGCGAAAGCGTGGGG |
| P-asv6414 | GCTAGCGTTGCTCGGAATTACTGGGCGTAAAGCGAGCGCAGACGGTTATTTAAGCAAGATGTGAAAGCCCCGGGCTCAACCTGGGAACTGCGTTTTGAACTGGGTAACTAGAGTGTGTCAGAGGGGGGTAGAATTCCACGTGTAGCAGTGAAATGCGTAGAGATGTGGAGGAATACCGATGGCGAAGGCAGCCCCCTGGGATAACACTGACGTTCATGCTCGAAAGCGTGGGT |
| P-asv6415 | GCGAGCGTTGTCCGGATTTATTGGGTTTAAAGGGTGCGTAGGTGGCTTTATAAGTCAGTGGTGAAATACAGCCGCTCAACGGTTGAGGTGCCATTGATACTGTAGAGCTTGAAACAATTGGAGGCTGCCGGAATGGATGGTGTAGCGGTGAAATGCATAGATATCATCCAGAACACCGATTGCGAAGGCAGGTGGCTACGTTTGATTTGACACTGAGGCGCGAAAGCATGGGG |
| P-asv6416 | GCGAGCGTTAATCGGAATTACTGGGCGTAAAGCGCATGTAGGCGGATTAGTAAGTTGGAGGTGAAATCCCAGAGCTCAACTCTGGAACTGCCTTCAAAACTGCTAGTCTGGAACACGGTAGAGGAAAGCGGAATACCTAGTGGAGAGGTGAAATTCGTAGATATTAGGTGGAACACCAGAGGCGAAGGCGGCTTTCTGGACCGTTGTTGGCGCTGAGATGCGAAAGCGTGGGG |
| P-asv6417 | GCGAGCGTTAATCGGAATTACTGGGCGTAAAGCGTGCGCAGGCGGCTTTGCAAGACAGATGTGAAATCCCCGGGCTCAACCTGGGAACTGCATTTGTGACTGCAAGGCTAGAGTACGGTAGAGGGGGATGGAATTCCGCGTGTAGCAGTGAAATGCGTAGATATTCGGAGGAACACCAGTGGCGAAGGCGGCTTTCTGGACAGTAACTGACGCTGAGGAACGAAAGCGTGGGG |
| P-asv6418 | GCGAGCGTTAATCGGAATTACTGGGCGTAAAGCGCGCGTAGGCGGTTTGTTAAGTCAGATGTGAAAGCCCCGGGCTCAACCTGGGAATTGCATTTGAAACTGGCAGGCTAGAGTGCAGTAGAGGGAGGTGGAATTTCCGGTGTAGCGGTGAAATGCGTAGAGATCGGAAGGAACACCAGTGGCGAAGGCGGCCTCCTGGACTGACACTGACGCTGAGGTGCGAAAGCGTGGGG |
| P-asv6419 | CCAAGCGTTAATCGGAATTACTGGGCGTAAAGCGTGCGCAGGCGGTTGTGCAAGACCGATGTGAAATCCCCGAGCTTAACTTGGGAATTGCATTGGTGACTGCACGGCTAGAGTGTGTCAGAGGGGGGTAGAATTCCACGCGTAGCAGTGAAATGCATAGATATGACATAGAACACCTATTGCGAAGGCAGCTTACTACGCCTATATTGACGCTGAGGCACGAAAGCGTGGGG |
| P-asv6420 | GCGAGCGTTATCCGGATTTATTGGGTGTAAAGGGTGCGTAGGCGGTTTGATAAGTCTGTGGTTTAAGCCCGAAGCTTAACTTCGGTTCGCCACAGAAACTGTCAGACTTGAGTATGGTAGAGGCAAGTGGAATTTCTAGTGTAGCGGTTAAATGCGTAGATATTAGAAGGAACACCAGTGGCGAAGGCGACTTGCTGGGCCATTACTGACGCTGAGGCACGAAAGCGTGGGG |
| P-asv6421 | GCAAGCGTTGTCCGGATTTATTGGGTTTAAAGGGTGCGTAGGCGGATGTTTAAGTCAGTGGTGAAAGCCGGCAGCTCAACTGTCGAACTGCCATTGATACTGAATATCTTGAGTATAGTTGAGGTAGGCGGAATGGAAGGTGTAGCGGTGAAATGCATAGATATCTTCCAGAACACCGATAGCGTAGGCAGCTTACTAAGCTATAACTGACGCTGAGGCACGAAAGCATGGGG |
| P-asv6422 | GCAAGCGTTGTCCGGATTTACTGGGCGTAAAGCGAACGCAGGCGGATGGTTAAGTCCAAAGCGAAAGGTCAGAGCTCAACTCTGGCACTGCTTGGGATACTGGCCGTCTTGAGTACCGGAGAGGAAAGCGGAACGACACGTGTAGCGGTGAAATGCGTTGATATGTGTCGGAACACCAATGGCGAAAGCAGCTTTCTGGACGGAAACTGACGCTCATGTTCGAAAGCCAAGGT |
| P-asv6423 | GCGAGCGTTAATCGGAATTACTGGGCGTAAAGGGCGCGTAGGCGGTTGTATAAGTTAGGTGTGAAATCCCCGGGCTTAACCTGGGAATGGCATTTAATACTGTATGACTAGAGTAGAGTAGAGGGAAGTGGAATTTCCGGTGTAGCGGTGAAATACGTAGATATCGGAAAGAACACCAGTGGCGAAGGCGGCTTCCTGGACTCATACTGACGTTGAGGCGCGAAAGCGTGGGT |
| P-asv6424 | CCGAGCGTTGTTCGGATTCACTGGGCGTAAAGGGTGCGTAGGTGGTTGGTTAAGTTTGAGGTGAAAGCTCCGAGCTCAACTCGGAAAATGCCTTGAAGACTATCTGACTAGAGGATTGGAGGGGAGATTGGAATTCTCGGTGTAGCAGTGAAATGCGTAGATATCGAGAGGAACACCAGTGGCGAAGGCGAATCTCTGGACAATTCCTGACACTGAGGCACGAAAGCCAAGGG |
| P-asv6425 | GCAAGCGTTATCCGGATTCACTGGGTTTAAAGGGTGCGTAGGTGGATTAATAAGTCAGTGGTGAAAGCCCCGAGCTCAACTTGGGAATTGCCATTGATACTGTTAGTCTTGAGTTCGGTTGAGGTAGGCGGAATAAGTCATGTAGCGGTGAAATGCTTAGATATGACTTAGAACACCGATTGCGAAGGCAGCTTACCAAGCCGACACTGACACTGAGGCACGAAAGCGTGGGG |
| P-asv6426 | GCGAGCGTTATTCGGAATTACTAGGCGTAAAGCGAGTGTAGGCGGACTCTTGAGTCTGACGTTAAATCTCCCGGCTTAACTGGGAGCTGTCGTCGGATACTGAGAGCCTCGAATGGGGTAGGGGGCAGCGGAATTCCCGGTGTAGCGGTGAAATGCGTAGATATCGGGAGGAACACCTATGGCGAAAGCAGCTGCCTGGGCCTCTATTGACGCTAAGACTCGAAAGCTGGGGG |
| P-asv6427 | TCGAGCGTTGTCCGGAGTTACTGGGCGTAAAGCGCGTGCAGGTGGCTCATTACGCCCGGCGTGAAAGCCCCCGGCTCAACCGGGGAGGGTCGTCGGGGACGGGTGAGCTTGAGGGTCGCAGGGGCTGGTAGAATTCCCGGTGTAGTGGTGAAATGCGTAGAGATCGGGAGGAATACCCGTGGCGAAGGCGGCCAGCTGGGCGACACCTGACACTGAGACGCGAAGGCGTGGGG |
| P-asv6428 | ACAAGCGTTGTCCGGAATTACTGGGTGTAAAGGGAGCGCAGGCGGGAAGGCAAGTTGGAAGTGAAATCCATGGGCTCAACCCATGAACTGCTTTCAAAACTGTTTTTCTTGAGTAGTGCAGAGGTAGGCGGAATTCCCGGTGTAGCGGTGGAATGCGTAGATATCGGGAGGAACACCAGTGGCGAAGGCGGCCTACTGGGCACCAACTGACGCTGAGGCTCGAAAGTGTGGGT |
| P-asv6429 | ACAAACGTTATTCGGAATCACTGGGCTTAAAGGGTGCGTAGGCGGCGCGATAAGTTGGATGTGAAATGCCTCGGCTCAACCGAGGAACTGCGTCCAAAACTATCGTGCTTGAGGAAGGTAGAGGTGAGCGGAACTTAGGGTGGAGCGGTGAAATGCGTTGATATCCTAAGGAACACCGGTGGCGAAAGCGGCTCACTGGACCTTTTCTGACGCTGAGGCACGAAAGCTAGGGG |
| P-asv6430 | GCGAACGTTATTCGGAATCACTGGGCTTAAAGCGCGTGTAGGCGGATGGCCGCGTCGGTCGCTCAAAGCCCCCGGCTCAACCGGGGGTGTGGCGCCGAAACGGGTCGTCTGGAGGGGCGTAGGGGGAGCTGGAACTCACGGTGGAGCGGTGATGTGCGTTGAGATCGTGAGGAACGCCCGCGGCGAAAGCGAGCTCCTGGACGCCTACTGACGCTGAGACGCGAAAGCCAGGGG |
| P-asv6431 | GCGAGCGTTAATCGGAATTACTGGGCGTAAAGGGTGCGTAGGTGGTTTGATAAGTTGGTTGTGAAATTCCTGGGCTCAACCTGGGCATTGCAACTAAGACTGTTGAACTCGAGTATGGGAGAGGGTAGTGGAATTTCCGGTGTAGCGGTGAAATGCGTAGAGATCGGAAGGAACACCAGTGGCGAAGGCGACTACCTGGCCCAATACTGACACTGAGGCACGAAAGCGTGGGG |
| P-asv6432 | GCAAGCGTTAATCGGAATGACTGGGCGTAAAGAGTGCGTAGGTGGCAAAATAAGTTAGTTGTGAAATCCCTGGGCTTAACCTAGGAACTGCAATTGATACTGTTTAGCTAGAGTATAGTAGAGGTAAGTGGAATTTCCGGTGTAGCGGTGAAATGCGTAGATATCGGAAGGAACACCAGTGGCGAAGGCGGCTTACTGGGCTAATACTGACACTGAGGCACGAAAGCGTGGGG |
| P-asv6433 | GCGAACGTTGTTCGGATTCACTGGGCTTAAAGGGCGCGTAGGCGGCTTGCCAAGTCAGGGGTGAAATCTTTCGGCTCAACCGGAAAACGGCCCCTGATACTGGCGGGCTGGAGGGAGGTAGGGGCAGATGGAACTTCCGGTGGAGCGGTGAAATGCGTAGATATCGGAAGGAACGCCGGTGGCGAAAGCGATCTGCTGGACCTCTTCTGACGCTGAGGCGCGAAAGCCAGGGG |
| P-asv6434 | GCAAGCGTTGTCCGGAATTACTGGGCGTAAAGGGCGTGTAGGCGGTCGCACACGTAGCGGTTTTCAGCCGCCGGCTCACCCGGCGGAGGGCGGCTAAACGGTGCGGCTGGAGGGCAGGAGAGGTGCACGGAATTCCTGGTGGAGCGGTGAAATGCGTAGAGATCAGGAAGAACACCAGTGGCGAAGGCGGTGCACTGGCCCGACCCTGACGCTGAGGCGCGATAGCGTGGGG |
| P-asv6435 | TCGAGCGTTATCCGGATTTATTGGGTTTAAAGGGTCCGTAGGCTGTATAATAAGTCAGAGGTGAAATCCCATAGCTTAACTATGGAACTGCCTTTGAAACTATTTTACTTGAATTTGGCGGCGGTATGCGGAATACGTTATGTAGCGGTGAAATGCATAGATATAACGTGGAACCCCAATTGCGAAGGCAGCATACCATACCAACATTGACGCTGATGGACGAAAGCATGGTT |
| P-asv6436 | GCAAGCGTTGTTCGGAATCACTGGGCGTAAAGAGTTCGTAGGCGGACTTGATAGTACGGGGTTAAAGACCAGGGCCCAACCTTGGGAATGCTTCGTAAACGTCAAGTCTAGAGTGTCGGAGAGGATAGCGGAATTCCTGGTGTAGAAGTGAAATTCGTAGATATCAGGAGGAACACCTGAGGCGAAGGCGGCTATCTGGCCTAATACTGACGCTGAGGTACGAAAGCATGGGG |
| P-asv6437 | GCAAGCGTTGTCCGGAATTATTGGGCGTAAAGCGCGCGCAGGCGGTTTGTTAAGTTGTGGGTAAAAGGCATTGGCTCAACCAATGTAAGCCTGCAAAACTGGCAGACTTGAGTGCAGGAGAGGGGAGTGGAATTCCCGGTGTAGCGGTGGAATGCGTAGATATCGGGAGGAACACCAGTGGCGAAGGCGACTCTCTGGCCTGTAACTGACGCTGAGGCGCGAAAGCGTGGGT |
| P-asv6438 | GCAAGCGTTGTTCGGAATAACTGGGCGTAAAGGGTGCGTAGGCTGCCCGATAAGTCTGATGTGAAAGCCCTGGGCTTAACCTAGGAATAGCATTGGATACTGTTGGGCTAGAGTGTTAAAGGGGATGATGGAATTATGCATGGAGAGGTGGAATTCATAGATATGCATAGGAACACCGGAGGCGAAGGCGATCATCTGGATAACAACTGACGCTGAGGCACGAAAGCGTGGGG |
| P-asv6439 | GCAAGCGTTAATCGGAATTACTGGGCGTAAAGGGCGCGTAGGTTGTTTGGTAAGTTGGCTGTGAAAGCCCCGGGCTTAACCTGGGAATGCCAGTCAAAACTGCCAAACTAGAGTACAGTAGAGGGTGGTGGAATTTCCAGTGTAGCGGTGAAATGCGTAGAGATTGGAAGGAACACCAGTGGCGAAGGCGGCCACCTGGACTGATACTGACACTGAGGCGCGAAAGCGTGGGT |

| | |
|---|---|
| P-asv6440 | GCAAGCGTTATCCAGATTTACTGGGTGTAAAGGGAGCGTAGGTGGCATGGCAAGTCCGATGTGAAAGCCCGGGGCTCAACCCCGGGATTGCATTAGAAACTGTCAGGCTAGAGTGCAGGAGAGGTAAGTGGAATTCCTAGTGTAGCGGTGAAATGCGTAGATATTAGGAGGAACACCAGTGGCGAAGGCGACCACCTGGACTGATACTGACACTGAGGTACGAAAGCATGGGG |
| P-asv6441 | GCAAGCGTTATTCGGAATTATTGGGCGTAAAGGGCGCGTAGGCGGTCTTGTCCGTCAGGTGTGAAAGCCCGGGGCTCAACCCCGGAAGTGCACTTGAAACAGCAAGACTTGAATACGGGAGAGGAAAGCGGAATTCCTGGTGTAGAGGTGAAATTCGTAGATATCAGGAGGAACACCGATGGCGAAGGCAGCTTTCTGGACCGATATTGACGCTGAGGCACGAAAGCGTGGGG |
| P-asv6442 | GCAAGCGTTAATCGGAATTACTGGGCGTAAAGCGAGCGTAGGTGGCTTGATAAGTCAGATGTGAAAGCCCCGGGCTTAACCTGGGAACGGCATCTGATACTGTTGAGCTAGAGTAGGTGAGAGGAAGGTAGAATTCCAGGTGTAGCGGTGAAATGCGTAGAGATCTGGAGGAATACCGATGGCGAAGGCAGCCTTCTGGCATCATACTGACACTGAGGCTCGAAAGCGTGGGT |
| P-asv6443 | GCTAGCGTTATTCGGAATTACTGGGCGTAAAGCGCTCGTAGGCGGCTTTGTAAGTTAGAGGTGAAAGCCTGGAGCTCAACTCCAGAACTGCCTTTGAAACTAGGTGACTAGAATCTTGGAGAGGTCAGTGGAATTCCGAGTGTAGAGGTGAAATTCGTAGATATTCGGAAGAACACCAGTGGCGAAGGCGACTGACTGGACAAGTATTGACGCTGAGGTGCGAAAGCGTGGGG |
| P-asv6444 | CCAAGCGTTGTTCGGATTCACTGGGCGTAAAGGGAGCGTAGGCTGTGCGGTAAGTCAGATGTGAAATCTCGAGGCTCAACCTCGAAACTGCATCCGATACTGCCGTACTAGAGGATTGGAGAGGTAACAGGAATTCACGGTGTAGCAGTGAAATGCGTAGATATCGTGAGGAAGACCAATGGCGAAAGCATGTTACTGGACAACTCCTGACGCTGAGGCTCGAAGGCTAGGGT |
| P-asv6445 | GCGAGCGTTATCCGGAATCATTGGGTTTAAAGGGTCCGTAGGCGGGCTAATAAGTCAGAGGTGAAAGCGCTCAGCTCAACTGAGCAACTGCCTTTGAAACTGTTAGTCTTGAATGGTTGTGAAGTAGTTGGAATGTGTAGTGTAGCGGTGAAATGCTTAGATATTACACAGAACACCGATAGCGAAGGCATATTACTAACAATTAATTGACGCTGATGGACGAAAGCGTGGGG |
| P-asv6446 | GCAAGCGTTGCTCGGAATTACTGGGCGTAAAGGGCGCGTAGGCGGGATGGCAAGTCGGTCGTGAAATCCCTCGGCTCAACTGAGGAGCGTCGATCGATACTGCTGTTCTTGAGGATTGCAGAGGAGACTGGAATTCCCGGTGTAGCGGTGAAATGTGTAGAGATCGGGAGGAACACCGGTGGCGAAGGCGGGTCTCTGGGCAATTCCTGACGCTGAGGCGCGAAAGCGTGGGG |
| P-asv6447 | GCGAGCGTTAATCGGAATTACTGGGCGTAAAGCGTGCGCAGGCGGCTTTGCAAGACAGAGGTGAAATCCCCGGGCTCAACCTGGGAACTGCCTTTGTGACTGCAAGGCTAGAGTACGGCAGAGGGGGATGGAATTCCGCGTGTAGCAGTGAAATGCGTAGAGATGTGGAAGAATACCGATGGCGAAGGCAGCCCCCTGGGATAACACTGACGCTCATGCACGAAAGCGTGGGG |
| P-asv6448 | GCTAGCATTGTTCGGAATCATTGGGCGTAAAGCGTATGTAGGCGGCTTGGCAAGTTGGTTGTGAAATCCCGGGGCTCAACCTCGGAATTGCAATCAAAACTGCTTCGCTAGAGTATAGTAGGGGATAGTGGAATTCCTAGTGTAGGGGTGAAATTCGTAGATATTAGGAGGAACATCGGCGGCGAAAGCGGCTATCTAGGCTATTACTGACGCTGAGATACGAAAGCGTGGGG |
| P-asv6449 | CCAAGCGTTGTTCGGAATAACTGGGCGTAAAGCGTGTGTAGGCTGCGCGGAAAGTCAAATGTGAAAGCCAAGGGCTCAACCCTTGAACTGCATTCGATACTCCCGTGCTAGAGTAATGGAGAGGTAAGTGGAATTCTCGGTGTAGCAGTGAAATGCGTAGATATCGAGAGGAACATCAATGGCGAAGGCAACTTACTGGACATTTACTGACGCTCAGACACGAAGGCTAGGGG |
| P-asv6450 | GCGAGCGTTGTTCGGAATTACTGGGCGTAAAGCGCGCGCAGGCGGCTCTTTAAGTCTATGGGTCAAAGACCGGGGCTCAACCCCGAGTTCGCCCGGGATACTGGAGAGCTAGGGACATGTAGGGGCGAGCGGAATTCCGGGTGTAGCGGTGGAATGCGTAGAGATCCGGGAGAACACCGGGGGCGAAGGCGGCTCGCTGGGCATGGTCCGACGCTGAGGCGCGAAAGCGTGGGG |
| P-asv6451 | GCTAGCGTTGCTCGGAATCACTGGGCGTAAAGGGTGCGTAGGTGGATCTTTAAGTCAGGGGTGAAATCCTGGAGCTCAACTCCAGAACTGCCTTTGATACTGAGGATCTTGAGTTCGGGAGAGGTAAGTGGAACTGCGAGTGTAGGGGTGAAATTCGTAGATATGCGGAGGAACACCGATGGCGAAGGCAATCCCCTGGACCTGTACTGACGCTCATGCACGAAAGCATGGGG |
| P-asv6452 | GCAAGCGTTGTTCGGAATTACTGGGCGTAAGGCGTGCGTAGTCGGTATTGAGAGTCACGGGTGAAATCCCAGGGCTTAACCCTGGAACTGCCTTTGAGACTGCATCGCTTGAATCCAGGAGAGGTGAGTGGAATTCCGAGTGTAGAGGTGAAATTCGTAGATATTCGGAAGAACACCAGTGGCGAAGGCGGCTCACTGGACTGGTATTGACGCTGAAGTGCGAAAGCGTGGGG |
| P-asv6453 | GCAAGCGTTGTTCGGAATTACTGGGAGTAAAGCGTGCGTAGTCGGTATTGAGAGTCACGGGTGAAATCCCAGGGCTCAACCTTGGAATTGCCTTTGATACTGGCAGTCTTGAGTACGGAAGAGGTATGTGGAACTCCGAGTGTAGAGGTGAAATTCGTAGATATTCGGAAGAACACCAGTGGCGAAGGCGACATACTGGTCCGTTACTGACGCTGAGGCTCGAAAGCGTGGGG |
| P-asv6454 | GCAAGCGTTGTTCGGAATTACTGGGCGTAAAGGGCGCGTAGGCGGTCTGTCAAGTCGGATGTGAAATCCCCGGGCTCAACCTGGGAACTGCAGGCGAAACTGCCGTGCTGGAGTATGGGAGAGGTGCGTGGAATTCCCGGTGTAGCGGTGAAATGCGTAGATATCGGGAGGAACACCTGTGGCGAAAGCGGCGCACTGGACCATAACTGACGCTGAGGCGCGAAAGCTAGGGG |
| P-asv6455 | GCAAGCGTTGTTCGGAATTATTGGGCGTAAAGGGCGCGTAGGCGGTCGGCTAAGTCAGCTGTGAAAGCCCGGGGCTCAACCCCGGAGGGTCGGCTGATACTGGTTGACTAGAGTACGGAAGAGGTAGCTGGAATTCCCGGTGTAGCGGTGAAATGCGTAGATATCGGGAGGAACACCTGAGGCGAAGGCGGGCTACTGGGCCGATACTGACGCTGAGGCGCGAAAGCCAGGGG |
| P-asv6456 | GCAAGCGTTGTCCGGATTTATTGGGCGTAAAGAGCTCGTAGGTGGTTTGGTAAGTCGGATGTGAAATCTCCAGGCTCAACCTGGAGGGGTCATTCGATACTGCCATGACTAGAGGTCGGTAGGGGAGTGTGGAATTCCTGGTGGAGCGGTGAAATGCGCAGATATCAGGAGGAACACCCGTAGCGAAGGCGGCACTCTGGGCCGATTCTGACACTGAGGAGCGAAAGCGTGGGG |
| P-asv6457 | GCGAGCGTTGCCCGGATTTATTGGGCGTAAAGGGCGCGTAGGCGGCCACGCACGTCCGTTGTGAAATCTCCCGGCTCAACCGGGAGGGTGCGCCGAATACTGAGGGACTAGAGTGTGGGAGAGGAAAGTGGAATTCCTGGTGTAGCGGTGAAATGCGTAGATATCAGGAGGAACACCGGTGGTGTAGACGGCTTTCTGGACCATAACTGACGCTGAGACACGAAAGCGTGGGT |
| P-asv6458 | GCAAGCGTTACTCGGAATTACTGGGCGTAAAGCGTGCGTAGGTGATTCGCCAAGTCAGGTGTGAAATCCCCGGGCTTAACCTGGGAATTGCGCTTGAAACTAGCGGGCTTGAGTGTGGCAGAGGGGGGTGGAATTCCACGTGTAGCAGTGAAATGCGTAGAGATGTGGAGGAACACCGATGGCGAAGGCAGCCCCCTGGGCTAACACTGACGCTCATGCACGAAAGCGTGGGG |
| P-asv6459 | GCGAGCGTTGTTCGGAATTATTGGGCGTAAGGCGCTGGTAGGTGGCTTGTTAAGTCGCGTGTGAAATCTCCAGGCTCAACCTGGAAAGTGCGCGCGAAACTGGCAGGCTTGAGTGCCGGAGAGGGTCGCGGAATTCCCGGTGTAGAGGTGAAATTCGTAGATATCGGGAGGAACACCAGAGGCGAAGGCGGCGACCTGGAAGGTAACTGACACTGAGCAGCGAAAGCGTGGGG |
| P-asv6460 | GCTAGCGTTGTTCGGAATCACTGGGCGTAAAGCGCACGTAGGCGGACCATTAAGTCAGGGGTGAAAGCCTGGAGCTCAACTCCAGAACTGCCTTTGATACTGATGGTCTCGAGTATGGAAGAGGTTGGTGGAACTGCGAGTGTAGAGGTGAAATTCGTAGATATTCGCAAGAACACCAGTGGCGAAGGCGGCCAACTGGTCCATTACTGACGCTGAGGTTCGAAAGCGTGGGG |
| P-asv6461 | GCAAGCGTTGTCCGGATTTACTGGGCGTAAAGCGAGCGCAGGCGGAATGATAAGTCTGATGTGAAAGCCCACGGCTCAACCGTGGAACTGCATCGGAAACTGTCATTCTTGAGTGCAGAAGAGGAGAGTGGAATTCCATGTGTAGCGGTGGAATGCGTAGATATATGGAAGAACACCAGTGGCGAAGGCGGCTCTCTGGTCTGCAACTGACGCTGAGGCTCGAAAGCATGGGT |
| P-asv6462 | GCGAGCGTTGTTCGGATTTACTGGGCGTAAAGAGCTCGTAGGCGGTCTGCTACGTCCGCTGTAAAAACCTAGGGCTTAACCCTGGGCTTGCAGTAGATACGGGCAGACTAGAGGTAGGTAGGGGAGAATGGAATTCCCGGTGTAGCGGTGAAATGCGCAGATATCGGGAGGAACACCGGTGGCGAAGGCGGTTCTCTGGGCCTTACCTGACGCTGAGGAGCGAAAGCGTGGGG |
| P-asv6463 | GCGAGCGTTGTCCGGAGTTACTGGGCGTAAAGGGCATGCAGGCGGTCGTACGCGTTTTGTGTGACAGCGGGTGGCTTAACTACCCGAGTATGCGAAAGACGGTATGACTTGAGGGCTCGAGAGGGACACGGAATTCCGGGTGGAGTGGTGAAATGCGTAGAGATCCGGAAGAACACCGACGGCGAAGGCAGCGTCCTGGCCGAGGCCTGACGCTCAGGCACGAAAGCGCGGGG |
| P-asv6464 | GCAAACGTTATTCGGAATCACTGGGCATAAAGAGCGCGTAGGCGGCTTCATAAGTCAGGGGTGAAATCCGGCAGCTTAACTGTCGCAGTGCCTTTGATACTGTGGAGCTCGAGGGAGGTAGGGGTCTGTGGAACTTCCGGTGGAGCGGTGAAATGCGTTGATATCGGAAGGAACGCCGGTGGCGAAAGCGACGGACTGGATCTCTTCTGACGCTGAGGCGCGAAAGCCAGGGG |
| P-asv6465 | GCAAGCGTTGTCCGGATTTATTGGGTTTAAAGGGTGCGTAGGCGGCCCTGTAAGTCAGTGGTGAAAGCCTCCAGCTCAACTGGAGAACTGCCATTGATACTGCAGGGCTTGAGTACAGACGAGGTAGGCGGAATTGATTGTGTAGCGGTGAAATGCATAGATACAATCAAGAACTCCGATTGCGTAGGCAGCTTACTAGGCTGTAACTGACGCTGAGGCACGAAAGCGTGGGG |
| P-asv6466 | GCGAGCGTTGTTCGGAATCACTGGGCGTAAAGGGCGCGTAGGCGGCCCTCGTTTGTCGCGTGTGAAAGCCCCGGGCTCAACCTGGGAACGGCGCGCGAGACGGCGGGGCTGGAGTTCGGGAGAGGGATGCGGAATTCCAGGTGTAGCGGTGAAATGCGTAGATATCTGGAGGAACACCGGTGGCGAAGGCGGCGTTCTGGACTGCAACTGACACTGAGAAACGAAAGCCAGGGG |
| P-asv6467 | GCGAGCGTTGCTCGGAATCACTGGGCGTAAAGCGTACGCAGGCGGATCGTTAAGTCGGGTGTGAAATCCCGGGGCTCAACCCGGAACTGCATTCGAAACTGGCGGTCTTGAGTACGGGAGAGGGAGGTGGAATTCCTGGTGTAGAGGTGAAATTCGTAGATATCAGGAGGAACACCGGTGGCGAAAGCGGCCTCCTGGCCCGATACTGACGCTCATGTACGAGAGCGTGGGG |
| P-asv6468 | CCAAGCGTTAGGCGGAATCACTGGGCTTAAAGCGCGTGTAGGTGGCGGCCTAAGTACCTTGTGAAATCCTGCGGCTCAACCGCAGAACTGCTTGGTATACTGGGTCGCTTGAGCCATCCAGGGGCAACTGGAACAAGCGGTGGAGCGGTGAAATGCGTAGATATCGCTTGGAACGCCAAAGGTGAAACCAGGTTGCTGGGGATGTGCTGACACTGAGACGCGAAAGCCAGGGG |
| P-asv6469 | GCGAGCGTTGTTCGGAATTACTGGGCGTAAAGCGCGCGCAGGCGGCCTTTTAAGTCTCACCGTTCAAGCCCGGGGCTCAACTCCGGTCCGCGGGGGATACTGGAAGGCTAGGGACATGTAGGGGCAAGCGGAATTCCGGGTGTAGCGGTGGAATGCGTAGAGATCCGGAAGAACACCGGAGGCGAAGGCGGCTTGCTGGGCATGGTCCGACGCTGAGGCGCGAAAGCGTGGGG |
| P-asv6470 | GCGAGCGTTGTCCGGAATCACTGGGCGTAAAGGGCGCGTAGGCGGCTCGGTAAGGTTGCGGTGAAAGCCCGGGGCTCAACCCCGGGTCGGCCGTGGCAACTGCTGGGCTCGAGCACTGTAGAGGCAGGTGGAATTCCGGGTGTAGCGGTGGAATGCGTAGAGATCCGGAAGAACACCGGTGGCGAAGGCGGCCTGCTGGGCAGTAGCTGACGCTGATGCGCGACAGCGCGGGG |
| P-asv6471 | GCAAGCGTTGTCCGGAATCATTGGGTGTAAAGGGTGCGCAGGCTGGCATTTAAGTCTGGGGTGAAATCTTGCCGCTTAACGGTAAAATGGCCTTAGATACTGAATGTCTTGAATCTGACAGAGGTCGGTGGAATTCGTAGTGTAGCGGTGAAATGCATAGATATTACGAAGAACATCAGTGGCGAAGGCGGCCGGCTGGGTCAGTATTGACGCTGAGGTACGAGAGCGTGGGG |
| P-asv6472 | GCGAGCGTTGTCCGGATTTATTGGGCGTAAAGGGCTCGTAGGTGGTTGATCGCGTCGGAAGTGTAATCTTGGGGCTTAACCCTGAGCGTGCTTTCGATACGGGTTGACTTGAGGAAGGTAGGGGAGAATGGAATTCCTGGTGGATCGGTGGAATGCGCAGATATCAGGAGGAACACCAGTGGCGAAGGCGGTTCTCTGGGCCTTTCCTGACGCTGAGGAGCGAAAGCGTGGGG |
| P-asv6473 | GCTAGCGTTGCTCGGAATCACTGGGCGTGAAGGGTGCGTAGGTGGATCTTTAAGTCAGGGGTGAAATCCTGGAGCTCAACTCCAGAACTGCCTTTGATACAGCATATCTCGAGTCCGAGAGAGGTGAGTGGAATTCCTAGTGTAGAGGTGAAATTCGTAGATATTAGGAAGAACACCGGTGGCGAAGGCGGCTCACTGGCTCGGTACTGACGCTGAGGTGCGAAAGCGTGGGG |
| P-asv6474 | GCAAGCGTTATCCGGATTCACTGGGTTTAAAGGGTGCGTAGGCGGGTATTTAAGTCAGTGGCGAAATCCTAGAGCTTAACTCCAGAACTGCCTTTGATACTGAGGATCTTGAGTTCGGGAGAGGTAAGTGGAACTGCGAGTGTAGAGGTGAAATTCGTAGATATTCGCAAGAACACCAGTGGCGAAGGCGGCTTACTGGCCCGATACTGACACTGAGGCACGAAAGCGTGGGG |
| P-asv6475 | GCTAGCGTTGCTCGGAATTACTGGGCGTAAAGGGCGCGTAGGCGGACAGTTAAGTTGGGGGTGAAAGCCCGGGGCTCAACCTCGGAAATGCCTTCAATACTGGCTGTCTTGAGTACGGGAGAGGTGAGTGGAACTCCGAGTGTAGAGGTGAAATGCTTAGATATGACGTAGTACACCTATTGCGAAGGCAGCTTACTACGCATATATTGACGCTGAGGCACGAAAGCGTGGGG |
| P-asv6476 | GCAAGCGTTAATCGGAATTACTGGGCGTAAAGCGTGCGTAGACGGTTACATAAGTCGGGTGTGAAAGCCCCGGGCTCAACCTGGGAATTGCATTCGAGACTGCGTAGCTAGGGTGCGGAAGAGGGAAGCTGAATTTCCGGTGTAGCGGTGAAATGCGTAGATATCGGAAGGAACACCAGTGGCGAAAGCGGCTTCCTGGTCCAGCACCGACGTTCAGGCACGAAAGCGTGGGG |
| P-asv6477 | TCTCTCTTTTCTCTTCCTTCCTTTTCTTAAAGGTCGCGTAGGCGGACAGTTAAGTTGGGGGTGAAAGCCCGGGGCTCAACCTCGGAAATGCCTTCAATACTGGCTGTCTTGAGTACGGGAGAGGTGAGTGGAACTCCGAGTGTAGAGGTGAAATTCGTAGATATTCGGAAGAACACCAGTGGCGAAGGCGACTCACTGGCCCGTTACTGACGCTGAGGCGCGAAAGCGTGGGG |
| P-asv6478 | GCGAGCGTTGTCCGGAATTACTGGGCGTAAAGAGCTCGTAGGTGGTTTGTCGCGTCGTTTGTGTAAGCCCGCAGCTTAACTGCGGGACTGCAGGCGATACGGGCATAACTTGAGTGCTGTAGGGGAGACTGGAATTCCTGGTGTAGCGGTGAAATGCGCAGATATCAGGAGGAACACCGATGGCGAAGGCAGGTCTCTGGGCAGTAACTGACGCTGAGGAGCGAAAGCGTGGGT |

| | |
|---|---|
| P-asv6479 | GCAAGCGTTGTCCGGAATGACTGGGCGTAAAGGGCGTGTAGGCGGTGCGTCACGTAGCGGTTTTCAGCCGTCGGCTCACCCGACGGAGGGCGGCTAAACGGGCGCGCTGGAGGGCAGGAGAGGTGCACGGAATTCCTGGTGGAGCGGTGAAATGCGTAGAGATCAGGAAGAACACCCGTGGCGAAGGCGGTGCACTGGCCTGACCCTGACGCTGAGGCGCGACAGCGTGGGG |
| P-asv6480 | GCAAGCGTTGTTCGGAATTACTGGGCGTAAAGGGCTCGTAGGCGGCCAACTAAGTCACACGTGAAATCCCTCGGCTCGACCGGGGAACTGCGTGTGAGACTGGATGGCTTGAGTTTGGGAGAGGAATGCGGAATTCCAGGTGTAGCGGTGAAATGCGTAGATATCTGGAGGAACACCGGTGGCGAAGGCGGCATTCTGGACCAACACTGACGCTGAGGAGCGAAAGCCAGGGG |
| P-asv6481 | ACAAGCGTTGTCCGGATTTATTGGGCGTAAAGGGAGCGCAGGCGGTCTGTTTAGTCTAATGTGAAAGCCCACGGCTTAACCGTGGAACGGCATTGGAAACTGACAGACTTGAATGTAGAAGAGGAAAATGGAATTCCAAGTGTAGCGGTGGAATGCGTAGATATTTGGAGGAACACTAGTGGCGAAGGCGATTTTCTGGTCTAACATTGACGCTGAGGCTCGAAAGCGTGGGG |
| P-asv6482 | GCAAGCGTTAATCGGAATTACTGGGCGTAAAGCGTGCGCAGGCGGCTGATTAAGTCAAATGTGAAAGCCCCGGGCTTAACCTGGGAATGGCATTCGATACTGGTCAGCTAGAGTTTGGTAGAGGTAAGCGGAATTCCGGGTGTAGCGGTGAAATGCGTAGATATCCGGAAGAACACCAGTGGCGAAGGCGGCTCACTGGCTCGATACTGACGCTGAGGTGCGAAAGTGTGGGG |
| P-asv6483 | GCTAGCGTTATCCGGATTTACTGGGCGTAAAGGGTGCGTAGGTGGTCTTTCAAGTCGGTGGTTAAAGGCTACGGCTCAACCGTAGTTAGCCTCCGAAACTGGAAGACTTGAGTGCAGGAGAGGAAAGTGGAATTCCCAGTGTAGCGGTGAAATGCGTAGATATTGGGAGGAACACCAGTAGCGAAGGCGGCTTTCTGGACTGCAACTGACACTGAGGCACGAAAGCGCGGGT |
| P-asv6484 | GCTAGCGTTGCTCGGAATCACTGGGCGTAAAGGGCGCGTAGGCGGCTGACTTAGTCGAGGGTGAAAGCCCGTGGCTCAACCACGGAATGGCCTTCGATACTGGTTGGCTTGAGACCGGAAGAGGACAGCGGAACTGCGAGTGTAGAGGTGAAATTCGTAGATATTCGCAAGAACACCAGTGGCGAAGGCGGCTGTCTGGTCCGGTTCTGACGCTGAGGCGCGAAAGCGTGGGG |
| P-asv6485 | GCAAGCGTTAATCGGAATTACTGGGCGTAAAGCGCGCGTAGGCGGGCATTTAAGTTGGGTGTGAAAGCCCCGGGCTTAACCTGGGAATTGCACTCAAGACTGGATGCCTGGAGTATGGTAGAGGAAAGTGGAATTCCCGGTGTAGCGGTGAAATGCGTAGATATCGGGAGGAACACCAGTGGCGAAGGCGGCTTTCTGGACCAATACTGACGCTGAGGCGCGAAAGCGTGGGG |
| P-asv6486 | GCAAGCGTTGTCCGGAATTATTGGGCGTAAAGCGCGCGCAGGCGGTCCTTTAAGTCTGATATGAAAGCCCACAGCTCAACCGTAGAGGGTCATTAGAAACTGGAAGACTTAAGTGCAGAAGAGAAGAGTAGAATTCCACATGTAGCGGTGAAATGCGTAGAGATGTAGAGGAACACCAGTGGCGAAGGCGACTCTTTGGTCTGTAACTGACACTGAGGCGCGAAAGCGTGGGG |
| P-asv6487 | GCAAGCGTTATCCGGATTTACTGGGTGTAAAGGGAGCGCAGGCGGTGCGGCAAGTCTGATGTGAAAGCCCGGGGCTCAACCCCGGTACTGCATTGGAAACTGTCGTACTAGAGTGTCGGAGGGGTAAGTGGAATTCCTAGTGTAGCGGTGAAATGCGTAGATATTAGGAGGAACACCAGTGGCGAAGGCGGCTTACTGGACGATAACTGACGCTGAGGCTCGAAAGCGTGGGG |
| P-asv6488 | GCAAGCGTTGTTCGGAGTTACTGGGCGTAAAGAGTCTGTAGGTGGTCTGTCAAGTCTTTGGTGAAAGGCCGTGGCTTAACCATGGGAATGCCAAAGAGACTGGCAGACTGGAGGCTGGGAGAGGGAAGCGGAATTTCTGGTGTAGCGGTGAAATGCGTAGATATCAGAAGGAAGGCCGGTGGCGAAGGCGGCTTCCTGGACCAGACCTGACACTGAGAGACGAAAGCGTGGGG |
| P-asv6489 | GCTAGCGTTACTCGGAATTACTGGGCGTAAAGGGCGCGTAGGCGGCGCTGCAAGTTAGGCGTGAAAGTCCTGGGCTCAACCTGGGAACTGCGCTTAAGACTGTGGTGCTTGAGGATGGAAGAGGCTCGTGGAATTCCCAGTGTAGAGGTGAAATTCGTAGATATTGGGAAGAACACCGGTGGCGAAGGCGGCGAGCTGGTCCATTACTGACGCTGAGGCGCGATAGCGTGGGG |
| P-asv6490 | ACGAGCGTTATCCGGAATTATTGGGTTTAAAGGGTGCGTAGGTGGCGTATTAAGTCAGTGGTGAAAAGCTGCAGCTCAACTGTAGTCTTGCCGTTGAAGCTGATATGCTAGAGAGGAGACGAGGTATGCGGAATGTGTGGTGTAGCGGTGAAATGCATAGATATCACACAGAACGCCGATTGCGAAGGCAGCGTACCAGGCTCCGTCTGACACTGAAGCACGAAAGCGTGGGG |
| P-asv6491 | CCAAACGTTATTCGGAATTATTGGGCTTAAAGGGTGCGTAGGCTGTGCAGTAAGTTGGGTGTGAAATCCCTCGGCTCAACCGAGGAACTGCGCCGAATACTGCTGTGCTAGAGGGAGATAGGGGTAAGCGGAACTGATGGTGGAGCGGTGAAATGCGTTGATATCATCAGGAACACCGGTGGCGAAGGCGGCTTACTGGATCTTTTCTGACGCTGATGCACGAAAGCCAGGGG |
| P-asv6492 | GCGAGCGTTGTCCGGAATTACTGGGCGTAAAGGGCGTGTAGGCGGTTTTTTAAGTTAGGAGTGAAAACCCAGGGCTTAACTCTGGGAGTGCTTCTAAAACTGGAAGACTTGAGTACAGGAGAGGAAGGTAGTATTCCTGGTGTAGCGGTGAAATGCGTAGATATCAGGAGGAATACTCGTGGCGAAGGCGGCCTTCTGGCCTGTAACTGACGCTGAGGCGCGAAAGCGTGGGG |
| P-asv6493 | CCAAGCGTTGTCCGGATTTACTGGGTTTAAAGGGTGCGCAGGCGGAAAATTAAGTCAGTGGTGAAAGCCCGCAGCTCAACTGTGGAACTGCCATTGAAACTGGTTTTCTTGAATATAGCTGAGGCAGATGGAATATAACATGTAGCGGTGAAATGCTTAGATATGTTATAGAACACCGATTGCGAAGGCAGTCTGCTAAACTATTATTGACGCTGAGGCACGAAAGCGTGGGG |
| P-asv6494 | GCGAGCGTTATCCGGAATCATTGGGCGTAGAGAGGGAGCAGGCGGCCGCAAGGGTCTGTGGTGAAAGACCGAAGCTAAACTTCGGTAAGCCATGGAAACCGGGCGGCTAGAGTGCGGAAGAGGATCGTGGAATTCCATGTGTAGCGGTGAAATGCGTAGATATATGGAGGAACACCAGTGGCGAAGGCGACGGTCTGGGCCGCAACTGACGCTCATTCCCGAAAGCGTGGGG |
| P-asv6495 | ACAAGCGTTGTCCGGATTTGTTGGGCGTAAAGGGAGCGCAGGCGGTTTGTTAAGTCTGATGTGAAAGCACACGGCTCAACCGTGGAACGGCATTGGAAACTGGCAAACTTGAGTATAGAAGAGGAAAGTGGAATTCCATGTGTAGCGGTGAAATGCGTAGATATATGGAGGAACACCGGTGGCGAAGGCGACTTTCTGGTCTATAACTGACGCTGAGGCTCGGAAGCGTGGGG |
| P-asv6496 | GCAAACGTTGTTCGGAATTACTGGGCGTAAAGCGCGTGTAGGCTGCTTGGAAAGTCGGATGTGAAAGCCCTGGGCTCAACCTAGGAAGTGCATTCGAAACTTCCAAGCTCGAGTTTCGGAGAGGAAGGCGGAATTTTCGGTGTAGAGGTGTAATTCGTAGATATCGAGAGGAACATCGGTGGCGAAGGCGGCCTTCTGGACGATGACTGACGCTGAGACGCGAAAGCGTGGGG |
| P-asv6497 | GCAAGCGTTATCCGGATTTATTGGGTTTAAAGGGTCCGTAGGCGGACTTATAAGTCAGTGGTGAAAGCCTGTCGCTTAACGATAGAACTGCCATTGATACTGTAAGTCTTGAGTATATTTGAGGTAGCTGGAATAAGTAGTGTAGCGGTGAAATGCATAGATATTACTTAGAACACCAATTGCGAAGGCAGGTTACCAAGATATAACTGACGCTGAGGGACGAAAGCGTGGGT |
| P-asv6498 | GCAAGCGTTACCCGGAATCAATGGGCGTAAAGGGCGTGTAGGCGGATCAGTCAGTCTGGCTTTAAAGACCGGGGCTCAACCCCGGGAGTGGGCTGGATACTGCTGATCTAGACCTCTGGAGAGAGAACCGGAATTCCTGGTGTAGCGGTGGAATGCGTAGATACCAGGAAGAACACCGATGGCGAAGGCAGGTTCTTGGACAGAAGGTGACGCTGAGGCGCGAAAGTGTGGGG |
| P-asv6499 | GCAAGCGTTACTCGGAATCACTGGGCGTAAAGCGCGCGCAGGCGGTCTTTTAAGTTGGATGTGAAAGCCTACGGCTCAACCGTAGAACTGCATCCAAAACTATTAGACTAGAGTGTGGGAGAGGAAGATGGAATTAGTTGTGTAGGGGTAAAATCCGTAGAGATAACTAGGAATACCAAAAGCGAAGGCAATCTTCTGGAACACTACTGACGCTGAGGCGCGAAAGCGTGGGG |
| P-asv6500 | CCGAGCGTTATTCGGAATCACTGGGCGTAAAGAGTACGTAGGCGGTTTGGTAAGTCAGGTGTGAAATCCCGGAGCTCAACTCCGGAACTGCACCCGATACTGCTAGACTAGAGTAATGGAGAGGCAGGTGGAATTCTCGGTGTAGCAGTGAAATGCGTGGATATCGAGAGGAAGACCAATGGCGAAGGCAGCCTGCTGGACATTTACTGACGCTGAGGTACGAAGGCCAGGGT |
| P-asv6501 | GCGAGTCTTATTTAGATTGAATGGGTGTAAAGGGTAATAAGGCGGTTTAGAAAAGTTGAAAGTGAAAGAGTCACAAGAAGTGTCAACAGCTTTCAAAACTATCAGACTAGAGGATCTGAGAGGACTATAGAATTTCGTAAGGAGCGATAAAATGCGAAGAAATACGAAGGAATATCAATGGCGAAGGCAATAGTCTAGCAGATAACTGACGCTGGGTCACGAAGGTTTGGGG |
| P-asv6502 | GCAAGCGTTATCCGGATTCATTGGGTTTAAAGGGTGCGCAGGCGGAATGATAAGTCAGTGGTGAAATCTCCCGGCTTAACTGGGAAACTGCCATTGATACTATCGTTCTTGAGTACAGTTGAGGTGGGCGGAATGTGTCATGTAGCGGTGAAATGCATAGATATGACACAGAACACCGATAGCGAAGGCAGCTCGCCAAATTGTAACTGACGCTCATGCACGAAAGCGTGGGG |
| P-asv6503 | GCAAGCGTTGTCCGGAATTATTGGGCGTAAAGAGCTCGTAGGCGGTTTGTCGCGTCTGCTGTGAAAACGCGGGGCTCAACTCCGCGCGTGCAGTGGGTACGGGCAGACTAGAGTGCAGTAGGGGAGACTGGAATTCCTGGTGTAGCGGTGAAATGCGCAGATATCAGGAGGAACACCGATGGCGAAGGCAGGTCTCTGGGCTGTAACTGACGCTGAGGAGCGAAAGCATGGGG |
| P-asv6504 | GCGAGCGTTACCCGGAATGACTGGGCGTAAAGGGTCCGTAGGTTGTATTTTTTGTTAAAGGTGAAAGATAAAAGCTTAACTTTTTTTAAGCCTTTAAAACGTTTATACTTTGAGTTTGAAAGAGGATAGCAGAATTTCACATGGAGGGGTGATATCCTTAGAGATGTGAAGGAATACCAGAGGCGAAGGCGGCTATCTAGTTCAATACTGACACTGAGGGACGAAAGCATGGGT |
| P-asv6505 | TCGAGTGATGGCCGATCTTATTGGGCCTAGAGCGTCCGTAGCTGGCCTGGCAAGTTCGTCGGGAAATCCACCCGCTCAACGGGAGGGCGTCCGGTAGAAACTGCCCGGCTTGGAACCGGAAGGCTCAGAGAGTACGTCCGGGGTAGGAGTGAAATCCCGTAATCCCGGACGGACTACCGATGGCGAAAGCACTCTGAGAAGACGGCTTCGACAGTGAGGGACGAAAGCTAGGGT |
| P-asv6506 | GCAGGCGTTGTCCGGATTTATTGGGTTTAAAGGGTGCGTAGGCGGGTCATTAAGTCAGTGGTGAAAGCCGACAGCTCAACTGTCGAACTGCAATTGATACTGGTGACCTTGAGTGTAGTTGAGGCAGGCGGAATTTACGATGTAGCGGTGAAATGCATAGATATCGTAAAGAACACCTATAGCGAAGGCAGCTTGCTAAGCTACAACTGACGCTGAGGCACGAAAGCATGGGG |
| P-asv6507 | GCGAGCGTTAATCGGAATTACTGGGCGTAAAGCGCATGCAGGTGGTTTGTTAAGTCAGATGTGAAAGCCCGGGGCTCAACCTCGGAATAGCATTTGAAACTGGCAGACTAGAGTACTGTAGAGGGGGGTAGAATTTCAGGTGTAGCGGAGAAATGCGTAGAGATCTGAAGGAATACCGGTGGCGAAGGCGGCCCCCTGGACAGATACTGACACTCAGATGCGAAAGCGTGGGG |
| P-asv6508 | CCGAGCGTTATCCGGATTTATTGGGCGTAAAGCGAGCGCAGACGGTTTATTAAGTCTGATGTGAAAGCCCGGAGCTCAACTCCGGAATGGCATTGGAAACTGGTTAACTTGAGTGCAGTAGAGGTAAGTGGAACTCCATGTGTAGCGGTGGAATGCGTAGATATATGGAAGAACACCAGTGGCGAAGGCGGCTTACTGGACTGCAACTGACGTTGAGGCTCGAAAGTGTGGGT |
| P-asv6509 | GCAAGCGTTACTCGGAATCACTGGGCGTAAAGAGCGTGTAGGCGGATAGATAAGTCAGAAGTGAAATCCAATAGCTTAACTATTGAACTGCTTTTGAAACTGTCTATCTAGAGTATGGGAGAGGTAGATGGAATTTCTGGTGTAGGGGTAAAATCCGTAGAGATCGGAAGGAATACCGATTGCGAAGGCGATCTACTGGAACATAACTGACGCTGAGACGCGAAAGCGTGGGG |
| P-asv6510 | GCAAGCGTTATCCGGAATTATTGGGCATAAAGGGCATCTAGGCGGATATACAAGTCAGGGGTGAAAACTTAGGGCTCAACTCAAAGCTTGCCCTTGAAACTGTATATCTAGAGTGCTGGAGAGGTGGACGGAACTACACGAGTAGAGGTGAAATTCGTAGATATGTGTAGGAATGCCGATGATGAAGATAGTCCACTGGACAGCAACTGACGCTGAAGTGCGAAAGCTAGGGG |
| P-asv6511 | GCAAGCGTTACTCAGAATTACTGGGCGTAAAGCGTGCGTAGGTGGTTTGTTAAGTCTGATGTGAAAGCCCTGGGCTCAACCTGGGAATGGCATTGGATACTGGCTTACTAGAGTGCGGTAGAGGGTGGCGGAATTCCCGGTGTAGCAGTGAAATGCGTAGATATCGGGAGGAACATCCGTGGCGAAGGCGGCCACCTGGACCAGCACTGACACTGAGGCACGAAAGCGGGGGG |
| P-asv6512 | GCAAGCGTTAATCGGAATTATTGAGCGTAAAGCGAGCGTAGGTGGCTTGATAAGTCAGATGTGAAAGCCCCGGGCTCAACCTGGGAACGGCATCTGATACTGTTAGGCTAGAGTAGGTGAGAGGAAGGTAGAATTCCAGGTGTAGCGGTGAAATGCGTAGAGATCTGGAGGAATACCGATGGCGAAGGCAGCCTTCTGGCATCATACTGACACTGAGGTTCGAAAGCGTGGGT |
| P-asv6513 | GCAAGCGTTAATCGGAATTACTGGGCGTAAAGCGTGCGCAGGCGGTTCGGAAAGAAAGATGTGAAATCCCAGAGCTCAACTTTGGAACTGCATTTTTAACTACCGAGCTAGAGTGTGTCAGAGGGAGGTGGAATTCCGCGTGTAGCAGTGAAATGCGTAGATATGCGGAGGAACACCGATGGCGAAGGCAGCCTCCTGGGATAACACTGGCGCTCAGGCACGAAAGCGTGGGG |
| P-asv6514 | GCAAACGTTGCTCGGAATCATTGGGCGTAAAGCGCACGTAGGCGGTTCGTTAAGTCGGGTGTGAAAGTCCAGGGCCCAACCCTGGAAGTGCACTCGAAACTGGCGAACTTGAATATGGAAGAGGGTCGCGGAATTCCCGGTGTAGAGGTGAAATTCGTAGATATCGGGAGGAACACCAGTGGCGAAGGCGGCGACCTGGGCCAATATTGACGCTGAGGTGCGAAAGCGTGGGG |
| P-asv6515 | GCTAGCGTTGTTCGGAATCACTGGGCGTAAAGGGCGCGTAGGCGGCTGTCTAAGTCGGAGGTGAAAGCCCGGGGCTCAACCCCGGAACGGCCTTCGAGACTGGGCGGCTCGAGACCGGAAGAGGAGGGTGGAATACCCAGTGTAGAGGTGAAATTCGTAGATATTGGGTGGAACACCAGTGGCGAAAGCGGCTCTCTGGTCCGGATCTGACGCTGAGGCGCGAAAGCGTGGGG |
| P-asv6516 | GCAAGCGTTATCCGGATTCACTGGGCGTATAGAGCGTTGAGGCGGTCCTGTAAGTTGGGCGTGAAAGTCCCGGGCCTAACCCGGGGAGGCCGTTCAATACTGCAGGACTTGAGGACAGCAGAGGGAGGTGGAATTCCCGGTGTAGCGGTGAAATGCGTAGATATCGGGAGGAACACCAGTGGCGAAGGCGGCCTCCTGGGCTGTTCCTGACGCTGAATGCGAAAGCTAGGGG |
| P-asv6517 | GCAAGCGTTATTCGAAATTACTGGGCGTAAAGCGTATGTAGGCGGACTAATAAGTCAGGTGTGAAATCCTCGGTTTCAAGCTGAGAATTGCACTTGATACTGTTAGTCTTGAGTAAGATAGAGGACAATAGAATTCCGAGTGTAGAGGTGATATTCGTAGATATTCGGAGGAATACTAATGGCGAAGGCAGTTGTCTGGGTCTTTACTGACGCTGAGATACGAAAGCGTGGGG |

| | |
|---|---|
| P-asv6518 | CCTAGCGTTGTTCGGAATTATTGGGCGTAAAGCGGGTGTAGGTGGCTATGTAAGTCAAGTGTGAAAGCCCGGGGCTCAACCCCAGAAGTGCATTTGATACTGCGTAGCTTGAGTGCTGGAGAGGTTGCTAGAATT<br>GTTGGTGTAGTGGTGAAATACGTAGATATCAACAGGAATACCGGTGGCGAAGGCGGGTAACTGGCCAAGCACTGACACTCAGACCCGAAAGCGTGGGG |
| P-asv6519 | ACGAGCGTTATTCGGAGTTAATGGGCGTAAAGGGTATGTAGGTGGTCATTCGATGTAACTGAAAAAAGCTAGGGCGATCCCTTAGACAACCGGTTTCAAAGTCTGTGACTTGGGGAAAAAAGAGGTAGGGAGAAT<br>TTCCAGAGTAGAAGTCGAATTCTTAGATACTGGAAGGACTATCATTAGGCGAAGGCACCTTTCCATTTTTTTCCGACGCTGAGGTACGAAGGCATGGGG |
| P-asv6520 | GCAAGCGTTGTTCGGAATTACTGGGCGTAAAGGGCGAGCAGGCGGTCTGCTAAGTCTACTGTGAAAATCCGGAGCTCAACTCCGGAAATGCAGTAGATACTGGCAGTCTAGAGACCAGTAGAGGCAAGCGGAAT<br>TCCTGGTGTAGCGGTGGAATGCGTAGATATCAGGAAGAACACCTGTGACGAAGGCGGCTTGCTGGGCTGGATCTGACGCTGATCCGCGAAAGCGTGGGG |
| P-asv6521 | GCAAACGTTATTCGGAATCACTGGGCTTAAAGGGTGCGTAGGCGGCGATGCAGGTGGGGTGTGAAATCCCTCGGCTCAACCGAGGAACTGCGCTCCAAACCACATGGCTGGAGGGAGACAGAGGTGAGCGGAA<br>CTGATGGTGGAGCGGTGAAATGCGTTGATATCATCAGGAACACCGGTGGCGAAGGCGGCTCACTGGGTCTCTTCTGACGCTGAGGCACGAAAGCTAGGGG |
| P-asv6522 | GCAAGCGTTGTTCGGAATTACTGGGCGTAAAGCGCAGGTAGGCGGCCTGCTAAGTCTTCTGTGAAACTCCGGGGCTCAACTCCGGACATGCAGGAGATACTGGCAGGCTCGAGTCCTGTAGAGGCTAGCGGAAT<br>TCCTGGTGTAGCGGTGGAATGCGTAGATATCAGGAAGAACACCTGTGGCGAAGGCAGCTAGCTGGGCAGGAACTGACGCTGATCTGCGAAAGCGTGGGG |
| P-asv6523 | GCAAGCGTTGTTCGGAATCACTGGGCGTAAAGCGCGTGTAGGCGGTTCGACAAGTCGGATGTGAAAGCCCGGGGCCCAACCTCGGAAGTGCATTCGAAACTGTCGGGCTAGAGTACGGGAGAGGGAAGTGGAA<br>TTCCTGGTGTAGAGGTGAAATTCGTAGATATCAGGAGGAACACCGGTGGCGAAGGCGGCTTCCTGGACCGATACTGACGCTGAGACGCGAAAGCGTGGGG |
| P-asv6524 | GCTAGCGTTGTTCGGAATTACTGGGCGTAAAGCGTGCGCAGGCGGCTATGCAAGTCAGGGGTGAAATGCCAAGGCTCAACCTTGGAACTGCCTTTGAAACTGCATAGCTAGAGACTCGGAGGGGTTAGGGGAAT<br>TCCTAATGTAGCGGTGAAATGCGTAGATATTAGGAGGAACACCAGTGGCGAAGGCGCCTAACTGGACGAGTACTGACGCTCATGCACGAAAGCGTGGGG |
| P-asv6525 | GCGAGCGTTAATCGGAATAACTGGGCGTAAAGGGTGAGTAGGCGGAAATATAAGTTAGATGTGAAAGTCCCGAGCTTAACTTGGGGAGGCCATTTAATACTGTATTTCTGGAGTTAGTAAGAGGGTGGTAGAATT<br>CCTGGTGTAGCAGTGAAATGCGTAGATATCAGGAGGAATACCGAAGGCGAAGGCAGCCACCTGGGATATAACTGACGCTGAGTCACGAAAGCGTGGGT |
| P-asv6526 | GCGAGCGTTAATCGGAATTACTGGGCGTAAAGGGTGCGTAGGTGGTTTGATAAGTTGGCTGTGAAATACCGGGGCTTAAGCTTGGGGCGTCAGCCAAGACTGTCAGGCTTGAGTATAGGAGAGGGCAGTGGAAT<br>TTCCGGTGTAGCGGTGAAATGCGTAGAGATCGGAAGGAACACCAGTGGCGAAGGCGTCTGCCTGGCCTAATACTGACACTGAGGCACGAAAGCGTGGGT |
| P-asv6527 | TCAAGCGTTGTTCGGAATCACTGGGCGTAAAGGGTGCGTAGGTGGCGTGGTAAGTCAGATGTGAAAGCCCGGGGCTCAACCTCGGAACTGCATCCGATACTGCTGTGCTAGAGTACTGGAGAGGTGACTAGGAT<br>TCCCGGTGTAGCAGTGAAATGCGTAGATATCGAGAGGAATACCAAAGGCGTAGGCAGGTCACTGGACAGTTACTGACACTGAGGCACGAAGGCCAGGGG |
| P-asv6528 | TCAAGCGTTATCCGGAATCATTGGGTTTAAAGGGTTCGTAGGCGGGACAATAAGTCAGTGGTTAAAGTTTGTGGCTCAACCATAAAATTGCCATTGATACTGTTGTTCTTGAGTTATTGTGAAGTGGTTAGAATGTG<br>TAGTGTAGCGGTGAAATGCATAGATATTACACAGAACACCAATTGCGAAGGCATATCACTAACAATATACTGACGCTGATGAACGAAAGCGTAGGT |
| P-asv6529 | CCGAGTGATGGCCGATATTATTGGGCCTAAAGCGTCCGTAGCTGGCCGAGCAAGTTCGTTGGGAAATCCACGCGCCTAACGTGTGGGCGTCCGGCGAAAACTGTTCGGCTTGGGACCGGAAGACCCGAGGGGT<br>ACGTCCGGGGTAGGAGTGAAATCCCGTAATCCTGGACGGACCGCCGGTGGCGAAAGCGCCTCGGGAAGACGGATCCGACAGTGAGGGACGAAAGCTAGGGT |
| P-asv6530 | GCGAGCGTTAATCGGATTTACTGGGCGTAAAGCGTGCGTAGGCGGCTTCTTAAGTCGGATGTGAAATCCCTGAGCTTAACTTAGGAATTGCATTCGATACTGGGAAGCTAGAGTATGGGAGAGGATGGTAGAATT<br>CCAGGTGTAGCGGTGTAATGCGTAGAGATCTGGAGGAATACCGATGGCGAAGGCAACCTCCTGGGCCTGAATTGACACTGAGACACGAAAGCGTGGGG |
| P-asv6531 | GCAAGCGTTGTTCGGATTCACTGGGCGTACAGGGAGCGTAGGCGGTTGGGTAAGCCCTCCGTGAAATCTCCGGGCCTAACCCGGAAAGTGCAGAGGGGACTACTCGGCTAGAGGATGGGAGAGGAGCGCGGA<br>ATTCCCGGTGTAGCGGTGAAATGCGTAGAGATCGGGAGGAAGGCCGGTGGCGAAGGCGGCGCTCTGGAACATTTCTGACGCTGAGGCTCGAAAGCGTGGGG |
| P-asv6532 | GCTAGCGTTGTTCGGAATCACTGGGCGTAAAGGGCGCGTAGGCGGCTGTTTAAGTCGGAGGTGAAAGCCCGGGGCTCAACCTCGGAACGGCCTTCGAGACTGGACAGCTAGAGACCGGAAGAGGAGGGTGGA<br>ATACCCAGTGTAGAGGTGAAATTCGTAGATATTGGGTAGAACACCAGTGGCGAAAGCGGCTCTCTGGTCCGGTTCTGACGCTGAGGCGCGAAAGCGTGGGG |
| P-asv6533 | GCGAGCGTTGTTCGGAATTACTGGGCGTAAAGCGCGTGTAGGCGGCCATGTAAGTCAGATGTGAAAGCCTGGGGCTCAACCCCAGAAGTGCATCTGAAACTATGTGGCTTGAGTGTGGAAGAGGGCAGTGGAAT<br>TCCTGGTGTAGTGGTGAAATACGTAGATATCAGGAGGAACATCGGTGGCGAAGGCGGCTGCCTGGTCCAACACTGACGCTGAGACGCGAAAGCGTGGGT |
| P-asv6534 | GCAAGCGTTGTTCGGAATTATTGGGCGTAAAGCGCGTGTAGGCGGTTTGCTAAGTCTGGTGTGAAAGCCCGAGGCTTAACTTCGGAAGTGCACTGGATACTGGCAAACTAGAGGACGAGAGAGGAAAGTGGAAT<br>TCCTGGTGTAGGGGTGAAATCCGTAGATATCAGGAGGAACATCAATGGCGAAGGCAGCTTTCTGGCTCGTATCTGACGCTGAGACGCGAAAGCGTGGGT |
| P-asv6535 | TCGAGTGATGGCCGATCTTATTGGGCCTAAAGCGTCCGTAGCTGGCCGTGCAAGTCCGTCGGGAAATCCACGTGCTCAACGCGTGGGCGACCGGTGGAAACCATCCGGCTTGGGGCCGGAAGACCCAGGGGG<br>TACGTCCGGGGTAGGAGTGAAATCCCGTAATCCTGGAGGGACCGCCGATGGCGAAAGCACCCTGGGAAGACGGACCCGACGGTGAGGGACGAAAGCTGGGGT |
| P-asv6536 | GCAAGCGTTATTCGGAATCATTGGGCGTAAAGGGTGCGTAGACGGTGTGTTAAGTCTGTTGTTAAATTTTCCGGCCTAACCGGAAATCTGCGGTGGAAACTGGGGCACTAGAGGGTGGAAGAGAGAAGTGGAAT<br>TCTCGGAGTAGCGGTAAAATGCGTGGATCTCGAGAGGAACACCAATGGCGAAAGCAGCTTCTTGGTCCATACCTGACGTTGATGCACGAAAGCGTGGGG |
| P-asv6537 | CCAAGCGTTGTTCGGAATTACTGGGCGTAAAGCGTCTGTAGGCGGCGCGGTAAGTCAGTTGTGAAATCCCGGGGCTCAACCTCGGAACTGCAGCCGATACTGCCGTGCTAGAGGGTCGGAGGGGAGTCTGGAA<br>TTCTCGGTGTAGCAGTGAAATGCGTAGATATCGAGAGGAACACTAGTGGCGAAGGCGAGACTCTGGACGACACCTGACGCTGAGAGACGAAGGCCAGGGT |
| P-asv6538 | GCAACCGTTGTTCGGAATCACTGGGCTTAAAGGGCACGTAGGCGGAATGCTAAGTTAGATGTGAAAGCCCATGGCTTACCCATGGAATTGCATTTAAAACTGGTGTTCTTGAGAGTAGAAGGGGAGAGTGGAACC<br>CCTGGTGGAGCGGTGAAATGCGTTGATATTAGGAGGAACGCCGGTGGCGAAAGCGACTCTCTGGTCTGCGTCTGACGCTGAGGTGCGAAAGCTAGGGG |
| P-asv6539 | CCAAGCGTTATCCGGATTTATTGGGCGTAAAGGGCTCGTAGGAGGTTATGTGCGTCTCGTGTTAAATCGTACCGCTCAACGGTACGGCTGCGCGGGATACGGCATAACTAGAGGATGTCAGAGGTAAGCGGAAC<br>GCACGGTGTAGGGGTGAAATCCGTTGATATCGTGCGGAACACCAGAGGCGAAGGCGGCTTACTGGGGCACTCCTGACTCTGAGGAGCGAAAGCATGGGT |
| P-asv6540 | GCGAGCGTTGTCCGGATTCACTGGGCGTAAAGGGCTCGTAGGCGGCCTGTCGCGTCGGCTGTGAAAGTCCGGAGCTCAACTCCGGGATTGCAGCCGATACGGGCAGGCTAGAGGGCGGTAGAGGGGAGTGG<br>AATTCCCGGTGTAGCGGTGAAATGCGCAGATATCGGGAGGAACACCAGTGGCGAAGGCGACTCACTGGTCTGTAACTGACGCTGAGGCGCGAAAGCGTGGGG |
| P-asv6541 | GCAAGCGTTAATCAGAATTACTGGGCGTAAAGCGCGCGTAGGTGGTTTGTTAAGTTGAATGTGAAAGCCCCGGGCTCAACCTAGGAACTGCATTGGATACGGTCTGGCTAGAGTCCGCGAGAGGAAGATGGAAT<br>TGTGCGTGTAGAGGTGAAATTCGTAGATATGCACAAGAACACCGGTGGCGAAGGCGATCTTCTGGAGCGGTACTGACGCTAAGGCACGAAAGTGTGGGG |
| P-asv6542 | GCAAGCGTTGTTCGGAATTATTGGGCGTAAAGCGCGTGTAGGCGGCTATCTAAGTCTGGTGTGAAAGCCCCGGGCTCAACCCGGGAAGTGCACTGGATACTGGATAGCTAGAGGGCGAGAGAGGAGAATGGAA<br>TTCCTGGTGTAGGGGTGAAATCCGTAGATATCAGGAGGAACATCAGAAGCGAAGGCGGTTCTCTGGCTCGCAACTGACGCTGAGACGCGAAAGCGTGGGG |
| P-asv6543 | CCGAGCGTTATCCGGAATTATTGGGCGTAAAGGGTCCGCAGGCGGTCCTATTAGTCTTGTGTGAAATCTCCCGGCTTAACTGGGAAAACGCATAGGAAACGGTAGGACTAGAGGATGTGAGAGGTATGTGGAACT<br>CATGGTGTAGGGGTGAAATCCGTTGATATCATGGGGAACACCAAAAGCGAAGGCAACATACTGGCACATTGCTGACGCTCAGGGACGAAAGCATGGGT |
| P-asv6544 | CCAAGCGCTAATCGGAATTACTGGGCGTAAAGCGTGCGCAGGCGGTTGTGCAAGACCGATGTGAAATCCCCGAGCTTAACTTGGGAATTGCATTGGTGACTGCTCATCTAGAGTGTGGGAGAGGTAGATGGAAT<br>TTCTGGTGTAGGGGTAAAATCCGTAGAGATCAGAAGGAATACCGATTGCGAAGGCGATCTACTGGAACATAACTGACGCTGAGACGCGAAAGCGTGGGG |
| P-asv6545 | GCGAGCGTTAATCAGAATTACTGGGCGTAAAGGGTGCGTAGGTGGTTAAATAAGTTACATGTGAAATTCCTGAGCTCAACTTGGGGACTGCGTGTAATACTGTCTGACTAGAGTACTGTAGAGGGAAGTGGAACT<br>GCGAGTGTAGAGGTGAAATTCGTAGATATTCGCAAGAACACCAGTGGCGAAGGCGACTCACTGGCCCGATACTGACGCTGAGGCACGAAAGCGTGGGG |
| P-asv6546 | GCGAGCGCTAATCAGAATTACTGGGCGTAAAGCGCATGCAGGTGGTTTGTCAAGTCAGAAGTGAAAGCCTGGGGCTCAACCCCAGAACTGCTTTTGAAACTGGATGGCTAGAGTCCCGGAGAGGTTGGCGGAAT<br>TCCGAGTGTAGAGGTGAAATTCGCAGATATTCGGAGGAACACCAGTGGCGTAGGCGGCCAACTGGACGGGAACTGACGCTGAGGTACGAAAGCGTGGGG |
| P-asv6547 | GCAAGCGTTAATCAGAATTACTGGGCGTAAAGGGTGCGTAGGCGGCATGACAAGCTAGATGTGAAAGCCCTGGGCTTAACCTGGGAAGTGCATCTGGGACTGTTGAGCTAGAGTATGGTAGAGGAAAGTGGAAT<br>TTCTAGTGTAGCGGTGAAATGCGTAGATATTAGAAGGAACACCAGAGGCGAAGGCGACTTTCTGGACCGATACTGACGCTGAGGCACGAAAGCGTGGGG |
| P-asv6548 | GCTAGCGTTGTCCGGAATCATTGGGCGTAAAGCGCGTGTAGGCGGTTCGATTAGTCCGCTGTAAAAGTCGGGGGCTCAACCCTCGAAAGCCGGTGGATACTGTCGGGCTAGAGTGCGGAAGAGGCGAGTGGAA<br>TTCCTGGTGTAGCGGTGAAATGCGCAGATATCAGGAAGAACACCAATTGCGAAGGCAGCTCGCTGGGACGTTACTGACGCTGAGACGCGAAAGCGTGGGG |
| P-asv6549 | GCGAGCGTTAATCGGAATTACTGGGCGTAAAGGGTGCGTAGGTTGTTGAGTAAGTTATCTGTGAAATTCCTGGGCTCAACCTAGGGTGGTCAGATAAGACTGCATGACTCGAGTATAGGAGAGGGTAGTGGAATT<br>TCCGGTGTAGCGGTGAAATGCGTAGAGATCGGAAAGAACACCGATGGCGAAGGCAATCCCCTGGGCCTGCACCGACGCTCATGCACGAAAGCGTGGGG |
| P-asv6550 | GCGAGCGTTAATCGGAATTACTGGGCGTAAAGCGCGCGTAGGCGGTTTGATAAGCTGGATGTGAAAGCCCCGGGCTCAACCTGGGAACTGCATCCAGAACTGTCTGACTAGAGTACGGTAGAGGTGAGTGGAAT<br>TTCCTGTGTAGCGGTGAAATGCGTAGATATGCGGAGGAACACCGATGGCGAAGGCAATCCCCTGGGCCTGCACTGACGCTCATGCACGAAAGCGTGGGG |
| P-asv6551 | GCAAGCGTTACTCGGATTTACTGGGCGTAAAGCGCGCGCAGGCGGATCGGCAAGTCAGATGTGAAAGCCCGGGGCTCAACCTCGGGGTAGCATTTGAAACTGGCAGACTAGAGTACTGTAGAGGGGGGTAGAA<br>TTTCAGGTGTAGCGGTGAAATGCGTAGAGATCTGAAGGAATACCGGTGGCGAAGGCGGCCCCCTGGACAGATACTGACACTCAGATGCGAAAGCGTGGGG |
| P-asv6552 | GCTAGCGTTGTTCGGAATTACTGGGCGTAAAGCGCGCGTAGGCGGCGGCACAAGTCAGGGGTGAAATCCCACAGCTCAACTGTGGAATTGCCTTTGAAACTAGGTCGCTAGAGTGTCGGAGGGGATAGCGGAA<br>TTGCTAATGTAGAGGTGAAATTCGTAGATATTCGCAAGAACACCAGTGGCGAAGGCGGCTCACTGGCCCGATACTGACGCTGAGGCACGAAAGCGTGGGG |
| P-asv6553 | GCAAGTGTTATCCGGAATTATTGGGCGTAAAGGGTCTCTAGGTTGAAATTTAAGTCTTTTGTTAAATATCAGGGCTTAACCCTGACTCTGCAAAAGAAACTAGGTTTCTAGAGTATGGTAGAGGTTAAGGGAATTTC<br>CAGTGGAGCGGTGAAATGCGTAGATATTGGAAGGAACACCAAGAGCGAAGGCACTTAACTGGGCCATTACTGACACTGACAGACGAAAGCTAGGGG |
| P-asv6554 | GCAAGCGTTATCCGGATTTACTGGGTGTAAAGGGTGCGTAGCCGGAAATGTAAGTCAGATGTGAAAGCCCGAAGCTCAACTTCGGGACTGCATTTGAAACTGCATTTCTTGAGTGCAGGAGAGGAAAGTGGAATT<br>CCTAGTGTAGCGGTGAAATGCGTAGATATTTGGAGGAACACCAGTGGCGAAGGCGACTTTCTGGACTGTAACTGACGGTGAGGCACGAAAGCGTGGGG |
| P-asv6555 | ACAAGTGGGGATCGTATTTATTTGGCCTAAAGCGCCCGTAGCCGGGCAGCTAAATCTTTTGTGAAATCGTTGCGCTTAACGTAACGTCGTGCAGAAGACACTGGCTGCCTAGGGACTGGGAGGCGTCAGATGTAT<br>GCCGGGGGGAGCAGTAAAATGCTATAATCCCCGGTAGACAACCGGTGGCGAAGGCGTCTGACGAGAACAGGTCCGACGGTCAGGGGCGAAAGCTAGGGG |
| P-asv6556 | GCTAGCGTTGTTCGGATTTACTGGGCGTAAAGCGCGCGTAGGCGGCTTGGAAAGTTGGGAGTGAAAGCCCGGGGCTCAACCCCGGAACTGCTCTCAAAACTTCCAAGCTAGAGATCTGGAGAGGAGGGTGGAA<br>TTCCAAGTGTAGAGGTGAAATTCGTAGATGTTTGGAAGAACACCAGTGGCGAAGGCGACTCTCTGGCCAGATACTGACGCTGAGGTGCGAAAGCGTGGGG |

| | |
|---|---|
| P-asv6557 | GCAAGCGTTAATCGGAATTACTGGGCGTAAAGCGTACGTAGGCGGCTTGGCAAGTCAGAAGTGAAATCCCAGGGCTTAACCCTGGAACGGCTTTTGAAACTGCCTCGCTAGAATTCGGGAGAGGTTGGCGGAATTCCTAGTGTAGAGGTGAAATTCGTAGATATTAGGAGGAACACCAGTGGCGTAGGCGGCCAACTGGACCGACATTGACGCTGAGGTACGAAAGCGTGGGG |
| P-asv6558 | ACAAGTGGCAACCGCTTTTATTGGGCCTAAAGCGTTCGTAGCCGGCTCAGCAAATTTTCTGTGAAATCGTTGAACTTAATTTAACGACGTGCAGGAAACACTACTAAGCTAGGGACCGGGAGGCGTCAGAGGTATTCTGGAGGGAGCGGTAAAATGCTATAATCTCCAGAGGACCTCCTGTGGCGAAGGCGTCTGACGAGAACGGATCCGACGGTGAGGGACGAAAGCCAGGGG |
| P-asv6559 | GCAAGCGTTGTTCGGAATCATTGGGCGTAAAGCGCGTGTAGGCGGCCTGGCAAGTCGGATGTGAAAGCCCGGGGCCCAACCCCGGAATGTCATTCGATACTGCCTCGCTCGAGTCCCGGAGAGGAGAGCGGAATTCCCGGTGTAGGGGTGAAATCCGTAGATATCGGGAGGAACACCAGTGGCGAAGGCGGCTCTCTGGACGGTGACTGACGCTGAGACGCGAAAGCGTGGGG |
| P-asv6560 | GCTAGCGTTGTTCGGAATTATTGGGCGTAAAGCGTATGTAGGCGGTTTAGTAAGTCGGTTGTGAAATCCCCAGGCTTAACCTGGGAATTGCAGTCGAAACTACTTTGCTTGAATTTGGTGGGGGATAGTGGAATTCTTAGTGTAGGGGTGAAATCCGTAGATATTAAGAGGAACACCGGCGGCGAAAGCGACTATCTACACCAACATTGACGCTGAGATACGAAAGCGTGGGG |
| P-asv6561 | CCAAGCATTATCCGGATTTACTGGGCGTAAAGGATGTGTAGGTGGTGTGATTAGTCTTGCGTTAAATCTTCGGGCTCAACCCGGAACTCGCGTAAGAAACGGTCATGCTAGAGGAAGTGAGAGGTGTGCGGAACTCATGGTGTAGGGGTGAAATCCGTTGATATCATGGGGAACACCAAAAGCGAAGGCAGCACACTGGCACTCTCCTGACACTGAAACATGAAAGCGTGGGT |
| P-asv6562 | GCGAGCGTTGTCCGGAATTACTGGGCGTAAAGCGCACGCAGGCTGAATTATAAGTCGGTAGTAAAAGGCGGAGGCTCAACCTCCGTTTATCTATCGATACTGTAAATCTAGAGTATGTGAGAGGGAAGTGGAATTCCCGGTGTAGCGGTGAAATGCGTAGATATCGGGAGGAACACCAGTGGCGAAGGCGACTTCCTGGCACAATACTGACGCTCATGTGCGAAAGCTAGGGT |
| P-asv6563 | GCCAGCGTTGTTCGGAATTACTGGGCGTAAAGCGCACGTATGCGGCTTTTTAAGTCAGGGGTGAAATCCCGGGGCTCAACCAATGTAAGCCTTTGAAACTGTTGGGCTTGAGTGCAGGAGAGGTAAGTGGAATTCCTAGTGTAGAGGTGAAATTCGTAGATATTAGGAGGAACACCAGTGGCGAAGGCGACTTACTGGACTGTAACTGACGCTGAGGCACGAAAGCGTGGGG |
| P-asv6564 | GCAAGCGTTGCTCGGAATCATTGGGCGTAAAGGGCAGGTAGGTGGTCTCATTTGTCAGGGGTGAAAGCCTTGGGCTTAACCCAAGAAGTGCCTCTGAAACGGTGGGACTAGAGTACTGGAGAGGGTCGTAGAATTCCTGGTGTAGCGGTGAAATGCGTAGAGATCAGGAGGAATACCAGAGGCGAAGGCGGCGACCTGGACAGTGACTGACACTCAACTGCGAAAGCGTGGGG |
| P-asv6565 | GCTAGCGTTGTTCGGAATTACTGGGCGTAAAGAGCTCGTAGGCGGTTCAGTAAGTCGGGTGTGAAAGCTCCGAGCTCAACTCGGAGAGGCCACCCGATACTGCTGTGACTAGAATCCAGTAGGGGAGTGTGGAACTCCTGGTGTAGCGGTGAAATGCGCAGATATCAGGAAGAACACCGACGGCGAAGGCAGCACTCTGGGCTGGTATTGACGCTGAGGAGCGAAAGTGTGGGT |
| P-asv6566 | GTTAGCGTTGTTCGGAATTACTGGGCGTAAAGCGCACGTAGGCGGATCGGCAAGACAGATGTGAAATCCCCGGGCTTAACCTGGGAACTGCGTTTGTGACTGCAAGGCTAGAGTACGGCAGAGGGGGGTGGAATTCCTGGTGTAGCAGTGAAATGCGTAGATATCAGGAGGAACACCGATGGCGAAGGCAGCCCCCTGGGCCTGTACTGACGCTCATGCACGAAAGCGTGGGG |
| P-asv6567 | ACCAGTGTTATTCGGCGTAATTAGGTGTAAAGTGTATGTAGGTTGTTTATTAAAGTTTTTATTTAAAAATTAAATTTTAACTTTATTTATAAAAATATTTAATAAACTAGAGTTAAGTAGAGGATAGTAGAATTTTCGGTGTAGGGGTGAAATCCTTCGATATCGTCAGGAATGCCATCGTGCGAAAGCGACTATCTGGGCTTATACTGACATTGAGATACGTAGGTATGGGT |
| P-asv6568 | GCAAGCGTTACTCGGAATTACTAGGCGTAAAGCGCAGGTAGGTGGTTGGGTAAGTCCGCAATGAAATCTCCCGGCTTAACTGGGAGGGGTTTGTGGATACTGCCTGACTCGAGTGTGGGAGAGGTAACTGGAATTCCTGGTGTAGCGGTGAAATGCGTAGATATCAGGAGGAACACCAATGGCGAAAGCAGGTTACTGGCCCAATACTGACGCTGAGCTGCGAAAGCTAGGGG |
| P-asv6569 | GCGAGCGTTAATCGGAATTACTGGGCGTAAAGCGTGCGCAGGCGGATCGATCAGTCAGAGGTGAAATCCCGGAGCTCAACTTCGGAACTGCCTTTGACACTGTCGATCTGGAGTCCGGGAGAGGTGCATGGAATTCCGAGTGTAGAGGTGAAATTCGTAGATATTCGGAGGAACACCAGAGGCGAAGGCGATGCACTGGACCGGTACTGACGCTGAGGTGCGAAAGCGTGGGG |
| P-asv6570 | GCAAGCGTTATCCGGAATTACTGGGCGTAAAGAGAGTGTAGGTGGTTATGTAAGTCTTATGTGAGATCCCTCGGCTTAACCGAGGAAGTGCATGAGATACTGTGTAACTAGAGTGCAGGAGAGGTAAGTGGAATTCCTAGTGTAGCGGTAGAATGCGTAGAGATTAGGAGGAACACCAGAGGCGAAGGCGACTTACTGGACTGTAACTGACACTGAGACTCGAAAGCGTGGGG |
| P-asv6571 | CCAAGCGTTGTCCGGATTTATTGGGTTTAAAGGGTGCGTAGGCGGATCTTTAAGTCAGTGGTGAAAGCCTGCAGCTTAACTGTAGAACAGCCATTGAAACTGAAGATCTTGAATTTGGTTAAAGTAGGCGGAATGTATCATGTAGCGGTGAAATGCTTAGATATGATACAGAACACCGATAGCGAAGGCAGCTTGCTGAACCAATATTGACGCTGAGGCACGAAAGCGTGGGG |
| P-asv6572 | CCAAGCGTTATCCGGATTTACTGGGCGTAAAGCGTCTGTAGCCTGATATGTAAGTCTGTTTTCAAATACCTGGACTCAATCGAGGGAAGGGGATAGATACTGCATATCTAGAGATATTTGGGGGTTAGTGGAATTTCCGGTGGAGCGGTGAAATGCGTTGATATCGGAAAGAACGCCAAAAGCGAAAGCAGCTAACTACAATACATCTGACGGTGAGAGACGAAAGTTTGGGT |
| P-asv6573 | GCAAGCGTTAATCGGAATTACTGGGCGTAAAGGGCGCGTAGGCGGATGAATAAGTTAGGTGTGAAATCCCCGGGCTTAACCTGGGAATGGCATTTAATACTGTTTATCTGGAGTAGAGTAGAGGGAAGTGGAATTTCCGGTGTAGCGGTGAAATGCGTAGAGATCGGAAAGAACACCAGTGGCGAAGGCGGCTTCCTGGACTCATACTGACGCTGAGGCGCGAAAGCGTGGGT |
| P-asv6574 | GCTAGCGTTACTCGGATTTACTGGGTGTAAAGGGCGCGTAGGCGGCTATGTAAGCTCTGCCTGAAATGCTCTCGCTCAACGAGAGTATGCGGTGGAGGACTGCATGGCTAGAGGACTGGAGAGGCAGGCGGAATTCCCGGTGTAAGGGTGACATCTGTAGATATCGGGAGGAACACCAGTGGCGAAAGCGGCCTGCTGGCCAGTTTCTGACGCTGAGGTGCGAAAGCTAGGGG |
| P-asv6575 | GCGAGCGTTATCCGGATTTATTGGGTTTAAAGGGTTCGTAGGCGGGAGAATTAGTCAGTGGTGAAAGCCTGCAGCTTGACTGTAGAACTGCCATTGATACTGTTCTTCTTGAATATATATGAAGTGGGCGGAATGCGTCATGTAGCGGTGAAATGCACAGATATGACGCAGAACACCGATAGCGAAGGCAGCTCACTAATATATTATTGACGCTCAGGAACGAAAGCATGGGG |
| P-asv6576 | TCGAGTGATGGCCACTATTACTGGGTTTAAAGCGTCCGTAGCTTGACTGTTAGGTCCCTTGGGAAATCTTCACGCTTAACGTGAAGGCGTCTAAGAGATACCGGCAGTCTTGGAACTGGGAGAGGTAAACCGTACTTCGGGGGTAGGAGTGAAATCTTGTAATCCTCGAGGGACGACCTATGGCGAAGGCAGTTTACCAGAACAGCTTCGACAGTGAGGGACGAAAGCTGGGGG |
| P-asv6577 | GCAAGCGTTAGTCGGAGTTATTGGGCGTAAAGGGTGCGTAGGCGGCGTTTTAAGTTGGATATGAAATTTTGGGGCTCAACCTCAACGCTATATCCAAAACTGGGACGCTAGAGGGATGGCGGAGAAAACGGAATTCCATGTGTAGCGGTGAAATGCGTAGATATATGGAAGAACACCTGTGGCGAAAGCGGTTTTCTAGCCTCCTCCTGACGCTAAGGCACGAAAGCAAGGGG |
| P-asv6578 | ACGAACGTTATTCGGAATTATTGGGCTTAAAGGGTGCGTAGGCGGCTCAGAAAGTGGGGTGTGAAAGCCCTCGGCTCAACCGAGGAATTGCGCCCCAAACTACTGAGCTCGAGGGAGACAGGGGTGAGCGGAACTGATGGTGGAGCGGTGAAATGCGTTGATATCATCAGGAACACCGGTGGCGAAAGCGGCTCACTGGGTCTCTTCTGACGCTGAGGCACGAAAGCTAGGGG |
| P-asv6579 | CCAAGCGTTGTTCGGATTTACTGGGCGTAAAGGGTGCGTAGGTGGTGTGGTAAGTCAAATGTGAAATCCTGCAGCTCAACTGCAGAACTGCATTCGAAACTGCCACGCTGGAGTACTGGAGAGGAGATCGGAATTCTCGGTGTAGCAGTGAAATGCGTAGATATCGAGAGGAACACCAGTGGCGAAGGCGGATCTCTGGACAGTTACTGACACTGAGGCACGAAAGCCAGGGG |
| P-asv6580 | ACAAGTGGGAACCACGTTTATTGGGTCTAAAGCATCCGTAGCCGGCTTGTTAAATGCCCTGTGAAATCGTTCCGCTCAACGGGACGGCGTGCAGGGTAGACTGGCAAGCTTGGAACCAGGAAAGGTATGCGGTATTCTGTGGGGAGCGGTAAAATGCTATAATCCACAGAGGACCACCGATGGCGAAGGCAGCATACTTGAATGGATTCGACGGTGAGGGATGAAAGCCAGGGG |
| P-asv6581 | GCAAGCGTTATCCGGAATCACTGGGCGTAAAGGGTGCGTAGGCGGCCTAACAAGTCAGAGGTGAAAGGCATTGGCTCAATCAATGTAAGCCTTTGAAACTGTAGGGCTTGAATTTTTGTGAAGTAACTAGAATATGTAGTGTAGCGGTGAAATGCTTAGATATTACATGGAATACCAATTGCGAAGGCAGGTTACTAACAAACGATTGACGCTGATGGACGAAAGCGTGGGG |
| P-asv6582 | CCAAGCGTTGTTCGGATTTACTGGGCGTAAAGAGTCTGTAGGCGGTGGGGTAAGTCAGATGTGAAATCCCGGGGCTCAACCCCGGAACTGCATCCGATACTGCCTCACTTGAGGGTTGGAGGGGAGTCTGGAATTCTCGGTGTAGCAGTGAAATGCGTAGATATCGAGAGGAACACTAGTGGCGAAGGCGAGACTCTGGACAACACCTGACGCTGAGAGACGAAGGCCAGGGG |
| P-asv6583 | ACGAGCGTTGTTCGGATTTATTGGGCGTAAAGGGTCCGCAGGGGGTTAAGTAAGTCTGATGTGAAATCCTGCAGCTCAACTGCAGAACTGCATTGGAAACTACTTGACTAGAGCGTCTGAGAGGAGAATGGAATTCCTGGTGTAGCGGTGGAATGCGTAGATATCAGGAAGAACACCAAAAGCGAAGGCATTTCTCTGGAAGATTGCTGACCCTCAGGGACGAAAGCGTGGGG |
| P-asv6584 | GCAAGCGTTGCTCGGAATAATTGGGCGTAAAGGGTGCGCAGGCGGTTTTTTAAGTAAAACATTAAAACCTTGGACTTAATCTAAGAAATGTGTTTTATACTGAAAAACTAGAGTTTGGAAGAGGAAATTGGAATTTCTCAAGTAGAGGTGAAATTCGTAGATATGAGAAGGAACACCAGAGGCGGAGGCGAATTTCTGGGACAATACTGATGCTAAGGCACGAAAGCGTGGGG |
| P-asv6585 | GCAAGCGTTGTTCGGAATCACTGGGCATAAAGCGCGCGTAGGCGGTCTGCTAAGTCGGATGTGAAAGCCCCGGCCCAACCGGGGAATGGCGTCCGATACTGGCGGACTCGAGGATTGGAGAGGTGTGTGGAATTCCTGGTGGAGCGGTGAAATGCGTTGATATCAGGAGGAACACCGGAGGCGAAGGCGACACACTGGCCAATTTCTGACGCTGAGGTGCGAAAGCATGGGG |
| P-asv6586 | GCAAGCGTTGTTCAGAATAACTGGGCGTAAAGGGCACGTAGGCGGTGATACGCGTCAGATGTGAAAGACCTGAGCTTAACTCAGGAATAGCATCTGAAACGGTATGACTCGAGATCGAGAGGGGAAAAGGGAATTGTGCGTGTAGAGGTGAAATTCGTAGATATGCACAGGAACACCGGTGGCGATGGCGCTTTTCTGGCTCGATACTGACGCTAAGGTGCGAAAGCGTGGGG |
| P-asv6587 | GCAAGCGTTGTTCAGAATTACTGGGCGTAAAGAGCGCGTAGGCGGTCTACCAAGTCAGACGTGAAAGCCCTGGGCTCAACCCGGGAACTGCGTTTGAAACTGGTAAACTTGAGTGTCAGAGGGGTGAGTGGAATTCCTAGTGTAGAGGTGAAATTCGTAGATATTAGGAGGAACACCAGTGGCGAAGGCGACTCACTGGCTGAATACTGACGCTAAGGCGCGAAAGCGTGGGG |
| P-asv6588 | CCAAGTGATTGCCGCTGTTATTAGGTCTAAAGCGTCCGTAGCCGGTTCTTCAAGTCCCTGGTGAAATCTGACAGCAAACTGTCAGGCGTGCTGGGGTTACTGCAGGACTCGAGACCGGGAGAGGGCATAGATACCCGTGGGGTAGGGGTTAAATCCTATAATCCTACGGAGATCACCTGTGGCGAAGGCGTATGCCTGGAACGGGTCTGACGGTGAGGGACGAAAGCTAGGGG |
| P-asv6589 | CCAAACGTTATTCGGAATCACTGGGCTTAAAGGGTGCGTAGGCTGTCCGGTAAGTTGGGTGTGAAAGCCCTCGGCTCAACCGAGGAATTGCGTCCAAAACCGTCGGGCTTGAGGAAGATAGAGGTGAGCGGAACTAAGGGTAGAGCGGTGAAATGCGTTGATATCCTTAGGAACACCTGTAGCGAAAGCGGCTCACTGGGTCTTTTCTGACGCTGAGGCACGAAAGCTAGGGG |
| P-asv6590 | ACGAGCGTTAATCGGAATCACTGGGCTTAAAGGGTTCGCAGGCGGCTCGAAAAGCGTTCCTTGAAATCCCCCGGCTCAACCGGGGAACTGCGGAGCGAACTGCCGAGCTTGAGGCAAGTAGGGGCGGGCAGAACTGTAGGTGGAGCGGTGAAATGCGTAGATATCTACAGGAATGCCGATGGTGAAGACGGCCCGCTGGGCTTGACCTGACGCTCATGCACGAAAGCGTGGGG |
| P-asv6591 | GCAAGCGTTATTCGGAATTATTGGGCGTAAAGGGTGCGCAGGCGGTTTATTAAGTCAACTGTTTAATTCCTCGGCCTAACTGAGGGTCAGCGGTAGAAACTGATAAGCTTGAGTATGAGAGAGAGAAGTGGAATTCTCGGAGTAGCGGTAAAATGCGTAGATCTCGAGAGGAACACCGATGGCGAAGGCAGCTTCTTGGCTCATTACTGACGCTCATGCACGAAAGCGTGGGG |
| P-asv6592 | GCAAGCGTTAATCGGAATCACTGGGCGTAAAGCGCTCGTAGGCTGTTTGGTAAGTCAGACGTGAAAGCCCGCGGCTTAACCGCGGAACTGCGTTTGATACTGCCAGACTTGAGTGTCGGAGAGGGTGGCGGAATTCCGGGTGTAGGAGTGAAATCCGTAGATATCCGGAGGAACACCAGTGGCGAAGGCGGCCACCTGGACGACAACTGACGCTGAGGAGCGAAAGCGTGGGT |
| P-asv6593 | GCGAGCGTTATCCGGATTCATTGGGTTTAAAGGGTGCGTAGGCGGCCCGGTAAGTCAGCGGTGAAAGTTTGCGGCTCAACCATAAAATTGCCGTTGATACTGTCGGGCTTGAGTTCAGATGAAGCGGGCGGAATGAGTAGTGTAGCGGTGAAATGCTTAGATATTACTCAGAACACCGATTGCGAAGGCAGCCCACTAAACTGCGACTGACGCTGATGCACGAAAGCGTAGGG |
| P-asv6594 | GCAAGCGTTGTTCGGAATCATTGGGCGTAAAGCGCGCGCAGGCGGCCATTTAAGTCTGATGTGAAAGCCCGGGGCTCAACCCCGGAGGTGCATTGGAAACTGGATGGCTTGAGTCCAAGAGAGGTTAGTGGAATTCGTGGTGTAGGGGTGAAATCCGTAGATATCACGAGGAACACCAGTGGCGAAGGCGACTGACTGGATTGGTACTGACGCTGAGGCGCGAAAGCGTGGGG |
| P-asv6595 | CCGGGCGTTATCCGGATTTATTGGGTTTAAAGGGAGTGTAGGCGGCCTGTTAAGCGTGTTGTGAAATGTAGATGCTCAACATCTGAACTGCAGCGCGAACTGGCTGGCTTGAGTACACGCAACGTGGGCGGAATTCATGGTGTAGCGGTAAAATGCTTAGATATCATGAGGAACTCCTATTGCGAAGGCAGCTCACGGGAGTGTCACTGACGCTTAAGCTCGAAGGTGCGGGC |

| | |
|---|---|
| P-asv6596 | GCAAGCGTTATCCGGAATCATTGGGTTTAAAGGGTCCGCAGGCGGACTTTCAAGTCAGTGGTGAAATCTCACAGCTTAACTGTGAAACTGCCATTGAAACTGTTAGTCTTGAATTCGGTCGAAGTGGGCGGAATATGTCATGTAGCGGTGAAATGCTTAGATATGACATAGAACACCGATAGCGAAGGCAGCTCACTAGGCCTGGATTGACGCTCAGGGACGAAAGCGTGGGG |
| P-asv6597 | GCAGGCGTTGTTCGGAATTACTGGGCGTAAAGCGTGCGTAGTCGGTATTGAGAGTCACGGGTGAAATCCCAGGGCTTAACTTGGGACGTGCATTTGGAACTGGGTGACTAGAGTGTGTCAGAGGGAGGTAGAATTCCACATGTAGCGGTGGAATGCGTAGAGATGTGGAGGAATACCGATGGCGAAGGCAGCCTCCTGGGATAACACTGACGTTGAGGCTCGAAAGCGTGGGG |
| P-asv6598 | CCGAGCGTTATCCGGAATCATTGGGTTTAAAGGGTCCGTAGGCGGGACATTAAGTCAGGGGTGAAAGGTTACAGCTCAACTGTAGAATTGCCTTTGATACTGGTGTTCTTGAGTTATGTTGTTGTGGTCGGAACATGTGGTGTAGCGGTGAAATGCATAGATATCACATAGAACGCCGATTGCGAAGGCAGGTCACAAAACATATACTGACGCTGAGGGACGAAAGCGTGGGG |
| P-asv6599 | CCAAGCGTTGTTCGGAATCACTGGGCGTAAAGCGTGTGTAGGCGGCGTGGAAAGTCAGATGTGAAATCTCCGGGCTCAACCTGGAAACTGCATCCGATACTCCCACGCTAGAGTAGTGGAGGGGCATCTGGAATTCACGGTGTAGCAGTGAAATGCGTAGAGATCGTGAGGAACACTAGTGGCGAAGGCGAGATGCTGGACACTAACTGACGCTGAGACACGAAGGCTAGGGT |
| P-asv6600 | GCAAGCGTTGTCCGGATTTATTGGGTTTAAAGGGTGCGTAGGCGGGTGATTAAGTCAGTGGTGAAAGCCCACAGCTCAACTGTGGAACTGCCATTGATACTGGTTATCTTGAGTGTAGTTGAGGCAGGCGGAATTTACGATGTAGCGGTGAAATGCATAGATATCGTAAAGAACACCTATAGCGAAGGCAGCTTGCTAAGCTACTACTGACGCTGAGGCACGAAAGCATGGGT |
| P-asv6601 | GCAAGCGTTATTCGGAATTACTGGGCGTAAAGCGCGCGTAGGCGGATTGTTAAGTCAGTTGTGAAAGCCCTGGGCTCAACCTAGGAATGGCGATTGAAACTGGCAGTCTAGAGTTTAGTAGAGGGAAGGGGAATTTCTGGAGTAGCAGTGAAATGCGTAGATATCAGAAGGAACATCAGTGGCGAAGGCGCCTTCCTGGACTAGAACTGACGCTGAGGTGCGAAAGCGTGGGG |
| P-asv6602 | GCTAGCGTTGTTCGGAATCACTGGGCGTAAAGCGCGCGTAGGCGGACTGATCAGTTGGGGGTGAAAGCCCGGAGCTTAACTCCGGAACTGCCTCCAATACTGTCAGTCTTGAGGTCGAGAGAGGTAAGTGGAATTCCGAGTGTAGAGGTGAAATTCGTAGATATTCGGAAGAACACCAGTGGCGAAGGCGACTTACTGGCTCGATTCTGACGCTGAGGTGCGAAAGCGTGGGG |
| P-asv6603 | GCAAGCGTTATCCGGTTTTATTGGGTTTAAAGGGTGCGTAGGCGGCTTATTAAGTCAGTTGTGAAATCCTAGCGCTCAACGTTAGAACTGCGATTGATACTGATAGGCTTGAGTCAAGAAGAGGTAGGTAGAATTTATGGTGTAGTAGTGAAATGCTTAGATATCATAAGGAATAGCAATAGCGTAGGCAGCTTACTGGTCTTGAACTGACGCTGAGGCACGAAAGCGTGGGG |
| P-asv6604 | GCGAGCGTTAATCGGAATTACTGGGCGTAAAGGGTGCGTAGGTGGTTTGATAAGTTAGCTGTGAAATCCCCGGGCTTAACCTGGGCAGGTCAGCTAAGACTGTTGAACTCGAGTATAGGAGAGGGTAGTGGAATTTCCGGTGTAGCGGTGAAATGCGTGGAGATCGGAAGGAACACCAGTGGCGAAGGCGGCTACCTGGCCAGATACTGACACTGAGGCACGAAAGCGTGGGG |
| P-asv6605 | GCGAGCGTTAATCGGAATTACTGGGCGTAAAGGGTGCGTAGGTTGCTTGGCAAGTCAGATGTGAAAGCCCTGGGCTTAACCTGGGAATTGCATTTGAAACTGCTAGGCTAGAGTACAGGAGAGGGAAGTGGAATTTCCGGTGTAGCGGTGAAATGCGTAGATATCGGAAGGAACACCAGTGGCGAAGGCGACTTCCTGGACTGTTACTGACACTGAGGCACGAAAGCGTGGGG |
| P-asv6606 | GCGAGCGTTGTTCGGAATCATTGGGCGTAAAGGGCGTCTAGGTGGCCTGGCAAGTCAGATGTGAAATCCCTCGGCTCAACCGAGGACGTGCATTTGAAACTGCCCGGCTCGAGTACCGGAGGGGGCAGCGGAATTCCCGGTGTAGAGGTGAAATTCGTAGATATCGGGAGGAACACCAGTGGCGAAGGCGGCTGCCTGGCCGGATACTGACACTCAAGCGCGAAAGCATGGGG |
| P-asv6607 | GCGAGCGTTATCCGGATTTATTGGGTTTAAAGGGTGCGTAGGCGGTTTAGTAAGTCGGTGGTTAAAGGTAGTCGCTTAACGATTTTACATGCCACCGATACTGTTGAGCTTGAGTTATTTGGAGGCAGCTAGAATTTCCGGTGTAGCGGTGAAATGCATAGATACCGGAAGGAATACCAATTGCGAAGGCAGGTTGCTACAAATATACTGACGCTGATGCACGAAAGCGTGGGG |
| P-asv6608 | CCGAGTGATGGCCGATATTATTGGGCCTAAAGCGTCCGTAGCTGGCCGGGGAAGTCCGTCGGGAAATCCACTCGCTCAACGAGTGGGCGTCCGGCGGAAACTCTCCGGCTTGGGACCGGAAGACCGAAGGGGTACGTCCGGGGTAGGAGTGAAATCCTGTAATCCTGGACGGACCACCGGTGGCGAAAGCGCCTTCGGAAGACGGATCCGACAGTGAGGGACGAAAGCTAGGGT |
| P-asv6609 | GCAAGCGTTAATCGGAATTACTGGGCGTAAAGCGCACGCAGGCGGCTTTTTAAGTCGGATGTGAAAGCCCCGGGCTCAACCTGGGAATTGCATTCGATACTGTAAAGCTAGAGTATGTGAGAGGGGGGTAGAATTCCAAGTGTAGCGGTGAAATGCGTAGATATGCGGAGGAACACCAGTGGCGAAGGCGACTCGCTGGACAAGTATTGACGCTGAGGTGCGAAAGCGTGGGG |
| P-asv6610 | ACAAACGTTATTCGGAATCACTGGGCTTAAAGGGTGCGTAGGCGGCGACGCAAGTTGGGTGTGAAATCCCTCGGCTCAACCGAGGAACTGCGCCCAAAACTGCGTTGCTTGAGGGAGATAGAGGTGAGTGGAACTTAGGGTGGAGCGGTGAAATGCGTTGATATCCTAAGGAACACCAGTGGCGAAAGCGACTCACTGGGTCTCTTCTGACGCTGAGGCACGAAAGCTAGGGG |
| P-asv6611 | CCGAGCGTTAATCGGAATCACTGGGCTTAAAGAGTGTGTAGGCGGACATATAGGTACTTTGTGAAATCCCACGGCTCAACCGTGGAATAGCTTGGTAAACCATATGTCTTGAGGCAGGTAGGGGTGGTCGGAACGATAGGTGGAGCGGTGAAATGCGTAGATATCTATCGGAACGCCAACGGAGAAGTCAGGCCACTGGGCCTGTCCTGACGCTAAGACACGAAAGCATGGGT |
| P-asv6612 | GCAAGCGTTATCCGGAATGACTGGGCGTAAAGAGTTCGTAGGTGGTAATGAAAGTTTGCCGTTAAAGACTAAGGCTTAACCTTAGGAGAGCGGTGAAAACTACATAACTAGAGTATGGAAGGGGCAGAGGGAATTCCTGGTGTAGCGGTGAAATGCGTAGAGATCAGGAGGAACATCGGTGGCGAAAGCGCTCTGCTAGGCCAAAACTGACACTGAGGAACGAAAGCTAGGGG |
| P-asv6613 | GCGAGCGTTATCCGGATTCATTGGGTTTAAAGGGTGCGTAGGCGGCCCGGTAAGTCAGCGGTGAAAGTTTGCGGCTCAACCGTAAAATTGCCGTTGATACTGTCGGGCTTGAGTTCAGATGAAGTGGGCGGAATGAGTAGTGTAGCGGTGAAATGCTTAGATATTACCCAGAACACCGATTGCGAAGGCAGCCCACTAAACTGCGACTGACGCTGATGCACGAAAGCGTGGGG |
| P-asv6614 | GCAACCGTTGTTCGGAATCACTGGGCTTAAAGCGCACGTAGGCGGAATGTTAAGTTGGCTGTGAAATTTCTTGGCTTACCCAAGGAACGGCAGCCAAAACTGGCGTTCTCGAGGATGGCATGGGAGAGTGGAATTCCTGGTGGAGCGGTGAAATGCGTTGATATCAGGAGGAACACCGGAGGCGAAGGCGACTCTCTGGGCCAAATCCTGACGCTGAGGTGCGAAAGCCAGGGG |
| P-asv6615 | GCGAGCGTTGTCCGGAATTATTGGGCGTAAAGAGCGTGTAGGCGGTCTGGTAAGTCTGTCGTGAAAGGCTGGGGCTCAACCCCGGACGTGCGATGGATACTGCCAGACTAGAGGATGGTAGAGGCGAGTGGAATTCCCGGTGTAGCGGTGAAATGCGCAGATATCGGGAGGAACACCAGTAGCGAAGGCGGCTCGCTGGGCCATACCTGACGCTGAGACGCGAAAGCGTGGGG |
| P-asv6616 | GCAAGCGTTACTCGGAATCACTGGGCGTAAAGCGCGCGCAGGCGGCTATTTAAGTTGGATGTGAAAGCCTACGGCTCAACCGTAGAACTGCATCCAAAACTATTTAGCTAGAGTGTGGGAGAGGAAGATGGAATTAGTTGTGTAGGGGTAAAATCCGTAGAGATAACTAGGAATACCAAAAGCGAAGGCAATCTTCTGGAACACTACTGACGCTGAGGCGCGAAAGCGTGGGG |
| P-asv6617 | GCTAGCGTTGCTCGGAATTACTGGGCGTAAAGGGCGCGTAGGCGGCACGATTAGTCAGGCGTGAAATTCCTGGGCTCAACCTGGGGACTGCGCTTGATACGGTCGAGCTAGAGGATGGAAGAGGGTCGCGGAATTCCCAGTGTAGATGTGAAATTCGTAGATATTGGGAAGAACACCAGTGGCGAAGGCGGCTCACTGGACTGGTATTGACGCTGAGGTGCGAAAGCGTGGGG |
| P-asv6618 | CCAAGCGTTATCCGGATTTATTGGGTTTAAAGGGAGCGTAGGTTGACGTATAAGTCAGCTGTGAAAGTTTACGGCTCAACCGTGAAATTGCAGTTGATACTGTATGTCTTGAGTGTACAAGAGGTGGGCGGAATTCGTGGTGTAGCGGTGAAATGCTTAGATATCACGAAGAACTCCAATTGCGAAGGCAGCTCACTGGGGTACAACTGACAGTGAGGCTCGAAAGTGTGGGT |
| P-asv6619 | GCAAGCGTTGTCCGGAATTACTGGGCGTACAGAGCGCGTAGGCGGCCATGTAAGTCGCGTGTGAAAGACCGGGGCTCAACCCCGGTATTGCACGCGATACTGCATGGCTGGAGACAGGCAGGGGCGAGCGGAATTCCCGGTGTAGCGGTGGAATGCGTAGATATCGGGAGGAACACCCGTGGCGAAGGCGGCTCGCTGGGCCTGATCTGACGCTGCAGCGCGAAAGCGTGGGG |
| P-asv6620 | GCGAGCGTTAATCGGAATTACTGGGCGTAAAGCGCACGCAGGCGGTTTGTTAAGCTAGATGTGAAAGCCCCGGGCTCAACCTGGGATGGTCATTTAGAACTGGCAGACTAGAGTCTTGGAGAGGGGAGTGGAATTCCAGGTGTAGCGGTGAAATGCGTAGATATCTGGAGGAACATCAGTGGCGAAGGCGACTCCCTGGTCAAAGACTGACGCTCATGTGCGAAAGTGTGGGT |
| P-asv6621 | GCAAGCGTTGTTCAGAATCACTGGGCTTAAAGCGTATGTAGGCGGACTTGAAAGTCAGAAGTGAAATCCCCGGGCTTAACCCGGGAACTGCTTTTGAAACTCCAAGTCTAGAATCCTGGAGAGGTTGGGAGAATTCCGAGTGTAGAGGTGAAATTCGCAGATATTCGGAGGAATACCAGTGGCGTAGGCGCCCAACTGGACAGGTATTGACGCTCAGATACGAAAGCGTGGGG |
| P-asv6622 | GCGAACGTTATTCGGAATCACTGGGCTTAAAGGGTGCGTAGGCGGCTTTCCAGGTCTGATGTGAAATCTCACGGCTCAACCGTGAAATTGCGTTGGAAACCGGAAGGCTTGAGTGTGCTAGGGGTGTGTGGAACTTCCTGTGGAGCGGTGAAATGTGTTGATATGGAAGGAACGCCGGTGGCGAAGGCGACACACTGGGGCATAACTGACGCTGAGGTACGAAAGCTAAGGT |
| P-asv6623 | GCAAGCGTTAATCGGATTTACTGGGCGTAAAGCGCGCGTAGGCGGTTCGTTAAGTCGATTGTGAAAGCCCTGGGCTTAACCCTGGAACTGCCTGTGAGACCTCAGTACTAGAGTGTGAGAGGGGATAGTGGAATACCCAGTGTAGCGGTGAAATGCGTAGAGATTGGGTAGAACACCGGTGGCGAAGGCGGCTATCTGGCTCACAACTGACGATCAGGCACGAAAGCGTGGGG |
| P-asv6624 | GCTAGCGTTGCTCGGAATTACTGGGCGTAAAGGGCGCGTAGGCGGACAGTTAAGTTGGGGGTGAAAGCCCGGGGCTCAACCTCGGAAATGCCTTCAATACTGGCTGTCTTGAGTACGGGAGAGGTGAGTGGAACTCCGAGTGTAGAGGTGAAGTGCGTAGATATCTGGAGGAACATCAGTGGCGAAGGCGGCTTCCTGGTCCAACATTGACATTCAGGTGCGAAAGCGTGGGG |
| P-asv6625 | GCGAGCGTTGTCCGGAATCACTGGGCGTAAAGGGCGCGTAGGCGGTGACTTAAGCGTGTGGTGAAAGCCCGGGGCTCAACCCCGGGTCGGCCGTGCGAACTGGGTCACTCGAGCACTGTAGAGGCAGGTGGAATTCCGGGTGTAGCGGTGGAATGCGTAGAGATCCGGAAGAACACCAGTGGCGAAGGCGGCCTGCTGGGCAGTAGCTGACGCTGAGGCGCGACAGCGTGGGG |
| P-asv6626 | GCAAGCGTTATTCGGAATTACTGGGCGTAAAGGGCGCGTAGGCGGTCCGTTAAGTGAGAAGTGAAAGCCCATGGCTCAACCATGGATGTGCTTCTCATACTGGCGAACTAGAGTATGGGAGAGGAAAGTGGAATTCCCGGTGTAGCGGTGAAATGCGTAGAGATCGGGAGGAACACCAGTGGCGAAGGCGACTTTCTGGACCATAACTGACGCTGAGGCGCGAAAGCGTGGGG |
| P-asv6627 | GCTAGCGTTGTTCGGATTTACTGGGCGTAAAGCGCACGTAGGCGGTTTGTTAAGTGAGAGGTGAAATCCCAGAGCTCAACTCTGGAACTGCCTTTCATACTGGCAAGCTAGAGTCCGGAAGAGGTAAGTGGAACTCCTAGTGTAGAGGTGGAATCCGTAGATATTAGGAAGAACACCAGTGGCGAAGGCGACTCACTGGCCCGTTACTGACGCTGAGGCGCGAAAGCGTGGGG |
| P-asv6628 | GCGAGCGTTAATCGGAATTACTGGGCGTAAAGCGAGTGTAGGTGGCTCATTAAGTCACATGTGAAATCCCCGGGCTTAACCTGGGAACTGCATGTGATACTGGTGGTGCTAGAATATGTGAGAGGGAAGTAGAATTCCAGGTGTAGCGGTGAAATGCGTAGAGATCTGGAGGAATACCGATGGCGAAGGCAGCTTCCTGGCACAATATTGACATTGAGATTCGAAAGCGTGGGT |
| P-asv6629 | GCAAGCGTTGTTCGGAATTACTGGGCGTAAAGGGCGCGTAGGCGGCCTGCTAAGTCGAATGTGAAATCCCTGGGCTCAACCCAGGAACTGCGTCCGATACTGGTTGGCTTGAATTCGGGAGAGGGATGCGGAATTCCAGGTGTAGCGGTGAAATGCGTAGATATCAGGAAGAACACCCGTGGCTAAGGCGGCTCTCTAGTACGGTACTGACGCTGAGACGCGAAAGCGTGGGG |
| P-asv6630 | GCTAGCGTTGTTCGGAATCACTGGGCGTAAAGCGCACGTAGGCGGATATGTCAGTCAGGGGTGAAATCCCGGAGCTCAACTTCGGAACTGCCTTTGATACAGCATATCTCGAGTCCGAGAGAGGTGAGTGGAATTCCTAGTATAGAGGTGAAATTCGTAGATATTCGGAAGAACACCAGTGGCGAAGGCGACTCGCTGGACAAGTATTGACGCTGAGGTGCGAAAGCGTGGGG |
| P-asv6631 | GCTAGCGTTGTTCGGAATCACTGGGCGTAAAGCGCACGTAGGCGGACTTTTAAGTCAGAGGTGAAAGCCTGGAGCTCGACTTTAGAACTGCCATTGATACTATTTATCTTGAATATTGTGGAGGTAAGCGGAATATGTCATGTAGCGGTGAAATGCTTAGATATGACATAGAACACCTATTGCGAAGGCAGCTTACTACGCATATATTGACGCTGAGGCACGAAAGCGTGGGG |
| P-asv6632 | CCGAGCGTTGTCCGGATTTATTGGGCGTAAAGCGAGCGCAGGTGGTTACATAAGTCTGAAGTAAAAGGCAGTGGCTCAACCATTGTATGCTTTGGAAACTGTGTAACTTGAGTGCAGAAGGGGAGAGTGGAATTCCATGTGTAGTGGTGAAATGCGTAGATATATGGAGGAACACCGGTGGCGAAAGCGGCTCTCTGGTCTGTAACTGACACTGAGGCTCGAAAGCGTGGGG |
| P-asv6633 | GCGAGCGTTGTCCGGAATTACTGGGCGTAAAGAGCTCGTAGGCGGTTTGTCACGTCGTCTGTGAAGTCCTAGGGCTTAACCCTGGACGTGCAGGCGATACGGGCTGACTTGAGTACTACAGGGGAGACTGGAATTTCTGGTGTAGCGGTGGAATGCACAGATATCAGGAAGAACACCGATGGCGAAGGCAGGTCTCTGGGTAGTAACTGACGCTGAGGAGCGAAAGCATGGGT |
| P-asv6634 | GCAAGCGTTATCCGGATTCACTGGGTTTAAAGGGTGCGTAGGAGGGTGTGTAAGTCAGTGGTGAAATCTTCGAGCTTAACTCGGAAACTGCCGTTGATACTACATGTCTTGAATATCGTGGAGGTGAGCGGAATATGTCATGTAGCGGTGAAATGCTTAGATATGACATAGAACACCAATTGCGAAGGCAGCTCGCTACACGAATATTGACTCTGAGGCACGAAAGCGTGGGG |

| | |
|---|---|
| P-asv6635 | CCAAGCGTTAATCGGAATTACTGGGCGTAAAGCGTGCGCAGGCGGTTGTGCAAGACCGATGTGAAATCCCCGAGCTTAACTTGGGAATTGCATTGGTGACTGCACGGCTAGAGTGTGTCAGAGGGGGGTAGAATTCCACGCGTAGCAGTGAAATGCGTAGAGATGTGGAGGAATACCGATGGCGAAGGCAGCCTTCTGGACTGACACTAACACTGAGGTGCGAAAGCGTGGGT |
| P-asv6636 | GCTAGCGTTGCTCGGAATCACTGGGCGTAAAGGGTGCGTAGGCGGGTTTTTAAGTCAGAGGTGAAATCCTGGAGCTCAACTCCAGAACTGCCTTTGATACTGAAAGTCTTGAGTATGGGAGAGGTGAGTGGAACTGCGGGTGTAGAGGTGAAATTCGTAGATATTCGGAAGAACACCAGTGGCGAAGGCGGCTCACTGGACTGGTATTGACGCTGAGGTGCGAAAGCGTGGGG |
| P-asv6637 | CCAAGCGTTATCCGGATTCACTGGGTTTAAAGGGTGCGCAGGCGGTCAGGTAAGTCAGTGGTGAAAGCCTTCGGCTCAACCGAAGAATTGCCATTGAAACTACTTGACTTGAGTGTAGTTGAGGTGGGCGGAATGTAGCATGTAGCGGTGAAATGCATAGATATGCTACAGAACACCGATAGCGAAAGCAGCTCACCAGACTATGACTGACGCTGAGGCACGAAAGCATGGGT |
| P-asv6638 | GCAAGCGTTATCCGGATTTACTGGGTTTAAAGGGTGCGTAGGCGGGCCATTAAGTCAGATGTGAAAGCCTCAAGCTCAACTTGAGAACTGCGTTTGGTACTGGTGGCCTTGAATTCGATAGAAGTGGGCGGAACAAATCGTGTAGCGGTGAAATGCTTAGATATGATTTGGAACACCAATTGCGAAGGCAGCTCACTATGCCGATATTGACGCTGAGGCACGAAAGCGTGGGG |
| P-asv6639 | GCAAGCGTTGTCCGGAATTATTGGGCGTAAAGCGCGCGCAGGCGGCATTGTAAGTCTTGTGTTTAGACGCGGGGCTCAACCCCGTGTCGCATGGGAAACTGCAAGGCTTGAGTGCAGAAGAGGAAAGTGGAATTCCACGTGTAGCGGTGAAATGCGTAGAGATGTGGAGGAACACCAGTGGCGAAGGCGACTTTCTGGGCTGTAACTGACGCTGAGGCGCGAAAGCGTGGGG |
| P-asv6640 | GCAAGCGTTATCCAGATTTACTGGGTGTAAAGGGAGCGTAGACGGTGTGGCAAGTCTGGTGTGAAAGGCATGGGCTCAACCTGTGGACTGCATTGGAAACTGTCATACTTGAGTGCCGGAGGGGTAAGCGGAATTCCTAGTGTAGCGGTGAAATGCGTAGATATTAGGAGGAACACCAGTGGCGAAGGCGGCTTACTGGACGGTAACTGACGTTGAGGCTCGAAAGCGTGGGG |
| P-asv6641 | GCTAGCGTTGTTCGGAATTACTGGGCGTAAAGCGCACGTAGGCGGCTCCGCAAGTCAGAGGTGAAATCCCCAGGCTCAACCTGGGAATTGCCTTTGATACTGCGGGGCTAGAGGCCGGGAGAGGGTAGTGGAATTCCCAGTGTAGAGGTGAAATTCGTAGATATTGGGAAGAACACCGGTGGCGAAGGCGGCTACCTGGACCGGTACTGACGCTGAGGTGCGAAAGCGTGGGG |
| P-asv6642 | GCTAGCGTTATTCGGAATCACTGGGCGTAAAGAGCAAGTAGGCGGCTTGGAAAGCCAGGGGTGAAAGCCCGGAGCTTAACTCCGGAATTGCCTTTGGAACTATCAAGCTAGCATCTGGCAGAGGGTGGTGGAATTCCTAGTGTAGAGGTGAAATTCGTAGATATTAGGAGGAACACCGGTGGCGAAGGCGACCACCTGGGCCAGTATGGACGCTGAGGTGCGAAAGCGTGGGG |
| P-asv6643 | GCTAGCGTTATTCGGAATTATTGGGCGTAAAGAGCGCGTAGGCTGGTTAGTAAGTTAAAAGTGAAATCCCAAAGCTCAACTTTGGAATTGCTTTTAAAACTGCTAATCTAGAGATTGAAAGAGGATAGAGGAATTCCTAGTGTAGAGGTGAAATTCGTAAATATTAGGAGGAACACCAGTGGCGAAGGCGTCTATCTGGTTCAAATCTGACGCTGATGCGCGAAGGCGTGGGG |
| P-asv6644 | GCAAGCGTTACTCGGAATCACTGGGCGTAAAGAGCACGTAGGCGGCCTTGCAAGTCAGATGTGAAATCCAACGGCTTAACCGTTGAACTGCATTTGAAACTGCAAGGCTAGAGTATGGGAGAGGTAGGTGGAATTCTCGGTGTAGGGGTAAAATCCGTAGAGATCGAGAGGAATACTCATTGCGAAGGCGACCTGCTGGAACATTACTGACGCTGAGGTGCGAAAGCGTGGGG |
| P-asv6645 | GCAAGCGTTGTTCGGATTCACTGGGCATAAAGAGCACGTAGGCGGCTACGTAAGTCGGATGTGAAATCCCAGGGCTCAACCGTGGAATGGCATCCGATACTGCGTGGCTCGAGGTCGGGAGGGGTGAGCGGAACTCCAGGTGGAACGCTGAAATGTGTAGATATCTGGAGGAACACCGGTGGCGAAGGCGGCTCACTGGCCCGAATCTGACGCTGAGGTGCGAAAGCTAGGGG |
| P-asv6646 | GCAAGCGTTGTCCGGATTTATTGGGTTTAAAGGGTGCGTAGGCGGCCGATTAAGTCAGTGGTGAAAGCCTATAGCTTAACTATAGAATTGCCATTGATACTGGTTGGCTTGAGTTCGATTGAGGTAGGCGGAATTTGTGGTGTAGCGGTGAAATGCATAGATACCACAAGGAACACCGATAGCGTAGGCAGCTTACTAAGTCGACACTGACGCTGAGGCACGAAAGCGTGGGG |
| P-asv6647 | GCTAGCGTTGCTCGGAATCACTGGGCGTAAAGGGTGCGTAGGCGGCGCTTTAAGTCGGGTGTGAAATCCTTCGGCTCAACCGAGGAATTGCGCCCGATACTGGGGTGCTAGAGGGAGACAGGGGTGAGCGGAACAGATGGTGGAGCGGTGAAATGCGTTGATATCATCTGGAACACCGGTGGCGAAAGCGGCTCACTGGGTCTCTTCTGACGCTGAGGCACGAAAGCTAGGGT |
| P-asv6648 | GCGAGCGTTGTCCGGATTTACTGGGCGTAAAGGGCGCGTAGGCGGTCTGTTAAGTGAGAAGTGAAATCTCCAGAGCTCAACTCGGAAACAGCTTTTCATACTGGCAGACTGGAGGAACGCAGAGGTGACTGGAATTCCTGGTGTAACGGTGAAATGTGTTGATATCAGGAGGAACTCCAATGGCGAAGGCAGGTCACTGGGCGTTATCTGACGCTGAGGCGCGAAAGCGTGGGT |
| P-asv6649 | GCAAGCGTTGTCCGGATTTACTGGGTGTAAAGGGCGCGCAGGCGGGTTTGTAAGTCAGGGGTGAAATCTCACGGCTTAACCGTGAAACTGCCTTTGATACTGCAAATCTTGAGTTTGGGAGGGAGAGATGGAATTCCAGGTGTAGTGGTGAAATACGTAGATATCTGGAAGAACACCAGTGGCGAAGGCGATCTCTTGGCCCATAACTGACGCTGAGGCGCGAAAGCGTGGGT |
| P-asv6650 | CCGAGCGTTATCCGGATTTATTAGGTTTAAAGGGTGCGTAGGCGGGCTGATAAGTCAGTGGTGAAAGTTTGCAGCTTAACTGTAAAATTGCCATTGAAACTGTTAGTCTTGAGTACAGTAGAGGTAGGCGGAATGTGTAGTGTAGCGGTGAAATGCATAGATATTACACAGAACACCAATTGCGTAGGCAGCTTACTGGAGTGTCACTGACGCTGAGGCACGATAGCGTGGGG |
| P-asv6651 | GCGAGCGTTGTCCGAATTTATTGGGCGTAAAGCGCGTGTAGGCGGATATCCAAGCCTGGGGTGAAAGGTTGGGGCTCAACCCCAAGAGTGCCCTGGGGACTGGGTGTCTGGAGGCAGGCAGAGGAGTGTGGAATTCCTGGTGTAGCGGTGGAATGCGTAGATATCAGGAAGAACACCCATGGCGAAGGCAGCACTCTGGGCCTGTACTGACGCTGAGACGCGAAAGCGTGGGG |
| P-asv6652 | CCGAGCGTTATCCGGAATCATTGGGTTTAAAGGGTCCGTAGGCGGGACAATCAGTCAGTGGTGAAAGTTTGTGGCTCAACCATAAAATTGCCATTGATACTGTTGTTCTTGAGTAATTGTGAAGTGGTTAGAATGAGTAGTGTAGCGGTGAAATGCATAGATATTACTCAGAATACCGATTGCGAAGGCAGATCACTAACAATGTACTGACGCTGATGGACGAAAGCGTAGGT |
| P-asv6653 | GCAAGCGTTGTCCGGAATCATTGGGTGTAAAGGGTGCGCAGGCTGGCATTTAAGTCTGGGGTGAAATCTTGCCGCTTAACGGTAAAATGGCCTTAGATACTGAATGTCTTGAATCTGACAGAGGTCGGTGGAATTCGTAGTGTAGCGGTGAAATGCATAGATATTACGAAGAACATCAGTGGCGAAGGCGGCCGGCTGGGTCAGTATTGACGCTGAGGCACGAGAGCGTGGGG |
| P-asv6654 | GCGAGCGTTACCCGGAATTACTGGGTGTAAAGGGGGCGTAGGTGGGGTAGTAGGTCAGATGTTAAAGGATACGGCTTAACCGTAGAAATGCGTTTGAAACCGCTACTCTTGGAGATGGTAGAGGAAAAGGGAATTACCGGTGTAGGGGTGAAATCCGTAGATACCGGTAAGAAGGCCGGTGGCGAAGGCGCTTTTCTGGACCAAACTCGACGCTGATGCCCGAAAGCTAGGGG |
| P-asv6655 | GCTAGCGTTACTCGGAATTACTGGGCGTAAAGAATGCGTAGGCGGATTTGCAAGTTAGGAGTGAAATCCCTGGGCTCAACCTAGGAACTGCTTTTAAAACTGCAAGTCTTGAATGTTGGATTGATTAGGGGAATTCCTGGTGTAGGGGTAATATCCGTAGATATCAGGAGGAAGACCGGAAGTGAAAGCGCTTAATTCGCCAAATATTGACGCTGATGCATGAAAGCGTGGGG |
| P-asv6656 | GCAAGCGTTGTCCGGAATTATTGGGCGTAAAGCGCGCGCAGGCGGCGCTTTAAGTTGGGGGTTAAAGGCAATGGCTCAACCATTGTTCGCCTCCAAAACTGGAGCGCTTGAGTGCAGCAGAGGAAAGTGGAATTCCCGGTGTAGCGGTGGAATGCGTAGAGATCGGGAGGAACACCAGTGGCGAAGGCGACTTTCTGGGCTGTAACTGACGCTGAGGCGCGAAAGCGTGGGG |
| P-asv6657 | GCTAGCGTTGTTCGGAATCACTGGGCGTAAAGCGTGCGTAGGCGGCTATTCAAGTCAGAGGTGAAAGCCTGGGGCTCAACCCCAGAACTGCCTTTGAAACTAGGTAGCTCGAACACGGGAGAGGTGAGTGGAATTCCGAGTGTAGAGGTGAAATTCGTAGATATTCGGAAGAACACCAGTGGCGAAGGCGGCTCACTGGACCGTTGTTGACGCTGAGGCACGAAAGCGTGGGG |
| P-asv6658 | GCGAGCGTTGTCCGGAATCATTGGGCGTAAAGCGCGTGTAGGCGGCTTGGTAAGTCTGGTGTGAAATCCTGGGGCTTAACCCCAGACCTGCACTGGATACTGTCAAGCTTGAGTCCGGGAGGGGAGAGCGGAATTCCTAGTGTAGCGGTGAAATGCGCAGATATTAGGAAGAACACCGGTGGCGAAGGCGGCTCTCTGGAACGGTACTGACGCTGAGACGCGAAAGCGTGGGG |
| P-asv6659 | CCTGGCGTTATCCGGATTTATTGGGTTTAAAGGGAGCGCAGGCTGCACTTTAAGCGTGTTGTGAAATGTACCGGCTCAACCGGTAACGTGCAGCGCGAACTGGGGTGCTTGAGTACGAAGAGGGAAGGCGGAACTCGTGGTGTAGCGGTGAAATGCTTAGATATCACGAGGAACTCCGATCGCGAAGGCAGCTTTCCGTTTCGGAACTGACGCTGAGGCTCGAAAGTGCGGGT |
| P-asv6660 | GCAAGCGTTGCTCGGAATCATTGGGCGTAAAGGGGAGGTAGGTGGTCTCGCATGTCCGAAGTAAAATCCCTGGGCTTAACCCAGGACGGTCTTTGGAAACTGCGGGACTTGAGTCTTGGAGAGGGTCGCGGAATTCCCGGTGTAGCGGTGAAATGCGTAGAGATCGGGAGGAACACCAGAGGCGAAGGCGGCGACCTGGACAAGTACTGACACTAAACTCCGAAAGCGTGGGG |
| P-asv6661 | GCAAGCGTTGTTCGGAATCACTGGGCGTAAAGGGAGCGTAGGTGCAATTGCAAGTGGACCGTGCAAACCCGGGGCCCAACCCCGGACTCGCGGTCCAGACTGCAGTTCTTGAGTTATTCAGAGGTAGACGGAATTCCTGGTGTAGCGGTGGAATGCGTAGAGATCAGGAAGAACACCGATGGCGAAGGCAGTCTACTGGGGATCAACTGACACTGAGGCTCGAAAGCGTGGGT |
| P-asv6662 | GCAAGCGTTAATCGGAATGACTGGGCGTAAAGGGCACGTAGGCGGTTATTTAAGTTGGGTGTTAAAGACCAGGGCTTAACCTTGGAGATGCACTCAAGACTGGATAGCTAGAATATAGCAGAGGCTGATGGAATTTCCGGTGTAGCGGTGAAATGCGTAGAGATCGGAAGGAACACCAGTGGCGAAGGCGATTGGCTGGGCTAATATTGACGCTGAGGTGCGAAAGCGTGGGG |
| P-asv6663 | GCGAGCGTTGTTCGGAATTATTGGGTGTAAAGGGCGTCTAGGCGGGAAATTAAGTTAGATGTGAAAGGTGTGGGCTCAACCCGCGAACTGCATCTAAAACTGGTTTTCTTGAGTTCGGGAGAGGAGAGCGGAATTCCTGGTGTAGCGGTGAAATGCGTAGATATCAGGAAGAACACCAGTGGCGAAGGCGGCTCTCTGGCCTAGAACTGACGCTGAAGCGCGAAAGCGTGGGG |
| P-asv6664 | TCAAGCGTTACCCGGATTTATTGGGCGTAAAGCGTCCGCAGGTGGTTTAGTGCGTCTTATGTTAAATGTTGAGGCTCAACCTCAACACTGCGTGAGATACGGCTAGACTTGAGACTGGGAGAGGTAAGCGGAACTACCGGTGTAGTAGTAAAATGCGTTAATATCGGTATGAACACCAAAAGCGAAAGCAGCTTACTGGAACAGTTCTGACACTCATGGACGAAAGCGTGGGG |
| P-asv6665 | GCAAGCGTTGTCCGGAATTATTGGGCGTAAAGCGCGCGCAGGCGGTTTCTTAAGTCCATCTTAAAAGCGTGGGGCTCAACCCCATGAGGGGATGGAAACTGGGAAGCTGGAGTATCGGAGAGGAAAGTGGAATTCCTAGTGTAGCGGTGAAATGCGTAGAGATTAGGAAGAACACCGGTGGCGAAGGCGACTTTCTAGACGAAAACTGACGCTGAGGCGCGAAAGCGTGGGG |
| P-asv6666 | GCAAGCGTTATCCGGAATCACTGGGCGTAAAGGGTGCGTAGGCGGCCTAACAAGTCAGAGGTGAAAGGCATTGGCTCAACCAATGTAAGCCTTTGAAACTGTTGGGCTTGAGTGCAGGAGAGGTAAGTGGAATTCCTAGTGTAGAGGTGAAATACGTAGAGATTGGGAAGAACACCAGTGGCGAAGGCGGCTATCTGGACGGTTTCTGACGCTAAGACGCGAAAGCGTGGGG |
| P-asv6667 | TCCCTCTTTCCTCTTCCTTCCTTTTCTTCTCTCTCTCGTAGGTGGTTCAGTCCGTTGGAAGTGCAATCCCCGGGCTCAACCTGGGAACTGCTTTCAAAACTGCTGAGCTAGAGTACGGTAGAGGGTGGTGGAATTTCCTGTGTAGCGGTGAAATGCGTAGATATAGGAAGGAACACCAGTGGCGAAGGCGACCACCTGGACTGATACTGACACTGAGGTGCGAAAGCGTGGGG |
| P-asv6668 | GCGAACGTTATTCGGAATCACTGGGCTTAAAGCGCGTGTAGGTGGAAAGGTACGTCTGTCGTTGAAATCCCCCGGCTCAACCGGGGAACGGGCGTGGATACGGCCTTTCTTGAGGAACGTAGGGGGACCTGGAACTTCCGGTAGAGTGGTGAAATGCGTTGAGATCGGATGGAACGCCCGTGGCGAAAGCGAGGTCCTGGACGTTTTCTGACACTGAGACGCGAAAGCCAGGGG |
| P-asv6669 | GCAAGCGTTGTTCGGAATCACTGGGCTTAAAGAGCACGTAGGCGGCCTGTCAAGCGTCTTGTGAAATCCCTCGGCTCAACCGAGGAATGGCTGGGCGAACTGGCAGGCTTGAGGCAGGTAGGGGCACACGGAACTCTTGGTGGAGCGGTGGAATGCGTAGATATCAAGAGGAACGCCGGTGGCGAAGGCGGTGTGCTGGGCCTGTCCTGACGCTGAGGTGCGAAAGCCAGGGG |
| P-asv6670 | CCGAGCGTTATCCGGATTTATTGGGTTTAAAGGGTGCGTAGGCGGCTTTTTAAGTCAGGGGTGAAAACTTGCGGCTCAACCGTAAGCCTGCCTTTGATACTGAAGAGCTTGAATACACTTGAGGTAGGCGGAATGTGACAAGTAGCGGTGAAATGCATAGATATGTCACAGAACACCAATTGCGAAGGCAGCTTACTAAAGTGTGATTGACGCTGAGGCACGAAAGCGTGGGG |
| P-asv6671 | GCAAGCGTTGTCCGGAATCATTGGGCGTAAAGAGCTCGTAGGCGGTTCGTCGCGTCGGCTGTGAAATCCCGAGGCTCAACCTCGGGCCTGCAGTCGATACGGGCGGACTTGAGTTCGGCAGGGGAGACTGGAATTCCTGGTGCAGCGGTGAAATGCGCAGATATCAGGAAGAACACCGGTGGCGAAGGCGGGTCTCTGGGCCGAAACTGACGCTGAGGAGCGAAAGCGTGGGG |
| P-asv6672 | GCGAGCGTTAATCGGAATTACTGGGCGTAAAGCGCACGTAGGCGGTTCTGTAAGTGGGATGTGAAACCCCCGGGCTTAACCTGGGACTTGCATACCAGACTGCAAAGCTAGAGTATGAGAGAGGGTGGTGGAATTTCCGGTGTAGCGGTGAAATGCGTAGAGATCGGAAGGAACATCAGTGGCGAAGGCGGCCACCTGGCTTAATACTGACGCTGAGGTGCGAAAGCGTGGGG |
| P-asv6673 | GCGAGCGTTAATCGGATTTACTGGGCGTAAAGCGTGCGTAGGCGGCTTTTTAAGTCGGATGTGAAATCCCTGAGCTTAACTTAGGAGTTGCATTCGATACTGGGAAGCTAGAGTATGGGAGAGGATGGTAGAATTCCATGTGTAGCGGTGAAATGCGTAGATATATGGAAGGACACCGGTGGCGAAGGCGGCTCTCTGGTCTGTTACTGACGCTGAGGCTCGAAAGCATGGGT |

| | |
|---|---|
| P-asv6674 | GCAAGCGTTGTCCGGATTCATTGGGCGTAAAGAGCTCGTAGGCGGTTCAGTAAGTCGGGTGTGAAAGCTCCGAGCTCAACTCGGAGAGGCCACCCGATACTGCTGTGACTTGAGTCCAGTAGAGGAGCGTGGAACTCCTGGTGTAGCGGTGGAATGCGCAGATATCAGGAAGAACACCTGCAGCGAAGGCGGCGCTCTGGGCTGGAACTGACGCTGAGGAGCGAAAGCGTGGGT |
| P-asv6675 | GCAAGCGTTGTCCGGAATCACTGGGTGTAAAGGGTGTGTAGGCGGGATTGTAAGTCAGAGGTGAAATCTTGCGGCTCAACCGTAAAATTGCCTTTGATACTGCAGTCCTTGAGTCCCGGAGAGGTTGTTGGAATTCGTGGTGTAGCGGTGAAATGCGTAGATATCACGAGGAACACCAGAAGCGAAAGCGAACAACTGGACGGGTACTGACGCTAAGGCACGAAAGCGTGGGG |
| P-asv6676 | GCGAGCGTCGTCCGGAATGACTGGGCGTAAAGGGCGTGTAGGCGGCCGATTAAGTGTGGAGTGAAAGTCCATTTTTCAAGGATGGAATTGCTTTGCAGACTGGTTGGCTTGAGTGCGGAAGAGGTAAGTGGAATTCCCAGTGTAGCGGTGAAATGCGTAGAGATTGGGAGGAACACCAGTGGCGAAGGCGACTTACTGGGCCGTAACTGACGCTGAGGCGCGAAAGCGTGGGG |
| P-asv6677 | CCAAGCGTTAATCGGAATTACTGGGCGTAAAGCGTGCGCAGGCGGTTGTGCAAGACCGATGTGAAAGCCTGTAGCTCAACTCCAGAATTGCCTTTAAGACTGCATCGCTTGAATCCAGGAGAGGTGAGTGGAATTCCGAGTGTAGAGGTGAAATTCGTAGATATTCGGAAGAACACCAGTGGCGAAGGCGGCTCACTGGACTGGTATTGACGCTGAAGTGCGAAAGCGTGGGG |
| P-asv6678 | GCAAGCGTTATTCGGAGTGATTGGGCGTAAAGGGTGTGTAGGCGGTATGTTAAGTCAACTGTTAAATTTCTCAGCTCAACTGGGAGCAAGCGGTTGAAACTGGCGTGCTTGAGGATGGAAGAGAGAAGTGGAATTCTCGGAGTAGCGGTAAAATGCGTAGATCTCGAGAGGAACACCGATGGCGAAGGCAGCTTCTTGGTCCATTTCTGACGCTGAGGCACGAAAGCGTGGGG |
| P-asv6679 | CCAAGCGTTATCCGGAATCATTGGGTTTAAAGGGAGCGTAGGCGGGAGGTCAAGTCCGGTGTGAAAGACCCTGGCTCAACCAGGGATGAGGGCTGGAAACTGACAATCTTGAGTACAGCAGAGGAAGAGGGAATTACCGGTGTAGGGGTGAAATCCGTAGAGACCGGTAAGAAGGCCAGTGGCGAAGGCGCTTTTCTGGGCTGTTACTGACGCTGAGGCTCGAAAGCCAGGGG |
| P-asv6680 | CCGAGCGTTAATCAGAATCACTGGGCTTAAAGGGTGCGCAGGCGGATTGGTAGGTGTGCTGTGAAATCCCTCGGCTCAACCGAGGAATTGCGGTGCAAACCCCCAATCTTGAGGCAGACAGAGGTCGTCGGAACGTTCGGTGGAGCGGTGAAATGCGTAGATATCGAACGGAACGCCAAAGGGGTAGCCAGACGACTGGGTCTGTCCTGACGCTCAGGCACGAAAGCGTGGGG |
| P-asv6681 | GCAAGTGTTACTCGTCATAATTGGGCGTAAAGGGTGCGCAGGCGGTCTTTTAAGTTATGGGTGAAAGGTTAATTTTAGAAGTTAAAAAGACCTGTAATACTAAAGGAGTAGAGTATGAAAGAGGAAAATGGAACTTTTACAGTAGAGGTGGAATTCAGAGATAGTAAAGGGACGACCAAAGGCGAAGGCAATTTTCTGGGTCATTACTGACGCTGAAGCACGAAAGCATGGGG |
| P-asv6682 | GCGAGCGTTATCCGGAATTATTGGGCGTAAAGCGTCCGTATGCGGTTTTGTTAGTCCGTGGTTAAATCCCAGAGCTCAACTCTGGAATCGTCATGGAAACTGCAAAACTAGAGTAGTGCAGAGGCAGATGGAATTTCGGAAGTAGGGGTAAAATCCGTAGATCTCCGAAGGAACACCAAACGCGAAGGCATTCTGCTGGGCACTTACTGACGCTGAGGGACGAAAGCGTGGGG |
| P-asv6683 | GCGAGCGTTAATCGGAATTACTGGGCGTAAAGCGTGCGCAGGCTGTTTTGTAAGTCAGATGTGAAATCCCGGGGCCCAACCCCGGAACTGCCTTTGAAACTGGAGAGCTAGAATCTTGGAGAGGCGAGTGGAATTCCGAGTGTAGAGGTGAAATTCGTAGATATTCGGAAGAACACCAGTGGCGAAGGCGACTCGCTGGACAAGTATTGACGCTGAGGTGCGAAAGCGTGGGG |
| P-asv6684 | GCAAGCGTTACTCGGATTTACTGGGCGTAAAGGGTGCGCAGGCGGCCAGGTGTGTCTAGGGTGAAATACTGCAGCTTAACTGTAGAACTGCCTTAGAAACTACCTGGCTAGAGTATCGGAGAGGCAAGCGGAATTCCAGGTGTAGCAGTGAAATGCGTAGATATCTGGAGGAACACCGAAGGCGAAGGCAGCTTGCTAGACGATTACTGACGCTCAGGCACGAAAGCATGGGT |
| P-asv6685 | GCAAGCGTGGTCCGGAATCATTGGGCGTAAAGAGTTCGTAGGTGGTATGTTAAGTCTGGTGTTAAAGCCCGAAGCCCAACTTCGGTTCGGCACTGGATACTGGCAAACTTGAATGTGGTAGAGGTAAAGGGAATTCCTGGTGTAGCGGTGAAATGCGTAGATATCAGGAGGAACATCGGTGGCGTAAGCGCTTTACTGGGCCATAATTGACACTGAGGAACGAAAGCCGGGGT |
| P-asv6686 | GCAAGCGTTGTTCGGAATTATTGGGCGTAAAGGGCATGTAGGCGGTTTGGTAAGCCTGGTGTGAAATCCTGCAGCTTAACTGTAGAATTGCATTGGGTACTGCCAGACTTGAATCACAGAGGGGAAACTAGAATTCCAGGTGTAGGGGTGGAATCTGTAGATATCTGGAAGAATACCGGCGGCGAAGGCGGGTTTCTAGCTGATGATTGACGCTGAGGTGCGAAAGTGCGGGG |
| P-asv6687 | CCAAGCGTTGTTCGGATTCACTGGGCGTAAAGGGTGCGTAGGCGGTGGGATAAGTCTGGTGTGAAAGCTCGGAGCTTAACTCCGAAATTACATTGGAAACTATCTCACTTGAGGTTTGGAGGGGGGACTGGAATTCTCGGTGTAGCAGTGAAATGCGTAGATATCGAGAGGAACACCAGTGGCGAAGGCGAGTCCCTGGACAAATCCTGACGCTGAGGCACGAAAGCTAGGGG |
| P-asv6688 | GCAAGCGTTGTTCGGAATCACTGGGCTTAAAGCGTGCGCAGGCGGATTTGCAAGTTGGGAGTGAAATCCCCGGGCTCAACCCGGGAATTGCTCTCAAAACTGCAAATCTAGAATCCAGGAGAGGTTAGCAGAATTCCGAGTGTAGAGGTGAAATTCGCAGATATTCGGAGGAATACCAATGGCGAAGGCACTTTACTGGGCTATTACTGACACTCAGAGACGAAAGCTAGGGT |
| P-asv6689 | GCGAGCGTTGTTCGGAATCACTGGGCGTAAAGGGCGTGTAGGCGGTCCGCTAAGTCTGATGTGAAAGCCCGGGGCTCACCCCCGGAAGGGCATTGGAAACTGGCGGACTCGAGTCCCGGAGAGGAGGGTGGAATTCCTGGTGTAGCGGTGAAATGCGTAGATATCAGGAGGAACACCGGTGGCGAAGGCGGCCCTCTGGACGGTGACTGACGCTGAGACGCGAAAGCGTGGGG |
| P-asv6690 | GCAAGCGTTACTCGGAATCACTGGGCGTAAAGAGCGTGTAGGCGGGTATATAAGTCAGAAGTGAAATCCAATAGCTTAACTATTGAACTGCTTTTGAAACTGTATACCTAGAATGTGGGAGAGGTAGATGGAATTTCTGGTGTAGGGGTAAAATCCGTAGAGATCAGAAGGAATACCGATGGCGAAGGCAGCCCCCTGGGCCTGTACCGACGCTCATGCACGAAAGCGTGGGG |
| P-asv6691 | TCCCTCTTTCCTCGGAATTACTGGGCGTAAAGCGCACGCAGGCGGCTTTTTAAGTCAGGGGTGAAATCCCGGGGCTCAACCCCGGAACTGCCCTTGAAACTGGGAAGCTAGAATCTTGGAGAGGCGAGTGGAATTCCGAGTGTAGAGGTGAAATTCGTAGATATTCGGAAGAACACCAGTGGCGAAGGCGACTCGCTGGACAAGTATTGACGCTGAGGTGCGAAAGCGTGGGG |
| P-asv6692 | GCAAGCGTTGTTCGGATTCACTGGGCGTAAAGGGTGTGTAGGTGGTTTGGTAAGTTGGATGTGAAAGGCCACAGCTCAACTGGGGAATTGCATCCAAAACTACCTTACTAGAGGATTTGGGAAGTGAGCGGAATTCTAGGTGTAGCGGTGAAATGCGTAGATATCTAGAAGAACACCAGTGGCGTAGGCGGCTCACTATCATTCTCCTGACACTGAGACACGAAAGCATGGGG |
| P-asv6693 | GCAAGCGTTGTTCGGATTTACTGGGCGTAAAGCGCGCGCAGGCTGATTATTAAGCCAGATGTGAAATCCCTGAGCTTAACTTAGGAATTGCATTCGATACTGGGAAGCTAGAGTATGGGAGAGGATGGTAGAATTCCAGGTGTAGCGGTGAAATGCGTAGAGATCTGGAGGAATACCGATGGCGAAGGCAGCCATCTGGCCTAATACTGACGCTGAGGTACGAAAGCATGGGG |
| P-asv6694 | ACAAACGTTATTCGGAATTACTGGGCTTAAAGGGTGCGTAGGCGGCGCGGAAAGTTGGGTGTGAAATCCCTCGGCTCAACCGAGGAACTGCGCCCAAAACTACCGTGCTCGAGGAAGATAGAGGTGAGCGGAACTTACGGTGGAGCGGTGAAATGCGTTGATATCGTAAGGAACACCCGTGGCGAAAGCGGCTCACTGGGTCTTTTCTGACGCTGATGCACGAAAGCTAGGGG |
| P-asv6695 | GCAAGCGTTATCCGGATTTATTGGGCGTAAAGCGCGTCTAGGCGGAAAAGTAAGTCTGATGTGAAAATGCGGGGCTCAACTCCGTATTGCGTTGGAAACTGCTTTTCTAGAGTACTGGAGAGGTGGGCGGAACTACAAGTGTAGAGGTGAAATTCGTAGATATTTGTAGGAATGCCGATGGAGAAGTCAGCTCACTGGACAGATACTGACGCTGAGGTACGAAAGCATGGGG |
| P-asv6696 | GCAAGCGTTGTCCGGAATGATTGGGCGTAAAGGGCGCGTAGGCGGCCATGTAAGTCCGATGTGAAAGTCCTGCTTTTAAGGTGGGAATTGCATTGGAAACTGTGTGGCTTGAGTGCAGGAGAGGTAAGTGGAATTCCCGGTGTAGCGGTGAAATGCGTAGAGATCGGGAGGAACACCAGTGGCGAAGGCGACTTACTGGACTGTAACTGACGCTGAGGCGCGAAAGTGTGGGG |
| P-asv6697 | GCAAGCGTTATCCGGATTTATTGGGTTTAAAGGGTCCGTAGGCGGGCTGATAAGTCAGTGGTGAAAGCCGACAGCTTAACTGTCGAACTGCCATTGATACTGTCAGTCTTGAGTATATTTGAGGTAGCTGGAATAAGTAGTGTAGCGGTGAAATGCATAGATATTACTTAGAACACCAATTGCGAAGGCAGGTTACCAAGATATAACTGACGCTGATGGACGAAAGCGTGGGG |
| P-asv6698 | CCAAGCGTTGTTCGGATTCACTGGGCGTAAAGGGTGCGCAGGTGGCCAGGCAAGTCGGATGTGAAAGCTCCGGGCTCAACCCGGAAATGGCATTGGAAACTGCTTGGCTTGAGGTTTGGAGGGGGGACTGGAATTCTCGGTGTAGCAGTGAAATGCGTAGATATCGAGAGGAACACCAGTGGCGAAGGCGAGTCCCTGGACAAGACCTGACACTGAGGCACGAAAGCTAGGGG |
| P-asv6699 | GCAAACGTTGCTCGGAATTATTGGGCGTAAAGCGCGCATAGGCGGCCGCATAAGTCTGATGTGAAAGCCCAGGGCTCAACCCTGGAAAGCCATTGGAAACTGTGCGGCTTGAGAACAGGAGGGGTAGGTGGAATTCCTGGTGTAGGAGTGAAATCCGTAGAGATCAGGAGGAACACCGGTGGCGAAGGCGACCTACTGGACTGGATCTGACGCTGAGGCGCGAAAGCGTGGGG |
| P-asv6700 | GCAAGCGTTGTTCGGATTTATTGGGCGTAAAGCGCATGTAGGCGGTATAGTAAGTCGGATGTTAAAGACCCACGGCTCACCCGGGGGAATGCGTTCGATACTGCTATTCTTGAGTGCAGGAGAGGAAAGTGGAATTCCTGGTGTAGCGGTGAAATGCGTAGATATCAGGAGGAACACCTGTGGCGAAGGCGGCTTTCTGGCCTGCTACTGACGCTGAGATGCGAAAGCGTGGGG |
| P-asv6701 | GCTAGCGTTGCTCGGAATTACTGGGCGTAAAGGGCGCGTAGGCGGACAGTTAAGTTGGGGGCGAAATCCTAGAGCTTAACTCTAGAACTGCCATTGATACTATTTATCTTGAATATTGTGGAGGTAAGCGGAATATGTCATGTAGCGGTGAAATGCTTAGATATGACATAGAACACCTATTGCGAAGGCAGCTTACTACGCATATATTGACGCTGAGGCACGAAAGCGTGGGG |
| P-asv6702 | GCGAACGTTATTCGGATTCACTGGGCTTAAAGGGTGCGTAGGCGGTCTGTTAAGCAGGGTGTGAAATTTTTCGGCTCAACCGGAAAAGTGCATTCTGTACTGGCGGACTGGAGTTGGTCAGGGGTGTGTGGAACTGCCAGTGGAGCGGTGAAATGTGTTGATATTGGCAGGAACGCCGGTGGCGAAAGCGACACACTGGGGCCGTACTGACGCTGAGGCACGAAAGCCAGGGG |
| P-asv6703 | CCAAGCGTTGTTCGGATTTACTGGGCGTAAAGGGCGCGTAGGCGGCGTGTTAAGTCAGCTGTGAAATCTCCGAGCTTAACTCGGAACGGTCAGCTGATACTGATGTGCTAGAGTGCAGAAGGGGCAATCGGAATTCTTGGTGTAGCGGCGAAATGCGTAGATATCAAGAGGAACACCTGAGGTGAAGACGGGTTACTGGGCTGACACTGACGCTGAGGCGCGAAAGCCAGGGG |
| P-asv6704 | GCAAGCGTTATTCGGCATGACTGGGCGTAAAGGGTCCGTAGACGGTAGAGTAAGTTCTTGGTTAAATCTTAAAGCCTAACTTTAAAATAACTTAGAATACTGCTTTACTTGAGTTTTGTACAGAAGACTGGAATTTTATACGGAAGGGTAAAATCTGTAGATATATAAGGGAACACCAGTAGCGAAGGCGAGTCTTTGGTAAATACTGACGTTGAGGGACGAAAGTATGGGT |
| P-asv6705 | GCGAGCGTTGTTCAGATTTACTGGGCGTAAAGGGCACGTAGGCGGTCCTGTAAGCCTGACGTGAAAGGCTATGGCTCAACCATGGCACTGCGTTGGGAACTGCGGGACTTGAATCATGGAGAGGGAGCTGGAATTCCAGGTGTAGGGGTGAAATCTGTAGATATCTGGAAGAACACCAATGGCGAAGGCAAGCTCCTGGCCAATGATTGACGCTGAGGTGCGAAAGTGTGGGG |
| P-asv6706 | GCAAACGTTGTTCGGAATCACTGGGCATAAAGGGTGCGTAGGCGGCTTTGTAAGTCGGGTGTGAAAGCCCGGAGCTCAACTCCGGAATGGCGCCTGATACTGCAGGGCTTGAGGAATAAAGGGGTGCGGGGAACTCTTGGTGGAGCGGTGAAATGCGTAGAGATCAAGGGGAACGCCGGTGGCGAAAGCGCCGCACTGGTTATATCCTGACGCTGAGGCACGAAAGCTAGGGG |
| P-asv6707 | GCAAGCGTTATCCGGATTCGCTGGGTTTAAAGGGTGCGTAGGCGGGTAGGTAAGTCAGAGGTGAAAGCCTGGAGCTCAACTCCAGAACTGCCTTTGAGACTGCATCGCTTGAATCCAGGAGAGGTGAGTGGAATTCCGAGTGTAGAGGTGAAATTCGTAGATATTCGGAAGAACACCAGTGGCGAAGGCGGCTCACTGGACTGGTATTGACGCTGAAGTGCGAAAGCGTGGGG |
| P-asv6708 | GCAAGCGTTGTCCGGAATTATTGGGCGTAAAGGGCGCGCAGGCGGCTTCTTAAGTCTGTCTTAAAAGTGCGGGGCTTAACCCCGTGATGGGATGGAAACTGGGAAGCTCAGAGTATCGGAGAGGAAAGCGGAATTCCTAGTGTAGCGGTGAAATGCGTAGATATTCGGAAGAACACCAGTGGCGAAGGCGACTGACTGGACAAGTATTGACGCTGAGGTACGAAAGCGTGGGG |
| P-asv6709 | GCGAGCGTTAATCGGAATTACTGGGCGTAAAGGGCGCGTAGGCGGTGAAGTCAGTCGGGTGTGAAAGCCCCGGGCTCAACCTGGGAACTGCATTCGATACTGCTTCGCTAGAGTATGGTAGAGGGAAGCGGAATTCCGGGTGTAGCGGTGAAATGCGTAGATATCCGGAGGAACACCAGTGGCGAAGGCGGCTTCCTGGACCAATACTGACGCTGAGGCGCGAAAGCGTGGGG |
| P-asv6710 | GCGAGCGTTGTCCGGAATCATTGGGCGTAAAGCGCACGTAGGCGGCCCTGTAAGTCGAATGTGAAATCCCGAGGCTCAACCTCGGAACTGCATTCGGTACTGCAGGGCTTGAGGAAGGTAGAGGAGAGTGGAATTCCCGGTGTAGCGGTGGAATGCGCAGATATCGGGAGGAACACCAGCGGCGAAGGCGGCTCTCTGGGCCTTACCTGACGCTGAGGTGCGAAAGCGTGGGG |
| P-asv6711 | ACAAGCATTGTCCGGAATCACTGGGCGTAAAGCGTTTCGCAGGTGTTTCTCTAAGTTACGTTTGAAAGACCGGGGCTCAACCTCGGGAACGGGCGTAATACTGGAGGGATTGAAGGGTATCGGGGCTATTGGAACTGTCGGTGTAGGGGTGAAATCCGTTGATATCGACAGGAACTCCGAGGGCGTAGGCAGATAGCTAGGTACCTCTTGACACTCATGAACGAAAGCTAGGGG |
| P-asv6712 | GCAAGCGTTGTCCGGAATTATTGGGCGTAAAGAGCTCGTAGGTGGTTCGTCACGTCGGATGTGAAACTCTGGGGCTTAACCCCAGACCTGCATTCGATACGGGCGAGCTTGAGTATGGTAGGGGAGTCTGGAATTCCTGGTGTAGCGGTGGAATGCGCAGATATCAGGAGGAACACCAATGGCGAAGGCAAGACTCTGGGACATTACTGACACTGAGGAGCGAAAGCGTGGGG |

| | |
|---|---|
| P-asv6713 | GCAAGCGTTATCCGGATTCACTGGGTTTAAAGGGTGCGTAGGTGGCTTGTAAGTCAGTGGTGAAATCTCCGTGCTTAACATGGAAACTGCCGTTGATACTATAGGTCTTGAATTTTCTGGAGGTAAGCGGAATATGTCATGTAGCGGTGAAATGCTTAGATATGACATAGAACACCAATTGCGAAGGCAGCTTACTACAGGGATATTGACACTGAGGCACGAAAGCGTGGGG |
| P-asv6714 | GCAAGCGTTGTCCGGATTTACTGGGTGTAAAGGGAGCGCAGGCGGGAAATCAAGTCAGCTGTGAAAACTATGGGCTTAACTCATAGACTGCAGTTAAAACTGTTTTTCTTGAGTGAAGTAGAGGTAGGCGGAATTCCGAGTGTAGCGGTGAAATGCGTAGAGATTCGGAGGAACACCAGTGGCGAAGGCGGCCTACTGGGCTTTAACTGACGCTGAGGCTCGAAAGTGTGGGG |
| P-asv6715 | CAAGCGTTACCCGGAATTACTGGGCGTAAAGGGAGCGTAGGCGGGAGGTCAAGTCCGGTGTGAAAGACCCTGGCTCAACCAGGGATGAGGGCTGGAAACTGACAATCTTGAGTACAGCAGAGGAAGAGGGAATTACCGGTGTAGGGGTGAAATCCGTAGAGACCGGTAAGAAGGCCAGTGGCGAAGGCGCTTTTCTGGGCTGTTACTGACGCTGAGGCTCGAAAGCCAGGGG |
| P-asv6716 | GCGAACGTTATTCGGAATCACTGGGCTTACAGGGTGCGTAGGCGGCCAGACAAGTCAGGTGTGAAAGCCCTCGGCTTAACCGAGGAACTGCGCTTGAAACTGTTTGGCTCGAGTCAGCTAGAGGTACACGGAACTTCCAGTGGAGCGGTGAAATGTGTTGATATTGGAAGGAACACCGGAGGCGAAAGCGGTGTACTGGGGCTGGTCTGACGCTGAGGCACGAAAGCCAGGGG |
| P-asv6717 | GCAAGCGTTGTCCGGAATCACTGGGTGTAAAGGGTGTGTAGGCGGGACGGTAAGTCAGTGGTGAAAGCCCTCGGCTCAACCGAGGAATGGCCTCTGATACTGCCGTTCTTGAGTCCTGGAGGGGCCGTTGGAATTCGTGGTGTAGCGGTGAAATGCGTAGATATCACGAGGAGCACCAGAGGTGTAGACGGACGGCTGGACAGGCACTGACGCTCAGGCACGAAAGCGTGGGG |
| P-asv6718 | TCGAGCGTTGTCCGGAGTTACTGGGCGTAAAGCGCGCGCAGGTGGCTCATTACGCCCGGCGTGAAAGCCCCCGGCTCAACCGGGGAGGGTCGTCGGGGACGGGTGAGCTTGAGGGTATCAGGGGCTGGTAGAATTCCCGGTGTAGTGGTGAAATGCGTAGAGATCGGGAGGAATACCCGTGGCGAAGGCGGCCAGCTGGGATACACCTGACACTGAGGCGCGAAGGCGTGGGG |
| P-asv6719 | GCAAGCGTTGTCCGGATTTATTGGGTTTAAAGGGTGCGTAGGCGGCCTGTTAAGTCAGTGGTGAAATCCAACAGCTCAACTGTTGATGTGCCAATGATACTGACAGGCTTGAGTACAGACGAGGTAGGCGGAATTGACAGTGTAGCGGTGAAATGCATAGATATTGTCAAGAACACCGATAGCGAAGGCAGCTTGCTAGCCTGTAACTGACGCTGAGGCACGAAAGTATGGGG |
| P-asv6720 | CCGAGCGTTGTTCGGAATCACTGGGCGTAAAGGGAGCGTAGGCGGCGTGGTAAGTCAGATGTGAAATCCCGGGGCTCAACCCCGGAACTGCATCCGATACTGCCGTGCTAGAGTAATGGAGGGGTAGCTGGAATTCTCGGTGTAGCAGTGAAATGCGTGGATATCGAGAGGAACACTCGTGGCGAAAGCGAGCTACTGGACATTAACTGACGCTGAGGCTCGAAGGCCAGGGG |
| P-asv6721 | GCTAGCGTTATTCGGAATTACTGGGCGTAAAGCGCGCGTAGGCGGATCAGAAAGTTGGGGGTGAAATCCCAGGGCTCAACCCTGGAACTGCCTTTAATACTGGCAATCTCGAGTCCGAGAGAGGTGAGTGGAATTCCGAGTGTAGAGGTGAAATTCGTAGATATTCGGAGGAACACCAGTGGCGAAGGCGGCTCACTGGCTCGGTACTGACGCTGAGGTGCGAAAGCGTGGGG |
| P-asv6722 | GCGAGCGTTGTCCGGAATTATTGGGCGTAAAGGGTACGTAGGCGGCCTTTTAAGTCAGGTGTGAAAGCGTGAGGCTTAACCTCATTAAGCACTTGAAACTGGAAGGCTTGAGTGAAGGAGAGGAAAGTGGAATTCCTAGTGTAGCGGTGAAATGCGTAGATATTAGGAGGAATACCGGTGGCGAAGGCGACTTTCTGGACTTTTACTGACGCTCAGGTACGAAAGCGTGGGG |
| P-asv6723 | TCGAGCGTTGCGCGGAATTACTGGGCTTAAAGGGTGCGTAGGCTGATATGCAAGTGCTTTGTGAAATCCCAGAGCTTAACTCTGGAACTGCTTGGCAGACTGCATGTCTTGAGGTCGGTAGGGGTTGCTGGAACGGTTGGTGGAGCGGTGAAATGCGTAGATATCAACCGGAACGCCAAAGGTGAAGACAAGCAACTGGGCCGATTCTGACGCTGAGGCACGAAAGCGTGGGG |
| P-asv6724 | GCGAGCGTTGTTCGGAATTACTGGGCGTAAAGGGCGCGTAGGCGGTTCGGTAAGTAGGGGGTGAAATTCCACGGCTCAACCGTGGGACGGCCTTCTAGACTGCCGGACTTGAGCACGGTAGAGGCAGATGGAATTCCCGGTGTAGCGGTGGAATGCGTAGAGATCGGGAAGAACACCAGTGGCGAAGGCGGTCTGCTGGGCCGTTGCTGACGCTGAGGCGCGACAGCGTGGGG |
| P-asv6725 | ACGAGTGGGAATCACATTTATTTGGCCTAAAGCGTTCGTAGCCGGCCTGGTAAATCTCTTGTGAAATCGCTTCGTTCAAGCAAGCGACGTGCAGGAGACACTGCCAGGCTAGGGACTGGGAAGAGTCAGGGGTATGGATAGGGGAGCAGTAAAATGCTATAATCCTTTCCAGACCACCGGTGGCGAAGGCGCCTGACTAGAACAGATCCGACGGTGAGGGACGAAAGCTAGGGG |
| P-asv6726 | GCGAGCGTTAATCGGATTTACTGGGCGTAAAGCGTGCGTAGGCGGCTTTTTAAGTCGGGTGTGAAATCCCTGAGCTTAACTTAGGAATTGCATTCGATACTGGGAAGCTAGAGTATGGGAGAGGATGGTAGAATTCCAGGTGTAGCGGTGAAATGCTTAGATATCTTCAAGAACACCGATTGCGTAGGCAGCTTACTAGGCTGTAACTGACGCTGAGGCACGAAAGCGTGGGG |
| P-asv6727 | GCAAGCGTTATTCGGCATTATTGGGCGTAAAGGGTGTGTAGGCGGCATTTTAAGTCTGCCGTTAAATCCTTCGGCTTAACCGGAGGTTCGCAGTGGAAACTAAAGTGCTAGAGGATGGAAGAGAGAAGTGGAATTCCCGGAGTAGCGGTAAAATGCGTAGATCTCGGGAGGAACACCAATGGCGAAGGCAGCTTCTTGGTCCATTCCTGACGCTGAGACACGAAAGCGTGGGG |
| P-asv6728 | GCGAGCGTTAATCGGAATTACTGGGCGTAAAGCGTGCGCAGGCGGTTTTGTAAGACAGATGTGAAATCCCCGGGCTTAACCTGGGAACTGCGTTTGTGACTGCAAGGCTAGAGTACGGCAGAGGGGGGTGGAATTCCTGGTGTAGCAGTGAAATGCGTAGATATCAGGAGGAACATCAGTGGCGAAGGCGACACCCTGGACTCATACTGACGCTGAGGTACGAAAGCATGGGG |
| P-asv6729 | TCCCTCTTTCCTCTTCCTTCCTGGGCGTAAAGCGCACGTAGGCGGTGTGTTAAGTTGGATGTGAAAGCCCCGGGCTCAACCTGGGAATTGCATTCAAAACTGGCACGCTAGAGTATGTGAGAGGGGGGTAGAATTCCAAGTGTAGCGGTGAAATGCGTAGAGATCTGGAGGAATACCGATGGCGAAGGCAGCCATCTGGCCTAATACTGACGCTGAGGTACGAAAGCATGGGG |
| P-asv6730 | GCAAGCGTTACTCGGAATTACTGGGCGTAAAGCGTGCGTAGGCGGTTTGTTAAGTCTGATGTGAAAGCCCCGGGCTCAACCTGGGAATGGCATTGGATACTGGCAGGCTAGAATGTGGTAGAGGGTGGTGGAATTCCCGGTGTAGCGGTGAAATGCGTAGAGATCGGGAGGAACATCCGTGGCGAAGGCGGCCACCTGGACCAACATTGGCGCTGAGGCACGGAAGCGTGGGG |
| P-asv6731 | GCGAGCGTTAATCGGAATTACTGGGCGTAAAGCGCGCGTAGGTGGTCTGTTAAGTCGGATGTGAAAGCCCCGGGCTCAACCTGGGAACTGCATTCGATACTGGCAGACTAGAGTGCGAGAGAGGGAGGTAGAATTCCATGCGTAGCGGTGAAATGCGTAGATATGTGGAGGAATACCAGTGGCGAAGGCGGCCTCCTGGCTCGACACTGACACTGAGGTGCGAAAGCGTGGGG |
| P-asv6732 | GCAAGCGTTAATCGGAATTATTGGGCGTAAAGCGAGTGCAGACGGTTACTTAAGCCAGATGTGAAATCCCCAAGCTTAACTTGGGACGTGCATTTGGAACTGGGTGACTAGAGTGTGTCAGAGGGAGGTAGAATTCCACATGTAGCGGTGGAATGCGTAGATATAGGAAGGAACACCAGTGGCGAAGGCGACCACCTGGGCTAATACTGACACTGAGGTGCAAAAGCGTGGGG |
| P-asv6733 | CCAGGCGTTATCCGGATTTATTGGGTTTAAAGGGAGTGTAGGCGGTTTGTTAAGCGTGTTGTGAAATTTAGGTGCTCAACATTTAACTTGCAGCGCGAACTGTCAGACTTGAGTACACGCAACGTATGCGGAATTCATGGTGTAGCGGTGAAATGCTTAGATATCATGAAGAACTCCGATTGCGAAGGCAGCACACGGGAGTGTAACTGACGCTTAAGCTCGAAGGTGCGGGT |
| P-asv6734 | GCAAGCGTTGTCCGGAATTACCGGGCGTAAAGGGCGTGTAGGCGGTCGCACACGTAGCGGTTTTCAGCCGCCGGCTCACCCGGCGGAGGGCGGCTAAACGGTGCGGCTGGAGGGCAGGAGAGGTGCACGGGATTCCTGGTGGAGCAGTGAAATGCGTAGAGATCAGGAAGAACACCAGTGGCGAAGGCGGTGCACTGGCCTGACCCTGACGCTGAGGCGCGATAGCGTGGGG |
| P-asv6735 | GCGAGCGTCAATCGGAATTACTGGGCGTAAAGCGCGCGTAGGCGGTCTGATAAGCCGGTTGTGAAAGCCCCGGGCTCAACCTGGGAACTGCATTTGTGACTGCAAGGCTAGAGTACGGTAGAGGGGGATGGAATTCCGCGTGTAGCAGTGAAATGCGTAGATATGCGGAGGAACACCAATGGCGAAGGCAATCCCCTGGACCTGTACTGACGCTCATGCACGAAAGCGTGGGG |
| P-asv6736 | TCGAGTGGTAGTCACTTTTATTGAGCCTAAAGCGTTTGTAGCCGGATTTGTAAATCTCTAGGAAAATACTTCTGCTTAACTGAAGAAATTTTGGAGAGACTGCAAATCTAGGGACCGGGTGAGGTTGAATGTACTTCTGGGGTAGGGGTAAAATCCTGTAATCCTGGAAGGACGACCGGTGGCGAAGGCGTTCAACTAGAACGGGTCCGACGGTGAGGAACGAAGGCTAGGGG |
| P-asv6737 | GCGAGCGTTGTCCGGAATTATTGGGCGTAAAGGGTACGCAGGCGGTTTACCAAGTTGGATGTGAAATCTTGTAGCTCAACTACAAACGTGCATCCAAAACTGGCTAACTTGAGTTAAGGAGAGGTAAGTGGAATTCCTGGTGTAGCGGTGGAATGCGTAGATATCAGGAGGAATACCGGTGGCGAAGGCGACTTACTGGACTTATACTGACGCTGAGGTACGAAAGCGTGGGG |
| P-asv6738 | GCGAGCGTTAATCGGAATCACTGGGCGTAAAGGGCGCGTAGGCGGTTAGTTACGTCTGCCGTGAAATCCCCGGGCTTAACCCCGGAACTGCCTTTGAAACTAGATGACTGGAATCTTGGAGAGGCGAGTGGAATTCCGAGTGTAGAGGTGAAATTCGTAGATATTCGGAAGAACACCAGTGGCGAAGGCGACTCGCTGGACAAGTATTGACGCTGAGGTGCGAAAGCGTGGGG |
| P-asv6739 | GCTAGCGTTGTTCAGAATTACTGGGCGTAAAGCGCACGTAGGCGGCTTTGTAAGTCAGAGGTGAAAGCCTGGAGCTCAACTCCAGAACTGCCTTTGATACTGGCGACCTCGAGTTCGAGAGAGGTTGGTGGAACTCCGAGTGTAGAGGTGAAATTCGTAGATATTCGGAAGAACACCAGTGGCGAAGGCGGCCAACTGGCTCGATACTGACGCTGAGGTGCGAAAGCGTGGGG |
| P-asv6740 | GCAAGCGTTATCCGGATTCACTGGGTTTAAAGGGTGCGTAGGCGGGTATTAAGTCAGTGGTGAAATCCCCGAGCTTAACTTGGGAATTGCATTGGTGACTGCACGGCTAGAGTGTGTCAGAGGGGGGTAGAATTCCACGTGTAGCAGTGAAATGCGTAGAGATGTGGAGGAATACCGATGGCGAAGGCAGCCCCCTGGGATAACACTGACGCTCATGCACGAAAGCGTGGGG |
| P-asv6741 | GCAAGCGTTGTCCGGAATTATTGGGCGTAAAGCGCGCGCAGGCGGCCTCTTAAGTCTGATGTGAAAGCCCCCGGCTCAACCGGGGAGGGCCATTGGAAACTGGGAGGCTTGAGTATAGGAGAGAAGAGTGGAATTCCACGTGTAGCGATGAAATGCGTAGAGATGTGGAGGAACACCAGTGGCGAAGGCGACTCTTTGGCCTATAACTGACGCTGAGGCGCGAAAGCGTGGGG |
| P-asv6742 | GCAAGCGTTATCCGGATTCACTGGGTTTAAAGGGTGCGTAGGCGGGTATTTAAGTCAGTGGTGAAATCCTAGAGCTTAACTCTAGAACTGCCATTGATACTATTTATCTTGAATATTGTGGAGGTAAGCGGAATATGTCATGTAGCGGTGAAATGCGTAGAGATTGGGTGGAACACCGGTGGCGAAGGCGGCTATCTGGCTCACAGCTGACGATCAGGCACGAAAGCGTGGGG |
| P-asv6743 | ACGAGTAGTGGTCACTTTTATTGAGCCTAAAGCGTTCGTAGCCGGTTTTGTAAATCTTCAGATAAAGCCTGAAGCTTAACTCCAGAAAGTCTGAAGAGACTGCAAGACTTGAGATCGGGTGAGGTTAAACGTACTTTCAGGGTAGGGGTCAAATCCTGTAATCCCGGAAGGACGACCAGTGGCGAAGGCGACTCACTGGCCCGTTACTGACGCTGAGGCGCGAAAGCGTGGGG |
| P-asv6744 | GCTAGCGTTGCTCGGAATCACTGGGCGTAGAGGGCGCGTAGGTGGATCTTTAAGTCAGGGGTGAAATCCCCGAGCTCAACTTGGGAACTGCGTTTGAAACTCCAAAGCTAGAGTATAGGAGAGGGGGGTAGAATTCCACGTGTAGCAGTGAAATGCGTAGAGATGTGGAGGAATACCAATGGCGAAGGCAGCCCCCTGGCCTAATACTGACGCTCATGCACGAAAGCGTGGGG |
| P-asv6745 | GCGAGCGTTATTCGGATTCACTGGGCGTAAAGGGTGCGCAGGCGGCCAAGTGTGTTAGGCGTGAAAGCCCGGGGCTTAACCCCGGAATTGCACCTAAAACTACATGGCTAGAGCATTGGAGAGGGTAGCAGAATTCACGGTGTAGCAGTGAAATGCGTAGATATCGTGAGGAATACCAGAGGCGAAGGCGGCTACCTGGACAATTGCTGACGCTCAGGCACGAAAGCGTGGGG |
| P-asv6746 | GCTAGCGTTGTTCGGAATCACTGGGCGTAAAGCGCGCGTAGGCGGCTATACAAGTCAGAGGTGAAAGCCCGGGGCTCAACCCCGGAATTGCCTTTGAAACTGCATAGCTAGAGTGTCGGAGGGGATAGCGGAATTCCTAATGTAGAGGTGAAATTCGTAGATATTAGGAGGAACACCGGTGGCGAAGGCGGCTATCTGGACGACAACTGACGCTGAGGCGCGAAAGCGTGGGG |
| P-asv6747 | GCAAGCGTTGTTCGGAATTACTGGGCGTAAAGGGTTTGTAGGTGGTCTGTTAAGTCAGACGTGAAATCCCTCAGCTTAACTGGGGAACTGCGTCTGAGATTGATGGACTTGAGGGCAGGAGAGGAACGCGGAATTCCAGGTGTAGCGGTGAAATGCGTAGATATCTGGAGGAACACCGGTGGCGAAGGCGGCGTTCTGGACTGTTTCTGACACTGAAAAACGAAAGCCAGGGG |
| P-asv6748 | GCAAGCGTTGTCCGGAATTACTGGGCGTAAAGGGCGTGTAGGCGGTGCGATACGTAGCGGTTTTAAGTCTCCGGCTCACCCGGAGGAGGGCGGCTAAACGGTCGCGCTGGAGGGCAGGAGAGGTGCGTGGAATTCCTGGTGGAGCGGTGAAATGCGTAGAGATCAGGAAGAACACCCGTGGCGAAGGCGGCGCACTGGCCCGGCCCTGACGCTGAGGCGCGACAGCGTGGGG |
| P-asv6749 | GCTAGCGTTGTTCGGATTTACTGGGCGTAAAGCGCGTGTAGGCGGGTCGTTAAGTTGGGGGTGAAATCCCGGAGCTCAACTCCGGAACTGCCTTCAAAACTGGCGACCTGGACCTAAGTAGAGGCAAGTGGAATTGCGAGTGTAGAGGTGAAATTCGTAGATATTCGCGGGAACACCAGTGGCGAAGGCGACTTGCTGGACTTATGGTGACGCTGAGACGCGAAAGCGTGGGG |
| P-asv6750 | GCTAGCGTTGCTCGGAATCACTGGGCGTAAAGGGTGCGTAGGTGGATCTTTAAGTCAGGGGTGAAATCCTGGAGCTCAACTCCAGAACTGCCTTTGATACTGAGGATCTTGAGTTCGGGAGAGGTAAGTGGAACTGCGAGTGTAGAGGTGAAATGCGCAGATATCAGGAGGAACACCAATGGCGAAGGCAGCTCGCTGGGACGGTATTGACGCTGAGACGCGAAAGCGTGGGG |
| P-asv6751 | GCAAGCGTTAATCGGGATTACTGGGCGTAAAGCGCACGCAGGCGGTCTGTCAAGTCGGAGGTGAAAGGCATTGGCTCAACCAATGTAAGCCTTTGAAACTGTTGGGCTTGAGTGCAGGAGAGGTAAGTGGAATTCCTAGTGTAGAGGTGAAATTCGTAGATATTAGGAGGAACACCAGTGGCGAAGGCGACTTACTGGACTGTAACTGACGCTGAGGCACGAAAGCGTGGGG |

| | |
|---|---|
| P-asv6752 | GCAAACGTTGCTCGGAATCACTGGGCGTAAAGGGCGTGTAGGCGGGAAGGTAAGTCGGGCGTGAAATCCCTGGGCTCAACCCAGGAACTGCACTCGAAACTACTTTTCTTGAGTGCCGGAGAGGAAAGCGGAATTCCTGGTGTAGAGGTGAAATTCGTAGATATCAGGAGGAACACCAGTGGCGAAGGCGGCTTTCTGGACGGTAACTGACGCTGAGACGCGAAAGCGTGGGG |
| P-asv6753 | GCAAGCGTTACTCGGAATTACTAGGCGTAAAGCGCAGGTAGGCGGTTCGATAAGTCTTTTGTGAAAGCTCCCGGCTTAACTGGGAGAGGCCAAGGGAAACTGTTGGACTAGAGTGTGGTAGAGGATGCTGGAATTCCCGGTGTAGCGGTGAAATGCGTAGAGATCGGGAGGAATACCTGTGGCGAAAGCAGGCATCTGGGCCATCACTGACGCTGAGCTGCAAAAGCTAGGGG |
| P-asv6754 | GCGAGCGTTGTCCGGATTTATTGGGCGTAAAGGGCGCGTAGGCGGTGCGGTAAGTCTGTTGTGAAATCTCCCGGCTTAACCGGGAGGGTGCGGTGAATACTGAGGGACTAGAGTACGGGAGAGGAAAGTGGAATTCCTGGTGTAGCGGTGAAATGCGTAGATATCAGGAGGAACACCGGTGGTGTAGACGGCTTTCTGGACCGTAACTGACGCTGAGACACGAAAGCGTGGGT |
| P-asv6755 | GCAAGCGTTGTCCGGATTTATTGGGTTTAAAGGGTGCGTAGGCGGTTCTATAAGTCTGATGTGAAAGCCCCGGGCTCACCCCCGGAAGTGCATTGGAAACTGGTGGACTCGAGTCCCGGAGAGGAGAGTGGAATTCCCGGTGTAGCGGTGAAATGCGTAGATATCAGGAGGAACACCGGTGGCGAAGGCGGCTCTCTGGACGGTGACTGACGCTGAGACGCGAAAGCGTGGGG |
| P-asv6756 | GCAAGCGTTGTCCGGATTTATCGGGCGTAAAGCGAGCGCAGGCGGTCTGTCAAGTCGGATGTGAAATCCCCGGGCTCAACCTGGGAACTGCATCCGAAACTGGCAGGCTAGAGTCTTGTAGAGGGGGGTAGAATTCCAGGTGTAGCGGTGAAATGCGTAGAGATCTGGAGGAATACCGGTGGCGAAGGCGGCCCCCTGGACAAAGACTGACGCTCAGGTGCGAAAGCGTGGGG |
| P-asv6757 | GCAAGCGTTATCCGGAATTATTGGGCGTAAAGAGCTCGTAGGCGGTTTGTCGCGTCTGCTGTGAAAACCCGAGGCTCAACCTCGGGCCTGCAGTGGGTACGGGCAGACTAGAGTGCGGTAGGGGAGATTGGAATTCCTGGTGTAGCGGCGAAATGCGTAGATATCGGAAGGAACATCAGTGGCGAAAGCGGCTTCCTGGACCAGCACCGACGTTCAGGCACGAAAGCGTGGGG |
| P-asv6758 | CCAAGCGTTGTTCGGATTCATTGGGCGTAAAGGGTGTGTAGGTGGCGCCGTAAGTCGGGTGTGAAATCTCGGGGCTTAACTCCGAAACTGCATTCGATACTGCGGTGCTTGAGGACTGGAGAGGAGACTGGAATTCATGGTGTAGCAGTGAAATGCGTTGATATCGGCTGGAACTCCGGTGGCGAAGGCGAGGTGCTGGACCCCTTCTGACGCTGAGGCGCGAAAGCCAGGGG |
| P-asv6759 | GCGAGCGTTAATTGGAGTAACTGGGCGTAAAGGGTGCGTAGGCGGTTTAGCGAGTGTGATGTGAAAGCCCCGGGCTTAACCTGGGAAGTGCAACGCAAACGGCTAGACTAGAGTAGATGAGAGGGTGATGGAATTTTCCGGTGTAGCGGTGAAATGCGTAGAAATCGGAAGGAACGTCGATGGCGAAGGCAGTCACCTGGCATCATACTGACGCTGAGGCACGAAAGCGTGGGG |
| P-asv6760 | GCAAGCGTTGTCCGGATTTACTGGGCGTAAAGCGTGTGTAGGCGGACTTTTAAGTAGAAAGTGAAAGGTTGGTGCTCAACACCAACATTGCTCTCTATACTGGAGGTCTTGAGTGTCGGAGAGGAAGATGGAACTGCACGTGTAGCGGTGAAATGCGTTGAGATGTACAGGAACACCAATGGCGAAGGCAATCTTCTGGACGAAAATTGACGCTGAGACACGAAAGCACAAGGT |
| P-asv6761 | GCAAGCGTTGTTCGGAATTACTGGGCGTAAAGGGCTCGTAGGTGGCCAACTAAGTCAGACGTGAAATCCCCAGGCTTAACCTGGGAACTGCGTCTGATACTGGATGGCTTGAATTCGGGAGAGGGATGCAGAATTCCAGGTGTAGCGGTGAAATGCGTAGATATCTGGAGGAATACCGGTGGCGAAGGCGGCATCCTGGACCGACATTGACGCTGAGGAGCGAAAGCTAGGGG |
| P-asv6762 | GCAAGCGTTGTTCGGAATCACTGGGCGTAAAGGGCGTGTAGGCGGTTCGGTAAGTGGGATGTGAAATCCCTCGGCTCACCCGAGGACCTGCATCTCATACTGTCGAGCTCGAGTACGGGAGAGGATGGAGGAATTCCTGGTGTAGCGGTGAAATGCGTAGATATCAGGAGGAACACCGGTGGCGAAGGCGTCCATCTGGCCCGATACTGACGCTGAGGCGCGAAAGCTAGGGG |
| P-asv6763 | TCAAGCGTTGTTCGGATTCATTGGGCGTAAAGGGTGCGTAGGCGGCGCGGTAAGTCGGGTGTGAAATCTCGGAGCTTAACTCCGAAACTGCATTCGATACTGCCGTGCTTGAGGACTGGAGAGGAGACTGGAATTTACGGTGTAGCGGTGAAATGCTTAGATATGACACAGAACACCCATTGCGAAGGCAGCTCGCTAAGCCACCATTGACGCTGAGGCACGAAAGCGTGGGT |
| P-asv6764 | GCTAGCGTTAATCGGATTTACTGGGCGTAAAGCGTGCGTAGGCGGTTTGTTAAGTCGATTGTGAAAGCCCCGGGCTCAACCTGGGAATTGCGGTCGATACTGGCAGACTCGAGAACGGTAGAGGGAGGCGGAACTCCAGGTGTAGCGGTGAAATGCGTAGAGATCTGGAGGAATACCGGTGGCGAAGGCGGCCCCCTGGACAAAGACTGACGCTCAGGTGCGAAAGCGTGGGG |
| P-asv6765 | GCGAGCGTTAATCAGAATTACTGGGCGTAAAGCGCACGTAGGTGGCGACGTAAGTCGGGTGTGAAATCCCCGGGCTCAACCTGGGAACTGCATCCGAAACTGGCAGGCTAGAGTCTTGTAGAGGGGGGTAGAATTCCAGGTGTAGCGGTGAAATGCGTAGAGATCTGGAGGAATACCGGTGGCGAAGGCGGCCCCCTGGACAAAGACTGACGCTCAGGTGCGAAAGCGTGGGG |
| P-asv6766 | GCAAGCGTTATCCGGATTCATTGGGCGTAAAGCGCGCGTAGGCGGCCTGTTAGGTCGGGAGTCAAATCCGGGGGCTCAACCCCCGCCCGCTCCCGATACCGGCGGGCTTGAGTCTGGCAGGGGAAGGCGGAATTCCAAGTGTAGCGGTGGAATGCGCAGATATTTGGAAGAACACCGGTGGCGAAGGCGGCCTTCTGGGCCACGACTGACGCTGAGGCGCGAAAGCTAGGGG |
| P-asv6767 | GCAAGCGTTAATCGGAATTACTGGGCGTAAAGCGCACGCAGGCGGTCTGTCAAGTCGGGTGTGAAAGCCCCCGGCTCAACCTGGGAATTGCATTCGAGACTGCGTAGCTAGGGTGCGGAAGAGGGAAGCGGAATTTTCCGGTGTAGCGGTGAAATGCGTAGATATCGGAAGGAACACCAGTGGCGAAAGCGGCTTCCTGGTCCAGCACCGACGTTCAGGCACGAAAGCGTGGGG |
| P-asv6768 | GCAAGCGTTATCCGGAATCACTGGGCGTAAAGGGCGCGTAGGCGGTTTGTTAAGTCCGATGTTAAAGACCGAGGCTCAACCTCGACACGGCGTTGGATACTGGCAAGCTGGACGGTTGGAGAGGAAGGTAGAATTACCAGTGTAGCGGTGGAATGCGTAGATACTGGTAGGAATACCCATTGCGAAGGCAGCCTTCTGGACAACACGTGACGCTGAGGCGCGAAAGTGTGGGG |
| P-asv6769 | TCCCTCTTTTTCCTTCCTTCTTTTTCTTAAAGCGCGCGCAGGCGGTTCCTTAAGTCTGATGTGAAAGCCCCCGGCTCAACCGGGGAGGGTCATTGGAAACTGGGGAACTTGAGTGCAGAAGAGGAGAGCGGAATTCCACGTGTAGCGGTGAAATGCGTAGAGATGTGGAGGAACACCAGTGGCGAAGGCGGCTCTCTGGTCTGTAACTGACGCTGAGGCGCGAAAGCGTGGGG |
| P-asv6770 | ACGAGTAGTGGTCACTTTTATTGAGCCTAAAGCGTTCGTAGCCGGTTTTGTAAATCTTCAGATAAAGCCTGAAGCTTAACTCCAGAAAGTCTGAAGAGACTGCAAGACTTGAGATCGGGTGAGGTTAAACGTACTTTCAGGGTAGGGGTAAAATCCTGTAATCCTGGAAGGACGACCGGTGGCGAAGGCGCTTAACTAGAACGAATCTGACGGTGAGGAACGAAGGCTAGGGT |
| P-asv6771 | GCGAGCGTTGTCCGGAGTTACTGGGCGTAAAGGGCCCGCAGGCGGTCGTGTAAGTTTGGGGTTACAGCTCCCGGCTCAACTGGGAGAGCGCCTCGAAGACTGCAGGACTTGAGGGCCAGAGAGGGACACAGAATTCCAGGTGTAGCGGTGAAATGCGTAGGGATGTGGAGGAACACCAGTGGCGAAGGCGACTCTTTGGTCTGTAACTGACGCTGAGGCGCGAAAGCGTGGGG |
| P-asv6772 | GCAAGCGTTGTTCGGTATCACTGGGCGTAAAGGGCGCGTAGGCGGCGCGACAAGTCAGGTGTGAAAGCCCCGGGCTCAACCCGGGAATTGCACTTGATACTGTCGTGCTTGAGGCCGGGAGAGGATGGCGGAATTCCCAGTGTAGAGGTGAAATTCGTAGATATTGGGAAGAACACCAGTGGCGAAGGCGACTCGCTGGACAAGTATTGACGCTGAGGTGCGAAAGCGTGGGG |
| P-asv6773 | TCTCTCTTTTCTCTTCCTTCCTTTTCTTCAAGGGCGCGTAGGCGGACAGTTAAGTTGGGGGTGAAAGCCCGGGGCTCAACCTCGGAAATGCCTTCAATACTGGCTGTCTTGAGTACGGGAGAGGTGAGTGGAACTCCGAGTGTAGAGGTGAAATTCGTAGATATTCGGAAGAACACCAGTGGCGAAGGCGACTCACTGGCCCGTTACTGACGCTGAGGCGCGAAAGCGTGGGG |
| P-asv6774 | GCAAGCGTTAATCGGAATTACTGGGCGTGAAGCGCACGCAGGCGGCTTTTTAAGTCAGGGGTGAAATCCTGGAGCTCAACTCCAGAACTGCCTCTGATACTGAAAGGCTTGAGTCCGGGAGAGGTGAGTGGAACTGCGAGTGTAGAGGTGAAATTCGTAGATATTCGCAAGAACACCAGTGGCGAAGGCGGCTCACTGGCCCGGTACTGACGCTGAGGTGCGAAAGCGTGGGG |
| P-asv6775 | CCGAGCGTTGTCCGGAGTTACTGGGCGTAAAGCGCGCGCAGGCGGTTTCGCGCGTCGGGTGTGAAAGCCCCCCGCTCAACGGGGGAGGGTCATTCGAAACGGCGGAACTGGAGGCAGGGAGAGGTCGGTGGAATTCCCGGTGTAGTGGTGAAATGCGTAGAGATCGGGAGGAACACCAGTGGCGAAGGCGGCCGACTGGACCTGGCCTGACGCTGAGGCGCGAAGGCGTGGGG |
| P-asv6776 | GCAAGCGTTGTCCGGAATTATTGGGCGTAAAGCGCGCGCAGGCGGCATTGTAAGTCTTGTGTTTAAACGCGGGGCTCAACCCCGTGTCGCATGGGAAACTGCAAGGCTTGAGTGCAGAAGAGGAAAGTGGAATTCCACGTGTAGCGGTGAAGTGCGTAGAGATGTGGAGGAACACCAGTGGCGAAGGCGGCTCACTGGACTGGTATTGACGCTGAAGTGCGAAAGCGTGGGG |
| P-asv6777 | TCTCTCTTTTTTCTTCCTTCCTTTTCTTAAAGCGCACGTAGGCGGTTATTCAAGTCAGAGGTGAAAGCCCGGGGCTCAACCCCGGAACTGCCTTTGAAACTAGATGACTGGAATCTTGGAGAGGCGAGTGGAATTCCGAGTGTAGAGGTGAAATTCGTAGATATTCGGAAGAACACCAGTGGCGAAGGCGACTCGCTGGACAAGTATTGACGCTGAGGTGCGAAAGCGTGGGG |
| P-asv6778 | GCAAGCGTTAATCGGAATTACTGGGCGTAAAGCGCACGCAGGCGGTCTGTCAAGTCGGATGTGAAATCCCCGGGCTCAACCTGGGAACTGCATCCGAAACTGGCAGGCTAGAGTCTTGTAGAGGGGGGTAGAATTCCAGGTGTAGAGGTGAAATTCGTAGATATCCGCAAGAACACCAGTGGCGAAGGCGGCTTACTGGCCCGATACTGACACTGAGGCACGAAAGCGTGGGG |
| P-asv6779 | GCTAGCGTTGTTCGGAATTACTGGGCGTAAAGCGCACGTAGGCGGCTTTGTAAGTCAGAGGTGAAAGCCTGGAGCTCAACTCCAGAACTGCCTTTGAGACTGCATCGCTTGAATCCAGGAGAGGTGAGTGGAATTCCGAGTGTAGCAGTGAAATGCGTAGGTATGCGGAGGAACACCGATGGCGAAGGCAATCCCCTGGACCTGTACTGACGCTCATGCACGAAAGCGTGGGG |
| P-asv6780 | GCGAGCGTTGTCCGGAATCATTGGGCGTAAAGGGGAGCGCAGGCGGGCATGTAAGTCTTTCTTAAAAGTTCGGGGCTCAACCCCGTGATGGGAAAGAAACTATGTGTCTTGAGTACAGGAGAGGAAAGCGGAATTCCCAGTGTAGCGGTGAAATGCGTAGATATTGGGAGGAACACCAGTGGCGAAGGCGGCTTTCTGGACTGCAACTGACGCTGAGGCTCGAAAGCCAGGGG |
| P-asv6781 | GCTAGCGTTGTTCGGAATCACTGGGCGTAAAGCGCACGTAGGCGGATATGTCAGTCAGGGGTGAAATCCTGGAGCTCAACCCCAGAACTGCCTTTGATACTGAAGATCTTGAGTCCGGGAGAGGTGAGTGGAACTGCGAGTGTAGAGGTGAAATTCGTAGATATTCGCAAGAACACCAGTGGCGAAGGCGGCTCACTGGCCCGGTACTGACGCTGAGGCACGAAAGCGTGGGG |
| P-asv6782 | TCTCTCTTTTCTCTTCCTTCCTTTTCTTAAAGGTCGCGTAGGCGGACAGTTAAGTTGGGGGTGAAAGCCCGGGGCTCAACCTCGGAAATGCCTTCAATACTGGCTGTCTTGAGTACGGGAGAGGTGAGTGGAACTCCGAGTGTAGAGGTGAAATTCGTAGATATTCGGAAGAACACCAGTGGCGAAGGCGACTCGCTGGACAAGTATTGACGCTGAGGTGCGAAAGCGTGGGG |
| P-asv6783 | GCAAGCGTTATCCGGAATTATTGGGCGTAAAGCGTCCTGAGGTGGCTTACCCAGTCGACTGTGAAAGCTCAAGGCTTAACCTTGAAACGGCAGTCGAAACTTGTAAGCTTGAGTACGGTAGGGGCAGAGGGAATTCCCGGAGGAGAGGTGAAATTCGTAGATATCGGGAGGAAGGCCAAAGGCGAAAGCACTCTGCTGGGCCGATTACTGACACTGACAGACGAAAGCTAGGGG |
| P-asv6784 | GCGAGCGTTATCCGGAATTATTGGGTTTAAAGGGTGCGCAGGTTGCAAGGGAAGTCAGGGGTGAAAAGCTGTAGCTCAACTATGGTCTTGCCTTTGAAACTCTCTAGCTAGAGTGTACTGGAGGTACGTGGAACGTGTGGTGTAGCGGTGAAATGCATAGATATCACACAGAACTCCGATTGCGCAGGCAGCGTACTACATTACAACTGACACTGAAGCACGAAAGCGTGGGT |
| P-asv6785 | GCGAGCGTTGTCCGGAATTATTGGGCGTAAAGGGCTCGTAGGCGGTTTGTCGCGTCGGGAGTGAAAACGCCGTGCTTAACACGGCGCTTGCTTTCGATACGGGCAGACTTGAGGTATTCAGGGGAGAACGGAATTCCTGGTGTAGCGGTGAAATGCGCAGATATCAGGAGGAACACCGGTGGCGAAGGCGGTTCTCTGGGAATGACCTGACGCTGAGGAGCGAAAGTGTGGGG |
| P-asv6786 | GCAAGCGTTGTCCGGAATTATTGGGCGTAAAGGGCACGCAGGCGGTCTTTTAAGTCTGATGTGAAATCCCGCGGCTTAACCGCGGGTGGTCATTGGAAACTGGAGGACTTGAGTACAGAAGAGGAGAGTGGAATTCCACGTGTAGCGGTGAAATGCGTAGATATGTGGAGGAACACCAGTGGCGAAGGCGACTCTCTGGTCTGTCACTGACGCTGAGGTGCGAAAGCGTGGGT |
| P-asv6787 | GCGAGCGTTGTCCGGATTTATTGGGCGTAAAGAGCTCGTAGGCGGCTTGTCGCGTCGACCGTGAAAACTTGGGGCTCAACTCCAAGCCTGCGGTCGATACGGGCAGGCTAGAGTTCGGTAGGGGAGACTGGAATTCCTGGTGTAGCGGTGAAATGCGCAGATATCAGGAGGAACACCGGTGGCGAAGGCGGGTCTCTGGGCCGATACTGACGCTGAGGAGCGAAAGCGTGGGG |
| P-asv6788 | GCTAGCGTTGTTCGGAATTACTAGGCGTAAAGGGCGTGTAGGCGGACGATCGTGTCAGGGGTGAAAGGCCAGGGCTCAACCTTGGAACGGCCCTTGAAACTGGTTGTCTAGAGTCCGGGAGAGGATGGCGGAATTCCCAGTGTAGAGGTGAAATTCGTAGATATTGGGAAGAACACCGATGGCGAAGGCGGCCATCTGGACCGGCACTGACGCTGAGGCGCGAAAGCGTGGGG |
| P-asv6789 | GCAAGCGTTGTTCGGAATTATTGGGCGTAAAGGGCTCGCAGGTGGTTTGTTAAGTTGGTGGTTTAATCTCTGGGCTCGACCCAGAGTCAGCCTCCAAAACTGGCGAACTTGAGTACGATAGGGGATAGCGGAATTCTCGGTGTAGCGGTGGAATGCGTAGATATCGAGAGGAACACCAATGGCGAAGGCAGCTATCTGGATCGTAACTGACACTCATGAGCGAAAGTGCGGGG |
| P-asv6790 | GCAAGCGTTATCCGGATTTACTGGGTGTAAAGGGAGCGTAGACGGCTGTGTAAGTCTGAAGTGAAAGCCCGGGGCTCAACCCCGGGACTGCTTTGGAAACTATGCAGCTAGAGTGTCGGAGAGGTAAGTGGAATTCCCAGTGTAGCGGTGAAATGCGTAGATATTGGGAGGAACACCAGTGGCGAAGGCGGCTTACTGGACGATGACTGACGTTGAGGCTCGAAAGCGTGGGG |

| | |
|---|---|
| P-asv6791 | GCGAGCGTTAATCGGATTTACTGGGCGTAAAGCGTGCGTAGGCGGCTTTTTAAGTCGGATGTGAAATCCCTGAGCTTAACTTAGGAATTGCATTCGATACTGGGAAGCTAGAGTATGGGAGAGGATGGTAGAATTCCAGGTGTAGCAGTGAAATGCGTAGATATAGGAAGGAACACCAGTGGCGAAGGCGACCACCTGGACTGATACTGACACTGAGGTGCGAAAGCGTGGGG |
| P-asv6792 | GCGAGCGTTAATCGGAATTACTGGGCGTAAAGGGTGCGTAGGTGGTTAATTAAGTTATCTGTGAAATCCCTGGGCTCAACCTGGGCAGGTCAGATAAAACTGTTTAACTCGAGTATGGGAGAGGGTAGTGGAATTTCCGGTGTAGCGGTGAAATGCGTAGAGATCGGAAGGAACACCAGTGGCGAAGGCGGCTACCTGGCCTAATACTGACACTGAGGCACGAAAGCGTGGGG |
| P-asv6793 | GCAAGCGTTGTCCGGAATTATTGGGCGTAAAGGGCTTGTAGGCGGCGTGTCGCGTCTGTCGTGAAAACCAATGGCTTAACTGTTGGCTTGCGGTGGGTACGGGCATGCTTGAGTGCGGTAGGGGAGACTGGAATTCCTGGTGTAGCGGTGGAATGCGCAGATATCAGGAGGAACACCGGTGGCGAAGACGGGTCTCTGGGCCGTTACTGACGCTGAGGAGCGAAAGCGTGGGG |
| P-asv6794 | GCAAGCGTTGTTCGGATTGACTGGGCGTAAAGGGCGTGTAGGCGGTTTGTCAAGTCGGGTGTGAAAGCCTGGGGCTCAGCCCCAGAAGTGCATCCGAAACTAGCAAGCTAGAGCGCGATAGAGGAGGGTGGAATTCCTGGTGTAGCGGTGAAATGCGTAGATATCAGGAGGAACACCGGCGGCGAAGGCGGCCCTCTGGATCGTTGCTGACGCTGAGACGCGAAAGCGTGGGG |
| P-asv6795 | TCGAGTGATGTCCCATATTATTGGGCCTAAAGCGTCCGTAGCTGGCCAGGCAAGTCCGTCGGGAAATCCACCCGCTCAACGGGTGGGCGGCCGACGGAAACTGCGAGGCTTGGGACCGAGAGGCTTGAAGGGTACGTCCGGGGTAGGAGTGAAATCCCGTAATCCTGGACGGACCACCGATGGGGAAACCACTTCAAGAGAACGGATCCGACAGTGAGGGACGAAAGCTAGGGT |
| P-asv6796 | GCAAACGTTGTTCGGAATTACTGGGCGTAAAGCGCGTGCAGGCTGCTCTGCAAGTCGGGTGTGTAAGCCCTGGGCTCAACCTAGGAAGTGCACTCGAAACTGCAGAGCTTGAGTCCTGGAGAGGATCGCGGAATTCTCGGTGTAGAGGTGAAATTCGTAGATATCGAGAGGAACACCGGTGGCGAAGGCGGCGATCTGGACAGTGACTGACGCTGAGACGCGAAAGCGTGGGG |
| P-asv6797 | GCTAGCGTTGCTCGGAATCACTGGGCGTAAGGGGCGCGTAGGCGGCGTTTTAAGTCGGGGGTGAAAGCCTGTGGCTCAACCACAGAATGGCCTTTGATACTGAAGATCTTGAGTTCGGGAGAGGTGAGTGGAACTGCGAGTGTAGAGGTGAAATTCGTAGATATTCGCAAGAACACCAGTGGCGAAGGCGGCTCACTGGCCCGATACTGACGCTGAGGCACGAAAGCGTGGGG |
| P-asv6798 | GCTAGCGTTGTTCGGAATTACTGGGCGTAAAGCGCACGTAGGCGGACATTTAAGTCAGGGGTGAAATCCCAGAGCTCAACTCTGGAACTGCCTTTGATACTGGGTGTCTTGAGTGTGGAAGAGGTAAGTGGAATTGCGAGTGTAGAGGTGAAATTCGTAGATATTCGCAGGAACACCAGTGGCGAAGGCGGCTTACTGGTCCACAACTGACGCTGAGGTGCGAAAGCGTGGGG |
| P-asv6799 | GCGAGCGTTGTCCGGAATTATTGGGCGTAAAGAGCGTGTAGGCGGTCTGGTAAGTCTGTCGTGAAAGGCTGGGGCTCAACCCCGGACGTGCGATGGATACTGCCAGACTAGAGGATGGTAGAGGCGAGTGGAATTCCCGGTGTAGCGGTGAAATGCGCAGATATCGGGAGGAACACCAGTAGCGAAGGCGGCTCGCTGGGCCATACCTGACGCTGAGGCGCGAAAGCGTGGGG |
| P-asv6800 | GCAAGCGTTGTCCGGAATTATTGGGCGTAAAGAGCTCGTAGGCGGTCTGTCGCGTCTGCTGTGAAAACTCAGGGCTTAACCCTGAGCTTGCAGTGGGTACGGGCAGACTAGAGTGCGGTAGGGGAGACTGGAATTCCTGGTGTAGCGGTGGAATGCGCAGATATCAGGAGGAACACCGATGGCGAAGGCAGGTCTCTGGGCCGTTACTGACGCTGAGGAGCGAAAGCATGGGG |
| P-asv6801 | GCGAACGTTGCTCGGAATGACTGGGCGTAAAGCGCGTGTAGGCGGCTCCGCAAGTCCGATGTGAAAGCCCAGGGCTCAACCCTGGAAGTGCATTGGAAACTACGAAGCTGGAGTCCTGGAGAGGAGAGCGGAATTCTCGGTGTAGAGGTGAAATTCGTAGATATCGGGAGGAACACCAGTGGCGAAGGCGGCTCTCTGGACAGGTACTGACGCTGAGACGCGAAAGCGTGGGT |
| P-asv6802 | GCTAGCGTTGCTCGGAATTACTGGGCGTAAAGCGCGTGCAGGCGGTTTGGTAAGTCAAATGTGAAATCCCAGGGCTCAACCCTGGAACTGCATTCGATACTGCCAGACTAGAGTGTAAGAGAGGATTGTGGAATTCCGGGTGTAGCGGTGAAATGCGTAGATATCCGGAGGAACATCAGTGGCGAAGGCGACAATCTGGCTTAACACTGACGCTCAGATGCGAAAGCGTGGGG |
| P-asv6803 | ACGAACGTTATCCGGAGTTACTGGGCGTAAAGGGCGTGTAGGCGGTATGCTAAGTCTGGCGTGAAAGCTCTCGGCTAAACCGGGAGAGGTCGTTGGATACTGGCAAACTAGAGGATGGTAGAGGCGTGTGGAATTCCGGGTGTAGTGGTGAAATGCGTAGAGATCCGGAGGAACACCAGTGGCGAAGGCGACACGCTGGGCCATACCTGACGCTAAAGCGCGACAGCATGGGG |
| P-asv6804 | GCAAGCGCTGTCCGGATTTATTGGGCGTAAAGGGAGCGCAGGTGGTTTCTTAAGTCTGATGTGAAAGCCCACGGCTTAACCGTGGAGGGTCATTGGAAACTGGGAAACTTGAGTACAGAAGAGGAATGTGGAACTCCATGTGTAGCGGTGGAATGCGTAGATATATGGAAGAACACCAGTGGCGAAGGCGACATTCTGGTCTGTTACTGACACTGAGGCTCGAAAGCGTGGGG |
| P-asv6805 | GCGAGCGTTAATCGGAATTACTGGGCGTAAAGCGCGCGTAGGCGGCTTTTCAAGTCGGATGTGAAATCCCCGGGCTTAACCTGGGAACTGCATTAGAGACTGTCTAGCTAGAGTATGGGAGAGGGAAGTGGAATTTCAGGTGTAGCGGTGAAATGCGTAGATATCTGAAGGAACATCAGTGGCGAAAGCGACTTCCTGGACCAATACTGACGCTCATGTGCGAAAGCGTGGGG |
| P-asv6806 | GCAAGCGTTATCCGGATTTACTGGGTTTAAAGGGTGTGTAGGCGGACTTTTAAGTCAGTGGTGAAATCTCCGGGCTCAACCCGGAAACTGCCATTGATACTATTAGTCTTGAATGTTGTGGAGGTGGGCGGAACGGGTCATGTAGCGGTGAAATGCATAGATATGACCCGGAACACCGATTGCGAAGGCAGCCCGCTACGCAACGATTGACGCTGAGGCACGAAAGCGTGGGG |
| P-asv6807 | CCGAGCGTTGTTCGGATTCATTGGGCGTAATGGGTGCGTAGGCGGCGGGGTAAGTTTGATGTGAAATCCCGGGGCTCAACCCTGGAATGGCATTGAATACTGCCTTGCTAGAGTACCGGAGAGGAAACTGGAATTTACGGTGTAGCAGTGAAATGCGTAGAGATCGTAAGGAAGGCCAGTGGCGAAGGCGAGTTTCTGGACGGTAACTGACGCTGAGGCACGAAGGCCAGGGG |
| P-asv6808 | GCAAGCGTTATCCGGAATCATTGGGCGTAAAGCGTCTGTAGGTGGCTTGGTAAGTCGACTGTTAAATCGTAGAGCTCAACTCTATATCGGCAGTCGAAACTACTTCGCTTGAGTGTGGTAGGGGCAGAGGGAATTCCCGGTGTAGCGGTGAAATGCGTAGATATCGGGAAGAACACCAACGGCGAAAGCACTCTGCTGGGCCAACACTGACACTGAGAGACGAAAGCTAGGGT |
| P-asv6809 | CCAAGCGTTATCCAGAATTATTGGGCGTAAAGGGTGCGTAGGCGGCCTTATAAGTCTTGTTTGAAAGCCCACGGCTCAACCGTGGAGGGTGACAAGAAACTGTAAGGCTTGAATGAAGCAGAGGTAAGTAGAATTCTGTGTGGAGCGGTGGAATGCGTAGATATACGGAAGAATACCAGTGGCGAAGGCGGCTTACTGGGCTTTAATTGACGCTGAGGCACGAAAGCGTGGGT |
| P-asv6810 | GCAAGCGTTGTCCGGAATTATTGGGCGTAAAGCGCGCGCAGGCGGCATTGTAAGTCTTGTGTTTAAACGCGGGGCTCAACCCCGTGTCGCATGGGAAACTGCAAGGCTTGAGTGCAGAAGAGGAAAGTGGAATTCCACGTGTAGCGGTGAAATGCGTAGATATAGGAAGGAACACCAGTGGTGAAGGCGACCACCTGGACTGATACTGACACTGAGGTGCGAAAGCGTGGGG |
| P-asv6811 | CCAAGCGTTATCCGGATTCATTGGGTTTAAAGGGTGCGTAGGCGGCCTGTTAAGTCAGTGGTGAAATTTTGTGGCTCAACCATAAAACTGCCATTGAAACTGACAGGCTTGAATACAGTTGAAGTAGGCGGAATAGGTAGTGTAGCGGTGAAATGCTTAGATATTACCTAGAACACCGATTGCGAAGGCAGCTTACTAAAGTGTCATTGACGCTGATGCACGAAAGCGTGGGT |
| P-asv6812 | GCAAGCGTTATCCGGATTCACTGGGTTTAAAGGGTGCGTAGGCGGGCTTGTAAGTCCGTGGTGAAATCTCTGAGCTTAACTCGGAAACTGCCATGGATACTATGAGTCTTGAATACTCGGGAGGTGAGCGGAATATGTCATGTAGCGGTGAAATGCTTAGATATGACATAGAACACCAATTGCGAAGGCAGCTTGCTACAGAGTTATTGACGCTGAGGCACGAAAGCGTGGGG |
| P-asv6813 | GCAAGCGTTGTTCGGAATTATTGGGTGTAAAGCGTGCGTAGGCGGACTGTTAAGTCTGGTGTGAAATCCCTCGGCTCAACCGAGGACGTGCATTGGATACTGGCAGTCTTGAGTCCTGGAGAGGAGAGCGGAATTCCTAGTGTAGAGGTGAAATTCGTAGATATTAGGAGGAACACCAGCGGCGAAAGCGGCTCTCTGGACAGTGACTGACGCTGAGGCACGAAAGCATAGGT |
| P-asv6814 | CCAGGCGTTATCCGGATTTATTGGGTTTAAAGGGAGTGTAGGCGGTTTGTTAAGCGTGTTGTGAAATTTAGATGCTCAACATTTAACTTGCAGCGCGAACTGGCGAACTTGAGTGCACACAACGTATGCGGAATTCATGGTGTAGCGGTGAAATGCTTAGATATCATGAAGAACTCCGATTGCGAAGGCAGCATACGGGAGTGTAACTGACGCTTAAGCTCGAAGGTGCGGGT |
| P-asv6815 | GCAAGCGTTAATCGGAATTACTGGGCGTAAAGCGCGCGTAGGTGGTTCGTTAAGTTGGATGTGATAGCCCCGGGCTCAACCTGGGAACTGCATCCAAAACTGGCGAGCTAGAGTACGGTAGAGGGTGGTGGAATTTCCTGTGTAGCGGTGAAATGCGTAGATATAGGAAGGAACACCAGTGGCGAAGGCGACCACCTGGACTGATACTGACACTGAGGTGCGAAAGCGTGGGG |
| P-asv6816 | GCAAGCGTTAATCGGAATCACCGGGCGTAAAGGGTGCGTAGGCGGTACGTTAGGTCTGTCGTGAAATCCCCAAGCTTAACTTGGGACGTGCATTTGGAACTGGGTGACTAGAGTGTGTCAGAGGGAGGTAGAATTCCACATGTAGCGGTGGAATGCGTAGAGATGTGGAGGAATACCGATGGCGAAGGCAGCCTCCTGGGATAACACTGACGTTGAGGCTCGAAAGCGTGGGG |
| P-asv6817 | GCAAGCGTTACTCGGAATCACTGGGCGTAAAGGACGCGTAGGCGGATTATCAAGTCTCTTGTGAAATCTAACGGCTTAACCGTTAAACTGCTTGGGAAACTGATAATCTAGAGTAAGGGAGAGGCAGATGGAATTCTTGGTGTAGGGGTAAAATCCGTAGATATCAAGAAGAATACCTATTGCGAAAGCGATCTGCTGGAACTTAACTGACGCTAATGCGTGAAAGCGTGGGG |
| P-asv6818 | CCAAGCGTTATCCGGATTCATTGGGTTTCAAGGGTGCGTAGGCGGCCTGTTAAGTCAGTGGTGAAATTTTGTGGCTCAACCATAAAACTGCCATTGAAACTGACAGGCTTGAATACAGTTGAAGTAGGCGGAATAGGTAGTGTAGCGGTGAAATGCTTAGATATTACCTAGAACACCGATTGCGAAGGCAGCTTACTAAAGTGTCATTGACGCTGATGCACGAAAGCGTGGGT |
| P-asv6819 | GCGAGCGTTAATCGGAATTACTGGGCGTAAAGGGTACGTGGGCGGCTTGATAAGTCAGATGTGAAATCCCCGGGCTTAACCTGGGAACTGCATTTGAAACTGTTTAGCTAGAGTGAGTGAGAGGTTAGTGGAATTCAAGGCGTAGCGGTGAAATGCGTAGAGGTCTTGAGGAACATCAGTGGCGAAGGCGACTGACTGGCACTACACTGACGCTGAGGTACGAAAGCGTGGGG |
| P-asv6820 | CCGAGCGTTAGGCGGAATCACTGGGCTTAAAGCGCGTGTAGGTGGATGCCTAAGTGTCTTGTGAAATCCCACGGCTTAACCGTGGAACTGCTTGACATACTGGGTGTCTTGAGCCACTCAGGGGCTACCGGAACAAATGGTGGAGCGGTGAAATGCGTAGATATCATTTGGAACGCCAATGGTGAAAACAGGTAGCTGGGGGTGTGCTGACACTGAGACACGAAAGCCAGGGG |
| P-asv6821 | ACGAACGTTAATCGGAATTACTGGGCGTAAAGCGCGCGTAGGCGGTTTGATAAGTGGGATGTGAAAGCCCCGGGCTCAACCTGGGAACTGCATTCCAAACTGTCAGACTAGAGTGTGGTAGAGGGAGGTAGAATTTCCTGTGTAGCGGTGAAATGCGTAGATATAGGAAGGAATACCGATGGCGAAGGCAGCCTCCTGGACCAACACTGACGCTGAGGTGCGAAAGCGTGGGG |
| P-asv6822 | TCGAGTGGTGATCACTTTTATTGAGTCTAAAGCGTCCGTAACCGGTCTTATAAATCTTCAGATAAATTCTTCCGCTTAATGGAAGAACTTCTGAAGAGACTGTATGACTTGGGACCGGGTGAGGTTGAATGTACTTTCAGGGTAGGGGTAAAATCCTGTAATCCTGAAAGGACGACCGGTGGCGAAAGCGTTCAACTAGAACGGATCCGACGGTGAGGGACGAAGGCTAGGGG |
| P-asv6823 | GCAAGCGTTGTCCGGAATCACTGGGCGTAAAGGGTACGCAGGCGGATATACAAGTCAAATGTGAAATGTACCGGCTCAACCGGTAAGCTGCATTTGAAACTGTATATCTTGAGTCTAGGAGAGGAGAGCGGAATTCCTGGTGTAGCGGTGGAATGCGTAGATATCAGGAAGAACATCAGTGGCGAAGGCGGCTCTCTGGCCTAAGACTGACGCTCAGGTACGAAAGCTGGGGG |
| P-asv6824 | GCAAGCGTTATCCGGATTCACTGGGTTTAAAGGGTGCGTAGGCGGATTGGTAAGTCAGTGGTGAAAGCTCTCAGCTCAACTGAGAAATTGCCATTGATACTGCTGATCTTGAATATAGACGAAGTGGGCGGAATGTGGCGTGTAGCGGTGAAATGCTTAGATATGCCACAGAACACCTATTGCGAAGGCAGCTCGCTAGACTATTATTGACGCTGAGGCACGAAAGCGTGGGG |
| P-asv6825 | GCAAGCGTTGCTCGGAATTACTGGGCGTAAAGGGCGTGTAGGCGGTTTCGTATGTCTGGCGTGAAAGCCCCGGGCTCACCTCGGGAAGTGCGTCGGAAACTGCGAGACTAGAGTCCCGGAGAGGGAGGTGGAATTCCAGGTGTAGCGGTGAAATGCGTAGATATCTGGAAGAACACCGGTGGCGAAAGCGGCCTCCTGGACGGTGACTGACGCTGAGACGCGAAAGCGTGGGG |
| P-asv6826 | TCAAGTGATGTCCCATATTATTGGGCCTAAAGCGTCCGTAGCTGGCCAGGCAAGTCTATCGGGAAATCCACCTGCCCAACGGGCAGGCGTCCAGCGGAAACTGTTCGGCTTGGGACCGGAAGACCTGAGGGGTACGTCCGGGGTAGGAGTGAAATCCCGTAATCCTGGACGGACCACCGGTGGCGAAAGCGCCTCAGGAGGACGGATCCGACAGTGAGGAACGAAAGCTAGGGT |
| P-asv6827 | GCAAGCGTTATCCGGAATTATTGGGCGTAAAGCGCGCGTAGGCGGTTTTTTAAGTCTGATGTGAAAGCCCACGGCTCAACCGTGGAGGGTCATTGGAAACTGGAAAACTTGAGTGCAGAAGAGGAAAGTGGAATTCCATGTGTAGCGGTGAAATTCGTAGATATTCGGAAGAACACCAGTGGCGAAGGCGGCTCACTGGACTGGTATAGACGCTGAAGTGCGAAAGCGTGGGG |
| P-asv6828 | GCGAGCGTTATCCGGAATTACTGGGTTTAAAGGGTGCGTAGGCGGCGTAATAAGTCAGAGGTGAAAGGCACGAGCTTAACTGGTGAATTGCCTTTGAAACTGTTATGCTAGAATTGAGTTGAGGTTAGCGGAATGTGACATGTAGCGGTGAAATGCATAGATATGTCATAGAACACCAATTGCGTAGGCAGCTAGCTAGGCTTTAATTGACGCTGAGGCACGAAAGCGTGGGG |
| P-asv6829 | GCAAGCGTTAATCGGATTTACTGGGCGTAAAGCGTACGCAGGCGGCTTGGTAAGACAGATGTGAAATCCCCGGGCTTAACCTGGGAACTGCATTTGTGACTGCCTGGCTCGAGTCTGTCAGAGGGGGGTGGAATTCCACATGTAGCAGTGAAATGCGTAGAGATGTGGAGGAACACCGATGGCGAAGGCAGCCCCCTGGGATGAGACTGACGCTCATGTACGAAAGCGTGGGG |

| | |
|---|---|
| P-asv6830 | GCTAGCGTTACTCGGAATCACTGGGCGTAAAGGGCGCGTAGGCGGCTGATTTAGTCGAGGGTGAAAGCCCGTGGCTCAACCACGGAATGGCCTTCGATACTGGTTGGCTTGAGACCGGAAGAGGACAGCGGAACTGCGAGTGTAGAGGTGAAATTCGTAGATATTCGCAAGAACACCAGTGGCGAAGGCGGCTGTCTGGTCCGGTTCTGACGCTGAGGCGCGAAAGCGTGGGG |
| P-asv6831 | GCAAGCGTTACTCGGATTTACTGGGCGTAAAGCGCGCGCAGGCGGATCGGCAAGTCAGGTGTGAAATCTCGGGGCCCAACTCCGAAACTGCGCCTGAAACTATCGATCTTGAGTCTTGGCGGGGGAGGTGGAATATCGCATGTAGGGGTAAAATCCGTAGAGATGCGATGGAACACCAGAGGCGAAGGCGACCTCCTGGACAAGTACTGACGCTGAGGCGCGAAAGCGTGGGG |
| P-asv6832 | GCTAGCGTTGTTCGGAATCACTGGGCGTAAAGGGCGCGTAGGCGGCCTTTGGTGTCGGATGTGGAAGCCCGGGGCTCAACTCCGGAGGTGCATCCGAAACTCAGGGGCTCGTGGCCGGAAGAGGAGGGTGGAATACCCAGTGTAGAGGTGAAATTCGTAGATATTGGGTGGAACACCGGTGGCGAAAGCGGCTCTCTGGTCCGGTCCTGACGCTGAGGCGCGAAAGCGTGGGG |
| P-asv6833 | GCTAGCGTTGCTCGGAATCACTGGGCGTAAAGCGCACGTAGGCGGATCGTTAAGTCAGGGGTGAAAGCCTGGAGCTCAACTCCAGAACTGCCCTTGATACTGGCGATCTCGAGTTCGAGAGAGGTTGGTGGAACTCCGAGTGTAGAGGTGAAATTCGTAAATATTCGGAAGAACACCAGTGGCGAAGGCGGCCAACTGGCTCGATACTGACGCTGAGGTGCGAAAGCGTGGGG |
| P-asv6834 | GCAAGCGTTGTCCGGAATCATTGGGTGTAAAGGGTGCGTAGGCGGAGGCCTAAGTCGGAGGTGAAATCCTGCCGCTCAACGGTAGAATGGCCTTCGATACTGGGTCTCTTGAGTTCGGAAGAGGTTGGTGGAATTCGTGGTGTAGCGGTGAAATGCATAGAGATCACGAAGAACACCGGTGGCGAAGGCGGCCAACTGGTCCGACACTGACGCTGAGGCACGAGAGCGTGGGG |
| P-asv6835 | ACAAGCGTTGTCCGGAATCACTGGGCGTAAAGGGCGCGCAGGCGGTGATTTAAGCAGGATGTGAAAGGCTACGGCTCAACCGTAGACATGCACCCTGAACTGGGTCACTTGAGTATTGGAGAGGCAGGCAGAATTCCTGGTGTAGCGGTGAAATGCGTAGATATCAGGAGGAATACCGGAGGCGAAGGCGGCCTGCTGGACAAATACTGACGCTGAGGCGCGAAAGCGTGGGG |
| P-asv6836 | TCTCTCTTTTCTCTTCCTTCTTTTTCTTAAAGCGAGTGCAGACGGTTGTTTAAGCCAGATGTGAAATCCCCGAGCTTAACTTGGGACGTGAATTTGGAACTGGATAACTAGAGTGTGTCAGAGGGAGGTAGAATTCCACATGTAGCGGTGGAATGCGTAGAGATGTGGAGGAATACCGATGGCGAAGGCAGCCTCCTGGGATAACACTGACGTTGAGGCTCGAAAGCGTGGGG |
| P-asv6837 | GCGAGTGTTATCCGGAATTATTGGGCGTAAAGCGTCTGTAGGTTGAATTCAGAGTCTTTTGTAAAATCTTTAAGCTCAACTTAAAAACCGCAAAAGTTACTAGTTTTCTTGAGTACGGTAGAGGTAAAGGGAATTCCCAGTGGAGCGGTGAAATGCGTAGAGATTGGAAGGAACACCAAAAGCGAAGGCACTTTACTGGGCCATTACTGACACTGAGAGACGAAAGCTAAGGG |
| P-asv6838 | GCAAGCGTTATCCGGAATGATTGGGCGTAAAGCGTCCGCAGGTGGCAGTTCAAGTCTGCTGTCAAAGACCGGGGCTTAACTTCGGAAAGGCAGTGGAAACTGAACAGCTAGAGTATGGTAGGGGCAGAGGGAATTCCTGGTGTAGCGGTGAAATGCGTAGAGATCAGGAAGAACATCGGTGGCGAAGGCGCTCTGCTGGACCATAACTGACACTCAGGGACGAAAGCTAGGGG |
| P-asv6839 | GCAAGCGTTATCCGGAATCACTGGGTTTAAAGGGTGCGTAGGCGGGCCTGTAAGTCAGGGGTGAAATCCTACAGCTTAACTGTAGAATTGCCTTTGGTACTGCAGGTCTTGAATTAGGTTGAAGTCAGCGGAATGTGGCATGTAGCGGTGAAATGCATAGATATGCCATAGAACACCGATTGCGAAGGCAGCTGGCTGGGCCTTAATTGACGCTGAGGCACGAAAGCGTGGGG |
| P-asv6840 | GCAAGCGTTAATCGGAATTACTGGGCGTAAAGGGCGCGTAGGCGGTTCGATAAGTCGGGTGTGAAAGCCCCGGGCTCAACCCGGGAATTGCACTCGATACTGTCGAGCTGGAGTGTGGTAGAGGGAAGCGGAATTCCGTGTGTAGCGGTGAAATGCGTAGATATCCGGAGGAACACCAGTGGCGAAGGCGGCTTCCTGGACCAACACTGACGCTGAGGCGCGAAAGCGTGGGG |
| P-asv6841 | GCAAGCGTTGTCCGGATTTATTGGGCGTAAAGCGAGCGCAGGCGGAAGAATAAGTCTGATGTGAAAGCCCTCGGCTTAACCGAGGAACTGCATCGGAAACTGTTTTTCTTGAGTGCAGAAGAGGAGAGTGGAACTCCATGTGTGGCGGTGGAATGCGTAGATATATGGAAGAACACCAGTGGCGAAGGCGGCTCTCTGGTCTGCAACTGACGCTGAGGCTCGAAAGCATGGGT |
| P-asv6842 | GCAAGCGTTAATCAGAATTATTGGGCGTAAAGCGAGTGCAGACGGTTACTTAAGCCAGATGTGAAATCCCCGGGCTCAACCTGGGAACTGCATTGGAGACTGCAAGGCTAGAGTGTGTCAGAGGGGGGTAGAATTCCACGTGTAGCAGTGAAATGCGTAGAGATGTGGAGGAATACCGATGGCGAAGGCAGCCACCTGGGATAACACTGACGCTCATGCACGAAAGCGTGGGG |
| P-asv6843 | GCAAGCGTTAATCAGAATTATTGGGCGTAAAGCGAGTGCAGACGGTTACTTAAGCCAGATGTGAAATCCCCGGGCTCAACCTGGGAACTGCATTGGAGACTGCAAGGCTAGAGTGTGTCAGAGGGGGGTAGAATTCCACGTGTAGCAGTGAAATGCGTAGAGATGTGGAGGAATACCGATGGCGAAGGCAGCCCCCTGGGATAACACTGACGCTCATGCACGAAAGCGTGGGG |
| P-asv6844 | GCGAGCGTTAATCGGAATTACTGGGCGTAAAGCGCGCGTAGGTGGTTCGTTAAGTTGGATGTGAAATCCCCGGGCTCAACCTGGGAACTGCCTTTGTGACTGCATAGCTAGAGTACGGTAGAGGGGGATGGAATTCCGCGTGTAGCAGTGAAATGCGTAGATATGCGGAGGAACACCGATGGCGAAGGCAATCCCCTGGACCTGTACTGACGCTCATGCACGAAAGCGTGGGG |
| P-asv6845 | GCTAGCGTTATTCGGAATTACTGGGCGTAAAGCGCGCGTAGGCGGCTATGTAAGTCGGATGTGAAAGCCCTGGGCTTAACCTGGGAATTGCATTCGATACTGCGTGGCTGGAGTATGGTAGAGGGTAGTGGAATTCCTGGTGTAGCGGTGAAATGCGTAGATATCAGGAGGAACATCAGTGGCGAAGGCGGCTACCTGGACCAATACTGACGCTGAGGTGCGAAAGCGTGGGG |
| P-asv6846 | GCAAGCGTTATCCGGAATTATTGGGCGTAAAGCGTCCGCAGGCGGCCTTTCAAGTCTGCTGTTAAAGCGTGGAGCTTAACTCCATTAAGGCAGTGGAAACTGATCGGCTTGAGTATGGTAGGGGCAGAGGGAATTCCCGGTGTAGCGGTGAAATGCGTAGATATCGGGAAGAACACCAGTGGCGAAGGCGCTCTGCTGGGCCATTACTGACGCTCATGGACGAAAGCCAGGGG |
| P-asv6847 | GCAAGCGTTGTCCGGATTCATTGGGCGTAAAGAGTTCGTAGGCGGTAGCCTAAGTCTGATGTTAAATCCCGAAGCTTAACTTCGGCATGGCATTGGAAACTGGGTTGCTAGAGTGTGATAGAGGTAACTGGAATTCTCGGTGTAGCGGTGAAATGCGTAGATATCGGGAGGAACACCAGTGGCGAAAGCGAGTTACTGGATCATTACTGACGCTGAGGAACGAAAGCCAGGGT |
| P-asv6848 | GCGAACGTTATTCGGAATCACTGGGCTTAAAGGGTGCGTAGGCGGCTTATCAAGTCAGGTGTGAAAGGCCCCGGCTCAACCGGGGCACTGCGCTTGAAACTGATTGGCTTGAGTGAGCTAGGGGTATGCGGAACTTCCAGTGGAGCGGTGAAATGTGTTGAGATTGGAAGGAACACCGGTGGCGTAAGCGGCATACTGGGGCTTAACTGACGCTGAGGCACGAAAGCCAGGGG |
| P-asv6849 | GCGAGCGTTGCTCGGAATCATTGGGCTTAAAGCGGATGTAGGCGGCTTGGCAAGTGTGACGTGAAAGCTCCCGGCTCAACCGGGGAATTGCGTTGCAAACTGCCAGGCTTGAGGCCGGTAGGGGTGCCTGGAACTCTAGGTGGAGCGGTGAAATGCGTAGATATCTAGAGGAACGCCAGAGGCGAAAGCGAGGCACTGGGCCGGTTCTGACGCTGAGATCCGAAAGCGTGGGG |
| P-asv6850 | GCAAGCGTTATTCGTCGTAACTGGGCGTAAAGAGTTTCTAGACGGTAAAATAAGTTATAAGTTTCAAATTTTAAAGCTCAACTTTAAATTAGCTTTTAAAACTGTTTTACTAGAGTTTTGTACAGAAAAGTAGAATTTTATGTGTAAGGGTAAAATCTAAATATACGTAAAGGAATTCCAATAGTGAAGACAACTTTTTAGTAAAAACTGACGTTGAGAAACGAAAGTATGGGG |
| P-asv6851 | GCAAGCGTTATCCGGATTCACTGGGTTTAAAGGGTGCGTAGGCGGGTATTTAAGTCAGTGGTGAAATCCTAGAGCTTAACTCTAGAACTGCCATTGATACTATTTATCTTGAATATTGTGGAGGTAAGCGGAATATGTCATGTAGCGGTGAAATGCTTAGATATCGCGAGGAACACCCGTGGCGAAAGCGGCCTACTGGATAGTCCCTGACGCTGAGGCACGATAGCGTGGGT |
| P-asv6852 | GCTAGCGTTGTTCGGAATTACTGGGCGTAAAGGGCGCGTAGGCGGTTCGGTAAGTCAGATGTGAAAGCCCTGGGCTCAACCCGGGAACTGCGTTTGATACTGCCGTACTTGAATTCGGTAGAGGTGAGTGGAATTCCCAGTGTAGAGGTGAAATTCGTAGATATTGGGAAGAACACTAGTGGCGAAGGCGGCTCACCGGACCGACATTGACGCTGAGGCGCGAAAGCGTGGGG |
| P-asv6853 | GCAAGCGTTGTCCGGATTTACTGGGTTTAAAGGGTGCGTAGGCGGGCCGATAAGTCAGTGGTGAAAGCCCTCTGCTTAACGGAGGAACTGCCATTGATACTGTTGGTCTTGAGATCAGTAGAGGTATGTGGAATTGATGGTGTAGCGGTGAAATGCATAGATACCATCAAGAACACCGAAAGCGAAGGCAGCATACTGGACTGAAGCTGACGCTGAGGCACGAAAGCGTGGGG |
| P-asv6854 | CCGAGCGTTAATCGGAATTACTGGGCGTAAAGGGTCTGTAGGTGGTTTGTTAAGTCAGATGTGAAAGCCCAGGGCTCAACCTTGGAACTGCATTTGATACTGGCAAACTAGAGTATGGTAGAGGAATGGGGAATTTCTGGTGTAGCGGTGAAATGCGTAGAGATCAGAAGGAACACCAATGGCGAAGGCAACACTCTGGACCAATACTGACACTGAGGGACGAAAGCGTGGGG |
| P-asv6855 | GCAAGCGTTGTCCGGAATTACTGGGCGTAAAGAGTTCGTAGGCGGTCAAATAAGTCAGGAGTGAAAGGCTACGGCTCAACCGTAGTGAGCTTTTGAAACTGTTTGACTTGAGTACTGGAGAGGAAAGTGGGATTCCTAGTGTAGCGGTGAAATGCGTAGATATTAGGAGGAACACCAGTGGCGAAGGCGGCTTTCTGGACAGTAACTGACGCTGAGGAACGAAAGCGTGGGG |
| P-asv6856 | ACGAGTGATGGCCGATATTATTGGGCCTAAAGCGTCCGTAGCTTGCTGTGTAAGTCCATTGGGAGATCCACGCGCTCAACACGTCGGCGTCCAGCGGAAACTGTCTGGCTTGGGGCCGGAAGACTCGAGGGGTACGTCCGGGGTAGGAGTGAAATCCCGTAATCCTGGACGGACCGCCGGTGGCGAAAGCGCCTCGAGAGGACGGACCCGACGGTCAGGGACGAAAGCTAGGGT |
| P-asv6857 | GCAAGCGTTGTCCGGATTTATTGGGTTTAAAGGGTGCGTAGGCGGCTCATTAAGTCTGGGGTGAAAGCCCGTTGCTCAACAACGGAACTGCCCTGGAAACTGGTGAGCTTGAGTACAGACGAGGGTGGCGGAATGGATGCTGTAGCGGAGAAATGCATAGATAGCATCCAGAACCCCGATTGCGAAGGCAGCTGCCTAGACTGTAACTGACGCTGAGGCACGAAAGCGTGGGG |
| P-asv6858 | GCGAGCGTTGTCCGGAATCATTGGGCGTAAAGCGCGTGTAGGCGGCTTGGCAAGTCTGGTGTGAAAACTTGGGGCTCAACCCCAAGCGTGCACTGGAAACTGCCTCGCTAGAGTCCGGGAGAGGAGAGCGGAATTCCCAGTGTAGCGGTGAAATGCGCAGATATTGGGAAGAACACCGATGGCGAAGGCAGCTCTCTGGAACGGTACTGACGCTGAGACGCGAAAGCGTGGGG |
| P-asv6859 | GCTAGCGTTGCTCGGAATCACTGGGCGTAAAGGGTGCGTAGGCGGGTCTTTAAGTCAGGGGTGAAATCCTGGAGCTCAACTCCAGAACTGCCTTTGATACTGAAGATCTTGAGTATGGGAGAGGTGAGTGGAACTGCGAGTGTAGAGGTGAAATTCGTAGATATCAGGAGGAACACCGGTGGCGAAGGCGACTTTCTGGACCAATACTGACGCTGAGGCGCGAAGGCGTGGGG |
| P-asv6860 | GCTAGCGTTGTTCGGAATTACTGGGCGTAAAGAGTACGTAGGCGGATATTTAAGCCAGATGTGAAATCCCAGGGCTCAACCTTGGAACTGCATTTGGAACTGGATACCTAGAGTCCGTTAGAGGCAAGTGGAATACCTAGTGTAGGGGTGAAATTCGTAGATATTAGGAGGAACACCAGTGGCGAAGGCGACTTACTGGACTGTGACTGACGCTGAGGCACGAAAGCGTGGGG |
| P-asv6861 | GCAAGCGTTGTCCGGAATCATTGGGTGTAAAGGGTGCGTAGGCGGGCGTCTGCGTCGGGGGTGAAATCTTGCCGCTCAACGGTAAAATGGCCTTCGATACGGGGCGTCTTGAGTACAGCAGAGGCTGGTAGAATTCGTGGTGTAGCGGTGAAATGCATAGAGATCACGAAGAATACCGGTGGCGAAGGCGGCCAGCCAGTCTGTAACTGACGCTGAGGCACGAGAGCGTGGGG |
| P-asv6862 | GCAAGCGTTAATCGGAATTATTGGGCGTAAAGCGCGCGTAGGTGGTTCAGTAAGTTGGAAGTGAAATCCCCGGGCTCAACCTGGGAATTGCATTGGTGACTGCACGGCTAGAGTGTGTCAGAGGGGGGTAGAATTCCACGTGTAGCAGTGAAATGCGTAGAGATGTGGAGGAATACCGATGGCGAAGGCAGCCCCCTGGGATAACACTGACGCTCATGCACGAAAGCGTGGGG |
| P-asv6863 | GCAAGCGTTAATCGGAATTACTGGGCGTAAAGCGCGCGTAGGTGGCAAAGTAAGATGGGTGTGAAATCCCCGGGCTCAACCTGGGAACTGCATCCATAACTGCTTCGCTAGAGTACGGTAGAGGGTAGTGGAATTTCCTGTGTAGCGGTGAAATGCGTAGATATAGGAAGGAACACCAGTGGCGAAGGCGACTTTCTGGTCTGTAACTGACACTGAGGCGCGAAAGCGTGGGG |
| P-asv6864 | ACAAGCGTTGTCCGGATTTACTGGGCGTAAAGAGCGCGCAGGCGGTCGATCAAGTCGAGTGTGAAAGCCCCCGGCTCAACTGGGGAGGGTCATTCGAAACTGGTCGACTCGAAGGCAGGAGAGGGTAGTGGAATTCCCGGTGTAGTGGTGAAATGCGTAGATATCGGGAGGAACACCAGTGGCGAAGGCGACTACCTGGCCTGTTCTTGACGCTGAGGCGCGAAAGCTAGGGG |
| P-asv6865 | GCGAGCGTTGTCCGGAGTTACTGGGCGTAAAGAGCGTGTAGGCGGTTGAAAAAGTCAGGTGTGTAAATTTTCGGCTCAACCGAGAAACTGCGCCTGAAACTTTTCGACTGGAGTTTGGGAGGGGCAACTGGAATTCCTGGTGTAACGGTGAAATGTGTAGATATCAGGAGGAACACCCGTGGCGAAGGCGGGTCGCTGGACCAAAACTGACGCTGAGACGCGAAAGCGTGGGT |
| P-asv6866 | GCAAGCGTTGTTCGCAATTACTGGGCGTAAAGCGCGCGTAGGCGGCGATGATAGTCAGAGGTGAAATGCCCGGGCTCAACCCGGGAGGTGCCTTTGATACTTCATTGCTAGGATTCAGGAGAGGTGAGCGGAATTCCGAGTGTAGAGGTGAAATTCGTAGATATTCGGAAGAACACCAGTGGCGAAGGCGGCTCACTGGCCTGAGATCGACGCTGAGGTGCGAAAGCGTGGGG |
| P-asv6867 | GCAAGCGTTGTTCGGAATCATTGGGCGTAAAGGGCGTGTAGGCGGTTTGTTAAGTCATGTGTGAAATCCCTCGGCTCAACCGGGGAACGACGCATGAAACTGACAAGCTAGAGTACCAAAGAGGGGGGTGGAATTCCCGGTGTAGCGGTGAAATGCGTAGATATCGGGAAGAACACCTGTGGCGAAGGCGACCCCCTGGTTGGATACTGACGCTGATACGCGAAAGCGTGGGG |
| P-asv6868 | GCAAGCGTTAATCGGAATTACTGGGCGTAAAGCGCGCGTAGGTGGTTTGTTAAGTTGAATGTGAAAGCCCCGGGCTCAACCTGGGAGCTGCATTTGATACTGCAAGGCTAGAGATCGAGAGAGGAAAGTGGAATTACGAGTGTAGAAGTGAAATTCGTAGATATTCGTAGGAACACCAGTGGCGAAGGCGACTTTCTGGCTCGATACTGACGCTAAAGCGCGAAAGCGTGGGG |

| | |
|---|---|
| P-asv6869 | GCAAGCGTTATCCGGATTCATTGGGTTTAAAGGGAGCGTAGGTGGCCAAATAAGTCAGTGGTGAAATCTTGCAGCTTAACTGTAAAATTGCCATTGATACTGTATGGCTTGAGTATAGATGCCGTTGGCGGAATAT<br>GACATGTAGTGGTGAAATACATAGAGATGTCATAGAACACCGATTGCGAAGGCAGCTGACGAAACTATAACTGACACTGAGGCTCGAAAGTGCGGGG |
| P-asv6870 | GCGAGCGTTAATCGGAATTACTGGGCGTAAAGCGCGCGTAGGCGGCTATTTAAGTCTGATGTGAAATCCCCGGGCTTAACCCGGGAACTGCTTTTGAAACTCCAAGTCTAGAATCCTGGAGAGGTTGGGAGAATT<br>CCGAGTGTAGAGGTGAAATTCGCAGATATTCGGAGGAATACCAGTGGCGTAGGCGCCCAACTGGACAGGTATTGACGCTCAGATACGAAAGCGTGGGG |
| P-asv6871 | CCGGGCGTTATCCGGAATTATTGGGTTTAAAGGGAGCGCAGGCGGGAGTATAAGTCAGCTGTTAAATATCAGGGCTCAACTCTGTTATGCAGTTGAAACTATATTTCTTGAGTACGCACAGGGATGGCGGAATTCA<br>GGGTGTAGCGGTGAAATGCTTAGATATCCTGAAGAACTCCGATCGCGAAGGCAGCCATCCGGAGCGTAACTGACGCTGAGGCTCGAAGGTGCGGGT |
| P-asv6872 | GCAAGCGTTATCCGGATTTATTGGGTTTAAAGGGTCCGCAGGCGGATCGTTAAGTCAGTGGTGAAAGCCTACCGCTCAACGGTAGAACTGCCATTGATACTGGCGATCTTGAATATGTATGAAGTTGGCGGAATG<br>TGTGGTGTAGCGGTGAAATGCTTAGAAATCACACAGAACACCGATTGCGAAGGCAGCTGACTAATACAGTATTGACGCTGAGGGACGAAAGTGTGGGG |
| P-asv6873 | CCGAGTGATTGCCGCTATTATTGGGTCTAAAGCGCCCGTAGCCGGTCCTGTAAGTCCTTGGTGAGATCTGGCAGCAGACTGTCAGGCGTGCTGAGGTTACTGCAGGACTCGAGACCGGGAGAGGGTGCAGATAC<br>TCGTAGGGTAGGGGTTAAATCCTATAATCCTACGGGGATCACCTGTGGCGAAGGCGTGCACCTGGAACGGGTCTGACGGTGAGGGACGAAAGCTAGGGG |
| P-asv6874 | GCGAATGTTGCTCGGATTCACTGGGCGTAAAGTGTCTGTAGGTGGCTTGTTAAGTTAGAGGTGAAAGCCCAGCGCTCAACGTTGGAATTGCCTTTAAAACTGGCGAGCTTGAGAGTATCAGAGGTAGGCGGAATT<br>GCTGGTGTAGAGGTAAAATTCATAGATATCAGCAGGAACATCCATGGCGAAGGCAGCTTACTGGGGTATTTCTGACACTGAGAGACGAAAGCGTGGGG |
| P-asv6875 | GCAAGCGTTATCCGGAATTATTGGGCGTAAAGCGTCCGCAGGTGGCACCTCAAGTCTGCTGTCAAAGACCGGGGCTTAACCCCAGAAAGGCAGTGGAAACTGAAGTGCTAGAGTGCAGTAGGGGTAGAGGGAAT<br>TCCCGGTGTAGCGGTGAAATGCGTAGAGATCGGGAAGAACATCGGTGGCGAAAGCGCTCTGCTGGACTGCAACTGACACTCAGGGACGAAAGCTAGGGG |
| P-asv6876 | CCTAGCGTTGTTCAGAATTATTGGGCGTAAAGCGGATGTAGGTGGCCTGTTAAGTCAGAAGTGAAAGCCCTGGGCTCAACCCAGGAAGGGCTTTTGATACTGGCAGGCTTGAGTGCAGGAGAGGTAAGTGGAAT<br>TTCTGGTGTAGTGGTGAAATACGTAGATATCAGAAGGAACATCGGTGGGGAAGCCGGCTTACTGGCCTGACACTGACACTAAGATCCGAAAGCGTGGGG |
| P-asv6877 | GCGAGCGTTGTTCGGAATTATTGGGCGTAAAGGGCATGTAGGCGGATGAGCAAGCTTGGTGTGAAATGCTACAGCTTAACTGTGGGACAGCATTGAGAACTGCACATCTTGAATTACTGAAGGGTAACTAGAATT<br>CCACGTGTAGGGGTGAAATCTGTAGATATGTGGAAGAATACCAATGGCGAAGGCAAGTTACCGGCAGATAATTGACGCTGAGGTGCGAAAGTGCGGGG |
| P-asv6878 | GCGAGCGTTAATCGGAATAACTGGGCGTAAAGGGCACGCAGGCGGTGACTTAAGTGAGATGTGAAAGCCCCGGGCTTAACCTGGGAATTGCATTTCATACTGGGTCGCTAGAGTACTTTAGGGAGGGGTAGAAT<br>TCCACGTGTAGCGGTGAAATGCGTAGATATGCGGAGGAACACCGATGGCGAAGGCAATCCCCTGGGCCTGCACTGACGCTCATGCACGAAAGCGTGGGG |
| P-asv6879 | GCAAGCGTTAATCGGAATTACTGGGCGTAAAGCGCACGTAGGCGGTTTTTTAAGTCAGATGTGAAAGCCCCGGGCTCAACCTGGGAATTGCATTTGAAACTCCCAGTCTTGAGGTCGAGAGAGGTGAGTGGAATT<br>CCGAGTGTAGGGGTGAAATTCGTAGATATATGGAAGAACACCCGTGGCGAAGGCGGTTTTCTAGCTTAGACCTGACGCTGAAGCGCGAAAGCAAGGGG |
| P-asv6880 | CCAAGCGTTATCCAGAATCACTGGGTGTAAAGGGTACGCAGGCGGACATGTAGGTCTAATGTGGAAGTCCTGGGCTTAACCCAGGAATTGCATCCAAAACTGGCAAGCTAGAGTATGGCAGAGGGTGGTGGAAT<br>TTCCTGTGTAGCGGTGAAATGCGTAGATATAGGAAGGAACACCAGTGGCGAAGGCGACCACCTGGGCTAATACTGACACTGAGGTGCGAAAGCGTGGGG |
| P-asv6881 | GCTAGCGTTGCTCAGAATCACTGGGCGTAAAGGGTGCGTAGGCGGGTCTTTAAGTCAGGGGTGAAATCCTGGAGCTCAACTCCAGAACTGCCTTTGATACTGGAGACCTTGAGTGTACTAGAGGTAGGCGGAAT<br>TTATGGTGTAGCGGTGAAATGCATAGATACCATAAAGAACACCGATAGCGTAGGCAGCTTACTGGAGTACAACTGACGCTGAGGCACGAAAGCATGGGG |
| P-asv6882 | GCAAGCGTTAATCGGAATTACTGGGCGTAAAGCGCACGCAGGCGGCTTTTTAAGTCGGATGTGAAAGCCCCGGGCTCAACCTGGGAATTGCATTCGATACTGTAAAGCTAGAGTATGTGAGAGGGGGGTAGAAT<br>TCCAGGTGTAGAGGTGAAATTCGTAGATATTCGCAAGAACACCAATGGCGAAGGCAGCCCCCTGGGCCTGTACTGACGCTCATGCACGAAAGCGTGGGG |
| P-asv6883 | GCAAGCGTTGTCCGGAATGACTGGGCGTAAAGCGCGCGCAGGCGGCTTCTTAAGTTGGATGTGAAATCCCCGGGCTCAACCTGGGAACTGCATTGGAGACTGCAAGGCTAGAGTGTGTCAGAGGGGGGTAGAA<br>TTCCACGTGTAGCAGTGAAATGCGTAGAGATGTGGAGGAATACCGATGGCGAAGGCAGCCCCCTGGGATAACACTGACGCTCATGCACGAAAGCGTGGGG |
| P-asv6884 | GCAAGCGTTGTCCGGAATCATTGGGCGTAAAGGGTGCGTAGGCGGTTTGTTAAGTCTGCATTGCAAGTTGAGGGCTCAACCCTGAAATTGGTGCGGAAACTGGCAGACTGGAGTACTGCAGGGGAAAGCGGAAT<br>ACCCGGTGTAGAGGTGAAATTCGTAGATATCGGGTGGAACACCAAACGCGAAGGCAGCTTTCTGGGCAGTAACTGACGCTGAGGCACGAAAGCGTGGGG |
| P-asv6885 | GCTAGCGTTATTCGGTGTCACTGGGCTTAAAGCGTATGTAGGCGGAAATGTAAGCGTTTTGTGAAATCCCTCGGCTTAACCGAGGAATTGCTTGGCGAACTACATTTCTTGAGGCAAGTAGGGGTGCATGGAACT<br>CTAGGTGGAGCGGTGGAATGCGTAGATATCTAGAGGAACGCCGATGGAGAAATCAGTGCACTGGGCTTGTCCTGACGCTCAGATACGAAAGCGTGGGG |
| P-asv6886 | GCAAGCGTTGTCCGGAATTATTGGGCGTAAAGCGCGCGCAGGTGGTTCTTTAAGTCTGATGTGAAAGCCCACGGCTCAACCGTGGAGGGTCATTGGAAACTGGGGAACTTGAGTGCAGAAGAGGAAAGTGGAAT<br>TCCACGTGTAGCGGTGAAATGTGTAGAGATGTGGAGGAATACCGATGGCGAAGGCAGCCCCCTGGGATAACACTGACGCTCATGCACGAAAGCGTGGGG |
| P-asv6887 | CCGAGCGTTAATCGGAATCACTGGGCTTAAAGCGTACGTAGGCGGACATGTAGGCATTTTGTGAAATCCCTCGGCTCAACCGGGGGATTGCAGAGCGAACCACATGTCTTGAGGCGAGTAGAGGCGGTCGGAAC<br>GATAGGTGGAGCGGTGAAATGCGTAGATATCTATCGGAACGCCAAAGGTGAAGACAGGCCGCTGGGCTCGTCCTGACACTGAGGTACGAAAGCGTGGGT |
| P-asv6888 | GCAAGCGTTAATCGGAATTACTGGGCGTAAAGGGCGCGTAGGTGGAAGTTTAAGTCAGAGGTGAAATCCCTGGGCTTAACCTAGGAATTGCCTCTGATACTGGACTTCTAGAGTACAGTAGAGGAAAGTGGAACT<br>TCCGGTGTAGCGGTGAAACGCGTAGATATCGGAAGGAACATCAGTGGCGAAGGCGACTTTCTGGACTGACACTGACACTGAGGTGCGAAAGCGTGGGT |
| P-asv6889 | GCGAGCATTATTCGGAATGATTAGGCGTAAAGAGTTTGTAGGTGGTTTTTTAAGTTGAAAAAAACAAATTAAAGCTCAACTTTATATATTTTTTCAAAACTGAGAAGCTAGAGTATAAATAGAGGATAATGGAATTTCT<br>ATTGGAGTGATAAAATACGTTGATAATAGAAGGAAGACCAAGAGCGAAGGCAATTATCTGGGTATATACTGACACTGAGAAACGAAAGCTTGGGT |
| P-asv6890 | CCAAGCGTTGTTCAGATTCACTGGGCGTAAAGGGTGCGTAGGTGGTGAGGTAAGTTTGATGTGAAATCTCCAGGCTTAACCTGGAAAAGGCATTGAATACTATCTTGCTTGAGGGTTGGAGGGGAGACTGGAATG<br>CTCGGTGTAGCAGTGAAATGCGTAGATATCGAGCGGAACACCAGTGGCGAAGGCGAGTCTCTGGACAACTCCTGACACTGAGGCACGAAAGCTAGGGG |
| P-asv6891 | GCAAGCGTTATCCGGATTTATTGGGTTTAAAGGGTCCGCAGGCGGGAGAATAAGTCAGTGGTGAAATCCTATCGCTTAACGATAGAACTGCCATTGATACTGTTTTTCTTGAGTGTAGTTGAAGTTGGCGGAATGT<br>GTCATGTAGCGGTGAAATGCATAGATATGACACAGAACACCGATTGCGAAGGCAGCTAACTAAACTATAACTGACGCTCATGGACGAAAGCGTGGGG |
| P-asv6892 | CCGAGCGTTATCCGGATTTACTGGGCGTAAAGCGTGTTCAGGCGGTCTAGCAAGTCGGGCATGAAATCTCTCGGCTCAACCGGGAGGGGTTGTCCGATACTGCTGGACTGGAGGGCAGCAGAGGGTGGTGGAA<br>TTCCGGGTGTAGTGGTGAAATGCGTAGATATCCGGAGGAACACCAGTGGCGAAGGCGGCCACCTGGGCTGTACCTGACGCTAACACACGAAAGCTAGGGG |
| P-asv6893 | GCAAGCGTTATCCGGATTTATTGGGTTTAAAGGGTCCGTAGGCGGATCTGTAAGTCAGTGGTGAAATCTCACAGCTTAACTGTGAAACTGCCATTGATACTGTAGGGCTTGAATTTTTGTGAAGTAACTAGAATATG<br>TAGTGTAGCGGTGAAATGCTTAGATATTACATGGAATACCAATTGCGAAGGCAGGTTACTAACAAACGATTGACGCTGATGGACGAAAGCGTGGGG |
| P-asv6894 | TCAAGCGTTGTTCGGAATCACTGGGCGTAAAGGGTGCGTAGGTGGCGTGGCAAGTCAGATGTGGAAGCCCGGGGCTCAACCTCGGAACTGCATCCGATACTGCCATGCTGGAGCGCTGAAGGGGTGACTGGAA<br>TTCTCGGTGTAGCAGTGAAATGCGTGGATATCGAGAGGAACGCCAAAGGCGAAGGCAGGTCACTGGGCAGTTGCTGACACTGAGGCACGAGGGCCAGGGT |
| P-asv6895 | GCGAACGTTGTCCGGATTCACTGGGCGTAAAGCGCGCGCAGGCGGACTGGTAAGTCCGAGGTGAAAGCTCCCGGCTCAACTGGGAGAAGCCCTTGGATACTGCCCGTCTTGAGACATCTAGAGGAGAGTGGAA<br>TTCCCGGTGTAGTGGTGGAATGCGTAGATATCGGGAGGAACACCAGTGGCGAAAGCGGCTCTCTGGGGATGTTCTGACGCTGAGGCGCGAAAGCGTGGGG |
| P-asv6896 | CCAAGCGTTATCCGGAATTATTGGGCGTAAAGGGTGCGTAGGTGGTCCTGTTAGTCAGGGGTCAAAGACCGGTGCTTAACATCGGGAAGGCCTTTGAAACGGCAGGACTTGAGGGTGAGAGAGGCATAGGGAA<br>CTCTAGGTGTAGGGGTGAAATCCGTTGATATCTAGGGGAACACCAAAGGCGAAGGCACTATGCTAGCTCATTCCTGACACTCATGCACGAAACCCTGGGT |
| P-asv6897 | GCTAGCGTTGTTCGGAATTACTGGGCGTAAAGCGCGCGCAGGCGGCTTGTCAAGTCAGATGTGAAATCCCGGGGCTTAACCTCGGAATTGCATTTGAAACTGGCTCGCTAGAGATAGGTAGAGGAGAGTGGAAT<br>TCCCAGTGTAGAGGTGAAATTCGTAGATATTGGGAGGAACACCGGAGGCGAAGGCGGCTCTCTGGACCTATTCTGACGCTGAGGCGCGAAAGCGTGGGG |
| P-asv6898 | CCTAGCGTTGCTCGGAATCATTGGGCGTAAAGCGGGGGTAGGCGGACAACCAAGTCAGATGTGAAAGCCCAGGGCTTAACCCTGGAATTGCTTTTGAGACTGTATGGCTAGAGATCTAGAGAGGTTGGGGGAAT<br>TCCGAGTGTAGAGGTGAAATTCGCAGATATTCGGAGGAACACCAGTGGCGTAGGCGCCCAACTGGCTAGATACTGACGCTGAGGCACGAAAGCGTGGGG |
| P-asv6899 | GCAAGCGTTATCCGGAATCACTGGGCGTAAAGGGTGCGTAGGCGGTGTGATTAGTCGGGTGTTAAATCCCGGGGCTCAACCCCGGAACTGCCCTTGAAACTGGAAAGCTAGAATACTGGAGAGGTGAGTGGAAT<br>TCCGAGTGTAGAGGTGAAATTCGTAGATATTCGGAAGAACACCAGTGGCGAAGGCGACTCACTGGACAGTTATTGACGCTGAGGTGCGAAAGCGTGGGG |
| P-asv6900 | GCAAGCGTTACTCGGATTTACTGGGCGTAAAGCGCGCGCAGGCGGATCGGCAAGTCAGGTGTGAAATCCCGGGGCTTAACTCCGGAATTGCACTTGAAACTAGAAAGCCGGAGGACGAGAGAGGATAGTAGAAT<br>TCCTAGTGTAGGAGTGAAATCCGTAGATATTAGGAGGAATACCAGTGGCGAAGGCGGCTATCTGGCTCGTTTCTGACGCTGAGGCTCGAAAGCGTGGGG |
| P-asv6901 | CCAAGCGTTGTCCGGATTTATTGGGTTTAAAGGGTGCGTAGGCGGATTAATAAGTCAGTGGTGAAAGCTTTCGGCTCAACCGGAAAATTGCCATTGAAACTGTTAATCTTGAGTATAGTTGGGGCGAATGGAATGT<br>TGCATGTAGCGGTGAAATGCTTAGATATGCAACAGAACACCGATCGCGAAGGCAGTTCACTAAACTATAACTGACGCTGAGGCACGAAAGCGTGGGT |
| P-asv6902 | GCTAGCGTCGTTCGGAATTACTGGGCGTAAAGCGCGCGTAGGCGGTTTTGCAAGCCAGAGGTGAAATGCCTGAGCTCAACTTAGGAATTGCCTTTGGGACTGCGAGGCTTTGAGGACTGGAGAGGTGAGTGGAA<br>TTCCCAGTGTAGAGGTGAAATTCGTAGATATTGGGAAGAACACCGGTGGCGAAGGCGGCTCACTGGCCAGTTTCTGACGCTGCAGCGCGAAAGCGTGGGG |
| P-asv6903 | TCAAGCGTTGCTCGGTGTCACTGGGCTTAAAGGGCGCGTAGGCTGTGCAGCACGGTTCGGGTGAAATCCCTCGGCTCAACCGAGGAACTGCCCGGGCGACGGCTGCGCTGGACTGCATCAGGGGCGGATGGA<br>ACTCCCGGTGGAGCGGTGAAATGCGTAGATATCGGGAGGAACGCCGGTGGCGAAAGCGATCCGCTGGGGTGCAAGTGACGCTGAGGCGCGAAAGCCAGGGG |
| P-asv6904 | GCTAGCGTTGTTCGGAATTACTGGGCGTAAAGCGCGCGTAGGCGGTCTATTAAGTCAGAGGTGAAATCCCGGGGCTCAACCCCGGAACTGCCTTTGAAACTGCTAGACTAGAATCTTGGAGAGGTCAGTGGAATT<br>CCGAGTGTAGAGGTGAAATTCGTAGATATCGGGAGGAACACCTATGGCGAAAGCAGCTGCCTGGGCCTTTACTGACGCTAAGACTCGAAAGCTGGGGG |
| P-asv6905 | GCGAACGTTACTCGGAATCATTGGGCTTAAAGCGTGCGTAGGCGGCGACATAAGTTGTAGGTGAAATCCCAACGGCTCAACCGTGGAACTGCCTTCAATACTGTGTTGCTTGAGGGAGAAAGGGGCATTCGGAA<br>CTTCCGGTGGAGCGGTGAAATGTGTTGATATCGGAAGGAACGCCGGTGGCGAAAGCGGGATGCTAGGTCTCTTCTGACGCTGAGGCACGAAAGCTAGGGT |
| P-asv6906 | GCGAGCGTTGTTCGGATTTACTGGGCGTAAAGGGTGCGTAGGCGGTTGCGTGTGTCAGATGTGAAAGCCCGGGGCTCAACCCCGGGACTGCATTCGAAACTGCGTGACTTGAGTATCGGAGAGGTATGTGGAA<br>TTCTTAGTGTAGCGGTGAAATGCGTAGATATTAAGAAGAACACCGGCGGCGAAAGCGGCATACTGGACGAATACTGACGCTGAGGCACGAAAGCCAGGGG |
| P-asv6907 | CCAAGCGTTAATCGGAATTACTGGGCTTAAAGCGTGCGTAGGCGGATGTGCAAGCGTCCTGTGAAAGCCCCCGGCTCAACCGGGGAACAGCGGGGCGAACTGCTCGTCTTGAGGCAGGTAGGGGCAGACGGAA<br>CCGTCGGTGGAGCGGTGAAATGCGTAGAGATCGACGGGAACGCCGATGGCGAAGGCAGTCTGCTGGGCCTGTCCTGACGCTGAGGTACGAAAGCGTGGGG |

| | |
|---|---|
| P-asv6908 | GCAAGCGTTACTCGGATTCACTGGGCGTAAAGGGTGCGTAGGCCGCTAAGCGTGTCAGGTGTGAAATCTCGGGGCTTAGCCTCGAAACTGCGCCTGAAACTGTTTAGCTTGAGTATTGGAGAGGTAAGCGGAAT<br>TTCTGGTGTAGCGGTGAAATGCGTAGATATCAGAAGGAACACCAATGGCGAAGGCAGCTTACTGGACAATAACTGACGCTGAGGCACGAAAGCGTGGGT |
| P-asv6909 | GCAAGCGTTGTCCGGAATTATTGGGCGTAAAGAGTTCGTAGGCGGTATGTCAAGTCTGGTGTTAAAGACTGGGGCTCAACCCCAGAAGGGCACTGGATACTGGCAAACTGGAGTGCAGAAGAGGCAAGGGGAAT<br>TCCCAGTGTAGCGGTGAAATGCGTAGATATTCGGAAGAACACCAGTGGCGAAGGCGACTCGCTGGACAAGTATTGACGCTGAGGTGCGAAAGCGTGGGG |
| P-asv6910 | CCGAACGTTATTCGGAATTACTGGGCTTAAAGGGTGCGTAGGCGGCCATACAAGTCAGATGTGAAATCCCACGGCTCAACCGTGGAACTGCGTTTGAAACTGGGAGTCTGGAGATCAGGAGAGGTTAGCGGAAT<br>ACCGAGTGTAGAGGTGAAATTCGTAGATATTCGGTGGAACACCAGTGGCGAAGGCGGCTAACTGGACTGATACTGACGCTGAGGTGCGAAAGTGTGGGG |
| P-asv6911 | GCGAGCGTTGTTCGGATTCACTGGGTGTAAAGGTCATGTAGGCGTCTCTTTAAGTCGGGTGTGAAATCCCTTGGCTCAACCAAGGAACTGCGCCCGATACTGGGGAGATTGAGTACTTGAGAGGTAAGCGGAATT<br>CCGGAAGTAAGGGTGAAATCTGTAGATATCCGGAAGAACACCATTAGCGAAGGCGGCTTACTGGCAAGATACTGACGCTGAAATGAGAAAGCTAGGGG |
| P-asv6912 | CCGAGTGGGAATCACATATTGGGCCTAAAGCGTTCGTAGTCGGCTTTGTAAATTCCTTGTGAAATCGTTGTGCTTAACGCAACGACGTGCGGGGGACACTGCATAGCTAGGGACCGGGAGGGGTAAGAAGTATTC<br>CTTGGGGAGCGGTAAAATGTGATAATCCAAGGGGGACTACCTGTGGCGAAGGCGTCTTACCAGAACGGGTCCGACGATCAGGAACGAAGGCTAGGGG |
| P-asv6913 | GCTAGCGTTGTTCGGAATTACTGGGCGTAAAGCGCATGAAGGTGGTGAAGTAAGTCGGATGTGAAATCCCGGGGCTTAACCCCGGAAGTGCATCCGGGACTGCTTTGCTCGAGTCCGGAAGAGGAGGGCGGAA<br>TTCCCAGTGTAGAGGTGAAATTCGTAGATATTGGGAAGAACACCGGTGGCGAAGGCGGCTCTCTGGTCCGGTACTGACACTATAGTGCGAAAGCGTGGGG |
| P-asv6914 | GCTAGCGTTGTTCGGAATTACTGGGCGTAAAGCGCACGTAGGCGGCTTTTTAAGTCAGGGGTGAAAGCCCGGGGCTCAACCCCGGGTCTGCAGTGGGTACGGGCAGACTAGAGTGCAGTAGGGGAGACTGGAA<br>TTCCTGGTGTAGCGGTGAAATGCGCAGATATCAGGAAGAACACCGATGGCGAAGGCAGGTCTCTGGGCTGTTACTGACGCTGAGGATCGAAAGCGTGGGG |
| P-asv6915 | GCAAGCGTTAATCGGAATTACTGGGCGTAAAGGGTTCGTAGGTGGTTTGATAAGTTATCTGTAAAAGACCTTGGCTCAACCCCGGAACGGCCCTCGATACTGGCAGCCTAGAGTTCGAGAGAGGTGAGTGGAATT<br>CCGAGTGTAGAGGTGAAATTCGTAGATATTCGGAGGAACACCAGTGGCGAAGGCGGCTCACTGGCTCGATACTGACGCTGAGGTGCGAAAGCGTGGGG |
| P-asv6916 | GCAAGCGTTGTTCGGATTTACTGGGCGTAAAGGGCGTGTAGGCGGACTGTTAAGCTGGGGGTGAAAGCCCGGGGCCCAACCCCGGAGGGTCTCCCAGAACTGGCAGTCTAGAGAAAGTCAGAGGCAGGTGGA<br>ATTCCAGGTGTAGAGGTGAAATTCGTAGAGACCTGGAAGAACACCGGCGGCGAAGGCGGCCTGCTGGGGCTTTTCTGACGCTGAGGCGCGAAAGCGTGGGT |
| P-asv6917 | GCGAGCGTTAATCGGAATTACTGGGCGTAAAGCGTGCGCAGGCGGTTGTGTAAGACAGGTGTGAAAGCCCCGGGCTCAACCTGGGAGGTGCGTTTGATACTGCGTTGCTTGAGAGCGGAAGAGGGTAGTGGAA<br>TTCCCAGTGTAGAGGTGAAATTCGTAGATATTCGCAAGAACACCAGTGGCGAAGGCGGCTCACTGGCCCATAACTGACGCTGAGGCACGAAAGCGTGGGG |
| P-asv6918 | GCTAGCGTTGTTCGGAATTACTGGGCGTAAAGCGCACGTAGGCGGACCAGAAAGTCAGAGGTGAAATCCCAGGGCTTAACCCCGGAATTGCTCTTGAAACTCCGGATCTAGAATATGGTAGAGGTTAGGAGAATT<br>CCTAGTGTAGAGGTGAAATTCGCAGATATTAGGAGGAATACCAGTGGCGTAGGCGCCTAACTGGACCATTATTGACGCTCATGTACGAAAGCGTGGGG |
| P-asv6919 | GCGAGCGTTGTTCGGAATTATTGGGCGTAAAGGGTGCGCAGGCGGTTTTACAAGTCTGGTGTTAAAGCCCGCAGCTTAACTGCGGAAAGGCATTGGAAACTGTGAGACTAGAGTTCTGGAGGGGTAGCTGGAAT<br>TCCAGGTGTAGGGGTGAAATCTGTAGATATCTGGAAGAACATCAATGGCGAAGGCAAGCTACTGGCCATGAACTGACGCTGAGGCACGAAAGCGTGGGG |
| P-asv6920 | GCAAGCGTTACTCGGAATCACTGGGCGTAAAGAGCGTGTAGGCGGATATTTAAGTCAGAAGTGAAAGCCTGGGGCTCAACCCCAGAACTGCTTTTGAAACTGGATGGCTAGAGTCCCGGAGAGGTTGGCGGAAT<br>TCCGAGTGTAGAGGTGAAATTCGCAGATATTCGGAGGAACACCAGTGGCGTAGGCGGCCAACTGGACGGGAACTGACGCTGAGGTACGAAAGCGTGGGG |
| P-asv6921 | GCGAGCGTTAATCGGAGTTACTGGGCGTAAAGGGCGCGTAGGCGGTGACTTAAGTCAGTTGTGAAATCCCAGGGCTTAACCTTGGAACTGCAATTGAAACTGGGTGACTAGAGTCGAAGAGAGGGAAGCGGAAT<br>TTCCGGTGTAGCGGTGAAATGCGTAGATATCGGAAGGAACACCCGTGGCGAAGGCGGCTTCCTGGCTTCAGACCGACGCTGAGGCGCGAAAGCGTGGGT |
| P-asv6922 | GCAAGCGTTAATCGGAATTACTGGGCGTAAAGCGTGCGTAGGCGGTTGTTTAAGTCAGATGTGAAAGCCCCGGGCTTAACCTGGGAACTGCATTTGATACTGGATGACTAGAGTTGAGTAGAGGGGAGTGGAATT<br>TCAGGTGTAGCGGTGAAATGCGTAGAGATCTGAAGGAACACCAGTGGCGAAGGCGGCTCCCTGGACTCAAACTGACGCTGAGGTACGAAAGCGTGGGT |
| P-asv6923 | GCAAGCGTTATCCGGAATTATTGGGCGTAAAGAGCTCGTAGGCGGTTTGTCGCGTCTGCTGTGAAATCCGGAGGCTCAACCTCCGGCCTGCAGTGGGTACGGGCAGACTAGAGTGCGGTAGGGGAGATTGGAA<br>TTCCTGGTGTAGCGGTGGAATGCGCAGATATCAGGAGGAACACCGATGGCGAAGGCAGATCTCTGGGCCGTAACTGACGCTGAGGAGCGAAAGGGTGGGG |
| P-asv6924 | GCAAGCGTTGTCCGGATTTATTGGGCGTAAAGAGCGTGTAGGCGGCCAAGCAGGTCCGTTGTGAAAACTGAAGGCTCAACCTTCAGAGGTCGATGGAAACCGCTTGGCTAGAGTCCGGAAGAGGGGAGTGGAA<br>TTCCTGGTGTAGCGGTGAAATGCGCAGATATCAGGAAGAACACCCGTGGCGAAGGCGGCTCTCTGGGACGGTACTGACGCTGAGACGCGAAAGCGTGGGG |
| P-asv6925 | GCGAGCGTTATTCGGAATCACTGGGCGTAAAGCGTGCGCAGGCGGTTTGGGAAGTCAGAAGTGAAAGCCCGAGGCTCAACCTCGGAATTGCTTTTGAAACTCCCTCACTAGAGTCCGGGAGAGGCCAGCAGAAT<br>TCCTAGTGTAGAGGTGAAATTCGTAGATATTAGGAGGAATACCAGTGGCGTAGGCGGCTGGCTGGACCGGTACTGACGCTCAGGCACGAAAGCGTGGGG |
| P-asv6926 | GCAAGCGTTGTTCGGAATTACTGGGCGTAAAGGGTTCGTAGGTGGCCAACTAAGTCAGATGTGAAATCCCTCAGCTTAACTGGGGAACTGCTTCTGATACTGGTTGGCTTGAGTGCAGGAGAGGAATGCGGAATT<br>CCAGGTGTAGCGGTGAAATGCGTAGATATCTGGAGGAACACCGGTGGCGAAGGCGGCATTCTGGACTGTAACTGACACTGAGGAACGAAAGCCAGGGG |
| P-asv6927 | GCTAGCGTTACTCGGAATTACTGGGCGTAAAGGATGCGTAGGCGGATTATCAAGTTGATAGTGAAATCCCCAAGCTTAACTTGGGAACTGCTGTCAAAACTGGTAATCTAGAATGCAGGATGGAGTAAGGGAATTC<br>TTGGTGTAGGGGTAAAATCCGTAGATATCAGGAGGAAGACCGGAAGCGAAAGCGCTTACTTGGACTGTTATTGACGCTGAGGCATGAAAGCGTGGGG |
| P-asv6928 | GCTAGCGTTGTTCGGATTTACTGGGCGTAAAGGGCGCGTAGGCGGCCTGATCAGTCGGGTGTGAAAGCCCGGGGCTCAACCCCGGAACGGCACCCGAGACGGTCAGGCTGGAGACCGGGAGAGGAGGGTGG<br>AATAGCCAGTGTAGAGGTGAAATTCGTAGATATTGGCTGGAACACCGGTGGCGAGAGCGGCTCTCTGGCCCGGTTCTGACGCTGAGGCGCGAAAGCGTGGGG |
| P-asv6929 | CCAAGCGTTATCCGGATTTATTGGGTTTAAAGGGTGCGTAGGCGGACTGGTAAGTCAGTGGTGAAATCCTGCAGCTTAACTGTAGAACTGCCATTGATACTGCTGGTCTTGAGTACAGTTGAGGTAGGCGGAGCG<br>AGTAGTGTAGCGGTGAAATGCTTAGATATTACTCAGAACACCGATTGCGAAGGCAGCTTGCCAAACTGACACTGACGCTGATGCACGAAAGCGTGGGG |
| P-asv6930 | GCAAGCGTTATCCGGAATGATTGGGCGTAAAGAGTCCGTAGGTAGTCATCCAAGTCTGCTGTTAAAGAGCGAGGCTTAACCTCGTAAAGGCAGTGGAAACTGGAAGACTAGAGTGTAGTAGGGGCAGAGGGAAT<br>TCCTGGTGTAGCGGTGAAATGCGTAGAGATCAGGAAGAAGACCGGTGGCGAAGGCGACTCACTGGCCCGTTACTGACGCTGAGGCGCGAAAGCGTGGGG |
| P-asv6931 | GCAAGCGTTAATCGGAATTACTGGGCGTAAAGCGCGCGTAGGCGGCTTGTTAAGCCAGATGTGAAAGCCCCGGGCTCAACCTGGGAACTGCATTTGAGACTGCATCGCTTGAATCCAGGAGAGGTGAGTGGAAT<br>TCCGAGTGTAGAGGTGAAATTCGTAGATATTCGGAAGAACACCAGTGGCGAAGGCGGCTCACTGGCCCGATACTGACGCTGAGGCACGAAAGCGTGGGG |
| P-asv6932 | CCGAGTGACGGCCGATCTTATTGGGCCTAAAGCGTCCGTAGCCGGCCGAGCAAGTCCGTCGGGAAATCCACGCGCTCAACGCGTGGGCGTCCGGCGGAAACTGTTCGGCTTGGGACCGGGAGACCAGAGGGG<br>TACGTCCGGGGTAGGAGTGAAATCCCGTAATCCTGGACGGACCACCGGTGGCGAAAGCGCCTCTGGAGAACGGATCCGACGGTGAGGGACGAAAGCCAGGGT |
| P-asv6933 | GCGAGCGTTAATCGGAATTACTGGGCGTAAAGCGTGCGTAGGCGGTTATGTAAGTCGGATGTGAAAGCCCCGGGCTTAACCTGGGAATTGCATTCGAAACTGCATGACTAGAGTGTGGTAGAGGAAGATGGAAT<br>TCCACGTGTAGCGGTGAAATGCGTAGATATGTGGAGGAACACCAATGGCGAAGGCAGTCTTCTGGGCCAACACTGACGCTGAGGCACGAAAGCGTGGGT |
| P-asv6934 | CCAAGCGTTATCCGGAATCATTGGGTTTAAAGGGTCCGTAGGTGGATAATTAAGTCAGAGGTGAAATCTTGCAGCTCAACTGTAAAATTGCCTTTGATACTGGTTATCTTGAGTTATTATGAAGTAGTTAGAATATGT<br>AGTGTAGCGGTGAAATGCATAGATATTACATAGAATACCAATTGCGAGGGCAGATTACTAATAATCAACTGACACTGATGGACGAAAGCGTGGGT |
| P-asv6935 | GCAAGCGTTGTTCGGATTTACCGGGCGTACAGGGAGCGTAGGCGGTTAGGTAAGCCCTCCGTGAAATCTCCGGGCCTAACCCGGAAAGTGCGGAGGGGACTGCTTAGCTAGAGGATGGGAGAGGAGCGCGGA<br>ATTCCCGGTGTAGCGGTGAAATGCGTAGAGATCGGGAGGAAGGCCGGTGGCGAAGGCGGCGCTCTGGAACATTTCTGACGCTGAGGCTCGAAAGCGTGGGG |
| P-asv6936 | GCAAGCGTTAATCGGAATTACTGGGCGTAAAGCGCGCGTAGGCGGCTGTTTAAGTCGGATGTGAAAGCCCTGGGCTCAACCTGGGAACTGCATTCGATACTGGGCAGCTAGAGTATAGGAGAGGAGTGTGGAAT<br>TTCAGGTGTAGCGGTGAAATGCGTAGAGATCTGAAGGAACACCAGTGGCGAAGGCGACACTCTGGCCTAATACTGACGCTGAGGGGCGAAAGCGTGGGG |
| P-asv6937 | GCAAGCGTTATCCGGAATGATTGGGCGTAAAGCGTCTGTAGGTGGTTTATTAAGTCTGCTGTTAAAGCACAGGGCTTAACCCTGTATAGGCGGTAGAAACTGGTAGGCTTGAGTATGGTAGGGGTAGAGGGAATT<br>CCCGATGTAGTGGTGAAATGCGTTGATATCGGGAAGAACACCAAAGGCGAAAGCACTCTGCTGGGCCAATACTGACACTAAGAGACGAAAGCTAGGGG |
| P-asv6938 | GCAAGCGTTATTCGGAATCATTGGGCTTAAAGAGCGCGTAGGCGGCTGGGTAAGTTAGAGGTGAAAGCCCATCGGCTCAACCGTGGAATTGCCTCTAATACTGCTTAGCTTGAGGGAGACAGGGGCATGTGGAA<br>CTTCCGGTGGAGCGGTGAAATGTGTTGAGATCGGAAGGAACGCCGGTGGCGAAAGCGGCATGCTGGGGCTCTTCTGACGCTGAGGCGCGAAAGCTAGGGT |
| P-asv6939 | GCAAGCGTTGTTCGGAATCACTGGGCTTAAAGCGTGCGTAGGCGGATTGCCAAGTCAGGAGTGAAATCCCGGGGCTCAACCCCGGAATTGCTCTTGAAACTGGCAGTCTAGAATCCGATAGAGGTCAGGAGAAT<br>TCCTAGTGTAGAGGTGAAATTCGCAGATATTAGGAGGAATACCAGTGGCGTAGGCGCCTGACTGGGTCGGTATTGACGCTGAGGTGCGAAAGCGTGGGG |
| P-asv6940 | GCTAGCGTTGTTCGGAATTACTGGGCGTAAAGCGCGCGTAGGCGGCTTTTTAAGTCGGATGTGAAAGCCCCGGGCTCAACCTGGGAACTGCATCCGATACTGGAAGGCTAGAGTGCGAGAGAGGGAGGTAGAA<br>TTCCATGTGTAGCGGTGAAATGCGTAGATATATGGAGGAATACCAGTGGCGAAGGCGGCCTCCTGGCTCGACACTGACGCTGAGGTGCGAAAGCGTGGGG |
| P-asv6941 | GCTAGCGTTGCTCAGAATTACTGGGCGTAAAGGGTCCGTAGGCGGATGCGTAAGTCAGTGGTGAAATCTCACAGCTCAACTGTGAAACTGCCATTGATACTGCATGTCTTGAGTAAGGTAGAAGTGGCTGGAATA<br>AGTAGTGTAGCGGTGAAATGCATAGATATTACTTAGAACACCAATTGCGAAGGCAGGTCACTATGTCTTAACTGACGCTGATGGACGAAAGCGTGGGG |
| P-asv6942 | CCGAGCGTTGTCCGGAGTTACTGGGCGTAAAGCGTACGCAGGCGGCGGCGTTGGCCCGGCGTGAAAGCCCCCGGCTCAACCGGGGAGGGTCGTCGGGGACCGCGTCGCTTGAGGATAGCAGGGGTTGGTGG<br>AATGCCCGGTGTAGTGGTGAAATGCGTAGGGATCGGGCAGAACACCCGTGGCGAAGGCGGCCAACTGGGCTATTCCTGACGCTGAGGTACGAAGGCGTGGGG |
| P-asv6943 | GCAAGCGTTGTCCGGATTTATTGGGCGTAAAGAGCTCGTAGGCGGTTAGACACGTCGGATGTGAAAACTGGTGGCTCAACCACCAGCCTGCATTCGATACGGTCTGACTTGAATGTGGTAGGGGTAACTGGAATT<br>CCTGGTGTAGCGGTGAAATGCGCAGATATCAGGAGGAACACCGATGGCGAAGGCAGGTCTCTGGGCTGTAACTGACGCTGAGAAGCGAAAGCATGGGG |
| P-asv6944 | GCAAGCGTTATCCGGATTTATTGGGCGTAAAGCGCGTCTAGGCGGAAAAGTAAGTCTGATGTGAAAATGCGGGGCTCAACTCCGTATTGCGTTGGAAACTGCTTTTCTAGAGTACTGGAGAGGTGGGCGGAACTA<br>CAAGTGTAGAGGTGAAATTCGTAGATATTTGTAGGAATGCCGATGGAGAAGTCAGCTCACTGGACAGATACTGACGCTGAAGCGCGAAAGCGTGGGG |
| P-asv6945 | GCAAGCGTTAATCGGAATTACTGGGCGTAAAGCGTGCGCAGGCGGTTTTGTAAGTCTGATGTGAAATCCCCGGGCTCAACCTGGGAACTGCATTGGAGACTGCAAGGCTAGAGTGTGTCAGAGGGGGGTAGAAT<br>TCCACGTGTAGCAGTGAAATGCGTAGAGATGTGGAGGAATACCGATGGCGAAGGCGACTAGCTGGGCCATAACTGACGCTGAGGAACGAAAGCGAGGGG |
| P-asv6946 | GCTAGCGTTGTCCGGAATCACTGGGCGTAAAGGGTTCGCAGGCGGAAATGCAAGTCAGATGTAAAAGGCAGTAGCTTAACTACTGTAAGCATTTGAAACTGCATATCTTGAGAAGAGTAGAGGTAAGTGGAATTTT<br>TAGTGTAGCGGTGAAATGCGTAGATATTAAAAAGAATACCGGTGGCGAAGGCGACTTACTGGCCCGGTACTGACGCTGAGGCACGAAAGCGTGGGG |

| | |
|---|---|
| P-asv6947 | CCAAGCGTTGTTCGGATTCACTGGGCGTAAAGGGTGCGTAGGCGGTAGGGTAAGTTTGAGGTGAAATCTCCGGGCTCAACCCGGAAAATGCCTTGAAGACTATCCTGCTAGAGGATCGGAGGGGAGACTGGAATTCTCGGTGTAGCAGTGAAATGCGTAGATATCGAGAGGAACACCAGTGGCGAAGGCGAGTCTCTGGACGATTCCTGACGCTGAGGCACGAAAGCCAGGGG |
| P-asv6948 | GCAAGCGTTATCCGGAATCACTGGGTTTAAAGGGTGCGTAGGTGGACCAATAAGTCAGAGGTGAAAGCCCGTCGCTTAACGACGGAACTGCCTTTGATACTGTTGGTCTTGAATCAGGTTGAGGTATGCGGAATGTGGCATGTAGCGGTGAAATGCGTAGATATGCCATAGAACACCGATTGCGAAGGCAGCATACTGGCCCTGTATTGACACTGAGGCACGAAAGCGTGGGG |
| P-asv6949 | GCAAGCGTTATCCGGAATTATTGGGCGTAAAGCGCGCGTAGGCGGTTTTTTAAGTCTGATGTGAAAGCCCACGGCTCAACCGTGGAGGGTCATTGGAAACTGGAAAACTTGAGTGCAGAAGAGGAAAGTGGAATTCCATGTGTGGCGGTGAAATGCGCAGAGATATGGAGGAACACCAGTGGCGAAGGCAATCCCCTGGACCTGTACTGACGCTCATGCACGAAAGCGTGGGG |
| P-asv6950 | GCAAACGTTATTCGGAATCACTGGGCTTAAAGGGTGCGTAGGCGGCTATGAAGGTGGGATGTGAAATCCCTCAGCTCAACTGAGGAACTGCGTCCCAAACCACACAGCTCGAGGAAGGCAGGGGTGAGTGGAACTGTAGGTGGAGCGGTGAAATGCGTTGATATCTACAGGAACACCGGTGGCGAAAGCGACTCACTGGGCCTTTTCTGACGCTGAGGCACGAAAGCTAGGGG |
| P-asv6951 | GCAAGCGTTGTCCGGAATTATTGGGCGTAAAGAGCTCGTAGGCGGTTTGTCGCGTCTGCTGTGAAAACCCGAGGCTCAACCTCGGGCCTGCAGTGGGTACGGGCAGACTAGAGTGTGGTAGGGGAGACTGGAATTCCTGGTGTAGCGGTGGAATGCGCAGATATCAGGAGGAACACCAATGGCGAAGGCAGGTCTCTGGGCCACGACTGACGCTGAGGAGCGAAAGCGTGGGG |
| P-asv6952 | GCTAGCGTTGTTCGGAATTACTGGGCGTAAAGCGCACGTAGGCGGTGACGTAAGTCGGGGGTGAAAGCCCGGGGCTTAACCCTGGAATTGCCTTCGAGACTGCGTCGCTGGAGTATGGAAGAGGAGAGCGGAATTCCCAGTGTAGAGGTGAAATTCGTAGATATTGGGAAGAACACCGGTGGCGAAGGCGGCTCTCTGGTCCATTACTGACGCTGAGGTGCGAAAGCGTGAGG |
| P-asv6953 | GCAAGCGTTAATCGGAATTACTGGGCGTAAAGCGCGCGTAGGCGGACTGGTGAGTCAGGGGTGAAATCCCGAGGCTTAACCTCGGAACTGCCTTTGATACTGCCAGTCTCGAGTCCGGAAGAGGTGAGTGGAATTCCTAGTGTAGAGGTGAAATTCGTAGATATTAGGAAGAACACCGGTGGCGAAGGCGGCTCACTGGTCCGGTACTGACGCTGAGGCGCGAAAGCGTGGGG |
| P-asv6954 | GCGAGCGTTGTCCGGATTTATTGGGCGTAAAGGGCATGTAGGTGGTCTTTTAAGTCAGTTGTTAAAACTATCGGCTTAACCGGTAATTGCAATTGATACTGGAGGACTTGAATTTGAGAGGGGATAGTGGAATATCAGGTGTAGCGGTGGAATGCGTAGATATTTTATAGAACACCCGAGGTGCAGACGACTATCTGGCTCATGATTGACACTGAGATGCGAAAGTATGGGG |
| P-asv6955 | GCTAGCGTTGCTCGGAATCACTGGGCGTAAAGGGTGCGTAGGTGGATCTTTAAGTCAGGGGTGAAATCCTGGAGCTCAACTCCAGAACTGCCTTTGATACTGAGGATCTTGAGTTCGGGAGAGGTAAGTGGAACTGCGAGTGTAGAGGTGAAATTCGTAGATATTCGCAAGAACACCAGTGGCGAAGGCGGCTTCCTAGTCCAACATTGACATTCAGGTGCGAAAGCGTGGGG |
| P-asv6956 | GCAAGCGTTGTTCGGAATTATTGGGCGTAAAGGGCATGTAGGCGGTTTCGTAAGCCTGGTGTGAAAGGCTGCAGCTTAACTGTAGCACTGTATTGGGTACTGCGAGACTTGAATCACGGAGGGGAAACTAGAATTCCAGGTGTAGGGGTGGAATCTGTAGATATCTGGAAGAATACCGGTGGCGAAGGCGGGTTTCTGGCCGATGATTGACGCTGAGGTGCGAAAGTGCGGGG |
| P-asv6957 | GCTAGCGTTATTCAGAATCATTGGGCGTAAAGGGAGCGCAGGCGGGCATGTAAGTCTTTCTTAAAAGTGCGGGGCTCAACCCCGTGATGGGAAAGAAACTATGTGTCTTGAGTACAGGAGAGGAAAGCGGAATTCCCAGTGTAGCGGTGAAATGCGTAGATATTGGGAGGAACACCAGTGGCGAAGGCGGCTTTCTGGACTGCAACTGACGCTGAGGCTCGAAAGCCAGGGG |
| P-asv6958 | GCGAGCGTTGATCGGAATTACTGGGCGTAAAGCGTGCGCAGGCGGTTTCGTAAGCTGGAGGTGAAATCCCCGGGCTTAACCATGGAAGTGCGCAGGAAACTGATGGACTTGAGTGCCGGAGAGGGTCGCGGAATTCCCGGTGTAGAGGTGAAATTCGTAGATATCGGGAGGAACACCAGTGGCGAAGGCGGCGACCTGGACGGACACTGACGCTGAGACGCGAAAGCGTGGGG |
| P-asv6959 | GCGAGCGTTGTCCGGATTTATTGGGTTTAAAGGGTGCGTAGGCGGTTTATTAAGTCAGTGGTGAAAGACGGTCGCTCAACGATTGCAGTGCCATTGATACTGGTAGACTTGAGTTCGTATGAGGTAGCTGGAATGGATAGTGTAGCTGTGAAATGCATAGATATTATCCAGAACTCCAATTGCGCAGGCAGGTTACTAATACGATACTGACGCTGATGCACGAAAGTGTGGGG |
| P-asv6960 | CCAAGCGTTATCCGGATTTATTGGGTTTAAAGGGTGCGTAGGCGGGTTTGTAAGTCAGTGGTGAAATCCTGCGGCTCAACCGTAGCACTGCCATTGAAACTGCAGATCTTGAATACAGATGAGGTAGGCGGAATATAGTATGTAGCGGTGAAATGCTTAGATATACTATAGAACACCGATTGCGAAGGCAGCTTACTAGGCTTGTATTGACGCTGATGCACGAAAGCGTGGGT |
| P-asv6961 | GCAAGCGTTAATCGGAATCACTGGGCGTAAAGGGCGCGTAGGCGGTACGTTAGGTCTGTCGTGAAATCTCCAGGGCTCAACCCGGAAACTGCTTCGCATACTGGCAGACTGGAGGGATGCAGAGGTTTGTGGAATTCCTGGTGTAGCGGTGATATGCATAGATATCAGGAGGAACACCCATGGCGAAGGCAGCAAACTGGGCATCATCTGACGCTGAGGCGCGAAAGCGTGGGT |
| P-asv6962 | GCAAGCGTTATTCGGAATTATTGGGCATAAAGCGCATGTAGGTGGTTTGATAAGTCTGATGTGAAATCCCCGGACTCAATTCGGGAAGTGCATTGGATACTGTCAGGCTAGAGTATGGGAGAGGTGAGCGGAATTCCAGGTGTAGAGGTGAAATTCGTAGATATCTGGAGGAACAACAGAGGCGAAGGCGGCTCACTGGACCATTACTGACACTGAGATGCGAAAGCGTGGGG |
| P-asv6963 | CCAAGCGTTGTTCGGATTCACTGGGCGTAAAGGGTGCGTAGGTGGCCGGGTAAGTTTGACGTGAAATCTCCGGGCTTAACTCGGAAAATGCGTTGAATACTATCCGGCTGGAGGTTCGGAGGGGGGACTGGAATTCTCGGTGTAGCAGTGAAATGCGTAGATATCGAGAGGAACACCAGTGGCGAAGGCGAGTCCCTGGACGACACCTGACACTGAGGCACGAAAGCCGGGGG |
| P-asv6964 | GCTAGCGTTGCTCGGAATTACTGGGCGTAAAGGGAGCGTAGGCGGACTGTTTAGTCAGAGGTGAAATCCCGGGGCTTAACCTCGGAACAGCCTTTGATACTGCCAGTCTAGAGTCCGGCAGAGGTGAGTGGAATTCCTAGTGTAGAGGTGAAATTCGTAGATATTAGGAAGAACACCAGTGGCGAAGGCGGCTCACTGGGCCGAGACTGACGCTGAGGCGCGAAAGCGTGGGG |
| P-asv6965 | GCAAGCGTTATCCGAAATCATTGGGCGTAAAGCGCCTGTAGGTTGTTTATTAAGTCTGTTGTTAAAGCCTAGGGCTTAACCCTAGAACAGCAATGGAAACTGGTAAACTTGAGTATGGCAGGGGTAGAGGGAATTTCTAGTGTAGCGGTGAAATGCGTAGATATTAGAAAGAACACCAGTGGCGAAAGCGCTCTACTGGACCATTACTGACACTCAGAGGCGAAAGCTAGGGT |
| P-asv6966 | GCAGGCGTTATCCGGAATTATTGGGCGTAAAGCGTCTGTAGGCGATCTAGCAAGTTTTCTGTGAAATTCCAATGCTCAACATTGGGGCTGCAGAAAATACTGCTAGAATTGAGGAAGATAGAGGTAAGTGGAATTCCCGGTGTAGCAGTGAAATGCGTTAATATCGGGAGGAACACCAATAGCGAAGGCAGCTTACTGGATCTTTCCTGACGCTCAGAGACGAAAGCGTGGGT |
| P-asv6967 | GCGAGCGTTGTCCGGAATCACTGGGCGTAAAGGGTCCGCAGGCGGATAAACAAGTCAAGAGTAAAAGGCATGGGCTTAACCCATGTAAGCTATTGAAACTGTTTATCTTGAGATGTGGAGAGGTAAGTGGAATTACTAGTGTAGCGGTGAAATGCGTAGATATTAGTAGGAATACCAGTAGCGAAGGCGACTTACTGGACACAAACTGACGCTGAGGAACGAAAGCGTGGGT |
| P-asv6968 | GCAAGCGTTGTCCGGATTTATTGGGCGTAAAGCGAGCGCAGGCGGAAGAATAAGTCTGATGTGAAAGCCCTCGGCTTAACCGAGGAACTGCATCGGAAACTGTTTTTCTTGAGTGCAGAAGAGGAGAGTGGAACTCCATGTGTAGCGGTGGAATGCGTAGATATTCGGAAGGACACCAGTGGCGAAGGCGGCTCACTGGACTGGTATTGACGCTGAAGTGCGAAAGCGTGGGG |
| P-asv6969 | GCTAGCGTTGCTCGGAATTACTGGGCGTAAAGGGCGCGTAGGCGGACTGTTTAGTCAGAGGTGAAAGCCCAGGGCTCAACCTTGGAATTGCATTCGATACTGGGAAGCTAGAGTATGGGAGAGGATGGTAGAATTCCAGGTGTAGCGGTGAAATGCGTAGAGATCTGGAGGAATACCGATGGCGAAGGCAGCCATCTGGCCTAATACTGACGCTGAGGTACGAAAGCATGGGG |
| P-asv6970 | GCAAGCGTTATCCGGAATCATTGGGTTTAAAGGGTCCGTAGGTGGATTGGTCAGTCAGAGGTGAAATCCTGCAGCTCAACTGTAGAACTGCCTTTGATACTGCCGGTCTTGAATTATTGTGAAGTGGTTAGAATATGTAGTGTAGCGGTGAAATGCATAGATATTACATAGAATACCGATTGCGAAGGCAGATCACTAACAATATATTGACACTGATGGACGAAAGCGTGGGG |
| P-asv6971 | GCAAGCGTTACCCGGATTTACTGGGTGTAAAGGGCGTGTAGGCGGTCTCTCAAGTCCGATGCTAAAGACCAAGGCTCAACCTCGGGAGTGCGTTGGATACTGTGAGGCTAGACGGTCGGAGAGGGTAGCGGAATTTCCGGAGTAGCGGTGGAATGCGCAGATACCGGAAGGAACGCCAATAGCGAAAGCAGCTACCTGGACGATTTGTGACGCTGAGGCGCGAAAGCGTGGGG |
| P-asv6972 | GCGAGCGTTATCCGGAATCACTGGGTTTAAAGGGTGCGCAGGCGGCGTGCTAAGTCAGGGGTGAAAGGCCACCGCTCAACGGTGGGGCTGCCTCTGATACTGGCATGCTCGAACGGGCGCGAGGCGGGCGGAATGTGGCATGTAGCGGTGAAATGCTTAGATATGCCATAGAACACCGATTGCGAAGGCAGCCCGCCAGAGCCTAGTTGACGCTCATGCACGAAAGCGTGGGG |
| P-asv6973 | GCTAGCGTTGTTCGGAATTACTGGGCGTAAAGCGCACGCAGGCGGTGCGGTCAGTTGGATGTGAAATCCCAGGGCTCAACCCTGGAATTGCATTCGATACTGCCGCGCTGGAGGACGGAAGAGGAGAGTGGAATTCCCAGTGTAGAGGTGAAATTCGTAGATATTGGGAAGAACACCAGTGGCGAAGGCGGCTCTCTGGTCCGTTTCTGACGCTCATGTGCGAAAGCGTGGGG |
| P-asv6974 | GCTAGCGTTGCTCGGAATTACTGGGCGTAAAGGGTGCGTAGGCGGGTCTTTAAGTCAGGGGTGAAATCCTGGAGCTCAACTCCAGAACTGCCTTTGATACTGAAGATCTTGAGTTCGGGAGAGGTGAGTGGAACTGCGAGTGTAGAGGTGAAATTCGCAGATATTCGGAGGAACACCAGTGGCGTAGGCGGCCAACTGGACGGGAACTGACGCTGAGGTACGAAAGCGTGGGG |
| P-asv6975 | GCAAGCGTTGTCCGGATTTACTGGGTGTAAAGGGCGTGTAGGCGGAGCAGCAAGTCAGATGTGAAATCTCCGGGCTTAACCCGGAAACTGCATTTGAAACTGTTGCCCTTGAGTATCGGAGAGGCAGGCGGAATTCCTAGTGTAGCGGTGAAATGCGTAGATATTAGGAAGAACACCAGTGGCGAAGGCGGTCTGCTGGACGACAACTGACGCTGAGGCGCGAAAGCGTGGGG |
| P-asv6976 | GCTAGCGTTGTTCGGAATTACTGGGCGTAAAGCGCACGTAGGCGGCTTTGTAAGTCAGAGGTGAAAGCCTGGAGCTCAACTCCAGAACTGCCTTTGAGACTGCATCGCTTGAATCCAGGAGAGGTGAGTGGAATTCCAGGTGTAGCGGTGATATGCATAGATATCAGGAGGAACACCCATGGCGAAGGCAGCAAACTGGGCATCATCTGACACTGAGGCACGAAAGCGTGGGG |
| P-asv6977 | GCAAGCGTTGTTCGGAATTACTGGGCATAAAGCGTGCGTAGTCGGTATTGAGAGTCACGGGTGAAATCCCGGAGCTCAACTTCGGAACTGCCTTTGATACTGGCGATCTTGAGTGTGAGAGAGGTATGTGGAACTCCGAGTGTAGAGGTGAAATTCGTAGATATTCGGAAGAACACCAGTGGCGAAGGCGACATACTGGCTCATTACTGACGCTGAGGCGCGAAAGCGTGGGG |
| P-asv6978 | GCAAGCGTTAATCGGAATTACTGGGCGTAAAGGGTGCGTAGGTGGTTATATAAGTTATCTGTGAAATCCCCGGGCTTAACCTGGGCAGGTCAGATAAGACTGTGTGACTAGAGTATGGGAGAGGGTAGTGGAATTTCCGGTGTAGCGGTGAAATTCGTAGATATTCGCAAGAACACCAGTGGCGAAGGCGGCTCACTGGCCCGGTACTGACGCTGAGGCACGAAAGCGTGGGG |
| P-asv6979 | GCAAGCGTTAATCGGAATTACTGGGCGTAAAGCGCACGTAGGCGGTTTATTAAGTCGGATGTGAAAGCCCTGGGCTCAACCTGGGAACTGCACTCGATACTGGTAAACTAGAATTTGGGAGAGGTGAGTGGAATTTCCGGTGTAGCGGTGAAATGCATAGAGATCGGAAGGAACACCAGTGGCGAAGGCGACTCACTGGACCAAAATTGACGCTGAGGTGCGAAAGCGTGGGT |
| P-asv6980 | GCAAGCGTTGTCCAGAATTATTGGGCGTAAAGAGCTCGTAGGCGGTTTGTCGCGTCTGTCGTGAAAGCCCACGGCTTAACCGTGGGTCTGCGGTGGATACGGGCAGACTAGAGGCAGGTAGGGGAGAATGGAATTCCCGGTGTAGCGGTGAAATGCGCAGATATCGGGAGGAACACCGGTGGCGAAGGCGGTTCTCTGGGCCTGTACTGACGCTGAGGAGCGAAAGCGTGGGG |
| P-asv6981 | GCAAGCGTTAATCGGAATCACTGGGCGTAAAGGGTGCGTAGGCGGTGCATTAGGTCTGTCGTGAAATCCCCGGGCTCAACTTGGGGATTGCATTGGTGACTGCACGGCTAGAGTGTGTCAGAGGGGGGTAGAATTCCACGTGTAGCAGTGAAATGCGTAGAGATGTGGAAGAATACCGATGGCGAAGGCAGCCCCCTGGGATAACACTGACGCTCATGCACGAAAGCGTGGGG |
| P-asv6982 | GCTAGCGTTGCTCGGAATCACTGGGCGTAAAGGGTGCGTAGGCGGCGTTTTAAGTCGGGGGTGAAAGCCTGTGGCTCAACCCCGGAACTGCCTTTGAAACTAGATGACTGGAATCTTGGAGAGGCGAGTGGAATTCCGAGTGTAGAGGTGAAATTCGTAGATATTCGGAAGAACACCAGTGGCGAAGGCGACTCGCTGGACAAGTATTGACGCTGAGGTGCGAAAGCGTGGGG |
| P-asv6983 | GCTAGCGTTGCTCGGAATCACTGGGCGTAAAGGGCGCGTAGGCGGCGTTTTAAGTCGGGGGTGAAAGCCTGTGGCTCAACCACAGAATGGCCTTCGATACTGGGACGCTTGAGTATGGTAGAGGTTGGTGGAACTTCGAGTGTAGAGGTGAAATTCGTAGATATTCGCAAGAACACCGGTGGCGAAGGCAGCCCCCTGGGATAACACTGACGCTCATGCACGAAAGCGTGGGG |
| P-asv6984 | GCAAGCGTTATCCGGAATCATTGGGTTTAAAGGGTCCGTAGGCGGACTGATAAGTCAGGGGTGAAATCCCACAGCTCAACTGTGGAACTGCCTTTGATAGTGTTAGTCTTGAGTTATATGGAAGTAGATAGAATATGTAGTGTAGCGGTGAAATGCTTAGAGATTACATAGAATACCGATTGCGAAGGCAGTCTACTACGTATATACTGACGCTAATGGACGAAAGCGTGGGG |
| P-asv6985 | CCGAGTGGTAGTCGTTATTATTGGGCTTAAAGTGTTCGTAGCCTGCCGTGTAAGTCCGCTGTGAAACCCACGCGCTCAACGTGTGGATGTGCGGCGGATACTGCATGGCTAGGGACCGGACGGCGTCGAGGCTATTCCAGGAGTAGGAGTGAAATCCAAAAATTCCTGGAGGAGCACCTGCGGCGAAAGCACTCTGAGAAGACGGCTTCGACAGTGAGGGACGAAAGCTAGGGT |

| | |
|---|---|
| P-asv6986 | GCAAGCGTTAATCGGAATTACTGGGCGTAAAGCGCGCGTAGGTGGTTAGTTAAGTTGGACGTGAAATCCCAGGGCTCAACCCTGGAACGGCCTCCAAAACTCCTGGTCTTGAGTTCGAGAGAGGTGAGTGGAAT<br>TCCGAGTGTAGAGGTGAAATTCGTAGATATTCGGAGGAACACCAGTGGCGAAGGCGGCTCACTGGCTCGATACTGACGCTGAGGTGCGAAAGTGTGGGG |
| P-asv6987 | TCAAGTGATGTCCAATATTATTGGGCCTAAAGCGTCCGTAGCTGGCCGCGCAAGTCCGTTGGGAAATCCGCGCGCTTAACGCGCGGGCGTCCAGCGGAAACTGCCTGGCTTGGGACCGGAAGGCTCAGAGAGT<br>ACGTCCGGGGTAGGAGTGAAATCCCGTAATCCCGGACGGACTACCGATGGCGAAAGCACTCTGAGAAGACGGCTCCGACAGTGAGGGACGAAAGCTCGGGT |
| P-asv6988 | GCTAGCGTCGTTCGGAATTACTGGGCGTAAAGGGCGCGTAGGCGGCTCTGTAAGTCAGATGTGAAAGCCCCGGGCTTAACCTGGGAACTGCATTTGATACTGCAGGGCTAGAGATCGAGAGAGGAAAGTGGAAT<br>TACGAGTGTAGAAGTGAAATTCGTAGATATTCGTAGGAACACCGGTGGCGAAGGCGACTTTCTGGCTCGATACTGACGCTAAAGCGCGAAAGCGTGGGG |
| P-asv6989 | TCGAGTGATGACCGATTTTATTGGGCCTAAAGCGTCCGTAGCTGGCCGCGCAAGTCCGTCGGGAAATCCACCCGCTTAACGGGTGGGCGTCCGGCGGAAACTGTGCGGCTTGGGACAGGAAGGCTCGAGGGGT<br>ACGTTCGGGGTAGGAGTGAAATCCTAAAATCCTGAGAGGAGCACCTGTGGCGGAAGCGCTCGACGAGTACGGATCCGACGGTGAGGAACGAAAGCTAGGGG |
| P-asv6990 | GCGAGCGTTAATCGGAATTACTGGGCGTAAAGCGAGTGTAGGTGGCTCATTAAGTCACATGTGAAATTCCCGGGCTTAACCTGGGAACTGCATGTGATACTGGTGGTGCTAGAATATGTGAGAGGGAAGTAGAAT<br>TCCAGGTGTAGCGGTGAAATGCGTAGAGATCTGGAAGAATACCGATGGCGAAGGCAGCTTCCTGGCATAATATTGACACTGAGATTCGAAAGCGTGGGT |
| P-asv6991 | GCAAGCGTTAATCGGAATTATTGGGCGTAAAGCGCGCGTAGGCGGCCGCATAAGTCTGATGTGAAAGCCCAGGGCTCAACCCTGGAAAGCCATTGGAAACTGTGCGGCTTGAGAACAGGAGGGGTAGGTGGAA<br>TTCCTGGTGTAGGAGTGAAATCCGTAGAGATCAGGAGGAACACCGGTGGCGAAGGCGACCTACTGGACTGGATCTGACGCTGAGGCGCGAAAGCGTGGGG |
| P-asv6992 | GCAAACGTTGTTCGGAATCACTGGGCTTAAAGGGCGCGTAGGCGGAGGACCAAGTCCGGGGTGAAATCTTTCGGCTCAACCGGAAAACTGCCTTGGATACTGGTCTTCTTGAGGGAGGTAGGGGCAGGTGGAA<br>CTTCCGGTGGAGCGGTGAAATGCGTTGATATCGGAAGGAACGCCGGTGGCGAAAGCGGCCTGCTGGATCTCTTCTGACGCTGAGGCGCGAAAGCTAGGGG |
| P-asv6993 | CCGAACGTTATTCGGATTCACTGGGCATAAAGGGCGCGTAGGCGGGCTGCCAAGTCAAGGGTGAAATCCTCCGGCTTAACCGGAGAACTGCCTTTGATACTGGTGGTCTTGAGGGAGGTAGGGGCGTGTGGAA<br>TTAAGGGTGGAGCGGTGAAATGCGTTGATATCCTTAGGAACTCCGGTGGCGAAGGCGACACGCTGGACCTTTTCTGACGCTGAGGCGCGAAAGCCAGGGG |
| P-asv6994 | ACAAGCGTTAATCGGAATTACTGGGCGTAAAGCGTGCGTAGACGGTTACATAAGTCGGGTGTGAAAGCCCCGGGCTCAACCTGGGAATTGCATTCGATACTGGGAAGCTAGAGTATGGGAGAGGATGGTAGAAT<br>TCCAGGTGTAGCGGTGAAATGCGTAGAGATCTGGAGGAATACCGATGGCGAAGGCAGCCATCTGGCCTAATACTGACGCTGAGGTACGAAAGCATGGGG |
| P-asv6995 | GCAAGCGTTGTTCGGAATTACTGGGCGTAAAGGGCGCGTAGGCGGCTTTTCAAGTCAGCCGTGAAATCTCCGGGCTTAACCTGGAAGGGTCGGTTGATACTGTCGAGCTAGAGTGTGGAAGAGGCAACTGGAAT<br>TCCGTGTGTAGCGGTGAAATGCGTAGATATACGGAGGAACACCTGCGGCGAAGGCGGGTTGCTGGGCCAACACTGACGCTGAGGCGCGAAAGCTAGGGT |
| P-asv6996 | GCAAGCGTTACTCGGAATCACTGGGCGTAAAGAGCGTGTAGGCGGATTTGTAAGTTGGAAGTGAAATCCTATGGCTCAACCATAGAACTGCTTCCAAAACTGCTCATCTAGAGTGTGGGAGAGGTAGATGGAATT<br>TCTGGTGTAGGGGTAAAATCCGTAGAGATGTGGAAGAATACCGATGGCGAAGGCAGCCCCCTGGCCAATACTGACGCTCATGCACGAAAGCGTGGGG |
| P-asv6997 | GCGAGCGTTGTTCGGAATCACTGGGCGTAAAGCGTGCGTAGGCGGTCTTGTAAGACAGAAGTGAAAGCCTGGAGCTCAACTCCAGAATTGCTTTTGTGACTGCAAGGCTTGAATCCAGGAGAGGTTGGGGGAAT<br>TCCGAGTGTAGAGGTGAAATTCGCAGATATTCGGAGGAACACCAGTGGCGTAGGCGCCCAACTGGACTGGTATTGACGCTGAGGCACGAAAGCGTGGGG |
| P-asv6998 | GCGAGCGTTGTTCGGAATTATTGGGCGTAAAGGGTTCGTAGGAGGTACATTAAGATTGGTGTGAAATCCCAAGGCTCAACCTTGGAACTGCATTGATTACTGATGGACTAGAGTATCGTAGAGGAGAGCGGAATT<br>CCAGGTGTAGCGGTGGAATGCGTAGATATCTGGAAGAACACCGACAGCGAAGGCAGCTCTCTGGGCGAGAACTGACTCTGAGGAACGAAAGCGTGGGT |
| P-asv6999 | TCCCTCTTTCCTCTTCCTTCCTTTTCTTAAAGCGCGCATAGGTGGTTTGTTAAGTTGAATGTGAAAGCCCCGGGCTCAACCTGGGAACTGCATCCAAAACTGGCAAGCTAGAGTATGGCAGAGGGTGGTGGAATTT<br>CCTGTGTAGCGGTGAAATGCGTAGATATAGGAAGGAACACCAGTGGCGAAGGCGACCACCTGGGCTAATACTGACACTGAGGTGCGAAAGCGTGGGG |
| P-asv7000 | GCGAGCGTTACCCGGATTTACTGGGCGTAAAGGGCGTGTAGGCGGCCTGGGGCGTCCCATGTGAAAGGCCACGGCTCAACCGTGGAGGAGCGTGGGATACGCTCAGGCTAGAGGGTGGGAGAGGGTGGTGG<br>AATTCCCGGAGTAGCGGTGAAATGCGCAGATACCGGGAGGAACGCCGATGGCGAAGGCAGCCACCTGGTCCACTTCTGACGCTGAGGCGCGAAAGCATGGGG |
| P-asv7001 | TCGAGCGTTATCCGGAATTATTGGGCGTAAAGGGTCCGTAGGTGGTTTTATTAGTCAGTCGTGAAAGATCCGTGCTCAACACGGGAACCGCGATTGAAACGGTAAGACTTGAGTGTGCGAGAGGTGAACAGAACT<br>CATGGTGTAGGGGTGAAATCCGTTGATATTATGGGGAATACCAAAAGCGAAGGCAGTTCACTGGCGCATTACTGACACTGAGGGACGAAACCCTGGGT |
| P-asv7002 | GCTAGCGTTGTTCGGAATTACTGGGCGTAAAGCGCACGTAGGCGGCTTTTTAAGTCGGATGTGAAATCCCTGAGCTTAACTTAGGAATTGCATTTCGAACTGGCAAACTAGAGTCTTGTAGAGGGGGGTAGAATTC<br>CAGGTGTAGCGGTGAAATGCGTAGAGATCTGGAGGAATACCGGTGGCGAAGGCGGCCCCCTGGACAAAGACTGACGCTCAGGCACGAAAGCGTGGGG |
| P-asv7003 | GTTAGCGTTGTTCGGAATTACTGGGCGTAAAGCGCGCGTAGGCGGATTGGAAAGTTGGGGGTGAAATCCCAGGGCTCAACCTGGGAACTGCATTAGTGACTGCATAGCTGGAGTACGGCAGAGGGGGATGGAA<br>TTCCGCGTGTAGCAGTGAAATGCGTAGATATGCGGAGGAACACCGATGGCGAAGGCAATCCCCTGGGCCTGTACTGACGCTCATGCACGAAAGCGTGGGG |
| P-asv7004 | GCGAGCGTTAATCGGGATAACTGGGCGTAAAGGGCACGCAGGCGGTGACTTAAGTGAGGTGTGAAAGCCCCGGGCTTAACCTGGGAATTGCATTCGATACTGGGAAGCTAGAGTATGGGAGAGGATGGTAGAA<br>TTCCAGGTGTAGCGGTGAAATGCGTAGAGATCTGGAGGAATACCGATGGCGAAGGCAGCCATCTGGCCTAATACTGACGCTGAGGTACGAAAGCATGGGG |
| P-asv7005 | GCAAGCGTTAATCGGAATTACTGGGCGTAAGGCGCGCGTAGGTGGTTTGTTAAGTTGAATGTGAAAGCCCCGGGCTCAACCTTGGAACTGCCTTTGAAACTATCAGTCTGGAGTTCGAGAGAGGTGAGTGGAATT<br>CCGAGTGTAGAGGTGAAATTCGTAGATATTCGGAGGAACACCAGTGGCGAAGGCGGCTCACTGGCTCGATACTGACGCTGAGGTGCGAAAGCGTGGGG |
| P-asv7006 | GCTAGCGTTGCTCGGAATCACTGGGCGTAAAGGGTGCGTAGGCGGGTCTTTAAGTCAGAGGTGAAAGGCATTGGCTCAACCGATGTAAGCCTTTGAAACTGTTGGGCTTGAGTGCAGGAGAGGTAAGTGGAATT<br>CCTAGTGTAGAGGTGAAATTCGTAGATATTAGGAGGAACACCAGTGGCGAAGGCGACTTACTGGACTGTAACTGACGCTGAGGCACGAAAGCGTGGGG |
| P-asv7007 | GCTAGCGTTGCTCGGAATCACTGGGCGTAAAGGGTGCGTAGGCGGGTCTTTAAGTCAGGGGTGAAATCCTGGAGCTCAACTCCAGAACTGCCTTTGATACTGAAGATCTTGAGTATGGGAGAGGTGAGTGGAAC<br>TGCGAGTGTAGAGGTGAAATTCGTAGATATTCGCAAGAACACCAGTGGCGAAGGCGATCTACTGGAACATTACTGGCGCTCAGATGCGAAAGCGTGGGG |
| P-asv7008 | GCTAGCGTTGTTCGGAATTACTGGGCGTAAAGCGCACGTAGGCGGACCATTAAGTCAGGGGTGAAATCCCGGGGCTCAACCTGGGAACGGCATTTCGAACTGTCAGGCTAGAGTGTGGTAGAGGGTAGTGGAAT<br>TTCCTGTGTAGCGGTGAAATGCGTAGATATAGGAAGGAACACCAGTGGCGAAGGCGGCTACCTGGACCAACACTGACACTGAGGTGCGAAAGCGTGGGG |
| P-asv7009 | GCGAACGTTATTCGGAATCACTGGGCTTAAAGAGCACGTAGGCGGCTATGTAAGTTGGAGGTGAAATCCCATCGGCTCAACCGTGGAACTGCCTTCAATACTGCATGGCTTGAGGGAGGTAGGGGCACGTGGAA<br>CTTCCAGTGGAGCGGTGAAATGCGTTGAGATTGGAAGGAACGCCGGTGGCGAAAGCGACGTGCTGGATCTCTTCTGACGCTGAGGTGCGAAAGCTAGGGT |
| P-asv7010 | CCAAGCGTTGTTCGGAATCATTGGGCGTAAAGCGGGAGTAGGTGGTTGGATATGTCAGAAGTGAAAGCCCTGGGCTTAACCCAGGAAGTGCTTTTGAAACTATTCAACTTGAGTGCCGGAGAGGTTGCTAGAATA<br>CCTGGTGTAGTGGTGAAATACGTAGATATCAGGTGGAATTCCGGAGAGGAAGCTGGGTAACTGGCCGGACACTGACACTCATCCCCGAAAGCGTGGGG |
| P-asv7011 | ACAAGTGGAGATCATGATTACTGGGTCTAAAGTGTTCGTAGCTGGCTTGTTAAGTCTCTTGTGAAAGTTCTGTGGCTCAACTCAGAAATTGCAAGAGATACTGGCAGGCTTGGGACCGGGAGAGACAAGAAGTATT<br>GCTTGGGTAGCGGTAAAATGCTATAATCCTAGCAAGACTAACGAAGGCGTAGGCATCTTGTTTGAACGGATCTGACAGTGAGGAACGAGAGCTGGGGG |
| P-asv7012 | GCAAGCGTTATCCAGATTTATTGGGTTTAAAGGGTGCGCAGGCGGCTTTATAAGTCAGTGGTGAAATCTTCCCGCTCAACGGGAAAACTGCCATTGATACTGTAGAGCTAGAGTACAGATGGGGTAGGCGGAATG<br>TAGCATGTAGCGGTGAAATGCTTAGATATGCTACAGAACACCAATCGCGAAGGCAGCTTACCAAGCTGGTACCGACGCTCAGGCACGAAAGCGTGGGT |
| P-asv7013 | GCGAGCATTATTCGGAATGATTAGGCGTAAAGGGTTTGTAGGTGGTTTTTTAAGTTGAAAAAGACGAATTAAAGCTCAACTTTATAAATTTTTTCAAAACTGAAAAACTTGAGTATACATAGAGGATAATGGAATTTCT<br>ATTGGAGGGATAAAATATGTTGATAATAGAAGGAAGGTCGATAGCGAAGGCAATTATCTGGGTATATACTGACACTGAAAAACGAAAGCTTGGGT |
| P-asv7014 | GCTAGCGTTATTCGGAATTACTGGGCGTAAAGCGCACGTAGGCGGACTGGAAAGTCAGAGGTGAAATCCCAGGGCTCAACCTTGGAACTGCCTTTGAAACTCCCAGTCTTGAGGTCGAGAGAGGTGAGTGGAAT<br>TCCGAGTGTAGCAGTGAAATGCGTAGAGATGTGGAGGAATACCGATTGCGAAGGCGATCTACTGGAACATTATTGACGCTGAGACGCGAAAGCGTGGGG |
| P-asv7015 | CCGAGCGTTGTCCGGAGTTACTGGGCGTAAAGCGCGCGCAGGCGGCGGTGTAAGCATGGCGTGAAAGCCCCAGGCTCAACCTGGGAGGGTCGTCGTGGACTGCACCGCTTGAGGGCGGTAGGGGCTGGTGG<br>AATGCCTGGTGTAGTGGTGAAATGCGTAGAGATCAGGCGGAACACCCGTGGCGAAGGCGGCCAGCTGGGCCGTCCCTGACGCTGAGGCGCGAAGGCGTGGGG |
| P-asv7016 | CCGAGCGTTAGGCGGAGTCACTGGGCTTAAAGGGTGTGTAGGCGGATCAGTAAGCATTTTGTGAAATCCCTCGGCTCAACCGAGGAATTGCAGAGTGAACTGCTGATCTTGAGGTAGGCAGAGGTCATCGGAAC<br>GTACGGTGGAGCGGTGAAATGCGTAGATATCGTACGGAACGCCGAAGGGGTAGCCAGGTGACTGGGCCTTACCTGACGCTGAGGCACGAAAGCGTGGGG |
| P-asv7017 | GCGAGCGTTAATCGGAATTACTGGGCGTAAAGCGCATGCAGGTGGTTTGTTAAGTCAGATGTGAAAGCCCGGGGCTCAACCTCGGAATAGCATTTGAAACTGGCAGACTAGAGTACTGTAGAGGGGGGTAGAAT<br>TTCAGGTGTAGCGGTGAAATGCGTAGAGATCTGAAGGAATACCGGTGGCGAAGGCAGTCCCCTGGGCCTGCACGGACGCTCATGCACGAAAGCGTGGGG |
| P-asv7018 | GCAAGCGTTGTCCGGATTTATTGGGTTTAAAGGGTGCGTAGGCGGCCCCTTAAGTCAGTGGTGAAAGCCCGCGGCTCAACCGCGGAACTGCCATTGATACTGGGGGGCTTGAGTACAGACGAGGTAGGCGGAA<br>TTGAAGATGTAGCGGTGAAATGCTTAGATATCTTCAAGAACCCCGATTGCGTAGGCAGCTTACTAGGCTGTAACTGACGCTGAGGCACGAAAGCGTGGGG |
| P-asv7019 | CCGAGCGTTATCCGGATTTATTGGGTTTAAAGGGTGCGTAGGCGGGATAATAAGTCAGGGGTGAAATCCTGGAGCTCAACTCCAGAACTGCCTTTGATACTGAAGATCTTGAGTTCGGGAGAGGTGAGTGGAACT<br>GCGAGTGTAGAGGTGAAATTCGTAGATATTCGCAAGAACACCAGTGGCGAAGGCGGCTCACTGGCCCGATACTGACGCTGAGGCACGAAAGCGTGGGG |
| P-asv7020 | GCAAGCGTTATCCAAATTGATTGGGCGTAAAGGGTTTGTAGGCGGTATTTTTTCGCGGTTTGCAAAAAAATGAGATTTAATCTTGTTATGACATTCTGTACTTTAATACTTGAGTCTAATTGAGGATAACAGAATTCTA<br>TAGGGAAGAGTAAAATCTATTAATCTATGGAGGAATGCCAGAGGCGAAAGCGGTTATCTGGGTTAGTACTGACGCTGAGAAGCGAAAGCGTAGGG |
| P-asv7021 | GCGAGCGTTGTCCGGAATTACTGGGCGTAAAGGGCGCGTAGGCGGCCGATTAAGTTAGATGTGAAAAACCGGGGCTCAACTCCGGGATTGCATTTAAGACTGGTCGGCTTGAGGGTGAGAGAGGAAAGTGGAA<br>TTCCTAGTGTAGCGGTGAAATGCGTAGATATTAGGAGGAACACCAGTGGCGAAGGCGACTTTCTGGCTCATACCTGACGCTGAGGCGCGAAAGCGTGGGG |
| P-asv7022 | GCTAGCGTTATTCGGAATTACTGGGCGTAAAGGGCGCGTAGGCGGTCTTGCAAGCCTGACGTGAAAGCCTGGGGCTTAACTCCAGAACTGCGTTGGGAACTGCATGACTTGAATCATGGAAGGGAAACTGGAAT<br>TCCAGGTGTAGGGGTGAAATCTGTAGATATCTGGAAGAACACCGGTGGCGAAGGCGAGTTTCTGGCCAATGATTGACGCTGAGGCGCGAAAGTGCGGGG |
| P-asv7023 | GCTAGCGTTGTTCGGAATTACTGGGCGTAAAGCGCACGTAGGCGGTTGACACAGTCAGATGTGAAAGCCCCGGGCTCAACCCGGGAATAGCATTTGATACTGGTCGACTCGAGGTCGGAAGAGGAGAGTGGAAT<br>TCCCAGTGTAGAGGTGAAATTCGTAGATATTGGGAAGAACACCAGTGGCGAAGGCGGCTCTCTGGTCCGATACTGACGCTGAGGTGCGAAAGCGTGGGG |
| P-asv7024 | GCGAACGTTACTCGGAATCACTGGGCTTACTGGGTGCGTAGGTGGCTTGTTAAGCAGGGTGTGAAATCCCTGGGCTCAACCGAGGAACTGCGCTCTGAACTGGCGAGCTGGAGTATGCTAGGGGTGTATGGAAC<br>ATCTAGTGGAGCGGTGAAATGTGTTGATATTAGATGGAACACCGGAGGCGAAGGCGATACACTGGGGCATAACTGACACTGAGGCACGAAAGCCAGGGG |

| | |
|---|---|
| P-asv7025 | GCGAGCGTTAATCGGATTTACTGGGCGTAAAGCGTGCGTAGGCGGCTTTTTAAGTCGGATGTGAAATCCCTGAGCTTAACTTAGGAATTGCATTCGATACTGGGAAGCTAGAGTATGGGAGAGGATGGTAGAATTCCAGGTGTAGCGGTGAAGTGCGTAGATATCAGGAGGAATACCTGTGGCGAGAGCGCTCTGCTGGATGGTTCCTGACGCTGAGGCACGAAAGCGTGGGT |
| P-asv7026 | GCAAGCGTTGTTCGGATTTACTGGGCGTAAAGCGCGCGCAGGCGGATCTGCAAGTCAGATGTGAAATCTCGGGGCTTAACTCCGAAATTGCGTCTGAAACCGCAAATCTAGAGTGTCGGAGAGGGATGGGGAATTTCCTGTGTAGGGGTAAAATCCGTAGAGATAGGAAGGAGCACCAGAGGCGAAGGCGCCATCCTGGAAGATAACTGACGCTGAGGCGCGAAAGCGTGGGG |
| P-asv7027 | CCTAGCGTTGTTCGGAATTATTGGGCATAAAGCGGGTGTAGGCGGACTTATAAGTCAGGTGTGAAATCCCTGAGCTTAACTTAGGAATTGCATTCGATACTGGGAAGCTAGAGTATGGGAGAGGATGGTAGAATTCCAGGTGTAGCGGTGAAATGCGTAGAGATCTGGAGGAATACCGATGGCGAAGGCAGCCATCTGGCCTAATACTGACGCTGAGGTACGAAAGCATGGGG |
| P-asv7028 | GCAAGCGCTGTCCGGAATCACTGGGCGTAAAGGGAGCGTAGGCGGGAAATTAAGTCAGATGTGAAAGTTTGCGGCTCAACCGTAAAATTGCATTTGATACTGGTTTTCTAGAGTATCGGAGGGGGAAGCGGAATTCCTTGTGTAGCGGTGAAATGCGCAGATATAAGGAAGAACACCAATGGCGAAGGCAGCTTTCTGGACGATAACTGACGCTGATGCTCGAAGGTGTGGGG |
| P-asv7029 | GCGAGCGTTAATCGGATTTACTGGGCGTAAAGCGTGCGTAGGCGGCTTTTTAAGTCGGACGTGAAAGCCTTGGGCTCAACCTAGGAATCGCATTTGAAACTGACAAGCTAGAGTACTGTAGAGGGGGGTAGAATTTCAGGTGTAGCGGTGAAATGCGTAGAGATCTGAAGGAATACCGGTGGCGAAGGCGGCCCCCTGGACAGATACTGACACTCAGATGCGAAAGCGTGGGG |
| P-asv7030 | GCTAGCGTTACTCGGAATTACTGGGCGTAAAGGGCGCGTAGGCGGCGTTCCAAGTTAGGCGTGAAAGTCCTGGGCTCAACCTGGGAACTGCGCTTAAGACTGGAGTGCTAGAGGATGGAAGAGGGTTGTGGAATTCCCAGTGTAGAGGTGAAATTCGTAGATATTGGGAAGAACACCGGTGGCGAAGGCGGCAACCTGGTCCATTTCTGACGCTGAGGCGCGATAGCGTGGGG |
| P-asv7031 | GCGAGCGTTATTCGGAATCGCTGGGCGTAAAGAGTGCGTAGGCGGTTAGATAAGTTAGGAGTGAAATCCCGGGGCTCAACCTCGGAATTGCTTTTAAAACTATTTAACTAGAATTCGGTATAGGATAGTGGAATTATCAGTGTAGCAGTGAAATGCGTAGATATTGATAGGAACACCAGTGGCGTAGGCGACTATCTGGACCGACATTGACGCTGAGGCACGAAAGTGTGGGG |
| P-asv7032 | GCAAGCGTTAATCGGAATTACTGGGCGTAAAGCGTGCGCAGGCGGTTTTGTAAGACAGGCGTGAAATCCCCGGGCTCAACCTGGGAATTGCGCTTGTGACTGCAAGGCTGGAGTGCGGCAGAGGGGGATGGAATTCCGCGTGTAGCAGTGAAATCCTGTAATCCCGGACGGACTACCGATGGCGAAAGCACTCTGAGAAGACGGCTTCGACAGTGAGGGACGAAAGCTAGGGT |
| P-asv7033 | CCAAGTTGCATCCTATTATTGGGTCTAAAACATCCGTAGCTCGTTTATTAAGTCTCTTGTGAAATCCCGTGTCTCAAGCGCGGGGCGAGCGGGAGATACTGACAAACTAGAGATCGGTAGACGTAAGGAGTACGCTGGAGGTAGTGGTTAAATACGTTAATCTTCGGTGGACTGACAATTTAGCGAAGGCACCTTACGAGGACGAGTCTGACAGTGAGGGATGAAGGCCAGGGG |
| P-asv7034 | CCAAGCGTTAATCAGAATTACTGGGCGTAGGGCGCACGTAGGCGGTTTTTAAGTCAGATGTGAAAGCCCCGGGCTCAACCTGGGAATTGCATTTAAAACTGTCCAGCTAGAGTCTTGTAGAGGGGGGTAGAATTCCAGGTGTAGCGGTGAAATGCGTAGAGATCTGGAGGAATACCGGTGGCGAAGGCGGCCCCCTGGACAAAGACTGACGCTCAGGTGCGAAAGCGTGGGG |
| P-asv7035 | GCGAGCGTTAATCGGAATTACTGGGCGTAAAGCGTGCGCAGGCGGTTTTTTAAGATAGGCGTGAAATCCCCGGGCTCAACCTGGGAACTGCGCTTATGACTGGAAGACTCGAGTATGGCAGAGGGGGGTGGAATTCCACGTGTAGCAGTGGAATGCGTAGATATTCGGAAGAACACCAGTGGCGAAGGCGGCTCACTGGACTGGTATTGACGCTGAGGTGCAAAAGCGTGGGG |
| P-asv7036 | GCAAGCGTTGCTCGGATTCATTGGGTGTAAAGGGCAAGTAGGCGGTTTGGTAAGTCATATGTGAAATCCCCGGGCTCAACCTGGGAACTGCATCTGAGACTGCCGGACTTGAGGACGGAAGAGGAGAGTGGAATATCCGGTGTAAGGGTGAAATCTGTAGATATCGGATAGAACACCAGTGGCGAAGGCGACTCTCTGGTCCGTTTCTGACGCTGAGAGGCGAAAGCTAGGGG |
| P-asv7037 | GCAAGCGTTATCCGGTTTTATTGGGTTTAAAGGGTGCGTAGGCGGCTTATTAAGTCAGTTGTGAAATCCTAGCGCTCAACGTTAGAACTGCGATTGATACTGATAGGCTTGAGTCTAGAAGAGGTAGGTAGAATTTATGGTGTAGTAGTGAAATGCTTAGATATCATGAGGAATACCAATAGCGTAGGCAGCTTACTGGTCTTGAACTGACGCTGAGGCACGAAAGCGTGGGG |
| P-asv7038 | CCAAGCGTTATCCGGATTTATTGGGTTTAAAGGGTCCGTAGGCGGGTGATTAAGTCAGAGGTGAAATCCTGCAGCTCAACTGTAGCATTGCCTTTGATACTGGGTATCTTGAGTATGGAAGAGGTGAGTGGAATTCCGAGTGTAGAGGTGAAATTCGTAGATATTCGGAGGAACACCAGTGGCGAAGGCGGCTCACTGGTCCATTACTGACGCTGAGGTGCGAAAGCGTGGGG |
| P-asv7039 | GCAAGCGTTGTCCGGATTTATTGGGTTTAAAGGGTGCGTAGGCGGCTTGTTAAGTCAGCGGTGAAAGTTTTCGGCTCAACCGGAAAAATGCCGTTGATACTGATGAGCTTGAGTCACGGAAGGGTACATGGAATTGGTGGTGTAGCGGTGAAATGCATAGATATAGGAAGGAACACCAGTGGCGAAGGCGACCACCTGGACTGATACTGACACTGAGGTGCGAAAGCGTGAGG |
| P-asv7040 | TCGAGCGTTGTCCGGAGTTACTGGGCGTAAAGCGCGTGCAGGTGGCTCATTACGCCCGGCGTGAAAGCCCCCGGCTCAACCGGGGAGGGTCGTCGGGGACGGGTGAGCTTGAGGGTCGCAGGGGCTGGTGGAATTCCCGGTGTAGTGGTGAAATGCGTAGAGATCGGGAGGAACACCCGTGGCGAAGGCGGCCAGCTGGGCGACACCTGACACTGAGACGCGAAGGCGTGGGG |
| P-asv7041 | GCAAGCGTTGTCCGGATTTATTGGGCGTAAAGAGTTTGTAGGCGGCTTTGTAAGTCTGAAGTGAAAGAATGGAGCTCAACTCCATCCACGCTTCTGAAACTGCAAGGCTAGAGAATGTTAGGGGTGAGTGGAATTTCCAGTGTAGCGGTGAAATGCGTAGATATTGGATAGAACACCGGAGGCGATAGCGGCTCACTGGGACATTTCTGACGCTGAGAAACGAAAGCTAGGGT |
| P-asv7042 | GTTAGCGTTGTTCGGAATTACCGGGCGTAAAGCGCGCGTAGGCGGACTGGAAAGTTGGGGGTGAAATCCCGGGGCTCAACTCCAGAACTGCCTTTGATACTGAAGATCTTGAGTTCGGGAGAGGTGAGTGGAACTGCGAGTGTAGAGGTGAAATTCGTAGATATTCGCAAGAACACCAGTGGCGAAGGCGGCTCACTGGCCCGATACTGACGCTGAGGCACGAAAGCGTGGGG |
| P-asv7043 | GCGAGCGTTAATCGGATTTACTGGGCGTAAAGGGTCCGTAGGCGGCCCGAAAAGTCGGATGTGAAATCCCACAGCTCAACTGTGGAACTGCATTCGAAACTCTCGGGCTTGAGTATGAGAGGGGTAAAGGGAATTCCCGGTGTAAGGGTGAAATCTGCAGAGATCGGGAGGAACACCAGTCGCGAAGGCGCTTTACTGGCTCATTACTGACGCTCAGGGACGAAAGCTAGGGT |
| P-asv7044 | GCTAGCGTTAATCGGAATTACTGGGCGTAAAGCGTGTGTAGGTGGTTAGATAAGTTAGATGTGAAATCCCCGGGCTTAACCTGGGAATTGCGTTTAAGACTGTTTAGCTTGAGTACTGTAGAGGATAGTGGAATTTCCAGTGTAGCGGTGAAATGCGTAGATATTGGAAGGAACACCAGTGGCGAAGGCGACTATCTGGATAGATACTGACACTGAGACACGAAAGCGTGGGG |
| P-asv7045 | CCGAGCGTTGTCCGGAATTATTGGGCGTAAAGAGCTCGTAGGCGGCGTGCCGCGTCTGCTGTGAGAGACCGGGGCTTAACTCCGGTTCTGCAGTGGATACGGGCATGCTAGGGGTAGGTAGGGGAGACTGGAATTCCTGGTGTAGCGGTGAAACGCGCAGATATCAGGAGGAACACCGGTGGCGAAGGCGGGTCTCTGGGCCTTACCTGACGCTGAGGAGCGAAAGCATGGGG |
| P-asv7046 | GCGAGCGTTGTCCGGAATCACTGGGCGTAAAGGGCGCGTAGGTGGCTTGGTAAGTAGGGGGTGAAATCTTGCGGCTCAACCGCAAGACTGCCTTCTAAACTGCCTGGCTTGAGCACAGTAGAGGCAGGTGGAATTCCCGGTGTAGCGGTGGAATGCGTAGAGATCGGGAAGAACATCGGTGGCGAAGGCGGCCTGCTGGGCTGTTGCTGACACTGAGGCGCGACAGCGTGGGG |
| P-asv7047 | GCAAGCGTTAATCGGAATTACTGGGCGTAAAGCGCGCGTAGGTGGTTTGTTAAGTTGGGGGCGAAAGCCCGGGGCTCAACCTCGGAAATGCCCTCAATACTGGCTGTCTTGAGTACGGGAGAGGTGAGTGGAACTCCGAGTGTAGAGGTGAAATTCGTAGATATTCGGAAGAACACCAGTGGCGAAGGCGACTCACTGGCCCGTTACTGACGCTGAGGCGCGAAAGCGTGGGG |
| P-asv7048 | GCGAGCGTTGTCCGGAATTATTGGGCGTAAAGGGCGCGTAGGCGGCTTGGTAAGTCTGCTGTGAAACCTTGGGGCTCAACCCCAAGCGTGCAGTGGATACTGTCAAGCTAGAGGATGGTAGAGGCCAGTGGAATTCCCAGTGTAGCGGTGAAATGCGCAGATATTGGGAGGAACACCAGTAGCGAAGGCGGCTGGCTGGGCCATTCCTGACGCTGAGGCGCGAAAGCTAGGGG |
| P-asv7049 | GCTAGCGTTGTTCGGAATCACTGGGCGTAAAGGGCGCGTAGGCGGCCTTGTAAGTTAGATGTGAAAGCCCTGGGCTTAACCCGGGAATTGCATTTAATACTGCAAAGCTTTGAGTAAGGTAGGGGATAGCGGAATACCCAGTGTAGAAGTGAAATTCGTAGATATTGGGTGGAACACCTATGGCGAAGGCAGCTATCTGGACCTTTACTGACGCTGAGGCGCGAAAGCGTGGGG |
| P-asv7050 | GCGAGCGTTATCCGGATTTATTGGGTTTAAAGGGTGCGTAGGCGGCGATTTAAGTCGGTGGTGAAAGGTAGCAGCTTAACTGTTTTACATGCTTCCGATACTGAATTGCTTGAGTTACCAAGAGGCAAGTAGAATTTCCGGTGTAGCGGTGAAATGCATAGATACCGGAAGGAATACCGATTGCGAGGGCCACTTGCTATAGGTAAACTGACGCTGATGCACGAAAGCGTGGGG |
| P-asv7051 | GCAAGCGTTGTCCGGATTTACTGGGTGTAAAGGGTGCGCAGGCGGGACTGCAAGTTGGATGTGAAATACCGAGGCTTAACCTCGGAACTGCATCCAAAACTGTGGTTCTTGAGTGGAGTAGAGGTAAGCGGAATTCCGAGTGTAGCGGTGAAATGCGTAGATATTCGGAGGAACACCAGTGGCGAAGGCGGCTTACTGGGCTCTAACTGACGCTGAGGCACGAAAGTGTGGGT |
| P-asv7052 | GCTAGCGTTATTCGGAATTACTGGGCGTAAAGCGCACGTAGGCGGCTTTGTAAGTTAGAGGTGAAAGCCTGGAGCTCAACTCCAGAATTGCCTTTAAGACTGCATCGCTTGAATCCAGGAGAGGTGAGTGGAATTCCGAGTGTAGAGGTGAAATTCGTCGAAACTGGAAGGAACACCAAGGGCGAAAGCAATTCTCGACAATATTCTGACGTTGAGGCACGAAGGGATGGGT |
| P-asv7053 | GCAAGCGTTATTCGGATTTACTGGGCGTAAAGGGCGCGTAGGCGGCCTGATAAGTCAGATGTGAAAGCCCGGGGCTCAACCCCGGAAGTGCATTTGAAACTGTTGGGCTTGAGTATGGTAGAGGGAAGTGGAATTCCAGGTGTAGCGGTGAAATGCGTAGATATCTGGAGGAACACCGGTGGCGAAGGCGACTTCCTGGACCAAAACTGACGCTGAGGCGCGAAGGTGTGGGT |
| P-asv7054 | GCGAGCGTTATTCGGATTTACTGGGTGTAAAGGGCGTGTAGGTGGTTGGGTGTGTTGGATGTGAAAACCTTGGGCTCACCCCAAGAACGGCATCTAAAACTACCCGACTTGAGTGTTGGAGGGGAAAGTGGAATTCCTGGTGTAGCGGTGAAATGCGTAGATATCAGGAGGAACGCCGGTGGCGAAAGCGACTTTCTAGCCAAACACTGACACTGAGGCGCGAAAGCCAGGGG |
| P-asv7055 | TCGAGCGTTGCGCGGAATCACTGGGCTTAAAGGGTGCGTAGGCGGATGTACAAGTGCCTTGTGAAATCCCCGGGCTCAACCCGGGAACTGCTTGGTAGACTGTATGTCTTGAGGTCGGTAGGGGTTGCTGGAACAGTTGGTGGAGCGGTGAAATGCGTAGATATCAGGAGGAACACCGATGGCGAAGGCAGCCCCCCTGGGCCTGTACTGACGCTCATGCACGAAAGCGTGGGG |
| P-asv7056 | GCGAGCGTTAATCGGATTTACTGGGCGTAAAGCGTGCGTAGGCGGTCTTGTGCGTCGAAGGTTAAAGGCCATGGCTCAACCATGGAAATGCCCTCGATACGGCAAGACTTGAGTATGAGAGAGGAAGATGGAATTCCTGGTGTAGCAGTGAAATGCGTAGATATCAGGAGGAACACCGGTGGCGAAGGCGGTCTTCTGGCTCATTACTGACGCTGAGGCACGAAAGTGTGGGG |
| P-asv7057 | GCAAGCGTTAATCGGAATTACTGGGCGTAAAGCGCACGTAGGCGGTTTTTTAAGTCAGATGTGAAAGCCCCGGGCTCAACCTGGGAATTGCATTTGAAACTGGAAAACTAGAGTGTGTGAGAGGGGGGTAGAATTCCAAGTGTAGCGGTGAAGTGCGTAGAGATTTGGAGGAACACCGATGGCGAAGGCACTTTACTGGGCTATTACTGACACTCAGAGACGAAAGCTAGGGT |
| P-asv7058 | CCAAGCGTTATCCGGATTTATTGGGCGTAAAGAGCTCGTAGGCGGTTTGGTAAGTCGGGTGTGAAAACTCTGGGCTCAACCCAGAGAGGCCACCCGATACTGCTATGACTAGAGTACGGTAGGGGAGCGGGGAATTCATGGTGTAGCGGTGAAATGCGCAGATATCATGAGGAACACCAGCGGCGAAGGCGCCGCTCTGGGCCGTAACTGACGCTGAGGAGCGAAAGCATGGGT |
| P-asv7059 | GCGAGCGTTATTCGGAATCACTGGGCGTAAAGCGTGCGTAGGCGGTTTTATAAGTTGGAAGTGAAAGCCCAGGGCTCAACCCTGGAATTGCTTTCAAAACTACAAGACTAGAAATCGGGAGAGGATAGCGGAATTGTCAGTGTAGCAGTGAAATGCGTAGATATTGACAGGAACACCAGTGGCGTAGGCGGCTATCTGGACCGACTTTGACGCTGAGGCACGAAAGCGTGGGG |
| P-asv7060 | GCTAGCGTTGTTCGGAATTACTGGGCGTAAAGCGCACGTAGGCGGACTGGAAAGTTGGGGGTGAAATCCCGGGGCTCAACCTCGGAACTGCCTTCAAAACTATCAGTCTGGAGTTCGAGAGAGGTGAGTGGAATTGCGAGTGTAGAGGTGAAATTCGTAGATATTCGCAGGAACACCAGTGGCGAAGGCGGCTCACTGGCTCGATACTGACGCTGAGGTGCGAAAGTGTGGGG |
| P-asv7061 | GCGAACGTTATTCGGAATCACTGGGCTTAAAGGGTGCGTAGGCGGCTTGAAGCGCCAGGTGTGAAATTTTCCGGCTCAACCGGGAAACTGCGCCTGGGACGATCAGGCTTGAGTCGATCGGGGGTGTGCGGAACTGCCAGTGGAGCGGTGAAATGTGTTGAGATTGGCAGGAACGCCGGTGGCGAAAGCGGCACACTGGGATCGTTCTGACGCTGAGGCACGAAAGCTAGGGG |
| P-asv7062 | GCGAGCGTTAATCGGAATTACTGGGCGTAAAGGGTGCGTAGGTGGTTTGATAAGTTGGCTGTGAAATACCGGGGCTTAACCTTGGGGCGTCAGCCAAGACTGTCAGGCTTGAGTATAGGAGAGGGCAGTGGAATTTCCGGTGTAGCGGTGAAATGCGTAGAGATCGGAAGGAACACCAGTGGCGAAGGCGTCTGCCTGGCCTAATACTGACACTGAGGCACGAAAGCGTGGGT |
| P-asv7063 | CCGAGTGGTCGGGACGATTATTGGGCCTAAAGCATCCGTAGCTTGTTCTGCAAGTCTTCCGTTAAATCCACCTGCTCAACAGATGGGCTGCGGGAGATACTACAGAGCTAGGAGGCGGGAGAGGCAAGCGGTACTCAGTGGGTAGGGGTAAAATCCTCTGATCCACTGAAGACCACCAGTGGCGAAGGCGGCTTGCCAGAACGCGCTCGACGGTGAGGGATGAAAGCTGGGGG |

| | |
|---|---|
| P-asv7064 | GCAAGTGTTACTCATAATAACTGGGCGTAAAGGGTACGTAGATGGTTTATATGGTTACTTTGTGAAAGAATAATGTAATGGTTATTTTGCTAGTAATACGTGTAAACTAGAGTATGACAGAGGAAAGTGGAATTTTTAATGTAAAGGTGGTATTTATCGATATTAAAAAGACGATCTAAGGCGAAGGCAACTTTCTGGGTTATTACTGACATTGAGGTACTGAAGCATGGGG |
| P-asv7065 | GCAAGTGTTATCCGGATTTATTGGGTTTAAAGGGTGCGTAGGCGGGGTATTAAGTCAGAGGTGAAATCCAGTCGCTTAACGATTGAACTGCCTTTGATACTGGTTGACTTGAGTCATATGGAAGTAGATAGAATGTGTAGTGTAGCGGTGAAATGCTTAGATATTACACAGAATACCGATTGCGAAGGCAGTCTACTACGTATGTACTGACGCTCATGGACGAAAGCGTGGGG |
| P-asv7066 | GCTAGCGTTGTTCGGAATTACTGGGCGTAAAGCGAGCGTAGGTTGCGCCGCAAGTCAGGGGTGAAATCCCGGGGCTCAACCTCGGAACTGCCCCTGAGACTGTTGGTCTAGAGTCTGTGAGAGGATGACGGAATACCCAGTGTAGAGGTGAAATTCGTAGATATTGGGTAGAACACCGGTGGCGAAGGCGGTCATCTGGCGCAGCACTGACGCTGAGGCGCGAAAGCGTGGGG |
| P-asv7067 | GCAAGCGTTAATCGGAATTACTGGGCGTAAAGCGCACGTAGGTGGTTTGTTAAGCCAGCTGTGAAATCCCCGGGCTTAACCTGGGAACTACGTTTGTGACTGCAAGGCTAGAGTACGGCAGAGGGGGGTGGAATTCCTGGTGTAGCAGTGAAATGCGTAGATATCAGGAGGAACACCGATGGCGAAGGCAGCCCCCTGGGCCTGTACTGACGCTCATGCACGAAAGCGTGGGG |
| P-asv7068 | GCAAGCGTTGTCCGGAATCATTGGGCGTAAAGCGCGTGTAGGCGGTTACGTAAGTCTGCTGTGAAAGTCGGAGGCTCAACCTTCGAATGCCAGTGGATACTGCGTGACTAGAGTTCGGAAGAGGCAAGTGGAATTCCTGGTGTAGCGGTGAAATGCGCAGATATCAGGAGGAACACCAACGGCGAAGGCAGCTTGCTGGTACGACACTGACGCTGAGACGCGAAAGCGTGGGG |
| P-asv7069 | GCAAGCGTTGTCCGGAATTATTGGGCGTAAAGCGCGCGCAGGCGGGTTTTTAAGTCTGTCTTAAAAGTTCGGGGCTTAACCCCGTGAGGGGATGGAAACTGGAAACCTAGAGTATCGGAGAGGAAAGCGGAATTCCTAGTGTAGCGGTGAAATGCGTAGATATTGGAAAGAACACCGATGGCGAAGGCACTTTACTGGGCTATTACTGACACTCAGAGACGAAAGCTAGGGT |
| P-asv7070 | GCGAGCGTTAATCAGATTTACTGGGCGTAAAGCGTGCGTAGGCGGCTTTTTAAGTCGGATGTGAAATCCCTGAGCTTAACTTAGGAATTGCATTTGATACTGCAAGGCTAGAGATCGAGAGAGGAAAGTGGAATTACGAGTGTAGAAGTGAAATTCGTAGATATTCGTAGGAACACCAGTGGCGAAGGCGACTTTCTGGCTCGATACTGACGCTAAAGCGCGAAAGCGTGGGG |
| P-asv7071 | GCTAGCGTTGTTCAGAATTACTGGGCGTAAAGCGCACGTAGGCGGCTTTTTAAGTCAGGGGTGAAATCCCGGGGCTCAACCCCGGAACTGCCTCCAAAACTATCAGTCTGGAGTTCGAGAGAGGTGAGTGGAATTGCGAGTGTAGAGGTGAAATTCGTAGATATTCGCAGGAACACCAGTGGCGAAGGCGGCTCACTGGCTCGATACTGACGCTGAGGTGCGAAAGTGTGGGG |
| P-asv7072 | TCAAGCGTTATCCGGAATTACTGGGCGTAAAGGGTGTGTAGGCGGTCGTAATTAGTCTTTCGTTAAAGCCCCGGGCTTAACTCGGGATCTGCGAAAGAAACGGTACGACTAGAGAATGCGAGGGGTATGCGGAACTCAAAGTGTAGGGGTGAAATCCGTTGATATTTTGGAGAACACCAAAAGCGAAAGCAGCATACTGGCGCATTTCTGACGCTGAGACACGAAAGCGTGGGT |
| P-asv7073 | CCGAGCGTTATCCGGATTTATTGGGCGTAAAGGGCGCGTAGGCGGCTTGGTGCGTCTCATGTTAAATCTCACTGCTCAACGGTGAAATCGCATGAGATACGATCAAGCTTGAGAGTGTTAGAGGTAGATGGAACTCATGGTGTAGGGGTGACATCCGTTGATATCATGGGGAACACCAAAGGCGAAGGCATTCTACTGGGACATTTCTGACGCTGAGGCGCGAAAGCGTGGGG |
| P-asv7074 | GCAAGCGTTAATCGGAATTACTGGGCGTAAAGCGCACGTAGGTGGTTTGTTAAGCCAGCTGTGAAATCCCCGGGCTTAACCTGGGAACTACGTTTGTGACTGCAAGGCTAGAGTACGGCAGAGGGGGGTGGAATTCCTGGTGTAGCAGTGAAATGCGTAGATATCAGGAAGAACACCGATGGCGAAGGCAGCCCCCTGGGCCTGTACTGACGCTCATGCACGAAAGCGTGGGG |
| P-asv7075 | TCGAGCGTTGTTCGGATTTACCGGGCGTAAAGGGAGCGTAGGCGGTTTGGTGTGTCGAATGTGAAAGCCCGCAACTCAACTGCGGAATGGCATTCGAAACTACCAGGCTAGAGTACTGGAGAGGAGAGCGGAATTCTTGGTGTAGCGGTGAAATGCGTAGATATCAAGAGGAACACCGGTGGCGAAGGCGGCTCTCTGGACAGATACTGACGCTGAGGCTCTAAAGTTAGGGG |
| P-asv7076 | GCAAGCGTTGTCCGGAATCATTGGGCGTAAAGGGCGTGTAGGTGGTTTCGTAAGTCGAATGTGAAATCTCACGGCTCAACCGTGAAATTGCGTTCGAAACTACGAGACTTGAGTTAGGGAGAGGTGTACGGAATTCCTGGTGTAGCGGTGGAATGCGTAGATATCAGGAAGAACACCAATGGCGAAGGCAGTACACTGGCCTGTAACTGACACTGAGGCGCGAAAGCGTGGGG |
| P-asv7077 | GCGAACGTTGTTCGGAATCACTGGGCTTAAAGGGCGCGTAGGCGGCTCATCAAGTCGGGGGTGAAATCTTTCAGCTTAACTGGAAAAGTGCCTTCGATACTGGTGGGCTAGAGGGAGGTAGGGGTCTGTGGAACTTCTGGTGGAGCGGTGAAATGCGTTGATATCAGAAGGAACGCCGGTGGCGAAAGCGACAGACTGGACCTCTTCTGACGCTGAGGCGCGAAAGCTAGGGG |
| P-asv7078 | CCAAGCGTCATCCGGATTTATTGGGTTTAAAGGGTTCGTAGGCGGAATATTAAGTCAGTGGTGAAATCCTACAGCTCAACTGTAGAACTGCCATTGATACTGGAGACCTTGAGTGTACTAGAGGTAGGCGGAATTTATGGTGTAGCGGTGAAATGCATAGATACCATAAAGAACACCGATAGCGTAGGCAGCTTACTGGAGTACAACTGACGCTGAGGCACGAAAGCATGGGG |
| P-asv7079 | GCAAGCATTAATCGGAATTATTGGGCGTAAAGGGCGTGTAGGCGGATCTGTAAGTCAGATGTGAAATTTCGGGACTCAACCCCGGAGCCGCATTTGAGACTGCAGGTCTAGAGGATAGGCGGAGAAAACGGAATTCCACATGTAGCGGTGAAATGCGTGGATATGTGGAAGAACACCGGTGGCGAAGGCGGTTTACTGGGCTATTACTGACACTCTGAGACGAAAGCTAGGGT |
| P-asv7080 | GCGAGCGTTAATCGGATTTACTGGGCGTAAAGCGTGCGTAGGCGGCTTTTTAAGTCGGGTGTGAAATCCCTGAGCTTAACTTAGGAATTGCATTCGATACTGGGAAGCTAGAGTATGGGAGAGGATGGTAGAATTCCAGGTGTAGCGGTGAAATGCGTAGATATCGGGAGGAACACCAGTGGCGAAGGCGACTTTCTGGCACACTCCTGACGCTGAGGAGCGAAAGCGTGGGT |
| P-asv7081 | GCAAGCGTTGTTCGGATTTACTGGGTGTAAAGGTCATGTAGGCGTTCCGTTAAGTCGAGTGTGAAATCCTATGGCTCAACCATAGAACTGCGCTCGATACTGGCGGAATTGAGTACGGGAGAGGTAAGCGGAATTCCAGAAGTAAGGGTGAAATCTGTAGATATCTGGAGGAACACCATTAGCGAAGGCGGCTTACTGGTCCGATACTGACGCTCAAATGAGAAAGCTAGGGG |
| P-asv7082 | GCAAGCGTTAATCGGAATGACTGGGCGTAAAGGGCGCGTAGGTGGATATGTAAGTAAATTGTGAAATACCCGGGCTTAACCTGGGGCGGTCAATTTAAACGGCATATCTGGAGTGGAGTAGAGGGAAGTAGAATTTCCGGTGTAGCGGTGAAATGCGTAGATATCGGAAGGAATACCGGTGGCGAAGGCGTCTTCCTGGACTCTAACTGACACTGAGGCGCGAAAGCATGGGG |
| P-asv7083 | CCGAGCGTTATCCGGATTTATTGGGTTTAAAGGGTGCGTAGGCGGAAATTTAAGTCAGTGGTGAAAGTTTAGGGCTCAACCTTAAAATTGCCATTGAAACTGGGTTTCTTGAGTACGGTAGAGGTAGGCGGAATTTGTGGTGTAGCGGTGAAATGCTTAGATATCACGAAGAACTCCGATTGCGAAGGCAGCTTACCGGACCGTAACTGACGCTGAGGCACGAAAGTGTGGGG |
| P-asv7084 | GCTAGCGTTGTTCGGAATTACTGGGCGTAAAGCGCACGTAGGCGGCTTTGTAAGTCAGAGGTGAAAGCCTGGAGCTCAACTCCAGAACTGCCTTTGAGACTGCATCGCTTGAATCCAGGAGAGGTGAGTGGAATTCCGAGTGTAGGGGTGAAATGCGTAGATATCTGGAGGAACATCAGTGGCGAAGGCGACACCCTGGACTCATACTGACACTGAGGTGCGAAAGCGTGGGG |
| P-asv7085 | GCTAGCGTTATTCGGAATTACTGGGCGTAAAGGGTGCGTAGGCGGGTCTTTAAGTCAGGGGTGAAATCCTGGAGCTCAACTCCAGAACTGCCTTTGGTACTGGCGAGCTAGAGTACGGTAGAGGGTGGTGGAATTCCCGGTGTAGCGGTGAAATGCGTAGAGATCGGGAGGAACATCTGTGGCGAAGGCGGCCACCTGGACCAGTACTGACGCTGAGGCACGAAAGCGTGGGG |
| P-asv7086 | GCGAGCGTTAATCGGAATTACTGGGCGTAAAGGGTGCGTAGGCGGTTCGTCGTGTCTGGTGTGAAATCCCCGGGCTCAACCTGGGAATGGCATTGGAAACTGGCGGACTAGAGTGTGGCAGAGGGAGGTGGAATTCCGCGTGTAGCAGTGAAATGCGTAGAGATGCGGAGGAACACCGATGGCGAAGGCAGCCTCCTGGGCTAACACTGACGCTGAGGCACGAAAGCGTGGGG |
| P-asv7087 | CCAAGCGTTATCCGGAATTACTGGGCGTAAAGCGTCTGTAGGTGGTTTCAAAGGTTTCGTGTGAAACTTCAGGGCTCAACCCTGAATTGTGCGGAAAACCATGGAACTTGAGTGTGGGAGAGGCTAGCAGAATGGTAAGAGTAGGGGTGCAATCCGTTGATACTTACCAGAATACCAAAAGCGAAGGCAGCTAGCTGGAACATTACTGACACTGAGAGACGAAAGCGTGGGG |
| P-asv7088 | GCGAGCGTTGTCCGGATTTACTGGGCGTAGAGAGCACGTAGGCGGTCTGTTAAGTGAGAAGTGAAATCTCCAGCGCTCAACGCGGAAACGGCTTTTCATACTGGCAGACTTGAGGGATGCAGAGGCAAGTAGAATTCCTGGTGTACCGGTGACATGGGTTGATATCAGGAGGAATACCAATGGCGAAGGCAGCTTGCTGGGCATCATCTGACGCTGAGGTGCGAAAGCGTGGGT |
| P-asv7089 | GCTAGCGTTGTTCGGAATTACTGGGCGTAAAGGGCGCGTAGGCGGATTAGTAAGTTGGGAGTGAAAGCCCGGGGCTTAACCTCGGAACTGCTTTCAAAACTGCTAGTCTTGAGTGAAGTAGGGGATGATGGAATTCCTAGTGTAGAGGTGAAATTCTTAGATATTAGGAGGAACACCGGTGGCGAGGGCGGTCATCTGGACTTCAACTGACGCTGAGGCGCGAAAGCGTGGGG |
| P-asv7090 | GCTAGCGTTGCTCGGAATCACTGGGCGTAAAGGGCGCGTAGGCGGACATTTAAGTCGGGGGTGAAAGCCCAGGGCTCAACCCTGGAATTGCCTCCGATACTGGGTGTCTAGAGACCGGAAGAGGTGAGTGGAACTGCGAGTGTAGAGGTGAAATTCGTAGATATTCGCAAGAACACCAGTGGCGAAGGCGGCTCACTGGTCCGGTTCTGACGCTGAGGCGCGAAAGCGTGGGG |
| P-asv7091 | GCAAGCGTTGTCCGGAATTACTGGGTGTAAAGGGCGTGTAGGCGGGAATATAAGTCAGATGTGAAATACCTGGGCTTAACTCAGGAACTGCAACTGAAACTATATTTCTTGAGTACTGGAGAGGAAAGTGGAATTCCTAGTGTAGCGGTGAAATGCGTAGATATTAGGAGGAACACCAGTGGCGAAGGCGACTTTCTGGACAGAAACTGACGCTGAGGCGCGAAAGCGTGGGG |
| P-asv7092 | CCGAGCGTTACCCGGATTTACTGGGCGTAAAGGGTGCGTAGGCGGTTATATTAGTCTTGTGTTAAATATTCGGGCTTAACCTGAATGCTGCATAGGAAACGGTATAACTAGAAGAAGCGAGAGGTGAACAGAACTCACGGTGTAGGGGTGAAATCCGTTGATATCGTGGGGAATACCAAATGCGAAGGCAGTTCACTGGCGCTCTCTTGACGCTGAGGCACGAAAGCGTGGGT |
| P-asv7093 | GCTAGCGTTGCTCGGAATTACTGGGCGTAAAGGACGCGTAGGCGGACATTTAAGTCAGATGTGAAATACCCGAGCTTAACTTGGGTGCTGCATTTGAGACTGGGTGTCTAGAGTGCAGGAGAGGTGAGTGGAATTCCTAGTGTAGCGGTGAAATGCGTAGAGATTAGGAAGAACACCAGTGGCGAAGGCGACTTACTGGACTGTAACTGACGCTGAGGCATGAAAGCGTGGGG |
| P-asv7094 | GCAAGCGTTAATCGGAATTACTGGGCGTAAAGGGTGCGTAGGTGGTCTGATAAGTTAGCTGTGAAATTCCTGGGCTTAACCTGGGGTGGTCAGCTAAGACTGTTAGACTGGAGTATAGGAGAGGGTAGTGGAATTTCCGGTGTAGCGGTGAAATGCGTAGAGATCTGGAGGAATACCGATGGCGAAGGCAGCCATCTGGCCTAATACTGACGCTGAGGTACGAAAGCATGGGG |
| P-asv7095 | GCAAGCGTTGTCCGGAATTATTGGGCGTAAAGAGCTCGTAGGCGGTTTGTCGCGTCTGCTGTGAAAGCCCGGGGCTCAACCCCGGGTCTGCAGTGGGTACGGGCAGACTAGAGTGCAGTAGGGGAGACTGGAATTCCTGGTGTAGCGGTGAAATGCGCAGATATCAGGAGGAACACCGATGGCGAAGGCAGCCATCTGGCCTAATACTGACGCTGAGGTACGAAAGCATGGGG |
| P-asv7096 | GCAAGCGTTACTCGGAATTACTGGGCGTAAAGCGTGCGTAGGCGGTTTGTTAAGTCAGAAGTGAAATCCCCGGGCTTAGCCCGGGAACTGCTTTTGAAACTCCAAGTCTAGAATCCTGGAGAGGTTGGGAGAATTCCGAGTGTAGAGGTGAAATTCGCAGATATTCGGAGGAATACCAGTGGCGTAGGCGCCCAACTGGACAGGTATTGACGCTCAGATACGAAAGCGTGGGG |
| P-asv7097 | CCGAGCGTTGTCCGGATTTATTGGGCGTAAAGCGAGCGCAGGCGGTTAGATAAGTCTGAAGTTAAAGGCTGTGGCTTAACCATAGTACGCTTTGGAAACTGTTTAACTTGAGTGCAAGAGGGGAGAGTGGAATTCCATGTGTAGCGGTGAAATGCGTAGATATGTGGAGGAGCACCAATGGCGAAGGCAGCCCCCTGGGATAACACTGACGCTCATGCACGAAAGCGTGGGG |
| P-asv7098 | GCTAGCGTTATTCGGAATTACTGGGCGTAAAGCGCACGTAGGCGGCTTTGTAAGTTAGAGGTGAAAGCCTGGAGCTCAACTCCAGAATTGCCTTTAAGACTGCATCGCTTGAATCCAGGAGAGGTGAGTGGAATTCCGAGTGTAGGGGTGAAATGCGTAGATATCTGGAGGAACATCAGTGGCGAAGGCGACACCCTGGACTCATACTGACACTGAGGTGCGAAAGCGTGGGG |
| P-asv7099 | GCAAACGTTGTCCAGAATTACTGGGTGTAAAGGGTGCACAGGCGGTTTTGTGCGTCAGAGGTAAAATCCCACAGCTCAACTGTGGACCTGCCTTTGATACGGCGAGACTTGAGTACAAGAGAGGATGATAGAATTCCTGGTGTAGCGGTGAAATGCGTAGATATCAGGAGGAATACCAGTGGCGAAGGCGGTCATCTGGCTTGTAACTGACGCTTAAGCACGAAAGCATGGGT |
| P-asv7100 | GCTAGCGTTATTCAGAATTACTGGGCGTAAAGCGCACGTAGGCGGCTTTGTAAGTTAGAGGTGAAAGCCTGGAGCTCAACTCCGGAACTGCCTTTGATACAGCATATCTCGAGTCCGAGAGAGGTGAGTGGAATTCCTAGTGTAGAGGTGAAATTCGTAGATATTAGGAAGAACACCGGTGGCGAAGGCGGCTCACTGGACTGGTATTGACGCTGAGGTGCGAAAGCGTGGGG |
| P-asv7101 | CCAAGCGTTATCCGGATTCATTGGGTTTAAAGGGTGCGTAGGCGGGTTTGTAAGTCAGTGGTGAAAGCCGGTCGCTTAACGATCGAACTGCCGTTGATACTGCAGACCTAGAATATAGATGATGTTGGCGGAATATGACATGTAGCGGTGAAATGCATAGATATGTCATAGAACACCGATTGCGAAGGCAGCTGACAAAACTATAATTGACGCTGATGCACGAAAGTGCGGGG |
| P-asv7102 | GCAAGCGTTGTCCGGACTCACTGGGCGTAAAGGGTGCGCAGGCGGTTTGGAGCGTTCGCGCTGAAATCACGGGGCTTAACTCCGTGGCGCGTGCGAATACGACCAAACTGGAGGCAGAGAGAGGATGGGGAACTCCTGGTGTAGTGGTGAAATGCGTAGATATCAGGAAGAACACCCGTGGCGTAAGCGCCCATCTGGCTCTGACCTGACGCTGAGGCACGAAAGCGTGGGG |

| | |
|---|---|
| P-asv7103 | GCAAGCGTTGTCCGGAATTATTGGGCGTAAAGAGTTCGTAGGCGGTTATCCAAGTCTGATGTTAAATACTGGCGCTCAACGTCAGCATGGCATTGGAAACTGGGTGGCTAGAGTATGGTAGAGGTCACGGGAATT<br>CCTGGTGTAGCGGTGAAATGCGTAGATATCAGGAGGAACACCGGTGGCGAAAGCGCGTGACTGGGCCATAACTGACGCTGAGGAACGAAAGCCAGGGT |
| P-asv7104 | GCGAGCGTTGTCCGGAATTATTGGGCGTAAAGCGCGCGCAGGCGGTCTTTTAAGTCTGATGTGAAATCTTACGGCTTAACCGTAAGCGGTCATTGGAAACTGGGAGACTTGAGTGCAGAAGAGGAGAGTGGAAT<br>TCCACGTGTAGCGGTGAAATGCGTAGAGATGTGGAGGAACACCAGTGGCGAAGGCGACTCTCTGGTCTGTAACTGACGCTGAGGAGCGAAAGCGTGGGG |
| P-asv7105 | GCGAGCGTTACTCGGAATCACTGGGCGTAAAGGGCGTGTAGGCGGCCCATCAAGTCAAATGTAAAATACAACGGCTCAACCGTTGGCCTGCGTTTGAAACTGAAGGGCTAGAACTTGGTAGAGGGTGATGGAAT<br>TCCCGGTGTAGCGGTGAAATGCGTAGATATCGGGAAGAACACCAAAGGCGAAGGCAGTCATCCATACCGAAGTTGACGCTGAGGCGCGAAAGCGTGGGG |
| P-asv7106 | GCAAGCGTTGCCCGGAATTATTGGGCGTAAAGCGCGCGCAGGCGGTCCTTTAAGTCTGATGTGAAATCTTGCGGCTCAACCGCAAGCGGTCATTGGAAACTGGGGGACTTGAGTACAGAAGAGGAAAGCGGAAT<br>TCCACGTGTAGCGGTGAAATGCGTAGAGATGTGGAGGAACACCAGTGGCGAAGGCGGCTTTCTGGTCTGTAACTGACGCTGAGGCGCGAAAGCGTGGGG |
| P-asv7107 | GCGAGCGTTATCCGGATTCATTGGGTTTAAAGGGTGCGTAGGCGGTTTCTTAAGTCAGTGGTGAAATCCTGCGGCTCAACCGTAGAACTGCCATTGATACTGAGGAACTTGAATTTAGTTGAGGTAGGCGGAATG<br>AGTAGTGTAGCGGTGAAATGCTTAGATATTACTCAGAACACCGATTGCGAAGGCAACTTGCCAAGCTACGATTGACGCTGATGCACGAAAGCGTGGGG |
| P-asv7108 | GCTAGCGTTGTTCGGAATCACTGGGCGTAAAGCGCGCGTAGGCGGTGACACAAGTCAGGGGTGAAATCCCAAGGCTCAACCTTGGAATTGCCTTTGAAACTAGGTCGCTGGAGTGTCGGAGGGGATAGCGGAA<br>TTGCTAATGTAGCGGTGAAATGCGTAGATATTAGCAGGAACACCGGTGGCGAAGGCGGCTATCTGGACGACAACTGACGCTGAGGCGCGAAAGCGTGGGG |
| P-asv7109 | GCAAACGTTGTTCGGTGTCACTGGGCTTAAAGAGCACGTAGGTGGCTGTGCAAGTAAGGTGTGAAAGCCCTCGGCTTACCCGGGGAATTGCACTTTAAACTGCATGGCTTGAGGGTCAGAGGGGTAAGAAGAAC<br>TCCAGGTGTAGCGGTGAAATGCGTAGATATCTGGGGGAAGGCCGATGGCGAAGGCGTCTTACTGGCTGATACCTGACACTGAGGTGCGAAAGCGTGGGG |
| P-asv7110 | GCTAGGGTTGTTCGGAATTACTGGGCGTAAAGCGCACGTAGGCGGATATTTAAGTCAGGGGTGAAATCCCAGAGCTCAACTCTGGAACTGCCTTTGATACTGGGTATCTTGAGTATGGAAGAGGTGAGTGGAATT<br>GCGAGTGTAGAGGTGAAATTCGTAGATATTCGCAGGAACACCAGTGGCGAAGGCGGCTCACTGGTCCATAACTGACGCTGAGGTGCGAAAGCGTGGGG |
| P-asv7111 | GCAAGCGTTGTTCGGAATCATTGGGCGTAAAGCGCGTGTAGGCGGCCAATTAAGTCTGATGTGAAAGCCCGGGGCTTAACCCCGGAAGTGCATTGGATACTGGTTGGCTAGAGTACTGTAGGGGGAAGTGGAAT<br>TCCTGGTGTAGGGGTGAAATCCGTAGATATCAGGAGGAACATCGGAGGCGAAGGCGACTTCCTGGGCAGTAACTGACGCTGAGACGCGAAAGCGTGGGG |
| P-asv7112 | CCAAGCGTTATCCGGATTTACTGGGCGTAAAGCGTCTGTAGCCTGATATGTAAGTCCGTTTTCAAATACCTGGACTCAATCGAGGGAAGGGGATAGATACTGCATATCTAGAGATATTTGGGGGTTAGTGGAATTT<br>CCGGTGGAGCGGTGAAATGCGTTGATATCGGAAAGAACGCCAAAAGCGAAAGCAGCTAACTACAATACATCTGACGGTGAGAGACGAAAGTTTGGGT |
| P-asv7113 | GCGAGCGTTAATCGGATTTACTGGGCGTAAAGGGTCCGTAGGCGGCCTTGTGTGTCGGGTGTGAAATCCCACGGCTTAACCGTGGAACTGCGCCCGAAACTACAAGGCTAGAGTATTGGAGAGGTGAGTGGAAT<br>TCCAGGTGTAGCGGTGAAATGCGTAGATATCTGGAGGAACACCAACGGCGAAGGCAGCTCACTGGACAATTACTGACGCTGAGGGACGAAAGTGTGGGG |
| P-asv7114 | GCGAACGTTGTTCGGAATCACTGGGCTTAAAGGGCGCGTAGGCGGGTCATCAAGTCCGGGGTGAAATCCTCCAGCTTAACTGGAGAATAGCCTTGGATACTGGTGGTCTCGAGTGAGGTAGGGGCAGCTGGAA<br>CTTCCGGTGGAGCGGTGAAGTGCGTTGATATCGGAAGAACACCAGTGGCGAAGGCGGCTCACTGGACTGGTATTGACGCTGAGGTGCGAAAGCGTGGGG |
| P-asv7115 | GCGAGCGTTAATCGGATTTACTGGGCGTAAAGCGTGCGTAGGCGGCTTTTTAAGTCGGATGTGAAATCCCTGAGCTTAACTTAGGAGTTGCATTCGATACTGGGAAGCTAGAGTATGGGAGAGGATGGTAGAATT<br>CCAGGTGTAGCGGTGAAATGCTTAGATATCTTCAAGAACACCGATTGCGTAGGCAGCTTACTAGGCTGTAACTGACGCTGAGGCACGAAAGCGTGGGG |
| P-asv7116 | GCAAGCGTTATCCGGAATCACTGGGTGTAAAGGGTGTGTAGGCGGTAATTTAAGTTTTGCGTTTAATTTCAGGGCTCAACCCTGAATGTGCGTGAAATACTGAATTGCTAGAGTGATGAAGAGGAAGTTGGAACTC<br>CAGGTGAAGCGGTGAAATGTGTAGATATCTGGGGGAACACCAATGGCGAGGGCAGGCTTCTGGTCATCAACTGACGCTGAGACACGAAAGCGTGGGG |
| P-asv7117 | CCGAGCGTTAATCGGAATCACTGGGCTTAAAGGGTGCGCAGGCTGGTCCGTGTGTGTGTCGTGAAATCCCCCGGCTCAACCGGGGAATTGCGGCGCAAACTGCGGACCTTGAGGCAGGTAGGGGTCGTTGGAA<br>CGATCGGTGGAGCGGTGAAATGCGTTGAGATCGATCGGAACGCCGAAGGTGAAGACAAGCGACTGGGCCTGACCTGACGCTCATGCACGAAAGCGTGGGG |
| P-asv7118 | GCTAGCGTTGCTCGGAATTACTGGGCGTAAAGGGCGCGTAGGCGGACAGTTAAGTTGGGGGTGAATGCCCGGGGCTCAACCACAGAATTGCCTTCGATACTGGTTGGCTTGAGACCGGAAGAGGACAGCGGAA<br>CTGCGAGTGTAGAGGTGAAATTCGTAGATATTCGCAAGAACACCAGTGGCGAAGGCGGCTGTCTGGTCCGGTTCTGACGCTGAGGCGCGAAAGCGTGGGG |
| P-asv7119 | GCGAGCGTTAATCGGAATCACTGGGCGTAAAGCGTGCGTAGGCTGCGTGGCAAGTCAGGCGTGAAAGCCCTCGGCTCAACCGAGGAATTGCGTTTGAAACTGCCATGCTAGAGTCTCGGAGAGGTTGGCGGAA<br>TTCCAGGTGTAGGAGTGAAATCCGTAGATATCTGGAGGAACACCGGCGGCGAAGGCGGCCAACTGGACGAGTACTGACGCTGAGGTACGAAAGCGTGGGT |
| P-asv7120 | TCGAGCGTTATCCGGAATTACTGGGCGTAGAGGGCACGCAGGCGGCTATTCAAGTCTGGCGTGAAAGCTCCCGGCTTAACTGGGAGAGGCCGTTGGAAACTGAGTAGCTTGAGGTGATGAGAGGGATGCGGAA<br>TTCCCGGTGTAGTGGTGGAATGCGTAGATATCGGGAAGAACACCAGTGGCGAAAGCGGCATCCTGGCATTGACCTGACGCTGAGGTACGAAAGCATGGGG |
| P-asv7121 | GCAAGCGTTGTCCGGAATGATTGGGCGTAAAGGGCGCGTAGGCGGCTTTATAAGTCTGGAGTGAAAGTCCTCTTTTCAAGGGAGGAATTGCTTTGGAAACTATAGAGCTTGAGTGCAGGAGAGGTTATCGGAATT<br>CCCGGTGTAGCGGTGAAATGCGTAGATATCGGGAGGAACATCGGTGGCGAAGGCGGGTAACTGGACTGCAACTGACGCTGAGGCGCGAAAGTGTGGGG |
| P-asv7122 | GCGAGCGTTAATCGGATTTACTGGGCGTAAAGCGTGCGTAGGCGGCTTCTTAAGTCGGATGTGAAATCCCTGAGCTTAACTTAGGAATTGCATTCGATACTGGGAAGCTAGAGTATGGGAGAGGATGGTAGAATT<br>CCATGTGTAGCGGTAAAATGCGTAAATATATGGAGGAACACCAGTGGCGAAGGCGGCTTGCTGGGCCACAACTGACACTGAGGCACGAAAGCGTGGGG |
| P-asv7123 | GCTAGCGTTGCTCGGAATTACTGGGCGTAAAGGGCGCGTAGGCGGTTTATCAAGTCAGGCGTGAAAGCCCCGGGCTCAACCTGGGAACTGCTTTCAAAACTGCTGAGCTAGAGTACGGTAGAGGGTGGTGGAAT<br>TTCCTGTGTAGCGGTGAAATGCGTAGATATAGGAAGGAACACCAGTGGCGAAGGCGACCACCTGGACTGATACTGACACTGAGGTGCGAAAGCGTGGGG |
| P-asv7124 | GCTAGCGTTGTTCGGAATCACTGGGCGTAAAGCGCACGTAGGCGGACTTTTAAGTCGGAGGTGAAAGCCTGGAGCTCAACTAGGAAACTGCCATTGATACTGTAGGGCTTGAATTTTTGTGAAGTAACTAGAATAT<br>GTAGTGTAGCGGTGAAATGCTTAGATATTACATGGAATACCAATTGCGAAGGCAGGTTACTAACAAACGATTGACGCTGATGGACGAAAGCGTGGGG |
| P-asv7125 | GCAAGCGTTGTCCGGATTCATTGGGCGTAAAGAGCTCGTAGGCGGTTCGGTAAGTCAGATGTCAAAACTCCAGGCTCAACTTGGAGCGGCCATCTGATACTGCCGTGACTAGAGTCCGGTAGAGGAGCATGGAA<br>TTCCTGGTGTAGCGGTGAAATGCGCAGATATCAGGAAGAACACCAGCGGCGAAGGCGGTGCTCTGGGCCGGAACTGACGCTGAGGCACGAAAGCGTGGGG |
| P-asv7126 | GCAAGCGTTATCCGGAATTATTGGGCGTAAAGCGCGCGTAGGCGGTTTTTTAAGTCTGATGTGAAAGCCCCCGGCTCAACCGGGGAGGGTCATTGGAAACTGGGGAACTTGAGTGCAGAAGAGGAGAGTGGAAT<br>TCCATGTGTAGCGGTGAAATGCGCAGAGATATGGAGGAACACCAGTGGCGAAGGCGGCTCTCTGGCCTAATACTGACGCTGAGGTACGAAAGCATGGGG |
| P-asv7127 | GCGAGCGTCAATCGGAATTACTGGGCGTAAAGCGTGCGTAGACGGTCGTGCAAGTCGGGTGTGAAAGCCCCAGGCTCAACCTGGGAATTGCATTCGATACTGCTCGGCTAGGGTGCGGAAGAGGGAAGCGGAA<br>TTTCCGGTGTAGCGGTGAAATGCGTAGATATCGGAAGGAACACCGGTGGCAAAAGCGGCTTCCTGGTCCGACACCGACGTTCAGGCACGAAAGCGTGGGG |
| P-asv7128 | GCGAGCGTTGTCCGGAATTACTGGGCGTAAAGGGCTCGTAGGTGGTTTGTCGCGTCGTCTGTGAAATTCTGGGGCTTAACTCCGGGCGTGCAGGCGATACGGGCATAACTTGAGTGCTGTAGGGGTAACTGGAA<br>TTCCTGGTGTAGCGGTGAAATGCGTAGATATAGGAAGGAACATCAGTGGCGAAGGCGACCACCTGGGCTAATACTGACACTGAGGTGCGAAAGCGTGGGG |
| P-asv7129 | CCGAGCGTTGTCCGGAATTATTGGGCGTAAAGGGCTCGTAGGCGGTTTGTCGCGTCGGGAGTGAAAACTCAGGGCTCAACCCTGAGCGTGCTTCCGATACGGGCAGACTAGAGGTATGCAGGGGAGAACGGAA<br>TTCCTGGTGTAGCGGTGGAATGCGCAGATATCAGGAGGAACACCGGTGGCGAAGGCGGTTCTCTGGGCATTACCTGGCGCTGAGGAGCGAAAGCATGGGG |
| P-asv7130 | GCAAGCGTTAATCAGAATTACTGGGCGTAAAGCGTGCGCAGGCCGCGTTTTAAGCGTGTTGTGAAATGTAGGCGCCCAACGTCTGCATCGCAGCGCGAACTGGAACGCTTGAGTACGCGCAACGTTGGCGGAAT<br>TCGTCGTGTAGCGGTGAAATGCTTAGATATGACGAAGAACTCCGATTGCGAAGGCAGCTGACGGGAGCGGCACTGACGCTTAAGCTCGAAGGTGCGGGT |
| P-asv7131 | GCAAGCGTTATCCGGATTTATTGGGTTTAAAGGGTCCGCAGGCGGATCGTTAAGTCAGTGGTGAAAGCCTACCGCTCAACGGTAGAACTGCCATTGATACTGGCGATCTTGAATATGTATGAAGTTGGCGGAATG<br>TGTGATGTAGCGGTGGTATGCGTAGAGATGTGGAGGAATACCGATGGCGAAGGCAGCCTCCTGGGATAACACTGACGTTGAGGCTCGAAAGCGTGGGG |
| P-asv7132 | CCAAGCGTTATTCGGAATCATTGGGTTTAAAGGGTCCGTAGGCGGGTGCGTAAGTCAGTGGTGAAAGTCTGCAGCTCAACTGTAGAACTGCCATTGATACTGCACACCTTGAGTAATTATGAAGTGGTTAGAATAT<br>GTAGTGTAGCGGTGAAATGCATAGATATTACATAGAATACCGATTGCGAAGGCAGATCACTAATAATTTACTGACGCTGATGGACGAAAGCGTAGGT |
| P-asv7133 | GCTAGTGTTGCTCGGAATTACTGGGCGTAAAGGGCGCGTAGGCGGACAGTTTAGTCAGAGGTGAAAGCCCAGGGCTCAACCTTGGAATTGCCTTTGATACTGGGAAGCTAGAGTATGGGAGAGGATGGTAGAAT<br>TCCAGGTGTAGCGGTGAAATGCGTAGAGATCTGGAGGAATACCGATGGCGAAGGCAGCCATCTGGCCTAATACTGACGCTGAGGTACGAAAGCATGGGG |
| P-asv7134 | GCGAGCGTTAATCGGAATCACTGGGCGTAAAGGGCGCGTAGGCGGTTAGTTACGTCTGCCGTGGAATCCCCGGGCTCAACCTGGGAATTGCATTCAAAACTGGCTAACTAGAGTATGTGAGAGGGGGGTAGAAT<br>TCCAAGTGTAGCGGTGAAATGCGTAGAGATTTGGAGGAATACCAGTGGCGAAGGCGGCCCCCTGGCACAATACTGACGCTGAGGTGCGAAAGCGTGGGG |
| P-asv7135 | GCAAGCGTTATCCGGATTTATTGGGTTTAAAGGGTCCGTAGGCGGATCTGTAAGTCAGTGGTGAAATCTCACTGTTTAACTGTGAAACTGCCATTGATACTGCAGGTCTTGAGTAAATTTGAAGTGGCTGGAATAA<br>GTAGTGTAGCGGTGAAATGCATAGATATTACTTAGAACACCAATTGCGAAGGCAGGTCACTAAGATTTAACTGACGCTGATGGACGAAAGCGTGGGG |
| P-asv7136 | GCAAGCGTTGTTCGGAGTTACTGGGCGTAAAGAGTCTGTAGGTGGTCTGTCAAGTCTTTGGTGAAATCCTAGAGCTTAACTCTAGAGCTGCCATTGATACTATTTATCTTGAATATTGTGGAGGTAAGCGGAATATG<br>TCATGTAGCGGTGAAATGCTTAGATATGACATAGAACACCTATTGCGAAGGCAGCTTACTACGCATATATTGACGCTGAGGCACGAAAGCGTGGGG |
| P-asv7137 | GCAAGCGTTAATCGGAATTATTGGGCGTAAAGCGAGTGCAGACGGTTACCTAAGCCAGATGTGAAATCCCCAAGCTTAACTTAGGAATTGCATTCGATACTGGGAAGCTAGAGTATGGGAGAGGATGGTAGAATT<br>CCAGGTGTAGCGGTGAAATGCGTAGAGATCTGGAGGAATACCGATGGCGAAGGCAGCCATCTGGCCTAATACTGACGCTGAGGTACGAAAGCATGGGG |
| P-asv7138 | GCGAGCGTTGTCCGAAGTTACTGGGCGTAAAGCGCGCGTAGGCGGCTCTTTAAGTCCGGGGTGAAAGGTCCAGCGCTCAACGCGGACAGTGCCCTGGATACTGGGGGGCTTGAGTGTGGCAGAGGAGAGTGG<br>AATGGCTGGTGTAGCGGTGAAATGCGTAGATATCAGTCGGAACACCCGTGGCGAAGGCGGCTCTCTGGGCTAACACTGACGCTGAGGTGCGAAAGCGTGGGG |
| P-asv7139 | GCTAGCGTTGTCCGGAATCATTGGGCGTAAAGAGCGTGTAGGCGGCCCGGTAAGTCCGCTGTAAAAGTCAAAGGCTCAACCTTTGAATGTCAGTGGATACTGCCGGGCTAGAGTCCGGAAGAGGCGAGTGGAAC<br>TCCTGGTGTAGCGGTGGAATGCGCAGATATCAGGAAGAACACCAACGGCGAAGGCAGCTCGCTGGGACGGAACTGACGCTAAGGCGCGAAAGCGTGGGG |
| P-asv7140 | GCGAGCGTTAATCGGAATTACTGGGCGTAAAGCGCACGTAGGTGGCGACGTAAGTCGGGTGTGAAAGCCCCGGGCTCAACCTGGGAACTGCATTCGAGACTGCGTTGCTGGAGTACGGAAGAGGAGGGCGGA<br>ATTCCCGGTGTAGCGGTGAAATGCGTAGATATCGGGAGGAACACCAGTGGCGAAGGCGAGTCTCTGGGATGATGCTGACGCTGAGGCGCGAAAGCCAGGGG |
| P-asv7141 | GCTAGCGTTGCTCGGAATTACTGGGCGTAAAGGGAGCGTAGGCGGACTGTTAAGTTAGAGGTGAAAGCCCAGGGCTCAACCTTGGAATTGCCTTTGATACTGGCAGTCTTGAGTACGGAAGAGGTATGTGGAAC<br>TCCGAGTGTAGAGGTGAAATTCGTAGATATCAGGAGGAACACCGGTGGTGTAGACGGCTCTCTGGACCGTAACTGACACTGAAACACGAAAGCGTGGGT |

| | |
|---|---|
| P-asv7142 | GCAAGCGTTGTCCGGAATTATTGGGCGTAAAGCGAACGTAGGCGGTTTCTTAAGTTTGGTCTTAAAGACCGGGGCTCAACCCCGGGAATGGACCGAAAACTGGGAGACTCGAGGATGTCAGAGGAAAGTGGAATTCCGAGTGTAGCGGTGAAATGCGTAGATATTCGGAGGAACACCAGTGGCGAAAGCGACTTTCTGGGGCGTTCCTGACGCTGAGGTTCGAAAGCTCGGGG |
| P-asv7143 | GCAAGCGTTGTTCGGAATTATTGGGCGTAAAGGGTGCGTAGGCGGTTTGGTAAGTCTTGTGTGAAATCTTCGGGCTCAACTCGAAGTCTGCACAAGAAACTGCCGGGCTTGAGTATGGGAGAGGTGAGTGGAATTTCCGGTGTAGCGGTGAAATGCGTAGATATCGGAAGGAACACCTGTGGCGAAAGCGGCTCACTGGACCATAACTGACGCTGATGCACGAAAGCTAGGGG |
| P-asv7144 | GCAAGCGTTAATCGGAATTACTGGGCGTAAAGCGTGCGTAGGCGGATTCTTAAGTCGGTGGTGAAATCCTGGAGCTCAACTCCAGAACTGCCTCCGATACTGGGAGTCTTCGAGTCCGGGAGAGGTGAGTGGAACTGCGAGTGTAGAGGTGAAATTCGTAGATATTCGCAAGAACACCAGTGGCGAAGGCGGCTCACTGGCCCGGTACTGACGCTGAGGTGCGAAAGCGTGGGG |
| P-asv7145 | GCAAGCGTTGCTCAGAATTATTGGGCGTAAAGGGTAGGTAGGTGGTTACGTATGTCTGGGGTGAAATCCCTGAGCTCAACTCAGGAAGTGCCTTGGAAACGGCGTAACTAGAGTGCTAGAGAGGTTCGTGGAATTCCCAGTGTAGCGGTGAAATGCGTAGAGCTTGGGAGGAACACCAGAGGCGAAGGCGGCGAACTGGATAGCAACTGACACTGAACTACGAAGGCGTGGGT |
| P-asv7146 | GCAAGCGTTGTTCGGAATTACTGGGCGTAAAGGGCTCGTAGGCGGCCAACTAAGTCAGACGTGAAATCCCTCGGCTTAACCGGGGAACTGCGTCTGATACTGGATGGCTTGAGTTTGGGAGAGGGATGCGGAATTCCAGGTGTAGCGATGAAATGCGTAGATATCAGGAGGAACACCGGTGGCGAAGGCGTCCATCTGGCCCGATACTGACGCTGAGGCGCGAAAGCTAGGGG |
| P-asv7147 | CCGAGCGTTACTGAGAGTCACTGGGTTTAAAGGGTGCGTAGGTGGCGCGTTAAGTTCGTTGCGAAATCCCTGGGCTCAACCCAGGAACCGCTTCGAATACTGGCGTGCTTGAGGCTAGTAGAGGTCACTGGAACTGACGGTGGAGCGGTGAAATGCGTAGATATCGTCAGGAACGCCGGTGGTGAGGACGGGTGACTGGGCTAGTTCTGACACTGAGGCACGAAAGCGTGGGG |
| P-asv7148 | GCAAGCGTTGTCCGGAATCATTGGGCGTAAAGAGCGCGTAGGCGGTCCGGTAAGTCTGCCGTGAAAGCCGGGGGCTCAACCCTCGGATGCCGGTGGATACTGTCGGGCTAGAGTCCGGAAGAGGCGAGTGGAATTCCTGGTGTAGCGGTGAAATGCGCAGATATCAGGAGGAACACCAATGGCGAAGGCAGCTCGCTGGGACGGTACTGACGCTGAGGCGCGAAAGCGTGGGG |
| P-asv7149 | GCGAGCGTTAATCGGAATTACTGGGCGTAAAGGGCGCGTAGGCGGTTATTTAAGTTGGATGTGAAATCCCTGGGCTTAACCTAGGAACTGCATACAATACTGGATAACTGGAGTGCAGTAGAGGGTAGTGGAATTTCCGGTGTAGCGGTGAAATGCGTAGATATCGGAAGGAACACCAGTGGCGAAGGCGGCTACCTGGACTGACACTGACGCTGAGGCGCGAGAGCGTGGGG |
| P-asv7150 | GCTAGCGTTGTTCGGAATTACTGGGCGTAAAGGGCGCGTAGGCGGCCCGATAAGTCTGGTGTGAAATCTTGGGGCTCAACCCCAAGCCTGCACTGGATACTGTCGAGCTTGAGTCCGGGAGGGGAGAGCGGAATTCCTAGTGTAGCGGTGAAATGCGCAGATATTAGGAAGAACACCGGTGGCGAAGGCGGCTCTCTGGAACGGTACTGACGCTGAGACGCGAAAGCGTGGGG |
| P-asv7151 | GCAAGCGTTGTTCGGAATTACTGGGCGTAAAGAGTGTGTAGGCGGTGCTCCAAGTCTGGTGTGAAATCTCCCGGCTTAACTGGGAGGGTGCATTAGAAACTGGAATGCTAGAGGATGGGAGAGGTAAGCGGAATTCCTGGTGTAGCGGTGAAATGCGTAGATATCAGGAGGAACACCTGTGGTGTAGACAGCTTACTGGACCATTTCTGACGCTGAGACACGAAAGCGTGGGT |
| P-asv7152 | GCTAGCGTTGTTCGGAATTACTGGGCGTAAAGCGCACGTAGGCGGCTTTGTAAGTCAGAGGTGAAAGCCTGGAGCTCAACTCCAGAACTGCCTTTGAGACTGCATCGCTTGAATCCAGGAGAGGTGAGTGGAATTCCGAGTGTAGAGGTGAGATGCGTAGATATCGGGAAGAATACCAGTGGCGAAAGCGGCTTCCTGGTCCAGCACCGACGTTCAGGTACGAAAGCATGGGG |
| P-asv7153 | GCAAGCGTTACTCGGAGTTACTAGGCGTAAAGGGCAGGTAGGTGGTTTGATAAGTCTATTGTGAAAGCTCCCGGCTCAACTGGGAGAGGCCAACAGAAACTATCAGACTTGAGTATGGGAGAGGGTACTGGAATTCCCGGTGTAGCGGTGAAATGCGCAGAGATCGGGAGGGACACCAATGGCGAAAGCAGGTACCTGGACCATCACTGACACTCAGCTGCGAAAGCTAGGGG |
| P-asv7154 | GCGAGCGTTGTTCGGAATTATTGGGCGTAAAGGGCGCGTAGGCGGTGCCACAAGTCACTTGTGAAATCTCCGGGCTTAACTCGGAGCCTGCAGGCGAAACTGTGGTGCTGGAGTGTGGGAGAGGTGCGTGGAATTCCCGGTGTAGCGGTGAAATACGTAGATATCGGGAGGAACACCAGTGGCGAAGGCGGCGATCTGGGAACATACTGACGCTGAGGTGCGAAAGCGTGGGG |
| P-asv7155 | GCGAGCGTTGTCCGGATTTACTGGGCGTAAAGAGCGCGCAGGCGGCTGGTCAAGTCCGATGTGAAAGCTCCCGGCTTAACTGGGAAATTGCATCGGAAACTGATCGGCTGGAAGGTGGGAGAGGGTAGCGGAATTCCCGGTGTAGTGGTGAAATGCGTAGATATCGGGAGGAACACCAGTGGCGAAGGCGCTCCTCTGGCCCAATACTGACGCTGAGGCGCGAAAGCTAGGGG |
| P-asv7156 | GCAAGCGTTAATCGGAATTACTGGGCGTAAAGCGCACGCAGGCGGTCTGTCAAGTCGGATGTGAAATCCCCGGGCTCAACCTGGGAACTGCATCCGAAACTGGCAGGCTAGAGTCTTGTAGAGGGGGGTAGAATTCCAGGTGTAGCGGTGAAATGCGTAGAGATGCGGAGGAGCACCGATGGCGAAGGCAGCCTCCTGGGCCAATACTGACGCTCATGCACGAAAGCGTGGGG |
| P-asv7157 | GCTAGCGTTGCTCGGAATTACTGGGCGTAAAGGGCGCGTATGCGGACAGTTAAGTTGGGGGTGAAAGCCCGGAGCTCAACTCCGGAACTGCCATTGAGACTGTTTAGCTTGAGTATGAGAGAGGTGAGTGGAATTCCCAGTGTAGAGGTGAAATTCGTAGATATTGGGAAGAACACCGGTGGCGAAGGCGGCTCACTGGCTCATTACTGACGCTCAGGCGCGACAGCGTGGGG |
| P-asv7158 | ACGAGTAGTGGTCACTTTTATTGAGCCTAAAGCGTTCGTAGCCGGTTTTGTAAATCTTCAGATAAGGCCTGAAGCTTAACTCCAGAACTGCCTTTGATACTGAAGATCTTGAGTTCGGGAGAGGTGAGTGGAACTGCGAGTGTAGAGGTGAAATTCGTAGATATTCGCAAGAACACCAGTGGCGAAGGCGGCTCACTGGCCCGATACTGACGCTGAGGCACGAAAGCGTGGGG |
| P-asv7159 | GCTAGCGTTGCTCGGAATTACTGGGCGTAAAGGGCGCGTAGGCGGACAGTTAAGTTGGGGGTGAAAGCCCGGGGCTCAACCTCGGAAATGCCTTCAATACTGGCTGTCTTGAGTACGGGAGAGGTGAGTGGAACTCCGAGTGTAGAGGTGAAATTCGTAGATATCGGGAGGAACACCAGTGGCGAAGGCGGCTTCCTGGACCAGTACTGACGCTGAGGCGCGTAAGCTAGGGG |
| P-asv7160 | GCTAGCGTTGCTCGGAATCACTGGGCGTAAAGGGTGTGTAGGTGGATCTTTAAGTCAGGGGTGAAATCCTGGAGCTCAACTTCGGAACTGCCTCTGATACTGCATGTCTTGAGTCCGGAAGAGGTGGGTGGAATTCCTAGTGTAGAGGTGAAATTCGTAGATATTAGGAAGAACACCGGTGGCGAAGGCGACCCACTGGTCCGGTACTGACGCTGAGGTGCGAAAGCGTGGGG |
| P-asv7161 | ACTAGCGTTGTCCGGAATTATTGGGCGTAAAGGGCGCGTAGGCGGCGCGGTAAGTCACCTGTGGAATCTCCTGGCTCAACTGGGAGCTTGCAGGCGAAACTGCCGTGCTGGAGTGTGGGAGAGGTGCGTGGAATTCCCGGTGTAGCGGTGAAATGCGTAGATATCGGGAGGAACACCTGTGGCGAAAGCGGCGCACTGGACCACAACTGACGCTGAGGCGCGAAAGCTAGGGG |
| P-asv7162 | GCAAGCGTTGTTCGGAATTACTGGGCGTAAAGGGTGCGTAGGCGGCTTTCTAAGTCGGATGTGAAAACTCTGGGCTCAACCCAGAGCCTGCATCCGAAACTGGAGAGCTAGGGTTCTGGAGGGGAAAGCGGAATTCCTGGTGTAGCGGTGAGATGCGTAGATATCAGGAGGAACACCGGTGGCGAAGGCGGCTTTCTGGACAGATACCGACGCTGAGGCACGAGAGCTAGGGG |
| P-asv7163 | GCGAGCGTTGTCCGGAATTATTGGGCGTAAAGAGCGTGTAGGCGGTTTGGTAAGTCTGCCGTGAAAACCCGGGGCTCAACCCCGGGCGTGCGGTGGATACTGCCAGGCTAGAGGATGGTAGAGGCGAGTGGAATTCCCGGTGTAGCGGTGAAATGCGCAGATATCGGGAGGAACACCAGTAGCGAAGGCGGCTCACTGGCCCGATACTGACGCTGAGGCACGAAAGCGTGGGG |
| P-asv7164 | GCAAGCGTTGTCCGGAATTATTGGGCGTAAAGCGCGCGCAGGCGGCTTTGTAAGTCTTGTGTTTAAACGCGGGGCTCAACCCCGTGTCGCATGGGAAACTGCAAGGCTTGAGTGCAGAAGAGGAAAGTGGAATTCCACGTGTAGCGGTGAAGTGCGTAGAGATGTGGAGGAACACCAGTGGCGAAGGCGGCTCACTGGACTGGTATTGACGCTGAAGTGCGAAAGCGTGGGG |
| P-asv7165 | GCAAGCGTTAATCGGAATTACTGGGCGTAAAGCGTGCGTAGGTGGTTTATTAAGTCAGATGTGAAAGCCCCGGGCTTAACCTGGGAACTGCATTTGAAACTGGTAAACTAGAGTATGGTAGAGGAAAGTGGAATTTCTGGTGTAGCGGTGAAATGCGTAGATATCAGAAGGAACATCAATGGCGAAGGCAGCTTTCTGGACCAATGCTGACACTGAGGTACGAAAGCGTGGGT |
| P-asv7166 | GCCCTCTTTCTCCTTCCTTCTTGTTCTTAAAGCGTCTGTAGGTGGCTTTTTAAGTCCGCCGTCAAATCCCAGGGCTCAACCCTGGACAGGCGGTGGAAACTACCAAGCTGGAGTACGGTAGGGGCAGAGGGAATTTCCGGTGGAGCGGTGAAATGCGTAGAGATCGGAAAGAACACCAACGGCGAAAGCACTCTGCTGGGCCGACACTGACACTGAGAGACGAAAGCTAGGGG |
| P-asv7167 | GCTAGCGTTGCTCGGAATCACTGGGCGTAAAGGGCGCGTAGGCGGCTTGCCAAGTCGGGGGTGAAAGCCCGTGGCTCAACCACGGAATGGCCTTCGATACTGGCAGGCTTGAGACCGGAAGAGGACAGCGGAACTGCGAGTGTAGAGGTGAAATTCGTAGATATTCGCATGAACACCAGTGGCGAAGGCGGCTTACTGGCCCGATACTGACACTGAGGCACGAAAGCGTGGGG |
| P-asv7168 | GCAAGCGTTGTTCGGAATTACTGGGCGTAAAGCGTGCGTAGTCGGTATTGAGAGTCACGGGTGAAATCCCAGGGCTTAACCCTGGAACTGCCTGTGAGACCTCAGTACTAGAGTGTGAGAGGGGATAGTGGAATACCCAGTGTAGCGGTGAAATGCGTAGAGATTGGGTGGAACACCTATGGCGAAAGCAGCTGCTTGGGCCTCTATTGACGCTAAGACTCGAAAGCTTGGGG |
| P-asv7169 | GCGAGCGTTAATCGGAATTACTGGGCGTAAAGCGCATGCAGGTGGTTTGTTAAGTCAGATGTGAAAGCCCGGGGCTCAACCTCGGAATAGCATTTGAAACTGGCAGACTAGAGTACTGTAGAGGGGGGTAGAATTCCAGGTGTAGCGGTGAAATGCGTAGAGATCTGGAGGAATACCGATGGCGGAGGCAGCCATCTGGCCTAACACTGACGCTGAGGTGCGAAAGCATGGGG |
| P-asv7170 | GCTAGCGTTGTTCGGAATTACTGGGCGTAAAGGGCGCGTAGGCGGTTTATCAAGTCAGGCGTGAAAGCCCCGGGCTCAACCTGGGAATTGCGCTTGATACTGGTAGACTTGAGTTCGGGAGAGGTGAGTGGAATTCCCAGTGTAGAGGTGAAATTCGTAGATATTGGGAAGAACACCGGTGGCGAAGGCGACTCTCTGGTCTGTAACTGACGCTGAGGTGCGAAAGCGTGGGT |
| P-asv7171 | GCAAGCGTTATCCAGAATCACTGGGCGTAAAGGGTGCGTAGGCGGCCCAACAAGTCAGGTGTGAAAGCCTTGAGCTCAACTCAAGAAATGCACTTGATACTGGGCGACTAGAGGACCGGAGAGGATAGTGGAATTCCCAGTGTAGTGGTGAAATACGTAGAGATTGGGAAGAACACCAGTGGCGAAGGCGGCTATCTGGACGGTTTCTGACACTAAGACTCGAAAGCGTGGGG |
| P-asv7172 | GCAAGCGTTATCCGGAATCACTGGGCGTAAAGGGTGCGTAGGCGGCCTAACAAGTCAGAGGTGAAAGGCATTGGCTCAACCAATGTAAGCCTTTGAAACTGTTAGGCTTGAGTGCAGGAGAGGTAAGTGGAATTCCCAGTGTAGTGGTGAAATACGTAGAGATTGGGAAAAACACCAGTGGCGAAGGCGGCTATCTGGACGGTTTCTGACACTAAGACTCGAAAGCGTGGGG |
| P-asv7173 | GCTAGCGTTATTCGGAATTACTGGGCGTAAAGCGCACGTAGGCGGCTTTGTAAGTTAGAGGTGAAAGCCTGGAGCTCAACTCCAGAATTGCCTTTAAGACTGCATCGCTTGAATCCAGGAGAGGTGAGTGGAATTCCGAGTGTAGAGTTAAAATTCGTCGAAACTGGAAGGAACACCAAGGGCGAAAGCAATTCTCGACAATATTCTGACGTTGAGGCACGAAGGGATGGGT |
| P-asv7174 | CCAAGCGCTGTCCGGATTTATTGGGCGTAAAGAGCTCGTAGGTGGTTCGGTAAGTCAGCTGTGAAATCTCCGCGCTCAACGCGGAGAGGTCAGCTGATACTGCCGTGACTCGAGTCCGGTAGGGGAGTGTGGAATTCCCGGTGTAGCGGTGAAATGCGCAGATATCGGGAGGAACACCAGTAGCGAAGGCGGCACTCTGGGCCGGTACTGACGCTGAGGAGCGAAAGCGTGGGG |
| P-asv7175 | GCAAGCGTTGTCCGGAATTATTGGGCGTAAAGAGCTCGTAGGCGGCTTGTCGCGTCGGATGTGAAAATCCAAGGCTCAACCTTGGACCTGCATTCGATACGGGCAGGCTAGAGTTCGGTAGGGGAGACTGGAATTCCTGGTGTAGCGGTGAAATGCGCAGATATCAGGAGGAACACCGGTGGCGAAGGCGGGTCTCTGGGCCGATACTGACGCTGAGGAGCGAAAGCGTGGGG |
| P-asv7176 | GCAAGCGTTATCCGGATTCACTGGGTTTAAAGGGTGCGTAGATGGGCTGTTAAGTCAGTGGTGAAATCTTTGGGCTTAACCCGGAAACTGCCATTGATACTATCAGTCTTGAATATTGTGGAGGCCTGCGGAATATGTCATGTAGCGGTGAAATGCTTAGATATGACATAGAACACCAATTGCGAAGGCAGCAGGCTACGCATATATTGACGCTGAGGCACGAAAGCGTGGGG |
| P-asv7177 | GCAAGCGTTAATCGGAATTACTGGGCGTAAAGCGTGCGTAGGTGGTTTGTTAAGTCAGATGTGAAAGCCCCGGGCTCAACCTGGGAACTGCATTTGAAACTGGCAAACTAGAGTATAGGAGAGGAAAGTGGAATTTCTGGTGTAGCGGTGAAATGCGTAGATATCAGAAGGAACATCAATGGCGAAGGCAGCTTTCTGGACTAATACTGACACTGAGGTACGAAAGCGTGGGT |
| P-asv7178 | CCGAGCGTTATCCGGATTTATTGGGTTTAAAGGGTGCGCAGGCGGGATGTTAAGTCGGCGGTGAAATTTTGCGGCTCAACCGTAAAAGTGCCTTCGAAACTGGCATTCTTGAGTATAGATGAAGTAGGCGGAATTTGTGGTGTAGCGGTGAAATGCATAGATATCACGAGGAACTCCGATTGCGCAGGCAGCTTACTAACTATAACTGACGCTCATGCACGAAGGCGTGGGG |
| P-asv7179 | GCGAGCGTTATTCGGAATTACTAGGCGTAAAGCGAGTGTAGGCGGACTCTTAAGTCTGACGTTAAATCTCCTGGCTTAACTAGGAGCTGTCGTCGGATACTGAGAGCCTCGAATGGGGTAGGGGGCAGCGGAATTCCCGGTGTAGCGGTGAAATGCGTAGATATCGGGAGGAACACCTATGGCGAAAGCAGCTGCTTGGGCCTCTATTGACGCTAAGACTCGAAAGCTGGGGG |
| P-asv7180 | GCAAGCGTTAATCGGAATTACTGGGCGTAAAGCGCGCGTAGGTGGTTCGTTAAGTTGGATGTGAAATCCCCGGGCTCAACCTGGGAACTGCATTCAAAACTGACGAGCTAGAGTATGGTAGAGGGTGGTGGAATTTCCTGTGTGGCGGTGAAATGCGTAGATATAGGAAGGAACACCAGTGGCGAAGGCGACTTACTGGACTGTAACTGACGCTGAGGCACGAAAGCGTGGGG |

| | |
|---|---|
| P-asv7181 | GCAAGCGTTATCCGGAATCACTGGGCGTAAAGGGTGCGTAGGCGGCCTAACAAGTCAGAGGTGAAAGGCATTGGCTCAACCAATGTAAGCCTTTGAAACTGTTGGGCTTGAGTGCAGGAGAGGTAAGTGGAATTCCTAGTGTAGAGGTGGAATGCGTAGAGATGTGGAGGAACACCAGTGGCGAAGGCGACTCTTTGGTCTGTAACTGACGCTGAGGCGCGAAAGCGTGGGG |
| P-asv7182 | GCAAGCGTTAATCGGAATTACTGGGCGTAAAGCGCGCGTAGGTGGCTTGATAAGTTGGATGTGAAATCCCCGGGCTCAACCTGGGAACTGCATCCAGAACTGTCTGGCTAGAGTGCGGTAGAGGGTAGTGGAATTTCCAGTGTAGCGGTGAAATGCGTAGATATGTGGAGGAACACCAGTGGCGAAGGCGACTCTTTGGTCTGTAACTGACGCTGAGGCGCGAAAGCGTGGGG |
| P-asv7183 | CCAAGCGTTGTCCGGATTTACTGGGCGTAAAGAGCGCGTAGGCGGTCTGTTAAGTGTGGAGTGAAATCTCCAGGGCTCAACCCGGAAACTGCTTCGCATACTGGCAGACTGGAGGGATGCAGAGGTTTGTGGAATTCCTGGTGTAGCGGTGATATGCATAGATATCAGGAGGAACACCCATGGCGAGGGCAGCAAACTGGGCATCATCTGACGCTGAGGCGCGAAAGCGTGGGT |
| P-asv7184 | GCTAGCGTTGCTCGGAATCACTGGGCGTAAAGGGTGCGTAGGCGGGTCTTTAAGTCAGGGGTGAAATCCTGGAGCTCAACTCCAGAACTGCCTTTGATACTGAAGATCTTGAGTTCGGGAGAGGTGAGTGGAACTGCGAGTGTAGAGGTGAAATGCGTAGAGATTGGGTAGAACACCGGTGGCGAAGGCGGCTATCTGGCTCACAACTGACGATCAGGCACGAAAGCGTGGAG |
| P-asv7185 | GCAAGCGTTGTCCGGATTTACTGGGCGTACAGCGCGCGCAGGCGGCTCGCCAGGAACCCGGTGACAGTCTCCCGCTCAATGGGAGGAGGTCTGGGTTGACCGGCGGGCTGGAGGCCGTGAGAGGGTGATGGAACGCCCGGTGGAGCGGTGAAATGCGTAGAGATCGGGCAGAACACCAGTGGGGAAGCCGGTCACCTGGCACGGACCTGACGCTGAGGCGCGACAGCGTGGGG |
| P-asv7186 | GCAAGCGTTATTCGGAATTATTGGGCGTAAAGGGTGCGTAGGCTGCGTACTAAGTCAACTGTTAAATTTCTCGGCCTAACTGAGAATCTGCAGTAGATACTGGTATGCTAGAGGGTGGAAGAGAGAAGTGGAATTCTCGGAGTAGCGGTAAAATGCGTAGATCTCGAGAGGAACACCAATGGCGAAGGCAGCTTCTTGGTCCACACCTGACGCTGAGGCACGAAAGCGTGGGG |
| P-asv7187 | GCGAACGTTATTCGGAATCACTGGGCTTACAGGGTGCGTAGGCGGCCTGTCAAGTAGGGTGTGAAAGCCCCCGGCTCAACCGGGGAATGGCGCCCTAAACTGGCGGGCTGGAGTGAGATAGGGGTGTGCGGAACTTCCGGTGGAGCGGTGAAATGTGTTGATATCGGAAGGAACGCCGGTGGCGAAAGCGGCACACTGGATCTCAACTGACGCTGAGGCACGAAAGCCAGGGG |
| P-asv7188 | GCGGACGTTGTTCGGAATCACTGGGCTTAAAGGGCGCGTAGGCGGGCTGACAAGTCTGGGGTGAAATCCCACGGCTCAACCGTGGAACTGCCTTAGATACTGTCGGCCTCGAGGAAGGTAGGGGCATGCGGAACTGTAGGTGGAGCGGTGAAATGCGTTGATATCTACAGGAACTCCGGTGGCGAAAGCGGCGTGCTGGACCTTTTCTGACGCTGAGGCGCGAAAGCTAGGGG |
| P-asv7189 | GCTAGCGTTGTTCGGATTTACTGGGCGTAAAGCGCGTGTAGGCGGGTCGTTAAGTTGGGGGTGAAATCCCGAGGCTCAACCTCGGAACTGCCTTCAAAACTGGCGACCTGGACTCAAGCAGAGGCAAGTGGAATTGCGAGTGTAGAGGTGAAATTCGTAGATATTCGCAGGAACACCAGTGGCGAAGGCGACTTGCTGGGCTTGTAGTGACGCTGAGACGCGAAAGCGTGGGG |
| P-asv7190 | GCGAGCGTTGTCCGGATTTATTGGGTTTAAAGGGTGCGCAGGTGGCCCGGCCAGTCAGTGGTGAAATGCGGCCGCTCAACGGTCGAACTGCCATTGATACTACCGGGCTTGAGAGGAGTGGAGGCTGCCGGAACGGACGGTGTAGCGGTGAAATGCATAGATATCGTCCAGAACGCCGATTGCGAAGGCAGGTGGCTACGCTTCTTCTGACACTGAGGCACGAAAGCGTGGGG |
| P-asv7191 | GCAAGCGTTACTCGGAATCACTGGGCGTAAAGGACGCGTAGGCGGATTATCAAGTCTCTTGTGAAATCTAACGGCTTAACCGTTAAACTGCTTGAGAAACTGATAATCTAGAGTAAGGGAGAGGCAGATGGAATTCTTGGTGTAGGGGTAAAATCCGTAGAGATCAAGAAGAATACCCATTGCGAAAGCGATCTGCTGGAACTTAACTGACGCTAATGCGTGAAAGCGTGGGG |
| P-asv7192 | CCTAGCGTTGTTCGGAATTATTGGGCGTAAAGCGGGCGTAGGTGGCTATGTAAGTCAAGTGAGAAAGCCTGGGGCTCAACCCCAGAAGTGCATTTGATACTGCGTAGCTTGAGTGCTAGAGAGGATAGTAGAATTGTTGGTGTAGTGGTGAAATACGTAGATATCAACAGGAATACCGGTGGCGAAGGCGGCTATCTGGCTAAGCACTGACACTCAGACCCGAAAGCGTGGGG |
| P-asv7193 | ACAAACGTTACTCGGAATTACTGGGCTTAAAGGGTGCGTAGGCTGCGCGGAAAGTTGGATGTGAAAGCCCTCAGCTCAACTGGGGAATTGCATTCAAAACCACCGTGCTCGAGGGAGACAGAGGTAAGCGGAACTCAAGGTGGAGCGGTGAAATGCGTTGATATCTTGAGGAACACCGGTGGCGAAAGCGGCTTACTGGGTCTCTTCTGACGCTGAGGCACGAAAGCTAAGGT |
| P-asv7194 | GCGAGCGTTGTCCGGAATTATTGGGCGTAAAGGGTACGCAGGCGGTTTAATAAGTCGAATGTTAAAGATCGGGGCTCAACCCCGTAAAGCATTGGAAACTGATAAACTTGAGTAGTGGAGAGGAAAGTGGAATTCCTAGTGTAGTGGTGAAATACGTAGATATTAGTAGGAATACCAGTAGCGAAGGCGACTTTCTGGACACAAACTGACGCTGAGGTACGAAGGCGTGGGG |
| P-asv7195 | GCAAGCGTTATTCGGAATCACTGGGCGTAAAGGACGCGTAGGCGGGTTGCCAAGTCAGGTGTGAAATCCTACAGCTTAACTGTAGAACTGCACTTGAAACTGGTTACCTAGAGTATGGGAGGGGGAGATGGAATTAGTGGTGTAGGGGTAAAATCCGTAGATATCACTAGGAATACCTAAAGCGAAGGCGATCTCCTGGAACATAACTGACGCTAAGGCGTGAAAGCGTGGGG |
| P-asv7196 | GCAAGCGTTATCCGGAATTATTGGGCGTAAAGCGTCCGTAGGTGGCTCATCAAGTCTTTTGTCAAATCGTGCAGCTTAACTGCATACGGGCGGAGGAAACTGAAGAGCTAGAGTGCGATAGGGGTCACAGGAATTCCCAGTGTAGCGGTGAAATGCGTAGATATTGGGAAGAACACCAGCGGCGAAAGCGTGTGACTGGGTCTGCACTGACACTGAGGGACGAAAGCTAGGGG |
| P-asv7197 | GCAAGCGTTGTCCGGAATTACTGGGCATAAAGCGCACGTAGGTGGTCCGCTAAGTCGCGTGCGAAATCCCTCGGCTCAACTGGGGAATTGCGGGCGATACTGGCGGGCTAGAGATCGGGAGGGGTGAGTGGAACTCCAGGTGGAGCGGTGAAATGCGTAGATATCTGGAGGAACACCGGTGGCGAAAGCGACTCACTGGCCCGATACTGACACTGAGGTGCGAAAGCTAGGGT |
| P-asv7198 | GCAAACGTTGTTCGGAATTACTGGGCGTAAAGCGTGTGTAGGCGGTCTTGTAAGTCGGATGTGAAAGCCCCGGGCTCAACCCGGGAAGTGCACTCGATACTGCGAGACTTGAGTATCGGAGAGGTTGGTGGAATTCTCGGTGTAGTGGTGAAATTCGTAGATATCGAGAGGAACACCGGTGGCGAAAGCGGCCAACTGGACGAATACTGACGCTGAGACACGAAAGCGTGGGG |
| P-asv7199 | GCAAACGTTGTTCGGAATCACTGGGCGTAAAGGGCGCGTAGGCGGACCTGTAAGTCAGGAGTGAAAACTCGGGGCTCAACCCCGAGCCTGCTTCTGAAACTGCATGTCTTGAGGACCGGAGAGGAAAGTGGAATTCCTAGTGTAGCGGTGGAATGCGTAGATATTAGGAAGAACACCAGCGGCGAAGGCGACTTTCTGGACGGTTCCTGACGCTGAGGCGCGAAAGCTAGGGG |
| P-asv7200 | GCGAGCGTTGTCCGGATTTATTGGGCGTAAAGAGCTCGTAGGCGGCTCGGTAAGTCGACTGTGAAAGCTCCAGGCTCAACCTGGAGAGGCCAGTCGATACTGCTGTGGCTAGAGTCCGGTAGGGGAGTGTGGAATTCCTGGTGTAGCGGTGGAATGCGCAGATATCAGGAGGAACACCAATGGCGAAGGCAGCACTCTGGGCCGGTACTGACGCTGAGGAGCGAAAGCGTGGGT |
| P-asv7201 | GCTAGCGTTGTCCGGAATCATTGGGCGTAAAGCGCGTGTAGGCGGTCCGGTAAGTCCGCTGTGAAAGTCGGGGGCTCAACCCTCGAATGCCGGTGGATACTGTCGGGCTAGAGTGCGGAAGAGGCGAGTGGAATTCCTGGTGTAGCGGTGAAATGCGCAGATATCAGGAGGAACACCAATTGCGAAGGCAGCTCGCTGGGACGTAACTGACGCTGAGACGCGAAAGCGTGGGG |
| P-asv7202 | GCAAGCGTTAACCGGAATTACTGGGCGTAAAGCGTGTGTAGGCGGTTCGGAAAGAAAGATGTGAAATCCCAGGGCTTAACCTTGGAACTGCATTTTTAACTGCCGAGCTAGAGTATGTCAGAGGGGGGTAGAATTCCACGTGTAGCAGTGAAATGCGTAGATATGTGGAGGAATACCGATGGCGAAGGCAGCCCCCTGGGATAATACTGACGCTCAGACACGAAAGCGTGGGG |
| P-asv7203 | GCTAGCGTTAATCGGATTTATTGGGCGTAAAGGGCGCGTAGGCGGGAATGAAAGTCAGATGTGAAATTCCGGGGCTCAACCTCGGAGCTGCATTTGAAACTTCATTTCTTGAGGATAGGCGGAGAAAACGGAATTCCACAAGTAGCGGTGAAATGCGTAGATATGTGGAAGAACACCGGTGGCGAAGGCGGTTTTCTAGCTTATTCCTGACGCTGAGGCGCGAAAGCATGGGG |
| P-asv7204 | GCAAGCGTTGTCCGGAATTATTGGGCGTAAAGAGCTCGTAGGCGGTCTGTCGCGTCTGCTGTGAAAACCTGGGGCTTAACCCCGGGCTTGCAGTGGGTACGGGCAGACTAGAGTGCGGTAGGGGAGACTGGAATTCCTGGTGTAGCGGTGGAATGCGCAGATATCAGGAGGAACACCGGTGGCGAAGGCGGGTCTCTGGGCCGTTACTGACGCTGAGGAGCGAAAGCATGGGG |
| P-asv7205 | GCAAGCGTTGTTCGGATTTACTGGGCGTAAAGGGCACGCAGGCGGTCTTGCCAGTCAGGGGTGAAAGCCCGGGGCTCAACCCCGGAACTGCCTCTGATACTGCGAGACTAGAGACTAGGAGAGGGTGGTGGAATTCCCAGTGTAGAGGTGAAATTCGTAGATATTGGGAGGAACACCAGAGGCGAAGGCGGCCACCTGGACTAGATCTGACGCTCAGGTGCGAAAGCGTGGGG |
| P-asv7206 | GCAAGCGTTGTCCGGATTTATCGGGTTTAAAGGGTGCGTAGGCGGCTGATTAAGTCAGTGGTGAAAGGCAGCAGCTCAACTGTTAGCACGGCCATTGATACTGGTTAGCTTGAGTACAGTCGAGGTAGGCGGAATTGAAGGTGTAGCGGTGAAATGCATAGATATCTTCAAGAACACCTATTGCGAAGGCAGCTTACTAGGCTGTAACTGACGCTGATGCACGAAAGCGTGGGG |
| P-asv7207 | GCGAGCGTTGTCCGGAATTATTGGGCGTAAGGGGCGTGTAGGCGGCTTTGCAAGTCGGAGGTGAAAGCCTGGGGCTTAACCCCAGCGAGCCTACGAAACTGCAGAGCTTGAGGGCAGTAGGGGAAAGCGGAATTCCCGGTGTAGCGGTGAAATGCGTAGATATCGGGAGGAACACCAGTGGCGAAGGCGGCTTTCTGGGCTGACCCTGACGCTGAGGCGCGAAAGCCAGGGG |
| P-asv7208 | GCGAGCGTTACCCGGATTTACTGGGTGTAAAGGGCGTGTAGGCGGTCTTTCAAGTCCGATGCTAAAGACCGAAGCTCAACTTCGGGGGTGCGTTGGATACTGTGAGGCTAGACGGTCGGAGAGGGTAGCGGAATTTCCGGAGTAGCGGTGAAATGCGCAGATACCGGAAGGAACGCCGATAGCGAAAGCAGCTACCTGGACGATTTGTGACGCTGAGGCGCAAAAGCGTGGGG |
| P-asv7209 | GCAAGCGTTATCCGGATTTACTGGGTGTAAAGGGAGCGCAGACGGCCAAGCAAGTCTGAAGTGAAATGCATGGGCTCAACCCATGAATTGCTTTGGAAACTGTTAGGCTTGAGTGTCGGAGGGGTAAGCGGAATTCCTAGTGTAGCGGTGAAATGCGTAGATATTAGGAGGAACACCGGAGGCGAAGGCGGCTTACTGGACGACAACTGACGTTGAGGCTCGAAGGCGTGGGG |
| P-asv7210 | GCTAGCGTTGCTCAGAATTACTGGGCGTAAAGGGCGCGTAGGCGGACAGTTAAGTTGGGGGTGAAGGCCCGGGGCTCAACCAATGTAAGCCCTTGAAACTGTTGGGCTTGAGTGCAGGAGAGGTAAGTGGAATTCCTAGTGTAGAGGTGAAATTCGTAGATATTAGGAGGAACACCAGTGGCGAAGGCGACTTACTGGACTGTAACTGACGCTGAGGCACGAAAGCGTGGGG |
| P-asv7211 | GCAAGCGTTATCCGGAATTATTGGGCGTAAAGGGTGCGTAGGTGGTTATTTAAGTCTTTGGTGTAAGTGCAGTGCTTAACGCTGTGATGCTAAAGAAACTGGGTAACTTGAGTGTGGTAGAGGCAAGTGGAATTCCATGTGTAGCGGTAAAATGCGTAAATATATGGAGGAACACCAGTGGCGAAGGCGGCTTGCTGGGCCACAACTGACACTGAGGCACGAAAGCGTGGGG |
| P-asv7212 | GCAAGCGTTAATCGGAATTATTGGGCGTAAAGGGCGTGTAGGTGGTCTTTTAAGTCAGGAGTTAAATTTTTGAGCTCAACTCAAAAACTGCTTTTGAAACTAATTGACTAGAGTTTAGGAGAAGAAGGTGGAATTATCAGAGTAGCGGTGAAATGCGTAGATCCTGATAGGAAGGCCAAAGACGAAGGTAGCCTTCTGGACTAATACTGACACTGAGGCGCGAAAGCTAGGGG |
| P-asv7213 | GCTAGCGTTAATCGGATTTACTGGGCGTAAAGCGTGCGTAGGCGGTTTGATAAGTCGATTGTGAAAGCCCCGGGCTTAACCTGGGAATTGCAGTTGATACTGTCAGACTCGAAAACGGTAGAGGGAGGCGGAACTCCAGGTGTAGCGGTGGAATGCGTAGATATCTGGAAGAACACCGATGGCGAAGGCAGCCTCCTGGGCCTGTTTTGACGCTGAGGCACGAAAGCGTGGGT |
| P-asv7214 | GCGAGCGTTATCCGGATTTACTGGGCGTAAAGCGCCTGTAGGCGGTTAGATAAGTCGGATGTGAAAGCCCCGTGCTCAACGCAGGGACGCCATTCGATACTGTACAACTTGAGAGCGGCAGAGGGAAACGGAATTTCTGGTGTAGCGGTAAAATGCGCAGAGATCAGAAGGAACACCGATGGCGAAGGCAGTTTCCTGGGCCGCATCTGACGCTGAGAGGCGAAAGCTAGGGG |
| P-asv7215 | ACAAACGTTGTTCGGAATCACTGGGCTTAAAGGGCGTGTAGGCGGAGCAATAGGTGTCTGGTGAAAACCCCGGTCTCAACCCGGGAATTGCTGGGCAAACCGGTGCTCTTGAGGCAGGTAGAGGTGACTGGAACAACTGGTGGAGCGGTGAAATGCGTTGATATCAGTTGGAACGCCGGCGGCGAAAGCGGGTCACTGGGCCTGTCCTGTCGCTGAGGCGCGAAAGCGTGGGG |
| P-asv7216 | GCAAGCGTTAATCGGATTTATTGGGCGTAAAGAGCACGTAGGCGGCAATGCAAGTTAGATGTGAAATTTCGGGGCTCAACCCCGAAGGTGCATTTAAAACTGCGTCGCTAGAGGATAGGCGGAGAAAGAGGAATTCCAAGTGTAGCGGTGAAATGCGTAGATATTTGGAAGAACACCTGTGGCGAAAGCGTCTCTCTAGCTTATACCTGACGCTGAGGTGCGAGAGCAAGGGG |
| P-asv7217 | GCTAGCGTTATTCGGAATCATTGGGCGTAAAGCGCATGTAGGTGGCTTTGTAAGTCAGGTGTGAAATCCCCGGGCTCAACCCGGGAAGGGCACTTGAAACTACTTAGCTTGAGTACGGAAGAGGTGAGTGGAATTCCTGGTGTAGTGGTGAAACACGTAGATATCAGGAGGAACACCGGTGGCGAAGGCGGCTCACTGGTCCTGTACTGACACTGAGATGCGAAAGCATGGGG |
| P-asv7218 | GCAAGCGTTGTTCGGAATCACTGGGCGTAAAGAATGCGTAGGCGGCTAGACAAGTCAGGAGTGAAATCCCAGGGCTCAACCCTGGAACTGCTTTTGAAACTGTCTTGCTAGAGTCCCGGAGGGGTTGGCGGAATTCCGAGTGTAGAGGTGAAATTCGCAGATATCAGGAGGAACACCGATGGCGAAGGCAGCCCCCTGGGCCTGTACTGACGCTCATGCGCGAAAGCGTGGGG |
| P-asv7219 | GCTAGCGTTGTTCGGAATTACTGGGCGTAAAGGGCGCGTAGGCTGCCATGTGCGTCAGGTGTGAAATCCCAAGGCTTAACCTTGGAACTGCATCTGAAACTGCATGGCTTGAGTGAGTAAGAGGCGGACGGAATTCCGAGTGTAGCAGTGAAATGCGTAGATATTCGGAGGAACATCGGTGGCGAAGGCGGTTCGCTGGTACTCTACTGACGCTGAGGCGCGAAAGCGTGGGG |

| | |
|---|---|
| P-asv7220 | GCGAACGTCATTCGGAATCACTGGGCTTAAAGGGTGCGTAGGCGGTTTTTCAAGTCAGGTGTGAAATCTTCCGGCTCAACCGGGAAACTGCGCTTGAAACTGATGAACTGGAGTATGCTAGGGGTGTGCGGAGCTGCCAGTGGAGCGGTGAAATGTGTTGAGATTGGCAGGAACGCCGGTGGCGAAAGCGGCACACTGGGGCATAACTGACGCTGAGGCACGAAAGCCAGGGT |
| P-asv7221 | CCGACTGGTGACCATGTTTATTGGGCCTAAAGCGTTCGTAGTCGGACGAGTAGATTTTTGGTGAAATTCCCTCGCTTAACGGGGGAGCGTGCTAAAAACACCGCTCGTCTTGGGACCGGGAGAGGTAAGGGGTACGCGATGAGTAGGGGCAAAATCCTATAATTCATTGAGGACCACCGGTGGCGAAGGCGCCTTACTTGAACGGGTCCGACGATGAGGGACGAAGGCCAAGGT |
| P-asv7222 | GCAAGCGTTATCCGGAATCATTGGGCGTAAAGGGCGCGTAGGCGGTATGTTAAGTCAGAGGTGAAATCAGCGGGCTCAACTCGTTGGCTGCCTTTGAAACTGGCGTGCTAGAGTTCGGGAGAGGAAAGCGGAATTCCTGGTGTAGTGGTGAAATGCGTAGATATCAGGAAGAACACCTATGGCGAAAGCAGCTTTCTGGACCGACACTGACGCTAATGCGCGAAAGCGTGGGG |
| P-asv7223 | GCGAGCGTTACTCGGAATCACTAGGCGTAAAGCGCAGGTAGGTTGTTGGGTAAGTCTGTTGTGAAAGCTCCTGGCTCAACTGGGAGAGGTCAACGGAAACTGCCTAGCTAGAGTATGGCAGAGGAAGCTGGAATTCCCGGTGCAGCGGTGAAATGCGTAGATATCGGGAGGAACACCTATGGCTAAAGCAGGCTTCTGGGCCATTGCTGACGCTGAGCTGCGAAAGCTAGGGG |
| P-asv7224 | GCAAGCGTTATTCGGAATTACTGGGCGTAAAGCGCGTGTAGGCGGTTTAGCAAGTCTGGTGTGAAATTCCTCGACTCAATCGAGGAAGTGCATTGGATACTGCTAGACTTGAGTACCGGAGGGGAGAACGGAATTCCTGGTGTAGAGGTGAAATTCGTAGATATGTGGAAGAACACCGGTGGCGAAGGCGGTTTTCTAGATTAGACCTGACGCTAAGGCCCGAAAGCGAGGGG |
| P-asv7225 | GCGAACGTTATTCGGTATCACTGGGCTTACAGGGTGCGTAGGCGGTCGATGAAGTAGGGTGTGAAATGCCTTGGCTCAACCAGGGCACAGCGCTCTAAACTGATTGACTGGAGCGAGCTAGGGGTGTACGGAACTTCCGGTGGAGCGGTGAAATGTGTTGATATCGGAAGGAACGCCGGTGGCGAAAGCGGTACACTGGGGCTTGTCTGACGCTGAGGCACGAAAGCCAGGGG |
| P-asv7226 | GCTAGCGTTGTTCGGAATTACTGGGCGTAAAGCGCACGTAGGCGGCGACACAAGTCAGAGGTGAAAGCCCGGGGCTCAACCCCGGAACTGCCTTTGAAACTAGGTTGCTAGAATCTTGGAGAGGTCAGTGGAATTCCGAGCGTAGAGGTGAAATTCGTAGATATTCGGAAGAACACCAATGGCGAAGGCAGCTTGCTGGGATACTCCTGACGCTAAATAGCGAAAGCGTGGGT |
| P-asv7227 | GCAAGCGTTGTTCGGATTTACTGGGCGTAAAGCGCGCGCAGGCTGATTATTAAGCCAGATGTGAAATCCCAAGGCTCAACCCTGGAACGGCCTCCAAAACTCCTGGTCTTGAGTTCGAGAGAGGTGAGTGGAATTCCGAGTGTAGAGGTGAAATTCGTAGATATTCGGAGGAACACCAGTGGCGAAGGCGGCTCACTGGCTCGATACTGACGCTGAGGTGCGAAAGTGTGGGG |
| P-asv7228 | GCGAGCGTTATCCGGATTCACTGGGCGTAAAGAGCGCGTAGGCGGTTCCGCAAGTCACCCGTGAAAGCCCCCGGCTCAACCGGGGAGGGCCGGGTGATACTGCGGGGCTTGAGGGCGGAAGAGGGAAGCGGAATTCCCGGTGTAGCGGTGAAATGCGTAGATACCGGGAGGAACACCGGTGGCGAAGGCGGCTTCCTGGTCCGTTCCTGACGCTGAGGCGCGAAGGCTAGGGG |
| P-asv7229 | TCGAGCGTTATCCGGAATTACTGGGTGTAAAGAGCTCGTAGGCGGCGAATTAAGTTAGGTATGAAAGACCGGAGCTCAACTCCGAGTTTGTGCTTAAAACTGGTTTGCTAGAATTAGGGAGAGGTAAGCGGAATTCTAAGTGTAGGGGTGCAATCCGTAGATACTTAGAGGAACACCAAAAGCGAAGGCAGCTTACTGGAACTATATTGACGCTGAGGAACGAAAGCGTGGGG |
| P-asv7230 | GCAAGCGTTGTTCGGAATAACTGGGCGTAAAGGGCACGTAGGCGGTGATACGCGTCAGATGTGAAAGACCTGAGCTTAACTCAGGAATAGCATCTGAAACGGTATGACTCGAGATCGAGAGGGGAAAAGGGAATTGTGCGTGTAGAGGTGAAATTCGTAGATATGCACAGGAACACCGGTGGCGAAGGCGCTTTTCTGGCTCGATACTGACGCTAAGGTGCGAAAGCGTGGGG |
| P-asv7231 | TCGAGCGTTGTCCGGAGTTACTGGGCGTAAAGCGCGCGCAGGTGGCTCATTGCGCCCGGCGTGAAAGCCCCCGGCCTAACCGGGGAGGGTCGTCGGGGACGGTGAGCTTGAGGGTATCAGGGGCTGGTAGAATTCCCGGTGTAGTGGTGAAATGCGTAGATATTGGAAGGAATACCAGTGGCGAAGGCGGCCACCTGGTCAGAGACTGACGCTGAGGTACGAAAGCGTGGGG |
| P-asv7232 | GCAAGCGTTGCTCGGAATTATTGGGCGTAAAGGGTAGGTAGGTGGTTACGTATGTCTGGGGTGAAATCCCTGAGCTCAACTCAGGAAGTGCCTTGGAAACGGCGTAACTAGAGTGCTAGAGAGGTTCGTAGAATTCCCAGTGTAGCGGTGAAATGCGTAGAGATTGGGAGGAATACCAGAGGCGAAGGCGGCGAACTGGATAGCAACTGACACTAAACTACGAAAGCGTGGGG |
| P-asv7233 | GCAAGCGTTATTCGGAATGACTGGGCTTAAAGGGCGCGTAGGCGGTTTGGCAAGTCAGGTGTGAGATCCCGGGGCTCAACTCCGGAACTGCGCTTGATACTGTTGGGCTAGAGTAAGGTATGGGGTCGTGGAACTCACAGTGGAGCGGTGAAATGCGTAGATATTGTGGGGAACACCGGTGGCGAAGGCGACGACCTGGGCCTTTACTGACGCTGAGGCGCGAAAGCATGGGT |
| P-asv7234 | ACAAATGTTATCCGGAATTACTGGGCGTAAAGGGTCCGCAGGCGGCTTTTTAGGTCTGATGTGACAACTCCCGGCTTAACTGGGAGAGCGCATTGGAAACCGGAAGGCTGGAATCACGTTGAGGTCGGTGGAATTCCCGGTGTAGTGGTGGAATGCGTAGAGATCGGGAGGAACCTCGAAAGCGAAGGCAGCCGGCTAAACGATGATTGACGCTGAGGGACGATAGCGTGGGT |
| P-asv7235 | GCAAGCGTTACTCGGAATCACTGGGCGTAAAGAGCGTGTAGGCGGTTTGATAAGTTGGAAGTGAAATCCTATAGCTTAACTATAGAATTGCTTCCAAAACTGTTTAACTAGAATGTGGGAGAGGTAGATGGAATTTCTGGTGTAGGGGTAAAATCCGTAGATATCAGAAGGAATACCTATTGCGAAGGCAATCCCCTGGGCCTGCACTGACGCTCATGCACGAAAGCATGGGG |
| P-asv7236 | GCTAGCGTTGTTCGGAATCACTGGGCTTAAAGCGCGTGTAGGCGGGACGTAGGGTGTCTTGTGAAAGCCCCCGGCTCAACCGGGGAATGGCTCGGCAAACCGACGTCCTTGAGGCAGGTAGAGGTAACTGGAACCTTTGGTGGAGCGGTGAAATGCGTAGAGATCAAAGGGAACGCCGGTGGCGAAAGCGGGTTACTGGACCTGTCCTGACGCTGAGACGCGAAAGCGTGGGT |
| P-asv7237 | GCTAGCGTTATTCGGTGTCACTGGGCTTAAGGCGTATGTAGGCGGAAATGTAAGTGCTTTGTGAAATCCCTCAGCTCAACTGAGGAATAGCTTGGCATACTGCGTTTCTTGAGGCAAGTAGGGGTGCATGGAACTCTAGGTGGAGCGGTGGAATGCGTAGATATCTAGAGGAACGCCGATGGAGAAATCAGTGCACTGGGCTTGTCCTGACGCTCAGATACGAAAGCGTGGGG |
| P-asv7238 | CCTAGCGTTGTTCGGAATCACTGGGCGTAAAGCGTATGTAGGCGGCCTTGCAAGTCAGGTGTGAAATCCCATGGCTTAACCATGGAACTGCATTTGAAACTGCAAGGCTTGAATATCAGAGAGGGTAGTAGAATTCCTGGTGTAGTGGTGAAATACGTAGATATCAGGAGGAACTCCGGTGGCGAAGGCGGCTACCTGGCTGAATATTGACGCTGAGATACGAAAGCGTGGGG |
| P-asv7239 | CCGAGCGTTGTCCGGAATTACTGGGTGTAAAGGGCGTGTAGGTGGGTCTCTAAGTCGATGGTGAAATCCTGCAGCTCAACTGTAGAATTGCCGTCGATACTGGAGGTCTTGAGTGCGGCAGAGGGAGACGGAATTCATGGTGTAGCGGTGAAATGTGTAGATATCATGAAGAACACCGGTTGCGAAGGCGGTCTCCTGGTCCGTTACTGACACTGAGGCGCGAAAGCGTGGGG |
| P-asv7240 | GCGAGCGTTAATCGGAATAACTGGGCGTAAAGGGTGAGTAGGCGGAAATATAAGTTAGATGTGAAAGTCCCGAGCTTAACTTGGGGAGGCCATTTAATACTGTATTTCTGGAGTTAGTGAGAGGGTGGTAGAATTCCTGGTGTAGCAGTGAAATGCGTAGATATCAGGAGGAATACCGAAGGCGAAGGCAGCCACCTGGGATATAACTGACGCTGAGTCACGAAAGCGTGGGT |
| P-asv7241 | CCGAGCGTTGTTCGGAATTACTGGGCGTAAAGGGCGCGTAGGCGGCCGTGCAAGTTGAAAGTCAAATATTGCGGCTCAACCGCAGGTCGCTTTCAAGACTGCATGGCTAGAGGTTTGCAGAAGAAACTGGAATTTTCGGTGTAGCAGTGAAATGCGTAGATATCGAAAGGAACATCGGTGGCGAAAGTGAGTTTCTAGGCAACACCAGACGCTGAGGCGCGAAAGCCAAGGT |
| P-asv7242 | GTTAGCGTTGTTCGGAATTACTGGGCGTAAAGCGATCGTAGGCGGATATTTAAGCTAGATGTGAAAGCCCTGGGCTCAACCCGGGAATTGCATTTAGAACTGGGTATCTAGAGTGTTGGAGAGGATGACGGAATTCCAAGTGTAGAGGTGAAATTCGCAGATATTTGGAGGAACATCAGTGGCGAAGGCGGTTGTCTGGCCAAACACTGACGCTGAGGGTCGAAAGCGTGGGT |
| P-asv7243 | CCGAACGTTATTCGGATTCACTGGGCATAAAGGGCGCGTAGGCGGGCTGCCAAGTCAAGGGTGAAATCCTCCGGCTTAACCGGAGAACTGCCTTTGATACTGGTGGTCTTGAGGGAGGTAGGGGCGTGTGGAACTAAGGGTGGAGCGGTGAAATGCGTTGATATCCTTAGGAACTCCGGTGGCGAAGGCGACACGCTGGACCTTTTCTGACGCTGAGGCGCGAAAGCCAGGGG |
| P-asv7244 | GCGAGCGTTGTCCGGATTTATTGGGCGTAAAGGGCATGTAGGTGGTCTTTTAAGTCAGTTGTTAAAACTATCGGCTTAACCGGTAATTGCAATTGATACTGGAGGACTTGAATTTGAGAGGGGATAGTGGAATATCAAGTGTAGCGGTGGAATGCGTAGATATTTGATAGAACACCCGAGGTGCAGACGACTATCTGGCTCATGATTGACACTGAGATGCGAAAGTATGGGG |
| P-asv7245 | GCTAGCGTTGTTCGGAATCACTGGGCGTAAAGGGAGTGTAGGCGGGGGTGCAAGTCAGAAGTGTAATCTATAGGCTTAACCTGTAGTTTGCTTCTGAGACTGTATCTCTTGAGTAGTGGAGAGGGAGATGGAATTCCAGGTGTAGCGGTGGAATGCGTAGAGATCTGGAAGAACACCAGTAGCGAAGGCGGTCTCCTGGCCACATACTGACGCTGAGGCTCGAAAGCTAGGGG |
| P-asv7246 | GCTAGCGTTGTTCGGAATTACTGGGCTTAAAGCGCATGTAGGCGGCTTGCCAAGTCAGGAGTGAAATCCCAGGGCTTAACCCTGGAATTGCTTTTGATACTAGCAGGCTAGAAGATGGTAGAGGATAATGGAATCCTTAGTGTAGGAGTGAAATCCGTAGATATTAAGAGGAACACCAGTGGCGAAGGCGGTTATCTGAGCCATTCTTGACGCTGAGATGCGAAAGTGTGGGG |
| P-asv7247 | GCTAGCGTTATTCGGAATTACTGGGCCTAAAGCGTCCGTAGCTGGCCTCGCAAGTCCGTCGGGAAATCCAGCAGCTCAACTGCTGGGCGTCCGGCGGAAACTCCGAGGCTTGGGACCGGAAGGCTCGAGGGGTACGTCCGGGGTAGGAGTGAAATCCCGTAATCCCGAACGGACCACCGATGGCGAAAGCACCTCGAGAAGACGGATCCGACGGTAAGGGACGAAAGCCAGGGT |
| P-asv7248 | GCAAGCGTTATCCGGAATCATTGGGTTTAAAGGGTCCGCAGGCGGTCAAGTAAGTCAGTGGTGAAATCTCCGAGCTTAACTCGGAAACTGCCTTTGATACTATAAGTCTTGAATACTCTGGAGGTTAGCGGAATATGTCATGTAGCGGTGAAATGCTTAGATATGACATAGAACACCTATTGCGAAGGCAGCTAGCTACGGAGTCATTGACGCTGAGGCACGAAAGCGTGGGG |
| P-asv7249 | GCAAGCATTAATCGGATTTACTGGGCGTAAAGGGTGCGTAGGCGGATTGATAAGTCAGGTGTGAAATTCAGGTACTCAATACCTGAGCTGCACTTGAAACTGTCTTTCTAGAGGTCAGGCGGAGAGAACGGAATTCCATATGTAGCGGTGAAATGCGTAGATATATGGAAGAACACCTGTGGCGAAAGCGGTTCTCTAGCTTGATACTGACGCTGAGGCACGAGAGCGTGGGG |
| P-asv7250 | TCGAGCGTTATCCGGAATTATTGGGCGTAAAGGGTGTGTAGGCGGTAGTGTTAGTCTTGTGTTAAATCTGTCGGCTCAACCGATAGGCTGCATGGGAAACGGCACAACTAAGAGGATGGAAGGGGTCTGCGGAACTCATAGTGTAGCGGTGAAATGCGTTGATATTATGGGGAACACCGAAAGCGAAGGCAGCAGACTGGTCCACTCCTGACGCTGAAACACGAAAGCGTGGGT |
| P-asv7251 | TCAAGCGTTGTTCAGATTCATTGGGCGTAAAGGGTGCGCAGGCTGTGTGGTAAGTCGGGTGTGAAATTTAGGGGCTCAACCTCTAAACTGCACCTGATACTGCCATGCTAGAGGACTGTAGAGGAGATTGGAATTCACGGTGTAGCAGTGAAATGCGTAGATATCGTGAGGAAGACCAGTGGCGAAGGCGAATCTCTGGGCAGTTCCTGACGCTCATGCACGAAGGCTAGGGG |
| P-asv7252 | CCAAGCGTTGTTCAGAATCACTGGGCTTAAAGCGCACGCAGGTGGATTATTAAGTGTCTTGTGAAATCCCACGGCTTAACTGTGGAACTGCAGGGCATACTGGTAATCTTGAGGCAAGTAGGGGTACATGGAACTCTGAGTGGAGCGGTGAAATGCGTAGATATTCAGAGGAACGCCAGAGGTGAAAACGGTGTACTGGGCTTGTTCTGACACTGAGGTGCGAAAGCGTGGGT |
| P-asv7253 | GCTAGCGTTATTCGGAATTACTGGGCGTAAAGCGCACGTAGGCGGATCAGAAAGTCAGAGGTGAAATCCCGGGGCTCAACCCCGGAACGGCCTTTGAAACTCCTGATCTTGAGTTCGAGAGAGGTGAGTGGAATTCCGAGTGTAGAGGTGAAATCCGTTGATATCGAGGGGAATACCAAAAGCGAAGGCAGCTTGCTGGAACGGTACTGACACTGAGGCATGAAACCCTGGGT |
| P-asv7254 | GCAAGCGTTGTTCGGAATCACTGGGCGTAAAGCGAACGTAGGCGGGGAAGTAAGTTAGGAGTGAAATGTATCGGCTCAACCGGTAATCTGCTTTTAATACTGCTTTTCTTGAGTATGGGAGAGGATGATGGAATTCCAGGTGTAGCGGTGGAATGCGTAGATATCTGGAAGAACGCCAGTTGCGAAGGCGGTCATCTGGCCCAATACTGACGCTGAGGTTCGAAAGTTGGGGG |
| P-asv7255 | CCGAGCGTTAATCGGAATCACTGGGCTTAAAGCGTGCGCAGGCGGACCAGTAAGCATCTCGTGAAATCCCACGGCTCAACCGTGGAACTGCGGGGTGAACTGCTGGTCTTGAGACAGGTAGGGGCAGACGGAACTATGGGTGGAGCGGTGAAATGCATAGATATTACATAGAATACCAATTGCAAAGGCAGGTTACTAACAAACGATTGACGCCGATGGACGAAAGCGTGGGG |
| P-asv7256 | GCAAGCGTTAATCGGAATTACTGGGCGTAAAGGGTCTGTAGGTGGTTTGATAAGTCAGATGTGAAAGCCCAGGGCTCAACCTTGGAACTGCATTGGATACGGTCTGGCTAGAGTCCGCGAGAGGAAGATGGAATTGTGCGTGTAGAGGTGAAATTCGTAGATATGCACAAGAACACCGGTGGCGAAGGCGATCTTCTGGAGCGGTACTGACGCTAAGGCACGAAAGTGTGGGG |
| P-asv7257 | GCAAGCGTTGTTCGGATTTACTGGGTGTAAAGGGTGTGTAGGCGGCATGGTAAGTGGGATGTGAAATCTCCGGGCTCAACCCGGAAACTGCATTCCATACTGCTAAGCTAGAGTACCGCAGAGGTAAATGGAATTCTCGGTGTAAGGGTGAAATCTGCAGATATCGAGAAGAACACCAGTGGCGAAGGCGATTTACTGGGCGGATACTGACGCTGATACACGAAAGCTAGGGT |
| P-asv7258 | ACAAGTGGTCATCACGTCTATTGGGCCTAAAGCGTTCGTAGCCGGTTTGGTAAATCTTTTGTGAAATTGTTCGGCTTAACCGGACAGCGCGCAGGGGACACTGCCAAACTCGAGACCGGGAGGCGCAAGAAGTATGCCGTGGGGACTGGTAAAATGGGATAATCCACGGTAGACTACCGATGGCGAAGGCATCTTGCGAGAACGGATCTGACGGTGAGGAACGAAAGCTAGGGG |

| | |
|---|---|
| P-asv7259 | GCAAGCGTTATTCGGAATTACTGGGCGTAAAGCGCGCGCAGGTGGCTTGATAAGTGGCAGGTGAAATCCCTCGGCTCAACCGAGGAACTGCCTGCCAAACTGTCGAGCTTGAGGCTGGGAGGGGAGAGCGGAATTCCTGGTGTAGCGGTGAAATGCGTAGATACCAGGAGGAACACCGGTGGCGAAGGCGGCTCTCTGGACCAGTTCTGACACTAAGGCGCGAAAGCGTGGGG |
| P-asv7260 | GCAAGCGTTAATCGGAATTACTGGGCGTAAAGGGCGCGTAGGCGGATAATCAAGCTGGATGTGAAAGACCTGGGCTTAACCTGGGGGTGTCATTCATAACTGGTTATCTAGAGTAGGGGAGAGGCGAGTGGAATTCACAGTGTAGCGGTGAAATGCGTAGATATTGTGAGGAACATCGATGGCGAAGGCAGCTCGCTGGCCTACTACTGACGCTGAGGCGCGAAAGCATGGGT |
| P-asv7261 | CCAAGTGTTATCCGGAGTTACTGGGCGTAAAGAGTCTGTAGCTTGATTTGTAAGTCTGTTTTCAAATATTTCGACTCAATCGAAAAAAGGGGATAGATACTGCAAATCTGGAGGTTATGAGAGGTTAGTAGAATTTCCAGTGGAGCGGTGAAATGTGTTGATATTGGAAAGAATACCAAAAGCGAAGGCAGCTAACTATTGTAAACCTGACAGTGAGAGACGAAAGCTTGGGT |
| P-asv7262 | GCAAGTGTCAATCGGAGTTATTGGGCGTAAAGCGTGCGTAGGCGGTTCGGAAAGTTAGGTGTGAAATCCCAGGGCTTAACCTTGGAATTGCGCTTAAAACTCCCGAGCTAGAGTACTGTAGAGGATAGTGGAATTTCCGGTGTAGCGGTGAAATGCGTAGAGATCGGAAGGAACACCAGTGGCGAAGGCGACTATCTGGACAGTAACTGACGCTGAGGTACGAAAGCGTGGGT |
| P-asv7263 | CCGAGCGTTGATCGGAATCACTGGGCTTAAAGGGTTCGTAGGCGGATTGACAAGTGCTTTGTGAAATCCCACGGCTCAACCGTGGAATTGCTTGGCAGACTGTCAGTCTTGAGCACGACAGGGGCCATCAGAACGCTAGGTGGAGCGGTGAAATGCGTAGATATCTAGCGGAATGCCGAAGGTGAAGACAGATGGCTGGGTCGTCGCTGACGCTCAGGAACGAAAGCATGGGT |
| P-asv7264 | GCGAGCGTTGTCCGGAATCACTAGGCGTAAAGGGCAAGTAGGTGGAGCTATAAGTCAGATGTTAAAGGCATCGGCTCAACCGATGTACGGCATTCGATACTGTAGTTCTAGAGTGGGGTAGAGGAAGACGGAATTCCTGGTGTAGCGGTGGAATGCGCAGATATCGGGAAGAACACCGATGGCGAAGGCAGTCTTCTGGGCCTCAACTGACGCTAAATTGCGAAGGTGTGGGT |
| P-asv7265 | GCGAGCGTTGTCCGGATTCATTGGGCTTAAAGGGTACGTAGGCGGCGGCCTAAGTCAGCGGTGAAAGGCCCCGGCTCAACCGGGGGAGCGCCGTTGATACTGGGTCGCTTGAGTGCCGCGTGGGTGGGCGGAATTCCTCATGTAGCGGTGAAATGCATAGATATGAGGAGGAACCCCGATAGCGAAGGCAGCCCACTGCACGGTTACTGACGCTGAGGTACGAAAGCGTGGGG |
| P-asv7266 | GCAAGCGTTAATCGGAATTACTGGGCGTAAAGCGTGCGTAGGTGGTCTGTTAAGTCCGATGTTAAAGCCCGGGGCTCAACCCCGGCACGGCGTTGGATACTGGCAGGCTTGACGGCTGGAGAGGCGAGTGGAATTACCAGTGTAGCGGTGGAATGCGTAGATACTGGTAGGACCACCTATGGCGAAGGCAGCTCGCTGGACAGCACGTGACGCTGAGGCGCGAAAGTGTGGGG |
| P-asv7267 | GCAAGCGTTATTCGGAATTATTGGGCGTAAAGGGCGCGCAGGCGGTCTTGTCCGTCAGGTGTGAAAGCTCGGGGCTCAACCCCGGAAGTGCACTTGAAACAGCAAGACTTGAATACGGGAGAGGAAAGCGGAATTCCTGGTGTAGAGGTGAAACTCGTAGATATCAGGAGGAACACCGATGGCGAAGGCAGCTTTCTGGACCGATATTGACGCTGAGGCGCGAAGGCGTGGGT |
| P-asv7268 | GCAAGCGTTACTCGGAATCACTGGGCGTAAAGCGTCTGTAGGCGGTGTGATGCGTCTGATGTGAAAACACTGGGCTCAACCCAGTATCCGCATTGGAAACGGTCACGCTCGAGAGACTCAGAGGCACCTGGAATGCTGTGTGTAGGGGTAAAATCCGTAGATACACAGTAGAACGCTCAAAGCGAAAGCAGGGTGCTGGGGGTTTTCTGACGCTGAGAGACGAAAGCGTGGGG |
| P-asv7269 | GCGAACGTTGTTCGGAATCACTGGGTTTAAAGGGCGCGTAGGCGGACTGTCAAGTCCAAGTTGAAATCCCCCAGCTCAACTGGGGAACAGGCCTGGATACTGACAGTCTCGAGGGAGGCAGGGGCGTGTGGAACGCTTGGTGGAGCGGTGAAATGTGTTGATATCAAGCGGAACTCCGGTGGCGAAGGCGACACGCTGGACCTCTTCTGACGCTGAGGCGCGAAAGCGTGGGG |
| P-asv7270 | GCTAGCGTTGCTCGGAATCACTGGGCTTACAGGGTGCGTAGGCTGCCATTCAGGTCCGATGTGAAATCCCACAGCTCAACTGTGGAATGGCATTGGAAACCGATTGGCTTGAGTGAGCTAGAGATGCACGGAACTTCCGGTGGAGCGGTGAAATGTGTTGAGATCGGAAGGAACACCAGAGGCGAAGGCGGTGCATTGGGGCTTGTCTGACGCTGAGGCACGAAAGCCAGGGG |
| P-asv7271 | GCAAGCGTTGTCCGGATTCATTGGGTTTAAAGGGTGCGTAGGCGGTTTATTAAGTCAGTGGTGAAAGTCTGCGGCTCAACCGTAGAATTGCCACTGATACTGATAGACTTGAGTTAAGTTGAGGCAGGCGGAATTCGTGGTGTAGCGGTGAAATGCATAGATACCACGAAGAACACCGATAGCGTAGGCAGCTTGCTAAGCTTATACTGACGCTGAGGCACGAAAGCGTGGGG |
| P-asv7272 | GCAAGCGTTAATCGGAATTATTGGGCGTAAAGCGCGCGCAGGTGGTTCGTCGCGTCCGAGGTGAAATCTCCCGGCTCAACTGGGAGGGGTCTTCGGATACGGGCGGACTTGAGGAGAGCAGAGGTAGGCGGAATGCTGGGTGTAGTGGTGAAATGCGTAGATATCCAGCGGAACACCAGTGGCGAAGGCGGCCTACTGGGCTCCACCTGACACTGAGGCGCGAAAGCGTGGGG |
| P-asv7273 | GCTAGCGTTGCTCGGAATTACTGGGCGTAAAGGGCATGTAGGCGGATGAGCAAGCTTGGTGTGAAATGCTACAGCTTAACTGTGGGACAGCATTGAGAACTGCACATCTTGAATTACTGAAGGGTAACTAGAATTCCACGTGTAGGGGTGAAATCTGTAGATATGTGGAAGAATACCAATGGCGAAGGCAAGTTACCGGCAGATGATTGACGCTGAGGTGCGAAAGTGCGGGG |
| P-asv7274 | GCAAGCGTTGTTCGGAATCACTGGGCGTAAAGGGTCCGTAGGCGGATACACAAGTCAGATGTTTAATTTCGAGGCCTAACTTCGAGCTTGCATTTGAAACTGTGTATCTTGAATAGATCAGAGGTTGATGGAATACTTGGTGTAGCGGTGGAATGCGTAGATATCAAGTAGAACACCAATGGCGAAGGCAGTCAACTGGGATCATATTGACGCTGAGGGACGAAAGCGTGGGT |
| P-asv7275 | GCTAGCGTTGTTCGGAATTACTGGGCGTAAAGCGCACGTAGGCGGCTTTGTAAGTCAGAGGTGAAAGCCTGGAGCTCAACTCCAGAACTGCCTTTGAGACTGCATCGCTTGAATCCAGGAGAGGTGAGTGGAATTCCGAGTGTAGAGGTGAAATTCGTAGATATTAGGAGGAACACCGGTGGCGAAGGCGACTCTCTGGGCCGATACTGACGCTCAAGTGCGAAAGCGTGGGG |
| P-asv7276 | GCAAGCGTTATTCGGAATTACTGGGCGTAAAGGGCGCGTAGGCGGCTATTTAAGTTATTTGTTAAAGACCACTGCATACCGGTGGGGAAGCAAGTAAAACTGGATGGCTAGAGGGTAAGAGAGAGAAGTAGAATTCTCGGAGTAGCGGTAAAATGCGTAGATCTCGAGAGGAATACCGGTGGCGAAGGCGGCTTCTTGGCTTATCCCTGACGCTGAGGTGCGAAAGCGTGGGT |
| P-asv7277 | CCAAGCGTTGTTCGGAATCATTGGGCGTAAAGCGGGTGTAGGCGGCTTCATAAGTCAGGAGTGAAAGCCCTGGGCTCAACCCAGGAAGTGCTTTTGATACTGTGAAGCTTGAGTGTGGGAGAGGATAGTGGAATTCCAGGTGTAGTGGTGAAATACGTAGATATCTGGAGGAACACCGGAGGCGAAGGCGGCTATCTGGCCCAACACTGACGCTCAGACCCGAAAGCGTGGGG |
| P-asv7278 | GCAAATGTTGTTCGGAATCACTGGGCGTAAAGGGAGCGTAGGCGGGTTTTTAAGTTAGGTGTTTAATCCCGGGGCCCAACCCCGGACAGGCACCTAAAACTGGGAATCTTGAATAGTGGAGAGGACGGCGGAATTGGGGGTGTAGCGGTGGAATGCGTAGAGATCCCCAAGAACACCAGTTGCGAAGGCGGCCGTCTGGACACATATTGACGCTGAGGCTCGAAAGCATGGGG |
| P-asv7279 | GCTAGCGTTATTCGGAATTACTGGGCGTAAAGCGCGCGTAGGCGGCGAAGCAAGTCAGGTGTGAAATCCCTCGGCTCAACCGAGGAACTGCATCTGATACTGCTTTGCTTGAGTCCGGTAGGGGGTAGCAGAATTCCCGGTGTAGCGGTGGAATGCGTAGATATCGGGAGGAATACCGATGGCGAAGGCGGCTACCTGGGCCGGTACTGACGCTGAGGCGCGAAAGCTAGGGG |
| P-asv7280 | GCAAGCGTTATTCGGAGTTACTGGGCGTAAAGAGTATGTAGGCGGTTTATTAAGTTAGTTGTTAAATACCACGGCTCAACTGTGGAAAGGCAGCTGATACTGATAGGCTAGAGCGCAGAAGAGAGGAGTGGAATTCCCGGAGTAGCGGTAAAATGCGTAGATCTCGGGAGGAACACCGGTTGCGAAGGCGGCTTCTTGGTCTGTGGCTGACGCTAAAGTACGAAAGCGTGGGT |
| P-asv7281 | GCAAGCGTTATTCGGAATTATTGGGCGTAAAGGGCGCGTAGGCGGTCTTGTCGGTCAGATGTGAAAGCCCGGGGCTCAACCCTGGAAGTGCATTTGAAACAGCAAGACTTGAGTACGGGAGAGGAAAGCGGAATTCCTGGTGTAGAGGTGAAATTCGTAGATATCAGGAGGAACACCGATGGCGAAGGCAGCTTTCTGGACCGATACTGACGCTGAGGCGCGAAGGCGTGGGT |
| P-asv7282 | GCAAGCGTTGTTCGGAATCATTGGGCGTAAAGCGCGCGCAGGCGGCCGAGCGGGTCCGTTGTGAAACCCCGGGGCTTAACTCCGGAGGCGCATCGGAAACTGTTCGGCTTGAGTCCGGGAGGGGCCTGTGGAATTCCTGGTGTAGGGGTGAAATCCGTAGAGATCAGGAGGAACACCGGCGGCGAAAGCGACAGGCTGGACCGGGACTGACGCTGAGGCGCGAAAGCGCGGGT |
| P-asv7283 | ACAAGTGTTATTCATCATAAATGGGTTTAAAGCGTACCTAGACGGCCTGTTAAGTCAGAGGTGAAAGACGGTAGCTCAACTATCGCAGTGCCTTTGATACTGACGGGCTTGAATAAACTAGAGGTAGGCGGAATGAGACAAGTAGCGGTGAAATGCATAGATATGTCTCAGAACACCGATTGCGAAGGCAGCTTACTATGGTTTTATTGACGCTGAGGCACGAAAGCGTGGGG |
| P-asv7284 | CCAAGCGTTGTTCGGAATCACTGGGCGTGAAGCGCATGTAGGTGGCTTTTCAAGTCAACTGTGAAAGCCCCGGGCCTAACCCGGGAAGTGCAGTTGATACTGTTGAGCTTGAGTACGAGAGAGGATAGTGGAATTCCAGGTGTAGTGGTGAAATACGTAGATATCTGGAGGAACATCGGTGGCGAAGGCGGCTATCTGGCTCGATACTGACACTGAGATGCGAAAGCGTGGGG |
| P-asv7285 | GCGAGCGTTATCCGGATTTATTGGGTTTAAAGGGAGCGCAGACGGGAGATTAAGTCAGTTGTGAAAGTTTGCGGCTCAACCGTAAAATTGCAGTTGATACTGATTTCCTTGAGTGCAGTTGAGGCAGGCGGAATTCGTGGTGTAGCGGTGAAATGCTTAGATATCACGAAGAACCCCGATTGCGAAGGCAGCTTGCTAAACTGTAACTGACGTTCATGCTCGAAAGTGTGGGT |
| P-asv7286 | GCGAGCGTTATCCGGATTTATTGGGTTTAACGGGTGCGTAGGCGGCTTTCTAAGTCAGCGGTGAAAGGTTCGGGCTCAACCCGGACATTGCCGTTGAAACTGGGGAGCTTGAGTATGGATGAGGTAGGCGGAATGCGTGGTGTAGCGGTGAAATGCATAGATGTCACGCAGAACTCCGATTGCGAAGGCAGCTTACTAGGGCACTACTGACGCTGAGGCGCGAAAGCGTGGGG |
| P-asv7287 | GCTAGCGTTATTCGGAATTACTGGGCGTAAAGCGCACGTAGGCGGCTTTGTAAGTTAGAGGTGAAAGCCTGGAGCTCAACTCCAGAATTGCCTTTAAGACTGCATCGCTTGAATCCAGGAGAGGTGAGTGGAATTCCAGGTGTAGCGGTGATATGCATAGATATCAGGAGGAACACCCATGGCGAAGGCAGCAAACTGGGCATCATCTGACACTGAGGCACGAAAGCGTGGGG |
| P-asv7288 | GCTAGCGTTGCTCGGAATCACTGGGCGTAAAGGGTGCGTAGGTGGTTTTAAAAGTCTTTTGTTAAATCTTTAAGCTTAACTTAAATGCTGCAAAAGATACTCTATTACTTGAGTATAGGAGAGGTAAAGGTAATTCCCAGTGGAGCGGTGAAATGCGTAGATATTGGGAGGAGCACCAAAGGCGAAGGCACTTTACTGGAATATTACTGACACTGAGAGACGAAAGCTAGGGG |
| P-asv7289 | GCAAGCGTTATCCGGGATTATTGGGCGTAAAGCGTTCGTAGGCGGTCTATTAAGTCTGAAGTTAAAGCCCGGGGCTCAACCCCGGCCCGCTTTGGATACTGATAGACTAGAGTTATAGAGAGGTAAGTGGAACTCCATGTGAAGCGGTGGAATGCGTAGATATATGGAAGAACACCAATGGCGAAGGCAACTTACTGGCTATACACTGACGCTGAGGAACGAAAGCGTGGGG |
| P-asv7290 | GCGAACGTTACTCAGATTCACTGGGCTTAAAGCGCGTGTAGGCGGACACGCACGTCCGAGGCTGAAAGCCCGGGGCTCAACCTCGGAATAGCATTTGGAACTGGCAGACTAGAGTACTGTAGAGGGGGGTAGAATTTCAGGTGTAGCGGTGAAATGCGTAGAGATTGGGTGGAACACCGGTGGCGAAGGCGGCTATCTGGCTCACAACTGACGATCAGGCACGAAAGCGTGGGG |
| P-asv7291 | GCAAGCGTTGTCCGGATTTATTGGGTTTAAAGGGTGCGTAAGCGGATAGATAAGTTCGTGGTGAAATCGTTCGGCTTAACCGGATACCCGCCAAGGATACTGTTTATCTTGAGATTGGATGAGGAAGTTAGAATGCGTGGTGTAGCGGTGAAATGCATAGATATCACGCAGAATACCGATTGCGAAGGCAGACTTCTGGTCCATATCTGACGCTGAGGCACGAGAGCGTGGGT |
| P-asv7292 | CCGAGTGATGGCCGATCTTATTGGGCCTTAAGCGTCCGTAGCCAGCCGAACAGGTCCGTTGGGAAATCTGCTCGCTCAACGAGCAGGCGTCCGGCGGAAACCAGTCGGCTTGGGACCGAGAGGACCGAAGGGTACGTCCGGGGTAGGAGTGAAATCCCGTAATCCTGGACGGACCACCGATGGCGAAAGCACTTCGGTAGATCGGATCCGACGGTGAGGGACGAAAGCTAGGGT |
| P-asv7293 | GCAAGCGTTGTTCGGAATCATTGGGCGTAAAGGGCTCGTAGGCGGACTGGCAAGTCTGACGTGAAATCTCCACCGCTCACCGGGGAGAGGTCATTGGATACTACTAGTCTTGAGTTCGGTAGAGGGTCGTGGAATTTCGGGTGTAGCGGTGGAATGCGTAGATATCCGAAAGAACACCGGTGGCGAAGGCGGCGACCTGGGCCTGAACTGACGCTGAGGAGCGAAAGCTAGGGT |
| P-asv7294 | GCAAGCGTTGTCCGGAATCATTGGGTGTAAAGGGTGCGCAGGCGGGCGTCTGCGTCGGGGGTGAAATCTTGCCGCTTAACGGCAAAATGGCCTTCGATACCAGGCGTCTTGAGTACAGGAGAGGCTGGTAGAATTCGTGGTGTAGCGGTGTAATGCATAGAGATCACGAAGAATACCGGTGGCGAAGGCGGCCAGCCAGTCTGTAACTGACGCTGAGGCACGAGAGCGTGGGG |
| P-asv7295 | ACGAGTGGTGGTCATTATTATTGGGCCTAAAGCGTCCGTAACCGGTCCTGTAAATGCCTGGGTAAATCGATGCGCTTAACGTATCGAATTCCGGGTAGACTGCAGGACTTGAGACCGGGAGAGGTGCGGGGTACTCGGGGGGTAGGGGTAAAATCCTGTAATCCTCCGGGGACGACCAGTGGCGAAGGCGCCGCACTGGAACGGCTCTGACGGTGAGGGACGAAGCCCTGGGG |
| P-asv7296 | GCAAGCGTTAATCGGAATCACTGGGCGTAAAGGGTGCGTAGGCGGTGCATTAGGTCTGTCGTGAAATCCCCGGGCTCAACTTGGGGATTGCATTGGTGACTGCACGGCTAGAGTGTGTCAGAGGGGGGTAGAATTCCACGTGTAGCAGTGAAATGCGTAGAGATGTGGAGGAATACCGATGGCGAAGGCAGCCCCCTGGGATAACACTGACGCTCATGCACGAAAGCGTGGGG |
| P-asv7297 | GCAAACGTTATTCGGAATCACTGGGCTTAAAGGGTGCGCAGGCGGTCTGTTAAGCAGGGTGTGAAAGCCCCCGGCTCAACCGGGGAATTGCGCCCTGAACTGGCAGGCTTGAGTGAGATAGAGGTGTGCGGAACTTCCGGTGGAGCGGTGAAATGTGTTGATATCGGAAGGAACGCCAGAGGCGAAAGCGGCACACTGGGTCTCAACTGACGCTGAGGCACGAAAGCCAGGGG |

| | |
|---|---|
| P-asv7298 | GCGAGCGTTGTCCGGAATTATTGGGCGTAAAGGGTACGCAGGCGGTTTAATAAGTCGAATGTTAAAGATCGGGGCTCAACCCCGTAAGGCATTGGAAACTGATAAACTTGAGTAGTGGAGAGGAAAGTGGAATTC<br>CTAGTGTAGTGGTGAAATACGTAGATATTAGGAGGAATACCAGTAGCGAAGGCGACTTTCTGGACACAAACTGACGCTGAGGTACGAAAGCGTGGGG |
| P-asv7299 | GCGAGCGTTATCCGGAATCACTGGGCGTAAAGGGCGCGTAGGCGGTTTGTTAAGTCCGATGTTAAAGCGTGAGGCTCAACCTCATCACGGCGTTGGATACTGGCAAGTTGGACGGTTAGAGAGGAAGGTAGAAT<br>TACCAGTGTAGCGGTGGAATGCGTAGATACTGGTAGGAATACCCATTGCGAAGGCAGCCTTCTGGACAACACGTGACGCTGAGGCGCGAAAGTGTGGGG |
| P-asv7300 | GCAAGCGTTGTCCGGAATTACTGGGTGTAAAGGGAGCGTAGGCGGGAATGCAAGTTGAATGTTAAATCTATGGGCTCAACCCATATACGCGTTCAAAACTGCATTTCTTGAATGAAGTAGAGGTAGGCGGAATTC<br>CTAGTGTAGCGGTGAAATGCGTAAATATTAGGAGGAACACCAGTGGCGAAGGCGGCCTACTGGGCTTTAATTGACGCTGAGGCTCGAAAGCGTGGGT |
| P-asv7301 | GCGAGCGTTGTTCGGAATTATTGGGCGTTAAGGGTACGTAGGCGGCTTGTTAAGTTTGGCGTTAAATTCTGGGGCTCAACCCCAGCAAGCGTTGAAAACTGGCGAGCTTGAGTAGTAGAGGGGACAGTGGAATT<br>CCTAGTGTAGCGGTGAAATGCGTAGAGATTAGGAAGAATACCGGTGGCGAAGGCGACTGTCTGGATACATACTGACGCTGAGGTACGAAAGCGTGGGG |
| P-asv7302 | GCAAGCGTTGTTCGGAATCACTGGGCTTAAAGCGTGCGCAGGCGGATCTGAAAGTCAGGAGTGAAATCCCGGGGCTCAATCCCGGAATTGCTCTTGAAACTCCGGATCTAGAATATGGTAGAGGTTAGGAGAATT<br>CCTAGTGTAGAGGTGAAATTCGCAGATATTAGGAGGAATACCAGTGGCGTAGGCGCCTAACTGGACCATTATTGACGCTCATGTACGAAAGCGTGGGG |
| P-asv7303 | CCAAGCGTTGCCCGGATTTATTGGGTTTAAAGGGTGCGCAGGCGGCTCAATAAGTCAGTGGTGAAAGCCGGCAGCTTAACTGTCGAATTGCCATTGATACTGTTGAGCTTGAGTATAGTTGAGGTTGGCGGAATG<br>TATCATGTAGCGGTGAAATGCTTAGATATGATACAGAACACCGATTGCGAAGGCAGCTGACTAAGCTATTACTGACGCTCATGCACGAAAGCGTGGGG |
| P-asv7304 | ACAAGTGGGAACCACGTTTATTGGGTCTAAAGCATCCGTAGCCGGTTTGGCAAATGCCCTGTGAAATCGATTCGCTCAACGGATCGGCGTGCAGGGTAGACTACCAAACTAGGGACCAGGGGAGGGAAGCGGTA<br>TTCCTGAGGGAGCGGTAAAATGTGATAATCTCAGGAGGACCACCGATGGCGAAGGCAGCTTCCTAAAATGGATCCGACGGTGAGGGATGAAAGCTAGGGG |
| P-asv7305 | CCGAGCGTTATCCGGAATCATTGGGTTTAAAGGGTCCGTAGGCGGCTATATAAGTCAGTGGTGAAATCCGTTAGCTCAACTAACGAATTGCCATTGATACTGTATAGCTTGAATAATTGTGAAGTAGCTAGAATATG<br>TAGTGTAGCGGTGAAATGCGTAGATATAGGAAGGAACACCAGTGGCGAAGGCGACCACCTGGACTGATACTGACACTGAGGTGCGAAAGCGTGGGG |
| P-asv7306 | CCGGGCGTTATCCGGAATTATTGGGTTTAAAGGGAGCGCAGGCGGGAGTATAAGTCAGCTGTTAAATATCAGAGCCCAACTCTGTTATGCAGTTGAAACTATATTTCTTGAGTACGCACAGGGATGGTGGAATTCA<br>GGGTGTAGCGGTGAAATGCTTAGATATCCTGAAGAACTCCGATCGCGAAGGCAGCCATCCGGAGCGTAACTGACGCTGAGGCTCGAAGGTGCGGGT |
| P-asv7307 | GCAAGCGTTATCCGGAATTATTGGGCGTAAAGCGTCCGTAGGTGGTTTTTCAAGTCAGTTGTTAAAGCGTGGAGCTTAACTCCATAAGGGCAATTGAAACTGAGAGGCTAGAGTGCGATAGGGGCAAGGGGAATT<br>CCCAGTGTAGCGGTGAAATGCGTAGATATTGGGAAGAACATCGGTGGCGAAAGCGCCTTGCTGGGTCTGCACTGACACTGAGGGACGAAAGCTAGGGG |
| P-asv7308 | GCAAGCGTTATCCGGAATTACTGGGCGTAAAGGGCGTGCAGGCGGTCTTTTAAGTTAGGCGTGAAAGCTCCCGGCTAAACCGGGAGAGGCCGTCTGATACTGGGAGACTAGAGTATGGTAGAGGCGTGTAGAAT<br>TCCGCGTGTAGTGGTGAAATGCGTAGAGATGCGGAGGAATACCAGTGGCGAAGGCGGCACGCTGGGCCATAACTGACGCTAAGACGCGAAAGCATGGAT |
| P-asv7309 | GCAAGCGTTGTCCGGATTCATTGGGTTTAAAGGGTGCGTAGGCGGCTCTTTAAGTCAGCAGTGAAATACCCCGGCTTAACCGGGGGGGTGCTGTTGATACTGATAAGCTTGAGTACAGACGAGGCAGGCGGAAT<br>TGACGGTGTAGCGGTGAGATGCATAGATATCGTCAAGAACACCTATAGCGAAGGCAGCTTGCTAGACTGGCACTGACGCTGAGGCACGAAAGCGTGGGG |
| P-asv7310 | GCAAGCGTTGCTCGGAATTACTGGGCGTAAAGGGTCCGCAGGCGGTCAAGCAAGTTAGGTGTGAAAGCTCAAGGCTTAACCTTGGAATTGCATCTAAAACTGCATGACTAGAGTACAAGAGAGGTTGGCGGAATT<br>CCCGGTGTAGCGGTGAAATGCGTAGATATCGGGAGGAACACCAGTGGCGAAGGCGGCCAACTGGCTTGGTACTGACGCTCAGGGACGAAAGCGTGGGG |
| P-asv7311 | GCGAGCGTTATCCGGATTTACTGGGTGTAAAGCGCATGCAGGCGGTTCGTTAAGTTGGGTGTGAAAGCTCCTGGCTCAACTGGGAGAGGTCGCTCAAGACTGGCGAACTAGAGGAAGGTAAGGGAAAGCGGAA<br>TTCCGGGTGTAGTGGTGAAATGCGTAGATATCCGGAGGAACACCAGTGGCGAAGGCGGCTTTCTGGACCCCCCCTGACGCTCAGATGTGAAAGCTAGGGT |
| P-asv7312 | GCAAGCGTTGTCCGGAATAACTGGGCGTAAAGCGCATGTAGACGGCATGATAAGCCAAATGTGAAATGTAGGGGCTCAACCCCTGCACTGCGATTGGAACTGTCAAGCTTGAGTGCAGGAGAGGCAAATGGAAT<br>TCCCAGTGTAGCAGTGAAATGCGTAGAGATTGGGAGGAACACCAGTGGCGAAGGCGATTTGCTGGACTGCAACTGACGTTGAGATGCGAAAGCTAGGGG |
| P-asv7313 | ACAAGCGTTATCCGGAATCATTGGGTTTAAAGGGTCCGTAGGCGGCCTTATAAGTCAGTGGTGAAATCTCCTAGCTCAACGGGTGGGCGACCGACGGAAACTGCGAGGCTTGGGACCGGGAGGCTTGAAGGGT<br>ACGTCCGGGGTAGGAGTGAAATCCCGTAATCCTGGACGGACCACCGATGGGGAAACCACTTCAAGAGAACGGATCCGACAGTGAGGGACGAAAGCTAGGGT |
| P-asv7314 | GCAAGCGTTAATCGGAATTACTGGGCGTAAAGCGCACGTAGGCGGTTTTTTAAGTCAGATGTGAAAGCCCCGGGCTCAACCTGGGAATTGCATTTGAAACTGGAAAACTAGAGTGTGTGAGAGGGGGGTAGAATT<br>CCAAGTGTAGCGGTGAAATGCGTAGATATTAGGAGGAACACCAGTGGCGAAGGCGGCTTTCTGGCCTGGTCCTGACGCTGAGGAGCGAAAGCCAGGGG |
| P-asv7315 | GCGAGCGTTATCCGGAATTATTGGGTTTAAAGGGTGCGTAGGTTGCAAGGGAAGTCAGGGGTGAAAAGCTATGGCTCAACTATGGTCTTGCCTTTGAAACTCTCTAGCTAGAGTGTACTGGAGGTACGTGGAACG<br>TGTGGTGTAGCGGTGAAATGCATAGATATCACACAGAACTACGATTGCGCAGGCAGCGTACTACATTACAACTGACACTGAAGCACGAAAGCGTGGGT |
| P-asv7316 | GCTAGTGTTACTCAACTTGACTGGGCGTAAAGGGCGTGAAGGTGGTATGGTACGTTTTATTGTAAATACTCAAACATAATTTGAGGAGCTTTAAAATATGGCTATGCTTGAGTTTATTTGAAGAAAGTTGTACTTCTA<br>GTGTAGAGGTGAAATTTGCAAAAATTAGAAGGACAGTCCACGGGCGAAAGCGACTTTCTACTATAAACTGACACTATAGCGCGAAAGCATAGGT |
| P-asv7317 | GCGAACGTTATTCGGAATCACTGGGCTTAAAGCGCGCGTAGGCGGGCCGGAACGTCCGTAGCTGAAATCCCCCGGCTCAACCGGGGAAGTGGCGTGGAAACGACCGGCCTGGAGTGGAGGAGGGGGATCCGG<br>AACTCCCGGTGGAGCGGTGAAATGCGTTGAGATCGGGAGGAACGCCGGCGGCGAAAGCGGGATCCTGGACTTCGACTGACGCTGAGGCGCGAAAGCCAGGGG |
| P-asv7318 | TCGAGCGATGGCCGATCTTATTGGGCCTAAAGCGTCCGTAGCTGGCCTGGCAAGTCCGTCGGGAAATCCACCCGCTCAACCTGGGAATTGCGCTTGTGACTGCAAAGCTGGAGTGCGGCAGAGGGGGATGGAA<br>TTCCGCGTGTAGCAGTGAAATGCGTAGATATGCGGAGGAACACCGATGGCGAAGGCAATCCCCTGGGCCTGCACTGACGCTCATGCACGAAAGCGTGGGG |
| P-asv7319 | CCGAGCGTTATCCGGATTTATTGGGTTTAAAGGGTGCGTAGGCGGAAATTTAAGTCAGTGGTGAAAGTTTAGGGCTCAACCTTAAAATTGCCATTGAAACTGGGTTTCTTGAGTACGGTAGAGGTAGGCGGAATTT<br>GTGGTGTAGCGGTGAAATGCTTAGATATCACGAAGAACTCCGATTGCGAAGGCAGCTTACCGGACCGTAACTGACGCTGAGGCACGAGAGTGTGGGG |
| P-asv7320 | GCAAGCGTTATCCGGATTTACTGGGTGTAAAGGGAGCGCAGACGGTTGCGCAAGTCTGAAGTGAAATCCCGAGGCTTAACCACGGGACTGCTTTGGGAACTGTGCGACTTGAGTATCGGAGGGGCAGGCGGAA<br>TTCCTAGTGTAGCGGTGAAATGCGTAGATATTAGGAAGAACACCGGTGGCGAAGGCGGCCTGCTGGACGAAAACTGACGTTGAGGCTCGAAGGCGTGGGG |
| P-asv7321 | GCAAGCGTTGTCCGGATTTATTGGGTTTAAAGGGTGCGCAGGCGGCTCATTAAGTTAGTGGTGAAAGCGTCCTGCTTAACGGGATACTCGCCATTAATACTGATGAGCTTGAGTGTACGTGAGGAAGTTGGAATG<br>CGTGGTGTAGCGGTGAAATGCTTAGATATCACGCAGAACACCGATTGCGAAGGCAGACTTCTGGCGTAATACTGACGCTCATGCACGAAAGCGTGGGG |
| P-asv7322 | GCGAGCGTTGTCCGGATTTATTGGGCGTAAAGGGCGTGTAGGCGGTTTGGTAAGTCAGAGGTGAAAACCCGGGGCTCAACCCCGAGGACTGCCTCTGATACTGCTGAACTTGAGTGCTCTAGAGGGCAGTGGA<br>ATTCCAGGTGTAGCAGTGAAATGCGTAGATATCTGGAGGAACACCAGTGGCGAAGGCGACTGCCTGGGGAGTAACTGACGCTGAGGCGCGAAAGCTGGGGG |
| P-asv7323 | GCGAGCGTTATCCGGAATCACTGGGCGTAAAGGGCGCGTAGGCGGTTTGTTAAGTCCGATGTCAAAGCGTGAGGCTCAACCTCATAGAGGCGTTGGGTACTGGCGAGCTGGACGGTTGGAGAGGAAGGTAGAA<br>TTACCAGTGTAGCGGTGGAATGCGTAGATACTGGTAGGAATACCCATTGCGAAGGCAGCCTTCTGGACAACACGTGACGCTGAGGCGCGAAAGTGTGGGG |
| P-asv7324 | GCTAGCGTTGCTCGGAATCACTGGGCGTAAAGGGTGCGTAGGCGGTTTGTCGCGTCGTCTGTGAAATTCTGCAACTCAATTGCAGGCGTGCAGGCGATACGGGCAGACTTGAGTACTACAGGGGAGACTGGAAT<br>TCCTGGTGTAGCGGTGAAATGCGCAGATATCAGGAGGAACACCGGTGGCGAAGGCGGGTCTCTGGGTAGTAACTGACGCTGAGGAGCGAAAGCATGGGT |
| P-asv7325 | CCTGGCGTTATCCGGATTTATTGGGTTTAAAGGGAGCGCAGGCTGCACTTTAAGCGTGTTGTGAAATGTACCGGCTCAACCGGTAACGTGCAGCGCGAACTGGGGTGCTTGAGTACGAAGAGGGAAAGCGGAAC<br>TCGTGGTGTAGCGGTGAAATGCTTAGATATCACGAGGAACTCCGATCGCGAAGGCAGCTTTCCGTTTCGGTACTGACGCTGAGGCTCGAAAGTGCGGGT |
| P-asv7326 | GCAAGCGTTAATCGGAATTACTGGGCGTAAAGCGTGCGCAGGCGGTTATGCAAGACAGAGGTGAAATCCCCGGGCTCAACCTGGGAACTGCCTTTGTGACTGCATAGCTAGAGTACGGTAGAGGGGGATGGAAT<br>TCCGCGTGTAGCAGTGAAATGCGTAGATATCTAGAGGAACACCGGTGGCGAAGGCGGCTCTCTGGTCCGTTTCTGACGCTGAGGTGCGAAAGCTAGGGT |
| P-asv7327 | GCAAGCGTTATCCGGAATTATTGGGCGTAAAGAGTCTGTAGGCGGCATGTTAAGTCGGGTATTAAATTCCGGGGCTCAACCCCGGAGCTGTATCCGAAACTGGCAAGCTAGAGTATTACAGGGGTAGGTAGAGTT<br>TCCGGTGTAGGGGTAAAATCCGTAGATATCGGAAGGAATACCAAAAGCGAAGGCAGCCTACTGGAGTAATACTGACGCTGAGAGACGAAAGCGTGGGG |
| P-asv7328 | ACGAGTGGCAATCGCATATTGGGCCTAAAGCGTTCGTAGTTGGTTTGGTAAATTTCTCGTGAAACTGTACGGCTTAACCGTACAACGTGCGGGAAACACTGCCTTACTTGGAACCGGGAAGGGTCAGAAGTATTTT<br>CGAGGGAGCGGTAAAATGTGATAATCTCGAAAGGACTACCTGTGGCGAAGGCGTCTGACTAGAACGGATTCGACAATGAGGAACGAAGGCTAGGGG |
| P-asv7329 | GCGAGCGTTAATCGGAATTACTGGGCGTAAAGCGCACGTAGGCGGCGACACAAGTCAGAGGTGAAATCCCGGGGCTCAACCCCGGAACGGCCCTCGATACTGGCAGCCTAGAGTTCGAGAGAGGTGAGTGGAA<br>TTCCGAGTGTAGAGGTGAAATTCGTAGATATTCGGAGGAACACCAGTGGCGAAGGCGGCTCACTGGCTCGATACTGACGCTGAGGTGCGAAAGTGTGGGG |
| P-asv7330 | ACAAGTGGAAATCATGTTTACTGGGTCTAAAGTGACCGTAGCCGGCTGAGTAAGTCTTTTGTGAAAGTTCCGTAGCTTAACTCGGAATTTGCAAAAGATACTGCTTAGCTTGAGGCCGGGGGGGGTATGAGGTATT<br>GTTTGGGTAGCGGTAAAATGTTATAATCCTTACAAGACCAACTAAGGCGAAGGCATCATACTTGAACGGACCTGACGGTGAGGGTCGAGAGCTGGGGG |
| P-asv7331 | GCAAGCGTTATCCGGATTCATTGGGTTTAAAGGGAGCGTAGGCGGTTTAGTAAGTCAGTGGTGAAAGCCGGTCGCTTAACGATCGAACTGCCATTGATACTGCTGAACTTGAGTACAGATGCTGTGGGCGGAATA<br>TGACATGTAGTGGTGAAATACTTAGAGATGTCATAGAACACCGATTGCGAAGGCAGCTCACAAAACTGTAACTGACGCTGAGGCTCGAAAGTGTGGGG |
| P-asv7332 | GCGAGCGTTGCTCGGAATTACTGGGCGTAAAGGGTCCGCAGGTGGCCTCGTAAGTTAGATGTGAAATCTTAGGGCTTAACCCTGAAACTGCATCTAATACTGCAAGGCTTGAGTCCAGGAGAGGTTGGCGGAATT<br>CCAGGTGTAGCGGTGAAATGCGTAGATATCTGGAGGAACACCAGTGGCGAAGGCGGCCAACTGGCCTGGTACTGACACTCAGGGACGAAAGCGTGGGT |
| P-asv7333 | TCAAGCGTTATCCGGAGTTACTGGGCGTAAAGGGCTCGCAGGCGGTTTGACAAGTTTGACGTGAAATCTCTCGGCTTAACTGGGAGGGGACGTTGGAGACTGTTAGACTTGAGGTAGTGAGAGGGAGGCGGAAT<br>TCCCGGTGTAGTGGTGGAATGCGTAGATATCGGGAAGAACACCAGTGGCGAAAGCGGCCTTCTGGCACTGACCTGACGCTGAGGAGCGAAAGCGTGGGT |
| P-asv7334 | CCAAGCGTTGTTCGGATTCATTGGGCGTAAAGGGTACGTAGGCGGTGCCGTTAGTCGAATGTGAAATCTCGGAGCTCAACTCCGAAACTGCATTCGATACTGCGGTGCTCGAGGATTGTAGAGGAGAGTGGAATT<br>CACGGTGTAGCAGTGAAATGCGTAGATATCGTGAGGAAGACCAGTGGCGAAGGCGACTCTCTGGGCAATTCCTGACGCTGAGGCACGAAGGCTAGGGG |
| P-asv7335 | GCAAGTGTTATCCGGAATTACTGGGCGTAAAGCGCGCGTCGGCGGCTTGCCAAGTCAGGTGTGAAAGCCCGGGGCTTAACCCCGGAATTGCATTTGAAACTGGCAGGCTAGAGTCTGGTAGAGGGTGGTGGAA<br>TTCCACGTGTAGAGGTGAAATTCGTAGATATGTGGAGGAACACCGGTGGCGAAGGCGACCACCTGGGCCAGTACTGACGCTATAGCGCGAAAGCGTGGGG |
| P-asv7336 | GCAAGCGTTAATCGGAATTACTGGGCGTAAAGCGTGCGTAGGCGGATGTCTAAGTCGAATGTGAAAGCCCCGGGCTTAACCTGGGAACTGCATTCGATACTAAACAGTCTTGAGTATAGTAGAGGAAAGTGGAAT<br>TCCCGGTGTAGCAGTGAAATGCGTAGATATACATCGGAACGCCAATGGCGAAGGCAGCTGGCTGGGCCTGTACTGACGCTCATGTACGAAAGCGTGGGG |

| | |
|---|---|
| P-asv7337 | TCAAGCGTTATCCGGAATCATTGGGCGTAAAGGATGTGTAGGTGGTTTTGTTAGTCTCTTGTTAAATTCTTCGGCTCAACCGGGGGCATGCAAGGGAAACGGCAGAACTAGAGGATGCAAGGGGTTTCTGGAACTCTAGGTGTAGCGGTGAAATGCGTTGATATCTAGGGGAACACCAAAAGCGAAGGCAAGAAACTGGGGCATTCCTGACACTGAGACATGAAAGCGTGGGT |
| P-asv7338 | GCGAACGTTGTTCGGAATCACTGGGCGTAAAGGGCGCGCAGGCGGCCTGGTAAGTGGGATGTGAAAGCCCCTGGCTCAATCAGGGAACTGCATCCCAGACTGCCGGGCTTGAGGACGGGAGAGGAAACGGGAATTCCCGGTGTAGCGGTGACATGCACTGATATCGGGAGGAACACCAGTGGCGAAGGCGCGTTTCTGGCCCGTTCCTGACGCTCAGGCGCGAAAGCTGGGGG |
| P-asv7339 | GCAAGCGTTATTCGGAATCATTGGGCGTACAGGGTGTGTAGGCGGTGTGTTAAGTCTGTTGTTAAATACTTCGGCCCAACTGAAGAAAGGCAGCGGAAACTGGCGCACTAGAGGATGGAAGAGAGAAGTGGAATTCTCGGTGTAGCGGTAAAATGCGTAGATATCGAGAGGAACACCGATGGCGAAGGCGACATACTGGTCCGTTACTGACGCTGAGGCTCGAAAGCGTGGGG |
| P-asv7340 | TCAAGCGTTATCCGGATTGATTGGGTGTAAAGCGTGCGCAGGCGGCTTATTAAGCTGGTGGTTAAATTCTTCGGCTCAACCGAAGAACCGCCATCAGAACTGATTAGCTAGAGTGTGGAAGAGGTTAGCGGAATTGCCGGTGTAACGGTAAAATGTGTTAATATCGGCAAGAACACCAGAGGCGAAGGCGGCTAACTGGTACACAACTGACGCTGTTTGCACGAAAGCGTGGGG |
| P-asv7341 | GCAAGCGTTATCCGGAGTTACTGGGCGTAAAGGGCAGGCAGGCGGGTCGATAAGTCGGATGTGAAATCTCCCTGGCTTAACCGGGAGGGTGCATTTGAGACTGTCGACCTGGAGGGCAGCAGAGGAGCGTGGAATTCCGGGTGGAGTGGTGAAATGCGTAGAGATCCGGAGGAACACCAGAGGCGAAAGCGGCGCTCTGGGCTGCACCTGACGCTGAACTGCGAAAGCCAGGGG |
| P-asv7342 | GCAAGCGTTGTTCGGAATTACTGGGCGTAAAGCGCATGCAGGTGGCTGAGTAAGTCGAATGTGAAAGCCCCGGGCTCAACCCGGGAACTGCATCCGAAACTGCTTGGCTAGAGTCCCGGAGAGGAAGGCAGAATTCCCAATGTAGGGGTGAAATCCGTAGATATTGGGAGGAATACCGGTGGCGAAGGCGGCCTTCTGGACGGTGACTGACACTCAGATGCGAAAGCGTGGGG |
| P-asv7343 | GCAAGCGTTGTCCGGATTTATTGGGTTTAAAGGGTGCGTAGGCGGCCCGTTAAGTCAGTGGTGAAAACCAGCAGCTCAACTGTTGATGTGCCAATGATACTGACGGGCTAGAGTACAGACGAGGTAGGCGGAATTGACAGTGTAGCGGTGAAATGCATAGATATTGTCAAGAACACCGATAGCGAAGGCACTTTACTGGGCTATTACTGACACTCAGAGACGAAAGCTAGGGT |
| P-asv7344 | ACTAGCGTTGTTCGGATTTACTGGGCGTAAAGAGCGCGTAGGCGGGTTATTAAGTTAGAAGTGAAAGCCCAGGGCTCAACCCTGGAACTGCTTCTAAAACTGGTAACCTAGAGTATGGAGGGGACAGTGGAATTCCTAGTGTAGAGGTGAAATTCTTAGATATTAGGAGGAACACCGGTGGCGAAAGCGACTGTCTGGAACATTACTGACGCTGAGGTGCGAAAGCGTGGGG |
| P-asv7345 | GCGAGCGTTAATCGGAATTACTGGGCGTAAAGCGGGTGCAGGTGGCTTGGTAAGTCAGATGTGAAATCCCGGGGCTCAACCTCAGAATCGCATTCGAAACGGTCATACTAGAAGAAGTCAGAGGTAAACAGAACTCTCGGTGTAGGGGTGAAATCCGTTGATATCGAGGGGAATACCAAATGCGAAGGCAGTTTACTGGGACTTTCTTGACGCTGAGGCACGAAAGCATGGGT |
| P-asv7346 | GCAAGCGTTGTTCGGAATTATTGGGCGTAAAGCGCGTGTAGGCGGCCATGAAAGTCTGACGTGAAAGTCCGGGGCTTAACCCCGGAAGTGCGTTGGATACTACATGGCTAGAGGATCGGAGGGGAGATCGGAATTCCTGGTGTAGGGGTGAAATCCGTAGATATCAGGAGGAACACCGATGGCGAAGGCAGATCTCTGGACGATTTCTGACGCTGAGACGCGAAAGCGTGGGT |
| P-asv7347 | GCAAGCGTTACCCGGAATCATTGGGCGTGAAGCGTCCGCAGCCGGTTATTTAAGTTGCTTATTAAATCTTGTAGCCCAACTATGAGGCTGTAAGCAAAACTGGATAACTAGAGACTTTTAGAGGCACGTGGAATTCTGTGTGTAGGGGTAAAATCCGTAGATACACAGAAGAACACCAAAAGCGAAGGCAGCGTGCTGGGAAAGTTCTGACGGTCAGGGACGAAAGCGTAGGT |
| P-asv7348 | GCGAGCGTTACCCGGATTTACTGGGCGTAAAGGGCGTGTAGGCGGTCCCTCAAGTCCGATGCTAAAGACCGGGGCTCAACCCCGGGGGTGCGTTGGATACTGTCGGGCTAGACGGTCGGAGGGGGCAGCGGAATTTCCGGAGTAGCGGTGAAATGCGCAGATACCGGAAGGAACGCCGATAGCGAAGGCAGCTGCCTGGACGACTTGTGACGCTGAGGCGCGAAAGCGTGGGG |
| P-asv7349 | GCAAGCGTTATCCGGATTCATTGGGTTTAAAGGGTACGTAGGTGGCCCAGTAAGTCAGTGGTGAAATCCTGCCGCTTAACGGTAGAACTGCCATTGAAACTGCTGGGCTTGAGTACAGTTGAGGTAGGCGGAATGTGTAGTGTAGCGGTGAAATGCTTAGATATTACACAGAACCCCGATTGCGAAGGCAGCTTACTAAACTGTAACTGACACTGAGGTACGAAAGCGTGGGG |
| P-asv7350 | GCAAGCGTTGTTCGGATTTATTGGGCGTAAAGCGTCCGTAGGCCGCTTGGAAAGTTGGATGTGAAATCCTGCAGCTTAACTGCAGAACTGCATTCAAAACTCCCAGGCTAGAGGATGAGAGGGGTAGAGGGAATTCCCGGTGTAAGGGTGAAATCTGCAGAGATCGGGAAGAACACCAGTCGCGAAGGCGCTCTACTGGCTCATATCTGACGCTGAGGGACGAAAGCTAGGGG |
| P-asv7351 | TCGGGCGTTGTTCGGATTTACTGGGCGTAAAGGGAGCGTAGGCGGTTCGGTATGTCGAATGTGAAAGCCCACAACTCAACTGTGGAAATGCATTCGAAACTGCCGGGCTAGAGTACTGGAGAGGAGAGCGGAATTCTTGGTGTAGCGGTGAAATGCGTAGATATCAAGAGGAACACCGGTGGCGAAGGCGGCTCTCTGGACAGATACTGACGCTGAGGCTCTAAAGTTAGGGG |
| P-asv7352 | GCGAGCGTTGTTCGGATTTACTGGGCGTAAGGAGCACGTAGGCGGTTTAGTAAGCCCCTTGTGAAATCTCCAGGCTTAACCTGGAAGGGTCGAGGGGGACTGCTAGGCTAGAGGGCGGGAGAGGAGCGCGGAATTCCCGGTGTAGCGGTGAAATGCGTAGATATCGGGAGGAAGGCCGGTGGCGAAGGCGGCGCTCTGGAACGTCTCTGACGCTGAGGTGCGAAAGCGTGGGG |
| P-asv7353 | GCTAGCGTTGTCCGGATTTACTGGGCGTAAAGGGTACGTAGGCGTTTTAATAAGTTGTATGTTAAATATCTTAGCTTAACTAAGAAAGTGCATCCAAAACTGTTAAAATAGAGTTTGAGAGAGGAACGCAGAATTCATGGTGGAGCGGTGACATGCGTAGATATCATGAGGAAAGTCAAATGCGAAGGCAGCCTTCTGGCTCAAAACTGACGCTGAGGTACGAAAGCGTGGGG |
| P-asv7354 | GCAAGCGTTATCCGGAATCACTGGGCGTAAAAGGCACGTAGGCGGACCTTTAAGTTGGGTTTTAAAGACAACAGCTTAACTGTTGATGTGGATCCAAAACTGGAGGTCTTGGATGCCAAAGAGGAAAGCGGAATTCCTGGTGTAGCGGTGAAATGCGTAGATATCAGGAGGAACACCGGTGGCGAAGGCGGCTTTCTGGTTGGTGATTGACGCTAATGTGCGAAAGCGTGGGT |
| P-asv7355 | GCAAGCGTTGTCCGGATTTACTAGGCGTAAAGGGCAGGTAGGCGGATCTTCAAGTCAGGCGTTAAAGGCATCGGCTCAACCGGTGTACGGCGTTTGAAACTGGAAATCTAGAGTCAGGTAGAGGTAGATGGAATTCCTAGTGTAGCGGTGGAACGCGCAGAGATTAGGAAGAACACCAAAGGCGAAGGCAATCTACTGGGCCATGACTGACGCTAAACTGCGAAGGTGTGGGT |
| P-asv7356 | GCAAGCGTTGTTCGGATTTATTGGGTGTAAAGGGCAGGTAGGTGGCCTTGCAAGTCCGGTGTGAAATCCCACGGCTCAACCGTGGAACTGCATTGGAAACTGCTTGGCTTGAATACCGGAGAGGTTAGGGGAATTCTCGGTGTAAGGGTGAAATCTGTAGATATCGAGAGGAACACCAGTGGCGAAGGCGCCTAACTGGCCGGTTATTGACACTGAGCTGCGAAAGCAAGGGG |
| P-asv7357 | CCTAGCGTTGCTCGGAATCATTGGGCGTAAAGCGGGCGTAGGTGGCTTAATAAGTCAGGAGTGAAAGCCCTGGGCTCAACCCGGGAAGTGCTCTTGAAACTGTTTGGCTTGAGTGTAGGAGAGGGCAGTAGAATTCCAGGTGTAATGGTGAAATACGTAGATATCTGGAGGAATACCGGAGGGGAAGCCGGCTGCCTGGCCTAACACTGACACTCAGGCCCGAAAGCGTGGGG |
| P-asv7358 | GCGAGCGTTGTTCGGATTTATTGGGTGTAAAGGGAGCGTAGGCGGCGCAGTCAGTGAGATGTGAAATCTTCGAGCTTAACTCGGAAACTGCATTTCATACTGCTGTGCTAGAGTACTAAAGAGGCGCATGGAATTCTTGGTGTAACGGTGAAATGTGCAGATATCAAGAAGAACACCTGTGGCGAAGGCGATGCGCTGGGTGGATACTGACGCTGAGGCTCGAAAGCAAGGGT |
| P-asv7359 | CCGAACGTTATTCGGTATCACTGGGCTTAAAGGGTGCGCAGGCGGTCCGGTAAGTGAGGTGTGAAATCCCACGGCTCAACCGTGGAACTGCGCTTCATACTGTCGGACTTGAGGGAAGCAGAGGTGAGCGGAACTAATGGTGGAGCGGTGAAATGCGTTGATATCATTAGGAACACCGGTGGCGAAGGCGGCTCACTGGGCTTCTTCTGACGCTCAGGCACGAAAGCCAGGGG |
| P-asv7360 | GCTAGTGTTACTCAACTTGACTGGGCGTAAAGGGCGTGAAGGTGGTATGGTACGTTTTATTGTAAATACTCAAACATAATTTGAGGAGCTTTAAAATACGGCTATGCTTGAGTTTATTTGAAGAAAGTTGTACTTCTAGTGTAGAGGTGAAATTTGCAAAAATTAGAAAGACAGTCCACGGGCGAAAGCGACTTTCTACTATAAACTGACACTATAGCGCGAAAGCATAGGT |
| P-asv7361 | GCTAGCGTTATTCAGAATTACTGGGCGTAAAGCGCACGTAGGCGGCTTTGTAAGTTAGAGGTGAAAGCCTGGAGCTCAACTCCGGAACTGCCTTTGATACAGCATATCTCGAGTCCGAGAGAGGTGAGTGGAATTCCTAGTGTAGAGGTGAAATTCGTAGATATTAGGAAGAACACCGGTGGCGAAGGCGGCTCACTGGCTCGGTACTGACGCTGAGGTGCGAAAGCGTGGGG |
| P-asv7362 | GCTAGCGTTGTCCGGAATCACTGGGCGTAAAGGGTTCGCAGGCGGAAATGCAAGTCAGATGTAAAAGGCAGTAGCTTAACTACTGTAAGCATTTGAAACTGCATATCTTGAGAAGAGTAGAGGTAAGTGGAATTTGTAGTGTAGCGGTGAAATGCGTAGATATTAAAAAGAATACCGGTGGCGAAGGCGACTTACTGGGCTCATTCTGACGCTGAGGAACGAAAGCGAGGGT |
| P-asv7363 | GCTAGCGTTGTTCGGAATTACTGGGCGTAAAGCGCACGTAGGCGGATCAGAAAGTCAGAGGTGAAATCCCAGGGCTCAACCTTGGAACTGCCTTTGATACTGGAAGTCTTGAGTCCGGTAGAGGTGAGTGGAACTGCTAGTGTAGAGGTGAAATTCGTAGATATTAGCAAGAACACCAGTGGCGAAGGCGGCTCACTGGACCGGTACTGACGCTGAGGTACGAAAGCATGGGG |
| P-asv7364 | GCAAGCGTTATCCGGAATTATTGGGCGTAAAGCGTCAGTAAACTGTTTGTTAAGTCGAATATTAAATATATGGGCTCAACCCATATGCTGTATTCGATACTGGCAGACTTGAGGTCGGTAGAGGTAACAGGAATTCCTGGTGTAGTAGTGAAATGCGTTAATATCAGGAAGAACGCCAATTGTGTAAACTTGTTACTGGCCGCACCTGACGTCGAATGACGAAAGCGTGGGG |
| P-asv7365 | GCAAGCGTTATCCGGATTTATTAGGTTTAAAGGGTTCGCAGGCGGAATTTTAAGTCAGTGGTGAAAGCCTATAGCTTAACTATAGAAATGCCACTGATACTGATATTCTTGAGTGTAGATGAAGTGGGCGGAATATGTCATGTAGCGGTGAAATGCATAGATATGACATGGAACACCAATTGCGAAGGCAGCTCACTAAACTATTACTGACGCTCATGAACGAAAGCATGGGG |
| P-asv7366 | GCAAGCGTTGTTCGGAATTACTGGGCGTAAAGCGCACGTAGGCGGGCTCGCAAGTCAATTGTGAAATCCCCGGGCTTAACCTGGGAATGGCAGTTGAAACTGTGGGTCTTGAATGCCTGAGAGGATGGGGGAATTCCTGGTGGAGAAGTGAAATTCGTAGATATCAGGAGGAACACCTGAGGCGAAGGCGCCTGTCTGGCAGTGCATTGACGCTGAGGTGCGAAAGCGTGGGG |
| P-asv7367 | TCGAGCGTTATCCGGAATTACTGGGCGTAAAGGGCACGCAGGCGGCTATTCAAGTCTGGCGTGAAAGCTCCCGGCTTAACTGGGAGAGGCCGTTGGAAACTGAGTAGCTTGAGGTGATGAGAGGGATGCGGAATTCCCGGTGTAGTGGTGTAATGCGTAGATATCGGGAAGAACACCAGTGGCGAAAGCGGCATCCTGGCATTGACCTGACGCTGAGGTACGAAAGCGTGGGG |
| P-asv7368 | GCAAACGTTATCCGGAATTACTGGGCGTAAAGGGCGCGCAGGCGGATCGATAAGTGAGGCATGAAAGCTTCGGGCTAAACCTGGAGAGGCTGTCTGAGACTGTCGATCTGGAGTACGATAGAGGGGTGTAGAATTCCGCGTGTAGTGGTGAAATGCGTAGAGATGCGGAAGAATACCAGAGGCGAAGGCGGCACCCTGGATCGTAACTGACGCTGAGGCGCGAAAGCATGGGG |
| P-asv7369 | GCGAACGTTATTCGGAATCACTGGGCTTAAAGCGCGTGTAGGCGGCCCGACACGTCGGTTGCTGAAATCCCCCGGCTCAACCGGGGAACGGGCATCGATACGATCGGGCTGGAGGGGGGTAGGGGGATCGGGAACTTCCGGTGGAGCGGTGAAATGCGTTGAGATCGGAAGGAACGCCCGCGGCGAAAGCGCGGTCCTGGACCCCACCTGACGCTGAGACGCGAAAGCTAGGGG |
| P-asv7370 | GCAAGCATTGATCGGAATTACTGGGCGTAAAGGGCGCGTAGGCGGCTAGGAAAGTCGGATGTGAAATTCCGGGGCTCAACCCCGGAGCTGCATTTGAAACTCTCTAGCTAGAGTTCAGTTAGGGTAAACGGAATTCCACGTGTAGCGGTGAAATGCGTAGATATGTGGAAGAACACCGGTGGTGAAGACGGTTTACTGGGCTGGTACTGACGCTGAGGCGCGAAAGCATGGGG |
| P-asv7371 | CCGAGCGTTGTCCGGAATCATTGGGCGTAAAGGGTACGTAGGCGGCCTAGTAAGTTAGAAGTGAAAGAATATAGCTCAACTATATAAAGCTTTTAAAACTGCTAGGCTTGAGAGATGAAAGGGAAAGTGGAATTCCTAGTGTAGCGGTGAAATGCGCAGATATTAGGAAGAATACCGGTGGCGAAGGCGACTTTCTGGTCATCATCTGACGCTGAGGTACGAAAGCGTGGGT |
| P-asv7372 | GCAGGCGTTGTCCGGAATTATTGGGCGTAAAGCGCGCGCAGGCGGATCAGTCAGTCTGTCTTAAAAGTTCGGGGCTTAACCCCGTGATGGGATGGAAACTGCTGATCTAGAGTATCGGAGAGGAAAGTGGAATTCCTAGTGTAGCGGTGAAATGCGTAGATATTAGGAAGAACACCAGTGGCGAAGGCGACTTTCTGGACGAAAACTGACGCTGAGGCGCGAAAGCCAGGGG |
| P-asv7373 | GCGAGCGTTAATCGGAATTACTGGGCGTAAAGCGCACGCAGGCGGTTTGTTAAGCTAGATGTGAAAGCCCCGGGCTCAACCTGGGATGGTCACTTAGAACTGGCAGACTAGAGTCTTGGAGAGGGGAGTGGAATTCCAGGTGTAGCGGTGAAATGCGTAGATATCTGGAGGAACATCAGTGGCGAAGGCGACTCCCTGGCCAAAGACTGACGCTCATGTGCGAAAGTGTGGGT |
| P-asv7374 | CCAAGCGTTATCCGGAATCATTGGGTTTAAAGGGTCCGTAGGCGGGATGATAAGTCAGTGGTGAAAGTTTTCGGCTCAACCGAGAAATTGCCATTGAAACTGTTGTTCTTGAATTATTGTGAAGTGGTTAGAACGTGTAGTGTAGCGGTGAAATGCATAGATATTACACAGAATACCGATTGCGAAGGCAGATCACTAACAATATATTGACGCTGATGGACGAAAGCGTGGGT |
| P-asv7375 | TCTCTCTTTTCTCTTCCTCCCTTTTCTTAAAGGGTGCGTAGGCGGGTTTTTAAGTCAGGGGTGAAATCCTGGAGCTCAACTCCAGAACTGCCTTTGATACTGAAGATCTTGAGTCCGGGAGAGGTGAGTGGAACTGCGAGTGTAGAGGTGAAATTCGTAGATATTCGCAAGAACACCAGTGGCGAAGGCGGCTCACTGGCCCGGTACTGACGCTGAGGCACGAAAGCGTGGGG |

| | |
|---|---|
| P-asv7376 | GCAAGCGTTGTCCGGATTTATTGGGCGTAAAGAGCGTGTAGGCGGCCGCACAGGTCCGTTGTGAAAACTGGAGGCTTAGCCTTCAGACGTCGATGGAAACCGTGCGGCTAGAGTCCGGAAGAGGAGAATGGAATTCCTGGTGTAGCGGTGAAATGCGCAGATATCAGGAAGAACACCCGTGGCTAAGGCGGTTCTCTAGTACGGTACTGACGCTGAGACGCGAAAGCGTGGGG |
| P-asv7377 | CCAAGCGTTATCCGGATTCATTGGGTTTAAAGGGTGCGTAGGCGGCCTGTTAAGTCAGTGGTGAAATTTTGTGGCTCAACCATAAAACTGCCATTGAAACTGACAGGCTTGAATACAGTTGAAGTAGGCGGAATAGGTAGTGTAGCGGTGAAATGCTTAGATATTACCTAGAACACCGATTGCGAAGGCAGCCTACTAAAGTGTCATTGACGCTGATGCACGAAAGCGTGGGT |
| P-asv7378 | GCGAGCGTTAATCGGATTTACTGGGCGTAAAGCGTGCGTAGGCGGCTTTTTAAGTCGGATGTGAAATCCCTGAGCTTAACTTAGGAATTGCATTCGATACTGGGAAGCTAGAGTATGGGAGAGGATGGTAGAATTCCAGGTGTAGCGGTGAAATGCGTAGTAAAAAAAAAAAAAAAAAAAAAAAAAAAAAAAAAAAAAAAAAAAAAAAAAAAAAGCTGAGGTACGAAAGCATGGGG |
| P-asv7379 | GCATGCGTTAATCGGATTTACTGGGCGTAAAGCGTGCGTAGGCGGTTCGTTAAGTCGATTGTGAAAGCCCTGGGCTTAACCTGGGAATTGCAGTCGATACTGGCGGACTAGAGAACGGTAGAGGGAGGCGGAACTCCAGGTGTAGCGGTGAGATGCGTAGATATCTGGAAGAACACCGATGGCGAAGGCAACCTCCTGGGCCTGTTCTGACGCTGAGGCACGAAAGCGTGGGT |
| P-asv7380 | GCAAGCGTTGTCCGGAATTATTGGGCGTAAAGAGCTCGTAGGCGGTCTGTTACGTCCGCTGTGAAAACCCGGGGCTTAACCCTTGGCTTGCAGTGGATACGGACAGACTAGAGGTAGGTAGGGGAGAATGGAATTCCCGGTGTAGCGGTGAGATGCTTAGATATGACATAGAACACCAATTGCGAAGGCAGCTCACTGCGCAGTGATTGACGCTGAGGCACGAAAGCGTGGGG |
| P-asv7381 | GCTAGCGTTGCTCGGAATCACTGGGCGTAAAGCGCACGTAGGCGGATTCTTAAGTCGGTGGTGAAATCCTGGAGCTCAACTCCAGAACTGCCTTTGATACTGGAGAGCTAGAGTCCGGAAGAGGTGAGTGGAACTGCGAGTGTAGAGGTGAAATTCGTAGATATTCGCAAGAACACCAGTGGCGAAGGCGGCTCACTGGCCCGGTACTGACGCTGAGATACGAAAGCGTGGGG |
| P-asv7382 | GCGAGCGTTGTCCGGATTTACTGGGCGTAGAGAGCGCGCAGGCGGTCGATCAAGTCGGATGTGAAAGCCCCCGGCTCAACTGGGGAGGGTCATTCGATACTGATCGACTCGAAGGCAGGAGAGGGAAGCGGAATTCCCGGTGTAGTGGTGAAATGCGTAGATATCGGGAGGAACACCAGTGGCGAAGGCGGCTTCCTGGCCTGTTCCTGACGCTGAGGCGCGAAAGCGTGGGG |
| P-asv7383 | GCAAACGTTGTTCGGAATTACTGGGCATAAAGCGCACGTAGGTGGTCCGCTAAGTCGCGTGTGAAATCCCCCGGCTCAACCGGGGAAATGCGCGTGATACTGGCTGTCTTGAGTACGGGAGAGGTGAGTGGAACTCCGAGTGTAGAGGTGAAATTCGTAGATATTCGGAAGAACACCAGTGGCGAAGGCGACTCACTGGCCCGTTACTGACGCTGAGGCGCGAAAGCGTGGGG |
| P-asv7384 | GCAAGCGTTGTTCGGAATTATTGGGCGTAAAGGGTGCGTAGGCGGTGCGGTAAGTCTTGTGTGAAATCTACAGACTCAACTTGTAGTCTGCACAAGAAACTGCCGGGCTTGAGTGTGGGAGAGGTGAGTGGAATTTCCGGTGTAGCGGTGAAATGCGTAGATATCGGAAGGAACACCTGTGGCGAAAGCGGCTCACTGGACCACAACTGACGCTGATGCACGAAAGCTAGGGG |
| P-asv7385 | GCGAGCGTTGTTCGGAATTATTGGGCGTAAAGGGCATGCAGGCGGGTCGCCAAGCTTGGTAAGAAATACCGGGGCTCAACTCCGGAGCTATATTGAGAACTGGCGAGCTAGAGTTGCCGAAGGGTATCCGGAATTCCGCGTGAAGGGGTGAAATCTGTAGATATGCGGAAGAACACCGATGGCGAAGGCAGGATACCGGCGGACGACTGACGTTGAGGTGCGAAGGTGCGGGG |
| P-asv7386 | GCTAGCGTTGTTCGGAATTACTGGGCGTAAAGCGCACGTAGGCGGCTTTGTAAGTCAGAGGTGAAAGCCTGGAGCTCAACTCCAGAACTGCCTTTGAGACTGCATCGCTTGAATCCAGGAGAGGTGAGTGGAATTCCGAGTGTAGAGTTAAAATTCGTCGAAACTGGAAGGAACACCAAGGGCGAAAGCAATTCTCGACAATATTCTGACGTTGAGGCACGAAGGGATGGGT |
| P-asv7387 | GCAAGCGTTAATCGGAATTACTGGGCGTAAAGCGTGCGCAGGCGGCTGATTAAGTCAAATGTGAAAGCCCCGGGCTTAACCTGGGGATGGCATTCGATACTGGTCAGCTAGAGTTTGGTAGAGGTAAGCGGAATTCCGGGTGTAGCGGTGAAATGCGTAGATATCCGGAGGAACATCAGTGGCGAAGGCGGCTTACTGGACCAAAACTGACGCTCAGGCACGAAAGCGTGGGT |
| P-asv7388 | GCGAGCGTTGTCCGGATTTATTGGGTTTAAAGGGTGCGTAGGCTGGGAGGTAAGTCAGCGGTGAAAAGCACACGCTTAACGGGTGTCTTGCCGCAGATACTACTTTTCTTGGATGAGGAGGAGGTCACTGGAACGAATGGTGTAGCGGTGAAATGCATAGATATCATCCAGAACACCGATTGCGAAGGCAGGTGACTGTACCTTAATCGACGCTGAGGCACGAAAGCGTGGGG |
| P-asv7389 | GCAAGCGTTAATCGGAATTATTGGGCGTAAAGCGGGTGCAGGTGGCTTGGTAAGTCAGATGTGAAAGCCTGGGGCTTAACCCCAGAAGTGCATTTGATACTGCTGAGCTTGAGTGCTGGAGAGGCTATTAGAATTCCTGGTGTAGTGGTGAAGTACGTAGATATCAGGAGGAATACCGGAGGCGAAGGCGGATAGCTGGCCAGACACTAACACTCAGACCCGAAAGCGTGGGG |
| P-asv7390 | ACGAACGTTATTCGGAATTACTGGGCTTAAAGGGTGCGTAGGCTGCACGGAAAGTTGGGTGTGAAAGCCCTCAGCTCAACTGAGGAATTGCATCCAAAACTGCCGTGCTTGAGGGAGACAGAGGTGAGCGGAACTCAAGGTGGAGCGGTGAAATGCGTTGATATCTTGAGGAACACCGGTGGCGAAGGCGGCTCACTGGGTCTCTTCTGACGCTGAGGCACGAAAGCTAAGGT |
| P-asv7391 | GCGAGCGTTGTCCGGATTTACTGGGCGTAAAGGGCTCGTAGGTGGTTTGTCGCGTCGTCTGTGAAATCCCCGGGCTCAACCTGGGAATTGCATTGGGGACTGCAAGGCTAGAATCTGGCAGAGGGGGGTAGAATTCCACGTGTAGCAGTGAAATGCGTAGATATGTGGAGGAACACCGATGGCGAAGGCAGCCCCCTGGGTCAAGATTGACGCTCATGCACGAAAGCGTGGGG |
| P-asv7392 | GCGAGCGTTATCCGGATTCACTGGGCGTAAAGAGCGCGTAGGCGGTTCCGCAAGTCACCCGTGAAAGCCCCCGGCTCAACCGGGGAGGGCCGGGTGATACTGCGGGGCTTGAGGGCGGAAGAGGGAAGCGGAATTCCCGGTGTAGCGGTGAAATGCGTAGATACCGGGAAGAACACCGGTGGCGAAGGCGGCTTCCTGGTCCGTTCCTGACGCTGAGGCGCGAAGGCTAGGGG |
| P-asv7393 | GCAAGCGTTGTTCGGAATTACTGGGCGTAAAGGGCTCGTAGGCGGCCAACTAAGTCAGACGTGAAATCCCCAGGCTTAACCTGGGAACTGCGTCTGATACTGGATGGCTTGAATTCGGGAGAGGGATGCAGAATTCCAGGTGTAGCGGTGAAATGCGTAGATATCTGGAGGAATACCGGTGGCGAAGGCGGCATCCTGGACCGACATTGATGCTGAGGAGCGAAAGCGTGGGG |
| P-asv7394 | GCTAGCGTTGTCCGGAATTACTGGGCGTAAAACGAGCGTAGGTTGCGCCGCAAGTCAGATGTGAAATCCCAGGGCTCAACCGTGGAACTGCGTTTGAAACTGCATGGCTTGAGGGAGATAGGGGTGAGCGGAACTGATGGTGGAGCGGTGAAATGCGTTGATATCATCAGGAACACCGGTGGCGAAGGCGGCTCACTGGGTCTCTTCTGACGCTGAGGCACGAAAGCTAGGGG |
| P-asv7395 | GCGAGCGTTAATCGGAATTACTGGGCGTAAAGCGCGCGTAGGTGGTTTGCTAAGCGATATGTGAAATCCCCGGGCTTAACCTGGGAACTGCATGTCGAACTGGCAGGCTAGAGTATGGTAGAGGGGTGTGGAATTTCCTGTGTAGCGGTGAAATGCGTAGATATAGGAAGGAACATCAGTGGCGAAGGCGACACCCTGGACTAATACTGACACTGAGGTGCGAAAGCGTGGGG |
| P-asv7396 | GCGAGCGTTAATCGGAATTACTGGGCGTAAAGCGAGCGCAGACGGTTACTTAAGCAGGATGTGAAATCCCCGGGCTCAACCTGGGAACTGCGTTCTGAACTGGGTGACTAGAGTGTGTCAGAGGGAGGTAGAATTCCACATGTAGCGGTGGAATGCGTAGAGATGTGGAGGAATACCGATGGCGAAGGCAGCCTCCTGGGATAACACTGACGCTCATGCACGAAAGCGTGGGG |
| P-asv7397 | GCAAGCGTTAATCGGAATTACTGGGCGTAAAGAGCTCGTAGGTGGTTTGTCGCGTTGTTCGTGAAAACCGAGGGCTTAACCCTGGAACTGCCTGCGAGACCTCAGTACTAGAGTGTGAGAGGGGATAGTGGAATACCCAGTGTAGCGGTGAAATGCGTAGAGATTGGGTGGAACACCGGTGGCGAAGGCGGCTATCTGGCTCACAACTGACGATCAGGCACGAAAGCGTGGGG |
| P-asv7398 | GCAAGCGTTAATCGGAATTACTGGGCGTAAAGAGCTCGTAGGTGGTTTGTCGCGTTGTTCGTGAAAACCGAGGGCTTAACCCTGGAACTGCCTGCGAGACCTCAGTACTAGAGTGTGAGAGGGGATAGTGGAATACCCAGTGTAGCGGTGAAATGCGTAGAGATTGGGTAGAACACCGGTGGCGAAGGCGGCTATCTGGCTCACAACTGACGATCAGGCACGAAAGCGTGGGG |
| P-asv7399 | GCAAGCGTTAATCGGAATTACTGGGCGTAAAGAGCTCGTAGGTGGTTTGTCGCGTTGTTCGTGAAAACCGAGGGCTTAACCCTGGAACTGCCTGCGAGACCTCAGTACTAGAGTGTGAGAGGGGATAGTGGAATACCCAGTGTAGCGGTGAAATGCGTAGAGATGTGGAGGAATACCGATGGCGAAGGCAGCCCCCTGGGATAACACTGACGCTCATGCACGAAAGCGTGGGG |
| P-asv7400 | GCGAGCGTTAATCGGATTTACTGGGCGTAAAGCGTGCGTAGGCGGCTTTTTAAGTCGGATGTGAAATCCCTGAGCTTAACTTAGGAATTGCATTCGATACTGGGAAGCTAGAGTATGGGAGAGGATGGTAGAATTCCAGGTGTAGCGGTGAAATGCGTAGATATTGGGAGGAACACCAGTGGCGAAGGCGACTTTCTGGACTGTAACTGACACTGAGGCACGAAAGCGTGGGG |
| P-asv7401 | GCGAGCGTTAATCGGATTTACTGGGCGTAAAGCGTGCGTAGGCGGCTTTTTAAGTCGGATGTGAAATCCCTGAGCTTAACTTAGGAGTTGCATTCGATACTGGGAAGCTAGAGTATGGGAGAGGATGGTAGAATTCCAGGTGTAGCGGTGAAATGCGTAGATATTGGGAGGAACACCAGTGGCGAAGGCGACTTTCTGGACTGTAACTGACACTGAGGCACGAAAGCGTGGGG |
| P-asv7402 | GCAAGCGTTGTTCGGAATTACTGGGCGTAAAGGGCTCGTAGGTGGCCAACTCAGTCAGACGTGAAATCCCTCGGCTTAACCGGGGAACTGCGTCTGATACTGGATGGCTTGAGGTTGGGAGAGGGATGCGGAATTCCAGGTGTAGCGGTGAAATGCGTAGATATCTGGAGGAACACCGGTGGCGAAGGCGGCATCCTGGACCAATTCTGACGCTGAGGAGCGAAAGCCAGGGG |
| P-asv7403 | CAAGCGTTGTTCGGAATTACTGGGCGTAAAGGGCTCGTAGGCGGCCAACTAAGTCGGATGTGAAATCCCCAGGCTTAACCTGGGAACTGCATCCGATACTGGATGGCTTGAATCCGGGAGAGGGATGCAGAATTCCAGGTGTAGCGGTGAAATGCGTAGATATCTGGAGGAATACCGGTGGCGAAGGCGGCATCCTGGACCGGCATTGACGCTGAACGGCGAAAGCTAGGGG |
| P-asv7404 | GCAAGCGTTGTCCGGAATCATTGGGCGTAAAGCGCGTGTAGGCGGTTCGGTAAGTCGGCTGTGAAAGTCCAGGGCTCAACCCTGGGATGCCGGTCGATACTGCCGGGCTAGAGGGTGGTAGAGGCGAGTGGAATTCCCGGTGTAGCGGTGAAATGCGCAGATATCGGGAGGAACACCAGTAGCGAAGGCGGCTCGCTGGGCCATACCTGACGCTGAGACGCGAAAGCGTGGGG |
| P-asv7405 | GCAAGCGTTGTCCGGAATTATTGGGCGTAAAGGGCTCGCAGGCGGTTTCTTAAGTCTGATGTGAAAGCCCCCGGCTCAACCGGGGAGGGTCATTGGAAACTGGGAAACTTGAGTGCAGAAGAGGAGAGTGGAATTCCATGTGTAGCGGTGAAATGCGCAGAGATATGGAGGAACACCAGTGGCGAAGGCGGCTCTCTGGTCTGTAACTGACGCTGATGTGCGAAAGCGTGGGG |
| P-asv7406 | GCAAGCGTTGTCCGGAATCATTGGGCGTAAAGAGCGCGTAGGCGGCCCTGTAAGTCCGCTGTGAAAGTCAAGGGCTCAACCCTCGGATGCCGGTGGATACTGCAGGGCTAGAGTCCGGAAGAGGCGAGTGGAATTCCTGGTGTAGCGGTGAAATGCGCAGATATCAGGAGGAACACCGATGGCGAAGGCAGCTCGCTGGGACGGTACTGACGCTGAGGCGCGAAAGCGTGGGG |
| BH-asv1 | GCTAGCGTTGTTCGGAATTACTGGGCGTAAAGGGCGCGTAGGCGGTTTATCAAGTCAGGCGTGAAAGCCCCGGGCTCAACCTGGGAATTGCGTTTGATACTGGTAGACTTGAGTTCGGGAGAGGTGAGTGGAATTCCCAGTGTAGAGGTGAAATTCGTAGATATTGGGAAGAACACCGGTGGCGAAGGCGGCTCACTGGACCGATACTGACGCTGAGGCGCGAAAGCGTGGGG |
| BH-asv2 | GCTAGCGTTGCTCGGAATCACTGGGCGTAAAGGGTGCGTAGGCGGGTCTTTAAGTCAGGGGTGAAATCCTGGAGCTCAACTCCAGAACTGCCTTTGATACTGAAGATCTTGAGTTCGGGAGAGGTGAGTGGAACTGCGAGTGTAGAGGTGAAATTCGTAGATATTCGCAAGAACACCAGTGGCGAAGGCGGCTCACTGGCCCGATACTGACGCTGAGGCACGAAAGCGTGGGG |
| BH-asv3 | GCTAGCGTTGCTCGGAATCACTGGGCGTAAAGCGCACGTAGGCGGATTCTTAAGTCGGTGGTGAAAGCCTGGAGCTCAACTCCAGAACTGCCTTCGATACTGAGAGTCTTGAGTCCGGGAGAGGTGAGTGGAACTGCGAGTGTAGAGGTGAAATTCGTAGATATTCGCAAGAACACCAGTGGCGAAGGCGGCTCACTGGCCCGGTACTGACGCTGAGGTGCGAAAGCGTGGGG |
| BH-asv4 | GCTAGCGTTGTTCGGAATTACTGGGCGTAAAGCGAGTGTAGGTGGTTGTCATAGTCAGGTGTGAAAGCCTTGAGCTCAACTCAAGAAATGCACTTGATACTGGGCGACTAGAGGACCGGAGAGGATAGTGGAATTCCCAGTGTAGTGGTGAAATACGTAGAGATTGGGAAGAACACCAGTGGCGAAGGCGGCTATCTGGACGGTTTCTGACACTAAGACTCGAAAGCGTGGGG |
| BH-asv5 | GCAAGCGTTATCCGGAATTATTGGGCGTAAAGCGCGCGTAGGCGGTTTTTTAAGTCTGATGTGAAAGCCCACGGCTCAACCGTGGAGGGTCATTGGAAACTGGAAAACTTGAGTGCAGAAGAGGAAAGTGGAATTCCATGTGTAGCGGTGAAATGCGCAGAGATATGGAGGAACACCAGTGGCGAAGGCGACTTTCTGGTCTGTAACTGACGCTGATGTGCGAAAGCGTGGGG |
| BH-asv6 | GCAAGCGTTGTCCGGAATTATTGGGCGTAAAGGGCTCGCAGGCGGTTCCTTAAGTCTGATGTGAAAGCCCCCGGCTCAACCGGGGAGGGTCATTGGAAACTGGGGAACTTGAGTGCAGAAGAGGAGAGTGGAATTCCACGTGTAGCGGTGAAATGCGTAGAGATGTGGAGGAACACCAGTGGCGAAGGCGACTCTCTGGTCTGTAACTGACGCTGAGGAGCGAAAGCGTGGGG |
| BH-asv7 | GCAAGCGTTGTCCGGATTTATTGGGCGTAAAGCGCGCGCAGGCGGTCTTTTAAGTCTGATGTGAAAGCCCCCGGCTTAACCGGGGAGGGTCATTGGAAACTGGAAGACTGGAGTGCAGAAGAGGAGAGTGGAATTCCACGTGTAGCGGTGAAATGCGTAGATATGTGGAGGAACACCAGTGGCGAAGGCGACTCTCTGGTCTGTAACTGACGCTGAGGCGCGAAAGCGTGGGG |
| BH-asv8 | GCAAGCGTTAATCGGAATTACTGGGCGTAAAGCGCACGCAGGCGGTCTGTCAAGTCGGATGTGAAATCCCCGGGCTCAACCTGGGAACTGCATTCGAAACTGGCAGGCTTGAGTCTTGTAGAGGGGGGTAGAATTCCAGGTGTAGCGGTGAAATGCGTAGAGATCTGGAGGAATACCGGTGGCGAAGGCGGCCCCCTGGACAAAGACTGACGCTCAGGTGCGAAAGCGTGGGG |
| BH-asv9 | GCAAGCGTTAATCGGAATTACTGGGCGTAAAGCGTGCGCAGGCGGTCCGCTAAGACAGATGTGAAATCCCCGGGCTTAACCTGGGAACTGCATTTGTGACTGGCGGGCTAGAGTATGGCAGAGGGGGGTAGAATTCCACGTGTAGCAGTGAAATGCGTAGAGATGTGGAGGAATACCGATGGCGAAGGCAGCCCCCTGGGCCAATACTGACGCTCATGCACGAAAGCGTGGGG |

| | |
|---|---|
| BH-asv10 | ACGAGCATTATCCGGATTTATTGGGCGTAAAGGAAGCGTAGGCGGGGAGGTTGATCCATTGTTAAAAGCATTTGCTTAACAAATGTGTGCATTGGAGATCGCCTCCCTAGAGTTAATCAGGGGGTACTGGAATTCAATGTGTAGCGGTGGAATGCGTAGATATATTGAGGAACACCAGAGGCTAAGGCGAGTGCCTAGGATATAACTGACGCTGAGGCTTGAAAGCGTGGGG |
| BH-asv11 | GCTAGCGTTGCTCGGAATTACTGGGCGTAAAGGGCGCGTAGGCGGACAGTTAAGTTGGGGGTGAAAGCCCGGGGCTCAACCTCGGAAATGCCTTCAATACTGGCTGTCTTGAGTACGGGAGAGGTGAGTGGAACTCCGAGTGTAGAGGTGAAATTCGTAGATATTCGGAAGAACACCAGTGGCGAAGGCGACTCACTGGCCCGTTACTGACGCTGAGGCGCGAAAGCGTGGGG |
| BH-asv14 | GCAAGCGTTGTCCGGATTTATTGGGCGTAAAGCGAGCGCAGGCGGTTTCTTAAGTCTGATGTGAAAGCCCCCGGCTCAACCGGGGAGGGTCATTGGAAACTGGAGACTTGAGTGCAGAAGAGGAGAGTGGAATTCCATGTGTAGCGGTGAAATGCGTAGATATATGGAGGAACACCAGTGGCGAAGGCGGCTCTCTGGTCTGTAACTGACGCTGAGGCTCGAAAGCGTGGGG |
| BH-asv15 | GCTAGCGTTGCTCGGAATTACTGGGCGTAAAGCGCACGTAGGCGGGTTCTTAAGTCGGTGGTGAAATCCTGGAGCTCAACTCCAGAACTGCCTTCGATACTGGGAATCTTGAGTCCGGGAGAGGTGAGTGGAACTGCGAGTGTAGAGGTGAAATTCGTAGATATTCGCAAGAACACCAGTGGCGAAGGCGGCTCACTGGCCCGGTACTGACGCTGAGGTGCGAAAGCGTGGGG |
| BH-asv16 | GCAAGCGTTAATCGGAATTACTGGGCGTAAAGCGTGCGTAGACGGTTACATAAGTCGGGTGTGAAAGCCCCGGGCTCAACCTGGGAATTGCATTCGAGACTGCGTAGCTAGGGTGCGGAAGAGGGAAGCGGAATTTCCGGTGTAGCGGTGAAATGCGTAGATATCGGAAGGAACACCAGTGGCGAAAGCGGCTTCCTGGTCCAGCACCGACGTTCAGGCACGAAAGCGTGGGG |
| BH-asv17 | GCAAGCGTTATCCGGATTTATTGGGCGTAAAGAGAGTGCAGGCGGTTTTCTAAGTCTGATGTGAAAGCCTTCGGCTTAACCGGAGAAGTGCATCGGAAACTGGATAACTTGAGTGCAGAAGAGGGTAGTGGAACTCCATGTGTAGCGGTGGAATGCGTAGATATATGGAAGAACACCAGTGGCGAAGGCGGCTACCTGGTCTGCAACTGACGCTGAGACTCGAAAGCATGGGT |
| BH-asv18 | GCAAGCGTTAATCGGAATTACTGGGCGTAAAGCGTGCGCAGGCGGTTATGTAAGACAGTTGTGAAATCCCCGGGCTCAACCTGGGAACTGCATCTGTGACTGCATAGCTAGAGTACGGTAGAGGGGGATGGAATTCCGCGTGTAGCAGTGAAATGCGTAGATATGCGGAGGAACACCGATGGCGAAGGCAATCCCCTGGACCTGTACTGACGCTCATGCACGAAAGCGTGGGG |
| BH-asv19 | GCAAGCGTTATCCGGATTCACTGGGTTTAAAGGGTGCGTAGGCGGTATGTAAGTCAGTGGTGAAATACCGGAGCTTAACTTCGGAACTGCCATTGATACTATATATCTTGAATATTGTGGAGGTAAGCGGAATATGTCATGTAGCGGTGAAATGCTTAGAGATGACATAGAACACCGATTGCGAAGGCAGCTTGCTACGCAAATATTGACGCTGAGGCACGAAAGCGTGGGG |
| BH-asv21 | GCAAGCGTTAATCGGAATTACTGGGCGTAAAGCGTGCGTAGACGGTTACATAAGTCGGGTGTGAAAGCCCCGGGCTCAACCTGGGAATTGCATTCGAGACTGCGTAGCTAGGGTACGGAAGAGGGAGGCGGAATTTCAGGTGTAGCGGTGAAATGCGTAGATATCTGAAGGAACACCAGTGGCGAAAGCGGCCTCCTGGTCCAGTACCGACGTTCAGGCACGAAAGCGTGGGG |
| BH-asv23 | GCTAGCGTTGCTCGGAATTACTGGGCGTAAAGGGCGCGTAGGCGGACAGTTAAGTTGGGGGTGAAAGCCCGGGGCTCAACCTCGGAATTGCCTTCAATACTGGCTGTCTTGAGTACGGGAGAGGTGAGTGGAACTCCGAGTGTAGAGGTGAAATTCGTAGATATTCGGAAGAACACCAGTGGCGAAGGCGACTCACTGGCCCGTTACTGACGCTGAGGCGCGAAAGCGTGGGG |
| BH-asv24 | GCAAGCGTTAATCGGAATTACTGGGCGTAAAGCGCGCGTAGGTGGTTCAGCAAGTTGGATGTGAAATCCCCGGGCTCAACCTGGGAACTGCATCCAAAACTACTGAGCTAGAGTACGGTAGAGGGTGGTGGAATTTCCTGTGTAGCGGTGAAATGCGTAGATATAGGAAGGAACACCAGTGGCGAAGGCGACCACCTGGACTGATACTGACACTGAGGTGCGAAAGCGTGGGG |
| BH-asv26 | GCAAGCGTTAATCGGATTTACTGGGCGTAAAGCGTGCGTAGGCGGTTCGTTAAGTCGATTGTGAAAGCCCTGGGCTTAACCTGGGAATTGCAGTCGATACTGGCGGACTAGAGAACGGTAGAGGGAGGCGGAACTCCAGGTGTAGCGGTGAAATGCGTAGATATCTGGAAGAACACCGATGGCGAAGGCAACCTCCTGGGCCTGTTCTGACGCTGAGGCACGAAAGCGTGGGT |
| BH-asv27 | CCAAGCGTTAATCGGAATTACTGGGCGTAAAGCGTGCGCAGGCGGTTGTGCAAGACCGATGTGAAATCCCCGAGCTTAACTTGGGAATTGCATTGGTGACTGCACGGCTAGAGTGTGTCAGAGGGGGGTAGAATTCCACGTGTAGCAGTGAAATGCGTAGAGATGTGGAGGAATACCGATGGCGAAGGCAGCCCCCTGGGATAACACTGACGCTCATGCACGAAAGCGTGGGG |
| BH-asv28 | GCTAGCGTTGCTCGGAATCACTGGGCGTAAAGGGTGCGTAGGCGGGTTTTTAAGTCAGAGGTGAAATCCTGGAGCTCAACTCCAGAACTGCCTTTGATACTGAAAGTCTTGAGTATGGGAGAGGTGAGTGGAACTGCGAGTGTAGAGGTGAAATTCGTAGATATTCGCAAGAACACCAGTGGCGAAGGCGGCTCACTGGCCCATAACTGACGCTGAGGCACGAAAGCGTGGGG |
| BH-asv32 | ACGAGCATTATCCGGATTTATTGGGCGTAAAGGAAGCGTAGGCGGGGAGGTTGATCCATTGTTAAAAGCATTTGCTTAACAAATGTGTGCATTGGAGATCGCCTCCCTAGAGTTAATCAGGGGGTACTGGAATTCAATGTGTAGCGGTGGAATGCGTAGATATATTGAGGAACACCAGAGGCTAAGGCGAGTACCTAGGATATAACTGACGCTGAGGCTTGAAAGCGTGGGG |
| BH-asv35 | CCGAGTGATGACCGTGATTATTGGGCCTAAAGCGTCCGTAGCCGGCCGGACAAGTCCGTTGGGAAATCGACGCGCTCAACGCGTCGGCGTCCAGCGGAAACTGTTCGGCTTGGGACCGGAAGGCTCGAGGAGTACGTCCGGGGTAGGAGTGAAATCCCGTAATCCTGGACGGACTACCGATGGGGAAACCACCTCGAGGAGGACGGATCCGACGGTCAGGGACGAAAGCTGAGAT |
| BH-asv37 | GCTAGCGTTGCTCGGAATCACTGGGCGTAAAGCGCACGTAGGCGGACTCTTAAGTCGGTGGTGAAATCCTGGAGCTCAACTCCAGAACTGCCTTCGATACTGAGAGTCTCGAGTCCGGGAGAGGTGAGTGGAACTGCGAGTGTAGAGGTGAAATTCGTAGATATTCGCAAGAACACCAGTGGCGAAGGCGGCTCACTGGCCCGGTACTGACGCTGAGGTGCGAAAGCGTGGGG |
| BH-asv41 | ACGAGCATTATCCGGATTTATTGGGCGTAAAGGAAGCGTAGGTGGGGAGGTTGATCCATTGTTAAAGGCATTTGCTTAACAAATGTGTGCGATGGAGATCGCCTCCCTAGAGTTAATCAGGGGGTACTGGAATTCAATGTGTAGCGGTGGAATGCGTAGATATATTGAGGAACACCAGAGGCTAAGGCGAGTACCTAGGATATAACTGACACTGAGGCTTGAAAGCGTGGGG |
| BH-asv42 | CCGAGTGGTGGCCACTATTATTGGGCTTAAAGTGTCCGTAGCTGGCCTGGTAAGTTCTCTGTGAAATCCGGCCGCTCAACGGTCGGGCGCGCAGGGAATACTGCCGGGCTTGGGGCCGGACGGCGTCGGGGCTATTCCCGGGGTAGGAGTGAAATCCTAAAATCCTGGGAGGAGCACCTGTGGCGAAAGCGCCCGGCGAGGACGGACCCGACGGTCAGGGACGAAAGCCAGGGG |
| BH-asv44 | GCAAGCGTTAATCGGAATTATTGGGCGTAAAGCGAGTGCAGACGGTTACTTAAGCCAGATGTGAAATCCCCAAGCTTAACTTGGGACGTGCATTTGGAACTGGGTGACTAGAGTGTGTCAGAGGGAGGTAGAATTCCACATGTAGCGGTGGAATGCGTAGAGATGTGGAGGAATACCGATGGCGAAGGCAGCCTCCTGGGATAACACTGACGTTGAGGCTCGAAAGCGTGGGG |
| BH-asv47 | CCAAGTGGTGGCCGCTATTATTGGGCTTAAAGAGTTCGTAGCCGGCCCGGTAAGTTCTCTGTGAAATCCGGCCGCTCAACGGTCGGGCGCGCAGGGAATACTGCCGGGCTTGGGGCCGGACGGCGTCGGGGCTATTCCCGGGGTAGGAGTGAAATCCTGAAATCCCGGGAGGAGCACCTGTGGCGAAAGCGCCCGGCGAGGACGGACCCGACGGTCAGGAACGAAAGCCAGGGG |
| BH-asv48 | GCTAGCGTTGTTCGGAATTACTGGGCGTAAAGCGCACGTAGGCGGATACTTAAGTCAGGGGTGAAATCCCGGGGCTCAACCCCGGAACTGCCTTTGATACTGGGTATCTCGAGTCCGGAAGAGGTGAGTGGAATTCCGAGTGTAGAGGTGAAATTCGTAGATATTCGGAGGAACACCAGTGGCGAAGGCGGCTCACTGGTCCGGTACTGACGCTGAGGTGCGAAAGCGTGGGG |
| BH-asv53 | CCAAGCGTTATCCGGAATCACTGGGCGTAAAGGGTGCGCAGGCGGGCACGTAAGTCCGACGTGAAAGCCCCGGGCTCAACCCGGGAATTGCGCCGGAAACTACGTGTCTTGAGGACCAGAGAGGCAGGGAGAATTCCCGGTGTAGCGGTGAAATGCGTAGATATCGGGAGGAATACCTGTGGCGAAAGCGCCCTGCTGGATGGTTCCTGACGCTGAGGCACGAAAGCGTGGGT |
| BH-asv54 | ACGAGCATTATCCGGATTTATTGGGCGTAAAGGAAGCGTAGGTGGGGAGGTTGATCCATTGTTAAAGGCATTTGCTTAACAAATGTGTGCGATGGAGATCGCCTCCCTAGAGTTAATCAGGGGGTACTGGAATTCAATGTGTAGCGGTGGAATGCGTAGATATATTGAGGAACACCAGAGGCTAAGGCGAGTGCCTAGGATATAACTGACACTGAGGCTTGAAAGCGTGGGG |
| BH-asv56 | CCAAGTGGTGGCCGCTATTATTGGGCTTAAAGAGTTCGTAGCCGGCCCGGTAAGTTCTCTGTGAAATCCGGCCGCTCAACGGTCGGGCGCGCGGGGAATACTGCCGGGCTTGGGGCCGGACGGCGTCGGGGCTATTCCCGGGGTAGGAGTGAAATCCTGAAATCCCGGGAGGAGCACCTGTGGCGAAAGCGCCCGGCGAGGACGGACCCGACGGTCAGGAACGAAAGCCAGGGG |
| BH-asv61 | GCTAGCGTTGCTCGGAATCACTGGGCGTAAAGGGCGCGTAGGCGGACTCTTAAGTCGGGGGTGAAAGCCCAGGGCTCAACCCTGGAATTGCCTTCGATACTGGGAGTCTTGAGTTCGGAAGAGGTTGGTGGAACTGCGAGTGTAGAGGTGAAATTCGTAGATATTCGCAAGAACACCAGTGGCGAAGGCGGCCAACTGGTCCGAAACTGACGCTGAGGCGCGAAAGCGTGGGG |
| BH-asv65 | GCAAGCGTTATCGGAATTACTGGGCGTAAAGCGTACGTAGGCGGTGTGTTAAGCTAGATGTGAAAGCCCCGGGCTCAACCTGGGAATAGCATTTAGAACTGGCACGCTAGAGTCCTGAAGAGGGTGGTAGAATTTCCAGTGTAGCGGTGAAATGCGTAGATATTGGAAGGAATACCGGTGGCGAAGGCGGCCACCTGGTCAGAGACTGACGCTGAGGTACGAAAGCGTGGGG |
| BH-asv66 | GCTAGCGTTGTTCGGAATTACTGGGCGTAAAGCGCACGTAGGCGGCTACTCAAGTCAGAGGTGAAAGCCCGGGGCTCAACCCCGGAACTGCCTTTGAAACTAGGTGGCTAGAATCTTGGAGAGGCGAGTGGAATTCCGAGTGTAGAGGTGAAATTCGTAGATATTCGGAAGAACACCAGTGGCGAAGGCGACTCGCTGGACAAGTATTGACGCTGAGGTGCGAAAGCGTGGGG |
| BH-asv68 | CCAAGTGGTGACCACTATTATTGGGCTTAAAGAGTTCGTAGCCGGCCCGGTAAGTTCTCTGTGAAATCCGGCCGCTCAACGGTCGGGCGCGCAGGGAATACTGCCGGGCTTGGGGCCGGACGGCGCTGGAGCTATTCCCGGGGTAGGAGTGAAATCCTAAAATCCTGGGAGGAGCACCTGTGGCGAAAGCGTCCAGCGAGGACGGACCCGACGGTGAGGGACGAAAGCCAGGGG |
| BH-asv69 | GCTAGCGTTGCTCGGAATCACTGGGCGTAAAGGGCGCGTAGGCGGCGTTTTAAGTCGGGGGTGAAAGCCTGTGGCTCAACCACAGAATGGCCTTCGATACTGGGACGCTTGAGTATGGTAGAGGTTGGTGGAACTGCGAGTGTAGAGGTGAAATTCGTAGATATTCGCAAGAACACCGGTGGCGAAGGCGGCCAACTGGACCATTACTGACGCTGAGGCGCGAAAGCGTGGGG |
| BH-asv71 | GCTAGCGTTGCTCGGAATCACTGGGCGTAAAGGGTGCGTAGGCGGGTCTTTAAGTCAGAGGTGAAATCCTGGAGCTCAACTCCAGAACTGCCTTTGATACTGAAGGTCTTGAGTTCGAGAGAGGTGAGTGGAACTGCGAGTGTAGAGGTGAAATTCGTAGATATTCGCAAGAACACCAGTGGCGAAGGCGGCTCACTGGCTCGATACTGACGCTGAGGCACGAAAGCGTGGGG |
| BH-asv72 | GCAAGCGTTGTCCGGAATCACTGGGCGTAAAGGGTCCGCAGGCGGCCTGACAACTCAGGCGTGAAATCCCACGGCTCAACCGTGGAATTGCGTCTGAAAGTGTCTGGCTCGAGGGCAGGAAGGGAGAGTGGAACTCCGGGTGGAGCGGTGAAATGCGCAGATATCCGGAGGAACACCGGAGGCGAAAGCGACTCTCTAGTCTGCTCCTGACGCTGAGGGACCAAAGCCAGGGT |
| BH-asv74 | GCGAGCGTTGTCCGGATTTATTGGGCGTAAAGAGTTCGTAGGCGGTTTGTTAAGTCTGATGTTAAAGATCGGGGCTCAACCCTGGGAGTGCATTGGATACTGGCAGACTGGAGTGTGGTAGAGGCTAGTGGAATTCCTAGTGTAGCGGTGAAATGCGTAGATATTAGGAAGAACACCAGTGGCGTAGGCGACTAGCTGGGCCATAACTGACGCTGAGGAACGAAAGCCAGGGG |
| BH-asv75 | GCAAGCGTTGTTCGGAGTTACTGGGCGTAAAGAGTCTGTAGGTGGTCTGTCAAGTCTTTGGTGAAAGCCGTGGCTTAACCATGGGAATGCCAAAGAGACTGGCAGACTGGAGGCTGGGAGAGGGAAGCGGAATTTTCTGGTGTAGCGGTGAAATGCGTAGATATCAGAAGGAAGGCCGGTGGCGAAGGCGGCTTCCTGGAACAGACCTGACACTGAGAGACGAAAGCGTGGGG |
| BH-asv76 | CCAAGTGGTGGCCGCTATTATTGGGCTTAAAGAGTCCGTAGCCGGCCCGGTAAGTTCTCTGTGAAATCCGGCCGCTCAACGGTCGGGCGCGCAGGGAATACTGCCGGGCTTGGGGCCGGACGGCGTCGGGGCTATTCCAGGGGTAGGAGTGAAATCCTGAAATCCCTGGAGGAGCACCTGTGGCGAAAGCGCCCGGCGAGGACGGACCCGACGGTCAGGGACGAAAGCCAGGGG |
| BH-asv79 | GCAAGTGTTATCCGGAATCACTGGGCGTAAAGCGTCTGTAGGTGGTTTGATAAGTCAACTGTTAAATCTTGAAGCTCAACTTCAAAATCGCAGTCGAAACTATTAGACTTGAGTATAGTAGGGGTAAAGGGAATTTCCAGTGGAGCGGTGAAATGCGTAGAGATTGGAAAGAACACCGATGGCGAAGGCACTTTACTGGGCTATTACTGACACTCAGAGACGAAAGCTAGGGT |
| BH-asv83 | CCAAGCGTTATCCGGAATTACTGGGCGTAAAGGGTGCGCAGGCGGACACGTAAGTCCGACGTGAAAGCCCCGGGCTCAACCCGGGAAGGGCGCCGGAAACTACGTGTCTTGAGGACCAGAGAGGCAGAGAGAATTCCTGGTGTAGCGGTGAAATGCGTAGATATCAGGAGGAATGCCAGTGGCGAAAGCGCTCTGCTGGATGGTTCCTGACGTTGAGGCACGAAAGCGTGGGT |
| BH-asv85 | GCTAGCGTTGTTCGGAATTACTGGGCGTAAAGCGCACGTAGGCGGATTGTTAAGTTAGGGGTGAAATCCCAGGGCTCAACCCTGGAACTGCCTTTAATACTGGCAATCTCGAGTCCGAGAGAGGTGAGTGGAATTCCGAGTGTAGAGGTGAAATTCGTAGATATTCGGAGGAACACCAGTGGCGAAGGCGGCTCACTGGCTCGGTACTGACGCTGAGGTGCGAAAGCGTGGGG |
| BH-asv89 | CCAAGCGTTATCCGGAATCACTGGGCGTAAAGGGTGCGCAGGCGGGCACGTAAGTCCGACGTGAAAGCCCCGGGCTCAACCCGGGAAGTGCGCCGGAAACCACGTGTCTTGAGGACCAGAGAGGCAGGGAGAATTCCCGGTGTAGCGGTGAAATGCGTAGATATCGGGAGGAATACCTGTGGCGAAAGCGCCCTGCTGGATGGTTCCTGACGCTGAGGCACGAAAGCGTGGGT |
| BH-asv90 | GCAAGCGTTAATCGGAATTACTGGGCGTAAAGCGCGCGTAGGTGGTTTGATAAGCGAGATGTGAAAGCCCCGGGCTTAACCTGGGAACGGCATTTCGAACTGTCAGGCTAGAGTATGGTAGAGGAGTGTGGAATTTCCTGTGTAGCGGTGAAATGCGTAGATATAGGAAGGAACACCAGTGGCGAAGGCGGCACTCTGGACCAATACTGACACTGAGGTGCGAAAGCGTGGGG |
| BH-asv93 | GCTAGCGTTGCTCGGAATCACTGGGCGTAAAGGGTGCGTAGGCGGGTCTTTAAGTCAGGGGTGAAATCCTGGAGCTCAACTCCAGAACTGCCTTTGATACTGAGGATCTTGAGTTCGGAAGAGGTGAGTGGAACTGCGAGTGTAGAGGTGAAATTCGTAGATATTCGCAAGAACACCAGTGGCGAAGGCGGCTCACTGGTCCGATACTGACGCTGAGGCACGAAAGCGTGGGG |
| BH-asv95 | CCAAGTGGTGGCCGCTATTATTGGGCTTAAAGAGTCCGTAGCCGGCCCGGTAAGTTCTCTGTGAAATCCGGCCGCTCAACGGTCGGGCGCGCAGGGAATACTGCCGGGCTTGGGGCCGGACGGCGTCGAGGCTATTCCCGGGGTAGGAGTGAAATCCTGAAATCCCGGGAGGAGCACCTGTGGCGAAAGCGCTCGGCGAGGACGGACCCGACGGTCAGGGACGAAAGCCAGGGG |
| BH-asv99 | CCAAGCGTTATCCGGAATTACTGGGCGTAAAGGGTGCGCAGGCGGACACGTAAGTCCGACGTGAAAGCCCCGGGCTCAACCCGGGAAGGGCGCCGGAAACTACGTGTCTTGAGGACCAGAGAGGCAGAGAGAATTCCTGGTGTAGCGGTGAAATGCGTAGATATCAGGAGGAATGCCAGTGGCGAAAGCGCTCTGCTGGATGGTTCCTGACGCTGAGGCACGAAAGCGTGGGT |
| BH-asv104 | GCTAGCGTTGCTCGGAATCACTGGGCGTAAAGGGTGCGTAGGCGGGTTTTTAAGTCAGGGGTGAAATCCTGGAGCTCAACTCCAGAACTGCCTTTGATACTGAAGATCTTGAGTCCGGGAGAGGTGAGTGGAACTGCGAGTGTAGAGGTGAAATTCGTAGATATTCGCAAGAACACCAGTGGCGAAGGCGGCTCACTGGCCCGGTACTGACGCTGAGGCACGAAAGCGTGGGG |
| BH-asv106 | GCAAGCATTATTCGGATTTATTGGGCGTAAAGCAAGCGCAGGCGGATGTGTAAGTTCTGTGTTAAAAGTAGCTACTTAATAGTTGTTTGCACCGAATACTACATGTCTAGAATGTGGTAGGAAGTTTTGGAATTAAACATGGAGCGGTGGAATGTGTAGATATGTTTAAGAACACCGGAGGCGAAGGCGAAAACTTAGGCCATAATTGACGCTTAGGCTTGAAAGTGTGGGG |

| | |
|---|---|
| BH-asv108 | CCGAGTGGTGGCCGCTATTATTGGGCTTAAAGAGTCCGTAGCCGGCCCGGTAAGTTCTCTGTGAAATCCGGCCGCTCAACGGTCGGGCGCGCAGGGAATACTGCCGGGCTTGGGGCCGGACGGCGTCGAGGCTATTCCCGGGGTAGGAGTGAAATCCTGAAATCCCGGGAGGAGCACCTGTGGCGAAAGCGCTCGGCGAGGACGGACCCGACGGTCAGGGACGAAAGCCAGGGG |
| BH-asv111 | GCAAGCGTTAATCGGAATTACTGGGCGTAAAGCGTGCGCAGGCGGTGATGTAAGACAGTTGTGAAATCCCCGGGCTCAACCTGGGAACTGCATCTGTGACTGCATCGCTGGAGTACGGCAGAGGGGGATGGAATTCCGCGTGTAGCAGTGAAATGCGTAGATATGCGGAGGAACACCGATGGCGAAGGCAATCCCCTGGGCCTGTACTGACGCTCATGCACGAAAGCGTGGGG |
| BH-asv115 | GCAAGCGTTAATCGGAATTACTGGGCGTAAAGCGTGCGCAGGCGGTTATGCAAGACAGAGGTGAAATCCCCGGGCTCAACCTGGGAACTGCCTTTGTGACTGCATAGCTAGAGTACGGTAGAGGGGGATGGAATTCCGCGTGTAGCAGTGAAATGCGTAGATATGCGGAGGAACACCGATGGCGAAGGCAATCCCCTGGACCTGTACTGACGCTCATGCACGAAAGCGTGGGG |
| BH-asv116 | GCAAGCGTTAATCGGATTTACTGGGCGTAAAGGGTGCGTAGGTGGTTGTATAAGTCGGATGTGAAAGCCCGGGGCTTAACCTCGGAACTGCATTCGATACTGTATGACTAGAGTTCGGTAGAGGGCGGCGGAATTTCCGGTGTAGCGGTGAAATGCGTAGATATCGGAAGGAACATCAATGGCGAAGGCAGCCGCCTGGGCCTGAACTGACACTGAGGCACGAAAGCGTGGGT |
| BH-asv124 | CCAAGTGGTGGCCACGTTTATTGGGCTTAAAGTGTTCGTAGCCTGCTCTGTGAGTCCGCTGTGAAATCTGGCAGCTCAACTGTCAGGCGTGCAGCGGATACCACAGGGCTTGGGACCGGAAGGCGTCGAGGCTATTCCTGGGGTAGGGGTGAAATCCTGAAATCCCAGGAGGAGCACCAGTGGCGAAAGCTCTCGGCGATGACGGGTCCGACGGTGAGGGACGAAAGCCAGGGG |
| BH-asv128 | GCAAGCGTTATCCGAAATCACTGGGCGTAAAGGGTGCGCAGGCGGCACGCAGGTCCGACGTGAAAGCCCCGGGCTTAACCCGGGAAGGGCGCCGGAAACCACGTGTCTTGAGGACCAGAGAGGCAGGGAGAATTCCCGGTGTAGCGGTGAAATGCGTAGATATCGGGAGGAATACCAGTGGCGAAAGCGCCCTGCTGGATGGTTCCTGACGCTGAGGCACGAAAGCGTGGGT |
| BH-asv140 | GCGAGCGTTAATCGGAATTATTGGGCGTAAAGCGAGTGCAGACGGTTGTTTAAGCCAGATGTGAAATCCCCGAGCTTAACTTGGGACGTGCATTTGGAACTGGATAACTAGAGTGTGTCAGAGGGAGGTAGAATTCCACATGTAGCGGTGGAATGCGTAGAGATGTGGAGGAATACCGATGGCGAAGGCAGCCTCCTGGGATAACACTGACGTTGAGGCTCGAAAGCGTGGGG |
| BH-asv141 | CCAAGCGTTATCCGGAATTACTGGGCGTAAAGGGTGCGCAGGCGGCACATAAGTCCGACGTGAAAGCCCCGGGCTTAACCCGGGAATGGCGCCGGAAACTATGTGTCTTGAGGACCAGAGAGGCAGGGAGAATTCCTGGTGTAGCGGTGAAATGCGTAGATATCAGGAGGAATACCTGTGGCGAAAGCGCCCTGCTGGATGGTTTCTGACGCTGAGGCACGAAAGCGTGGGG |
| BH-asv142 | GCAAGCGTTGTCCGGAATTACTGGGCGTAAAGAGCTCGTAGGTGGTTTGTCACGTCGTCTGTGAAATTCCACAGCTTAACTGTGGGCGTGCAGGCGATACGGGCTGACTTGAGTACTGTAGGGGTAACTGGAATTCCTGGTGTAGCGGTGAAATGCGCAGATATCAGGAGGAACACCGATGGCGAAGGCAGGTTACTGGGCAGTTACTGACGCTGAGGAGCGAAAGCATGGGT |
| BH-asv144 | CCAAGCGTTATCCGGAATTACTGGGCGTAAAGGGTGCGCAGGCGGACACGTAAGTCCGACGTGAAAGCCCCGGGCTCAACCCGGGAAGGGCGCCGGAAACTACATGTCTTGAGGACCAGAGAGGCAGAGAGAATTCCTGGTGTAGCGGTGAAATGCGTAGATATCAGGAGGAATACCAGTGGCGAAAGCGCTCTGCTGGATGGTTCCTGACGCTGAGGCACGAAAGCGTGGGT |
| BH-asv152 | CCGAGTGGTGGCCGCTATTATTGGGCTTAAAGAGTCCGTAGCCGGCCCGGTAAGTTCTCTGTGAAATCCGGCCGCTCAACGGTCGGGCGCGCAGGGAATACTGCCGGGCTTGGGGCCGGACGGCGTCGGGGCTATTCCCGGGGTAGGAGTGAAATCCTGAAATCCCGGGAGGAGCACCTGTGGCGAAAGCGCCCGGCGAGGACGGACCCGACGGTCAGGGACGAAAGCCAGGGG |
| BH-asv153 | GCAAGCGTTAATCGGAATTACTGGGCGTAAAGCGTGCGCAGGCGGTTGTGTAAGTCAGATGTGAAATCCCCGGGCTCAACCTGGGAATTGCATTTGAGACTGCACGGCTAGAGTGTGTCAGAGGGGGGTAGAATTCCACGTGTAGCAGTGAAATGCGTAGATATGTGGAGGAATACCGATGGCGAAGGCAGCCCCCTGGGATAACACTGACGCTCATGCACGAAAGCGTGGGG |
| BH-asv156 | CCGAGTGGTGGCCGCTGTTATTGGGCTTAAAGTGTCCGTAGCCGGCCCGGTAAGTTCTCTGTGAAATCCGGCCGCTCAACGGTCGGGCGCGCAGGGAATACTGCCGGGCTTGGGGCCGGACGGCGCTGGGGCTATTCCCGGGGTAGGAGTGAAATCCTGAAATCCCGGGAGGAGCACCTGTGGCGAAAGCGCCCAGCGAGGACGGACCCGACGGTCAGGGACGAAAGCCAGGGG |
| BH-asv158 | CCAAGTGGTAGCCACTTTTATTGGGCTTAAAGTGTTCGTAGCCGGCCGTGTAAGTCTGCTGTGAAATCCACGCGCTCAACGTGTGGAAGTGCAGCGGATACTGCATGGCTAGGAACCGGATGGCGTCGAGGCTATTCCAGGGGTAGGAGTGAAATCCAAAAATCCCTGGAGGAGCACCTGTGGCGAAAGCGCTCGACGAATACGGATTCGACGGTGAGGAACGAAAGCCAGGGG |
| BH-asv159 | GCAAGCGTTAATCGGATTTACTGGGCGTAAAGCGTGCGTAGGCGGTTCGTTAAGTCGATTGTGAAAGCCCTGGGCTCAACCTGGGAATTGCAGTCGATACTGATGGACTAGAGAACGGTAGAGGGAGGCGGAACTCCAGGTGTAGCGGTGAAATGCGTAGATATCTGGAAGAACACCGATGGCGAAGGCAACCTCCTGGGCCTGTTCTGACGCTGAGGCACGAAAGCGTGGGT |
| BH-asv162 | GCAAGCGTTGTCCGGAATCACTGGGCGTAAAGGGTCCGCAGGCGGATCAGCAACTCAGGCGTGAAATCCCGCGGCTTAACCGCGGAGGTGCGTCTGAAAGTGTTGATCTCGAGGACAGGAAGGGAGAGTGGAACTCCGGGTGGAGCGGTGAAATGCGCAGATATCCGGAGGAACACCGGAGGCGAAAGCGACTCTCTAGCCTGTACCTGACGCTGAGGGACCAAAGCCAGGGT |
| BH-asv172 | GCAAGCGTTATCCGGAATTATTGGGCGTAAAGAGCTCGTAGGCGGTTTGTCGCGTCTGTCGTGAAAGTCCGGGGCTCAACTCCGGATCTGCGGTGGGTACGGGCAGACTAGAGTGATGTAGGGGAGACTGGAATTCCTGGTGTAGCGGTGGAATGCGCAGATATCAGGAGGAACACCGATGGCGAAGGCAGGTCTCTGGGCATTAACTGACGCTGAGGAGCGAAAGCATGGGG |
| BH-asv173 | CCGAGCGTTAATCGGAATTACTGGGCGTAAAGGGTCTGTAGGTGGTTTGTTAAGTCAGATGTGAAAGCCCAGGGCTCAACCTTGGAACTGCATTTGATACTGGCAAACTAGAGTATGGTAGAGGAATGGGGAATTTCTGGTGTAGCGGTGAAATGCGTAGAGATCAGAAGGAACACCAATGGCGAAGGCAACATTCTGGACCAATACTGACACTGAGGGACGAAAGCGTGGGG |
| BH-asv174 | GCTAGCGTTGTTCGGAATTACTGGGCGTAAAGCGCACGTAGGCGGCTTTGTAAGTCAGAGGTGAAAGCCTGGAGCTCAACTCCAGAACTGCCTTTGAGACTGCATCGCTTGAATCCAGGAGAGGTGAGTGGAATTCCGAGTGTAGAGGTGAAATTCGTAGATATTCGGAAGAACACCAGTGGCGAAGGCGGCTCACTGGACTGGTATTGACGCTGAGGTGCGAAAGCGTGGGG |
| BH-asv175 | ACAAGCGTTGTCCGGATTTATTGGGCGTAAAGGGAGCGCAGGCGGTCTGTTTAGTCTAATGTGAAAGCCCACGGCTTAACCGTGGAACGGCATTGGAAACTGACAGACTTGAATGTAGAAGAGGAAAATGGAATTCCAAGTGTAGCGGTGGAATGCGTAGATATTTGGAGGAACACCAGTGGCGAAGGCGATTTTCTGGTCTAACATTGACGCTGAGGCTCGAAAGCGTGGGG |
| BH-asv179 | GCTAGCGTTGCTCGGAATCACTGGGCGTAAAGGGTGCGTAGGCGGGTCTTTAAGTCAGAGGTGAAAGCCTGGAGCTCAACTCCAGAACTGCCTTTGATACTGAGGATCTTGAGTTCGGGAGAGGTGAGTGGAACTGCGAGTGTAGAGGTGAAATTCGTAGATATTCGCAAGAACACCAGTGGCGAAGGCGGCTCACTGGCCCGATACTGACGCTGAGGCACGAAAGCGTGGGG |
| BH-asv180 | ACGAGCATTATCCGGATTTATTGGGCGTAAAGGAAGCGTAGGCGGGGAGGTTGATCCATTGTTAAAGGCATTTGCTTAACAAATGTGTGCGATGGAGATCGCCTCCCTAGAGTTAATCAGGGGGTACTGGAATTCAATGTGTAGCGGTGGAATGCGTAGATATATTGAGGAACACCAGAGGCTAAGGCGAGTACCTAGGATATAACTGACGCTGAGGCTTGAAAGCGTGGGG |
| BH-asv190 | GCAAGCGTTGTCCGGAATCACTGGGCGTAAAGGGTCCGCAGGCGGATCAGCAACTCAGGCGTGAAATCCCGCGGCTTAACCGCGGAGGTGCGTCTGAAAGTGTTGATCTCGAGGACAGGAAGGGAGAGTGGAACTCCGGGTGGAGCGGTGAAATGCGCAGATATCCGGAGGAACACCGGAGGCGAAAGCGACTCTCTAGCCTGTTCCTGACGCTGAGGGACCAAAGCCAGGGT |
| BH-asv193 | GCTAGCGTTGCTCGGAATTACTGGGCGTAAAGGGCGCGTAGGCGGATGTTTAAGTCAGAGGTGAAAGCCCAGGGCTCAACCTTGGAAATGCCTTTGATACTGAACATCTTGAGTATGGGAGAGGTGAGTGGAACTCCGAGTGTAGAGGTGAAATTCGTAGATATTCGGAAGAACACCAGTGGCGAAGGCGACTCACTGGCCCATTACTGACGCTGAGGCGCGAAAGCGTGGGG |
| BH-asv194 | GCTAGCGTTGCTCGGAATTACTGGGCGTAAAGGGCGCGTAGGCGGACAGTTAAGTCAGAAGTCAAAGCCCAGGGCTCAACCTTGGAATCGCTTTTGATACTGGCTGTCTTGAGTGCGGGAGAGGTGTGTGGAACTCCGAGTGTAGAGGTGAAATTCGTAGATATTCGGAAGAACACCAGTGGCGAAGGCGACACACTGGCCCGTTACTGACGCTGAGGCGCGAAAGCGTGGGG |
| BH-asv196 | GCAAGCGTTATCCGGCATGACTGGGCGTAAAGAGTCCGTAGACGGTAAAGTAAGTTCTTTGTTAAATTTTAAGACTCAATCTTAACACAGCAATTAATACTACTTTACTTTGAGTTTAATACAGAAAAGTAGAATTTTATACGGTAGAGGTGAAATTTGCAGATATATAAAGGAATGCCAATAGCGAAGGCAACTTTTTGGTATAAACTGACGTTGAGGGACGAAAGCGTGGGT |
| BH-asv206 | ACGAGTAGTGGTCACTTTTATTGAGCCTAAAGCGTTCGTAGCCGGTTTTGTAAATCTTCAGATAAAGCCTGAAGCTTAACTCCAGAAAGTCTGAAGAGACTGCAAGACTTGAGATCGGGTGAGGTTAAACGTACTTTCAGGGTAGGGGTAAAATCCTGTAATCCCGGAAGGACGACCAGTGGCGAAAGCGTTTAACTAGAACGAATCTGACGGTAAGGAACGAAGGCTAGGGT |
| BH-asv207 | CCAAGCGTTATCCGGAATCACTGGGTGTAAAGGGTACGCAGGCGGACATGTAGGTCCAATGTGAAAGTCCTGGGCTCAACCCAGGAATTGCATTGGAAACCCCATGTCTTGAGGATCAGAGAGGCAGGCGCAATTCGTGGTGTAGCGGTGAAATGCGTAGATATCACGAGGAACACCAGTGGTGTAGACGGCCTGCTGGATGAATCCTGACGCTGAGGTACGAAAGCGTGGGG |
| BH-asv210 | GCTAGCGTTGTTCGGAATTACTGGGCGTAAAGGGCGCGTAGGCGGTTTATCAAGTCAGGCGTGAAAGCCCCGGGCTCAACCTGGGAATTGCGTTTGATACTGGTAGACTTGAGTTCTGGAGAGGTGAGTGGAATTCCCAGTGTAGAGGTGAAATTCGTAGATATTGGGAAGAACACCGGTGGCGAAGGCGGCTCACTGGACCGATACTGACGCTGAGGCGCGAAAGCGTGGGG |
| BH-asv217 | GCAAGCGTTAATCGGAATTACTGGGCGTAAAGCGTGCGCAGGCGGTTTGTTAAGTCTGTTGTGAAATCCCCGAGCTCAACTTGGGAATTGCAATGGATACTGGCAAGCTGGAGTATGGTAGAGGAAGGTGGAATTCCCGGTGTAGCAGTGAAATGCGTAGAGATCGGGAGGAACACCAGTGGCGAAGGCGGCCTTCTGGACCAATACTGACGCTCATGCACGAAAGCGTGGGG |
| BH-asv222 | GCGAGCGTTAATCGGAATAACTGGGCGTAAAGGGCACGCAGGCGGTGACTTAAGTGAGGTGTGAAAGCCCCGGGCTTAACCTGGGAATTGCATTTCATACTGGGTCGCTAGAGTACTTTAGGGAGGGGTAGAATTCCACGTGTAGCGGTGAAATGCGTAGAGATGTGGAGGAATACCGAAGGCGAAGGCAGCCCCTTGGGAATGTACTGACGCTCATGTGCGAAAGCGTGGGG |
| BH-asv225 | CCGAGCGTTATCCGGATTTATTGGGCGTAAAGCGAGCGCAGGCGGTTAGATAAGTCTGAAGTTAAAGGCTGTGGCTTAACCATAGTACGCTTTGGAAACTGTTTAACTTGAGTGCAGAAGGGGAGAGTGGAATTCCATGTGTAGCGGTGAAATGCGTAGATATATGGAGGAACACCGGTGGCGAAAGCGGCTCTCTGGTCTGTAACTGACGCTGAGGCTCGAAAGCGTGGGG |
| BH-asv228 | GCAAGCGTTAATCGGAATCACTGGGCGTAAAGGGTGCGTAGGCGGTACGTTAGGTCTGTCGTGAAATCCCCGGGCTCAACCTGGGAATGGCGGTAGAAACCGGCGCACTAGAGTATGGGAGAGGGTGGTGGAATTCCAGGTGTAGCGGTGAAATGCGTAGAGATCTGGAGGAACATCAGTGGCGAAGGCGGCCACCTGGCCCAATACTGACGCTGAGGCACGAAAGCGTGGGG |
| BH-asv231 | CCAAGCGTTATCCGGAATCACTGGGCGTAAAGGGTGAGCAGGCGGGCATGCAAGTTCGACGTGAAAGCCCCGGGCTTAACCCGGGAAGTGCGCCGAAAACTGCATGTCTTGAGGACCAGAGAGGCAGGGAGAATTCCTGGTGTAGCGGTGAAATGCGTAGATATCAGGAGGAATACCTGTGGCGAAAGCGCCCTGCTGGATGGTTCCTGACGCTGAGTCACGAAAGCGTGGGT |
| BH-asv235 | CCAAGCGTTATCCGGAATTACTGGGCGTAAAGGGTGCGCAGGCGGGCACGTAAGTCCGACGTGAAAGCCCCGAGCTCAACTCGGGAAGTGCGCCGGAAACTACGTGTCTTGAGGACCAGAGAGGCAGGGAGAATTCCTGGTGTAGCGGTGAAATGCGTAGATATCAGGAGGAATACCTGTGGCGAAAGCGCCCTGCTGGATGGTTCCTGACGCTGAGGCACGAAAGCGTGGGT |
| BH-asv237 | ACTAGCGTTGTTCGGAATCACTGGGCGTAAAGCGCACGTAGGCGGATTTGTAAGTCAGGGGTGAAATCCCGGGGCTCAACCTCGGAACTGCCTTTGATACTGCGAATCTTGAGTCCGATAGAGGTGGGTGGAATTCCTAGTGTAGAGGTGAAATTCGTAGATATTAGGAAGAACACCGGTGGCGAAGGCGGCCCACTGGATCGGTACTGACGCTGAGGTGCGAAAGCGTGGGG |
| BH-asv244 | GCAAGCGTTGTCCGGAATTATTGGGCGTAAAGAGCTCGTAGGCGGCTTGTCGCGTCGGCTGTGAAAACCTGGGGCTCAACCCCGGGCCTGCAGCCGATACGGGCAAGCTAGAATTCGGTAGGGGAGACTGGAATTCCTGGTGTAGCGGTGAAATGCGCAGATATCAGGAGGAACACCGGTGGCGAAGGCGGGTCTCTGGGCCGATATTGACGCTGAGGAGCGAAAGCGTGGGG |
| BH-asv247 | CCAAGCGTTATCCGGAATTACTGGGCGTAAAGGGTGCGCAGGCGGACACGTAAGTCCGACGTGAAAGCCCCGGGCTCAACCCGGGAAGGGCGCCGGAAACTACGTGTCTTGAGGACCAGAGAGGCAGAGAGAATTCTTGGTGTAGCGGTGAAATGCGTAGATATCAGGAGGAATGCCAGTGGCGAAAGCGCTCTGCTGGATGGTTCCTGACGTTGAGGCACGAAAGCGTGGGT |
| BH-asv251 | GCGAGCGTTAATCGGAATTACTGGGCGTAAAGCGAGTGTAGGTGGCTCATTAAGTCACATGTGAAATCCCCGGGCTTAACCTGGGAACTGCATGTGATACTGGTGGTGCTAGAATATGTGAGAGGGAAGTAGAATTCCAGGTGTAGCGGTGAAATGCGTAGAGATCTGGAGGAATACCGATGGCGAAGGCAGCTTCCTGGCATAATATTGACACTGAGATTCGAAAGCGTGGGT |
| BH-asv257 | ACGAGCATTATCCGGATTTATTGGGCGTAAAGGAAGCGTAGGTGGGAAGGTTGATCCATTGTTAAAGGCATTTGCTTAACAAATGTGTGCGATGGAGATCGCCTTCCTAGAGTTAATCAGGGGGTACTGGAATTCAATGTGTAGCGGTGGAATGCGTAGATATATTGAGGAACACCAGAGGCTAAGGCGAGTACCTAGGATATAACTGACACTGAGGCTTGAAAGCGTGGGG |
| BH-asv260 | GCTAGCGTTGTTCGGAATTACTGGGCGTAAAGCGCACGTAGGCGGCTTTGCAAGTTAGAGGTGAAAGCCTGGGGCTCAACCCCAGAATTGCCTTTAAGACTGCATCGCTAGAATCCAGAAGAGGTGAGTGGAATTCCGAGTGTAGAGGTGAAATTCGTAGATATTCGGAAGAACACCAGTGGCGAAGGCGGCTCACTGGTCTGGTATTGACGCTGAGGTGCGAAAGCGTGGGG |
| BH-asv262 | GCTAGCGTTGTTCGGAATTACTGGGCGTAAAGCGCATGTAGGCGGATATTTAAGTCAGGGGTGAAATCCCGCAGCTCAACTGCGGAACTGCCTTTGATACTGGGTATCTTGAGTATGGAAGAGGTGAGTGGAATTCCGAGTGTAGAGGTGAAATTCGTAGATATTCGGAGGAACACCAGTGGCGAAGGCGGCTCACTGGTCCATTACTGACGCTGAGGTGCGAAAGCGTGGGG |
| BH-asv264 | CCAAGCGTTATCCGGAATCACTGGGCGTAAAGGGTGAGCAGGCGGGCACGTAGGTCCGACGTGAAAGCCCCGAGCTTAACTCGGGAATGGCGCCGGAAACCACGTGTCTTGAGGACCAGAGAGGCAGGGAGAATTCCCGGTGTAGCGGTGAAATGCGTAGATATCGGGAGGAATACCAGTGGCGAAAGCGCCCTGCTGGATGGTTCCTGACGCTGAGTCACGAAAGCGTGGGT |
| BH-asv265 | CCGAGCGTTGTCCGGAATCATTGGGCGTAAAGGGTACGTAGGCGGGTAAGCAAGTTAGAAGTGAAATCCTATAGCTCAACTATAGTAAGCTTTTAAAACTGCTCATCTTGAGGTATGGAAGGGAAAGTGGAATTCCTAGTGTAGCGGTGAAATGCGCAGATATTAGGAGGAATACCGGTGGCGAAGGCGACTTTCTGGCCATAACCTGACGCTGAGGTACGAAAGCGTGGGT |
| BH-asv274 | GCTAGCGTTGTTCGGAATCACTGGGCGTAAAGGGCGCGTAGGCGGCTGTGGCAGCCGGGTGTGAAAGCCCGGAGCTCAACTCCGGAATGGCACTCGAGACGCCACGGCTGGAGGCCGGAAGAGGAGGGTGGAATACCCAGTGTAGAGGTGAAATTCGTAGATATTGGGTGGAACACCGGTGGCGAAAGCGGCTCTCTGGTCCGGACCTGACGCTGAGGCGCGAAAGCGTGGGG |

| | |
|---|---|
| BH-asv275 | GCAAGCGTTATCCGGAATTATTGGGCGTAAAGCGCGCGCAGGCGGTTTCTTAAGTCTGATGTGAAAGCCCACGGCTCAACCGTGGAGGGTCATTGGAAACTGGGGAACTTGAGTGCAGAAGAGAAAAGCGGAATTCCACGTGTAGCGGTGAAATGCGTAGAGATGTGGAGGAACACCAGTGGCGAAGGCGGCTTTTTGGTCTGTAACTGACGCTGAGGCGCGAAAGCGTGGGG |
| BH-asv278 | TCGAGTGGTAGCCGATTTTATTGGGTCTAAAGCGTCCGTAGCCGGACCGGCAAGTCCCTTGTGAAATCGGGCGGCTTAACCGTTCGGCGTGCAGGGGAAACTGCCAGTCTTGGGGCCGGCAGGCGCCGAGGGTACTCCGGGGGTAGCGGTGAAATGCTATAATCCCTGGGGGACCGACAGTGGCGAAAGCGCTCGGCGAGGCCGGACCCGACGGTGAGGGACGAAAGCCAGAAG |
| BH-asv284 | GCAAGCGTTGTCCGGAATTATTGGGCGTAAAGAGCTCGTAGGTGGCCTGTCGCGTCGGCTGTGAAAACTCGGGGCTCAACTCCGAGCTTGCAGTCGATACGGGCAGGCTAGAGTTCGGCAGGGGAGACTGGAATTCCTGGTGTAGCGGTGAAATGCGCAGATATCAGGAGGAACACCGGTGGCGAAGGCGGGTCTCTGGGCCGATACTGACACTGAGGAGCGAAAGCGTGGGG |
| BH-asv286 | CCGAGTGGTGACCAGTATTATTGGGCTTAAAGTGTCCGTAGCTTGCCGTGTAAGTCCGCTGTGAAATCCACACGCTCAACGTGTGGCCGTGCAGCGGATACTGCATGGCTTGGGACCAGACGGCGTCGAGGCTATTCCAGGGGTAGGAGTGAAATCCTAAAATCCCTGGAGGAGCACCTGTGGCGAAAGCGCTCGGCGAGTATGGATCCGACGGTGAGGGACGAAAGCCAGGGG |
| BH-asv288 | GCGAACGTTGTTCGGATTCACTGGGCTTAAAGGGCGCGTAGGCGGACAAGCAAGTCGGGGGTGAAATCTTCCAGCTCAACTGGAAACGTGCCTTCGATACTGTTTGTCTCGAGGGAGGTAGGGGCATGTGGAACTTCCGGTGGAGCGGTGAAATGCGTAGATATCGGAAGGAACGCCGGTGACGAAAGCGGCGTGCTGGACCTCTTCTGACGCTGAGGCGCGAAAGCTAGGGT |
| BH-asv291 | CCAAGTGGTGGCCATTATTATTGGGCTTAAAGTGTTCGTAGCCGGCCGTGCAAGTCCGCTGTGAAATCCACGCGCTCAACGTGTGGGCGTGCAGCGGATACTGCATGGCTTGGGACCGGACGGCGTCGAGGCTATTCCCGGGGTAGGAGTGAAATCCTAAAATCCCGGGAGGAGCACCTGTGGAGAAATCGCTCGACGAGTACGGATCCGACGGTGAGGAACGAAAGCCAGGGG |
| BH-asv293 | CCAAGTGGTGGCCACTGTTATTGGGCTTAAAGAGTCCGTAGCCGGCCCGGTAAGTTCTCTGTGAAATCCGGCCGCTCAACGGTCGGGCGCGCAGGGAATACTGCCGGGCTTGGGGCCGGACGGCGTCGGGGCTATTCCCGGGGTAGGAGTGAAATCCTGAAATCCCGGGAGGAGCACCTGTGGCGAAAGCGCCCGGCGAGGACGGACCCGACGGTCAGGGACGAAAGCCAGGGG |
| BH-asv296 | GCAAGCGTTAATCGGATTTACTGGGCGTAAAGAGTGTGTAGGTGGCTAGCTAAGTCGGTTGTGAAAGCCCTGGGCTCAACCTGGGAATTGCAGTCGATACTGGCAAGCTCGAGTTCGGTAGAGGGTGACGGAATTCCCGGTGTAGCGGTGAAATGCGTAGATATCGGGAGGAACATCGATGGCGAAGGCAATCACCTGGGCCTGAACTGACACTGTGACACGAAAGCGTGGGT |
| BH-asv308 | CCGAGTGATGGCCGATATTATTGGGCCTAAAGCGTCCGTAGCCGGCTGTGTAAGTCTATTGGGAAATCGACCAGCTCAACTGGTCGGCGTCCGGTGGAAACTACACAGCTTGGGGCCGAGAGACTCAACGGGTACGTCCGGGGTAGGAGTGAAATCCTATAATCCTGGACGGACCACCAATGGGGAAACCACGTTGAGAGACCGGACCCGACGGTGAGGGACGAAAGCCAGGGT |
| BH-asv310 | GCTAGCGTTGTCCGGAATCATTGGGCGTAAAGCGCGTGTAGGCGGCCCGGTAAGTCCGCTGTGAAAGTCGGGGGCTCAACCCCCGAAAGCCGGTGGAAACTGTCGGGCTAGAGTACTGAAGAGGCGAGTGGAATTCCTGGTGTAGCGGTGAAATGCGCAGATATCAGGAGGAACACCAATTGCGAAGGCAGCTCGCTGGGACGTCACTGACGCTGAGACGCGAAAGCGTGGGG |
| BH-asv311 | GCGAGCGTTGTCCGGAATTACTGGGCGTAAAGAGCTCGTAGGTGGTTTGTCGCGTTGTCCGTGAAATTCCCTGGCTTAACTGGGGGCGTGCGGGCGATACGGGCAGACTGGAGTACTGCAGGGGAGACTGGAATTCCTGGTGTAGCGGTGGAATGCGCAGATATCAGGAGGAACACCGGTGGCGAAGGCGGGTCTCTGGGCAGTAACTGACGCTGAGGAGCGAAAGCGTGGGG |
| BH-asv312 | GCAGGCGTTAATCGGAATTACTGGGCGTAAAGCGTACGCAGGCGGTCTGTAAAGTCAGATGTGAAAGCCCCGGGCTCAACCTGGGAATTGCATTTGAAACTGGCAGGCTAGAGTGTGGCAGAGGGGGGTGGAATTCCACGTGTATCAGTGAAATGCGTAGAGATGTGGAGGAACACCGATGGCGAAGGCAGCCCCCTGGGCCAACACTGACGCTCAGGCACGAAAGCGTGGGG |
| BH-asv314 | GCGAGCGTTGTCCGGAATTACTGGGCGTAAAGAGCTCGTAGGTGGTTTGTCGCGTTGTTCGTGAAAACTCACAACTCAATTGTGGGCGTGCGGGCGATACGGGCAGACTAGAGTACTGCAGGGGAGACTGGAATTCCTGGTGTAGCGGTGGAATGCGCAGATATCAGGAGGAACACCGGTGGCGAAGGCGGGTCTCTGGGCAGTAACTGACGCTGAGGAGCGAAAGCGTGGGG |
| BH-asv315 | CCGAGTGATGGCCGCGATTATTGGGCCTAAAGCGTCCGTAGCCGGCCGGACAAGTCCGTTGGGAAATCTGCTCGCCTAACGAGCAGGCGTCCAGCGGAAACTGTTCGGCTTGGGGCCGGAAGACCCGAGGGGTACGTCCAGGGTAGGAGTGAAATCCCGTAATCCTGGACGGACCACCGATGGCGAAAGCACCTCGGGAGGACGGACCCGACGGTGAGGGACGAAAGCTGGGGT |
| BH-asv316 | ACGAGCATTATCCGGATTTATTGGGCGTAAAGGAAGCGTAGGTGGGGAGGTTGATCCATTGTTAAAGGCATTTGCTTAACAAATGTGTGCGATGGAGATCGCCTTCCTAGAGTTAATCAGGGGGTACTGGAATTCAATGTGTAGCGGTGGAATGCGTAGATATATTGAGGAACACCAGAGGCTAAGGCGAGTACCTAGGATATAACTGACACTGAGGCTTGAAAGCGTGGGG |
| BH-asv317 | GCAAACGTTGTTCGGAATTACTGGGCGTAAAGCGCGTGTAGGCTGCTTCGCAAGTCGGATGTGAAAGCCCTGGGCTCAACCCAGGAAGTGCATTCGAAACTGCAGAGCTGGAGTCCTGGAGAGGAAGGCGGAATTCTCGGTGTAGAGGTGAAATTCGTAGATATCGAGAGGAACACCGGTGGCGAAGGCGGCCTTCTGGACAGTGACTGACGCTGAGACGCGAAAGCGTGGGG |
| BH-asv321 | GCAAGCGTTATCCGGAATCACTGGGTGTAAAGGGTGCGCAGGCGGGCATGTAGGTCTGATGTGAAAGTCCTGGGCTTAACCCAGGAATTGCATCGGAAACCCCATGTCTTGAGGGCCAGAGAGGCAGGCGCAATTCGCGGTGTAGCGGTGAAATGCGTAGATATCGCGAGGAACACCAGTGGTGTAGACGGCCTGCTGGATGGACCCTGACGCTGAGGCACGAAAGCGTGGGG |
| BH-asv322 | GCAAGCATTATTCGGATTTATTGGGCGTAAAGCAAGCGTAGGCGGATGTGTAAGTTCTGTGTTAAAAGTAGCTACTTAATAGTTGTTTGCACCGAATACTACATGTCTAGAATGTGGTAGGAAGTTTTGGAATTAAACATGGAGCGGTGGAATGTGTAGATATGTTTAAGAACACCGGAGGCGAAGGCGAAAACTTAGGCCATAATTGACGCTTAGGCTTGAAAGTGTGGGG |
| BH-asv325 | GCAAGCGTTGTCCGGAATTATTGGGCGTAAAGCGCGCGCAGGCGGCTTACTAAGTCTGGTGTGAAAGCCCACGGCTCAACCGTGGATGGCCATTGGAAACTGGTAGGCTTGAGTGCAGGAGAGGAGAGCGGAATTCCCGGTGTAGCGGTGAAATGCGTAGATATCGGGAGGAACACCCGTGGCGAAGGCGGCCAGCTGGGCCGGCCCTGACGCTGAGGCACGAAGGCGTGGGG |
| BH-asv332 | GCAAGCGTTGTCCGGAATTATTGGGCGTAAAGCGCGCGCAGGCGGCCTTTTAAGTCTGATGTGAAAGCCCACGGCTCAACCGTGGAGGGTCATTGGAAACTGGAAGGCTTGAGTGCAGAAGAGAAGAGCGGAATTCCACGTGTAGCGGTGAAATGCGTAGAGATGTGGAGGAACACCAGTGGCGAAGGCGGCTCTTTGGTCTGTAACTGACGCTGAGGCGCGAAAGCGTGGGG |
| BH-asv333 | GCAAGCGTTGTCCGGAATTATTGGGCGTAAAGCGCGCGCAGGCGGTCCTTTAAGTCTGATGTGAAAGCCCACGGCTCAACCGTGGAGGGTCATTGGAAACTGGGGACTTGAGTGCAGGAGAGAAGAGTGGAATTCCACGTGTAGCGGTGAAATGCGTAGAGATGTGGAGGAACACCAGTGGCGAAGGCGACTCTTTGGCCTGTAACTGACGCTGAGGCGCGAAAGCGTGGGG |
| BH-asv334 | CCGAGCGTTGTCCGGAGTTATTGGGCGTAAAGAGCTCGTAGGCGGTTCGTCGCGTCTGCCGTGAAATCTTCAGGCTCAACCTGAGGCGTGCGGTGGGTACGGGCGAGCTAGAGTGCGGTAGGGGAGAATGGAACTCCTGGTGTAGCTGTGGAATGCGCAGATATCAGGAAGAACACCGATGGCGAAGGCAGTTCTCTGGGCCGTAACTGACGCTGAGGAGCGAAAGCGTGGGG |
| BH-asv335 | CTGAGTGGTGGCCACGTTTATTGGGCTTAAAGTGTCCGTAGCCGGCTCTGTGAGTCTGCTGTGAAATCTGGCAGCTTAACTGTCAGGCGTGCAGCAGATACCATAGGGCTTGGGACCGGAAGGCGTTGAGGCTATTCCTGGGGTAGGGGTGAAATCCTGAAATCCCAGGAGGAGCACCAGTGGCGAAGGCTCTCAACGATGACGGGTCCGACGGTGAGGGACGAAAGCCAGGGG |
| BH-asv336 | GCGAGCGTTGTCCGGAATTATTGGGCGTAAAGAGCTTGTAGGCGGTTTGTCGCGTCTGCTGTGAAAGGCCGGGGCTTAACTCCGTGTATTGCAGTGGGTACGGGCAGACTAGAGTGCAGTAGGGGAGACTGGAATTCCTGGTGTAGCGGTGGAATGCGCAGATATCAGGAGGAACACCGATGGCGAAGGCAGGTCTCTGGGCTGTAACTGACGCTGAGAAGCGAAAGCATGGGG |
| BH-asv337 | GCGAGCGTTGTCCGGAATCACTGGGCGTAAAGGGCGCGTAGGCGGCCTGGTAAGTAGGGGGTGAAATCCTGCGGCTTAACCGCAGGGCTTCCTTCTAAACTGTCAGGCTCGAGCACAGTAGAGGCAGGTGGAATTCCCGGTGTAGCGGTGGAATGCGTAGAGATCGGGAAGAACATCAGTGGCGAAGGCGGCCTGCTGGGCTGTTGCTGACGCTGAGGCGCGACAGCGTGGGG |
| BH-asv340 | GCGAGCGTTGTCCGGAATTATTGGGCGTAAAGAGCGTGTAGGCGGTTTGGTAAGTCTGCCGTGAAAACCCGGGGCTCAACCCCGGTCGTGCGGTGGATACTGCCAGGCTAGAGGATGGTAGAGGCGAGTGGAATTCCCGGTGTAGCGGTGAAATGCGCAGATATCGGGAGGAACACCAGTAGCGAAGGCGGCTCGCTGGGCCATTCCTGACGCTGAGACGCGAAAGCTAGGGG |
| BH-asv343 | CCGAGTGGTGGCCACTGTTATTGGGCTTAAAGAGTCCGTAGCCGGCCCGGTAAGTTCTCTGTGAAATCCGGCCGCTCAACGGTCGGGCGCGCAGGGAATACTGCCGGGCTTGGGGCCGGACGGCGTCGGGGCTATTCCCGGGGTAGGAGTGAAATCCTGAAATCCCGGGAGGAGCACCTGTGGCGAAAGCGCCCGGCGAGGACGGACCCGACGGTCAGGGACGAAAGCCAGGGG |
| BH-asv344 | GCAAGCGTTATCCGGAATTATTGGGCGTAAAGAGCTCGTAGGCGGTTTGTCGCGTCTGCCGTGAAAGTCCGGGGCTCAACTCTGGATCTGCGGTGGGTACGGGCAGGCTAGAGTGATGTAGGGGAGACTGGAATTCCTGGTGTAGCGGTGAAATGCGCAGATATCAGGAGGAACACCGATGGCGAAGGCAGGTCTCTGGGCATTAACTGACGCTGAGGAGCGAAAGCATGGGG |
| BH-asv345 | GCTAGCGTTGTTCGGAATTACTGGGCGTAAAGCGCACGTAGGCGGGTCGTTAAGTCGGGGGTGAAATCCCAGGGCTCAACCCTGGAACTGCCTTCGATACTGGCGACCTAGAGTGTGGGAGAGGTTGGTGGAATGCCGAGTGTAGAGGTGAAATTCGTAGATATTAGGAAGAACACCGGTGGCGAAGGCGGCCCACTGGATCGGTACTGACGCTGAGGTGCGAAAGCGTGGGG |
| BH-asv348 | GCAAGCGTTGTCCGGATTTATTGGGCGTAAAGAGCGCGTAGGCGGCCAGGTAAGTCCGACGTGAAATCTGGAGGCTCAACCTCCAGCTGTCGTTGGAAACTACCTGGCTAGAGTCCGGAAGAGGAGAGTGGAATTCCTGGTGTAGCGGTGAAATGCGCAGATATCAGGAAGAACACCAATGGCGAAAGCAGCTCTCTGGGACGGTACTGACGCTGAGGCGCGAAAGCGTGGGG |
| BH-asv350 | GCGAGCGTTGTCCGGAATTATTGGGCGTAAAGGGCTCGTAGGCGGCTTGTCGCGTCTGCTGTGAAAACGCGGGGCTCAACTCCGCGCGTGCAGTGGGTACGGGCAGGCTGGAGTGCGGCAGGGGTGGCTGGAATTCCTGGTGTAGCGGTGGAATGCGCAGATATCAGGAAGAACACCGATGGCGAAGGCAGGCCACTGGGCCGTCACTGACGCTGAGGAGCGAAAGCGTGGGG |
| BH-asv355 | GCAAGCGTTAATCGGAATTACTGGGCGTAAAGCGTGCGCAGGCGGTGATGTAAGACAGGTGTGAAATCCCCGGGCTTAACCTGGGAACTGCATTTGTGACTGCATCGCTGGAGTGCGGCAGAGGGGGATGGAATTCCGCGTGTAGCAGTGAAATGCGTAGATATGCGGAGGAACACCGATGGCGAAGGCAATCCCCTGGGCCTGCACTGACGCTCATGCACGAAAGCGTGGGG |
| BH-asv356 | GCTAGCGTTGTTCGGAATTACTGGGCGTAAAGGGCGCGCAGGCGGACTGGTAAGTCTGGGGTGAAATCTCACGGCTCTACTGTGAGGGGTCCCCGGATACTGCCGGTCTTGAGGGGCTCAGAGGAGAGTGGAATTCCCGGTGTAGTGGTGGAATGCGTAGATATCGGGAGGAACACCAGTGGCGAAAGCGACTCTCTGGGGGCAACCTGACGCTGAGGCGCGAAAGCGTGGGG |
| BH-asv359 | GCGAGCGTTGTCCGGATTCATTGGGCGTAAAGAGCTCGTAGGCGGCTTGCTAAGTCGGGTGTGAAAACTCCGGGCTCAACCCGGAGACGCCACCCGATACTGGCATGGCTAGAGTCCGGTAGGGGAGCGTGGAATTCCTGGTGTAGCGGTGAAATGCGCAGATATCAGGAGGAACACCAGCGGCGAAGGCGGCGCTCTGGGCCGGAACTGACGCTGAGGAGCGAAAGCGTGGGT |
| BH-asv361 | TGCGAGCGTTGTCCGGAATTATTGGGCGTAAAGGGCTCGTAGGCGGTTTGTCACGTCGGGGGTGAAAGCCTGTGGCTCAACCACAGAATTGCCTTCGATACTGCATGGCTTGAGACCGGAAGAGGTTAGTGGAACTGCGAGTGTAGAGGTGAAATTCGTAGATATTCGCAAGAACACCAGTGGCGAAGGCGGCTGACTGGTCCGGTTCTGACGCTGAGGCGCGAAAGCGTGGGG |
| BH-asv362 | GCAAGCGTTGCTCGGAATCACTGGGCGTATAGGGTCCGCAGGCGGCCCTGCAACTCAGGCGTGAAAACCTGTGGCTTAACCACAGGCGTGCGTCCGAAAGTGTAGGGCTCGAGTGCGGGAAGGGAGAGTGGAACTGCGGGTGGAGCGGTGAAATGCGTAGATATCCGCAGGAACACCGGAGGCGAAAGCGACTCTCTAGCCCGTCACTGACGCTCATGGACAAAAGCCAGGGT |
| BH-asv368 | GCGAGCGTTGTCCGGATTTATTGGGCGTAAAGAGCTCGTAGGCGGCTTGTCGCGTCGACTGTGAAAACCCGCGGCTCAACCGCGGGCCTGCAGCCGATACGGGCAGGCTAGAGTTCGGTAGGGGAGACTGGAATTCCTGGTGTAGCGGTGAAATGCGCAGATATCAGGAGGAACACCGGTGGCGAAGGCGGGTCTCTGGGCCGATACTGACGCTGAGGAGCGAAAGCGTGGGG |
| BH-asv370 | GCGAGCGTTGTCCGGAATTATTGGGCGTAAAGGGCTTGTAGGCGGTTGGTCGCGTCTGCCGTGAAATCCTCTGGCTTAACTGGGGGCGTGCGGTGGGTACGGGCTGACTTGAGTGCGGTAGGGGAGACTGGAACTCCTGGTGTAGCGGTGGAATGCGCAGATATCAGGAAGAACACCGGTGGCGAAGGCGGGTCTCTGGGCCGTTACTGACGCTGAGGAGCGAAAGCGTGGGG |
| BH-asv371 | GCAGGCGTTGTTCGGAATTACTGGGCGTAAAGCGCGCGTAGGCGGCTTTTCAAGTCGAGTGTGAAAGCTCCAGGCTTAACCTGGGAATTGCGCTCGATACTGTTGAGCTAGAGTGTGGAAGGGGCTACTGGAATTCCGTGTGTAGCGGTGAAATGCGTAGATATACGGAGGAACACCAGCGGCGAAGGCGGGTAGCTGGGCCAACACTGACGCTGATGTGAGAAAGCTAGGGG |
| BH-asv375 | GCTAGCGTTGTTCGGAATTACTGGGCGTAAAGCGCGCGTAGGCGGACTGAAAAGTTGGGGGTGAAATCCCGGGGCTCAACCTCGGAACTGCCTTCAAAACTATCAGTCTGGAGTTCGAGAGAGGTGAGTGGAATTCCGAGTGTAGAGGTGAAATTCGTAGATATTCGGAGGAACACCAGTGGCGAAGGCGGCTCACTGGCTCGATACTGACGCTGAGGTGCGAAAGCGTGGGG |
| BH-asv376 | GCGAGCGTTAATCGGAATTACTGGGCGTAAAGCGTGCGTAGACGGTAGCGTAAGTCGGGTGTGAAAGCCCCGGGCCCAACCTGGGAATTGCATTCGAGACTGTGCTGCTAGGGTGCGGAAGAGGGAAGCGGAATTTCCGGTGTAGCGGTGGAATGCGTAGATATCGGAAGGAACATCAGTGGCGAAAGCGGCTTCCTGGTCCAGCACCGACGTTCAGGCACGAAAGCGTGGGG |
| BH-asv377 | CCAAGCGTTATCCGGAATTACTGGGCGTAAAGGGTGCGCAGGCGGCATGCAAGTCTGACGTGAAAGCCCCGAGCTCAACTCGGGAAGTGCGCCGGAAACTGCATGTCTTGAGGACCAGAGAGGCAGAGGGAATTCCTGGTGTAGCGGTGAAATGCGTAGATATCAGGAGGAACACCTGTGGCGAAAGCGCTCTGCTGGATGGTTCCTGACGCTGAGGCACGAAAGCGTGGGT |
| BH-asv378 | GCAAGCGTTAATCGGATTTACTGGGCGTAAAGCGTGTGTAGGTGGTCGGGAAAGTCAATTGTGAAAGCCCAGGGCTTAACTCTGGAACTGCTTTTGATACTGTCTGACTAGAATTCGGTAGAGGGAGGCGGAACTCCAGGTGTAGCGGTGAAATGCGTAGATATCTGGAAGAACACCAATGGCGAAGGCAACCTCCTGGGCCTGAATTGACACTGAGACACGAAAGCGTGGGG |
| BH-asv379 | GCGAGCGTTGTCCGGATTTATTGGGCGTAAAGCGCGTGCAGGCGGTTCGTTAAGTCCGTTGTGAAAGCTCCTGGCTCAACTGGGAGAGGTCTTCGGATACTGGCGAACTCGAGGGAGGCAGAGGAAGGTGGAATTCCGGGTGTAGTGGTGAAATGCGTAGAGATCCGGAGGAACACCAGTGGCGAAGGCGGCCTTCTGGTCTGTAACTGACGCTGAGGCGCGAAAGCGTGGGG |
| BH-asv380 | GCAAGCGTTGTCCGGAATCATTGGGCGTAAAGCGCGCGTAGGTGGCCATGTAAGTCCGCTGTGAAAGTCAAAGGCTCAACCTTTGAGAGCCGGTGGATACTGCATGGCTAGAGTACGGAAGAGGCGAGTGGAATTCCTGGTGTAGCGGTGGAATGCGCAGATATCAGGAGGAACACCAATAGCGAAGGCAGCTCGCTGGGACGTTACTGACACTGAGGCGCGAAAGCGTGGGG |

| | |
|---|---|
| BH-asv381 | GCAAGCGTTAATCGGAATCACTGGGCGTAAAGCGCTCGTAGGTGGCTTTGTAAGTCAGGTGTGAAAGCCCACGGCTCAACCGTGGAATTGCACTTGATACTGCATTGCTTGAGTTCGGGAGAGGGTAGTGGAATT<br>CCAGGTGTAGGAGTGAAATCCGTAGAGATCTGGAGGAACACCAGTGGCGAAGGCGACTACCTGGACCGATACTGACACTGAGGAGCGAAAGCGTGGGG |
| BH-asv382 | CCGAGTGGTGACCACGTTTACTGGGCTTAAAGCGCCCGTAGCCGGCTTAGTAAGTACTGTGTGAAATCTGGCGGCTTAACCGTCAGGCGGGCACAGTATACTACTAGGCTAGGGACCGGACGGCGTTAGGGGTA<br>TTCCTGGGGTAGGGGTGAAATCCTAAAATCCCAGGAGGACCACCGGTGGCGAAAGCGCCTAACGAGTACGGATCCGACGGTGAGGGACGAAAGCCAGGGG |
| BH-asv383 | GCGAGCGTTGTCCGGAATTACTGGGCGTAAAGGGCTCGTAGGTGGTTTGTCGCGTCGTCTGTGAAATTCCGGGGCTTAACTCCGGGCGTGCAGGCGATACGGGCATAACTTGAGTACTGTAGGGGTAACTGGAA<br>TTCCTGGTGTAGCGGTGAAATGCGCAGATATCAGGAGGAACACCGATGGCGAAGGCAGGTTACTGGGCAGTTACTGACGCTGAGGAGCGAAAGCATGGGT |
| BH-asv384 | GCAAGCGTTGCCCGGAATCACTGGGCGTAAAGGGTCCGCAGGCGGTCTGGCAACTCAGGCGTGAAATCCCGCGGCTTAACCGTGGACCTGCGTCTGAAAGTGTCAGGCTCGAGTGCGGGAAGGGGGAGTGGA<br>ACTCCGGGTGGAGCGGTGAAATGCGCAGATATCCGGAGGAACACCGGAGGCGAAAGCGACTCCCTAGCCCGCTACTGACGCTCAGGGACAAAAGCCAGGGT |
| BH-asv391 | GCAAGCGTTGTTCGGATTTACTGGGCGTAAAGGGCGCGTAGGCGGATATTCAAGTCAGAGGTGAAAGCCCTGGGCTCAACCCGGGAACTGCCTTTGAAACTGGATGTCTTGAGTATGGGAGAGGGTGGCGGAA<br>TACCCAGTGTAGAGGTGAAATTCGTAGATATTGGGTAGAACACCAGTGGCGAAGGCGGCCACCTGGCCCATAACTGACGCTGAGGCGCGAAAGCGTGGGG |
| BH-asv395 | GCAAGCGTTGTCCGGAATTACTGGGCGTAAAGGGCGTGTAGGCGGTGTCATACGTAGCGGTTTTAAGCCGCCGGCTCACCCGGCGGAGGGTCGCTAAACGGTGGCGCTGGAGGGCAGGAGAGGTGCGTGGAA<br>TTCCTGGTGGAGCGGTGAAATGCGTAGAGATCAGGAAGAACACCAGTGGCGAAGGCGGCGCACTGGCCTGGCCCTGACGCTGAGGCGCGACAGCGTGGGG |
| BH-asv399 | CCAAGTGGTGGCCGATGTTACTGGGCTTAAAGTGTTCGTAGCCGGCCGTGTAAGTCTGCTGTGAAATCCACGCGCTTAACGTGTGGGCGTGCAGCGGATACTGCATGGCTTGGGACCGGACGGCGTCGAGGCT<br>ATTCCCGGGGTAGGAGTGAAATCCTAAAATCCCGGGAGGAGCACCTGTGGCGAAAGCGCTCGGCGAGTACGGATCCGACGGTGAGGAACGAAAGCCAGGGG |
| BH-asv404 | ACGAGCATTATCCGGATTTATTGGGCGTAAAGGAAGCGTAGGCGGGGAGGTTGATCCATTGTTAAAAGCATTTGCTTAACAAATGTGTGCATTGGAGATCGCCTCCCTAGAGTTAATCAGGGGGTACTGGAATTCA<br>ATGTATAGCGGTGGAATGCGTAGATATATTGAGGAACACCAGAGGCTAAGGCGAGTACCTAGGATATAACTGACGCTGAGGCTTGAAAGCGTGGGG |
| BH-asv406 | GCAAGCGTTTTTCGGAATTACTGGGCGTAAAGGGCGCGTAGGCGGCCTGCTTAGTCGGACGTGAAATCCCTCGGCTCAACCGGGGAACTGCGTCCGATACTGGCGGGCTCGAATCCGGGAGAGGGATGTGGAA<br>TTCCAGGTGTAGCGGTGAAATGCGTAGATATCTGGAGGAACACCAGTGGCGAAGGCGGCATCCTGGACCGGCATTGACGCTGATGAGCGAAAGCTAGGGG |
| BH-asv407 | GCGAGCGTTAATCGGAATTACTGGGCGTAAAGCGTGCGCAGGCGGTTGTGCAAGACAGATGTGAAATCCCCGGGCTCAACCTGGGAACTGCATTTGTGACTGCACGGCTGGAGTGCGGCAGAGGGGGATGGAA<br>TTCCGCGTGTAGCAGTGAAATGCGTAGATATGCGGAGGAACACCGATGGCGAAGGCAATCCCCTGGGCCTGCACTGACGCTCATGCACGAAAGCGTGGGG |
| BH-asv408 | GCAAGCGTTGTTCGGAATTATTGGGCGTAAAGCGCGTGTAGGCGGCTTTGCAAGTCGGGTGTGAAAGCCCTCGGCTCAACCGAGGAAGTGCGCCCGAAACTGCAGAGCTCGAGTGCCAGAGAGGGTGGCGGAA<br>TTCCCCAAGTAGAGGTGAAATTCGTAGATATGGGGAGGAACACCAGTGGCGAAGGCGGCCACCTGGATGGTGACTGACGCTGAGACGCGAAAGCGTGGGG |
| BH-asv413 | GCAAGCGTTGTCCGGAATCACTGGGCGTAAAGGGTCCGCAGGCGGATCAGCAACTCAGGCGTGAAACCCTGTGGCTTAACCACGGAGGTGCGTCTGAAAGTGTTGATCTCGAGGACAGGAAGGGAGAGTGGAA<br>CTCCGGGTGGAGCGGTGAAATGCGCAGATATCCGGAGGAACACCGGAGGCGAAAGCGACTCTCTAGCCTGTTCCTGACGCTGAGGGACCAAAGCCAGGGG |
| BH-asv414 | CCAAGCGTTATCCGGAATTACTGGGCGTAAAGGGTGCGCAGGCGGGCACATAAGTCCGACGTGAAAGCCCCGAGCTTAACTCGGGAATGGCGCCGGAAACTATGTGTCTTGAGGACCAGAGAGGCAGGGAGAA<br>TTCCTGGTGTAGCGGTGAAATGCGTAGATATCAGGAGGAATACCTGTGGCGAAAGCGCCCTGCTGGATGGTTTCTGACGCTGAGGCACGAAAGCGTGGGG |
| BH-asv417 | GCAAGCGTTGTCCGGAATCACTGGGCGTAAAGGGTCCGCAGGCGGATCAGCAACTCAGGCGTGAAATCCCGCGGCTTAACCGCGGAGGTGCGCCTGAAAGTGTTGATCTCGAGGACAGGAAGGGAGAGTGGA<br>ACTCCGGGTGGAGCGGTGAAATGCGCAGATATCCGGAGGAACACCGGAGGCGAAAGCGACTCTCTAGCCTGTTCCTGACGCTGAGGGACCAAAGCCAGGGT |
| BH-asv424 | GCGAGCGTTGTTCGGAATTATTGGGCGTAAAGGGTGCGTAGGCGGTTCAGTAAGTCTTGTGTGAAATCTTCGGGCTCAACTCGAAGTCTGCATGAGAAACTGCTGGGCTTGAGTGTGGGAGAGGTGAGTGGAAT<br>TCCCGGTGTAGCGGTGAAATGCGTAGATATCGGGAGGAACACCTGTGGCGAAAGCGGCTCACTGGACCACAACTGACGCTGATGCACGAAAGCTAGGGG |
| BH-asv426 | GCAAGCGTTGTCCGGAATTATTGGGCGTAAGGCGCGCGCAGGCGGTTCCTTAAGTCTGATGTGAAAGCCCACGGCTCAACCGTGGAGGGTCATTGGAAACTGGGGAACTTGATTGCAGAAGAGGAAAGCGGAA<br>TTCCACGTGTAGCGGTGAAATGCGTAGAGATGTGGAGGAACACCAGTGGCGAAGGCGGCTTTCTGGTCTGTAACTGACGCTGAGGCGCGAAAGCGTGGGG |
| BH-asv436 | GCAAGCGTTGTCCGGAATTATTGGGCGTAAAGCGCGCGCAGGCGGTTTCTTAAGTCTGATGTGAAAGCCCACGGCTCAACCGTGGAGGGTCATTGGAAACTGGGAAACTTGAGTGCAGAAGAGGAGAGTGGAAT<br>TCCACGTGTAGCGGTGAAATGCGTAGAGATGTGGAGGAACACCAGTGGCGAAGGCGACTCTCTGGTCTGTAACTGACGCTGAGGCGCGAAAGCGTGGGG |
| BH-asv439 | CCAAGCGTTATCCGGAATCACTGGGCGTAAAGGGTGCGCAGGCGGCCTGCAAGTCCGACGTGAAAGCCCCGGGCTCAACCCGGGAAGAGCGCCGGAAACTGCAGGTCTTGAGGACCAGAGAGGCAGGGAGA<br>ATTCCCGGTGTAGCGGTGAAATGCGTAGATATCGGGAGGAATACCAGTGGCGAAAGCGCCCTGCTGGATGGTTCCTGACGCTGAGGCACGAAAGCGTGGGT |
| BH-asv440 | GCAAGCGTTAATCGGAATTACTGGGCGTAAAGTGCGCGTAGGCGGCTGGGTAAGTCGGATGTGAAAGCCCTGGGCTCAACCTAGGAATTGCATTCGATACTGCTCGGCTAGAGTGCAGTAGAGGGGAGTGGAAT<br>TCCCGGTGTAGCGGTGAAATGCGTAGATATCGGGAGGAATACCAGTGGCGAAGGCGACTCCCTGGACTGACACTGACGCTGACAGCGCGAAAGCGTGGGG |
| BH-asv443 | GCGAGCGTTGTCCGGAATTACTGGGCGTAAAGAGCTCGTAGGTGGTTTGTTGCGTCGTCTGTGAAATTCCGGGGCTTAACTTCGGGGTGGCAGGCGATACGGGCATAACTAGAGTGCTGTAGGGGAGACTGGA<br>ATTCCTGGTGTAGCGGTGGAATGCGCAGATATCAGGAGGAACACCGATGGCGAAGGCAGGTCTCTGGGCAGTAACTGACGCTGAGGAGCGAAAGCATGGGG |
| BH-asv444 | GCAAGCGTTGTCCGGAATTATTGGGCGTAAAGAGCTCGTAGGCGGTTCGTCGCGTCGGCTGTGAAAACCTGAGGCTCAACCTCAGGCCTGCAGTCGATACGGGCGGACTTGAGTACTGCAGGGGAGACTGGAA<br>TTCCTGGTGTAGCGGTGAAATGCGCAGATATCAGGAGGAACACCGGTGGCGAAGGCGGGTCTCTGGGCAGTAACTGACGCTGAGGAGCGAAAGCGTGGGG |
| BH-asv449 | GCAAGCGTTGTCCGGAATCACTGGGCGTAAAGGGTCCGCAGGCGGACCGGCAACTCAGGCGTGAAATCCCACGGCTCAACCGTGGAACTGCGTCTGAAAGTGTTGGTCTCGAGGACAGTAAGGGAGAGTGGAA<br>CTCCGGGTGGAGCGGTGAAATGCGCAGATATCCGGAGGAACACCGGAGGCGAAAGCGACTCTCTAGCCTGTTCCTGACGCTGAGGGACCAAAGCCAGGGT |
| BH-asv452 | CCGAGTGATGGCCGCTATTATTGGGCCTAAAGCGTCCGTAGCCGGCCGGGCAAGTCCGTCGGGAAATCCACCCGCTTAACGGGTGGGCGACCGGCGGAAACTGCTCGGCTTGGGACCGGAGGACCCGAGGGG<br>TACGTCCGGGGTAGGAGTGAAATCCCGTAATCCTGGACGGACCACCGATGGGGAAACCACTTCAAGAGAACGGATCCGACAGTGAGGGACGAAAGCTAGGGT |
| BH-asv457 | CCGAGTGATGGCCGCTATTATTGGGCCTAAAGCGTCCGTAGCCGGCCGGGCAAGTCCGTCGGGAAATCCACCCGCTTAACGGGTGGGCGTCCGGCGGAAACTGTTCGGCTTGGGACCGGAAGACCCGAGGGG<br>TACGTCCGGGGTAGGAGTGAAATCCCGTAATCTCGGACGGACCGCCGGTGGCGAAAGCGCCTCGGGAAGACGGATCCGACGGTGAGGGACGAAAGCTAGGGT |
| BH-asv458 | CCGAGTTGTCGGGACGATTATTGGGCCTAAAGCATCCGTAGCCTGTTCTGCAAGTCCTCCGTTAAATCCACCTGCTCAACGGATGGGCTGCGGAGGAAACCGCAGAGCTAGGAGGCGGGAGAGGCAAACGGTA<br>CTCAGTGGGTAGGGGTAAAATCCATTGATCTACTGAAGACCACTAGTGGCGAAGGCGGTTTGCCAGAACGCGCTCGACGGTGAGGGATGAAAGCTGGGGG |
| BH-asv464 | GCTAGCGTTGTTCGGAATTACTGGGCGTAAAGCGCACGTAGGCGGACATTTAAGTCAGGGGTGAAATCCCGGGGCTCAACCCCGGAACTGCCTTTGATACTGGGTGTCTTGAGTGTGGTAGAGGTGAGTGGAAT<br>TGCGAGTGTAGAGGTGAAATTCGTAGATATTCGCAGGAACACCAGTGGCGAAGGCGGCTCACTGGACCACAACTGACGCTGAGGTGCGAAAGCGTGGGG |
| BH-asv465 | TCAAGCGTTGTTCGGAATTACTGGGCGTAAAGGGCGCGTAGGCGGCCTGTTAAGTCGGACGTGAAATCCCTCGGCTCAACCGAGGAACTGCGTCCGATACTGGTGGGCTTGAGTTCGGGAGAGGGATGTGGAA<br>TTCCAGGTGTAGCGGTGAAATGCGTAGATATCTGGAGGAACACCGGTGGCGAAGGCGGCATCCTGGACCGACACTGACGCTGAGGCGCGAAAGCCAGGGG |
| BH-asv466 | CCAAGCGTTATCCGGATTTACTGGGCGTAAAGCATTCGTAGGCGGCTTGGCACGTCTCCGGTTAAAGCTCTTGGCTCAACCAAGAGAATGCCGGGGATACGGCCTGGCTAGAGGATGTCAGAGGCCGGCGGAAC<br>GTAGGGAGTAGGGATGAAATCCGTCGATACCCTGCGGAACACCGAAGGCGAAAGCAGCCGGCTGGGGCACTCCTGACGCTCAGGGATGTAAGCCAGGGT |
| BH-asv472 | GCAAGCGTTGTCCGGAATTATTGGGCGTAAAGCGCGCGCAGGTGGTTCCTTAAGTCTGATGTGAAAGCCCACGGCTCAACCGTGGAGGGTCATTGGAAACTGGGGAACTTGAGTGCAGAAGAGGAAAGTGGAAT<br>TCCAAGTGTAGCGGTGAAATGCGTAGAGATTTGGAGGAACACCAGTGGCGAAGTCGACTTTCTGGTCTGTAACTGACACTGAGGCGCGAAAGCGTGGGG |
| BH-asv477 | CCAAGCGTTATCCGGAATTACTGGGCGTAAAGGGTGCGCAGGCGGGCACATAAGTCCGACGTGAAAGCCCCGGGCTTAACCCGGGAATGGCGCCGGAAACTATGTGTCTTGAGGACCAGAGAGGCAGGCAGAA<br>TTCCTGGTGTAGCGGTGAAATGCGTAGATATCAGGAGGAATACCTGTGGCGAAAGCGGCCTGCTGGATGGTTTCTGACGCTGAGGCACGAAAGCGTGGGG |
| BH-asv492 | GCAAGCGTTAATCGGAACTACTGGGCGTAAAGCGTGCGCAGGCGGTTTTGTAAGTCTGTCGTGAAAGCCCCGGGCTCAACCTGGGAATTGCGATGGAGACTGCAAGGCTTGAATCTGGCAGAGGGGGGTAGAA<br>TTCCACGTGTAGCAGTGAAATGCGTAGAGATGTGGAGGAACACCGATGGCGAAGGCAGCCCCCTGGGTCAAGATTGACGCTCATGCACGAAAGCGTGGGG |
| BH-asv493 | ACGAGTGATGGCCGATCTTATTGGGCCTAAAGCGTCCGTAGCCGGCCGGACAAGTCCGTTGGGAAATCGACGCGCCTAACGCGTCGGCGTCCAGCGGAAACTGTCCGGCTTGGGGCCGGAAGACTCGAGGGG<br>TACGTCCGGGGTAGGAGTGAAATCCCGTAATCCTGGACGGACCACCGGTGGCGAAAGCGCCTCGAGAGGACGGACCCGACGGTCAGGGACGAAAGCTAGGGT |
| BH-asv494 | ACGAGTGATGGCCGCTCTTATTGGGCCTAAAGCGTTCGTAGCCGGCCACGCAAGTTCGTCGGGAAATCTGCCCGCTTAACGGGCAGGCGTCCGGCGAAAACTGCGTGGCTTGGGGCCGGAAGATCCGAGGGGT<br>ACGTCCGGGGTAGGAGTGAAATCCCGTAATCCTGGACGGACCACCGGTGGCGAAAGCGCCTCGGAAAGACGGACCCGACGGTGAGGGACGAAAGCTAGGGT |
| BH-asv497 | GCAAGCGTTGTCCGGAATGATTGGGCGTAAAGGGCGCGTAGGCGGCTTTATAAGTCTGGAGTGAAAGTCCTCTTTTCAAGGGAGGAATTGCTTTGGAAACTATAGAGCTTGAGTGCAGGAGAGGTTATCGGAATT<br>CCCGGTGTAGCGGTGAAATGCGTAGATATCGGGAGGAACATCGGTGGCGAAGGCGGGTAACTGGACTGCAACTGACGCTGAGGCGCGAAAGTGTGGGG |
| BH-asv502 | CTGAGTGGTGGCCACGTTTATTGGGCTTAAAGTGTCCGTAGCCGGCTCTGTGAGTCTGCTGTGAAATCTGGCAGCTTAACTGTCAGGCGTGCAGCAGATACCATAGGGCTTGGGACCGGAAGGCGTCGAGGCTA<br>TTCCTGGGGTAGGGGTGAAATCCTGAAATCCCAGGAGGAGCACCAGTGGCGAAGGCTCTCGACGATGACGGGTCCGACGGTGAGGGACGAAAGCCAGGGG |
| BH-asv509 | GCGAGCGTTGTCCGGAATTATTGGGCGTAAAGAGCTTGTAGGCGGTTGGTCGCGTCTGCTGTGAAAGGCTGGGGCTTAACCCTGGTTTTGCAGTGGGTACGGGCTAACTAGAGTGCAGTAGGGGAGACTGGAA<br>TTCCTGGTGTAGCGGTGGAATGCGCAGATATCAGGAGGAACACCGATGGCGAAGGCAGGTCTCTGGGCTGTAACTGACGCTGAGAAGCGAAAGCATGGGG |
| BH-asv510 | GCAAGCGTTGTCCGGAATTATTGGGCGTAAAGAGCTCGTAGGCGGCTTGTCGCGTCGGATGTGAAAGCCCGGGGCTTAACTCCGGGTCTGCATTCGATACGGGCAGGCTAGAGTGTGGTAGGGGAGATCGGAA<br>TTCCTGGTGTAGCGGTGAAATGCGCAGATATCAGGAGGAACACCGGTGGCGAAGGCGGATCTCTGGGCCATTACTGACGCTGAGGAGCGAAAGCGTGGGG |
| BH-asv511 | GCAAGCGTTGTTCGGAATTACTGGGCGTAAAGGGCGCGTAGGCGGCCTGTTAAGTCGAACGTGAAATCCCTGGGCTCAACCCAGGAACTGCGTCCGATACTGGCAGGCTTGAATCCGGGAGAGGGATGCGGAA<br>TTCCTGGTGTAGCGGTGAAATGCGCAGATATCAGGAGGAACACCGATTGCGAAGGCAGCTCGCTGGGACGTAACTGACGCTGAGACGCGAAAGCGTGTGG |
| BH-asv515 | GCAAGCGTTATCCGGATTCATTGGGTTTAAAGGGTGCGTAGGCGGGGTGGTAAGTCAGTGGTGAAATCCTGCAGCTTAACTGTAGAATTGCCATTGATACTGCCATTCTTGAGTGCGCTTGAAGTGGGCGGAATG<br>TGCCGTGTAGCGGTGAAATGCTTAGATATGGCACAGAACACCAATTGCGAAGGCAGCTCACTAAGGCGATACTGACGCTGAGGCACGAAAGCGTGGGG |
| BH-asv517 | GCAAGCGTTAATCGGAATTACTGGGCGTAAAGCGTGCGTAGGTGGTTTGTTAAGTCAGATGTGAAAGCCCTGGGCTCAACCTGGGAACTGCATTTGATACTGGCAGGCTAGAGTGCGGTAGAGGGTGGTGGAAT<br>TCCCGGTGTAGCGGTGAAATGCGTAGAGATCGGGAGGAACATTCGTGGCGAAGGCGGCCACCTGGACCAGCACTGACACTGAGGCACGAAAGCGTGGGG |
| BH-asv518 | CCAAGCGTTAATCGGAATTACTGGGCGTAAAGCGTGCGCAGGCGGTTGTGCAAGACCGATGTGAAATCCCCGGGCTTAACCTGGGAATTGCATTGGTGACTGCACGGCTAGAGTGTGTCAGAGGGGGGTAGAAT<br>TCCACGTGTAGCAGTGAAATGCGTAGAGATGTGGAGGAATACCGATGGCGAAGGCAGCCCCCTGGGATAACACTGACGCTCATGCACGAAAGCGTGGGG |
| BH-asv519 | GCAAGCGTTAATCGGAATTACTGGGCGTAAAGCGTGCGCAGGCGGTTTTGTAAGTCTGACGTGAAATCCCCGGGCTTAACCTGGGAATTGCGTTGGAGACTGCAAGGCTGGAGTCTGGCAGAGGGGGGTAGAA<br>TTCCACGTGTAGCAGTGAAATGCGTAGAGATGTGGAGGAACACCGATGGCGAAGGCAGCCCCCTGGGTCAAGACTGACGCTCATGCACGAAAGCGTGGGG |
| BH-asv525 | GCGAGCGTTAATCGGAATTACTGGGCGTAAAGCGTGCGCAGGCGGTTATGTAAGACAGATGTGAAATCCCCGGGCTCAACCTGGGAACTGCATTAGTGACTGCATAGCTGGAGTACGGCAGAGGGGGATGGAAT<br>TCCGCGTGTAGCAGTGAAATGCGTAGATATGCGGAGGAACACCGATGGCGAAGGCAATCCCCTGGGCCTGTACTGACGCTCATGCACGAAAGCGTGGGG |
| BH-asv528 | GCAAGCGTTGTCCGGAATCACTGGGCGTAAAGGGTGTGCAGGCGGGGCAGCAAGTCGGAGGTGAAATCCCACGGCTTAACCGTGGAAGTGCCTTCGATACTGTTGCTCTTGAGTCCCGGAGAGGCTGCCGGAA<br>TTCGTGGTGTAGCGGTGAAATGCGTAGATATCACGAGGAACACCAGAGGCGAAAGCGGGCAGCTGGACGGGTACTGACGCTCAGGCACGAAAGCGTGGGG |
| BH-asv531 | CCGAGTGGTAGTCGTTGTTATTGGGCTTAAAGTGTTCGTAGCCTGTCGTGTAAGTCCGCTGTGAAACCCACGCGCTCAACGTGTGGATGCGCAGCGGATACTGCATGACTAGGGACCGGACGGCGTCGAGGCTA<br>TTCCAGGGGTAGGAGTGAAATCCAAAAATCCCTGGAGGAGCACCTGTGGCGAAAGCGCTCGACGAGTACGGATCCGACGGTGAGGAACGAAAGCCGGGGG |

| | |
|---|---|
| BH-asv534 | CCAAGCGTTATCCGGAATTACTGGGCGTAAAGGGTGCGCAGGCGGCATGCAAGTCTGACGTGAAAGCCCCGAGCTCAACTCGGGAAGTGCGCCGGAAACTGCATGTCTTGAGGACCAGAGAGGCAGAGAGAA<br>TTCCTGGTGTAGCGGTGAAATGCGTAGATATCAGGAGGAATACCTGTGGCGAAAGCGCTCTGCTGGATGGTTTCTGACGCTGAGGCACGAAAGCGTGGGT |
| BH-asv537 | GCAAGCGTTATCCGGAATCACTGGGTTTAAAGGGTGCGTAGGCGGTTGCGTAAGTCAGAAGTGAAAGCTCACAGCTTAACTGTGGAATTGCTTTTGATACTGCGCGACTAGAATTAGGTTGAGGTTGGCGGAATG<br>TGACATGTAGCGGTGAAATGCATAGATATGTCATAGAACACCAATTGCGAAGGCAGCTGACTGGACCTATATTGACGCTGAGGCACGAAAGCGTGGGG |
| BH-asv540 | GCAAGCGTTAATCGGAATTACTGGGCGTAAAGCGCGCGTAGGTGGTTCAGTAAGTTGGAAGTGAAATCCCCGGGCTCAACCTGGGAACTGCTTTCAAAACTGCTGAGCTAGAGTACGGTAGAGGGTGGTGGAAT<br>TTCCTGTGTAGCGGTGAAATGCGTAGATATAGGAAGGAACACCAGTGGCGAAGGCGACCACCTGGACTGATACTGACACTGAGGTGCGAAAGCGTGGGG |
| BH-asv542 | CCGAGTGATAGCCAATTTTATTGGGCCTAAAGCGTCCGTAGCCGGCCGAACGGGTCCGTCGGGAAATCCACCCGCTTAACGGGTGGGCGTCCGGCGGAAACCAGTCGGCTTGGGGCCGGGAGACCAGAGGGG<br>TACGTCCGGGGTAGGAGTGAAATCCTGTAATCCTGGACGGACCACCGGTTGCGAAAGCGCCTCTGGAGAACGGACCCGACGGTGAGGGACGAAAGCTAGGGT |
| BH-asv544 | GCAAGCGTTACTCGGAATTACTGGGCGTAAAGCGTGCGTAGGTGGTTCGTTAAGTCCGATGTGAAAGCCCTGGGCTCAACCTGGGAATGGCATTGGATACTGGCGGGCTAGAGTGCGGTAGAGGATAGTGGAAT<br>TCCTGGTGTAGCAGTGAAATGCGTAGAGATCAGGAGGAACATCCGTGGCGAAGGCGACTGTCTGGACCAGCACTGACACTGAGGCACGAAAGCGTGGGG |
| BH-asv545 | GCAAGCGTTATCCGGAATTACTGGGCGTAAAGCGCACGCAGGTGGCTGCTTAAGTCTGACGTGAAAGCTCCTGGCTTAACTGGGAGAGGTCGTTGGAGACTGGGCGGCTTGAGGCGAGGAGAGGGGTGTAGAA<br>TTCCCGGTGTAGTGGTGGAATGCGTAGATATCGGGAGGAATACCAGTGGCGAAAGCGGCACCCTGGCCTTGGCCTGACGCTCAGGTGCGAAAGCGTGGGG |
| BH-asv547 | GCAAGCGTTGCTCGGAATTACTGGGCGTAAAGGGTCCGCAGGCGGCCTGGCAACTCAGGCGTGAAATCCCACCGCTTAACGGTGGACGTGCGTCCGAAAGTGCCAGGCTCGAGTACGGGAAGGGAGAGTGGA<br>ACTGTGGGTGGAGCGGTGAAATGCGTAGATATCCACAGGAACACCGGAGGCGAAAGCGACTCTCTAGCCCGTTACTGACGCTGAGGGACAAAAGCCAGGGT |
| BH-asv548 | CCAAGTGGTAGCCAGTTTTATTGGGCTTAAAGTGTTCGTAGCCTGCCGTGTAAGTCCGCTGTGAAACCCAGGCGCTCAACGTCTGGATGTGCAGCGGATACTGCATGGCTTGGACCGGACGGCGTCGAGGCTA<br>TTCCAGGAGTAGGAGTGAAATCCAAAAATTCCTGGAGGAGCACCTGTGGCGAAAGCGCTCGACGAGTACGGATCCGACGGTGAGGAACGAAAGCCGGGGG |
| BH-asv549 | GCAAGCGTTATCCGGATTCACTGGGTTTAAAGGGTGCGTAGGCGGGCAGGTAAGTCAGTGGTGAAATCCTGGAGCTTAACTCCAGAACTGCCATTGATACTATCTGTCTTGAATATTGTGGAGGTAAGCGGAATA<br>TGTCATGTAGCGGTGAAATGCTTAGATATGACATAGAACACCTATTGCGAAGGCAGCTTACTACGCATATATTGACGCTGAGGCACGAAAGCGTGGGG |
| BH-asv550 | GCAAGCGTTGTTCGGAATTACTGGGCGTAAAGCGCGTGTAGGCGGACTCAGAAGTCGGGTGTGAAAGCCCCGGGCTCAACCCGGGAAGTGCGCTCGAAACCATGAGTCTTGAGGTCCGGAGGGGATCGGGGA<br>ATTCCTGGTGTAGAGGTGAAATTCGTAGATATCAGGAGGAACACCGGTGGCGAAGGCGCCGATCTGGACGGATTCTGACGCTGAGACGCGAAAGCGTGGGT |
| BH-asv555 | GCGAGCGTTATTCGGATTTATTGGGCGTAAAGGGTCCGCAGGAGGTTCTTTAAGTCTGATGTGAAATCCCGGGGCTCAACCCCGGAACTGCATTGGAAACTAAAGAACTAGAGTATCGGAGAGGTAAGTGGAATT<br>TCAGGTGTAGCGGTGGAATGCGTAGATATCTGAAAGAACACCAATAGCGAAGGCAGCTTACTGGACGATTACTGACTCTCAGGGACGAAAGCGTGGGG |
| BH-asv561 | GCGAGCGTTGTTCGGAATCACTGGGCTTAAAGAGTGCGTAGGTGGATTTGCAAGTCAGGAGTGAAATCCCGGGGCTCAACCCCGGAACTGCTTTTGAAACTGCAAGTCTGGAGTCCCGGAGAGGTTAGGAGAAT<br>TCCGAGTGTAGAGGTGAAATTCGCAGATATTCGGAGGAATACCAGTGGCGTAGGCGCCTAACTGGACGGGTACTGACACTGAGGCACGAAAGCGTGGGG |
| BH-asv563 | GCAAGCGTTATCCGGATTCACTGGGTTTAAAGGGTGCGTAGGCGGGTATGTAAGTCAGTGGTGAAATACCGGAGCTTAACTTCGGAACTGCCATTGATACTATATATCTTGAATATTGTGGAGGTAAGCGGAATAT<br>GTCATGTAGCGGTGAAATGCTTAGAGATGACATAGAACACCGATTGCGAAGGCAGCTTACTACGCAAATATTGACGCTGAGGCACGAAAGCGTGGGG |
| BH-asv565 | TCAAGCGCTACCCGGAGTTACTGGGCGTAAAGGGCGTGTAGGGGGCTTTGTAAGTTCATTGTTAAATCTCAGGGCTTAACCCTGAGGCTGCAATGAAAACTGCAAGGCTAGAGGACGTCAGAGGAGAGTGGAAC<br>GCACGGTGTAGAGGTGAAATTCGTTGATATCGTGCGGAACATCAAAAGCGAAGGCATCTCTCTGGGGCGTTCCTGACCCTAAGGCGCTAAAGCTTGGGT |
| BH-asv566 | CCAAGTGGTAGTCGTTATTATTGGGCTTAAAGTGTTCGTAGCCGGCCATGCAAGTCCGCTGTGAAATCCACGCGCTCAACGTGTGGGCGTGCAGCGGATACTGCATGGCTAGGGACCGGACGGCGTCGAGGCTA<br>TTCCAGGGGTAGGGGTGAAATCCTAAAATCCCTGGAGGAGCACCTGTGGCGAAAGCGCTCGACGAGTACGGATCCGACGGTGAGGAACGAAAGCCAGAGG |
| BH-asv569 | GCAAGCGTTATCCGGAATCACTGGGCGTAAAGAGCTCGTAGGCGGCCCTGTAAGTCGAATGTTAAATATCCGGGCTCAACCCGGAAAATGCGTTCGATACTGCTTGGCTAGAGAGTGGGAGAGGAAAGTGGAAT<br>TCCTGGTGTAGCGGTGAAATGCATAGATATCAGGAAGAACACCAGTGGCGAAGGCGGCTTTCTGGACCATTTCTGACGCTGAGGAGCGAAAGCCAGGGG |
| BH-asv572 | GCGAGCGTTGTCCGGAATTATTGGGCGTAAAGGGCTTGTAGGCGGCTGGTCGCGTCTGCCGTGAAATCCTCTGGCTCAGCTGGGGGCGTGCGGTGGGTACGGGCTGGCTTGAGTGCGGTAGGGGAGACTGGA<br>ACTCCTGGTGTAGCGGTGGAATGCGCAGATATCAGGAGGAACACCGGTGGCGAAGGCGGGTCTCTGGGCCGTTACTGACGCTGAGGAGCGAAAGCGTGGGG |
| BH-asv573 | GCAAGCGTTGTCCGGAATTATTGGGCGTAAAGAGCTCGTAGGCGGCTAGTCGCGTCGGCTGTGAAAACCTGGGGCTCAACCCCGGGCCTGCAGACGATACGGGCTAGCTAGAATTCGGTAGGGGAGACTGGAA<br>TTCCTGGTGTAGCGGTGAAATGCGCAGATATCAGGAGGAACACCGGTGGCGAAGGCGGGTCTCTGGGCCGATATTGACGCTGAGGAGCGAAAGCGTGGGG |
| BH-asv574 | GCTAGCGTTGTTCGGAATCACTGGGCGTAAAGGGCGCGTAGGCGGCTTTGTAAGTCGGGGGTGAAAGCCTGTGGCTCAACCACAGAATTGCCTTCGATACTGCATGACTTGAGACCGGAAGAGGTAGTGGAACT<br>GCGAGTGTAGAGGTGAAATTCGTAGATATTCGCAAGAACACCAGTGGCGAAGGCGGCTGACTGGTCCGGTTCTGACGCTGAGGCGCGAAAGCGTGGGG |
| BH-asv575 | GCAAGCGTTATCCGGAATTATTGGGCGTAAAGCGCGCGTAGGCGGTTTTTTAAGTCTGATGTGAAAGCCCACGGCTCAACCGTGGAGGGTCATTGGAAACTGGAAAACTTGAGTGCAGAAGAGGAAAGTGGAATT<br>CCATGTGTAGCGGTGAAATGCGCAGAGATATGGAGGAACACCAGTGGCGAAGGCGACTTTCTGGTCTGCAACTGACGCTGATGTGCGAAAGCGTGGGG |
| BH-asv576 | TCGAGTGGTAGTCACTTTTATTGAGCCTAAAGCGTTCGTAGCCTGATTTGTAAATCTCTAGGAAAATACTTCTGCTTAACTGAAGAAATTTTGGAGAGACTGCGAATCTAGGGACCGGGTGAGGTTGAATGTACTTC<br>TGGGGTAGGGGTAAAATCCTGTAATCCTGGAAGGACGACCGGTGGCGAAGGCGTTCAACTAGAACGGATCCGACGGTGAGGAACGAAGGCTAGGGG |
| BH-asv578 | GCAAGCGTTATCCGGAATGATTGGGCGTAAAGCGTCTGTAGGTGGCTTTTTAAGTCCGCCGTCAAATCCCAGGGCTCAACCCTGGACAGGCGGTGGAAACTACCAAGCTGGAGTACGGTAGGGGCAGAGGGAAT<br>TTCCGGTGGAGCGGTGAAATGCGTAGAGATCGGAAAGAACACCAACGGCGAAAGCACTCTGCTGGGCCGACACTGACACTGAGAGACGAAAGCTAGGGG |
| BH-asv580 | GCAAGCGTTAATCGGAATTACTGGGCGTAAAGCGCGCGTAGGTGGTTTGATAAGCGAGATGTGAAAGCCCCGGGCTTAACCTGGGAACGGCATTTCGAACTGTCAGGCTAGAGTATGGTAGAGGGTAGTGGAAT<br>TTCCTGTGTAGCGGTGAAATGCGTAGATATAGGAAGGAACACCAGTGGCGAAGGCGGCTACCTGGACCAATACTGACACTGAGGTGCGAAAGCGTGGGG |
| BH-asv583 | CCAAGCGTTATCCGGAATTACTGGGCGTAAAGGGTGCGCAGGCGGCACATAAGTCCGACGTGAAAGCCCCGGGCTCAACCCGGGAAAGGCGCCGGAAACTATGTGTCTTGAGGACCAGAGAGGCAAGGAGAA<br>TTCCTGGTGTAGCGGTGAAATGCGTAGATATCAGGAGGAATGCCAGTGGCGAAAGCGCCTTGCTGGATGGTTCCTGACGCTGAGGCACGAAAGCGTGGGT |
| BH-asv588 | GCGAGCGTTATCCGGATTTATTGGGTGTAAAGGGTGCGTAGACGGAAATGCAAGTTAGTTGTGAAATACCTCGGCTTAACTGAGGAACTGCAACTAAAACTATATTTCTTGAGTATCGGAGGGGTTTGTGGAATTC<br>CTAGTGTAGCGGTGAAATGCGTAGATATTAGGAAGAACACCGGTGGCGAAGGCGACAAACTGGACGATAACTGACGTTGAGGCACGAAAGTGTGGGG |
| BH-asv589 | GCTAGCGTTGTCCGGAATCATTGGGCGTAAAGCGCGTGTAGGCGGCCCGGTAAGTCCGCTGTGAAAGTCGGGGGCTCAACCCTCGAAAGCCGGTGGATACTGTCGGGCTAGAGTACGGAATAGGCGAGTGGAA<br>TTCCTGGTGTAGCGGTGAAATGCGCAGATATCAGGAGGAACACCGGCGGCGAAGGCAGCTCGCTAGGACGGTACTGACGCTGAGGCGCGAAAGCGTGGGG |
| BH-asv590 | GCAAGCGTTGTTCGGAATCATTGGGCGTAGAGGGCGCGCAGGCGGCCGGGCGGGTCCGGTGTGAAAGCCCGGGGCTCAACCCCGGAAGCGCACTGGAAACTGTCTGGCTTGAGTCCGGGAAGGGCCAGTGG<br>AATTCCTGGTGTAGGGGTGAAATCCGTAGAGATCAGGAGGAACACCAGAGGCGAAAGCGACTGGCTGGACCGGAACTGACGCTGAGGCGCGAAAGCACGGGT |
| BH-asv591 | GCGAGCGTTGTCCGGAATTATTGGGCGTAAAGAGTGCGTAGGCGGCAAATTAAGTCAGATGTGAAAACTAAGGGCTCAACCCATAGATTGCATCTGAAACTGATATGCTTGAGTCAAGGAGAGGAAAGTGGAATT<br>CCTAGTGTAGCGGTGGAATGCGTAGATATTAGGAGGAATACCGGTGGCGAAGGCGACTTTCTGGACTTGAACTGACGCTGAGGCACGAAAGCGTGGGG |
| BH-asv594 | GCGAGCGTTATCCGGATTTATTGGGTTTAAAGGGAGCGTAGACGGGGGATTAAGTCAGTTGTGAAAGGCTGCGGCTCAACCGCAGCACTGCAGTTGATACTGGTTTCCTTGAGTGCGGTTGAGGTGTATGGAATT<br>CGTGGTGTAGCGGTGAAATGCTTAGATATCACGAAGAACTCCGATTGCGAAGGCAGTACACTAAGCCGTAACTGACGTTGAGGCTCGAAAGTGTGGGT |
| BH-asv595 | GCCAGCGTTGCTCGGAATTACTGGGTGTAAAGGGTTCGTAGGCGGTGTGGCAAGTCGGGAGTGAAATCTCTGGGCTCAACCCAGAGTCTGCTTCCGAAACTGCTGTGCTTGAGTGTGGGAGAGGCGCGTGGAA<br>TTGCAGGTGTAGCGGTGAAATGCGTAGATATCTGCAGGAACACCCGTGGCGAAAGCGGCGCGCTGGACCACTACTGACGCTGAGGAACGAAAGCTAGGGG |
| BH-asv601 | GCTAGCGTTGTTCGGAATTACTGGGCGTAAAGCGCACGTAGGCGGCTTTCCAAGTCAGGGGTGAAATCCCGGGGCTCAACCCCGGAACTGCCCTTGAAACTGGATGGCTAGAATACTGGAGAGGTGAGTGGAAT<br>TCCGAGTGTAGAGGTGAAATTCGTAGATATTCGGAAGAACACCAGTGGCGAAGGCGACTCACTGGACAGTTATTGACGCTGAGGTGCGAAAGCGTGGGG |
| BH-asv603 | CCGAGCGTTATCCGGATTTATTGGGTTTAAAGGGTGCGTAGGTGGTTTGATAAGTCAGCGGTGAAAGTTTGCAGCTTAACTGTAAAAATGCCGTTGAAACTGTCGGACTTGAGTGTAAATGAGGTAGGCGGAATG<br>CGTGGTGTAGCGGTGAAATGCATAGATATCACGCAGAACTCCGATTGCGAAGGCAGCTTACTAAGCTACAACTGACACTGAAGCACGAAAGCGTGGGG |
| BH-asv606 | GCAAGCGTTATCCGGATTTATTGGGTTTAAAGGGTCCGTAGGCGGACTTATAAGTCAGTGGTGAAAGCCTGTCGCTTAACGATAGAACTGCCATTGATACTGTAAGTCTTGAGTATATTTGAGGTAGCTGGAATAA<br>GTAGTGTAGCGGTGAAATGCATAGATATTACTTAGAACACCAATTGCGAAGGCAGGTTACCAAGATATAACTGACGCTGAGGGACGAAAGCGTGGGG |
| BH-asv609 | GCGAGCGTTGTCCGGAATTACTGGGCGTAAAGAGCTCGTAGGTGGTTTGTCGCGTCGTTTGTGTAAGCCCGCAGCTTAACTGCGGGACTGCAGGCGATACGGGCATAACTTGAGTGCTGTAGGGGAGACTGGA<br>ATTCCTGGTGTAGCGGTGGAATGCGCAGATATCAGGAGGAACACCGATGGCGAAGGCAGGTCTCTGGGCAGTAACTGACGCTGAGGAGCGAAAGCATGGGG |
| BH-asv612 | CCAAGTGGTAGTCGTTATTATTGGGCTTAAAGTGTTCGTAGCCTGCCGTGTAAGTCCGCTGTGAAATCCACGCGCTCAACGTGTGGACGTGCAGCGGATACTGCATGGCTAGGGACCGGACGGCGTCGAGGCTA<br>TTCCAGGAGTAGGAGTGAAATCCAAAAATTCCTGGAGGAGCACCTGTGGCGAAAGCGCTCGACGAGTACGGATCCGACGGTGAGGAACGAAAGCCGGGGG |
| BH-asv613 | GCTAGCGTTGTTCGGAATTACTGGGCGTAAAGCGCACGTAGGCGGACCGGAAAGTCAGAGGTGAAATCCCAGGGCTCAACCTTGGAACTGCCTTTGAAACTCCCGGTCTTGAGTTCGAGAGAGGTGAGTGGAAT<br>TCCGAGTGTAGAGGTGAAATTCGTAGATATTCGGAGGAACACCAGTGGCGAAGGCGGCTCACTGGCTCGATACTGACGCTGAGGTGCGAAAGCGTGGGG |
| BH-asv620 | TCGAGTGATGGCCGATATTATTGGGCCTAAAGCGTTCGTAGCTGGCCGAGCAAGTCTGTCGGGAAATCTGTTCGCTCAACGAGCAGGCGTCCGGCAGAAACTGCTCGGCTTGGGACCGGAAGACTCGAGGGGT<br>ACGTCCGGGGTAGGAGTGAAATCCCGTAATCCTGGACGGACCACCGATGGCGAAAGCACCTCGAGAGCACGGATCCGACAGTGAGGAACGAAAGCTGGGGT |
| BH-asv623 | GCTAGCGTTGTTCGGATTTACTGGGCGTAAAGCGCACGTAGGCGGATTGTTAAGTCGAGGGTGAAATCCTGGAGCTCAACTCCAGAACTGCCTTCGATACTGGCAATCTAGAGTCCGGAAGAGGTGAGTGGAAC<br>TCCTAGTGTAGAGGTGGAATTCGTAGATATTAGGAAGAACACCAGTGGCGAAGGCGACTCACTGGTCCGGTACTGACGCTGAGGTGCGAAAGCGTGGGG |
| BH-asv624 | GCAAGCGTTAATCGGAATTACTGGGCGTAAAGGGTGCGTAGGCGGTTAGTTAAGTCTGTTGTGAAATCCCCGGGCTCAACCTGGGAATGGCAATGGATACTGGCTAGCTAGAGTGTGTCAGAGGATGGTGGAAT<br>CCCCGGTGTAGCGGTGAAATGCGTAGAGATCGGGAGGAACATCAGTGGCGAAGGCGGCCATCTGGGACAACACTGACGCTGAAGCACGAAAGCGTGGGG |
| BH-asv627 | CCGAGTGATGGCCGATATTATTGGGCCTAAAGCGTCCGTAGCTGGCCGGACAAGTCCGTTGGGAAATCTGCTCGCTTAACGAGCAGGCGTCCAGCGGAAACTGTTCGGCTTGGGACCGGAAGACCTGAGGGGT<br>ACGTCCGGGGTAGGAGTGAAATCCCGTAATCCTGGACGGACCACCGGTGGCGAAAGCGCCTCAGGAAGACGGATCCGACAGTCAGGGACGAAAGCTAGGGT |
| BH-asv633 | GCAAGCGTTGTTCGGATTTATTGGGCGTAAAGGGTGCCTAGGTGGCATTGTAAGTCGGGTGTGAAATATATTGGCTCAACCAATAAAGTGCACTCGATACTGCTTTGCTAGAGGGTAGGAGAGGTAAGCGGAATT<br>CTTGGTGTAGCGGTGAAATGCGTAGATATCAAGAAGAACACCGATGGCGAAGGCAGCTTACTGGCCTATTTCTGACACTGACGCACGAAAGTGTGGGG |
| BH-asv637 | CCAAGCGTTATCCGGAATCACTGGGCGTAAAGGGTGCGCAGGCGGCACGTAAGTCCGACGTGAAAGCCCCGGGCTTAACCCGGGAAGGGCGCCGGAAACTACGTGTCTTGAGGACCAGAGAGGCAGGGAGA<br>ATTCCTGGTGTAGCGGTGAAATGCGTAGATATCAGGAGGAATACCAGTGGCGAAAGCGCCCTGCTGGATGGTTCCTGACGCTGAGGCACGAAAGCGTGGGT |
| BH-asv644 | GCTAGCGTTGTTCGGAATTACTGGGCGTAAAGCGCACGTAGGCGGGTCGTTAAGTCAGGGGTGAAATCCCGGAGCTCAACTCCGGAACTGCCTTTGATACTGGCGATCTCGAGATCGGAAGAGGTGAGTGGAAT<br>TCCCAGTGTAGAGGTGAAATTCGTAGATATTGGGAAGAACACCAGTGGCGAAGGCGGCTCACTGGTCCGATACTGACGCTGAGGTGCGAAAGCGTGGGG |
| BH-asv645 | GCTAGCGTTGCTCGGAATTACTGGGCGTAAAGGGCGCGTAGGCGGACAGTTTAGTCAGAGGTGAAAGCCCAGGGCTCAACCTTGGAATTGCCTTTGATACTGGCTGTCTTGAGTACGGGAGAGGTGAGTGGAAC<br>TCCGAGTGTAGAGGTGAAATTCGTAGATATTCGGAAGAACACCAGTGGCGAAGGCGACTCACTGGCCCGTTACTGACGCTGAGGCGCGAAAGCGTGGGG |
| BH-asv646 | CCAAGCGTTGTCCGGATTTACTGGGCGTAAAGAGCGCGTAGGCGGTCTGTTAAGTGTGAAGTGAAATCTCCAGGGCTCAACCCGGAAACTGCTTCGCATACTGGCAGACTGGAGGGATGCAGAGGTTTGTGGAA<br>TTCCTGGTGTAGCGGTGATATGCATAGATATCAGGAGGAACACCCATGGCGAAGGCAGCAAACTGGGCATCATCTGACGCTGAGGCGCGAAAGCGTGGGT |

| | |
|---|---|
| BH-asv649 | GCTAGCGTTGTCCGGAATCATTGGGCGTAAAGCGCGTGTAGGCGGTTCGGTAAGTCCGCTGTGAAAGTCGGGGGCTCAACCCTCGAAAGCCGGTGGATACTGTCGGGCTAGAGTACGGAAGAGGCGAGTGGAATTCCTGGTGTAGCGGTGAAATGCGCAGATATCAGTAGGAACACCAATAGCGAAGGCAGCTCGCTGGGACGTTACTGACACTGAGGCGCGAAAGCGTGGGG |
| BH-asv652 | GCGAGCGTTAATCGGAATTACTGGGCGTAAAGCGCATGCAGGTGGTTTGTTAAGTCAGATGTGAAAGCCCGGGGCTCAACCTCGGAATAGCATTTGAAACTGGCAGACTAGAGTACTGTAGAGGGGGGTAGAATTTCAGGTGTAGCGGTGAAATGCGTAGAGATCTGAAGGAATACCGGTGGCGAAGGCGGCCCCCTGGACAGATACTGACACTCAGATGCGAAAGCGTGGGG |
| BH-asv656 | CCTAGCGTTAAGCGGAATCACTGGGCTTAAAGGGTGCGTAGGCGGACTTGAAAGTACCTTGTGAAATCCCACGGCTCAACCGTGGAAGGGCATGGTAAACTTCAAGTCTTGAGCACGGTAGGGGCTGTCGGAACGCTGGGTGTAGCGGTGAAATGCGTAGATATCCAGCGGAACGCCAAAAGCGAAAGCAGACAGCTGGGCCGTTGCTGACGCTGAGGCACGAAAGCGTGGGG |
| BH-asv663 | GCAAGCGTTATCCGGAATTATTGGGCGTAAAGCGTCTGTAGGTTGTTAATTAAGTCTATTGTTAAATATTAAAGCTTAACTTTGAATAAGCAATAGAAACTACTTAGCTAGAGTTTAGAAGAGGTAGAGGGAATTCCTAGTGTAGCGGTGAAATGTGTAGATATTAGGAAGAACACCGGTGGCGAAAGCGCTCTACTGGGCTAATACTGACACTGAGAGACGAAAGCTAGGGG |
| BH-asv664 | ACGAGTGATGGCCAATCTTATTGGGCCTAAAGCGTCCGTAGCCGGCCAGACAAGTCTGTTGGGAAATCGGCGCGCTCAACGCGTCGGCGTCCAGCGGAAACTGTCTGGCTTGGGGCCGGAGGACCTGAGGGGTACGTCCGGGGTAGGAGTGAAATCCCGTAATCCTGGACGGACCACCGGTGGCGAAAGCGCCTCAGGAGGACGGACCCGACGGTGAGGGACGAAAGCTAGGGT |
| BH-asv665 | CCAAGCGTTATCCGGATTTATTGGGTTTAAAGGGTGCGTAGGTGGCTTTTTAAGTCAGGGGTGAAAGACGGCAGCTTAACTGTCGCAGTGCCCTTGATACTGAAGAGCTTGAATCCACTAGAGGTAGGCGGAATGTGACAAGTAGCGGTGAAATGCATAGATATGTCACAGAACACCAATTGCGAAGGCAGCTTACTATGGTGGTATTGACACTGAGGCACGAAAGCGTGGGG |
| BH-asv666 | GCAAGCGTTATCCGGATTTACTGGGTGTAAAGGGAGCGTAGACGGATGGACAAGTCTGATGTGAAAGGCTGGGGCTCAACCCCGGGACTGCATTGGAAACTGCCCGTCTTGAGTGCCGGAGAGGTAAGCGGAATTCCTAGTGTAGCGGTGAAATGCGTAGATATTAGGAGGAACACCAGTGGCGAAGGCGGCTTACTGGACGGTAACTGACGTTGAGGCTCGAAAGCGTGGGG |
| BH-asv671 | CAAGCGTTAATCGGAATCACTGGGCGTAAAGGGTGCGTAGGCGGTACGTTAGGTCTGTCGTGAAATCCCCGGGCTCAACCTGGGAATGGCGGTAGAAACCGGCGCACTAGAGTATGGGAGAGGGTGGTGGAATTCCAGGTGTAGCGGTGAAATGCGTAGAGATCTGGAGGAACATCAGTGGCGAAGGCGGCCACCTGGCCCAATACTGACGCTGAGGCACGAAAGCGTGGGG |
| BH-asv676 | GCAAGCGTTATCCGGAATTATTGGGCGTAAAGAGCTCGTAGGCGGTTTGTCGCGTCTGTCGTGAAAGTCCGGGGCTCAACTCCGGATCTGCGGTGGGTACGGGCAGACTAGAGTGATGTAGGGGAGACTGGAATTCCTGGTGTAGCGGTGGAATGCCCAGATATCAGGAGGAACACCGATGGCGAAGGCAGGTCTCTGGGCATTAACTGACGCTGAGGAGCGAAAGCATGGGG |
| BH-asv677 | CCGAGTGATGGCCACTATTATTGGGCTTAAAGCGCCCGTAGCCGGCCAGGTAAGTCCTCGGTGAAATCTGCCTGCTCAACAGGTAGGAGCGCCGAGGAAACTGCGTGGCTTGGGGCCGGACGGCGTCGGGGGTATTCCCGGGGTAGGGGTAAAATCCAAAAATCCCGGGAGGACCGCCCGTGGCGAAAGCGCCCGACGAGGACGGACCCGACGGTGAGGGACGAAAGCCAGGGG |
| BH-asv679 | GCAAGCGTTATCCGGATTCATTGGGTTTAAAGGGTGCGCAGGTGGGAATGTAAGTCAGTGGTGAAATACTGCCGCTTAACGGTAGAGCTGCCATTGATACTGCGTTTCTTGAGTATGGTTGAGGTAGGTGGAATGTGTTGTGTAGCGGTGAAATGCATAGATATGACACATAACGCCGATTGCGAAGGCAACTTACTAAGCCATTACTGACGCTGAGGCACGAAAGCGTGGGT |
| BH-asv681 | GCAAGCGTTAATCGGAATCACTGGGCGTAAAGGGTGCGTAGGCGGTGCGTTAGGTCTGTCGTGAAATCCCCGGGCTCAACCTGGGAATGGCGGTGGAAACCGGCGCGCTAGAGTATGGGAGAGGGTGGTGGAATTCCAGGTGTAGCGGTGAAATGCGTAGAGATCTGGAGGAACATCAGTGGCGAAGGCGGCCACCTGGCCCAATACTGACGCTGAGGCACGAAAGCGTGGGG |
| BH-asv683 | TCGAGTGATGACCGATCTTATTGAGCCTAAAGCGTCCGTAGCCGGCCACGCAAGTCCATCGGGAAATCCGCACGCCCAACGTGCGGGCGTCCGGTGGAAACTGCCTGGCTTGGGACCGGAAGACTCGACGGGTACGTTCGGGGTAGGAGTGAAATCCCGTAATCCTGAACGGACCACCGATGGCGAAAGCACGTCGAGAAGACGGATCCGACGGTGAGGGACGAAAGCTAGGGT |
| BH-asv684 | CCGAGTGGTGGCCACTTTTATTGGGCTTAAAGTGTTCGTAGCCTGCTCTGTGAGTCTGCTGTGAAATCTGGCAGCTCAACTGTCAGGCGTGCAGCGGATACCACAGGGCTTGGGACCGGAAGGCGTTGAGGCTATTCCTGGGGTAGGGGTGAAATCCTGAAATCCCAGGAGGAGCACCAGTGGCGAAAGCTCTCAACGATGACGGGTCCGACGGTGAGGGACGAAAGCCAGGGG |
| BH-asv685 | CCAAGTGGTAGCCGTTATTATTGGGCTTAAAGTGTTCGTAGCCGGCCGTGTAAGTCCGCTGTGAAATCCAAGCGCTCAACGTTTGGAAGCGCAGCGGATACTGCATGGCTAGGGACCGGACGGCGTCGAGGCTATTCCAGGGGTAGGAGTGAAATCCAAAAATCCTTGGAGGAGCACCTGTGGCGAAAGCGCTCGGCGAGTACGGATCCGACGGTGAGGAACGAAAGCCAGAGG |
| BH-asv687 | GCGAACGTTATTCGGAATCACTGGGCTTACAGGGTGCGTAGGCGGTGAGACAAGTCAGATGTGAAATCCCACAGCTCAACTGTGGAACTGCGTTTGAAACTGTCTTACTTGAGGAATCTAGAGGTATGCGGAACTTCCGGTGGAGCGGTGAAATGTGTTGATATCGGAAGGAACACCAGAGGCGAAGGCGGCATACTGGGGATTATCTGACGCTGAGGCACGAAAGCCAGGGG |
| BH-asv689 | GCGAGCGTTGTTCGGAATTATTGGGCGTAAAGGGCATGTAGGCGGTTATGTAAGCCTGATGTGAAATCCTGGGGCTTAACCCTAGAATAGCATTGGGTACTGTGTAACTTGAATTACGGAAGGGAAACTGGAATTCCAAGTGTAGGGGTGGAATCTGTAGATATTTGGAAGAACACCGGTGGCGAAGGCGGGTTTCTGGCCGATAATTGACGCTGAGATGCGAAAGTGTGGGG |
| BH-asv690 | GCAAGCGTTATCCGGAATCACTGGGTGTAAAGGGTGCGCAGGCGGACGTGTAGGTCTAATGTGAAAGTCCCGGGCTCAACCCGGGAATTGCATTGGAAACCCCACGTCTTGAGGGCCAGAGAGGCAGGCGCAATTCGTGGTGTAGCGGTGAAATGCGTAGATATCACGAGGAACATCAGTGGTGTAGACGGCCTGCTGGATGGACCCTGACGCTGAGGCACGAAAGCGTGGGG |
| BH-asv691 | GCAAGCGTTATCCGGAATCACTGGGCGTACAGGGAGCGCAGGCGGACGGGCAGGTCTGATGTGAAAGCCCCGGGCTCAACCCGGGAAGTGCATCGGAAACCTCCCGTCTTGAGGACCAGAGAGGGAGGCGCAATTCGCGGTGTAGCGGTGAAATGCGTAGATATCGCGAGGAACACCAGTGGCGAAAGCGGCCTCCTGGATGGTTCCTGACGCTGAGGCTCGAAAGCGTGGAT |
| BH-asv698 | GCGAGCGTTAATCGGAATTACTGGGCGTAAAGCGTGCGCAGGCGGTGATGTAAGACAGATGTGAAATCCCCGGGCTCAACCTGGGACCTGCATTTGTGACTGCATCGCTAGAGTACGGTAGAGGGGGATGGAATTCCGCGTGTAGCAGTGAAATGCGTAGATATGCGGAGGAACACCGATGGCGAAGGCAATCCCCTGGACCTGTACTGACGCTCATGCACGAAAGCATGGGG |
| BH-asv699 | CCAGGCGTTATCCGGATTCATTGGGTTTAAAGGGAGCGTAGGCCGCCTTTTAAGCGTGTTGTGAAATGTAGGGGCTCAACCCCTGCACTGCAGCGCGAACTGGAGGGCTTGAGTACACACAAAGTAGGCGGAATTCATGGTGTAGCGGTGAAATGCTTAGATATCATGAAGAACCCCGATTGCGAAGGCAGCTTACTGGAGTGTAACTGACGCTGAAGCTCGAAAGCGCGGGT |
| BH-asv704 | GCAAGCGTTATCCGGAATTACTGGGCGTAAAGCGCACGCAGGTGGTTGTCTAAGACGGACGTGAAAGCTCCTGGCTTAACTGGGAGAGGTCGTTGGTGACTGGATGACTTGAGGCGAGGAGAGGGGTGTAGAATTCCCGGTGTAGTGGTGGAATGCGTAGAGATCGGGAGGAATACCAGTGGCGAAAGCGACACCCTGGCCTTGGCCTGACGCTCAGGTGCGAAAGCGTGGGG |
| BH-asv707 | GCAAGCGTTAATCGGAATTACTGGGCGTAAAGCGTGCGTAGGTGGTTGATTAAGTTATCTGTGAAATTCCTGGGCTTAACCTAGGCAGGTCAGATGATACTGATTGACTCGAGTATGGGAGAGGGTAGTGGAATTTCCGGTGTAGCGGTGAAATGCGTAGAGATCGGAAGGAACACCAGTGGCGAAGGCGGCTACCTGGCCTAATACTGACACTGAGGCACGAAAGCGTGGGG |
| BH-asv708 | GTTAGCGTTGTTCGGAATTACTGGGCGTAAAGCGTACGTAGGCGGATCAGAAAGTTAGGGGTGAAATCCCAGGGCTCAACCCTGGAACTGCCTCTAAAACTCCTGGTCTTGAGTTCGAGAGAGGTGAGTGGAATTCCGAGTGTAGAGGTGAAATTCGTAGATATTCGGAGGAACACCAGTGGCGAAGGCGGCTCACTGGCTCGATACTGACGCTGAGGTACGAAAGTGTGGGG |
| BH-asv709 | GCGAGCGTTGTCCGGAATTATTGGGCGTAAAGAGCTCGTAGGCGGCTGGTCGCGTCTGTCGTGAAATCCTCTGGCTTAACTGGGGGCTTGCGGTGGGTACGGGCCGGCTTGAGTGCGGTAGGGGAGACTGGAACTCCTGGTGTAGCGGTGGAATGCGCAGATATCAGGAAGAACACCGGTGGCGAAGGCGGGTCTCTGGGCCGTTACTGACGCTGAGGAGCGAAAGCGTGGGG |
| BH-asv715 | GCAAGCGTTGTCCGGAATTATTGGGCGTAAAGCGCGCGCAGGCGGCCTTTTAAGTCTGATGTGAAAGCCCCCGGCTCAACCGGGGAGGGTCATTGGAAACTGGAAGGCTTGAGTGCAGAAGAGAAGAGCGGAATTCCACGTGTAGCGGTGAAATGCGTAGAGATGTGGAGGAACACCAGTGGCGAAGGCGGCTCTTTGGTCTGTAACTGACGCTGAGGCGCGAAAGCGTGGGG |
| BH-asv722 | CCGAGTGGTGGCCACGATTATTGGGCTTAAAGTGTTCGTAGCCGGCCATACAAGTTCTCTGTGAAATCCAGCCGCTTAACGGTTGGACGCGCAGAGAATACTGTATGGCTTGGGGCCGGACGGCGTTGGGGCTATTCCAGGGGTAGGAGTGAAATCCTGAAATCCCTGGAGGAGCACCGGTGGCGAAAGCGCCCAGCGAGGACGGACCCGACGGTGAGGAACGAAAGCCAGGGG |
| BH-asv723 | GCGAGCGTTAATCGGATTTACTGGGCGTAAAGCGCGCGTAGGTGGTTAATTAAGTCAAATGTGAAATCCCCGAGCTTAACTTGGGAATTGCATTCGATACTGGTTAGCTAGAGTATGGGAGAGGATGGTAGAATTCCAGGTGTAGCGGTGAAATGCGTAGAGATCTGGAGGAATACCGATGGCGAAGGCAGCCATCTGGCCTAATACTGACACTGAGGTGCGAAAGCATGGGG |
| BH-asv724 | GCAAGCGTTAATCGGAATTACTGGGCGTAAAGCGCGCGTAGGTGGTTTTGTAAGTTGGAGGTGAAATCCCCGGGCTCAACCTGGGAACTGCCTCCAAAACTGCATGACTAGAGTACGGTAGAGGGTAGTGGAATTTCCTGTGTAGCGGTGAAATGCGTAGATATAGGAAGGAACACCAGTGGCGAAGGCGACTACCTGGACTGATACTGACACTGAGGTGCGAAAGCGTGGGG |
| BH-asv725 | GCAAGCGTTATTCGGAATTATTGGGCGTAAAGGGCGCGTAGGCGGCCAAGCAAGTCAGGGGTGAAATCCCTCGGCTCAACCGAGGAATTGCCTCTGAAACTGCATGGCTTGAGACCGGGAGAGGGTGGTGGAATTCCCAGTGTAGCGGTGAAATGCGTAGATATTGGGAGGAACACCGGTGGCGAAGGCGGCCACCTGGACCGGTTCTGACGCTGAGGCGCGAAAGTGTGGGG |
| BH-asv727 | GCAAGCGTTGTCCGGAATTATTGGGCGTAAAGCGCGCGCAGGCGGCTTTGTAAGTCTTGTGTTTAAACGCGGGGCTCAACCCCGTGTCGCATGGGAAACTGCAAGGCTTGAGTGCAGAAGAGGAAAGTGGAATTCCACGTGTAGCGGTGAAATGCGTAGAGATGTGGAGGAACACCAGTGGCGAAGGCGACTTTCTGGGCTGTAACTGACGCTGAGGCGCGAAAGCGTGGGG |
| BH-asv728 | GCTAGCGTTGTTCGGAATTACTGGGCGTAAATGGCGCGTAGGCGGCTCTTTCAGTCGGATGTGAAAGCCCGGGGCTCAACCCCGGAACGGCATCCGAGACGGGAGGGCTCGAGACCGGGAGAGGAGGGTGGAATAGCCAGTGTAGAGGTGAAATTCGTAGATATTGGCTGGAACACCGGTGGCGAAAGCGGCTCTCTGGCCCGGTTCTGACGCTGAGGCGCGAAAGCGTGGGG |
| BH-asv729 | CCGAGTGATGGCCGCTATTATTGGGCCTAAAGCGTCCGTAGCCGGCCGGTCAAGTCCGTCGGGAAATCCACTCGCTTAACGAGTGGGCGTCCGGCGGAAACTGATCGGCTTGGGACCGGAAGACCCGAGGGGTACGTCCAGGGTAGGAGTGAAATCCCGTAATCCTGGACGGACCGTCGGTGGCGAAAGCGCCTCGGGAAGACGGATCCGACGGTGAGGGACGAAAGCTAGGGT |
| BH-asv730 | CCAAGCGTTGTCCGGAATTATTGGGTGTAAAGGGTCCGCAGGCGGGACAGTAAGTCGGAGGTGAAAGCCCTCGGCTCAACCGAGGAATTGCCTTCGATACTGCTGTTCTTGAGTCCCGGAGAGGCCACCGGAATTCGTGGTGTAGCGGTGAAATGCGTAGATATCACGAGGAACACCAGAAGTGAAAACGGGTGGCTGGACGGGTACTGACGCTCAGGGACGAAAGCGTGGGG |
| BH-asv733 | GCTAGCGTTGTTCGGAATTACTGGGCGTAAAGCGCACGTAGGCGGATATTTAAGTCAGGGGTGAAATCCCAGAGCTCAACTCTGGAACTGCCTTTGATACTGGGTATCTCGAGTATGGAAGAGGTGAGTGGAATTCCGAGTGTAGAGGTGAAATTCGTAGATATTCGGAGGAACACCAGTGGCGAAGGCGGCTCACTGGTCCATTACTGACGCTGAGGTGCGAAAGCGTGGGG |
| BH-asv734 | GCTAGCGTTATTCGGAATTACTGGGCGTAAAGCGCACGTAGGCGGACCAGAAAGTCAGAGGTGAAATCCCGGGGCTCAACCCCGGAACTGCCTTTGAAACTCCTGGTCTTGAGGTCGAGAGAGGTGAGTGGAATTCCGAGTGTAGAGGTGAAATTCGTAGATATTCGGAGGAACACCAGTGGCGAAGGCGGCTCACTGGCTCGATACTGACGCTGAGGTGCGAAAGCGTGGGG |
| BH-asv737 | CCGAGTGGTGGCCGCTGTTATTGGGCTTAAAGTGTCCGTAGCCGGCCCGGTAAGTTCTCTGTGAAATCCGGCCGCTCAACGGTCGGGCGCGCAGAGAATACTGCTGGGCTTGGGGCCGGATGGCGTCGGGGCTATTCCCGGGGTAGGAGTGAAATCCTGAAATCCCGGGAGGAGCACCTGTGGCGAAAGCGCCCGGCGAAGACGGACCCGACGGTCAGGGACGAAAGCCAGGGG |
| BH-asv738 | GCTAGCGTTGTTCGGAATTACTGGGCGTAAAGCGCACGTAGGCGGACGATCAAGTTGGGGGTGAAAGCCCAAGGCTCAACCTTGGAATTGCCCCCAAAACTGATCATCTAGAGGCCAGGAGAGGGCAGTGGAATTCCCAGTGTAGAGGTGAAATTCGTAGATATTGGGAAGAACACCAGTGGCGAAGGCGGCTGCCTGGACTGGACCTGACGCTGAGGTGCGAAAGCGTGGGG |
| BH-asv739 | CCAAGTGGTAGCCACTTTTATTGGGCTTAAAGTGTTCGTAGCCTGCCGTGTAAGTCCGCTGTGAAACCCACGCGCTCAACGTGTGGATGTGCAGCGGATACTGCATGGCTAGGGACCGGACGGCGTCGAGGCTATTCCAGGGGTAGGAGTGAAATCCAAAAATCCCTGGAGGAGCACCTGTGGCGAAAGCGCTCGGCGAGTACGGATCCGACGGTGAGGAACGAAAGCCAGGGG |
| BH-asv740 | GCGAGCGTTAATCGGAATTACTGGGCGTAAAGCGTGCGCAGGCGGTTATATAAGACAGATGTGAAATCCCCGGGCTCAACCTGGGAACTGCATTAGTGACTGTATAGCTGGAGTGCGGCAGAGGGGGATGGAATTCCGCGTGTAGCAGTGAAATGCGTAGATATGCGGAGGAACACCGATGGCGAAGGCAATCCCCTGGGCCTGCACTGACGCTCATGCACGAAAGCGTGGGG |
| BH-asv741 | GCAAACGTTGTTCGGAATTACTGGGCGTAAAGCGCGTGTAGGCTGCCCCGCAAGTCGGATGTGAAAGCCCTGGGCTCAACCTAGGAAGTGCATTCGAAACTGCGGGGCTGGAGTCCTGGAGAGGATAGCGGAATTCTCGGTGTAGAGGTGAAATTCGTAGATATCGAGAGGAACACCGGTGGCGAAGGCGGCTATCTGGACAGTGACTGACGCTGAGACGCGAAAGCGTGGGG |
| BH-asv743 | GCGAGCGTTAATCGGAATTACTGGGCGTAAAGGGCGCGTAGGCGGCCGATCAAGTCAGATGTGAAAGCCCACGGCTTAACCGTGTGAAGTGCATCTGAAACTGGTTGGCTAGAGTATCGGAGGGGAAAGTGGAATTCCCGGTGTAGAGGTGAAATTCGTAGATATCGGGAGGAATACCGGTGGCGAAGGCGACTTTCTGGACGAATACTGACGCTGAGGCGCGAAAGCGTGGGG |
| BH-asv744 | GCAAGCGTTGTCCGGAATTATTGGGCTTAAAGCGCGCGCAGGCGGTTCCTTAAGTCTGATGTGAAAGCCCACGGCTCAACCGTGGAGGGTCATTGGAAACTGGGGAACTTGAGTGCAGAAGAGGAAAGCGGAATTCCACGTGTAGCGGTGAAATGCGTAGAGATGTGGAGGAACACCAGTGGCGAAGGCGGCTTTCTGGTCTGTAACTGACGCTGAGGCGCGAAAGCGTGGGG |
| BH-asv746 | CCAAGCGTTATCCGGAATTACTGGGCGTAAAGGGTGCGCAGGCGGGCACGTAAGTCCGACGTGAAAGCCCCGAGCTTAACTCGGGAAGTGCGCCGGAAACTACGTGTCTTGAGGACCAGAGAGGCAGGGAGAATTCCTGGTGTAGCGGTGAAATGCGTAGATATCAGGAGGAATACCTGTGGCGAAAGCGCCCTGCTGGATGGTTCCTGACGCTGAGGCACGAAAGCGTGGGT |
| BH-asv747 | GCAAGCGTTAATCGGAATTACTGGGCGTAAAGAGCACGTAGGCGGTAGGGCAAGTCGGATGTGAAAGCCCTGGGCTCAACCTGGGAATTGCATTCGATACTGCTCTACTAGAGTTCGGGAGAGGAGAGCGGAATTCCCGGTGTAGCGGTGAAATGCGTAGATATCGGGAGGAACACCAGTGGCGAAGGCGGCTCTCTGGCCCGAAACTGACGCTGAGGTGCGAAAGCGTGGGG |

| | |
|---|---|
| BH-asv749 | GCAAGCGTTAATCGGAATCACTGGGCGTAAAGGGCACGCAGGCGGTTGGATAAGTCGGGTGTGAAAGCCCCGGGCTCAACCTGGGAATGGCATTCGATACTGTCTAGCTAGAGTCTGGTAGAGGGAAGTGGAATTTCCGGTGTAGCGGTGAAATGCGTAGATATCGGAAGGAACACCAGTGGCGAAGGCGACTTCCTGGACCAAGACTGACGCTGAGGTGCGAAAGCGTGGGT |
| BH-asv750 | GCAAGCGTTAATCGGAATTACTGGGCGTAAAGCGCACGTAGGCTGTAGTGTAAGTCAGGGGTGAAATCCCACGGCTCAACCGTGGAACTGCCTTTGATACTGCACAACTTGAATCCGGGAGAGGGTGGCGGAATTCCAGGTGTAGGAGTGAAATCCGTAGATATCTGGAGGAACATCAGTGGCGAAGGCGGCCACCTGGACCGGTATTGACGCTGAGGTGCGAAAGCGTGGGG |
| BH-asv752 | GCTAGCGTTGTTCGGAATTACTGGGCGTAAAGCGCACGTAGGCGGATCGATAAGTAAGGGGTGAAATCCCAGAGCTCAACTCTGGAACTGCCTTTTAAACTGTCGATCTGGAGCAGAGTAGAGGTGAGTGGAATTCCGAGTGTAGAGGTGAAATTCGTAGATATTCGGAGGAACACCAGTGGCGAAGGCGGCTCACTGGACTCTTGCTGACGCTGAGGTGCGAAAGCGTGGGG |
| BH-asv755 | GCAAGCGTTAATCGGAATTACTGGGCGTAAAGCGTGCGCAGGCGGTGATGTAAGACAGTTGTGAAATCCCCGGGCTCAACCTGGGAACTGCATCTGTGACTGCATCGCTGGAGTGCGGCAGAGGGGGATGGAATTCCGCGTGTAGCAGTGAAATGCGTAGATATGCGGAGGAACACCGATGGCGAAGGCAATCCCCTGGGCCTGCACTGACGCTCATGCACGAAAGCGTGGGG |
| BH-asv757 | GCAAGCGTTATCCGGATTCACTGGGTTTAAAGGGTGCGTAGGCGGGCGTATAAGTCAGTGGTGAAATCCTGGAGCTTAACTCCAGAACTGCCATTGATACTATATGTCTTGAATATGGTGGAGGTAAGCGGAATATGTCATGTAGCGGTGAAATGCATAGATATGACATAGAACACCTATTGCGAAGGCAGCTTACTACGCCTATATTGACGCTGAGGCACGAAAGCGTGGGG |
| BH-asv761 | CCAAGCGTTAGTCGGAGTCACTGGGCTTAAAGGGTGCGCAGGCGGGGATGTGCGTGTCTTGTGAAAGCCCACGGCTCAACCGTGGAATGGCAGGGCATACGACATCCCTTGAGGCCGGCAGAGGGTGCCGGAACTCTAGGTGTAGCGGTGAAATGCGTAGATATCTAGAGGAACACCGATGGCGAAAGCAGGCACCTGGGCCGGACCTGACGCTGAGGCACGAAAGCGTGGGG |
| BH-asv762 | GCAAGCGTTGTCCGGATTTATTGGGCGTAAAGAGCTCGTAGGCGGTTCCGTAAGTCGAGTGTGAAAAATCTGGGCTCAACCCAGTGGAGCACTCGATACTGCGGTGACTAGAGTACGGTAGAGGAGTGTGGAATTCCTGGTGTAGCGGTGAAATGCGCAGATATCAGGAGGAACACCAACGGCGAAGGCAGCACTCTGGGCCGGTACTGACGCTGAGGAGCGAAAGCGTGGGG |
| BH-asv764 | GCAAGCGTTGTCCGGAATTATTGGGCGTAAAGATCTCGTAGGCGGTTTGTCGCGTCGATTGTGAAAGCTCGGGGCTTAACCCCGGGTCTGCAGTCGATACGGGCAGGCTAGAGTTCGGTAGGGGAGACTGGAATTCCTGGTGTAGCGGTGAAATGCGCAGATATCAGGAGGAACACCGGTGGCGAAGGCGGGTCTCTGGGCCGATACTGACGCTGAGGAGCGAAAGCGTGGGG |
| BH-asv765 | CCTAGCGTTGCTCGGAATCATTGGGCGTAAAGCGGGTGTAGGTGGACGACCAAGTCAGATGTGAAAGCCCCGGGCTTAACCCGGGAAGTGCATTTGAAACTGGTTGTCTTGAGTGCAGTAGAGGAAGGTGGAATTCCAAGTGTAGTGGTGAAATACGTAGATATTTGGAGGAACATCGGTGGCGAAGGCGACCTTCTGGACTGTTACTGACACTGAGACCCGAAAGCGTGGGG |
| BH-asv766 | GCGAGCGTTAATCGGAATTACTGGGCGTAAAGCGTGCGCAGGCGGTTGTGTAAGACAGATGTGAAATCCCCGGGCTCAACCTGGGAACTGCGTTTGTGACTGCACAACTAGAGTACGGCAGAGGGAGGTGGAATTCCGCGTGTAGCAGTGAAATGCGTAGAGATGCGGAGGAACACCGATGGCGAAGGCAGCCTCCTGGGCCAGTACTGACGCTCATGCACGAAAGCGTGGGG |
| BH-asv767 | GCAAGCGTTAATCGGAATTACTGGGCGTAAAGCGTGCGTAGGCGGTTTCGTAAGTCGGGTGTGAAAGCCCCGGGCTCAACCTGGGAATTGCATTCGAGACTGCGAAGCTGGAGTGCGGAAGAGGGAAGTGGAATTTCCGGTGTAGCGGTGAAATGCGTAGATATCGGAAGGAACATCAGTGGCGAAAGCGACTTCCTGGTCCAGCACTGACGCTCAGGCACGAAAGCGTGGGG |
| BH-asv768 | GCAAGCGTTAATCGGAATTACTGGGCGTAAAGCGTGCGCAGGCGGTTATGCAAGACAGATGTGAAATCCCCGGGCTCAACCTGGGAACTGCATTTGTGACTGCATGGCTAGAGTGCGGCAGAGGGGGATGGAATTCCGCGTGTAGCAGTGAAATGCGTAGATATGCGGAGGAACACCGATGGCGAAGGCAATCCCCTGGGCCTGCACTGACGCTCATGCACGAAAGCGTGGGG |
| BH-asv769 | GCGAGCGTTGTCCGGAATTACTGGGCGTAAAGGGAGCGTAGGCGGTCGATTAAGTTAGATGTGAAACCCCCGGGCTTAACTTGGGGACTGCATCTAATACTGGTTGACTTAGAGTACAGGAGAGGGAAGCGGAATTCCTAGTGTAGCGGTGAAATGCGTAGATATTAGGAGGAACACCAGTGGCGAAGGCGGCTTTCTGGACTGACACTGACGCTGAGGCTCGAAAGCGTGGGG |
| BH-asv770 | GCAAGCGTTGTCCGGAATCACTGGGCGTAAAGGGTTCGCAGGCGGCCTGACAACTCAGGCGTGAAATCCCACGGCTCAACCGTGGAATTGCGTCTGAAAGTGTCCGGCTCGAGGACAGGAAGGGGGAGTGGAACTCCGGGTGGAGCGGTGAAATGCGCAGATATCCGGAGGAACACCGGAGGCGAAAGCGACTCCCTAGCCTGTTCCTGACGCTGAGGGACCAAAGCCAGGGT |
| BH-asv771 | GCAAGCGTTAATCGGAATTACTGGGCGTAAAGGGCGTGTAGGCGGCTGGGTAAGTCGGGTGTGAAAGCCCCGGGCTCAACCCGGGAATTGCATTCGATACTGCTCAGCTAGAGTCCAGGAGAGGAAGGCGGAATTCCCGGTGTAGCGGTGAAATGCGTAGATATCGGGAGGAACACCAGTGGCGAAGGCGGCCTTCTGGACTGAGACTGACGCTGAGGCGCGAAAGCGTGGGG |
| BH-asv772 | CCGAGTGGTCGGGACGATTATTGGGCCTAAAGCATCCGTAGCCGGTGCTGCAAGTCTTCCGTTAAATCCACCTGCTTAACAGTTGGGCTGCGGGGGATACTACAGCGCTAGGAGGCGGGAGAGGCAAGCGGTACTCAATGGGTAGGGGTAAAATCCGCTGATCCATTGAAGACCACCAGTGGCGAAGGCGGCTTGCCAGAACGCGCTCGACGGTGAGGGATTAAAGCTGGGGG |
| BH-asv773 | GCGAGCGTTATCCGGAATTACTGGGCGTAAAGCGCGCGTAGGCGGTTATTTAAGTGGTGCGTGAAAGCTCCTGGCTCAACTGGGAGAGGTCGTACCAAACTGGGTAACTAGAGACAGATAGAGGTCAGTGGAATTCCGGGTGGAGTGGTGAAATGCGTAGAGATCCGGAGGAACACCAGAGGCGAAGGCGACTGGCTGGGTCTGTTCTGACGCTGAGGCGCGAAAGCGTGGGG |
| BH-asv775 | GCAAGCGTTACCCGGAATCACTGGGCGTAAAGGGCGTGTAGGCGGTTATTTAAGTCTGGTTTTAAAGACCGGGGCTCAACCCCGGGAGTGGACTGGATACTGGATGACTTGACCTCTGGAGAGGGAACTGGAATTCCTGGTGTAGCGGTGGAATGCGTAGATACCAGGAGGAACACCAATGGCGAAGGCAAGTTCCTGGACAGAAGGTGACGCTGAGGCGCGAAAGTGTGGGG |
| BH-asv779 | CCGAGTGGTGGCCGCTATTATTGGGCTTAAAGAGTCCGTAGCCGGCCCGGTAAGTTCTCTGTGAAATCCGGCCGCTCAACGGTCGGGCGCGCAGGGAATACTGCCGGGCTTGGGGCCGGACGGCGTCGGGGCTATTCCCGGGGTAGGAGTGAAATCCCAAAATCCCGGGAGGAGCACCTGTGGCGAAAGCGCCCGGCGAGGACGGACCCGACGGTCAGGGACGAAAGCCAGGGG |
| BH-asv780 | GCAAGCGTTGTTCGGAATCATTGGGCGTACAGGGCGCGCAGGCGGCCGGGCAAGTCCGATGTGAAAGCCCGGGGCTCAACCCCGGAGTTGCATTGGAAACTGTCTGGCTTGAGTCCGAGAAGGGTCAGTGGAATTCCTGGTGTAGGGGTGAAATCCGTAGATATCAGGAGGAACACCGGCGGCGAAAGCGACTGACTGGATCGGGACTGACGCTGAGGCGCAAAAGCGCGGGT |
| BH-asv781 | GCAAGCGTTGTCCGGAATTATTGGGCGTAAAGCGCTCGCAGGCGGTCTTTTAAGTCTGATGTGAAAGCCCACGGCTTAACCGTGGAGGGTCATTGGAAACTGGAGGACTTGAGTACAGAAGAGGAGAGTGGAATTCCACGTGTAGCGGTGAAATGCGTAGAGATGTGGAGGAACACCAGTGGCGAAGGCGACTCTCTGGTCTGTAACTGACGCTGAGGAGCGAAAGCGTGGGT |
| BH-asv783 | TCGAGTGATGGCCGATTTTATTGGGCCTAAAGCGTCCGTAGCTGGCCTGGCAAGTCCGTCGGGAAATCCACCCGCCCAACGGGTGGGCGTCCGGCGGAAACTGCCGGGCTTGGGACCGGGAGGCTTGAGGGGTACGTCCGGGGTAGGAGTGAAATCCCGTAATCCTGGACGGACCACCGATGGGGAAACCACCTCAAGAGAACGGATCCGACAGTGAGGGACGAAAGCTAGGGT |
| BH-asv785 | GCGAGCGTTGTCCGGATTTATTGGGCGTAAAGCGCGCGCAGGCGGTTCCTTAAGTCTGATGTGAAAGCCCACGGCTCAACCGTGGAGGGTCATTGGAAGCTGGGGAACTTGAGTGCAGAAGAGGAGAGCGGAATTCCACGTGTAGCGGTGAAATGCGTAGAGATGTGGAGGAACACCAGTGGCGAAGGCGGCTCTCTGGTCTGTAACTGACGCTGAGGCGCGAAAGCGTGGGG |
| BH-asv786 | GCGAGCGTTAATCGGAATTACTGGGCGTAAAGCGTGCGCAGGCGGTGATGTAAGACAGGTGTGAAATCCCCGGGCTTAACCTGGGAACTGCATTTGTGACTGCATCTCTGGAGTGCGGCAGAGGGGGATGGAATTCCGCGTGTAGCAGTGAAATGCGTAGATATGCGGAGGAACACCGATGGCGAAGGCAATCCCCTGGGCCTGCACTGACGCTCATGCACGAAAGCGTGTGG |
| BH-asv787 | CCGAGTGGTGGCCACGTTTATTGGGCTTAAAGAGTCCGTAGCCGGCCTGGTAAGTTCTCTGTGAAATCCGGTCGCTCAACGATCGGGCGTGCAGGGAATACTGTCAGGCTTGGGGCCGGACGGCGCTGGAGCTATTCCCGGAGTAGGAGTGAAATCCTAAAATTCCGGGAGGAGCGCCTGTGGCGAAAGCGTCCAGCGAGGACGGACCCGACGGTGAGGGACGAAAGCCAGGGG |
| BH-asv788 | ACAAGCGGGAACCGCGTTTATTGGGTCTAAAGCGTTCGTAGCCGGTCCAGTAAATACCTTGTGAAATCGTTCCGCTCAACGGAACGGCGTGCGAGGTACACTGGTGGACTAGGGACCGGGATACGTAAGCGGTATTCCCGGGGGAGCGGTAAAATGCAATAATCCCGGGAGGACCACCAACGGCGAAAGCAGCTTACGAGAACGGCTCCGACGGTCAGGAACGAAGGCTAGGGG |
| BH-asv789 | CCGAGTGATGGCCGATATTATTGGGCCTAAAGCGTCCGTAGCTTGCCGAGTAAGTCCGTCGGGAAATCCACTCGCCTAACGAGTGGGCGTCCGGCGGAAACTGTTCGGCTTGGGACCGGAAGACCCAGGGGGTACGTCCGGGGTAGGAGTGAAATCCCGTAATCCCGGACGGACCACCGGTGGCGAAAGCGCCCTGGGAAGACGGATCCGACAGTGAGGGACGAAAGCTAGGGT |
| BH-asv793 | GCAAGCGTTGTCCGGAATTACTGGGCGTAAAGCGCACGCAGGCGGATTAAAAAGTCAGCCGTAAAATGCACGGGCTCAACCTGTGTTAGTCAGCTGATACTATTAGTCTAGAGTATGTGAGAGGGAAGTGGAATTCCCGGTGTAGCGGTGAAATGCGTAGATATCGGGAGGAACACCAGTGGCGAAGGCGGCTTCCTGGCACAAAACTGACGCTCATGTGCGAAAGCTAGGGC |
| BH-asv796 | GCGAGCGTTAATCGGAATTACTGGGCGTAAAGCGTGCGCAGGCGGTCTTGCAAGACAGGTGTGAAATCCCCGGGCTTAACCTGGGAACTGCACTTGTGACTGCAAGGCTGGAGTACGGCAGAGGGGGATGGAATTCCGCGTGTAGCAGTGAAATGCGTAGATATGCGGAGGAACACCGATGGCGAAGGCAATCCCCTGGGCCTGTACTGACGCTCATGCACGAAAGCGTGGGG |
| BH-asv797 | GCTAGCGTTGTTCGGAATTACTGGGCGTAAAGCGCACGTAGGCGGCTTTGTAAGTTAGAGGTGAAAGCCTGGAGCTCAACTCCAGAACTGCCTTTAAGACTGCATCGCTTGAATCCAGGAGAGGTGAGTGGAATTCCGAGTGTAGAGGTGAAATTCGTAGATATTCGGAAGAACACCAGTGGCGAAGGCGGCTCACTGGACTGGTATTGACGCTGAGGTGCGAAAGCGTGGGG |
| BH-asv798 | GCTAGCGTTGTTCGGAATTACTGGGCGTAAAGCGCACGTAGGCGGCTGTTCAAGTCAGAGGTGAAAGCCCGGAGCTCAACTTCGGAACTGCCTTTGAAACTAGACAGCTAGAATCCTGGAGAGGTGAGTGGAATTCCGAGTGTAGAGGTGAAATTCGTAGATATTCGGAAGAACACCAGTGGCGAAGGCGGCTCACTGGACAGGTATTGACGCTGAGGTGCGAAAGCGTGGGG |
| BH-asv799 | GCAAGCGTTGTCCGGATTTATTGGGCGTAAAGCGAGCGCAGGCGGAAGAATAAGTCTGATGTGAAAGCCCTCGGCTTAACCGAGGAACTGCATCGGAAACTGTTTTTCTTGAGTGCAGAAGAGGAGAGTGGAACTCCATGTGTAGCGGTGGAATGCGTAGATATATGGAAGAACACCAGTGGCGAAGGCGGCTCTCTGGTCTGCAACTGACGCTGAGGCTCGAAAGCATGGGT |
| BH-asv800 | GCGAGCGTTGTCCGGAATTATTGGGCGTAAAGGGTACGTAGGCGGTTTAACAAGTCAGATGTTAAAACTTGAGGCTTAACCTCATGTTGCATTTGAAACTGTCAAACTTGAGTAGTGGAGAGGAAAGTGGAATTCCTAGTGTAGCGGTGAAATGCGTAGATATTAGGAGGAATACCAGTAGCGAAGGCGACTTTCTGGACACAAACTGACGCTGAGGTACGAAAGCGTGGGG |
| BH-asv801 | CCGAGTAGTCGGGACGATTATTGGGCCTAAAGCATCCGTAGCCTGTTCTGCAAGTCCTCCGTTAAATCCACCCGCTTAACGGATGGGCTGCGGGGGATACTGCAGAGCTAGGAGGCGGGAGAGGCAAACGGTACTCAGTGGGTAGGGGTAAAATCCATTGATCTACTGAAGACCACCAGTGGCGAAGGCGGTTTGCCAGAACGCGCTCGACGGTGAGGGATGAAAGCTGGGGG |
| BH-asv804 | TCGAGTGATGGCCGATCTTATTGGGCCTAAAGCGTTCGTAGCTGGCTAGTCAAGTCCGTTGGGAAATCGACGTGCCTAACACGTCGGCGTCCAGCGGAAACTGACTTGCTTGGGACCGGAAGACTCGAGGGGTACGTCCGGGGTAGGAGTGAAATCCTGTAATCCTGGACGGACCACCAATGGCGAAAGCACCTCGAGAGCACGGATCCGACAGTGAGGAACGAAAGCTGGGGT |
| BH-asv805 | CCGAGTGGTGGCCACGTTTATTGGGCTTAAAGTGTCCGTAGCCGGCCCTGTAAGTCCGCTGTGAAATCTGGCAGCTTAACTGTCAGGCGCGCGGTGGATACTACAGGGCTTGGGACCGGAAGGCGCCGAGGCTATTCCTGGGGTAGGGGTGAAATCCAAAAATCCTAGGAGGAGCACCAGTGGCGAAAGCTCTCGGCGATGACGGGTCCGACGGTCAGGGACGAAAGCTAGGGG |
| BH-asv806 | GCTAGCGTTGTTCGGAATTACTGGGCGTAAAGCGCACGTAGGCGGCTTTTTAAGTCAGGGGTGAAATCCCGGGGCTCAACCCCGGAACTGCCCTTGAAACTGGGAAGCTAGAATCTTGGAGAGGCGAGTGGAATTCCGAGTGTAGAGGTGAAATTCGTAGATATTCGGAAGAACACCAGTGGCGAAGGCGACTCGCTGGACAAGTATTGACGCTGAGGTGCGAAAGCGTGGGG |
| BH-asv808 | CCAAGTGGTAGCCATTTTTATTGGGCTTAAAGTGTTCGTAGCCGGCCATGTAAGTCCGCTGTGAAACCCAGGCGCTCAACGTCTGGATGCGCAGCGGATACTGCATAGCTAGGGACCGGACGGCGTCGAGGCTATTCCCGGGGTAGGAGTGAAATCCAAAAATCCCGGGAGGAGCACCTGTGGCGAAAGCGCTCGACGAGTACGGATCCGACGGTGAGGAACGAAAGCCAGGGG |
| BH-asv809 | GCAAGCGTTGTCCGGAATTATTGGGCGTAAAGCGCGCGCAGGCGGCCTTTTAAGTCTGATGTGAAAGCCCACGGCTTAACCGTGGAAGGTCATTGGAAACTGGAAGGCTTGAGTACAGAAGAGGAAAGCGGAATTCCACGTGTAGCGGTGAAATGCGTAGAGATGTGGAGGAACACCAGTGGCGAAGGCGGCTTTCTGGTCTGTAACTGACGCTGAGGCGCGAAAGCATGGGG |
| BH-asv812 | GCAAGCGTTAATCGGAATAACTGGGCGTAAAGCGCACGTAGGCGGCGCGGTCAGTCAGATGTGAAATCCCCGGGCTCAACCTGGGAACTGCATTTGATACTGCCGTGCTCGAGTCTGAGAGAGGGGGGTGGAATTCCAGGTGTAGCGGTGAAATGCGTAGATATCTGGAGGAACACCAGTGGCGAAGGCGGCCCCCTGGCTCAAGACTGACGCTGAGGTGCGAAAGCGTGGGG |
| BH-asv813 | GCGAGCGTTAATCGGAATTACTGGGCGTAAAGCGTGCGCAGGCGGTTTTTTAAGATAGGCGTGAAATCCCCGGGCTCAACCTGGGAACTGCGCTTATGACTGGAAGACTCGAGTATGGCAGAGGGGGGTGGAATTCCACGTGTAGCAGTGAAATGCGTAGAGATGTGGAGGAACACCGATGGCGAAGGCAGCCCCCTGGGCCAATACTGACGCTCATGCACGAAAGCGTGGGG |
| BH-asv814 | GCAAGCGTTGTTCGGAATTATTGGGCGTAAAGCGCGTGTAGGCGGCTTCGCAAGTCGGGTGTGAAATCCCCCAGCTCAACTGGGGAAGTGCGCTCGAAACTGCGGAGCTCGAGTACCGGAGAGGATGGCGGAATTCCCCAAGTAGAGGTGAAATTCGTAGATATGGGGAGGAACACCGGTGGCGAAGGCGGCCATCTGGACGGATACTGACGCTGAGACGCGAAAGCGTGGGG |
| BH-asv815 | GCGAGCGTTGTCCGGAATTATTGGTCGTAAAGAGCTCGTAGGCGGTTTGTCGCGTCGACTGTGAAAACCTGGGGCTTAACTCTGGGCTTGCAGTCGATACGGGCAGACTTGAGTTCGGTAGGGGAGACTGGAATTCCTGGTGTAGCGGTGAAATGCGCAGATATCAGGAGGAACACCGGTGGCGAAGGCGGGTCTCTGGGCCGATACTGACGCTGAGGAGCGAAAGCGTGGGG |
| BH-asv822 | GCTAGCGTTGTTCGGAATTACTGGGCGTAAAGCGCGCGTAGGCGGATTGGAAAGTTGGGGGTGAAATCCCGGGGCTCAACCCCGGAACTGCCTCCAAAACTATCAGTCTGGAGTTCGAGAGAGGTGAGTGGAATTGCGAGTGTAGAGGTGAAATTCGTAGATATTCGCAGGAACACCAGTGGCGAAGGCGGCTCACTGGCTCGATACTGACGCTGAGGTGCGAAAGTGTGGGG |
| BH-asv824 | CCGGGCGTTATCCGGATTTATTGGGTTTAAAGGGAGCGTAGGCCGGAGATTAAGCGTGTTGTGAAATGTAGACGCTCAACGTCTGCACTGCAGCGCGAACTGGTTTCCTTGAGTACGCACAAAGTGGGCGGAATTCGTGGTGTAGCGGTGAAATGCTTAGATATCACGAAGAACTCCGATTGCGAAGGCAGCTCACTGGAGCGCAACTGACGCTGAAGCTCGAAAGTGCGGGT |

| | |
|---|---|
| BH-asv826 | GCGAGCGTTATCCGGATTCATTGGGTTTAAAGGGTGCGTAGGCGGTTTCTTAAGTCAGTGGTGAAATCCTGCGGCTCAACCGTAGAACTGCCATTGATACTGAGGAACTTGAATTTAGTTGAGGTAGGCGGAATGAGTAGTGTAGCGGTGAAATGCTTAGATATTACTCAGAACACCGATTGCGAAGGCAGCTTACCAAGCTACGATTGACGCTGATGCACGAAAGCGTGGGG |
| BH-asv828 | GCAAGCGTTATCCGGATTTACTGGGTGTAAAGGGAGCGTAGACGGTGTGGCAAGTCTGATGTGAAAGGCAGGGGCTCAACTCCTGGACTGCATTGGAAACTGCCAGACTTGAGTGCCGGAGGGGTAAGCGGAATTCCTAGTGTAGCGGTGAAATGCGTAGATATTAGGAGGAACACCAGTGGCGAAGGCGGCTTACTGGACGGTAACTGACGTTGAGGCTCGAAAGCGTGGGG |
| BH-asv829 | CCGAGCGTTGTCCGGATTCATTGGGCGTAAAGAGCTCGTAGGCGGCCACTCAAGTCGAATGTGAAACCTCGAGGCTTAACCTCGAGCCTGCATTCGATACTGGGCGGCTAGAGTCTGGTAGGGGGATCTGGAATTCCTGGTGTAGCGGTGGAATGCGCAGATATCAGGAGGAACACCGGTGGCGAAGGCGGGATCCTGGGCCAGTACTGACGCTGAGGAGCGAAAGCGTGGGG |
| BH-asv830 | GCAAGCGTTGTCCGGAATTATTGGGCGTAAAGCGCGCGCAGGCGGTTTCTTAAGTCTGATGTGAAAGCCCACGGCTCAACCGTGGAGGGTCATTGGAAACTGGGGAACTTGAGTGCAGAAGAGGAGAGTGGAATTCCACGTGTAGCGGTGAAATGCGTAGAGATGTGGAGGAACACCAGTGGCGAAGGCGACTCTCTGGTCTGTAACTGACGCTGAGGCGCGAAAGCGTGGGG |
| BH-asv834 | CCGAGCGTTATCCGGAATTATTGGGTTTAAAGGGTCCGTAGGCGGGCCGGTAAGTCAGGGGTGAAAGTCCGTAGCTCAACTATGGAACTGCCTTTGATACTGCTGGTCTTGAGTTATGGTGAAGTTGGCGGAATATGTAGTGTAGCGGTGAAATGCATAGATATTACATAGAACACCGATTGCGAAGGCAGCTGGCTAACCATATACTGACGCTGATGGACGAAAGCGTGGGG |
| BH-asv835 | GCAAGCGTTATCCGGAATTACTGGGCGTAAAGGGTGCGTAGGCGGTTTATCAAGTCAGGGGTGAAAGGCTACGGCTCAACCGTAGTGAGCCTCTGAAACTGGTGAACTTGAGTGTAGGAGAGGAAAGTGGAATTCCCAGTGTAGCGGTGAAATGCGTAGATATTGGGAGGAACACCAGTGGCGAAGGCGACTTTCTGGACTACAACTGACGCTGAGGCACGAAAGCGTGGGG |
| BH-asv837 | GCAAGCGTTATCCGGATTTACTGGGTGTAAAGGGAGCGTAGACGGCATCACAAGTCAGAAGTGAAAATCCGGGGCTCAACCCCGGAACTGCTTTTGAAACTGTGGAGCTGGAGTGCAGGAGAGGTAAGCGGAATTCCTAGTGTAGCGGTGAAATGCGTAGATATTAGGAGGAACACCAGTGGCGAAGGCGGCTTACTGGACTGTAACTGACGTTGAGGCTCGAAAGCGTGGGG |
| BH-asv838 | GCAAACGTTGTTCGGAATTACTGGGCGTAAAGCGCGTGTAGGCTGCTCGGAAAGTCGGATGTGAAAGCCCTGGGCTCAACCTAGGAAGTGCATTCGAAACTTCCGAGCTGGAGTGCTGGAGAGGAAGGCGGAATTCTCGGTGTAGAGGTGAAATTCGTAGATATCGAGAGGAACACCGGTGGCGAAGGCGGCCTTCTGGACAGCTACTGACGCTGAGACGCGAAAGCGTGGGG |
| BH-asv840 | GCGAGCGTTAATCGGAATTACTGGGCGTAAAGCGTGCGCAGGCGGTTTCGTAAGACAGAGGTGAAATCCCCGGGCTCAACCTGGGAACTGCCTTTGTGACTGCGAGGCTAGAGTATGGCAGAGGGGGGTGGAATTCCACGTGTAGCAGTGAAATGCGTAGAGATGTGGAGGAACACCGATGGCGAAGGCAGCCCCCTGGGCCAATACTGACGCTCATGCACGAAAGCGTGGGG |
| BH-asv841 | GCAAGCGTTAATCGGAATTACTGGGCGTAAAGCGTACGTAGGCGGCTTGTTAAGCAAGATGTGAAAGCCCCGGGCTCAACCTGGGAATGGCATTTTGAACTGGCAGGCTCGAGTTCTGAAGAGGGTGGTAGAATTTCCTGTGTAGCGGTGAAATGCGTAGATATAGGAAGGAATACCGGTGGCGAAGGCGGCCACCTGGTCAGAAACTGACGCTGAGGTACGAAAGCGTGGGG |
| BH-asv842 | GCAAGCGTTATCCGGATTCATTGGGTTTAAAGGGTGCGTAGGCGGCTTTTTAAGTCAGTGGTGAAATCTTGCCGCTTAACGGTAAAATTGCCATTGATACTGAAGAGCTTGAGTACAATTGAGGTAAGCGGAATGTGTAGTGTAGCGGTGAAATGCTTAGATATTACACAGAACACCAATTGCGAAGGCAGCTTACTAAACTGTCACTGACGCTGATGCACGAAAGCGTGGGG |
| BH-asv843 | GCAAGCGTTAATCGGAATTACTGGGCGTAAAGGGCGCGTAGGTGGTTTGTTAAGTCGGGTGTGAAAGCCCCGGGCTCAACCTGGGAACTGCATCCGAGACTGGCTGACTTGAGTATGGTAGAGGGAAGCGGAATTCCCGGTGTAGCGGTGAAATGCGTAGATATCGGGAGGAACATCAGTGGCGAAGGCGGCTTCCTGGACCAATACTGACACTGAGGCGCGAAAGCGTGGGG |
| BH-asv845 | GCTAGCGTTGCTCGGAATTACTGGGCGTAAAGGGAGCGTAGGCGGACAGTTAAGTTAGAGGTGAAAGCCCAGGGCTCAACCTTGGAATTGCCTTTGATACTGGCTGTCTTGAGTATGGGAGAGGTGTGTGGAACTCCGAGTGTAGAGGTGAAATTCGTAGATATTCGGAAGAACACCAGTGGCGAAGGCGACACACTGGCCCATTACTGACGCTGAGGCTCGAAAGCGTGGGG |
| BH-asv847 | CCGAGTGGTGGCCGATGTTACTGGGCTTAAAGCGTTCGTAGCCGGCCATGTAAGTCCGCTGTGAAATCCGCGCGCTCAACGTGCGGCCGTGCAGCGGATACTGCATGGCTAGGGACCGGACGGCGTCGAGGCTATTCCAGGGGTAGGAGTGAAATCCCGAAATCCCTGGAGGAGCACCTGTGGCGAAAGCGCTCGGCGAGTACGGATCCGACGGTGAGGGACGAAAGCCAGGGG |
| BH-asv848 | CCAAGTGATGGCCGATATTATTGGGCCTAAAGCGTTCGTAGCTGGCCAGACAAGTCCGTTGGGAAATCCGCATGCTTAACGTGCGGGCGTCCAGCGGAAACTGTCCGGCTTGGGACCGGAAGACCCGAGGGGTACGTCCGGGGTAGGAGTGAAATCCTGTAATCTTGGACGGACCACCTGTGGCGAAAGCGCCTCGGGAAGACGGATCCGACAGTGAGGAACGAAAGCCAGGGT |
| BH-asv849 | GCTAGCGTTATTCGGATTTACTGGGCATAAAGCGCGCGTAGGCGGCTTGGTAAGTCGGAGGTGAAAGCCCGCAGCTCAACTGCGGAACTGCCCCCGATACTGTCGAGCTTGAGTGCAGGAGGGGTGACTGGAACTCCAGGTGTAGCGGTGAAATGCGTAGATATCTGGAGGAACACCGGAGGCGAAGGCGGGTCACTGGACTGCAACTGACGCTGAGGCGCGAAAGCTAGGGG |
| BH-asv851 | GCAAGCGTTGTTCGGAATTACTGGGCGTAAAGGGCGCGTAGGCGGTTTGGTAAGTCAGATGTGAAATTCACCGGCTCAACCGGTTGAACTGCATCCGATACTGCCAGACTTGAGATCAGTAGAGGAGAGTGGAATTCCCAGTGTAGCGGTGGAATGCGTAGATATTGGGAAGAACACCGGTGGCGAAGGCGGCTCTCTGGACTGATTCTGACGCTGAGGCGCGAAAGCTAGGGG |
| BH-asv852 | CCGAGCGTTGTCCGGATTTATTGGGCGTAAAGCGAGCGCAGGCGGTTTGATAAGTCTGAAGTAAAAGGCTGTGGCTTAACCATAGTATGCTTTGGAAACTGTCAAACTTGAGTGCAGAAGGGGAGAGTGGAATTCCATGTGTAGCGGTGAAATGCGTAGATATATGGAGGAACACCGGTGGCGAAAGCGGCTCTCTGGTCTGTAACTGACGCTGAGGCTCGAAAGCGTGGGG |
| BH-asv853 | GCAAGCGTTGTCCGGAATTATTGGGCGTAAAGAGCTCGTAGGCGGTTTGTTGCGTCGGCTGTGAAATCCCGAGGCTCAACCTCGGGTCTGCAGTCGATACGAGCAAACTAGAGTGTTGCAGGGGAGACTGGAATTCCTGGTGTAGCGGTGAAATGCGCAGATATCAGGAGGAACACCGGTGGCGAAGGCGGGTCTCTGGGCAACAACTGACGCTGAGGAGCGAAAGCGTGGGG |
| BH-asv855 | CCGAGCGTTGTCCGGATTTATTGGGCGTAAAGCGAGCGCAGGCGGTTTAATAAGTCTGAAGTTAAAGGCAGTGGCTTAACCATTGTTCGCTTTGGAAACTGTTAGACTTGAGTGCAGAAGGGGAGAGTGGAATTCCATGTGTAGCGGTGAAATGCGTAGATATATGGAGGAACACCGGTGGCGAAAGCGGCTCTCTGGTCTGTAACTGACGCTGAGGCTCGAAAGCGTGGGG |
| BH-asv858 | GCGAACGTTACTCGGAATCACTGGGCTTACAGGGTGCGTAGGTGGCTTGTTAAGCAGGGTGTGAAATCCCTCGGCTCAACCGAGGAACTGCGCTCTGAACTGGCGAGCTAGAGTATGCTAGAGGTGAATGGAACATCCAGTGGAGCGGTGAAATGTGTTGATATTGGATGGAACACCAGAGGCGAAGGCGATTCACTGGGGCATAACTGACACTGAGGCACGAAAGCCAGGGG |
| BH-asv859 | GCGAGCGTTGTTCGGAATTACTGGGCGTAAAGGGCATGTAGGCGGTTTGGTAAGCCTGACGTGAAAGGCTATGGCTCAACCATGGCATGGCGTTGGGAACTGCTAGACTTGAATCACGGAGAGGGAGCTGGAATTCCAGGTGTAGGGGTGAAATCTGTAGATATCTGGAAGAACACCAGTGGCGAAGGCGAGCTCCTGGCCGATGATTGACGCTGAGGTGCGAAAGTGTGGGG |
| BH-asv860 | TCAAGTGATGACCGATTTTATTGGGCCTAAAGCGTCCGTAGCTGGCCGCGCAAGTCCGTCGGGAAATCCACCCGCTTAACGGGTGGGCGTCCGGCGGAAACTGTGCGGCTTGGGACAGGAAGGTTCGAGGGGTACGTTCGGGGTAGGAGTGAAATCCCGTAATCCTGAACGGACCACCGATGGCGAAAGCACCTCAAGAGAACGGATCCGACAGTGAGGGACGAAAGCTAGGGT |
| BH-asv861 | GCTAGCGTTGTTCGGAATCACTGGGCGTAAAGGGCGCGTAGGCGGCCGAGGCAGTTGGATGTGAAAGCCCGGGGCTCAACCCCGGAGGTGCATCCGATACTCCTTGGCTGGAGGCCGGAAGAGGAGGGTGGAATACCCAGTGTAGAGGTGAAATTCGTAGATATTGGGTGGAACACCGGTGGCGAAAGCGGCTCTCTGGTCCGGTCCTGACGCTGAGGCGCGAAAGCGTGGGG |
| BH-asv862 | GCTAGCGTTAATCGGATTTACTGGGCGTAAAGCGTGCGCAGGCGGTTTGTTAAGTCTGTTGTGAAATCCCCGAGCTCAACTTGGGAATTGCAATGGATACTGGCAAGCTGGAGTATGGTAGAGGAAGGTGGAATTCCCGGTGTAGCAGTGAAATGCGTAGAGATCGGGAGGAACACCAGTGGCGAAGGCGGCCTTCTGGACCAATACTGACGCTCATGCACGAAAGCGTGGGG |
| BH-asv865 | GCAAGCGTTGTCCGGAATCACTGGGCGTAAAGGGTCCGCAGGCGGATCAGCAACTCAGGCGTGAAACCCCATGGCTTAACCATGGAGGCGCGTCTGAAAGTGTTGATCTCGAGGACAGGAAGGGAGAGTGGAACTCCGGGTGGAGCGGTGAAATGCGCAGATATCCGGAGGAACACCGGAGGCGAAAGCGACTCTCTAGCCTGTACCTGACGCTGAGGGACCAAAGCCAGGGT |
| BH-asv867 | CCAAGTGGTCACCGATATTATTGGGCCTAAAGCGTCCGTAGCCGGCCTGGTAAATGCCTGGGCAAATCGATGGGCTCAACCCATCGAAATCCGGGCAGACTGCCAGGCTTGGGACCGGGAGAGGCCAGAGGTACTCCGGGGGTAGAGGTGAAATTCCGTAATCCCCGGGGGACCACCAGTGGTGAAGACGTCTGGCTTGAACGGCTCCGACGGTGAGGGACGAAGCCCTGGGG |
| BH-asv868 | GCAAGCGTTGTCCGGAATCACTGGGCGTAAAGGGTCCGCAGGCGGCCTGGCAACTCAGGCGTGAAATCCCACGGCTTAACCGTGGAACTGCGTCTGAAAGTGTCGGGCTCGAGGGCAGGAAGGGAGAGTGGAACTCCGGGTGGAGCGGTGAAATGCGCAGATATCCGGAGGAACACCGGAGGCGAAAGCGACTCTCTAGCCTGCTCCTGACGCTGAGGGACCAAAGCCAGGGT |
| BH-asv871 | CCTAACGTTGTTCGGATTCACTGGGCTTAAAGGGCGCGTAGGCGGGTGAGCAAGTCCGGGGTGAAATCTTTCGGCTTAACCGGAAAATGGCCTTGGATACTGCTCGTCTCGAGGGAGGTAGGGGTCTGCGGAACTTCCGGTGGAGCGGTGAAATGCGTTGATATCGGAAGGAACGCCGGTGGCGAAAGCGGCGGACTGGACCTCTTCTGACGCTGAGGCGCGAAAGCTAGGGT |
| BH-asv872 | GCAAGCGTTAATCGGAATTACTGGGCGTAAAGCGCACGTAGGTGGCCAAACAAGTTAGATGTGAAAGCCCTGGGCTTAACCTAGGAAAGTCATCTAAGACTGTGTGGCTAGAGTACAGTAGAGGAAAGTAGAATTTCCGGTGTAGCGGTGAAATGCGTAGATATCGGAAGGAATACCAGTGGCGAAGGCGACTTTCTGGACTGAAACTGACACTGAGGTGCGAAAGCGTGGGG |
| BH-asv873 | GCAAGCGTTACTCGGAATCACTGGGCGTAAAGAGCATGTAGGCGGATAGATAAGTTGGAAGTGAAATCCACTGGCTTAACCAGTGAACTGCTTCCAAAACTGTCTATCTAGAATATGGGAGAGGTAGATGGAATTTCTGGTGTAGGGGTAAAATCCGTAGAGATCAGAAGGAATACCGATTGCGAAGGCGATCTACTGGAACATTATTGACGCTCAGATGCGAAAGCGTGGGG |
| BH-asv876 | GCTAGCGTTGTTCGGAATTACTGGGCGTAAAGCGCACGTAGGCGGCGATTTAAGTCAGGGGTGAAAGCCCAGTGCTCAACACTGGAACTGCCCTTGAGACTGGATTGCTTGAATCCGGGAGAGGTGAGTGGAATTCCGAGTGTAGAGGTGAAATTCGTAGATATTCGGAAGAACACCAGTGGCGAAGGCGGCTCACTGGACCGGCATTGACGCTGAGGTGCGAAAGCGTGGGG |
| BH-asv878 | GCAAGCGTTGCTCGGAATTACTGGGCGTAAAGGGTCCGCAGGTGGCCTCGTAAGTTGAATGTGAAATCTCAGGGCTTAACCCTGAAACTGCATCCAATACTGCGGGGCTTGAGTCCAAGAGAGGTTGGCGGAATTCCCGGTGTAGCGGTGAAATGCGTAGATATCGGGAGGAACACCAGTGGCGAAGGCGGCCAACTGGCTTGGAACTGACACTCAGGGACGAAAGCGTGGGT |
| BH-asv880 | GCAAGCGTTGTCCGGAATCACTGGGCGTAAAGGGTCCGCAGGCGGATCAGCAACTCAGGCGTGAAATCCCATAGCTTAACTATGGAAGTGCGTCTGAAAGTGTTGATCTCGAGGGCAGGAAGGGAGAGTGGAACTCCGGGTGGAGCGGTGAAATGCGCAGATATCCGGAGGAACACCGGAGGCGAAAGCGACTCTCTAGCCTGTTCCTGACGCTGAGGGACCAAAGCCAGGGT |
| BH-asv881 | CCAAGCGTTGTCCGGAATCACTGGGCGTAAAGGGTGTGCAGGTGGGGCAGCAAGTCAGGGGTGAAAGCCCGCGGCTTAACCGCGGAAGTGCCCTTGAAACTGCTGCTCTTGAGTCCCGGAGAGGCTGCCGGAATTCGTGGTGTAGCGGTGAAATGCGTAGATATCACGAGGAACACCAGAGGCGAAAGCGGGCAGCTGGACGGGTACTGACACTGAGGCACGAAAGCGTGGGG |
| BH-asv882 | TCGAGTGATGGCCGATATTATTGGGCCTAAAGCGTTCGTAGCTGGCCGAGCAAGTCTGTCGGGAAATCTGTTCGCTCAACGAGCAGGCGTCCGGCAGAAACTGCCCGGCTTGGGACCGGAAGACTCGAGGAGTACGTCTGGGGTAGGAGTGAAATCCCGTAATCCGAGACGGACTACCGATGGCGAAAGCACCTCGAGAGCACGGATCCGACAGTGAGGAACGAAAGCTGGGGT |
| BH-asv888 | GCGAGCGTTATCCGGATTTATTGGGTTTAAAGGGTGCGTAGGCGGATTAATAAGTCAGTGGTGAAATCCTAGGGCTCAACTTTAGAATTGCCATTGATACTGTTAATCTTGAATGCAGTTGAGGTATGCGGAATGTGTAATGTAGCGGTGAAATGCTTAGATATTACACAGAACACCGATTGCGAAGGCAGCATGCTAAGCTGTGATTGACGCTGATGCACGAAAGCGTGGGG |
| BH-asv889 | GCAAGCGTTAATCGGAATTACTGGGCGTAAAGCGCGCGTAGGCGGCTTGTTAAGTCGGATGTGAAATCCCCGGGCTCAACCTGGGAATTGCCTCCGATACTGGCGAGCTGGAGTACGGGAGAGGAGAGTAGAATTCCAGGTGTAGCGGTGAAATGCGTAGATATCTGGAGGAATACCGGTGGCGAAGGCGGCTCTCTGGCCCGATACTGACGCTGAGGTGCGAAAGCGTGGGG |
| BH-asv891 | GCAAGCGTTGTCCGGAATTATTGGGCGTAAAGCGCGCGCAGGCGGTCCTTTAAGTCTGATGTGAAAGCCCACGGCTCAACCGTGGAGGGTCATTGGAAACTGGGGGACTTGAGTGCAGAAGAGGAAAGCGGAATCCCACGTGTAGCGGTGAAATGCGTAGAGATGTGGAGGAACACCAGTGGCGAAGGCGGCTTTCTGGTCTGTAACTGACGCTGAGGCGCGAAAGCGTGGGG |
| BH-asv892 | GCAAGCGTTGTTCGGAATCATTGGGCGTAAAGCGCGCGCAGGCGGATCAGCAAGTCAGGTGTGAAATCTCGGGGCTCAACCCCGAAACTGCGCCTGAAACTGCTAGTCTAGAATGTCGGAGGGGGCAGGGGAATTTCACGTGTAGGGGTAAAATCCGTAGAGATGTGAAGGAACACCGGAGGCGAAGGCGCCTGCCTGGACGACTATTGACGCTGAGGCGCGAAAGCGTGGGG |
| BH-asv893 | GCAAGCGTTGTCCGGAATTATTGGGCGTAAAGCGCGCGCAGGCGGTTCCTTAAGTCTGATGTGAAAGCCCACGGCTCAACCGTGGAGGGTCATTGGAAACTGGGGAACTTGAGTGCAGAAGAGGAGAGCGGAATTCCACGTGTAGCGGTGAAATGCGTAGAGATGTGGAGGAACACCAGTGGCGAAGGCGGCTCTCTGGTCTGTAACTGACGCTGAGGCGCGAAAGCGTGGGG |
| BH-asv894 | GCTAGCGTTGTTCGGAATTGCTGGGCGTAAAGGGCGCGTAGGCGGTGTGCCAAGTCAGGCGTGAAAGCCCCGGGCTCAACCTGGGAACCGCGCTTGAGACTGGCACGCTCGAGTTCGGGAGAGGATGGTGGAATTCCCAGTGTAGAGGTGAAATTCGTAGATATTGGGAAGAACACCGATGGCGAAGGCAGCCATCTGGACCGGCACTGACGCTGAGGCGCGAAAGCGTGGGG |
| BH-asv895 | ACAAGTGGTGGTCACTTTTATTGAGCCTAAAGCGTTCGTAGCCGGTTTTGTAAATCTTCAGGTAAATCCTGAAGCCTAACTTCAGAAGGCCTGAAGAGACTGCAAAACTTGAGATCGGGTGAGGTTAAACGTACTTTCAGGGTAGGGGTAAAATCCTGTAATCCTGGAAGGACGACCGGTGGCGAAGGCGTTTAACTAGAACGAATCTGACGGTGAGGAACGAAGGCTAGGGT |
| BH-asv897 | GCAAGCGTTGTTCGGAATCATTGGGCGTAAAGCGCGCGCAGGTGGCCAGACAAGTCCGGTGTGAAAGCCCGGGGCTTAACCCCGGAGGTGCATCGGAAACTGTGTGGCTTGAGTCCGGGAGGGGCCAGTGGAATTCGTGGTGTAGGGGTGAAATCCGTAGATATCACGAAGAACACCGACGGCGAAAGCAACTGGCTGGACCGGGACTGACGCTGAGGCGCGAAAGCGTAGGT |
| BH-asv898 | CCAAGTGGTAGCCATTTTTATTGGGCTTAAAGTGTTCGTAGCCGGCCATGTAAGTCCGCTGTGAAACCCAGGCGCTCAACGTCTGGATGTGCAGCGGATACTGCATAGCTAGGGACCGGACGGCGTCGAGGCTATTCCCGGGGTAGGAGTGAAATCCAAAAATCCCGGGAGGAGCACCTGTGGCGAAAGCGCTCGACGAGTACGGATCCGACGGTGAGGAACGAAAGCCAGGGG |

| | |
|---|---|
| BH-asv899 | CCAAGTGGTGGTCACGATTATTGGGCCTAAAGCGTCCGTAGCCGGCCGGATGGGTCCTCGGTGAAATCCCACGTCAAAGCGTGGGGCTGGCCGGGGAAACTATCTGGCTTGGGACCGGGAGAGGTCGGGGCTACTCCTGGGGTAGTGGTAAAATACTGTAATCCTAGGAGGAGCACCTGTGGCGAAAGCGCCCGGCTGGAACGGCTCCGACGGTGAGGGACGAAAGCTAGGGG |
| BH-asv901 | GCAAGCGTTGCCCGGAATCACTGGGCGTAAAGGGTCCGCAGGCGGCCTGGCAACTCAGGCGTGAAAGCCCATGGCTTAACCATGGAATTGCGTTTGAAAGTGTCAGGCTCGAGTACGGGAAGGGGGAGTGGAACTCCGGGTGGAGCGGTGAAATGCGCAGATATCCGGAGGAACACCGGAGGCGAAAGCGACTCCCTAGCCCGTTACTGACGCTGAGGGACAAAAGCCAGGGT |
| BH-asv903 | GCAAGCGTTAATCGGAATTACTGGGCGTAAAGCGTGCGCAGGCGGTTTTGTAAGACAGGTGTGAAATCCCCGAGCTCAACTTGGGAACTGCACTTGTGACTGCAAGACTAGAGTGTGTCAGAGGGAGGTAGAATTCCACGTGTAGCAGTGAAATGCGTAGAGATGTGGAGGAATACCGGTGGCGAAGGCGGCCTCCTGGGATAACACTGACGCTCATGTACGAAAGCGTGGGG |
| BH-asv904 | GCAAGCGTTAATCGGAATTACTGGGCGTAAAGCGCGCGTAGGCGGTTGTGTAAGTTGGATGTGAAATCCCCGGGCTTAACCTGGGCACTGCATTCAAAACTGCACGGCTAGAGTATAGGAGAGGAAGGTAGAATTCCAGGTGTAGCGGTGAAATGCGTAGAGATCTGGAGGAATACCGATGGCGAAGGCAGCCTTCTGGCCTAATACTGACGCTGAGGTGCGAAAGCATGGGG |
| BH-asv906 | GCAAGCGTTACTCGGAATTACTGGGCGTAAAGCGTGCGTAGGTGGTTGTTTAAGTCTGTCGTGAAAGCCCTGGGCTCAACCTGGGAATTGCGATGGATACTGGGCGACTAGAATGTGGTAGAGGGTGGCGGAATTCCCGGTGTAGCAGTGAAATGCGTAGAGATCGGGAGGAACATCCGTGGCGAAGGCGGCCACCTGGACCAACATTGACACTGAGGCACGAAAGCGTGGGG |
| BH-asv907 | GCTAGCGTTGCTCGGAATTACTGGGCGTAAAGGGCGCGTAGGCGGACATTTAAGTCAGGGGTGAAATCCCAGAGCTCAACTCTGGAACTGCCTTTGATACTGGGTGTCTTGAGTGTGAGAGAGGTATGTGGAACTCCGAGTGTAGAGGTGAAATTCGTAGATATTCGGAAGAACACCAGTGGCGAAGGCGACATACTGGCTCATTACTGACGCTGAGGCGCGAAAGCGTGGGG |
| BH-asv908 | GCAAGCGTTATCCGGAATTATTGGGCGTAAAGCGCGCGTAGGCGGTTCGTTAAGTCTGATGTGAAAGCCCCCGGCTCAACCGGGGAGGGTCATTGGAAACTGGCGGACTTGAGTGCAGAAGAGGAGAGTGGAATTCCATGTGTAGCGGTGAAATGCGCAGAGATATGGAGGAACACCAGTGGCGAAGGCGGCTCTCTGGTCTGCAACTGACGCTGAGGTGCGAAAGCGTGGGG |
| BH-asv909 | GCTAGCGTTGCTCAGAATCACTGGGCGTAAAGGGCGCGTAGGCGGATCTTTAAGTCAGGGGTGAAAGCCCAAGGCTCAACCTTGGAATTGCCTTTGATACTGAGGATCTCGAGACCGGAAGAGTTAAGTGGAACTGCGAGTGTAGAGGTGAAATTCGTAGATATTCGCAAGAACACCAGTGGCGAAGGCGGCTTACTGGTCCGGTTCTGACGCTGAGGCGCGAAAGCGTGGGG |
| BH-asv910 | GCAAGCGTTGTTCGGAATCATTGGGCGTAAAGAGCGTGTAGGCGGCCAGACAGGTCCGTTGTGAAAACTCGAGGCTCAACCTCGAGACGTCGGTGGAAACCGTCTGGCTAGAGTCCGGAAGAGGAGAGTGGAATTCCTGGTGTAGCGGTGAAATGCGCAGATATCAGGAGGAACACCCGTGGCGAAGGCGGCTCTCTGGGACGGTACTGACGCTGAGACGCGAAAGCGTGGGG |
| BH-asv911 | CCAAGCGTTAGGCGGAATCACTGGGCTTAAAGCGTGTGTAGGCGGCTTCTAAGTACCTTGTGAAATCCCACGGCTCAACCGTGGAACTGCTTGGTATACTGGTTATCTTGAGCCACTTAGGGGCGACCGGAACAAATGGTGGAGCGGTGAAATGCGTAGATATCATTTGGAACGCCAATGGTGAAAACAGGTCGCTGGGAGTGTGCTGACGCTGAGGTACGAAAGCCAGGGG |
| BH-asv913 | GCGAGCGTTGTCCGGATTTATTGGGCGTAAAGAGTTCGTAGGTGGTTTGTTAAGTTTGGTGTTAAAGATCGGGGCTCAACCCTGGGACTGCACTGAATACTGGCAGACTCGAGTGTGGTAGAGGCTAGTGGAATTCCCAGTGTAGCGGTGAAATGCGTAGATATTGGGAAGAACACCGGTGGCGTAGGCGACTAGCTGGGCCATAACTGACGCTGAGGAACGAAAGCCAGGGG |
| BH-asv914 | GCGAGCGTTATCCGGATTTATTGGGTTTAAAGGGAGCGTAGATGGGCTTTTAAGTCAGTTGTGAAATTCTGCGGCTCAACCGTAGGACTGCAGTTGAAACTGGAGCCTTGAGTGCAGTTGAGGTGCATGGAATTCGTGGTGTAGCGGTGAAATGCTTAGATATCACGAAGAACTCCGATTGCGAAGGCAGTGCACCAAGCTGTAACTGACATTGATGCTCGAAAGTGTGGGT |
| BH-asv915 | GCAAGCGTTATCCGGAATTATTGGGCGTAAAGCGCGCGTAGGCGGTTTTTTAAGTCTGATGTGAAAGCCCACGGCTCAACCGTGGAGGGTCATTGGAAACTGTAAAACTTGAGTGCAGAAGAGGAAAGTGGAATTCCATGTGTAGCGGTGAAATGCGCAGAGATATGGAGGAACACCAGTGGCGAAGGCGACTTTCTGGTCTGTAACTGACGCTGATGTGCGAAAGCGTGGGG |
| BH-asv916 | GCTAGCGTTGTTCGGAATTACTGGGCGTAAAGCGTGCGCAGGTTGTGAGATCAGTTGGATGTGAAAGCCCGGGGCTTAACCTCGGATGTGCATCCAATACTGTCTCGCTGGAGGTTGGAAGAGGAGAGTGGAATTCCCAGTGTAGAGGTGAAATTCGTAGATATTGGGAAGAACACCAGTGGCGAAGGCGGCTCTCTGGTCCATACCTGACACTCATGCACGAAAGCGTGGGG |
| BH-asv917 | GCGAGCGTTGTTCGGATTTACTGGGCGTAAAGAGCACGTAGGCGGTTGGGTAAGCCTCTTGGGAAATCTCCCGGCTTAACCGGGAAAGGTCGAGGGGAACTACTCAGCTAGAGGACGGGAGAGGAGCGCGGAATTCCCGGTGTAGCGGTGAAATGCGTAGATATCGGGAAGAAGGCCGGTGGCGAAGGCGGCGCTCTGGAACGTACCTGACGCTGAGGTGCGAAAGCGTGGGG |
| BH-asv918 | GCTAGCGTTGTTCGGAATTACTGGGCGTAAAGCGCGCGTAGGCGGCTATTCAAGTCAGGGGTGAAATCCCGGGGCTCAACCTCGGAACTGCCTTTGAAACTGGATAGCTAGAATCCTGGAGAGGCGAGTGGAATTCCGAGTGTAGAGGTGAAATTCGTAGATATTCGGAAGAACACCAGTGGCGAAGGCGACTCGCTGGACAGGTATTGACGCTGAGGTGCGAAAGCGTGGGG |
| BH-asv919 | GCAAGCGTTATCCGGATTCATTGGGTTTAAAGGGTGCGCAGGTGGGAAAATAAGTCAGTGGTGAAATACTGCCGCTTAACGGTAGAATTGCCATTGATACTGTTTTTCTTGAGTATGGTTGAGGTAGGTGGAATGTGTTGTGTAGCGGTGAAATGCATAGATATGACACAGAACGCCAATTGCGAAGGCAGCTTACTAAGCCATTACTGACGCTGAGGCACGAAAGCGTGGGG |
| BH-asv920 | GCAAGCGTTATCCGGATTTACTGGGTGTAAAGGGAGCGCAGGCGGTGCGGCAAGTCTGATGTGAAAGCCCGGGGCTCAACCCCGGTACTGCATTGGAAACTGTCGTACTAGAGTGTCGGAGGGGTAAGCGGAATTCCTAGTGTAGCGGTGAAATGCGTAGATATTAGGAGGAACACCAGTGGCGAAGGCGGCTTACTGGACGATAACTGACGATGAGGCTCGAAAGCGTGGGG |
| BH-asv921 | GCTAGCGTTGTTCGGAATTACTGGGCGTAAAGCGCACGTAGGCGGACTGGAAAGTTGGGGGTGAAATCCCGGGGCTCAACCCCGGAACTGCCTTCAAAACTCCCAGTCTGGAGTTCGAGAGAGGTGAGTGGAATTCCGAGTGTAGAGGTGAAATTCGTAGATATTCGGAGGAACACCAGTGGCGAAGGCGGCTCACTGGCTCGATACTGACGCTGAGGTGCGAAAGCGTGGGG |
| BH-asv922 | GCTAGCGTTGTTCGGAATTACTGGGCGTAAAGCGCGCGCAGGCGGCTATTCCAGTCAGAGGTGAAAGGCCCGGGCTCAACCTGGGAACTGCCTTTGATACTGAATGGCTTGAGTTCGGAAGAGGGTAGTGGAATTCCCAGTGTAGAGGTGAAATTCGTAGATATTGGGAAGAACACCGGTGGCGAAGGCGGCTACCTGGTCCGATACTGACGCTGAGGCGCGAAAGCGTGGGG |
| BH-asv923 | GCAAGCGTTAATCGGAATTACTGGGCGTAAAGGGTTCGTAGGTGGTTTAGTAAGTTATGTGTGAAATTCCTGGGCTCAACCTGGGGTGGTCAGATAATACTGCTAGACTAGAGTATAGGAGAGGGTAGTGGAATTTCCGGTGTAGCGGTGAAATGCGTAGATATCGGAAGGAACACCAGTGGCGAAGGCGACTACCTGGCCGGATACTGACACTGAGGGACGAAAGCGTGGGG |
| BH-asv924 | GCTAGCGTTGTTCGGAATTACTGGGCGTAAAGCGCACGTAGGCGGCTTTTCAAGTCAGAGGTGAAAGCCTGGAGCTTAACTCCAGAACTGCCTTTGAAACTAGAAAGCTAGAATCATGGAGAGGTGAGTGGAATTCCGAGTGTAGAGGTGAAATTCGTAGATATTCGGAAGAACACCAGTGGCGAAGGCGGCTCACTGGACATGTATTGACGCTGAGGTGCGAAAGCGTGGGG |
| BH-asv925 | GCAAGCGTTGTCCGGAATTATTGGGCGTAAAGAGCGCGTAGGCGGTATGATAAGTCTGTCTTAGAAGTGCGGGGCTTAACCCCGTGAGGGGACAGAGACTGTCATACTAGAGTCTCGGAGAGGAAAGCGGAATTCCTAGTGTAGCGGTGAAATGCGTAGAGATTAGGAAGAACACCGGTGGCGAAGGCGGCTTTCTGGACGAGCACTGACGCTGAGGCGCGAAAGCGTGGGG |
| BH-asv926 | GCGAGCGTTGTCCGGAATCACTGGGCGTAAAGGGCGCGTAGGTGGCGTGGTTAGCGTGCGGTGAAAGCCCGGGGCTCAACTCCGGGTCGGCCGTGCGAACTGCCATGCTTGAGCACGGTAGAGGCCAATGGAATTCCGTGTGTAGCGGTGGAATGCGTAGAGATCCGGAGGAACACCGGTGGCGAAGGCGGTTGGCTGGGCCGTTGCTGACACTGATGCGCGACAGCGTGGGG |
| BH-asv927 | GCTAGCGTTGCTCGGAATCACTGGGCGTAAAGGGCGCGTAGGCGGCCGGCCAAGTCGGGGGTGAAAGCCTGTGGCTCAACCACAGAATGGCCCTCGATACTGGACGGCTCGAGACCGGAAGAGGTTGGTGGAATTGCGAGTGTAGAGGTGAAATTCGTAGATATTCGCAAGAACACCGGTGGCGAAGGCGGCCAACTGGTCCGGATCTGACGCTGAGGCGCGAAAGCGTGGGG |
| BH-asv928 | ACAAGCGTTGTCCGGAATTATTGGGCGTAAAGCGCGCGCAGGTGGTTCCTTAAGTCTGATGTGAAAGCCCACGGCTCAACCGTGGAGGGTCATTGGAAACTGGGGAACTTGAGTGCAGAAGAGGAAAGTGGAATTCCAAGTGTAGCGGTGAAATGCGTAGAGATTTGGAGGAACACCAGTGGCGAAGGCGACTTTCTGGTCTGTAACTGACACTGAGGCGCGAAAGCGTGGGG |
| BH-asv929 | GCGAGCGTTGTCCGGAATCATTGGGCGTAAAGAGCTCGTAGGCGGCTTGGTAAGTCGGATGTGAAACCTCCAGGCTCAACCTGGAGTCGCCATCCGATACTGCCATGGCTAGAGTCCGGTAGGGGCCCACGGAACTCCTGGTGTAGCGGTGAAATGCGCAGATATCAGGAAGAACACCGGTGGCGAAGGCGGTGGGCTGGGCCGGCACTGACGCTGAGGAGCGAAAGCGTGGGG |
| BH-asv930 | CCAAGCGTTATCCGGATTTACTGGGCTTACAGGGTCCGCAGGCTGTCTGGAAAGTTGGTGGTTAAACCTTCAGGCTCAACCTGAAGCTTGCTATCAATACTGCCAGACTTGAGGCTGTAACAGGCGGACGGAATTCCCGGTGTAGTCGTAAAAAGCGCTGATATCGGGAGGAACGTCGAAAGCGAAAGCAGTCCGCTGGGACAAGCCTGACGCTCAGGGACGAAAGCGTGGGT |
| BH-asv931 | GCAAGCGTTATCCGGAATGATTGGGCGTAAAGCGTCTGTAGGTGGCTTTTTAAGTTCGCCGTCAAATCCCAGGGCTCAACCCTGGACAGGCGGTGGAAACTACCAAGCTGGAGTACGGTAGGGGCAGAGGGAATTTCCGGTGGAGCGGTGAAATGCGTAGAGATCGGAAGAACACCAACGGCGAAGGCACTCTGCTGGGCCGAACGACTGACGCTGAGAGACGAAAGCTAGGGG |
| BH-asv932 | GCAAGCGTTGTTCGGAATCATTGGGCGTACAGGGCGCGCAGGCGGCCGTTCAAGTCCGGTGTGAAAGCCCGGGGCTCAACCCCGGAGGTGCATCGGAAACTGTCCGGCTTGAGTCCGAGAGGGGCCCGTGGAATTCCTGGTGTAGGGGTGAAATCCATAGAGATCAGGAGGAACACCGGAGGTGAAAGCGACGGGCTGGATCGGGACTGACGCTGAGGCGCGAAAGCGCGGGT |
| BH-asv933 | TCGAGTGATGGCCGCTATTATTGGGCCTAAAGCGTCCGTAGCTGGCCGTGTTAGTCCATCGGGAAATCCAGGCGCTTAACGTCTGGACGTCCGGTGGAAACTGCACGGCTTGGGACCGGAAGACTCGAGGGGTACGTCCGGGGTAGGAGTGAAATCCCGTAATCCTGGACGGACCACCGAAGGCGAAAGCACCTCGAGAGCACGGATCCGACAGTGAGGAACGAAAGCTGGGGT |
| BH-asv934 | TCAAGCGTTATCCGGATTTATTGGGCGTAAAGGGCGCGTAGGCGGCCTATCAGGTTCTCTGTAAAATCTGGAGGGCTCAACCCTCCTGCTGCAGGGAAAACCGGTAGGCTTGAGGACGTCAGAGGCTGACGGAACGCACGGTGTAGAGGTGAAATTCGTTGATATCGTGCGGACACCAAAAGCGAAGGCAGTCGGCTGGGACGTTCCTGACGCTGAGGCGCGATACCTCGGGT |
| BH-asv935 | GCAAGCGTTGTCCGGAATCACTGGGCGTAAAGGGTCCGCAGGCGGCCTGACAACTCAGGCGTGAAATCCCACGGCTTAACCGTGGAATTGCGTTTGAAAGTGTCAGGCTCGAGGACAGTAAGGGGGAGTGGAACTCCGGGTGGAGCGGTGAAATGCGCAGATATCGGAGGAACACCGGAGGCGAAAGCGACTCCCTAGCCTGTTCCTGACGCTGAGGGACCAAAGCCAGGGT |
| BH-asv936 | GCTAGCGTTGCTCGGAATCACTGGGCGTAAAGGGTGCGTAGGTGGATCTTTAAGTCAGGGGTGAAATCCTGGAGCTCAACTCCAGAACTGCCTTTGATACTGAGGATCTTGAGTTCGGGAGAGGTAAGTGGAACTGCGAGTGTAGAGGTGAAATTCGTAGATATTCGCAAGAACACCAGTGGCGAAGGCGGCTTACTGGCCCGATACTGACACTGAGGCACGAAAGCGTGGGG |
| BH-asv937 | GCTAGCGTTGTTCGGAATCACTGGGCGTAAAGCGCACGTAGGCGGATCGTTAAGTCAGAGGTGAAATCCCGGAGCTCAACTTCGGAACTGCCTTTGATACTGGCGATCTCGAGTCTGGAAGAGGTTAGTGGAACTGCGAGTGTAGAGGTGAAATTCGTAGATATTCGCAAGAACACCAGTGGCGAAGGCGGCTAACTGGTCCAGTACTGACGCTGAGGTGCGAAAGCGTGGGG |
| BH-asv938 | GCAAGCGTTGTTCGGAATTACTGGGCGTAAAGCGCGCGTAGGCGGTTTGTCAAGTCAGCTGTGAAAGTCCATGGCTTAACCATGGAAGTGCACTTGAAACTGACAGACTTGAGTACTGGAGGGGGTGGTGGAATTCCCGGTGTAGAGGTGAAATTCGTAGATATCGGGAGGAATACCGGTGGCGAAGGCGACCACCTGGCCAGATACTGACGCTGAGGTGCGAAAGCGTGGGG |
| BH-asv939 | GCAGGCGTTAATCGGAATTACTGGGCGTAAAGCGTGCGCAGGCGGTCTGCTAAGACAGGTGTGAAATCCCCGGGCTTAACCTGGGAACTGCGCTTGTTACTGGCAGGCTTGAGTGTGGCAGAGGGGGGTGGAATTCCACGTGTAGCAGTGAAATGCGTAGAGATGTGGAGGAACACCGATGGCGAAGGCAGCCCCCTGGGCTAACACTGACGCTCATGCACGAAAGCGTGGGG |
| BH-asv940 | GCAAGCGTTGTCCGGATTTATTGGGTTTAAAGGGTGCGTAGGCGGCCTTATAAGCCAGTGCTGAAATATCCCGGCTTAACCGGGAGGGTGGCATTGATACTGTAGGGCTTGAGTGAAATCGAGGTAGGCGGAATTGACGGTGTAGCGGTGAAATGCTTAGATATCGTCAAGAACACCGATAGTGAAGACAGCTTACTAGGCTTAAACTGACGCTGAGGCACGAAAGTGTGGGG |
| BH-asv941 | GCGAGCGTTGTCCGGAATTATTTGGCGTAAAGAGCGTGTAGGCGGTTCGGTAAGTCTGCCGTGAAAACCCAGGGCTCAACCCTGGGCGTGCGGTGGATACTGCCGGGCTAGAGGGTGGTAGAGGCGAGTGGAATTCCCGGTGTAGCGGTGAAATGCGCAGATATCGGGAGGAACACCAGTAGCGAAGGCGGCTCGCTGGGCCATACCTGACGCTGAGACGCGAAAGCGTGGGG |
| BH-asv942 | GCTAGCGTTGTCCGGAATTACTGGGCGTAAAGCGCGCGTAGGCGGTCCACCAAGTCGGGGGTGAAAGCCCGGGGCTCAACCCCGGAATGTCCTCCGAGACTGATGGACTGGAGTTCGGGAGAGGAGAGCGGAATTCCCAGTGTAGAGGTGAAATTCGTAGATATTGGGAAGAACACCGGTGGCGAAGGCGGCTCTCTGGCCCGCAACTGACGCTGAGGCGCGAAAGCGTGGGG |
| BH-asv945 | TCGAGTGATGACCGATATTATTGGGCCTAAAGCGTCCGTAGCTGGCTGGGCAAGTCTGTTGGGAAATCTGCTCGCTCAACGAGCAGGCGTCCGGCAGAAACTGCTCAGCTCGGGACCGAGCGGTCCGAGGGGTACGTCCGGGGTAGGAGTGAAATCCCGTAATCCTGGACGGACCACCGATGGCGAAAGCACCTCGGAAGATCGGATCCGACAGTGAGGGACGAAAGCTGGGGT |
| BH-asv946 | GCGAGTGGTGGCCACGATTATTGGGCTTAAAGCGTCCGTAGCTGGCCTCATAAGTCTGTTGTGAAATCTGGGGGCTCAACCTTCAGAGTTGCAACGGATACTATGAGGCTTGGGACCGGAGGGCGTCGAGGCTATTCCCGGGGTAGGGGTGAAATCCAAAAATCCCGGGAGGAGCACCAGTGGCGAAAGCGCTCGGCGACTACGGGTCCGACGGTAAGGGACGAAAGCCAGGGG |
| BH-asv947 | CCGAGTGATGGCCGTGATTATTGGGCCTAAAGCGTCCGTAGCCGGCCGCACAAGTCCGTCGGGAAATCCGCCCGCTCAACGGGTGGGCGACCGGCGGAAACTGTGCGGCTTGGGGCCGGAAGACCCGAGGGGTACGTCCGGGGTAGGAGTGAAATCCCGTAATCCCGGACGGACCACCGGTGGCGAAAGCGCCTCGGGAGGACGGACCCGACGGTCAGGGACGAAAGCTAGGGT |
| BH-asv948 | GCAAACGTTGCTCGGATTTACTGGGCGTAAAGCGCGTGTAGGCGGGTTCGCAAGTCGGACGTGAAATCCCTGGGCTCAACCCAGGAACTGCGTTCGAAACTGCGAATCTAGAGTAGTGGAGAGGAAAGTAGAATTCCCGGTGTAGAGGTGAAATTCGTAGATATCGGGAGGAATACCAGTGGCGAAGGCGGCTTTCTGGACACTTACTGACGCTGAGACGCGAAAGCGTGGGG |
| BH-asv949 | GCAAGCGTTGTTCGGAATTACTGGGCGTAAAGCGCGCGTAGGCGGTTTTATAAGTCAGATGTGAAAGTCCACGGCTCAACCGTGGAAGTGCATTTGAAACTGTAATACTTGAGTACTGGAGGGGGTGGTGGAATTCCCGGTGTAGAGGTGAAATTCGTAGATATCGGGAGGAATACCGGTGGCGAAGGCGACCACCTGGCCAGATACTGACGCTGAGGTGCGAAAGCGTGGGG |

| | |
|---|---|
| BH-asv950 | GCAAGCGTTGTTCGGAATTATTGGGCGTAAAGCGCGCGCAGGCGGCTCGCAAAGTCCGATGTGAAATCCCTCGGCTCAACCGAGGACGTGCATTGGATACTGGCGAACTTGAGTCTCGGAGAGGGTGGCGGAA<br>TTCCCAGTGTAGAGGTGAAATTCGTAGATATTGGGAGGAACACCAGTGGCGAAGGCGGCCACCTGGACGATGACTGACGCTGAGGCGCGAAAGCGTGGGT |
| BH-asv952 | GCAAGCGTTGTTCGGATTCACTGGGTATAAAGGGTGCGCAGGCGGCCTGGTAAGTCAGGGGTGAAATATGACGGCTCAACCGTCAAACTGCCTCTGAAACTGCCTGGCTTGAGTCCGGGAGAGGTAGGTGGAAT<br>TCCAGGTGTAGCGGTGAAATGCGTAAATATCTGGAGGAACACCGGTGGCGAAGGCGGCCTACTGGCCCGGTACTGACGCTCAGGCACGAAAGCTAGGGG |
| BH-asv953 | GCAAACGTTGTTCGGATTTACTGGGCTTAAAGGGCGCGTAGGCGGCAGCGTAAGTCAAGGTTGAAATCCGGCGGCTCACCCGTCGAAGTGGCCTCGATACTGCGTTGCTAGAATCATGGAGGGGTGCGCGGAA<br>CTTGTGGTGGAGCGGTGAAATGCTCAGATATCACAGGGAACACCGGAGGCGAAGGCGGCGCACTGGCCATGTATTGACGCTGAGGCGCGAAAGCGTGGGG |
| BH-asv954 | GCTAGCGTTGTTCGGAATCACTGGGCGTAAAGCGCATGCAGGCGGATCGTTAAGTCGGGGGTGAAATCCCGAGGCTCAACCTCGGAACTGCCTCCGATACTGGCGATCTCGAGTCCGGAAGAGGTGAGTGGAA<br>TTGCGAGTGTAGAGGTGAAATTCGTAGATATTCGCAAGAACACCAGTGGCGAAGGCGGCTCACTGGTCCGGTACTGACGCTGAGATGCGAAAGCGTGGGG |
| BH-asv956 | GCAAGCGTTATCCGGATTCACTGGGTTTAAAGGGAGCGTAGGCGGGCATGTAAGTCCGTGGTGAAAGCCCTCAGCTTAACTGAGGAACTGCCATGGATACTATATGTCTTGAATGTCGTTGAGGTTTGCGGAATA<br>GGTCATGTAGCGGTGAAATGCTTAGATATGACCTGGAACACCTATTGCGAAGGCAGCAGGCTAAACGAATATTGACGCTGAGGCTCGAAAGCGTGGGG |
| BH-asv957 | CCGAGCGTTATCCGGATTTATTGGGTTTAAAGGGAGCGTAGATGGATGTTTAAGTCAGTTGTGAAAGTTTGCGGCTCAACCGTAAAATTGCAGTTGATACTGGATATCTTGAGTGCAGTTGAGGCAGGCGGAATTC<br>GTGGTGTAGCGGTGAAATGCTTAGATATCACGAAGAACTCCGATTGCGAAGGCAGCCTGCTAAGCTGCAACTGACATTGAGGCTCGAAAGTGTGGGT |
| BH-asv958 | GCGAGCGTTAATCGGAATCACTGGGCGTAAAGCGCACGTAGGCTGTTTGGTAAGTCAGGGGTGAAATCCCGCGGCTCAACCGCGGAATTGCCCTTGATACTGCTGGACTTGAGTCCGGGAGAGGGTGGCGGAA<br>TTCCAGGTGTAGGAGTGAAATCCGTAGATATCTGGAGGAACATCAGTGGCGAAGGCGGCCACCTGGACCGGTACTGACGCTGAGGTGCGAAAGCGTGGGG |
| BH-asv959 | GCAAGCGTTATCCGGAATCATTGGGTTTAAAGGGTCCGTAGGCGGATAATTAAGTCAGGGGTGAAAGTCTGCAGCTCAACTGTAGGACTGCCCTTGATACTGGTTATCTTGAATCGTTGTGAAGTGGCTAGAATAT<br>GTAGTGTAGCGGTGAAATGCATAGATATTACATAGAATACCGATTGCGAAGGCAGGTCACTAACAACGTATTGACGCTGAGGGACGAAAGCGTGGGG |
| BH-asv960 | GCGAGCGTTGTCCGGAATTACTGGGCGTAAAGAGCTCGTAGGTGGTCTGTCGCGTCATTTGTGAAAGCCCGGTGCTTAACTCCGGGTTGGCAGGTGATACGGGCATGACTGGAGTACTGTAGGGGAGACTGGA<br>ATTCCTGGTGTAGCGGTGAAATGCGCAGATATCAGGAGGAACACCGGTGGCGAAGGCGGGTCTCTGGGCAGTAACTGACGCTGAGGAGCGAAAGCATGGGT |
| BH-asv961 | GCAAGCGTTAATCGGAATTACTGGGCGTAAAGCGCACGTAGGCGGTTTGCTAAGTCTGCTGTGAAAGCCCCGGGCTCAACCTGGGAATTGCAGTGGATACTGGCAAGCTCGAGTTTGGTAGAGGGAAGTGGAAT<br>TCCAGGTGTAGCGGTGAAATGCGTAGATATCTGGAGGAACATCAGTGGCGAAGGCGACTTCCTGGCCCAAAACTGACGCTGAGGTGCGAAAGCGTGGGG |
| BH-asv962 | GCGAGCGTTGTCCGGAATCACTGGGCGTAAAGGGCGCGTAGGCGGTGGCTTAAGCGTGTGGTGAAAGCCCGGGGCTCAACCTCGGGTCGGCCATGCGAACTGGGTCACTCGAGCACTGTAGAGGCAGGTGGA<br>ATTCCGGGTGTAGCGGTGGAATGCGTAGAGATCCGGAAGAACACCGGTGGCGAAGGCGGCCTGCTGGGCAGTAGCTGACGCTGAGGCGCGACAGCGTGGGG |
| BH-asv963 | GCTAGCGTTGTTCGGAATTACTGGGCGTAAAGCGCGCGTAGGCGGCTTGGTAAGTCGATTGTGAAATCCCCGGGCTCAACCTGGGAATTGCAGTCGAGACTGTCAGGCTAGAGTGTGGGAGAGGGCAGTGGAA<br>TTCCCGGTGTAGCGGTGAAATGCGTAGATATCGGGAGGAACATCAGTGGCGAAGGCGACTGCCTGGCCCAACACTGACGCTGAGGTGCGAAAGCGTGGGT |
| BH-asv964 | GCTAGCGTTGTCCGGAATCATTGGGCGTAAAGCGCGTGTAGGCGGCCCGGTAAGTCCGCTGTGAAAGTCGGGGGCTCAACCCTCGGATGCCGGTGGATACTGTCGGGCTCGAGTACGGAAGAGGCGAGTGGA<br>ATTCCTGGTGTAGCGGTGAAATGCGCAGATATCAGGAGGAACACCGATTGCGAAGGCAGCTCGCTGGGACGTTACTGACGCTGAGACGCGAAAGCGTGGGG |
| BH-asv965 | GCAAGCGTTGTCCGGAATTATTGGGCGTAAAGCGCGCGCAGGCGGTCCTTTAAGTCTGATGTGAAAGCCCACGGCTCAACCGTGGAGGGTCATTTGAAACTGGGGGACTTGAGTACAGAAGAGGAAAGCGGAAT<br>TCCACGTGTAGCGGTGAAATGCGTAGAGATGTGGAGGAACACCAGTGGCGAATGCGGCTTTCTGGTCTGTAACTGACGCTGAGGCGCGAAAGCGTGGGG |
| BH-asv966 | GCAAGCGTTGTTCGGAATTACTGGGCGTAAAGGGCGCGTAGGCGGTCGCCTGAGTCGGACGTGAAATCCCCAAGCTTAACTTGGGAACTGCGTCCGATACTGGGTGACTTGAGTTCGGGAGAGGAATGTGGAAT<br>TCCAGGTGTAGCGGTGAAATGCGTAGATATCTGGAGGAACACCGGTGGCGAAGGCGGCATTCTGGACCGAAACTGACGCTGAGGCGCGAAAGCTAGGGG |
| BH-asv967 | GCTAGCGTTGTTCGGAATCACTGGGCGTAAAGGGCGCGTAGGCGGCCTTTGGTGTCGGATGTGAAAGCCCGGGGCTCAACTCCGGAGGTGCATCCGAAACTCAGGGGCTCGAGGCCGGAAGAGGAGGGTGGA<br>ATACCCAGTGTAGAGGTGAAATTCGTAGATATTGGGTGGAACACCGGTGGCGAAAGCGGCTCTCTGGTCCGGTCCTGACGCTGAGGCGCGAAAGCGTGGGG |
| BH-asv968 | GCGAGCGTTAATCGGAATTACTGGGCGTAAAGCGCGCGTAGGCGGCTTTGCAAGTCGGGTGTGAAATCCCCAGGCTTAACCTGGGAACTGCACTCGAGACTGCATTGCTAGAGTATGGGAGAGGGAAGTGGAAT<br>TTCAGGTGTAGCGGTGAAATGCGTAGATATCTGAAGGAACATCAGTGGCGAAAGCGACTTCCTGGACCAATACTGACGCTCATGTGCGAAAGCGTGGGG |
| BH-asv970 | TCGAGTGATGGCCGATATTATTGGGCCTAAAGCGTTCGTAGCTGGCCGAGCAAGTCTGTCGGGAAATCCACGCGCTTAACGCGTGGGCGTCCGGCAGAAACTGTTCGGCTTGGGACCGGAAGACTCGGGGAGT<br>ACGTCTTGGGTAGGAGTGAAATCCCGTAATCCTAGACGGACTACCGATGGCGAAAGCACCCCGAGAGCACGGATCCGACAGTGAGGAACGAAAGCTGGGGT |
| BH-asv971 | CCAAGCGTTATCCGGAATCATTGGGTTTAAAGGGTCCGTAGGCGGAATAATAAGTCAGTGGTGAAAGTCTGCAGCTCAACTGTAGAATTGCCATTGATACTGTTGTTCTTGAATTAGTATGAAGTGGTTGGAATGA<br>GTAGTGTAGCGGTGAAATGCATAGATATTACTCAGAACACCTATTGCGAAGGCAGATCACTAATACTATATTGACGCTGAGGGACGAAAGCGTAGGT |
| BH-asv972 | GCAAGCGTTGTCCGGAATCACTGGGCGTAAAGGGTGCGCAGGCGGGGCAGCAAGTCGGAGGTGAAAGCCCATGGCTCAACCATGGAAGTGCCTTCGAAACTGTTGCTCTTGAGTCCCGGAGAGGTTGTCGGAA<br>TTCGTGGTGTAGCGGTGAAATGCGTAGAGATCACGAGGAACACCAGAGGCGAAAGCGGGCAGCTGGACGGGTACTGACGCTCAGGCACGAAAGCGTGGGG |
| BH-asv973 | CCGAGTGGTAGCCACTATTATTGGGCTTTAAGGGCCCGTAGCCGGCCATGTAAGTCCCTAGTGAAATCTGCCCGCTTAACGGGTAGAATTGCTGGGGAAACTGCGTGGCTTGGGGCCGGACGGCGCCGAGGGT<br>ATTCCCGGGGTAGGGGTAAAATCCAAAAATCCTGGGAGGACCGCCCGTGGCGAAAGCGCTCGGCGAGGACGGACCCGACGGTGAGGGACGAAAGCCAGGGG |
| BH-asv974 | CCAAGCGTTATCCGGATTCACTGGGTTTAAAGGGTGCGCAGGCGGTCAGGTAAGTCAGTGGTGAAAGCCTTCGGCTCAACCGAAGAATTGCCATTGAAACTACTTGACTTGAGTGTAGTTGAGGTGGGCGGAATG<br>TAGCATGTAGCGGTGAAATGCATAGATATGCTACAGAACACCGATAGCGAAAGCAGCTCACCAGACTATAACTGACGCTGAGGCACGAAAGCATGGGT |
| BH-asv976 | GCGAGTGGTGGCCACGATTATTGGGCTTAAAGTGTTCGTAGCCGGCTTCGTGAGTCCGCTGTGAAATCTAGCGGCTCAACCGTTAGAAGCGCAGCGGATACCACGAGGCTAGGGGCCGGAGGGCGTCGAGGCT<br>ATTCCTGGAGTAGGGGTGAAATCCCAAAATTCCAGGAGGAGCACCAGTGGCGAAAGCGCTCGGCGACTACGGACCCGACGGTGAGGAACGAAAGCCAGGGG |
| BH-asv977 | GCAAGCGTTGTCCGGATTTATTGGGCGTAAAGAGTTCGTAGGCGGTTTACTAAGTCTGATGTTAAAGACCGGGGCCCAACCCTGGGACTGCATTGGATACTGGTAGGCTTGAGTGTGGTAGAGGCAAGTGGAATT<br>CCCAGTGTAGCGGTGAAATGCGTAGATATTGGGAAGAACACCAGTGGCGTAGGCGACTTGCTGGGCCATAACTGACGCTGAGGAACGAAAGCCAGGGG |
| BH-asv978 | GCAAGCGTTGTCCGGAATTATTGGGCGTAAAGAGCTCGTAGGCGGTTTGTCGCGTCTGCTGTGAAAGCCCGGGGCTCAACCCCGGGTCTGCAGTGGGTACGGGCAGACTAGAGTGCAGTAGGGGAGACTGGAA<br>TTCCTGGTGTAGCGGTGAAATGCGCAGATATCAGGAGGAACACCGATGGCGAAGGCAGGTCTCTGGGCTGTTACTGACGCTGAGGAGCGAAAGCATGGGG |
| BH-asv979 | CCGAGTGATGGCCGATATTATTGGGCCTAAAGCGTCCGTAGCTGGCCGCGGAGGTCCGTCGGGAAATCTACTGGCCTAACCAGTAGGCGTCCGGCGGAAACCCCGTGGCTTGGGACCGGAGACCTGAGGGG<br>TACGTCCGGGGTAGGAGTGAAATCCCGTAATCCTGGACGGACCACCGGTGGCGAAAGCGCCTCAGGAGAACGGATCCGACAGTGAGGGACGAAAGCTAGGGT |
| BH-asv980 | GCAAGCGTTAATCGGATTTACTGGGCGTAAAGCGTGCGTAGGCGGTTTGTTAAGTCGGATGTGAAATCCCTGGGCTCAACCTGGGAACTGCATTCGATACTGGCAGGCTAGAGAACGGTAGAGGGAGGCGGAAC<br>TCCAGGTGTAGCAGTGAAATGCGTAGATATCTGGAAGAACACCGATGGCGAAGGCAACCTCCTGGGCCTGTTCTGACGCTGAGGCACGAAAGCGTGGGG |
| BH-asv981 | GCAAGCGTTATCCGGAATCATTGGGTTTAAAGGGTGCGTAGGCGGAGCAGTAAGTCAGTGGTGAAAGCCTACGGCTTAACCGTGGAATTGCCATTGATACTGCTGCTCTTGAGTACGGCCGGAGTGGGCGGAAT<br>GCGTCATGTAGCGGTGAAATGCATAGATATGACGCAGAACACCGATCGCGAAAGCAGCTCACTAGACCGTAACTGACGCTGAGGCACGAAAGCGTGGGG |
| BH-asv982 | GCAAGCGTTAATCGGAATTACTGGGCGTAAAGCGCGCGTAGGTGGTTTGATAAGTTGGATGTGAAAGCCCCGGGCTCAACCTGGGAATTGCATCCAAAACTGTCTGACTAGAGTATGGCAGAGGGTGGTGGAAT<br>TTCCTGTGTAGCGGTGAAATGCGTAGATATAGGAAGGAACACCAGTGGCGAAGGCGACCGCCTGGGCTAATACTGACACTGAGGTGCGAAAGCGTGGGG |
| BH-asv983 | GCAAGCGTTGTCCGGATTTACTGGGCGTAAAGGGGGTGCAGGCGGCTGGGTCCGTCGGTAGTGAAAGCTCCGGGCTCAACCTGGAGAGGTCTACCGATACCACCCAGCTTGAGGGCGGTAGAGGAGCACGGA<br>ATTCCTGGTGTAGTGGTGAAATGCGTAGATATCAGGAGGAACACCAGTGGCGAAAGCGGTGCTCTGGGCCGTTCCTGACGCTGAGGCCCGAAAGCGTGGGG |
| BH-asv984 | GCAAGCGTTAACCGGAATTACTGGGCGTAAAGCGTGCGTAGGCGGCTCCTCATGTCGGGTGTGAAAGCCCCGGGCTCAACCTGGGAATGGCACTCGAAACTGCGGGGCTAGAGTGTGGTAGAGGAAGATGGAA<br>TTCCACGTGTAGCGGTGAAATGCGTAGATATGTGGAGGAACACCGATGGCGAAGGCAGTCTTCTGGGCCAACACTGACGCTGAGGTACGAAAGCGTGGGT |
| BH-asv985 | GCTAGCGTTGTTCGGAATCACTGGGCGTAAAGCGTACGTAGGCGGATGCTTAAGTCAGAGGTGAAAGCCTGGAGCTCAACTCCAGAACTGCCTTTGATACTGGGCGTCTTGAGTCCGGGAGAGGTGAGTGGAAC<br>TGCTAGTGTAGAGGTGAAATTCGTAGATATTAGCAAGAACACCAGTGGCGAAGGCGGCTCACTGGCCCGGTACTGACGCTGAGGTACGAAAGCGTGGGG |
| BH-asv986 | GCAAGCGTTACTCGGAATCACTGGGCGTAAAGGGTGCGTAGGCGGCTTGTTAAGTTGGATGTGAAAGCCCTCGGCTCAACTGAGGAACTGCGTCCAAAACTGGCAAGCTCGAGGATGGTAGGGGACAGTGGAA<br>CTCCAGGTGGAGCGGTGAAATGCGTAGAGATCTGGAGGAACGCCGGTGGCGAAAGCGATTGTCTGGGCCATTCCTGACGCTGAGGCACGAAAGCCAGGGG |
| BH-asv989 | CCGAGCGTTAGGCGGAATCACTGGGCTTAAAGCGTGCGTAGGTGGATGAGAAAGCGTCTTGTGAAAGCCCTCAGCTCAACTGAGGAATTGCGGGGCGAACTGCTTGTCTTGAGGCAGGTAGAGGTCGTCGGAA<br>CGTTAGGTGGAGCGGTGAAATGCGTTGATATCTAACGGAACACCGAAGGGGTAGCCAGGCGACTGGGCCTGTCCTGACACTGAGGCACGAAAGCGTGGGG |
| BH-asv990 | GCTAGCGTTGTTCGGAATTACTGGGCGTAAAGCGCACGTAGGCGGATATTTAAGTCAGGGGTGAAATCCCGGAGCTCAACTCCGGAACTGCCTTTGATACTGGATATCTAGAGACCGAGAGAGGTAAGTGGAACT<br>CCTAGTGTAGAGGTGGAATTCGTAGATATTAGGAAGAACACCAGTGGCGAAGGCGGCTTACTGGCTCGGAACTGACGCTGAGGTGCGAAAGCGTGGGG |
| BH-asv992 | GCAAGCGTTGTCCGGAATTATTGGGCGTAAAGCGCGTGTAGGCGGCTTTGCAAGTCGGGTGTGAAAGCCCTCAGCTCAACTGAGGAAGTGCGCCCGAAACTGCAGAGCTTGAGTGCCGGAGAGGGTGGCGGAA<br>TTCCCCAAGTAGAGGTGAAATTCGTAGATATGGGGAGGAACACCGGTGGCGAAGGCGGCCACCTGGACGGTAACTGACGCTGAGACGCGAAAGCGTGGGG |
| BH-asv993 | GCAAGCGTTGTCCGGAATTATTGGGCGTAAAGCGCGCGCAGGCGGTCCTTTAAGTCTGGTGTGAAAGCCCACGGCTCAACCGTGGAGGGTCATTGGAAACTGGGGGACTTGAGTACAGAAGAGGAAAGCGGAA<br>TTCCACGTGTAGCGGTGAAATGCGTAGAGATGTGGAGGAACACCAGTGGCGAAGGCGGCTTTCTGGTCTGTAACTGACGCTGAGGCGCGAAAGCGTGGGG |
| BH-asv994 | GCAAGCGTTATCCGGATTCACTGGGTTTAAAGGGTGCGTAGGAGGGCAGGTAAGTCAGTGGTGAAATCTCCAAGCTTAACTTGGAAACTGCCGTTGATACTATCTGTCTTGAATATTGTGGAGGTGAGCGGAATAT<br>GTCATGTAGCGGTGAAATGCTTAGATATGACATAGAACACCAATTGCGAAGGCAGCTCGCTACTCAATTATCGACTCTGAGGCACGAAAGCGTGGGG |
| BH-asv995 | GCAAGCGTTGTTCGGAATTATTGGGCGTAAAGCGCGTGTAGGCGGCTTTGCAAGTCGGGTGTGAAAGCCCTCAGCTCAACTGAGGAAGTGCGCCCGAAACTGCAGAGCTTGAGTGCCGGAGAGGGTGGCGGAA<br>CTCCCCAAGTAGAGGTGAAATTCGTAGATATGGGGAGGAACACCGGTGGCGAAGGCGGCCACCTGGACGGTAACTGACGCTGAGACGCGAAAGCGTGGGG |
| BH-asv996 | GCTAGCGTTGTTCGGAATTACTGGGCGTAAAGCGCGCGTAGGCGGACCTGCTAGTCGGGGGTGAAAGCCCGAGGCTCAACCTCGGAACTGCCTTCGATACTGGATGGCTTGAATTCGGGAGAGGGATGCAGAA<br>TTCCAGGTGTAGCGGTGAAATGCGTAGATATCTGGAGGAATACCGGTGGCGAAGGCGGCATCCTGGACCGACATTGACGCTGAGGAGCGAAAGCTAGGGG |
| BH-asv997 | GCAAGCGTTGTCCGGAATTATTGGGCGTAAAGAGCTCGTAGGCGGTTTGTTGCGTCGGCTGTGAAATCCCGAGGCTCAGCCTCGGGTCTGCAGTCGATACGAGCAAACTAGAGTGTTGCAGGGGAGACTGGAAT<br>TCCTGGTGTAGCGGTGAAATGCGCAGATATCAGGAGGAACACCGGTGGTGAAGGCGGGTCTCTGGGCAGCAACTGACGCTGAGGAGCGAAAGCGTGGGG |
| BH-asv998 | GCTAGCGTTGTTTGGAATTACTGGGCGTAAAGCGCGCGTAGGCGGCTTTGTAAGTTAGAGGTGAAAGCCCGGAGCTCAACTCCGGAACTGCCTTTAAGACTGCATCGCTAGAATTGTGGAGAGGTAAGTGGAATT<br>CCGAGTGTAGAGGTGAAATTCGTAGATATTCGGAAGAACACCAGTGGCGAAGGCGACTTACTGGACACATATTGACGCTGAGGTGCGAAAGCGTGGGG |
| BH-asv999 | GCAAGCGTTAATCGGAATTACTGGGCGTAAAGCGTGCGCAGGCGGCTTTGCAAGACAGATGTGAAATCCCCGGGCTTAACCTGGGAACTGCATTTGTGACTGCAAGGCTGGAGTGCGACAGAGGGGGATGGAA<br>TTCCGCGTGTAGCAGTGAAATGCGTAGATATGCGGAGGAACACCGATGGCGAAGGCAATCCCCTGTGCCTGCACTGACGCTCATGCACGAAAGCGTGGGG |
| BH-asv1000 | CCGAGCGTTGTCCGGAATTATTGGGCGTAAAGAGCTCGTAGGCGGTTTGTCGCGTCTGCTGTGAAATCCCGAGGCTCAACCTCGGGCCTGCAGTGGGTACGGGCAGACTGGAGTGCGGTAGGGGAGAATGGAA<br>TTCCTGGTGTAGCGGTGGAATGCGCAGATATCAGGAGGAACACCGATGGCGAAGGCAGTTCTCTGGGCCGTAACTGACGCTGAGGAGCGAAAGCGTGGGG |
| BH-asv1001 | GCAAGCGTTAATCGGATTTACTGGGCGTAAAGCGCGCGTAGGTGGCCAATTAAGTCAAATGTGAAATCCCCGAGCTTAACTTGGGAATTGCATTCGATACTGGTTGGCTAGAGTATGGGAGAGGATGGTAGAATT<br>CCAGGTGTAGCGGTGAAATGCGTAGAGATCTGGAGGAATACCGATGGCGAAGGCAGCCATCTGGCCTAATACTGACACTGAGGTGTGAAAGCATGGGG |

| | |
|---|---|
| BH-asv1002 | GCAAACGTTGCTCGGAATCATTGGGCGTAAAGCGCGCGTAGGCGGCTGGACAAGTTGGATGTGAAAGCCCTGGGCTCAACCCAGGATGTGCATTCAAAACTGTCTGGCTTGAATCTCGGAGGGGGTCAGGGAAT<br>TCCCGGTGTAGAGGTGAAATTCGTAGATATCGGGAGGAACACCAGTGGCGAAGGCGCTGACCTGGACGATGATTGACGCTGAGGCGCGAAAGCGTGGGG |
| BH-asv1003 | CCAAGCGTTATCCGGATCTACTGGGCGTAAAGGGTGTGTAGGCGGTCGTGTTAGTCCATCGTTAAAGCCCTCGGCTTAACCGAGGGGTTGCGGCGGATACGGCACGACTTGAGGGCGCGAGAGGCCAACGGAA<br>CTCATGGTGTAGGGGTGAAATCCGTTGATATCATGGGGAACGCCAAAGGCGAAGGCAGTTGGCTGGCGCGTACCTGACGCTCAAACACGAAAGCCAGGGG |
| BH-asv1004 | GCGAGCGTTAATCGGAATTACTGGGCGTAAAGCGTGCGCAGGCGGTTTCGTAAGACAGACGTGAAATCCCCGGGCTCAACCTGGGAACTGCGTTTGTGACTGCGAGGCTAGAGTACGGCAGAGGGGGGTAGAA<br>TTCCACGTGTAGCAGTGAAATGCGTAGAGATGTGGAGGAATACCGATGGCGAAGGCAGCCCCCTGGGTTAGTACTGACGCTCATGCACGAAAGCGTGGGG |
| BH-asv1005 | GCTAGCGTTGTTCGGAATCACTGGGCGTAAAGCGCACGTAGGCGGATGTGTCAGTCAGGGGTGAAATCCCGGAGCTCAACTTCGGAACTGCCCTTGATACAGCACGTCTCGAGTCCGAGAGAGGTGAGTGGAAT<br>TCCTAGTGTAGAGGTGAAATTCGTAGATATTAGGAAGAACACCGGTGGCGAAGGCGGCTCACTGGCTCGGTACTGACGCTGAGGTGCGAAAGCGTGGGG |
| BH-asv1006 | GCAAGCGTTGTTCGGAATAACTGGGCTTAAAGCGTGCGTAGGCGGCCAAGCAAGTCAGGAGTGAAATCCCCGGGCTCAACCCGGGAATTGCTCTTGAAACTGCATGGCTAGAATCCGGTAGAGGTTAGCAGAAT<br>TCCCAGTGTAGAGGTGAAATTCGCAGATATTGGGAGGAATACCAGTGGCGTAGGCGGCTAACTGGACCGGTTTTGACGCTGAGGTACGAAAGCGTGGGG |
| BH-asv1007 | TCAAGCGTTGTCCGGATTTATTGGGCGTAAAGCGCCCGCAGGCGGCTTGCCAAGTTCGAGGTGAAATCTCCCGGCTCAACTGGGAGGGTGCCTCGAAAACTGGCAGGCTTGAGGCAGGGAGAGGGATGTGGAA<br>TTCCAGGTGTAGTGGTGAAATGCGTAGATATCTGGAGGAACACCAGTGGCGAAGGCGGCATCCTGGCCCTGTCCTGACGCTCAGGGGCGAAAGCGTGGGG |
| BH-asv1008 | GCAAGCGTTAATCGGAATTACTGGGCGTAAAGCGCGCGTAGGCGGTTTGGTAAGCGGGTTGTGAAATCCCCGGGCTTAACCTGGGAATTGCATCCCGAACTGCCAGGCTAGAGTACGATAGAGGTGAGTGGAAT<br>TTCCTGTGTAGCGGTGAAATGCGTAGATATAGGAAGGAACATCAGTGGCGAAGGCGACTCACTGGATCGATACTGACGCTGAGGTGCGAAAGCGTGGGT |
| BH-asv1009 | GCAAGCGTTGTTCGGATTTACTGGGCGTAAAGGGCGCGTAGGCGGCGTATCAAGTCAATTGTGAAATCTTCGGGCTTAACTCGGAACGGTCAATTGATACTGATATGCTAGAGTGCAGAAAGGGCAACTGGAATT<br>CTTGGTGTAGCGGTGAAATGCGTAGATATCAAGAGGAACACCTGAGGCGAAGGCGAGTTGCTAGGCTGACACTGACGCTGAGGCGCGAAAGCTAGGGG |
| BH-asv1011 | GCAAGCGTTGTTCGGAATCACTGGGCGTAAAGCGCGCGTAGGCGGTTGATTAAGTCGAATGTGAAAGCCCAGGGCTCAACCCTGGAACTGCATCCGAAACTGGTCAACTTGAGTCCCGGAGAGGGTGGTGGAAT<br>TCCTGGTGTAGAGGTGAAATTCGTAGATATCAGGAGGAACACCGGTGGCGAAGGCGACCACCTGGACGGTGACTGACGCTGAGGTGCGAAAGCGTGGGT |
| BH-asv1014 | GCAAGCGTTGTTCGGAATTACTGGGCGTAAAGGGCTCGTAGGCGGCCAACTAAGTCGGATGTGAAATCCCCAGGCTCAACCTGGGAACTGCATCCGATACTGGAAGGCTCGAATCCGGGAGAGGGATGCAGAAT<br>TCCAGGTGTAGCGGTGAAATGCGTAGATATCTGGAGGAATACCGGTGGCGAAGGCGGCATCCTGGACCGGCATTGACGCTGAGGAGCGAAAGCCAGGGG |
| BH-asv1015 | GCAAGCGTTGTCCGGAATTATTGGGCGTAAAGCGCGCGCAGGTGGTTTCTTAAGTCTGATGTGAAAGCCCACGGCTCAACCGTGGAGGGTCATTGGAAACTGGGGAACTTGAGTGCAGAAGAGAAGAGTGGAAT<br>TCCACGTGTAGCGGTGAAATGCGTAGAGATGTGGAGGAACACCAGTGGCGAAGGCGACTCTTTGGTCTGTAACTGACGCTGAGGAGCGAAAGCGTGGGG |
| BH-asv1016 | GCTAGCGTTGTTCGGATTTACTGGGCGTAAAGCGCACGTAGGCGGATCGTTAAGTCGGGGGTGAAATCCCGGGGCTCAACCCCGGAACTGCCTTCGATACTGGCGATCTTGAGTCCGGAAGAGGTGAGTGGAA<br>CTCCTAGTGTAGAGGTGGAATTCGTAGATATTAGGAAGAACACCAGTGGCGAAGGCGGCTCACTGGTCCGGTACTGACGCTGAGGTGCGAAAGCGTGGGG |
| BH-asv1017 | GCGAGCGTTGTCCGGAATCACTGGGCGTAAAGGGCGCGTAGGCGGCAGGCCAAGCGCGCGGTGAAAGTCCGGGGCTCAACCCCGGATCTGCCGTGCGGACTGGTTTGCTGGAGCACTGTAGAGGCGAGTGG<br>AATTCCGGGTGTAGCGGTGGAATGCGTAGAGGTCCGGAAGAACACCGGTGGCGAAGGCGGCTCGCTGGGCAGTGGCTGACGCTGAGGCGCGACAGCGTGGGG |
| BH-asv1018 | GCAAGCGTTGTCCGGAATTATTGGGCGTAAAGAGCTCGTAGGCGGTTTGTCGCGTCTGCTGTGAAAGCCCGGGGCTCAACCCCGGGTCTGCAGTGGGTACGGGCAGACTAGAGTGCAGTAGGGGAGACTGGAA<br>TTCCTGGTGTAGCGGTGAAATGCGCAGATATCAGGAGGAACACCGATGGCGAAGGCAGGTCTCTGGGCTGTAACTGACGCTGAGGAGCGAAAGCATGGGG |
| BH-asv1019 | GCAAGCGTTGCTCGGAATTACTGGGCGTAAAGCGTGCGTAGGTGGTTCGTTAAGTCTGATGTGAAAGCCCTGGGCTCAACCTGGGAATGGCATTGGATACTGGCGACCTAGAGTGCGGTAGAGGGTAGTGGAAT<br>TCCCGGTGTAGCAGTGAAATGCGTAGAGATCGGGAGGAACATCTGTGGCGAAGGCGACTGCCTGGACCAGCACTGACACTGAGGCACGAAAGCGTGGGG |
| BH-asv1020 | GCAAGCGTTGTCCGGAATTATTGGGCGTAAAGAGCGTGTAGGCGGTCCGGTAAGTCGGCTGTGAAAGTCCAGGGCTCAACCCTGGGATGCCGGTCGATACTGCCGGACTAGAGTCCGGAAGAGGCGAGTGGAA<br>TTCCCGGTGTAGCGGTGAAATGCGCAGATATCGGGAGGAACACCTATGGCGAAGGCAGCTCGCTGGGACGGTACTGACGCTGAGACGCGAAAGCGTGGGG |
| BH-asv1021 | GCAAACGTTGCTCGGAATCATTGGGCGTAAAGCGCACGTAGGCGGCGAAGCAAGTTGGGTGTGAAAGCCCTGGGCTCAACCCAGGAAGTGCATTCAAAACTGCTTCGCTTGAATCTCGGAGGGGGTCAGAGAAT<br>TCCCGGTGTAGAGGTGAAATTCGTAGATATCGGGAAGAATACCAGTGGCGAAGGCGCTGACCTGGACGAAGATTGACGCTGAGGTGCGAAAGCGCGGGG |
| BH-asv1022 | GCAAGCGTTACTCGGAATTACTGGGCGTAAAGCGTGCGTAGGTGGTTTGTTAAGTCCGATGTGAAAGCCCTGGGCTCAACCTGGGAACGGCATTGGATACTGGCAGGCTAGAGTGCGGTAGAGGGGTGCGGAA<br>TTCCCGGTGTAGCAGTGAAATGCGTAGATATCGGGAGGAACATCCGTGGCGAAGGCGGCACCCTGGACCAGCACTGACACTGAGGCACGAAAGCGTGGGG |
| BH-asv1023 | GCAAGCGTTAATCGGAATTATTGGGCGTAAAGAGCTCGTAGGCGGTTTGTTGCGTCGGCTGTGAGATCCCGAGGCTCAACCTCGGGTCTGCAGTCGATACGAGCAAACTAGAGTGTTGCAGGGGAGACTGGAAT<br>TCCTGGTGTAGCGGTGAAATGCGCAGATATCAGGAGGAACACCGGTGGCGAAGGCGGGTCTCTGGGCAACAACTGACGCTGAGGAGCGAAAGCGTGGGG |
| BH-asv1024 | GCAAGCGTTGTCCGGAATTATTGGGCGTAAAGCGCGCGCAGGCGGTCCCTTAAGTCTGATGTGAAAGCCCACGGCTCAACCGTGGAGGGTCATTGGAAACTGGGGGACTTGAGTGCAGAAGAGGAAAGCGGAA<br>TTCCACGTGTAGCGGTGAAATGCGTAGAGATGTGGAGGAACACCAGTGGCGAAGGCGGCTTTCTGGTCTGTAACTGACGCTGAGGCGCGAAAGCGTGGGG |
| BH-asv1025 | GCTAGCGTTGTTCGGAATTACTGGGCGTAAAGGGCGCGTAGGCGGCCGTTCGCGTCGGATGTGAAATCCCCAGGCTCAACTTGGGAACTGCATCCGATACTGGATGGCTTGAATCCGGGAGAGGGATGCAGAA<br>TTCCAGGTGTAGCGGTGAAATGCGTAGATATCTGGAGGAACACCGGTGGCGAAGGCGGATCTCTGGGCCGATACTGACGCTGAGGAGCGAAAGCATGGGG |
| BH-asv1028 | GCAAGCGTTGCTCGGAATCACTGGGCGTAAAGGGTCCGCAGGCGGTGAATCAACTCGGACGTGAAAACCCATGGCCCAACTATGGAAGTGCGTTCGAAAGTGTTTCACTTGAGTGCGGGAAGGGAGAGTGGAA<br>CTCCGGGTGGAGCAGTGAAATGCGTAGATATCCGGAGGAACGCCGGAGGCGAAAGCGACTCTCTTGCCCGCTACTGACGCTGAGGGACGAAAGCCAGGGT |
| BH-asv1029 | GCAAGCGTTGCCCGGAATCACTGGGCGTAAAGGGTCCGTAGGCGGCCTTAAAACTCAGGCGTGAAATCCCACGGCTTAACCGTGGAACTGCGTCTGAAAGTGTAAGGCTCGAGTCCGGGAAGGGGAAGTGGAA<br>CTCCGGGTGGAGCGGTGAAATGCGCAGATATCCGGAGGAACACCGGAGGTGAAAACGACTTCCTAGCCCGGTACTGACGCTGTGGGACGAAAGCCAGGGT |
| BH-asv1030 | GCGAGCGTTGTCCGGAATTACTGGGCGTAAAGAGCTCGTAGGCGGTTTGTCACGTCGTCTGTGAAATCCTAGGGCTTAACCCTGGACGTGCAGGCGATACGGGCTGACTTGAGTACTACAGGGGAGACTGGAAT<br>TTCTGGTGTAGCGGTGGAATGCACAGATATCAGGAAGAACACCGATGGCGAAGGCAGGTCTCTGGGTAGTAACTGACGCTGAGGAGCGAAAGCATGGGT |
| BH-asv1031 | GCAAGCGTTATCCGGAATTACTGGGCGTAAAGCGCGTGTAGGTGGATATTTAAGTCACTTGTGAAAGCTCCCGGCTAAACTGGGAGAGGTCAGGTGATACTGGATGTCTGGAGGATGTCAGAGGGAAGCAGAAT<br>TCCGGGTGTAGTGGTGAAATGCGTAGAGATCCGGAGGAATACCAGAGGCGAAGGCGGCTTCCTGGGACATACCTGACACTGAGACGCGAAAGCACGGGG |
| BH-asv1032 | GCAAGCGTTATCCGGATTCATTGGGTTTAAAGGGTGCGTAGGCGGGAAGTTAAGTCAGTGGTGAAATCCTGCAGCTCAACTGTAGACTTGCCATTGATACTGGCTTTCTTGAGTGCGCTTGAAGTAGGCGGAATG<br>TGTCGTGTAGCGGTGAAATGCTTAGATATGACACAGAACACCAATTGCGAAGGCAGCTTACTAAGGCGATACTGACGCTGAGGCACGAAAGCGTGGGG |
| BH-asv1034 | GCAAGCGTTGTTCGGAATAACTGGGCGTAAAGGGGGCGCAGGCGGCTTATTAAGTCTGAAGTGAAAGTCCGGGGCTTAACCTCGGAAGGCCTTTGGATACTGGTAAGCTAGAGTACGGAAGAGGAGAGTGGAAT<br>TCCAGGTGTAGCGGTGAAATGCGTAGATATCTGGAGGAACACCGGTGGCGAAAGCGGCTCTCTGGTCCGATACTGACGCTCAGGCCCGAAAGCGTGGGG |
| BH-asv1035 | GCGAGCGTTGTCCGGATTTACTGGGCGTAAAGGATGCGTAGGTGGTAACTTAAGTGGGATGTGAAATCCCCGGGCTCAACCCGGGAACTGCATTCCAAACTGAGTAACTAGAGTGCAGGAGAGGAAAGCGGAAT<br>TCCTAGTGTAGCGGTGAAATGCGTAGATATTAGGAGGAACACCAGTGGCGAAGGCGGCTTTCTGGACTGTAACTGACACTGAGGCATGAAAGCGTGGGG |
| BH-asv1036 | GCAAGCGTTAATCGGAATTACTGGGCGTAAAGCGCGCGTAGGCGGCTCGGTCAGTCGGGTGTGAAATCCCTGGGCTCAACCTGGGAACTGCATTCGATACTGCCGGGCTAGAGTGTGGGAGAGGGTAGTGGAA<br>TTCCCGGTGTAGCGGTGAAATGCGTAGATATCGGGAGGAACATCAGTGGCGAAGGCGACTGCCTGGCCCAACACTGACGCTGAGGTGCGAAAGCGTGGGT |
| BH-asv1037 | CCAAGCGTTATCCGGAATCATTGGGTTTAAAGGGTGCGCAGGCGGTTTTGTAAGTCAGTGGTGAAATCTCTCGGCTCAACCGAGAAACTGCCATTGATACTGCAGAACTTGAATACAGTTGAAGTAGGCGGAATGT<br>GTAGTGTAGCGGTGAAATGCTTAGATATTACACAGAACACCGATAGCGAAGGCAGCTTACTAAACTGAGATTGACGCTCATGCACGAAAGCGTGGGG |
| BH-asv1038 | GCAAGCGTTATCCGGAATCACTGGGCGTAAAGGGTGCGTAGGTGGCTTATCAAGCCAGAGGTGAAAGGCATTGGCTCAACCAATGTAAGCCTTTGGAACTGATTAGCTTGAGTGCAGGAGAGGTAAGTGGAATTC<br>CTAGTGTAGAGGTGAAATTCGTAGATATTAGGAGGAACACCAGTGGCGAAGGCGACTTACTGGACTGTAACTGACACTGAGGCACGAAAGCGTGGGT |
| BH-asv1039 | GCAAGCGTTGTCCGGATTTATTGGGCGTAAAGAGCTCGTAGGCGGTTTCGTAAGTCGAGTGTGAAAAATCTGGGCTTAACCCAGTGGAGCACTTGATACTGCGATGACTTGAGTTTGGTAGAGGAGTGTGGAATT<br>CCTGGTGTAGCGGTGAAATGCGCAGATATCAGGAGGAACACCAACGGCGAAGGCAGCACTCTGGGCCAGTACTGACGCTGAGGAGCGAAAGCGTGGGG |
| BH-asv1041 | GCTAGCGTTGTTCGGAATTACTGGGCGTAAAGCGCGCGTAGGCGGATTGTTAAGTCGGGGGTGAAATCCCGGGGCTCAACCCCGGAACTGCCTTCGATACTGGCAATCTTGAGGTCGAGAGAGGTGAGTGGAA<br>TTCCGAGTGTAGAGGTGAAATTCGTAGATATTCGGAGGAACACCAGTGGCGAAGGCGGCTCACTGGCTCGATACTGACGCTGAGGTGCGAAAGCGTGGGG |
| BH-asv1042 | GCAAGCGTTAATCGGAATTACTGGGCGTAAAGCGCACGTAGGCGGTTTTTTAAGTCAGATGTGAAAGCCCCGGGCTCAACCTGGGAATTGCATTTGAAACTGGAAAACTAGAGTGTGTGAGAGGGGGGTAGAATT<br>CCAAGTGTAGCGGTGAAATGCGTAGAGATTTGGAGGAATACCAGTGGCGAAGGCGGCCCCCTGGCACAACACTGACGCTCAGGTGCGAAAGCGTGGGG |
| BH-asv1043 | CCTAGCGTTGTTCGGAATCACTGGGCGTAAAGCGGATGTAGGTGGATTAGCAAGTCAGGTGTGAAATCCCGGGGCTTAACCTCGGAACTGCATTTGATACTGCTGATCTTGAGTACTGGAGAGGGTAGTAGAATT<br>CCTGGTGTAGTGGTGAAATACGTAGATATCAGGAGGAATACCGGTGGCGAAGGCGGCTACCTGGCCAGATACTGACACTCAGATCCGAAAGCGTGGGG |
| BH-asv1044 | AAGAGCGTTGTTCGGATTTACTGGGCGTAAAGGGCGCGTAGGCGGCATTGCGTGTCAAGTGTGAAATCTCCGGGCTCAACCCGGAAAGTGCGCTTGAAACTACAATGCTCGAGTCTGGGAGAGGGAATCGGAAT<br>GCAGGGTGTAGCGGTGAAATGCGTTGATATCCTGCAGAACACCGGAGGCGAAGGCGGATTCCTTGAACAGCACTGACGCTGAGGCGCGAAAGTCGGGGG |
| BH-asv1045 | GCTAGCGTTGTTCGGAAATACTGGGCGTAAAGCGCACGTAGGCGGCGCAGCAAGTCAGGGGTGAAATCCCGGGGCTCAACCCCGGAACTGCCTCTGAAACTGCAGTGCTAGAATCTTGGAGAGGCGAGTGGAA<br>TTCCGAGTGTAGAGGTGAAATTCGTAGATATTCGGAAGAACACCAGTGGCGAAGGCGACTCGCTGGACAAGTATTGACGCTGAGGTGCGAAAGCGTGGGG |
| BH-asv1046 | GCTAGCGTTGTTCGGAATCACTGGGCGTAAAGGGCGCGTAGGCGGCTTTATAAGTCGGGGGTGAAAGCCTGTGGCTCAACCACAGAATTGCCTTCGATACTGTAGAGCTTGAGACCGGAAGAGGTAAGTGGAAC<br>TGCGAGTGTAGAGGTGAAATTCGTAGATATTCGCAAGAACACCAGTGGCGAAGGCGGCTTACTGGTCCGGTTCTGACGCTGAGGCGCGAAAGCGTGGGG |
| BH-asv1047 | GCGAGCGTTGTCCGGAATTATTGGGCGTAAAGGGCTCGTAGGCGGTTTGTCGCGTCGGGAGTGAAAACTCAGGGCTTAACCCTGAGCCTGCTTCCGATACGGGCAGACTAGAGGTATGCAGGGGAGAACGGAA<br>TTCCTGGTGTAGCGGTGAAATGCGCAGATATCAGGAGGAACACCGGTGGCGAAGGCGGTTCTCTGGGCATTACCTGACGCTGAGGCGCGAAAGCGTGGGG |
| BH-asv1048 | GCAAGCGTTACTCGGAATTACTGGGCGTAAAGCGTGCGTAGGTGGTTGTTTAAGTCTGCTGTGAAAGCCCTGGGCTCAACCTGGGAATTGCAGTGGATACTGGGCGACTAGAGTGTGGTAGAGGATAGTGGAAT<br>TTCCGGTGTAGCAGTGAAATGCGTAGAGATCGGAAGGAACATCTGTGGCAAAGGCGACTATCTGGGCCAACACTGACACTGAGGCACGAAAGCGTGGGG |
| BH-asv1049 | GCAAGCGTTGTCCGGAATTATTGGGCGTAGAGAGCTCGTAGGCGGTTTGTCGCGTCTGTTGTGAAAACGCGAGGCTCAACCTCGCGCCTGCAGTGGGTACGGGCAGACTAGAGTGCGGTAGGGGAGATTGGAA<br>TTCCTGGTGTAGCGGTGGAATGCGCAGATATCAGGAGGAACACCGATGGCGAAGGCAGATCTCTGGGCCGTAACTGACGCTGAGTAGCGAAATCATGGGG |
| BH-asv1050 | GCAGGCGTTAATCGGAATTACTGGGCGTAAAGCGTGCGCAGGCGGTGATGTAAGACAGGTGTGAAATCCCAGGGCTTAACCTTGGAACTGCATTTGTGACCGCATCGCTGGAGTGCGGCAGAGGGGGATGGAA<br>TTCCGCGTGTAGCAGTGAAATGCGTAGATATGCGGAGGAACACCGATGGCGAAGGCAATCCCCTGGGCCTGCACTGACGCTCATGCACGAAAACGTGGGG |
| BH-asv1051 | GCGAGCGTTGTCCGGAATTATTGGGCGTAAAGGGCTCGTAGGCGGTCTGTCACGTCTGCCGTGAAAGCCCAGAGCTTAACTCCAGGTCTGCGGTGGATACGGGCAGACTAGAGTTCGGTAGGGGAGACTGGAA<br>TTCCTGGTGTAGCGGTGAAATGCGCAGATATCAGGAGGAACACCGGTGGCGAAGGCGGGTCTCTGGGCCGATACTGACGCTGAGGAGCGAAAGCGTGGGG |
| BH-asv1052 | CCAAGCATTAGGCGGAATCACTGGGCTTAAAGCGTGTGTAGGCGGATGCCTAAGTACCTTGTGAAATCCCACGGCTCACCCGTGGAACTGCTGGGTATACTGGGTGTCTTGAGCCAACTAGGGGCAACCGGAAC<br>AAACGGTGGAGCGGTGAAATGCGTAGATATCGTTTGGAACGCCAATGGTGAAAACAGGTTGCTGGGGTTGTGCTGACGCTGAGACACGAAAGCCAGGGG |
| BH-asv1053 | CCTAGCGTTGTTCGGAATCATTGGGCGTAAAGCGTGTGTAGGCGGTCTTGTAAGTCAGGTGTGAAAGCCCCGGGCTCAACCCGGGAAGTGCATCTGATACTGCTTGACTTGAGTACTGGAGAGGATAGTGGAAT<br>TCCTGGTGTAGTGGTGAAATACGTAGATATCAGGAGGAACACCGGTGGCGAAGGCGGCTATCTGGCCATGTACTGACGCTGAGACACGAAAGCGTGGGG |

| | |
|---|---|
| BH-asv1054 | GCGAGCGTTGTTCGGAGTTACTGGGCGTAAAGGGTGTGTAGGCGGCTCTCCAAGTTTGGTGTGAAATCTCCCGGCTCAACCGGGAGGGTGCGCCGAAAACTGGAAGGCTAGAGTGCGGGATGGGAGAGTGGA<br>ATTCCTGGTGTAGCGGTGAAATGCGTAGATATCAGGAGGAACACCGGTGGTGTAGACGGCTCTCTGGACCGTAACTGACGCTGAGACACGAAAGCGTGGGT |
| BH-asv1055 | GCTAGCGTTGTTCGGAATCACTGGGCGTAAAGGGCGCGTAGGCGGCCGTTCAAGTCGGGGGTGAAAGCCTGTGGCTCAACCACAGAATGGCCTTCGATACTGGACGGCTCGAGACCGGAAGAGGTCAGTGGAA<br>CTGCGAGTGTAGAGGTGAAATTCGTAGATATTCGCAAGAACACCAGTGGCGAAGGCGGCTGACTGGTCCGGTTCTGACGCTGAGGCGCGAAAGCGTGGGG |
| BH-asv1056 | GCAAGCGTTGTCCGGAATTATTGGGCGTAAAGAGCTCGTAGGCGGTTTGTCGCGTCCGTCGTGAAAGCCCACAGCTTAACTGTGGGTCTGCGGTGGATACGGGCAGACTAGAGGCAGGTAGGGGAGAACGGAA<br>TTCCCGGTGTAGCGGTGAAATGCGCAGATATCGGGAGGAACACCGGTGGCGAAGGCGGTTCTCTGGGCCTGTACTGACGCTGAGGAGCGAAAGCGTGGGG |
| BH-asv1058 | CCAAACGTTATCCGGAATTATTGGTGCGTATAGGGTCCGCAGGCGGTCTGATAAGTTGGTAGTTAAAGTTTCGAGCTTAACTCGAAAATTGCTGCCAAAACTGTCAGACTTGAGGCTGGAAGAGGCGAGTGGAATT<br>CCCGGTGTAGTCGTAATAAGCGCTGATATCGGGAGGAACGCCAAAGGTGAAAACAACTCGCTGGTACAAGCCTGACGCTCAGGGACGAAAGCATGGGT |
| BH-asv1059 | GCTAGCGTTAATCGGATTTACTGGGCGTAAAGCGTGCGTAGGCGGTTTGATAAGTCGATTGTGAAAGCCCCGGGCTTAACCTGGGAATTGCAGTTGATACTGTCAGACTCGAAAACGGTAGAGGGAGGCGGAAC<br>TCCAGGTGTAGCGGTGAAATGCGTAGATATCTGGAAGAACACCGATGGCGAAGGCAACCTCCTGGGCCTGTTTTGACGCTGAGGCACGAAAGCGTGGGT |
| BH-asv1060 | GCAAGCGTTAATCGGAATTACTGGGCGTAAAGCGAGCGTAGGTGGCTTGATAAGTCAGATGTGAAAGCCCCGGGCTTAACCTGGGAACGGCATCTGATACTGTTAGGCTAGAGTAGGTGAGAGGAAGGTAGAAT<br>TCCAGGTGTAGCGGTGAAATGCGTAGAGATCTGGAGGAATACCGATGGCGAAGGCAGCCTTCTGGCATCATACTGACACTGAGGTTCGAAAGCGTGGGT |
| BH-asv1061 | GCTAGCGTTGTTCGGAATGACTGGGCGTAAAGGGCGCGTAGGCGGTTTTTTAAGTGAGGCGTGAAAGCCCTGGGCTTAACCCAGGAGGTGCGTTTCATACTGGAAGACTTGAGTGCGAGAGAGGAAAGTGGAAT<br>TCCTAGTGTAGAGGTGAAATTCGTAGATATTAGGAAGAACACCCGTGGCGAAGGCGGCTTTCTGGCTCGTAACTGACGTTGAGGCGCGAAAGCGTGGGG |
| BH-asv1063 | GCAAGCGTTGTCCGGATTTACTGGGTGTAAAGGGTGCGCAGGCGGACTTGTAAGTCAGGGGTGAAATCCTACAGCTCAACTGTAGAACTGCCTTTGATACTGCAAGTCTTGAGTTCGGAAGAGAGAGGCGGAATT<br>CCAGGTGTAGTGGTGAAATACGTAGATATCTGGAAGAACACCAGAGGCGAAGGCGGTCTCTTGGTCCGTAACTGACGCTGAGGCACGAAAGCGTGGGG |
| BH-asv1064 | GCAAGCGTTATCCGGATTTACTGGGTTTAAAGGGTGCGTAGGCGGGCCATTAAGTCAGATGTGAAAGCCTCAAGCTCAACTTGAGAACTGCGTTTGATACTGGTGGCCTTGAATTCGATAGAAGTGGGCGGAACA<br>AATCGTGTAGCGGTGAAATGCTTAGATATGATTTGGAACACCAATTGCGAAGGCAGCTCACTATGCCGATATTGACGCTGAGGCACGAAAGCGTGGGG |
| BH-asv1065 | CCGAGCGTTGTTCGGAATCACTGGGCGTAAAGGGAGCGTAGGCGGCGTGGCAAGTCAGATGTGAAATCCCGGGGCTCAACCCCGGAACCGCATCCGATACTGCCGCGCTTGAGGATTGGAGAGGTAGCTGGAA<br>TTCTTGGTGTAGCAGTGAAATGCGTGGAGATCAAGAGGAACACTCGTGGCGAAAGCGAGCTACTGGACAATATCTGACGCTGAGGCTCGAAGGCCGGGGT |
| BH-asv1066 | GCAAGCGTTGCCCGGAATTACTGGGCGTAAAGAGCTCGTAGGCGGGGGCGTGCGTCCGGGAAGAAATCTCAAGGCTCAACCTTGGGGCTATCTCGGATACGGCGCTTCTTGAGGGTGAGAGAGGAAAGTAGAA<br>TTCCCGGTGTAGCGGTGAAATGCGTAGATATCGGGAGGAATACCAGCGGCGAAGGCGACTTTCTGGCTCATTCCTGACGCTGAGGAGCGAAAGCGTGGGT |
| BH-asv1068 | GCAAGCGTTATTCGGATTCACTGGGCGTAAAGGGTGCGCAGGCGGTCGTGTGTGTCAGGTGTGAAATCTCGGAGCTTAACTCCGAAACTGCGCCTGAAACTACACGACTAGAGTATCGGAGAGGTGAGCGGAAT<br>TCACGGTGTAGCAGTGAAATGCGTAGATATCGTGAGGAACACCAGAGGCGAAGGCGGCTCACTGGACGATTACTGACGCTCAGGCACGAAAGCGTGGGG |
| BH-asv1071 | GCGAGCGTTAATCGGAATTACTGGGCGTAAAGCGTGCGCAGGCGGTTTTGAAAGTCAGATGTGAAATCCCCGAGCTCAACTTGGGAACTGCGTTTGAAACTCCAAAGCTAGAGTATAGGAGAGGGGGGTAGAAT<br>TCCACGTGTAGCAGTGAAATGCGTAGAGATGTGGAGGAATACCAATGGCGAAGGCAGCCCCCTGGCCTAATACTGACGCTCATGCACGAAAGCGTGGGG |
| BH-asv1072 | GCAAGCGTTAATCGGAATTACTGGGCGTAAAGCGCGCGTAGGTGGTTTATTAAGTTGGATGTGAAATCCCCGGGCTCAACCTGGGAACTGCATTCAAAACTGACTGACTAGAGTATGGTAGAGGGTGGTGGAATT<br>TCCTGTGTAGCGGTGAAATGCGTAGATATAGGAAGGAACACCAGTGGCGAAGGCGACCACCTGGACTGATACTGACACTGAGGTGCGAAAGCGTGGGG |
| BH-asv1073 | GCAAGCGTTAATCGGAATTACTGGGCGTAAAGCGTGCGCAGGCGGTCTGTAAAGTCAGATGTGAAAGCCCCGGGCTCAACCTGGGAATTGCATTTGAAACTGGCAGGCTAGAGTGTGGCAGAGGGGGGTGGAA<br>TTCCACGTGTAGCAGTGAAATACGTAGAGATGTGGAGGAACACCGATGGCGAAGGCAGCCCCCTGGGCCAACACTGACGCTCAGGCACGAAAGCGTGGGG |
| BH-asv1074 | GCGAGCGTTGTCCGGAATCACTGGGCGTAAAGGGCGCGTAGGTGGCGCGGAAAGTCAACGGTGAAAGCTTGGGGCTCAACCCCGAGTCGGCCGTCGATACTTTCGTGCTCGAGCACTGTAGAGGCAGATGGAA<br>TTCCGGGTGTAGCGGTGGAATGCGTAGAGATCCGGAAGAACACCGGTGGCGAAGGCGGTCTGCTGGGCAGTTGCTGACACTGATGCGCGACAGCGTGGGG |
| BH-asv1075 | GCGAGCGTTAATCGGAATAACTGGGCGTAAAGGGCACGCAGGCGGTGATTTAAGTGAGGTGTGAAAGCCCCGGGCTTAACCTGGGAATTGCATTTCATACTGGGTCGCTAGAGTACTTTAGGGAGGGGTAGAAT<br>TCCACGTGTAGCGGTGAAATGCGTAGAGATGTGGAGGAATACCGAAGGCGAAGGCAGCCCCTTGGGAATGTACTGACGCTCATGTGCGAAAGCGTGGGG |
| BH-asv1076 | GCGAGCGTTAATCGGAATTACTGGGCGTAAAGTGTGCGCAGGCGGCCGAGCAAGTCGAGTGTGAAATCTCCGGGCTTAACCCGGAACGGTCAGCCGATACTGCAGGGCTAGAGTGCAGAAGGGGCAACTGGAA<br>TTCTCGGTGTAGCGGTGAAATGCGTAGATATCGAGAGGAACACCTGCGGCGAAGGCGGGTTGCTGGGCTGACACTGACGCTGAGGCGCGAAAGCTAGGGG |
| BH-asv1077 | GCTAGCGTTGTTCGGAATTACTGGGCGTAAAGCGCGCGTGGGCGGCTTCGCAAGTCAGAGGTGAAAGCCCGGGGCTCAACCCCGGAATTGCCTTTGAGACTGCGTCGCTAGAATCCGGGAGAGGTGAGTGGAA<br>TTCCGAGTGTAGAGGTGAAATTCGTAGATATTCGGAAGAACACCAGTGGCGAAGGCGGCTCACTGGACCGGTATTGACGCTGAGGTGCGAAAGCGTGGGG |
| BH-asv1078 | GCAAGCGTTGTCCGGATTTATTGGGCGTAAAGAGCGTGTAGGCGGCCAGACAGGTCCGTTGTGAAAACTGGAGGCTTAACCTCCAGACGTCGATGGAAACCGTCTGGCTAGATTCCGGAGGAGGAGAGTGGAAT<br>TCCTGGTGTAGCGGTGAAATGCGCAGATATCAGGAAGAACACCCGTGGCTAAGGCGGCTCTCTAGTACGGTACTGACGCTGAGACGCGAAAGCGTGGGG |
| BH-asv1079 | GCGAGCGTTGTCCGGATTTACTGGGCGTAAAGGGAGCGTAGGCGGACTTTTAAGTGAGATGTGAAATACCCGGGCTCAACTTGGGTGCTGCATTTCAAACTGGAAGTCTAGAGTGCAGGAGAGGAGAATGGAAT<br>TCCTAGTGTAGCGGTGAAATGCGTAGAGATTAGGAAGAACACCAGTGGCGAAGGCGATTCTCTGGACTGTAACTGACGCTGAGGCTCGAAAGCGTGGGG |
| BH-asv1081 | GCTAGCGTTGCTCGGAATTACTGGGCGTAAAGGGAGCGTAGGCGGACATTTAAGTCAGGGGTGAAATCCCGGGGCTCAACCTCGGAATTGCCTTTGATACTGGGTGTCTTGAGTATGAGAGAGGTGTGTGGAAC<br>TCCGAGTGTAGAGGTGAAATTCGTAGATATTAGGAAGAACACCAGTGGCGAAGGCGACACACTGGCTCATTACTGACGCTGAGGCTCGAAAGCGTGGGG |
| BH-asv1082 | GCAAGCGTTATCCGGATTTATTGGGTTTAAAGGGTCCGTAGGCGGACTTATAAGTCAGTGGTGAAATCCTGTCGCTTAACGATAGAACTGCCATTGATACTGTAAGTCTTGAGTATATTTGAGGTAGCTGGAATAAG<br>TAGTGTAGCGGTGAAATGCATAGATATTACTTAGAACACCAATTGCGAAGGCAGGTTACCAAGATATAACTGACGCTGAGGGACGAAAGCGTGGGT |
| BH-asv1083 | GCGAACGTTGCTCGGAATCATTGGGCGTAAAGCGCGCGTAGGCGGCCGGACAAGTTGGATGTGAAAGCCCTGGGCTCAACCCGGGATGTGCATTCAAAACTGTTCGGCTTGAATCTCGGAGGGGGTCAGGGAA<br>TTCCCGGTGTAGAGGTGAAATTCGTAGATATCGGGAGGAACACCAGTGGCGAAGGCGCTGACCTGGACGATGATTGACGCTGAGGCGCGAAAGCGTGGGG |
| BH-asv1084 | GCAAGCGTTGTCCGGATTTACTGGGTGTAAAGGGCGCGTAGGCTGGTTTGTTAGTCAGAGGTGAAATCCTGCAGCTCAACTGTAGAATTGCCTTTGATACTGCAAGCCTTGAGTGCGGAAGAGAGAGACGGAATT<br>CCAGGTGTAGTAGTGAAATACGTAGATATCTGGAAGAACACCAGAGGCGAAGGCGGTCTCTTGGTCCGTTACTGACGCTGATGCGCGAAAGCGTGGGG |
| BH-asv1085 | GCAAGCGTTAATCGGAATTACTGGGCGTAAAGCGTGCGCAGGCGGATGATTAAGTTAGGAGTGAAATCCCCGGGCTCAACCTGGGAACTGCTTTTAAGACTGGTCATCTAGAGTTTGTCAGAGGGGGGTGGAAT<br>TCCAAGTGTAGCAGTGAAATGCGTAGAGATTTGGAGGAACACCGATGGCGAAGGCAGCCCCCTGGGATAAAACTGACGCTCATGCACGAAAGCGTGGGT |
| BH-asv1086 | GCAAGCGTTGTCCGGATTTATTGGGCGTAAAGAGCTCGTAGGTGGCTGAGTAAGTCGGGTGTCAAATCCCCAGGCTCAACCTGGGAACCCACCCGATACTGCTCTGGCTAGAGTCCGGTAGGGGAGCGTGGAAT<br>TCCTGGTGTAGCGGTGAAATGCGCAGATATCAGGAGGAACACCAGCGGCGAAGGCGGCGCTCTGGGCCGGTACTGACACTGAGGAGCGAAAGCGTGGGT |
| BH-asv1087 | GCGAACGTTATTCGGAATCACTGGGCTTAAAGGGTGCGTAGGCGGCGAACTAAGTAGGGTGTGAAATCCCATCGCTCAACGATGGAACTGCGCTCTAAACTGGTTTGCTTGAGTAAGATAGGGGTGTGCGGAAC<br>TTCAAGTGGAGCGGTGAAATGTGTAGATATTTGAAGGAACACCGGTGGCGAAAGCGGCACACTAGGTCTTAACTGACGCTGAGGCACGAAAGCCAGGGT |
| BH-asv1088 | GCAAGCGTTAATCGGAATTACTGGGCGTAAAGCGCGCGTAGGTGGTTTGTTAAGTTGGATGTGAAAGCCCTGGGCTCAACCTAGGAACTGCATCCAAAACTAACTCACTAGAGTACGATAGAGGGAGGTAGAATT<br>CATAGTGTAGCGGTGGAATGCGTAGATATTATGAAGAATACCAGTGGCGAAGGCGGCCTCCTGGATCTGTACTGACACTGAGGTGCGAAAGCGTGGGT |
| BH-asv1089 | GCTAGCGTTGTTCGGAATGACTGGGCGTAAAGGGCGCGTAGGCGGCCCTCGCCGTCGGATGTGAAAGCCCGGGGCTCAACCCCGGAACGGCATCCGAGACGCGGGGGCTGGAGACCGGGAGATGAGGGTGG<br>AATACCCAGTGTAGAGGTGAAATTCGTAGATATTTGGTGGAACACCGGTGGCGAAAGCGGCTCTCTGGCCCGGTTCTGACGCTGAGGCGCGAAAGCGTGGGG |
| BH-asv1090 | GCAAGCGTTGTCCGGATTTACTGGGCGTACAGCGCGCGCAGACGGCACGCGACGTGCCCTGTGACAGCGCACCGCTTAACGGTGCGTCGCCAGGGTACACGAGCGCGCTGGAGGCAGGGAGAGGGATGTGG<br>AATGCCCGGTGGAGCGGTGAAATGCGTAGAGATCGGGCGGAACACCAGCGGCGAAGGCGGCATCCTGGCCCTGACCTGACGTTGAGGCGCGAGAGCGTGGGG |
| BH-asv1091 | GCAAGCGTTAATCGGAATTACTGGGCGTAAAGGGTGCGTAGGCGGTTAGTTAAGTCTGTTGTGAAATCCCCGGGCTCAACCTGGGAATGGCAATGGATACTGGCTAGCTAGAGTGTGTCAGAGGATGGTGGAAT<br>TCCCGGTGTAGCGGTGAAATGCGTAGAGATCGGGAGGAACATCAGTGGCGAAGGCGGCCATCTGGGACAACACTGACGCTGAAGCACGAAAGCGTGGGG |
| BH-asv1092 | GCAAGCGTTGTTCGGAATTACTGGGCGTAAAGGGCTCGTAGGCGGCCAACTAAGTCGGATGTGAAATCCCCAGGCTCAACCTGGGAACTGCATCCGATACTGGATGGCTTGAATCCGGGAGAGGGATGCAGAAT<br>TCCAGGTGTAGCGGTGAAATGCGTAGATATCTGGAGGAATACCGGTGGCGAAGGCGGCATCCTGGACCGGCATTGACGCTGAGGAGCGAAAGCCAGGGG |
| BH-asv1093 | GCAAGCGTTGTTCGGAATTACTGGGCGTAAAGGGCTCGTAGGCGGCCAACTAAGTCGGATGTGAAATCCCCAGGCTCAACTTGGGAACTGCATCCGATACTGGATGGCTTGAATCCGGGAGAGGGATGCAGAAT<br>TCCAGGTGTAGCGGTGAAATGCGTAGATATCTGGAGGAATACCGGTGGCGAAGGCGGCATCCTGGACCGGTATTGACGCTGAGGCGCGAAAGCTAGGGG |
| BH-asv1094 | GCAAGCGTTAATCGGAATTACTGGGCGTAAAGCGTGCGTAGGCGGTTATTTAAGTCAGATGTGAAAGCCCCGGGCTTAACCTGGGAACTGCATTTGATACTGGGTAACTAGAGTTTAATAGAGGAGAGTGGAATT<br>TCAGGTGTAGCGGTGAAATGCGTAGAGATCTGAAGGAACACCAGTGGCGAAGGCTACTCTCTGGATTAAAACTGACGCTGAGGTACGAAAGCGTGGGT |
| BH-asv1095 | GCAAGCGTTAATCGGATTTACTGGGCGTAAAGCGTGCGTAGGCGGCCAGACAAGTCGGATGTGAAATCCCCGAGCTTAACTTGGGAACTGCATCCGAAACTATCTGGCTAGAGTGTGGTAGAGGGCGGTGGAAT<br>TCCCGGTGTAGCGGTGAAATGCGTAGATATCGGGAGGAACATCAATGGCGAAGGCAGCTGCTTGGGCCAACACTGACGCTGAGGTACGAAAGCGTGGGG |
| BH-asv1096 | GCGAGCGTTGTTCGGAATTACTGGGCGTAAAGAGTGTGTAGGCGGTGCTCTAAGTTTGGTGTGAAATCTCCCGGCTTAACTGGGAGGGTGCGCCGAAGACTGGAGTGCTCGAGCGTGGGAGAGGAAAGCGGAA<br>TTCCTGGTGTAGCGGTGAAATGCGTAGATATCAGGAGGAACACCTGCGGTGTAGACGGCTTTCTGGACCATTGCTGACGCTGAGACACGAAAGCGTGGGT |
| BH-asv1098 | GCAAGAGTTATCCGGAATTATTGGGCGTAAAGCGCGCGTAGGCGGTTCGTTAAGTCTGATGTGAAAGCCCCCGGCTCAACCGGGGAGGGTCATTGGAAACTGGCGGACTTGAGTGCAGAAGAGGAGAGTGGAA<br>TTCCATGTGTAGCGGTGAAATGCGCAGAGATATGGAGGAACACCAGTGGCGAAGGCGGCTCTCTGGTCTGCAACTGACGCTGAGGTGCGAAAGCGTGGGG |
| BH-asv1099 | GCTAGCGTTGCTCGGAATTACTGGGCGTAAAGGGAGCGTAGGCGGACTGTTTAGTCAGAGGTGAAAGCCCAGGGCTCAACCTTGGAATTGCCTTTGATACTGGCAGTCTTGAGTACGGAAGAGGTATGTGGAAC<br>TCCGAGTGTAGAGGTGAAATTCGTAGATATTCGGAAGAACACCAGTGGCGAAGGCGACATACTGGTCCGTTACTGACGCTGAGGCTCGAAAGCGTGGGG |
| BH-asv1100 | GCGAGCGTTAATCGGAATTACTGGGCGTAAAGCGCGCGTAGGTGGTTAATTAAGTCAGATGTGAAAGCCCAGGGCTTAACCTTGGAACTGCATTTAATACTGATTAACTAGAGTTTTGTAGAGGGTGGTAGAATTT<br>CAGGTGTAGCGGTGAAATGCGTAGAGATCTGAAGGAATACCAGTGGCGAAGGCGGCCACCTGGACAAAGACTGACACTGAGGCGCGAAGGCGTGGGG |
| BH-asv1101 | GCAAGCGTTGTTCGGAATTATTGGGCGTAAAGGGCGCGCAGGCGGCCTCGTATGTCTGCTGTAAAAGCACACAGCTCAACTGTGTAACGTCAGTGGAAACTGCAAGGCTTGAGGTTGGGAGAGGCTGGTGGAAT<br>TCCCTGTGTAGCGGTGAAATGCGTAGATATAGGGGAGAACACTCGTGGCGAAGGCGGCCAGCTGGACCAGACCTGACGCTGAGGCGCGATAGCGTGGGG |
| BH-asv1102 | CCAAGCGTTATCCGGATTTATTGGGTTTAAAGGGTGCGTAGGTGGCTTTTTAAGTCAGGAGTGAAAGACGGCAGCTTAACTGTCGCAGTGCCCTTGATACTGAAGAGCTTGAATCCACTAGAGGTAGGCGGAATG<br>TGACAAGTAGCGGTGAAATGCATAGATATCGCGAGGAACACCCGTGGCGAAAGCGGCCTGCTGGATGGTTCCTGACGCTGAGGCTCGAAAGCGTGGGT |
| BH-asv1103 | CCAAGCGTTATCCGGATTTATTGGGTTTAAAGGGTGCGTAGGCGGCCTGTTAAGTCAGGGGTGAAAGACGGTAGCTCAACTATCGCAGTGCCCTTGATACTGATGGGCTTGAATGGACTAGAGGTAGGCGGAAT<br>GAGACAAGTAGCGGTGAAATGCATAGATATGTCTCAGAACACCGATTGCGAAGGCAGCTTACTATGGTCTTATTGACGCTGAGGCACGAAAGCGTGGGG |
| BH-asv1104 | GCTAGCGTTGTCCGGAATCACTGGGCGTAAAGGGTTCGCAGGCGGAAATGCAAGTCAGATGTAAAAGGCAGTAGCTTAACTACTGTAAGCATTTGAAACTGCATATCTTGAGAAGAGTAGAGGTAAGTGGAATTTT<br>TAGTGTAGCGGTGAAATGCGTAGATATTAAAAAGAATACCGGTGGCGAAGGCGACTTACTGGGCTCATTCTGACGCTGAGGAACGAAAGCGTGGGT |
| BH-asv1105 | GCGAGCGTTGTTCGGAATTATTGGGCGTAAAGCGCTTGTAGGCGGTCAATCAAGTCCTGTGTGAAAGCCCATGGCTTAACCATGGAAGTGCGCAGGAAACTGATGGACTTGAGTGCCGGAGAGGGTCGCGGAAT<br>TCCCGGTGTAGAGGTGAAATTCGTAGATATCGGGAGGAACACCAGTGGCGAAGGCGGCGACCTGGACGGACACTGACGCTGAGACGCGAAAGCGTGGGG |

| | |
|---|---|
| BH-asv1107 | GCTAGCGTTGTTCGGAATTACTGGGCGTAAAGGGCGCGTAGGCGGCTTTGTAAGTCAGATGTGAAAGCCCCGGGCTCAACCTGGGAACTGCATTTGATACTGCAAGGCTAGAGATCGAGAGAGGAAAGTGGAATTACGAGTGTAGAAGTGAAATTCGTAGATATTCGTAGGAACACCAGTGGCGAAGGCGACTTTCTGGCTCGATACTGACGCTAAAGCGCGAAAGCGTGGGG |
| BH-asv1108 | CCAGGCGTTATCCGGATTTATTGGGTTTAAAGGGAGCGTAGGCTGGAGATTAAGTGTGTTGTGAAATGTAGACGCTCAACGTCTGAATTGCAGCGCATACTGGTTTCCTTGAGTACGCACAACGTTGGCGGAATTCGTCGTGTAGCGGTGAAATGCTTAGATATGACGAAGAACTCCGATTGCGAAGGCAGCTGACGGGAGCGCAACTGACGCTTAAGCTCGAAGGTGCGGGT |
| BH-asv1109 | ACGAGCGTTGTCCGGATTCATTGGGCGTAAAGCGCGCGTAGGCGGCTCGGAAAGTCGGTCGTGAAATCCTGGGGCTCAACCCCGGAACTGCGTCCGATACTTCCGAGCTGGAGGCAGGTAGGGGAGATCGGAATTCCTGGTGTAGCGGTGAAATGCGCAGATATCAGGAGGAACACCGGTGGCGAAGGCGGATCTCTGGGCCTTGCCTGACGCTGAGGCGCGAAAGCTAGGGG |
| BH-asv1110 | GCAAGCGTTGTCCGGAATCACTGGGCGTAAAGCGCGCGCAGGCGGCTGTCTACGTCCGGGGTGAAAGCCCAGAGCTCAACTCTGGGACTGCCTTGGATACGGGATGGCTTGAGGATCGGAGAGGCAAGGGGAATTCCACGTGTAGCGGTGAAATGCGTAGAGATGTGGAGGAACACCTGTGGCGAAGGCGCCTTGCTGGCCGATTTCTGACGCTGAGGCGCGAAAGCGTGGGG |
| BH-asv1111 | GCAAGCGTTAATCGGAATTACTGGGCGTAAAGCGTGCGTAGGCGGTCCGCTAAGACAGATGTGAAATCCCCGGGCTTAACCTGGGAACTGCATTTGTGACTGGCGGGCTAGAGTATGGCAGAGGGGGGTAGAATTCCACGTGTAGCAGTGAAATGCGTAGAGATGTGGAGGAATACCGATGGCGAAGGCAGCCCCCTGGGCCAATACTGACGCTCATGCACGAAAGCGTGGGG |
| BH-asv1112 | GCAAGCGTTAATCGGAATTACTGGGCGTAAAGCGTGCGCAGGCGGTTTGTTAAGTCTGTTGTGAAATCCCCGAGCTCAACTTGGGAATTGCAATGGATACTGGCAAGCCGGAGTATGGTAGAGGAAGGTGGAATTCCCGGTGTAGCAGTGAAATGCGTAGAGATCGGGAGGAACACCAGTGGCGAAGGCGGCCTTCTGGACCAATACTGACGCTCATGCACGAAAGCGTGGGG |
| BH-asv1113 | CCAAGTGGCATTCGGAGTTACTAGGTCTTAAGCGTTCGTAGCCGGCCGTGTAAGTTCTGGGTGAAATCGTTCCACTCAACGGAACGACGTGCTCGGAATACTGCATGGCTTGGGACCGGGAAGCGTCAGAGGTATTCTAGGGGGGAGCGGTAAAATGCTATAATCCTTAGAAGACCACCTATGGCGAAAGCATCTGACGAGAACGGGTCCGACGGTCAGGAACAAAAGCTAGGGG |
| BH-asv1114 | GCAAACGTTGTTCGGAATTACTGGGCGTAAAGCGCGTGTAGGCGGCTATGCAAGTCGGGCGTGAAATCCCACGGCTCAACCGTGGAAGTGCACTCGAAACTGCGTAGCTAGAGTCCCGGAGAGGAAGGTGGAATACTTGGTGTAGAGGTGAAATTCGTAGATATCAAGTGGAACACCGGTGGCGAAGGCGGCCTTCTGGACGGTGACTGACGCTGAGACGCGAAAGCGTGGGG |
| BH-asv1115 | GCTAGCGTTGCTCGGAATCACTGGGCGTAAAGGGCGCGTAGGCGGCTTGCCAAGTCGGGGGTGAAAGCCCGTGGCTCAACCACGGAATGGCCTTCGATACTGGCAGGCTTGAGACCGGAAGAGGACAGCGGAACTGCGAGTGTAGAGGTGAAATTCGTAGATATTCGCAAGAACACCAGTGGCGAAGGCGGCTGTCTGGTCCGGTTCTGACGCTGAGGCGCGAAAGCGTGGGG |
| BH-asv1116 | GCAAGCGTTAATCGGAATGACTGGGCGTAAAGGGCGAGTAGGCGGTTGTATAGGTGTGGTGTGAAAGACACGGGCTTAACCTGTGGGTGTCACCGCAAACGGTATGACTAGAGTGCAGTAGAGGGGAGTGGAATTTCCGGAGGAGCGGTGAAATGCGTAGATCTCGGAAGGAACACCGGTGGCGAAGGCGACTCTCTGGACTGTAACTGACGCTGAGGCGCGAAGGCGTGGGG |
| BH-asv1117 | GCAAGCGTTATCCGGATTTATTGGGTTTAAAGGGTCCGTAGGCGGAACTTTAAGTCAGTGGTGAAAGCCTACAGCTCAACTGTAGAACTGCCATTGATACTGAAGTTCTTGAATATATATGAAGTGGGCGGAATGCGTCATGTAGCGGTGAAATGCACAGATATGACGCAGAACACCAATTGCGAAGGCAGCTCACTAATGTATTATTGACGCTGAGGGACGAAAGCATGGGG |
| BH-asv1118 | GCAAGCGTTATCCGGAATCATTGGGCGTAAAGGGTCCTGAGGTGGCCTGCCCAGTCTACTGTGAAAGATCAGGGCTCAACCCTGAAGATGCGGTAGAAACTAGCAGGCTAGGATACGGTAGGGGCAGGGGGAATTCCCGTTGTAGCGGTGAAATGCATAGATATCGGGAGGAACACCAAAGGCGAAGGCACCCTGCTGGGCCGATATTGACACTGACAGACGAGAGCTAGGGG |
| BH-asv1119 | GCAAGCGTTGTCCGGATTTATTGGGTTTAAAGGGTGCGTAGGCGGCTTAATAAGTCAGTGGTGAAAGCCGGTAGCTTAACTATCGAAATGCCATTGATACTGCTAGGCTTGAGTACAGTTGAGGTAGGCGGAATTGACAGTGTAGCGGTGAAATGCTTAGATATTGTCAAGAACACCAATTGCGAAGGCAGCTTACTAAAGTGTAACTGACGCTGAGGCACGAAAGTGCGGGG |
| BH-asv1120 | CCGAGCGTTATCCGGATTTATTGGGTTTAAAGGGTGCGTAGGCGGTTTTATAAGTCAGCGGTGAAATTTTCCGGCTCAACCGGGATACTGCCGTTGATACTGTAAGGCTAGGATATGGTTGCTGTGGGTGGAATGTGTGGTGTAGCGGTGAAATGCTTAGAGATCACACAGAATATCGATTGCGAAGGCAGCTCACAAATCCATTATTGACGCTGAGGCACGAAAGTGTGGGG |
